# RNA-sequencing of butyrate-treated human lymphoblastoid cell lines shows dysregulated genes of multiple disease-related pathways

**DOI:** 10.64898/2026.10.21.760620

**Authors:** Hadas Volkov, Tehila Cohen, Rani Shlayem, Elad Shapira, Irena Voinsky, Hana Klin, Meitar Grad, Noam Shomron, David Gurwitz

**Author notes:** Authors for correspondence: Noam Shomron David Gurwitz.

## Abstract

Butyrate, a 4-carbon fatty acid produced from insoluble starch by the gut microbiome, plays key roles in mitochondrial respiration, activation of cell-surface G-protein-coupled receptors, and protein acetylation by inhibiting histone deacetylases (HDACs). Butyrate (as sodium salt) added to drinking water was shown to improve pathological symptoms in numerous model mouse disorders, including models for cancer metastasis, neuroinflammation, autism spectrum disorders, dementia, insulin resistance, and colorectal cancer. Over 5000 PubMed-listed studies feature butyrate in their title, yet its genome-wide transcriptomic effects have not been reported in human cell lines from multiple healthy donors. Here, we report the genome-wide transcriptomic effects of 1 mM sodium butyrate on human lymphoblastoid cell lines (LCLs) generated from blood samples of eight unrelated healthy donors. Paired bioinformatics analysis comparing the butyrate-treated LCLs to the same untreated cell line identified thousands of differentially expressed gene (DEG) transcripts with padj<0.05, of which over half were upregulated by butyrate. Pathway analysis showed that butyrate exposure of LCLs upregulated extracellular matrix, cell adhesion, junction organization, Wnt signaling, axonogenesis, and neurodevelopment pathways. Downregulated pathways included cell cycle, G2/M checkpoint, E2F/MYC targets, DNA replication and repair, mitotic spindle integrity, mTOR signaling, oxidative phosphorylation, unfolded-protein response, and translation/ribosome-related pathways. Our RNA-seq study, utilizing LCLs from healthy donors and paired bioinformatic analysis, opens venues for assessing butyrate effects on cellular pathways and potential therapeutics for cancer, metabolic, immune, and CNS-related disorders.

## 1. Introduction

The human intestinal microbiome composition has gained wider attention from the biomedical research community over the past two decades. Among the best-studied metabolites influencing host physiology is the 4-carbon fatty acid butyrate, generated primarily through bacterial fermentation of dietary carbohydrates that escape digestion in the upper gastrointestinal tract (Koe et al. 2026; Louis et al. 2027; Zhang el al. 2026). Butyrate has gained particular interest owing to its prominent effects on intestinal epithelial metabolism, immune regulation, chromatin organization, and gene transcription. The quantity of butyrate produced in the intestine depends on dietary fiber (starch) consumption, microbial community composition, intestinal transit, and host gut absorption and metabolism (Zhang el al. 2026). Major butyrate-producing bacteria include members of the families *Lachnospiraceae* and *Ruminococcaceae*, in particular, *Faecalibacterium*, *Roseburia*, *Eubacterium*, and *Anaerobutyricum* species (Louis et al. 2027).

Butyrate is among the key energy sources for the gut colonocytes (Donohoe et al, 2011, 2012). Studies in germ-free mice demonstrated that the absence of microbial butyrate was associated with reduced colonocyte mitochondrial respiration and increased autophagy, which were relieved by dietary butyrate supplementation, indicating its key roles for gut health (Donohoe et al. 2011).

Butyrate has emerged as both a gut metabolic substrate, and a key regulator of protein acetylation through inhibition of histone deacetylases (HDACs) enzymes. Butyrate is known to inhibit zinc-dependent HDAC enzymes (Boffa et al. 1978; Candido et al. 1978; Davie et al. 2003, Steliou et al. 2012). Histone acetylation is controlled by the opposing activities of histone acetyltransferase and HDAC enzymes and includes histone butyrylation by microbiome-derived butyrate (Gates et al. 2024). These enzymes affect chromatin accessibility to transcription factors and thereby their transcriptional activity. Butyrate was established as a low-potency HDAC inhibitor capable of altering acetylation by histones, transcription factors, and non-histone proteins (Davie et al. 2003, Steliou et al. 2012). As individual HDACs participate in distinct transcriptional complexes, butyrate exposure does not simply produce generalized transcriptional activation. Rather, butyrate may increase or decrease the transcription of different genes, with effects depending on local chromatin structure, transcription-factor availability, and each cell’s metabolic status. Notably, most studies on the metabolic, epigenomic, or clinical effects of butyrate applied in vivo butyrate supplementation studies in rodent disease models, or in vitro studies in human or rodent cell lines. However, metabolic and epigenomic effects of butyrate associated with HDAC inhibition were reported in insects, including Drosophila (St Laurent et al. 2013; Zhou et al. 2021) and honeybees (Liu et al. 2026).

Genome-wide studies in human cells, cell lines, or tissues have shown that butyrate may alter the transcription of thousands of genes, depending on cell type, its metabolic status, treatment duration, and other experimental conditions. However, most published RNA-seq studies in human cells applied a single cell type, including human colorectal cancer cell lines or transfected cells. Results from genome-wide transcriptomics studies with an individual cell line may thus reflect its genetic, epigenetic, metabolic, or oncogenic characteristics and may not distinguish between widely reproducible responses compared with donor-specific effects. A paired design in which each cell line is compared with the same cell line exposed to butyrate can substantially reduce this source of variation and improve the identification of consistent treatment-associated transcriptional changes.

Our study applied human lymphoblastoid cell lines (LCLs) that offer several advantages for paired transcriptomic experiments, where each butyrate-treated LCL is compared to the same immortalized cells grown under control conditions. This allows LCLs from unrelated donors to be cultured in parallel under similar conditions. This pairing analysis helps to distinguish treatment effects from stable inter-individual differences. LCLs have been widely employed in population genomics, expression quantitative trait locus mapping, pharmacogenomics, toxicogenomics, and studies of gene-by-environment interactions. Analyses of HapMap and 1000 Genomes LCL collections helped establish that inherited genetic variants contribute substantially to inter-individual differences in basal gene expression and transcript structure (Stranger et al. 2007; Choy et al. 2008; Jiang et al, 2017; Pickrell et al. 2010; Montgomery et al. 2010; Battle et al. 2014). LCLs have also been used to investigate genetically determined variation in cellular responses to drugs, metabolic stressors, and other experimental perturbations (Choy et al. 2008; Montgomery et al. 2010).

Keeping these considerations in mind, our genome-wide RNA-seq study designed to explore the transcriptomic effects of butyrate using RNA-seq applied a panel of eight human LCLs grown in parallel under similar conditions (including passage numbers, and batches of medium, FBS, and antibiotics) harvested and processed for RNA extraction with similar RNA extraction kit, and sequenced on the same machine by the same commercial RNA-seq provider. Moreover, the RNA-seq data were analyzed using the paired analysis design described above, allowing the detection of 10,056 transcripts as differentially expressed genes (DEGs) following the LCLs exposure to sodium butyrate (1 mM, 48 h) compared with the matched control LCLs according to the DEG cutoff criteria (see Methods).

## 2. Materials and Methods

### 2.1 Human Lymphoblastoid Cell Lines (LCLs)

Our genome-wide RNA-seq study applied human LCLs representing eight unrelated donors from the large human diversity collection of LCLs from healthy adult donors maintained by the National Laboratory for the Genetics of Israeli Populations (NLGIP; https://www.nlgip-yoran.sites.tau.ac.il). All donors reported being free from diabetes, cancer, or other severe disorders. Access to the medical records of the blood donors of the NLGIP biobanked LCLs was not possible. Notably, many of the Israeli human diversity collection LCLs are also available from the Coriell Institute for Medical Research (Camden, NJ, USA; https://www.coriell.org/).

### 2.2 Cell culturing and sodium butyrate treatment

LCLs were grown under optimal conditions as described in our recent transcriptomic studies. In short, cells were grown in RPMI supplemented with 10% FBS, 2 mM extra glutamine, 100 U/ml penicillin, and 100 ug/ml streptomycin (Shapira et al. 2025). LCLs were grown in upright T-25 flasks at cell densities of 0.5 × 10^6^ to 1 × 10^6^ cells/mL for up to 10 passages from thawing a frozen vial. Sodium butyrate (B5887; Sigma-Aldrich, USA) was added for the last 48 h before cell harvesting (∼5 ml) by centrifugation in parallel with control and butyrate-treated LCL cultures (5 min; 1700 RPM; 15 ml tubes). Cell pellets were gently resuspended in 1.5 ml cold PBS and re-centrifuged (5 min; 1700 RPM; 2 ml Eppendorf tubes). The supernatants were discarded, and the washing step was repeated. The final pellets were stored at -80^0^C in marked tubes until their parallel RNA extractions.

The concentration of 1 mM butyrate and the exposure period of 48 h were chosen for studying butyrate’s transcriptomic effects as they allowed consistent gene expression effects on several selected genes (Shapira et al. 2025). Treating LCLs with 0.5 mM or 2 mM sodium butyrate resulted in less consistent effects. Additionally, treatments with 1 mM sodium butyrate for 24 h resulted in smaller effects, and treatments for 72 h resulted in closely similar transcriptomic effects as for 48 h treatments, while the density of treated cells was reduced compared with 48 h (not shown).

### 2.3 RNA Extraction

RNAs were extracted in parallel from each of the 16 LCL pellets (8 LCLs, each without or with 1 mM sodium butyrate treatment for 48 h). RNAs were extracted from the thawed cell pellets using Zymo Research Quick-RNA MiniPrep kit #R1054 (Irvine, CA, USA). RNA quality and yields were determined by a NanoDrop Microvolume Spectrophotometer (Thermo Fisher Scientific, MA, USA), to ensure both 260/280 nm and 260/230 nm parameters > 2.0 and RIN>8.5 for RNA-seq and stored at −80°C

RNA samples were quantified by NanoDrop spectrophotometer (ThermoFisher Scientific, MA, USA), with both 260/280 nm and 260/230 nm parameters > 2.0 and stored at −80°C until analysis (Voinsky et al. 2024).

### 2.4 RNA-sequencing (RNA-seq)

RNA samples were poly-A mRNA sequenced. Libraries were prepared using a TrueSeq stranded total RNA LT sample prep kit (Illumina, San Diego, CA, USA). RNA sequencing was performed as paired end reads on the Truseq stranded mRNA + Novaseq X 2x100bp platform. Sequencing depth was 40 M reads/sample. Raw sequencing data were trimmed using fastp 0.20.0 (Chen et al. 2018) and aligned to the GRCm38 assembly using STAR 2.6.0c (Dobin et al. 2023). DESeq2 1.30.1 (Love et al. 2014) and R 3.6 were used for normalization of count data.

### 2.5 RNA-seq differential gene expression analysis and pathway analysis

Raw paired-end FASTQ files were preprocessed, and quantification was performed with nf-core/rnaseq v3.26.0 using Docker containers. Reads were processed as stranded mRNA-seq data and aligned to the human GRCh38 primary assembly with HISAT2. Gene-level quantification was generated with Salmon using GENCODE v50 annotation.

Differential transcription analysis was performed with DESeq2 (Love et al. 2014) using gene-level length-scaled counts from Salmon. The primary model was a paired donor-blocked design: “expression ∼ donor_id + condition” where condition compares 1 mM sodium butyrate-treated LCLs (48 h) with their matched untreated control LCLs. This design controls donor-specific baseline expression and provides the main statistical power of the study (see Discussion).

Positive fold-change values indicate higher expression after butyrate treatment. P-values were adjusted for multiple testing using the Benjamini-Hochberg false discovery rate (FDR) procedure (Ferreira 2007). Genes were defined as differentially expressed (DEGs) using padj < 0.05 and absolute linear fold difference >= 1.2. Genes encoding mitochondrial proteins were identified using the MitoCarta3.0 database (Li et al. 2024).

Pathway analysis was performed using preranked Gene Set Enrichment Analysis (GSEA) with GSEApy v1.3.0. All genes tested in the paired DESeq2 model were ranked by the DESeq2 Wald statistics (Terra Machado et al. 2023). Positive normalized enrichment scores (NES) indicate enrichment among genes upregulated by butyrate; negative NES values indicate enrichment among genes downregulated by butyrate. Gene sets were taken from MSigDB Hallmark (Liberzon et al. 2015), Reactome Pathways 2024, and GO Biological Process Gene sets with 10-500 overlapping genes were tested using 10,000 permutations. Pathways were considered significant at FDR q-value < 0.05.

### 2.6 Real-Time qPCR measurements

Nine genes were selected from the RNA-seq for validation using real-time quantitative PCR (RT-qPCR). These genes included three butyrate upregulated genes, three butyrate downregulated genes, and three genes dysregulated in blood samples of children diagnosed with autism spectrum disorder (ASD) compared with neurotypically developing age-matched children: *BATF2, LY6E*, and *MT2A* (Voinsky et al. 2022, 2023; Frye et al. 2024).

RT-qPCRs were performed with 1 μg RNA samples converted to cDNA using the qScript cDNA Synthesis Kit (Quanta Bio, Beverly, MA, USA). Reverse transcription (RT) was performed using a thermal cycler over three steps (22°C for 5 min, followed by 42°C for 30 min, and 85°C for 5 min). RT-qPCR reactions were performed in mixtures containing 10 ng of cDNA, PerfeCTa SYBR® Green FastMix Kit (Quanta Bio, Beverly, MA, USA), and Integrated DNA Technologies, Inc. (Leuven, Belgium) Primers. Forward and reverse primer sequences for RT-qPCR are listed in Supplementary Table S3.

## 3. Results

### 3.1 Differentially expressed genes (DEGs) in butyrate-treated LCLs

Our RNA-seq analysis applied GENCODE v50 GRCh38 and examined the transcriptomics of eight LCLs from unrelated individuals following butyrate treatment compared with untreated controls. In the paired analysis, the most informative and powerful analysis, each of the eight butyrate-treated LCLs was compared to its untreated control (see Methods). As shown, this paired analysis identified 10,056 differentially expressed genes (DEGs) in butyrate-treated LCLs with FD>1.2 or <0.8 and padj <0.05 (Supplementary Table S1). This table presents the RNA-seq paired analysis. Figure 1 shows a Volcano plot for the DEGs identified using the paired analysis (see Methods). Notably, about 25% fewer DEGs were detected using the group analysis compared with the paired analysis (with similar FD and padj cutoff criteria). The DEGs identified by paired analysis included 645 genes coding for mitochondrial proteins, of which 495 (76.7%) were downregulated in butyrate-treated LCLs (Supplementary Table S2). This contrasts with the slightly higher rate of butyrate-upregulated DEGs in Supplementary Table S1.

**Figure 1:**
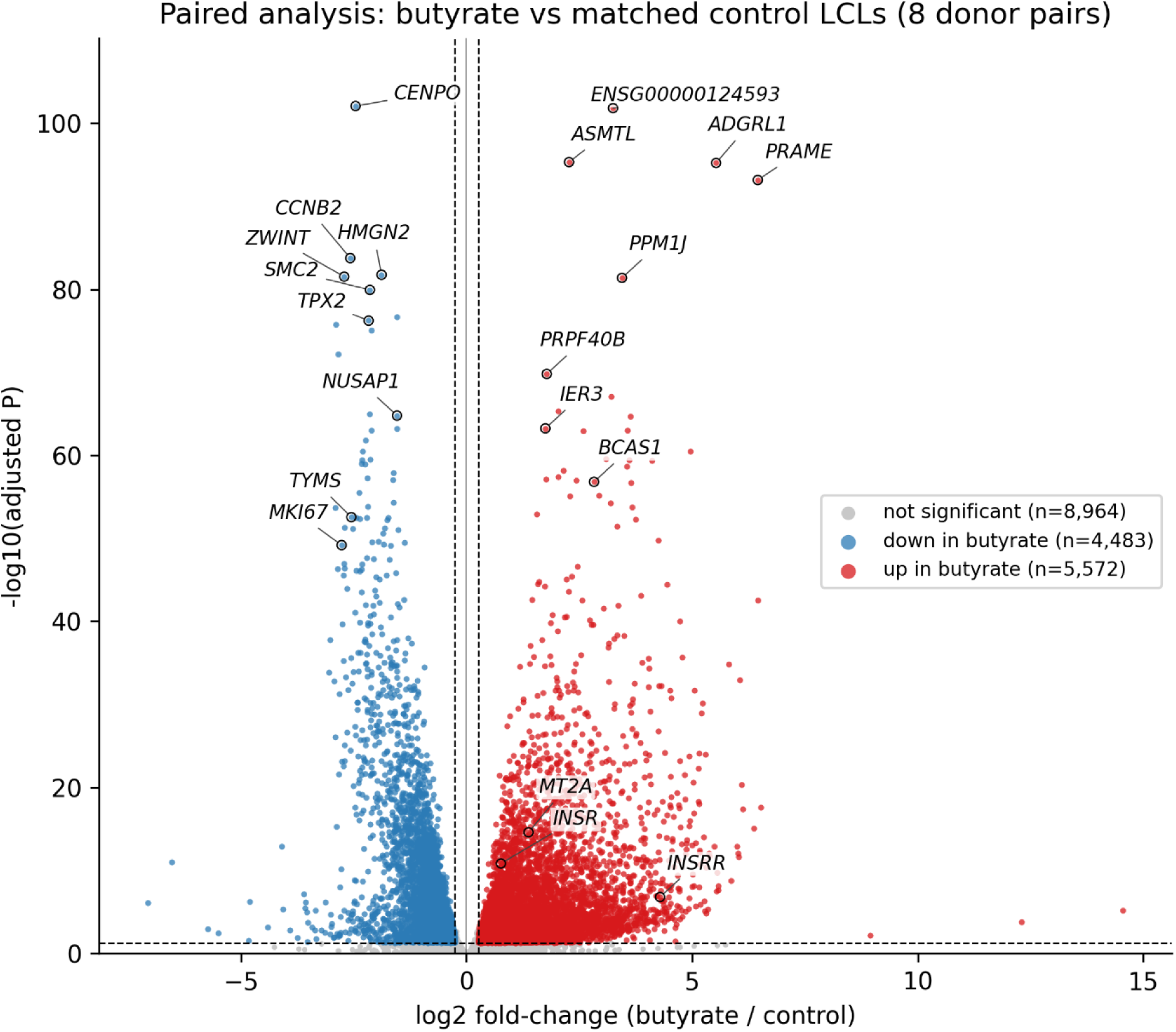
A Volcano plot for DEGs identified by RNA-seq using paired analysis. The transcriptome of each butyrate-treated LCL (N=8 donors) was compared to its untreated control. The x-axis represents the log2 fold difference between butyrate-treated and control samples, while the y-axis represents −log10 of the adjusted p-value. Upregulated and downregulated genes are shown in red and blue, respectively. The gene symbols of selected upregulated and downregulated DEGs are labelled.

### 3.2 Real-time qPCR validation (RT-qPCR)

Nine DEGs were selected for RT-qPCR validation (Table 1). Three DEGs were genes of interest for ASD diagnostics (see Introduction and its citations). The other six DEGs were arbitrarily selected for RT-qPCR validation from the top DEGs (three butyrate upregulated and three butyrate downregulated genes). RNA-seq and RT-qPCR findings are compared in Table 1. RT-qPCR findings (Figure 2) indicate general validation for eight of the nine selected genes.

**Figure 2.**
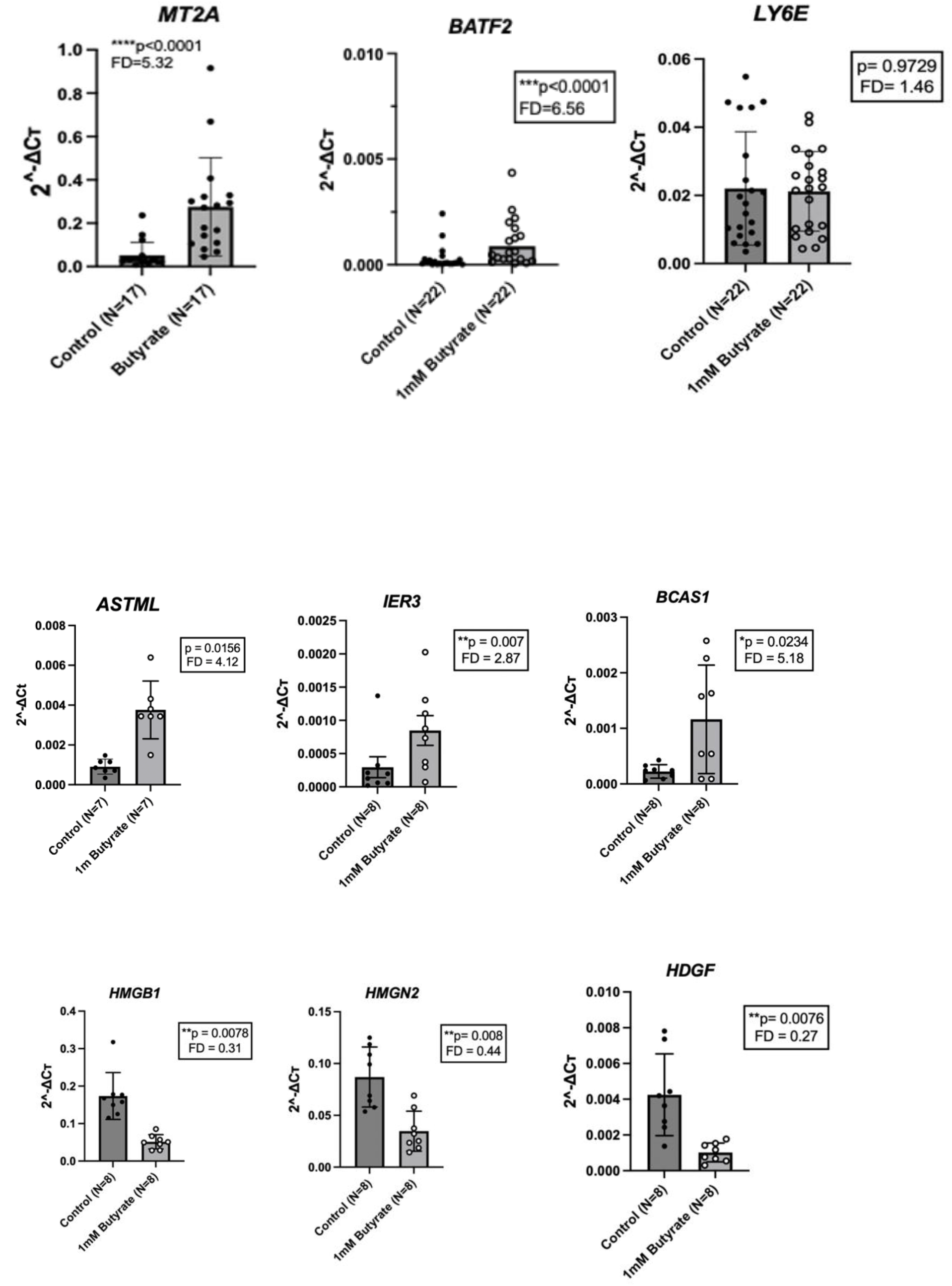
RT-qPCR validation of selected RNA-seq findings. The sodium butyrate (1 mM; 48 h) effects on the transcription of selected genes are shown in dot-plots in up to 22 LCLs. Butyrate effects on the RNA-seq paired analysis transcription of selected genes are shown in Table 1 and Supplementary Fig. S1. See Table 1 for further information.

**Table 1.**
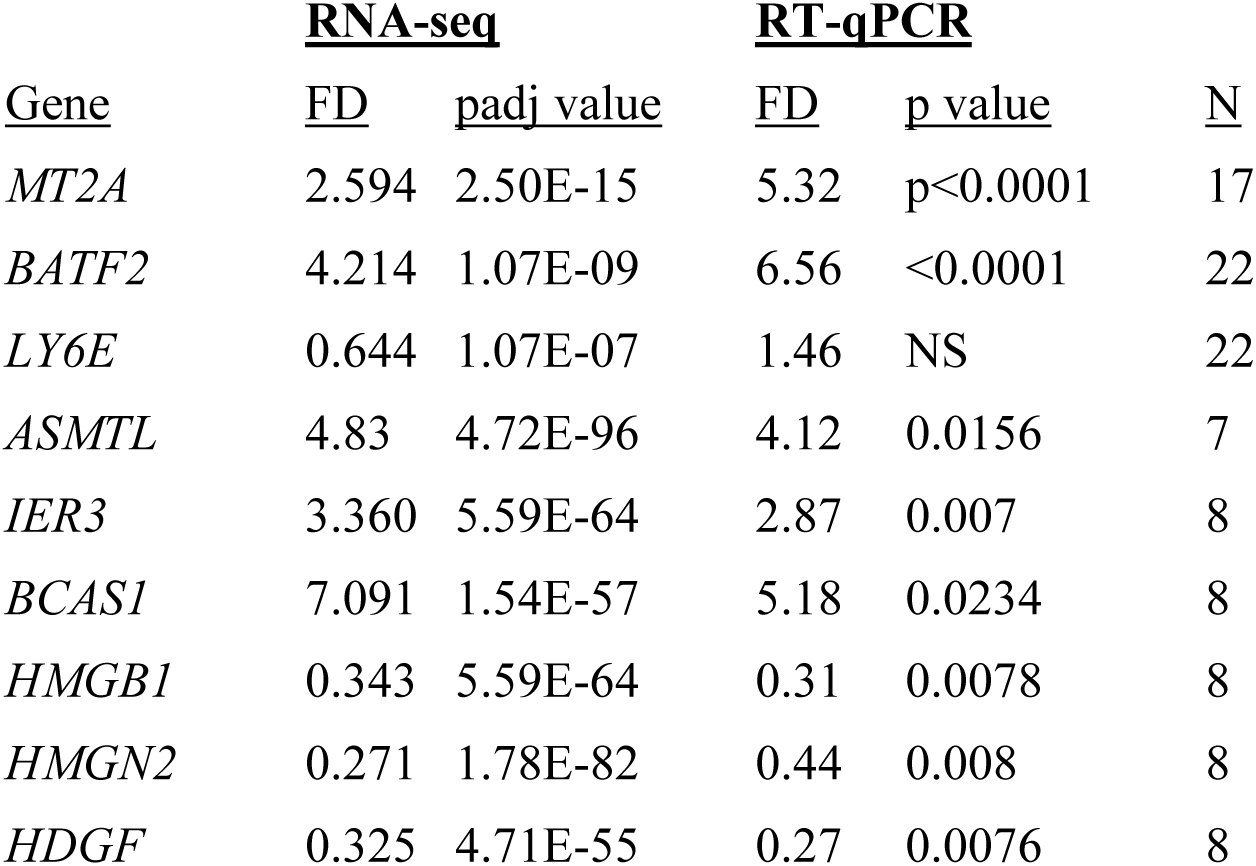
Nine DEGs selected for RT-qPCR validation of RNA-seq findings. The RT-qPCR findings FD and p values were calculated using a two-tailed Wilcoxon matched-pairs signed-rank test (N = 8 paired LCLs). The RT-qPCR findings are displayed as dot-plots in Fig. 2 and were obtained from the same 8 LCLs as in Table 1, except for *MT2A*, *BATF2* and *LY6E* findings that were obtained from larger LCL cohorts. The complete list of DEGs FD and padj values from the RNA-seq paired analysis of butyrate-treated vs. control LCLs (N=8 pairs) is shown in Supplementary Table S1.

### 3.3 Pathway analysis

GENCODE v50 GRCh38 (https://www.gencodegenes.org/human/) was applied for pathway analysis of DEGs in butyrate-treated LCLs. The analysis indicated that the upregulated pathways included extracellular matrix, cell adhesion, junction organization, Wnt signaling, axonogenesis, and neurodevelopment. The downregulated pathways included cell cycle, G2/M checkpoint, E2F/MYC targets, DNA replication and repair, mitotic spindle integrity, mTOR signaling, oxidative phosphorylation, unfolded-protein response, and translation/ribosome-related pathways (Figure 3).

**Figure 3.**
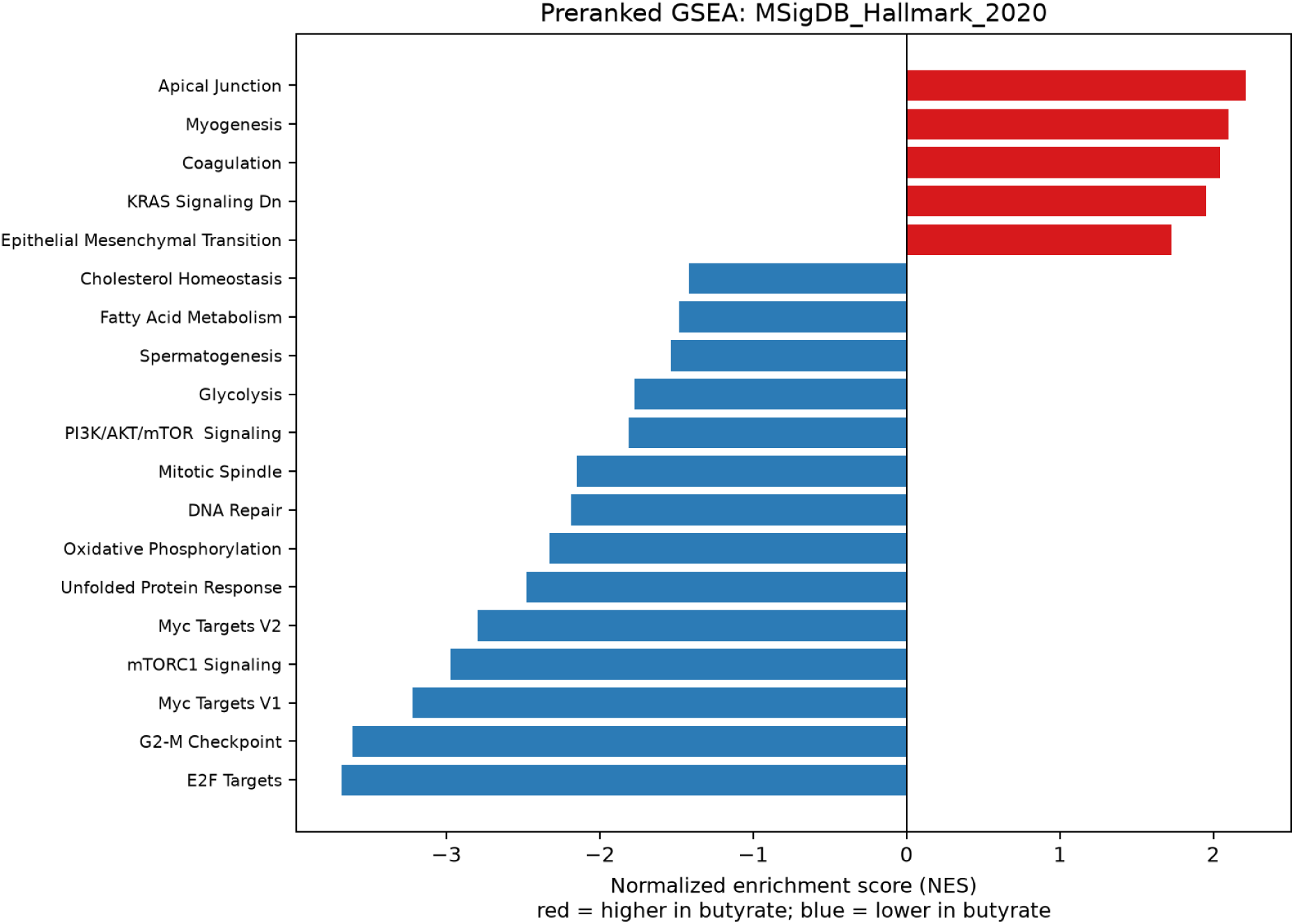
Reactome pathway analysis of butyrate response in human LCLs. Reactome preranked GSEA showed broad downregulation of mitotic cell-cycle checkpoint, chromatid-segregation, DNA-replication, and DNA-repair pathways following butyrate treatment. Upregulated pathways included extracellular matrix organization, collagen/laminin interactions, integrin signaling, and related adhesion pathways. Butyrate treatment downregulated cell-cycle, E2F/MYC, DNA repair, mTOR, oxidative phosphorylation, and unfolded-protein-response programs.

GO Biological Process preranked GSEA confirms strong downregulation of mitotic cell-cycle, chromosome segregation, DNA replication/repair, translation, and mitochondrial translation pathways after butyrate treatment. Upregulated terms include cell-cell junction organization, axonogenesis, and canonical Wnt signaling.

### 3.4 Chromosomal DEG distribution

As shown in Figure 5 and Figure 6, the analysis of the chromosomal distribution of butyrate-dysregulated genes using 10 Mb windows showed nearly equal distribution across the human genome. There were slightly higher numbers of upregulated vs. downregulated genes in most chromosomes (Figure 5). No autosome showed strong whole-chromosome DEG enrichment after FDR correction. Using 10 Mb windows, only one window, chr15:40-50 Mb, showed significant DEG enrichment after FDR correction (Figure 6). Studies with smaller windows may possibly identify local DEG clusters. Chromosome Y had only 24 DEGs, hence, it is not included in the chromosomal distribution analysis.

**Figure 4.**
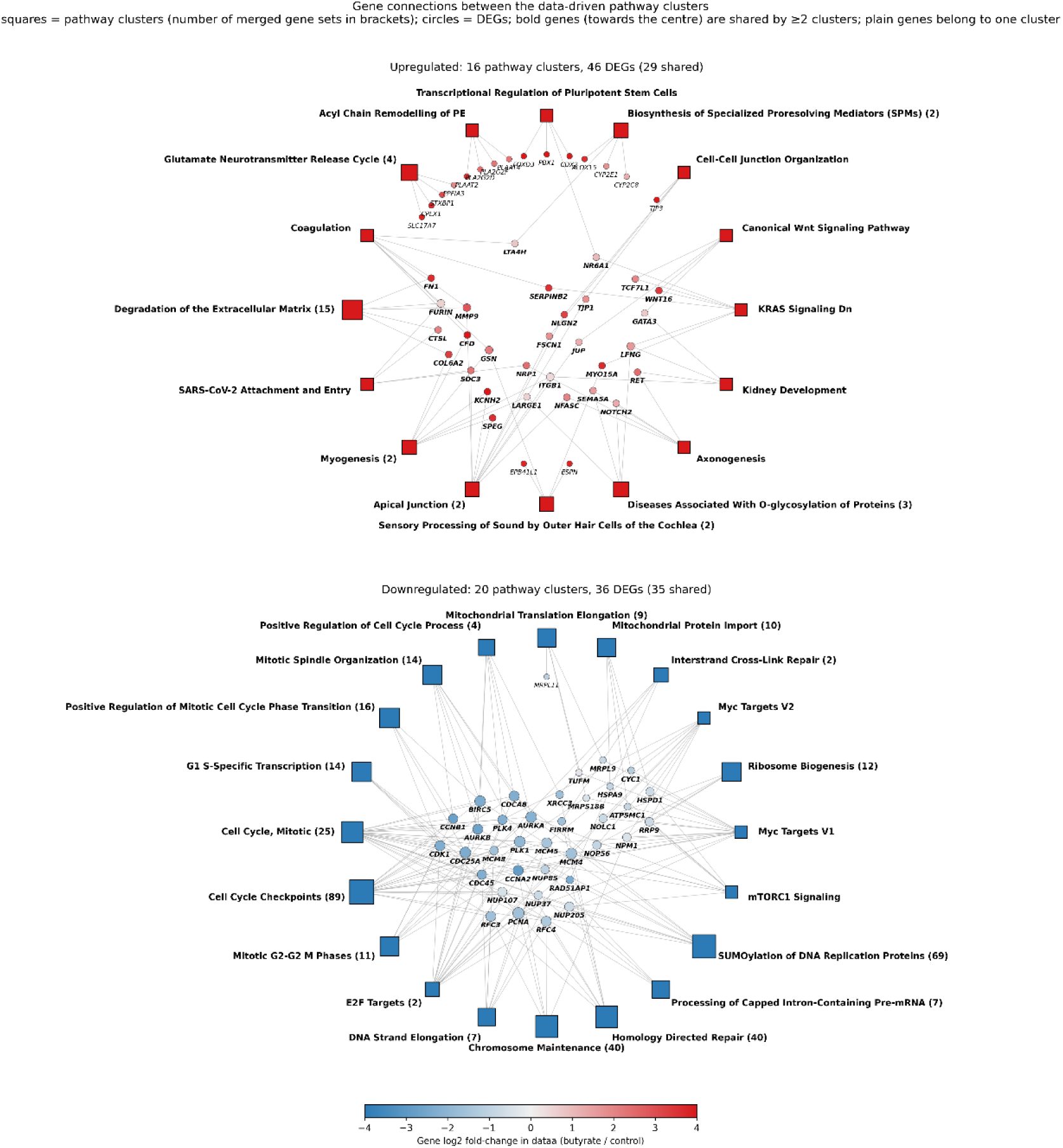
Gene connections between butyrate-regulated pathway clusters. Pathway clusters (squares; the number of merged gene sets is shown in brackets) are linked to the paired-analysis DEGs (circles) found in the leading edge of their strongest gene set. For each cluster, up to four DEGs are shown, preferring genes shared with other clusters and then genes with the largest fold change; circle color shows each gene’s log2 FD after butyrate treatment. Genes shown in bold connect two or more clusters.

**Figure 5.**
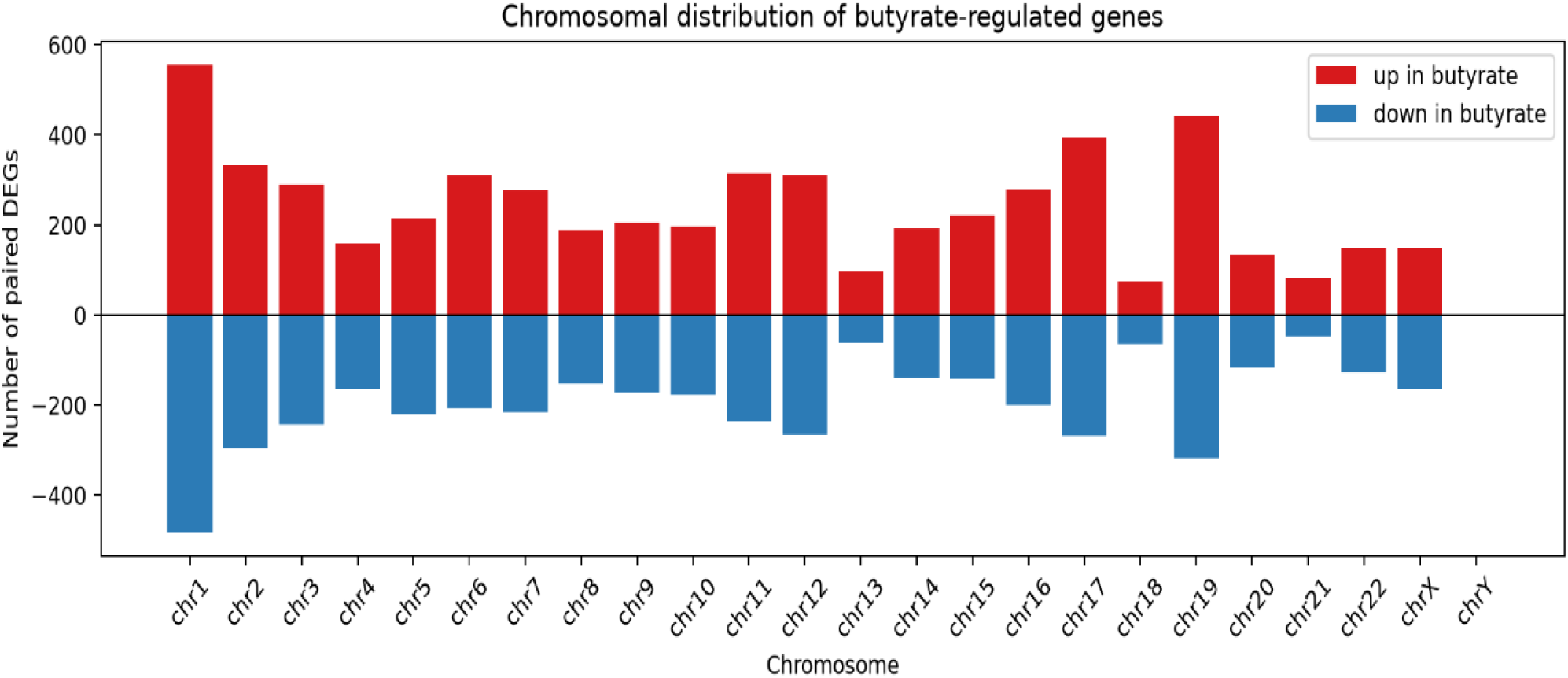
Chromosomal distribution of butyrate-regulated genes. Genes were assigned to chromosomal positions using GENCODE v50 annotation. Bar plots show the number of paired-analysis DEGs per chromosome, separated into upregulated and downregulated genes. This analysis is exploratory and should be interpreted relative to the background of expressed/tested genes on each chromosome.

**Figure 6.**
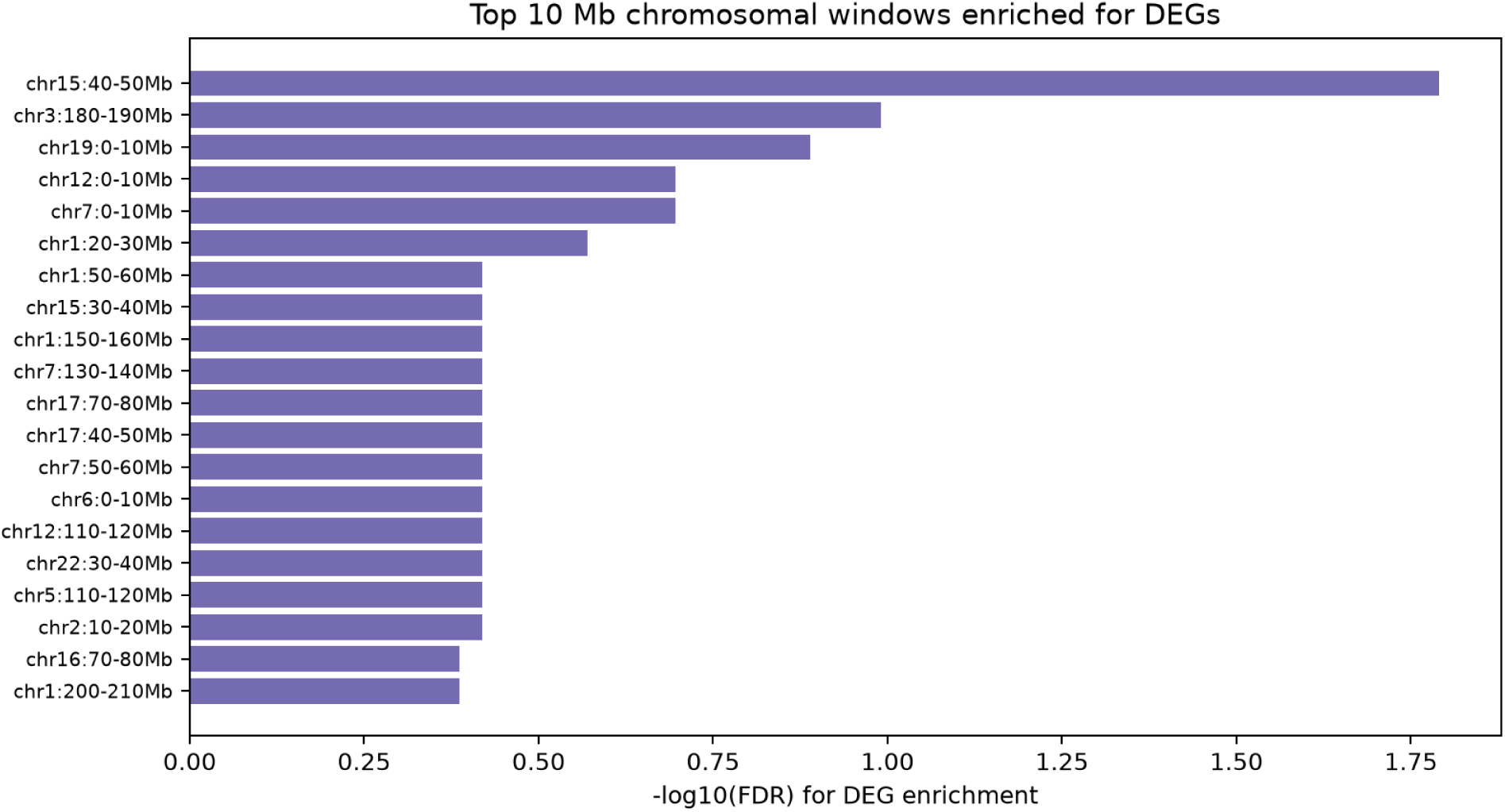
Top 10 Mb chromosomal windows enriched for DEGs. As shown, the distribution of DEGs was nearly even across the human genome, with a single DEG-enriched window at chr15:40-50 Mb.

### 3.5. Histone deacetylase (*HDAC*) transcription

The human genome includes 11 *HDAC* genes. RNA-seq paired analysis (N=8 LCLs) indicated that the transcription of five of these genes was dysregulated following butyrate exposure: *HDAC2*, *HDAC3*, *HDAC7, HDAC10*, and *HDAC11.* Of these five HDAC family genes, the largest effect of butyrate treatment was found for *HDAC11* (FD=4.75; padj=2.95E-32; mean reads=99). The second largest effect was for *HDAC3* (FD=1.58; padj=4.25E-11; mean reads=1663). *HDAC2, HDAC7*, and *HDAC10* showed relatively small upregulation or downregulation following butyrate treatment (see Supplementary Table S1). Further studies are required for elucidating the transcriptional effects of butyrate treatments in other human cell types.

## 4. Discussion

### 4.1 Advantages of paired bioinformatics analyses of RNA-seq data from treated vs. control human LCLs

This study applied RNA-seq of human LCLs from eight unrelated healthy donors, each sequenced both under control optimal growth conditions and immediately following treatment with 1 mM sodium butyrate for 48 h. The bioinformatics approach compared each butyrate-treated sample to its control LCL grown in parallel. This paired bioinformatics analysis allows extra power, owing to less noise arising from the genomic and epigenomic inter-individual differences between donors whose B lymphocytes were used for LCL generation. Paired analysis also minimizes confounders when compared with group analysis, such as passage number, growth medium batch, incubator temperature, etc. Hence, paired analysis yields extra statistical power compared with group analysis, as evident from the supplementary RNA-seq Excel file. This extra statistical power allows savings on RNA-seq costs thanks to smaller sample sizes.

Notable published examples for the advantages of paired analysis of RNA-seq data include breast cancer detection (Yaqoob et al. 2024), mapping the immune landscape of lung adenocarcinoma (Renaut et al. 2024), and gaining insights into the impact of HIV on alveolar macrophages (Staitieh et al. 2023).

The extra power of applying paired analysis for the effects of a given compound (or growth conditions) is exemplified by the detection of ∼10^4^ DEGs with fold-difference >1.2 or <0.8 following exposure to 1 mM butyrate (with padj <0.05. For comparison, the group analysis of the same eight butyrate-treated LCLs panel with their control LCL panel detected ∼25% fewer DEGs (7550 compared with 10054 for the paired and the group analyses, respectively).

The use of LCLs has several advantages for paired transcriptomic experiments. Cells from different donors can be cultured in parallel under similar conditions and periods, and repeated experiments can be easily performed. Because treated and untreated cultures are derived from the same donor, they share the same inherited genome and epigenome, and each LCL can serve as its own control.

A paired RNA-seq design is particularly advantageous when investigating treatment effects in cells derived from multiple individuals, while potential inter-individual differences can be examined separately. Although an RNA-seq study with LCLs from eight donors is not powered for DNA-seq studies, replication across eight unrelated genomes can help distinguish consistent from chance effects, as evident from our robust RNA-seq paired analysis findings (Table 1).

### 4.2 Pathway and pathway network analyses of butyrate-affected DEGs

The Reactome pathway analysis of the RNA-seq paired analysis DEGs (Figure 3) showed that sodium butyrate exposure of human LCLs (1 mM; 48 h) upregulated pathways of extracellular matrix, cell adhesion, junction organization, Wnt signaling, axonogenesis, and neurodevelopment pathways. Downregulated pathways included cell cycle, G2/M checkpoint, E2F/MYC targets, DNA replication and repair, mitotic spindle integrity, mTOR signaling, oxidative phosphorylation, unfolded-protein response, and translation/ribosome-related pathways.

The pathway network analyses of the same DEG findings (Figure 4) showed that the downregulated DEG clusters are tightly interconnected through a common core of cell-cycle and DNA-replication genes. For example, these genes included *PCNA, PLK1, AURKA, CDK1*, and *MCM4*, indicating a single coordinated proliferation program. By comparison, the upregulated DEG clusters are more loosely connected and represent more distinct responses, such as extracellular matrix remodeling (*MMP9, FN1, COL6A2*) and Wnt signaling (*WNT16, TCF7L1*).

Notably, the detected DEGs – and hence the butyrate-affected pathways and pathway networks – may be different in other cell types. Researchers are advised to keep in mind that our findings were obtained by RNA-seq of human LCLs, cell lines that were derived from immortalized B lymphocytes.

### 4.3 Butyrate-affected DEGs associated with CNS disorders

#### 4.3.1 Autism spectrum disorder (ASD)

Autism Spectrum Disorder (ASD) is the most common neurodevelopmental disorder. It has strong genetic components but is also affected by epigenetic factors. Among the latter, the gut microbiome was shown to differ between children with ASD and neurotypically developing (NT) children (Coretti et al. 2018), leading to the assumption that it is implicated in ASD. Indeed, transfer of gut microbiome from ASD individuals (but not from NT individuals) to germ-free mice was shown to induce ASD-like phenotypes, such as repetitive behaviour, in recipient mice (Sharon et al. 2019).

In 2022 we reported four DEGs from an RNA-seq study comparing whole blood samples from children diagnosed with ASD compared to blood samples from neurotypically-developing (NT) children and validated by RT-qPCR in 73 Israeli and American children with ASD compared to 26 NT children (Voinsky et al 2022). These findings were applied for building an ML tool as an aid for ASD diagnosis and demonstrated closely similar accuracy and specificity reported for behavioral scores-based ASD diagnostics, suggesting that they may supplement current protocols for ASD diagnosis (Voinsky et al 2023). Findings from the current RNA-seq study that explored butyrate-affected DEGs demonstrated that the expression levels of three of the four dysregulated genes reported in blood samples from ASD children (Voinsky et al. 2022) were affected in the opposite direction by in vitro butyrate exposure of LCLs (Fig. 2 and Table 1). These three genes included butyrate-mediated upregulation of *MT2A* and *ISG15* and downregulation of *LY6E*. In contrast, *BATF2*, a transcription factor whose expression was upregulated in blood samples of children with ASD, was also upregulated in butyrate-treated LCLs (Fig. 2). This finding does not necessarily mean that butyrate supports autism phenotypic symptoms. First, human LCLs are generated by immortalization of B lymphocytes, which are a minority (under 10%) of white blood cells (WBC) in whole blood, used for the RNA-seq study that detected higher *BATF2* mRNA in children with ASD (Voinsky et al. 2022). It could be that BATF2 transcription is differently dysregulated in specific WBC cell types of children with ASD. Additionally, the increased *BATF2* transcription observed in blood samples from ASD children may reflect an attempt of an epigenetic mechanism to ‘correct’ an autism phenotype associated with a dysregulated immune system (Estes et al. 2015), and this ‘corrective’ attempt is enhanced by butyrate. Notably, *BATF2* codes for basic leucine zipper ATF-like transcription factor 2, which modulates immune responses and inflammation (van der Geest et al. 2025). Its upregulation by butyrate also supports the anti-cancer capacities of butyrate (see section 4.3). Among the butyrate-upregulated genes whose expression was downregulated in ASD blood samples, *MT2A* codes for metallothionein 2A, a cysteine-rich metal-binding protein acting as an antioxidant and key suppressor of free radical damage (Frye et al. 2024). The butyrate-mediated upregulation of *MT2A* expression was recently demonstrated in LCLs from 17 unrelated donors in the context of the anti-diabetic effects of butyrate (Shapira et al. 2025)

#### 4.3.2 Neuroinflammation and neurodegenerative disorders

The RNA-seq analysis of the butyrate-associated DEGs included genes coding for proteins that reduce neuroinflammation. Reduced neuroinflammation was associated with the beneficial effects of butyrate observed in autism mouse models (Bouelkhair et al. 2026) and in Alzheimer’s disease mouse models (Brossaud et al. 2026). Neuroinflammation is among the hallmarks of Alzheimer’s disease (Heneka et al. 2025). The beneficial effects of butyrate were shown to reduce aggregated amyloid-beta accumulation in Alzheimer’s disease mouse models (Govindarajan et al. 2011; Sun et al. 2020; Fernando et al. 2020; Wang et al. 2022; Brossaud et al. 2026).

Neuroinflammation and synaptic loss were also implicated in the tau pathology observed in Alzheimer’s disease temporal cortex (Buchanan et al. 2020), while the effects of butyrate on tau pathology remain unclear. Additionally, sodium butyrate was shown to reduce neuroinflammation and tissue damage in spinal cord injury mouse and rat models (Cui et al. 2025; Ganggang et al. 2025). Sodium butyrate also reduced neuroinflammation and improved neurological dysfunction and post-traumatic epilepsy in a brain injury mouse model (Golub et al. 2026).

Progressive hearing loss is another CNS disorder where butyrate seems to be protective. Butyrate protected against hearing loss mediated in Guinea pig models by the chemotherapy cisplatin (Drottar et al. 2006), the antibiotic gentamycin (Wang et al. 2025), or by noise (Yang el al. 2017). A recent study showed that the *FMN1* gene coding for formin-1 is essential for correct cochlear microtubule architecture <u>x</u>required for hearing in both mice and humans (Kamal et al. 2026). Indeed, our RNA-seq showed that butyrate dramatically upregulated *FMN1* in human LCLs (FD=7.11; padj=7.11E-07; Supplementary Table 1).

### 4.4 DEGs associated with cancer growth and metastasis

Many of the DEGs in our RNA-seq findings belong to pathways affecting the growth and metastasis of cancer cells (Figure 3), as supported by studies of colorectal cancer (CRC). Among the genes robustly downregulated by butyrate is *HDGF* (heparin binding growth factor; Table 1), as validated by RT-qPCR (Figure 2). Notably, the progression of CRC and its drug resistance were suppressed in *HDGF* knockout mice (Su et al. 2025), a finding that may participate in the protective effect of butyrate against CRC progression (Kang et al. 2023).

Numerous studies described beneficial effects of butyrate or butyrate-producing gut microbiome in human cancer cells and rodent cancer models, mostly for colorectal cancer (Hassig et al. 1997; Wu et al. 2018; Karim et al. 2024). A notable RNA-seq study of two human colorectal cancer cell lines, HCT116 and SW480, reported that in vitro exposure to 2 mM sodium butyrate inhibited cell proliferation and induced apoptosis (Zhou et al, 2019). Notably, a recent study suggested that the active form of vitamin D, 1,25(OH)2D, increased the abundance of butyrate-producing gut bacteria, thereby reducing breast cancer metastasis (Gong et al. 2026). Clinical trials should clarify the utility of sodium butyrate or butyrate-producing gut microbiome taxa in cancer therapy.

### 4.5 Study Limitations

Along with the study advantages, it has limitations that should be addressed in future studies on the health benefits of gut microbiome-generated butyrate. An obvious limitation is that our study examined only RNA expression dysregulation following in vitro butyrate exposure. Future studies should include, when feasible, measurements of proteins generated from the translation of dysregulated gene transcripts. At present, the high costs of proteomics compared with RNA-seq transcriptomics are prohibitive, as evident from the extensive list of thousands of butyrate-mediated DEGs (supplementary Excel file) compared with hundreds of proteins that can typically be measured in biosamples by protein arrays or other proteomic technologies (Du et al. 2025; Hu et al. 2026). LCLs are derived from freshly isolated B lymphocytes using Epstein–Barr virus (EBV) transformation with a variant that does not release mature virions, thus resulting in cell immortalization (Gurwitz 2016). This immortalization method may affect cell proliferation, metabolism, mitochondrial respiration, transcription-factor activity, and immune signaling. Hence, for planning preclinical studies on the potential of butyrate (or other HDAC inhibitors), our detected DEGs and DEG pathways should ideally be examined using cell types (or tissues) relevant for the disease/disorder (e.g., in rodent disease models). For example, we suggest that future in vitro studies assessing the transcriptional effects of butyrate may benefit from using panels of iPSC-generated cell types of interest. Nevertheless, due to the wide availability of LCL collections, and the involvement of immune cells in many human diseases/disorders, LCLs remain a valuable tool for identifying transcriptional responses and for precision medicine studies (Chouchana et al. 2015; Gurwitz 2016). Interesetingly, butyrate-mediated DEGs coding for mitochondrial proteins were by far more likely to be downregulated by butyrate compared with non-mitochondrial proteins (Supplementary Table S2). This finding should be explored in other cell types and may give clues for treating mitochondrial disorders.

### 4.6 Clinical trials and forward look

Dietary supplementation with butyrate, or transfer of microbiome species known to generate butyrate from dietary starch, are likely to remain a hot research topic for basic and applied pharmacology. This is evident for the large number of relevant clinical trials listed on clinicaltrails.gov (for any disease or condition). As of October 2026, over 1800 clinical trials with “butyrate” as the treatment/intervention were listed in clinicaltrials.gov; the majority (>1,100) were concluded, 139 trials were recruiting, and 57 were “not yet recruiting”. The few published studies on butyrate supplementations were small (<50 patients), and no meta-analysis studies have been published. As an example for an encouraging clinical trial, a double-blind placebo-controlled Dutch trial in patients with joint osteoarthritis reported improvements in inflammatory blood markers following four to five weeks of 600 mg/kg per day sustained-release sodium butyrate supplementation (Korsten et al. 2024).

Clinical trials are required for stablishing the therapeutic value and possible side-effects of butyrate, butyrate analogues, or its precursor tributyrin, a triglyceride with three butyrate residues (Heidor et al. 2012). Knowledge is missing on the long-term metabolic, hormonal, and immunologic effects of butyrate-based therapeutics (Golea et al. 2026). Microencapsulated butyrate formulations are also being studied as drug candidates for further development and tentative clinical trials (Panufnik et al, 2026, Caban et al. 2026, Xie et al. 2026). The promising therapeutic potential of butyrate and butyrate-producing gut microbiome will require better knowledge for identifying the most suitable model for preclinical and clinical studies. Our current report on the genome-wide in vitro transcriptomic effects of butyrate on human B lymphocyte-derived cell lines is a small step in the long road toward this goal.

## Acknowledgements

The study was funded in part by a grant from The Center for AI & Data Science (TAD) at Tel Aviv University to DG, and a separate TAD grant to NS. The study was done as part of the requirements of HK for the M.Sc. degree at the Gray Faculty of Medical and Health Sciences Graduate School at Tel Aviv University. The authors are grateful for the individuals who donated blood samples for the generation of the lymphoblastoid cell lines, which made this study possible.

## Data Availability Statement

Table S1 (Supplementary file 1) is available as Excel file with additional data by request from Prof. Noam Shomron or Prof. David Gurwitz.

### Supplementary file 1

**Table S1:** RNA-seq Paired Analysis DEGs (butyrate-treated vs. control). The DEGs Excell file was printed to PDF and covers all DEGs with padj<0.05 (arranged by increasing padj values). The Excell spreadsheet with additional tabs and information (including average reads per sample) is available from Prof. Noam Shomron or Prof. David Gurwitz. Please consult the Methods section of the main text.

### Supplementary file 2

**Table S2:** DEGs encoding mitochondrial proteins according to Human MitoCarta3.0 (n = 645; 150 up, 495 down). See Table S1 for further information.

### Supplementary file 3

**Table S3:** List of primers for RT-qPCR of selected genes (see Table 1).

## References

Battle A, Mostafavi S, Zhu X, et al. Characterizing the genetic basis of transcriptome diversity through RNA sequencing of 922 individuals. Genome Res. 2014;24:14–24.

Boffa LC, Vidali G, Mann RS, Allfrey VG. Suppression of histone deacetylation in vivo and in vitro by sodium butyrate. J Biol Chem. 1978;253:3364–3366.

Bouelkhair M, Kacimi FE, Azzaoui FZ, Ramchoun M, Berrougui H, Boulbaroud S. Short-chain fatty acids, neuroinflammation, and autism spectrum disorders: A mechanistic systematic review. J Neuroimmunol. 2026 Jul;416:578914. doi: 10.1016/j.jneuroim.2026.578914.

Brossaud R, Oullier T, Bessard A, Aubert P, Brossard L, Mahé MM, Caillaud M, Delfino G, Paillusson S, Falentin H, Naveilhan P, Le-Loir Y, Paillé V, Neunlist M, Cissé M. The short-chain fatty acid butyrate prevents gut-brain amyloid-beta pathology and neuroinflammation in an Alzheimer mouse model. Mol Psychiatry. 2026 Jul;31(7):3910–3928. doi: 10.1038/s41380-026-03522-6.

Buchanan H, Mackay M, Palmer K, Tothová K, Katsur M, Platt B, Koss DJ. Synaptic Loss, ER Stress and Neuro-Inflammation Emerge Late in the Lateral Temporal Cortex and Associate with Progressive Tau Pathology in Alzheimer’s Disease. Mol Neurobiol. 2020 Aug;57(8):3258–3272. doi: 10.1007/s12035-020-01950-1.

Caban M, Pikus E, Czarnecka-Chrebelska K, Oszajca K, Witusik-Perkowska M, Dudek J, Szemraj J, Brzeziańska-Lasota E, Talar-Wojnarowska R. Use of Sodium Butyrate and Its Microencapsulated Forms in Intestinal Diseases-Current Clinical Approach. Dig Dis Sci. 2026 May;71(5):1626–1639. doi: 10.1007/s10620-025-09536-4.

Candido EPM, Reeves R, Davie JR. Sodium butyrate inhibits histone deacetylation in cultured cells. Cell. 1978;14:105–113.

Chen, S., Zhou, Y., Chen, Y., & Gu, J. (2018). fastp: An ultra-fast all-in-one FASTQ preprocessor. Bioinformatics, 34, i884–i890. 10.1093/bioinformatics/bty560

Chouchana L, Fernández-Ramos AA, Dumont F, Marchetti C, Ceballos-Picot I, Beaune P, Gurwitz D, Loriot MA. Molecular insight into thiopurine resistance: transcriptomic signature in lymphoblastoid cell lines. Genome Med. 2015 Apr 18;7(1):37. doi: 10.1186/s13073-015-0150-6.

Choy E, Yelensky R, Bonakdar S, et al. Genetic analysis of human traits in vitro: drug response and gene expression in lymphoblastoid cell lines. PLoS Genet. 2008;4:e1000287.

Coretti L, Paparo L, Riccio MP, Amato F, Cuomo M, Natale A, Borrelli L, Corrado G, Comegna M, Buommino E, Castaldo G, Bravaccio C, Chiariotti L, Berni Canani R, Lembo F. Gut Microbiota Features in Young Children With Autism Spectrum Disorders. Front Microbiol. 2018 Dec 19;9:3146. doi: 10.3389/fmicb.2018.03146. eCollection 2018.

Cui Y, Cen Q, Feng J, Wei J, Wang L, Chang C, Pang R, Wang J, Zhang A. Sodium butyrate alleviates spinal cord injury via inhibition of NLRP3/Caspase-1/GSDMD-mediated pyroptosis. Metab Brain Dis. 2025 Mar 24;40(4):157. doi: 10.1007/s11011-025-01589-8.

Davie JR. Inhibition of histone deacetylase activity by butyrate. J Nutr. 2003;133:2485S–2493S. Donohoe DR, Garge N, Zhang X, et al. The microbiome and butyrate regulate energy metabolism and autophagy in the mammalian colon. Cell Metab. 2011;13:517–526.

Dobin, A., Davis, C. A., Schlesinger, F., Drenkow, J., Zaleski, C., Jha, S., Batut, P., Chaisson, M., & Gingeras, T. R. (2013). STAR: Ultrafast universal RNA-seq aligner. Bioinformatics, 29, 15–21. 10.1093/bioinformatics/bts635

Donohoe DR, Collins LB, Wali A, Bigler R, Sun W, Bultman SJ. The Warburg effect dictates the mechanism of butyrate-mediated histone acetylation and cell proliferation. Mol Cell. 2012;48:612–626.

Drottar M, Liberman MC, Ratan RR, Roberson DW. The histone deacetylase inhibitor sodium butyrate protects against cisplatin-induced hearing loss in guinea pigs. Laryngoscope. 2006 Feb;116(2):292–6. doi: 10.1097/01.mlg.0000197630.85208.36.

Du M, Zeng F, Wang Y, Li Y, Chen G, Jiang J, Wang Q. Assembly and Functionality of 2D ProteinArrays. Adv Sci (Weinh). 2025 Apr;12(15):e2416485. doi: 10.1002/advs.202416485.

Estes ML, McAllister AK. Immune mediators in the brain and peripheral tissues in autism spectrum disorder. Nat Rev Neurosci. 2015 Aug;16(8):469–86. doi: 10.1038/nrn3978.PMID: 26189694

Ferreira JA. The Benjamini-Hochberg method in the case of discrete test statistics. Int J Biostat. 2007;3(1):Article 11. doi: 10.2202/1557-4679.1065.

Frye RE, Rose S, Voinsky I, Gurwitz D. Nitrosative Stress in Autism: Supportive Evidence and Implications for Mitochondrial Dysfunction. Adv Sci (Weinh). 2024 Apr;11(16):e2304439. doi: 10.1002/advs.202304439.

Ganggang K, Cheng G, Gang L, Wenwu Z, Di Z, Yan L, Baoshu X, Yiqin W. From Molecular Docking to Rat Models: Butyrate From a High-Fiber Diet Inhibits HDAC1 and NLRP3 Inflammasome to Alleviate Oxidative Stress and Inflammation After Spinal Cord Injury. J Neurochem. 2025 Oct;169(10):e70250. doi: 10.1111/jnc.70250.

Gates LA, Reis BS, Lund PJ, Paul MR, Leboeuf M, Djomo AM, Nadeem Z, Lopes M, Vitorino FN, Unlu G, Carroll TS, Birsoy K, Garcia BA, Mucida D, Allis CD. Histone butyrylation in the mouse intestine is mediated by the microbiota and associated with regulation of gene expression. Nat Metab. 2024 Apr;6(4):697–707. doi: 10.1038/s42255-024-00992-2.

Golea GI, Ilieș RA, Caziuc A, Andraș D, Ilie-Ene A, Seicean R, Neagoe RM, Dăscălescu Ș, Simionescu L, Vesa ȘC, Dindelegan GC, Zaharie F. Sodium Butyrate Therapy in Ulcerative Colitis: A Systematic Review of Preclinical and Clinical Evidence. Med Sci (Basel). 2026 Aug 9;14(4):469. doi: 10.3390/medsci140

Golub VM, Ramakrishnan S, Reddy DS. Epigenetic histone deacetylase inhibition by sodium butyrate reduces neuroinflammation, improves neurological dysfunction and promotes disease modification of epileptogenesis following traumatic brain injury. Exp Neurol. 2026 Oct;404:115857. doi: 10.1016/j.expneurol.2026.115857.

Gong F, Liu N, Miao J, Wang C, Lin M, Luan B, Chen J, Chen Y, Chen X, Ge F, Chen W. Vitamin D regulation of gut microbiota-derived butyrate as a potential inhibitor of breast cancer proliferation. Gene. 2026 Jan 15;976:149872. doi: 10.1016/j.gene.2025.149872.

Govindarajan N, Agis-Balboa RC, Walter J, Sananbenesi F, Fischer A. Sodium butyrate improves memory function in an Alzheimer’s disease mouse model when administered at an advanced stage of disease progression. J Alzheimers Dis. 2011;26(1):187–97. doi: 10.3233/JAD-2011-110080.

Gurwitz D. Human iPSC-derived neurons and lymphoblastoid cells for personalized medicine research in neuropsychiatric disorders. Dialogues Clin Neurosci. 2016 Sep;18(3):267–276. doi: 10.31887/DCNS.2016.18.3/dgurwitz.

Hassig CA, Tong JK, Schreiber SL. Fiber-derived butyrate and the prevention of colon cancer. Chem Biol. 1997 Nov;4(11):783–9. doi: 10.1016/s1074-5521(97)90111-3.

Heidor R, Ortega JF, de Conti A, Ong TP, Moreno FS. Anticarcinogenic actions of tributyrin, a butyric acid prodrug. Curr Drug Targets. 2012 Dec;13(14):1720–9. doi: 10.2174/138945012804545443.

Hu R, Montes C, Walley JW. Mass spectrometry-based single-cell proteomics technologies, trends, and biological insights. Trends Biochem Sci. 2026 Jun;51(6):542–557. doi: 10.1016/j.tibs.2026.04.011.

Jiang S, Willox B, Zhou H, Holthaus AM, Wang A, Shi TT, et al. The Epstein–Barr virus regulome in lymphoblastoid cells. Cell Host Microbe. 2017;22:561–573.e4.

Heneka, et al. Neuroinflammation in Alzheimer disease. Nat Rev Immunol. 2025 May;25(5):321–352. doi: 10.1038/s41577-024-01104-7.

Kamal L, Aburayyan A, Hahn R, Taiber S, Gulsuner S, Kanaan MN, King MC, Avraham KB. Formin-1 maintains cochlear microtubule architecture required for hearing in humans and mice Proc Natl Acad Sci U S A. 2026 Sep 15;123(37):e2622920123. doi: 10.1073/pnas.2622920123.

Kang J, Sun M, Chang Y, Chen H, Zhang J, Liang X, Xiao T. Butyrate ameliorates colorectal cancer through regulating intestinal microecological disorders. Anticancer Drugs. 2023 Feb 1;34(2):227–237. doi: 10.1097/CAD.0000000000001413.

Karim MR, Iqbal S, Mohammad S, Morshed MN, Haque MA, Mathiyalagan R, Yang DC, Kim YJ, Song JH, Yang DU. Butyrate’s (a short-chain fatty acid) microbial synthesis, absorption, and preventive roles against colorectal and lung cancer. Arch Microbiol. 2024 Mar 4;206(4):137. doi: 10.1007/s00203-024-03834-7.

Koh A, De Vadder F, Kovatcheva-Datchary P, Bäckhed F. From dietary fiber to host physiology: short-chain fatty acids as key bacterial metabolites. Cell. 2016;165:1332–1345.

Korsten SGPJ, Hartog M, Berends AJ, Koenders MI, Popa CD, Vromans H, Garssen J, van de Ende CHM, Vermeiden JPW, Willemsen LEM. A Sustained-Release Butyrate Tablet Suppresses Ex Vivo T Helper Cell Activation of Osteoarthritis Patients in a Double-Blind Placebo-Controlled Randomized Trial. Nutrients. 2024 Oct 4;16(19):3384. doi: 10.3390/nu16193384.

Li S, Li X, Jiang S, Wang C, Hu Y. Identification of sepsis-associated mitochondrial genes through RNA and single-cell sequencing approaches. BMC Med Genomics. 2024 May 3;17(1):120. doi: 10.1186/s12920-024-01891-x.

Liberzon A, Birger C, Thorvaldsdóttir H, Ghandi M, Mesirov JP, Tamayo P. The Molecular Signatures Database (MSigDB) hallmark gene set collection. Cell Syst. 2015 Dec 23;1(6):417–425. doi: 10.1016/j.cels.2015.12.004.

Liu J, Wu Y, Li Z, Tang J, Zhou X, Luo S. Gut microbiota-derived butyrate primes systemic immunity in honey bees by mediating lipid metabolic reprogramming. Nat Commun. 2026 Feb 2;17(1):2924. doi: 10.1038/s41467-026-69073-0.

Louis P, Flint HJ. Formation of propionate and butyrate by the human colonic microbiota. Environ Microbiol. 2017;19:29–41.

Love MI, Huber W, Anders S. (2014). Moderated estimation of fold change and dispersion for RNA-seq data with DESeq. 2. Genome Boilogy, 15, 550. doi.org/10.1186/s13059-014-0550-8

Montgomery SB, Sammeth M, Gutierrez-Arcelus M, et al. Transcriptome genetics using second-generation sequencing in a Caucasian population. Nature. 2010;464:773–777.

Panufnik P, Więcek M, Szwarc P, Szydlarska D, Kadaj-Lipka R, Franek E, Rydzewska G. Effect of microencapsulated sodium butyrate on abdominal symptoms and carbohydrate metabolism in patients with type 2 diabetes: a randomized placebo-controlled trial. Sci Rep. 2026 Apr 13;16(1):17404. doi: 10.1038/s41598-026-47309-9.

Pickrell JK, Marioni JC, Pai AA, et al. Understanding mechanisms underlying human gene-expression variation with RNA sequencing. Nature. 2010;464:768–772.

Renaut S, Saavedra Armero V, Boudreau DK, Gaudreault N, Desmeules P, Thériault S, Mathieu P, Joubert P, Bossé Y. Single-cell and single-nucleus RNA-sequencing from paired normal-adenocarcinoma lung samples provide both common and discordant biological insights. PLoS Genet. 2024 May 30;20(5):e1011301. doi: 10.1371/journal.pgen.1011301.

Shapira E, Voinsky I, Klin H, Gurwitz D. Butyrate-Mediated Upregulation of Insulin Pathway Gene Expression Suggests Potential Antidiabetic Effects. Drug Dev Res. 2025 Dec;86(8):e70203. doi: 10.1002/ddr.70203.

Sharon G, Cruz NJ, Kang DW, Gandal MJ, Wang B, Kim YM, Zink EM, Casey CP, Taylor BC, Lane CJ, Bramer LM, Isern NG, Hoyt DW, Noecker C, Sweredoski MJ, Moradian A, Borenstein E, Jansson JK, Knight R, Metz TO, Lois C, Geschwind DH, Krajmalnik-Brown R, Mazmanian SK. Human Gut Microbiota from Autism Spectrum Disorder Promote Behavioral Symptoms in Mice. Cell. 2019 May 30;177(6):1600–1618.e17. doi: 10.1016/j.cell.2019.05.004.

St Laurent R, O’Brien LM, Ahmad ST. Sodium butyrate improves locomotor impairment and early mortality in a rotenone-induced Drosophila model of Parkinson’s disease. Neuroscience. 2013 Aug 29;246:382–90. doi: 10.1016/j.neuroscience.2013.04.037.

Staitieh BS, Hu X, Yeligar SM, Auld SC. Paired ATAC- and RNA-seq offer insight into the impact of HIV on alveolar macrophages: a pilot study. Sci Rep. 2023 Sep 15;13(1):15276. doi: 10.1038/s41598-023-42644-7.

Steliou K, Boosalis MS, Perrine SP, Sangerman J, Faller DV. Butyrate histone deacetylase inhibitors. Biores Open Access. 2012;1:192–198.

Stranger BE, Nica AC, Forrest MS, et al. Population genomics of human gene expression. Nat Genet. 2007;39:1217–1224.

Su R, Wang Q, Hu Q, Wendurige, Li K, Wang C, Tao L. HDGF Knockout Suppresses Colorectal Cancer Progression and Drug Resistance by Modulating the DNA Damage Response. Biomolecules. 2025 Feb 14;15(2):282. doi: 10.3390/biom15020282.

Terra Machado D, Bernardes Brustolini OJ, Côrtes Martins Y, Grivet Mattoso Maia MA, Ribeiro de Vasconcelos AT. Inference of differentially expressed genes using generalized linear mixed models in a pairwise fashion. PeerJ. 2023 Apr 3;11:e15145. doi: 10.7717/peerj.15145.

van der Geest R, Lee JS. Role of the basic leucine zipper transcription factor BATF2 in modulating immune responses and inflammation in health and disease. J Leukoc Biol. 2025 Mar 14;117(3):qiae245. doi: 10.1093/jleuko/qiae245.PMID: 39504573

Voinsky I, Zoabi Y, Shomron N, Harel M, Cassuto H, Tam J, Rose S, Scheck AC, Karim MA, Frye RE, Aran A, Gurwitz D. Blood RNA Sequencing Indicates Upregulated BATF2 and LY6E and Downregulated ISG15 and MT2A Expression in Children with Autism Spectrum Disorder. Int J Mol Sci. 2022 Aug 30;23(17):9843. doi: 10.3390/ijms23179843.

Voinsky I, Fridland OY, Aran A, Frye RE, Gurwitz D. Machine Learning-Based Blood RNA Signature for Diagnosis of Autism Spectrum Disorder. Int J Mol Sci. 2023 Jan 20;24(3):2082. doi: 10.3390/ijms24032082.

Voinsky I, Goldenberg-Bogner O, Israel-Elgali I, Volkov H, Puzianowska-Kuźnicka M, Shomron N, Gurwitz D. RNA sequencing comparing centenarian and middle-aged women lymphoblastoid cell lines identifies age-related dysregulated expression of genes encoding selenoproteins, heat shock proteins, CD99, and BID. Drug Dev Res. 2024 Nov;85(7):e70011. doi: 10.1002/ddr.70011.

Wang C, Zheng D, Weng F, Jin Y, He L. Sodium butyrate ameliorates the cognitive impairment of Alzheimer’s disease by regulating the metabolism of astrocytes. Psychopharmacology (Berl). 2022 Jan;239(1):215–227. doi: 10.1007/s00213-021-06025-0.

Wang J, Wang Y, Chen X, Zhang PZ, Shi ZT, Wen LT, Qiu JH, Chen FQ. Histone deacetylase inhibitor sodium butyrate attenuates gentamicin-induced hearing loss in vivo. Am J Otolaryngol. 2015 Mar-Apr;36(2):242–8. doi: 10.1016/j.amjoto.2014.11.003.

Wu X, Wu Y, He L, Wu L, Wang X, Liu Z. Effects of the intestinal microbial metabolite butyrate on the development of colorectal cancer. J Cancer. 2018 Jun 15;9(14):2510–2517. doi: 10.7150/jca.25324. eCollection 2018.

Xie G, Fu Z, Zhang Y, Xu Z, Xue K, Dai H, Wu F, Lin S, Liu J. Oral delivery of enterocyte-mimetic butyrate-loaded nanoparticles repairs the gut barrier and inhibits growth of digestive system tumors. Nat Commun. 2026 May 9;17(1):6288. doi: 10.1038/s41467-026-72843-5.

Yaqoob A, Verma NK, Aziz RM, Shah MA. RNA-Seq analysis for breast cancer detection: a study on paired tissue samples using hybrid optimization and deep learning techniques. J Cancer Res Clin Oncol. 2024 Oct 10;150(10):455. doi: 10.1007/s00432-024-05968-z.

Yang DH, Xie J, Liu K, Peng Z, Guo JY, Yu SK, Wang GP, Gong SS. The histone deacetylase inhibitor sodium butyrate protects against noise-induced hearing loss in Guinea pigs. Neurosci Lett. 2017 Nov 1;660:140–146. doi: 10.1016/j.neulet.2017.09.036.

Zhang F, Zhang W, Ren X, Liu B, Zhou X. Butyrate-Producing Bacteria in Intestinal Disease Therapy: Potential and Challenges. Biotechnol J. 2026 Jun;21(6):e70260. doi: 10.1002/biot.70260.

Zhou Q, Li G, Zuo S, Zhu W, Yuan X. RNA Sequencing Analysis of Molecular Basis of Sodium Butyrate-Induced Growth Inhibition on Colorectal Cancer Cell Lines. Biomed Res Int. 2019 Feb 27;2019:1427871. doi: 10.1155/2019/1427871.

Zhou F, Liu B, Liu X, Li Y, Wang L, Huang J, Luo G, Wang X. The Impact of Microbiome and Microbiota-Derived Sodium Butyrate on Drosophila Transcriptome and Metabolome Revealed by Multi-Omics Analysis. Metabolites. 2021 May 6;11(5):298. doi: 10.3390/metabo11050298.

