## Supplemental Table 1 for "RNA-sequencing of butyrate-treated human lymphoblastoid cell lines shows dysregulated genes of multiple disease-related pathways"

| Ensembl_gene_id | Gene_symbol | NCBI_Gene_ID | Gene_type | mean_expression_baseMean | log2_FD_Butyrate_vs_Control | log10_FD_Butyrate_vs_Control | FD_linear_Butyrate_over_Control | pvalue | padj |
|---|---|---|---|---|---|---|---|---|---|
| ENSG00000138092.13 | **CENPO** | 79172 | protein_coding | **317.317** | -2.458 | -0.740 | **0.182** | 4.39E-107 | **8.35E-103** |
| ENSG00000124593.16 | **ENSG00000124593** | | protein_coding | **494.117** | 3.247 | 0.977 | **9.493** | 1.57E-106 | **1.49E-102** |
| ENSG00000169093.18 | **ASMTL** | 8623 | protein_coding | **2951.979** | 2.272 | 0.684 | **4.830** | 7.44E-100 | **4.72E-96** |
| ENSG00000072071.19 | **ADGRL1** | 22859 | protein_coding | **521.290** | 5.531 | 1.665 | **46.230** | 1.26E-99 | **6.01E-96** |
| ENSG00000185686.18 | **PRAME** | 23532 | protein_coding | **272.814** | 6.455 | 1.943 | **87.703** | 1.77E-97 | **6.75E-94** |
| ENSG00000157456.10 | **CCNB2** | 9133 | protein_coding | **613.842** | -2.575 | -0.775 | **0.168** | 5.52E-88 | **1.75E-84** |
| ENSG00000198830.15 | **HMGN2** | 3151 | protein_coding | **4965.710** | -1.882 | -0.567 | **0.271** | 6.54E-86 | **1.78E-82** |
| ENSG00000122952.19 | **ZWINT** | 11130 | protein_coding | **927.264** | -2.709 | -0.816 | **0.153** | 1.25E-85 | **2.96E-82** |
| ENSG00000155367.18 | **PPM1J** | 333926 | protein_coding | **220.638** | 3.446 | 1.037 | **10.896** | 1.99E-85 | **4.21E-82** |
| ENSG00000136824.21 | **SMC2** | 10592 | protein_coding | **434.985** | -2.140 | -0.644 | **0.227** | 6.10E-84 | **1.16E-80** |
| ENSG00000136938.11 | **ANP32B** | 10541 | protein_coding | **2646.116** | -1.546 | -0.465 | **0.342** | 1.29E-80 | **2.22E-77** |
| ENSG00000088325.18 | **TPX2** | 22974 | protein_coding | **1120.710** | -2.168 | -0.653 | **0.223** | 3.74E-80 | **5.93E-77** |
| ENSG00000154839.12 | **SKA1** | 220134 | protein_coding | **285.577** | -2.889 | -0.870 | **0.135** | 1.24E-79 | **1.82E-76** |
| ENSG00000146918.22 | **NCAPG2** | 54892 | protein_coding | **424.307** | -2.104 | -0.633 | **0.233** | 6.81E-79 | **9.25E-76** |
| ENSG00000145386.13 | **CCNA2** | 890 | protein_coding | **373.154** | -2.839 | -0.855 | **0.140** | 4.82E-76 | **6.12E-73** |
| ENSG00000110844.16 | **PRPF40B** | 25766 | protein_coding | **692.581** | 1.779 | 0.535 | **3.431** | 1.31E-73 | **1.56E-70** |
| ENSG00000169992.11 | **NLGN2** | 57555 | protein_coding | **329.565** | 3.203 | 0.964 | **9.211** | 7.83E-71 | **8.76E-68** |
| ENSG00000293178.2 | **ENSG00000293178** | | lncRNA | **271.714** | 2.033 | 0.612 | **4.093** | 4.72E-69 | **4.99E-66** |
| ENSG00000149554.17 | **CHEK1** | 1111 | protein_coding | **306.271** | -2.149 | -0.647 | **0.225** | 1.02E-68 | **1.02E-65** |
| ENSG00000137804.16 | **NUSAP1** | 51203 | protein_coding | **1440.755** | -1.539 | -0.463 | **0.344** | 1.69E-68 | **1.61E-65** |
| ENSG00000021300.15 | **PLEKHB1** | 58473 | protein_coding | **243.023** | 3.634 | 1.094 | **12.415** | 2.40E-68 | **2.17E-65** |
| ENSG00000189403.17 | **HMGB1** | 3146 | protein_coding | **4590.175** | -1.542 | -0.464 | **0.343** | 6.72E-67 | **5.59E-64** |
| ENSG00000137331.14 | **IER3** | 8870 | protein_coding | **2114.917** | 1.749 | 0.526 | **3.360** | 6.77E-67 | **5.59E-64** |
| ENSG00000144554.15 | **FANCD2** | 2177 | protein_coding | **818.154** | -2.106 | -0.634 | **0.232** | 1.26E-66 | **1.00E-63** |
| ENSG00000152192.8 | **POU4F1** | 5457 | protein_coding | **116.833** | 3.568 | 1.074 | **11.857** | 1.36E-66 | **1.03E-63** |
| ENSG00000177469.15 | **CAVIN1** | 284119 | protein_coding | **311.757** | 2.582 | 0.777 | **5.987** | 1.66E-66 | **1.22E-63** |

| | | | | | | | | | |
|---|---|---|---|---|---|---|---|---|---|
| ENSG00000080986.15 | **NDC80** | 10403 | protein_coding | **498.705** | -2.241 | -0.675 | **0.212** | 2.32E-65 | **1.64E-62** |
| ENSG00000170312.19 | **CDK1** | 983 | protein_coding | **621.009** | -2.307 | -0.694 | **0.202** | 4.65E-64 | **3.16E-61** |
| ENSG00000108387.17 | **SEPTIN4** | 5414 | protein_coding | **121.916** | 4.962 | 1.494 | **31.162** | 5.16E-64 | **3.38E-61** |
| ENSG00000108821.16 | **COL1A1** | 1277 | protein_coding | **584.564** | 3.089 | 0.930 | **8.511** | 4.37E-63 | **2.77E-60** |
| ENSG00000153044.12 | **CENPH** | 64946 | protein_coding | **403.056** | -2.137 | -0.643 | **0.227** | 5.41E-63 | **3.32E-60** |
| ENSG00000007516.17 | **BAIAP3** | 8938 | protein_coding | **363.904** | 3.600 | 1.084 | **12.129** | 5.75E-63 | **3.42E-60** |
| ENSG00000164045.14 | **CDC25A** | 993 | protein_coding | **263.299** | -2.315 | -0.697 | **0.201** | 6.00E-63 | **3.46E-60** |
| ENSG00000062524.18 | **LTK** | 4058 | protein_coding | **320.233** | 4.107 | 1.236 | **17.234** | 7.63E-63 | **4.27E-60** |
| ENSG00000186185.16 | **KIF18B** | 146909 | protein_coding | **447.144** | -2.322 | -0.699 | **0.200** | 1.65E-62 | **8.97E-60** |
| ENSG00000134690.13 | **CDCA8** | 55143 | protein_coding | **579.312** | -2.252 | -0.678 | **0.210** | 2.04E-62 | **1.08E-59** |
| ENSG00000133328.6 | **PLAAT2** | 54979 | protein_coding | **292.166** | 3.559 | 1.071 | **11.784** | 4.46E-62 | **2.29E-59** |
| ENSG00000148180.24 | **GSN** | 2934 | protein_coding | **824.684** | 2.154 | 0.648 | **4.450** | 1.29E-61 | **6.46E-59** |
| ENSG00000168393.15 | **DTYMK** | 1841 | protein_coding | **446.428** | -1.617 | -0.487 | **0.326** | 2.55E-61 | **1.24E-58** |
| ENSG00000243926.4 | **TIPARP-AS** | 730091 | lncRNA | **373.895** | 2.028 | 0.611 | **4.079** | 8.38E-61 | **3.98E-58** |
| ENSG00000126787.15 | **DLGAP5** | 9787 | protein_coding | **381.683** | -2.198 | -0.662 | **0.218** | 1.15E-60 | **5.33E-58** |
| ENSG00000109689.21 | **STIM2** | 57620 | protein_coding | **630.812** | 1.763 | 0.531 | **3.394** | 1.72E-60 | **7.80E-58** |
| ENSG00000149503.16 | **INCENP** | 3619 | protein_coding | **511.495** | -1.632 | -0.491 | **0.323** | 1.88E-60 | **8.33E-58** |
| ENSG00000293413.2 | **ENSG00000293413** | | lncRNA | **523.226** | 2.437 | 0.733 | **5.413** | 2.43E-60 | **1.05E-57** |
| ENSG00000064787.16 | **BCAS1** | 8537 | protein_coding | **1244.108** | 2.826 | 0.851 | **7.091** | 3.64E-60 | **1.54E-57** |
| ENSG00000091536.20 | **MYO15A** | 51168 | protein_coding | **233.023** | 3.647 | 1.098 | **12.531** | 5.09E-60 | **2.10E-57** |
| ENSG00000164109.16 | **MAD2L1** | 4085 | protein_coding | **264.411** | -2.384 | -0.718 | **0.192** | 7.63E-59 | **3.09E-56** |
| ENSG00000273066.5 | **ENSG00000273066** | | lncRNA | **164.594** | 2.941 | 0.885 | **7.681** | 1.85E-58 | **7.34E-56** |
| ENSG00000130304.19 | **SLC27A1** | 376497 | protein_coding | **344.651** | 2.287 | 0.689 | **4.882** | 2.01E-58 | **7.81E-56** |
| ENSG00000143321.19 | **HDGF** | 3068 | protein_coding | **2291.259** | -1.621 | -0.488 | **0.325** | 1.24E-57 | **4.71E-55** |
| ENSG00000008323.18 | **PLEKHG6** | 55200 | protein_coding | **239.510** | 3.192 | 0.961 | **9.140** | 1.48E-57 | **5.54E-55** |
| ENSG00000198826.13 | **ARHGAP11** | 9824 | protein_coding | **329.779** | -2.185 | -0.658 | **0.220** | 4.20E-57 | **1.54E-54** |
| ENSG00000116661.13 | **FBXO2** | 26232 | protein_coding | **167.166** | 3.672 | 1.105 | **12.743** | 5.02E-57 | **1.80E-54** |
| ENSG00000168078.12 | **PBK** | 55872 | protein_coding | **167.175** | -2.904 | -0.874 | **0.134** | 5.85E-57 | **2.06E-54** |
| ENSG00000293491.2 | **TMEM198** | 440104 | lncRNA | **1511.353** | 1.559 | 0.469 | **2.947** | 3.47E-56 | **1.20E-53** |
| ENSG00000176890.18 | **TYMS** | 7298 | protein_coding | **1573.321** | -2.551 | -0.768 | **0.171** | 8.07E-56 | **2.74E-53** |

| | | | | | | | | | |
|---|---|---|---|---|---|---|---|---|---|
| ENSG00000163507.17 | **CIP2A** | 57650 | protein_coding | **151.822** | -2.440 | -0.735 | **0.184** | 8.80E-56 | **2.94E-53** |
| ENSG00000165304.10 | **MELK** | 9833 | protein_coding | **544.441** | -1.738 | -0.523 | **0.300** | 9.49E-56 | **3.11E-53** |
| ENSG00000142731.13 | **PLK4** | 10733 | protein_coding | **228.784** | -2.213 | -0.666 | **0.216** | 1.05E-55 | **3.39E-53** |
| ENSG00000146670.11 | **CDCA5** | 113130 | protein_coding | **572.886** | -2.363 | -0.711 | **0.194** | 1.32E-55 | **4.20E-53** |
| ENSG00000169884.15 | **WNT10B** | 7480 | protein_coding | **239.747** | 3.753 | 1.130 | **13.483** | 1.52E-55 | **4.74E-53** |
| ENSG00000094916.18 | **CBX5** | 23468 | protein_coding | **2846.090** | -1.762 | -0.530 | **0.295** | 2.12E-55 | **6.51E-53** |
| ENSG00000230628.1 | **ENSG00000230628** | | lncRNA | **209.683** | 3.340 | 1.005 | **10.126** | 1.23E-54 | **3.70E-52** |
| ENSG00000138778.16 | **CENPE** | 1062 | protein_coding | **338.205** | -2.695 | -0.811 | **0.154** | 1.61E-54 | **4.79E-52** |
| ENSG00000173207.15 | **CKS1B** | 1163 | protein_coding | **709.570** | -1.806 | -0.544 | **0.286** | 2.08E-54 | **6.07E-52** |
| ENSG00000111206.15 | **FOXM1** | 2305 | protein_coding | **738.962** | -2.520 | -0.758 | **0.174** | 2.35E-54 | **6.77E-52** |
| ENSG00000111602.14 | **TIMELESS** | 8914 | protein_coding | **796.459** | -1.514 | -0.456 | **0.350** | 3.46E-54 | **9.83E-52** |
| ENSG00000138160.9 | **KIF11** | 3832 | protein_coding | **535.314** | -1.851 | -0.557 | **0.277** | 9.06E-54 | **2.53E-51** |
| ENSG00000121753.13 | **ADGRB2** | 576 | protein_coding | **129.038** | 4.252 | 1.280 | **19.059** | 6.24E-53 | **1.72E-50** |
| ENSG00000085840.15 | **ORC1** | 4998 | protein_coding | **212.536** | -1.951 | -0.587 | **0.259** | 9.18E-53 | **2.50E-50** |
| ENSG00000087586.20 | **AURKA** | 6790 | protein_coding | **555.914** | -2.097 | -0.631 | **0.234** | 1.01E-52 | **2.72E-50** |
| ENSG00000109685.21 | **NSD2** | 7468 | protein_coding | **2256.833** | -1.371 | -0.413 | **0.387** | 1.16E-52 | **3.05E-50** |
| ENSG00000119403.17 | **PHF19** | 26147 | protein_coding | **1302.011** | -1.832 | -0.552 | **0.281** | 2.01E-52 | **5.23E-50** |
| ENSG00000148773.16 | **MKI67** | 4288 | protein_coding | **2791.728** | -2.768 | -0.833 | **0.147** | 2.51E-52 | **6.45E-50** |
| ENSG00000103995.15 | **CEP152** | 22995 | protein_coding | **321.694** | -1.677 | -0.505 | **0.313** | 2.98E-52 | **7.55E-50** |
| ENSG00000164104.14 | **HMGB2** | 3148 | protein_coding | **3657.104** | -2.211 | -0.666 | **0.216** | 8.03E-51 | **2.01E-48** |
| ENSG00000090889.14 | **KIF4A** | 24137 | protein_coding | **224.309** | -2.057 | -0.619 | **0.240** | 1.22E-50 | **3.02E-48** |
| ENSG00000155115.9 | **GTF3C6** | 112495 | protein_coding | **544.374** | -1.712 | -0.515 | **0.305** | 1.36E-50 | **3.31E-48** |
| ENSG00000171320.16 | **ESCO2** | 157570 | protein_coding | **111.506** | -2.719 | -0.818 | **0.152** | 4.79E-50 | **1.15E-47** |
| ENSG00000089685.17 | **BIRC5** | 332 | protein_coding | **625.081** | -2.320 | -0.698 | **0.200** | 5.09E-50 | **1.21E-47** |
| ENSG00000079337.18 | **RAPGEF3** | 10411 | protein_coding | **154.220** | 2.462 | 0.741 | **5.511** | 1.10E-49 | **2.57E-47** |
| ENSG00000134057.17 | **CCNB1** | 891 | protein_coding | **807.336** | -2.701 | -0.813 | **0.154** | 1.73E-49 | **4.02E-47** |
| ENSG00000183856.13 | **IQGAP3** | 128239 | protein_coding | **375.687** | -2.859 | -0.861 | **0.138** | 2.04E-49 | **4.68E-47** |
| ENSG00000101945.19 | **SUV39H1** | 6839 | protein_coding | **286.246** | -2.330 | -0.701 | **0.199** | 2.55E-49 | **5.78E-47** |
| ENSG00000196550.13 | **FAM72A** | 729533 | protein_coding | **158.505** | -2.441 | -0.735 | **0.184** | 3.21E-49 | **7.19E-47** |
| ENSG00000112742.12 | **TTK** | 7272 | protein_coding | **203.850** | -2.365 | -0.712 | **0.194** | 3.92E-49 | **8.67E-47** |

| | | | | | | | | | |
|---|---|---|---|---|---|---|---|---|---|
| ENSG00000109046.17 | **WSB1** | 26118 | protein_coding | **2410.521** | -1.565 | -0.471 | **0.338** | 7.04E-49 | **1.54E-46** |
| ENSG00000101003.13 | **GINS1** | 9837 | protein_coding | **226.356** | -1.769 | -0.532 | **0.293** | 1.33E-48 | **2.87E-46** |
| ENSG00000139193.6 | **CD27** | 939 | protein_coding | **2568.326** | -2.727 | -0.821 | **0.151** | 1.38E-48 | **2.95E-46** |
| ENSG00000075399.16 | **VPS9D1** | 9605 | protein_coding | **525.894** | 2.330 | 0.701 | **5.027** | 1.85E-48 | **3.91E-46** |
| ENSG00000178078.14 | **STAP2** | 55620 | protein_coding | **198.408** | -2.007 | -0.604 | **0.249** | 2.27E-48 | **4.74E-46** |
| ENSG00000160323.21 | **ADAMTS13** | 11093 | protein_coding | **408.926** | 2.218 | 0.668 | **4.654** | 3.84E-48 | **7.93E-46** |
| ENSG00000094804.14 | **CDC6** | 990 | protein_coding | **375.653** | -1.933 | -0.582 | **0.262** | 4.38E-48 | **8.95E-46** |
| ENSG00000019582.19 | **CD74** | 972 | protein_coding | **171736.274** | 1.605 | 0.483 | **3.042** | 8.10E-48 | **1.64E-45** |
| ENSG00000204580.16 | **DDR1** | 780 | protein_coding | **715.590** | 1.586 | 0.478 | **3.003** | 1.52E-47 | **3.05E-45** |
| ENSG00000178038.19 | **ALS2CL** | 259173 | protein_coding | **96.302** | 4.449 | 1.339 | **21.839** | 1.71E-47 | **3.40E-45** |
| ENSG00000107551.23 | **RASSF4** | 83937 | protein_coding | **1320.374** | 1.741 | 0.524 | **3.343** | 3.15E-47 | **6.18E-45** |
| ENSG00000123485.13 | **HJURP** | 55355 | protein_coding | **780.458** | -2.646 | -0.797 | **0.160** | 6.24E-47 | **1.21E-44** |
| ENSG00000196584.5 | **XRCC2** | 7516 | protein_coding | **135.254** | -2.031 | -0.611 | **0.245** | 1.03E-46 | **1.97E-44** |
| ENSG00000133321.14 | **PLAAT4** | 5920 | protein_coding | **646.520** | 2.259 | 0.680 | **4.785** | 1.40E-46 | **2.67E-44** |
| ENSG00000111247.17 | **RAD51AP1** | 10635 | protein_coding | **183.518** | -2.322 | -0.699 | **0.200** | 1.46E-46 | **2.75E-44** |
| ENSG00000277363.5 | **SRCIN1** | 80725 | protein_coding | **129.265** | 3.868 | 1.165 | **14.606** | 4.11E-46 | **7.67E-44** |
| ENSG00000171848.18 | **RRM2** | 6241 | protein_coding | **1564.021** | -2.000 | -0.602 | **0.250** | 5.07E-46 | **9.37E-44** |
| ENSG00000127419.19 | **TMEM175** | 84286 | protein_coding | **1278.364** | 1.456 | 0.438 | **2.743** | 1.29E-45 | **2.36E-43** |
| ENSG00000076003.7 | **MCM6** | 4175 | protein_coding | **928.250** | -1.388 | -0.418 | **0.382** | 1.37E-45 | **2.48E-43** |
| ENSG00000105419.20 | **MEIS3** | 56917 | protein_coding | **131.047** | 6.460 | 1.945 | **88.039** | 1.51E-45 | **2.71E-43** |
| ENSG00000204681.13 | **GABBR1** | 2550 | protein_coding | **1024.406** | 2.567 | 0.773 | **5.925** | 1.68E-45 | **2.98E-43** |
| ENSG00000147536.13 | **GINS4** | 84296 | protein_coding | **321.206** | -1.928 | -0.580 | **0.263** | 2.94E-45 | **5.17E-43** |
| ENSG00000111445.16 | **RFC5** | 5985 | protein_coding | **469.733** | -1.533 | -0.461 | **0.346** | 4.04E-45 | **7.05E-43** |
| ENSG00000051180.19 | **RAD51** | 5888 | protein_coding | **259.677** | -2.239 | -0.674 | **0.212** | 5.60E-45 | **9.68E-43** |
| ENSG00000168280.19 | **KIF5C** | 3800 | protein_coding | **108.423** | 3.360 | 1.012 | **10.270** | 6.94E-45 | **1.19E-42** |
| ENSG00000068489.13 | **PRR11** | 55771 | protein_coding | **522.171** | -2.064 | -0.621 | **0.239** | 7.35E-45 | **1.25E-42** |
| ENSG00000161800.15 | **RACGAP1** | 29127 | protein_coding | **538.624** | -1.489 | -0.448 | **0.356** | 1.22E-44 | **2.05E-42** |
| ENSG00000105290.16 | **APLP1** | 333 | protein_coding | **113.520** | 3.040 | 0.915 | **8.227** | 1.62E-44 | **2.70E-42** |
| ENSG00000215784.8 | **FAM72D** | 728833 | protein_coding | **146.359** | -2.471 | -0.744 | **0.180** | 5.37E-44 | **8.88E-42** |
| ENSG00000135476.14 | **ESPL1** | 9700 | protein_coding | **692.021** | -2.051 | -0.617 | **0.241** | 7.74E-44 | **1.27E-41** |
| ENSG00000012822.18 | **CALCOCO1** | 57658 | protein_coding | **2479.173** | 1.903 | 0.573 | **3.741** | 9.48E-44 | **1.54E-41** |

| | | | | | | | | | |
|---|---|---|---|---|---|---|---|---|---|
| ENSG00000146410.14 | **MTFR2** | 113115 | protein_coding | **193.023** | -2.133 | -0.642 | **0.228** | 1.44E-43 | **2.32E-41** |
| ENSG00000105270.17 | **CLIP3** | 25999 | protein_coding | **272.805** | 2.212 | 0.666 | **4.632** | 1.65E-43 | **2.63E-41** |
| ENSG00000084636.20 | **COL16A1** | 1307 | protein_coding | **357.561** | 2.174 | 0.654 | **4.512** | 2.02E-43 | **3.19E-41** |
| ENSG00000120616.17 | **EPC1** | 80314 | protein_coding | **303.060** | -1.688 | -0.508 | **0.310** | 2.03E-43 | **3.19E-41** |
| ENSG00000157653.14 | **C9orf43** | 257169 | protein_coding | **124.944** | 2.735 | 0.823 | **6.656** | 4.63E-43 | **7.23E-41** |
| ENSG00000254815.7 | **LMNTD2-A** | 692247 | lncRNA | **170.726** | 4.734 | 1.425 | **26.610** | 6.30E-43 | **9.74E-41** |
| ENSG00000164850.18 | **GPER1** | 2852 | protein_coding | **193.707** | -2.362 | -0.711 | **0.194** | 8.17E-43 | **1.25E-40** |
| ENSG00000184635.18 | **ZNF93** | 81931 | protein_coding | **143.199** | -2.189 | -0.659 | **0.219** | 9.56E-43 | **1.45E-40** |
| ENSG00000271646.2 | **IRF2-DT** | 117134601 | lncRNA | **226.111** | 1.859 | 0.560 | **3.627** | 1.06E-42 | **1.60E-40** |
| ENSG00000167513.11 | **CDT1** | 81620 | protein_coding | **660.746** | -2.133 | -0.642 | **0.228** | 1.25E-42 | **1.87E-40** |
| ENSG00000066279.21 | **ASPM** | 259266 | protein_coding | **300.961** | -2.756 | -0.830 | **0.148** | 1.50E-42 | **2.23E-40** |
| ENSG00000127585.15 | **FBXL16** | 146330 | protein_coding | **238.503** | 2.770 | 0.834 | **6.821** | 1.65E-42 | **2.43E-40** |
| ENSG00000213889.13 | **PPM1N** | 147699 | protein_coding | **230.742** | 2.802 | 0.844 | **6.975** | 1.77E-42 | **2.59E-40** |
| ENSG00000012048.28 | **BRCA1** | 672 | protein_coding | **330.518** | -2.190 | -0.659 | **0.219** | 2.00E-42 | **2.90E-40** |
| ENSG00000169679.17 | **BUB1** | 699 | protein_coding | **708.109** | -1.897 | -0.571 | **0.269** | 3.49E-42 | **5.02E-40** |
| ENSG00000168026.21 | **TTC21A** | 199223 | protein_coding | **377.713** | 2.018 | 0.607 | **4.049** | 9.55E-42 | **1.37E-39** |
| ENSG00000141510.21 | **TP53** | 7157 | protein_coding | **1087.179** | -1.876 | -0.565 | **0.272** | 2.14E-41 | **3.03E-39** |
| ENSG00000109881.19 | **CCDC34** | 91057 | protein_coding | **188.980** | -2.095 | -0.631 | **0.234** | 2.32E-41 | **3.27E-39** |
| ENSG00000103316.14 | **CRYM** | 1428 | protein_coding | **235.755** | 3.339 | 1.005 | **10.118** | 3.50E-41 | **4.90E-39** |
| ENSG00000167964.15 | **RAB26** | 25837 | protein_coding | **147.561** | 3.490 | 1.051 | **11.237** | 3.59E-41 | **4.99E-39** |
| ENSG00000035141.12 | **FAM136A** | 84908 | protein_coding | **875.212** | -1.296 | -0.390 | **0.407** | 7.36E-41 | **1.01E-38** |
| ENSG00000142235.14 | **LMTK3** | 114783 | protein_coding | **131.834** | 3.265 | 0.983 | **9.612** | 8.51E-41 | **1.17E-38** |
| ENSG00000124635.12 | **H2BC11** | 8970 | protein_coding | **231.982** | -2.248 | -0.677 | **0.211** | 1.04E-40 | **1.42E-38** |
| ENSG00000183684.10 | **ALYREF** | 10189 | protein_coding | **715.734** | -1.535 | -0.462 | **0.345** | 1.16E-40 | **1.57E-38** |
| ENSG00000126860.14 | **EVI2A** | 2123 | protein_coding | **301.984** | -3.018 | -0.909 | **0.123** | 1.29E-40 | **1.72E-38** |
| ENSG00000213366.15 | **GSTM2** | 2946 | protein_coding | **1282.618** | 1.676 | 0.505 | **3.195** | 1.34E-40 | **1.78E-38** |
| ENSG00000138821.16 | **SLC39A8** | 64116 | protein_coding | **440.403** | -1.623 | -0.489 | **0.325** | 1.64E-40 | **2.16E-38** |
| ENSG00000140525.22 | **FANCI** | 55215 | protein_coding | **1152.686** | -2.267 | -0.682 | **0.208** | 1.94E-40 | **2.54E-38** |
| ENSG00000123416.17 | **TUBA1B** | 10376 | protein_coding | **13917.730** | -2.239 | -0.674 | **0.212** | 2.42E-40 | **3.16E-38** |
| ENSG00000185664.17 | **PMEL** | 6490 | protein_coding | **177.945** | 3.150 | 0.948 | **8.879** | 3.28E-40 | **4.24E-38** |
| ENSG00000164466.15 | **SFXN1** | 94081 | protein_coding | **797.016** | -1.219 | -0.367 | **0.430** | 3.48E-40 | **4.47E-38** |

| | | | | | | | | | |
|---|---|---|---|---|---|---|---|---|---|
| ENSG00000185189.20 | **NRBP2** | 340371 | protein_coding | **589.910** | 1.419 | 0.427 | **2.674** | 6.45E-40 | **8.24E-38** |
| ENSG00000096996.17 | **IL12RB1** | 3594 | protein_coding | **250.164** | -2.153 | -0.648 | **0.225** | 6.62E-40 | **8.39E-38** |
| ENSG00000184524.8 | **CEND1** | 51286 | protein_coding | **177.133** | 3.137 | 0.944 | **8.800** | 1.00E-39 | **1.26E-37** |
| ENSG00000077514.11 | **POLD3** | 10714 | protein_coding | **319.653** | -1.744 | -0.525 | **0.298** | 1.19E-39 | **1.49E-37** |
| ENSG00000000460.19 | **FIRRM** | 55732 | protein_coding | **348.239** | -1.572 | -0.473 | **0.336** | 2.53E-39 | **3.14E-37** |
| ENSG00000071539.16 | **TRIP13** | 9319 | protein_coding | **241.609** | -2.399 | -0.722 | **0.190** | 4.30E-39 | **5.31E-37** |
| ENSG00000213402.3 | **PTPRCAP** | 5790 | protein_coding | **1818.499** | -2.359 | -0.710 | **0.195** | 4.37E-39 | **5.36E-37** |
| ENSG00000276043.7 | **UHRF1** | 29128 | protein_coding | **552.404** | -2.006 | -0.604 | **0.249** | 4.93E-39 | **6.01E-37** |
| ENSG00000136813.17 | **ECPAS** | 23392 | protein_coding | **1473.719** | -1.360 | -0.410 | **0.389** | 5.57E-39 | **6.75E-37** |
| ENSG00000142945.15 | **KIF2C** | 11004 | protein_coding | **827.134** | -2.215 | -0.667 | **0.215** | 7.26E-39 | **8.74E-37** |
| ENSG00000104833.13 | **TUBB4A** | 10382 | protein_coding | **426.182** | 2.539 | 0.764 | **5.810** | 9.76E-39 | **1.17E-36** |
| ENSG00000017483.18 | **SLC38A5** | 92745 | protein_coding | **730.266** | -1.917 | -0.577 | **0.265** | 1.05E-38 | **1.24E-36** |
| ENSG00000131398.15 | **KCNC3** | 3748 | protein_coding | **263.148** | 1.855 | 0.558 | **3.617** | 1.22E-38 | **1.44E-36** |
| ENSG00000114626.20 | **ABTB1** | 80325 | protein_coding | **665.508** | 1.487 | 0.448 | **2.802** | 1.54E-38 | **1.81E-36** |
| ENSG00000183248.13 | **PRR36** | 80164 | protein_coding | **96.160** | 4.778 | 1.438 | **27.441** | 1.85E-38 | **2.15E-36** |
| ENSG00000165501.19 | **LRR1** | 122769 | protein_coding | **204.466** | -1.700 | -0.512 | **0.308** | 1.85E-38 | **2.15E-36** |
| ENSG00000117724.16 | **CENPF** | 1063 | protein_coding | **1020.532** | -2.555 | -0.769 | **0.170** | 2.30E-38 | **2.65E-36** |
| ENSG00000065717.18 | **TLE2** | 7089 | protein_coding | **74.013** | 4.035 | 1.215 | **16.388** | 2.69E-38 | **3.08E-36** |
| ENSG00000058804.14 | **NDC1** | 55706 | protein_coding | **353.315** | -1.367 | -0.412 | **0.388** | 3.98E-38 | **4.53E-36** |
| ENSG00000188610.14 | **FAM72B** | 653820 | protein_coding | **191.347** | -2.732 | -0.822 | **0.151** | 5.08E-38 | **5.75E-36** |
| ENSG00000259781.1 | **HMGB1P6** | | processed_pseud | **902.589** | -1.637 | -0.493 | **0.321** | 6.93E-38 | **7.79E-36** |
| ENSG00000164087.10 | **POC1A** | 25886 | protein_coding | **218.106** | -1.821 | -0.548 | **0.283** | 7.84E-38 | **8.77E-36** |
| ENSG00000105255.13 | **FSD1** | 79187 | protein_coding | **116.760** | 3.889 | 1.171 | **14.813** | 7.89E-38 | **8.77E-36** |
| ENSG00000066923.21 | **STAG3** | 10734 | protein_coding | **1051.269** | -1.678 | -0.505 | **0.312** | 8.87E-38 | **9.81E-36** |
| ENSG00000156970.15 | **BUB1B** | 701 | protein_coding | **833.006** | -2.230 | -0.671 | **0.213** | 9.00E-38 | **9.89E-36** |
| ENSG00000167280.19 | **ENGASE** | 64772 | protein_coding | **1344.999** | 1.394 | 0.420 | **2.629** | 1.06E-37 | **1.15E-35** |
| ENSG00000246982.8 | **BRPF3-AS1** | | lncRNA | **251.897** | 2.419 | 0.728 | **5.349** | 1.08E-37 | **1.18E-35** |
| ENSG00000109805.12 | **NCAPG** | 64151 | protein_coding | **848.617** | -2.239 | -0.674 | **0.212** | 1.09E-37 | **1.18E-35** |
| ENSG00000182272.14 | **B4GALNT4** | 338707 | protein_coding | **114.797** | 5.816 | 1.751 | **56.343** | 1.33E-37 | **1.43E-35** |
| ENSG00000139625.15 | **MAP3K12** | 7786 | protein_coding | **583.561** | 1.880 | 0.566 | **3.680** | 1.61E-37 | **1.72E-35** |
| ENSG00000159055.6 | **MIS18A** | 54069 | protein_coding | **193.124** | -1.663 | -0.501 | **0.316** | 1.62E-37 | **1.72E-35** |

| | | | | | | | | | |
|---|---|---|---|---|---|---|---|---|---|
| ENSG00000174428.21 | **GTF2IRD2E** | 389524 | protein_coding | **258.370** | -1.534 | -0.462 | **0.345** | 1.83E-37 | **1.93E-35** |
| ENSG00000125885.15 | **MCM8** | 84515 | protein_coding | **353.325** | -1.563 | -0.470 | **0.339** | 2.04E-37 | **2.14E-35** |
| ENSG00000167325.17 | **RRM1** | 6240 | protein_coding | **827.563** | -1.598 | -0.481 | **0.330** | 2.14E-37 | **2.24E-35** |
| ENSG00000167702.15 | **KIFC2** | 90990 | protein_coding | **1712.556** | 1.733 | 0.522 | **3.325** | 2.38E-37 | **2.48E-35** |
| ENSG00000006607.16 | **FARP2** | 9855 | protein_coding | **791.881** | 1.182 | 0.356 | **2.270** | 2.46E-37 | **2.55E-35** |
| ENSG00000101361.19 | **NOP56** | 10528 | protein_coding | **3060.882** | -0.928 | -0.279 | **0.525** | 2.94E-37 | **3.02E-35** |
| ENSG00000166851.17 | **PLK1** | 5347 | protein_coding | **563.774** | -1.954 | -0.588 | **0.258** | 3.11E-37 | **3.18E-35** |
| ENSG00000113810.18 | **SMC4** | 10051 | protein_coding | **1313.255** | -1.679 | -0.505 | **0.312** | 3.81E-37 | **3.88E-35** |
| ENSG00000167103.13 | **PIP5KL1** | 138429 | protein_coding | **112.622** | 4.046 | 1.218 | **16.519** | 4.25E-37 | **4.30E-35** |
| ENSG00000178202.15 | **POGLUT3** | 143888 | protein_coding | **117.809** | -2.192 | -0.660 | **0.219** | 5.12E-37 | **5.15E-35** |
| ENSG00000108828.18 | **VAT1** | 10493 | protein_coding | **908.980** | 2.378 | 0.716 | **5.197** | 9.68E-37 | **9.69E-35** |
| ENSG00000293495.2 | **C19orf48P** | 84798 | lncRNA | **1593.059** | -1.534 | -0.462 | **0.345** | 1.01E-36 | **1.01E-34** |
| ENSG00000171241.11 | **SHCBP1** | 79801 | protein_coding | **335.684** | -2.387 | -0.719 | **0.191** | 1.09E-36 | **1.08E-34** |
| ENSG00000092621.15 | **PHGDH** | 26227 | protein_coding | **1691.756** | -2.102 | -0.633 | **0.233** | 1.12E-36 | **1.11E-34** |
| ENSG00000129173.15 | **E2F8** | 79733 | protein_coding | **137.280** | -2.505 | -0.754 | **0.176** | 1.30E-36 | **1.27E-34** |
| ENSG00000121769.10 | **FABP3** | 2170 | protein_coding | **144.779** | -3.053 | -0.919 | **0.121** | 1.31E-36 | **1.28E-34** |
| ENSG00000117632.25 | **STMN1** | 3925 | protein_coding | **3097.093** | -1.889 | -0.569 | **0.270** | 2.47E-36 | **2.40E-34** |
| ENSG00000186638.19 | **KIF24** | 347240 | protein_coding | **132.796** | -2.107 | -0.634 | **0.232** | 2.73E-36 | **2.64E-34** |
| ENSG00000165480.18 | **SKA3** | 221150 | protein_coding | **289.802** | -2.200 | -0.662 | **0.218** | 4.23E-36 | **4.06E-34** |
| ENSG00000075218.21 | **GTSE1** | 51512 | protein_coding | **377.943** | -2.676 | -0.806 | **0.156** | 5.00E-36 | **4.78E-34** |
| ENSG00000291006.1 | **ENSG0000(** | 730101 | lncRNA | **247.349** | 1.997 | 0.601 | **3.991** | 6.63E-36 | **6.31E-34** |
| ENSG00000137310.15 | **TCF19** | 6941 | protein_coding | **1083.694** | -1.541 | -0.464 | **0.344** | 6.77E-36 | **6.40E-34** |
| ENSG00000072571.22 | **HMMR** | 3161 | protein_coding | **217.490** | -1.821 | -0.548 | **0.283** | 6.92E-36 | **6.51E-34** |
| ENSG00000100749.11 | **VRK1** | 7443 | protein_coding | **622.923** | -1.339 | -0.403 | **0.395** | 9.43E-36 | **8.84E-34** |
| ENSG00000105048.20 | **TNNT1** | 7138 | protein_coding | **58.770** | 6.057 | 1.823 | **66.582** | 1.13E-35 | **1.06E-33** |
| ENSG00000006625.20 | **GGCT** | 79017 | protein_coding | **727.984** | -1.093 | -0.329 | **0.469** | 1.38E-35 | **1.28E-33** |
| ENSG00000169857.11 | **AVEN** | 57099 | protein_coding | **175.505** | -1.759 | -0.530 | **0.295** | 1.43E-35 | **1.32E-33** |
| ENSG00000163462.20 | **TRIM46** | 80128 | protein_coding | **148.536** | 2.009 | 0.605 | **4.025** | 1.59E-35 | **1.46E-33** |
| ENSG00000130513.8 | **GDF15** | 9518 | protein_coding | **91.037** | -2.929 | -0.882 | **0.131** | 1.72E-35 | **1.57E-33** |
| ENSG00000095539.18 | **SEMA4G** | 57715 | protein_coding | **72.563** | 3.151 | 0.949 | **8.883** | 1.94E-35 | **1.77E-33** |
| ENSG00000168496.6 | **FEN1** | 2237 | protein_coding | **871.577** | -1.866 | -0.562 | **0.274** | 2.14E-35 | **1.94E-33** |

| | | | | | | | | | |
|---|---|---|---|---|---|---|---|---|---|
| ENSG00000141854.11 | **MISP3** | 113230 | protein_coding | **87.785** | 2.361 | 0.711 | **5.137** | 2.32E-35 | **2.09E-33** |
| ENSG00000143228.15 | **NUF2** | 83540 | protein_coding | **350.262** | -2.528 | -0.761 | **0.173** | 3.29E-35 | **2.95E-33** |
| ENSG00000182809.13 | **CRIP2** | 1397 | protein_coding | **129.154** | 3.724 | 1.121 | **13.218** | 3.50E-35 | **3.13E-33** |
| ENSG00000197451.14 | **HNRNPAB** | 3182 | protein_coding | **3007.173** | -1.485 | -0.447 | **0.357** | 4.74E-35 | **4.22E-33** |
| ENSG00000168676.12 | **KCTD19** | 146212 | protein_coding | **131.476** | 4.316 | 1.299 | **19.916** | 6.04E-35 | **5.34E-33** |
| ENSG00000159214.15 | **CCDC24** | 149473 | protein_coding | **259.843** | 2.229 | 0.671 | **4.688** | 6.37E-35 | **5.61E-33** |
| ENSG00000108587.18 | **GOSR1** | 9527 | protein_coding | **700.421** | -1.171 | -0.352 | **0.444** | 6.72E-35 | **5.89E-33** |
| ENSG00000186862.22 | **PDZD7** | 79955 | protein_coding | **51.683** | 4.265 | 1.284 | **19.222** | 6.98E-35 | **6.09E-33** |
| ENSG00000123384.15 | **LRP1** | 4035 | protein_coding | **198.573** | 2.395 | 0.721 | **5.259** | 7.81E-35 | **6.77E-33** |
| ENSG00000171793.19 | **CTPS1** | 1503 | protein_coding | **601.855** | -1.720 | -0.518 | **0.304** | 7.83E-35 | **6.77E-33** |
| ENSG00000197766.11 | **CFD** | 1675 | protein_coding | **177.286** | 4.028 | 1.213 | **16.312** | 9.41E-35 | **8.10E-33** |
| ENSG00000225783.10 | **MIAT** | 440823 | lncRNA | **1655.502** | 2.004 | 0.603 | **4.011** | 9.77E-35 | **8.37E-33** |
| ENSG00000167670.18 | **CHAF1A** | 10036 | protein_coding | **624.621** | -1.268 | -0.382 | **0.415** | 9.86E-35 | **8.41E-33** |
| ENSG00000121211.10 | **MND1** | 84057 | protein_coding | **207.914** | -2.116 | -0.637 | **0.231** | 1.10E-34 | **9.37E-33** |
| ENSG00000132646.13 | **PCNA** | 5111 | protein_coding | **2306.095** | -1.650 | -0.497 | **0.319** | 1.23E-34 | **1.04E-32** |
| ENSG00000166508.21 | **MCM7** | 4176 | protein_coding | **2318.390** | -1.158 | -0.349 | **0.448** | 1.43E-34 | **1.20E-32** |
| ENSG00000170855.6 | **TRIAP1** | 51499 | protein_coding | **453.552** | -1.350 | -0.406 | **0.392** | 1.66E-34 | **1.39E-32** |
| ENSG00000161692.19 | **DBF4B** | 80174 | protein_coding | **533.727** | -1.227 | -0.369 | **0.427** | 2.06E-34 | **1.72E-32** |
| ENSG00000178999.15 | **AURKB** | 9212 | protein_coding | **1247.884** | -2.623 | -0.790 | **0.162** | 2.08E-34 | **1.73E-32** |
| ENSG00000119969.18 | **HELLS** | 3070 | protein_coding | **664.565** | -1.182 | -0.356 | **0.441** | 2.10E-34 | **1.74E-32** |
| ENSG00000196968.12 | **POFUT4** | 170384 | protein_coding | **662.814** | -1.845 | -0.555 | **0.278** | 2.22E-34 | **1.83E-32** |
| ENSG00000126583.13 | **PRKCG** | 5582 | protein_coding | **70.716** | 5.050 | 1.520 | **33.126** | 2.32E-34 | **1.90E-32** |
| ENSG00000165905.20 | **LARGE2** | 120071 | protein_coding | **83.386** | 4.506 | 1.357 | **22.728** | 2.33E-34 | **1.90E-32** |
| ENSG00000180921.9 | **FAM83H** | 286077 | protein_coding | **184.096** | 1.966 | 0.592 | **3.907** | 2.34E-34 | **1.90E-32** |
| ENSG00000186301.8 | **MST1P2** | | transcribed_unprc | **143.940** | 2.047 | 0.616 | **4.132** | 2.48E-34 | **2.01E-32** |
| ENSG00000160117.18 | **ANKLE1** | 126549 | protein_coding | **259.906** | -1.539 | -0.463 | **0.344** | 2.49E-34 | **2.01E-32** |
| ENSG00000127418.17 | **FGFRL1** | 53834 | protein_coding | **270.411** | 2.365 | 0.712 | **5.151** | 2.63E-34 | **2.11E-32** |
| ENSG00000163517.17 | **HDAC11** | 79885 | protein_coding | **99.218** | 2.249 | 0.677 | **4.754** | 3.69E-34 | **2.95E-32** |
| ENSG00000142230.15 | **SAE1** | 10055 | protein_coding | **1294.799** | -1.074 | -0.323 | **0.475** | 3.87E-34 | **3.08E-32** |
| ENSG00000084764.14 | **MAPRE3** | 22924 | protein_coding | **165.500** | 2.473 | 0.744 | **5.550** | 4.26E-34 | **3.38E-32** |
| ENSG00000151575.17 | **TEX9** | 374618 | protein_coding | **264.916** | -1.698 | -0.511 | **0.308** | 5.00E-34 | **3.94E-32** |

| | | | | | | | | | |
|---|---|---|---|---|---|---|---|---|---|
| ENSG00000173531.17 | **MST1** | 4485 | protein_coding | **1382.949** | 1.696 | 0.511 | **3.240** | 5.26E-34 | **4.13E-32** |
| ENSG00000163535.20 | **SGO2** | 151246 | protein_coding | **151.681** | -1.947 | -0.586 | **0.259** | 5.95E-34 | **4.66E-32** |
| ENSG00000113460.15 | **BRIX1** | 55299 | protein_coding | **563.393** | -1.072 | -0.323 | **0.476** | 7.18E-34 | **5.59E-32** |
| ENSG00000152253.11 | **SPC25** | 57405 | protein_coding | **107.667** | -2.822 | -0.849 | **0.141** | 7.44E-34 | **5.77E-32** |
| ENSG00000123473.19 | **STIL** | 6491 | protein_coding | **245.946** | -1.906 | -0.574 | **0.267** | 7.75E-34 | **5.99E-32** |
| ENSG00000172638.15 | **EFEMP2** | 30008 | protein_coding | **1285.584** | 2.082 | 0.627 | **4.234** | 8.17E-34 | **6.29E-32** |
| ENSG00000120539.17 | **MASTL** | 84930 | protein_coding | **490.160** | -1.611 | -0.485 | **0.327** | 8.31E-34 | **6.37E-32** |
| ENSG00000166451.16 | **CENPN** | 55839 | protein_coding | **303.425** | -1.768 | -0.532 | **0.294** | 1.13E-33 | **8.64E-32** |
| ENSG00000166145.17 | **SPINT1** | 6692 | protein_coding | **415.836** | 1.574 | 0.474 | **2.978** | 1.20E-33 | **9.13E-32** |
| ENSG00000128050.11 | **PAICS** | 10606 | protein_coding | **3531.072** | -1.490 | -0.448 | **0.356** | 1.34E-33 | **1.02E-31** |
| ENSG00000293339.1 | **ENSG00000293339** | | lncRNA | **282.530** | -1.968 | -0.592 | **0.256** | 1.50E-33 | **1.13E-31** |
| ENSG00000139428.14 | **MMAB** | 326625 | protein_coding | **707.034** | -1.652 | -0.497 | **0.318** | 1.79E-33 | **1.35E-31** |
| ENSG00000241489.8 | **ENSG00000241489** | | protein_coding | **259.791** | 2.053 | 0.618 | **4.149** | 1.85E-33 | **1.38E-31** |
| ENSG00000137547.11 | **MRPL15** | 29088 | protein_coding | **716.898** | -1.116 | -0.336 | **0.461** | 1.98E-33 | **1.48E-31** |
| ENSG00000158402.23 | **CDC25C** | 995 | protein_coding | **188.885** | -2.258 | -0.680 | **0.209** | 2.15E-33 | **1.60E-31** |
| ENSG00000185761.13 | **ADAMTSL5** | 339366 | protein_coding | **44.125** | 4.539 | 1.366 | **23.248** | 2.32E-33 | **1.72E-31** |
| ENSG00000068308.17 | **OTUD5** | 55593 | protein_coding | **2839.982** | 1.256 | 0.378 | **2.388** | 3.13E-33 | **2.31E-31** |
| ENSG00000168993.17 | **CPLX1** | 10815 | protein_coding | **71.984** | 3.682 | 1.108 | **12.832** | 3.35E-33 | **2.46E-31** |
| ENSG00000085999.15 | **RAD54L** | 8438 | protein_coding | **463.277** | -2.223 | -0.669 | **0.214** | 3.99E-33 | **2.92E-31** |
| ENSG00000175130.9 | **MARCKSL1** | 65108 | protein_coding | **11163.642** | 1.261 | 0.380 | **2.396** | 7.23E-33 | **5.27E-31** |
| ENSG00000129810.17 | **SGO1** | 151648 | protein_coding | **161.273** | -2.422 | -0.729 | **0.187** | 7.66E-33 | **5.56E-31** |
| ENSG00000011426.13 | **ANLN** | 54443 | protein_coding | **184.176** | -2.128 | -0.640 | **0.229** | 7.80E-33 | **5.64E-31** |
| ENSG00000123219.15 | **CENPK** | 64105 | protein_coding | **170.998** | -2.413 | -0.726 | **0.188** | 8.03E-33 | **5.79E-31** |
| ENSG00000176208.11 | **ATAD5** | 79915 | protein_coding | **177.681** | -2.073 | -0.624 | **0.238** | 8.55E-33 | **6.13E-31** |
| ENSG00000105613.11 | **MAST1** | 22983 | protein_coding | **105.959** | 3.677 | 1.107 | **12.790** | 9.24E-33 | **6.60E-31** |
| ENSG00000105289.17 | **TJP3** | 27134 | protein_coding | **58.198** | 5.234 | 1.576 | **37.641** | 1.03E-32 | **7.31E-31** |
| ENSG00000136960.15 | **ENPP2** | 5168 | protein_coding | **193.710** | 3.290 | 0.990 | **9.781** | 1.22E-32 | **8.62E-31** |
| ENSG00000163918.14 | **RFC4** | 5984 | protein_coding | **447.730** | -1.296 | -0.390 | **0.407** | 1.32E-32 | **9.30E-31** |
| ENSG00000100983.14 | **GSS** | 2937 | protein_coding | **845.723** | -1.386 | -0.417 | **0.383** | 1.90E-32 | **1.34E-30** |
| ENSG00000213213.15 | **CCDC183** | 84960 | protein_coding | **68.553** | 3.624 | 1.091 | **12.325** | 1.98E-32 | **1.39E-30** |
| ENSG00000151849.18 | **CPAP** | 55835 | protein_coding | **552.529** | -1.354 | -0.408 | **0.391** | 2.10E-32 | **1.47E-30** |

| | | | | | | | | | |
|---|---|---|---|---|---|---|---|---|---|
| ENSG00000122966.20 | **CIT** | 11113 | protein_coding | **500.781** | -1.981 | -0.596 | **0.253** | 2.79E-32 | **1.95E-30** |
| ENSG00000112029.11 | **FBXO5** | 26271 | protein_coding | **169.697** | -2.093 | -0.630 | **0.234** | 3.53E-32 | **2.45E-30** |
| ENSG00000006118.16 | **TMEM132/** | 54972 | protein_coding | **394.768** | 2.491 | 0.750 | **5.621** | 3.65E-32 | **2.52E-30** |
| ENSG00000163898.12 | **LIPH** | 200879 | protein_coding | **369.516** | 1.879 | 0.566 | **3.679** | 3.70E-32 | **2.55E-30** |
| ENSG00000188747.11 | **NOXA1** | 10811 | protein_coding | **217.738** | 3.275 | 0.986 | **9.683** | 4.18E-32 | **2.87E-30** |
| ENSG00000139636.18 | **LMBR1L** | 55716 | protein_coding | **1073.754** | 1.146 | 0.345 | **2.213** | 4.48E-32 | **3.07E-30** |
| ENSG00000159259.10 | **CHAF1B** | 8208 | protein_coding | **327.167** | -1.339 | -0.403 | **0.395** | 4.97E-32 | **3.39E-30** |
| ENSG00000163808.19 | **KIF15** | 56992 | protein_coding | **245.224** | -2.420 | -0.728 | **0.187** | 6.11E-32 | **4.15E-30** |
| ENSG00000102879.19 | **CORO1A** | 11151 | protein_coding | **11792.410** | -2.197 | -0.661 | **0.218** | 6.52E-32 | **4.41E-30** |
| ENSG00000139880.22 | **CDH24** | 64403 | protein_coding | **187.495** | 2.380 | 0.717 | **5.207** | 7.22E-32 | **4.87E-30** |
| ENSG00000182459.6 | **TEX19** | 400629 | protein_coding | **45.836** | 4.074 | 1.226 | **16.844** | 9.57E-32 | **6.43E-30** |
| ENSG00000162433.17 | **AK4** | 205 | protein_coding | **1195.311** | -1.593 | -0.479 | **0.332** | 1.07E-31 | **7.15E-30** |
| ENSG00000121621.9 | **KIF18A** | 81930 | protein_coding | **129.416** | -2.204 | -0.664 | **0.217** | 1.26E-31 | **8.44E-30** |
| ENSG00000213551.9 | **DNAJC9** | 23234 | protein_coding | **727.954** | -1.059 | -0.319 | **0.480** | 1.32E-31 | **8.79E-30** |
| ENSG00000140545.18 | **MFGE8** | 4240 | protein_coding | **209.080** | 1.885 | 0.567 | **3.693** | 1.36E-31 | **9.00E-30** |
| ENSG00000131153.11 | **GINS2** | 51659 | protein_coding | **659.197** | -1.699 | -0.512 | **0.308** | 1.39E-31 | **9.21E-30** |
| ENSG00000265972.8 | **TXNIP** | 10628 | protein_coding | **4823.360** | 2.668 | 0.803 | **6.354** | 1.43E-31 | **9.39E-30** |
| ENSG00000271046.2 | **ENSG0000(** | 105376412 | lncRNA | **150.960** | 3.648 | 1.098 | **12.540** | 1.51E-31 | **9.91E-30** |
| ENSG00000104738.21 | **MCM4** | 4173 | protein_coding | **1337.872** | -1.598 | -0.481 | **0.330** | 1.61E-31 | **1.05E-29** |
| ENSG00000181085.17 | **MAPK15** | 225689 | protein_coding | **85.576** | 5.207 | 1.567 | **36.927** | 1.79E-31 | **1.17E-29** |
| ENSG00000237520.3 | **LINC02971** | 123706537 | lncRNA | **80.981** | 2.625 | 0.790 | **6.169** | 1.97E-31 | **1.28E-29** |
| ENSG00000224660.2 | **SH3BP5-AS** | 100505696 | lncRNA | **447.104** | 1.861 | 0.560 | **3.632** | 2.08E-31 | **1.34E-29** |
| ENSG00000143476.20 | **DTL** | 51514 | protein_coding | **544.026** | -2.108 | -0.635 | **0.232** | 2.09E-31 | **1.35E-29** |
| ENSG00000136732.18 | **GYPC** | 2995 | protein_coding | **1396.780** | -1.403 | -0.422 | **0.378** | 2.13E-31 | **1.37E-29** |
| ENSG00000113368.14 | **LMNB1** | 4001 | protein_coding | **1544.457** | -1.396 | -0.420 | **0.380** | 2.16E-31 | **1.38E-29** |
| ENSG00000237649.10 | **KIFC1** | 3833 | protein_coding | **806.743** | -2.044 | -0.615 | **0.243** | 2.40E-31 | **1.53E-29** |
| ENSG00000185480.14 | **PARPBP** | 55010 | protein_coding | **127.115** | -1.915 | -0.577 | **0.265** | 2.42E-31 | **1.54E-29** |
| ENSG00000110848.8 | **CD69** | 969 | protein_coding | **1607.347** | 1.582 | 0.476 | **2.993** | 2.76E-31 | **1.75E-29** |
| ENSG00000111196.13 | **MAGOHB** | 55110 | protein_coding | **261.681** | -1.635 | -0.492 | **0.322** | 3.21E-31 | **2.03E-29** |
| ENSG00000100379.20 | **KCTD17** | 79734 | protein_coding | **226.741** | 1.457 | 0.439 | **2.746** | 3.25E-31 | **2.05E-29** |
| ENSG00000168918.15 | **INPP5D** | 3635 | protein_coding | **2311.977** | -1.177 | -0.354 | **0.442** | 3.70E-31 | **2.32E-29** |

| | | | | | | | | | |
|---|---|---|---|---|---|---|---|---|---|
| ENSG00000186648.17 | **CARMIL3** | 90668 | protein_coding | **98.738** | 2.687 | 0.809 | **6.440** | 3.73E-31 | **2.33E-29** |
| ENSG00000104972.18 | **LILRB1** | 10859 | protein_coding | **4163.169** | 0.967 | 0.291 | **1.955** | 3.76E-31 | **2.34E-29** |
| ENSG00000145220.16 | **LYAR** | 55646 | protein_coding | **527.997** | -1.354 | -0.408 | **0.391** | 3.93E-31 | **2.45E-29** |
| ENSG00000060688.15 | **SNRNP40** | 9410 | protein_coding | **528.518** | -1.105 | -0.333 | **0.465** | 4.01E-31 | **2.48E-29** |
| ENSG00000089847.15 | **ANKRD24** | 170961 | protein_coding | **351.915** | 2.009 | 0.605 | **4.025** | 4.93E-31 | **3.04E-29** |
| ENSG00000275418.2 | **LINC03110** | 10272474[illegible] | lncRNA | **454.851** | 1.784 | 0.537 | **3.444** | 5.30E-31 | **3.26E-29** |
| ENSG00000132563.19 | **REEP2** | 51308 | protein_coding | **148.395** | 2.108 | 0.635 | **4.312** | 5.50E-31 | **3.37E-29** |
| ENSG00000253893.3 | **FAM85B** | | lncRNA | **196.377** | 1.917 | 0.577 | **3.776** | 6.73E-31 | **4.12E-29** |
| ENSG00000138180.18 | **CEP55** | 55165 | protein_coding | **342.045** | -2.120 | -0.638 | **0.230** | 1.18E-30 | **7.18E-29** |
| ENSG00000102390.14 | **PBDC1** | 51260 | protein_coding | **407.341** | -1.214 | -0.366 | **0.431** | 1.48E-30 | **9.00E-29** |
| ENSG00000167851.18 | **CD300A** | 11314 | protein_coding | **1696.582** | -1.944 | -0.585 | **0.260** | 1.67E-30 | **1.01E-28** |
| ENSG00000070814.24 | **TCOF1** | 6949 | protein_coding | **1357.139** | -0.942 | -0.284 | **0.521** | 1.73E-30 | **1.04E-28** |
| ENSG00000051341.17 | **POLQ** | 10721 | protein_coding | **434.649** | -1.583 | -0.476 | **0.334** | 2.41E-30 | **1.45E-28** |
| ENSG00000102393.16 | **GLA** | 2717 | protein_coding | **406.308** | -1.290 | -0.388 | **0.409** | 2.84E-30 | **1.70E-28** |
| ENSG00000154262.15 | **ABCA6** | 23460 | protein_coding | **296.656** | -1.525 | -0.459 | **0.348** | 3.38E-30 | **2.02E-28** |
| ENSG00000176749.11 | **CDK5R1** | 8851 | protein_coding | **263.186** | -1.616 | -0.487 | **0.326** | 3.40E-30 | **2.03E-28** |
| ENSG00000156802.15 | **ATAD2** | 29028 | protein_coding | **523.816** | -1.496 | -0.450 | **0.355** | 3.62E-30 | **2.15E-28** |
| ENSG00000135424.21 | **ITGA7** | 3679 | protein_coding | **152.140** | 2.414 | 0.727 | **5.329** | 3.78E-30 | **2.24E-28** |
| ENSG00000216490.6 | **IFI30** | 10437 | protein_coding | **6207.950** | -1.389 | -0.418 | **0.382** | 3.79E-30 | **2.24E-28** |
| ENSG00000115875.21 | **SRSF7** | 6432 | protein_coding | **2693.967** | -0.960 | -0.289 | **0.514** | 4.13E-30 | **2.43E-28** |
| ENSG00000204248.14 | **COL11A2** | 1302 | protein_coding | **163.355** | 2.238 | 0.674 | **4.716** | 4.66E-30 | **2.74E-28** |
| ENSG00000188312.16 | **CENPP** | 401541 | protein_coding | **184.724** | -1.746 | -0.526 | **0.298** | 4.81E-30 | **2.82E-28** |
| ENSG00000228716.9 | **DHFR** | 1719 | protein_coding | **1099.524** | -2.007 | -0.604 | **0.249** | 6.01E-30 | **3.51E-28** |
| ENSG00000184216.17 | **IRAK1** | 3654 | protein_coding | **842.881** | -1.369 | -0.412 | **0.387** | 6.15E-30 | **3.58E-28** |
| ENSG00000128928.12 | **IVD** | 3712 | protein_coding | **1509.651** | 0.902 | 0.271 | **1.868** | 6.44E-30 | **3.73E-28** |
| ENSG00000035499.15 | **DEPDC1B** | 55789 | protein_coding | **136.358** | -1.973 | -0.594 | **0.255** | 6.88E-30 | **3.98E-28** |
| ENSG00000102384.16 | **CENPI** | 2491 | protein_coding | **202.238** | -2.463 | -0.742 | **0.181** | 7.79E-30 | **4.49E-28** |
| ENSG00000112486.20 | **CCR6** | 1235 | protein_coding | **180.076** | -2.129 | -0.641 | **0.229** | 9.00E-30 | **5.17E-28** |
| ENSG00000023697.15 | **DERA** | 51071 | protein_coding | **317.670** | -1.300 | -0.391 | **0.406** | 9.24E-30 | **5.29E-28** |
| ENSG00000042088.16 | **TDP1** | 55775 | protein_coding | **525.469** | -1.129 | -0.340 | **0.457** | 1.10E-29 | **6.29E-28** |
| ENSG00000174080.14 | **CTSF** | 8722 | protein_coding | **104.953** | 2.832 | 0.853 | **7.121** | 1.39E-29 | **7.92E-28** |

| | | | | | | | | | |
|---|---|---|---|---|---|---|---|---|---|
| ENSG00000082014.17 | **SMARCD3** | 6604 | protein_coding | **62.491** | 3.665 | 1.103 | **12.688** | 1.58E-29 | **8.97E-28** |
| ENSG00000091651.11 | **ORC6** | 23594 | protein_coding | **575.025** | -1.860 | -0.560 | **0.275** | 1.58E-29 | **8.97E-28** |
| ENSG00000184602.8 | **SNN** | 8303 | protein_coding | **735.506** | 2.207 | 0.665 | **4.619** | 1.66E-29 | **9.39E-28** |
| ENSG00000137135.20 | **ARHGEF39** | 84904 | protein_coding | **299.132** | -2.314 | -0.697 | **0.201** | 2.06E-29 | **1.16E-27** |
| ENSG00000104889.9 | **RNASEH2A** | 10535 | protein_coding | **568.528** | -1.682 | -0.506 | **0.312** | 2.45E-29 | **1.37E-27** |
| ENSG00000245532.13 | **NEAT1** | 283131 | lncRNA | **17327.231** | 1.387 | 0.417 | **2.615** | 2.85E-29 | **1.60E-27** |
| ENSG00000121152.12 | **NCAPH** | 23397 | protein_coding | **568.356** | -1.975 | -0.595 | **0.254** | 2.87E-29 | **1.60E-27** |
| ENSG00000197903.11 | **H2BC12** | 85236 | protein_coding | **1720.965** | -1.957 | -0.589 | **0.258** | 2.95E-29 | **1.64E-27** |
| ENSG00000162062.17 | **TEDC2** | 80178 | protein_coding | **211.579** | -1.946 | -0.586 | **0.260** | 3.07E-29 | **1.70E-27** |
| ENSG00000285448.1 | **LNCARGI** | | lncRNA | **67.842** | 4.253 | 1.280 | **19.066** | 3.15E-29 | **1.74E-27** |
| ENSG00000131724.13 | **IL13RA1** | 3597 | protein_coding | **154.857** | 2.080 | 0.626 | **4.227** | 3.90E-29 | **2.15E-27** |
| ENSG00000175063.19 | **UBE2C** | 11065 | protein_coding | **1287.018** | -2.645 | -0.796 | **0.160** | 4.22E-29 | **2.32E-27** |
| ENSG00000181938.16 | **GINS3** | 64785 | protein_coding | **168.567** | -1.529 | -0.460 | **0.347** | 4.73E-29 | **2.59E-27** |
| ENSG00000140534.16 | **TICRR** | 90381 | protein_coding | **335.648** | -2.064 | -0.621 | **0.239** | 4.97E-29 | **2.72E-27** |
| ENSG00000104888.12 | **SLC17A7** | 57030 | protein_coding | **51.127** | 3.763 | 1.133 | **13.573** | 5.65E-29 | **3.08E-27** |
| ENSG00000137285.12 | **TUBB2B** | 347733 | protein_coding | **562.139** | 1.938 | 0.583 | **3.832** | 6.07E-29 | **3.30E-27** |
| ENSG00000029993.17 | **HMGB3** | 3149 | protein_coding | **569.322** | -1.848 | -0.556 | **0.278** | 6.09E-29 | **3.30E-27** |
| ENSG00000242732.6 | **RTL5** | 340526 | protein_coding | **91.023** | 2.558 | 0.770 | **5.890** | 6.59E-29 | **3.56E-27** |
| ENSG00000237886.3 | **NALT1** | | lncRNA | **315.793** | 1.745 | 0.525 | **3.353** | 6.73E-29 | **3.63E-27** |
| ENSG00000169715.16 | **MT1E** | 4493 | protein_coding | **155.304** | 2.917 | 0.878 | **7.554** | 6.99E-29 | **3.76E-27** |
| ENSG00000138101.22 | **DTNB** | 1838 | protein_coding | **294.894** | 1.703 | 0.513 | **3.256** | 7.69E-29 | **4.12E-27** |
| ENSG00000263513.8 | **FAM72C** | 554282 | protein_coding | **150.401** | -2.455 | -0.739 | **0.182** | 9.25E-29 | **4.94E-27** |
| ENSG00000135446.19 | **CDK4** | 1019 | protein_coding | **1802.664** | -1.314 | -0.395 | **0.402** | 1.10E-28 | **5.86E-27** |
| ENSG00000187017.19 | **ESPN** | 83715 | protein_coding | **85.624** | 3.751 | 1.129 | **13.461** | 1.10E-28 | **5.86E-27** |
| ENSG00000010803.20 | **SCMH1** | 22955 | protein_coding | **744.399** | -1.526 | -0.459 | **0.347** | 1.18E-28 | **6.23E-27** |
| ENSG00000255008.6 | **LINC02739** | 10536931[illegible] | lncRNA | **39.520** | 3.467 | 1.044 | **11.058** | 1.23E-28 | **6.50E-27** |
| ENSG00000149781.16 | **FERMT3** | 83706 | protein_coding | **2899.709** | -1.999 | -0.602 | **0.250** | 1.24E-28 | **6.54E-27** |
| ENSG00000162889.13 | **MAPKAPK[illegible]** | 9261 | protein_coding | **2574.852** | 1.578 | 0.475 | **2.985** | 1.30E-28 | **6.84E-27** |
| ENSG00000107819.17 | **SFXN3** | 81855 | protein_coding | **346.997** | 1.441 | 0.434 | **2.715** | 1.54E-28 | **8.05E-27** |
| ENSG00000139734.20 | **DIAPH3** | 81624 | protein_coding | **127.977** | -2.906 | -0.875 | **0.133** | 1.59E-28 | **8.30E-27** |
| ENSG00000233276.10 | **GPX1** | 2876 | protein_coding | **2799.269** | -1.085 | -0.327 | **0.471** | 2.06E-28 | **1.08E-26** |

| | | | | | | | | | |
|---|---|---|---|---|---|---|---|---|---|
| ENSG00000099901.20 | **RANBP1** | 5902 | protein_coding | **1808.575** | -1.215 | -0.366 | **0.431** | 2.25E-28 | **1.17E-26** |
| ENSG00000128944.15 | **KNSTRN** | 90417 | protein_coding | **469.419** | -1.310 | -0.394 | **0.403** | 2.44E-28 | **1.27E-26** |
| ENSG00000101057.18 | **MYBL2** | 4605 | protein_coding | **2191.312** | -2.205 | -0.664 | **0.217** | 3.04E-28 | **1.57E-26** |
| ENSG00000076382.19 | **SPAG5** | 10615 | protein_coding | **1274.083** | -2.351 | -0.708 | **0.196** | 3.55E-28 | **1.83E-26** |
| ENSG00000174371.19 | **EXO1** | 9156 | protein_coding | **239.632** | -2.275 | -0.685 | **0.207** | 3.77E-28 | **1.94E-26** |
| ENSG00000184988.10 | **TMEM106/** | 113277 | protein_coding | **189.205** | -1.462 | -0.440 | **0.363** | 5.74E-28 | **2.94E-26** |
| ENSG00000140092.18 | **FBLN5** | 10516 | protein_coding | **70.445** | 2.747 | 0.827 | **6.715** | 6.30E-28 | **3.22E-26** |
| ENSG00000133134.14 | **BEX2** | 84707 | protein_coding | **194.817** | 2.390 | 0.719 | **5.240** | 6.86E-28 | **3.50E-26** |
| ENSG00000120437.11 | **ACAT2** | 39 | protein_coding | **1348.452** | -1.935 | -0.582 | **0.262** | 7.42E-28 | **3.77E-26** |
| ENSG00000130775.18 | **THEMIS2** | 9473 | protein_coding | **4821.222** | 1.242 | 0.374 | **2.365** | 7.83E-28 | **3.97E-26** |
| ENSG00000277283.2 | **RAB35-AS1** | 12713886 | lncRNA | **218.536** | 1.275 | 0.384 | **2.420** | 8.03E-28 | **4.06E-26** |
| ENSG00000231233.4 | **CFAP58-DT** | 10050586 | lncRNA | **45.317** | 3.846 | 1.158 | **14.378** | 8.45E-28 | **4.26E-26** |
| ENSG00000160188.12 | **RSPH1** | 89765 | protein_coding | **77.805** | 2.771 | 0.834 | **6.827** | 9.58E-28 | **4.82E-26** |
| ENSG00000151503.16 | **NCAPD3** | 23310 | protein_coding | **797.186** | -1.537 | -0.463 | **0.345** | 1.14E-27 | **5.74E-26** |
| ENSG00000275395.8 | **FCGBP** | 8857 | protein_coding | **124.450** | 2.049 | 0.617 | **4.137** | 1.34E-27 | **6.69E-26** |
| ENSG00000271122.2 | **HERPUD2-/** | 10193008 | lncRNA | **231.374** | 1.497 | 0.451 | **2.822** | 1.39E-27 | **6.92E-26** |
| ENSG00000160185.16 | **UBASH3A** | 53347 | protein_coding | **90.426** | 2.384 | 0.718 | **5.220** | 1.41E-27 | **7.01E-26** |
| ENSG00000293268.2 | **ENSG0000** | 12490626 | protein_coding | **505.929** | 1.234 | 0.371 | **2.351** | 1.47E-27 | **7.31E-26** |
| ENSG00000112984.14 | **KIF20A** | 10112 | protein_coding | **457.102** | -2.703 | -0.814 | **0.154** | 1.51E-27 | **7.45E-26** |
| ENSG00000108175.21 | **ZMIZ1** | 57178 | protein_coding | **986.469** | -1.212 | -0.365 | **0.432** | 1.51E-27 | **7.45E-26** |
| ENSG00000085733.18 | **CTTN** | 2017 | protein_coding | **1526.740** | 1.749 | 0.527 | **3.362** | 1.53E-27 | **7.55E-26** |
| ENSG00000166803.16 | **PCLAF** | 9768 | protein_coding | **498.803** | -2.236 | -0.673 | **0.212** | 1.54E-27 | **7.59E-26** |
| ENSG00000112992.20 | **NNT** | 23530 | protein_coding | **637.797** | -1.264 | -0.380 | **0.417** | 1.61E-27 | **7.88E-26** |
| ENSG00000165891.18 | **E2F7** | 144455 | protein_coding | **190.483** | -2.030 | -0.611 | **0.245** | 1.63E-27 | **7.99E-26** |
| ENSG00000068028.20 | **RASSF1** | 11186 | protein_coding | **660.225** | -1.381 | -0.416 | **0.384** | 1.92E-27 | **9.37E-26** |
| ENSG00000103034.17 | **NDRG4** | 65009 | protein_coding | **145.040** | 3.523 | 1.060 | **11.492** | 2.15E-27 | **1.05E-25** |
| ENSG00000263004.4 | **AKAP1-DT** | 11913990 | lncRNA | **228.051** | 1.609 | 0.484 | **3.050** | 3.44E-27 | **1.67E-25** |
| ENSG00000101210.17 | **EEF1A2** | 1917 | protein_coding | **305.993** | 3.089 | 0.930 | **8.511** | 3.65E-27 | **1.76E-25** |
| ENSG00000174547.15 | **MRPL11** | 65003 | protein_coding | **880.724** | -1.119 | -0.337 | **0.461** | 5.12E-27 | **2.47E-25** |
| ENSG00000111615.16 | **KRR1** | 11103 | protein_coding | **625.699** | -1.070 | -0.322 | **0.476** | 6.67E-27 | **3.21E-25** |
| ENSG00000105479.18 | **ODAD1** | 93233 | protein_coding | **87.088** | 2.538 | 0.764 | **5.808** | 6.71E-27 | **3.22E-25** |

| | | | | | | | | | |
|---|---|---|---|---|---|---|---|---|---|
| ENSG00000196517.15 | **SLC6A9** | 6536 | protein_coding | **147.614** | -2.570 | -0.774 | **0.168** | 7.53E-27 | **3.61E-25** |
| ENSG00000213886.6 | **UBD** | 10537 | protein_coding | **523.024** | 2.869 | 0.864 | **7.305** | 7.57E-27 | **3.62E-25** |
| ENSG00000167748.12 | **KLK1** | 3816 | protein_coding | **1016.114** | -1.756 | -0.529 | **0.296** | 8.40E-27 | **4.00E-25** |
| ENSG00000181513.17 | **ACBD4** | 79777 | protein_coding | **850.720** | 1.412 | 0.425 | **2.661** | 8.93E-27 | **4.25E-25** |
| ENSG00000147852.19 | **VLDLR** | 7436 | protein_coding | **322.153** | -1.115 | -0.336 | **0.462** | 1.09E-26 | **5.16E-25** |
| ENSG00000137807.18 | **KIF23** | 9493 | protein_coding | **390.823** | -1.770 | -0.533 | **0.293** | 1.17E-26 | **5.53E-25** |
| ENSG00000107331.20 | **ABCA2** | 20 | protein_coding | **476.846** | 2.220 | 0.668 | **4.658** | 1.21E-26 | **5.72E-25** |
| ENSG00000111671.12 | **SPSB2** | 84727 | protein_coding | **176.476** | 1.774 | 0.534 | **3.420** | 1.33E-26 | **6.24E-25** |
| ENSG00000105696.11 | **TMEM59L** | 25789 | protein_coding | **47.426** | 4.391 | 1.322 | **20.988** | 1.33E-26 | **6.24E-25** |
| ENSG00000101986.14 | **ABCD1** | 215 | protein_coding | **331.848** | 1.508 | 0.454 | **2.845** | 1.48E-26 | **6.95E-25** |
| ENSG00000182179.15 | **UBA7** | 7318 | protein_coding | **2299.324** | -1.386 | -0.417 | **0.383** | 1.51E-26 | **7.04E-25** |
| ENSG00000171811.15 | **CFAP46** | 54777 | protein_coding | **89.920** | 3.973 | 1.196 | **15.709** | 1.51E-26 | **7.05E-25** |
| ENSG00000221946.10 | **FXYD7** | 53822 | protein_coding | **47.711** | 5.159 | 1.553 | **35.740** | 1.76E-26 | **8.16E-25** |
| ENSG00000101460.15 | **MAP1LC3A** | 84557 | protein_coding | **76.804** | 2.250 | 0.677 | **4.756** | 1.90E-26 | **8.82E-25** |
| ENSG00000125170.13 | **DOK4** | 55715 | protein_coding | **310.258** | 2.762 | 0.832 | **6.785** | 1.92E-26 | **8.89E-25** |
| ENSG00000101412.15 | **E2F1** | 1869 | protein_coding | **523.811** | -1.828 | -0.550 | **0.282** | 1.93E-26 | **8.92E-25** |
| ENSG00000077152.14 | **UBE2T** | 29089 | protein_coding | **529.618** | -1.534 | -0.462 | **0.345** | 2.23E-26 | **1.03E-24** |
| ENSG00000088280.23 | **ASAP3** | 55616 | protein_coding | **56.156** | 5.290 | 1.593 | **39.132** | 2.42E-26 | **1.11E-24** |
| ENSG00000073111.17 | **MCM2** | 4171 | protein_coding | **1258.908** | -1.321 | -0.398 | **0.400** | 2.43E-26 | **1.12E-24** |
| ENSG00000162430.19 | **SELENON** | 57190 | protein_coding | **291.760** | 2.006 | 0.604 | **4.017** | 2.60E-26 | **1.19E-24** |
| ENSG00000083720.15 | **OXCT1** | 5019 | protein_coding | **754.286** | -1.383 | -0.416 | **0.383** | 2.77E-26 | **1.26E-24** |
| ENSG00000171368.14 | **TPPP** | 11076 | protein_coding | **86.654** | 4.045 | 1.218 | **16.504** | 2.82E-26 | **1.28E-24** |
| ENSG00000198331.13 | **HYLS1** | 219844 | protein_coding | **178.153** | -1.592 | -0.479 | **0.332** | 2.83E-26 | **1.28E-24** |
| ENSG00000247315.4 | **ZCCHC3** | 85364 | protein_coding | **685.607** | -0.884 | -0.266 | **0.542** | 2.86E-26 | **1.30E-24** |
| ENSG00000179091.7 | **CYC1** | 1537 | protein_coding | **1662.207** | -1.192 | -0.359 | **0.438** | 2.93E-26 | **1.33E-24** |
| ENSG00000122515.18 | **ZMIZ2** | 83637 | protein_coding | **9427.026** | 1.038 | 0.312 | **2.053** | 3.01E-26 | **1.36E-24** |
| ENSG00000117643.18 | **MAN1C1** | 57134 | protein_coding | **111.923** | 2.733 | 0.823 | **6.647** | 3.29E-26 | **1.48E-24** |
| ENSG00000149547.17 | **EI24** | 9538 | protein_coding | **746.660** | -1.008 | -0.303 | **0.497** | 3.46E-26 | **1.55E-24** |
| ENSG00000161888.13 | **SPC24** | 147841 | protein_coding | **428.155** | -2.398 | -0.722 | **0.190** | 3.58E-26 | **1.60E-24** |
| ENSG00000187535.16 | **IFT140** | 9742 | protein_coding | **291.402** | 1.249 | 0.376 | **2.377** | 3.69E-26 | **1.65E-24** |
| ENSG00000286403.1 | **ENSG00000286403** | | lncRNA | **53.806** | 4.934 | 1.485 | **30.574** | 3.73E-26 | **1.66E-24** |

| | | | | | | | | | |
|---|---|---|---|---|---|---|---|---|---|
| ENSG00000140451.15 | **PIF1** | 80119 | protein_coding | **1053.613** | -1.430 | -0.430 | **0.371** | 4.39E-26 | **1.95E-24** |
| ENSG00000141012.15 | **GALNS** | 2588 | protein_coding | **431.778** | 1.375 | 0.414 | **2.594** | 4.40E-26 | **1.95E-24** |
| ENSG00000042286.17 | **AIFM2** | 84883 | protein_coding | **687.415** | -1.261 | -0.379 | **0.417** | 4.65E-26 | **2.06E-24** |
| ENSG00000259316.12 | **ENSG00000259316** | | protein_coding | **194.530** | -2.292 | -0.690 | **0.204** | 5.36E-26 | **2.37E-24** |
| ENSG00000124098.11 | **FAM210B** | 116151 | protein_coding | **112.779** | 1.863 | 0.561 | **3.639** | 5.96E-26 | **2.62E-24** |
| ENSG00000138658.19 | **ZGRF1** | 55345 | protein_coding | **395.943** | -1.474 | -0.444 | **0.360** | 5.98E-26 | **2.63E-24** |
| ENSG00000197238.5 | **H4C11** | 8363 | protein_coding | **193.145** | 1.697 | 0.511 | **3.243** | 6.18E-26 | **2.71E-24** |
| ENSG00000248554.2 | **C5orf34-AS1** | | lncRNA | **101.963** | 1.802 | 0.542 | **3.486** | 7.72E-26 | **3.37E-24** |
| ENSG00000053372.7 | **MRTO4** | 51154 | protein_coding | **839.546** | -1.221 | -0.367 | **0.429** | 8.85E-26 | **3.86E-24** |
| ENSG00000196668.5 | **LINC00173** | 10028756[illegible] | lncRNA | **408.133** | 1.802 | 0.542 | **3.487** | 8.97E-26 | **3.91E-24** |
| ENSG00000130653.18 | **PNPLA7** | 375775 | protein_coding | **343.912** | 2.079 | 0.626 | **4.226** | 9.23E-26 | **4.01E-24** |
| ENSG00000115163.17 | **CENPA** | 1058 | protein_coding | **195.698** | -2.259 | -0.680 | **0.209** | 1.03E-25 | **4.45E-24** |
| ENSG00000124507.13 | **PACSIN1** | 29993 | protein_coding | **47.200** | 3.153 | 0.949 | **8.894** | 1.07E-25 | **4.63E-24** |
| ENSG00000145050.20 | **MANF** | 7873 | protein_coding | **1065.141** | -1.359 | -0.409 | **0.390** | 1.08E-25 | **4.67E-24** |
| ENSG00000111752.13 | **PHC1** | 1911 | protein_coding | **564.554** | 1.600 | 0.482 | **3.032** | 1.12E-25 | **4.81E-24** |
| ENSG00000204314.14 | **PRRT1** | 80863 | protein_coding | **43.845** | 3.986 | 1.200 | **15.843** | 1.22E-25 | **5.22E-24** |
| ENSG00000100526.23 | **CDKN3** | 1033 | protein_coding | **398.684** | -2.089 | -0.629 | **0.235** | 1.22E-25 | **5.23E-24** |
| ENSG00000189057.13 | **FAM111B** | 374393 | protein_coding | **333.130** | -1.386 | -0.417 | **0.383** | 1.24E-25 | **5.28E-24** |
| ENSG00000127946.19 | **HIP1** | 3092 | protein_coding | **301.692** | 1.381 | 0.416 | **2.604** | 1.36E-25 | **5.82E-24** |
| ENSG00000143624.16 | **INTS3** | 65123 | protein_coding | **5311.821** | 0.951 | 0.286 | **1.934** | 1.39E-25 | **5.91E-24** |
| ENSG00000105427.11 | **CNFN** | 84518 | protein_coding | **34.679** | 4.729 | 1.424 | **26.520** | 1.43E-25 | **6.07E-24** |
| ENSG00000066294.17 | **CD84** | 8832 | protein_coding | **202.047** | -1.890 | -0.569 | **0.270** | 1.46E-25 | **6.18E-24** |
| ENSG00000024526.19 | **DEPDC1** | 55635 | protein_coding | **74.155** | -2.696 | -0.812 | **0.154** | 1.48E-25 | **6.26E-24** |
| ENSG00000171960.14 | **PPIH** | 10465 | protein_coding | **960.476** | -1.311 | -0.395 | **0.403** | 1.67E-25 | **7.05E-24** |
| ENSG00000255375.4 | **ENSG0000[illegible]** | 10798432[illegible] | lncRNA | **322.345** | 3.093 | 0.931 | **8.533** | 1.70E-25 | **7.15E-24** |
| ENSG00000297561.1 | **ENSG00000297561** | | lncRNA | **70.176** | 3.724 | 1.121 | **13.216** | 1.70E-25 | **7.15E-24** |
| ENSG00000166881.12 | **NEMP1** | 23306 | protein_coding | **724.014** | -1.171 | -0.352 | **0.444** | 1.77E-25 | **7.41E-24** |
| ENSG00000067992.19 | **PDK3** | 5165 | protein_coding | **942.227** | -1.497 | -0.451 | **0.354** | 1.82E-25 | **7.60E-24** |
| ENSG00000093009.13 | **CDC45** | 8318 | protein_coding | **452.748** | -2.262 | -0.681 | **0.208** | 1.85E-25 | **7.70E-24** |
| ENSG00000139269.5 | **INHBE** | 83729 | protein_coding | **109.543** | -2.764 | -0.832 | **0.147** | 1.87E-25 | **7.79E-24** |
| ENSG00000144231.13 | **POLR2D** | 5433 | protein_coding | **754.889** | -1.120 | -0.337 | **0.460** | 2.00E-25 | **8.32E-24** |

| | | | | | | | | | |
|---|---|---|---|---|---|---|---|---|---|
| ENSG00000188290.13 | **HES4** | 57801 | protein_coding | **396.588** | 2.065 | 0.621 | **4.183** | 2.02E-25 | **8.35E-24** |
| ENSG00000257242.10 | **LINC01619** | 256021 | lncRNA | **830.458** | 2.732 | 0.822 | **6.643** | 2.04E-25 | **8.45E-24** |
| ENSG00000247121.9 | **ENSG00000247121** | | lncRNA | **123.562** | 1.991 | 0.599 | **3.976** | 2.26E-25 | **9.30E-24** |
| ENSG00000293185.2 | **TPM3P9** | 147804 | lncRNA | **141.209** | -1.696 | -0.511 | **0.309** | 2.27E-25 | **9.34E-24** |
| ENSG00000137193.16 | **PIM1** | 5292 | protein_coding | **967.778** | 1.124 | 0.338 | **2.180** | 2.29E-25 | **9.41E-24** |
| ENSG00000097046.15 | **CDC7** | 8317 | protein_coding | **325.347** | -1.205 | -0.363 | **0.434** | 2.43E-25 | **9.96E-24** |
| ENSG00000169660.19 | **HEXD** | 284004 | protein_coding | **1021.424** | 1.134 | 0.341 | **2.194** | 2.44E-25 | **9.96E-24** |
| ENSG00000136010.16 | **ALDH1L2** | 160428 | protein_coding | **514.177** | -2.199 | -0.662 | **0.218** | 2.47E-25 | **1.01E-23** |
| ENSG00000143882.14 | **ATP6V1C2** | 245973 | protein_coding | **87.303** | 3.240 | 0.975 | **9.447** | 2.51E-25 | **1.02E-23** |
| ENSG00000130598.18 | **TNNI2** | 7136 | protein_coding | **53.196** | 2.994 | 0.901 | **7.969** | 2.53E-25 | **1.03E-23** |
| ENSG00000132780.19 | **NASP** | 4678 | protein_coding | **3508.704** | -1.092 | -0.329 | **0.469** | 2.54E-25 | **1.03E-23** |
| ENSG00000100307.15 | **CBX7** | 23492 | protein_coding | **789.109** | 1.539 | 0.463 | **2.906** | 2.67E-25 | **1.08E-23** |
| ENSG00000152795.20 | **HNRNPDL** | 9987 | protein_coding | **4698.110** | -0.801 | -0.241 | **0.574** | 2.87E-25 | **1.16E-23** |
| ENSG00000291179.2 | **LCMT1P1** | 554206 | lncRNA | **135.536** | 2.285 | 0.688 | **4.873** | 2.96E-25 | **1.19E-23** |
| ENSG00000182578.16 | **CSF1R** | 1436 | protein_coding | **93.496** | 3.600 | 1.084 | **12.130** | 3.19E-25 | **1.28E-23** |
| ENSG00000143375.17 | **CGN** | 57530 | protein_coding | **80.970** | 2.660 | 0.801 | **6.318** | 3.40E-25 | **1.36E-23** |
| ENSG00000136811.18 | **ODF2** | 4957 | protein_coding | **996.068** | -0.889 | -0.268 | **0.540** | 3.47E-25 | **1.39E-23** |
| ENSG00000005882.14 | **PDK2** | 5164 | protein_coding | **171.845** | 1.614 | 0.486 | **3.060** | 3.48E-25 | **1.39E-23** |
| ENSG00000259803.9 | **SLC22A31** | 146429 | protein_coding | **118.707** | 3.506 | 1.055 | **11.359** | 3.57E-25 | **1.42E-23** |
| ENSG00000166165.16 | **CKB** | 1152 | protein_coding | **505.093** | 1.996 | 0.601 | **3.989** | 3.65E-25 | **1.45E-23** |
| ENSG00000129255.18 | **MPDU1** | 9526 | protein_coding | **622.963** | -1.448 | -0.436 | **0.367** | 3.70E-25 | **1.47E-23** |
| ENSG00000169508.9 | **GPR183** | 1880 | protein_coding | **362.352** | -1.798 | -0.541 | **0.288** | 4.11E-25 | **1.63E-23** |
| ENSG00000147274.16 | **RBMX** | 27316 | protein_coding | **3030.510** | -1.056 | -0.318 | **0.481** | 4.13E-25 | **1.63E-23** |
| ENSG00000198393.10 | **ZNF26** | 7574 | protein_coding | **417.908** | -1.237 | -0.372 | **0.424** | 4.36E-25 | **1.72E-23** |
| ENSG00000162572.22 | **SCNN1D** | 6339 | protein_coding | **234.864** | 2.009 | 0.605 | **4.024** | 4.41E-25 | **1.74E-23** |
| ENSG00000115257.17 | **PCSK4** | 54760 | protein_coding | **202.332** | 2.236 | 0.673 | **4.711** | 5.21E-25 | **2.05E-23** |
| ENSG00000198856.15 | **OSTC** | 58505 | protein_coding | **845.843** | -1.019 | -0.307 | **0.494** | 5.25E-25 | **2.06E-23** |
| ENSG00000164620.11 | **RELL2** | 285613 | protein_coding | **548.312** | 1.376 | 0.414 | **2.596** | 5.27E-25 | **2.06E-23** |
| ENSG00000235706.10 | **DICER1-AS** | 400242 | lncRNA | **473.017** | 1.501 | 0.452 | **2.830** | 5.34E-25 | **2.09E-23** |
| ENSG00000158286.15 | **RNF207** | 388591 | protein_coding | **451.400** | 1.536 | 0.462 | **2.900** | 5.55E-25 | **2.16E-23** |
| ENSG00000287539.2 | **ENSG00000287539** | | lncRNA | **89.615** | 2.228 | 0.671 | **4.685** | 6.08E-25 | **2.36E-23** |

| | | | | | | | | | |
|---|---|---|---|---|---|---|---|---|---|
| ENSG00000115233.14 | **PSMD14** | 10213 | protein_coding | **1084.303** | -0.964 | -0.290 | **0.513** | 6.58E-25 | **2.55E-23** |
| ENSG00000119242.11 | **CCDC92** | 80212 | protein_coding | **383.289** | 1.357 | 0.409 | **2.562** | 6.74E-25 | **2.61E-23** |
| ENSG00000151725.14 | **CENPU** | 79682 | protein_coding | **560.227** | -1.684 | -0.507 | **0.311** | 7.17E-25 | **2.77E-23** |
| ENSG00000172817.6 | **CYP7B1** | 9420 | protein_coding | **172.021** | -2.732 | -0.823 | **0.150** | 7.21E-25 | **2.78E-23** |
| ENSG00000006634.10 | **DBF4** | 10926 | protein_coding | **318.095** | -1.486 | -0.447 | **0.357** | 7.58E-25 | **2.92E-23** |
| ENSG00000073605.21 | **GSDMB** | 55876 | protein_coding | **2885.375** | 1.354 | 0.408 | **2.556** | 7.74E-25 | **2.97E-23** |
| ENSG00000168237.20 | **GLYCTK** | 132158 | protein_coding | **136.967** | -2.107 | -0.634 | **0.232** | 8.87E-25 | **3.40E-23** |
| ENSG00000143126.8 | **CELSR2** | 1952 | protein_coding | **103.150** | 1.873 | 0.564 | **3.663** | 8.88E-25 | **3.40E-23** |
| ENSG00000203760.11 | **CENPW** | 387103 | protein_coding | **384.305** | -2.316 | -0.697 | **0.201** | 9.93E-25 | **3.79E-23** |
| ENSG00000204257.17 | **HLA-DMA** | 3108 | protein_coding | **7162.553** | 1.208 | 0.364 | **2.311** | 1.02E-24 | **3.88E-23** |
| ENSG00000114942.17 | **EEF1B2** | 1933 | protein_coding | **9264.130** | -0.901 | -0.271 | **0.536** | 1.07E-24 | **4.07E-23** |
| ENSG00000178295.17 | **GEN1** | 348654 | protein_coding | **548.269** | -1.547 | -0.466 | **0.342** | 1.20E-24 | **4.57E-23** |
| ENSG00000128040.14 | **SPINK2** | 6691 | protein_coding | **58.507** | 5.144 | 1.548 | **35.349** | 1.33E-24 | **5.03E-23** |
| ENSG00000143222.14 | **UFC1** | 51506 | protein_coding | **1288.220** | -1.065 | -0.321 | **0.478** | 1.37E-24 | **5.19E-23** |
| ENSG00000095383.21 | **TBC1D2** | 55357 | protein_coding | **48.720** | 2.765 | 0.832 | **6.799** | 1.38E-24 | **5.19E-23** |
| ENSG00000087077.16 | **TRIP6** | 7205 | protein_coding | **600.798** | -1.520 | -0.458 | **0.349** | 1.40E-24 | **5.28E-23** |
| ENSG00000182796.17 | **TMEM198B** | | transcribed_unita | **65.553** | 2.334 | 0.703 | **5.044** | 1.46E-24 | **5.49E-23** |
| ENSG00000228594.4 | **FNDC10** | 643988 | protein_coding | **131.568** | 2.271 | 0.684 | **4.828** | 1.50E-24 | **5.61E-23** |
| ENSG00000133119.15 | **RFC3** | 5983 | protein_coding | **388.581** | -1.650 | -0.497 | **0.319** | 1.63E-24 | **6.10E-23** |
| ENSG00000143401.17 | **ANP32E** | 81611 | protein_coding | **982.532** | -0.930 | -0.280 | **0.525** | 1.90E-24 | **7.09E-23** |
| ENSG00000204899.7 | **MZT1** | 440145 | protein_coding | **241.414** | -1.512 | -0.455 | **0.351** | 1.98E-24 | **7.40E-23** |
| ENSG00000154134.16 | **ROBO3** | 64221 | protein_coding | **181.801** | 1.859 | 0.559 | **3.627** | 1.99E-24 | **7.42E-23** |
| ENSG00000074855.12 | **ANO8** | 57719 | protein_coding | **221.038** | 1.623 | 0.489 | **3.080** | 2.08E-24 | **7.73E-23** |
| ENSG00000006451.10 | **RALA** | 5898 | protein_coding | **837.693** | -0.863 | -0.260 | **0.550** | 2.14E-24 | **7.94E-23** |
| ENSG00000183196.12 | **CHST6** | 4166 | protein_coding | **74.816** | 2.037 | 0.613 | **4.103** | 2.19E-24 | **8.12E-23** |
| ENSG00000035862.15 | **TIMP2** | 7077 | protein_coding | **65.106** | 3.380 | 1.017 | **10.408** | 2.31E-24 | **8.54E-23** |
| ENSG00000092470.14 | **WDR76** | 79968 | protein_coding | **294.440** | -1.281 | -0.386 | **0.412** | 2.63E-24 | **9.68E-23** |
| ENSG00000135451.15 | **TROAP** | 10024 | protein_coding | **725.963** | -2.364 | -0.712 | **0.194** | 2.81E-24 | **1.03E-22** |
| ENSG00000174326.17 | **SLC16A11** | 162515 | protein_coding | **103.893** | 2.469 | 0.743 | **5.537** | 3.14E-24 | **1.15E-22** |
| ENSG00000010671.20 | **BTK** | 695 | protein_coding | **1919.368** | -1.079 | -0.325 | **0.474** | 3.30E-24 | **1.21E-22** |
| ENSG00000177084.21 | **POLE** | 5426 | protein_coding | **1779.347** | -1.438 | -0.433 | **0.369** | 3.36E-24 | **1.23E-22** |

| | | | | | | | | | |
|---|---|---|---|---|---|---|---|---|---|
| ENSG00000198832.12 | **SELENOM** | 140606 | protein_coding | **214.317** | 1.792 | 0.539 | **3.463** | 3.47E-24 | **1.27E-22** |
| ENSG00000126215.16 | **XRCC3** | 7517 | protein_coding | **714.322** | -1.708 | -0.514 | **0.306** | 3.66E-24 | **1.33E-22** |
| ENSG00000234771.3 | **SLC25A25-AS1** | | lncRNA | **730.328** | 1.156 | 0.348 | **2.229** | 3.69E-24 | **1.34E-22** |
| ENSG00000137070.21 | **IL11RA** | 3590 | protein_coding | **656.196** | 1.341 | 0.404 | **2.533** | 4.16E-24 | **1.51E-22** |
| ENSG00000065328.19 | **MCM10** | 55388 | protein_coding | **162.336** | -2.100 | -0.632 | **0.233** | 4.80E-24 | **1.74E-22** |
| ENSG00000162063.15 | **CCNF** | 899 | protein_coding | **390.761** | -1.425 | -0.429 | **0.372** | 4.88E-24 | **1.76E-22** |
| ENSG00000266714.11 | **MYO15B** | 80022 | protein_coding | **209.227** | 2.248 | 0.677 | **4.749** | 4.98E-24 | **1.80E-22** |
| ENSG00000125319.17 | **HROB** | 78995 | protein_coding | **196.149** | -2.054 | -0.618 | **0.241** | 5.24E-24 | **1.89E-22** |
| ENSG00000117115.15 | **PADI2** | 11240 | protein_coding | **109.246** | 1.822 | 0.548 | **3.535** | 5.69E-24 | **2.04E-22** |
| ENSG00000164032.14 | **H2AZ1** | 3015 | protein_coding | **4225.788** | -0.974 | -0.293 | **0.509** | 5.78E-24 | **2.07E-22** |
| ENSG00000167992.16 | **VWCE** | 220001 | protein_coding | **198.907** | -2.256 | -0.679 | **0.209** | 5.84E-24 | **2.09E-22** |
| ENSG00000117009.14 | **KMO** | 8564 | protein_coding | **129.351** | -2.827 | -0.851 | **0.141** | 6.56E-24 | **2.35E-22** |
| ENSG00000148834.15 | **GSTO1** | 9446 | protein_coding | **1548.153** | -1.080 | -0.325 | **0.473** | 6.72E-24 | **2.40E-22** |
| ENSG00000272325.4 | **NUDT3** | 11165 | protein_coding | **1833.292** | -1.444 | -0.435 | **0.367** | 6.81E-24 | **2.42E-22** |
| ENSG00000167695.16 | **TLCD3A** | 79850 | protein_coding | **224.403** | -1.506 | -0.453 | **0.352** | 6.84E-24 | **2.43E-22** |
| ENSG00000188917.17 | **TRMT2B** | 79979 | protein_coding | **189.477** | -1.316 | -0.396 | **0.402** | 6.97E-24 | **2.47E-22** |
| ENSG00000166578.12 | **DRC10** | 115811 | protein_coding | **62.260** | 2.596 | 0.781 | **6.046** | 7.59E-24 | **2.69E-22** |
| ENSG00000176533.15 | **GNG7** | 2788 | protein_coding | **2630.455** | 0.988 | 0.297 | **1.983** | 7.61E-24 | **2.69E-22** |
| ENSG00000169607.14 | **CKAP2L** | 150468 | protein_coding | **162.655** | -2.109 | -0.635 | **0.232** | 7.73E-24 | **2.73E-22** |
| ENSG00000160471.14 | **COX6B2** | 125965 | protein_coding | **54.701** | 4.833 | 1.455 | **28.511** | 8.29E-24 | **2.92E-22** |
| ENSG00000090447.14 | **TFAP4** | 7023 | protein_coding | **334.097** | -1.311 | -0.395 | **0.403** | 8.44E-24 | **2.96E-22** |
| ENSG00000235688.4 | **SNTG2-AS1** | | lncRNA | **240.737** | 2.335 | 0.703 | **5.044** | 8.44E-24 | **2.96E-22** |
| ENSG00000060558.5 | **GNA15** | 2769 | protein_coding | **378.481** | -1.817 | -0.547 | **0.284** | 8.55E-24 | **3.00E-22** |
| ENSG00000172244.11 | **C5orf34** | 375444 | protein_coding | **74.463** | -2.326 | -0.700 | **0.199** | 8.69E-24 | **3.04E-22** |
| ENSG00000205116.4 | **TMEM278** | 643965 | protein_coding | **58.278** | 4.055 | 1.221 | **16.624** | 9.58E-24 | **3.34E-22** |
| ENSG00000148341.20 | **SH3GLB2** | 56904 | protein_coding | **1947.769** | 0.729 | 0.219 | **1.658** | 1.02E-23 | **3.56E-22** |
| ENSG00000135605.15 | **TEC** | 7006 | protein_coding | **806.261** | -1.311 | -0.395 | **0.403** | 1.06E-23 | **3.70E-22** |
| ENSG00000154920.17 | **EME1** | 146956 | protein_coding | **384.297** | -2.138 | -0.644 | **0.227** | 1.07E-23 | **3.71E-22** |
| ENSG00000110321.20 | **EIF4G2** | 1982 | protein_coding | **3615.591** | -1.338 | -0.403 | **0.395** | 1.09E-23 | **3.76E-22** |
| ENSG00000309132.1 | **ENSG00000309132** | | lncRNA | **62.894** | 3.162 | 0.952 | **8.950** | 1.11E-23 | **3.84E-22** |
| ENSG00000154640.17 | **BTG3** | 10950 | protein_coding | **444.620** | -1.154 | -0.347 | **0.449** | 1.11E-23 | **3.84E-22** |

| | | | | | | | | | |
|---|---|---|---|---|---|---|---|---|---|
| ENSG00000106733.23 | **NMRK1** | 54981 | protein_coding | **528.984** | 1.123 | 0.338 | **2.178** | 1.16E-23 | **3.99E-22** |
| ENSG00000166816.17 | **LDHD** | 197257 | protein_coding | **60.193** | 4.523 | 1.361 | **22.983** | 1.33E-23 | **4.58E-22** |
| ENSG00000124145.8 | **SDC4** | 6385 | protein_coding | **300.913** | 2.034 | 0.612 | **4.096** | 1.43E-23 | **4.91E-22** |
| ENSG00000196456.15 | **ZNF775** | 285971 | protein_coding | **476.880** | 1.063 | 0.320 | **2.089** | 1.51E-23 | **5.18E-22** |
| ENSG00000177679.17 | **SRRM3** | 222183 | protein_coding | **28.142** | 3.940 | 1.186 | **15.346** | 1.62E-23 | **5.56E-22** |
| ENSG00000157379.16 | **DHRS1** | 115817 | protein_coding | **609.873** | 1.022 | 0.308 | **2.031** | 1.63E-23 | **5.58E-22** |
| ENSG00000187109.17 | **NAP1L1** | 4673 | protein_coding | **11323.522** | -0.942 | -0.284 | **0.520** | 1.75E-23 | **5.98E-22** |
| ENSG00000177181.17 | **RIMKLA** | 284716 | protein_coding | **204.716** | -1.559 | -0.469 | **0.339** | 1.80E-23 | **6.12E-22** |
| ENSG00000288919.2 | **ENSG00000288919** | | lncRNA | **86.923** | 2.385 | 0.718 | **5.222** | 1.85E-23 | **6.27E-22** |
| ENSG00000133895.21 | **MEN1** | 4221 | protein_coding | **1548.664** | 0.874 | 0.263 | **1.832** | 1.91E-23 | **6.47E-22** |
| ENSG00000173559.17 | **NABP1** | 64859 | protein_coding | **1223.373** | 1.760 | 0.530 | **3.386** | 2.16E-23 | **7.30E-22** |
| ENSG00000138172.13 | **CALHM2** | 51063 | protein_coding | **163.598** | -1.608 | -0.484 | **0.328** | 2.41E-23 | **8.13E-22** |
| ENSG00000196459.17 | **TRAPPC2** | 6399 | protein_coding | **552.129** | 0.950 | 0.286 | **1.932** | 2.60E-23 | **8.76E-22** |
| ENSG00000204227.7 | **RING1** | 6015 | protein_coding | **1056.323** | 0.861 | 0.259 | **1.817** | 2.66E-23 | **8.96E-22** |
| ENSG00000196155.15 | **PLEKHG4** | 25894 | protein_coding | **159.846** | 2.502 | 0.753 | **5.665** | 2.67E-23 | **8.98E-22** |
| ENSG00000144580.17 | **CNOT9** | 9125 | protein_coding | **1543.194** | -0.735 | -0.221 | **0.601** | 2.79E-23 | **9.37E-22** |
| ENSG00000099308.15 | **MAST3** | 23031 | protein_coding | **1577.073** | 0.957 | 0.288 | **1.941** | 2.83E-23 | **9.48E-22** |
| ENSG00000175592.11 | **FOSL1** | 8061 | protein_coding | **80.621** | -2.380 | -0.716 | **0.192** | 3.07E-23 | **1.03E-21** |
| ENSG00000136449.18 | **MYCBPAP** | 84073 | protein_coding | **107.267** | 3.155 | 0.950 | **8.905** | 3.27E-23 | **1.09E-21** |
| ENSG00000143499.16 | **SMYD2** | 56950 | protein_coding | **317.511** | -1.148 | -0.345 | **0.451** | 3.29E-23 | **1.10E-21** |
| ENSG00000115935.21 | **WIPF1** | 7456 | protein_coding | **2132.036** | -1.318 | -0.397 | **0.401** | 3.30E-23 | **1.10E-21** |
| ENSG00000185885.19 | **IFITM1** | 8519 | protein_coding | **10070.033** | -1.023 | -0.308 | **0.492** | 3.36E-23 | **1.12E-21** |
| ENSG00000069275.15 | **NUCKS1** | 64710 | protein_coding | **2058.644** | -0.927 | -0.279 | **0.526** | 3.37E-23 | **1.12E-21** |
| ENSG00000158158.14 | **CNNM4** | 26504 | protein_coding | **722.770** | 1.142 | 0.344 | **2.207** | 3.54E-23 | **1.17E-21** |
| ENSG00000138182.17 | **KIF20B** | 9585 | protein_coding | **310.393** | -1.738 | -0.523 | **0.300** | 3.71E-23 | **1.22E-21** |
| ENSG00000105371.12 | **ICAM4** | 3386 | protein_coding | **89.670** | 2.322 | 0.699 | **5.001** | 3.74E-23 | **1.23E-21** |
| ENSG00000100299.20 | **ARSA** | 410 | protein_coding | **1030.335** | 0.899 | 0.271 | **1.865** | 3.81E-23 | **1.25E-21** |
| ENSG00000159433.13 | **STARD9** | 57519 | protein_coding | **1443.799** | 1.027 | 0.309 | **2.038** | 3.93E-23 | **1.29E-21** |
| ENSG00000242114.8 | **MTFP1** | 51537 | protein_coding | **1043.127** | -1.545 | -0.465 | **0.343** | 4.02E-23 | **1.32E-21** |
| ENSG00000184661.16 | **CDCA2** | 157313 | protein_coding | **233.154** | -1.978 | -0.595 | **0.254** | 4.72E-23 | **1.55E-21** |
| ENSG00000062282.17 | **DGAT2** | 84649 | protein_coding | **434.176** | 1.553 | 0.468 | **2.935** | 5.00E-23 | **1.63E-21** |

| | | | | | | | | | |
|---|---|---|---|---|---|---|---|---|---|
| ENSG00000133433.13 | **GSTT2B** | 653689 | protein_coding | **76.257** | 3.038 | 0.914 | **8.213** | 5.12E-23 | **1.67E-21** |
| ENSG00000142224.18 | **IL19** | 29949 | protein_coding | **190.260** | -1.604 | -0.483 | **0.329** | 5.27E-23 | **1.72E-21** |
| ENSG00000167476.12 | **JSRP1** | 126306 | protein_coding | **193.788** | 3.068 | 0.923 | **8.384** | 5.69E-23 | **1.85E-21** |
| ENSG00000167749.12 | **KLK4** | 9622 | protein_coding | **93.823** | 3.318 | 0.999 | **9.972** | 5.71E-23 | **1.85E-21** |
| ENSG00000120254.18 | **MTHFD1L** | 25902 | protein_coding | **617.753** | -1.241 | -0.374 | **0.423** | 7.10E-23 | **2.30E-21** |
| ENSG00000288234.2 | **ENSG00000288234** | | lncRNA | **28.542** | 5.024 | 1.513 | **32.547** | 7.44E-23 | **2.41E-21** |
| ENSG00000102172.18 | **SMS** | 6611 | protein_coding | **1525.745** | -0.832 | -0.250 | **0.562** | 7.45E-23 | **2.41E-21** |
| ENSG00000106976.24 | **DNM1** | 1759 | protein_coding | **152.211** | 2.955 | 0.890 | **7.755** | 7.96E-23 | **2.57E-21** |
| ENSG00000100629.19 | **CEP128** | 145508 | protein_coding | **84.769** | -2.394 | -0.721 | **0.190** | 8.53E-23 | **2.74E-21** |
| ENSG00000120802.17 | **TMPO** | 7112 | protein_coding | **1566.598** | -1.031 | -0.310 | **0.489** | 8.55E-23 | **2.75E-21** |
| ENSG00000253140.3 | **LINC02947** | 10537582 | lncRNA | **139.739** | 1.811 | 0.545 | **3.509** | 8.71E-23 | **2.79E-21** |
| ENSG00000130066.18 | **SAT1** | 6303 | protein_coding | **3926.432** | 1.477 | 0.445 | **2.785** | 9.58E-23 | **3.07E-21** |
| ENSG00000136492.12 | **BRIP1** | 83990 | protein_coding | **145.829** | -1.872 | -0.563 | **0.273** | 9.60E-23 | **3.07E-21** |
| ENSG00000172803.20 | **SNX32** | 254122 | protein_coding | **85.635** | 2.025 | 0.609 | **4.069** | 9.88E-23 | **3.15E-21** |
| ENSG00000161642.20 | **ZNF385A** | 25946 | protein_coding | **617.595** | 1.910 | 0.575 | **3.759** | 1.02E-22 | **3.26E-21** |
| ENSG00000126458.6 | **RRAS** | 6237 | protein_coding | **438.395** | 1.722 | 0.518 | **3.300** | 1.03E-22 | **3.27E-21** |
| ENSG00000272927.2 | **ENSG00000272927** | | lncRNA | **134.164** | 1.605 | 0.483 | **3.042** | 1.07E-22 | **3.38E-21** |
| ENSG00000123600.23 | **METTL8** | 79828 | protein_coding | **598.723** | -1.260 | -0.379 | **0.418** | 1.17E-22 | **3.71E-21** |
| ENSG00000092853.16 | **CLSPN** | 63967 | protein_coding | **187.798** | -1.790 | -0.539 | **0.289** | 1.20E-22 | **3.81E-21** |
| ENSG00000106665.18 | **CLIP2** | 7461 | protein_coding | **2327.940** | 1.166 | 0.351 | **2.245** | 1.21E-22 | **3.81E-21** |
| ENSG00000274180.2 | **NATD1** | 256302 | protein_coding | **141.857** | 2.535 | 0.763 | **5.796** | 1.21E-22 | **3.81E-21** |
| ENSG00000132313.16 | **MRPL35** | 51318 | protein_coding | **627.867** | -1.080 | -0.325 | **0.473** | 1.24E-22 | **3.89E-21** |
| ENSG00000186907.9 | **RTN4RL2** | 349667 | protein_coding | **60.579** | 2.700 | 0.813 | **6.500** | 1.29E-22 | **4.04E-21** |
| ENSG00000167371.23 | **PRRT2** | 112476 | protein_coding | **30.137** | 3.719 | 1.119 | **13.166** | 1.32E-22 | **4.14E-21** |
| ENSG00000197785.16 | **ATAD3A** | 55210 | protein_coding | **724.246** | -0.961 | -0.289 | **0.514** | 1.32E-22 | **4.14E-21** |
| ENSG00000170734.14 | **POLH** | 5429 | protein_coding | **646.284** | -1.035 | -0.312 | **0.488** | 1.35E-22 | **4.21E-21** |
| ENSG00000139194.10 | **RBP5** | 83758 | protein_coding | **47.832** | 4.079 | 1.228 | **16.902** | 1.37E-22 | **4.26E-21** |
| ENSG00000264451.1 | **ENSG00000264451** | | lncRNA | **32.384** | 6.100 | 1.836 | **68.574** | 1.40E-22 | **4.37E-21** |
| ENSG00000182180.16 | **MRPS16** | 51021 | protein_coding | **1222.204** | -1.067 | -0.321 | **0.477** | 1.50E-22 | **4.65E-21** |
| ENSG00000130751.12 | **NPAS1** | 4861 | protein_coding | **68.487** | 2.402 | 0.723 | **5.284** | 1.50E-22 | **4.66E-21** |
| ENSG00000161714.14 | **PLCD3** | 113026 | protein_coding | **62.018** | 2.802 | 0.844 | **6.976** | 1.50E-22 | **4.67E-21** |

| | | | | | | | | | |
|---|---|---|---|---|---|---|---|---|---|
| ENSG00000211584.16 | **SLC48A1** | 55652 | protein_coding | **201.814** | 1.498 | 0.451 | **2.825** | 1.52E-22 | **4.72E-21** |
| ENSG00000136425.16 | **CIB2** | 10518 | protein_coding | **93.490** | 2.446 | 0.736 | **5.449** | 1.91E-22 | **5.90E-21** |
| ENSG00000135363.14 | **LMO2** | 4005 | protein_coding | **361.924** | -1.665 | -0.501 | **0.315** | 2.04E-22 | **6.31E-21** |
| ENSG00000109674.6 | **NEIL3** | 55247 | protein_coding | **50.762** | -2.508 | -0.755 | **0.176** | 2.10E-22 | **6.47E-21** |
| ENSG00000141994.18 | **DUS3L** | 56931 | protein_coding | **332.202** | -1.563 | -0.470 | **0.338** | 2.26E-22 | **6.96E-21** |
| ENSG00000163482.14 | **STK36** | 27148 | protein_coding | **747.237** | 0.804 | 0.242 | **1.746** | 2.35E-22 | **7.21E-21** |
| ENSG00000157916.22 | **RER1** | 11079 | protein_coding | **2206.694** | -0.822 | -0.248 | **0.565** | 2.60E-22 | **7.98E-21** |
| ENSG00000137822.15 | **TUBGCP4** | 27229 | protein_coding | **516.002** | -1.566 | -0.472 | **0.338** | 2.65E-22 | **8.12E-21** |
| ENSG00000229474.10 | **PATL2** | 197135 | protein_coding | **635.981** | 1.329 | 0.400 | **2.512** | 2.87E-22 | **8.77E-21** |
| ENSG00000183527.14 | **PSMG1** | 8624 | protein_coding | **604.963** | -1.157 | -0.348 | **0.448** | 3.40E-22 | **1.04E-20** |
| ENSG00000114423.25 | **CBLB** | 868 | protein_coding | **1204.994** | 1.398 | 0.421 | **2.635** | 3.46E-22 | **1.06E-20** |
| ENSG00000144867.15 | **SRPRB** | 58477 | protein_coding | **561.314** | -0.928 | -0.279 | **0.526** | 3.61E-22 | **1.10E-20** |
| ENSG00000272288.9 | **ILRUN-AS1** | 10192924[illegible] | lncRNA | **98.408** | 1.644 | 0.495 | **3.126** | 3.82E-22 | **1.16E-20** |
| ENSG00000136854.28 | **STXBP1** | 6812 | protein_coding | **63.975** | 3.070 | 0.924 | **8.398** | 3.84E-22 | **1.17E-20** |
| ENSG00000165801.12 | **ARHGEF40** | 55701 | protein_coding | **82.174** | 2.398 | 0.722 | **5.272** | 3.89E-22 | **1.18E-20** |
| ENSG00000185215.13 | **TNFAIP2** | 7127 | protein_coding | **3303.639** | 1.660 | 0.500 | **3.160** | 4.25E-22 | **1.29E-20** |
| ENSG00000211450.11 | **SELENOH** | 280636 | protein_coding | **1376.248** | -1.083 | -0.326 | **0.472** | 4.66E-22 | **1.41E-20** |
| ENSG00000160949.20 | **TONSL** | 4796 | protein_coding | **563.699** | -1.681 | -0.506 | **0.312** | 4.79E-22 | **1.44E-20** |
| ENSG00000105281.15 | **SLC1A5** | 6510 | protein_coding | **2559.488** | -1.546 | -0.465 | **0.342** | 5.03E-22 | **1.51E-20** |
| ENSG00000088367.25 | **EPB41L1** | 2036 | protein_coding | **72.555** | 3.723 | 1.121 | **13.209** | 6.01E-22 | **1.81E-20** |
| ENSG00000237883.5 | **DGUOK-AS1** | | lncRNA | **171.928** | 1.612 | 0.485 | **3.057** | 6.03E-22 | **1.81E-20** |
| ENSG00000300510.1 | **ENSG00000300510** | | protein_coding | **138.266** | 2.196 | 0.661 | **4.583** | 6.40E-22 | **1.92E-20** |
| ENSG00000100271.19 | **TTLL1** | 25809 | protein_coding | **164.036** | 1.836 | 0.553 | **3.570** | 6.44E-22 | **1.93E-20** |
| ENSG00000186918.16 | **ZNF395** | 55893 | protein_coding | **1402.742** | -1.065 | -0.321 | **0.478** | 6.95E-22 | **2.07E-20** |
| ENSG00000162755.16 | **KLHDC9** | 126823 | protein_coding | **121.928** | 2.210 | 0.665 | **4.628** | 7.44E-22 | **2.22E-20** |
| ENSG00000246089.6 | **MCPH1-DT** | 10028701[illegible] | lncRNA | **218.675** | 1.231 | 0.371 | **2.347** | 7.60E-22 | **2.26E-20** |
| ENSG00000169896.20 | **ITGAM** | 3684 | protein_coding | **351.102** | 2.086 | 0.628 | **4.247** | 7.71E-22 | **2.29E-20** |
| ENSG00000079999.16 | **KEAP1** | 9817 | protein_coding | **600.034** | -0.993 | -0.299 | **0.503** | 7.73E-22 | **2.29E-20** |
| ENSG00000101400.8 | **SNTA1** | 6640 | protein_coding | **162.159** | 2.292 | 0.690 | **4.897** | 7.78E-22 | **2.30E-20** |
| ENSG00000105750.17 | **ZNF85** | 7639 | protein_coding | **116.298** | -1.741 | -0.524 | **0.299** | 8.03E-22 | **2.38E-20** |
| ENSG00000132879.16 | **FBXO44** | 93611 | protein_coding | **936.873** | 0.968 | 0.292 | **1.957** | 9.13E-22 | **2.70E-20** |

| | | | | | | | | | |
|---|---|---|---|---|---|---|---|---|---|
| ENSG00000095397.20 | **WHRN** | 25861 | protein_coding | **83.913** | 2.376 | 0.715 | **5.192** | 9.31E-22 | **2.75E-20** |
| ENSG00000155980.15 | **KIF5A** | 3798 | protein_coding | **72.429** | 2.746 | 0.827 | **6.707** | 9.39E-22 | **2.77E-20** |
| ENSG00000234132.3 | **ENSG0000(** | 10192930! | lncRNA | **95.445** | 3.448 | 1.038 | **10.915** | 9.88E-22 | **2.90E-20** |
| ENSG00000171792.13 | **RHNO1** | 83695 | protein_coding | **569.000** | -1.071 | -0.322 | **0.476** | 1.03E-21 | **3.02E-20** |
| ENSG00000198901.16 | **PRC1** | 9055 | protein_coding | **859.116** | -1.724 | -0.519 | **0.303** | 1.08E-21 | **3.18E-20** |
| ENSG00000286760.4 | **ENSG0000(** | 10028949! | lncRNA | **109.061** | 1.782 | 0.537 | **3.440** | 1.09E-21 | **3.19E-20** |
| ENSG00000272512.1 | **ENSG00000272512** | | lncRNA | **37.270** | 3.135 | 0.944 | **8.785** | 1.09E-21 | **3.19E-20** |
| ENSG00000111906.20 | **HDDC2** | 51020 | protein_coding | **615.202** | -1.034 | -0.311 | **0.488** | 1.10E-21 | **3.22E-20** |
| ENSG00000258376.3 | **PAPLN-AS1** | | lncRNA | **78.340** | 1.821 | 0.548 | **3.533** | 1.11E-21 | **3.22E-20** |
| ENSG00000135736.8 | **CCDC102A** | 92922 | protein_coding | **213.046** | 1.441 | 0.434 | **2.715** | 1.27E-21 | **3.69E-20** |
| ENSG00000112655.18 | **PTK7** | 5754 | protein_coding | **247.798** | 2.205 | 0.664 | **4.612** | 1.37E-21 | **3.97E-20** |
| ENSG00000110031.14 | **LPXN** | 9404 | protein_coding | **1875.035** | -1.591 | -0.479 | **0.332** | 1.39E-21 | **4.03E-20** |
| ENSG00000118193.14 | **KIF14** | 9928 | protein_coding | **164.648** | -1.700 | -0.512 | **0.308** | 1.44E-21 | **4.17E-20** |
| ENSG00000108556.11 | **CHRNE** | 1145 | protein_coding | **54.805** | 2.868 | 0.863 | **7.302** | 1.44E-21 | **4.17E-20** |
| ENSG00000100626.19 | **GALNT16** | 57452 | protein_coding | **28.791** | 4.665 | 1.404 | **25.361** | 1.44E-21 | **4.17E-20** |
| ENSG00000120697.12 | **ALG5** | 29880 | protein_coding | **305.538** | -1.046 | -0.315 | **0.484** | 1.46E-21 | **4.21E-20** |
| ENSG00000101079.23 | **NDRG3** | 57446 | protein_coding | **504.482** | 0.844 | 0.254 | **1.795** | 1.47E-21 | **4.24E-20** |
| ENSG00000119782.15 | **FKBP1B** | 2281 | protein_coding | **45.553** | 3.425 | 1.031 | **10.742** | 1.52E-21 | **4.37E-20** |
| ENSG00000125445.13 | **MRPS7** | 51081 | protein_coding | **788.042** | -0.913 | -0.275 | **0.531** | 1.57E-21 | **4.49E-20** |
| ENSG00000182199.13 | **SHMT2** | 6472 | protein_coding | **3764.986** | -1.406 | -0.423 | **0.377** | 1.59E-21 | **4.56E-20** |
| ENSG00000171865.12 | **RNASEH1** | 246243 | protein_coding | **554.800** | -1.005 | -0.303 | **0.498** | 1.68E-21 | **4.81E-20** |
| ENSG00000179598.7 | **PLD6** | 201164 | protein_coding | **128.069** | -1.356 | -0.408 | **0.391** | 1.69E-21 | **4.82E-20** |
| ENSG00000117266.18 | **CDK18** | 5129 | protein_coding | **93.071** | 2.343 | 0.705 | **5.075** | 1.69E-21 | **4.83E-20** |
| ENSG00000153208.21 | **MERTK** | 10461 | protein_coding | **98.095** | 2.282 | 0.687 | **4.865** | 1.72E-21 | **4.91E-20** |
| ENSG00000099282.12 | **TSPAN15** | 23555 | protein_coding | **101.654** | 2.383 | 0.717 | **5.215** | 1.74E-21 | **4.94E-20** |
| ENSG00000170949.20 | **ZNF160** | 90338 | protein_coding | **1095.569** | -0.938 | -0.282 | **0.522** | 1.77E-21 | **5.03E-20** |
| ENSG00000268001.1 | **CARD8-AS:** | 10050581: | lncRNA | **77.307** | -1.977 | -0.595 | **0.254** | 1.86E-21 | **5.29E-20** |
| ENSG00000151466.14 | **SCLT1** | 132320 | protein_coding | **437.073** | -1.104 | -0.332 | **0.465** | 1.88E-21 | **5.33E-20** |
| ENSG00000127564.19 | **PKMYT1** | 9088 | protein_coding | **507.881** | -2.397 | -0.721 | **0.190** | 1.96E-21 | **5.55E-20** |
| ENSG00000135486.21 | **HNRNPA1** | 3178 | protein_coding | **22595.756** | -0.913 | -0.275 | **0.531** | 2.21E-21 | **6.23E-20** |
| ENSG00000095066.14 | **HOOK2** | 29911 | protein_coding | **378.582** | 1.309 | 0.394 | **2.478** | 2.36E-21 | **6.65E-20** |

| | | | | | | | | | |
|---|---|---|---|---|---|---|---|---|---|
| ENSG00000127415.15 | **IDUA** | 3425 | protein_coding | **342.758** | 1.442 | 0.434 | **2.717** | 2.55E-21 | **7.16E-20** |
| ENSG00000163902.14 | **RPN1** | 6184 | protein_coding | **2024.294** | -0.726 | -0.218 | **0.605** | 2.57E-21 | **7.23E-20** |
| ENSG00000100479.16 | **POLE2** | 5427 | protein_coding | **111.010** | -2.311 | -0.696 | **0.202** | 2.65E-21 | **7.44E-20** |
| ENSG00000129195.18 | **PIMREG** | 54478 | protein_coding | **171.254** | -1.422 | -0.428 | **0.373** | 2.69E-21 | **7.52E-20** |
| ENSG00000138134.14 | **STAMBPL1** | 57559 | protein_coding | **316.521** | -1.251 | -0.376 | **0.420** | 2.75E-21 | **7.69E-20** |
| ENSG00000160460.18 | **SPTBN4** | 57731 | protein_coding | **56.086** | 2.245 | 0.676 | **4.740** | 2.76E-21 | **7.72E-20** |
| ENSG00000123159.18 | **GIPC1** | 10755 | protein_coding | **591.241** | 1.402 | 0.422 | **2.642** | 2.77E-21 | **7.72E-20** |
| ENSG00000048828.20 | **FAM120A** | 23196 | protein_coding | **1190.547** | -0.897 | -0.270 | **0.537** | 2.87E-21 | **7.99E-20** |
| ENSG00000070404.12 | **FSTL3** | 10272 | protein_coding | **567.108** | 2.605 | 0.784 | **6.085** | 3.05E-21 | **8.47E-20** |
| ENSG00000050344.11 | **NFE2L3** | 9603 | protein_coding | **623.060** | 0.962 | 0.289 | **1.948** | 3.09E-21 | **8.58E-20** |
| ENSG00000196365.14 | **LONP1** | 9361 | protein_coding | **2535.253** | -1.028 | -0.309 | **0.491** | 3.28E-21 | **9.09E-20** |
| ENSG00000134574.14 | **DDB2** | 1643 | protein_coding | **2205.603** | -1.210 | -0.364 | **0.432** | 3.59E-21 | **9.94E-20** |
| ENSG00000105483.20 | **CARD8** | 22900 | protein_coding | **905.154** | -1.133 | -0.341 | **0.456** | 3.60E-21 | **9.95E-20** |
| ENSG00000149929.18 | **HIRIP3** | 8479 | protein_coding | **486.879** | -1.251 | -0.377 | **0.420** | 3.69E-21 | **1.02E-19** |
| ENSG00000005189.22 | **REXO5** | 81691 | protein_coding | **157.523** | -1.284 | -0.387 | **0.411** | 3.78E-21 | **1.04E-19** |
| ENSG00000100985.9 | **MMP9** | 4318 | protein_coding | **56.105** | 2.821 | 0.849 | **7.066** | 3.99E-21 | **1.10E-19** |
| ENSG00000176485.16 | **PLAAT3** | 11145 | protein_coding | **493.480** | 1.823 | 0.549 | **3.538** | 4.64E-21 | **1.27E-19** |
| ENSG00000148229.15 | **POLE3** | 54107 | protein_coding | **439.689** | -0.982 | -0.296 | **0.506** | 4.66E-21 | **1.28E-19** |
| ENSG00000255337.2 | **TMEM123-** | 10192842 | lncRNA | **72.460** | 2.128 | 0.641 | **4.371** | 4.70E-21 | **1.29E-19** |
| ENSG00000276900.1 | **AMN1-AS1** | 138842261 | lncRNA | **132.777** | 1.537 | 0.463 | **2.901** | 4.76E-21 | **1.30E-19** |
| ENSG00000143891.19 | **GALM** | 130589 | protein_coding | **192.772** | 2.256 | 0.679 | **4.776** | 4.87E-21 | **1.33E-19** |
| ENSG00000230798.8 | **FOXD3-AS1** | 100996301 | lncRNA | **62.023** | 4.537 | 1.366 | **23.223** | 4.94E-21 | **1.35E-19** |
| ENSG00000134955.14 | **SLC37A2** | 219855 | protein_coding | **334.441** | -1.443 | -0.434 | **0.368** | 5.03E-21 | **1.37E-19** |
| ENSG00000176490.7 | **DIRAS1** | 148252 | protein_coding | **162.411** | 2.515 | 0.757 | **5.717** | 5.09E-21 | **1.38E-19** |
| ENSG00000117650.15 | **NEK2** | 4751 | protein_coding | **197.233** | -1.759 | -0.529 | **0.296** | 5.31E-21 | **1.44E-19** |
| ENSG00000111665.14 | **CDCA3** | 83461 | protein_coding | **861.845** | -1.915 | -0.576 | **0.265** | 5.57E-21 | **1.51E-19** |
| ENSG00000160957.17 | **RECQL4** | 9401 | protein_coding | **865.386** | -2.217 | -0.668 | **0.215** | 5.62E-21 | **1.52E-19** |
| ENSG00000180198.19 | **RCC1** | 1104 | protein_coding | **1615.023** | -1.124 | -0.338 | **0.459** | 5.87E-21 | **1.59E-19** |
| ENSG00000165244.8 | **ZNF367** | 195828 | protein_coding | **79.457** | -2.022 | -0.609 | **0.246** | 5.97E-21 | **1.61E-19** |
| ENSG00000169188.7 | **APEX2** | 27301 | protein_coding | **619.967** | -0.979 | -0.295 | **0.507** | 6.30E-21 | **1.70E-19** |
| ENSG00000111669.17 | **TPI1** | 7167 | protein_coding | **12874.859** | -1.093 | -0.329 | **0.469** | 6.34E-21 | **1.71E-19** |

| | | | | | | | | | |
|---|---|---|---|---|---|---|---|---|---|
| ENSG00000137868.21 | **STRA6** | 64220 | protein_coding | **89.972** | 2.546 | 0.766 | **5.839** | 6.60E-21 | **1.77E-19** |
| ENSG00000205786.9 | **LINC01531** | 100128682 | lncRNA | **63.645** | 3.589 | 1.080 | **12.033** | 7.19E-21 | **1.93E-19** |
| ENSG00000014138.12 | **POLA2** | 23649 | protein_coding | **732.206** | -1.409 | -0.424 | **0.376** | 7.68E-21 | **2.06E-19** |
| ENSG00000166813.19 | **KIF7** | 374654 | protein_coding | **72.352** | 2.029 | 0.611 | **4.081** | 7.93E-21 | **2.12E-19** |
| ENSG00000229839.7 | **ENSG0000(** | 12490601З | lncRNA | **175.764** | 1.940 | 0.584 | **3.836** | 8.03E-21 | **2.15E-19** |
| ENSG00000233308.3 | **OSTN-AS1** | 10648073ε | lncRNA | **159.714** | 2.271 | 0.684 | **4.827** | 8.23E-21 | **2.20E-19** |
| ENSG00000160867.17 | **FGFR4** | 2264 | protein_coding | **178.179** | 1.966 | 0.592 | **3.906** | 8.48E-21 | **2.26E-19** |
| ENSG00000163807.8 | **KIAA1143** | 57456 | protein_coding | **533.761** | -1.686 | -0.508 | **0.311** | 8.99E-21 | **2.39E-19** |
| ENSG00000109089.9 | **CDR2L** | 30850 | protein_coding | **30.444** | 4.840 | 1.457 | **28.638** | 9.14E-21 | **2.43E-19** |
| ENSG00000063244.16 | **U2AF2** | 11338 | protein_coding | **3563.710** | -0.747 | -0.225 | **0.596** | 9.32E-21 | **2.47E-19** |
| ENSG00000123329.22 | **ARHGAP9** | 64333 | protein_coding | **1215.662** | -1.654 | -0.498 | **0.318** | 9.72E-21 | **2.58E-19** |
| ENSG00000105011.11 | **ASF1B** | 55723 | protein_coding | **909.632** | -1.611 | -0.485 | **0.327** | 9.91E-21 | **2.63E-19** |
| ENSG00000273272.3 | **ENSG00000273272** | | lncRNA | **158.246** | -1.954 | -0.588 | **0.258** | 1.04E-20 | **2.76E-19** |
| ENSG00000146232.19 | **NFKBIE** | 4794 | protein_coding | **1751.872** | -1.180 | -0.355 | **0.441** | 1.05E-20 | **2.79E-19** |
| ENSG00000138834.16 | **MAPK8IP3** | 23162 | protein_coding | **4897.544** | 0.777 | 0.234 | **1.713** | 1.09E-20 | **2.88E-19** |
| ENSG00000072501.21 | **SMC1A** | 8243 | protein_coding | **1148.411** | -1.385 | -0.417 | **0.383** | 1.20E-20 | **3.16E-19** |
| ENSG00000280734.5 | **LINC01232** | | lncRNA | **122.806** | 1.717 | 0.517 | **3.288** | 1.25E-20 | **3.29E-19** |
| ENSG00000173540.16 | **GMPPB** | 29925 | protein_coding | **515.755** | -1.424 | -0.429 | **0.373** | 1.26E-20 | **3.32E-19** |
| ENSG00000160193.14 | **WDR4** | 10785 | protein_coding | **336.411** | -1.205 | -0.363 | **0.434** | 1.45E-20 | **3.80E-19** |
| ENSG00000173272.18 | **MZT2A** | 653784 | protein_coding | **1379.246** | -1.114 | -0.335 | **0.462** | 1.48E-20 | **3.88E-19** |
| ENSG00000101255.15 | **TRIB3** | 57761 | protein_coding | **870.895** | -1.322 | -0.398 | **0.400** | 1.53E-20 | **4.01E-19** |
| ENSG00000143851.19 | **PTPN7** | 5778 | protein_coding | **1225.338** | -1.419 | -0.427 | **0.374** | 1.55E-20 | **4.05E-19** |
| ENSG00000095932.8 | **SMIM24** | 284422 | protein_coding | **26.674** | 3.165 | 0.953 | **8.969** | 1.59E-20 | **4.14E-19** |
| ENSG00000179409.12 | **GEMIN4** | 50628 | protein_coding | **428.553** | -1.108 | -0.334 | **0.464** | 1.60E-20 | **4.17E-19** |
| ENSG00000215375.9 | **MYL5** | 4636 | protein_coding | **636.078** | 1.168 | 0.352 | **2.247** | 1.61E-20 | **4.20E-19** |
| ENSG00000230091.7 | **TMEM254-AS1** | | lncRNA | **68.287** | 2.242 | 0.675 | **4.731** | 1.62E-20 | **4.21E-19** |
| ENSG00000186871.8 | **ERCC6L** | 54821 | protein_coding | **88.100** | -1.996 | -0.601 | **0.251** | 1.79E-20 | **4.64E-19** |
| ENSG00000047932.16 | **GOPC** | 57120 | protein_coding | **743.560** | 1.034 | 0.311 | **2.048** | 1.91E-20 | **4.96E-19** |
| ENSG00000185862.9 | **EVI2B** | 2124 | protein_coding | **1027.668** | -1.549 | -0.466 | **0.342** | 1.99E-20 | **5.14E-19** |
| ENSG00000102109.11 | **PCSK1N** | 27344 | protein_coding | **56.443** | 5.142 | 1.548 | **35.304** | 2.00E-20 | **5.16E-19** |
| ENSG00000158793.16 | **NIT1** | 4817 | protein_coding | **1037.306** | 0.863 | 0.260 | **1.818** | 2.05E-20 | **5.29E-19** |

| | | | | | | | | | |
|---|---|---|---|---|---|---|---|---|---|
| ENSG00000293488.3 | **CLECL1** | 160365 | protein_coding | **412.685** | -1.840 | -0.554 | **0.279** | 2.06E-20 | **5.31E-19** |
| ENSG00000105205.8 | **CLC** | 1178 | protein_coding | **70.493** | 4.340 | 1.306 | **20.251** | 2.07E-20 | **5.33E-19** |
| ENSG00000215039.10 | **CD27-AS1** | 678655 | lncRNA | **576.372** | 1.935 | 0.583 | **3.824** | 2.09E-20 | **5.36E-19** |
| ENSG00000162522.13 | **NHSL3** | 57648 | protein_coding | **54.378** | 4.029 | 1.213 | **16.324** | 2.13E-20 | **5.47E-19** |
| ENSG00000164105.6 | **SAP30** | 8819 | protein_coding | **334.026** | -1.259 | -0.379 | **0.418** | 2.17E-20 | **5.56E-19** |
| ENSG00000105223.23 | **PLD3** | 23646 | protein_coding | **1723.878** | 0.881 | 0.265 | **1.842** | 2.43E-20 | **6.22E-19** |
| ENSG00000111344.14 | **RASAL1** | 8437 | protein_coding | **94.117** | 2.126 | 0.640 | **4.365** | 2.44E-20 | **6.23E-19** |
| ENSG00000130159.17 | **ECSIT** | 51295 | protein_coding | **519.349** | -1.226 | -0.369 | **0.428** | 2.46E-20 | **6.29E-19** |
| ENSG00000060656.22 | **PTPRU** | 10076 | protein_coding | **65.321** | 1.949 | 0.587 | **3.861** | 2.60E-20 | **6.63E-19** |
| ENSG00000102317.20 | **RBM3** | 5935 | protein_coding | **3137.737** | -0.859 | -0.259 | **0.551** | 2.63E-20 | **6.70E-19** |
| ENSG00000108179.16 | **PPIF** | 10105 | protein_coding | **1439.246** | -1.067 | -0.321 | **0.477** | 2.83E-20 | **7.19E-19** |
| ENSG00000133466.16 | **C1QTNF6** | 114904 | protein_coding | **459.515** | 1.307 | 0.393 | **2.474** | 2.90E-20 | **7.36E-19** |
| ENSG00000172725.16 | **CORO1B** | 57175 | protein_coding | **1331.288** | 0.881 | 0.265 | **1.842** | 2.98E-20 | **7.55E-19** |
| ENSG00000153551.16 | **CMTM7** | 112616 | protein_coding | **95.162** | 2.065 | 0.622 | **4.184** | 3.12E-20 | **7.90E-19** |
| ENSG00000146216.14 | **TTBK1** | 84630 | protein_coding | **31.823** | 3.214 | 0.968 | **9.282** | 3.27E-20 | **8.26E-19** |
| ENSG00000163449.13 | **TMEM169** | 92691 | protein_coding | **43.375** | 2.551 | 0.768 | **5.859** | 3.36E-20 | **8.47E-19** |
| ENSG00000069849.13 | **ATP1B3** | 483 | protein_coding | **1452.062** | -0.939 | -0.283 | **0.522** | 3.36E-20 | **8.47E-19** |
| ENSG00000289074.3 | **ENSG00000289074** | | lncRNA | **50.375** | 3.251 | 0.979 | **9.517** | 3.38E-20 | **8.53E-19** |
| ENSG00000128534.10 | **LSM8** | 51691 | protein_coding | **1037.701** | -0.862 | -0.259 | **0.550** | 3.41E-20 | **8.57E-19** |
| ENSG00000165490.15 | **DDIAS** | 220042 | protein_coding | **85.654** | -1.805 | -0.543 | **0.286** | 3.93E-20 | **9.86E-19** |
| ENSG00000160224.18 | **AIRE** | 326 | protein_coding | **32.247** | 5.159 | 1.553 | **35.722** | 4.01E-20 | **1.01E-18** |
| ENSG00000126266.5 | **FFAR1** | 2864 | protein_coding | **147.315** | 1.524 | 0.459 | **2.875** | 4.03E-20 | **1.01E-18** |
| ENSG00000183010.19 | **PYCR1** | 5831 | protein_coding | **944.709** | -1.180 | -0.355 | **0.441** | 4.16E-20 | **1.04E-18** |
| ENSG00000198722.17 | **UNC13B** | 10497 | protein_coding | **659.819** | 1.563 | 0.471 | **2.955** | 4.19E-20 | **1.05E-18** |
| ENSG00000008394.15 | **MGST1** | 4257 | protein_coding | **420.132** | -1.489 | -0.448 | **0.356** | 4.37E-20 | **1.09E-18** |
| ENSG00000117411.20 | **B4GALT2** | 8704 | protein_coding | **487.357** | -1.309 | -0.394 | **0.404** | 4.85E-20 | **1.21E-18** |
| ENSG00000227117.7 | **HORMAD2** | 10192966 | lncRNA | **45.938** | 2.342 | 0.705 | **5.071** | 4.94E-20 | **1.23E-18** |
| ENSG00000169169.17 | **CPT1C** | 126129 | protein_coding | **50.528** | 4.701 | 1.415 | **26.019** | 4.94E-20 | **1.23E-18** |
| ENSG00000172081.16 | **MOB3A** | 126308 | protein_coding | **5174.777** | -1.081 | -0.325 | **0.473** | 5.09E-20 | **1.26E-18** |
| ENSG00000196169.17 | **KIF19** | 124602 | protein_coding | **22.915** | 4.503 | 1.355 | **22.672** | 5.32E-20 | **1.32E-18** |
| ENSG00000119411.13 | **BSPRY** | 54836 | protein_coding | **155.994** | 1.891 | 0.569 | **3.708** | 5.46E-20 | **1.35E-18** |

| | | | | | | | | | |
|---|---|---|---|---|---|---|---|---|---|
| ENSG00000187609.18 | **EXD3** | 54932 | protein_coding | **438.661** | 1.074 | 0.323 | **2.105** | 5.66E-20 | **1.40E-18** |
| ENSG00000116679.19 | **IVNS1ABP** | 10625 | protein_coding | **1356.270** | -0.945 | -0.284 | **0.520** | 5.77E-20 | **1.42E-18** |
| ENSG00000132274.19 | **TRIM22** | 10346 | protein_coding | **3500.704** | -1.194 | -0.360 | **0.437** | 5.80E-20 | **1.43E-18** |
| ENSG00000005961.21 | **ITGA2B** | 3674 | protein_coding | **42.901** | 2.936 | 0.884 | **7.650** | 5.95E-20 | **1.47E-18** |
| ENSG00000166126.13 | **AMN** | 81693 | protein_coding | **94.748** | 2.451 | 0.738 | **5.469** | 6.49E-20 | **1.60E-18** |
| ENSG00000109084.15 | **TMEM97** | 27346 | protein_coding | **525.623** | -1.556 | -0.468 | **0.340** | 6.95E-20 | **1.71E-18** |
| ENSG00000146963.20 | **LUC7L2** | 51631 | protein_coding | **1523.977** | -0.648 | -0.195 | **0.638** | 7.55E-20 | **1.85E-18** |
| ENSG00000115255.14 | **REEP6** | 92840 | protein_coding | **43.329** | 2.612 | 0.786 | **6.112** | 7.65E-20 | **1.87E-18** |
| ENSG00000301565.1 | **ENSG00000301565** | | lncRNA | **310.106** | 1.237 | 0.372 | **2.356** | 8.16E-20 | **2.00E-18** |
| ENSG00000213015.11 | **ZNF580** | 51157 | protein_coding | **897.786** | 1.028 | 0.309 | **2.039** | 8.73E-20 | **2.13E-18** |
| ENSG00000007968.9 | **E2F2** | 1870 | protein_coding | **193.381** | -1.980 | -0.596 | **0.254** | 8.81E-20 | **2.15E-18** |
| ENSG00000040275.19 | **SPDL1** | 54908 | protein_coding | **366.763** | -1.298 | -0.391 | **0.407** | 9.46E-20 | **2.31E-18** |
| ENSG00000305420.1 | **ENSG00000305420** | | lncRNA | **24.325** | 4.524 | 1.362 | **23.009** | 9.58E-20 | **2.33E-18** |
| ENSG00000188833.12 | **ENTPD8** | 377841 | protein_coding | **29.694** | 6.522 | 1.963 | **91.882** | 1.04E-19 | **2.53E-18** |
| ENSG00000167740.11 | **CYB5D2** | 124936 | protein_coding | **346.131** | 1.084 | 0.326 | **2.119** | 1.05E-19 | **2.55E-18** |
| ENSG00000128342.6 | **LIF** | 3976 | protein_coding | **47.120** | -2.468 | -0.743 | **0.181** | 1.06E-19 | **2.57E-18** |
| ENSG00000130202.12 | **NECTIN2** | 5819 | protein_coding | **27.415** | 3.833 | 1.154 | **14.253** | 1.06E-19 | **2.57E-18** |
| ENSG00000102007.13 | **PLP2** | 5355 | protein_coding | **1940.646** | -1.383 | -0.416 | **0.383** | 1.08E-19 | **2.62E-18** |
| ENSG00000159147.20 | **DONSON** | 29980 | protein_coding | **335.706** | -1.459 | -0.439 | **0.364** | 1.11E-19 | **2.68E-18** |
| ENSG00000154473.20 | **BUB3** | 9184 | protein_coding | **1771.758** | -0.602 | -0.181 | **0.659** | 1.16E-19 | **2.79E-18** |
| ENSG00000075702.20 | **WDR62** | 284403 | protein_coding | **417.578** | -2.069 | -0.623 | **0.238** | 1.16E-19 | **2.79E-18** |
| ENSG00000131037.17 | **EPS8L1** | 54869 | protein_coding | **103.894** | 1.785 | 0.537 | **3.447** | 1.18E-19 | **2.83E-18** |
| ENSG00000226200.8 | **SGMS1-AS1** | | lncRNA | **245.318** | 1.507 | 0.454 | **2.842** | 1.21E-19 | **2.91E-18** |
| ENSG00000266964.8 | **FXYD1** | 5348 | protein_coding | **31.409** | 4.722 | 1.421 | **26.387** | 1.21E-19 | **2.91E-18** |
| ENSG00000121101.18 | **TEX14** | 56155 | protein_coding | **76.198** | 2.183 | 0.657 | **4.542** | 1.24E-19 | **2.97E-18** |
| ENSG00000145241.13 | **CENPC** | 1060 | protein_coding | **293.580** | -1.218 | -0.367 | **0.430** | 1.34E-19 | **3.21E-18** |
| ENSG00000182903.17 | **ZNF721** | 170960 | protein_coding | **417.058** | -0.867 | -0.261 | **0.548** | 1.35E-19 | **3.24E-18** |
| ENSG00000158828.10 | **PINK1** | 65018 | protein_coding | **164.730** | 1.777 | 0.535 | **3.427** | 1.37E-19 | **3.27E-18** |
| ENSG00000198089.18 | **SFI1** | 9814 | protein_coding | **2248.356** | 0.901 | 0.271 | **1.868** | 1.53E-19 | **3.66E-18** |
| ENSG00000157343.10 | **ARMC12** | 221481 | protein_coding | **72.113** | 1.935 | 0.583 | **3.825** | 1.59E-19 | **3.79E-18** |
| ENSG00000132026.17 | **RTBDN** | 83546 | protein_coding | **44.598** | 6.121 | 1.843 | **69.599** | 1.59E-19 | **3.79E-18** |

| | | | | | | | | | |
|---|---|---|---|---|---|---|---|---|---|
| ENSG00000156127.8 | **BATF** | 10538 | protein_coding | **1164.412** | -0.882 | -0.266 | **0.542** | 1.65E-19 | **3.92E-18** |
| ENSG00000048162.24 | **NOP16** | 51491 | protein_coding | **839.722** | -1.188 | -0.358 | **0.439** | 1.70E-19 | **4.03E-18** |
| ENSG00000188191.18 | **PRKAR1B** | 5575 | protein_coding | **300.469** | -1.296 | -0.390 | **0.407** | 1.70E-19 | **4.04E-18** |
| ENSG00000028277.24 | **POU2F2** | 5452 | protein_coding | **1804.340** | -1.218 | -0.367 | **0.430** | 1.74E-19 | **4.11E-18** |
| ENSG00000099330.11 | **OCEL1** | 79629 | protein_coding | **463.568** | 1.614 | 0.486 | **3.062** | 1.76E-19 | **4.15E-18** |
| ENSG00000155438.14 | **NIFK** | 84365 | protein_coding | **760.835** | -0.846 | -0.255 | **0.556** | 1.76E-19 | **4.15E-18** |
| ENSG00000167900.15 | **TK1** | 7083 | protein_coding | **1010.691** | -1.514 | -0.456 | **0.350** | 1.79E-19 | **4.21E-18** |
| ENSG00000130038.12 | **CRACR2A** | 84766 | protein_coding | **149.844** | -1.489 | -0.448 | **0.356** | 1.84E-19 | **4.33E-18** |
| ENSG00000184381.22 | **PLA2G6** | 8398 | protein_coding | **1527.342** | 0.973 | 0.293 | **1.963** | 1.93E-19 | **4.55E-18** |
| ENSG00000065054.16 | **NHERF2** | 9351 | protein_coding | **62.370** | 4.232 | 1.274 | **18.787** | 2.09E-19 | **4.91E-18** |
| ENSG00000156042.20 | **CFAP70** | 118491 | protein_coding | **91.866** | 2.072 | 0.624 | **4.204** | 2.11E-19 | **4.96E-18** |
| ENSG00000061938.22 | **TNK2** | 10188 | protein_coding | **3210.625** | 0.915 | 0.276 | **1.886** | 2.13E-19 | **4.99E-18** |
| ENSG00000139083.13 | **ETV6** | 2120 | protein_coding | **2571.527** | 1.290 | 0.388 | **2.445** | 2.34E-19 | **5.47E-18** |
| ENSG00000125656.14 | **CLPP** | 8192 | protein_coding | **967.382** | -0.930 | -0.280 | **0.525** | 2.35E-19 | **5.51E-18** |
| ENSG00000075426.14 | **FOSL2** | 2355 | protein_coding | **91.146** | -2.287 | -0.688 | **0.205** | 2.41E-19 | **5.64E-18** |
| ENSG00000185112.6 | **FAM43A** | 131583 | protein_coding | **457.855** | 1.768 | 0.532 | **3.406** | 2.43E-19 | **5.68E-18** |
| ENSG00000035403.20 | **VCL** | 7414 | protein_coding | **498.761** | 1.369 | 0.412 | **2.583** | 2.45E-19 | **5.71E-18** |
| ENSG00000112118.22 | **MCM3** | 4172 | protein_coding | **2465.115** | -0.938 | -0.282 | **0.522** | 2.53E-19 | **5.89E-18** |
| ENSG00000187790.14 | **FANCM** | 57697 | protein_coding | **126.317** | -1.361 | -0.410 | **0.389** | 2.53E-19 | **5.89E-18** |
| ENSG00000184900.18 | **SUMO3** | 6612 | protein_coding | **765.433** | -1.088 | -0.327 | **0.470** | 2.75E-19 | **6.39E-18** |
| ENSG00000105655.21 | **ISYNA1** | 51477 | protein_coding | **847.618** | 1.910 | 0.575 | **3.757** | 2.81E-19 | **6.52E-18** |
| ENSG00000099814.19 | **CEP170B** | 283638 | protein_coding | **75.042** | 2.404 | 0.724 | **5.294** | 2.88E-19 | **6.66E-18** |
| ENSG00000089006.19 | **SNX5** | 27131 | protein_coding | **2843.538** | -0.930 | -0.280 | **0.525** | 2.88E-19 | **6.66E-18** |
| ENSG00000161980.8 | **POLR3K** | 51728 | protein_coding | **390.230** | -0.898 | -0.270 | **0.537** | 2.97E-19 | **6.86E-18** |
| ENSG00000163346.19 | **PBXIP1** | 57326 | protein_coding | **1687.007** | 0.854 | 0.257 | **1.807** | 2.98E-19 | **6.87E-18** |
| ENSG00000058668.18 | **ATP2B4** | 493 | protein_coding | **87.517** | 2.064 | 0.621 | **4.180** | 2.98E-19 | **6.88E-18** |
| ENSG00000105376.7 | **ICAM5** | 7087 | protein_coding | **328.205** | 1.361 | 0.410 | **2.568** | 2.99E-19 | **6.88E-18** |
| ENSG00000111275.15 | **ALDH2** | 217 | protein_coding | **1433.684** | 1.666 | 0.502 | **3.174** | 3.01E-19 | **6.92E-18** |
| ENSG00000087086.18 | **FTL** | 2512 | protein_coding | **31867.962** | -0.981 | -0.295 | **0.506** | 3.06E-19 | **7.02E-18** |
| ENSG00000134748.15 | **PRPF38A** | 84950 | protein_coding | **781.931** | -1.138 | -0.343 | **0.454** | 3.20E-19 | **7.35E-18** |
| ENSG00000112137.20 | **PHACTR1** | 221692 | protein_coding | **1321.992** | -1.461 | -0.440 | **0.363** | 3.21E-19 | **7.36E-18** |

| | | | | | | | | | |
|---|---|---|---|---|---|---|---|---|---|
| ENSG00000177943.16 | **MAMDC4** | 158056 | protein_coding | **258.982** | -1.374 | -0.414 | **0.386** | 3.47E-19 | **7.94E-18** |
| ENSG00000069974.19 | **RAB27A** | 5873 | protein_coding | **449.879** | -1.369 | -0.412 | **0.387** | 3.50E-19 | **8.00E-18** |
| ENSG00000173020.13 | **GRK2** | 156 | protein_coding | **3877.660** | -0.818 | -0.246 | **0.567** | 3.77E-19 | **8.60E-18** |
| ENSG00000184916.11 | **JAG2** | 3714 | protein_coding | **72.331** | 2.240 | 0.674 | **4.724** | 3.82E-19 | **8.72E-18** |
| ENSG00000148660.24 | **CAMK2G** | 818 | protein_coding | **507.562** | 1.254 | 0.377 | **2.385** | 3.90E-19 | **8.88E-18** |
| ENSG00000145911.9 | **N4BP3** | 23138 | protein_coding | **467.032** | -1.231 | -0.370 | **0.426** | 3.90E-19 | **8.88E-18** |
| ENSG00000184992.14 | **BRI3BP** | 140707 | protein_coding | **583.461** | -1.328 | -0.400 | **0.398** | 4.11E-19 | **9.33E-18** |
| ENSG00000104147.11 | **OIP5** | 11339 | protein_coding | **127.278** | -1.447 | -0.436 | **0.367** | 4.14E-19 | **9.40E-18** |
| ENSG00000291176.2 | **ENSG00000291176** | | lncRNA | **384.347** | 1.190 | 0.358 | **2.282** | 4.18E-19 | **9.48E-18** |
| ENSG00000149809.19 | **TM7SF2** | 7108 | protein_coding | **906.106** | 1.010 | 0.304 | **2.015** | 4.24E-19 | **9.59E-18** |
| ENSG00000141736.16 | **ERBB2** | 2064 | protein_coding | **309.343** | 0.929 | 0.280 | **1.904** | 4.29E-19 | **9.70E-18** |
| ENSG00000171861.13 | **MRM3** | 55178 | protein_coding | **311.650** | -0.911 | -0.274 | **0.532** | 4.30E-19 | **9.70E-18** |
| ENSG00000180828.3 | **BHLHE22** | 27319 | protein_coding | **191.747** | -1.639 | -0.493 | **0.321** | 4.32E-19 | **9.75E-18** |
| ENSG00000126749.19 | **EMG1** | 10436 | protein_coding | **734.055** | -1.178 | -0.355 | **0.442** | 4.40E-19 | **9.91E-18** |
| ENSG00000135069.16 | **PSAT1** | 29968 | protein_coding | **1332.499** | -1.932 | -0.581 | **0.262** | 4.78E-19 | **1.08E-17** |
| ENSG00000287097.2 | **ENSG0000(** | 10798663( | lncRNA | **68.860** | 2.701 | 0.813 | **6.503** | 4.84E-19 | **1.09E-17** |
| ENSG00000124802.14 | **EEF1E1** | 9521 | protein_coding | **279.828** | -1.112 | -0.335 | **0.463** | 5.03E-19 | **1.13E-17** |
| ENSG00000236790.8 | **LINC00299** | 339789 | lncRNA | **135.690** | 1.599 | 0.481 | **3.030** | 5.11E-19 | **1.15E-17** |
| ENSG00000100373.12 | **UPK3A** | 7380 | protein_coding | **32.306** | 4.007 | 1.206 | **16.074** | 5.14E-19 | **1.15E-17** |
| ENSG00000183763.11 | **TRAIP** | 10293 | protein_coding | **222.135** | -2.112 | -0.636 | **0.231** | 5.16E-19 | **1.15E-17** |
| ENSG00000005022.8 | **SLC25A5** | 292 | protein_coding | **2478.351** | -1.149 | -0.346 | **0.451** | 5.49E-19 | **1.23E-17** |
| ENSG00000159164.12 | **SV2A** | 9900 | protein_coding | **257.612** | 1.371 | 0.413 | **2.586** | 5.51E-19 | **1.23E-17** |
| ENSG00000148840.13 | **PPRC1** | 23082 | protein_coding | **780.888** | -0.904 | -0.272 | **0.534** | 5.57E-19 | **1.24E-17** |
| ENSG00000136982.8 | **DSCC1** | 79075 | protein_coding | **96.054** | -2.333 | -0.702 | **0.198** | 5.67E-19 | **1.26E-17** |
| ENSG00000163958.15 | **ZDHHC19** | 131540 | protein_coding | **51.894** | 5.035 | 1.516 | **32.787** | 5.78E-19 | **1.29E-17** |
| ENSG00000172794.22 | **RAB37** | 326624 | protein_coding | **170.319** | 1.603 | 0.483 | **3.038** | 5.91E-19 | **1.31E-17** |
| ENSG00000167642.15 | **SPINT2** | 10653 | protein_coding | **344.042** | 2.525 | 0.760 | **5.755** | 6.00E-19 | **1.33E-17** |
| ENSG00000130518.20 | **IQCN** | 80726 | protein_coding | **53.575** | 1.898 | 0.571 | **3.726** | 6.08E-19 | **1.35E-17** |
| ENSG00000205356.12 | **TECPR1** | 25851 | protein_coding | **522.293** | 0.979 | 0.295 | **1.971** | 6.27E-19 | **1.39E-17** |
| ENSG00000154265.18 | **ABCA5** | 23461 | protein_coding | **1047.012** | -0.867 | -0.261 | **0.548** | 7.26E-19 | **1.61E-17** |
| ENSG00000133069.19 | **TMCC2** | 9911 | protein_coding | **32.822** | 3.335 | 1.004 | **10.091** | 7.33E-19 | **1.62E-17** |

| | | | | | | | | | |
|---|---|---|---|---|---|---|---|---|---|
| ENSG00000139131.15 | **YARS2** | 51067 | protein_coding | **305.233** | -1.246 | -0.375 | **0.421** | 7.36E-19 | **1.62E-17** |
| ENSG00000112996.13 | **MRPS30** | 10884 | protein_coding | **466.895** | -0.980 | -0.295 | **0.507** | 7.71E-19 | **1.70E-17** |
| ENSG00000127423.12 | **AUNIP** | 79000 | protein_coding | **85.368** | -2.440 | -0.735 | **0.184** | 7.89E-19 | **1.74E-17** |
| ENSG00000116221.17 | **MRPL37** | 51253 | protein_coding | **1225.845** | -1.030 | -0.310 | **0.490** | 8.07E-19 | **1.77E-17** |
| ENSG00000019144.22 | **PHLDB1** | 23187 | protein_coding | **40.012** | 4.162 | 1.253 | **17.898** | 8.07E-19 | **1.77E-17** |
| ENSG00000046651.18 | **OFD1** | 8481 | protein_coding | **2407.678** | 0.992 | 0.299 | **1.988** | 8.32E-19 | **1.83E-17** |
| ENSG00000134463.17 | **ECHDC3** | 79746 | protein_coding | **262.643** | 1.423 | 0.428 | **2.682** | 8.46E-19 | **1.85E-17** |
| ENSG00000063127.16 | **SLC6A16** | 28968 | protein_coding | **122.640** | 1.588 | 0.478 | **3.007** | 8.46E-19 | **1.85E-17** |
| ENSG00000124140.16 | **SLC12A5** | 57468 | protein_coding | **71.724** | 3.130 | 0.942 | **8.755** | 8.59E-19 | **1.88E-17** |
| ENSG00000159708.19 | **LRRC36** | 55282 | protein_coding | **32.764** | 4.445 | 1.338 | **21.779** | 8.79E-19 | **1.92E-17** |
| ENSG00000273702.2 | **ENSG00000273702** | | lncRNA | **94.710** | 2.479 | 0.746 | **5.576** | 8.91E-19 | **1.94E-17** |
| ENSG00000235419.6 | **ENSG00000235419** | | lncRNA | **102.529** | 2.413 | 0.726 | **5.325** | 9.29E-19 | **2.02E-17** |
| ENSG00000013810.23 | **TACC3** | 10460 | protein_coding | **1841.773** | -1.638 | -0.493 | **0.321** | 9.41E-19 | **2.05E-17** |
| ENSG00000182307.16 | **C8orf33** | 65265 | protein_coding | **347.759** | -1.094 | -0.329 | **0.469** | 1.04E-18 | **2.26E-17** |
| ENSG00000162512.18 | **SDC3** | 9672 | protein_coding | **85.227** | 2.263 | 0.681 | **4.798** | 1.04E-18 | **2.27E-17** |
| ENSG00000128245.17 | **YWHAH** | 7533 | protein_coding | **985.663** | -0.962 | -0.290 | **0.513** | 1.07E-18 | **2.33E-17** |
| ENSG00000139112.13 | **GABARAPL** | 23710 | protein_coding | **215.431** | 1.469 | 0.442 | **2.769** | 1.08E-18 | **2.33E-17** |
| ENSG00000104361.12 | **NIPAL2** | 79815 | protein_coding | **313.046** | -1.221 | -0.368 | **0.429** | 1.08E-18 | **2.33E-17** |
| ENSG00000163328.16 | **GPR155** | 151556 | protein_coding | **206.329** | 1.345 | 0.405 | **2.541** | 1.10E-18 | **2.37E-17** |
| ENSG00000291113.3 | **ENSG00000291113** | | lncRNA | **544.277** | -1.421 | -0.428 | **0.374** | 1.14E-18 | **2.47E-17** |
| ENSG00000093000.21 | **NUP50** | 10762 | protein_coding | **1218.038** | -0.788 | -0.237 | **0.579** | 1.31E-18 | **2.83E-17** |
| ENSG00000187595.20 | **ZNF385C** | 201181 | protein_coding | **36.852** | 2.473 | 0.744 | **5.550** | 1.37E-18 | **2.94E-17** |
| ENSG00000167733.16 | **HSD11B1L** | 374875 | protein_coding | **109.100** | 2.082 | 0.627 | **4.235** | 1.39E-18 | **3.00E-17** |
| ENSG00000218510.9 | **LINC00339** | 29092 | lncRNA | **386.951** | 1.147 | 0.345 | **2.214** | 1.42E-18 | **3.04E-17** |
| ENSG00000165175.19 | **MID1IP1** | 58526 | protein_coding | **897.137** | -0.998 | -0.300 | **0.501** | 1.43E-18 | **3.07E-17** |
| ENSG00000206077.15 | **ZDHHC11B** | 653082 | protein_coding | **118.724** | 3.179 | 0.957 | **9.058** | 1.47E-18 | **3.16E-17** |
| ENSG00000074842.10 | **MYDGF** | 56005 | protein_coding | **745.041** | -0.914 | -0.275 | **0.531** | 1.50E-18 | **3.22E-17** |
| ENSG00000261211.3 | **ENSG00000261211** | | lncRNA | **42.859** | 4.591 | 1.382 | **24.102** | 1.61E-18 | **3.45E-17** |
| ENSG00000145022.7 | **TCTA** | 6988 | protein_coding | **627.325** | 0.782 | 0.235 | **1.719** | 1.67E-18 | **3.58E-17** |
| ENSG00000135596.20 | **MICAL1** | 64780 | protein_coding | **6234.424** | 0.792 | 0.239 | **1.732** | 1.71E-18 | **3.65E-17** |
| ENSG00000166448.17 | **TMEM130** | 222865 | protein_coding | **26.093** | 3.868 | 1.164 | **14.601** | 1.75E-18 | **3.73E-17** |

| | | | | | | | | | |
|---|---|---|---|---|---|---|---|---|---|
| ENSG00000100077.18 | **GRK3** | 157 | protein_coding | **1088.976** | -0.819 | -0.247 | **0.567** | 1.77E-18 | **3.76E-17** |
| ENSG00000059915.19 | **PSD** | 5662 | protein_coding | **64.142** | 2.507 | 0.755 | **5.686** | 1.83E-18 | **3.89E-17** |
| ENSG00000157193.20 | **LRP8** | 7804 | protein_coding | **569.801** | -1.568 | -0.472 | **0.337** | 1.87E-18 | **3.97E-17** |
| ENSG00000011028.15 | **MRC2** | 9902 | protein_coding | **219.702** | 1.591 | 0.479 | **3.012** | 1.90E-18 | **4.02E-17** |
| ENSG00000149485.21 | **FADS1** | 3992 | protein_coding | **1085.228** | -1.177 | -0.354 | **0.442** | 2.08E-18 | **4.40E-17** |
| ENSG00000130475.18 | **FCHO1** | 23149 | protein_coding | **1156.670** | 1.004 | 0.302 | **2.005** | 2.08E-18 | **4.40E-17** |
| ENSG00000172889.18 | **EGFL7** | 51162 | protein_coding | **71.039** | 2.787 | 0.839 | **6.900** | 2.09E-18 | **4.42E-17** |
| ENSG00000253958.3 | **CLDN23** | 137075 | protein_coding | **36.804** | 2.599 | 0.782 | **6.057** | 2.11E-18 | **4.47E-17** |
| ENSG00000272273.4 | **IER3-AS1** | | lncRNA | **49.697** | 1.886 | 0.568 | **3.697** | 2.14E-18 | **4.52E-17** |
| ENSG00000156869.16 | **FRRS1** | 391059 | protein_coding | **297.668** | 1.169 | 0.352 | **2.248** | 2.22E-18 | **4.68E-17** |
| ENSG00000290831.2 | **NSUN5P2** | 260294 | lncRNA | **1159.176** | 0.801 | 0.241 | **1.743** | 2.31E-18 | **4.86E-17** |
| ENSG00000126803.10 | **HSPA2** | 3306 | protein_coding | **94.494** | 2.613 | 0.787 | **6.116** | 2.35E-18 | **4.94E-17** |
| ENSG00000182253.16 | **SYNM** | 23336 | protein_coding | **130.603** | 2.162 | 0.651 | **4.476** | 2.47E-18 | **5.18E-17** |
| ENSG00000163435.17 | **ELF3** | 1999 | protein_coding | **116.386** | 1.584 | 0.477 | **2.999** | 2.56E-18 | **5.38E-17** |
| ENSG00000161981.12 | **SNRNP25** | 79622 | protein_coding | **703.071** | -1.370 | -0.412 | **0.387** | 2.63E-18 | **5.52E-17** |
| ENSG00000274523.7 | **RCC1L** | 81554 | protein_coding | **872.684** | -1.056 | -0.318 | **0.481** | 2.80E-18 | **5.86E-17** |
| ENSG00000152455.18 | **SUV39H2** | 79723 | protein_coding | **133.345** | -1.539 | -0.463 | **0.344** | 2.90E-18 | **6.07E-17** |
| ENSG00000275302.3 | **CCL4** | 6351 | protein_coding | **894.934** | -1.877 | -0.565 | **0.272** | 2.96E-18 | **6.18E-17** |
| ENSG00000101190.15 | **TCFL5** | 10732 | protein_coding | **450.152** | 1.268 | 0.382 | **2.408** | 3.05E-18 | **6.37E-17** |
| ENSG00000080839.14 | **RBL1** | 5933 | protein_coding | **214.246** | -1.246 | -0.375 | **0.422** | 3.12E-18 | **6.50E-17** |
| ENSG00000113648.19 | **MACROH2.** | 9555 | protein_coding | **2155.936** | -0.633 | -0.191 | **0.645** | 3.14E-18 | **6.54E-17** |
| ENSG00000051596.12 | **THOC3** | 84321 | protein_coding | **546.751** | -1.279 | -0.385 | **0.412** | 3.16E-18 | **6.59E-17** |
| ENSG00000233834.8 | **GIRGL** | | lncRNA | **181.412** | 1.306 | 0.393 | **2.473** | 3.21E-18 | **6.67E-17** |
| ENSG00000204560.12 | **DHX16** | 8449 | protein_coding | **1035.433** | 1.038 | 0.312 | **2.053** | 3.32E-18 | **6.88E-17** |
| ENSG00000143819.15 | **EPHX1** | 2052 | protein_coding | **87.624** | 1.607 | 0.484 | **3.046** | 3.38E-18 | **7.01E-17** |
| ENSG00000120334.18 | **CENPL** | 91687 | protein_coding | **142.644** | -1.584 | -0.477 | **0.334** | 3.39E-18 | **7.02E-17** |
| ENSG00000163528.15 | **CHCHD4** | 131474 | protein_coding | **174.442** | -1.234 | -0.372 | **0.425** | 3.44E-18 | **7.12E-17** |
| ENSG00000134030.16 | **CTIF** | 9811 | protein_coding | **518.813** | 1.535 | 0.462 | **2.898** | 3.51E-18 | **7.26E-17** |
| ENSG00000119650.15 | **IFT43** | 112752 | protein_coding | **410.617** | 1.026 | 0.309 | **2.036** | 3.61E-18 | **7.45E-17** |
| ENSG00000079616.16 | **KIF22** | 3835 | protein_coding | **1512.464** | -1.789 | -0.539 | **0.289** | 3.63E-18 | **7.49E-17** |
| ENSG00000102001.14 | **CACNA1F** | 778 | protein_coding | **154.407** | 1.911 | 0.575 | **3.761** | 3.88E-18 | **8.00E-17** |

| | | | | | | | | | |
|---|---|---|---|---|---|---|---|---|---|
| ENSG00000134061.6 | **CD180** | 4064 | protein_coding | **236.300** | -2.385 | -0.718 | **0.191** | 3.92E-18 | **8.07E-17** |
| ENSG00000131747.17 | **TOP2A** | 7153 | protein_coding | **1217.434** | -1.871 | -0.563 | **0.273** | 4.20E-18 | **8.63E-17** |
| ENSG00000285725.2 | **ENSG00000285725** | | lncRNA | **410.109** | -0.982 | -0.296 | **0.506** | 4.36E-18 | **8.95E-17** |
| ENSG00000136068.18 | **FLNB** | 2317 | protein_coding | **1129.131** | 1.161 | 0.350 | **2.237** | 4.39E-18 | **9.01E-17** |
| ENSG00000112249.16 | **ASCC3** | 10973 | protein_coding | **477.843** | -1.081 | -0.325 | **0.473** | 4.54E-18 | **9.31E-17** |
| ENSG00000071246.12 | **VASH1** | 22846 | protein_coding | **175.482** | 2.013 | 0.606 | **4.035** | 4.86E-18 | **9.94E-17** |
| ENSG00000070950.12 | **RAD18** | 56852 | protein_coding | **288.377** | -1.403 | -0.422 | **0.378** | 5.04E-18 | **1.03E-16** |
| ENSG00000163655.18 | **GMPS** | 8833 | protein_coding | **1193.508** | -0.954 | -0.287 | **0.516** | 5.11E-18 | **1.04E-16** |
| ENSG00000287064.1 | **ENSG00000287064** | | lncRNA | **45.943** | 4.953 | 1.491 | **30.967** | 5.27E-18 | **1.08E-16** |
| ENSG00000143847.18 | **PPFIA4** | 8497 | protein_coding | **528.583** | -1.315 | -0.396 | **0.402** | 5.31E-18 | **1.08E-16** |
| ENSG00000143179.18 | **UCK2** | 7371 | protein_coding | **857.404** | -1.120 | -0.337 | **0.460** | 5.53E-18 | **1.13E-16** |
| ENSG00000124466.9 | **LYPD3** | 27076 | protein_coding | **23.904** | 3.499 | 1.053 | **11.309** | 5.65E-18 | **1.15E-16** |
| ENSG00000183317.19 | **EPHA10** | 284656 | protein_coding | **22.116** | 4.370 | 1.315 | **20.677** | 5.82E-18 | **1.18E-16** |
| ENSG00000270659.3 | **ENSG00000270659** | | lncRNA | **100.551** | 1.491 | 0.449 | **2.811** | 5.83E-18 | **1.18E-16** |
| ENSG00000022267.21 | **FHL1** | 2273 | protein_coding | **63.229** | 2.360 | 0.710 | **5.133** | 5.84E-18 | **1.18E-16** |
| ENSG00000102996.7 | **MMP15** | 4324 | protein_coding | **36.790** | 3.291 | 0.991 | **9.785** | 5.94E-18 | **1.20E-16** |
| ENSG00000241769.7 | **EOLA1-DT** | 10013143[illegible] | lncRNA | **130.191** | 2.048 | 0.616 | **4.134** | 5.95E-18 | **1.20E-16** |
| ENSG00000150753.14 | **CCT5** | 22948 | protein_coding | **2495.342** | -0.870 | -0.262 | **0.547** | 6.08E-18 | **1.23E-16** |
| ENSG00000075826.19 | **SEC31B** | 25956 | protein_coding | **1426.720** | 0.892 | 0.269 | **1.856** | 6.12E-18 | **1.24E-16** |
| ENSG00000126603.11 | **GLIS2** | 84662 | protein_coding | **110.802** | 2.056 | 0.619 | **4.158** | 6.14E-18 | **1.24E-16** |
| ENSG00000237638.4 | **LINC02245** | 400958 | lncRNA | **541.355** | 1.106 | 0.333 | **2.152** | 6.18E-18 | **1.25E-16** |
| ENSG00000291151.2 | **NSUN5P1** | 155400 | lncRNA | **1798.908** | 0.937 | 0.282 | **1.915** | 6.27E-18 | **1.26E-16** |
| ENSG00000101856.12 | **PGRMC1** | 10857 | protein_coding | **263.396** | -1.023 | -0.308 | **0.492** | 6.58E-18 | **1.32E-16** |
| ENSG00000166313.22 | **APBB1** | 322 | protein_coding | **101.829** | 2.384 | 0.718 | **5.219** | 6.70E-18 | **1.35E-16** |
| ENSG00000161267.14 | **BDH1** | 622 | protein_coding | **439.921** | -0.871 | -0.262 | **0.547** | 6.73E-18 | **1.35E-16** |
| ENSG00000197498.15 | **RPF2** | 84154 | protein_coding | **741.021** | -1.072 | -0.323 | **0.476** | 6.78E-18 | **1.36E-16** |
| ENSG00000158106.17 | **RHPN1** | 114822 | protein_coding | **575.296** | 1.718 | 0.517 | **3.290** | 7.04E-18 | **1.41E-16** |
| ENSG00000025708.16 | **TYMP** | 1890 | protein_coding | **2714.951** | 0.986 | 0.297 | **1.981** | 7.07E-18 | **1.41E-16** |
| ENSG00000269044.2 | **ENSG00000269044** | | lncRNA | **270.024** | 1.033 | 0.311 | **2.046** | 7.40E-18 | **1.48E-16** |
| ENSG00000076770.18 | **MBNL3** | 55796 | protein_coding | **209.181** | -1.764 | -0.531 | **0.294** | 7.68E-18 | **1.53E-16** |
| ENSG00000049089.17 | **COL9A2** | 1298 | protein_coding | **4636.371** | 0.953 | 0.287 | **1.936** | 7.92E-18 | **1.58E-16** |

| | | | | | | | | | |
|---|---|---|---|---|---|---|---|---|---|
| ENSG00000125510.20 | **OPRL1** | 4987 | protein_coding | **43.103** | 2.040 | 0.614 | **4.111** | 8.03E-18 | **1.60E-16** |
| ENSG00000137364.7 | **TPMT** | 7172 | protein_coding | **323.307** | 1.046 | 0.315 | **2.065** | 8.13E-18 | **1.62E-16** |
| ENSG00000236591.2 | **UST-AS2** | | lncRNA | **291.594** | -1.951 | -0.587 | **0.259** | 8.42E-18 | **1.67E-16** |
| ENSG00000054967.15 | **RELT** | 84957 | protein_coding | **761.461** | -1.018 | -0.306 | **0.494** | 8.71E-18 | **1.73E-16** |
| ENSG00000105486.17 | **LIG1** | 3978 | protein_coding | **1276.372** | -1.277 | -0.385 | **0.413** | 8.72E-18 | **1.73E-16** |
| ENSG00000262528.2 | **ENSG00000262528** | | lncRNA | **106.822** | 2.314 | 0.696 | **4.971** | 8.81E-18 | **1.74E-16** |
| ENSG00000102038.18 | **SMARCA1** | 6594 | protein_coding | **33.126** | 3.879 | 1.168 | **14.717** | 9.22E-18 | **1.83E-16** |
| ENSG00000179403.14 | **VWA1** | 64856 | protein_coding | **227.201** | 1.849 | 0.557 | **3.603** | 9.95E-18 | **1.97E-16** |
| ENSG00000143158.13 | **MPC2** | 25874 | protein_coding | **580.566** | -0.799 | -0.240 | **0.575** | 1.02E-17 | **2.01E-16** |
| ENSG00000183833.18 | **CFAP91** | 89876 | protein_coding | **139.535** | 3.045 | 0.916 | **8.251** | 1.02E-17 | **2.01E-16** |
| ENSG00000171502.16 | **COL24A1** | 255631 | protein_coding | **490.861** | 1.524 | 0.459 | **2.875** | 1.03E-17 | **2.03E-16** |
| ENSG00000125450.13 | **NUP85** | 79902 | protein_coding | **889.945** | -1.050 | -0.316 | **0.483** | 1.03E-17 | **2.03E-16** |
| ENSG00000197380.12 | **DACT3** | 147906 | protein_coding | **72.315** | 2.885 | 0.868 | **7.386** | 1.08E-17 | **2.13E-16** |
| ENSG00000161996.21 | **WDR90** | 197335 | protein_coding | **597.293** | -1.583 | -0.476 | **0.334** | 1.11E-17 | **2.19E-16** |
| ENSG00000132376.22 | **INPP5K** | 51763 | protein_coding | **767.501** | 1.032 | 0.311 | **2.044** | 1.14E-17 | **2.25E-16** |
| ENSG00000204287.16 | **HLA-DRA** | 3122 | protein_coding | **60590.327** | 1.158 | 0.349 | **2.232** | 1.23E-17 | **2.42E-16** |
| ENSG00000131650.17 | **KREMEN2** | 79412 | protein_coding | **76.544** | 1.825 | 0.549 | **3.543** | 1.26E-17 | **2.46E-16** |
| ENSG00000137996.15 | **RTCA** | 8634 | protein_coding | **672.761** | -0.961 | -0.289 | **0.514** | 1.29E-17 | **2.53E-16** |
| ENSG00000160111.17 | **CPAMD8** | 27151 | protein_coding | **80.050** | 1.767 | 0.532 | **3.403** | 1.29E-17 | **2.53E-16** |
| ENSG00000065000.22 | **AP3D1** | 8943 | protein_coding | **3184.234** | 0.654 | 0.197 | **1.573** | 1.33E-17 | **2.60E-16** |
| ENSG00000175262.15 | **CIROZ** | 148345 | protein_coding | **161.693** | 1.993 | 0.600 | **3.981** | 1.35E-17 | **2.63E-16** |
| ENSG00000185033.17 | **SEMA4B** | 10509 | protein_coding | **640.660** | 0.822 | 0.247 | **1.767** | 1.36E-17 | **2.65E-16** |
| ENSG00000230148.9 | **HOXB-AS1** | 10087436? | lncRNA | **417.338** | 1.240 | 0.373 | **2.362** | 1.39E-17 | **2.70E-16** |
| ENSG00000213413.4 | **PVRIG** | 79037 | protein_coding | **135.900** | -2.137 | -0.643 | **0.227** | 1.46E-17 | **2.85E-16** |
| ENSG00000168005.11 | **SPINDOC** | 144097 | protein_coding | **581.491** | -1.030 | -0.310 | **0.490** | 1.48E-17 | **2.88E-16** |
| ENSG00000165716.13 | **DIPK1B** | 138311 | protein_coding | **192.386** | 2.966 | 0.893 | **7.813** | 1.50E-17 | **2.91E-16** |
| ENSG00000250506.11 | **CDK3** | 1018 | protein_coding | **153.617** | -1.630 | -0.491 | **0.323** | 1.54E-17 | **2.99E-16** |
| ENSG00000139835.16 | **GRTP1** | 79774 | protein_coding | **33.053** | 3.528 | 1.062 | **11.538** | 1.60E-17 | **3.09E-16** |
| ENSG00000178075.22 | **GRAMD1C** | 54762 | protein_coding | **36.133** | 2.827 | 0.851 | **7.094** | 1.62E-17 | **3.14E-16** |
| ENSG00000097021.23 | **ACOT7** | 11332 | protein_coding | **754.775** | -1.274 | -0.384 | **0.413** | 1.69E-17 | **3.26E-16** |
| ENSG00000147419.21 | **CCDC25** | 55246 | protein_coding | **567.715** | -0.953 | -0.287 | **0.517** | 1.72E-17 | **3.32E-16** |

| | | | | | | | | | |
|---|---|---|---|---|---|---|---|---|---|
| ENSG00000123975.7 | **CKS2** | 1164 | protein_coding | **907.186** | -1.731 | -0.521 | **0.301** | 1.75E-17 | **3.38E-16** |
| ENSG00000153879.11 | **CEBPG** | 1054 | protein_coding | **681.266** | -1.503 | -0.453 | **0.353** | 1.78E-17 | **3.43E-16** |
| ENSG00000180891.15 | **CUEDC1** | 404093 | protein_coding | **112.157** | 2.126 | 0.640 | **4.364** | 1.83E-17 | **3.52E-16** |
| ENSG00000188486.4 | **H2AX** | 3014 | protein_coding | **1634.837** | -1.579 | -0.475 | **0.335** | 1.93E-17 | **3.71E-16** |
| ENSG00000004866.24 | **ST7** | 7982 | protein_coding | **606.814** | -1.135 | -0.342 | **0.455** | 1.97E-17 | **3.79E-16** |
| ENSG00000121966.8 | **CXCR4** | 7852 | protein_coding | **395.301** | -1.235 | -0.372 | **0.425** | 1.98E-17 | **3.79E-16** |
| ENSG00000113739.12 | **STC2** | 8614 | protein_coding | **867.754** | -1.886 | -0.568 | **0.271** | 1.98E-17 | **3.79E-16** |
| ENSG00000182866.20 | **LCK** | 3932 | protein_coding | **1187.475** | -1.363 | -0.410 | **0.389** | 2.06E-17 | **3.95E-16** |
| ENSG00000112679.16 | **DUSP22** | 56940 | protein_coding | **4638.051** | 1.063 | 0.320 | **2.089** | 2.12E-17 | **4.06E-16** |
| ENSG00000128891.19 | **CCDC32** | 90416 | protein_coding | **1982.807** | 1.087 | 0.327 | **2.125** | 2.17E-17 | **4.14E-16** |
| ENSG00000196715.9 | **VKORC1L1** | 154807 | protein_coding | **446.817** | -1.342 | -0.404 | **0.395** | 2.20E-17 | **4.20E-16** |
| ENSG00000135406.16 | **PRPH** | 5630 | protein_coding | **19.584** | 4.941 | 1.487 | **30.713** | 2.27E-17 | **4.32E-16** |
| ENSG00000141385.14 | **AFG3L2** | 10939 | protein_coding | **966.367** | -0.666 | -0.200 | **0.630** | 2.35E-17 | **4.47E-16** |
| ENSG00000011600.14 | **TYROBP** | 7305 | protein_coding | **30.113** | 3.328 | 1.002 | **10.045** | 2.36E-17 | **4.49E-16** |
| ENSG00000300007.1 | **ENSG00000300007** | | lncRNA | **47.110** | 2.615 | 0.787 | **6.126** | 2.50E-17 | **4.75E-16** |
| ENSG00000107902.16 | **LHPP** | 64077 | protein_coding | **370.811** | 1.626 | 0.490 | **3.087** | 2.51E-17 | **4.76E-16** |
| ENSG00000261371.8 | **PECAM1** | 5175 | protein_coding | **227.143** | 1.414 | 0.426 | **2.665** | 2.62E-17 | **4.97E-16** |
| ENSG00000256073.5 | **URB1-AS1** | | lncRNA | **135.460** | 1.202 | 0.362 | **2.301** | 2.74E-17 | **5.19E-16** |
| ENSG00000112195.10 | **TREML2** | 79865 | protein_coding | **357.933** | -1.382 | -0.416 | **0.384** | 2.75E-17 | **5.21E-16** |
| ENSG00000163071.13 | **SPATA18** | 132671 | protein_coding | **32.438** | -2.886 | -0.869 | **0.135** | 2.85E-17 | **5.40E-16** |
| ENSG00000101298.17 | **SNPH** | 9751 | protein_coding | **46.510** | 3.410 | 1.026 | **10.628** | 2.93E-17 | **5.54E-16** |
| ENSG00000240771.10 | **ARHGEF25** | 115557 | protein_coding | **362.451** | 1.275 | 0.384 | **2.420** | 2.96E-17 | **5.60E-16** |
| ENSG00000175061.21 | **SNHG29** | 125144 | lncRNA | **9574.257** | -0.816 | -0.246 | **0.568** | 3.02E-17 | **5.69E-16** |
| ENSG00000135925.11 | **WNT10A** | 80326 | protein_coding | **1839.658** | -1.570 | -0.473 | **0.337** | 3.02E-17 | **5.70E-16** |
| ENSG00000148459.18 | **PDSS1** | 23590 | protein_coding | **140.363** | -1.348 | -0.406 | **0.393** | 3.10E-17 | **5.85E-16** |
| ENSG00000285958.2 | **ENSG00000285958** | | lncRNA | **23.589** | 4.739 | 1.427 | **26.711** | 3.15E-17 | **5.92E-16** |
| ENSG00000184371.16 | **CSF1** | 1435 | protein_coding | **527.371** | -1.128 | -0.340 | **0.457** | 3.22E-17 | **6.06E-16** |
| ENSG00000124207.19 | **CSE1L** | 1434 | protein_coding | **1144.336** | -0.944 | -0.284 | **0.520** | 3.28E-17 | **6.16E-16** |
| ENSG00000168961.19 | **LGALS9** | 3965 | protein_coding | **2382.106** | -1.303 | -0.392 | **0.405** | 3.41E-17 | **6.39E-16** |
| ENSG00000169403.14 | **PTAFR** | 5724 | protein_coding | **481.463** | -1.324 | -0.399 | **0.399** | 3.51E-17 | **6.58E-16** |
| ENSG00000137563.15 | **GGH** | 8836 | protein_coding | **233.228** | -1.516 | -0.457 | **0.350** | 3.64E-17 | **6.82E-16** |

| | | | | | | | | | |
|---|---|---|---|---|---|---|---|---|---|
| ENSG00000120699.16 | **EXOSC8** | 11340 | protein_coding | **1212.889** | -1.244 | -0.374 | **0.422** | 3.67E-17 | **6.87E-16** |
| ENSG00000006047.15 | **YBX2** | 51087 | protein_coding | **84.192** | 6.365 | 1.916 | **82.452** | 4.36E-17 | **8.14E-16** |
| ENSG00000136897.9 | **MRPL50** | 54534 | protein_coding | **444.413** | -1.103 | -0.332 | **0.465** | 4.38E-17 | **8.17E-16** |
| ENSG00000185291.14 | **IL3RA** | 3563 | protein_coding | **814.649** | 1.281 | 0.386 | **2.430** | 4.41E-17 | **8.23E-16** |
| ENSG00000115042.12 | **FAHD2A** | 51011 | protein_coding | **512.003** | -0.806 | -0.243 | **0.572** | 4.43E-17 | **8.25E-16** |
| ENSG00000135045.8 | **C9orf40** | 55071 | protein_coding | **203.791** | -1.280 | -0.385 | **0.412** | 4.54E-17 | **8.44E-16** |
| ENSG00000298169.1 | **ENSG00000298169** | | lncRNA | **56.114** | 2.343 | 0.705 | **5.075** | 4.64E-17 | **8.62E-16** |
| ENSG00000140263.17 | **SORD** | 6652 | protein_coding | **488.330** | -1.084 | -0.326 | **0.472** | 4.70E-17 | **8.73E-16** |
| ENSG00000116032.5 | **GRIN3B** | 116444 | protein_coding | **25.742** | 3.943 | 1.187 | **15.379** | 4.99E-17 | **9.25E-16** |
| ENSG00000145354.14 | **CISD2** | 493856 | protein_coding | **331.978** | -0.880 | -0.265 | **0.543** | 4.99E-17 | **9.25E-16** |
| ENSG00000183160.11 | **TMEM119** | 338773 | protein_coding | **47.918** | 2.913 | 0.877 | **7.532** | 5.00E-17 | **9.27E-16** |
| ENSG00000260805.4 | **ENSG00000260805** | | lncRNA | **44.943** | 1.964 | 0.591 | **3.902** | 5.08E-17 | **9.40E-16** |
| ENSG00000113649.16 | **TCERG1** | 10915 | protein_coding | **1428.843** | -0.797 | -0.240 | **0.575** | 5.40E-17 | **9.97E-16** |
| ENSG00000110104.14 | **CCDC86** | 79080 | protein_coding | **484.902** | -1.215 | -0.366 | **0.431** | 5.44E-17 | **1.01E-15** |
| ENSG00000239672.10 | **NME1** | 4830 | protein_coding | **2559.563** | -1.151 | -0.347 | **0.450** | 5.46E-17 | **1.01E-15** |
| ENSG00000125967.19 | **NECAB3** | 63941 | protein_coding | **293.308** | 1.176 | 0.354 | **2.260** | 5.49E-17 | **1.01E-15** |
| ENSG00000234028.5 | **EIF2AK3-DT** | | lncRNA | **167.458** | 1.929 | 0.581 | **3.807** | 5.51E-17 | **1.01E-15** |
| ENSG00000257167.4 | **TMPO-AS1** | 100128191 | lncRNA | **171.392** | -1.556 | -0.468 | **0.340** | 5.71E-17 | **1.05E-15** |
| ENSG00000124222.25 | **STX16** | 8675 | protein_coding | **4543.761** | 0.901 | 0.271 | **1.867** | 5.81E-17 | **1.07E-15** |
| ENSG00000132199.22 | **ENOSF1** | 55556 | protein_coding | **1174.875** | -1.191 | -0.358 | **0.438** | 5.82E-17 | **1.07E-15** |
| ENSG00000054148.20 | **PHPT1** | 29085 | protein_coding | **2016.211** | -0.962 | -0.290 | **0.513** | 5.98E-17 | **1.10E-15** |
| ENSG00000111145.10 | **ELK3** | 2004 | protein_coding | **147.028** | 1.748 | 0.526 | **3.358** | 6.02E-17 | **1.10E-15** |
| ENSG00000245904.7 | **BTG1-DT** | | lncRNA | **173.570** | 1.645 | 0.495 | **3.127** | 6.10E-17 | **1.12E-15** |
| ENSG00000174307.8 | **PHLDA3** | 23612 | protein_coding | **409.294** | -1.498 | -0.451 | **0.354** | 6.13E-17 | **1.12E-15** |
| ENSG00000170144.25 | **HNRNPA3** | 220988 | protein_coding | **6165.823** | -0.890 | -0.268 | **0.539** | 6.18E-17 | **1.13E-15** |
| ENSG00000163002.15 | **NUP35** | 129401 | protein_coding | **148.603** | -1.375 | -0.414 | **0.385** | 6.43E-17 | **1.17E-15** |
| ENSG00000233621.4 | **LITATS1** | 728431 | lncRNA | **71.722** | 1.833 | 0.552 | **3.563** | 6.43E-17 | **1.17E-15** |
| ENSG00000158042.11 | **MRPL17** | 63875 | protein_coding | **1269.620** | -0.996 | -0.300 | **0.501** | 6.60E-17 | **1.20E-15** |
| ENSG00000093100.13 | **ENSG00000093100** | | lncRNA | **362.218** | 2.094 | 0.630 | **4.270** | 6.64E-17 | **1.21E-15** |
| ENSG00000184557.6 | **SOCS3** | 9021 | protein_coding | **278.155** | 1.617 | 0.487 | **3.068** | 6.66E-17 | **1.21E-15** |
| ENSG00000101187.18 | **SLCO4A1** | 28231 | protein_coding | **367.799** | -1.485 | -0.447 | **0.357** | 6.90E-17 | **1.25E-15** |

| | | | | | | | | | |
|---|---|---|---|---|---|---|---|---|---|
| ENSG00000238045.10 | **MVP-DT** | | lncRNA | **197.466** | 1.447 | 0.436 | **2.726** | 7.15E-17 | **1.30E-15** |
| ENSG00000123191.20 | **ATP7B** | 540 | protein_coding | **42.705** | 3.022 | 0.910 | **8.125** | 7.24E-17 | **1.31E-15** |
| ENSG00000106144.22 | **CASP2** | 835 | protein_coding | **598.981** | -0.751 | -0.226 | **0.594** | 7.39E-17 | **1.34E-15** |
| ENSG00000102962.7 | **CCL22** | 6367 | protein_coding | **8017.249** | 1.721 | 0.518 | **3.296** | 7.99E-17 | **1.45E-15** |
| ENSG00000101638.16 | **ST8SIA5** | 29906 | protein_coding | **38.428** | 2.286 | 0.688 | **4.878** | 8.30E-17 | **1.50E-15** |
| ENSG00000278175.7 | **GLIDR** | 389741 | lncRNA | **73.175** | -1.606 | -0.483 | **0.329** | 8.57E-17 | **1.55E-15** |
| ENSG00000260192.4 | **LINC02240** | 10192746[illegible] | lncRNA | **82.685** | 2.598 | 0.782 | **6.054** | 8.62E-17 | **1.56E-15** |
| ENSG00000139190.18 | **VAMP1** | 6843 | protein_coding | **2372.099** | 1.066 | 0.321 | **2.094** | 8.92E-17 | **1.61E-15** |
| ENSG00000116128.13 | **BCL9** | 607 | protein_coding | **440.644** | 1.155 | 0.348 | **2.227** | 9.03E-17 | **1.63E-15** |
| ENSG00000243279.6 | **PRAF2** | 11230 | protein_coding | **379.338** | 1.169 | 0.352 | **2.248** | 9.08E-17 | **1.63E-15** |
| ENSG00000105402.9 | **NAPA** | 8775 | protein_coding | **3335.370** | 0.784 | 0.236 | **1.722** | 9.53E-17 | **1.71E-15** |
| ENSG00000116663.13 | **FBXO6** | 26270 | protein_coding | **390.163** | -1.103 | -0.332 | **0.465** | 9.84E-17 | **1.77E-15** |
| ENSG00000108671.13 | **PSMD11** | 5717 | protein_coding | **1174.287** | -0.767 | -0.231 | **0.588** | 1.01E-16 | **1.80E-15** |
| ENSG00000130635.19 | **COL5A1** | 1289 | protein_coding | **345.557** | -1.594 | -0.480 | **0.331** | 1.02E-16 | **1.82E-15** |
| ENSG00000107798.20 | **LIPA** | 3988 | protein_coding | **739.954** | 1.169 | 0.352 | **2.248** | 1.02E-16 | **1.83E-15** |
| ENSG00000153107.15 | **ANAPC1** | 64682 | protein_coding | **892.879** | -0.727 | -0.219 | **0.604** | 1.03E-16 | **1.85E-15** |
| ENSG00000115129.16 | **TP53I3** | 9540 | protein_coding | **223.401** | -1.428 | -0.430 | **0.372** | 1.08E-16 | **1.93E-15** |
| ENSG00000180329.16 | **CCDC43** | 124808 | protein_coding | **268.180** | -0.999 | -0.301 | **0.500** | 1.11E-16 | **1.99E-15** |
| ENSG00000001036.16 | **FUCA2** | 2519 | protein_coding | **381.403** | -0.984 | -0.296 | **0.505** | 1.12E-16 | **1.99E-15** |
| ENSG00000178700.11 | **DHFR2** | 200895 | protein_coding | **145.077** | 1.249 | 0.376 | **2.377** | 1.12E-16 | **1.99E-15** |
| ENSG00000135916.17 | **ITM2C** | 81618 | protein_coding | **3494.387** | 1.546 | 0.465 | **2.920** | 1.14E-16 | **2.03E-15** |
| ENSG00000173988.15 | **LRRC63** | 220416 | protein_coding | **23.960** | 3.327 | 1.001 | **10.034** | 1.14E-16 | **2.03E-15** |
| ENSG00000279092.1 | **ENSG00000279092** | | TEC | **134.673** | -1.606 | -0.483 | **0.329** | 1.20E-16 | **2.13E-15** |
| ENSG00000160131.16 | **VMA21** | 203547 | protein_coding | **750.404** | -1.718 | -0.517 | **0.304** | 1.21E-16 | **2.16E-15** |
| ENSG00000278817.1 | **ENSG00000278817** | | protein_coding | **51.496** | 2.828 | 0.851 | **7.100** | 1.23E-16 | **2.18E-15** |
| ENSG00000275549.1 | **STPG3-AS1** | 10012972[illegible] | lncRNA | **44.089** | 2.092 | 0.630 | **4.263** | 1.23E-16 | **2.18E-15** |
| ENSG00000214706.14 | **IFRD2** | 7866 | protein_coding | **599.725** | -1.066 | -0.321 | **0.478** | 1.25E-16 | **2.21E-15** |
| ENSG00000160211.22 | **G6PD** | 2539 | protein_coding | **1093.730** | 0.848 | 0.255 | **1.800** | 1.32E-16 | **2.34E-15** |
| ENSG00000116977.21 | **LGALS8** | 3964 | protein_coding | **1214.481** | -1.095 | -0.330 | **0.468** | 1.37E-16 | **2.41E-15** |
| ENSG00000125148.9 | **MT2A** | 4502 | protein_coding | **1073.613** | 1.375 | 0.414 | **2.594** | 1.42E-16 | **2.50E-15** |
| ENSG00000197142.13 | **ACSL5** | 51703 | protein_coding | **953.279** | -1.305 | -0.393 | **0.405** | 1.44E-16 | **2.55E-15** |

| | | | | | | | | | |
|---|---|---|---|---|---|---|---|---|---|
| ENSG00000152700.16 | **SAR1B** | 51128 | protein_coding | **767.966** | -1.028 | -0.310 | **0.490** | 1.45E-16 | **2.55E-15** |
| ENSG00000163412.15 | **EIF4E3** | 317649 | protein_coding | **42.357** | 3.260 | 0.981 | **9.581** | 1.46E-16 | **2.57E-15** |
| ENSG00000164237.11 | **CMBL** | 134147 | protein_coding | **186.597** | -1.289 | -0.388 | **0.409** | 1.54E-16 | **2.71E-15** |
| ENSG00000245694.13 | **CRNDE** | 10192748( | lncRNA | **181.998** | -1.245 | -0.375 | **0.422** | 1.62E-16 | **2.85E-15** |
| ENSG00000170779.13 | **CDCA4** | 55038 | protein_coding | **701.979** | -1.144 | -0.344 | **0.452** | 1.65E-16 | **2.89E-15** |
| ENSG00000289609.3 | **ENSG00000289609** | | lncRNA | **66.827** | 2.247 | 0.676 | **4.748** | 1.65E-16 | **2.89E-15** |
| ENSG00000090266.15 | **NDUFB2** | 4708 | protein_coding | **789.288** | -0.938 | -0.282 | **0.522** | 1.65E-16 | **2.90E-15** |
| ENSG00000023228.17 | **NDUFS1** | 4719 | protein_coding | **804.453** | -0.734 | -0.221 | **0.601** | 1.67E-16 | **2.92E-15** |
| ENSG00000106605.13 | **BLVRA** | 644 | protein_coding | **1087.195** | 0.896 | 0.270 | **1.861** | 1.68E-16 | **2.94E-15** |
| ENSG00000266208.2 | **GJD3-AS1** | | lncRNA | **75.844** | 1.716 | 0.516 | **3.284** | 1.86E-16 | **3.24E-15** |
| ENSG00000171316.16 | **CHD7** | 55636 | protein_coding | **826.316** | 1.034 | 0.311 | **2.048** | 1.87E-16 | **3.27E-15** |
| ENSG00000147576.19 | **ADHFE1** | 137872 | protein_coding | **154.568** | 1.978 | 0.595 | **3.938** | 1.90E-16 | **3.31E-15** |
| ENSG00000259207.11 | **ITGB3** | 3690 | protein_coding | **53.487** | -1.879 | -0.566 | **0.272** | 1.96E-16 | **3.41E-15** |
| ENSG00000126016.18 | **AMOT** | 154796 | protein_coding | **34.341** | 2.824 | 0.850 | **7.082** | 2.04E-16 | **3.55E-15** |
| ENSG00000255182.2 | **ENSG00000255182** | | lncRNA | **145.593** | 1.822 | 0.548 | **3.535** | 2.05E-16 | **3.57E-15** |
| ENSG00000197813.7 | **ENSG00000197813** | | lncRNA | **72.071** | 2.012 | 0.606 | **4.033** | 2.09E-16 | **3.64E-15** |
| ENSG00000186603.7 | **HPDL** | 84842 | protein_coding | **117.572** | -1.580 | -0.476 | **0.335** | 2.22E-16 | **3.85E-15** |
| ENSG00000238197.9 | **PAXBP1-AS** | 10050621! | lncRNA | **68.319** | 2.044 | 0.615 | **4.125** | 2.23E-16 | **3.86E-15** |
| ENSG00000100889.14 | **PCK2** | 5106 | protein_coding | **1568.306** | -1.297 | -0.390 | **0.407** | 2.28E-16 | **3.95E-15** |
| ENSG00000097033.17 | **SH3GLB1** | 51100 | protein_coding | **1040.311** | -0.830 | -0.250 | **0.562** | 2.33E-16 | **4.04E-15** |
| ENSG00000185340.18 | **GAS2L1** | 10634 | protein_coding | **37.083** | 3.282 | 0.988 | **9.730** | 2.36E-16 | **4.08E-15** |
| ENSG00000173638.21 | **SLC19A1** | 6573 | protein_coding | **317.443** | -1.245 | -0.375 | **0.422** | 2.50E-16 | **4.33E-15** |
| ENSG00000227502.5 | **MROCKI** | 285758 | lncRNA | **181.018** | 1.168 | 0.352 | **2.247** | 2.53E-16 | **4.36E-15** |
| ENSG00000100401.23 | **RANGAP1** | 5905 | protein_coding | **1134.814** | -0.896 | -0.270 | **0.538** | 2.59E-16 | **4.46E-15** |
| ENSG00000287104.3 | **ENSG00000287104** | | lncRNA | **155.125** | 1.392 | 0.419 | **2.624** | 2.60E-16 | **4.48E-15** |
| ENSG00000067141.19 | **NEO1** | 4756 | protein_coding | **59.516** | 2.394 | 0.721 | **5.256** | 2.68E-16 | **4.62E-15** |
| ENSG00000164654.18 | **MIOS** | 54468 | protein_coding | **715.587** | -0.807 | -0.243 | **0.572** | 2.70E-16 | **4.64E-15** |
| ENSG00000130309.14 | **COLGALT1** | 79709 | protein_coding | **807.326** | -0.665 | -0.200 | **0.630** | 2.72E-16 | **4.67E-15** |
| ENSG00000290851.2 | **ENSG00000290851** | | lncRNA | **104.058** | 1.821 | 0.548 | **3.533** | 2.72E-16 | **4.68E-15** |
| ENSG00000154237.14 | **LRRK1** | 79705 | protein_coding | **1585.778** | -0.941 | -0.283 | **0.521** | 2.74E-16 | **4.71E-15** |
| ENSG00000157353.19 | **FCSK** | 197258 | protein_coding | **620.394** | 0.781 | 0.235 | **1.718** | 2.92E-16 | **5.00E-15** |

| | | | | | | | | | |
|---|---|---|---|---|---|---|---|---|---|
| ENSG00000167969.15 | **ECI1** | 1632 | protein_coding | **712.242** | -0.879 | -0.265 | **0.544** | 3.00E-16 | **5.15E-15** |
| ENSG00000166582.13 | **CENPV** | 201161 | protein_coding | **244.584** | -1.433 | -0.432 | **0.370** | 3.04E-16 | **5.20E-15** |
| ENSG00000111087.12 | **GLI1** | 2735 | protein_coding | **102.180** | 2.284 | 0.688 | **4.872** | 3.19E-16 | **5.46E-15** |
| ENSG00000179111.10 | **HES7** | 84667 | protein_coding | **196.463** | 1.847 | 0.556 | **3.598** | 3.20E-16 | **5.46E-15** |
| ENSG00000115194.13 | **SLC30A3** | 7781 | protein_coding | **35.529** | 2.547 | 0.767 | **5.843** | 3.21E-16 | **5.48E-15** |
| ENSG00000162368.17 | **CMPK1** | 51727 | protein_coding | **1051.253** | -0.807 | -0.243 | **0.572** | 3.24E-16 | **5.52E-15** |
| ENSG00000180758.14 | **GPR157** | 80045 | protein_coding | **281.563** | 1.518 | 0.457 | **2.863** | 3.29E-16 | **5.60E-15** |
| ENSG00000101213.9 | **PTK6** | 5753 | protein_coding | **69.067** | 1.668 | 0.502 | **3.178** | 3.34E-16 | **5.68E-15** |
| ENSG00000205089.10 | **CCNI2** | 645121 | protein_coding | **696.332** | 1.164 | 0.350 | **2.241** | 3.40E-16 | **5.78E-15** |
| ENSG00000075188.11 | **NUP37** | 79023 | protein_coding | **626.210** | -0.795 | -0.239 | **0.576** | 3.41E-16 | **5.80E-15** |
| ENSG00000235162.11 | **C12orf75** | 387882 | protein_coding | **1617.874** | -0.927 | -0.279 | **0.526** | 3.50E-16 | **5.94E-15** |
| ENSG00000117399.16 | **CDC20** | 991 | protein_coding | **872.324** | -2.087 | -0.628 | **0.235** | 3.52E-16 | **5.96E-15** |
| ENSG00000108561.9 | **C1QBP** | 708 | protein_coding | **3322.368** | -0.888 | -0.267 | **0.540** | 3.62E-16 | **6.13E-15** |
| ENSG00000213337.11 | **ANKRD39** | 51239 | protein_coding | **264.229** | -1.101 | -0.331 | **0.466** | 3.62E-16 | **6.13E-15** |
| ENSG00000259132.1 | **ENSG00000259132** | | protein_coding | **244.724** | -1.459 | -0.439 | **0.364** | 3.64E-16 | **6.16E-15** |
| ENSG00000153574.11 | **RPIA** | 22934 | protein_coding | **494.145** | -0.842 | -0.253 | **0.558** | 3.66E-16 | **6.19E-15** |
| ENSG00000198915.14 | **RASGEF1A** | 221002 | protein_coding | **77.627** | 2.263 | 0.681 | **4.801** | 3.78E-16 | **6.39E-15** |
| ENSG00000169427.9 | **KCNK9** | 51305 | protein_coding | **23.407** | 4.260 | 1.282 | **19.164** | 3.82E-16 | **6.45E-15** |
| ENSG00000089472.20 | **HEPH** | 9843 | protein_coding | **59.269** | 2.514 | 0.757 | **5.713** | 3.86E-16 | **6.51E-15** |
| ENSG00000251364.10 | **SYT9-AS1** | 10050625ε | lncRNA | **221.637** | 1.327 | 0.400 | **2.509** | 3.94E-16 | **6.64E-15** |
| ENSG00000157873.20 | **TNFRSF14** | 8764 | protein_coding | **5249.678** | -0.872 | -0.262 | **0.546** | 4.06E-16 | **6.84E-15** |
| ENSG00000226067.8 | **LINC00623** | | lncRNA | **237.739** | -1.667 | -0.502 | **0.315** | 4.13E-16 | **6.95E-15** |
| ENSG00000151353.18 | **TMEM18** | 129787 | protein_coding | **506.876** | -0.885 | -0.267 | **0.541** | 4.25E-16 | **7.15E-15** |
| ENSG00000111732.13 | **AICDA** | 57379 | protein_coding | **1894.805** | 1.678 | 0.505 | **3.200** | 4.39E-16 | **7.38E-15** |
| ENSG00000033627.20 | **ATP6V0A1** | 535 | protein_coding | **2059.845** | 0.803 | 0.242 | **1.745** | 4.56E-16 | **7.65E-15** |
| ENSG00000174130.14 | **TLR6** | 10333 | protein_coding | **500.007** | -0.778 | -0.234 | **0.583** | 4.76E-16 | **7.98E-15** |
| ENSG00000065978.21 | **YBX1** | 4904 | protein_coding | **13138.362** | -0.854 | -0.257 | **0.553** | 4.81E-16 | **8.05E-15** |
| ENSG00000181418.8 | **DDN** | 23109 | protein_coding | **77.187** | -1.717 | -0.517 | **0.304** | 4.81E-16 | **8.05E-15** |
| ENSG00000105088.11 | **OLFM2** | 93145 | protein_coding | **26.189** | 5.263 | 1.584 | **38.394** | 5.38E-16 | **8.99E-15** |
| ENSG00000183762.15 | **KREMEN1** | 83999 | protein_coding | **53.301** | 2.351 | 0.708 | **5.103** | 5.40E-16 | **9.02E-15** |
| ENSG00000084234.20 | **APLP2** | 334 | protein_coding | **287.625** | 1.428 | 0.430 | **2.691** | 5.41E-16 | **9.03E-15** |

| | | | | | | | | | |
|---|---|---|---|---|---|---|---|---|---|
| ENSG00000172340.17 | **SUCLG2** | 8801 | protein_coding | **414.951** | -0.864 | -0.260 | **0.549** | 5.69E-16 | **9.48E-15** |
| ENSG00000172824.19 | **CES4A** | 283848 | protein_coding | **269.075** | 1.516 | 0.456 | **2.860** | 5.76E-16 | **9.60E-15** |
| ENSG00000166801.19 | **FAM111A** | 63901 | protein_coding | **1504.178** | -0.751 | -0.226 | **0.594** | 5.86E-16 | **9.75E-15** |
| ENSG00000142173.19 | **COL6A2** | 1292 | protein_coding | **40.074** | 3.296 | 0.992 | **9.823** | 5.96E-16 | **9.91E-15** |
| ENSG00000127337.9 | **YEATS4** | 8089 | protein_coding | **211.094** | -1.315 | -0.396 | **0.402** | 5.97E-16 | **9.91E-15** |
| ENSG00000125731.15 | **SH2D3A** | 10045 | protein_coding | **390.075** | 0.910 | 0.274 | **1.879** | 6.13E-16 | **1.02E-14** |
| ENSG00000130713.19 | **EXOSC2** | 23404 | protein_coding | **608.200** | -0.830 | -0.250 | **0.563** | 6.15E-16 | **1.02E-14** |
| ENSG00000132773.14 | **TOE1** | 114034 | protein_coding | **223.081** | -1.032 | -0.311 | **0.489** | 6.26E-16 | **1.04E-14** |
| ENSG00000086730.19 | **LAT2** | 7462 | protein_coding | **4232.627** | -0.962 | -0.290 | **0.513** | 6.31E-16 | **1.04E-14** |
| ENSG00000166004.16 | **CEP295** | 85459 | protein_coding | **350.583** | -1.131 | -0.340 | **0.457** | 6.44E-16 | **1.07E-14** |
| ENSG00000233429.9 | **HOTAIRM1** | | lncRNA | **75.571** | 1.838 | 0.553 | **3.576** | 6.45E-16 | **1.07E-14** |
| ENSG00000167615.20 | **LENG8** | 114823 | protein_coding | **13116.187** | 0.971 | 0.292 | **1.960** | 6.55E-16 | **1.08E-14** |
| ENSG00000291105.2 | **ENSG00000291105** | | lncRNA | **23.616** | 4.434 | 1.335 | **21.608** | 6.66E-16 | **1.10E-14** |
| ENSG00000140025.18 | **EFCAB11** | 90141 | protein_coding | **103.088** | -2.130 | -0.641 | **0.228** | 6.85E-16 | **1.13E-14** |
| ENSG00000287110.4 | **ACAD9-DT** | 12370652[illegible] | lncRNA | **137.651** | 1.606 | 0.483 | **3.044** | 6.89E-16 | **1.14E-14** |
| ENSG00000007866.23 | **TEAD3** | 7005 | protein_coding | **33.474** | 3.912 | 1.178 | **15.051** | 7.00E-16 | **1.15E-14** |
| ENSG00000132109.12 | **TRIM21** | 6737 | protein_coding | **751.253** | -0.936 | -0.282 | **0.523** | 7.10E-16 | **1.17E-14** |
| ENSG00000272831.3 | **ENSG00000272831** | | lncRNA | **223.416** | 1.095 | 0.330 | **2.136** | 7.31E-16 | **1.20E-14** |
| ENSG00000037042.11 | **TUBG2** | 27175 | protein_coding | **1037.416** | 1.095 | 0.330 | **2.136** | 7.33E-16 | **1.20E-14** |
| ENSG00000253368.6 | **TRNP1** | 388610 | protein_coding | **27.241** | 3.949 | 1.189 | **15.442** | 7.44E-16 | **1.22E-14** |
| ENSG00000123836.17 | **PFKFB2** | 5208 | protein_coding | **143.303** | 1.731 | 0.521 | **3.319** | 7.80E-16 | **1.28E-14** |
| ENSG00000206053.15 | **JPT2** | 90861 | protein_coding | **809.013** | -0.714 | -0.215 | **0.610** | 8.00E-16 | **1.31E-14** |
| ENSG00000100767.19 | **PAPLN** | 89932 | protein_coding | **990.647** | 1.255 | 0.378 | **2.386** | 8.16E-16 | **1.33E-14** |
| ENSG00000291232.2 | **PDE4DIPP[illegible]** | 10013205[illegible] | lncRNA | **63.955** | 1.750 | 0.527 | **3.363** | 8.48E-16 | **1.39E-14** |
| ENSG00000134809.11 | **TIMM10** | 26519 | protein_coding | **656.770** | -0.887 | -0.267 | **0.541** | 8.58E-16 | **1.40E-14** |
| ENSG00000104967.9 | **NOVA2** | 4858 | protein_coding | **25.350** | 4.554 | 1.371 | **23.495** | 8.68E-16 | **1.42E-14** |
| ENSG00000262814.10 | **MRPL12** | 6182 | protein_coding | **1728.567** | -1.022 | -0.308 | **0.492** | 8.77E-16 | **1.43E-14** |
| ENSG00000026036.23 | **RTEL1-TNFRSF6B** | | protein_coding | **263.477** | 1.048 | 0.316 | **2.068** | 8.81E-16 | **1.43E-14** |
| ENSG00000166130.16 | **IKBIP** | 121457 | protein_coding | **309.889** | -1.335 | -0.402 | **0.396** | 8.87E-16 | **1.44E-14** |
| ENSG00000105676.16 | **ARMC6** | 93436 | protein_coding | **525.669** | -1.053 | -0.317 | **0.482** | 9.02E-16 | **1.47E-14** |
| ENSG00000117595.15 | **IRF6** | 3664 | protein_coding | **46.898** | 2.391 | 0.720 | **5.246** | 9.19E-16 | **1.49E-14** |

| | | | | | | | | | |
|---|---|---|---|---|---|---|---|---|---|
| ENSG00000171703.19 | **TCEA2** | 6919 | protein_coding | **775.551** | 1.077 | 0.324 | **2.110** | 9.38E-16 | **1.52E-14** |
| ENSG00000189306.14 | **RRP7A** | 27341 | protein_coding | **1607.080** | -0.875 | -0.263 | **0.545** | 9.43E-16 | **1.53E-14** |
| ENSG00000166961.17 | **MS4A15** | 219995 | protein_coding | **36.863** | 2.782 | 0.837 | **6.876** | 9.93E-16 | **1.61E-14** |
| ENSG00000174705.15 | **SH3PXD2B** | 285590 | protein_coding | **39.164** | 2.989 | 0.900 | **7.938** | 1.00E-15 | **1.63E-14** |
| ENSG00000167173.21 | **C15orf39** | 56905 | protein_coding | **2517.492** | 0.772 | 0.232 | **1.708** | 1.03E-15 | **1.66E-14** |
| ENSG00000134594.7 | **RAB33A** | 9363 | protein_coding | **246.134** | -1.161 | -0.350 | **0.447** | 1.03E-15 | **1.67E-14** |
| ENSG00000132465.14 | **JCHAIN** | 3512 | protein_coding | **4507.394** | -1.879 | -0.566 | **0.272** | 1.04E-15 | **1.68E-14** |
| ENSG00000147251.18 | **DOCK11** | 139818 | protein_coding | **259.643** | -1.064 | -0.320 | **0.478** | 1.06E-15 | **1.71E-14** |
| ENSG00000149480.8 | **MTA2** | 9219 | protein_coding | **2335.500** | -0.597 | -0.180 | **0.661** | 1.09E-15 | **1.76E-14** |
| ENSG00000111726.15 | **CMAS** | 55907 | protein_coding | **544.549** | -0.657 | -0.198 | **0.634** | 1.09E-15 | **1.76E-14** |
| ENSG00000084463.10 | **WBP11** | 51729 | protein_coding | **1291.416** | -0.724 | -0.218 | **0.605** | 1.16E-15 | **1.87E-14** |
| ENSG00000185201.19 | **IFITM2** | 10581 | protein_coding | **2067.296** | -1.608 | -0.484 | **0.328** | 1.17E-15 | **1.88E-14** |
| ENSG00000196072.14 | **BLOC1S2** | 282991 | protein_coding | **531.347** | -0.747 | -0.225 | **0.596** | 1.20E-15 | **1.92E-14** |
| ENSG00000187840.7 | **EIF4EBP1** | 1978 | protein_coding | **1141.166** | -1.421 | -0.428 | **0.373** | 1.21E-15 | **1.94E-14** |
| ENSG00000126778.14 | **SIX1** | 6495 | protein_coding | **24.402** | 3.864 | 1.163 | **14.564** | 1.22E-15 | **1.96E-14** |
| ENSG00000136122.20 | **BORA** | 79866 | protein_coding | **355.523** | -0.972 | -0.293 | **0.510** | 1.24E-15 | **1.99E-14** |
| ENSG00000104892.19 | **KLC3** | 147700 | protein_coding | **22.304** | 4.877 | 1.468 | **29.384** | 1.32E-15 | **2.12E-14** |
| ENSG00000167721.13 | **TSR1** | 55720 | protein_coding | **856.266** | -0.926 | -0.279 | **0.526** | 1.37E-15 | **2.20E-14** |
| ENSG00000138801.11 | **PAPSS1** | 9061 | protein_coding | **768.914** | 1.045 | 0.314 | **2.063** | 1.40E-15 | **2.24E-14** |
| ENSG00000124596.19 | **OARD1** | 221443 | protein_coding | **557.010** | -0.825 | -0.248 | **0.565** | 1.42E-15 | **2.27E-14** |
| ENSG00000207751.5 | **ENSG0000(** | 107985532 | lncRNA | **101.332** | 1.446 | 0.435 | **2.725** | 1.44E-15 | **2.30E-14** |
| ENSG00000161929.16 | **SCIMP** | 388325 | protein_coding | **1584.930** | -1.405 | -0.423 | **0.378** | 1.46E-15 | **2.32E-14** |
| ENSG00000254995.4 | **STX16-NPEPL1** | | protein_coding | **219.207** | 1.480 | 0.445 | **2.789** | 1.51E-15 | **2.40E-14** |
| ENSG00000184867.16 | **ARMCX2** | 9823 | protein_coding | **285.323** | 1.196 | 0.360 | **2.291** | 1.54E-15 | **2.46E-14** |
| ENSG00000161265.16 | **U2AF1L4** | 199746 | protein_coding | **642.784** | 1.040 | 0.313 | **2.056** | 1.56E-15 | **2.48E-14** |
| ENSG00000138442.13 | **WDR12** | 55759 | protein_coding | **502.552** | -1.276 | -0.384 | **0.413** | 1.57E-15 | **2.49E-14** |
| ENSG00000166889.16 | **PATL1** | 219988 | protein_coding | **1147.047** | -0.733 | -0.221 | **0.602** | 1.61E-15 | **2.55E-14** |
| ENSG00000102096.11 | **PIM2** | 11040 | protein_coding | **7548.462** | 1.023 | 0.308 | **2.032** | 1.69E-15 | **2.69E-14** |
| ENSG00000212993.6 | **POU5F1B** | 5462 | protein_coding | **46.194** | 2.939 | 0.885 | **7.667** | 1.73E-15 | **2.75E-14** |
| ENSG00000138346.18 | **DNA2** | 1763 | protein_coding | **121.106** | -1.460 | -0.440 | **0.363** | 1.83E-15 | **2.89E-14** |
| ENSG00000182378.17 | **PLCXD1** | 55344 | protein_coding | **989.233** | -0.752 | -0.226 | **0.594** | 1.84E-15 | **2.91E-14** |

| | | | | | | | | | |
|---|---|---|---|---|---|---|---|---|---|
| ENSG00000205683.13 | **DPF3** | 8110 | protein_coding | **56.488** | 2.106 | 0.634 | **4.304** | 1.84E-15 | **2.91E-14** |
| ENSG00000009790.17 | **TRAF3IP3** | 80342 | protein_coding | **1370.180** | -1.252 | -0.377 | **0.420** | 1.88E-15 | **2.98E-14** |
| ENSG00000154719.16 | **MRPL39** | 54148 | protein_coding | **851.348** | -0.715 | -0.215 | **0.609** | 1.92E-15 | **3.02E-14** |
| ENSG00000125434.13 | **SLC25A35** | 399512 | protein_coding | **233.332** | 1.154 | 0.347 | **2.226** | 1.93E-15 | **3.04E-14** |
| ENSG00000100664.12 | **EIF5** | 1983 | protein_coding | **1724.078** | -0.982 | -0.296 | **0.506** | 1.93E-15 | **3.04E-14** |
| ENSG00000167608.14 | **TMC4** | 147798 | protein_coding | **27.613** | 4.563 | 1.374 | **23.642** | 1.98E-15 | **3.12E-14** |
| ENSG00000135698.12 | **MPHOSPH** | 10200 | protein_coding | **113.270** | -1.605 | -0.483 | **0.329** | 2.05E-15 | **3.23E-14** |
| ENSG00000188322.9 | **SBK1** | 388228 | protein_coding | **25.749** | 2.632 | 0.792 | **6.198** | 2.11E-15 | **3.32E-14** |
| ENSG00000278619.7 | **MRM1** | 79922 | protein_coding | **266.190** | -1.061 | -0.320 | **0.479** | 2.12E-15 | **3.32E-14** |
| ENSG00000146063.22 | **TRIM41** | 90933 | protein_coding | **1276.390** | 0.660 | 0.199 | **1.580** | 2.12E-15 | **3.33E-14** |
| ENSG00000259520.7 | **SLC28A2-A** | 10192841 | lncRNA | **48.091** | 1.933 | 0.582 | **3.818** | 2.13E-15 | **3.34E-14** |
| ENSG00000110852.7 | **CLEC2B** | 9976 | protein_coding | **174.125** | -1.643 | -0.495 | **0.320** | 2.13E-15 | **3.34E-14** |
| ENSG00000259877.3 | **ZNF778-DT** | 12370654 | lncRNA | **102.746** | 1.486 | 0.447 | **2.802** | 2.16E-15 | **3.38E-14** |
| ENSG00000260086.6 | **ENSG0000** | 10537124 | lncRNA | **56.541** | 1.674 | 0.504 | **3.191** | 2.23E-15 | **3.49E-14** |
| ENSG00000140403.15 | **DNAJA4** | 55466 | protein_coding | **104.091** | -1.383 | -0.416 | **0.383** | 2.29E-15 | **3.58E-14** |
| ENSG00000188825.17 | **LINC00910** | | lncRNA | **193.909** | 1.163 | 0.350 | **2.240** | 2.30E-15 | **3.59E-14** |
| ENSG00000115758.16 | **ODC1** | 4953 | protein_coding | **2760.403** | -0.897 | -0.270 | **0.537** | 2.33E-15 | **3.64E-14** |
| ENSG00000205045.14 | **SLFN12L** | 10050673 | protein_coding | **571.208** | -1.603 | -0.482 | **0.329** | 2.35E-15 | **3.67E-14** |
| ENSG00000178921.16 | **PFAS** | 5198 | protein_coding | **746.977** | -1.038 | -0.312 | **0.487** | 2.40E-15 | **3.74E-14** |
| ENSG00000205581.13 | **HMGN1** | 3150 | protein_coding | **4331.331** | -0.581 | -0.175 | **0.669** | 2.43E-15 | **3.79E-14** |
| ENSG00000142227.14 | **EMP3** | 2014 | protein_coding | **1383.504** | -1.283 | -0.386 | **0.411** | 2.48E-15 | **3.85E-14** |
| ENSG00000066827.18 | **ZFAT** | 57623 | protein_coding | **1442.267** | -1.170 | -0.352 | **0.444** | 2.48E-15 | **3.86E-14** |
| ENSG00000187837.5 | **H1-2** | 3006 | protein_coding | **3123.270** | -1.311 | -0.395 | **0.403** | 2.50E-15 | **3.88E-14** |
| ENSG00000132141.16 | **CCT6B** | 10693 | protein_coding | **85.288** | 1.860 | 0.560 | **3.630** | 2.51E-15 | **3.89E-14** |
| ENSG00000265778.5 | **ZNF516-AS** | 10192798 | lncRNA | **66.143** | 2.618 | 0.788 | **6.140** | 2.51E-15 | **3.89E-14** |
| ENSG00000308747.1 | **ENSG00000308747** | | lncRNA | **43.662** | 2.163 | 0.651 | **4.479** | 2.60E-15 | **4.02E-14** |
| ENSG00000063601.19 | **MTMR1** | 8776 | protein_coding | **552.336** | -0.826 | -0.249 | **0.564** | 2.60E-15 | **4.02E-14** |
| ENSG00000197746.17 | **PSAP** | 5660 | protein_coding | **5354.862** | 1.112 | 0.335 | **2.161** | 2.62E-15 | **4.05E-14** |
| ENSG00000125730.20 | **C3** | 718 | protein_coding | **112.111** | 2.700 | 0.813 | **6.497** | 2.70E-15 | **4.18E-14** |
| ENSG00000138709.22 | **LARP1B** | 55132 | protein_coding | **250.137** | -0.910 | -0.274 | **0.532** | 2.71E-15 | **4.18E-14** |
| ENSG00000132661.6 | **NXT1** | 29107 | protein_coding | **346.486** | -0.891 | -0.268 | **0.539** | 2.72E-15 | **4.20E-14** |

| | | | | | | | | | |
|---|---|---|---|---|---|---|---|---|---|
| ENSG00000075336.14 | **TIMM21** | 29090 | protein_coding | **281.108** | -1.080 | -0.325 | **0.473** | 2.76E-15 | **4.25E-14** |
| ENSG00000073050.14 | **XRCC1** | 7515 | protein_coding | **866.414** | -0.805 | -0.242 | **0.572** | 2.76E-15 | **4.25E-14** |
| ENSG00000131470.16 | **PSMC3IP** | 29893 | protein_coding | **262.990** | -1.574 | -0.474 | **0.336** | 2.80E-15 | **4.31E-14** |
| ENSG00000171202.9 | **TMEM126/** | 84233 | protein_coding | **263.491** | -1.105 | -0.333 | **0.465** | 2.91E-15 | **4.47E-14** |
| ENSG00000240972.4 | **MIF** | 4282 | protein_coding | **9983.872** | -1.392 | -0.419 | **0.381** | 2.97E-15 | **4.56E-14** |
| ENSG00000168385.20 | **SEPTIN2** | 4735 | protein_coding | **4858.270** | 0.695 | 0.209 | **1.619** | 3.04E-15 | **4.67E-14** |
| ENSG00000179364.17 | **PACS2** | 23241 | protein_coding | **595.425** | 1.040 | 0.313 | **2.056** | 3.07E-15 | **4.70E-14** |
| ENSG00000100413.19 | **POLR3H** | 171568 | protein_coding | **514.332** | -0.834 | -0.251 | **0.561** | 3.07E-15 | **4.70E-14** |
| ENSG00000082293.14 | **COL19A1** | 1310 | protein_coding | **259.157** | -1.523 | -0.458 | **0.348** | 3.08E-15 | **4.72E-14** |
| ENSG00000118363.15 | **SPCS2** | 9789 | protein_coding | **1084.891** | -0.839 | -0.253 | **0.559** | 3.10E-15 | **4.74E-14** |
| ENSG00000153291.17 | **SLC25A27** | 9481 | protein_coding | **29.405** | 5.076 | 1.528 | **33.728** | 3.14E-15 | **4.80E-14** |
| ENSG00000137274.16 | **BPHL** | 670 | protein_coding | **278.190** | 1.308 | 0.394 | **2.476** | 3.14E-15 | **4.80E-14** |
| ENSG00000250067.14 | **YJEFN3** | 374887 | protein_coding | **351.619** | 1.191 | 0.359 | **2.283** | 3.16E-15 | **4.82E-14** |
| ENSG00000285043.4 | **ENSG0000(** | 11269475( | protein_coding | **104.303** | -1.809 | -0.545 | **0.285** | 3.26E-15 | **4.98E-14** |
| ENSG00000213145.12 | **CRIP1** | 1396 | protein_coding | **3229.800** | 0.994 | 0.299 | **1.991** | 3.28E-15 | **4.99E-14** |
| ENSG00000151576.12 | **QTRT2** | 79691 | protein_coding | **317.618** | -1.027 | -0.309 | **0.491** | 3.36E-15 | **5.12E-14** |
| ENSG00000243678.14 | **NME2** | 4831 | protein_coding | **6760.981** | -0.961 | -0.289 | **0.514** | 3.37E-15 | **5.13E-14** |
| ENSG00000085465.15 | **OVGP1** | 5016 | protein_coding | **229.016** | 1.448 | 0.436 | **2.728** | 3.38E-15 | **5.14E-14** |
| ENSG00000162738.9 | **VANGL2** | 57216 | protein_coding | **122.957** | 1.518 | 0.457 | **2.864** | 3.40E-15 | **5.17E-14** |
| ENSG00000184441.4 | **ENSG00000184441** | | lncRNA | **252.629** | 1.287 | 0.387 | **2.440** | 3.43E-15 | **5.21E-14** |
| ENSG00000160360.14 | **GPSM1** | 26086 | protein_coding | **29.846** | 3.199 | 0.963 | **9.181** | 3.52E-15 | **5.34E-14** |
| ENSG00000185100.14 | **ADSS1** | 122622 | protein_coding | **25.047** | 3.127 | 0.941 | **8.738** | 3.53E-15 | **5.35E-14** |
| ENSG00000148218.18 | **ALAD** | 210 | protein_coding | **649.301** | 0.983 | 0.296 | **1.977** | 3.54E-15 | **5.36E-14** |
| ENSG00000101384.14 | **JAG1** | 182 | protein_coding | **339.233** | 1.601 | 0.482 | **3.033** | 3.57E-15 | **5.41E-14** |
| ENSG00000156509.15 | **FBXO43** | 286151 | protein_coding | **100.606** | -1.214 | -0.365 | **0.431** | 3.59E-15 | **5.43E-14** |
| ENSG00000145198.17 | **VWA5B2** | 90113 | protein_coding | **29.958** | 4.874 | 1.467 | **29.332** | 3.62E-15 | **5.47E-14** |
| ENSG00000100106.23 | **TRIOBP** | 11078 | protein_coding | **610.338** | -0.846 | -0.255 | **0.556** | 3.65E-15 | **5.50E-14** |
| ENSG00000160445.13 | **ZER1** | 10444 | protein_coding | **827.628** | 0.876 | 0.264 | **1.835** | 3.69E-15 | **5.56E-14** |
| ENSG00000253438.6 | **PCAT1** | 10075022! | lncRNA | **30.023** | 3.939 | 1.186 | **15.340** | 3.80E-15 | **5.73E-14** |
| ENSG00000120885.24 | **CLU** | 1191 | protein_coding | **64.715** | 2.325 | 0.700 | **5.011** | 3.92E-15 | **5.90E-14** |
| ENSG00000113845.12 | **TIMMDC1** | 51300 | protein_coding | **749.217** | 0.636 | 0.191 | **1.554** | 3.94E-15 | **5.93E-14** |

| | | | | | | | | | |
|---|---|---|---|---|---|---|---|---|---|
| ENSG00000126351.15 | **THRA** | 7067 | protein_coding | **133.906** | 1.376 | 0.414 | **2.596** | 3.95E-15 | **5.94E-14** |
| ENSG00000106355.12 | **LSM5** | 23658 | protein_coding | **637.540** | -1.071 | -0.323 | **0.476** | 4.03E-15 | **6.06E-14** |
| ENSG00000188163.8 | **CIMIP2A** | 401565 | protein_coding | **16.329** | 4.768 | 1.435 | **27.241** | 4.14E-15 | **6.22E-14** |
| ENSG00000145088.11 | **EAF2** | 55840 | protein_coding | **369.870** | -1.014 | -0.305 | **0.495** | 4.16E-15 | **6.23E-14** |
| ENSG00000268129.3 | **ENSG0000(** | 100289361 | lncRNA | **50.668** | 1.863 | 0.561 | **3.638** | 4.20E-15 | **6.30E-14** |
| ENSG00000175662.21 | **TOM1L2** | 146691 | protein_coding | **642.195** | 0.817 | 0.246 | **1.762** | 4.21E-15 | **6.30E-14** |
| ENSG00000152518.10 | **ZFP36L2** | 678 | protein_coding | **1115.557** | 1.297 | 0.390 | **2.457** | 4.24E-15 | **6.34E-14** |
| ENSG00000137312.17 | **FLOT1** | 10211 | protein_coding | **4936.178** | 0.785 | 0.236 | **1.724** | 4.34E-15 | **6.48E-14** |
| ENSG00000253125.1 | **EGR3-AS1** | | lncRNA | **119.649** | -1.471 | -0.443 | **0.361** | 4.35E-15 | **6.50E-14** |
| ENSG00000105369.11 | **CD79A** | 973 | protein_coding | **7111.108** | -1.061 | -0.319 | **0.479** | 4.36E-15 | **6.50E-14** |
| ENSG00000100297.18 | **MCM5** | 4174 | protein_coding | **2031.253** | -1.470 | -0.443 | **0.361** | 4.51E-15 | **6.73E-14** |
| ENSG00000147119.6 | **CHST7** | 56548 | protein_coding | **357.545** | 1.097 | 0.330 | **2.139** | 4.94E-15 | **7.37E-14** |
| ENSG00000005302.20 | **MSL3** | 10943 | protein_coding | **1871.531** | -0.839 | -0.253 | **0.559** | 4.95E-15 | **7.38E-14** |
| ENSG00000092036.21 | **HAUS4** | 54930 | protein_coding | **463.982** | -1.190 | -0.358 | **0.438** | 5.29E-15 | **7.87E-14** |
| ENSG00000108576.12 | **SLC6A4** | 6532 | protein_coding | **112.486** | 1.775 | 0.534 | **3.422** | 5.44E-15 | **8.08E-14** |
| ENSG00000196363.12 | **WDR5** | 11091 | protein_coding | **769.428** | -0.658 | -0.198 | **0.634** | 5.50E-15 | **8.18E-14** |
| ENSG00000020922.15 | **MRE11** | 4361 | protein_coding | **685.283** | -0.797 | -0.240 | **0.575** | 5.54E-15 | **8.23E-14** |
| ENSG00000262246.8 | **CORO7** | 79585 | protein_coding | **2918.301** | 0.755 | 0.227 | **1.688** | 5.58E-15 | **8.28E-14** |
| ENSG00000134779.17 | **TPGS2** | 25941 | protein_coding | **814.856** | -0.692 | -0.208 | **0.619** | 5.67E-15 | **8.40E-14** |
| ENSG00000139641.15 | **ESYT1** | 23344 | protein_coding | **1148.661** | -0.812 | -0.244 | **0.570** | 5.72E-15 | **8.47E-14** |
| ENSG00000289056.3 | **ENSG00000289056** | | lncRNA | **166.826** | 1.311 | 0.395 | **2.481** | 5.80E-15 | **8.59E-14** |
| ENSG00000140416.28 | **TPM1** | 7168 | protein_coding | **75.700** | 1.860 | 0.560 | **3.630** | 5.98E-15 | **8.85E-14** |
| ENSG00000244119.3 | **PDCL3P4** | | transcribed_proce | **132.463** | 1.419 | 0.427 | **2.674** | 5.99E-15 | **8.85E-14** |
| ENSG00000088836.16 | **SLC4A11** | 83959 | protein_coding | **64.720** | 2.118 | 0.638 | **4.341** | 6.10E-15 | **9.01E-14** |
| ENSG00000137106.20 | **GRHPR** | 9380 | protein_coding | **3213.650** | 0.835 | 0.251 | **1.784** | 6.40E-15 | **9.44E-14** |
| ENSG00000074660.19 | **SCARF1** | 8578 | protein_coding | **187.223** | -1.327 | -0.399 | **0.399** | 6.50E-15 | **9.59E-14** |
| ENSG00000168209.8 | **DDIT4** | 54541 | protein_coding | **4590.218** | -1.163 | -0.350 | **0.447** | 6.53E-15 | **9.62E-14** |
| ENSG00000092096.20 | **SLC22A17** | 51310 | protein_coding | **27.102** | 4.998 | 1.504 | **31.951** | 6.54E-15 | **9.63E-14** |
| ENSG00000203883.8 | **SOX18** | 54345 | protein_coding | **140.766** | 2.371 | 0.714 | **5.172** | 7.32E-15 | **1.08E-13** |
| ENSG00000104852.17 | **SNRNP70** | 6625 | protein_coding | **6951.820** | 0.605 | 0.182 | **1.521** | 7.38E-15 | **1.09E-13** |
| ENSG00000148672.12 | **GLUD1** | 2746 | protein_coding | **1642.951** | -0.643 | -0.193 | **0.641** | 7.68E-15 | **1.13E-13** |

| | | | | | | | | | |
|---|---|---|---|---|---|---|---|---|---|
| ENSG00000167196.16 | **FBXO22** | 26263 | protein_coding | **552.266** | -1.001 | -0.301 | **0.499** | 7.70E-15 | **1.13E-13** |
| ENSG00000002016.20 | **RAD52** | 5893 | protein_coding | **364.625** | 0.948 | 0.285 | **1.929** | 7.70E-15 | **1.13E-13** |
| ENSG00000003756.19 | **RBM5** | 10181 | protein_coding | **2986.793** | -0.637 | -0.192 | **0.643** | 7.71E-15 | **1.13E-13** |
| ENSG00000179862.7 | **CITED4** | 163732 | protein_coding | **30.162** | 3.494 | 1.052 | **11.270** | 7.76E-15 | **1.14E-13** |
| ENSG00000232104.6 | **RFX3-DT** | | lncRNA | **615.664** | 1.078 | 0.325 | **2.111** | 7.80E-15 | **1.14E-13** |
| ENSG00000227619.4 | **ENSG0000(** | 10537629Z | lncRNA | **66.887** | 1.857 | 0.559 | **3.622** | 7.98E-15 | **1.17E-13** |
| ENSG00000233850.2 | **MAL-AS1** | | lncRNA | **33.759** | 2.736 | 0.824 | **6.663** | 8.06E-15 | **1.18E-13** |
| ENSG00000130643.11 | **CALY** | 50632 | protein_coding | **19.938** | 5.493 | 1.654 | **45.033** | 8.14E-15 | **1.19E-13** |
| ENSG00000107371.16 | **EXOSC3** | 51010 | protein_coding | **442.501** | -0.872 | -0.262 | **0.547** | 8.21E-15 | **1.20E-13** |
| ENSG00000141560.17 | **FN3KRP** | 79672 | protein_coding | **602.676** | -0.700 | -0.211 | **0.616** | 8.39E-15 | **1.22E-13** |
| ENSG00000174695.11 | **TMEM167/** | 153339 | protein_coding | **604.662** | -1.006 | -0.303 | **0.498** | 8.40E-15 | **1.22E-13** |
| ENSG00000086475.17 | **SEPHS1** | 22929 | protein_coding | **758.756** | -0.776 | -0.234 | **0.584** | 8.46E-15 | **1.23E-13** |
| ENSG00000117122.16 | **MFAP2** | 4237 | protein_coding | **16.593** | 5.991 | 1.803 | **63.604** | 8.48E-15 | **1.23E-13** |
| ENSG00000005884.20 | **ITGA3** | 3675 | protein_coding | **244.985** | 1.390 | 0.419 | **2.622** | 8.51E-15 | **1.24E-13** |
| ENSG00000146826.19 | **TRAPPC14** | 55262 | protein_coding | **753.989** | 0.806 | 0.243 | **1.748** | 8.52E-15 | **1.24E-13** |
| ENSG00000010295.22 | **IFFO1** | 25900 | protein_coding | **1014.493** | 0.777 | 0.234 | **1.714** | 8.52E-15 | **1.24E-13** |
| ENSG00000114346.16 | **ECT2** | 1894 | protein_coding | **110.052** | -1.775 | -0.534 | **0.292** | 8.78E-15 | **1.27E-13** |
| ENSG00000186529.18 | **CYP4F3** | 4051 | protein_coding | **31.035** | -4.093 | -1.232 | **0.059** | 9.13E-15 | **1.32E-13** |
| ENSG00000117877.12 | **POLR1G** | 10849 | protein_coding | **157.813** | -1.274 | -0.384 | **0.413** | 9.17E-15 | **1.33E-13** |
| ENSG00000255529.10 | **POLR2M** | 81488 | protein_coding | **365.593** | -1.092 | -0.329 | **0.469** | 9.24E-15 | **1.34E-13** |
| ENSG00000106268.18 | **NUDT1** | 4521 | protein_coding | **432.387** | -1.470 | -0.443 | **0.361** | 9.29E-15 | **1.34E-13** |
| ENSG00000152082.17 | **MZT2B** | 80097 | protein_coding | **2454.783** | -1.012 | -0.305 | **0.496** | 9.34E-15 | **1.35E-13** |
| ENSG00000087088.23 | **BAX** | 581 | protein_coding | **2027.117** | -0.880 | -0.265 | **0.544** | 9.34E-15 | **1.35E-13** |
| ENSG00000171298.17 | **GAA** | 2548 | protein_coding | **437.339** | 0.892 | 0.268 | **1.855** | 9.35E-15 | **1.35E-13** |
| ENSG00000149257.18 | **SERPINH1** | 871 | protein_coding | **431.892** | -0.872 | -0.263 | **0.546** | 9.39E-15 | **1.35E-13** |
| ENSG00000006042.14 | **TMEM98** | 26022 | protein_coding | **21.853** | 3.627 | 1.092 | **12.356** | 9.77E-15 | **1.41E-13** |
| ENSG00000163933.13 | **RFT1** | 91869 | protein_coding | **448.883** | -0.896 | -0.270 | **0.537** | 1.00E-14 | **1.44E-13** |
| ENSG00000181885.20 | **CLDN7** | 1366 | protein_coding | **22.483** | 3.477 | 1.047 | **11.133** | 1.02E-14 | **1.47E-13** |
| ENSG00000125821.15 | **DTD1** | 92675 | protein_coding | **604.960** | -0.788 | -0.237 | **0.579** | 1.03E-14 | **1.48E-13** |
| ENSG00000160325.17 | **CACFD1** | 11094 | protein_coding | **183.592** | 1.367 | 0.412 | **2.580** | 1.03E-14 | **1.48E-13** |
| ENSG00000077454.19 | **LRCH4** | 4034 | protein_coding | **1601.644** | 0.847 | 0.255 | **1.799** | 1.04E-14 | **1.48E-13** |

| | | | | | | | | | |
|---|---|---|---|---|---|---|---|---|---|
| ENSG00000227268.5 | **KLLN** | 10014474[illegible] | protein_coding | **72.507** | 1.754 | 0.528 | **3.372** | 1.04E-14 | **1.50E-13** |
| ENSG00000213397.14 | **HAUS7** | 55559 | protein_coding | **316.677** | -1.542 | -0.464 | **0.343** | 1.06E-14 | **1.52E-13** |
| ENSG00000166831.11 | **RBPMS2** | 348093 | protein_coding | **169.994** | -1.232 | -0.371 | **0.426** | 1.09E-14 | **1.55E-13** |
| ENSG00000160255.21 | **ITGB2** | 3689 | protein_coding | **2459.386** | -1.049 | -0.316 | **0.483** | 1.11E-14 | **1.59E-13** |
| ENSG00000136878.16 | **USP20** | 10868 | protein_coding | **887.851** | 0.728 | 0.219 | **1.656** | 1.12E-14 | **1.61E-13** |
| ENSG00000168938.8 | **PPIC** | 5480 | protein_coding | **30.446** | 3.297 | 0.992 | **9.826** | 1.13E-14 | **1.61E-13** |
| ENSG00000206418.6 | **RAB12** | 201475 | protein_coding | **501.299** | 0.930 | 0.280 | **1.905** | 1.13E-14 | **1.62E-13** |
| ENSG00000132128.20 | **LRRC41** | 10489 | protein_coding | **1073.507** | -0.895 | -0.269 | **0.538** | 1.15E-14 | **1.63E-13** |
| ENSG00000011304.24 | **PTBP1** | 5725 | protein_coding | **4823.715** | -0.654 | -0.197 | **0.636** | 1.16E-14 | **1.65E-13** |
| ENSG00000127483.22 | **HP1BP3** | 50809 | protein_coding | **2119.377** | -1.101 | -0.332 | **0.466** | 1.18E-14 | **1.67E-13** |
| ENSG00000100504.19 | **PYGL** | 5836 | protein_coding | **139.001** | 1.874 | 0.564 | **3.666** | 1.18E-14 | **1.68E-13** |
| ENSG00000183486.16 | **MX2** | 4600 | protein_coding | **2260.737** | -1.013 | -0.305 | **0.496** | 1.19E-14 | **1.69E-13** |
| ENSG00000298833.1 | **ENSG0000[illegible]** | 10536990[illegible] | lncRNA | **86.962** | -1.521 | -0.458 | **0.348** | 1.23E-14 | **1.75E-13** |
| ENSG00000143418.22 | **CERS2** | 29956 | protein_coding | **883.024** | -0.733 | -0.221 | **0.602** | 1.25E-14 | **1.78E-13** |
| ENSG00000158815.12 | **FGF17** | 8822 | protein_coding | **51.580** | 2.161 | 0.650 | **4.472** | 1.26E-14 | **1.78E-13** |
| ENSG00000198771.13 | **RCSD1** | 92241 | protein_coding | **1405.952** | -0.866 | -0.261 | **0.549** | 1.27E-14 | **1.79E-13** |
| ENSG00000131652.16 | **THOC6** | 79228 | protein_coding | **517.055** | -0.960 | -0.289 | **0.514** | 1.30E-14 | **1.85E-13** |
| ENSG00000143479.19 | **DYRK3** | 8444 | protein_coding | **206.014** | 0.947 | 0.285 | **1.928** | 1.33E-14 | **1.88E-13** |
| ENSG00000115414.23 | **FN1** | 2335 | protein_coding | **69.887** | 3.600 | 1.084 | **12.128** | 1.34E-14 | **1.90E-13** |
| ENSG00000254531.4 | **PPP3CA-DT** | | lncRNA | **117.817** | 1.623 | 0.489 | **3.081** | 1.35E-14 | **1.90E-13** |
| ENSG00000196664.6 | **TLR7** | 51284 | protein_coding | **335.693** | -1.370 | -0.412 | **0.387** | 1.36E-14 | **1.92E-13** |
| ENSG00000267100.3 | **ILF3-DT** | 147727 | lncRNA | **288.163** | 0.970 | 0.292 | **1.959** | 1.37E-14 | **1.94E-13** |
| ENSG00000085563.17 | **ABCB1** | 5243 | protein_coding | **85.737** | 2.360 | 0.710 | **5.134** | 1.38E-14 | **1.95E-13** |
| ENSG00000125744.14 | **RTN2** | 6253 | protein_coding | **265.209** | 1.111 | 0.335 | **2.160** | 1.41E-14 | **1.99E-13** |
| ENSG00000198917.15 | **SPOUT1** | 51490 | protein_coding | **648.180** | -0.861 | -0.259 | **0.551** | 1.41E-14 | **1.99E-13** |
| ENSG00000198700.12 | **IPO9** | 55705 | protein_coding | **1286.742** | -0.904 | -0.272 | **0.534** | 1.46E-14 | **2.06E-13** |
| ENSG00000289019.2 | **ENSG00000289019** | | lncRNA | **18.868** | 3.694 | 1.112 | **12.938** | 1.48E-14 | **2.08E-13** |
| ENSG00000153944.14 | **MSI2** | 124540 | protein_coding | **816.122** | -0.882 | -0.265 | **0.543** | 1.54E-14 | **2.17E-13** |
| ENSG00000272269.3 | **NUP153-AS1** | | lncRNA | **140.192** | -1.580 | -0.476 | **0.334** | 1.55E-14 | **2.17E-13** |
| ENSG00000270964.1 | **MAP2K5-D** | 11873229[illegible] | lncRNA | **30.993** | 2.246 | 0.676 | **4.745** | 1.55E-14 | **2.18E-13** |
| ENSG00000277969.3 | **ENSG00000277969** | | lncRNA | **323.716** | 1.072 | 0.323 | **2.103** | 1.62E-14 | **2.27E-13** |

| | | | | | | | | | |
|---|---|---|---|---|---|---|---|---|---|
| ENSG00000177646.22 | **ACAD9** | 28976 | protein_coding | **686.119** | -0.819 | -0.247 | **0.567** | 1.63E-14 | **2.28E-13** |
| ENSG00000005379.18 | **TSPOAP1** | 9256 | protein_coding | **426.592** | 1.571 | 0.473 | **2.972** | 1.68E-14 | **2.35E-13** |
| ENSG00000127980.18 | **PEX1** | 5189 | protein_coding | **280.910** | 1.064 | 0.320 | **2.091** | 1.69E-14 | **2.37E-13** |
| ENSG00000166793.13 | **YPEL4** | 219539 | protein_coding | **75.225** | 1.714 | 0.516 | **3.281** | 1.71E-14 | **2.39E-13** |
| ENSG00000075407.21 | **ZNF37A** | 7587 | protein_coding | **465.047** | -0.915 | -0.275 | **0.531** | 1.72E-14 | **2.40E-13** |
| ENSG00000177548.15 | **RABEP2** | 79874 | protein_coding | **722.123** | -0.865 | -0.260 | **0.549** | 1.73E-14 | **2.42E-13** |
| ENSG00000254777.6 | **ENSG00000254777** | | lncRNA | **52.267** | 1.729 | 0.521 | **3.316** | 1.75E-14 | **2.44E-13** |
| ENSG00000197930.15 | **ERO1A** | 30001 | protein_coding | **1515.706** | -0.994 | -0.299 | **0.502** | 1.76E-14 | **2.45E-13** |
| ENSG00000137161.20 | **CNPY3** | 10695 | protein_coding | **2149.340** | -0.997 | -0.300 | **0.501** | 1.76E-14 | **2.45E-13** |
| ENSG00000115053.19 | **NCL** | 4691 | protein_coding | **9787.996** | -0.776 | -0.234 | **0.584** | 1.78E-14 | **2.47E-13** |
| ENSG00000113013.18 | **HSPA9** | 3313 | protein_coding | **5543.372** | -0.957 | -0.288 | **0.515** | 1.81E-14 | **2.52E-13** |
| ENSG00000167619.14 | **TMEM145** | 284339 | protein_coding | **89.636** | 2.580 | 0.777 | **5.979** | 1.82E-14 | **2.53E-13** |
| ENSG00000169228.16 | **RAB24** | 53917 | protein_coding | **1516.292** | 0.911 | 0.274 | **1.880** | 1.83E-14 | **2.54E-13** |
| ENSG00000182557.9 | **SPNS3** | 201305 | protein_coding | **49.600** | -2.269 | -0.683 | **0.207** | 1.83E-14 | **2.54E-13** |
| ENSG00000128872.12 | **TMOD2** | 29767 | protein_coding | **231.223** | 1.295 | 0.390 | **2.454** | 1.86E-14 | **2.58E-13** |
| ENSG00000087250.11 | **MT3** | 4504 | protein_coding | **30.344** | 4.874 | 1.467 | **29.321** | 1.91E-14 | **2.64E-13** |
| ENSG00000204252.16 | **HLA-DOA** | 3111 | protein_coding | **1872.150** | 1.192 | 0.359 | **2.284** | 1.91E-14 | **2.64E-13** |
| ENSG00000165475.17 | **CRYL1** | 51084 | protein_coding | **319.621** | 1.009 | 0.304 | **2.012** | 1.94E-14 | **2.68E-13** |
| ENSG00000128815.22 | **WDFY4** | 57705 | protein_coding | **2523.915** | -1.086 | -0.327 | **0.471** | 1.96E-14 | **2.71E-13** |
| ENSG00000129625.15 | **REEP5** | 7905 | protein_coding | **1265.294** | -0.695 | -0.209 | **0.618** | 1.97E-14 | **2.73E-13** |
| ENSG00000181026.17 | **AEN** | 64782 | protein_coding | **782.391** | -0.839 | -0.253 | **0.559** | 2.06E-14 | **2.85E-13** |
| ENSG00000103091.17 | **WDR59** | 79726 | protein_coding | **1745.983** | 0.718 | 0.216 | **1.645** | 2.07E-14 | **2.86E-13** |
| ENSG00000159399.12 | **HK2** | 3099 | protein_coding | **906.175** | -1.143 | -0.344 | **0.453** | 2.13E-14 | **2.93E-13** |
| ENSG00000183060.18 | **LYSMD4** | 145748 | protein_coding | **321.652** | 1.022 | 0.308 | **2.031** | 2.17E-14 | **2.99E-13** |
| ENSG00000177602.5 | **HASPIN** | 83903 | protein_coding | **71.090** | -1.625 | -0.489 | **0.324** | 2.21E-14 | **3.04E-13** |
| ENSG00000124574.18 | **ABCC10** | 89845 | protein_coding | **876.448** | 1.361 | 0.410 | **2.569** | 2.24E-14 | **3.08E-13** |
| ENSG00000145687.19 | **SSBP2** | 23635 | protein_coding | **598.789** | -0.958 | -0.288 | **0.515** | 2.25E-14 | **3.09E-13** |
| ENSG00000163945.21 | **UVSSA** | 57654 | protein_coding | **1015.819** | 0.743 | 0.224 | **1.674** | 2.34E-14 | **3.21E-13** |
| ENSG00000139508.17 | **SLC46A3** | 283537 | protein_coding | **284.396** | 1.011 | 0.304 | **2.016** | 2.36E-14 | **3.24E-13** |
| ENSG00000274070.3 | **CASTOR2** | 729438 | protein_coding | **73.628** | 1.716 | 0.517 | **3.285** | 2.42E-14 | **3.32E-13** |
| ENSG00000171219.11 | **CDC42BPG** | 55561 | protein_coding | **78.984** | 2.166 | 0.652 | **4.487** | 2.46E-14 | **3.37E-13** |

| | | | | | | | | | |
|---|---|---|---|---|---|---|---|---|---|
| ENSG00000262580.5 | **ENSG00000262580** | | lncRNA | **442.170** | 1.381 | 0.416 | **2.604** | 2.55E-14 | **3.50E-13** |
| ENSG00000274253.7 | **ENSG0000(** | 283683 | lncRNA | **21.138** | 3.510 | 1.057 | **11.392** | 2.58E-14 | **3.53E-13** |
| ENSG00000103067.16 | **ESRP2** | 80004 | protein_coding | **24.148** | 2.941 | 0.885 | **7.681** | 2.59E-14 | **3.54E-13** |
| ENSG00000143458.14 | **GABPB2** | 126626 | protein_coding | **257.132** | -0.984 | -0.296 | **0.506** | 2.59E-14 | **3.54E-13** |
| ENSG00000101160.17 | **CTSZ** | 1522 | protein_coding | **3419.448** | 0.606 | 0.183 | **1.522** | 2.61E-14 | **3.56E-13** |
| ENSG00000308666.1 | **ENSG00000308666** | | lncRNA | **17.762** | 5.163 | 1.554 | **35.826** | 2.61E-14 | **3.56E-13** |
| ENSG00000214279.14 | **SCART1** | 619207 | protein_coding | **202.471** | 1.774 | 0.534 | **3.420** | 2.70E-14 | **3.68E-13** |
| ENSG00000137171.17 | **KLC4** | 89953 | protein_coding | **550.099** | 0.913 | 0.275 | **1.882** | 2.73E-14 | **3.72E-13** |
| ENSG00000106105.17 | **GARS1** | 2617 | protein_coding | **3012.559** | -1.257 | -0.379 | **0.418** | 2.88E-14 | **3.92E-13** |
| ENSG00000151117.10 | **TMEM86A** | 144110 | protein_coding | **22.977** | 3.309 | 0.996 | **9.910** | 2.93E-14 | **3.98E-13** |
| ENSG00000020181.20 | **ADGRA2** | 25960 | protein_coding | **35.421** | 3.376 | 1.016 | **10.380** | 2.93E-14 | **3.99E-13** |
| ENSG00000018189.15 | **RUFY3** | 22902 | protein_coding | **718.795** | 1.034 | 0.311 | **2.048** | 3.02E-14 | **4.10E-13** |
| ENSG00000152240.15 | **HAUS1** | 115106 | protein_coding | **552.194** | -0.743 | -0.224 | **0.597** | 3.08E-14 | **4.18E-13** |
| ENSG00000177409.14 | **SAMD9L** | 219285 | protein_coding | **1143.002** | -1.415 | -0.426 | **0.375** | 3.16E-14 | **4.28E-13** |
| ENSG00000167123.22 | **CERCAM** | 51148 | protein_coding | **247.749** | 1.837 | 0.553 | **3.573** | 3.17E-14 | **4.30E-13** |
| ENSG00000143382.18 | **ADAMTSL4** | 54507 | protein_coding | **26.656** | 4.545 | 1.368 | **23.348** | 3.18E-14 | **4.31E-13** |
| ENSG00000105778.22 | **AVL9** | 23080 | protein_coding | **485.794** | -0.996 | -0.300 | **0.501** | 3.26E-14 | **4.41E-13** |
| ENSG00000105135.18 | **HACL2** | 10994 | protein_coding | **492.144** | -0.930 | -0.280 | **0.525** | 3.27E-14 | **4.42E-13** |
| ENSG00000113140.13 | **SPARC** | 6678 | protein_coding | **441.325** | -1.144 | -0.344 | **0.452** | 3.28E-14 | **4.44E-13** |
| ENSG00000107262.27 | **BAG1** | 573 | protein_coding | **731.752** | -0.800 | -0.241 | **0.574** | 3.40E-14 | **4.59E-13** |
| ENSG00000113621.17 | **TXNDC15** | 79770 | protein_coding | **543.802** | -1.122 | -0.338 | **0.460** | 3.41E-14 | **4.60E-13** |
| ENSG00000142459.11 | **EVI5L** | 115704 | protein_coding | **475.154** | 1.071 | 0.322 | **2.100** | 3.42E-14 | **4.61E-13** |
| ENSG00000096746.20 | **HNRNPH3** | 3189 | protein_coding | **2395.500** | -0.822 | -0.247 | **0.566** | 3.51E-14 | **4.73E-13** |
| ENSG00000187736.16 | **NHEJ1** | 79840 | protein_coding | **203.369** | -1.003 | -0.302 | **0.499** | 3.59E-14 | **4.84E-13** |
| ENSG00000229962.3 | **ENSG0000(** | 10537275: | lncRNA | **48.732** | 2.095 | 0.631 | **4.273** | 3.69E-14 | **4.97E-13** |
| ENSG00000286989.2 | **ENSG00000286989** | | lncRNA | **45.622** | 2.496 | 0.751 | **5.640** | 3.69E-14 | **4.97E-13** |
| ENSG00000214114.11 | **MYCBP** | 26292 | protein_coding | **448.301** | -1.254 | -0.378 | **0.419** | 3.71E-14 | **4.99E-13** |
| ENSG00000055130.20 | **CUL1** | 8454 | protein_coding | **1077.628** | -0.654 | -0.197 | **0.636** | 3.75E-14 | **5.03E-13** |
| ENSG00000076248.14 | **UNG** | 7374 | protein_coding | **482.719** | -1.252 | -0.377 | **0.420** | 3.79E-14 | **5.09E-13** |
| ENSG00000198157.13 | **HMGN5** | 79366 | protein_coding | **252.691** | -1.498 | -0.451 | **0.354** | 3.89E-14 | **5.22E-13** |
| ENSG00000011523.17 | **CEP68** | 23177 | protein_coding | **363.678** | 1.000 | 0.301 | **2.001** | 3.90E-14 | **5.23E-13** |

| | | | | | | | | | |
|---|---|---|---|---|---|---|---|---|---|
| ENSG00000257345.3 | **LINC02413** | | lncRNA | **73.689** | 1.739 | 0.524 | **3.339** | 3.95E-14 | **5.29E-13** |
| ENSG00000215788.13 | **TNFRSF25** | 8718 | protein_coding | **33.074** | 3.000 | 0.903 | **7.998** | 4.10E-14 | **5.48E-13** |
| ENSG00000105245.11 | **NUMBL** | 9253 | protein_coding | **278.803** | 1.219 | 0.367 | **2.328** | 4.11E-14 | **5.50E-13** |
| ENSG00000125657.6 | **TNFSF9** | 8744 | protein_coding | **2507.171** | 0.719 | 0.216 | **1.646** | 4.16E-14 | **5.56E-13** |
| ENSG00000166477.15 | **LEO1** | 123169 | protein_coding | **490.822** | -0.654 | -0.197 | **0.636** | 4.18E-14 | **5.58E-13** |
| ENSG00000135253.17 | **KCP** | 375616 | protein_coding | **213.523** | 0.943 | 0.284 | **1.922** | 4.18E-14 | **5.58E-13** |
| ENSG00000154059.13 | **IMPACT** | 55364 | protein_coding | **165.103** | 1.184 | 0.356 | **2.272** | 4.21E-14 | **5.61E-13** |
| ENSG00000107099.19 | **DOCK8** | 81704 | protein_coding | **1156.317** | -0.855 | -0.257 | **0.553** | 4.26E-14 | **5.67E-13** |
| ENSG00000115561.18 | **CHMP3** | 51652 | protein_coding | **385.167** | -0.751 | -0.226 | **0.594** | 4.27E-14 | **5.68E-13** |
| ENSG00000273230.1 | **ENSG00000273230** | | lncRNA | **43.075** | 2.105 | 0.634 | **4.303** | 4.30E-14 | **5.73E-13** |
| ENSG00000163170.14 | **BOLA3** | 388962 | protein_coding | **391.290** | -1.086 | -0.327 | **0.471** | 4.54E-14 | **6.04E-13** |
| ENSG00000294562.1 | **ENSG00000294562** | | lncRNA | **22.524** | 4.246 | 1.278 | **18.978** | 4.59E-14 | **6.10E-13** |
| ENSG00000174996.14 | **KLC2** | 64837 | protein_coding | **915.789** | 0.647 | 0.195 | **1.566** | 4.70E-14 | **6.24E-13** |
| ENSG00000067836.15 | **ROGDI** | 79641 | protein_coding | **507.560** | 1.019 | 0.307 | **2.026** | 4.82E-14 | **6.40E-13** |
| ENSG00000154889.19 | **MPPE1** | 65258 | protein_coding | **611.718** | 1.040 | 0.313 | **2.057** | 4.88E-14 | **6.47E-13** |
| ENSG00000102904.17 | **TSNAXIP1** | 55815 | protein_coding | **32.983** | 2.304 | 0.694 | **4.939** | 4.91E-14 | **6.51E-13** |
| ENSG00000141756.22 | **FKBP10** | 60681 | protein_coding | **196.631** | 2.029 | 0.611 | **4.080** | 5.01E-14 | **6.63E-13** |
| ENSG00000198783.6 | **ZNF830** | 91603 | protein_coding | **222.058** | -1.162 | -0.350 | **0.447** | 5.01E-14 | **6.64E-13** |
| ENSG00000227888.5 | **FAM66A** | | lncRNA | **83.226** | 1.364 | 0.411 | **2.574** | 5.06E-14 | **6.69E-13** |
| ENSG00000099341.14 | **PSMD8** | 5714 | protein_coding | **1741.699** | -0.751 | -0.226 | **0.594** | 5.09E-14 | **6.73E-13** |
| ENSG00000109917.13 | **ZPR1** | 8882 | protein_coding | **763.976** | -0.899 | -0.271 | **0.536** | 5.13E-14 | **6.77E-13** |
| ENSG00000168003.21 | **SLC3A2** | 6520 | protein_coding | **2570.790** | -1.329 | -0.400 | **0.398** | 5.13E-14 | **6.77E-13** |
| ENSG00000213988.12 | **ZNF90** | 7643 | protein_coding | **214.748** | -1.391 | -0.419 | **0.381** | 5.18E-14 | **6.83E-13** |
| ENSG00000118495.23 | **PLAGL1** | 5325 | protein_coding | **145.397** | -1.264 | -0.381 | **0.416** | 5.19E-14 | **6.84E-13** |
| ENSG00000010256.14 | **UQCRC1** | 7384 | protein_coding | **1891.583** | -0.872 | -0.263 | **0.546** | 5.46E-14 | **7.19E-13** |
| ENSG00000289009.3 | **ENSG00000289009** | | lncRNA | **81.186** | 1.403 | 0.422 | **2.645** | 5.49E-14 | **7.23E-13** |
| ENSG00000100304.14 | **TTLL12** | 23170 | protein_coding | **636.306** | -0.910 | -0.274 | **0.532** | 5.56E-14 | **7.31E-13** |
| ENSG00000136717.17 | **BIN1** | 274 | protein_coding | **701.006** | -0.748 | -0.225 | **0.595** | 5.69E-14 | **7.48E-13** |
| ENSG00000288859.2 | **H2AC19** | 723790 | protein_coding | **587.201** | -1.354 | -0.408 | **0.391** | 5.71E-14 | **7.50E-13** |
| ENSG00000162882.17 | **HAAO** | 23498 | protein_coding | **453.689** | -1.658 | -0.499 | **0.317** | 5.73E-14 | **7.52E-13** |
| ENSG00000159958.8 | **TNFRSF13(** | 115650 | protein_coding | **2744.600** | -1.098 | -0.331 | **0.467** | 5.81E-14 | **7.62E-13** |

| | | | | | | | | | |
|---|---|---|---|---|---|---|---|---|---|
| ENSG00000204044.6 | **SLC12A5-A** | 10972918[illegible] | lncRNA | **16.088** | 5.162 | 1.554 | **35.809** | 5.83E-14 | **7.63E-13** |
| ENSG00000174442.14 | **ZWILCH** | 55055 | protein_coding | **292.190** | -1.108 | -0.334 | **0.464** | 5.83E-14 | **7.63E-13** |
| ENSG00000119042.20 | **SATB2** | 23314 | protein_coding | **29.950** | 2.979 | 0.897 | **7.885** | 5.84E-14 | **7.64E-13** |
| ENSG00000099977.18 | **DDT** | 1652 | protein_coding | **1214.694** | -0.836 | -0.252 | **0.560** | 6.14E-14 | **8.03E-13** |
| ENSG00000117748.12 | **RPA2** | 6118 | protein_coding | **1025.577** | -0.703 | -0.212 | **0.614** | 6.15E-14 | **8.04E-13** |
| ENSG00000205436.10 | **EXOC3L4** | 91828 | protein_coding | **16.012** | 5.377 | 1.619 | **41.560** | 6.16E-14 | **8.04E-13** |
| ENSG00000129991.16 | **TNNI3** | 7137 | protein_coding | **17.120** | 6.018 | 1.811 | **64.783** | 6.45E-14 | **8.41E-13** |
| ENSG00000137218.12 | **FRS3** | 10817 | protein_coding | **183.401** | 1.195 | 0.360 | **2.289** | 6.48E-14 | **8.45E-13** |
| ENSG00000160783.21 | **PMF1** | 11243 | protein_coding | **726.685** | -1.015 | -0.306 | **0.495** | 6.54E-14 | **8.52E-13** |
| ENSG00000164284.17 | **GRPEL2** | 134266 | protein_coding | **209.047** | -1.192 | -0.359 | **0.438** | 6.57E-14 | **8.56E-13** |
| ENSG00000137871.23 | **ZNF280D** | 54816 | protein_coding | **235.764** | -1.221 | -0.367 | **0.429** | 6.75E-14 | **8.79E-13** |
| ENSG00000225062.3 | **CATIP-AS1** | 10192851[illegible] | lncRNA | **31.548** | 2.350 | 0.708 | **5.099** | 6.77E-14 | **8.81E-13** |
| ENSG00000176973.10 | **FAM89B** | 23625 | protein_coding | **217.440** | 1.473 | 0.443 | **2.776** | 6.80E-14 | **8.84E-13** |
| ENSG00000204262.16 | **COL5A2** | 1290 | protein_coding | **80.500** | 1.993 | 0.600 | **3.981** | 6.88E-14 | **8.93E-13** |
| ENSG00000172301.12 | **COPRS** | 55352 | protein_coding | **212.190** | -0.990 | -0.298 | **0.504** | 6.88E-14 | **8.93E-13** |
| ENSG00000077147.18 | **TM9SF3** | 56889 | protein_coding | **831.976** | -0.715 | -0.215 | **0.609** | 6.88E-14 | **8.93E-13** |
| ENSG00000198604.13 | **BAZ1A** | 11177 | protein_coding | **795.823** | -0.961 | -0.289 | **0.514** | 6.94E-14 | **8.99E-13** |
| ENSG00000145604.19 | **SKP2** | 6502 | protein_coding | **334.673** | -0.868 | -0.261 | **0.548** | 6.97E-14 | **9.03E-13** |
| ENSG00000101972.22 | **STAG2** | 10735 | protein_coding | **572.823** | -1.338 | -0.403 | **0.395** | 7.09E-14 | **9.18E-13** |
| ENSG00000124006.17 | **OBSL1** | 23363 | protein_coding | **52.737** | 2.737 | 0.824 | **6.668** | 7.29E-14 | **9.43E-13** |
| ENSG00000198554.14 | **WDHD1** | 11169 | protein_coding | **303.304** | -1.567 | -0.472 | **0.337** | 7.31E-14 | **9.46E-13** |
| ENSG00000142694.9 | **EVA1B** | 55194 | protein_coding | **161.664** | 1.194 | 0.359 | **2.287** | 7.36E-14 | **9.51E-13** |
| ENSG00000087448.13 | **KLHL42** | 57542 | protein_coding | **296.964** | -1.041 | -0.313 | **0.486** | 7.40E-14 | **9.56E-13** |
| ENSG00000167005.16 | **NUDT21** | 11051 | protein_coding | **1073.275** | -1.159 | -0.349 | **0.448** | 7.48E-14 | **9.65E-13** |
| ENSG00000103145.13 | **HCFC1R1** | 54985 | protein_coding | **1103.025** | 1.337 | 0.403 | **2.527** | 7.49E-14 | **9.66E-13** |
| ENSG00000172167.10 | **MTBP** | 27085 | protein_coding | **112.307** | -1.648 | -0.496 | **0.319** | 7.59E-14 | **9.78E-13** |
| ENSG00000305353.1 | **ENSG00000305353** | | lncRNA | **17.693** | 4.859 | 1.463 | **29.017** | 7.78E-14 | **1.00E-12** |
| ENSG00000166847.12 | **DCTN5** | 84516 | protein_coding | **965.465** | -0.764 | -0.230 | **0.589** | 7.83E-14 | **1.01E-12** |
| ENSG00000175482.11 | **POLD4** | 57804 | protein_coding | **1831.980** | 0.657 | 0.198 | **1.577** | 7.94E-14 | **1.02E-12** |
| ENSG00000114054.16 | **PCCB** | 5096 | protein_coding | **676.239** | -0.795 | -0.239 | **0.576** | 7.96E-14 | **1.02E-12** |
| ENSG00000293457.2 | **PTPRVP** | 148713 | lncRNA | **63.778** | 1.499 | 0.451 | **2.826** | 8.03E-14 | **1.03E-12** |

| | | | | | | | | | |
|---|---|---|---|---|---|---|---|---|---|
| ENSG00000114107.11 | **CEP70** | 80321 | protein_coding | **152.859** | 1.551 | 0.467 | **2.930** | 8.11E-14 | **1.04E-12** |
| ENSG00000184887.16 | **BTBD6** | 90135 | protein_coding | **395.861** | -0.877 | -0.264 | **0.545** | 8.20E-14 | **1.05E-12** |
| ENSG00000288924.1 | **ENSG00000288924** | | lncRNA | **1512.314** | 0.835 | 0.251 | **1.784** | 8.44E-14 | **1.08E-12** |
| ENSG00000110076.22 | **NRXN2** | 9379 | protein_coding | **37.092** | 3.181 | 0.957 | **9.067** | 8.57E-14 | **1.10E-12** |
| ENSG00000128271.24 | **ADORA2A** | 135 | protein_coding | **691.127** | -0.961 | -0.289 | **0.514** | 8.62E-14 | **1.10E-12** |
| ENSG00000198521.14 | **ZNF43** | 7594 | protein_coding | **187.085** | -1.210 | -0.364 | **0.432** | 8.65E-14 | **1.11E-12** |
| ENSG00000139631.22 | **CSAD** | 51380 | protein_coding | **592.878** | 0.845 | 0.254 | **1.796** | 8.65E-14 | **1.11E-12** |
| ENSG00000205220.14 | **PSMB10** | 5699 | protein_coding | **2998.585** | -1.212 | -0.365 | **0.432** | 8.78E-14 | **1.12E-12** |
| ENSG00000301519.1 | **ENSG00000301519** | | lncRNA | **23.767** | 2.451 | 0.738 | **5.468** | 8.79E-14 | **1.12E-12** |
| ENSG00000118655.10 | **DCLRE1B** | 64858 | protein_coding | **157.807** | -1.085 | -0.327 | **0.471** | 8.91E-14 | **1.14E-12** |
| ENSG00000115919.17 | **KYNU** | 8942 | protein_coding | **588.905** | -0.709 | -0.213 | **0.612** | 8.96E-14 | **1.14E-12** |
| ENSG00000175806.17 | **MSRA** | 4482 | protein_coding | **108.673** | -1.134 | -0.341 | **0.456** | 8.97E-14 | **1.14E-12** |
| ENSG00000286388.1 | **ENSG00000286388** | | lncRNA | **86.950** | 1.427 | 0.430 | **2.689** | 9.39E-14 | **1.20E-12** |
| ENSG00000176438.15 | **SYNE3** | 161176 | protein_coding | **967.667** | -0.855 | -0.257 | **0.553** | 9.45E-14 | **1.20E-12** |
| ENSG00000165923.17 | **AGBL2** | 79841 | protein_coding | **132.377** | 1.224 | 0.368 | **2.335** | 9.64E-14 | **1.23E-12** |
| ENSG00000176381.6 | **PRR18** | 285800 | protein_coding | **111.066** | -1.767 | -0.532 | **0.294** | 9.68E-14 | **1.23E-12** |
| ENSG00000130595.22 | **TNNT3** | 7140 | protein_coding | **96.839** | 1.485 | 0.447 | **2.799** | 9.70E-14 | **1.23E-12** |
| ENSG00000082805.23 | **ERC1** | 23085 | protein_coding | **99.592** | 1.674 | 0.504 | **3.190** | 9.83E-14 | **1.25E-12** |
| ENSG00000206561.16 | **COLQ** | 8292 | protein_coding | **174.324** | 1.186 | 0.357 | **2.275** | 1.00E-13 | **1.27E-12** |
| ENSG00000169410.13 | **PTPN9** | 5780 | protein_coding | **401.399** | -0.844 | -0.254 | **0.557** | 1.01E-13 | **1.27E-12** |
| ENSG00000138796.20 | **HADH** | 3033 | protein_coding | **840.346** | -0.737 | -0.222 | **0.600** | 1.03E-13 | **1.30E-12** |
| ENSG00000132341.14 | **RAN** | 5901 | protein_coding | **9002.788** | -0.934 | -0.281 | **0.523** | 1.05E-13 | **1.33E-12** |
| ENSG00000221968.11 | **FADS3** | 3995 | protein_coding | **582.444** | 1.369 | 0.412 | **2.583** | 1.05E-13 | **1.33E-12** |
| ENSG00000157680.18 | **DGKI** | 9162 | protein_coding | **102.149** | -1.868 | -0.562 | **0.274** | 1.06E-13 | **1.34E-12** |
| ENSG00000186868.20 | **MAPT** | 4137 | protein_coding | **33.784** | 3.462 | 1.042 | **11.020** | 1.11E-13 | **1.40E-12** |
| ENSG00000149927.20 | **DOC2A** | 8448 | protein_coding | **46.334** | 2.115 | 0.637 | **4.332** | 1.12E-13 | **1.42E-12** |
| ENSG00000136986.12 | **DERL1** | 79139 | protein_coding | **863.547** | -0.846 | -0.255 | **0.556** | 1.13E-13 | **1.42E-12** |
| ENSG00000152256.16 | **PDK1** | 5163 | protein_coding | **730.885** | -0.800 | -0.241 | **0.574** | 1.13E-13 | **1.42E-12** |
| ENSG00000082512.17 | **TRAF5** | 7188 | protein_coding | **958.317** | -1.118 | -0.336 | **0.461** | 1.14E-13 | **1.43E-12** |
| ENSG00000165424.9 | **ZCCHC24** | 219654 | protein_coding | **48.219** | 2.045 | 0.616 | **4.128** | 1.15E-13 | **1.45E-12** |
| ENSG00000130429.18 | **ARPC1B** | 10095 | protein_coding | **4966.969** | -0.856 | -0.258 | **0.552** | 1.15E-13 | **1.45E-12** |

| | | | | | | | | | |
|---|---|---|---|---|---|---|---|---|---|
| ENSG00000138185.22 | **ENTPD1** | 953 | protein_coding | **4240.313** | -0.840 | -0.253 | **0.559** | 1.19E-13 | **1.49E-12** |
| ENSG00000005801.20 | **ZNF195** | 7748 | protein_coding | **629.578** | -0.942 | -0.283 | **0.521** | 1.19E-13 | **1.49E-12** |
| ENSG00000213563.8 | **C8orf82** | 414919 | protein_coding | **240.093** | -0.988 | -0.297 | **0.504** | 1.21E-13 | **1.52E-12** |
| ENSG00000167930.19 | **FAM234A** | 83986 | protein_coding | **1095.477** | 0.869 | 0.261 | **1.826** | 1.22E-13 | **1.54E-12** |
| ENSG00000108515.20 | **ENO3** | 2027 | protein_coding | **1064.964** | 1.058 | 0.319 | **2.082** | 1.23E-13 | **1.54E-12** |
| ENSG00000285823.1 | **LINC02793** | | lncRNA | **56.144** | 1.629 | 0.490 | **3.092** | 1.25E-13 | **1.57E-12** |
| ENSG00000167549.21 | **CORO6** | 84940 | protein_coding | **261.542** | 1.649 | 0.496 | **3.136** | 1.26E-13 | **1.58E-12** |
| ENSG00000158856.20 | **DMTN** | 2039 | protein_coding | **34.769** | 2.272 | 0.684 | **4.828** | 1.29E-13 | **1.62E-12** |
| ENSG00000101447.16 | **FAM83D** | 81610 | protein_coding | **102.527** | -1.454 | -0.438 | **0.365** | 1.30E-13 | **1.62E-12** |
| ENSG00000159733.15 | **ZFYVE28** | 57732 | protein_coding | **234.992** | 1.975 | 0.595 | **3.932** | 1.33E-13 | **1.66E-12** |
| ENSG00000124570.23 | **SERPINB6** | 5269 | protein_coding | **61.372** | 1.874 | 0.564 | **3.667** | 1.33E-13 | **1.66E-12** |
| ENSG00000157020.20 | **SEC13** | 6396 | protein_coding | **1459.453** | -0.646 | -0.195 | **0.639** | 1.33E-13 | **1.66E-12** |
| ENSG00000170515.16 | **PA2G4** | 5036 | protein_coding | **4949.122** | -1.076 | -0.324 | **0.474** | 1.36E-13 | **1.70E-12** |
| ENSG00000108654.18 | **DDX5** | 1655 | protein_coding | **8132.625** | -0.667 | -0.201 | **0.630** | 1.39E-13 | **1.74E-12** |
| ENSG00000030582.21 | **GRN** | 2896 | protein_coding | **5256.086** | 0.778 | 0.234 | **1.714** | 1.42E-13 | **1.77E-12** |
| ENSG00000105122.15 | **RASAL3** | 64926 | protein_coding | **2194.473** | -0.982 | -0.296 | **0.506** | 1.42E-13 | **1.77E-12** |
| ENSG00000104907.15 | **TRMT1** | 55621 | protein_coding | **1216.952** | -1.139 | -0.343 | **0.454** | 1.42E-13 | **1.77E-12** |
| ENSG00000284543.2 | **LINC01226** | | lncRNA | **263.620** | -1.579 | -0.475 | **0.335** | 1.44E-13 | **1.78E-12** |
| ENSG00000073792.18 | **IGF2BP2** | 10644 | protein_coding | **24.793** | 5.384 | 1.621 | **41.756** | 1.44E-13 | **1.79E-12** |
| ENSG00000184470.22 | **TXNRD2** | 10587 | protein_coding | **363.235** | -1.079 | -0.325 | **0.473** | 1.45E-13 | **1.80E-12** |
| ENSG00000204569.12 | **PPP1R10** | 5514 | protein_coding | **1850.733** | 0.586 | 0.176 | **1.501** | 1.46E-13 | **1.82E-12** |
| ENSG00000164638.13 | **SLC29A4** | 222962 | protein_coding | **108.555** | 1.677 | 0.505 | **3.198** | 1.48E-13 | **1.83E-12** |
| ENSG00000111144.12 | **LTA4H** | 4048 | protein_coding | **1956.385** | 0.799 | 0.241 | **1.740** | 1.49E-13 | **1.85E-12** |
| ENSG00000099785.13 | **MARCHF2** | 51257 | protein_coding | **396.713** | 1.192 | 0.359 | **2.285** | 1.52E-13 | **1.88E-12** |
| ENSG00000136463.11 | **TACO1** | 51204 | protein_coding | **334.703** | -0.824 | -0.248 | **0.565** | 1.53E-13 | **1.89E-12** |
| ENSG00000113643.11 | **RARS1** | 5917 | protein_coding | **1486.822** | 0.610 | 0.184 | **1.526** | 1.56E-13 | **1.93E-12** |
| ENSG00000141540.13 | **TTYH2** | 94015 | protein_coding | **166.100** | 1.348 | 0.406 | **2.546** | 1.58E-13 | **1.96E-12** |
| ENSG00000236017.9 | **ASMTL-AS1** | | lncRNA | **943.396** | 0.974 | 0.293 | **1.964** | 1.58E-13 | **1.96E-12** |
| ENSG00000119681.14 | **LTBP2** | 4053 | protein_coding | **17.840** | 3.605 | 1.085 | **12.168** | 1.59E-13 | **1.96E-12** |
| ENSG00000134049.8 | **IER3IP1** | 51124 | protein_coding | **340.517** | -0.942 | -0.284 | **0.520** | 1.60E-13 | **1.98E-12** |
| ENSG00000112561.21 | **TFEB** | 7942 | protein_coding | **1634.461** | -0.884 | -0.266 | **0.542** | 1.62E-13 | **1.99E-12** |

| | | | | | | | | | |
|---|---|---|---|---|---|---|---|---|---|
| ENSG00000116455.15 | **WDR77** | 79084 | protein_coding | **977.339** | -0.710 | -0.214 | **0.611** | 1.62E-13 | **2.00E-12** |
| ENSG00000233175.3 | **FMNL1-AS** | 10798504( | lncRNA | **99.562** | 1.556 | 0.468 | **2.940** | 1.68E-13 | **2.07E-12** |
| ENSG00000103275.24 | **UBE2I** | 7329 | protein_coding | **2768.454** | -0.556 | -0.167 | **0.680** | 1.72E-13 | **2.11E-12** |
| ENSG00000138050.17 | **THUMPD2** | 80745 | protein_coding | **251.028** | -1.106 | -0.333 | **0.465** | 1.72E-13 | **2.11E-12** |
| ENSG00000257949.9 | **TEN1** | 10013493 | protein_coding | **176.703** | -1.110 | -0.334 | **0.463** | 1.73E-13 | **2.12E-12** |
| ENSG00000138400.14 | **MDH1B** | 130752 | protein_coding | **38.918** | 2.409 | 0.725 | **5.313** | 1.75E-13 | **2.15E-12** |
| ENSG00000196954.15 | **CASP4** | 837 | protein_coding | **784.718** | -1.116 | -0.336 | **0.461** | 1.76E-13 | **2.16E-12** |
| ENSG00000182054.12 | **IDH2** | 3418 | protein_coding | **2945.558** | -0.758 | -0.228 | **0.591** | 1.77E-13 | **2.17E-12** |
| ENSG00000087995.18 | **METTL2A** | 339175 | protein_coding | **273.093** | -0.987 | -0.297 | **0.505** | 1.78E-13 | **2.18E-12** |
| ENSG00000287207.2 | **ENSG00000287207** | | lncRNA | **13.115** | 6.026 | 1.814 | **65.164** | 1.78E-13 | **2.18E-12** |
| ENSG00000144485.13 | **HES6** | 55502 | protein_coding | **70.088** | 1.841 | 0.554 | **3.583** | 1.80E-13 | **2.20E-12** |
| ENSG00000174885.14 | **NLRP6** | 171389 | protein_coding | **31.853** | 2.455 | 0.739 | **5.482** | 1.86E-13 | **2.28E-12** |
| ENSG00000060642.14 | **PIGV** | 55650 | protein_coding | **443.206** | 0.900 | 0.271 | **1.866** | 1.87E-13 | **2.28E-12** |
| ENSG00000145850.10 | **TIMD4** | 91937 | protein_coding | **180.590** | 2.012 | 0.606 | **4.033** | 1.88E-13 | **2.29E-12** |
| ENSG00000243927.8 | **MRPS6** | 64968 | protein_coding | **1106.263** | -0.595 | -0.179 | **0.662** | 1.90E-13 | **2.32E-12** |
| ENSG00000134824.16 | **FADS2** | 9415 | protein_coding | **3061.670** | -1.108 | -0.333 | **0.464** | 1.91E-13 | **2.33E-12** |
| ENSG00000238164.7 | **TNFRSF14-AS1** | | lncRNA | **791.561** | -1.203 | -0.362 | **0.434** | 1.93E-13 | **2.35E-12** |
| ENSG00000131944.12 | **FAAP24** | 91442 | protein_coding | **56.517** | -1.844 | -0.555 | **0.279** | 1.97E-13 | **2.40E-12** |
| ENSG00000214826.6 | **DDX12P** | | transcribed_unpro | **61.170** | -1.603 | -0.483 | **0.329** | 1.97E-13 | **2.40E-12** |
| ENSG00000149782.15 | **PLCB3** | 5331 | protein_coding | **467.104** | 1.034 | 0.311 | **2.047** | 1.97E-13 | **2.40E-12** |
| ENSG00000293485.2 | **SERHL** | 94009 | lncRNA | **37.395** | 2.405 | 0.724 | **5.295** | 1.99E-13 | **2.42E-12** |
| ENSG00000179148.11 | **ALOXE3** | 59344 | protein_coding | **20.945** | 3.913 | 1.178 | **15.064** | 2.00E-13 | **2.43E-12** |
| ENSG00000056736.12 | **IL17RB** | 55540 | protein_coding | **1331.498** | 1.039 | 0.313 | **2.055** | 2.02E-13 | **2.45E-12** |
| ENSG00000144560.17 | **VGLL4** | 9686 | protein_coding | **706.286** | -0.733 | -0.221 | **0.602** | 2.02E-13 | **2.46E-12** |
| ENSG00000167617.5 | **CDC42EP5** | 148170 | protein_coding | **46.970** | 2.261 | 0.681 | **4.793** | 2.05E-13 | **2.49E-12** |
| ENSG00000154760.16 | **SLFN13** | 146857 | protein_coding | **1619.521** | -0.619 | -0.186 | **0.651** | 2.11E-13 | **2.56E-12** |
| ENSG00000182534.15 | **MXRA7** | 439921 | protein_coding | **115.007** | 2.851 | 0.858 | **7.213** | 2.12E-13 | **2.57E-12** |
| ENSG00000138443.19 | **ABI2** | 10152 | protein_coding | **680.422** | -1.007 | -0.303 | **0.498** | 2.15E-13 | **2.60E-12** |
| ENSG00000095585.20 | **BLNK** | 29760 | protein_coding | **2538.022** | -0.810 | -0.244 | **0.570** | 2.16E-13 | **2.62E-12** |
| ENSG00000127554.14 | **GFER** | 2671 | protein_coding | **409.728** | -0.739 | -0.222 | **0.599** | 2.17E-13 | **2.63E-12** |
| ENSG00000005844.21 | **ITGAL** | 3683 | protein_coding | **1551.584** | -1.525 | -0.459 | **0.347** | 2.21E-13 | **2.67E-12** |

| | | | | | | | | | |
|---|---|---|---|---|---|---|---|---|---|
| ENSG00000174780.19 | **SRP72** | 6731 | protein_coding | **860.029** | -0.780 | -0.235 | **0.582** | 2.25E-13 | **2.72E-12** |
| ENSG00000164611.15 | **PTTG1** | 9232 | protein_coding | **1834.090** | -1.186 | -0.357 | **0.439** | 2.32E-13 | **2.79E-12** |
| ENSG00000147403.20 | **RPL10** | 6134 | protein_coding | **36677.403** | -0.660 | -0.199 | **0.633** | 2.40E-13 | **2.90E-12** |
| ENSG00000129932.12 | **DOHH** | 83475 | protein_coding | **226.016** | -0.828 | -0.249 | **0.563** | 2.42E-13 | **2.91E-12** |
| ENSG00000166548.18 | **TK2** | 7084 | protein_coding | **183.470** | 1.282 | 0.386 | **2.431** | 2.64E-13 | **3.18E-12** |
| ENSG00000127948.19 | **POR** | 5447 | protein_coding | **901.654** | 0.975 | 0.294 | **1.966** | 2.65E-13 | **3.20E-12** |
| ENSG00000228526.9 | **MIR34AHG** | 106614088 | lncRNA | **602.110** | -0.924 | -0.278 | **0.527** | 2.71E-13 | **3.26E-12** |
| ENSG00000138031.17 | **ADCY3** | 109 | protein_coding | **1209.769** | -0.714 | -0.215 | **0.610** | 2.71E-13 | **3.26E-12** |
| ENSG00000272068.1 | **BCAN-AS1** | | lncRNA | **84.546** | 1.240 | 0.373 | **2.361** | 2.82E-13 | **3.39E-12** |
| ENSG00000188266.15 | **HYKK** | 123688 | protein_coding | **75.690** | 2.165 | 0.652 | **4.484** | 2.84E-13 | **3.41E-12** |
| ENSG00000246263.5 | **UBR5-DT** | | lncRNA | **216.458** | 1.061 | 0.319 | **2.086** | 2.84E-13 | **3.41E-12** |
| ENSG00000286214.1 | **COPG2IT1** | | lncRNA | **22.884** | 3.746 | 1.128 | **13.419** | 2.90E-13 | **3.48E-12** |
| ENSG00000295544.1 | **ENSG00000295544** | | lncRNA | **23.530** | 2.696 | 0.812 | **6.482** | 2.92E-13 | **3.50E-12** |
| ENSG00000186891.15 | **TNFRSF18** | 8784 | protein_coding | **207.461** | 1.194 | 0.359 | **2.288** | 2.92E-13 | **3.50E-12** |
| ENSG00000108106.16 | **UBE2S** | 27338 | protein_coding | **2127.137** | -1.072 | -0.323 | **0.476** | 2.93E-13 | **3.51E-12** |
| ENSG00000110046.16 | **ATG2A** | 23130 | protein_coding | **576.570** | 1.052 | 0.317 | **2.073** | 2.97E-13 | **3.55E-12** |
| ENSG00000267750.7 | **RUNDC3A-AS1** | | lncRNA | **35.518** | 2.556 | 0.769 | **5.879** | 2.97E-13 | **3.55E-12** |
| ENSG00000056558.13 | **TRAF1** | 7185 | protein_coding | **7819.424** | 0.996 | 0.300 | **1.994** | 2.99E-13 | **3.57E-12** |
| ENSG00000213967.13 | **ZNF726** | 730087 | protein_coding | **46.347** | -2.044 | -0.615 | **0.242** | 3.07E-13 | **3.66E-12** |
| ENSG00000136108.17 | **CKAP2** | 26586 | protein_coding | **318.662** | -1.038 | -0.313 | **0.487** | 3.14E-13 | **3.75E-12** |
| ENSG00000131778.22 | **CHD1L** | 9557 | protein_coding | **920.281** | -0.658 | -0.198 | **0.634** | 3.19E-13 | **3.81E-12** |
| ENSG00000180596.11 | **H2BC4** | 8347 | protein_coding | **88.207** | -1.931 | -0.581 | **0.262** | 3.21E-13 | **3.83E-12** |
| ENSG00000163714.20 | **U2SURP** | 23350 | protein_coding | **1767.900** | -0.793 | -0.239 | **0.577** | 3.21E-13 | **3.83E-12** |
| ENSG00000088387.24 | **DOCK9** | 23348 | protein_coding | **203.954** | 1.720 | 0.518 | **3.294** | 3.26E-13 | **3.88E-12** |
| ENSG00000136159.6 | **NUDT15** | 55270 | protein_coding | **163.393** | -0.970 | -0.292 | **0.511** | 3.28E-13 | **3.91E-12** |
| ENSG00000039068.22 | **CDH1** | 999 | protein_coding | **73.274** | 2.628 | 0.791 | **6.183** | 3.30E-13 | **3.92E-12** |
| ENSG00000196305.21 | **IARS1** | 3376 | protein_coding | **2338.690** | -0.868 | -0.261 | **0.548** | 3.41E-13 | **4.06E-12** |
| ENSG00000197136.7 | **PCNX3** | 399909 | protein_coding | **1338.741** | 0.673 | 0.203 | **1.595** | 3.47E-13 | **4.12E-12** |
| ENSG00000167414.7 | **GNG8** | 94235 | protein_coding | **88.788** | 1.851 | 0.557 | **3.607** | 3.50E-13 | **4.15E-12** |
| ENSG00000144036.18 | **EXOC6B** | 23233 | protein_coding | **137.479** | 1.579 | 0.475 | **2.988** | 3.52E-13 | **4.18E-12** |
| ENSG00000136104.24 | **RNASEH2B** | 79621 | protein_coding | **531.710** | -1.011 | -0.304 | **0.496** | 3.53E-13 | **4.18E-12** |

| | | | | | | | | | |
|---|---|---|---|---|---|---|---|---|---|
| ENSG00000116983.15 | **HPCAL4** | 51440 | protein_coding | **51.049** | 3.794 | 1.142 | **13.867** | 3.53E-13 | **4.18E-12** |
| ENSG00000139496.20 | **NUP58** | 9818 | protein_coding | **493.146** | -0.798 | -0.240 | **0.575** | 3.56E-13 | **4.21E-12** |
| ENSG00000120659.17 | **TNFSF11** | 8600 | protein_coding | **148.719** | -1.637 | -0.493 | **0.322** | 3.69E-13 | **4.36E-12** |
| ENSG00000100292.19 | **HMOX1** | 3162 | protein_coding | **525.917** | 1.150 | 0.346 | **2.219** | 3.70E-13 | **4.37E-12** |
| ENSG00000289398.2 | **ENSG00000289398** | | lncRNA | **90.661** | 2.131 | 0.642 | **4.381** | 3.85E-13 | **4.55E-12** |
| ENSG00000133302.15 | **SLF1** | 84250 | protein_coding | **94.092** | -1.200 | -0.361 | **0.435** | 3.89E-13 | **4.60E-12** |
| ENSG00000290088.2 | **ENSG00000290088** | | lncRNA | **18.859** | 3.999 | 1.204 | **15.994** | 3.97E-13 | **4.69E-12** |
| ENSG00000171798.21 | **KNDC1** | 85442 | protein_coding | **17.391** | 4.271 | 1.286 | **19.308** | 4.07E-13 | **4.80E-12** |
| ENSG00000160469.18 | **BRSK1** | 84446 | protein_coding | **100.912** | 2.074 | 0.624 | **4.211** | 4.18E-13 | **4.92E-12** |
| ENSG00000243081.3 | **ENSG0000(** | 105374041 | lncRNA | **285.688** | 1.546 | 0.465 | **2.920** | 4.26E-13 | **5.02E-12** |
| ENSG00000291130.2 | **ENSG00000291130** | | lncRNA | **106.302** | -1.454 | -0.438 | **0.365** | 4.28E-13 | **5.03E-12** |
| ENSG00000122694.18 | **GLIPR2** | 152007 | protein_coding | **87.427** | 2.421 | 0.729 | **5.357** | 4.35E-13 | **5.12E-12** |
| ENSG00000115165.12 | **CYTIP** | 9595 | protein_coding | **2800.689** | -0.778 | -0.234 | **0.583** | 4.36E-13 | **5.13E-12** |
| ENSG00000136240.12 | **KDELR2** | 11014 | protein_coding | **1316.313** | -0.938 | -0.282 | **0.522** | 4.44E-13 | **5.21E-12** |
| ENSG00000162894.14 | **FCMR** | 9214 | protein_coding | **2032.261** | -1.290 | -0.388 | **0.409** | 4.46E-13 | **5.23E-12** |
| ENSG00000171174.17 | **RBKS** | 64080 | protein_coding | **37.627** | 1.785 | 0.537 | **3.447** | 4.49E-13 | **5.27E-12** |
| ENSG00000291223.1 | **FAM86DP** | 692099 | lncRNA | **72.380** | -1.262 | -0.380 | **0.417** | 4.60E-13 | **5.40E-12** |
| ENSG00000295617.1 | **ENSG00000295617** | | lncRNA | **18.846** | 3.863 | 1.163 | **14.546** | 4.64E-13 | **5.44E-12** |
| ENSG00000111371.18 | **SLC38A1** | 81539 | protein_coding | **1635.080** | -0.980 | -0.295 | **0.507** | 4.72E-13 | **5.53E-12** |
| ENSG00000267390.4 | **KDSR-DT** | 118827811 | lncRNA | **96.023** | 1.607 | 0.484 | **3.046** | 4.75E-13 | **5.56E-12** |
| ENSG00000153060.8 | **TEKT5** | 146279 | protein_coding | **41.072** | 2.306 | 0.694 | **4.945** | 4.76E-13 | **5.57E-12** |
| ENSG00000165795.27 | **NDRG2** | 57447 | protein_coding | **18.499** | 4.691 | 1.412 | **25.837** | 4.80E-13 | **5.61E-12** |
| ENSG00000186204.17 | **CYP4F12** | 66002 | protein_coding | **34.704** | 2.857 | 0.860 | **7.247** | 4.87E-13 | **5.69E-12** |
| ENSG00000108830.13 | **RND2** | 8153 | protein_coding | **17.454** | 4.488 | 1.351 | **22.445** | 4.95E-13 | **5.78E-12** |
| ENSG00000270127.2 | **ENSG00000270127** | | lncRNA | **114.998** | 1.066 | 0.321 | **2.094** | 5.14E-13 | **6.00E-12** |
| ENSG00000189001.11 | **SBSN** | 374897 | protein_coding | **34.946** | 3.966 | 1.194 | **15.624** | 5.23E-13 | **6.09E-12** |
| ENSG00000288961.3 | **ENSG0000(** | 124904009 | lncRNA | **95.987** | 1.214 | 0.365 | **2.320** | 5.28E-13 | **6.16E-12** |
| ENSG00000286482.3 | **ENSG00000286482** | | lncRNA | **47.705** | 1.986 | 0.598 | **3.960** | 5.31E-13 | **6.18E-12** |
| ENSG00000235092.8 | **ID2-AS1** | 100506299 | lncRNA | **39.584** | 1.991 | 0.599 | **3.974** | 5.39E-13 | **6.27E-12** |
| ENSG00000213420.10 | **GPC2** | 221914 | protein_coding | **121.149** | 1.624 | 0.489 | **3.082** | 5.40E-13 | **6.28E-12** |
| ENSG00000140853.19 | **NLRC5** | 84166 | protein_coding | **4432.206** | 0.644 | 0.194 | **1.563** | 5.53E-13 | **6.43E-12** |

| | | | | | | | | | |
|---|---|---|---|---|---|---|---|---|---|
| ENSG00000296092.1 | **ENSG00000296092** | | lncRNA | **31.992** | 2.576 | 0.775 | **5.962** | 5.55E-13 | **6.45E-12** |
| ENSG00000159714.14 | **ZDHHC1** | 29800 | protein_coding | **37.034** | 2.449 | 0.737 | **5.462** | 5.74E-13 | **6.66E-12** |
| ENSG00000119326.17 | **CTNNAL1** | 8727 | protein_coding | **256.653** | -0.782 | -0.235 | **0.582** | 5.80E-13 | **6.73E-12** |
| ENSG00000205517.15 | **RGL3** | 57139 | protein_coding | **210.327** | 3.401 | 1.024 | **10.563** | 5.83E-13 | **6.76E-12** |
| ENSG00000070785.19 | **EIF2B3** | 8891 | protein_coding | **284.230** | -1.038 | -0.313 | **0.487** | 5.84E-13 | **6.76E-12** |
| ENSG00000111707.14 | **SUDS3** | 64426 | protein_coding | **1057.385** | 0.725 | 0.218 | **1.653** | 5.89E-13 | **6.82E-12** |
| ENSG00000257086.2 | **ENSG00000257086** | | lncRNA | **43.673** | 1.785 | 0.537 | **3.447** | 5.91E-13 | **6.84E-12** |
| ENSG00000105767.5 | **CADM4** | 199731 | protein_coding | **24.466** | 3.033 | 0.913 | **8.184** | 5.94E-13 | **6.87E-12** |
| ENSG00000128016.10 | **ZFP36** | 7538 | protein_coding | **4655.625** | 0.852 | 0.256 | **1.805** | 5.99E-13 | **6.93E-12** |
| ENSG00000196378.16 | **ZNF34** | 80778 | protein_coding | **174.617** | 1.057 | 0.318 | **2.080** | 6.13E-13 | **7.08E-12** |
| ENSG00000136574.21 | **GATA4** | 2626 | protein_coding | **31.092** | 3.383 | 1.019 | **10.436** | 6.19E-13 | **7.14E-12** |
| ENSG00000088899.18 | **LZTS3** | 9762 | protein_coding | **123.139** | 1.502 | 0.452 | **2.833** | 6.21E-13 | **7.16E-12** |
| ENSG00000139218.20 | **SCAF11** | 9169 | protein_coding | **1247.645** | -0.841 | -0.253 | **0.558** | 6.29E-13 | **7.25E-12** |
| ENSG00000151650.10 | **VENTX** | 27287 | protein_coding | **14.484** | 5.266 | 1.585 | **38.481** | 6.30E-13 | **7.26E-12** |
| ENSG00000055118.20 | **KCNH2** | 3757 | protein_coding | **15.651** | 4.169 | 1.255 | **17.987** | 6.34E-13 | **7.30E-12** |
| ENSG00000119285.13 | **HEATR1** | 55127 | protein_coding | **937.873** | -0.653 | -0.197 | **0.636** | 6.44E-13 | **7.42E-12** |
| ENSG00000260807.8 | **CEROX1** | 115804232 | lncRNA | **30.876** | 2.812 | 0.847 | **7.025** | 6.45E-13 | **7.42E-12** |
| ENSG00000233585.3 | **ACAA2P1** | | transcribed_proce | **23.504** | 2.663 | 0.802 | **6.332** | 6.46E-13 | **7.43E-12** |
| ENSG00000154358.24 | **OBSCN** | 84033 | protein_coding | **154.539** | 2.056 | 0.619 | **4.160** | 6.65E-13 | **7.64E-12** |
| ENSG00000165912.18 | **PACSIN3** | 29763 | protein_coding | **98.380** | 1.369 | 0.412 | **2.583** | 6.80E-13 | **7.81E-12** |
| ENSG00000177380.16 | **PPFIA3** | 8541 | protein_coding | **60.450** | 1.923 | 0.579 | **3.793** | 6.90E-13 | **7.92E-12** |
| ENSG00000287259.2 | **ENSG0000(** | 12490453( | lncRNA | **25.462** | 3.172 | 0.955 | **9.010** | 7.13E-13 | **8.18E-12** |
| ENSG00000267432.6 | **DNAH17-AS1** | | lncRNA | **126.012** | 1.760 | 0.530 | **3.386** | 7.21E-13 | **8.27E-12** |
| ENSG00000198752.13 | **CDC42BPB** | 9578 | protein_coding | **700.216** | 1.074 | 0.323 | **2.105** | 7.31E-13 | **8.38E-12** |
| ENSG00000139343.13 | **SNRPF** | 6636 | protein_coding | **1689.480** | -1.073 | -0.323 | **0.475** | 7.32E-13 | **8.39E-12** |
| ENSG00000167535.10 | **CACNB3** | 784 | protein_coding | **261.723** | 1.845 | 0.555 | **3.591** | 7.36E-13 | **8.42E-12** |
| ENSG00000085185.16 | **BCORL1** | 63035 | protein_coding | **155.160** | 1.197 | 0.360 | **2.293** | 7.39E-13 | **8.45E-12** |
| ENSG00000198435.4 | **NRARP** | 441478 | protein_coding | **119.043** | 2.153 | 0.648 | **4.446** | 7.40E-13 | **8.45E-12** |
| ENSG00000276070.6 | **CCL4L2** | 9560 | protein_coding | **432.048** | -1.815 | -0.546 | **0.284** | 7.44E-13 | **8.50E-12** |
| ENSG00000112208.13 | **BAG2** | 9532 | protein_coding | **232.627** | -1.057 | -0.318 | **0.481** | 7.46E-13 | **8.51E-12** |
| ENSG00000266947.4 | **ENSG00000266947** | | lncRNA | **62.082** | 1.399 | 0.421 | **2.638** | 7.64E-13 | **8.72E-12** |

| | | | | | | | | | |
|---|---|---|---|---|---|---|---|---|---|
| ENSG00000125084.13 | **WNT1** | 7471 | protein_coding | **13.207** | 4.061 | 1.222 | **16.691** | 7.80E-13 | **8.89E-12** |
| ENSG00000258655.3 | **ARHGAP5-AS1** | | lncRNA | **23.158** | 2.410 | 0.725 | **5.314** | 7.80E-13 | **8.89E-12** |
| ENSG00000037280.18 | **FLT4** | 2324 | protein_coding | **41.796** | 3.963 | 1.193 | **15.596** | 7.97E-13 | **9.08E-12** |
| ENSG00000105519.19 | **CAPS** | 828 | protein_coding | **480.868** | 1.002 | 0.302 | **2.003** | 8.02E-13 | **9.13E-12** |
| ENSG00000166012.19 | **TAF1D** | 79101 | protein_coding | **2133.585** | -0.745 | -0.224 | **0.597** | 8.04E-13 | **9.14E-12** |
| ENSG00000128284.23 | **APOL3** | 80833 | protein_coding | **1293.208** | 0.570 | 0.172 | **1.485** | 8.07E-13 | **9.18E-12** |
| ENSG00000084774.16 | **CAD** | 790 | protein_coding | **949.695** | -0.712 | -0.214 | **0.611** | 8.10E-13 | **9.20E-12** |
| ENSG00000171992.15 | **SYNPO** | 11346 | protein_coding | **1620.253** | 1.024 | 0.308 | **2.033** | 8.39E-13 | **9.53E-12** |
| ENSG00000166228.11 | **PCBD1** | 5092 | protein_coding | **720.072** | -1.090 | -0.328 | **0.470** | 8.64E-13 | **9.80E-12** |
| ENSG00000139514.15 | **SLC7A1** | 6541 | protein_coding | **1072.041** | -1.188 | -0.357 | **0.439** | 8.66E-13 | **9.82E-12** |
| ENSG00000159496.16 | **RGL4** | 266747 | protein_coding | **9.878** | -6.528 | -1.965 | **0.011** | 8.78E-13 | **9.95E-12** |
| ENSG00000120800.7 | **UTP20** | 27340 | protein_coding | **201.915** | -1.139 | -0.343 | **0.454** | 8.80E-13 | **9.97E-12** |
| ENSG00000120738.9 | **EGR1** | 1958 | protein_coding | **358.394** | 1.006 | 0.303 | **2.009** | 8.97E-13 | **1.02E-11** |
| ENSG00000183814.18 | **LIN9** | 286826 | protein_coding | **59.768** | -1.557 | -0.469 | **0.340** | 8.98E-13 | **1.02E-11** |
| ENSG00000090097.24 | **PCBP4** | 57060 | protein_coding | **539.866** | 0.880 | 0.265 | **1.841** | 9.22E-13 | **1.04E-11** |
| ENSG00000114859.18 | **CLCN2** | 1181 | protein_coding | **109.598** | 1.404 | 0.423 | **2.646** | 9.22E-13 | **1.04E-11** |
| ENSG00000137265.18 | **IRF4** | 3662 | protein_coding | **19675.326** | 0.989 | 0.298 | **1.985** | 9.34E-13 | **1.05E-11** |
| ENSG00000165916.11 | **PSMC3** | 5702 | protein_coding | **1579.143** | -0.695 | -0.209 | **0.618** | 9.37E-13 | **1.06E-11** |
| ENSG00000127586.19 | **CHTF18** | 63922 | protein_coding | **1212.584** | -1.617 | -0.487 | **0.326** | 9.39E-13 | **1.06E-11** |
| ENSG00000119333.14 | **DYNC2I2** | 89891 | protein_coding | **912.534** | -0.932 | -0.281 | **0.524** | 9.47E-13 | **1.07E-11** |
| ENSG00000185818.9 | **NAT8L** | 339983 | protein_coding | **21.637** | 3.251 | 0.979 | **9.524** | 9.49E-13 | **1.07E-11** |
| ENSG00000163832.18 | **ELP6** | 54859 | protein_coding | **435.536** | -0.842 | -0.253 | **0.558** | 9.60E-13 | **1.08E-11** |
| ENSG00000260001.10 | **TGFBR3L** | 10050758 | protein_coding | **33.146** | 3.764 | 1.133 | **13.587** | 9.62E-13 | **1.08E-11** |
| ENSG00000213390.13 | **ARHGAP19** | 84986 | protein_coding | **339.327** | -1.002 | -0.302 | **0.499** | 9.84E-13 | **1.11E-11** |
| ENSG00000092820.21 | **EZR** | 7430 | protein_coding | **5824.286** | 0.619 | 0.186 | **1.536** | 1.01E-12 | **1.13E-11** |
| ENSG00000101868.15 | **POLA1** | 5422 | protein_coding | **371.982** | -0.847 | -0.255 | **0.556** | 1.02E-12 | **1.14E-11** |
| ENSG00000166900.19 | **STX3** | 6809 | protein_coding | **63.232** | 2.440 | 0.735 | **5.427** | 1.02E-12 | **1.15E-11** |
| ENSG00000139546.13 | **TARBP2** | 6895 | protein_coding | **584.605** | 0.800 | 0.241 | **1.741** | 1.02E-12 | **1.15E-11** |
| ENSG00000148200.19 | **NR6A1** | 2649 | protein_coding | **319.614** | 1.023 | 0.308 | **2.032** | 1.03E-12 | **1.15E-11** |
| ENSG00000108511.12 | **HOXB6** | 3216 | protein_coding | **26.891** | 2.586 | 0.778 | **6.004** | 1.04E-12 | **1.16E-11** |
| ENSG00000182481.12 | **KPNA2** | 3838 | protein_coding | **1892.751** | -1.276 | -0.384 | **0.413** | 1.09E-12 | **1.23E-11** |

| | | | | | | | | | |
|---|---|---|---|---|---|---|---|---|---|
| ENSG00000141076.20 | **UTP4** | 84916 | protein_coding | **692.822** | -0.865 | -0.260 | **0.549** | 1.10E-12 | **1.23E-11** |
| ENSG00000168275.17 | **COA6** | 388753 | protein_coding | **325.224** | -0.856 | -0.258 | **0.552** | 1.10E-12 | **1.24E-11** |
| ENSG00000224616.5 | **RTCA-AS1** | | lncRNA | **81.988** | 1.513 | 0.455 | **2.854** | 1.11E-12 | **1.24E-11** |
| ENSG00000185504.19 | **FAAP100** | 80233 | protein_coding | **657.129** | -0.956 | -0.288 | **0.516** | 1.12E-12 | **1.25E-11** |
| ENSG00000134107.7 | **BHLHE40** | 8553 | protein_coding | **1712.876** | -0.782 | -0.235 | **0.582** | 1.16E-12 | **1.29E-11** |
| ENSG00000288095.3 | **ENSG00000288095** | | lncRNA | **45.588** | 2.067 | 0.622 | **4.191** | 1.18E-12 | **1.32E-11** |
| ENSG00000132259.15 | **CNGA4** | 1262 | protein_coding | **36.747** | 1.986 | 0.598 | **3.961** | 1.20E-12 | **1.34E-11** |
| ENSG00000139410.17 | **SDSL** | 113675 | protein_coding | **107.240** | 1.535 | 0.462 | **2.898** | 1.22E-12 | **1.35E-11** |
| ENSG00000100410.10 | **PHF5A** | 84844 | protein_coding | **540.343** | -0.712 | -0.214 | **0.611** | 1.25E-12 | **1.40E-11** |
| ENSG00000130768.17 | **SMPDL3B** | 27293 | protein_coding | **54.761** | 1.905 | 0.574 | **3.746** | 1.25E-12 | **1.40E-11** |
| ENSG00000165392.13 | **WRN** | 7486 | protein_coding | **351.457** | -0.884 | -0.266 | **0.542** | 1.26E-12 | **1.40E-11** |
| ENSG00000251143.1 | **NUMA1-AS** | 10012849 | lncRNA | **45.355** | 1.753 | 0.528 | **3.370** | 1.26E-12 | **1.40E-11** |
| ENSG00000100714.20 | **MTHFD1** | 4522 | protein_coding | **2454.075** | -0.921 | -0.277 | **0.528** | 1.28E-12 | **1.42E-11** |
| ENSG00000171105.16 | **INSR** | 3643 | protein_coding | **1511.559** | 0.763 | 0.230 | **1.697** | 1.29E-12 | **1.43E-11** |
| ENSG00000027847.16 | **B4GALT7** | 11285 | protein_coding | **356.782** | -0.862 | -0.259 | **0.550** | 1.29E-12 | **1.43E-11** |
| ENSG00000290931.2 | **ENSG00000290931** | | lncRNA | **212.412** | -0.879 | -0.265 | **0.544** | 1.31E-12 | **1.45E-11** |
| ENSG00000145569.8 | **OTULINL** | 54491 | protein_coding | **35.399** | 2.751 | 0.828 | **6.730** | 1.32E-12 | **1.46E-11** |
| ENSG00000090238.14 | **YPEL3** | 83719 | protein_coding | **1468.901** | 1.089 | 0.328 | **2.127** | 1.33E-12 | **1.47E-11** |
| ENSG00000154930.17 | **ACSS1** | 84532 | protein_coding | **873.472** | -0.668 | -0.201 | **0.629** | 1.33E-12 | **1.47E-11** |
| ENSG00000166924.11 | **NYAP1** | 222950 | protein_coding | **24.688** | 3.531 | 1.063 | **11.560** | 1.37E-12 | **1.52E-11** |
| ENSG00000100083.22 | **GGA1** | 26088 | protein_coding | **2451.876** | 0.481 | 0.145 | **1.396** | 1.39E-12 | **1.54E-11** |
| ENSG00000291224.2 | **ENSG00000291224** | | lncRNA | **125.070** | -1.290 | -0.388 | **0.409** | 1.39E-12 | **1.54E-11** |
| ENSG00000116815.18 | **CD58** | 965 | protein_coding | **1387.270** | -1.221 | -0.368 | **0.429** | 1.40E-12 | **1.54E-11** |
| ENSG00000249898.11 | **MCPH1-AS1** | | lncRNA | **148.941** | 1.657 | 0.499 | **3.154** | 1.40E-12 | **1.55E-11** |
| ENSG00000122483.20 | **CCDC18** | 343099 | protein_coding | **309.859** | -1.516 | -0.456 | **0.350** | 1.43E-12 | **1.57E-11** |
| ENSG00000185386.17 | **MAPK11** | 5600 | protein_coding | **567.078** | 0.876 | 0.264 | **1.835** | 1.44E-12 | **1.59E-11** |
| ENSG00000006747.17 | **SCIN** | 85477 | protein_coding | **69.791** | 3.385 | 1.019 | **10.445** | 1.47E-12 | **1.62E-11** |
| ENSG00000109519.16 | **GRPEL1** | 80273 | protein_coding | **623.409** | -0.677 | -0.204 | **0.626** | 1.48E-12 | **1.63E-11** |
| ENSG00000095319.16 | **NUP188** | 23511 | protein_coding | **1462.165** | -0.633 | -0.191 | **0.645** | 1.54E-12 | **1.70E-11** |
| ENSG00000144395.19 | **CCDC150** | 284992 | protein_coding | **126.279** | -1.756 | -0.528 | **0.296** | 1.62E-12 | **1.78E-11** |
| ENSG00000066379.17 | **POLR1H** | 30834 | protein_coding | **672.158** | -0.803 | -0.242 | **0.573** | 1.64E-12 | **1.80E-11** |

| | | | | | | | | | |
|---|---|---|---|---|---|---|---|---|---|
| ENSG00000101773.21 | **RBBP8** | 5932 | protein_coding | **467.148** | -1.074 | -0.323 | **0.475** | 1.65E-12 | **1.82E-11** |
| ENSG00000267706.3 | **ENSG00000267706** | | protein_coding | **18.266** | 5.177 | 1.558 | **36.170** | 1.66E-12 | **1.82E-11** |
| ENSG00000170412.20 | **GPRC5C** | 55890 | protein_coding | **324.387** | 0.979 | 0.295 | **1.971** | 1.69E-12 | **1.85E-11** |
| ENSG00000197568.16 | **ANKRD13C** | 11147 | lncRNA | **397.330** | 1.037 | 0.312 | **2.051** | 1.69E-12 | **1.85E-11** |
| ENSG00000105185.14 | **PDCD5** | 9141 | protein_coding | **1339.561** | -0.799 | -0.241 | **0.575** | 1.76E-12 | **1.94E-11** |
| ENSG00000089723.12 | **OTUB2** | 78990 | protein_coding | **24.360** | 2.941 | 0.885 | **7.678** | 1.78E-12 | **1.95E-11** |
| ENSG00000143297.21 | **FCRL5** | 83416 | protein_coding | **2441.934** | -1.659 | -0.499 | **0.317** | 1.84E-12 | **2.02E-11** |
| ENSG00000071967.14 | **CYBRD1** | 79901 | protein_coding | **559.520** | -0.917 | -0.276 | **0.530** | 1.84E-12 | **2.02E-11** |
| ENSG00000256262.4 | **USP30-AS1** | | lncRNA | **62.256** | -1.796 | -0.541 | **0.288** | 1.85E-12 | **2.02E-11** |
| ENSG00000145945.9 | **FAM50B** | 26240 | protein_coding | **197.050** | 1.370 | 0.412 | **2.585** | 1.90E-12 | **2.08E-11** |
| ENSG00000213619.13 | **NDUFS3** | 4722 | protein_coding | **1293.375** | -0.934 | -0.281 | **0.523** | 1.91E-12 | **2.09E-11** |
| ENSG00000185669.7 | **SNAI3** | 333929 | protein_coding | **92.168** | 1.168 | 0.352 | **2.247** | 1.92E-12 | **2.10E-11** |
| ENSG00000085982.18 | **USP40** | 55230 | protein_coding | **832.433** | 0.827 | 0.249 | **1.774** | 1.94E-12 | **2.12E-11** |
| ENSG00000162434.16 | **JAK1** | 3716 | protein_coding | **1178.760** | 0.832 | 0.250 | **1.780** | 1.94E-12 | **2.12E-11** |
| ENSG00000074416.18 | **MGLL** | 11343 | protein_coding | **1869.829** | -1.150 | -0.346 | **0.451** | 1.96E-12 | **2.13E-11** |
| ENSG00000088881.23 | **EBF4** | 57593 | protein_coding | **32.267** | 2.801 | 0.843 | **6.967** | 1.96E-12 | **2.13E-11** |
| ENSG00000196405.15 | **EVL** | 51466 | protein_coding | **3834.074** | -1.164 | -0.350 | **0.446** | 1.98E-12 | **2.15E-11** |
| ENSG00000289100.2 | **ENSG00000289100** | | lncRNA | **44.188** | 2.048 | 0.617 | **4.136** | 2.01E-12 | **2.19E-11** |
| ENSG00000089060.14 | **SLC8B1** | 80024 | protein_coding | **228.205** | 0.950 | 0.286 | **1.931** | 2.05E-12 | **2.23E-11** |
| ENSG00000205885.9 | **C1RL-AS1** | 283314 | lncRNA | **160.213** | -1.679 | -0.506 | **0.312** | 2.06E-12 | **2.24E-11** |
| ENSG00000130584.14 | **ZBTB46** | 140685 | protein_coding | **136.936** | 1.605 | 0.483 | **3.042** | 2.07E-12 | **2.25E-11** |
| ENSG00000087074.11 | **PPP1R15A** | 23645 | protein_coding | **2479.157** | -0.907 | -0.273 | **0.533** | 2.09E-12 | **2.27E-11** |
| ENSG00000175575.15 | **PAAF1** | 80227 | protein_coding | **500.326** | -0.710 | -0.214 | **0.611** | 2.13E-12 | **2.31E-11** |
| ENSG00000142156.18 | **COL6A1** | 1291 | protein_coding | **75.890** | 3.104 | 0.934 | **8.600** | 2.19E-12 | **2.38E-11** |
| ENSG00000111052.10 | **LIN7A** | 8825 | protein_coding | **189.631** | -1.559 | -0.469 | **0.339** | 2.20E-12 | **2.38E-11** |
| ENSG00000007312.16 | **CD79B** | 974 | protein_coding | **1770.696** | -0.804 | -0.242 | **0.573** | 2.21E-12 | **2.39E-11** |
| ENSG00000163909.10 | **HEYL** | 26508 | protein_coding | **16.465** | 4.085 | 1.230 | **16.969** | 2.22E-12 | **2.40E-11** |
| ENSG00000197448.16 | **GSTK1** | 373156 | protein_coding | **1830.912** | -0.602 | -0.181 | **0.659** | 2.22E-12 | **2.40E-11** |
| ENSG00000229162.1 | **RUNX3-AS1** | | lncRNA | **19.790** | 3.252 | 0.979 | **9.529** | 2.24E-12 | **2.42E-11** |
| ENSG00000184983.13 | **NDUFA6** | 4700 | protein_coding | **1283.328** | -0.893 | -0.269 | **0.538** | 2.25E-12 | **2.43E-11** |
| ENSG00000178922.20 | **HYI** | 81888 | protein_coding | **199.936** | -1.431 | -0.431 | **0.371** | 2.26E-12 | **2.44E-11** |

| | | | | | | | | | |
|---|---|---|---|---|---|---|---|---|---|
| ENSG00000131080.17 | **EDA2R** | 60401 | protein_coding | **239.802** | -0.939 | -0.283 | **0.522** | 2.26E-12 | **2.44E-11** |
| ENSG00000204311.17 | **PJVK** | 494513 | protein_coding | **154.048** | 1.196 | 0.360 | **2.291** | 2.27E-12 | **2.45E-11** |
| ENSG00000162521.20 | **RBBP4** | 5928 | protein_coding | **2689.414** | -0.582 | -0.175 | **0.668** | 2.30E-12 | **2.48E-11** |
| ENSG00000310496.1 | **ENSG00000310496** | | lncRNA | **217.094** | -1.262 | -0.380 | **0.417** | 2.32E-12 | **2.50E-11** |
| ENSG00000080608.12 | **PUM3** | 9933 | protein_coding | **576.315** | -0.983 | -0.296 | **0.506** | 2.32E-12 | **2.50E-11** |
| ENSG00000006062.20 | **MAP3K14** | 9020 | protein_coding | **960.169** | 0.870 | 0.262 | **1.827** | 2.33E-12 | **2.51E-11** |
| ENSG00000239306.7 | **RBM14** | 10432 | protein_coding | **917.284** | -0.905 | -0.273 | **0.534** | 2.35E-12 | **2.53E-11** |
| ENSG00000166888.15 | **STAT6** | 6778 | protein_coding | **5215.742** | -0.861 | -0.259 | **0.551** | 2.39E-12 | **2.57E-11** |
| ENSG00000165507.11 | **DEPP1** | 11067 | protein_coding | **234.036** | 1.557 | 0.469 | **2.942** | 2.41E-12 | **2.60E-11** |
| ENSG00000137198.12 | **GMPR** | 2766 | protein_coding | **341.445** | 1.604 | 0.483 | **3.041** | 2.45E-12 | **2.63E-11** |
| ENSG00000100162.18 | **CENPM** | 79019 | protein_coding | **559.957** | -1.233 | -0.371 | **0.425** | 2.51E-12 | **2.69E-11** |
| ENSG00000237248.7 | **LINC00987** | 10049940 | lncRNA | **25.212** | 2.651 | 0.798 | **6.280** | 2.62E-12 | **2.81E-11** |
| ENSG00000107281.12 | **NPDC1** | 56654 | protein_coding | **613.661** | 3.553 | 1.070 | **11.741** | 2.69E-12 | **2.89E-11** |
| ENSG00000166845.17 | **C18orf54** | 162681 | protein_coding | **69.378** | -1.527 | -0.460 | **0.347** | 2.71E-12 | **2.91E-11** |
| ENSG00000169894.20 | **MUC3A** | 4584 | protein_coding | **26.682** | 3.890 | 1.171 | **14.824** | 2.79E-12 | **2.99E-11** |
| ENSG00000118260.17 | **CREB1** | 1385 | protein_coding | **573.858** | -0.768 | -0.231 | **0.587** | 2.84E-12 | **3.04E-11** |
| ENSG00000133958.15 | **UNC79** | 57578 | protein_coding | **27.299** | 2.901 | 0.873 | **7.470** | 2.84E-12 | **3.04E-11** |
| ENSG00000223745.9 | **CCDC18-AS1** | | lncRNA | **1611.937** | 0.768 | 0.231 | **1.703** | 2.88E-12 | **3.08E-11** |
| ENSG00000197747.11 | **S100A10** | 6281 | protein_coding | **82.131** | 1.786 | 0.538 | **3.449** | 2.93E-12 | **3.14E-11** |
| ENSG00000277632.2 | **CCL3** | 6348 | protein_coding | **5063.458** | -1.197 | -0.360 | **0.436** | 3.00E-12 | **3.21E-11** |
| ENSG00000112685.16 | **EXOC2** | 55770 | protein_coding | **487.048** | 0.766 | 0.231 | **1.700** | 3.02E-12 | **3.22E-11** |
| ENSG00000167800.11 | **TBX10** | 347853 | protein_coding | **49.594** | 2.510 | 0.756 | **5.696** | 3.03E-12 | **3.23E-11** |
| ENSG00000286086.2 | **TMEM41A-DT** | | lncRNA | **31.310** | 1.958 | 0.589 | **3.884** | 3.06E-12 | **3.27E-11** |
| ENSG00000103550.16 | **KNOP1** | 400506 | protein_coding | **279.612** | -0.951 | -0.286 | **0.517** | 3.10E-12 | **3.30E-11** |
| ENSG00000076344.18 | **RGS11** | 8786 | protein_coding | **50.444** | 3.920 | 1.180 | **15.139** | 3.19E-12 | **3.39E-11** |
| ENSG00000198719.11 | **DLL1** | 28514 | protein_coding | **163.521** | 1.563 | 0.471 | **2.955** | 3.25E-12 | **3.46E-11** |
| ENSG00000187634.15 | **SAMD11** | 148398 | protein_coding | **19.589** | 4.596 | 1.383 | **24.182** | 3.28E-12 | **3.49E-11** |
| ENSG00000117899.13 | **MESD** | 23184 | protein_coding | **553.547** | -0.851 | -0.256 | **0.554** | 3.29E-12 | **3.50E-11** |
| ENSG00000186854.13 | **TRABD2A** | 129293 | protein_coding | **63.988** | 1.965 | 0.591 | **3.903** | 3.35E-12 | **3.56E-11** |
| ENSG00000022556.18 | **NLRP2** | 55655 | protein_coding | **576.707** | 1.386 | 0.417 | **2.614** | 3.37E-12 | **3.58E-11** |
| ENSG00000186265.13 | **BTLA** | 151888 | protein_coding | **608.056** | -1.479 | -0.445 | **0.359** | 3.41E-12 | **3.62E-11** |

| | | | | | | | | | |
|---|---|---|---|---|---|---|---|---|---|
| ENSG00000181104.8 | **F2R** | 2149 | protein_coding | **99.240** | 1.980 | 0.596 | **3.946** | 3.41E-12 | **3.62E-11** |
| ENSG00000149582.18 | **TMEM25** | 84866 | protein_coding | **175.307** | 1.333 | 0.401 | **2.520** | 3.50E-12 | **3.71E-11** |
| ENSG00000159140.24 | **SON** | 6651 | protein_coding | **2407.046** | -0.597 | -0.180 | **0.661** | 3.51E-12 | **3.72E-11** |
| ENSG00000166016.8 | **ABTB2** | 25841 | protein_coding | **210.658** | 1.676 | 0.505 | **3.196** | 3.52E-12 | **3.73E-11** |
| ENSG00000184515.13 | **BEX5** | 340542 | protein_coding | **60.711** | 2.394 | 0.721 | **5.255** | 3.55E-12 | **3.76E-11** |
| ENSG00000119431.12 | **HDHD3** | 81932 | protein_coding | **401.189** | 0.600 | 0.181 | **1.516** | 3.59E-12 | **3.80E-11** |
| ENSG00000225039.3 | **LINC01058** | | lncRNA | **53.840** | 1.391 | 0.419 | **2.623** | 3.60E-12 | **3.80E-11** |
| ENSG00000133627.20 | **ACTR3B** | 57180 | protein_coding | **228.737** | 0.954 | 0.287 | **1.937** | 3.74E-12 | **3.95E-11** |
| ENSG00000152689.21 | **RASGRP3** | 25780 | protein_coding | **2622.514** | -0.969 | -0.292 | **0.511** | 3.77E-12 | **3.99E-11** |
| ENSG00000113448.21 | **PDE4D** | 5144 | protein_coding | **56.105** | -1.418 | -0.427 | **0.374** | 3.81E-12 | **4.03E-11** |
| ENSG00000143367.19 | **TUFT1** | 7286 | protein_coding | **53.654** | 2.081 | 0.626 | **4.231** | 3.87E-12 | **4.09E-11** |
| ENSG00000161682.17 | **FAM171A2** | 284069 | protein_coding | **30.113** | 2.212 | 0.666 | **4.634** | 3.89E-12 | **4.11E-11** |
| ENSG00000170955.12 | **CAVIN3** | 112464 | protein_coding | **53.177** | -1.716 | -0.517 | **0.304** | 3.91E-12 | **4.13E-11** |
| ENSG00000144134.21 | **RABL2A** | 11159 | protein_coding | **400.293** | 1.082 | 0.326 | **2.117** | 3.95E-12 | **4.17E-11** |
| ENSG00000161082.15 | **CELF5** | 60680 | protein_coding | **13.516** | 5.312 | 1.599 | **39.718** | 3.98E-12 | **4.19E-11** |
| ENSG00000162129.17 | **CLPB** | 81570 | protein_coding | **478.340** | -0.859 | -0.259 | **0.551** | 3.99E-12 | **4.20E-11** |
| ENSG00000248538.11 | **PPP1R3B-D** | 10272488( | lncRNA | **24.382** | 2.663 | 0.801 | **6.331** | 4.00E-12 | **4.21E-11** |
| ENSG00000099999.17 | **RNF215** | 200312 | protein_coding | **366.604** | 0.955 | 0.288 | **1.939** | 4.04E-12 | **4.25E-11** |
| ENSG00000171720.12 | **HDAC3** | 8841 | protein_coding | **1663.514** | 0.665 | 0.200 | **1.585** | 4.05E-12 | **4.25E-11** |
| ENSG00000100697.19 | **DICER1** | 23405 | protein_coding | **505.234** | -0.943 | -0.284 | **0.520** | 4.05E-12 | **4.26E-11** |
| ENSG00000172006.14 | **ZNF554** | 115196 | protein_coding | **103.257** | 1.151 | 0.346 | **2.220** | 4.07E-12 | **4.27E-11** |
| ENSG00000087053.21 | **MTMR2** | 8898 | protein_coding | **299.020** | -1.113 | -0.335 | **0.462** | 4.10E-12 | **4.30E-11** |
| ENSG00000285972.2 | **CERNA2** | 642934 | lncRNA | **427.335** | -1.514 | -0.456 | **0.350** | 4.22E-12 | **4.43E-11** |
| ENSG00000274922.4 | **ENSG00000274922** | | lncRNA | **79.710** | 1.837 | 0.553 | **3.573** | 4.25E-12 | **4.46E-11** |
| ENSG00000120875.10 | **DUSP4** | 1846 | protein_coding | **1716.061** | -0.955 | -0.287 | **0.516** | 4.26E-12 | **4.46E-11** |
| ENSG00000101811.17 | **CSTF2** | 1478 | protein_coding | **375.605** | -0.811 | -0.244 | **0.570** | 4.28E-12 | **4.48E-11** |
| ENSG00000170049.12 | **KCNAB3** | 9196 | protein_coding | **180.830** | 1.061 | 0.319 | **2.086** | 4.36E-12 | **4.56E-11** |
| ENSG00000174718.14 | **RESF1** | 55196 | protein_coding | **600.700** | -0.920 | -0.277 | **0.529** | 4.36E-12 | **4.56E-11** |
| ENSG00000126217.24 | **MCF2L** | 23263 | protein_coding | **59.427** | 2.038 | 0.613 | **4.106** | 4.47E-12 | **4.67E-11** |
| ENSG00000107954.13 | **NEURL1** | 9148 | protein_coding | **23.371** | 2.558 | 0.770 | **5.888** | 4.55E-12 | **4.75E-11** |
| ENSG00000289526.3 | **ENSG00000289526** | | lncRNA | **78.342** | 1.804 | 0.543 | **3.491** | 4.56E-12 | **4.76E-11** |

| | | | | | | | | | |
|---|---|---|---|---|---|---|---|---|---|
| ENSG00000204389.11 | **HSPA1A** | 3303 | protein_coding | **580.874** | 1.369 | 0.412 | **2.583** | 4.72E-12 | **4.92E-11** |
| ENSG00000134243.14 | **SORT1** | 6272 | protein_coding | **348.878** | 1.606 | 0.483 | **3.044** | 4.83E-12 | **5.03E-11** |
| ENSG00000085415.18 | **SEH1L** | 81929 | protein_coding | **558.035** | -0.755 | -0.227 | **0.593** | 4.84E-12 | **5.05E-11** |
| ENSG00000091513.18 | **TF** | 7018 | protein_coding | **113.754** | 2.432 | 0.732 | **5.398** | 4.84E-12 | **5.05E-11** |
| ENSG00000245552.10 | **LNCRNA-IL** | 10192929! | lncRNA | **37.157** | 1.846 | 0.556 | **3.595** | 4.96E-12 | **5.17E-11** |
| ENSG00000235621.11 | **LINC00494** | 284749 | lncRNA | **347.830** | 0.907 | 0.273 | **1.875** | 4.97E-12 | **5.17E-11** |
| ENSG00000104320.17 | **NBN** | 4683 | protein_coding | **765.434** | -0.866 | -0.261 | **0.549** | 4.99E-12 | **5.19E-11** |
| ENSG00000164181.16 | **ELOVL7** | 79993 | protein_coding | **32.791** | 2.091 | 0.630 | **4.261** | 5.02E-12 | **5.21E-11** |
| ENSG00000171530.17 | **TBCA** | 6902 | protein_coding | **1636.971** | -0.758 | -0.228 | **0.591** | 5.33E-12 | **5.53E-11** |
| ENSG00000152154.14 | **TMEM178/** | 130733 | protein_coding | **46.299** | 1.772 | 0.533 | **3.415** | 5.35E-12 | **5.55E-11** |
| ENSG00000188559.18 | **RALGAPA2** | 57186 | protein_coding | **526.151** | 0.957 | 0.288 | **1.942** | 5.44E-12 | **5.65E-11** |
| ENSG00000179388.9 | **EGR3** | 1960 | protein_coding | **56.017** | -1.732 | -0.521 | **0.301** | 5.45E-12 | **5.65E-11** |
| ENSG00000174021.14 | **GNG5** | 2787 | protein_coding | **1705.967** | -0.869 | -0.262 | **0.548** | 5.56E-12 | **5.76E-11** |
| ENSG00000203778.10 | **FAM229B** | 619208 | protein_coding | **31.655** | 2.105 | 0.634 | **4.302** | 5.59E-12 | **5.79E-11** |
| ENSG00000137824.18 | **RMDN3** | 55177 | protein_coding | **765.627** | 0.710 | 0.214 | **1.636** | 5.61E-12 | **5.81E-11** |
| ENSG00000112245.15 | **PTP4A1** | 7803 | protein_coding | **678.640** | -0.663 | -0.200 | **0.632** | 5.69E-12 | **5.89E-11** |
| ENSG00000134802.20 | **SLC43A3** | 29015 | protein_coding | **2989.784** | -1.091 | -0.329 | **0.469** | 5.69E-12 | **5.89E-11** |
| ENSG00000177370.7 | **TIMM22** | 29928 | protein_coding | **272.558** | -0.730 | -0.220 | **0.603** | 5.83E-12 | **6.03E-11** |
| ENSG00000198932.15 | **GPRASP1** | 9737 | protein_coding | **77.251** | 1.409 | 0.424 | **2.655** | 5.85E-12 | **6.05E-11** |
| ENSG00000183615.8 | **FAM167B** | 84734 | protein_coding | **47.947** | 1.593 | 0.479 | **3.016** | 5.96E-12 | **6.15E-11** |
| ENSG00000120942.15 | **UBIAD1** | 29914 | protein_coding | **262.117** | -0.893 | -0.269 | **0.539** | 6.00E-12 | **6.19E-11** |
| ENSG00000233695.2 | **GAS6-AS1** | | lncRNA | **662.760** | 1.290 | 0.388 | **2.445** | 6.09E-12 | **6.28E-11** |
| ENSG00000124657.1 | **OR2B6** | 26212 | protein_coding | **26.434** | -2.777 | -0.836 | **0.146** | 6.17E-12 | **6.36E-11** |
| ENSG00000115507.12 | **OTX1** | 5013 | protein_coding | **24.129** | 2.757 | 0.830 | **6.761** | 6.25E-12 | **6.44E-11** |
| ENSG00000056972.22 | **TRAF3IP2** | 10758 | protein_coding | **545.537** | -0.711 | -0.214 | **0.611** | 6.26E-12 | **6.45E-11** |
| ENSG00000213066.16 | **CEP43** | 11116 | protein_coding | **707.369** | -0.725 | -0.218 | **0.605** | 6.27E-12 | **6.45E-11** |
| ENSG00000106012.21 | **IQCE** | 23288 | protein_coding | **206.730** | 0.996 | 0.300 | **1.995** | 6.28E-12 | **6.46E-11** |
| ENSG00000126953.10 | **TIMM8A** | 1678 | protein_coding | **141.142** | -1.524 | -0.459 | **0.348** | 6.34E-12 | **6.52E-11** |
| ENSG00000196247.15 | **ZNF107** | 51427 | protein_coding | **325.275** | -1.196 | -0.360 | **0.436** | 6.35E-12 | **6.53E-11** |
| ENSG00000106780.11 | **MEGF9** | 1955 | protein_coding | **136.564** | 1.271 | 0.383 | **2.413** | 6.51E-12 | **6.68E-11** |
| ENSG00000139160.15 | **ETFBKMT** | 254013 | protein_coding | **48.766** | 1.928 | 0.580 | **3.805** | 6.59E-12 | **6.76E-11** |

| | | | | | | | | | |
|---|---|---|---|---|---|---|---|---|---|
| ENSG00000119383.23 | **PTPA** | 5524 | protein_coding | **996.956** | -0.771 | -0.232 | **0.586** | 6.64E-12 | **6.81E-11** |
| ENSG00000151176.10 | **PLBD2** | 196463 | protein_coding | **784.112** | 0.660 | 0.199 | **1.580** | 6.66E-12 | **6.83E-11** |
| ENSG00000137814.14 | **HAUS2** | 55142 | protein_coding | **273.535** | -1.216 | -0.366 | **0.431** | 6.72E-12 | **6.89E-11** |
| ENSG00000274414.1 | **ENSG00000274414** | | lncRNA | **20.348** | 2.520 | 0.758 | **5.734** | 6.75E-12 | **6.92E-11** |
| ENSG00000025423.13 | **HSD17B6** | 8630 | protein_coding | **53.610** | 2.311 | 0.696 | **4.961** | 6.80E-12 | **6.96E-11** |
| ENSG00000106089.14 | **STX1A** | 6804 | protein_coding | **149.080** | 1.519 | 0.457 | **2.866** | 7.00E-12 | **7.16E-11** |
| ENSG00000089639.13 | **GMIP** | 51291 | protein_coding | **960.020** | -0.696 | -0.209 | **0.617** | 7.13E-12 | **7.29E-11** |
| ENSG00000160161.11 | **CILP2** | 148113 | protein_coding | **26.714** | 2.520 | 0.759 | **5.735** | 7.22E-12 | **7.38E-11** |
| ENSG00000225217.1 | **HSPA7** | | transcribed_unprc | **414.453** | 1.478 | 0.445 | **2.785** | 7.30E-12 | **7.45E-11** |
| ENSG00000135205.17 | **CCDC146** | 57639 | protein_coding | **147.795** | 1.114 | 0.335 | **2.164** | 7.40E-12 | **7.56E-11** |
| ENSG00000135299.20 | **ANKRD6** | 22881 | protein_coding | **146.010** | 2.110 | 0.635 | **4.318** | 7.41E-12 | **7.56E-11** |
| ENSG00000133985.5 | **TTC9** | 23508 | protein_coding | **457.819** | -0.839 | -0.253 | **0.559** | 7.91E-12 | **8.07E-11** |
| ENSG00000166136.19 | **NDUFB8** | 4714 | protein_coding | **2663.990** | -0.818 | -0.246 | **0.567** | 8.10E-12 | **8.25E-11** |
| ENSG00000071189.23 | **SNX13** | 23161 | protein_coding | **141.922** | -1.032 | -0.311 | **0.489** | 8.12E-12 | **8.27E-11** |
| ENSG00000133641.20 | **RLIG1** | 91298 | protein_coding | **142.080** | -1.414 | -0.426 | **0.375** | 8.21E-12 | **8.36E-11** |
| ENSG00000274528.1 | **ENSG00000274528** | | lncRNA | **48.038** | 1.885 | 0.567 | **3.693** | 8.24E-12 | **8.39E-11** |
| ENSG00000139180.13 | **NDUFA9** | 4704 | protein_coding | **1196.946** | -0.850 | -0.256 | **0.555** | 8.26E-12 | **8.40E-11** |
| ENSG00000103257.12 | **SLC7A5** | 8140 | protein_coding | **2164.770** | -1.309 | -0.394 | **0.403** | 8.28E-12 | **8.41E-11** |
| ENSG00000186283.16 | **TOR3A** | 64222 | protein_coding | **860.127** | -0.680 | -0.205 | **0.624** | 8.35E-12 | **8.48E-11** |
| ENSG00000085231.15 | **AK6** | 10215740 | protein_coding | **411.839** | -0.647 | -0.195 | **0.638** | 8.53E-12 | **8.66E-11** |
| ENSG00000290476.2 | **SUGT1P3** | 283507 | lncRNA | **177.099** | 0.929 | 0.280 | **1.905** | 8.55E-12 | **8.68E-11** |
| ENSG00000027869.15 | **SH2D2A** | 9047 | protein_coding | **119.608** | -1.494 | -0.450 | **0.355** | 8.60E-12 | **8.73E-11** |
| ENSG00000166965.16 | **RCCD1** | 91433 | protein_coding | **517.213** | -0.657 | -0.198 | **0.634** | 8.63E-12 | **8.75E-11** |
| ENSG00000172828.15 | **CES3** | 23491 | protein_coding | **243.382** | 1.140 | 0.343 | **2.204** | 8.74E-12 | **8.85E-11** |
| ENSG00000149212.13 | **SESN3** | 143686 | protein_coding | **95.640** | 1.977 | 0.595 | **3.936** | 8.77E-12 | **8.88E-11** |
| ENSG00000119906.15 | **SLF2** | 55719 | protein_coding | **349.529** | -0.841 | -0.253 | **0.558** | 9.09E-12 | **9.20E-11** |
| ENSG00000181856.17 | **SLC2A4** | 6517 | protein_coding | **18.154** | 2.788 | 0.839 | **6.907** | 9.16E-12 | **9.26E-11** |
| ENSG00000112796.12 | **ENPP5** | 59084 | protein_coding | **72.350** | 1.625 | 0.489 | **3.084** | 9.18E-12 | **9.28E-11** |
| ENSG00000308872.1 | **ENSG00000308872** | | lncRNA | **44.098** | 1.791 | 0.539 | **3.460** | 9.18E-12 | **9.28E-11** |
| ENSG00000171817.19 | **ZNF540** | 163255 | protein_coding | **75.535** | 1.223 | 0.368 | **2.335** | 9.42E-12 | **9.52E-11** |
| ENSG00000159753.16 | **CARMIL2** | 146206 | protein_coding | **1954.397** | -1.182 | -0.356 | **0.441** | 9.47E-12 | **9.56E-11** |

| | | | | | | | | | |
|---|---|---|---|---|---|---|---|---|---|
| ENSG00000147874.13 | **HAUS6** | 54801 | protein_coding | **355.458** | -0.937 | -0.282 | **0.522** | 9.50E-12 | **9.59E-11** |
| ENSG00000063241.10 | **ISOC2** | 79763 | protein_coding | **595.478** | -0.860 | -0.259 | **0.551** | 9.54E-12 | **9.62E-11** |
| ENSG00000131480.10 | **AOC2** | 314 | protein_coding | **87.892** | 1.316 | 0.396 | **2.490** | 9.57E-12 | **9.64E-11** |
| ENSG00000093010.17 | **COMT** | 1312 | protein_coding | **1018.082** | -0.853 | -0.257 | **0.554** | 9.57E-12 | **9.64E-11** |
| ENSG00000188997.10 | **KCTD21** | 283219 | protein_coding | **139.645** | 0.982 | 0.296 | **1.975** | 9.77E-12 | **9.83E-11** |
| ENSG00000178531.7 | **CTXN1** | 404217 | protein_coding | **17.067** | 3.183 | 0.958 | **9.082** | 9.80E-12 | **9.87E-11** |
| ENSG00000230438.10 | **SERPINB9P1** | | lncRNA | **201.137** | 1.136 | 0.342 | **2.198** | 1.01E-11 | **1.01E-10** |
| ENSG00000100804.21 | **PSMB5** | 5693 | protein_coding | **869.312** | -1.018 | -0.306 | **0.494** | 1.06E-11 | **1.06E-10** |
| ENSG00000173914.14 | **RBM4B** | 83759 | protein_coding | **340.526** | -0.795 | -0.239 | **0.576** | 1.06E-11 | **1.07E-10** |
| ENSG00000146386.10 | **ABRACL** | 58527 | protein_coding | **1304.048** | -0.956 | -0.288 | **0.516** | 1.06E-11 | **1.07E-10** |
| ENSG00000134153.12 | **EMC7** | 56851 | protein_coding | **524.377** | -0.702 | -0.211 | **0.615** | 1.11E-11 | **1.11E-10** |
| ENSG00000105607.16 | **GCDH** | 2639 | protein_coding | **496.822** | -0.656 | -0.197 | **0.635** | 1.13E-11 | **1.13E-10** |
| ENSG00000130766.7 | **SESN2** | 83667 | protein_coding | **840.168** | -1.268 | -0.382 | **0.415** | 1.19E-11 | **1.19E-10** |
| ENSG00000092929.15 | **UNC13D** | 201294 | protein_coding | **1354.849** | -1.032 | -0.311 | **0.489** | 1.20E-11 | **1.20E-10** |
| ENSG00000147155.13 | **EBP** | 10682 | protein_coding | **1233.181** | -1.163 | -0.350 | **0.447** | 1.24E-11 | **1.24E-10** |
| ENSG00000163050.20 | **COQ8A** | 56997 | protein_coding | **600.694** | 0.766 | 0.231 | **1.701** | 1.26E-11 | **1.26E-10** |
| ENSG00000204934.11 | **ATP6V0E2-** | 401431 | lncRNA | **57.470** | 1.419 | 0.427 | **2.674** | 1.27E-11 | **1.27E-10** |
| ENSG00000260563.3 | **ENSG00000260563** | | lncRNA | **271.239** | 0.910 | 0.274 | **1.879** | 1.31E-11 | **1.31E-10** |
| ENSG00000286264.3 | **ENSG0000(** | 11484103! | protein_coding | **321.420** | 0.773 | 0.233 | **1.709** | 1.32E-11 | **1.32E-10** |
| ENSG00000074317.13 | **SNCB** | 6620 | protein_coding | **22.361** | 2.772 | 0.834 | **6.831** | 1.36E-11 | **1.36E-10** |
| ENSG00000101197.14 | **BIRC7** | 79444 | protein_coding | **87.937** | -2.184 | -0.657 | **0.220** | 1.38E-11 | **1.37E-10** |
| ENSG00000213689.14 | **TREX1** | 11277 | protein_coding | **1093.009** | -0.951 | -0.286 | **0.517** | 1.38E-11 | **1.38E-10** |
| ENSG00000076662.13 | **ICAM3** | 3385 | protein_coding | **3057.328** | -0.827 | -0.249 | **0.564** | 1.40E-11 | **1.40E-10** |
| ENSG00000305512.1 | **ENSG00000305512** | | lncRNA | **66.316** | -1.838 | -0.553 | **0.280** | 1.43E-11 | **1.43E-10** |
| ENSG00000010810.20 | **FYN** | 2534 | protein_coding | **862.345** | 1.363 | 0.410 | **2.571** | 1.47E-11 | **1.46E-10** |
| ENSG00000229922.6 | **LINC02865** | | lncRNA | **35.085** | 2.543 | 0.766 | **5.829** | 1.48E-11 | **1.47E-10** |
| ENSG00000163565.22 | **IFI16** | 3428 | protein_coding | **4059.042** | -0.862 | -0.259 | **0.550** | 1.48E-11 | **1.48E-10** |
| ENSG00000134330.20 | **IAH1** | 285148 | protein_coding | **866.818** | -0.844 | -0.254 | **0.557** | 1.51E-11 | **1.50E-10** |
| ENSG00000105641.5 | **SLC5A5** | 6528 | protein_coding | **30.307** | 2.539 | 0.764 | **5.813** | 1.51E-11 | **1.51E-10** |
| ENSG00000147082.19 | **CCNB3** | 85417 | protein_coding | **20.269** | 3.352 | 1.009 | **10.208** | 1.54E-11 | **1.53E-10** |
| ENSG00000101546.14 | **RBFA** | 79863 | protein_coding | **536.717** | -0.678 | -0.204 | **0.625** | 1.55E-11 | **1.54E-10** |

| | | | | | | | | | |
|---|---|---|---|---|---|---|---|---|---|
| ENSG00000071205.14 | **ARHGAP1(** | 79658 | protein_coding | **970.945** | 0.765 | 0.230 | **1.699** | 1.55E-11 | **1.54E-10** |
| ENSG00000264548.1 | **ENSG00000264548** | | lncRNA | **48.644** | 1.678 | 0.505 | **3.201** | 1.58E-11 | **1.57E-10** |
| ENSG00000068438.17 | **FTSJ1** | 24140 | protein_coding | **670.523** | -0.744 | -0.224 | **0.597** | 1.61E-11 | **1.60E-10** |
| ENSG00000171204.14 | **TMEM126I** | 55863 | protein_coding | **337.673** | -0.687 | -0.207 | **0.621** | 1.63E-11 | **1.62E-10** |
| ENSG00000198951.14 | **NAGA** | 4668 | protein_coding | **646.934** | -0.838 | -0.252 | **0.559** | 1.64E-11 | **1.63E-10** |
| ENSG00000065150.23 | **IPO5** | 3843 | protein_coding | **1830.009** | -0.646 | -0.194 | **0.639** | 1.66E-11 | **1.64E-10** |
| ENSG00000170619.12 | **COMMD5** | 28991 | protein_coding | **300.774** | -0.688 | -0.207 | **0.621** | 1.68E-11 | **1.66E-10** |
| ENSG00000135972.11 | **MRPS9** | 64965 | protein_coding | **603.828** | -0.632 | -0.190 | **0.645** | 1.70E-11 | **1.68E-10** |
| ENSG00000269343.10 | **ZNF587B** | 10029351( | protein_coding | **204.312** | -0.762 | -0.229 | **0.590** | 1.73E-11 | **1.71E-10** |
| ENSG00000160862.15 | **AZGP1** | 563 | protein_coding | **23.934** | 4.184 | 1.260 | **18.182** | 1.73E-11 | **1.71E-10** |
| ENSG00000031691.9 | **CENPQ** | 55166 | protein_coding | **63.800** | -1.429 | -0.430 | **0.371** | 1.76E-11 | **1.74E-10** |
| ENSG00000256229.9 | **ZNF486** | 90649 | protein_coding | **71.610** | -1.337 | -0.403 | **0.396** | 1.77E-11 | **1.74E-10** |
| ENSG00000299246.1 | **ENSG00000299246** | | lncRNA | **22.954** | 2.825 | 0.850 | **7.085** | 1.78E-11 | **1.75E-10** |
| ENSG00000106344.10 | **RBM28** | 55131 | protein_coding | **600.088** | -0.775 | -0.233 | **0.584** | 1.79E-11 | **1.77E-10** |
| ENSG00000177606.9 | **JUN** | 3725 | protein_coding | **782.057** | 1.363 | 0.410 | **2.571** | 1.80E-11 | **1.77E-10** |
| ENSG00000309328.1 | **ENSG00000309328** | | lncRNA | **18.122** | 3.040 | 0.915 | **8.223** | 1.80E-11 | **1.78E-10** |
| ENSG00000136560.16 | **TANK** | 10010 | protein_coding | **1390.971** | 0.916 | 0.276 | **1.887** | 1.82E-11 | **1.79E-10** |
| ENSG00000295671.1 | **ENSG00000295671** | | lncRNA | **23.093** | 4.023 | 1.211 | **16.261** | 1.84E-11 | **1.81E-10** |
| ENSG00000151702.20 | **FLI1** | 2313 | protein_coding | **597.373** | -0.942 | -0.284 | **0.521** | 1.84E-11 | **1.81E-10** |
| ENSG00000179846.11 | **NKPD1** | 284353 | protein_coding | **30.705** | 2.213 | 0.666 | **4.637** | 1.84E-11 | **1.81E-10** |
| ENSG00000185522.11 | **LMNTD2** | 256329 | protein_coding | **403.665** | 1.052 | 0.317 | **2.073** | 1.88E-11 | **1.84E-10** |
| ENSG00000125895.8 | **TMEM74B** | 55321 | protein_coding | **19.810** | 2.872 | 0.864 | **7.319** | 1.88E-11 | **1.84E-10** |
| ENSG00000167105.10 | **TMEM92** | 162461 | protein_coding | **17.662** | 5.554 | 1.672 | **46.967** | 1.88E-11 | **1.84E-10** |
| ENSG00000095002.18 | **MSH2** | 4436 | protein_coding | **425.412** | -0.660 | -0.199 | **0.633** | 1.93E-11 | **1.89E-10** |
| ENSG00000135372.12 | **NAT10** | 55226 | protein_coding | **1793.109** | -0.719 | -0.216 | **0.608** | 1.93E-11 | **1.89E-10** |
| ENSG00000117318.11 | **ID3** | 3399 | protein_coding | **1218.998** | 0.932 | 0.281 | **1.909** | 2.00E-11 | **1.96E-10** |
| ENSG00000236682.5 | **MAP3K2-D** | 10050692: | lncRNA | **176.034** | 0.853 | 0.257 | **1.806** | 2.02E-11 | **1.98E-10** |
| ENSG00000181444.15 | **ZNF467** | 168544 | protein_coding | **21.885** | 2.789 | 0.839 | **6.910** | 2.04E-11 | **2.00E-10** |
| ENSG00000132478.12 | **UNK** | 85451 | protein_coding | **777.154** | -0.681 | -0.205 | **0.624** | 2.07E-11 | **2.02E-10** |
| ENSG00000154429.12 | **CCSAP** | 126731 | protein_coding | **241.657** | -1.004 | -0.302 | **0.499** | 2.09E-11 | **2.04E-10** |
| ENSG00000211772.11 | **TRBC2** | | TR_C_gene | **422.200** | -1.277 | -0.384 | **0.413** | 2.09E-11 | **2.04E-10** |

| | | | | | | | | | |
|---|---|---|---|---|---|---|---|---|---|
| ENSG00000149292.19 | **TTC12** | 54970 | protein_coding | **302.190** | 0.907 | 0.273 | **1.876** | 2.10E-11 | **2.05E-10** |
| ENSG00000058453.19 | **CROCC** | 9696 | protein_coding | **293.293** | 1.227 | 0.369 | **2.341** | 2.13E-11 | **2.08E-10** |
| ENSG00000186806.8 | **VSIG10L** | 147645 | protein_coding | **117.922** | 1.458 | 0.439 | **2.748** | 2.14E-11 | **2.09E-10** |
| ENSG00000100796.20 | **PPP4R3A** | 55671 | protein_coding | **1287.605** | -0.624 | -0.188 | **0.649** | 2.17E-11 | **2.12E-10** |
| ENSG00000238005.5 | **LNCATV** | 10192785[illegible] | lncRNA | **32.053** | 2.211 | 0.666 | **4.631** | 2.20E-11 | **2.14E-10** |
| ENSG00000063660.11 | **GPC1** | 2817 | protein_coding | **42.069** | 1.965 | 0.591 | **3.904** | 2.21E-11 | **2.16E-10** |
| ENSG00000125835.22 | **SNRPB** | 6628 | protein_coding | **4050.931** | -0.695 | -0.209 | **0.618** | 2.23E-11 | **2.17E-10** |
| ENSG00000107554.19 | **DNMBP** | 23268 | protein_coding | **919.015** | 0.741 | 0.223 | **1.671** | 2.26E-11 | **2.20E-10** |
| ENSG00000166405.17 | **RIC3** | 79608 | protein_coding | **400.002** | 0.833 | 0.251 | **1.782** | 2.28E-11 | **2.22E-10** |
| ENSG00000038002.11 | **AGA** | 175 | protein_coding | **181.467** | -1.317 | -0.396 | **0.401** | 2.32E-11 | **2.25E-10** |
| ENSG00000027697.18 | **IFNGR1** | 3459 | protein_coding | **1434.087** | 0.745 | 0.224 | **1.676** | 2.39E-11 | **2.32E-10** |
| ENSG00000141506.17 | **PIK3R5** | 23533 | protein_coding | **1186.714** | 0.792 | 0.238 | **1.731** | 2.42E-11 | **2.35E-10** |
| ENSG00000178773.17 | **CPNE7** | 27132 | protein_coding | **833.066** | -1.113 | -0.335 | **0.462** | 2.42E-11 | **2.35E-10** |
| ENSG00000076706.19 | **MCAM** | 4162 | protein_coding | **185.525** | 1.734 | 0.522 | **3.326** | 2.43E-11 | **2.36E-10** |
| ENSG00000188130.16 | **MAPK12** | 6300 | protein_coding | **1555.909** | 0.597 | 0.180 | **1.513** | 2.44E-11 | **2.37E-10** |
| ENSG00000198720.15 | **ANKRD13[illegible]** | 124930 | protein_coding | **128.967** | 1.174 | 0.353 | **2.256** | 2.48E-11 | **2.40E-10** |
| ENSG00000141030.15 | **COPS3** | 8533 | protein_coding | **822.940** | -0.685 | -0.206 | **0.622** | 2.49E-11 | **2.41E-10** |
| ENSG00000248884.3 | **ENSG00000248884** | | lncRNA | **19.966** | 3.086 | 0.929 | **8.493** | 2.50E-11 | **2.42E-10** |
| ENSG00000131788.18 | **PIAS3** | 10401 | protein_coding | **615.581** | 0.663 | 0.200 | **1.583** | 2.52E-11 | **2.43E-10** |
| ENSG00000187051.11 | **RPS19BP1** | 91582 | protein_coding | **1112.461** | -0.669 | -0.201 | **0.629** | 2.52E-11 | **2.44E-10** |
| ENSG00000161328.13 | **LRRC56** | 115399 | protein_coding | **68.017** | 1.209 | 0.364 | **2.312** | 2.53E-11 | **2.45E-10** |
| ENSG00000134333.17 | **LDHA** | 3939 | protein_coding | **19558.701** | -0.989 | -0.298 | **0.504** | 2.56E-11 | **2.47E-10** |
| ENSG00000204946.11 | **ZNF783** | 10028967[illegible] | protein_coding | **708.621** | 0.601 | 0.181 | **1.517** | 2.56E-11 | **2.47E-10** |
| ENSG00000111254.10 | **AKAP3** | 10566 | protein_coding | **50.752** | 2.111 | 0.635 | **4.319** | 2.58E-11 | **2.49E-10** |
| ENSG00000262155.3 | **LINC02175** | 283887 | lncRNA | **28.083** | 2.305 | 0.694 | **4.943** | 2.64E-11 | **2.54E-10** |
| ENSG00000100867.17 | **DHRS2** | 10202 | protein_coding | **221.784** | 3.476 | 1.046 | **11.130** | 2.69E-11 | **2.59E-10** |
| ENSG00000167600.16 | **CYP2S1** | 29785 | protein_coding | **43.514** | 2.979 | 0.897 | **7.886** | 2.73E-11 | **2.63E-10** |
| ENSG00000196923.16 | **PDLIM7** | 9260 | protein_coding | **1067.588** | 0.940 | 0.283 | **1.918** | 2.79E-11 | **2.69E-10** |
| ENSG00000156398.15 | **SFXN2** | 118980 | protein_coding | **223.308** | -0.770 | -0.232 | **0.587** | 2.81E-11 | **2.71E-10** |
| ENSG00000170684.11 | **ZNF296** | 162979 | protein_coding | **425.592** | -1.218 | -0.367 | **0.430** | 2.88E-11 | **2.77E-10** |
| ENSG00000168288.15 | **MMADHC** | 27249 | protein_coding | **818.692** | -0.914 | -0.275 | **0.531** | 3.01E-11 | **2.90E-10** |

| | | | | | | | | | |
|---|---|---|---|---|---|---|---|---|---|
| ENSG00000301550.1 | **ENSG00000301550** | | lncRNA | **11.870** | 5.015 | 1.510 | **32.335** | 3.04E-11 | **2.93E-10** |
| ENSG00000184445.13 | **KNTC1** | 9735 | protein_coding | **1498.683** | -0.927 | -0.279 | **0.526** | 3.05E-11 | **2.93E-10** |
| ENSG00000109756.12 | **RAPGEF2** | 9693 | protein_coding | **1298.470** | 0.841 | 0.253 | **1.791** | 3.12E-11 | **3.00E-10** |
| ENSG00000085871.11 | **MGST2** | 4258 | protein_coding | **249.084** | -0.997 | -0.300 | **0.501** | 3.15E-11 | **3.03E-10** |
| ENSG00000099783.15 | **HNRNPM** | 4670 | protein_coding | **2578.146** | -0.526 | -0.158 | **0.695** | 3.23E-11 | **3.10E-10** |
| ENSG00000273253.3 | **TRABD-AS:** | 12370651: | lncRNA | **84.389** | 1.109 | 0.334 | **2.157** | 3.28E-11 | **3.14E-10** |
| ENSG00000181450.19 | **ZNF678** | 339500 | protein_coding | **116.358** | -1.499 | -0.451 | **0.354** | 3.29E-11 | **3.15E-10** |
| ENSG00000116205.16 | **TCEANC2** | 127428 | protein_coding | **310.503** | -0.778 | -0.234 | **0.583** | 3.29E-11 | **3.15E-10** |
| ENSG00000204852.19 | **TCTN1** | 79600 | protein_coding | **534.240** | 0.953 | 0.287 | **1.937** | 3.36E-11 | **3.22E-10** |
| ENSG00000122566.25 | **HNRNPA2E** | 3181 | protein_coding | **14602.086** | -0.623 | -0.187 | **0.649** | 3.36E-11 | **3.22E-10** |
| ENSG00000156110.17 | **ADK** | 132 | protein_coding | **570.080** | -0.720 | -0.217 | **0.607** | 3.39E-11 | **3.24E-10** |
| ENSG00000304501.1 | **ENSG00000304501** | | lncRNA | **254.547** | 1.029 | 0.310 | **2.041** | 3.41E-11 | **3.26E-10** |
| ENSG00000130956.16 | **HABP4** | 22927 | protein_coding | **463.881** | 0.948 | 0.285 | **1.929** | 3.42E-11 | **3.27E-10** |
| ENSG00000184925.14 | **LCN12** | 286256 | protein_coding | **38.035** | 1.803 | 0.543 | **3.490** | 3.44E-11 | **3.29E-10** |
| ENSG00000111678.13 | **C12orf57** | 113246 | protein_coding | **2139.987** | -0.717 | -0.216 | **0.608** | 3.57E-11 | **3.40E-10** |
| ENSG00000269893.11 | **SNHG8** | | lncRNA | **1115.934** | -0.971 | -0.292 | **0.510** | 3.57E-11 | **3.40E-10** |
| ENSG00000130826.21 | **DKC1** | 1736 | protein_coding | **1415.759** | -0.687 | -0.207 | **0.621** | 3.57E-11 | **3.41E-10** |
| ENSG00000110881.14 | **ASIC1** | 41 | protein_coding | **34.874** | 2.429 | 0.731 | **5.385** | 3.61E-11 | **3.44E-10** |
| ENSG00000163781.16 | **TOPBP1** | 11073 | protein_coding | **797.700** | -0.898 | -0.270 | **0.537** | 3.62E-11 | **3.45E-10** |
| ENSG00000231770.9 | **TMEM44-AS1** | | lncRNA | **163.713** | 1.055 | 0.318 | **2.078** | 3.65E-11 | **3.48E-10** |
| ENSG00000168056.20 | **LTBP3** | 4054 | protein_coding | **336.926** | 1.119 | 0.337 | **2.172** | 3.67E-11 | **3.49E-10** |
| ENSG00000143612.23 | **LTAP1** | 25912 | protein_coding | **2102.359** | -0.664 | -0.200 | **0.631** | 3.70E-11 | **3.52E-10** |
| ENSG00000273356.1 | **LINC02019** | | lncRNA | **26.459** | 2.196 | 0.661 | **4.583** | 3.81E-11 | **3.62E-10** |
| ENSG00000127511.12 | **SIN3B** | 23309 | protein_coding | **1705.100** | 0.500 | 0.151 | **1.415** | 3.85E-11 | **3.66E-10** |
| ENSG00000305429.1 | **ENSG00000305429** | | lncRNA | **15.443** | 3.678 | 1.107 | **12.802** | 3.85E-11 | **3.66E-10** |
| ENSG00000173805.18 | **HAP1** | 9001 | protein_coding | **14.159** | 4.066 | 1.224 | **16.751** | 3.89E-11 | **3.69E-10** |
| ENSG00000168002.14 | **POLR2G** | 5436 | protein_coding | **997.592** | -0.811 | -0.244 | **0.570** | 3.92E-11 | **3.72E-10** |
| ENSG00000276085.1 | **CCL3L3** | 414062 | protein_coding | **1155.255** | -1.410 | -0.424 | **0.376** | 3.92E-11 | **3.72E-10** |
| ENSG00000297175.1 | **ENSG00000297175** | | lncRNA | **28.606** | 2.952 | 0.889 | **7.740** | 3.98E-11 | **3.77E-10** |
| ENSG00000249572.4 | **TARS1-DT** | | lncRNA | **107.157** | 1.189 | 0.358 | **2.280** | 3.99E-11 | **3.78E-10** |
| ENSG00000064687.13 | **ABCA7** | 10347 | protein_coding | **2380.740** | 1.368 | 0.412 | **2.582** | 4.02E-11 | **3.81E-10** |

| | | | | | | | | | |
|---|---|---|---|---|---|---|---|---|---|
| ENSG00000245849.9 | **RAD51-AS1** | 10050564 | lncRNA | **534.858** | 0.934 | 0.281 | **1.910** | 4.05E-11 | **3.84E-10** |
| ENSG00000122224.21 | **LY9** | 4063 | protein_coding | **1341.759** | 0.972 | 0.293 | **1.962** | 4.06E-11 | **3.84E-10** |
| ENSG00000285077.4 | **ARHGAP11** | 89839 | protein_coding | **101.777** | -1.167 | -0.351 | **0.445** | 4.14E-11 | **3.91E-10** |
| ENSG00000280206.1 | **ENSG00000280206** | | lncRNA | **372.393** | -1.075 | -0.324 | **0.475** | 4.22E-11 | **3.99E-10** |
| ENSG00000141503.19 | **MINK1** | 50488 | protein_coding | **2639.285** | 0.522 | 0.157 | **1.436** | 4.27E-11 | **4.03E-10** |
| ENSG00000256967.2 | **ENSG00000256967** | | lncRNA | **11.910** | 3.715 | 1.118 | **13.132** | 4.27E-11 | **4.04E-10** |
| ENSG00000158887.22 | **MPZ** | 4359 | protein_coding | **134.656** | 0.956 | 0.288 | **1.940** | 4.32E-11 | **4.08E-10** |
| ENSG00000253669.5 | **GASAL1** | | lncRNA | **62.393** | 1.754 | 0.528 | **3.373** | 4.34E-11 | **4.09E-10** |
| ENSG00000147804.11 | **SLC39A4** | 55630 | protein_coding | **302.811** | -1.015 | -0.306 | **0.495** | 4.37E-11 | **4.12E-10** |
| ENSG00000153066.15 | **TXNDC11** | 51061 | protein_coding | **2319.598** | 0.686 | 0.206 | **1.608** | 4.42E-11 | **4.17E-10** |
| ENSG00000268061.7 | **NAPA-AS1** | | lncRNA | **67.178** | -1.238 | -0.373 | **0.424** | 4.45E-11 | **4.19E-10** |
| ENSG00000167618.12 | **LAIR2** | 3904 | protein_coding | **18.993** | 4.675 | 1.407 | **25.547** | 4.47E-11 | **4.21E-10** |
| ENSG00000182628.15 | **SKA2** | 348235 | protein_coding | **643.010** | -1.076 | -0.324 | **0.474** | 4.50E-11 | **4.23E-10** |
| ENSG00000111845.7 | **PAK1IP1** | 55003 | protein_coding | **246.575** | -0.942 | -0.284 | **0.520** | 4.54E-11 | **4.27E-10** |
| ENSG00000147955.20 | **SIGMAR1** | 10280 | protein_coding | **697.795** | -0.774 | -0.233 | **0.585** | 4.54E-11 | **4.27E-10** |
| ENSG00000103005.15 | **USB1** | 79650 | protein_coding | **898.061** | 0.896 | 0.270 | **1.861** | 4.59E-11 | **4.31E-10** |
| ENSG00000122121.14 | **XPNPEP2** | 7512 | protein_coding | **49.351** | 2.216 | 0.667 | **4.647** | 4.59E-11 | **4.31E-10** |
| ENSG00000165476.16 | **REEP3** | 221035 | protein_coding | **287.052** | -0.878 | -0.264 | **0.544** | 4.60E-11 | **4.32E-10** |
| ENSG00000130649.12 | **CYP2E1** | 1571 | protein_coding | **58.428** | 1.733 | 0.522 | **3.323** | 4.60E-11 | **4.32E-10** |
| ENSG00000100372.17 | **SLC25A17** | 10478 | protein_coding | **705.251** | -0.707 | -0.213 | **0.612** | 4.67E-11 | **4.38E-10** |
| ENSG00000141738.16 | **GRB7** | 2886 | protein_coding | **35.558** | 2.724 | 0.820 | **6.608** | 4.67E-11 | **4.38E-10** |
| ENSG00000167553.19 | **TUBA1C** | 84790 | protein_coding | **3278.437** | -1.065 | -0.321 | **0.478** | 4.67E-11 | **4.38E-10** |
| ENSG00000164733.25 | **CTSB** | 1508 | protein_coding | **998.949** | 0.697 | 0.210 | **1.621** | 4.71E-11 | **4.41E-10** |
| ENSG00000133216.18 | **EPHB2** | 2048 | protein_coding | **35.427** | 2.190 | 0.659 | **4.562** | 4.77E-11 | **4.46E-10** |
| ENSG00000099866.18 | **MADCAM1** | 8174 | protein_coding | **35.254** | 2.355 | 0.709 | **5.116** | 4.79E-11 | **4.48E-10** |
| ENSG00000134291.14 | **TMEM106** | 79022 | protein_coding | **663.052** | -0.745 | -0.224 | **0.597** | 4.79E-11 | **4.48E-10** |
| ENSG00000168569.10 | **TMEM223** | 79064 | protein_coding | **214.717** | -0.963 | -0.290 | **0.513** | 4.79E-11 | **4.48E-10** |
| ENSG00000123453.21 | **SARDH** | 1757 | protein_coding | **53.077** | 1.849 | 0.557 | **3.602** | 4.80E-11 | **4.48E-10** |
| ENSG00000273061.4 | **CDC37L1-DT** | | lncRNA | **47.173** | 1.677 | 0.505 | **3.197** | 4.81E-11 | **4.49E-10** |
| ENSG00000089775.14 | **ZBTB25** | 7597 | protein_coding | **856.557** | -0.763 | -0.230 | **0.589** | 4.81E-11 | **4.49E-10** |
| ENSG00000132793.14 | **LPIN3** | 64900 | protein_coding | **79.728** | 1.816 | 0.547 | **3.522** | 4.85E-11 | **4.52E-10** |

| | | | | | | | | | |
|---|---|---|---|---|---|---|---|---|---|
| ENSG00000058404.23 | **CAMK2B** | 816 | protein_coding | **52.711** | 2.159 | 0.650 | **4.467** | 4.86E-11 | **4.53E-10** |
| ENSG00000114686.11 | **MRPL3** | 11222 | protein_coding | **1180.164** | -0.883 | -0.266 | **0.542** | 4.90E-11 | **4.57E-10** |
| ENSG00000163584.20 | **RPL22L1** | 200916 | protein_coding | **3659.591** | -1.142 | -0.344 | **0.453** | 4.93E-11 | **4.59E-10** |
| ENSG00000059804.20 | **SLC2A3** | 6515 | protein_coding | **1010.680** | -1.021 | -0.307 | **0.493** | 5.03E-11 | **4.68E-10** |
| ENSG00000225518.4 | **LINC01703** | 10537311 | lncRNA | **20.442** | 2.960 | 0.891 | **7.781** | 5.16E-11 | **4.80E-10** |
| ENSG00000171490.15 | **RSL1D1** | 26156 | protein_coding | **2714.502** | -0.674 | -0.203 | **0.627** | 5.19E-11 | **4.82E-10** |
| ENSG00000185634.13 | **SHC4** | 399694 | protein_coding | **18.383** | 2.688 | 0.809 | **6.446** | 5.28E-11 | **4.91E-10** |
| ENSG00000285804.2 | **ENSG00000285804** | | lncRNA | **58.649** | 1.433 | 0.431 | **2.701** | 5.29E-11 | **4.91E-10** |
| ENSG00000138100.16 | **TRIM54** | 57159 | protein_coding | **20.311** | 2.819 | 0.849 | **7.057** | 5.31E-11 | **4.93E-10** |
| ENSG00000161958.13 | **FGF11** | 2256 | protein_coding | **614.304** | -0.924 | -0.278 | **0.527** | 5.36E-11 | **4.97E-10** |
| ENSG00000226137.6 | **BAIAP2-DT** | 440465 | lncRNA | **22.276** | 3.574 | 1.076 | **11.910** | 5.46E-11 | **5.07E-10** |
| ENSG00000114770.19 | **ABCC5** | 10057 | protein_coding | **270.157** | 0.986 | 0.297 | **1.980** | 5.70E-11 | **5.28E-10** |
| ENSG00000099904.18 | **ZDHHC8** | 29801 | protein_coding | **903.892** | 0.671 | 0.202 | **1.592** | 5.73E-11 | **5.31E-10** |
| ENSG00000143155.15 | **TIPRL** | 261726 | protein_coding | **645.839** | -0.803 | -0.242 | **0.573** | 5.80E-11 | **5.37E-10** |
| ENSG00000267221.4 | **C17orf113** | 11080629 | protein_coding | **23.965** | 2.112 | 0.636 | **4.323** | 5.88E-11 | **5.44E-10** |
| ENSG00000177685.19 | **CRACR2B** | 283229 | protein_coding | **440.352** | 1.152 | 0.347 | **2.222** | 5.89E-11 | **5.45E-10** |
| ENSG00000169251.15 | **NMD3** | 51068 | protein_coding | **422.778** | -0.676 | -0.204 | **0.626** | 5.97E-11 | **5.52E-10** |
| ENSG00000153006.18 | **SREK1IP1** | 285672 | protein_coding | **312.008** | -1.177 | -0.354 | **0.442** | 5.99E-11 | **5.54E-10** |
| ENSG00000144712.15 | **CAND2** | 23066 | protein_coding | **59.994** | 1.645 | 0.495 | **3.128** | 6.24E-11 | **5.76E-10** |
| ENSG00000114745.17 | **GORASP1** | 64689 | protein_coding | **639.895** | 0.604 | 0.182 | **1.520** | 6.26E-11 | **5.78E-10** |
| ENSG00000105197.13 | **TIMM50** | 92609 | protein_coding | **1011.666** | -0.585 | -0.176 | **0.667** | 6.41E-11 | **5.91E-10** |
| ENSG00000004142.14 | **POLDIP2** | 26073 | protein_coding | **1651.694** | -0.537 | -0.162 | **0.689** | 6.47E-11 | **5.97E-10** |
| ENSG00000187583.13 | **PLEKHN1** | 84069 | protein_coding | **142.848** | 1.487 | 0.448 | **2.803** | 6.55E-11 | **6.04E-10** |
| ENSG00000164050.15 | **PLXNB1** | 5364 | protein_coding | **621.037** | 1.025 | 0.309 | **2.035** | 6.63E-11 | **6.12E-10** |
| ENSG00000116863.13 | **ADPRS** | 54936 | protein_coding | **659.241** | -0.737 | -0.222 | **0.600** | 6.70E-11 | **6.17E-10** |
| ENSG00000099954.21 | **CECR2** | 27443 | protein_coding | **27.256** | 2.761 | 0.831 | **6.780** | 6.73E-11 | **6.20E-10** |
| ENSG00000176871.11 | **WSB2** | 55884 | protein_coding | **618.427** | -0.609 | -0.183 | **0.656** | 6.78E-11 | **6.24E-10** |
| ENSG00000143727.18 | **ACP1** | 52 | protein_coding | **1401.167** | -0.766 | -0.231 | **0.588** | 6.88E-11 | **6.33E-10** |
| ENSG00000215769.9 | **ARHGAP27** | 10928655 | lncRNA | **171.956** | 1.323 | 0.398 | **2.501** | 6.90E-11 | **6.35E-10** |
| ENSG00000146858.9 | **ZC3HAV1L** | 92092 | protein_coding | **298.912** | -0.739 | -0.222 | **0.599** | 6.91E-11 | **6.35E-10** |
| ENSG00000142621.23 | **FHAD1** | 114827 | protein_coding | **21.035** | 3.219 | 0.969 | **9.310** | 6.99E-11 | **6.43E-10** |

| | | | | | | | | | |
|---|---|---|---|---|---|---|---|---|---|
| ENSG00000124209.6 | **RAB22A** | 57403 | protein_coding | **321.374** | -0.920 | -0.277 | **0.529** | 7.13E-11 | **6.54E-10** |
| ENSG00000121716.22 | **PILRB** | 29990 | protein_coding | **3498.080** | 0.690 | 0.208 | **1.613** | 7.17E-11 | **6.59E-10** |
| ENSG00000103569.13 | **AQP9** | 366 | protein_coding | **54.682** | 1.540 | 0.464 | **2.908** | 7.20E-11 | **6.60E-10** |
| ENSG00000144029.16 | **MRPS5** | 64969 | protein_coding | **1391.986** | -0.438 | -0.132 | **0.738** | 7.27E-11 | **6.67E-10** |
| ENSG00000166197.19 | **NOLC1** | 9221 | protein_coding | **1728.349** | -0.751 | -0.226 | **0.594** | 7.41E-11 | **6.80E-10** |
| ENSG00000167100.17 | **SAMD14** | 201191 | protein_coding | **90.188** | 1.741 | 0.524 | **3.343** | 7.48E-11 | **6.85E-10** |
| ENSG00000120915.17 | **EPHX2** | 2053 | protein_coding | **231.421** | 0.940 | 0.283 | **1.919** | 7.55E-11 | **6.91E-10** |
| ENSG00000154114.15 | **TBCEL** | 219899 | protein_coding | **319.623** | 0.834 | 0.251 | **1.782** | 7.56E-11 | **6.92E-10** |
| ENSG00000198825.17 | **INPP5F** | 22876 | protein_coding | **154.952** | 1.033 | 0.311 | **2.047** | 7.58E-11 | **6.93E-10** |
| ENSG00000213760.12 | **ATP6V1G2** | 534 | protein_coding | **60.190** | 1.524 | 0.459 | **2.876** | 7.65E-11 | **6.99E-10** |
| ENSG00000278743.3 | **ENSG00000278743** | | lncRNA | **50.192** | 1.736 | 0.523 | **3.331** | 7.65E-11 | **6.99E-10** |
| ENSG00000071994.12 | **PDCD2** | 5134 | protein_coding | **1124.756** | -0.627 | -0.189 | **0.648** | 7.67E-11 | **7.00E-10** |
| ENSG00000114125.15 | **RNF7** | 9616 | protein_coding | **799.158** | -0.877 | -0.264 | **0.545** | 7.72E-11 | **7.04E-10** |
| ENSG00000198176.15 | **TFDP1** | 7027 | protein_coding | **1316.454** | -0.774 | -0.233 | **0.585** | 7.78E-11 | **7.10E-10** |
| ENSG00000136861.21 | **CDK5RAP2** | 55755 | protein_coding | **947.247** | -0.860 | -0.259 | **0.551** | 7.90E-11 | **7.21E-10** |
| ENSG00000074964.19 | **ARHGEF10** | 55160 | protein_coding | **49.892** | 2.180 | 0.656 | **4.530** | 7.91E-11 | **7.21E-10** |
| ENSG00000162694.16 | **EXTL2** | 2135 | protein_coding | **246.469** | 1.088 | 0.328 | **2.126** | 7.92E-11 | **7.21E-10** |
| ENSG00000293855.1 | **ENSG00000** | 124909363 | lncRNA | **18.261** | 2.761 | 0.831 | **6.780** | 7.97E-11 | **7.26E-10** |
| ENSG00000214562.16 | **NUTM2D** | 728130 | protein_coding | **70.836** | 1.560 | 0.470 | **2.949** | 8.01E-11 | **7.29E-10** |
| ENSG00000291202.2 | **ENSG00000291202** | | lncRNA | **574.102** | 0.771 | 0.232 | **1.707** | 8.06E-11 | **7.34E-10** |
| ENSG00000287817.1 | **ENSG00000287817** | | lncRNA | **46.486** | 1.975 | 0.594 | **3.931** | 8.19E-11 | **7.45E-10** |
| ENSG00000067840.15 | **PDZD4** | 57595 | protein_coding | **14.466** | 3.601 | 1.084 | **12.135** | 8.43E-11 | **7.66E-10** |
| ENSG00000111679.19 | **PTPN6** | 5777 | protein_coding | **11422.335** | -1.211 | -0.364 | **0.432** | 8.46E-11 | **7.69E-10** |
| ENSG00000122687.21 | **MRM2** | 29960 | protein_coding | **742.141** | -0.582 | -0.175 | **0.668** | 8.47E-11 | **7.69E-10** |
| ENSG00000177556.14 | **ATOX1** | 475 | protein_coding | **1899.469** | 0.986 | 0.297 | **1.981** | 8.67E-11 | **7.87E-10** |
| ENSG00000155034.23 | **FBXL18** | 80028 | protein_coding | **285.990** | 0.887 | 0.267 | **1.850** | 8.93E-11 | **8.10E-10** |
| ENSG00000134508.15 | **CABLES1** | 91768 | protein_coding | **1015.573** | -0.679 | -0.204 | **0.625** | 8.93E-11 | **8.10E-10** |
| ENSG00000181804.17 | **SLC9A9** | 285195 | protein_coding | **212.785** | 1.156 | 0.348 | **2.228** | 9.12E-11 | **8.26E-10** |
| ENSG00000155629.17 | **PIK3AP1** | 118788 | protein_coding | **803.398** | -0.805 | -0.242 | **0.572** | 9.13E-11 | **8.27E-10** |
| ENSG00000056487.19 | **PHF21B** | 112885 | protein_coding | **13.643** | 3.301 | 0.994 | **9.859** | 9.23E-11 | **8.35E-10** |
| ENSG00000138668.22 | **HNRNPD** | 3184 | protein_coding | **5694.292** | -0.524 | -0.158 | **0.695** | 9.35E-11 | **8.46E-10** |

| | | | | | | | | | |
|---|---|---|---|---|---|---|---|---|---|
| ENSG00000144381.20 | **HSPD1** | 3329 | protein_coding | **6440.626** | -0.715 | -0.215 | **0.609** | 9.48E-11 | **8.58E-10** |
| ENSG00000134668.15 | **SPOCD1** | 90853 | protein_coding | **20.155** | 3.927 | 1.182 | **15.208** | 9.62E-11 | **8.70E-10** |
| ENSG00000177398.19 | **UMODL1** | 89766 | protein_coding | **39.358** | 3.025 | 0.911 | **8.143** | 9.79E-11 | **8.85E-10** |
| ENSG00000151692.17 | **RNF144A** | 9781 | protein_coding | **253.764** | 0.949 | 0.286 | **1.931** | 9.83E-11 | **8.88E-10** |
| ENSG00000076826.12 | **CAMSAP3** | 57662 | protein_coding | **248.445** | 1.420 | 0.427 | **2.676** | 9.92E-11 | **8.96E-10** |
| ENSG00000070759.19 | **TESK2** | 10420 | protein_coding | **435.489** | 0.990 | 0.298 | **1.986** | 9.96E-11 | **8.99E-10** |
| ENSG00000063180.11 | **CA11** | 770 | protein_coding | **197.088** | 1.291 | 0.388 | **2.446** | 1.00E-10 | **9.05E-10** |
| ENSG00000129534.16 | **MIS18BP1** | 55320 | protein_coding | **261.908** | -1.064 | -0.320 | **0.478** | 1.02E-10 | **9.23E-10** |
| ENSG00000113163.20 | **CERT1** | 10087 | protein_coding | **394.039** | -1.020 | -0.307 | **0.493** | 1.03E-10 | **9.25E-10** |
| ENSG00000133392.21 | **MYH11** | 4629 | protein_coding | **40.949** | 1.843 | 0.555 | **3.587** | 1.03E-10 | **9.30E-10** |
| ENSG00000099953.12 | **MMP11** | 4320 | protein_coding | **24.533** | 2.174 | 0.655 | **4.514** | 1.04E-10 | **9.37E-10** |
| ENSG00000303405.1 | **ENSG00000303405** | | lncRNA | **127.469** | -1.290 | -0.388 | **0.409** | 1.05E-10 | **9.49E-10** |
| ENSG00000129235.13 | **TXNDC17** | 84817 | protein_coding | **557.946** | -0.913 | -0.275 | **0.531** | 1.09E-10 | **9.81E-10** |
| ENSG00000250571.9 | **GLI4** | 2738 | protein_coding | **629.616** | 0.853 | 0.257 | **1.806** | 1.12E-10 | **1.01E-09** |
| ENSG00000169951.12 | **ZNF764** | 92595 | protein_coding | **111.019** | -0.972 | -0.293 | **0.510** | 1.13E-10 | **1.01E-09** |
| ENSG00000214517.12 | **PPME1** | 51400 | protein_coding | **970.859** | -0.674 | -0.203 | **0.627** | 1.14E-10 | **1.02E-09** |
| ENSG00000104870.15 | **FCGRT** | 2217 | protein_coding | **114.201** | 1.800 | 0.542 | **3.481** | 1.14E-10 | **1.03E-09** |
| ENSG00000108961.16 | **RANGRF** | 29098 | protein_coding | **562.219** | -0.673 | -0.202 | **0.627** | 1.14E-10 | **1.03E-09** |
| ENSG00000183048.14 | **SLC25A10** | 1468 | protein_coding | **251.863** | -1.279 | -0.385 | **0.412** | 1.15E-10 | **1.03E-09** |
| ENSG00000169413.5 | **RNASE6** | 6039 | protein_coding | **163.729** | -1.706 | -0.514 | **0.306** | 1.16E-10 | **1.04E-09** |
| ENSG00000291099.2 | **ENSG00000291099** | | lncRNA | **61.962** | 1.463 | 0.440 | **2.757** | 1.16E-10 | **1.04E-09** |
| ENSG00000231249.4 | **ITPR1-DT** | 10099653[illegible] | lncRNA | **34.139** | 1.735 | 0.522 | **3.329** | 1.17E-10 | **1.05E-09** |
| ENSG00000167641.13 | **PPP1R14A** | 94274 | protein_coding | **18.199** | 4.347 | 1.309 | **20.352** | 1.18E-10 | **1.06E-09** |
| ENSG00000168062.11 | **BATF2** | 116071 | protein_coding | **36.066** | 2.075 | 0.625 | **4.214** | 1.19E-10 | **1.07E-09** |
| ENSG00000253210.5 | **DENND3-A** | 10192796[illegible] | lncRNA | **104.478** | 1.089 | 0.328 | **2.127** | 1.19E-10 | **1.07E-09** |
| ENSG00000154582.20 | **ELOC** | 6921 | protein_coding | **999.845** | -0.542 | -0.163 | **0.687** | 1.20E-10 | **1.07E-09** |
| ENSG00000129128.14 | **SPCS3** | 60559 | protein_coding | **1556.360** | -0.574 | -0.173 | **0.672** | 1.21E-10 | **1.08E-09** |
| ENSG00000133561.17 | **GIMAP6** | 474344 | protein_coding | **109.433** | -1.346 | -0.405 | **0.393** | 1.21E-10 | **1.08E-09** |
| ENSG00000273148.3 | **LCDR** | 10027080[illegible] | lncRNA | **105.660** | 1.042 | 0.314 | **2.060** | 1.22E-10 | **1.09E-09** |
| ENSG00000269954.2 | **BLK-AS1** | | lncRNA | **33.632** | 1.859 | 0.560 | **3.627** | 1.25E-10 | **1.12E-09** |
| ENSG00000117500.15 | **TMED5** | 50999 | protein_coding | **439.332** | -0.888 | -0.267 | **0.540** | 1.27E-10 | **1.13E-09** |

| | | | | | | | | | |
|---|---|---|---|---|---|---|---|---|---|
| ENSG00000221829.13 | **FANCG** | 2189 | protein_coding | **594.778** | -1.199 | -0.361 | **0.436** | 1.27E-10 | **1.13E-09** |
| ENSG00000154914.19 | **USP43** | 124739 | protein_coding | **47.579** | 2.124 | 0.639 | **4.358** | 1.28E-10 | **1.14E-09** |
| ENSG00000242265.6 | **PEG10** | 23089 | protein_coding | **1049.878** | 1.336 | 0.402 | **2.525** | 1.28E-10 | **1.14E-09** |
| ENSG00000198169.11 | **ZNF251** | 90987 | protein_coding | **548.866** | 0.774 | 0.233 | **1.710** | 1.29E-10 | **1.15E-09** |
| ENSG00000232859.12 | **LYRM9** | 201229 | protein_coding | **218.467** | 0.838 | 0.252 | **1.788** | 1.30E-10 | **1.16E-09** |
| ENSG00000110218.11 | **PANX1** | 24145 | protein_coding | **156.066** | -0.950 | -0.286 | **0.518** | 1.31E-10 | **1.17E-09** |
| ENSG00000186815.15 | **TPCN1** | 53373 | protein_coding | **4695.782** | 0.684 | 0.206 | **1.607** | 1.32E-10 | **1.18E-09** |
| ENSG00000186132.17 | **C2orf76** | 130355 | protein_coding | **254.435** | 0.800 | 0.241 | **1.741** | 1.33E-10 | **1.18E-09** |
| ENSG00000120053.14 | **GOT1** | 2805 | protein_coding | **749.798** | -0.931 | -0.280 | **0.524** | 1.35E-10 | **1.20E-09** |
| ENSG00000119777.22 | **TMEM214** | 54867 | protein_coding | **1540.494** | -0.852 | -0.257 | **0.554** | 1.38E-10 | **1.23E-09** |
| ENSG00000267257.1 | **ENSG00000267257** | | lncRNA | **285.119** | 1.438 | 0.433 | **2.710** | 1.39E-10 | **1.24E-09** |
| ENSG00000175832.15 | **ETV4** | 2118 | protein_coding | **326.888** | -1.366 | -0.411 | **0.388** | 1.40E-10 | **1.24E-09** |
| ENSG00000188859.8 | **FAM78B** | 149297 | protein_coding | **93.172** | 1.639 | 0.493 | **3.114** | 1.40E-10 | **1.24E-09** |
| ENSG00000134285.13 | **FKBP11** | 51303 | protein_coding | **1207.332** | -1.100 | -0.331 | **0.466** | 1.42E-10 | **1.26E-09** |
| ENSG00000213722.12 | **DDAH2** | 23564 | protein_coding | **314.836** | 1.262 | 0.380 | **2.398** | 1.42E-10 | **1.26E-09** |
| ENSG00000224905.9 | **ENSG0000(** | 10537713( | lncRNA | **57.648** | -1.320 | -0.397 | **0.401** | 1.43E-10 | **1.26E-09** |
| ENSG00000140350.17 | **ANP32A** | 8125 | protein_coding | **3205.448** | -0.773 | -0.233 | **0.585** | 1.44E-10 | **1.27E-09** |
| ENSG00000156738.20 | **MS4A1** | 931 | protein_coding | **17559.434** | 0.673 | 0.203 | **1.594** | 1.44E-10 | **1.28E-09** |
| ENSG00000309324.1 | **ENSG0000(** | 12490785( | lncRNA | **110.442** | 1.511 | 0.455 | **2.851** | 1.46E-10 | **1.29E-09** |
| ENSG00000172687.16 | **ZNF738** | 148203 | protein_coding | **260.075** | -0.885 | -0.266 | **0.542** | 1.47E-10 | **1.30E-09** |
| ENSG00000287733.2 | **ENSG00000287733** | | lncRNA | **114.998** | -1.454 | -0.438 | **0.365** | 1.52E-10 | **1.35E-09** |
| ENSG00000132434.12 | **LANCL2** | 55915 | protein_coding | **254.910** | -0.887 | -0.267 | **0.541** | 1.54E-10 | **1.36E-09** |
| ENSG00000197134.14 | **ZNF257** | 113835 | protein_coding | **41.428** | -2.477 | -0.746 | **0.180** | 1.58E-10 | **1.39E-09** |
| ENSG00000064300.10 | **NGFR** | 4804 | protein_coding | **35.466** | 1.886 | 0.568 | **3.696** | 1.62E-10 | **1.43E-09** |
| ENSG00000146094.18 | **DOK3** | 79930 | protein_coding | **2900.878** | -1.099 | -0.331 | **0.467** | 1.64E-10 | **1.45E-09** |
| ENSG00000148225.19 | **WDR31** | 114987 | protein_coding | **30.711** | 1.721 | 0.518 | **3.297** | 1.66E-10 | **1.46E-09** |
| ENSG00000223960.10 | **CHROMR** | 10192702( | lncRNA | **355.661** | -0.874 | -0.263 | **0.545** | 1.69E-10 | **1.49E-09** |
| ENSG00000246339.7 | **EXTL3-AS1** | 10192940( | lncRNA | **29.883** | 1.930 | 0.581 | **3.810** | 1.70E-10 | **1.49E-09** |
| ENSG00000106723.19 | **SPIN1** | 10927 | protein_coding | **325.127** | 1.117 | 0.336 | **2.170** | 1.70E-10 | **1.50E-09** |
| ENSG00000169194.10 | **IL13** | 3596 | protein_coding | **31.573** | 3.290 | 0.990 | **9.781** | 1.75E-10 | **1.54E-09** |
| ENSG00000283959.3 | **ENSG00000283959** | | lncRNA | **130.777** | 1.362 | 0.410 | **2.570** | 1.79E-10 | **1.57E-09** |

| | | | | | | | | | |
|---|---|---|---|---|---|---|---|---|---|
| ENSG00000232162.3 | **USP12-AS1** | | lncRNA | **28.311** | 2.426 | 0.730 | **5.375** | 1.79E-10 | **1.57E-09** |
| ENSG00000072310.20 | **SREBF1** | 6720 | protein_coding | **1950.303** | 0.705 | 0.212 | **1.630** | 1.81E-10 | **1.59E-09** |
| ENSG00000084112.17 | **SSH1** | 54434 | protein_coding | **1150.410** | 0.730 | 0.220 | **1.658** | 1.81E-10 | **1.59E-09** |
| ENSG00000286646.3 | **MAP3K5-AS2** | | lncRNA | **383.880** | 1.454 | 0.438 | **2.740** | 1.83E-10 | **1.61E-09** |
| ENSG00000104951.18 | **IL4I1** | 259307 | protein_coding | **3273.383** | -1.056 | -0.318 | **0.481** | 1.84E-10 | **1.61E-09** |
| ENSG00000204308.10 | **RNF5** | 6048 | protein_coding | **910.698** | -0.733 | -0.221 | **0.602** | 1.85E-10 | **1.62E-09** |
| ENSG00000261949.8 | **GFY** | 10050700З | protein_coding | **9.079** | 5.861 | 1.764 | **58.126** | 1.85E-10 | **1.62E-09** |
| ENSG00000130810.22 | **PPAN** | 56342 | protein_coding | **1774.238** | -0.835 | -0.251 | **0.561** | 1.86E-10 | **1.63E-09** |
| ENSG00000164078.17 | **MST1R** | 4486 | protein_coding | **148.536** | 1.851 | 0.557 | **3.606** | 1.88E-10 | **1.65E-09** |
| ENSG00000106635.10 | **BCL7B** | 9275 | protein_coding | **633.560** | -0.762 | -0.229 | **0.590** | 1.91E-10 | **1.67E-09** |
| ENSG00000198346.13 | **ZNF813** | 126017 | protein_coding | **69.500** | -1.262 | -0.380 | **0.417** | 1.93E-10 | **1.69E-09** |
| ENSG00000121957.17 | **GPSM2** | 29899 | protein_coding | **135.426** | -1.420 | -0.427 | **0.374** | 1.94E-10 | **1.70E-09** |
| ENSG00000180370.12 | **PAK2** | 5062 | protein_coding | **971.023** | -0.657 | -0.198 | **0.634** | 1.94E-10 | **1.70E-09** |
| ENSG00000054179.14 | **ENTPD2** | 954 | protein_coding | **1287.262** | 3.628 | 1.092 | **12.366** | 1.94E-10 | **1.70E-09** |
| ENSG00000137821.14 | **LRRC49** | 54839 | protein_coding | **60.142** | 1.926 | 0.580 | **3.800** | 1.96E-10 | **1.71E-09** |
| ENSG00000196449.6 | **YRDC** | 79693 | protein_coding | **344.427** | -0.638 | -0.192 | **0.642** | 1.96E-10 | **1.71E-09** |
| ENSG00000182379.11 | **NXPH4** | 11247 | protein_coding | **90.710** | 1.780 | 0.536 | **3.434** | 1.98E-10 | **1.72E-09** |
| ENSG00000127533.5 | **F2RL3** | 9002 | protein_coding | **42.247** | -2.452 | -0.738 | **0.183** | 1.98E-10 | **1.72E-09** |
| ENSG00000047644.21 | **WWC3** | 55841 | protein_coding | **516.231** | 0.660 | 0.199 | **1.580** | 1.99E-10 | **1.73E-09** |
| ENSG00000184271.19 | **POU6F1** | 5463 | protein_coding | **172.052** | -1.021 | -0.307 | **0.493** | 2.01E-10 | **1.75E-09** |
| ENSG00000119514.9 | **GALNT12** | 79695 | protein_coding | **79.585** | 1.490 | 0.449 | **2.809** | 2.01E-10 | **1.75E-09** |
| ENSG00000140543.17 | **DET1** | 55070 | protein_coding | **57.596** | 1.668 | 0.502 | **3.178** | 2.03E-10 | **1.77E-09** |
| ENSG00000175611.15 | **ERCC6L2-A** | 10012878‹ | lncRNA | **143.987** | 1.069 | 0.322 | **2.098** | 2.05E-10 | **1.79E-09** |
| ENSG00000170476.17 | **MZB1** | 51237 | protein_coding | **4485.378** | -1.047 | -0.315 | **0.484** | 2.09E-10 | **1.81E-09** |
| ENSG00000234465.12 | **PINLYP** | 390940 | protein_coding | **97.523** | -1.592 | -0.479 | **0.332** | 2.09E-10 | **1.81E-09** |
| ENSG00000126878.15 | **AIF1L** | 83543 | protein_coding | **15.605** | 2.964 | 0.892 | **7.803** | 2.11E-10 | **1.84E-09** |
| ENSG00000149115.16 | **TNKS1BP1** | 85456 | protein_coding | **660.539** | 1.052 | 0.317 | **2.074** | 2.13E-10 | **1.85E-09** |
| ENSG00000167264.20 | **DUS2** | 54920 | protein_coding | **386.313** | -0.654 | -0.197 | **0.636** | 2.16E-10 | **1.87E-09** |
| ENSG00000287747.2 | **ENSG0000(** | 12490187‹ | lncRNA | **23.520** | 3.039 | 0.915 | **8.220** | 2.17E-10 | **1.88E-09** |
| ENSG00000058262.13 | **SEC61A1** | 29927 | protein_coding | **3062.054** | -0.919 | -0.277 | **0.529** | 2.19E-10 | **1.90E-09** |
| ENSG00000181472.7 | **ZBTB2** | 57621 | protein_coding | **257.933** | -1.004 | -0.302 | **0.499** | 2.20E-10 | **1.91E-09** |

| | | | | | | | | | |
|---|---|---|---|---|---|---|---|---|---|
| ENSG00000232053.8 | **ENSG0000(** | 105375523 | lncRNA | **48.031** | 1.816 | 0.547 | **3.521** | 2.23E-10 | **1.93E-09** |
| ENSG00000172508.13 | **CARNS1** | 57571 | protein_coding | **69.593** | 2.104 | 0.633 | **4.298** | 2.24E-10 | **1.94E-09** |
| ENSG00000054598.9 | **FOXC1** | 2296 | protein_coding | **24.849** | 2.514 | 0.757 | **5.712** | 2.26E-10 | **1.96E-09** |
| ENSG00000221883.6 | **ARIH2OS** | 646450 | lncRNA | **53.915** | 1.520 | 0.457 | **2.867** | 2.26E-10 | **1.96E-09** |
| ENSG00000100427.18 | **MLC1** | 23209 | protein_coding | **23.451** | -2.304 | -0.693 | **0.203** | 2.27E-10 | **1.96E-09** |
| ENSG00000164404.10 | **GDF9** | 2661 | protein_coding | **47.609** | 1.898 | 0.571 | **3.727** | 2.28E-10 | **1.97E-09** |
| ENSG00000121390.21 | **PSPC1** | 55269 | protein_coding | **1033.647** | -0.525 | -0.158 | **0.695** | 2.29E-10 | **1.98E-09** |
| ENSG00000101444.14 | **AHCY** | 191 | protein_coding | **2554.040** | -0.582 | -0.175 | **0.668** | 2.29E-10 | **1.98E-09** |
| ENSG00000167965.20 | **MLST8** | 64223 | protein_coding | **613.320** | -0.618 | -0.186 | **0.652** | 2.31E-10 | **1.99E-09** |
| ENSG00000065621.17 | **GSTO2** | 119391 | protein_coding | **34.078** | 1.725 | 0.519 | **3.306** | 2.31E-10 | **1.99E-09** |
| ENSG00000143119.17 | **CD53** | 963 | protein_coding | **6107.683** | -0.999 | -0.301 | **0.500** | 2.36E-10 | **2.03E-09** |
| ENSG00000204634.15 | **TBC1D8** | 11138 | protein_coding | **153.186** | 1.362 | 0.410 | **2.570** | 2.40E-10 | **2.07E-09** |
| ENSG00000261884.2 | **ENSG00000261884** | | protein_coding | **106.271** | -1.449 | -0.436 | **0.366** | 2.44E-10 | **2.10E-09** |
| ENSG00000273355.3 | **ENSG00000273355** | | lncRNA | **33.206** | 1.919 | 0.578 | **3.781** | 2.45E-10 | **2.11E-09** |
| ENSG00000213626.14 | **LBH** | 81606 | protein_coding | **873.564** | 1.041 | 0.313 | **2.057** | 2.46E-10 | **2.12E-09** |
| ENSG00000293465.2 | **ENSG00000293465** | | lncRNA | **351.437** | 0.856 | 0.258 | **1.810** | 2.54E-10 | **2.19E-09** |
| ENSG00000186020.15 | **ZNF529** | 57711 | protein_coding | **317.984** | -0.686 | -0.207 | **0.622** | 2.55E-10 | **2.19E-09** |
| ENSG00000006757.14 | **PNPLA4** | 8228 | protein_coding | **329.163** | 0.800 | 0.241 | **1.742** | 2.60E-10 | **2.24E-09** |
| ENSG00000246731.4 | **MGC1627!** | 85001 | lncRNA | **16.340** | 2.830 | 0.852 | **7.108** | 2.62E-10 | **2.25E-09** |
| ENSG00000015285.14 | **WAS** | 7454 | protein_coding | **1819.446** | -0.974 | -0.293 | **0.509** | 2.62E-10 | **2.25E-09** |
| ENSG00000180773.17 | **SLC36A4** | 120103 | protein_coding | **82.382** | -1.359 | -0.409 | **0.390** | 2.64E-10 | **2.26E-09** |
| ENSG00000197912.18 | **SPG7** | 6687 | protein_coding | **3538.970** | 0.757 | 0.228 | **1.691** | 2.64E-10 | **2.26E-09** |
| ENSG00000118997.15 | **DNAH7** | 56171 | protein_coding | **30.876** | 3.873 | 1.166 | **14.653** | 2.65E-10 | **2.27E-09** |
| ENSG00000162775.18 | **RBM15** | 64783 | protein_coding | **214.149** | -1.029 | -0.310 | **0.490** | 2.66E-10 | **2.28E-09** |
| ENSG00000290948.2 | **ENSG00000290948** | | lncRNA | **879.126** | 0.967 | 0.291 | **1.955** | 2.67E-10 | **2.29E-09** |
| ENSG00000266389.2 | **PIK3R5-DT** | 101928235 | lncRNA | **95.969** | 1.244 | 0.374 | **2.368** | 2.67E-10 | **2.29E-09** |
| ENSG00000161513.14 | **FDXR** | 2232 | protein_coding | **926.255** | -0.688 | -0.207 | **0.621** | 2.69E-10 | **2.31E-09** |
| ENSG00000280057.1 | **ENSG00000280057** | | TEC | **19.085** | 2.578 | 0.776 | **5.973** | 2.70E-10 | **2.31E-09** |
| ENSG00000088002.13 | **SULT2B1** | 6820 | protein_coding | **11.174** | 4.080 | 1.228 | **16.911** | 2.72E-10 | **2.33E-09** |
| ENSG00000161654.12 | **LSM12** | 124801 | protein_coding | **1070.188** | -0.481 | -0.145 | **0.716** | 2.81E-10 | **2.40E-09** |
| ENSG00000104219.15 | **ZDHHC2** | 51201 | protein_coding | **126.968** | -1.091 | -0.329 | **0.469** | 2.84E-10 | **2.43E-09** |

| | | | | | | | | | |
|---|---|---|---|---|---|---|---|---|---|
| ENSG00000184860.12 | **SDR42E1** | 93517 | protein_coding | **91.888** | -1.204 | -0.362 | **0.434** | 2.86E-10 | **2.44E-09** |
| ENSG00000204922.7 | **UQCC3** | 790955 | protein_coding | **266.522** | -0.944 | -0.284 | **0.520** | 2.89E-10 | **2.47E-09** |
| ENSG00000290577.2 | **ENSG00000290577** | | lncRNA | **25.355** | 3.222 | 0.970 | **9.330** | 2.93E-10 | **2.50E-09** |
| ENSG00000161547.19 | **SRSF2** | 6427 | protein_coding | **4350.919** | -0.781 | -0.235 | **0.582** | 2.93E-10 | **2.50E-09** |
| ENSG00000132819.18 | **RBM38** | 55544 | protein_coding | **3080.946** | 0.756 | 0.228 | **1.689** | 2.98E-10 | **2.54E-09** |
| ENSG00000136891.16 | **TEX10** | 54881 | protein_coding | **433.227** | -0.703 | -0.212 | **0.614** | 3.03E-10 | **2.58E-09** |
| ENSG00000107951.17 | **MTPAP** | 55149 | protein_coding | **314.446** | -0.775 | -0.233 | **0.584** | 3.04E-10 | **2.59E-09** |
| ENSG00000165886.7 | **UBTD1** | 80019 | protein_coding | **58.761** | 1.547 | 0.466 | **2.923** | 3.12E-10 | **2.66E-09** |
| ENSG00000235480.2 | **ENSG0000(** | 10537349( | lncRNA | **28.130** | 2.165 | 0.652 | **4.485** | 3.12E-10 | **2.66E-09** |
| ENSG00000110013.15 | **SIAE** | 54414 | protein_coding | **258.455** | 0.988 | 0.297 | **1.984** | 3.13E-10 | **2.66E-09** |
| ENSG00000104341.19 | **LAPTM4B** | 55353 | protein_coding | **313.169** | 1.038 | 0.312 | **2.053** | 3.14E-10 | **2.67E-09** |
| ENSG00000273124.2 | **ENSG0000(** | 12490247( | lncRNA | **32.482** | 2.503 | 0.754 | **5.670** | 3.19E-10 | **2.71E-09** |
| ENSG00000265690.7 | **ENSG00000265690** | | protein_coding | **107.298** | 1.245 | 0.375 | **2.370** | 3.20E-10 | **2.72E-09** |
| ENSG00000276231.7 | **PIK3R6** | 146850 | protein_coding | **4441.247** | -1.042 | -0.314 | **0.486** | 3.21E-10 | **2.72E-09** |
| ENSG00000175216.17 | **CKAP5** | 9793 | protein_coding | **1516.495** | -0.687 | -0.207 | **0.621** | 3.25E-10 | **2.76E-09** |
| ENSG00000163132.8 | **MSX1** | 4487 | protein_coding | **193.498** | 1.152 | 0.347 | **2.222** | 3.25E-10 | **2.76E-09** |
| ENSG00000243056.3 | **EIF4EBP3** | 8637 | protein_coding | **240.729** | -1.408 | -0.424 | **0.377** | 3.28E-10 | **2.78E-09** |
| ENSG00000295746.1 | **ENSG00000295746** | | lncRNA | **23.221** | 2.875 | 0.865 | **7.334** | 3.28E-10 | **2.78E-09** |
| ENSG00000120253.16 | **NUP43** | 348995 | protein_coding | **591.045** | -0.705 | -0.212 | **0.613** | 3.34E-10 | **2.83E-09** |
| ENSG00000181045.18 | **SLC26A11** | 284129 | protein_coding | **205.482** | 1.344 | 0.405 | **2.538** | 3.34E-10 | **2.83E-09** |
| ENSG00000170473.19 | **PYM1** | 84305 | protein_coding | **585.662** | -0.570 | -0.171 | **0.674** | 3.36E-10 | **2.84E-09** |
| ENSG00000154096.16 | **THY1** | 7070 | protein_coding | **41.671** | 2.954 | 0.889 | **7.752** | 3.39E-10 | **2.87E-09** |
| ENSG00000287595.4 | **ENSG00000287595** | | lncRNA | **11.992** | 4.186 | 1.260 | **18.204** | 3.39E-10 | **2.87E-09** |
| ENSG00000126453.13 | **BCL2L12** | 83596 | protein_coding | **508.490** | -1.189 | -0.358 | **0.439** | 3.41E-10 | **2.88E-09** |
| ENSG00000140678.19 | **ITGAX** | 3687 | protein_coding | **904.512** | 1.455 | 0.438 | **2.741** | 3.43E-10 | **2.90E-09** |
| ENSG00000135587.11 | **SMPD2** | 6610 | protein_coding | **221.917** | -0.858 | -0.258 | **0.552** | 3.46E-10 | **2.93E-09** |
| ENSG00000100522.12 | **GNPNAT1** | 64841 | protein_coding | **277.047** | -0.931 | -0.280 | **0.525** | 3.56E-10 | **3.01E-09** |
| ENSG00000164022.20 | **AIMP1** | 9255 | protein_coding | **1159.139** | -0.454 | -0.137 | **0.730** | 3.60E-10 | **3.04E-09** |
| ENSG00000176171.14 | **BNIP3** | 664 | protein_coding | **1969.901** | -0.794 | -0.239 | **0.577** | 3.61E-10 | **3.05E-09** |
| ENSG00000305656.1 | **ENSG00000305656** | | lncRNA | **43.554** | 1.570 | 0.473 | **2.969** | 3.65E-10 | **3.08E-09** |
| ENSG00000285578.2 | **LIRIL2R** | | lncRNA | **370.622** | 3.617 | 1.089 | **12.272** | 3.65E-10 | **3.08E-09** |

| | | | | | | | | | |
|---|---|---|---|---|---|---|---|---|---|
| ENSG00000105664.13 | **COMP** | 1311 | protein_coding | **18.890** | 4.336 | 1.305 | **20.203** | 3.67E-10 | **3.10E-09** |
| ENSG00000162377.8 | **COA7** | 65260 | protein_coding | **213.658** | -1.273 | -0.383 | **0.414** | 3.69E-10 | **3.11E-09** |
| ENSG00000101161.10 | **PRPF6** | 24148 | protein_coding | **3135.899** | 0.578 | 0.174 | **1.493** | 3.70E-10 | **3.11E-09** |
| ENSG00000196776.19 | **CD47** | 961 | protein_coding | **1627.760** | -0.662 | -0.199 | **0.632** | 3.70E-10 | **3.11E-09** |
| ENSG00000255521.4 | **CD44-DT** | 10050714 | lncRNA | **33.364** | 2.275 | 0.685 | **4.840** | 3.71E-10 | **3.12E-09** |
| ENSG00000060140.11 | **STYK1** | 55359 | protein_coding | **13.010** | 4.291 | 1.292 | **19.570** | 3.73E-10 | **3.13E-09** |
| ENSG00000126262.5 | **FFAR2** | 2867 | protein_coding | **68.721** | 2.381 | 0.717 | **5.210** | 3.77E-10 | **3.17E-09** |
| ENSG00000163938.19 | **GNL3** | 26354 | protein_coding | **2434.116** | -0.530 | -0.160 | **0.693** | 3.82E-10 | **3.21E-09** |
| ENSG00000173821.21 | **RNF213** | 57674 | protein_coding | **6077.412** | 0.810 | 0.244 | **1.754** | 3.85E-10 | **3.23E-09** |
| ENSG00000104826.15 | **LHB** | 3972 | protein_coding | **30.500** | 3.633 | 1.094 | **12.406** | 3.89E-10 | **3.27E-09** |
| ENSG00000168781.26 | **PPIP5K1** | 9677 | protein_coding | **617.443** | 0.686 | 0.206 | **1.609** | 3.91E-10 | **3.28E-09** |
| ENSG00000145214.16 | **DGKQ** | 1609 | protein_coding | **351.072** | 0.834 | 0.251 | **1.783** | 3.94E-10 | **3.31E-09** |
| ENSG00000119771.17 | **KLHL29** | 114818 | protein_coding | **393.499** | 0.945 | 0.284 | **1.925** | 3.96E-10 | **3.32E-09** |
| ENSG00000173442.15 | **EHBP1L1** | 254102 | protein_coding | **1792.799** | 0.710 | 0.214 | **1.636** | 4.02E-10 | **3.37E-09** |
| ENSG00000110888.21 | **CAPRIN2** | 65981 | protein_coding | **528.206** | 0.868 | 0.261 | **1.826** | 4.12E-10 | **3.45E-09** |
| ENSG00000107829.17 | **FBXW4** | 6468 | protein_coding | **687.066** | -0.789 | -0.238 | **0.579** | 4.19E-10 | **3.51E-09** |
| ENSG00000026652.17 | **AGPAT4** | 56895 | protein_coding | **251.707** | 0.872 | 0.262 | **1.830** | 4.29E-10 | **3.59E-09** |
| ENSG00000076513.19 | **ANKRD13A** | 88455 | protein_coding | **2163.837** | 0.744 | 0.224 | **1.674** | 4.39E-10 | **3.67E-09** |
| ENSG00000265241.10 | **RBM8A** | 9939 | protein_coding | **1558.813** | -0.813 | -0.245 | **0.569** | 4.39E-10 | **3.67E-09** |
| ENSG00000134900.15 | **TPP2** | 7174 | protein_coding | **1231.846** | -0.648 | -0.195 | **0.638** | 4.43E-10 | **3.70E-09** |
| ENSG00000035681.12 | **NSMAF** | 8439 | protein_coding | **531.414** | -0.922 | -0.278 | **0.528** | 4.44E-10 | **3.71E-09** |
| ENSG00000149084.15 | **HSD17B12** | 51144 | protein_coding | **594.381** | -0.847 | -0.255 | **0.556** | 4.50E-10 | **3.76E-09** |
| ENSG00000121900.21 | **TMEM54** | 113452 | protein_coding | **26.749** | 2.886 | 0.869 | **7.391** | 4.66E-10 | **3.89E-09** |
| ENSG00000129646.17 | **QRICH2** | 84074 | protein_coding | **53.424** | 1.400 | 0.421 | **2.639** | 4.76E-10 | **3.97E-09** |
| ENSG00000105401.12 | **CDC37** | 11140 | protein_coding | **6534.791** | 0.694 | 0.209 | **1.617** | 4.76E-10 | **3.97E-09** |
| ENSG00000271888.1 | **JARID2-DT** | | lncRNA | **32.452** | 1.831 | 0.551 | **3.558** | 4.77E-10 | **3.97E-09** |
| ENSG00000189367.17 | **KIAA0408** | 9729 | protein_coding | **52.928** | 1.846 | 0.556 | **3.595** | 4.81E-10 | **4.00E-09** |
| ENSG00000173141.7 | **MRPL57** | 78988 | protein_coding | **832.402** | -0.847 | -0.255 | **0.556** | 4.81E-10 | **4.00E-09** |
| ENSG00000113328.23 | **CCNG1** | 900 | protein_coding | **1241.017** | -0.884 | -0.266 | **0.542** | 4.82E-10 | **4.01E-09** |
| ENSG00000189266.15 | **PNRC2** | 55629 | protein_coding | **1627.391** | -0.694 | -0.209 | **0.618** | 4.83E-10 | **4.01E-09** |
| ENSG00000099203.9 | **TMED1** | 11018 | protein_coding | **172.278** | -0.914 | -0.275 | **0.531** | 4.84E-10 | **4.03E-09** |

| | | | | | | | | | |
|---|---|---|---|---|---|---|---|---|---|
| ENSG00000215421.12 | **ZNF407** | 55628 | protein_coding | **194.425** | 1.033 | 0.311 | **2.046** | 4.88E-10 | **4.06E-09** |
| ENSG00000175643.11 | **RMI2** | 116028 | protein_coding | **342.052** | -0.992 | -0.299 | **0.503** | 4.89E-10 | **4.06E-09** |
| ENSG00000162664.19 | **ZNF326** | 284695 | protein_coding | **330.002** | -1.001 | -0.301 | **0.500** | 4.90E-10 | **4.07E-09** |
| ENSG00000096063.18 | **SRPK1** | 6732 | protein_coding | **899.727** | -0.583 | -0.176 | **0.667** | 4.94E-10 | **4.10E-09** |
| ENSG00000171121.18 | **KCNMB3** | 27094 | protein_coding | **135.482** | 1.226 | 0.369 | **2.339** | 5.06E-10 | **4.20E-09** |
| ENSG00000272825.1 | **ENSG00000272825** | | lncRNA | **82.686** | -1.514 | -0.456 | **0.350** | 5.09E-10 | **4.22E-09** |
| ENSG00000175155.12 | **YPEL2** | 388403 | protein_coding | **149.112** | 1.177 | 0.354 | **2.262** | 5.11E-10 | **4.23E-09** |
| ENSG00000161960.18 | **EIF4A1** | 1973 | protein_coding | **10610.561** | -0.683 | -0.206 | **0.623** | 5.16E-10 | **4.27E-09** |
| ENSG00000053747.18 | **LAMA3** | 3909 | protein_coding | **92.094** | 1.335 | 0.402 | **2.522** | 5.19E-10 | **4.30E-09** |
| ENSG00000141295.16 | **SCRN2** | 90507 | protein_coding | **804.673** | 0.641 | 0.193 | **1.560** | 5.20E-10 | **4.31E-09** |
| ENSG00000172426.18 | **RSPH9** | 221421 | protein_coding | **14.391** | 3.471 | 1.045 | **11.092** | 5.33E-10 | **4.41E-09** |
| ENSG00000117322.21 | **CR2** | 1380 | protein_coding | **1158.070** | 1.040 | 0.313 | **2.056** | 5.34E-10 | **4.42E-09** |
| ENSG00000214548.20 | **MEG3** | 55384 | lncRNA | **47.922** | 2.569 | 0.773 | **5.936** | 5.42E-10 | **4.48E-09** |
| ENSG00000169519.23 | **METTL15** | 196074 | protein_coding | **202.420** | -0.805 | -0.242 | **0.572** | 5.43E-10 | **4.49E-09** |
| ENSG00000184708.20 | **EIF4ENIF1** | 56478 | protein_coding | **819.602** | 0.572 | 0.172 | **1.487** | 5.49E-10 | **4.54E-09** |
| ENSG00000116771.8 | **AGMAT** | 79814 | protein_coding | **53.912** | -1.447 | -0.436 | **0.367** | 5.50E-10 | **4.54E-09** |
| ENSG00000113569.19 | **NUP155** | 9631 | protein_coding | **320.229** | -1.000 | -0.301 | **0.500** | 5.52E-10 | **4.56E-09** |
| ENSG00000198324.16 | **PHETA1** | 144717 | protein_coding | **66.877** | 1.348 | 0.406 | **2.545** | 5.68E-10 | **4.69E-09** |
| ENSG00000184922.17 | **FMNL1** | 752 | protein_coding | **2245.806** | -0.587 | -0.177 | **0.666** | 5.69E-10 | **4.70E-09** |
| ENSG00000111271.17 | **ACAD10** | 80724 | protein_coding | **874.501** | 0.611 | 0.184 | **1.528** | 5.71E-10 | **4.70E-09** |
| ENSG00000244682.8 | **FCGR2C** | | protein_coding | **265.007** | 1.368 | 0.412 | **2.581** | 5.71E-10 | **4.70E-09** |
| ENSG00000285410.1 | **GABPB1-IT** | 55056 | lncRNA | **331.910** | 0.851 | 0.256 | **1.804** | 5.85E-10 | **4.82E-09** |
| ENSG00000081059.22 | **TCF7** | 6932 | protein_coding | **1626.761** | 1.142 | 0.344 | **2.207** | 5.94E-10 | **4.89E-09** |
| ENSG00000110002.18 | **VWA5A** | 4013 | protein_coding | **66.926** | 1.966 | 0.592 | **3.907** | 6.01E-10 | **4.95E-09** |
| ENSG00000135093.15 | **USP30** | 84749 | protein_coding | **241.257** | 0.928 | 0.279 | **1.903** | 6.07E-10 | **4.99E-09** |
| ENSG00000156469.11 | **MTERF3** | 51001 | protein_coding | **361.969** | -0.652 | -0.196 | **0.636** | 6.08E-10 | **5.00E-09** |
| ENSG00000138606.21 | **SHF** | 90525 | protein_coding | **191.461** | 1.210 | 0.364 | **2.313** | 6.18E-10 | **5.08E-09** |
| ENSG00000156876.11 | **SASS6** | 163786 | protein_coding | **183.079** | -0.961 | -0.289 | **0.514** | 6.21E-10 | **5.10E-09** |
| ENSG00000167799.12 | **NUDT8** | 254552 | protein_coding | **153.414** | -0.963 | -0.290 | **0.513** | 6.27E-10 | **5.15E-09** |
| ENSG00000172534.16 | **HCFC1** | 3054 | protein_coding | **1107.353** | -0.683 | -0.206 | **0.623** | 6.29E-10 | **5.17E-09** |
| ENSG00000142949.20 | **PTPRF** | 5792 | protein_coding | **120.216** | 1.148 | 0.346 | **2.216** | 6.33E-10 | **5.19E-09** |

| | | | | | | | | | |
|---|---|---|---|---|---|---|---|---|---|
| ENSG00000113387.15 | **SUB1** | 10923 | protein_coding | **3434.747** | -0.769 | -0.232 | **0.587** | 6.41E-10 | **5.26E-09** |
| ENSG00000165879.9 | **FRAT1** | 10023 | protein_coding | **47.542** | 1.444 | 0.435 | **2.721** | 6.42E-10 | **5.26E-09** |
| ENSG00000293621.2 | **UVRAG-DT** | | lncRNA | **20.454** | 2.813 | 0.847 | **7.028** | 6.96E-10 | **5.70E-09** |
| ENSG00000290660.2 | **ENSG00000290660** | | lncRNA | **12.686** | 4.079 | 1.228 | **16.895** | 6.99E-10 | **5.72E-09** |
| ENSG00000129244.11 | **ATP1B2** | 482 | protein_coding | **78.487** | 3.836 | 1.155 | **14.280** | 7.11E-10 | **5.82E-09** |
| ENSG00000092607.16 | **TBX15** | 6913 | protein_coding | **106.176** | -1.452 | -0.437 | **0.366** | 7.19E-10 | **5.89E-09** |
| ENSG00000067606.19 | **PRKCZ** | 5590 | protein_coding | **79.286** | 1.708 | 0.514 | **3.266** | 7.31E-10 | **5.98E-09** |
| ENSG00000115902.13 | **SLC1A4** | 6509 | protein_coding | **1710.615** | -0.699 | -0.210 | **0.616** | 7.37E-10 | **6.02E-09** |
| ENSG00000221955.13 | **SLC12A8** | 84561 | protein_coding | **2858.553** | 0.713 | 0.215 | **1.639** | 7.47E-10 | **6.10E-09** |
| ENSG00000081307.17 | **UBA5** | 79876 | protein_coding | **560.264** | -0.791 | -0.238 | **0.578** | 7.54E-10 | **6.16E-09** |
| ENSG00000139200.16 | **PIANP** | 196500 | protein_coding | **16.694** | 4.538 | 1.366 | **23.230** | 7.59E-10 | **6.20E-09** |
| ENSG00000299639.1 | **ENSG00000299639** | | lncRNA | **56.809** | 1.722 | 0.518 | **3.298** | 7.61E-10 | **6.21E-09** |
| ENSG00000053501.16 | **USE1** | 55850 | protein_coding | **392.746** | -0.657 | -0.198 | **0.634** | 7.63E-10 | **6.22E-09** |
| ENSG00000228789.11 | **HCG22** | 285834 | lncRNA | **70.549** | 1.688 | 0.508 | **3.223** | 7.64E-10 | **6.23E-09** |
| ENSG00000186575.22 | **NF2** | 4771 | protein_coding | **494.244** | -0.585 | -0.176 | **0.667** | 7.66E-10 | **6.24E-09** |
| ENSG00000167088.13 | **SNRPD1** | 6632 | protein_coding | **1923.646** | -0.774 | -0.233 | **0.585** | 7.67E-10 | **6.25E-09** |
| ENSG00000234215.4 | **CT66** | | lncRNA | **14.515** | 3.040 | 0.915 | **8.227** | 7.78E-10 | **6.33E-09** |
| ENSG00000143889.17 | **HNRNPLL** | 92906 | protein_coding | **112.744** | 1.421 | 0.428 | **2.678** | 7.82E-10 | **6.37E-09** |
| ENSG00000178860.11 | **MSC** | 9242 | protein_coding | **2689.019** | -0.742 | -0.223 | **0.598** | 7.90E-10 | **6.43E-09** |
| ENSG00000310471.1 | **ENSG00000310471** | | lncRNA | **469.102** | 1.312 | 0.395 | **2.483** | 7.91E-10 | **6.44E-09** |
| ENSG00000133773.14 | **CCDC59** | 29080 | protein_coding | **419.481** | -0.859 | -0.259 | **0.551** | 7.95E-10 | **6.47E-09** |
| ENSG00000088854.15 | **DNAAF9** | 25943 | protein_coding | **74.381** | 1.583 | 0.476 | **2.996** | 7.99E-10 | **6.49E-09** |
| ENSG00000183621.17 | **ZNF438** | 220929 | protein_coding | **137.432** | 0.858 | 0.258 | **1.813** | 8.02E-10 | **6.51E-09** |
| ENSG00000103540.19 | **CCP110** | 9738 | protein_coding | **160.409** | -1.131 | -0.341 | **0.456** | 8.05E-10 | **6.53E-09** |
| ENSG00000076258.12 | **FMO4** | 2329 | protein_coding | **145.200** | 1.157 | 0.348 | **2.230** | 8.09E-10 | **6.56E-09** |
| ENSG00000260534.1 | **ENSG00000260534** | | lncRNA | **37.791** | 1.734 | 0.522 | **3.328** | 8.09E-10 | **6.57E-09** |
| ENSG00000242574.11 | **HLA-DMB** | 3109 | protein_coding | **3157.385** | 0.856 | 0.258 | **1.810** | 8.12E-10 | **6.59E-09** |
| ENSG00000175029.19 | **CTBP2** | 1488 | protein_coding | **45.322** | 2.332 | 0.702 | **5.035** | 8.17E-10 | **6.62E-09** |
| ENSG00000167634.13 | **NLRP7** | 199713 | protein_coding | **45.697** | 1.872 | 0.563 | **3.660** | 8.29E-10 | **6.72E-09** |
| ENSG00000115183.17 | **TANC1** | 85461 | protein_coding | **209.278** | 1.009 | 0.304 | **2.012** | 8.49E-10 | **6.88E-09** |
| ENSG00000126259.22 | **KIRREL2** | 84063 | protein_coding | **21.948** | 3.123 | 0.940 | **8.711** | 8.63E-10 | **6.98E-09** |

| | | | | | | | | | |
|---|---|---|---|---|---|---|---|---|---|
| ENSG00000135766.11 | **EGLN1** | 54583 | protein_coding | **321.287** | -1.061 | -0.319 | **0.479** | 8.63E-10 | **6.98E-09** |
| ENSG00000283036.4 | **LINC01988** | 10050631[illegible] | lncRNA | **9.311** | 5.044 | 1.518 | **32.989** | 8.70E-10 | **7.04E-09** |
| ENSG00000133835.20 | **HSD17B4** | 3295 | protein_coding | **1115.949** | 0.632 | 0.190 | **1.550** | 8.71E-10 | **7.05E-09** |
| ENSG00000111639.10 | **MRPL51** | 51258 | protein_coding | **1698.082** | -0.682 | -0.205 | **0.623** | 8.81E-10 | **7.12E-09** |
| ENSG00000132522.18 | **GPS2** | 2874 | protein_coding | **1399.952** | 0.722 | 0.217 | **1.650** | 8.89E-10 | **7.18E-09** |
| ENSG00000106689.12 | **LHX2** | 9355 | protein_coding | **50.686** | 1.501 | 0.452 | **2.831** | 8.93E-10 | **7.21E-09** |
| ENSG00000092200.13 | **RPGRIP1** | 57096 | protein_coding | **11.494** | 4.426 | 1.332 | **21.493** | 8.96E-10 | **7.23E-09** |
| ENSG00000073060.19 | **SCARB1** | 949 | protein_coding | **3306.851** | -0.756 | -0.228 | **0.592** | 9.06E-10 | **7.31E-09** |
| ENSG00000290319.2 | **ENSG00000290319** | | lncRNA | **8.538** | 5.575 | 1.678 | **47.667** | 9.14E-10 | **7.37E-09** |
| ENSG00000203326.14 | **ZNF525** | 170958 | protein_coding | **230.845** | -0.737 | -0.222 | **0.600** | 9.15E-10 | **7.38E-09** |
| ENSG00000073350.16 | **LLGL2** | 3993 | protein_coding | **219.733** | 1.807 | 0.544 | **3.498** | 9.23E-10 | **7.44E-09** |
| ENSG00000166394.17 | **CYB5R2** | 51700 | protein_coding | **261.037** | 0.954 | 0.287 | **1.938** | 9.23E-10 | **7.44E-09** |
| ENSG00000108312.18 | **UBTF** | 7343 | protein_coding | **1533.259** | -0.560 | -0.168 | **0.678** | 9.24E-10 | **7.44E-09** |
| ENSG00000168724.20 | **DNAJC21** | 134218 | protein_coding | **627.669** | -0.682 | -0.205 | **0.623** | 9.28E-10 | **7.47E-09** |
| ENSG00000145715.17 | **RASA1** | 5921 | protein_coding | **688.430** | 0.862 | 0.259 | **1.818** | 9.30E-10 | **7.48E-09** |
| ENSG00000236778.10 | **INTS6-AS1** | | lncRNA | **135.191** | 1.003 | 0.302 | **2.004** | 9.33E-10 | **7.51E-09** |
| ENSG00000198870.9 | **STKLD1** | 169436 | protein_coding | **22.581** | 2.467 | 0.742 | **5.527** | 9.37E-10 | **7.53E-09** |
| ENSG00000233593.13 | **LINC02609** | 10537885[illegible] | lncRNA | **13.966** | 3.431 | 1.033 | **10.785** | 9.42E-10 | **7.57E-09** |
| ENSG00000136950.15 | **ARPC5L** | 81873 | protein_coding | **1525.559** | -0.786 | -0.237 | **0.580** | 9.43E-10 | **7.57E-09** |
| ENSG00000176371.16 | **ZSCAN2** | 54993 | protein_coding | **146.534** | 1.022 | 0.308 | **2.031** | 9.44E-10 | **7.58E-09** |
| ENSG00000006283.18 | **CACNA1G** | 8913 | protein_coding | **22.808** | 3.111 | 0.937 | **8.641** | 9.54E-10 | **7.65E-09** |
| ENSG00000275066.7 | **SYNRG** | 11276 | protein_coding | **778.129** | -0.790 | -0.238 | **0.578** | 9.63E-10 | **7.73E-09** |
| ENSG00000187867.12 | **PALM3** | 342979 | protein_coding | **9.717** | 4.994 | 1.503 | **31.867** | 9.84E-10 | **7.89E-09** |
| ENSG00000080824.21 | **HSP90AA1** | 3320 | protein_coding | **6285.303** | -0.955 | -0.287 | **0.516** | 9.88E-10 | **7.92E-09** |
| ENSG00000177989.16 | **CIMAP1B** | 440836 | protein_coding | **1076.994** | -0.844 | -0.254 | **0.557** | 9.93E-10 | **7.96E-09** |
| ENSG00000221963.8 | **APOL6** | 80830 | protein_coding | **338.840** | -1.352 | -0.407 | **0.392** | 9.98E-10 | **7.99E-09** |
| ENSG00000301234.1 | **ENSG00000301234** | | lncRNA | **45.006** | 1.525 | 0.459 | **2.879** | 1.00E-09 | **8.02E-09** |
| ENSG00000104524.16 | **PYCR3** | 65263 | protein_coding | **186.392** | -0.986 | -0.297 | **0.505** | 1.00E-09 | **8.04E-09** |
| ENSG00000273162.2 | **ENSG0000[illegible]** | 10798426[illegible] | lncRNA | **42.541** | -1.864 | -0.561 | **0.275** | 1.02E-09 | **8.13E-09** |
| ENSG00000245146.8 | **MALINC1** | 12490019[illegible] | lncRNA | **39.642** | 1.739 | 0.523 | **3.337** | 1.02E-09 | **8.16E-09** |
| ENSG00000114861.26 | **FOXP1** | 27086 | protein_coding | **72.809** | 1.412 | 0.425 | **2.661** | 1.02E-09 | **8.17E-09** |

| | | | | | | | | | |
|---|---|---|---|---|---|---|---|---|---|
| ENSG00000140009.20 | **ESR2** | 2100 | protein_coding | **99.099** | -1.195 | -0.360 | **0.437** | 1.04E-09 | **8.33E-09** |
| ENSG00000075884.16 | **ARHGAP15** | 55843 | protein_coding | **749.054** | -0.645 | -0.194 | **0.640** | 1.04E-09 | **8.34E-09** |
| ENSG00000215252.14 | **GOLGA8B** | 440270 | protein_coding | **5668.650** | 0.740 | 0.223 | **1.671** | 1.05E-09 | **8.37E-09** |
| ENSG00000157734.16 | **SNX22** | 79856 | protein_coding | **668.416** | -0.814 | -0.245 | **0.569** | 1.05E-09 | **8.38E-09** |
| ENSG00000242028.10 | **HYPK** | 25764 | protein_coding | **258.319** | -1.220 | -0.367 | **0.429** | 1.05E-09 | **8.40E-09** |
| ENSG00000227908.6 | **IL6ST-DT** | 441072 | lncRNA | **34.618** | 2.152 | 0.648 | **4.445** | 1.06E-09 | **8.42E-09** |
| ENSG00000081154.13 | **PCNP** | 57092 | protein_coding | **1010.362** | -0.680 | -0.205 | **0.624** | 1.06E-09 | **8.42E-09** |
| ENSG00000272888.9 | **CHASERR** | | lncRNA | **609.848** | -0.953 | -0.287 | **0.516** | 1.06E-09 | **8.42E-09** |
| ENSG00000180354.18 | **MTURN** | 222166 | protein_coding | **117.732** | 1.249 | 0.376 | **2.377** | 1.06E-09 | **8.43E-09** |
| ENSG00000116793.18 | **PHTF1** | 10745 | protein_coding | **463.980** | -0.667 | -0.201 | **0.630** | 1.07E-09 | **8.53E-09** |
| ENSG00000184897.7 | **H1-10** | 8971 | protein_coding | **3126.291** | 1.155 | 0.348 | **2.226** | 1.07E-09 | **8.53E-09** |
| ENSG00000198522.16 | **GPN1** | 11321 | protein_coding | **446.967** | -0.810 | -0.244 | **0.570** | 1.08E-09 | **8.61E-09** |
| ENSG00000272391.8 | **POM121C** | 100101267 | protein_coding | **1075.237** | -0.572 | -0.172 | **0.673** | 1.08E-09 | **8.61E-09** |
| ENSG00000073861.5 | **TBX21** | 30009 | protein_coding | **693.698** | 0.878 | 0.264 | **1.838** | 1.09E-09 | **8.67E-09** |
| ENSG00000290691.1 | **STRCP1** | 554225 | lncRNA | **49.314** | 2.363 | 0.711 | **5.143** | 1.09E-09 | **8.67E-09** |
| ENSG00000141314.15 | **RHBDL3** | 162494 | protein_coding | **20.236** | 2.316 | 0.697 | **4.979** | 1.09E-09 | **8.67E-09** |
| ENSG00000164758.9 | **MED30** | 90390 | protein_coding | **202.700** | -0.884 | -0.266 | **0.542** | 1.09E-09 | **8.67E-09** |
| ENSG00000224165.9 | **DNAJC27-A** | 729723 | lncRNA | **26.636** | 1.990 | 0.599 | **3.971** | 1.11E-09 | **8.78E-09** |
| ENSG00000086717.21 | **PPEF1** | 5475 | protein_coding | **24.017** | 2.735 | 0.823 | **6.657** | 1.14E-09 | **9.03E-09** |
| ENSG00000156787.19 | **TBC1D31** | 93594 | protein_coding | **168.426** | -0.981 | -0.295 | **0.507** | 1.15E-09 | **9.10E-09** |
| ENSG00000204568.14 | **MRPS18B** | 28973 | protein_coding | **1475.843** | -0.475 | -0.143 | **0.720** | 1.15E-09 | **9.12E-09** |
| ENSG00000291174.2 | **FAUP4** | 101927789 | lncRNA | **345.865** | 0.978 | 0.294 | **1.969** | 1.17E-09 | **9.23E-09** |
| ENSG00000231721.9 | **LINC-PINT** | 378805 | lncRNA | **596.509** | 0.801 | 0.241 | **1.743** | 1.17E-09 | **9.24E-09** |
| ENSG00000137812.22 | **KNL1** | 57082 | protein_coding | **218.349** | -1.450 | -0.436 | **0.366** | 1.18E-09 | **9.31E-09** |
| ENSG00000256039.5 | **LINC02446** | 10106003[illegible] | lncRNA | **11.325** | 5.151 | 1.550 | **35.520** | 1.18E-09 | **9.34E-09** |
| ENSG00000186834.4 | **HEXIM1** | 10614 | protein_coding | **1029.893** | 0.863 | 0.260 | **1.818** | 1.19E-09 | **9.37E-09** |
| ENSG00000148019.16 | **CEP78** | 84131 | protein_coding | **291.576** | -0.876 | -0.264 | **0.545** | 1.19E-09 | **9.38E-09** |
| ENSG00000114315.6 | **HES1** | 3280 | protein_coding | **359.555** | 0.961 | 0.289 | **1.946** | 1.21E-09 | **9.57E-09** |
| ENSG00000135838.16 | **NPL** | 80896 | protein_coding | **206.465** | 2.950 | 0.888 | **7.725** | 1.22E-09 | **9.60E-09** |
| ENSG00000270574.1 | **ENSG00000270574** | | lncRNA | **51.456** | 1.330 | 0.400 | **2.513** | 1.22E-09 | **9.61E-09** |
| ENSG00000162613.20 | **FUBP1** | 8880 | protein_coding | **1819.273** | -0.716 | -0.216 | **0.609** | 1.22E-09 | **9.62E-09** |

| | | | | | | | | | |
|---|---|---|---|---|---|---|---|---|---|
| ENSG00000106003.14 | **LFNG** | 3955 | protein_coding | **62.786** | 1.595 | 0.480 | **3.021** | 1.23E-09 | **9.70E-09** |
| ENSG00000173530.8 | **TNFRSF10D** | 8793 | protein_coding | **106.794** | -1.303 | -0.392 | **0.405** | 1.25E-09 | **9.85E-09** |
| ENSG00000198417.9 | **MT1F** | 4494 | protein_coding | **28.448** | 2.936 | 0.884 | **7.651** | 1.25E-09 | **9.85E-09** |
| ENSG00000176896.11 | **TCEANC** | 170082 | protein_coding | **74.197** | 1.246 | 0.375 | **2.371** | 1.26E-09 | **9.89E-09** |
| ENSG00000131899.15 | **LLGL1** | 3996 | protein_coding | **115.420** | 1.289 | 0.388 | **2.444** | 1.26E-09 | **9.91E-09** |
| ENSG00000160678.14 | **S100A1** | 6271 | protein_coding | **13.150** | 4.495 | 1.353 | **22.555** | 1.27E-09 | **9.99E-09** |
| ENSG00000270605.2 | **ATP5IF1-AS1** | | lncRNA | **725.203** | 0.916 | 0.276 | **1.886** | 1.27E-09 | **1.00E-08** |
| ENSG00000128965.14 | **CHAC1** | 79094 | protein_coding | **499.613** | -1.311 | -0.395 | **0.403** | 1.28E-09 | **1.01E-08** |
| ENSG00000217825.3 | **ENSG00000217825** | | lncRNA | **51.246** | 1.971 | 0.593 | **3.922** | 1.30E-09 | **1.02E-08** |
| ENSG00000246705.5 | **H2AJ** | 55766 | protein_coding | **548.617** | 0.723 | 0.218 | **1.651** | 1.31E-09 | **1.03E-08** |
| ENSG00000151929.12 | **BAG3** | 9531 | protein_coding | **43.234** | 1.814 | 0.546 | **3.517** | 1.31E-09 | **1.03E-08** |
| ENSG00000254470.5 | **AP5B1** | 91056 | protein_coding | **437.016** | -0.603 | -0.181 | **0.658** | 1.32E-09 | **1.04E-08** |
| ENSG00000129187.17 | **DCTD** | 1635 | protein_coding | **1227.225** | -0.608 | -0.183 | **0.656** | 1.33E-09 | **1.04E-08** |
| ENSG00000132388.15 | **UBE2G1** | 7326 | protein_coding | **1030.026** | 0.650 | 0.196 | **1.569** | 1.33E-09 | **1.04E-08** |
| ENSG00000227292.2 | **ENSG00000227292** | | lncRNA | **16.842** | 3.476 | 1.046 | **11.127** | 1.34E-09 | **1.05E-08** |
| ENSG00000179051.16 | **RCC2** | 55920 | protein_coding | **3584.813** | -0.565 | -0.170 | **0.676** | 1.34E-09 | **1.05E-08** |
| ENSG00000262877.6 | **ENSG00000262877** | | lncRNA | **30.486** | 2.626 | 0.791 | **6.174** | 1.37E-09 | **1.08E-08** |
| ENSG00000123609.13 | **NMI** | 9111 | protein_coding | **814.409** | -0.887 | -0.267 | **0.541** | 1.38E-09 | **1.08E-08** |
| ENSG00000286069.3 | **ENSG0000(** | 10537825( | lncRNA | **17.367** | -2.802 | -0.843 | **0.143** | 1.38E-09 | **1.08E-08** |
| ENSG00000228521.3 | **ENSG00000228521** | | lncRNA | **37.759** | 1.774 | 0.534 | **3.420** | 1.39E-09 | **1.09E-08** |
| ENSG00000140511.14 | **HAPLN3** | 145864 | protein_coding | **188.742** | 1.200 | 0.361 | **2.297** | 1.41E-09 | **1.10E-08** |
| ENSG00000285091.1 | **ENSG00000285091** | | lncRNA | **17.355** | 2.402 | 0.723 | **5.285** | 1.42E-09 | **1.11E-08** |
| ENSG00000224877.5 | **NDUFAF8** | 284184 | protein_coding | **665.618** | -0.902 | -0.272 | **0.535** | 1.42E-09 | **1.11E-08** |
| ENSG00000161243.12 | **FBXO27** | 126433 | protein_coding | **34.036** | 1.891 | 0.569 | **3.709** | 1.43E-09 | **1.11E-08** |
| ENSG00000112787.16 | **FBRSL1** | 57666 | protein_coding | **1349.248** | 0.687 | 0.207 | **1.609** | 1.43E-09 | **1.12E-08** |
| ENSG00000123992.23 | **DNPEP** | 23549 | protein_coding | **1007.075** | -0.620 | -0.187 | **0.651** | 1.43E-09 | **1.12E-08** |
| ENSG00000136827.13 | **TOR1A** | 1861 | protein_coding | **441.316** | -0.825 | -0.248 | **0.564** | 1.46E-09 | **1.14E-08** |
| ENSG00000120318.18 | **ARAP3** | 64411 | protein_coding | **9.599** | 4.487 | 1.351 | **22.420** | 1.47E-09 | **1.14E-08** |
| ENSG00000184575.14 | **XPOT** | 11260 | protein_coding | **1516.592** | -1.018 | -0.307 | **0.494** | 1.47E-09 | **1.14E-08** |
| ENSG00000136828.21 | **RALGPS1** | 9649 | protein_coding | **172.312** | 1.067 | 0.321 | **2.095** | 1.47E-09 | **1.15E-08** |
| ENSG00000281912.1 | **LINC01144** | 400752 | lncRNA | **46.910** | 1.542 | 0.464 | **2.912** | 1.47E-09 | **1.15E-08** |

| | | | | | | | | | |
|---|---|---|---|---|---|---|---|---|---|
| ENSG00000167984.20 | **NLRC3** | 197358 | protein_coding | **405.889** | -1.117 | -0.336 | **0.461** | 1.48E-09 | **1.15E-08** |
| ENSG00000117280.15 | **RAB29** | 8934 | protein_coding | **434.914** | 0.893 | 0.269 | **1.858** | 1.48E-09 | **1.15E-08** |
| ENSG00000157978.15 | **LDLRAP1** | 26119 | protein_coding | **651.081** | -0.803 | -0.242 | **0.573** | 1.49E-09 | **1.16E-08** |
| ENSG00000176393.13 | **RNPEP** | 6051 | protein_coding | **1183.484** | -0.708 | -0.213 | **0.612** | 1.49E-09 | **1.16E-08** |
| ENSG00000104731.16 | **KLHDC4** | 54758 | protein_coding | **1096.867** | -0.513 | -0.155 | **0.701** | 1.54E-09 | **1.20E-08** |
| ENSG00000133878.11 | **DUSP26** | 78986 | protein_coding | **15.282** | 3.332 | 1.003 | **10.067** | 1.56E-09 | **1.22E-08** |
| ENSG00000166483.14 | **WEE1** | 7465 | protein_coding | **196.002** | -0.964 | -0.290 | **0.512** | 1.59E-09 | **1.23E-08** |
| ENSG00000066336.15 | **SPI1** | 6688 | protein_coding | **432.902** | -1.154 | -0.347 | **0.449** | 1.60E-09 | **1.24E-08** |
| ENSG00000172922.13 | **RNASEH2C** | 84153 | protein_coding | **683.099** | -0.732 | -0.220 | **0.602** | 1.60E-09 | **1.24E-08** |
| ENSG00000279861.1 | **ENSG00000279861** | | TEC | **199.920** | 0.881 | 0.265 | **1.841** | 1.60E-09 | **1.24E-08** |
| ENSG00000079257.9 | **LXN** | 56925 | protein_coding | **110.745** | -0.927 | -0.279 | **0.526** | 1.61E-09 | **1.24E-08** |
| ENSG00000111057.13 | **KRT18** | 3875 | protein_coding | **63.705** | 2.266 | 0.682 | **4.810** | 1.62E-09 | **1.25E-08** |
| ENSG00000036448.12 | **MYOM2** | 9172 | protein_coding | **76.628** | 1.291 | 0.389 | **2.448** | 1.63E-09 | **1.26E-08** |
| ENSG00000111335.15 | **OAS2** | 4939 | protein_coding | **5115.022** | -0.854 | -0.257 | **0.553** | 1.66E-09 | **1.29E-08** |
| ENSG00000106333.15 | **PCOLCE** | 5118 | protein_coding | **129.110** | 1.474 | 0.444 | **2.778** | 1.67E-09 | **1.29E-08** |
| ENSG00000119698.14 | **PPP4R4** | 57718 | protein_coding | **26.624** | 3.117 | 0.938 | **8.675** | 1.68E-09 | **1.30E-08** |
| ENSG00000154917.13 | **RAB6B** | 51560 | protein_coding | **37.881** | 2.853 | 0.859 | **7.227** | 1.69E-09 | **1.31E-08** |
| ENSG00000162869.18 | **PPP1R21** | 129285 | protein_coding | **993.126** | 0.714 | 0.215 | **1.640** | 1.69E-09 | **1.31E-08** |
| ENSG00000197816.18 | **CCDC180** | 10049948: | protein_coding | **40.140** | 3.200 | 0.963 | **9.188** | 1.71E-09 | **1.32E-08** |
| ENSG00000104765.18 | **BNIP3L** | 665 | protein_coding | **1423.556** | -0.864 | -0.260 | **0.549** | 1.74E-09 | **1.34E-08** |
| ENSG00000142751.17 | **GPN2** | 54707 | protein_coding | **617.931** | -0.743 | -0.224 | **0.597** | 1.75E-09 | **1.35E-08** |
| ENSG00000291112.2 | **ENSG00000291112** | | lncRNA | **17.907** | 4.688 | 1.411 | **25.784** | 1.76E-09 | **1.36E-08** |
| ENSG00000169129.18 | **AFAP1L2** | 84632 | protein_coding | **51.839** | 1.853 | 0.558 | **3.614** | 1.77E-09 | **1.37E-08** |
| ENSG00000204531.22 | **POU5F1** | 5460 | protein_coding | **61.432** | 1.323 | 0.398 | **2.502** | 1.81E-09 | **1.40E-08** |
| ENSG00000257923.14 | **CUX1** | 1523 | protein_coding | **369.118** | 0.872 | 0.262 | **1.830** | 1.82E-09 | **1.40E-08** |
| ENSG00000205323.11 | **SARNP** | 84324 | protein_coding | **1344.828** | -0.455 | -0.137 | **0.729** | 1.82E-09 | **1.40E-08** |
| ENSG00000289582.2 | **ENSG00000289582** | | lncRNA | **10.192** | 3.792 | 1.141 | **13.849** | 1.82E-09 | **1.41E-08** |
| ENSG00000169689.17 | **CENPX** | 201254 | protein_coding | **322.779** | -0.910 | -0.274 | **0.532** | 1.85E-09 | **1.42E-08** |
| ENSG00000166681.16 | **BEX3** | 27018 | protein_coding | **248.661** | 1.201 | 0.361 | **2.299** | 1.86E-09 | **1.43E-08** |
| ENSG00000069493.16 | **CLEC2D** | 29121 | protein_coding | **2572.561** | -1.072 | -0.323 | **0.476** | 1.87E-09 | **1.44E-08** |
| ENSG00000160179.22 | **ABCG1** | 9619 | protein_coding | **465.196** | 1.409 | 0.424 | **2.656** | 1.88E-09 | **1.44E-08** |

| | | | | | | | | | |
|---|---|---|---|---|---|---|---|---|---|
| ENSG00000176340.5 | **COX8A** | 1351 | protein_coding | **2598.110** | -0.806 | -0.242 | **0.572** | 1.92E-09 | **1.47E-08** |
| ENSG00000143653.12 | **SCCPDH** | 51097 | protein_coding | **472.643** | -0.846 | -0.255 | **0.556** | 1.93E-09 | **1.48E-08** |
| ENSG00000197372.12 | **ZNF675** | 171392 | protein_coding | **152.599** | -1.130 | -0.340 | **0.457** | 1.93E-09 | **1.49E-08** |
| ENSG00000132382.17 | **MYBBP1A** | 10514 | protein_coding | **1222.463** | -0.618 | -0.186 | **0.651** | 1.96E-09 | **1.50E-08** |
| ENSG00000128585.20 | **MKLN1** | 4289 | protein_coding | **652.241** | -0.925 | -0.278 | **0.527** | 1.98E-09 | **1.52E-08** |
| ENSG00000105855.12 | **ITGB8** | 3696 | protein_coding | **105.642** | 1.165 | 0.351 | **2.242** | 1.99E-09 | **1.53E-08** |
| ENSG00000137177.21 | **KIF13A** | 63971 | protein_coding | **41.414** | 1.650 | 0.497 | **3.139** | 2.03E-09 | **1.55E-08** |
| ENSG00000132357.16 | **CARD6** | 84674 | protein_coding | **43.814** | -1.544 | -0.465 | **0.343** | 2.03E-09 | **1.56E-08** |
| ENSG00000259172.2 | **SNRPA1-D** | 12076613 | lncRNA | **53.695** | 1.322 | 0.398 | **2.499** | 2.05E-09 | **1.57E-08** |
| ENSG00000197894.14 | **ADH5** | 128 | protein_coding | **1081.326** | -0.543 | -0.163 | **0.686** | 2.08E-09 | **1.60E-08** |
| ENSG00000169084.16 | **DHRSX** | 207063 | protein_coding | **252.354** | -0.994 | -0.299 | **0.502** | 2.11E-09 | **1.61E-08** |
| ENSG00000184207.10 | **PGP** | 283871 | protein_coding | **512.615** | -0.754 | -0.227 | **0.593** | 2.13E-09 | **1.63E-08** |
| ENSG00000073150.15 | **PANX2** | 56666 | protein_coding | **131.195** | 1.660 | 0.500 | **3.161** | 2.14E-09 | **1.63E-08** |
| ENSG00000197978.11 | **GOLGA6L9** | 440295 | protein_coding | **571.066** | 0.976 | 0.294 | **1.966** | 2.14E-09 | **1.64E-08** |
| ENSG00000155008.18 | **APOOL** | 139322 | protein_coding | **151.410** | -1.064 | -0.320 | **0.478** | 2.15E-09 | **1.64E-08** |
| ENSG00000049541.13 | **RFC2** | 5982 | protein_coding | **850.297** | -0.627 | -0.189 | **0.648** | 2.15E-09 | **1.64E-08** |
| ENSG00000146143.21 | **PRIM2** | 5558 | protein_coding | **310.355** | -0.735 | -0.221 | **0.601** | 2.16E-09 | **1.65E-08** |
| ENSG00000223768.5 | **LINC00205** | 642852 | lncRNA | **156.272** | 1.148 | 0.346 | **2.216** | 2.17E-09 | **1.66E-08** |
| ENSG00000122507.23 | **BBS9** | 27241 | protein_coding | **126.069** | 0.931 | 0.280 | **1.907** | 2.18E-09 | **1.66E-08** |
| ENSG00000176092.18 | **CRYBG2** | 55057 | protein_coding | **14.719** | 4.712 | 1.418 | **26.208** | 2.20E-09 | **1.67E-08** |
| ENSG00000293489.2 | **CMAHP** | 8418 | lncRNA | **159.657** | -1.205 | -0.363 | **0.434** | 2.21E-09 | **1.69E-08** |
| ENSG00000090006.20 | **LTBP4** | 8425 | protein_coding | **1370.165** | 1.031 | 0.310 | **2.044** | 2.22E-09 | **1.69E-08** |
| ENSG00000249348.3 | **UGDH-AS1** | 10088577 | lncRNA | **72.097** | 1.122 | 0.338 | **2.176** | 2.22E-09 | **1.69E-08** |
| ENSG00000271503.8 | **CCL5** | 6352 | protein_coding | **923.392** | 1.238 | 0.373 | **2.358** | 2.23E-09 | **1.70E-08** |
| ENSG00000238260.4 | **ENSG00000238260** | | lncRNA | **29.022** | 2.155 | 0.649 | **4.454** | 2.23E-09 | **1.70E-08** |
| ENSG00000104093.16 | **DMXL2** | 23312 | protein_coding | **460.966** | 0.998 | 0.300 | **1.997** | 2.24E-09 | **1.70E-08** |
| ENSG00000182508.16 | **LHFPL1** | 340596 | protein_coding | **23.144** | 2.546 | 0.766 | **5.840** | 2.24E-09 | **1.71E-08** |
| ENSG00000291210.2 | **ENSG00000291210** | | lncRNA | **260.036** | 0.918 | 0.276 | **1.889** | 2.25E-09 | **1.71E-08** |
| ENSG00000276564.1 | **ENSG00000276564** | | lncRNA | **32.011** | 1.474 | 0.444 | **2.778** | 2.26E-09 | **1.72E-08** |
| ENSG00000254726.3 | **MEX3A** | 92312 | protein_coding | **80.525** | 1.577 | 0.475 | **2.984** | 2.28E-09 | **1.73E-08** |
| ENSG00000076242.20 | **MLH1** | 4292 | protein_coding | **633.349** | -0.693 | -0.209 | **0.619** | 2.30E-09 | **1.75E-08** |

| | | | | | | | | | |
|---|---|---|---|---|---|---|---|---|---|
| ENSG00000177875.5 | **CCDC184** | 387856 | protein_coding | **15.554** | 4.129 | 1.243 | **17.493** | 2.30E-09 | **1.75E-08** |
| ENSG00000174123.12 | **TLR10** | 81793 | protein_coding | **988.303** | -0.700 | -0.211 | **0.616** | 2.32E-09 | **1.76E-08** |
| ENSG00000165671.24 | **NSD1** | 64324 | protein_coding | **807.140** | -0.722 | -0.217 | **0.606** | 2.33E-09 | **1.76E-08** |
| ENSG00000189308.13 | **LIN54** | 132660 | protein_coding | **157.274** | -1.062 | -0.320 | **0.479** | 2.33E-09 | **1.77E-08** |
| ENSG00000204856.14 | **FAM216A** | 29902 | protein_coding | **503.802** | -0.631 | -0.190 | **0.646** | 2.33E-09 | **1.77E-08** |
| ENSG00000167700.11 | **SLC33A2** | 113655 | protein_coding | **236.678** | 0.713 | 0.215 | **1.639** | 2.34E-09 | **1.77E-08** |
| ENSG00000277117.6 | **ENSG0000(** | 10272399( | artifact | **127.075** | -1.537 | -0.463 | **0.345** | 2.34E-09 | **1.77E-08** |
| ENSG00000086589.14 | **RBM22** | 55696 | protein_coding | **609.893** | -0.681 | -0.205 | **0.624** | 2.34E-09 | **1.77E-08** |
| ENSG00000134222.18 | **PSRC1** | 84722 | protein_coding | **198.292** | -1.000 | -0.301 | **0.500** | 2.34E-09 | **1.77E-08** |
| ENSG00000100211.14 | **CBY1** | 25776 | protein_coding | **398.548** | 0.749 | 0.225 | **1.680** | 2.35E-09 | **1.78E-08** |
| ENSG00000270021.1 | **ENSG00000270021** | | lncRNA | **23.765** | 2.025 | 0.610 | **4.070** | 2.37E-09 | **1.79E-08** |
| ENSG00000148296.9 | **SURF6** | 6838 | protein_coding | **632.057** | -0.537 | -0.162 | **0.689** | 2.39E-09 | **1.80E-08** |
| ENSG00000011243.21 | **AKAP8L** | 26993 | protein_coding | **3254.267** | 0.646 | 0.194 | **1.565** | 2.39E-09 | **1.81E-08** |
| ENSG00000124074.14 | **ENKD1** | 84080 | protein_coding | **432.702** | 0.927 | 0.279 | **1.901** | 2.40E-09 | **1.81E-08** |
| ENSG00000128626.14 | **MRPS12** | 6183 | protein_coding | **622.488** | -0.671 | -0.202 | **0.628** | 2.40E-09 | **1.81E-08** |
| ENSG00000178149.19 | **DALRD3** | 55152 | protein_coding | **411.170** | 0.673 | 0.203 | **1.594** | 2.40E-09 | **1.81E-08** |
| ENSG00000146729.12 | **NIPSNAP2** | 2631 | protein_coding | **369.501** | -0.718 | -0.216 | **0.608** | 2.44E-09 | **1.84E-08** |
| ENSG00000087884.18 | **AAMDC** | 28971 | protein_coding | **203.499** | 0.811 | 0.244 | **1.755** | 2.45E-09 | **1.85E-08** |
| ENSG00000149150.11 | **SLC43A1** | 8501 | protein_coding | **427.402** | -0.858 | -0.258 | **0.552** | 2.45E-09 | **1.85E-08** |
| ENSG00000181027.13 | **FKRP** | 79147 | protein_coding | **230.586** | -0.841 | -0.253 | **0.558** | 2.47E-09 | **1.86E-08** |
| ENSG00000144655.17 | **CSRNP1** | 64651 | protein_coding | **544.158** | -0.649 | -0.195 | **0.638** | 2.48E-09 | **1.87E-08** |
| ENSG00000055332.21 | **EIF2AK2** | 5610 | protein_coding | **1577.048** | -0.812 | -0.245 | **0.569** | 2.49E-09 | **1.87E-08** |
| ENSG00000001630.20 | **CYP51A1** | 1595 | protein_coding | **902.725** | -1.067 | -0.321 | **0.477** | 2.49E-09 | **1.87E-08** |
| ENSG00000196372.15 | **ASB13** | 79754 | protein_coding | **51.181** | 2.322 | 0.699 | **5.000** | 2.50E-09 | **1.88E-08** |
| ENSG00000221990.5 | **EXOC3-AS1** | | lncRNA | **125.647** | 0.867 | 0.261 | **1.824** | 2.52E-09 | **1.89E-08** |
| ENSG00000137802.17 | **MAPKBP1** | 23005 | protein_coding | **562.586** | 0.801 | 0.241 | **1.743** | 2.53E-09 | **1.90E-08** |
| ENSG00000111653.22 | **ING4** | 51147 | protein_coding | **599.646** | 0.594 | 0.179 | **1.509** | 2.54E-09 | **1.91E-08** |
| ENSG00000112079.12 | **STK38** | 11329 | protein_coding | **960.663** | 0.613 | 0.185 | **1.530** | 2.55E-09 | **1.91E-08** |
| ENSG00000188070.11 | **ZFTA** | 65998 | protein_coding | **195.142** | 0.954 | 0.287 | **1.938** | 2.63E-09 | **1.98E-08** |
| ENSG00000126456.18 | **IRF3** | 3661 | protein_coding | **3783.888** | 0.738 | 0.222 | **1.668** | 2.65E-09 | **1.99E-08** |
| ENSG00000205903.9 | **ZNF316** | 100131017 | protein_coding | **1540.434** | 0.443 | 0.133 | **1.360** | 2.66E-09 | **1.99E-08** |

| | | | | | | | | | |
|---|---|---|---|---|---|---|---|---|---|
| ENSG00000177169.12 | **ULK1** | 8408 | protein_coding | **754.944** | 0.755 | 0.227 | **1.688** | 2.66E-09 | **2.00E-08** |
| ENSG00000116701.18 | **NCF2** | 4688 | protein_coding | **1155.070** | 0.792 | 0.238 | **1.731** | 2.67E-09 | **2.00E-08** |
| ENSG00000160703.19 | **NLRX1** | 79671 | protein_coding | **195.022** | -0.882 | -0.265 | **0.543** | 2.67E-09 | **2.00E-08** |
| ENSG00000165959.14 | **CLMN** | 79789 | protein_coding | **432.146** | 1.140 | 0.343 | **2.204** | 2.68E-09 | **2.01E-08** |
| ENSG00000260804.4 | **LINC01963** | 150967 | lncRNA | **121.758** | 1.239 | 0.373 | **2.360** | 2.71E-09 | **2.03E-08** |
| ENSG00000168453.18 | **HR** | 55806 | protein_coding | **15.325** | 4.049 | 1.219 | **16.552** | 2.74E-09 | **2.05E-08** |
| ENSG00000231999.9 | **LRRC8C-DT** | | lncRNA | **80.893** | 1.645 | 0.495 | **3.127** | 2.74E-09 | **2.05E-08** |
| ENSG00000047230.17 | **CTPS2** | 56474 | protein_coding | **412.867** | -0.967 | -0.291 | **0.512** | 2.74E-09 | **2.05E-08** |
| ENSG00000197361.9 | **FBXL22** | 283807 | protein_coding | **28.931** | 1.680 | 0.506 | **3.205** | 2.74E-09 | **2.05E-08** |
| ENSG00000050165.21 | **DKK3** | 27122 | protein_coding | **9.066** | 4.524 | 1.362 | **23.002** | 2.76E-09 | **2.07E-08** |
| ENSG00000197461.15 | **PDGFA** | 5154 | protein_coding | **229.969** | -1.145 | -0.345 | **0.452** | 2.78E-09 | **2.07E-08** |
| ENSG00000167566.19 | **NCKAP5L** | 57701 | protein_coding | **477.971** | 0.779 | 0.234 | **1.716** | 2.78E-09 | **2.07E-08** |
| ENSG00000134717.20 | **BTF3L4** | 91408 | protein_coding | **446.724** | -0.795 | -0.239 | **0.576** | 2.86E-09 | **2.14E-08** |
| ENSG00000165102.17 | **HGSNAT** | 138050 | protein_coding | **1012.434** | 0.890 | 0.268 | **1.853** | 2.87E-09 | **2.14E-08** |
| ENSG00000203867.11 | **RBM20** | 282996 | protein_coding | **37.199** | 2.190 | 0.659 | **4.564** | 2.88E-09 | **2.14E-08** |
| ENSG00000133316.18 | **WDR74** | 54663 | protein_coding | **1021.472** | -0.553 | -0.167 | **0.682** | 2.93E-09 | **2.18E-08** |
| ENSG00000187091.16 | **PLCD1** | 5333 | protein_coding | **146.666** | 1.329 | 0.400 | **2.513** | 2.93E-09 | **2.18E-08** |
| ENSG00000132004.15 | **FBXW9** | 84261 | protein_coding | **97.985** | -0.948 | -0.285 | **0.518** | 2.93E-09 | **2.18E-08** |
| ENSG00000138758.14 | **SEPTIN11** | 55752 | protein_coding | **1206.112** | -0.744 | -0.224 | **0.597** | 2.96E-09 | **2.20E-08** |
| ENSG00000104938.20 | **CLEC4M** | 10332 | protein_coding | **12.696** | 3.459 | 1.041 | **10.997** | 2.97E-09 | **2.21E-08** |
| ENSG00000173080.6 | **RXFP4** | 339403 | protein_coding | **7.386** | 5.579 | 1.679 | **47.790** | 3.02E-09 | **2.25E-08** |
| ENSG00000177025.3 | **C19orf18** | 147685 | protein_coding | **31.823** | 2.101 | 0.632 | **4.289** | 3.07E-09 | **2.28E-08** |
| ENSG00000229539.3 | **PANK2-AS1** | | lncRNA | **54.707** | 1.302 | 0.392 | **2.466** | 3.07E-09 | **2.28E-08** |
| ENSG00000104081.16 | **BMF** | 90427 | protein_coding | **2633.948** | 0.730 | 0.220 | **1.658** | 3.07E-09 | **2.28E-08** |
| ENSG00000173890.19 | **GPR160** | 26996 | protein_coding | **468.010** | 0.925 | 0.278 | **1.899** | 3.09E-09 | **2.30E-08** |
| ENSG00000309115.1 | **ENSG00000309115** | | lncRNA | **18.623** | 2.183 | 0.657 | **4.540** | 3.10E-09 | **2.30E-08** |
| ENSG00000115364.16 | **MRPL19** | 9801 | protein_coding | **556.949** | -0.772 | -0.232 | **0.586** | 3.11E-09 | **2.31E-08** |
| ENSG00000108813.12 | **DLX4** | 1748 | protein_coding | **25.906** | 1.863 | 0.561 | **3.637** | 3.12E-09 | **2.32E-08** |
| ENSG00000215440.14 | **NPEPL1** | 79716 | protein_coding | **1798.195** | 0.713 | 0.214 | **1.639** | 3.13E-09 | **2.32E-08** |
| ENSG00000134452.22 | **FBH1** | 84893 | protein_coding | **2058.879** | 0.799 | 0.241 | **1.740** | 3.13E-09 | **2.32E-08** |
| ENSG00000113456.22 | **RAD1** | 5810 | protein_coding | **257.727** | -0.761 | -0.229 | **0.590** | 3.15E-09 | **2.33E-08** |

| | | | | | | | | | |
|---|---|---|---|---|---|---|---|---|---|
| ENSG00000187775.17 | DNAH17 | 8632 | protein_coding | 72.925 | 1.661 | 0.500 | 3.162 | 3.16E-09 | 2.34E-08 |
| ENSG00000060566.16 | CREB3L3 | 84699 | protein_coding | 15.350 | 3.295 | 0.992 | 9.816 | 3.28E-09 | 2.43E-08 |
| ENSG00000092529.27 | CAPN3 | 825 | protein_coding | 528.026 | 0.773 | 0.233 | 1.709 | 3.29E-09 | 2.44E-08 |
| ENSG00000170545.19 | SMAGP | 57228 | protein_coding | 33.801 | 2.031 | 0.611 | 4.087 | 3.31E-09 | 2.45E-08 |
| ENSG00000105835.15 | NAMPT | 10135 | protein_coding | 1714.602 | -0.972 | -0.293 | 0.510 | 3.31E-09 | 2.45E-08 |
| ENSG00000165140.14 | FBP1 | 2203 | protein_coding | 14.661 | 3.376 | 1.016 | 10.383 | 3.32E-09 | 2.46E-08 |
| ENSG00000286122.2 | LINC02964 | | lncRNA | 19.587 | 2.787 | 0.839 | 6.900 | 3.34E-09 | 2.47E-08 |
| ENSG00000272990.1 | ENSG00000272990 | | lncRNA | 101.742 | 1.455 | 0.438 | 2.742 | 3.35E-09 | 2.48E-08 |
| ENSG00000144749.16 | LRIG1 | 26018 | protein_coding | 297.773 | 0.691 | 0.208 | 1.614 | 3.36E-09 | 2.48E-08 |
| ENSG00000111450.17 | STX2 | 2054 | protein_coding | 459.433 | 0.670 | 0.202 | 1.592 | 3.37E-09 | 2.49E-08 |
| ENSG00000100060.20 | MFNG | 4242 | protein_coding | 906.619 | -0.658 | -0.198 | 0.634 | 3.37E-09 | 2.49E-08 |
| ENSG00000271858.5 | ENSG0000( | 101928965 | lncRNA | 30.585 | 1.848 | 0.556 | 3.600 | 3.38E-09 | 2.49E-08 |
| ENSG00000072062.18 | PRKACA | 5566 | protein_coding | 1244.129 | 0.724 | 0.218 | 1.651 | 3.41E-09 | 2.52E-08 |
| ENSG00000114491.16 | UMPS | 7372 | protein_coding | 392.519 | -0.930 | -0.280 | 0.525 | 3.42E-09 | 2.52E-08 |
| ENSG00000261338.2 | ENSG00000261338 | | lncRNA | 27.120 | 1.717 | 0.517 | 3.288 | 3.45E-09 | 2.54E-08 |
| ENSG00000174938.17 | SEZ6L2 | 26470 | protein_coding | 17.301 | 4.300 | 1.294 | 19.700 | 3.49E-09 | 2.57E-08 |
| ENSG00000214063.14 | TSPAN4 | 7106 | protein_coding | 219.903 | 0.984 | 0.296 | 1.979 | 3.49E-09 | 2.57E-08 |
| ENSG00000154016.16 | GRAP | 10750 | protein_coding | 1324.716 | -0.804 | -0.242 | 0.573 | 3.49E-09 | 2.57E-08 |
| ENSG00000023572.11 | GLRX2 | 51022 | protein_coding | 151.738 | -0.839 | -0.252 | 0.559 | 3.50E-09 | 2.57E-08 |
| ENSG00000174004.8 | NRROS | 375387 | protein_coding | 634.533 | -0.921 | -0.277 | 0.528 | 3.52E-09 | 2.59E-08 |
| ENSG00000288993.2 | ENSG00000288993 | | lncRNA | 19.679 | 2.425 | 0.730 | 5.371 | 3.58E-09 | 2.63E-08 |
| ENSG00000108784.12 | NAGLU | 4669 | protein_coding | 506.467 | -0.610 | -0.184 | 0.655 | 3.61E-09 | 2.65E-08 |
| ENSG00000183087.17 | GAS6 | 2621 | protein_coding | 106.784 | 1.173 | 0.353 | 2.255 | 3.64E-09 | 2.68E-08 |
| ENSG00000100418.10 | DESI1 | 27351 | protein_coding | 2422.752 | -0.878 | -0.264 | 0.544 | 3.68E-09 | 2.70E-08 |
| ENSG00000111879.21 | FAM184A | 79632 | protein_coding | 24.542 | 2.245 | 0.676 | 4.740 | 3.68E-09 | 2.70E-08 |
| ENSG00000065060.19 | BLTP3A | 54887 | protein_coding | 387.120 | -0.949 | -0.286 | 0.518 | 3.70E-09 | 2.71E-08 |
| ENSG00000186314.14 | PRELID2 | 153768 | protein_coding | 61.624 | -1.625 | -0.489 | 0.324 | 3.71E-09 | 2.72E-08 |
| ENSG00000159720.15 | ATP6V0D1 | 9114 | protein_coding | 813.537 | 0.955 | 0.288 | 1.939 | 3.71E-09 | 2.72E-08 |
| ENSG00000071626.19 | DAZAP1 | 26528 | protein_coding | 3565.869 | -0.598 | -0.180 | 0.661 | 3.72E-09 | 2.73E-08 |
| ENSG00000071462.15 | BUD23 | 114049 | protein_coding | 1130.044 | -0.582 | -0.175 | 0.668 | 3.73E-09 | 2.74E-08 |
| ENSG00000232788.3 | ITGA6-AS1 | 101929947 | lncRNA | 68.060 | -1.641 | -0.494 | 0.321 | 3.77E-09 | 2.76E-08 |

| | | | | | | | | | |
|---|---|---|---|---|---|---|---|---|---|
| ENSG00000168916.18 | **ZNF608** | 57507 | protein_coding | **267.996** | 0.861 | 0.259 | **1.816** | 3.79E-09 | **2.77E-08** |
| ENSG00000145494.14 | **NDUFS6** | 4726 | protein_coding | **1012.972** | -0.679 | -0.204 | **0.624** | 3.82E-09 | **2.79E-08** |
| ENSG00000234184.9 | **LINC01781** | 10192741 | lncRNA | **62.524** | 1.519 | 0.457 | **2.867** | 3.83E-09 | **2.80E-08** |
| ENSG00000285292.1 | **ABCF2-H2E** | 11448383 | protein_coding | **30.813** | 2.580 | 0.777 | **5.979** | 3.85E-09 | **2.82E-08** |
| ENSG00000215018.11 | **COL28A1** | 340267 | protein_coding | **13.564** | 4.057 | 1.221 | **16.644** | 3.90E-09 | **2.85E-08** |
| ENSG00000116761.14 | **CTH** | 1491 | protein_coding | **618.335** | -1.423 | -0.428 | **0.373** | 3.94E-09 | **2.88E-08** |
| ENSG00000255152.8 | **MSH5-SAPCD1** | | protein_coding | **430.479** | -1.055 | -0.317 | **0.481** | 3.95E-09 | **2.89E-08** |
| ENSG00000284954.1 | **ENSG0000** | 10050754 | lncRNA | **21.238** | 2.619 | 0.788 | **6.145** | 3.96E-09 | **2.89E-08** |
| ENSG00000130204.15 | **TOMM40** | 10452 | protein_coding | **1587.556** | -0.689 | -0.207 | **0.620** | 3.97E-09 | **2.89E-08** |
| ENSG00000172613.10 | **RAD9A** | 5883 | protein_coding | **1444.104** | 0.813 | 0.245 | **1.756** | 3.97E-09 | **2.90E-08** |
| ENSG00000160094.17 | **ZNF362** | 149076 | protein_coding | **292.853** | 0.719 | 0.217 | **1.647** | 3.97E-09 | **2.90E-08** |
| ENSG00000214941.10 | **ZSWIM7** | 125150 | protein_coding | **235.439** | 0.773 | 0.233 | **1.709** | 3.98E-09 | **2.90E-08** |
| ENSG00000177150.15 | **FAM210A** | 125228 | protein_coding | **271.792** | -0.889 | -0.268 | **0.540** | 3.98E-09 | **2.90E-08** |
| ENSG00000137834.17 | **SMAD6** | 4091 | protein_coding | **12.944** | 3.072 | 0.925 | **8.410** | 4.01E-09 | **2.92E-08** |
| ENSG00000291233.2 | **ENSG00000291233** | | lncRNA | **155.005** | 1.002 | 0.302 | **2.002** | 4.02E-09 | **2.92E-08** |
| ENSG00000297379.1 | **ENSG0000** | 12490162 | lncRNA | **7.272** | 5.537 | 1.667 | **46.430** | 4.03E-09 | **2.93E-08** |
| ENSG00000169972.15 | **PUSL1** | 126789 | protein_coding | **200.358** | -0.971 | -0.292 | **0.510** | 4.08E-09 | **2.96E-08** |
| ENSG00000196230.16 | **TUBB** | 203068 | protein_coding | **23757.514** | -1.026 | -0.309 | **0.491** | 4.09E-09 | **2.97E-08** |
| ENSG00000109654.18 | **TRIM2** | 23321 | protein_coding | **43.367** | 2.052 | 0.618 | **4.148** | 4.09E-09 | **2.97E-08** |
| ENSG00000142552.10 | **RCN3** | 57333 | protein_coding | **74.038** | 2.059 | 0.620 | **4.167** | 4.11E-09 | **2.99E-08** |
| ENSG00000117395.15 | **EBNA1BP2** | 10969 | protein_coding | **818.721** | -0.676 | -0.204 | **0.626** | 4.11E-09 | **2.99E-08** |
| ENSG00000183137.18 | **CEP57L1** | 285753 | protein_coding | **199.009** | -1.090 | -0.328 | **0.470** | 4.14E-09 | **3.01E-08** |
| ENSG00000135677.13 | **GNS** | 2799 | protein_coding | **714.537** | 0.999 | 0.301 | **1.999** | 4.16E-09 | **3.02E-08** |
| ENSG00000225519.2 | **ENSG00000225519** | | lncRNA | **124.347** | -1.842 | -0.555 | **0.279** | 4.19E-09 | **3.04E-08** |
| ENSG00000231453.2 | **LINC01305** | 285084 | lncRNA | **10.500** | 4.720 | 1.421 | **26.349** | 4.20E-09 | **3.05E-08** |
| ENSG00000258727.2 | **AP1G2-AS1** | 10272481 | lncRNA | **543.813** | 1.035 | 0.312 | **2.050** | 4.21E-09 | **3.06E-08** |
| ENSG00000135537.19 | **AFG1L** | 246269 | protein_coding | **74.237** | -1.141 | -0.344 | **0.453** | 4.22E-09 | **3.06E-08** |
| ENSG00000151882.12 | **CCL28** | 56477 | protein_coding | **204.518** | -1.157 | -0.348 | **0.449** | 4.27E-09 | **3.09E-08** |
| ENSG00000132535.23 | **DLG4** | 1742 | protein_coding | **251.467** | 1.412 | 0.425 | **2.661** | 4.28E-09 | **3.10E-08** |
| ENSG00000185347.21 | **TEDC1** | 283643 | protein_coding | **318.298** | -0.962 | -0.290 | **0.513** | 4.28E-09 | **3.10E-08** |
| ENSG00000091483.10 | **FH** | 2271 | protein_coding | **1394.401** | -0.514 | -0.155 | **0.700** | 4.31E-09 | **3.12E-08** |

| | | | | | | | | | |
|---|---|---|---|---|---|---|---|---|---|
| ENSG00000182552.16 | **RWDD4** | 201965 | protein_coding | **273.073** | -0.914 | -0.275 | **0.531** | 4.32E-09 | **3.13E-08** |
| ENSG00000105971.17 | **CAV2** | 858 | protein_coding | **20.191** | 2.714 | 0.817 | **6.564** | 4.33E-09 | **3.13E-08** |
| ENSG00000113272.16 | **THG1L** | 54974 | protein_coding | **520.133** | -0.643 | -0.194 | **0.640** | 4.34E-09 | **3.14E-08** |
| ENSG00000297141.1 | **ENSG00000297141** | | lncRNA | **38.481** | 1.723 | 0.519 | **3.300** | 4.34E-09 | **3.14E-08** |
| ENSG00000141965.5 | **FEM1A** | 55527 | protein_coding | **479.470** | -0.643 | -0.193 | **0.641** | 4.37E-09 | **3.16E-08** |
| ENSG00000013288.11 | **MAN2B2** | 23324 | protein_coding | **450.523** | -0.577 | -0.174 | **0.670** | 4.38E-09 | **3.16E-08** |
| ENSG00000119446.16 | **RBM18** | 92400 | protein_coding | **284.845** | -0.793 | -0.239 | **0.577** | 4.41E-09 | **3.19E-08** |
| ENSG00000178761.17 | **FAM219B** | 57184 | protein_coding | **977.048** | -0.685 | -0.206 | **0.622** | 4.41E-09 | **3.19E-08** |
| ENSG00000205476.10 | **CCDC85C** | 317762 | protein_coding | **105.451** | 1.826 | 0.550 | **3.545** | 4.42E-09 | **3.19E-08** |
| ENSG00000236609.6 | **ZNF853** | 54753 | protein_coding | **16.023** | 3.660 | 1.102 | **12.638** | 4.50E-09 | **3.25E-08** |
| ENSG00000103051.23 | **COG4** | 25839 | protein_coding | **2085.283** | 0.638 | 0.192 | **1.556** | 4.50E-09 | **3.25E-08** |
| ENSG00000095587.11 | **TLL2** | 7093 | protein_coding | **43.969** | 1.880 | 0.566 | **3.682** | 4.53E-09 | **3.27E-08** |
| ENSG00000089902.12 | **RCOR1** | 23186 | protein_coding | **344.377** | 0.771 | 0.232 | **1.707** | 4.57E-09 | **3.29E-08** |
| ENSG00000291228.2 | **ENSG0000(** | 652276 | lncRNA | **107.290** | -1.362 | -0.410 | **0.389** | 4.59E-09 | **3.30E-08** |
| ENSG00000137747.18 | **TMPRSS13** | 84000 | protein_coding | **42.526** | 1.647 | 0.496 | **3.131** | 4.61E-09 | **3.32E-08** |
| ENSG00000123737.15 | **EXOSC9** | 5393 | protein_coding | **1095.548** | -0.689 | -0.207 | **0.620** | 4.63E-09 | **3.33E-08** |
| ENSG00000115392.15 | **FANCL** | 55120 | protein_coding | **739.367** | -0.648 | -0.195 | **0.638** | 4.66E-09 | **3.35E-08** |
| ENSG00000197771.15 | **MCMBP** | 79892 | protein_coding | **638.001** | -0.781 | -0.235 | **0.582** | 4.66E-09 | **3.35E-08** |
| ENSG00000177238.16 | **TRIM72** | 493829 | protein_coding | **24.172** | 2.261 | 0.681 | **4.793** | 4.69E-09 | **3.37E-08** |
| ENSG00000048462.12 | **TNFRSF17** | 608 | protein_coding | **807.718** | -1.151 | -0.347 | **0.450** | 4.70E-09 | **3.38E-08** |
| ENSG00000136718.12 | **IMP4** | 92856 | protein_coding | **1611.464** | -0.648 | -0.195 | **0.638** | 4.71E-09 | **3.38E-08** |
| ENSG00000112294.16 | **ALDH5A1** | 7915 | protein_coding | **359.557** | -1.119 | -0.337 | **0.460** | 4.75E-09 | **3.41E-08** |
| ENSG00000104976.15 | **SNAPC2** | 6618 | protein_coding | **397.804** | 0.723 | 0.218 | **1.650** | 4.76E-09 | **3.42E-08** |
| ENSG00000188342.15 | **GTF2F2** | 2963 | protein_coding | **510.677** | -0.617 | -0.186 | **0.652** | 4.78E-09 | **3.43E-08** |
| ENSG00000188372.16 | **ZP3** | 7784 | protein_coding | **124.604** | 1.323 | 0.398 | **2.503** | 4.83E-09 | **3.46E-08** |
| ENSG00000100281.15 | **HMGXB4** | 10042 | protein_coding | **266.666** | -0.750 | -0.226 | **0.595** | 4.84E-09 | **3.47E-08** |
| ENSG00000196218.15 | **RYR1** | 6261 | protein_coding | **60.765** | 1.627 | 0.490 | **3.088** | 4.85E-09 | **3.48E-08** |
| ENSG00000044459.17 | **CNTLN** | 54875 | protein_coding | **86.317** | -1.171 | -0.353 | **0.444** | 4.96E-09 | **3.55E-08** |
| ENSG00000128708.15 | **HAT1** | 8520 | protein_coding | **391.103** | -1.174 | -0.353 | **0.443** | 5.02E-09 | **3.59E-08** |
| ENSG00000138756.21 | **BMP2K** | 55589 | protein_coding | **669.956** | 0.574 | 0.173 | **1.489** | 5.04E-09 | **3.60E-08** |
| ENSG00000160953.19 | **PWWP3A** | 84939 | protein_coding | **1055.703** | 0.670 | 0.202 | **1.591** | 5.04E-09 | **3.60E-08** |

| | | | | | | | | | |
|---|---|---|---|---|---|---|---|---|---|
| ENSG00000116299.19 | **ELAPOR1** | 57535 | protein_coding | **65.514** | 1.873 | 0.564 | **3.662** | 5.05E-09 | **3.61E-08** |
| ENSG00000267751.7 | **BSG-AS1** | | lncRNA | **58.677** | -1.553 | -0.468 | **0.341** | 5.05E-09 | **3.61E-08** |
| ENSG00000168591.18 | **TMUB2** | 79089 | protein_coding | **873.165** | 0.554 | 0.167 | **1.468** | 5.07E-09 | **3.63E-08** |
| ENSG00000224066.1 | **IQCC-AS1** | | lncRNA | **15.515** | 2.589 | 0.779 | **6.017** | 5.10E-09 | **3.64E-08** |
| ENSG00000153015.18 | **CWC27** | 10283 | protein_coding | **427.294** | -0.701 | -0.211 | **0.615** | 5.12E-09 | **3.66E-08** |
| ENSG00000140931.22 | **CMTM3** | 123920 | protein_coding | **370.366** | 1.259 | 0.379 | **2.394** | 5.19E-09 | **3.70E-08** |
| ENSG00000110811.22 | **P3H3** | 10536 | protein_coding | **19.928** | 2.143 | 0.645 | **4.417** | 5.22E-09 | **3.72E-08** |
| ENSG00000102181.24 | **CD99L2** | 83692 | protein_coding | **650.638** | 0.623 | 0.187 | **1.540** | 5.31E-09 | **3.78E-08** |
| ENSG00000178297.16 | **TMPRSS9** | 360200 | protein_coding | **22.027** | 3.667 | 1.104 | **12.704** | 5.33E-09 | **3.80E-08** |
| ENSG00000171806.14 | **METTL18** | 92342 | protein_coding | **111.522** | -0.882 | -0.266 | **0.542** | 5.39E-09 | **3.84E-08** |
| ENSG00000126391.16 | **FRMD8** | 83786 | protein_coding | **718.156** | 0.682 | 0.205 | **1.605** | 5.42E-09 | **3.86E-08** |
| ENSG00000228801.9 | **PCMTD1-D** | 10272433( | lncRNA | **153.751** | 0.927 | 0.279 | **1.901** | 5.42E-09 | **3.86E-08** |
| ENSG00000170860.6 | **LSM3** | 27258 | protein_coding | **620.021** | -0.854 | -0.257 | **0.553** | 5.43E-09 | **3.86E-08** |
| ENSG00000117597.20 | **UTP25** | 27042 | protein_coding | **189.383** | -0.923 | -0.278 | **0.527** | 5.46E-09 | **3.88E-08** |
| ENSG00000136490.11 | **LIMD2** | 80774 | protein_coding | **4191.901** | -0.889 | -0.268 | **0.540** | 5.49E-09 | **3.91E-08** |
| ENSG00000141934.12 | **PLPP2** | 8612 | protein_coding | **31.734** | 3.464 | 1.043 | **11.032** | 5.51E-09 | **3.92E-08** |
| ENSG00000204991.13 | **SPIRE2** | 84501 | protein_coding | **19.612** | 3.052 | 0.919 | **8.293** | 5.60E-09 | **3.98E-08** |
| ENSG00000145555.16 | **MYO10** | 4651 | protein_coding | **14.445** | 3.620 | 1.090 | **12.297** | 5.60E-09 | **3.98E-08** |
| ENSG00000171466.12 | **ZNF562** | 54811 | protein_coding | **253.446** | -0.687 | -0.207 | **0.621** | 5.61E-09 | **3.99E-08** |
| ENSG00000197093.13 | **GAL3ST4** | 79690 | protein_coding | **215.186** | 1.843 | 0.555 | **3.588** | 5.70E-09 | **4.04E-08** |
| ENSG00000043462.15 | **LCP2** | 3937 | protein_coding | **231.881** | -1.019 | -0.307 | **0.493** | 5.70E-09 | **4.04E-08** |
| ENSG00000100099.24 | **HPS4** | 89781 | protein_coding | **1145.806** | -0.483 | -0.145 | **0.716** | 5.76E-09 | **4.09E-08** |
| ENSG00000181163.16 | **NPM1** | 4869 | protein_coding | **20514.979** | -0.454 | -0.137 | **0.730** | 5.78E-09 | **4.10E-08** |
| ENSG00000167118.13 | **URM1** | 81605 | protein_coding | **926.531** | -0.656 | -0.197 | **0.635** | 5.79E-09 | **4.11E-08** |
| ENSG00000179241.15 | **LDLRAD3** | 143458 | protein_coding | **30.139** | 1.828 | 0.550 | **3.550** | 5.84E-09 | **4.14E-08** |
| ENSG00000068885.17 | **IFT80** | 57560 | protein_coding | **478.353** | 0.905 | 0.272 | **1.872** | 5.89E-09 | **4.17E-08** |
| ENSG00000026025.18 | **VIM** | 7431 | protein_coding | **7287.478** | -0.734 | -0.221 | **0.601** | 5.91E-09 | **4.18E-08** |
| ENSG00000112701.20 | **SENP6** | 26054 | protein_coding | **865.201** | -0.631 | -0.190 | **0.646** | 5.93E-09 | **4.20E-08** |
| ENSG00000255717.10 | **SNHG1** | 23642 | lncRNA | **3682.184** | -1.050 | -0.316 | **0.483** | 5.95E-09 | **4.21E-08** |
| ENSG00000163935.16 | **SFMBT1** | 51460 | protein_coding | **110.752** | -0.811 | -0.244 | **0.570** | 5.99E-09 | **4.23E-08** |
| ENSG00000075239.17 | **ACAT1** | 38 | protein_coding | **718.340** | -0.555 | -0.167 | **0.681** | 5.99E-09 | **4.24E-08** |

| | | | | | | | | | |
|---|---|---|---|---|---|---|---|---|---|
| ENSG00000133460.22 | **SLC2A11** | 66035 | protein_coding | **326.585** | 0.752 | 0.226 | **1.684** | 6.06E-09 | **4.28E-08** |
| ENSG00000073803.16 | **MAP3K13** | 9175 | protein_coding | **384.409** | 0.597 | 0.180 | **1.513** | 6.07E-09 | **4.29E-08** |
| ENSG00000151876.15 | **FBXO4** | 26272 | protein_coding | **169.637** | -0.854 | -0.257 | **0.553** | 6.16E-09 | **4.35E-08** |
| ENSG00000198792.15 | **TMEM184I** | 25829 | protein_coding | **379.329** | 0.805 | 0.242 | **1.747** | 6.18E-09 | **4.36E-08** |
| ENSG00000231908.3 | **IDH1-AS1** | | lncRNA | **38.528** | 1.501 | 0.452 | **2.830** | 6.28E-09 | **4.43E-08** |
| ENSG00000130707.20 | **ASS1** | 445 | protein_coding | **700.696** | -1.569 | -0.472 | **0.337** | 6.44E-09 | **4.54E-08** |
| ENSG00000138378.22 | **STAT4** | 6775 | protein_coding | **120.051** | 1.351 | 0.407 | **2.552** | 6.45E-09 | **4.55E-08** |
| ENSG00000137225.13 | **CAPN11** | 11131 | protein_coding | **29.532** | 2.052 | 0.618 | **4.146** | 6.49E-09 | **4.58E-08** |
| ENSG00000110328.8 | **GALNT18** | 374378 | protein_coding | **132.959** | -1.365 | -0.411 | **0.388** | 6.51E-09 | **4.59E-08** |
| ENSG00000166173.12 | **LARP6** | 55323 | protein_coding | **39.196** | 2.067 | 0.622 | **4.190** | 6.63E-09 | **4.67E-08** |
| ENSG00000132275.13 | **RRP8** | 23378 | protein_coding | **411.737** | -0.631 | -0.190 | **0.646** | 6.70E-09 | **4.72E-08** |
| ENSG00000140320.14 | **BAHD1** | 22893 | protein_coding | **548.885** | 0.686 | 0.206 | **1.609** | 6.92E-09 | **4.87E-08** |
| ENSG00000174007.11 | **CEP19** | 84984 | protein_coding | **129.794** | 1.198 | 0.361 | **2.294** | 6.95E-09 | **4.89E-08** |
| ENSG00000073670.16 | **ADAM11** | 4185 | protein_coding | **27.490** | 2.008 | 0.604 | **4.021** | 7.06E-09 | **4.96E-08** |
| ENSG00000109919.12 | **MTCH2** | 23788 | protein_coding | **1142.887** | -0.827 | -0.249 | **0.564** | 7.07E-09 | **4.97E-08** |
| ENSG00000247556.8 | **OIP5-AS1** | 729082 | lncRNA | **1239.985** | -0.754 | -0.227 | **0.593** | 7.15E-09 | **5.02E-08** |
| ENSG00000187994.15 | **RINL** | 126432 | protein_coding | **686.115** | -0.897 | -0.270 | **0.537** | 7.20E-09 | **5.06E-08** |
| ENSG00000299955.1 | **ENSG00000299955** | | lncRNA | **25.904** | 2.514 | 0.757 | **5.714** | 7.26E-09 | **5.10E-08** |
| ENSG00000236287.10 | **ZBED5** | 58486 | protein_coding | **466.997** | -0.726 | -0.218 | **0.605** | 7.30E-09 | **5.13E-08** |
| ENSG00000135966.16 | **TGFBRAP1** | 9392 | protein_coding | **450.424** | -0.725 | -0.218 | **0.605** | 7.35E-09 | **5.16E-08** |
| ENSG00000214595.14 | **EML6** | 400954 | protein_coding | **34.574** | 3.461 | 1.042 | **11.011** | 7.36E-09 | **5.16E-08** |
| ENSG00000233828.6 | **MIR4280H** | 11775173( | lncRNA | **34.463** | 1.949 | 0.587 | **3.862** | 7.45E-09 | **5.22E-08** |
| ENSG00000065154.14 | **OAT** | 4942 | protein_coding | **179.305** | 1.331 | 0.401 | **2.516** | 7.48E-09 | **5.25E-08** |
| ENSG00000123131.15 | **PRDX4** | 10549 | protein_coding | **639.682** | -0.905 | -0.272 | **0.534** | 7.51E-09 | **5.26E-08** |
| ENSG00000104880.20 | **ARHGEF18** | 23370 | protein_coding | **1116.494** | 0.591 | 0.178 | **1.507** | 7.52E-09 | **5.27E-08** |
| ENSG00000149418.13 | **ST14** | 6768 | protein_coding | **31.098** | 2.067 | 0.622 | **4.189** | 7.58E-09 | **5.31E-08** |
| ENSG00000124772.13 | **CPNE5** | 57699 | protein_coding | **710.758** | 0.676 | 0.204 | **1.598** | 7.63E-09 | **5.34E-08** |
| ENSG00000140995.19 | **DEF8** | 54849 | protein_coding | **1141.605** | 0.565 | 0.170 | **1.479** | 7.67E-09 | **5.36E-08** |
| ENSG00000230107.2 | **ENSG00000230107** | | lncRNA | **165.494** | -1.231 | -0.370 | **0.426** | 7.68E-09 | **5.37E-08** |
| ENSG00000102178.14 | **UBL4A** | 8266 | protein_coding | **311.818** | -0.783 | -0.236 | **0.581** | 7.69E-09 | **5.38E-08** |
| ENSG00000011105.16 | **TSPAN9** | 10867 | protein_coding | **36.942** | 2.089 | 0.629 | **4.256** | 7.74E-09 | **5.41E-08** |

| | | | | | | | | | |
|---|---|---|---|---|---|---|---|---|---|
| ENSG00000143543.17 | **JTB** | 10899 | protein_coding | **2185.620** | -0.539 | -0.162 | **0.688** | 7.75E-09 | **5.41E-08** |
| ENSG00000245498.8 | **MSANTD2-** | 10050728 | lncRNA | **93.596** | 0.959 | 0.289 | **1.944** | 7.75E-09 | **5.42E-08** |
| ENSG00000213160.12 | **KLHL23** | 151230 | protein_coding | **139.017** | -0.957 | -0.288 | **0.515** | 7.82E-09 | **5.46E-08** |
| ENSG00000239789.8 | **MRPS17** | 51373 | protein_coding | **358.710** | -0.766 | -0.231 | **0.588** | 7.97E-09 | **5.56E-08** |
| ENSG00000249673.10 | **NOP14-AS1** | | lncRNA | **354.565** | 0.715 | 0.215 | **1.642** | 7.98E-09 | **5.57E-08** |
| ENSG00000034693.17 | **PEX3** | 8504 | protein_coding | **231.808** | -0.739 | -0.223 | **0.599** | 7.99E-09 | **5.57E-08** |
| ENSG00000160352.18 | **ZNF714** | 148206 | protein_coding | **129.057** | -1.325 | -0.399 | **0.399** | 8.00E-09 | **5.58E-08** |
| ENSG00000163950.15 | **SLBP** | 7884 | protein_coding | **999.396** | -0.690 | -0.208 | **0.620** | 8.02E-09 | **5.59E-08** |
| ENSG00000143801.20 | **PSEN2** | 5664 | protein_coding | **269.519** | 0.755 | 0.227 | **1.687** | 8.12E-09 | **5.65E-08** |
| ENSG00000276234.7 | **TADA2A** | 6871 | protein_coding | **457.863** | -0.805 | -0.242 | **0.572** | 8.12E-09 | **5.65E-08** |
| ENSG00000284308.3 | **C2orf81** | 388963 | protein_coding | **236.058** | 0.987 | 0.297 | **1.982** | 8.24E-09 | **5.73E-08** |
| ENSG00000120093.13 | **HOXB3** | 3213 | protein_coding | **375.587** | 0.829 | 0.250 | **1.777** | 8.27E-09 | **5.75E-08** |
| ENSG00000257727.8 | **CNPY2** | 10330 | protein_coding | **940.811** | -0.513 | -0.154 | **0.701** | 8.33E-09 | **5.79E-08** |
| ENSG00000286585.4 | **ENSG0000** | 10537417 | lncRNA | **21.257** | 2.475 | 0.745 | **5.558** | 8.33E-09 | **5.79E-08** |
| ENSG00000125124.16 | **BBS2** | 583 | protein_coding | **345.867** | 0.723 | 0.218 | **1.650** | 8.36E-09 | **5.81E-08** |
| ENSG00000186281.15 | **GPAT2** | 150763 | protein_coding | **174.575** | -1.676 | -0.505 | **0.313** | 8.40E-09 | **5.84E-08** |
| ENSG00000118246.16 | **FASTKD2** | 22868 | protein_coding | **361.263** | -0.723 | -0.218 | **0.606** | 8.44E-09 | **5.87E-08** |
| ENSG00000053524.14 | **MCF2L2** | 23101 | protein_coding | **27.156** | 3.082 | 0.928 | **8.470** | 8.52E-09 | **5.91E-08** |
| ENSG00000106477.22 | **CEP41** | 95681 | protein_coding | **148.195** | -0.997 | -0.300 | **0.501** | 8.52E-09 | **5.92E-08** |
| ENSG00000075415.17 | **SLC25A3** | 5250 | protein_coding | **6655.343** | -0.715 | -0.215 | **0.609** | 8.55E-09 | **5.93E-08** |
| ENSG00000290803.2 | **ENSG00000290803** | | lncRNA | **26.151** | 2.689 | 0.809 | **6.448** | 8.60E-09 | **5.97E-08** |
| ENSG00000175581.16 | **MRPL48** | 51642 | protein_coding | **563.022** | -0.668 | -0.201 | **0.629** | 8.62E-09 | **5.98E-08** |
| ENSG00000181924.10 | **COA4** | 51287 | protein_coding | **787.934** | -0.864 | -0.260 | **0.550** | 8.63E-09 | **5.98E-08** |
| ENSG00000116898.14 | **MRPS15** | 64960 | protein_coding | **1644.838** | -0.895 | -0.269 | **0.538** | 8.63E-09 | **5.98E-08** |
| ENSG00000089597.20 | **GANAB** | 23193 | protein_coding | **2821.923** | -0.495 | -0.149 | **0.710** | 8.65E-09 | **5.99E-08** |
| ENSG00000120889.15 | **TNFRSF10** | 8795 | protein_coding | **2603.927** | -0.594 | -0.179 | **0.662** | 8.67E-09 | **6.00E-08** |
| ENSG00000081377.19 | **CDC14B** | 8555 | protein_coding | **42.785** | 1.680 | 0.506 | **3.204** | 8.68E-09 | **6.01E-08** |
| ENSG00000141294.11 | **LRRC46** | 90506 | protein_coding | **23.722** | 1.858 | 0.559 | **3.626** | 8.71E-09 | **6.03E-08** |
| ENSG00000260231.3 | **KDM7A-DT** | 10013422 | lncRNA | **37.061** | 2.383 | 0.717 | **5.216** | 8.79E-09 | **6.08E-08** |
| ENSG00000113812.16 | **ACTR8** | 93973 | protein_coding | **455.803** | -0.673 | -0.203 | **0.627** | 8.79E-09 | **6.08E-08** |
| ENSG00000108826.18 | **MRPL27** | 51264 | protein_coding | **655.133** | -0.867 | -0.261 | **0.548** | 8.87E-09 | **6.13E-08** |

| | | | | | | | | | |
|---|---|---|---|---|---|---|---|---|---|
| ENSG00000164221.15 | **CCDC112** | 153733 | protein_coding | **93.222** | 1.424 | 0.429 | **2.684** | 8.90E-09 | **6.15E-08** |
| ENSG00000229299.3 | **ZBTB46-AS2** | | lncRNA | **15.486** | 2.796 | 0.842 | **6.945** | 8.91E-09 | **6.15E-08** |
| ENSG00000241288.9 | **LINC02614** | 101927056 | lncRNA | **20.712** | 2.608 | 0.785 | **6.095** | 9.00E-09 | **6.22E-08** |
| ENSG00000139624.16 | **CERS5** | 91012 | protein_coding | **859.665** | -0.788 | -0.237 | **0.579** | 9.10E-09 | **6.28E-08** |
| ENSG00000213139.9 | **CRYGS** | 1427 | protein_coding | **147.240** | 0.853 | 0.257 | **1.806** | 9.23E-09 | **6.37E-08** |
| ENSG00000105825.16 | **TFPI2** | 7980 | protein_coding | **61.216** | 1.853 | 0.558 | **3.612** | 9.26E-09 | **6.39E-08** |
| ENSG00000239900.16 | **ADSL** | 158 | protein_coding | **1441.644** | -0.705 | -0.212 | **0.613** | 9.29E-09 | **6.40E-08** |
| ENSG00000168772.12 | **CXXC4** | 80319 | protein_coding | **132.363** | 1.308 | 0.394 | **2.476** | 9.34E-09 | **6.43E-08** |
| ENSG00000141627.16 | **DYM** | 54808 | protein_coding | **526.673** | -0.749 | -0.226 | **0.595** | 9.39E-09 | **6.47E-08** |
| ENSG00000303840.1 | **ENSG00000303840** | | lncRNA | **12.094** | -3.598 | -1.083 | **0.083** | 9.41E-09 | **6.48E-08** |
| ENSG00000108469.17 | **RECQL5** | 9400 | protein_coding | **1310.209** | 0.732 | 0.220 | **1.661** | 9.51E-09 | **6.54E-08** |
| ENSG00000117394.26 | **SLC2A1** | 6513 | protein_coding | **1087.841** | -0.777 | -0.234 | **0.584** | 9.53E-09 | **6.56E-08** |
| ENSG00000159322.20 | **ADPGK** | 83440 | protein_coding | **1722.931** | -0.556 | -0.167 | **0.680** | 9.58E-09 | **6.59E-08** |
| ENSG00000102144.17 | **PGK1** | 5230 | protein_coding | **9598.482** | -0.819 | -0.247 | **0.567** | 9.59E-09 | **6.59E-08** |
| ENSG00000185236.14 | **RAB11B** | 9230 | protein_coding | **917.588** | -0.804 | -0.242 | **0.573** | 9.67E-09 | **6.64E-08** |
| ENSG00000170989.12 | **S1PR1** | 1901 | protein_coding | **401.292** | -0.902 | -0.272 | **0.535** | 9.72E-09 | **6.68E-08** |
| ENSG00000145016.19 | **RUBCN** | 9711 | protein_coding | **1800.277** | 0.741 | 0.223 | **1.671** | 9.74E-09 | **6.69E-08** |
| ENSG00000267364.5 | **ENSG00000267364** | | lncRNA | **27.031** | 2.146 | 0.646 | **4.425** | 9.79E-09 | **6.72E-08** |
| ENSG00000206417.9 | **H1-10-AS1** | | lncRNA | **123.798** | 1.510 | 0.455 | **2.848** | 9.87E-09 | **6.77E-08** |
| ENSG00000265752.4 | **LINC02958** | 107985181 | lncRNA | **16.349** | 2.805 | 0.844 | **6.986** | 9.90E-09 | **6.79E-08** |
| ENSG00000167595.18 | **PROSER3** | 148137 | protein_coding | **364.183** | -0.556 | -0.167 | **0.680** | 1.00E-08 | **6.86E-08** |
| ENSG00000066322.17 | **ELOVL1** | 64834 | protein_coding | **968.748** | -0.572 | -0.172 | **0.673** | 1.00E-08 | **6.86E-08** |
| ENSG00000165997.7 | **ARL5B** | 221079 | protein_coding | **173.483** | -0.982 | -0.296 | **0.506** | 1.01E-08 | **6.93E-08** |
| ENSG00000135837.20 | **CEP350** | 9857 | protein_coding | **542.379** | -0.554 | -0.167 | **0.681** | 1.01E-08 | **6.94E-08** |
| ENSG00000130396.24 | **AFDN** | 4301 | protein_coding | **897.170** | 1.153 | 0.347 | **2.224** | 1.02E-08 | **7.00E-08** |
| ENSG00000175984.18 | **DENND2C** | 163259 | protein_coding | **17.908** | 3.103 | 0.934 | **8.593** | 1.03E-08 | **7.03E-08** |
| ENSG00000227946.4 | **INO80D-AS1** | | lncRNA | **68.778** | -1.335 | -0.402 | **0.396** | 1.03E-08 | **7.04E-08** |
| ENSG00000165168.11 | **CYBB** | 1536 | protein_coding | **535.173** | -1.178 | -0.355 | **0.442** | 1.05E-08 | **7.21E-08** |
| ENSG00000261183.8 | **SPINT1-AS1** | | lncRNA | **35.619** | 1.572 | 0.473 | **2.972** | 1.06E-08 | **7.23E-08** |
| ENSG00000113318.13 | **MSH3** | 4437 | protein_coding | **205.475** | -0.769 | -0.231 | **0.587** | 1.07E-08 | **7.29E-08** |
| ENSG00000165678.24 | **GHITM** | 27069 | protein_coding | **1785.766** | -0.627 | -0.189 | **0.648** | 1.07E-08 | **7.33E-08** |

| | | | | | | | | | |
|---|---|---|---|---|---|---|---|---|---|
| ENSG00000122965.13 | **RBM19** | 9904 | protein_coding | **891.869** | -0.720 | -0.217 | **0.607** | 1.07E-08 | **7.33E-08** |
| ENSG00000167107.15 | **ACSF2** | 80221 | protein_coding | **423.930** | 0.760 | 0.229 | **1.693** | 1.08E-08 | **7.36E-08** |
| ENSG00000131471.8 | **AOC3** | 8639 | protein_coding | **88.781** | 1.366 | 0.411 | **2.578** | 1.08E-08 | **7.39E-08** |
| ENSG00000083312.21 | **TNPO1** | 3842 | protein_coding | **552.214** | -1.033 | -0.311 | **0.489** | 1.09E-08 | **7.47E-08** |
| ENSG00000249115.11 | **HAUS5** | 23354 | protein_coding | **708.727** | -0.587 | -0.177 | **0.666** | 1.10E-08 | **7.49E-08** |
| ENSG00000147383.13 | **NSDHL** | 50814 | protein_coding | **407.813** | -0.864 | -0.260 | **0.549** | 1.11E-08 | **7.54E-08** |
| ENSG00000099864.20 | **PALM** | 5064 | protein_coding | **32.093** | 2.854 | 0.859 | **7.228** | 1.13E-08 | **7.72E-08** |
| ENSG00000185187.16 | **SIGIRR** | 59307 | protein_coding | **295.078** | 0.909 | 0.274 | **1.878** | 1.13E-08 | **7.73E-08** |
| ENSG00000129595.15 | **EPB41L4A** | 64097 | protein_coding | **57.560** | 1.570 | 0.473 | **2.970** | 1.16E-08 | **7.88E-08** |
| ENSG00000162426.18 | **SLC45A1** | 50651 | protein_coding | **10.432** | 3.294 | 0.992 | **9.810** | 1.18E-08 | **8.05E-08** |
| ENSG00000127481.16 | **UBR4** | 23352 | protein_coding | **5953.149** | 0.506 | 0.152 | **1.421** | 1.19E-08 | **8.10E-08** |
| ENSG00000184743.15 | **ATL3** | 25923 | protein_coding | **549.713** | -0.539 | -0.162 | **0.688** | 1.19E-08 | **8.12E-08** |
| ENSG00000214944.12 | **ARHGEF28** | 64283 | protein_coding | **114.281** | 1.195 | 0.360 | **2.290** | 1.20E-08 | **8.16E-08** |
| ENSG00000111331.16 | **OAS3** | 4940 | protein_coding | **2526.526** | -0.942 | -0.284 | **0.521** | 1.20E-08 | **8.17E-08** |
| ENSG00000135521.11 | **LTV1** | 84946 | protein_coding | **452.266** | -0.596 | -0.180 | **0.661** | 1.21E-08 | **8.21E-08** |
| ENSG00000226853.6 | **GPR155-DT** | | lncRNA | **41.740** | 1.314 | 0.395 | **2.486** | 1.21E-08 | **8.21E-08** |
| ENSG00000181704.14 | **YIPF6** | 286451 | protein_coding | **230.577** | -0.940 | -0.283 | **0.521** | 1.23E-08 | **8.33E-08** |
| ENSG00000116062.21 | **MSH6** | 2956 | protein_coding | **868.892** | -0.555 | -0.167 | **0.681** | 1.23E-08 | **8.33E-08** |
| ENSG00000105464.5 | **GRIN2D** | 2906 | protein_coding | **23.696** | 2.253 | 0.678 | **4.767** | 1.26E-08 | **8.52E-08** |
| ENSG00000175727.17 | **MLXIP** | 22877 | protein_coding | **487.125** | 1.095 | 0.330 | **2.136** | 1.27E-08 | **8.60E-08** |
| ENSG00000135269.20 | **TES** | 26136 | protein_coding | **594.423** | -0.576 | -0.173 | **0.671** | 1.28E-08 | **8.67E-08** |
| ENSG00000134684.15 | **YARS1** | 8565 | protein_coding | **2815.569** | -0.841 | -0.253 | **0.558** | 1.28E-08 | **8.68E-08** |
| ENSG00000166166.15 | **TRMT61A** | 115708 | protein_coding | **303.097** | -0.680 | -0.205 | **0.624** | 1.28E-08 | **8.68E-08** |
| ENSG00000142303.16 | **ADAMTS1(** | 81794 | protein_coding | **15.056** | 3.673 | 1.106 | **12.758** | 1.28E-08 | **8.70E-08** |
| ENSG00000173457.13 | **PPP1R14B** | 26472 | protein_coding | **2159.405** | -0.947 | -0.285 | **0.519** | 1.29E-08 | **8.77E-08** |
| ENSG00000293486.2 | **ENSG00000293486** | | lncRNA | **92.909** | 1.245 | 0.375 | **2.371** | 1.30E-08 | **8.78E-08** |
| ENSG00000169231.15 | **THBS3** | 7059 | protein_coding | **511.333** | 1.099 | 0.331 | **2.142** | 1.31E-08 | **8.83E-08** |
| ENSG00000248015.7 | **NDUFS7-AS1** | | lncRNA | **48.956** | 1.519 | 0.457 | **2.865** | 1.31E-08 | **8.84E-08** |
| ENSG00000198055.13 | **GRK6** | 2870 | protein_coding | **1593.420** | -0.607 | -0.183 | **0.656** | 1.31E-08 | **8.89E-08** |
| ENSG00000188026.15 | **RILPL1** | 353116 | protein_coding | **95.518** | 1.642 | 0.494 | **3.121** | 1.32E-08 | **8.89E-08** |
| ENSG00000242259.11 | **C22orf39** | 128977 | protein_coding | **355.209** | -0.807 | -0.243 | **0.571** | 1.32E-08 | **8.92E-08** |

| | | | | | | | | | |
|---|---|---|---|---|---|---|---|---|---|
| ENSG00000137692.14 | **DCUN1D5** | 84259 | protein_coding | **385.576** | -0.930 | -0.280 | **0.525** | 1.34E-08 | **9.02E-08** |
| ENSG00000273783.3 | **GTF2A1-AS** | 10192850 | lncRNA | **25.694** | 2.017 | 0.607 | **4.046** | 1.34E-08 | **9.03E-08** |
| ENSG00000226777.10 | **FAM30A** | 9834 | lncRNA | **2399.165** | 0.894 | 0.269 | **1.859** | 1.34E-08 | **9.03E-08** |
| ENSG00000273001.2 | **PFKP-DT** | 10192782 | lncRNA | **102.670** | 1.137 | 0.342 | **2.199** | 1.37E-08 | **9.24E-08** |
| ENSG00000259494.4 | **MRPL46** | 26589 | protein_coding | **285.388** | -0.633 | -0.191 | **0.645** | 1.37E-08 | **9.27E-08** |
| ENSG00000120805.16 | **ARL1** | 400 | protein_coding | **794.033** | -0.666 | -0.201 | **0.630** | 1.40E-08 | **9.42E-08** |
| ENSG00000155957.20 | **TMBIM4** | 51643 | protein_coding | **374.792** | -0.786 | -0.237 | **0.580** | 1.41E-08 | **9.52E-08** |
| ENSG00000268916.8 | **CSAG3** | 389903 | protein_coding | **25.669** | 2.373 | 0.714 | **5.180** | 1.42E-08 | **9.54E-08** |
| ENSG00000287200.2 | **ENSG00000287200** | | lncRNA | **58.209** | 1.533 | 0.461 | **2.893** | 1.42E-08 | **9.55E-08** |
| ENSG00000125733.20 | **TRIP10** | 9322 | protein_coding | **2619.234** | 0.600 | 0.181 | **1.516** | 1.43E-08 | **9.60E-08** |
| ENSG00000166938.15 | **DIS3L** | 115752 | protein_coding | **322.580** | -0.756 | -0.228 | **0.592** | 1.43E-08 | **9.63E-08** |
| ENSG00000142166.17 | **IFNAR1** | 3454 | protein_coding | **573.310** | -0.595 | -0.179 | **0.662** | 1.43E-08 | **9.65E-08** |
| ENSG00000161395.15 | **PGAP3** | 93210 | protein_coding | **328.716** | 0.673 | 0.203 | **1.594** | 1.44E-08 | **9.67E-08** |
| ENSG00000131459.15 | **GFPT2** | 9945 | protein_coding | **35.346** | 1.606 | 0.483 | **3.043** | 1.44E-08 | **9.67E-08** |
| ENSG00000103047.10 | **TANGO6** | 79613 | protein_coding | **124.063** | -0.964 | -0.290 | **0.513** | 1.44E-08 | **9.69E-08** |
| ENSG00000124920.16 | **MYRF** | 745 | protein_coding | **471.656** | 0.837 | 0.252 | **1.787** | 1.45E-08 | **9.76E-08** |
| ENSG00000162191.16 | **UBXN1** | 51035 | protein_coding | **3038.908** | 0.555 | 0.167 | **1.470** | 1.45E-08 | **9.77E-08** |
| ENSG00000277258.7 | **PCGF2** | 7703 | protein_coding | **1068.623** | 1.150 | 0.346 | **2.219** | 1.46E-08 | **9.83E-08** |
| ENSG00000163319.13 | **MRPS18C** | 51023 | protein_coding | **312.338** | -0.735 | -0.221 | **0.601** | 1.47E-08 | **9.90E-08** |
| ENSG00000012124.19 | **CD22** | 933 | protein_coding | **9357.483** | -0.891 | -0.268 | **0.539** | 1.48E-08 | **9.93E-08** |
| ENSG00000081237.24 | **PTPRC** | 5788 | protein_coding | **876.757** | -0.752 | -0.227 | **0.594** | 1.48E-08 | **9.94E-08** |
| ENSG00000140030.8 | **GPR65** | 8477 | protein_coding | **133.448** | -0.957 | -0.288 | **0.515** | 1.48E-08 | **9.94E-08** |
| ENSG00000272668.3 | **ENSG0000(** | 12490443 | lncRNA | **11.878** | 3.864 | 1.163 | **14.560** | 1.50E-08 | **1.00E-07** |
| ENSG00000066651.22 | **TRMT11** | 60487 | protein_coding | **343.467** | -0.865 | -0.261 | **0.549** | 1.50E-08 | **1.00E-07** |
| ENSG00000298634.1 | **ENSG00000298634** | | lncRNA | **21.227** | 2.512 | 0.756 | **5.706** | 1.51E-08 | **1.01E-07** |
| ENSG00000224950.2 | **ENSG00000224950** | | lncRNA | **106.049** | -1.103 | -0.332 | **0.465** | 1.51E-08 | **1.01E-07** |
| ENSG00000126261.14 | **UBA2** | 10054 | protein_coding | **1228.068** | -0.729 | -0.220 | **0.603** | 1.52E-08 | **1.01E-07** |
| ENSG00000127125.11 | **PPCS** | 79717 | protein_coding | **670.543** | -0.542 | -0.163 | **0.687** | 1.52E-08 | **1.02E-07** |
| ENSG00000091136.17 | **LAMB1** | 3912 | protein_coding | **26.913** | 2.565 | 0.772 | **5.917** | 1.53E-08 | **1.02E-07** |
| ENSG00000112578.13 | **BYSL** | 705 | protein_coding | **307.881** | -0.746 | -0.225 | **0.596** | 1.53E-08 | **1.02E-07** |
| ENSG00000120820.15 | **GLT8D2** | 83468 | protein_coding | **37.245** | 1.618 | 0.487 | **3.069** | 1.55E-08 | **1.04E-07** |

| | | | | | | | | | |
|---|---|---|---|---|---|---|---|---|---|
| ENSG00000235531.12 | **MSC-AS1** | 100132891 | lncRNA | **412.639** | 0.760 | 0.229 | **1.693** | 1.57E-08 | **1.05E-07** |
| ENSG00000101224.20 | **CDC25B** | 994 | protein_coding | **1143.466** | -0.573 | -0.173 | **0.672** | 1.57E-08 | **1.05E-07** |
| ENSG00000228878.11 | **SEPTIN7-DT** | | lncRNA | **109.597** | 0.921 | 0.277 | **1.893** | 1.57E-08 | **1.05E-07** |
| ENSG00000172269.22 | **DPAGT1** | 1798 | protein_coding | **675.875** | -0.529 | -0.159 | **0.693** | 1.57E-08 | **1.05E-07** |
| ENSG00000179152.22 | **TCAIM** | 285343 | protein_coding | **243.570** | -0.664 | -0.200 | **0.631** | 1.57E-08 | **1.05E-07** |
| ENSG00000286610.2 | **ENSG00000286610** | | lncRNA | **15.606** | 2.369 | 0.713 | **5.166** | 1.59E-08 | **1.06E-07** |
| ENSG00000118898.18 | **PPL** | 5493 | protein_coding | **19.717** | 2.835 | 0.854 | **7.138** | 1.59E-08 | **1.06E-07** |
| ENSG00000108823.19 | **SGCA** | 6442 | protein_coding | **35.139** | 1.678 | 0.505 | **3.200** | 1.60E-08 | **1.07E-07** |
| ENSG00000105928.18 | **GSDME** | 1687 | protein_coding | **1607.446** | -0.779 | -0.234 | **0.583** | 1.60E-08 | **1.07E-07** |
| ENSG00000127445.16 | **PIN1** | 5300 | protein_coding | **770.308** | -0.543 | -0.163 | **0.686** | 1.61E-08 | **1.07E-07** |
| ENSG00000136147.20 | **PHF11** | 51131 | protein_coding | **1077.544** | -0.724 | -0.218 | **0.605** | 1.61E-08 | **1.07E-07** |
| ENSG00000136213.12 | **CHST12** | 55501 | protein_coding | **1102.007** | 0.980 | 0.295 | **1.972** | 1.61E-08 | **1.07E-07** |
| ENSG00000160932.13 | **LY6E** | 4061 | protein_coding | **4473.216** | -0.635 | -0.191 | **0.644** | 1.61E-08 | **1.07E-07** |
| ENSG00000122140.15 | **MRPS2** | 51116 | protein_coding | **961.146** | -0.608 | -0.183 | **0.656** | 1.63E-08 | **1.08E-07** |
| ENSG00000204498.12 | **NFKBIL1** | 4795 | protein_coding | **468.324** | 0.702 | 0.211 | **1.626** | 1.64E-08 | **1.09E-07** |
| ENSG00000171469.13 | **ZNF561** | 93134 | protein_coding | **229.105** | -0.743 | -0.224 | **0.598** | 1.65E-08 | **1.10E-07** |
| ENSG00000136731.15 | **UGGT1** | 56886 | protein_coding | **903.664** | -0.976 | -0.294 | **0.508** | 1.66E-08 | **1.10E-07** |
| ENSG00000163683.14 | **SMIM14** | 201895 | protein_coding | **1056.096** | 0.772 | 0.232 | **1.708** | 1.67E-08 | **1.11E-07** |
| ENSG00000136444.12 | **RSAD1** | 55316 | protein_coding | **683.499** | 0.572 | 0.172 | **1.487** | 1.69E-08 | **1.12E-07** |
| ENSG00000134352.23 | **IL6ST** | 3572 | protein_coding | **931.946** | 1.213 | 0.365 | **2.318** | 1.70E-08 | **1.13E-07** |
| ENSG00000198400.15 | **NTRK1** | 4914 | protein_coding | **10.892** | 4.226 | 1.272 | **18.720** | 1.72E-08 | **1.14E-07** |
| ENSG00000159648.13 | **SPMIP8** | 374739 | protein_coding | **12.049** | 2.587 | 0.779 | **6.010** | 1.73E-08 | **1.15E-07** |
| ENSG00000136933.19 | **RABEPK** | 10244 | protein_coding | **375.414** | -0.550 | -0.166 | **0.683** | 1.78E-08 | **1.18E-07** |
| ENSG00000183779.8 | **ZNF703** | 80139 | protein_coding | **26.518** | 2.139 | 0.644 | **4.404** | 1.80E-08 | **1.19E-07** |
| ENSG00000087157.21 | **PGS1** | 9489 | protein_coding | **974.645** | 0.667 | 0.201 | **1.588** | 1.82E-08 | **1.21E-07** |
| ENSG00000179163.14 | **FUCA1** | 2517 | protein_coding | **793.881** | 0.726 | 0.218 | **1.654** | 1.82E-08 | **1.21E-07** |
| ENSG00000142856.20 | **ITGB3BP** | 23421 | protein_coding | **296.587** | -0.948 | -0.285 | **0.518** | 1.82E-08 | **1.21E-07** |
| ENSG00000188505.7 | **NCCRP1** | 342897 | protein_coding | **9.597** | 4.205 | 1.266 | **18.449** | 1.84E-08 | **1.22E-07** |
| ENSG00000188177.17 | **ZC3H6** | 376940 | protein_coding | **195.017** | -1.007 | -0.303 | **0.498** | 1.84E-08 | **1.22E-07** |
| ENSG00000108557.22 | **RAI1** | 10743 | protein_coding | **602.702** | -0.575 | -0.173 | **0.671** | 1.84E-08 | **1.22E-07** |
| ENSG00000174446.14 | **SNAPC5** | 10302 | protein_coding | **258.574** | -0.816 | -0.246 | **0.568** | 1.85E-08 | **1.22E-07** |

| | | | | | | | | | |
|---|---|---|---|---|---|---|---|---|---|
| ENSG00000102871.18 | **TRADD** | 8717 | protein_coding | **1222.155** | 0.583 | 0.175 | **1.498** | 1.86E-08 | **1.23E-07** |
| ENSG00000213937.4 | **CLDN9** | 9080 | protein_coding | **9.374** | 3.929 | 1.183 | **15.232** | 1.86E-08 | **1.23E-07** |
| ENSG00000287126.1 | **ENSG0000(** | 12490772( | lncRNA | **80.108** | 0.996 | 0.300 | **1.995** | 1.86E-08 | **1.23E-07** |
| ENSG00000293256.2 | **ENSG00000293256** | | lncRNA | **41.900** | 1.312 | 0.395 | **2.483** | 1.87E-08 | **1.23E-07** |
| ENSG00000187951.11 | **ENSG0000(** | 100288637 | lncRNA | **239.275** | -1.455 | -0.438 | **0.365** | 1.88E-08 | **1.24E-07** |
| ENSG00000113597.20 | **TRAPPC13** | 80006 | protein_coding | **171.340** | -0.874 | -0.263 | **0.546** | 1.88E-08 | **1.24E-07** |
| ENSG00000279227.2 | **ENSG00000279227** | | TEC | **22.938** | 1.853 | 0.558 | **3.613** | 1.89E-08 | **1.25E-07** |
| ENSG00000092421.19 | **SEMA6A** | 57556 | protein_coding | **388.759** | 1.188 | 0.358 | **2.278** | 1.90E-08 | **1.26E-07** |
| ENSG00000127824.17 | **TUBA4A** | 7277 | protein_coding | **2614.859** | -0.652 | -0.196 | **0.636** | 1.90E-08 | **1.26E-07** |
| ENSG00000170854.20 | **RIOX2** | 84864 | protein_coding | **389.391** | -0.615 | -0.185 | **0.653** | 1.91E-08 | **1.26E-07** |
| ENSG00000151502.13 | **VPS26B** | 112936 | protein_coding | **623.792** | 0.565 | 0.170 | **1.480** | 1.93E-08 | **1.27E-07** |
| ENSG00000300559.1 | **ENSG00000300559** | | lncRNA | **12.334** | 3.290 | 0.990 | **9.782** | 1.93E-08 | **1.27E-07** |
| ENSG00000177879.19 | **AP3S1** | 1176 | protein_coding | **1651.426** | -0.776 | -0.234 | **0.584** | 1.94E-08 | **1.28E-07** |
| ENSG00000011376.14 | **LARS2** | 23395 | protein_coding | **319.783** | -0.703 | -0.212 | **0.614** | 1.94E-08 | **1.28E-07** |
| ENSG00000183570.19 | **PCBP3** | 54039 | protein_coding | **39.822** | 1.915 | 0.577 | **3.772** | 1.95E-08 | **1.28E-07** |
| ENSG00000288771.3 | **ENSG00000288771** | | lncRNA | **44.109** | -2.021 | -0.608 | **0.246** | 1.96E-08 | **1.29E-07** |
| ENSG00000124334.21 | **IL9R** | 3581 | protein_coding | **96.231** | -1.219 | -0.367 | **0.430** | 1.99E-08 | **1.31E-07** |
| ENSG00000112697.19 | **TMEM30A** | 55754 | protein_coding | **1298.383** | 0.501 | 0.151 | **1.415** | 2.02E-08 | **1.33E-07** |
| ENSG00000059377.21 | **TBXAS1** | 6916 | protein_coding | **160.558** | -0.999 | -0.301 | **0.500** | 2.02E-08 | **1.33E-07** |
| ENSG00000254718.7 | **ENSG0000(** | 10192770‡ | lncRNA | **46.894** | 1.742 | 0.524 | **3.346** | 2.05E-08 | **1.35E-07** |
| ENSG00000297975.1 | **ENSG00000297975** | | lncRNA | **66.171** | -1.124 | -0.338 | **0.459** | 2.08E-08 | **1.36E-07** |
| ENSG00000171700.16 | **RGS19** | 10287 | protein_coding | **726.095** | -0.716 | -0.215 | **0.609** | 2.08E-08 | **1.36E-07** |
| ENSG00000113645.17 | **WWC1** | 23286 | protein_coding | **46.389** | 2.208 | 0.665 | **4.621** | 2.08E-08 | **1.37E-07** |
| ENSG00000099800.10 | **TIMM13** | 26517 | protein_coding | **883.579** | -0.548 | -0.165 | **0.684** | 2.08E-08 | **1.37E-07** |
| ENSG00000139697.16 | **SBNO1** | 55206 | protein_coding | **1000.540** | 0.716 | 0.216 | **1.643** | 2.08E-08 | **1.37E-07** |
| ENSG00000166352.20 | **IFTAP** | 119710 | protein_coding | **81.650** | 1.290 | 0.388 | **2.446** | 2.10E-08 | **1.38E-07** |
| ENSG00000114767.9 | **RRP9** | 9136 | protein_coding | **634.868** | -0.720 | -0.217 | **0.607** | 2.11E-08 | **1.38E-07** |
| ENSG00000167771.8 | **RCOR2** | 283248 | protein_coding | **65.309** | 1.725 | 0.519 | **3.306** | 2.11E-08 | **1.38E-07** |
| ENSG00000124193.18 | **SRSF6** | 6431 | protein_coding | **2238.554** | -0.557 | -0.168 | **0.680** | 2.11E-08 | **1.38E-07** |
| ENSG00000141750.9 | **STAC2** | 342667 | protein_coding | **8.827** | 3.990 | 1.201 | **15.893** | 2.14E-08 | **1.40E-07** |
| ENSG00000204301.8 | **NOTCH4** | 4855 | protein_coding | **51.870** | -1.324 | -0.399 | **0.399** | 2.14E-08 | **1.40E-07** |

| | | | | | | | | | |
|---|---|---|---|---|---|---|---|---|---|
| ENSG00000185621.13 | **LMLN** | 89782 | protein_coding | **72.750** | 1.223 | 0.368 | **2.334** | 2.14E-08 | **1.40E-07** |
| ENSG00000188659.11 | **SAXO2** | 283726 | protein_coding | **32.775** | 2.162 | 0.651 | **4.475** | 2.15E-08 | **1.41E-07** |
| ENSG00000243155.1 | **ENSG00000243155** | | lncRNA | **57.580** | 1.258 | 0.379 | **2.391** | 2.16E-08 | **1.41E-07** |
| ENSG00000075131.12 | **TIPIN** | 54962 | protein_coding | **201.705** | -0.811 | -0.244 | **0.570** | 2.18E-08 | **1.42E-07** |
| ENSG00000164182.14 | **NDUFAF2** | 91942 | protein_coding | **366.773** | -0.713 | -0.215 | **0.610** | 2.19E-08 | **1.43E-07** |
| ENSG00000307459.1 | **ENSG00000307459** | | lncRNA | **10.569** | 4.488 | 1.351 | **22.446** | 2.22E-08 | **1.45E-07** |
| ENSG00000122557.13 | **HERPUD2** | 64224 | protein_coding | **498.046** | 0.588 | 0.177 | **1.503** | 2.22E-08 | **1.45E-07** |
| ENSG00000105063.21 | **PPP6R1** | 22870 | protein_coding | **3174.376** | -0.564 | -0.170 | **0.676** | 2.24E-08 | **1.46E-07** |
| ENSG00000129071.12 | **MBD4** | 8930 | protein_coding | **567.395** | -0.859 | -0.259 | **0.551** | 2.24E-08 | **1.46E-07** |
| ENSG00000141569.14 | **TRIM65** | 201292 | protein_coding | **397.029** | -0.604 | -0.182 | **0.658** | 2.27E-08 | **1.48E-07** |
| ENSG00000156171.17 | **DRAM2** | 128338 | protein_coding | **1617.576** | -0.566 | -0.170 | **0.676** | 2.28E-08 | **1.49E-07** |
| ENSG00000072849.13 | **DERL2** | 51009 | protein_coding | **712.383** | -0.597 | -0.180 | **0.661** | 2.28E-08 | **1.49E-07** |
| ENSG00000105737.12 | **GRIK5** | 2901 | protein_coding | **22.343** | 2.316 | 0.697 | **4.980** | 2.30E-08 | **1.50E-07** |
| ENSG00000303830.1 | **ENSG00000303830** | | lncRNA | **22.347** | -2.331 | -0.702 | **0.199** | 2.33E-08 | **1.52E-07** |
| ENSG00000104064.20 | **GABPB1** | 2553 | protein_coding | **299.454** | -0.943 | -0.284 | **0.520** | 2.33E-08 | **1.52E-07** |
| ENSG00000132801.9 | **ZSWIM3** | 140831 | protein_coding | **86.906** | -1.180 | -0.355 | **0.441** | 2.33E-08 | **1.52E-07** |
| ENSG00000184220.13 | **CMSS1** | 84319 | protein_coding | **440.573** | -0.914 | -0.275 | **0.531** | 2.34E-08 | **1.52E-07** |
| ENSG00000124243.21 | **BCAS4** | 55653 | protein_coding | **959.995** | -0.801 | -0.241 | **0.574** | 2.34E-08 | **1.52E-07** |
| ENSG00000125877.15 | **ITPA** | 3704 | protein_coding | **846.974** | -0.720 | -0.217 | **0.607** | 2.36E-08 | **1.53E-07** |
| ENSG00000062582.16 | **MRPS24** | 64951 | protein_coding | **739.549** | -0.675 | -0.203 | **0.626** | 2.39E-08 | **1.55E-07** |
| ENSG00000106123.15 | **EPHB6** | 2051 | protein_coding | **34.699** | 2.503 | 0.754 | **5.670** | 2.39E-08 | **1.55E-07** |
| ENSG00000234807.10 | **JUN-DT** | | lncRNA | **120.794** | 0.997 | 0.300 | **1.995** | 2.40E-08 | **1.56E-07** |
| ENSG00000075618.20 | **FSCN1** | 6624 | protein_coding | **8488.951** | 1.736 | 0.523 | **3.331** | 2.40E-08 | **1.56E-07** |
| ENSG00000099331.16 | **MYO9B** | 4650 | protein_coding | **5611.897** | 0.517 | 0.156 | **1.431** | 2.41E-08 | **1.56E-07** |
| ENSG00000027644.6 | **INSRR** | 3645 | protein_coding | **7.616** | 4.289 | 1.291 | **19.546** | 2.43E-08 | **1.57E-07** |
| ENSG00000120685.22 | **PROSER1** | 80209 | protein_coding | **1035.479** | -0.566 | -0.170 | **0.675** | 2.44E-08 | **1.58E-07** |
| ENSG00000132185.20 | **FCRLA** | 84824 | protein_coding | **2499.734** | -0.948 | -0.285 | **0.518** | 2.45E-08 | **1.59E-07** |
| ENSG00000258572.2 | **SYNE3-AS1** | 10192910[illegible] | lncRNA | **61.350** | -1.178 | -0.355 | **0.442** | 2.48E-08 | **1.60E-07** |
| ENSG00000197283.18 | **SYNGAP1** | 8831 | protein_coding | **876.622** | 1.001 | 0.301 | **2.002** | 2.48E-08 | **1.61E-07** |
| ENSG00000298216.1 | **ENSG00000298216** | | lncRNA | **29.610** | -1.744 | -0.525 | **0.299** | 2.49E-08 | **1.61E-07** |
| ENSG00000280202.1 | **ENSG00000280202** | | TEC | **22.804** | -1.683 | -0.507 | **0.312** | 2.50E-08 | **1.62E-07** |

| | | | | | | | | | |
|---|---|---|---|---|---|---|---|---|---|
| ENSG00000117593.14 | **DARS2** | 55157 | protein_coding | **408.452** | -0.652 | -0.196 | **0.637** | 2.56E-08 | **1.65E-07** |
| ENSG00000261408.7 | **TEN1-CDK3** | 100529145 | protein_coding | **43.878** | -1.402 | -0.422 | **0.378** | 2.56E-08 | **1.66E-07** |
| ENSG00000110171.22 | **TRIM3** | 10612 | protein_coding | **233.781** | 0.944 | 0.284 | **1.923** | 2.57E-08 | **1.66E-07** |
| ENSG00000101216.13 | **GMEB2** | 26205 | protein_coding | **458.936** | -0.543 | -0.163 | **0.686** | 2.59E-08 | **1.67E-07** |
| ENSG00000270504.2 | **ENSG00000270504** | | lncRNA | **21.054** | 2.448 | 0.737 | **5.457** | 2.62E-08 | **1.69E-07** |
| ENSG00000214655.12 | **ZSWIM8** | 23053 | protein_coding | **1886.512** | 0.637 | 0.192 | **1.555** | 2.62E-08 | **1.69E-07** |
| ENSG00000164031.19 | **DNAJB14** | 79982 | protein_coding | **578.328** | 0.878 | 0.264 | **1.838** | 2.67E-08 | **1.73E-07** |
| ENSG00000136870.12 | **ZNF189** | 7743 | protein_coding | **263.258** | -0.704 | -0.212 | **0.614** | 2.68E-08 | **1.73E-07** |
| ENSG00000189091.15 | **SF3B3** | 23450 | protein_coding | **2433.707** | -0.625 | -0.188 | **0.648** | 2.69E-08 | **1.73E-07** |
| ENSG00000197948.13 | **FCHSD1** | 89848 | protein_coding | **1279.879** | 0.566 | 0.170 | **1.481** | 2.69E-08 | **1.73E-07** |
| ENSG00000167920.13 | **KRT10-AS1** | 147184 | lncRNA | **203.590** | 0.665 | 0.200 | **1.586** | 2.72E-08 | **1.75E-07** |
| ENSG00000231789.5 | **PIK3CD-AS** | 101929074 | lncRNA | **374.734** | 0.883 | 0.266 | **1.844** | 2.73E-08 | **1.76E-07** |
| ENSG00000268069.2 | **ENSG00000268069** | | lncRNA | **13.069** | 2.966 | 0.893 | **7.812** | 2.74E-08 | **1.77E-07** |
| ENSG00000103064.17 | **SLC7A6** | 9057 | protein_coding | **1018.801** | -0.566 | -0.171 | **0.675** | 2.77E-08 | **1.78E-07** |
| ENSG00000291003.2 | **ENSG00000291003** | | lncRNA | **1821.091** | 0.674 | 0.203 | **1.595** | 2.77E-08 | **1.78E-07** |
| ENSG00000099949.24 | **LZTR1** | 8216 | protein_coding | **1250.625** | 0.543 | 0.163 | **1.457** | 2.77E-08 | **1.78E-07** |
| ENSG00000182566.16 | **CLEC4G** | 339390 | protein_coding | **63.248** | 1.749 | 0.526 | **3.361** | 2.78E-08 | **1.79E-07** |
| ENSG00000122188.15 | **LAX1** | 54900 | protein_coding | **180.684** | -1.305 | -0.393 | **0.405** | 2.80E-08 | **1.80E-07** |
| ENSG00000185624.18 | **P4HB** | 5034 | protein_coding | **9600.794** | -0.809 | -0.244 | **0.571** | 2.80E-08 | **1.80E-07** |
| ENSG00000223891.8 | **OSER1-DT** | 100505783 | lncRNA | **105.350** | 0.944 | 0.284 | **1.924** | 2.81E-08 | **1.81E-07** |
| ENSG00000277400.1 | **ENSG00000277400** | | protein_coding | **192.466** | 1.112 | 0.335 | **2.162** | 2.81E-08 | **1.81E-07** |
| ENSG00000163428.6 | **LRRC58** | 116064 | protein_coding | **354.403** | -0.627 | -0.189 | **0.648** | 2.82E-08 | **1.81E-07** |
| ENSG00000273287.3 | **ENSG00000273287** | | lncRNA | **9.669** | 3.193 | 0.961 | **9.145** | 2.82E-08 | **1.81E-07** |
| ENSG00000141956.15 | **PRDM15** | 63977 | protein_coding | **658.036** | -0.734 | -0.221 | **0.601** | 2.83E-08 | **1.82E-07** |
| ENSG00000034510.8 | **TMSB10** | 9168 | protein_coding | **33674.270** | 0.756 | 0.227 | **1.688** | 2.84E-08 | **1.82E-07** |
| ENSG00000180879.16 | **SSR4** | 6748 | protein_coding | **3683.730** | -0.737 | -0.222 | **0.600** | 2.85E-08 | **1.83E-07** |
| ENSG00000117090.18 | **SLAMF1** | 6504 | protein_coding | **3550.953** | -0.664 | -0.200 | **0.631** | 2.86E-08 | **1.84E-07** |
| ENSG00000139722.9 | **VPS37B** | 79720 | protein_coding | **1128.480** | 0.727 | 0.219 | **1.656** | 2.87E-08 | **1.84E-07** |
| ENSG00000171314.11 | **PGAM1** | 5223 | protein_coding | **10153.154** | -0.754 | -0.227 | **0.593** | 2.87E-08 | **1.84E-07** |
| ENSG00000163154.8 | **TNFAIP8L2** | 79626 | protein_coding | **232.493** | -0.896 | -0.270 | **0.538** | 2.87E-08 | **1.84E-07** |
| ENSG00000287165.2 | **ENSG00000287165** | | lncRNA | **25.247** | 1.740 | 0.524 | **3.341** | 2.89E-08 | **1.85E-07** |

| | | | | | | | | | |
|---|---|---|---|---|---|---|---|---|---|
| ENSG00000172175.18 | **MALT1** | 10892 | protein_coding | **1458.659** | -0.532 | -0.160 | **0.692** | 2.89E-08 | **1.85E-07** |
| ENSG00000168701.20 | **TMEM208** | 29100 | protein_coding | **528.954** | -0.637 | -0.192 | **0.643** | 2.92E-08 | **1.87E-07** |
| ENSG00000010361.16 | **FUZ** | 80199 | protein_coding | **270.181** | 0.757 | 0.228 | **1.690** | 2.92E-08 | **1.87E-07** |
| ENSG00000293945.1 | **ENSG00000293945** | | lncRNA | **42.456** | 1.669 | 0.502 | **3.180** | 2.92E-08 | **1.87E-07** |
| ENSG00000160818.18 | **GPATCH4** | 54865 | protein_coding | **321.443** | -0.999 | -0.301 | **0.500** | 2.93E-08 | **1.87E-07** |
| ENSG00000288852.2 | **ENSG00000288852** | | lncRNA | **70.110** | 1.060 | 0.319 | **2.085** | 2.95E-08 | **1.88E-07** |
| ENSG00000214456.10 | **PLIN5** | 440503 | protein_coding | **16.740** | 2.732 | 0.822 | **6.642** | 2.96E-08 | **1.89E-07** |
| ENSG00000185272.16 | **RBM11** | 54033 | protein_coding | **14.362** | 2.640 | 0.795 | **6.233** | 2.96E-08 | **1.89E-07** |
| ENSG00000100429.20 | **HDAC10** | 83933 | protein_coding | **1481.584** | 0.520 | 0.156 | **1.434** | 2.96E-08 | **1.89E-07** |
| ENSG00000150627.18 | **WDR17** | 116966 | protein_coding | **41.214** | 1.838 | 0.553 | **3.575** | 2.98E-08 | **1.90E-07** |
| ENSG00000174151.17 | **CYB561D1** | 284613 | protein_coding | **278.270** | 0.655 | 0.197 | **1.575** | 2.99E-08 | **1.91E-07** |
| ENSG00000198003.14 | **ODAD3** | 115948 | protein_coding | **263.300** | 1.542 | 0.464 | **2.912** | 3.00E-08 | **1.91E-07** |
| ENSG00000204387.15 | **SNHG32** | 50854 | lncRNA | **3147.457** | -0.864 | -0.260 | **0.549** | 3.00E-08 | **1.91E-07** |
| ENSG00000167291.18 | **TBC1D16** | 125058 | protein_coding | **71.089** | 1.962 | 0.590 | **3.895** | 3.00E-08 | **1.91E-07** |
| ENSG00000176884.18 | **GRIN1** | 2902 | protein_coding | **96.731** | 3.153 | 0.949 | **8.897** | 3.00E-08 | **1.91E-07** |
| ENSG00000279186.1 | **ENSG00000279186** | | TEC | **12.322** | 3.907 | 1.176 | **15.000** | 3.01E-08 | **1.92E-07** |
| ENSG00000239775.1 | **ENSG00000239775** | | lncRNA | **31.098** | 1.906 | 0.574 | **3.748** | 3.03E-08 | **1.93E-07** |
| ENSG00000101624.13 | **CEP76** | 79959 | protein_coding | **96.205** | -0.960 | -0.289 | **0.514** | 3.04E-08 | **1.94E-07** |
| ENSG00000249494.7 | **DMXL1-DT** | | lncRNA | **23.278** | 2.210 | 0.665 | **4.627** | 3.05E-08 | **1.94E-07** |
| ENSG00000106049.11 | **HIBADH** | 11112 | protein_coding | **423.241** | 0.562 | 0.169 | **1.476** | 3.06E-08 | **1.95E-07** |
| ENSG00000196821.12 | **ILRUN** | 64771 | protein_coding | **2315.125** | -0.578 | -0.174 | **0.670** | 3.07E-08 | **1.95E-07** |
| ENSG00000013725.16 | **CD6** | 923 | protein_coding | **38.612** | 2.127 | 0.640 | **4.368** | 3.08E-08 | **1.96E-07** |
| ENSG00000302470.1 | **ENSG00000302470** | | lncRNA | **14.517** | 3.868 | 1.164 | **14.598** | 3.08E-08 | **1.96E-07** |
| ENSG00000178685.16 | **PARP10** | 84875 | protein_coding | **2130.327** | -0.736 | -0.221 | **0.600** | 3.08E-08 | **1.96E-07** |
| ENSG00000131368.11 | **MRPS25** | 64432 | protein_coding | **1708.923** | -0.570 | -0.172 | **0.673** | 3.09E-08 | **1.96E-07** |
| ENSG00000197822.14 | **OCLN** | 10050665 | protein_coding | **171.148** | 1.136 | 0.342 | **2.198** | 3.09E-08 | **1.96E-07** |
| ENSG00000184014.11 | **DENND5A** | 23258 | protein_coding | **1076.645** | 1.035 | 0.312 | **2.049** | 3.12E-08 | **1.98E-07** |
| ENSG00000119402.20 | **FBXW2** | 26190 | protein_coding | **742.219** | -0.502 | -0.151 | **0.706** | 3.12E-08 | **1.98E-07** |
| ENSG00000231133.9 | **HAR1B** | 768097 | lncRNA | **11.684** | 3.015 | 0.908 | **8.082** | 3.17E-08 | **2.01E-07** |
| ENSG00000169758.15 | **TMEM266** | 123591 | protein_coding | **44.383** | 1.567 | 0.472 | **2.962** | 3.20E-08 | **2.03E-07** |
| ENSG00000259065.4 | **MIDEAS-A** | 10050647 | lncRNA | **109.224** | 0.896 | 0.270 | **1.860** | 3.20E-08 | **2.03E-07** |

| | | | | | | | | | |
|---|---|---|---|---|---|---|---|---|---|
| ENSG00000302611.1 | **ENSG00000302611** | | lncRNA | **34.266** | 1.744 | 0.525 | **3.349** | 3.25E-08 | **2.06E-07** |
| ENSG00000173535.17 | **TNFRSF10(** | 8794 | protein_coding | **36.585** | -1.878 | -0.565 | **0.272** | 3.26E-08 | **2.06E-07** |
| ENSG00000023445.19 | **BIRC3** | 330 | protein_coding | **3530.087** | 0.994 | 0.299 | **1.992** | 3.28E-08 | **2.08E-07** |
| ENSG00000169189.19 | **NSMCE1** | 197370 | protein_coding | **1419.424** | 0.809 | 0.243 | **1.752** | 3.30E-08 | **2.09E-07** |
| ENSG00000120694.22 | **HSPH1** | 10808 | protein_coding | **755.472** | -0.944 | -0.284 | **0.520** | 3.30E-08 | **2.09E-07** |
| ENSG00000069956.16 | **MAPK6** | 5597 | protein_coding | **550.172** | -0.796 | -0.240 | **0.576** | 3.31E-08 | **2.10E-07** |
| ENSG00000188629.16 | **ZNF177** | 7730 | protein_coding | **78.975** | 1.367 | 0.411 | **2.579** | 3.36E-08 | **2.12E-07** |
| ENSG00000267532.7 | **MIR497HG** | 10050671: | lncRNA | **213.009** | 0.796 | 0.240 | **1.736** | 3.38E-08 | **2.14E-07** |
| ENSG00000115297.11 | **TLX2** | 3196 | protein_coding | **14.242** | 3.737 | 1.125 | **13.336** | 3.41E-08 | **2.15E-07** |
| ENSG00000140836.18 | **ZFHX3** | 463 | protein_coding | **173.737** | 0.847 | 0.255 | **1.799** | 3.42E-08 | **2.16E-07** |
| ENSG00000100416.17 | **TRMU** | 55687 | protein_coding | **1153.096** | -0.645 | -0.194 | **0.640** | 3.43E-08 | **2.17E-07** |
| ENSG00000100065.17 | **CARD10** | 29775 | protein_coding | **6.923** | 4.940 | 1.487 | **30.700** | 3.44E-08 | **2.17E-07** |
| ENSG00000182575.9 | **NXPH3** | 11248 | protein_coding | **22.105** | 2.182 | 0.657 | **4.536** | 3.44E-08 | **2.17E-07** |
| ENSG00000244509.6 | **APOBEC3C** | 27350 | protein_coding | **2590.880** | -0.827 | -0.249 | **0.564** | 3.45E-08 | **2.18E-07** |
| ENSG00000112576.14 | **CCND3** | 896 | protein_coding | **917.545** | -0.597 | -0.180 | **0.661** | 3.46E-08 | **2.18E-07** |
| ENSG00000285230.4 | **RALY-AS1** | 10192688 | lncRNA | **131.832** | 1.079 | 0.325 | **2.112** | 3.47E-08 | **2.19E-07** |
| ENSG00000308692.1 | **ENSG00000308692** | | lncRNA | **79.301** | 2.144 | 0.645 | **4.420** | 3.48E-08 | **2.19E-07** |
| ENSG00000101109.16 | **STK4** | 6789 | protein_coding | **2567.316** | 0.782 | 0.235 | **1.720** | 3.54E-08 | **2.23E-07** |
| ENSG00000113360.19 | **DROSHA** | 29102 | protein_coding | **811.996** | -0.636 | -0.191 | **0.643** | 3.55E-08 | **2.23E-07** |
| ENSG00000164849.12 | **GPR146** | 115330 | protein_coding | **253.579** | -0.932 | -0.281 | **0.524** | 3.56E-08 | **2.24E-07** |
| ENSG00000255458.10 | **SMIM14-D** | 10537441! | lncRNA | **36.427** | 2.593 | 0.781 | **6.035** | 3.58E-08 | **2.25E-07** |
| ENSG00000157778.11 | **PSMG3** | 84262 | protein_coding | **708.995** | -0.718 | -0.216 | **0.608** | 3.59E-08 | **2.26E-07** |
| ENSG00000273611.7 | **ZNHIT3** | 9326 | protein_coding | **567.802** | -0.457 | -0.138 | **0.728** | 3.61E-08 | **2.27E-07** |
| ENSG00000261504.1 | **LINC01686** | 284648 | lncRNA | **23.693** | 1.708 | 0.514 | **3.267** | 3.61E-08 | **2.27E-07** |
| ENSG00000272686.2 | **WASL-DT** | | lncRNA | **63.199** | 1.207 | 0.363 | **2.308** | 3.61E-08 | **2.27E-07** |
| ENSG00000268516.5 | **ZNF8-DT** | 10537248: | lncRNA | **60.472** | -1.919 | -0.578 | **0.264** | 3.64E-08 | **2.29E-07** |
| ENSG00000168350.9 | **DEGS2** | 123099 | protein_coding | **28.341** | 1.860 | 0.560 | **3.630** | 3.65E-08 | **2.29E-07** |
| ENSG00000137497.21 | **NUMA1** | 4926 | protein_coding | **3405.416** | -0.650 | -0.196 | **0.637** | 3.66E-08 | **2.30E-07** |
| ENSG00000091640.10 | **SPAG7** | 9552 | protein_coding | **1115.437** | -0.509 | -0.153 | **0.703** | 3.66E-08 | **2.30E-07** |
| ENSG00000148344.12 | **PTGES** | 9536 | protein_coding | **12.250** | 2.830 | 0.852 | **7.111** | 3.67E-08 | **2.30E-07** |
| ENSG00000228649.12 | **SNHG26** | | lncRNA | **276.935** | -0.785 | -0.236 | **0.581** | 3.73E-08 | **2.34E-07** |

| | | | | | | | | | |
|---|---|---|---|---|---|---|---|---|---|
| ENSG00000120963.14 | **ZNF706** | 51123 | protein_coding | **1288.223** | -0.737 | -0.222 | **0.600** | 3.76E-08 | **2.36E-07** |
| ENSG00000014914.23 | **MTMR11** | 10903 | protein_coding | **31.188** | 1.707 | 0.514 | **3.264** | 3.81E-08 | **2.39E-07** |
| ENSG00000073536.20 | **NLE1** | 54475 | protein_coding | **304.705** | -0.875 | -0.263 | **0.545** | 3.81E-08 | **2.39E-07** |
| ENSG00000106153.16 | **CHCHD2** | 51142 | protein_coding | **5011.417** | -0.597 | -0.180 | **0.661** | 3.81E-08 | **2.39E-07** |
| ENSG00000290387.2 | **SORD2P** | 653381 | lncRNA | **130.935** | -1.097 | -0.330 | **0.467** | 3.89E-08 | **2.44E-07** |
| ENSG00000165899.12 | **OTOGL** | 283310 | protein_coding | **13.078** | 3.224 | 0.971 | **9.344** | 3.91E-08 | **2.45E-07** |
| ENSG00000164649.22 | **CDCA7L** | 55536 | protein_coding | **723.226** | 0.576 | 0.173 | **1.491** | 3.92E-08 | **2.45E-07** |
| ENSG00000105374.12 | **NKG7** | 4818 | protein_coding | **33.105** | -1.901 | -0.572 | **0.268** | 3.93E-08 | **2.46E-07** |
| ENSG00000101846.11 | **STS** | 412 | protein_coding | **112.282** | -1.359 | -0.409 | **0.390** | 3.95E-08 | **2.47E-07** |
| ENSG00000181408.7 | **UTS2R** | 2837 | protein_coding | **84.888** | 1.513 | 0.455 | **2.854** | 3.96E-08 | **2.48E-07** |
| ENSG00000035687.11 | **ADSS2** | 159 | protein_coding | **427.121** | -0.765 | -0.230 | **0.588** | 3.97E-08 | **2.48E-07** |
| ENSG00000198586.16 | **TLK1** | 9874 | protein_coding | **980.695** | -0.743 | -0.224 | **0.598** | 4.00E-08 | **2.50E-07** |
| ENSG00000131016.19 | **AKAP12** | 9590 | protein_coding | **24.649** | 3.111 | 0.937 | **8.642** | 4.00E-08 | **2.50E-07** |
| ENSG00000273008.2 | **CSGALNAC** | 118568825 | lncRNA | **35.025** | 1.395 | 0.420 | **2.629** | 4.01E-08 | **2.50E-07** |
| ENSG00000166411.16 | **IDH3A** | 3419 | protein_coding | **1079.891** | -0.574 | -0.173 | **0.672** | 4.04E-08 | **2.52E-07** |
| ENSG00000131094.6 | **C1QL1** | 10882 | protein_coding | **8.601** | 3.851 | 1.159 | **14.428** | 4.05E-08 | **2.53E-07** |
| ENSG00000108829.12 | **LRRC59** | 55379 | protein_coding | **905.958** | -0.718 | -0.216 | **0.608** | 4.09E-08 | **2.55E-07** |
| ENSG00000110446.14 | **SLC15A3** | 51296 | protein_coding | **707.020** | -1.055 | -0.318 | **0.481** | 4.16E-08 | **2.60E-07** |
| ENSG00000295453.1 | **ENSG00000295453** | | lncRNA | **63.088** | 1.239 | 0.373 | **2.360** | 4.16E-08 | **2.60E-07** |
| ENSG00000116209.14 | **TMEM59** | 9528 | protein_coding | **709.641** | -0.620 | -0.187 | **0.651** | 4.18E-08 | **2.60E-07** |
| ENSG00000128487.21 | **SPECC1** | 92521 | protein_coding | **389.674** | -0.692 | -0.208 | **0.619** | 4.18E-08 | **2.61E-07** |
| ENSG00000272112.1 | **ENSG00000272112** | | lncRNA | **24.513** | 1.934 | 0.582 | **3.822** | 4.21E-08 | **2.62E-07** |
| ENSG00000254966.2 | **IGSF22-AS1** | | lncRNA | **20.934** | 2.118 | 0.638 | **4.341** | 4.23E-08 | **2.64E-07** |
| ENSG00000155893.15 | **PXYLP1** | 92370 | protein_coding | **25.061** | 1.772 | 0.534 | **3.416** | 4.25E-08 | **2.65E-07** |
| ENSG00000205208.7 | **C4orf46** | 201725 | protein_coding | **294.735** | -0.756 | -0.227 | **0.592** | 4.27E-08 | **2.66E-07** |
| ENSG00000155755.21 | **TMEM237** | 65062 | protein_coding | **91.975** | -0.986 | -0.297 | **0.505** | 4.30E-08 | **2.68E-07** |
| ENSG00000164953.18 | **TMEM67** | 91147 | protein_coding | **78.175** | 0.907 | 0.273 | **1.875** | 4.30E-08 | **2.68E-07** |
| ENSG00000204410.17 | **MSH5** | 4439 | protein_coding | **897.581** | -0.775 | -0.233 | **0.584** | 4.33E-08 | **2.69E-07** |
| ENSG00000087008.18 | **ACOX3** | 8310 | protein_coding | **347.209** | -0.680 | -0.205 | **0.624** | 4.34E-08 | **2.69E-07** |
| ENSG00000185163.12 | **DDX51** | 317781 | protein_coding | **824.946** | -0.724 | -0.218 | **0.606** | 4.34E-08 | **2.69E-07** |
| ENSG00000099810.23 | **MTAP** | 4507 | protein_coding | **358.623** | -0.679 | -0.204 | **0.625** | 4.39E-08 | **2.72E-07** |

| | | | | | | | | | |
|---|---|---|---|---|---|---|---|---|---|
| ENSG00000081665.16 | **ZNF506** | 440515 | protein_coding | **534.082** | 0.626 | 0.188 | **1.543** | 4.39E-08 | **2.72E-07** |
| ENSG00000169398.22 | **PTK2** | 5747 | protein_coding | **308.673** | 1.067 | 0.321 | **2.095** | 4.40E-08 | **2.73E-07** |
| ENSG00000163872.18 | **YEATS2** | 55689 | protein_coding | **1200.954** | -0.464 | -0.140 | **0.725** | 4.41E-08 | **2.73E-07** |
| ENSG00000133606.13 | **MKRN1** | 23608 | protein_coding | **1833.869** | 0.542 | 0.163 | **1.456** | 4.43E-08 | **2.75E-07** |
| ENSG00000176473.16 | **WDR25** | 79446 | protein_coding | **146.526** | 0.878 | 0.264 | **1.838** | 4.44E-08 | **2.75E-07** |
| ENSG00000165271.19 | **NOL6** | 65083 | protein_coding | **614.565** | -0.729 | -0.219 | **0.603** | 4.48E-08 | **2.78E-07** |
| ENSG00000236935.4 | **ENSG0000(** | 10272387ٰ8 | lncRNA | **214.925** | 0.779 | 0.234 | **1.716** | 4.50E-08 | **2.79E-07** |
| ENSG00000161920.13 | **MED11** | 400569 | protein_coding | **425.081** | 0.648 | 0.195 | **1.567** | 4.52E-08 | **2.80E-07** |
| ENSG00000279108.1 | **ENSG00000279108** | | TEC | **11.147** | 3.306 | 0.995 | **9.891** | 4.52E-08 | **2.80E-07** |
| ENSG00000135316.21 | **SYNCRIP** | 10492 | protein_coding | **1977.739** | -0.714 | -0.215 | **0.610** | 4.54E-08 | **2.81E-07** |
| ENSG00000289141.3 | **ENSG00000289141** | | lncRNA | **14.511** | 2.470 | 0.743 | **5.539** | 4.57E-08 | **2.83E-07** |
| ENSG00000124491.18 | **F13A1** | 2162 | protein_coding | **724.078** | -0.844 | -0.254 | **0.557** | 4.60E-08 | **2.85E-07** |
| ENSG00000198113.5 | **TOR4A** | 54863 | protein_coding | **324.687** | -0.675 | -0.203 | **0.626** | 4.61E-08 | **2.85E-07** |
| ENSG00000128268.13 | **MGAT3** | 4248 | protein_coding | **736.558** | -0.811 | -0.244 | **0.570** | 4.68E-08 | **2.89E-07** |
| ENSG00000146054.20 | **TRIM7** | 81786 | protein_coding | **280.735** | 1.440 | 0.433 | **2.713** | 4.68E-08 | **2.89E-07** |
| ENSG00000068097.18 | **HEATR6** | 63897 | protein_coding | **391.404** | 0.816 | 0.245 | **1.760** | 4.71E-08 | **2.91E-07** |
| ENSG00000205930.12 | **EPCIP-AS1** | | lncRNA | **49.053** | 1.127 | 0.339 | **2.184** | 4.74E-08 | **2.93E-07** |
| ENSG00000163617.13 | **CCDC191** | 57577 | protein_coding | **190.052** | 0.885 | 0.266 | **1.846** | 4.75E-08 | **2.93E-07** |
| ENSG00000231389.10 | **HLA-DPA1** | 3113 | protein_coding | **22678.118** | 0.784 | 0.236 | **1.722** | 4.76E-08 | **2.94E-07** |
| ENSG00000175322.13 | **ZNF519** | 162655 | protein_coding | **98.967** | -1.251 | -0.377 | **0.420** | 4.80E-08 | **2.96E-07** |
| ENSG00000286373.2 | **ENSG00000286373** | | lncRNA | **15.814** | 2.544 | 0.766 | **5.830** | 4.81E-08 | **2.97E-07** |
| ENSG00000185561.11 | **TLCD2** | 727910 | protein_coding | **20.860** | 2.410 | 0.725 | **5.314** | 4.83E-08 | **2.98E-07** |
| ENSG00000100104.15 | **SRRD** | 402055 | protein_coding | **332.683** | -0.630 | -0.190 | **0.646** | 4.87E-08 | **3.01E-07** |
| ENSG00000145103.17 | **ILDR1** | 286676 | protein_coding | **392.817** | 0.878 | 0.264 | **1.838** | 4.89E-08 | **3.02E-07** |
| ENSG00000254338.3 | **MAFA-AS1** | | lncRNA | **33.986** | 2.615 | 0.787 | **6.124** | 4.92E-08 | **3.03E-07** |
| ENSG00000187741.18 | **FANCA** | 2175 | protein_coding | **1139.641** | -1.070 | -0.322 | **0.476** | 4.92E-08 | **3.03E-07** |
| ENSG00000197472.16 | **ZNF695** | 57116 | protein_coding | **37.115** | -1.474 | -0.444 | **0.360** | 4.95E-08 | **3.04E-07** |
| ENSG00000215533.9 | **LINC00189** | 193629 | lncRNA | **10.278** | 3.073 | 0.925 | **8.415** | 5.02E-08 | **3.09E-07** |
| ENSG00000272031.4 | **ANKRD34A** | 284615 | protein_coding | **94.754** | 1.069 | 0.322 | **2.099** | 5.02E-08 | **3.09E-07** |
| ENSG00000158526.10 | **TSR2** | 90121 | protein_coding | **716.550** | -0.561 | -0.169 | **0.678** | 5.02E-08 | **3.09E-07** |
| ENSG00000130844.21 | **ZNF331** | 55422 | protein_coding | **332.743** | -1.163 | -0.350 | **0.447** | 5.03E-08 | **3.09E-07** |

| | | | | | | | | | |
|---|---|---|---|---|---|---|---|---|---|
| ENSG00000164815.13 | **ORC5** | 5001 | protein_coding | **232.359** | -0.719 | -0.216 | **0.607** | 5.05E-08 | **3.11E-07** |
| ENSG00000100412.19 | **ACO2** | 50 | protein_coding | **2081.719** | -0.567 | -0.171 | **0.675** | 5.07E-08 | **3.12E-07** |
| ENSG00000186496.14 | **ZNF396** | 252884 | protein_coding | **31.409** | 1.549 | 0.466 | **2.927** | 5.08E-08 | **3.12E-07** |
| ENSG00000153902.16 | **LGI4** | 163175 | protein_coding | **12.203** | 3.236 | 0.974 | **9.421** | 5.10E-08 | **3.13E-07** |
| ENSG00000113108.22 | **APBB3** | 10307 | protein_coding | **1140.802** | 0.617 | 0.186 | **1.534** | 5.11E-08 | **3.14E-07** |
| ENSG00000296294.1 | **ENSG00000296294** | | lncRNA | **46.515** | 1.226 | 0.369 | **2.339** | 5.12E-08 | **3.14E-07** |
| ENSG00000112081.20 | **SRSF3** | 6428 | protein_coding | **2675.588** | -0.697 | -0.210 | **0.617** | 5.18E-08 | **3.18E-07** |
| ENSG00000184574.12 | **LPAR5** | 57121 | protein_coding | **159.217** | -0.981 | -0.295 | **0.506** | 5.20E-08 | **3.19E-07** |
| ENSG00000131171.15 | **SH3BGRL** | 6451 | protein_coding | **925.233** | -0.580 | -0.175 | **0.669** | 5.22E-08 | **3.20E-07** |
| ENSG00000072195.16 | **SPEG** | 10290 | protein_coding | **15.573** | 3.593 | 1.082 | **12.065** | 5.24E-08 | **3.21E-07** |
| ENSG00000134058.14 | **CDK7** | 1022 | protein_coding | **534.379** | -0.497 | -0.150 | **0.709** | 5.27E-08 | **3.23E-07** |
| ENSG00000104980.10 | **TIMM44** | 10469 | protein_coding | **1005.325** | -0.938 | -0.282 | **0.522** | 5.28E-08 | **3.24E-07** |
| ENSG00000164430.19 | **CGAS** | 115004 | protein_coding | **161.836** | -0.805 | -0.242 | **0.573** | 5.29E-08 | **3.24E-07** |
| ENSG00000168421.15 | **RHOH** | 399 | protein_coding | **2100.548** | 0.793 | 0.239 | **1.733** | 5.30E-08 | **3.25E-07** |
| ENSG00000067057.21 | **PFKP** | 5214 | protein_coding | **2745.523** | -0.747 | -0.225 | **0.596** | 5.31E-08 | **3.25E-07** |
| ENSG00000237753.3 | **SLC20A1-DT** | | lncRNA | **51.943** | 1.551 | 0.467 | **2.930** | 5.32E-08 | **3.26E-07** |
| ENSG00000131238.20 | **PPT1** | 5538 | protein_coding | **877.912** | -0.632 | -0.190 | **0.645** | 5.36E-08 | **3.28E-07** |
| ENSG00000162706.15 | **CADM3** | 57863 | protein_coding | **19.283** | 2.533 | 0.762 | **5.787** | 5.39E-08 | **3.30E-07** |
| ENSG00000248508.11 | **SRP14-DT** | 10013108 | lncRNA | **330.421** | 0.696 | 0.209 | **1.619** | 5.41E-08 | **3.31E-07** |
| ENSG00000173456.5 | **RNF26** | 79102 | protein_coding | **456.080** | -0.725 | -0.218 | **0.605** | 5.42E-08 | **3.31E-07** |
| ENSG00000170703.17 | **TTLL6** | 284076 | protein_coding | **23.638** | 3.486 | 1.049 | **11.206** | 5.48E-08 | **3.35E-07** |
| ENSG00000095321.20 | **CRAT** | 1384 | protein_coding | **223.821** | 0.769 | 0.232 | **1.704** | 5.51E-08 | **3.36E-07** |
| ENSG00000291317.3 | **TMEM276** | 84773 | protein_coding | **246.371** | 0.699 | 0.210 | **1.623** | 5.54E-08 | **3.38E-07** |
| ENSG00000169021.8 | **UQCRFS1** | 7386 | protein_coding | **1018.611** | -0.804 | -0.242 | **0.573** | 5.54E-08 | **3.38E-07** |
| ENSG00000177302.18 | **TOP3A** | 7156 | protein_coding | **576.535** | -0.695 | -0.209 | **0.618** | 5.54E-08 | **3.38E-07** |
| ENSG00000101898.9 | **MCTS2** | 10010149 | protein_coding | **699.964** | -0.849 | -0.256 | **0.555** | 5.55E-08 | **3.38E-07** |
| ENSG00000213189.5 | **BTF3L4P2** | | transcribed_proce | **88.198** | -1.063 | -0.320 | **0.479** | 5.55E-08 | **3.38E-07** |
| ENSG00000241468.10 | **ATP5MF** | 9551 | protein_coding | **2238.686** | -0.796 | -0.240 | **0.576** | 5.58E-08 | **3.40E-07** |
| ENSG00000161010.17 | **MRNIP** | 51149 | protein_coding | **1578.973** | 0.925 | 0.278 | **1.898** | 5.60E-08 | **3.41E-07** |
| ENSG00000287593.1 | **ENSG00000287593** | | lncRNA | **116.316** | 1.228 | 0.370 | **2.342** | 5.61E-08 | **3.42E-07** |
| ENSG00000095139.17 | **ARCN1** | 372 | protein_coding | **1520.975** | -0.539 | -0.162 | **0.688** | 5.62E-08 | **3.42E-07** |

| | | | | | | | | | |
|---|---|---|---|---|---|---|---|---|---|
| ENSG00000197226.15 | **TBC1D9B** | 23061 | protein_coding | **1790.959** | -0.421 | -0.127 | **0.747** | 5.62E-08 | **3.42E-07** |
| ENSG00000256514.1 | **ENSG00000256514** | | protein_coding | **1864.116** | 0.509 | 0.153 | **1.423** | 5.67E-08 | **3.45E-07** |
| ENSG00000178188.18 | **SH2B1** | 25970 | protein_coding | **1905.952** | 0.656 | 0.197 | **1.576** | 5.67E-08 | **3.45E-07** |
| ENSG00000167384.13 | **ZNF180** | 7733 | protein_coding | **116.222** | -0.835 | -0.251 | **0.561** | 5.68E-08 | **3.45E-07** |
| ENSG00000175147.13 | **TMEM51-A** | 200197 | lncRNA | **28.435** | 3.173 | 0.955 | **9.019** | 5.69E-08 | **3.46E-07** |
| ENSG00000243789.13 | **JMJD7** | 100137047 | protein_coding | **95.717** | -1.024 | -0.308 | **0.492** | 5.70E-08 | **3.47E-07** |
| ENSG00000178809.13 | **TRIM73** | 375593 | protein_coding | **71.671** | 1.358 | 0.409 | **2.564** | 5.71E-08 | **3.47E-07** |
| ENSG00000184702.22 | **SEPTIN5** | 5413 | protein_coding | **44.708** | 1.413 | 0.425 | **2.663** | 5.71E-08 | **3.47E-07** |
| ENSG00000293535.3 | **ENSG00000293535** | | lncRNA | **28.455** | 1.675 | 0.504 | **3.194** | 5.73E-08 | **3.48E-07** |
| ENSG00000253633.3 | **LINC03047** | | lncRNA | **302.905** | 0.982 | 0.296 | **1.975** | 5.74E-08 | **3.48E-07** |
| ENSG00000144815.20 | **NXPE3** | 91775 | protein_coding | **348.182** | 0.835 | 0.251 | **1.784** | 5.77E-08 | **3.50E-07** |
| ENSG00000013364.21 | **MVP** | 9961 | protein_coding | **3408.342** | 0.571 | 0.172 | **1.486** | 5.95E-08 | **3.61E-07** |
| ENSG00000131668.15 | **BARX1** | 56033 | protein_coding | **7.961** | 3.952 | 1.190 | **15.480** | 6.05E-08 | **3.67E-07** |
| ENSG00000112367.14 | **FIG4** | 9896 | protein_coding | **472.963** | 0.659 | 0.198 | **1.579** | 6.09E-08 | **3.69E-07** |
| ENSG00000130830.17 | **MPP1** | 4354 | protein_coding | **353.205** | 0.588 | 0.177 | **1.503** | 6.10E-08 | **3.70E-07** |
| ENSG00000105355.11 | **PLIN3** | 10226 | protein_coding | **1247.109** | 0.628 | 0.189 | **1.545** | 6.15E-08 | **3.72E-07** |
| ENSG00000266904.9 | **LINC00663** | | lncRNA | **42.293** | 1.679 | 0.505 | **3.203** | 6.15E-08 | **3.73E-07** |
| ENSG00000198934.5 | **MAGEE1** | 57692 | protein_coding | **45.605** | 1.271 | 0.383 | **2.413** | 6.16E-08 | **3.73E-07** |
| ENSG00000157782.12 | **CABP1** | 9478 | protein_coding | **183.377** | -1.227 | -0.369 | **0.427** | 6.17E-08 | **3.73E-07** |
| ENSG00000310539.1 | **DDX11L2** | 84771 | lncRNA | **24.123** | 4.304 | 1.296 | **19.758** | 6.31E-08 | **3.82E-07** |
| ENSG00000170899.13 | **GSTA4** | 2941 | protein_coding | **100.497** | 1.486 | 0.447 | **2.802** | 6.31E-08 | **3.82E-07** |
| ENSG00000166707.14 | **ZCCHC18** | 644353 | protein_coding | **38.582** | 1.501 | 0.452 | **2.831** | 6.34E-08 | **3.83E-07** |
| ENSG00000109534.19 | **GAR1** | 54433 | protein_coding | **196.301** | -0.739 | -0.223 | **0.599** | 6.44E-08 | **3.89E-07** |
| ENSG00000222020.6 | **HDAC4-AS:** | 101928111 | lncRNA | **45.652** | 1.200 | 0.361 | **2.298** | 6.44E-08 | **3.89E-07** |
| ENSG00000083093.14 | **PALB2** | 79728 | protein_coding | **218.164** | -0.697 | -0.210 | **0.617** | 6.48E-08 | **3.91E-07** |
| ENSG00000102543.16 | **CDADC1** | 81602 | protein_coding | **108.678** | -1.198 | -0.361 | **0.436** | 6.53E-08 | **3.94E-07** |
| ENSG00000253404.2 | **CTNNA1-A:** | 10537919 | lncRNA | **15.191** | 2.347 | 0.707 | **5.089** | 6.54E-08 | **3.95E-07** |
| ENSG00000259462.3 | **CPEB1-AS1** | 283692 | lncRNA | **22.415** | 2.040 | 0.614 | **4.112** | 6.55E-08 | **3.95E-07** |
| ENSG00000288930.1 | **ENSG0000(** | 124906471 | lncRNA | **9.675** | 4.377 | 1.318 | **20.775** | 6.56E-08 | **3.96E-07** |
| ENSG00000128272.21 | **ATF4** | 468 | protein_coding | **11977.854** | -0.771 | -0.232 | **0.586** | 6.60E-08 | **3.98E-07** |
| ENSG00000170759.13 | **KIF5B** | 3799 | protein_coding | **553.177** | -0.790 | -0.238 | **0.578** | 6.60E-08 | **3.98E-07** |

| | | | | | | | | | |
|---|---|---|---|---|---|---|---|---|---|
| ENSG00000143162.11 | **CREG1** | 8804 | protein_coding | **273.272** | 0.779 | 0.235 | **1.716** | 6.62E-08 | **3.99E-07** |
| ENSG00000125901.8 | **MRPS26** | 64949 | protein_coding | **710.225** | -0.605 | -0.182 | **0.657** | 6.64E-08 | **4.00E-07** |
| ENSG00000147548.19 | **NSD3** | 54904 | protein_coding | **1008.141** | -0.602 | -0.181 | **0.659** | 6.66E-08 | **4.01E-07** |
| ENSG00000273899.7 | **NOL12** | 79159 | protein_coding | **418.357** | -0.689 | -0.207 | **0.620** | 6.71E-08 | **4.04E-07** |
| ENSG00000156958.17 | **GALK2** | 2585 | protein_coding | **338.439** | -0.742 | -0.223 | **0.598** | 6.72E-08 | **4.04E-07** |
| ENSG00000178307.12 | **TMEM11** | 8834 | protein_coding | **438.119** | -0.664 | -0.200 | **0.631** | 6.72E-08 | **4.05E-07** |
| ENSG00000105472.16 | **CLEC11A** | 6320 | protein_coding | **27.306** | 1.983 | 0.597 | **3.953** | 6.73E-08 | **4.05E-07** |
| ENSG00000094914.16 | **AAAS** | 8086 | protein_coding | **1967.904** | 0.617 | 0.186 | **1.533** | 6.75E-08 | **4.06E-07** |
| ENSG00000177383.6 | **MAGEF1** | 64110 | protein_coding | **567.156** | 0.678 | 0.204 | **1.600** | 6.78E-08 | **4.08E-07** |
| ENSG00000079739.19 | **PGM1** | 5236 | protein_coding | **446.473** | -1.074 | -0.323 | **0.475** | 6.81E-08 | **4.09E-07** |
| ENSG00000308454.1 | **ENSG00000308454** | | lncRNA | **9.156** | 3.461 | 1.042 | **11.015** | 6.84E-08 | **4.11E-07** |
| ENSG00000099958.17 | **DERL3** | 91319 | protein_coding | **449.992** | -2.065 | -0.622 | **0.239** | 6.87E-08 | **4.13E-07** |
| ENSG00000130414.15 | **NDUFA10** | 4705 | protein_coding | **1564.098** | -0.512 | -0.154 | **0.701** | 6.94E-08 | **4.17E-07** |
| ENSG00000250510.10 | **GPR162** | 27239 | protein_coding | **34.119** | 3.250 | 0.978 | **9.513** | 6.94E-08 | **4.17E-07** |
| ENSG00000177666.19 | **PNPLA2** | 57104 | protein_coding | **1200.875** | 0.867 | 0.261 | **1.824** | 6.96E-08 | **4.18E-07** |
| ENSG00000142185.20 | **TRPM2** | 7226 | protein_coding | **221.893** | -1.241 | -0.374 | **0.423** | 6.97E-08 | **4.18E-07** |
| ENSG00000126001.19 | **CEP250** | 11190 | protein_coding | **791.379** | -0.666 | -0.200 | **0.630** | 7.03E-08 | **4.21E-07** |
| ENSG00000225032.5 | **ENSG0000(** | 10272356€ | lncRNA | **42.665** | 1.495 | 0.450 | **2.819** | 7.07E-08 | **4.24E-07** |
| ENSG00000137965.13 | **IFI44** | 10561 | protein_coding | **2618.394** | -0.709 | -0.213 | **0.612** | 7.08E-08 | **4.24E-07** |
| ENSG00000146909.10 | **NOM1** | 64434 | protein_coding | **322.621** | -0.809 | -0.244 | **0.571** | 7.09E-08 | **4.25E-07** |
| ENSG00000081870.14 | **IFT25** | 51668 | protein_coding | **582.750** | -0.706 | -0.212 | **0.613** | 7.11E-08 | **4.26E-07** |
| ENSG00000136783.12 | **NIPSNAP3/** | 25934 | protein_coding | **159.674** | -0.802 | -0.242 | **0.573** | 7.17E-08 | **4.29E-07** |
| ENSG00000269404.8 | **SPIB** | 6689 | protein_coding | **1674.209** | 0.786 | 0.237 | **1.725** | 7.23E-08 | **4.33E-07** |
| ENSG00000170791.20 | **CHCHD7** | 79145 | protein_coding | **769.774** | -0.660 | -0.199 | **0.633** | 7.24E-08 | **4.33E-07** |
| ENSG00000291077.2 | **ENSG00000291077** | | lncRNA | **32.944** | 1.452 | 0.437 | **2.736** | 7.35E-08 | **4.40E-07** |
| ENSG00000198959.14 | **TGM2** | 7052 | protein_coding | **231.072** | 1.181 | 0.356 | **2.267** | 7.37E-08 | **4.40E-07** |
| ENSG00000069424.17 | **KCNAB2** | 8514 | protein_coding | **3973.433** | -0.621 | -0.187 | **0.650** | 7.37E-08 | **4.40E-07** |
| ENSG00000178694.11 | **NSUN3** | 63899 | protein_coding | **194.870** | -0.647 | -0.195 | **0.639** | 7.38E-08 | **4.41E-07** |
| ENSG00000154165.5 | **GPR15** | 2838 | protein_coding | **1569.230** | 1.205 | 0.363 | **2.305** | 7.50E-08 | **4.48E-07** |
| ENSG00000078804.15 | **TP53INP2** | 58476 | protein_coding | **468.996** | 0.839 | 0.253 | **1.789** | 7.53E-08 | **4.50E-07** |
| ENSG00000152669.10 | **CCNO** | 10309 | protein_coding | **5.971** | 5.276 | 1.588 | **38.737** | 7.56E-08 | **4.51E-07** |

| | | | | | | | | | |
|---|---|---|---|---|---|---|---|---|---|
| ENSG00000097007.22 | **ABL1** | 25 | protein_coding | **804.128** | -0.612 | -0.184 | **0.654** | 7.67E-08 | **4.58E-07** |
| ENSG00000172115.11 | **CYCS** | 54205 | protein_coding | **1822.211** | -0.596 | -0.180 | **0.661** | 7.77E-08 | **4.64E-07** |
| ENSG00000299747.1 | **ENSG00000299747** | | lncRNA | **28.617** | 1.711 | 0.515 | **3.275** | 7.78E-08 | **4.64E-07** |
| ENSG00000280079.1 | **ENSG00000280079** | | TEC | **20.054** | -2.479 | -0.746 | **0.179** | 7.79E-08 | **4.64E-07** |
| ENSG00000300700.1 | **ENSG00000300700** | | lncRNA | **19.162** | 1.765 | 0.531 | **3.399** | 7.84E-08 | **4.67E-07** |
| ENSG00000109133.16 | **TMEM33** | 55161 | protein_coding | **305.174** | -0.794 | -0.239 | **0.577** | 7.86E-08 | **4.69E-07** |
| ENSG00000183019.11 | **MCEMP1** | 199675 | protein_coding | **7.740** | 4.340 | 1.306 | **20.252** | 7.87E-08 | **4.69E-07** |
| ENSG00000115415.23 | **STAT1** | 6772 | protein_coding | **10851.716** | 1.150 | 0.346 | **2.219** | 7.88E-08 | **4.69E-07** |
| ENSG00000290366.2 | **ENSG00000290366** | | lncRNA | **56.385** | 1.525 | 0.459 | **2.878** | 7.94E-08 | **4.73E-07** |
| ENSG00000132405.21 | **TBC1D14** | 57533 | protein_coding | **803.641** | 0.526 | 0.158 | **1.440** | 8.06E-08 | **4.79E-07** |
| ENSG00000136699.22 | **SMPD4** | 55627 | protein_coding | **1354.184** | -0.371 | -0.112 | **0.773** | 8.09E-08 | **4.81E-07** |
| ENSG00000196730.15 | **DAPK1** | 1612 | protein_coding | **2079.048** | 0.786 | 0.237 | **1.725** | 8.17E-08 | **4.86E-07** |
| ENSG00000280287.1 | **ENSG00000280287** | | TEC | **223.280** | 1.192 | 0.359 | **2.285** | 8.19E-08 | **4.87E-07** |
| ENSG00000159640.19 | **ACE** | 1636 | protein_coding | **12.278** | 2.896 | 0.872 | **7.445** | 8.20E-08 | **4.88E-07** |
| ENSG00000124126.16 | **PREX1** | 57580 | protein_coding | **3594.111** | 0.503 | 0.151 | **1.417** | 8.25E-08 | **4.90E-07** |
| ENSG00000144802.12 | **NFKBIZ** | 64332 | protein_coding | **726.776** | 1.167 | 0.351 | **2.246** | 8.26E-08 | **4.91E-07** |
| ENSG00000251396.10 | **LINC01301** | | lncRNA | **17.520** | 2.257 | 0.680 | **4.781** | 8.28E-08 | **4.92E-07** |
| ENSG00000132603.17 | **NIP7** | 51388 | protein_coding | **458.299** | -0.587 | -0.177 | **0.666** | 8.32E-08 | **4.94E-07** |
| ENSG00000198612.13 | **COPS8** | 10920 | protein_coding | **563.831** | -0.539 | -0.162 | **0.688** | 8.41E-08 | **4.99E-07** |
| ENSG00000187961.17 | **KLHL17** | 339451 | protein_coding | **637.001** | 0.619 | 0.186 | **1.536** | 8.42E-08 | **4.99E-07** |
| ENSG00000123179.16 | **EBPL** | 84650 | protein_coding | **323.073** | -0.568 | -0.171 | **0.675** | 8.42E-08 | **4.99E-07** |
| ENSG00000078140.16 | **UBE2K** | 3093 | protein_coding | **1265.355** | -0.522 | -0.157 | **0.696** | 8.44E-08 | **5.01E-07** |
| ENSG00000051128.21 | **HOMER3** | 9454 | protein_coding | **101.113** | 1.662 | 0.500 | **3.164** | 8.47E-08 | **5.02E-07** |
| ENSG00000126243.10 | **LRFN3** | 79414 | protein_coding | **73.436** | 1.046 | 0.315 | **2.065** | 8.49E-08 | **5.03E-07** |
| ENSG00000185900.13 | **POMK** | 84197 | protein_coding | **203.095** | -0.740 | -0.223 | **0.599** | 8.57E-08 | **5.08E-07** |
| ENSG00000103599.22 | **IQCH** | 64799 | protein_coding | **19.168** | 2.154 | 0.648 | **4.451** | 8.62E-08 | **5.10E-07** |
| ENSG00000161179.16 | **YDJC** | 150223 | protein_coding | **664.830** | -0.948 | -0.285 | **0.518** | 8.66E-08 | **5.12E-07** |
| ENSG00000158457.8 | **TSPAN33** | 340348 | protein_coding | **3829.895** | 0.734 | 0.221 | **1.663** | 8.67E-08 | **5.13E-07** |
| ENSG00000203782.6 | **LORICRIN** | 4014 | protein_coding | **14.041** | 2.890 | 0.870 | **7.415** | 8.71E-08 | **5.15E-07** |
| ENSG00000173825.9 | **TIGD3** | 220359 | protein_coding | **31.474** | 1.456 | 0.438 | **2.744** | 8.78E-08 | **5.19E-07** |
| ENSG00000187607.18 | **ZNF286A** | 57335 | protein_coding | **175.780** | -0.920 | -0.277 | **0.529** | 8.78E-08 | **5.19E-07** |

| | | | | | | | | | |
|---|---|---|---|---|---|---|---|---|---|
| ENSG00000245466.1 | **TTC9-DT** | 10192807! | lncRNA | **33.092** | -1.502 | -0.452 | **0.353** | 8.78E-08 | **5.19E-07** |
| ENSG00000166086.15 | **JAM3** | 83700 | protein_coding | **24.953** | 2.274 | 0.685 | **4.837** | 8.84E-08 | **5.22E-07** |
| ENSG00000229931.4 | **ATXN1-AS1** | | lncRNA | **233.844** | 0.835 | 0.251 | **1.784** | 8.84E-08 | **5.22E-07** |
| ENSG00000017427.19 | **IGF1** | 3479 | protein_coding | **125.625** | -1.079 | -0.325 | **0.473** | 8.85E-08 | **5.22E-07** |
| ENSG00000198892.9 | **SHISA4** | 149345 | protein_coding | **16.235** | 2.669 | 0.803 | **6.359** | 8.90E-08 | **5.25E-07** |
| ENSG00000130856.19 | **ZNF236** | 7776 | protein_coding | **424.620** | 0.932 | 0.280 | **1.907** | 8.97E-08 | **5.29E-07** |
| ENSG00000054523.22 | **KIF1B** | 23095 | protein_coding | **209.101** | 0.729 | 0.219 | **1.657** | 9.01E-08 | **5.32E-07** |
| ENSG00000118308.18 | **IRAG2** | 4033 | protein_coding | **3651.662** | -0.880 | -0.265 | **0.543** | 9.02E-08 | **5.32E-07** |
| ENSG00000123908.13 | **AGO2** | 27161 | protein_coding | **843.242** | -1.125 | -0.339 | **0.458** | 9.05E-08 | **5.34E-07** |
| ENSG00000107968.12 | **MAP3K8** | 1326 | protein_coding | **1504.483** | 0.836 | 0.252 | **1.786** | 9.09E-08 | **5.36E-07** |
| ENSG00000136689.21 | **IL1RN** | 3557 | protein_coding | **12.780** | 3.024 | 0.910 | **8.134** | 9.16E-08 | **5.40E-07** |
| ENSG00000051825.17 | **MPHOSPH!** | 10198 | protein_coding | **368.788** | -0.955 | -0.287 | **0.516** | 9.19E-08 | **5.41E-07** |
| ENSG00000106603.22 | **COA1** | 55744 | protein_coding | **1186.127** | 0.397 | 0.119 | **1.317** | 9.44E-08 | **5.56E-07** |
| ENSG00000136830.14 | **NIBAN2** | 64855 | protein_coding | **51.092** | 1.946 | 0.586 | **3.852** | 9.50E-08 | **5.59E-07** |
| ENSG00000124733.6 | **MEA1** | 4201 | protein_coding | **1051.796** | -0.592 | -0.178 | **0.663** | 9.50E-08 | **5.59E-07** |
| ENSG00000147224.14 | **PRPS1** | 5631 | protein_coding | **426.351** | -0.876 | -0.264 | **0.545** | 9.51E-08 | **5.60E-07** |
| ENSG00000236352.2 | **BCAS1-AS1** | | lncRNA | **98.193** | 1.036 | 0.312 | **2.050** | 9.54E-08 | **5.61E-07** |
| ENSG00000261360.4 | **OTULIN-DT** | 728178 | lncRNA | **69.400** | 1.528 | 0.460 | **2.885** | 9.55E-08 | **5.62E-07** |
| ENSG00000197375.16 | **SLC22A5** | 6584 | protein_coding | **99.048** | 1.274 | 0.383 | **2.418** | 9.57E-08 | **5.63E-07** |
| ENSG00000116747.15 | **RO60** | 6738 | protein_coding | **274.442** | -0.824 | -0.248 | **0.565** | 9.64E-08 | **5.67E-07** |
| ENSG00000261618.2 | **LINC02605** | | lncRNA | **81.170** | 1.530 | 0.461 | **2.888** | 9.68E-08 | **5.69E-07** |
| ENSG00000172575.15 | **RASGRP1** | 10125 | protein_coding | **169.702** | -0.803 | -0.242 | **0.573** | 9.72E-08 | **5.71E-07** |
| ENSG00000274422.1 | **ENSG00000274422** | | lncRNA | **28.433** | 1.511 | 0.455 | **2.850** | 9.77E-08 | **5.74E-07** |
| ENSG00000123545.8 | **NDUFAF4** | 29078 | protein_coding | **282.532** | -0.631 | -0.190 | **0.646** | 9.84E-08 | **5.78E-07** |
| ENSG00000157764.16 | **BRAF** | 673 | protein_coding | **480.151** | -0.714 | -0.215 | **0.610** | 9.87E-08 | **5.79E-07** |
| ENSG00000167550.13 | **RHEBL1** | 121268 | protein_coding | **238.663** | -0.718 | -0.216 | **0.608** | 9.95E-08 | **5.83E-07** |
| ENSG00000112715.26 | **VEGFA** | 7422 | protein_coding | **3174.451** | -0.685 | -0.206 | **0.622** | 1.00E-07 | **5.89E-07** |
| ENSG00000197275.16 | **RAD54B** | 25788 | protein_coding | **380.378** | -0.704 | -0.212 | **0.614** | 1.01E-07 | **5.89E-07** |
| ENSG00000141179.15 | **PCTP** | 58488 | protein_coding | **280.621** | 0.970 | 0.292 | **1.959** | 1.01E-07 | **5.90E-07** |
| ENSG00000183018.11 | **SPNS2** | 124976 | protein_coding | **50.050** | 1.940 | 0.584 | **3.837** | 1.01E-07 | **5.92E-07** |
| ENSG00000091436.19 | **MAP3K20** | 51776 | protein_coding | **178.881** | -0.995 | -0.299 | **0.502** | 1.01E-07 | **5.92E-07** |

| | | | | | | | | | |
|---|---|---|---|---|---|---|---|---|---|
| ENSG00000145293.19 | **ENOPH1** | 58478 | protein_coding | **515.894** | -0.635 | -0.191 | **0.644** | 1.01E-07 | **5.92E-07** |
| ENSG00000182004.15 | **SNRPE** | 6635 | protein_coding | **947.023** | -0.838 | -0.252 | **0.559** | 1.01E-07 | **5.92E-07** |
| ENSG00000137656.14 | **BUD13** | 84811 | protein_coding | **316.704** | -0.612 | -0.184 | **0.654** | 1.01E-07 | **5.93E-07** |
| ENSG00000023902.15 | **PLEKHO1** | 51177 | protein_coding | **2064.552** | -0.589 | -0.177 | **0.665** | 1.02E-07 | **5.94E-07** |
| ENSG00000005249.16 | **PRKAR2B** | 5577 | protein_coding | **9.659** | 3.651 | 1.099 | **12.566** | 1.02E-07 | **5.96E-07** |
| ENSG00000271913.10 | **TAGAP-AS1** | | lncRNA | **60.206** | 1.027 | 0.309 | **2.037** | 1.02E-07 | **5.97E-07** |
| ENSG00000120992.20 | **LYPLA1** | 10434 | protein_coding | **240.373** | -0.753 | -0.227 | **0.593** | 1.03E-07 | **6.02E-07** |
| ENSG00000188985.6 | **DHFRP1** | | processed_pseud | **112.662** | -4.800 | -1.445 | **0.036** | 1.03E-07 | **6.04E-07** |
| ENSG00000050426.18 | **LETMD1** | 25875 | protein_coding | **1293.173** | -0.439 | -0.132 | **0.738** | 1.03E-07 | **6.04E-07** |
| ENSG00000178947.10 | **SMIM10L2** | 399668 | protein_coding | **9.046** | 3.721 | 1.120 | **13.190** | 1.04E-07 | **6.05E-07** |
| ENSG00000235703.8 | **EOLA2-DT** | | lncRNA | **163.955** | 0.903 | 0.272 | **1.870** | 1.04E-07 | **6.08E-07** |
| ENSG00000188511.15 | **MIR3667H** | 348645 | lncRNA | **168.750** | -0.911 | -0.274 | **0.532** | 1.05E-07 | **6.11E-07** |
| ENSG00000140471.19 | **LINS1** | 55180 | protein_coding | **266.002** | -0.626 | -0.189 | **0.648** | 1.05E-07 | **6.11E-07** |
| ENSG00000148187.20 | **MRRF** | 92399 | protein_coding | **1300.854** | -0.548 | -0.165 | **0.684** | 1.05E-07 | **6.12E-07** |
| ENSG00000124380.13 | **SNRNP27** | 11017 | protein_coding | **338.744** | -0.700 | -0.211 | **0.615** | 1.05E-07 | **6.12E-07** |
| ENSG00000182287.17 | **AP1S2** | 8905 | protein_coding | **332.200** | -0.776 | -0.234 | **0.584** | 1.05E-07 | **6.12E-07** |
| ENSG00000261118.2 | **ENSG0000** | 10192786 | lncRNA | **22.299** | 1.742 | 0.524 | **3.344** | 1.05E-07 | **6.14E-07** |
| ENSG00000008283.18 | **CYB561** | 1534 | protein_coding | **186.722** | 0.919 | 0.277 | **1.891** | 1.06E-07 | **6.15E-07** |
| ENSG00000137955.18 | **RABGGTB** | 5876 | protein_coding | **1982.972** | -0.563 | -0.170 | **0.677** | 1.06E-07 | **6.15E-07** |
| ENSG00000164414.21 | **SLC35A1** | 10559 | protein_coding | **456.378** | 0.590 | 0.178 | **1.505** | 1.07E-07 | **6.20E-07** |
| ENSG00000118680.16 | **MYL12B** | 103910 | protein_coding | **4020.977** | 0.567 | 0.171 | **1.481** | 1.07E-07 | **6.20E-07** |
| ENSG00000131473.19 | **ACLY** | 47 | protein_coding | **2271.133** | -0.560 | -0.169 | **0.678** | 1.07E-07 | **6.21E-07** |
| ENSG00000184990.15 | **SIVA1** | 10572 | protein_coding | **806.525** | -0.587 | -0.177 | **0.666** | 1.07E-07 | **6.21E-07** |
| ENSG00000087303.20 | **NID2** | 22795 | protein_coding | **17.935** | 3.294 | 0.992 | **9.808** | 1.07E-07 | **6.21E-07** |
| ENSG00000100347.17 | **SAMM50** | 25813 | protein_coding | **723.263** | -0.595 | -0.179 | **0.662** | 1.08E-07 | **6.28E-07** |
| ENSG00000283674.5 | **ENSG0000** | 729732 | lncRNA | **54.585** | 1.490 | 0.448 | **2.809** | 1.09E-07 | **6.31E-07** |
| ENSG00000100575.16 | **TIMM9** | 26520 | protein_coding | **801.557** | -0.462 | -0.139 | **0.726** | 1.09E-07 | **6.32E-07** |
| ENSG00000152944.11 | **MED21** | 9412 | protein_coding | **152.737** | -0.795 | -0.239 | **0.576** | 1.09E-07 | **6.32E-07** |
| ENSG00000177453.9 | **NIM1K** | 167359 | protein_coding | **97.832** | 1.051 | 0.316 | **2.071** | 1.09E-07 | **6.35E-07** |
| ENSG00000147416.14 | **ATP6V1B2** | 526 | protein_coding | **655.186** | -0.537 | -0.162 | **0.689** | 1.09E-07 | **6.35E-07** |
| ENSG00000186226.9 | **LCE1E** | 353135 | protein_coding | **13.107** | -3.123 | -0.940 | **0.115** | 1.10E-07 | **6.35E-07** |

| | | | | | | | | | |
|---|---|---|---|---|---|---|---|---|---|
| ENSG00000166503.12 | **HDGFL3** | 50810 | protein_coding | **486.558** | -0.651 | -0.196 | **0.637** | 1.11E-07 | **6.42E-07** |
| ENSG00000267757.5 | **EML2-AS1** | 100287177 | lncRNA | **143.882** | -0.813 | -0.245 | **0.569** | 1.14E-07 | **6.59E-07** |
| ENSG00000141101.15 | **NOB1** | 28987 | protein_coding | **1409.329** | -0.663 | -0.200 | **0.631** | 1.15E-07 | **6.69E-07** |
| ENSG00000138111.16 | **SLC68A1** | 79847 | protein_coding | **100.355** | 0.966 | 0.291 | **1.954** | 1.16E-07 | **6.71E-07** |
| ENSG00000126264.12 | **HCST** | 10870 | protein_coding | **429.183** | -1.117 | -0.336 | **0.461** | 1.17E-07 | **6.77E-07** |
| ENSG00000181904.11 | **C5orf24** | 134553 | protein_coding | **163.576** | -1.113 | -0.335 | **0.462** | 1.17E-07 | **6.80E-07** |
| ENSG00000244038.12 | **DDOST** | 1650 | protein_coding | **3008.379** | -0.526 | -0.158 | **0.695** | 1.17E-07 | **6.80E-07** |
| ENSG00000187260.17 | **WDR86** | 349136 | protein_coding | **11.196** | 3.073 | 0.925 | **8.418** | 1.18E-07 | **6.80E-07** |
| ENSG00000280649.2 | **ENSG00000280649** | | TEC | **94.669** | 1.380 | 0.415 | **2.603** | 1.19E-07 | **6.87E-07** |
| ENSG00000077984.8 | **CST7** | 8530 | protein_coding | **10.885** | -3.570 | -1.075 | **0.084** | 1.19E-07 | **6.88E-07** |
| ENSG00000237017.1 | **PRPF31-AS1** | | lncRNA | **59.336** | 1.591 | 0.479 | **3.012** | 1.19E-07 | **6.89E-07** |
| ENSG00000187164.22 | **SHTN1** | 57698 | protein_coding | **22.382** | 2.031 | 0.611 | **4.088** | 1.22E-07 | **7.03E-07** |
| ENSG00000179583.22 | **CIITA** | 4261 | protein_coding | **2766.098** | 0.707 | 0.213 | **1.632** | 1.22E-07 | **7.04E-07** |
| ENSG00000291158.2 | **LINC00869** | 57234 | lncRNA | **193.014** | -0.942 | -0.283 | **0.521** | 1.22E-07 | **7.04E-07** |
| ENSG00000196290.19 | **NIF3L1** | 60491 | protein_coding | **454.640** | -0.503 | -0.151 | **0.706** | 1.23E-07 | **7.08E-07** |
| ENSG00000248905.11 | **FMN1** | 342184 | protein_coding | **31.762** | 2.830 | 0.852 | **7.111** | 1.23E-07 | **7.11E-07** |
| ENSG00000177595.20 | **PIDD1** | 55367 | protein_coding | **929.177** | -0.632 | -0.190 | **0.646** | 1.24E-07 | **7.15E-07** |
| ENSG00000100678.22 | **SLC8A3** | 6547 | protein_coding | **182.509** | -0.852 | -0.257 | **0.554** | 1.25E-07 | **7.18E-07** |
| ENSG00000167632.20 | **TRAPPC9** | 83696 | protein_coding | **413.287** | 0.805 | 0.242 | **1.747** | 1.25E-07 | **7.20E-07** |
| ENSG00000158525.16 | **CPA5** | 93979 | protein_coding | **25.612** | 2.637 | 0.794 | **6.222** | 1.26E-07 | **7.27E-07** |
| ENSG00000271780.3 | **ENSG00000271780** | | lncRNA | **76.603** | 0.951 | 0.286 | **1.933** | 1.27E-07 | **7.30E-07** |
| ENSG00000147443.15 | **DOK2** | 9046 | protein_coding | **132.301** | -0.933 | -0.281 | **0.524** | 1.27E-07 | **7.31E-07** |
| ENSG00000259456.5 | **ADNP-AS1** | 101927631 | lncRNA | **12.300** | -2.501 | -0.753 | **0.177** | 1.29E-07 | **7.44E-07** |
| ENSG00000167862.12 | **MRPL58** | 3396 | protein_coding | **408.413** | -0.528 | -0.159 | **0.693** | 1.31E-07 | **7.52E-07** |
| ENSG00000265118.5 | **ENSG00000265118** | | protein_coding | **26.373** | -7.064 | -2.126 | **0.007** | 1.31E-07 | **7.56E-07** |
| ENSG00000198945.10 | **L3MBTL3** | 84456 | protein_coding | **230.396** | 1.027 | 0.309 | **2.037** | 1.32E-07 | **7.60E-07** |
| ENSG00000132591.14 | **ERAL1** | 26284 | protein_coding | **916.104** | -0.660 | -0.199 | **0.633** | 1.33E-07 | **7.65E-07** |
| ENSG00000187833.8 | **C2orf78** | 388960 | protein_coding | **17.181** | 2.636 | 0.793 | **6.215** | 1.33E-07 | **7.66E-07** |
| ENSG00000178403.4 | **NEUROG2** | 63973 | protein_coding | **164.981** | -1.128 | -0.339 | **0.458** | 1.34E-07 | **7.70E-07** |
| ENSG00000143942.7 | **CHAC2** | 494143 | protein_coding | **53.885** | -1.410 | -0.424 | **0.376** | 1.34E-07 | **7.70E-07** |
| ENSG00000155660.13 | **PDIA4** | 9601 | protein_coding | **2516.903** | -0.612 | -0.184 | **0.654** | 1.35E-07 | **7.76E-07** |

| | | | | | | | | | |
|---|---|---|---|---|---|---|---|---|---|
| ENSG00000110492.19 | **MDK** | 4192 | protein_coding | **125.629** | 1.788 | 0.538 | **3.454** | 1.35E-07 | **7.78E-07** |
| ENSG00000213123.13 | **DYNLT2B** | 255758 | protein_coding | **114.047** | -0.735 | -0.221 | **0.601** | 1.36E-07 | **7.80E-07** |
| ENSG00000174233.14 | **ADCY6** | 112 | protein_coding | **207.894** | 1.035 | 0.312 | **2.049** | 1.37E-07 | **7.89E-07** |
| ENSG00000182872.18 | **RBM10** | 8241 | protein_coding | **1538.425** | -0.454 | -0.137 | **0.730** | 1.38E-07 | **7.92E-07** |
| ENSG00000182568.20 | **SATB1** | 6304 | protein_coding | **39.142** | 2.535 | 0.763 | **5.795** | 1.38E-07 | **7.94E-07** |
| ENSG00000161618.12 | **ALDH16A1** | 126133 | protein_coding | **688.968** | -0.786 | -0.237 | **0.580** | 1.39E-07 | **8.00E-07** |
| ENSG00000077713.21 | **SLC25A43** | 203427 | protein_coding | **190.049** | -0.637 | -0.192 | **0.643** | 1.42E-07 | **8.15E-07** |
| ENSG00000141905.21 | **NFIC** | 4782 | protein_coding | **461.102** | -0.722 | -0.217 | **0.606** | 1.42E-07 | **8.15E-07** |
| ENSG00000055163.23 | **CYFIP2** | 26999 | protein_coding | **2909.130** | -0.636 | -0.192 | **0.643** | 1.43E-07 | **8.22E-07** |
| ENSG00000128881.20 | **TTBK2** | 146057 | protein_coding | **359.394** | 0.695 | 0.209 | **1.619** | 1.44E-07 | **8.24E-07** |
| ENSG00000164053.24 | **ATRIP** | 84126 | protein_coding | **357.117** | -0.558 | -0.168 | **0.679** | 1.45E-07 | **8.30E-07** |
| ENSG00000203395.2 | **ENSG00000203395** | | lncRNA | **10.627** | 2.747 | 0.827 | **6.715** | 1.45E-07 | **8.30E-07** |
| ENSG00000267462.1 | **ENSG00000267462** | | lncRNA | **10.134** | 2.967 | 0.893 | **7.820** | 1.46E-07 | **8.36E-07** |
| ENSG00000268996.3 | **MAN1B1-D** | 100289341 | lncRNA | **39.348** | 1.262 | 0.380 | **2.398** | 1.46E-07 | **8.37E-07** |
| ENSG00000186716.23 | **BCR** | 613 | protein_coding | **409.737** | 1.038 | 0.312 | **2.053** | 1.47E-07 | **8.43E-07** |
| ENSG00000077420.19 | **APBB1IP** | 54518 | protein_coding | **1210.502** | -0.563 | -0.169 | **0.677** | 1.48E-07 | **8.45E-07** |
| ENSG00000188690.18 | **UROS** | 7390 | protein_coding | **1038.077** | 0.539 | 0.162 | **1.453** | 1.50E-07 | **8.55E-07** |
| ENSG00000167674.17 | **HDGFL2** | 84717 | protein_coding | **1858.376** | 0.429 | 0.129 | **1.347** | 1.50E-07 | **8.55E-07** |
| ENSG00000146263.14 | **MMS22L** | 253714 | protein_coding | **205.111** | -1.211 | -0.364 | **0.432** | 1.50E-07 | **8.56E-07** |
| ENSG00000188283.14 | **ZNF383** | 163087 | protein_coding | **148.041** | -1.034 | -0.311 | **0.488** | 1.51E-07 | **8.60E-07** |
| ENSG00000175193.16 | **PARL** | 55486 | protein_coding | **670.791** | -0.672 | -0.202 | **0.628** | 1.51E-07 | **8.61E-07** |
| ENSG00000160752.17 | **FDPS** | 2224 | protein_coding | **5106.453** | -1.035 | -0.311 | **0.488** | 1.51E-07 | **8.62E-07** |
| ENSG00000109861.19 | **CTSC** | 1075 | protein_coding | **2174.098** | -0.732 | -0.220 | **0.602** | 1.52E-07 | **8.66E-07** |
| ENSG00000166123.16 | **GPT2** | 84706 | protein_coding | **515.314** | -1.143 | -0.344 | **0.453** | 1.52E-07 | **8.66E-07** |
| ENSG00000185527.13 | **PDE6G** | 5148 | protein_coding | **89.251** | -1.241 | -0.373 | **0.423** | 1.54E-07 | **8.75E-07** |
| ENSG00000254413.8 | **CHKB-CPT1B** | | protein_coding | **323.065** | 0.723 | 0.218 | **1.651** | 1.54E-07 | **8.80E-07** |
| ENSG00000130560.11 | **UBAC1** | 10422 | protein_coding | **604.903** | -0.584 | -0.176 | **0.667** | 1.55E-07 | **8.86E-07** |
| ENSG00000134825.18 | **TMEM258** | 746 | protein_coding | **1536.214** | -0.786 | -0.237 | **0.580** | 1.58E-07 | **9.02E-07** |
| ENSG00000196335.14 | **STK31** | 56164 | protein_coding | **12.015** | 2.683 | 0.808 | **6.421** | 1.58E-07 | **9.03E-07** |
| ENSG00000227533.8 | **SLC2A1-DT** | 440584 | lncRNA | **58.804** | 1.442 | 0.434 | **2.718** | 1.59E-07 | **9.04E-07** |
| ENSG00000070371.18 | **CLTCL1** | 8218 | protein_coding | **19.077** | 2.273 | 0.684 | **4.833** | 1.60E-07 | **9.08E-07** |

| | | | | | | | | | |
|---|---|---|---|---|---|---|---|---|---|
| ENSG00000128683.16 | **GAD1** | 2571 | protein_coding | **157.449** | 1.405 | 0.423 | **2.647** | 1.60E-07 | **9.12E-07** |
| ENSG00000290723.2 | **ENSG00000290723** | | lncRNA | **13.393** | 2.532 | 0.762 | **5.785** | 1.60E-07 | **9.12E-07** |
| ENSG00000254087.10 | **LYN** | 4067 | protein_coding | **3600.599** | 0.531 | 0.160 | **1.445** | 1.60E-07 | **9.13E-07** |
| ENSG00000114735.12 | **HEMK1** | 51409 | protein_coding | **1005.266** | 0.532 | 0.160 | **1.446** | 1.62E-07 | **9.20E-07** |
| ENSG00000267493.4 | **CIRBP-AS1** | 148046 | lncRNA | **12.058** | 2.676 | 0.806 | **6.391** | 1.62E-07 | **9.21E-07** |
| ENSG00000234509.2 | **SOD1-DT** | | lncRNA | **6.524** | 4.517 | 1.360 | **22.903** | 1.62E-07 | **9.22E-07** |
| ENSG00000160753.19 | **RUSC1** | 23623 | protein_coding | **442.453** | 0.674 | 0.203 | **1.595** | 1.64E-07 | **9.30E-07** |
| ENSG00000111142.16 | **METAP2** | 10988 | protein_coding | **705.822** | -0.839 | -0.253 | **0.559** | 1.64E-07 | **9.31E-07** |
| ENSG00000112146.18 | **FBXO9** | 26268 | protein_coding | **597.609** | 0.587 | 0.177 | **1.502** | 1.64E-07 | **9.32E-07** |
| ENSG00000234964.4 | **FABP5P7** | | processed_pseudo | **193.962** | -0.820 | -0.247 | **0.566** | 1.65E-07 | **9.35E-07** |
| ENSG00000116350.20 | **SRSF4** | 6429 | protein_coding | **2648.359** | -0.425 | -0.128 | **0.745** | 1.65E-07 | **9.36E-07** |
| ENSG00000162607.15 | **USP1** | 7398 | protein_coding | **382.079** | -0.915 | -0.275 | **0.530** | 1.65E-07 | **9.37E-07** |
| ENSG00000198885.12 | **ITPRIPL1** | 150771 | protein_coding | **296.957** | -0.714 | -0.215 | **0.610** | 1.66E-07 | **9.43E-07** |
| ENSG00000143622.14 | **RIT1** | 6016 | protein_coding | **349.856** | -0.781 | -0.235 | **0.582** | 1.67E-07 | **9.44E-07** |
| ENSG00000185567.8 | **AHNAK2** | 113146 | protein_coding | **10.679** | 3.759 | 1.131 | **13.535** | 1.67E-07 | **9.46E-07** |
| ENSG00000213965.5 | **NUDT19** | 390916 | protein_coding | **331.886** | -0.655 | -0.197 | **0.635** | 1.67E-07 | **9.46E-07** |
| ENSG00000142089.19 | **IFITM3** | 10410 | protein_coding | **393.565** | -1.278 | -0.385 | **0.413** | 1.67E-07 | **9.48E-07** |
| ENSG00000108551.7 | **RASD1** | 51655 | protein_coding | **25.461** | 2.776 | 0.836 | **6.851** | 1.68E-07 | **9.49E-07** |
| ENSG00000164828.20 | **SUN1** | 23353 | protein_coding | **2271.152** | 0.636 | 0.191 | **1.554** | 1.68E-07 | **9.52E-07** |
| ENSG00000176225.17 | **RTTN** | 25914 | protein_coding | **478.275** | -0.558 | -0.168 | **0.679** | 1.69E-07 | **9.57E-07** |
| ENSG00000172336.7 | **POP7** | 10248 | protein_coding | **519.878** | -0.726 | -0.219 | **0.605** | 1.69E-07 | **9.59E-07** |
| ENSG00000114631.13 | **PODXL2** | 50512 | protein_coding | **108.603** | 1.155 | 0.348 | **2.226** | 1.72E-07 | **9.71E-07** |
| ENSG00000143537.16 | **ADAM15** | 8751 | protein_coding | **501.686** | 0.792 | 0.238 | **1.731** | 1.72E-07 | **9.72E-07** |
| ENSG00000117602.15 | **RCAN3** | 11123 | protein_coding | **18.334** | 2.101 | 0.633 | **4.290** | 1.73E-07 | **9.77E-07** |
| ENSG00000261790.2 | **ENSG00000261790** | | lncRNA | **15.360** | 2.065 | 0.621 | **4.183** | 1.76E-07 | **9.92E-07** |
| ENSG00000049449.12 | **RCN1** | 5954 | protein_coding | **1268.524** | -0.664 | -0.200 | **0.631** | 1.76E-07 | **9.92E-07** |
| ENSG00000134326.13 | **CMPK2** | 129607 | protein_coding | **150.696** | -1.240 | -0.373 | **0.423** | 1.76E-07 | **9.96E-07** |
| ENSG00000245910.10 | **SNHG6** | | lncRNA | **1501.950** | -0.750 | -0.226 | **0.595** | 1.78E-07 | **1.00E-06** |
| ENSG00000300824.1 | **ENSG00000300824** | | lncRNA | **27.100** | 1.573 | 0.474 | **2.976** | 1.78E-07 | **1.00E-06** |
| ENSG00000105671.14 | **DDX49** | 54555 | protein_coding | **825.248** | -0.649 | -0.195 | **0.638** | 1.78E-07 | **1.00E-06** |
| ENSG00000186318.20 | **BACE1** | 23621 | protein_coding | **26.329** | 1.588 | 0.478 | **3.006** | 1.79E-07 | **1.01E-06** |

| | | | | | | | | | |
|---|---|---|---|---|---|---|---|---|---|
| ENSG00000135638.16 | **EMX1** | 2016 | protein_coding | **200.208** | -0.840 | -0.253 | **0.559** | 1.80E-07 | **1.02E-06** |
| ENSG00000184986.14 | **TMEM121** | 80757 | protein_coding | **15.174** | 2.592 | 0.780 | **6.028** | 1.81E-07 | **1.02E-06** |
| ENSG00000278540.6 | **ACACA** | 31 | protein_coding | **702.417** | -0.647 | -0.195 | **0.639** | 1.81E-07 | **1.02E-06** |
| ENSG00000185519.11 | **FAM131C** | 348487 | protein_coding | **10.303** | 3.852 | 1.159 | **14.436** | 1.81E-07 | **1.02E-06** |
| ENSG00000180881.23 | **CAPS2** | 84698 | protein_coding | **86.297** | 0.863 | 0.260 | **1.819** | 1.81E-07 | **1.02E-06** |
| ENSG00000164038.18 | **SLC9B2** | 133308 | protein_coding | **297.689** | -0.735 | -0.221 | **0.601** | 1.82E-07 | **1.02E-06** |
| ENSG00000108963.20 | **DPH1** | 1801 | protein_coding | **733.314** | 0.631 | 0.190 | **1.549** | 1.82E-07 | **1.02E-06** |
| ENSG00000126067.14 | **PSMB2** | 5690 | protein_coding | **2357.370** | -0.703 | -0.212 | **0.614** | 1.82E-07 | **1.02E-06** |
| ENSG00000128298.20 | **BAIAP2L2** | 80115 | protein_coding | **71.402** | 1.106 | 0.333 | **2.153** | 1.83E-07 | **1.03E-06** |
| ENSG00000100473.20 | **COCH** | 1690 | protein_coding | **252.077** | -0.610 | -0.184 | **0.655** | 1.83E-07 | **1.03E-06** |
| ENSG00000156521.16 | **TYSND1** | 219743 | protein_coding | **485.999** | -0.647 | -0.195 | **0.639** | 1.85E-07 | **1.04E-06** |
| ENSG00000184792.18 | **OSBP2** | 23762 | protein_coding | **26.626** | 1.793 | 0.540 | **3.465** | 1.85E-07 | **1.04E-06** |
| ENSG00000157429.17 | **ZNF19** | 7567 | protein_coding | **76.962** | 0.948 | 0.285 | **1.929** | 1.85E-07 | **1.04E-06** |
| ENSG00000100505.16 | **TRIM9** | 114088 | protein_coding | **44.791** | 1.890 | 0.569 | **3.706** | 1.86E-07 | **1.05E-06** |
| ENSG00000171714.15 | **ANO5** | 203859 | protein_coding | **201.203** | 0.690 | 0.208 | **1.614** | 1.87E-07 | **1.05E-06** |
| ENSG00000300781.1 | **ENSG00000300781** | | lncRNA | **112.413** | -1.040 | -0.313 | **0.486** | 1.88E-07 | **1.06E-06** |
| ENSG00000116584.24 | **ARHGEF2** | 9181 | protein_coding | **1687.510** | 0.429 | 0.129 | **1.346** | 1.88E-07 | **1.06E-06** |
| ENSG00000232931.7 | **LINC00342** | | lncRNA | **460.299** | -0.682 | -0.205 | **0.623** | 1.89E-07 | **1.06E-06** |
| ENSG00000295628.1 | **ENSG0000(** | 105377582 | lncRNA | **26.879** | 2.148 | 0.647 | **4.432** | 1.90E-07 | **1.06E-06** |
| ENSG00000155561.17 | **NUP205** | 23165 | protein_coding | **1270.037** | -0.746 | -0.225 | **0.596** | 1.91E-07 | **1.07E-06** |
| ENSG00000065989.18 | **PDE4A** | 5141 | protein_coding | **718.263** | 0.629 | 0.189 | **1.546** | 1.91E-07 | **1.07E-06** |
| ENSG00000106028.13 | **SSBP1** | 6742 | protein_coding | **1799.997** | -0.515 | -0.155 | **0.700** | 1.92E-07 | **1.07E-06** |
| ENSG00000005448.19 | **WDR54** | 84058 | protein_coding | **1012.403** | -0.808 | -0.243 | **0.571** | 1.92E-07 | **1.07E-06** |
| ENSG00000266903.4 | **CEACAM16** | 107985305 | lncRNA | **10.773** | 3.033 | 0.913 | **8.183** | 1.94E-07 | **1.08E-06** |
| ENSG00000170291.19 | **ELP5** | 23587 | protein_coding | **822.480** | -0.584 | -0.176 | **0.667** | 1.95E-07 | **1.09E-06** |
| ENSG00000070882.15 | **OSBPL3** | 26031 | protein_coding | **804.335** | -0.794 | -0.239 | **0.577** | 1.95E-07 | **1.09E-06** |
| ENSG00000232065.3 | **LINC01063** | | lncRNA | **15.271** | 2.271 | 0.684 | **4.826** | 1.95E-07 | **1.09E-06** |
| ENSG00000170161.8 | **FAM88B** | | lncRNA | **51.484** | -1.374 | -0.414 | **0.386** | 1.96E-07 | **1.10E-06** |
| ENSG00000166926.11 | **MS4A6E** | 245802 | protein_coding | **26.046** | 2.680 | 0.807 | **6.408** | 1.96E-07 | **1.10E-06** |
| ENSG00000222041.13 | **CYTOR** | 112597 | lncRNA | **236.798** | -0.805 | -0.242 | **0.572** | 1.97E-07 | **1.10E-06** |
| ENSG00000132432.16 | **SEC61G** | 23480 | protein_coding | **564.751** | -0.927 | -0.279 | **0.526** | 1.97E-07 | **1.10E-06** |

| | | | | | | | | | |
|---|---|---|---|---|---|---|---|---|---|
| ENSG00000137571.13 | **SLCO5A1** | 81796 | protein_coding | **358.357** | -0.617 | -0.186 | **0.652** | 1.98E-07 | **1.11E-06** |
| ENSG00000124789.14 | **NUP153** | 9972 | protein_coding | **858.094** | -0.766 | -0.231 | **0.588** | 1.99E-07 | **1.11E-06** |
| ENSG00000225555.3 | **ENSG0000(** | 105372791 | lncRNA | **7.203** | 3.906 | 1.176 | **14.988** | 1.99E-07 | **1.11E-06** |
| ENSG00000224614.3 | **TNK2-AS1** | | lncRNA | **19.945** | -2.087 | -0.628 | **0.235** | 2.00E-07 | **1.11E-06** |
| ENSG00000142765.20 | **SYTL1** | 84958 | protein_coding | **1086.067** | -1.325 | -0.399 | **0.399** | 2.00E-07 | **1.12E-06** |
| ENSG00000289915.2 | **ENSG00000289915** | | lncRNA | **14.255** | -2.429 | -0.731 | **0.186** | 2.00E-07 | **1.12E-06** |
| ENSG00000295911.1 | **ENSG00000295911** | | lncRNA | **333.245** | -1.275 | -0.384 | **0.413** | 2.01E-07 | **1.12E-06** |
| ENSG00000106366.11 | **SERPINE1** | 5054 | protein_coding | **33.110** | 1.810 | 0.545 | **3.507** | 2.02E-07 | **1.13E-06** |
| ENSG00000259772.9 | **LINC03034** | 283710 | lncRNA | **2722.466** | 0.972 | 0.292 | **1.961** | 2.03E-07 | **1.13E-06** |
| ENSG00000184634.19 | **MED12** | 9968 | protein_coding | **1009.558** | -0.616 | -0.185 | **0.653** | 2.03E-07 | **1.13E-06** |
| ENSG00000258457.6 | **AJUBA-DT** | | lncRNA | **10.527** | 2.721 | 0.819 | **6.591** | 2.03E-07 | **1.13E-06** |
| ENSG00000144320.16 | **LNPK** | 80856 | protein_coding | **269.412** | -1.110 | -0.334 | **0.463** | 2.04E-07 | **1.14E-06** |
| ENSG00000167468.21 | **GPX4** | 2879 | protein_coding | **6042.587** | 0.590 | 0.178 | **1.506** | 2.07E-07 | **1.15E-06** |
| ENSG00000198668.16 | **CALM1** | 801 | protein_coding | **2373.225** | -0.468 | -0.141 | **0.723** | 2.07E-07 | **1.15E-06** |
| ENSG00000163882.12 | **POLR2H** | 5437 | protein_coding | **1219.686** | -0.598 | -0.180 | **0.661** | 2.08E-07 | **1.16E-06** |
| ENSG00000227507.4 | **LTB** | 4050 | protein_coding | **661.848** | -0.820 | -0.247 | **0.566** | 2.08E-07 | **1.16E-06** |
| ENSG00000104218.18 | **CSPP1** | 79848 | protein_coding | **262.223** | 0.794 | 0.239 | **1.734** | 2.08E-07 | **1.16E-06** |
| ENSG00000100201.24 | **DDX17** | 10521 | protein_coding | **7671.326** | -0.664 | -0.200 | **0.631** | 2.08E-07 | **1.16E-06** |
| ENSG00000142920.20 | **AZIN2** | 113451 | protein_coding | **24.328** | 2.837 | 0.854 | **7.148** | 2.10E-07 | **1.17E-06** |
| ENSG00000135930.17 | **EIF4E2** | 9470 | protein_coding | **1236.674** | -0.607 | -0.183 | **0.657** | 2.12E-07 | **1.18E-06** |
| ENSG00000143436.12 | **MRPL9** | 65005 | protein_coding | **856.784** | -0.839 | -0.252 | **0.559** | 2.13E-07 | **1.18E-06** |
| ENSG00000100490.12 | **CDKL1** | 8814 | protein_coding | **115.516** | -1.388 | -0.418 | **0.382** | 2.13E-07 | **1.18E-06** |
| ENSG00000305128.1 | **ENSG00000305128** | | lncRNA | **8.249** | 5.186 | 1.561 | **36.396** | 2.14E-07 | **1.19E-06** |
| ENSG00000300428.1 | **ENSG0000(** | 10798709! | lncRNA | **50.015** | 1.179 | 0.355 | **2.264** | 2.14E-07 | **1.19E-06** |
| ENSG00000104904.13 | **OAZ1** | 4946 | protein_coding | **19935.901** | 0.535 | 0.161 | **1.449** | 2.16E-07 | **1.20E-06** |
| ENSG00000105643.13 | **ARRDC2** | 27106 | protein_coding | **597.676** | 0.705 | 0.212 | **1.630** | 2.17E-07 | **1.20E-06** |
| ENSG00000176974.24 | **SHMT1** | 6470 | protein_coding | **503.369** | -0.684 | -0.206 | **0.622** | 2.17E-07 | **1.20E-06** |
| ENSG00000104998.6 | **IL27RA** | 9466 | protein_coding | **595.615** | -0.865 | -0.260 | **0.549** | 2.18E-07 | **1.21E-06** |
| ENSG00000070010.21 | **UFD1** | 7353 | protein_coding | **924.295** | -0.576 | -0.173 | **0.671** | 2.18E-07 | **1.21E-06** |
| ENSG00000294024.1 | **ENSG0000(** | 12490405( | lncRNA | **71.841** | 1.170 | 0.352 | **2.251** | 2.19E-07 | **1.21E-06** |
| ENSG00000106399.13 | **RPA3** | 6119 | protein_coding | **587.526** | -0.674 | -0.203 | **0.627** | 2.21E-07 | **1.22E-06** |

| | | | | | | | | | |
|---|---|---|---|---|---|---|---|---|---|
| ENSG00000126602.13 | **TRAP1** | 10131 | protein_coding | **2814.135** | -0.561 | -0.169 | **0.678** | 2.22E-07 | **1.23E-06** |
| ENSG00000253490.5 | **LINC02099** | 10192945( | lncRNA | **17.659** | 2.267 | 0.682 | **4.812** | 2.23E-07 | **1.23E-06** |
| ENSG00000123374.12 | **CDK2** | 1017 | protein_coding | **1079.770** | -0.913 | -0.275 | **0.531** | 2.23E-07 | **1.24E-06** |
| ENSG00000159231.8 | **CBR3** | 874 | protein_coding | **16.936** | 2.358 | 0.710 | **5.127** | 2.25E-07 | **1.24E-06** |
| ENSG00000248485.4 | **PCP4L1** | 654790 | protein_coding | **34.053** | 2.106 | 0.634 | **4.306** | 2.25E-07 | **1.25E-06** |
| ENSG00000288844.2 | **ENSG00000288844** | | lncRNA | **14.466** | 2.354 | 0.709 | **5.112** | 2.26E-07 | **1.25E-06** |
| ENSG00000103121.11 | **CMC2** | 56942 | protein_coding | **725.935** | -0.513 | -0.154 | **0.701** | 2.27E-07 | **1.26E-06** |
| ENSG00000173465.10 | **ZNRD2** | 10534 | protein_coding | **711.701** | 0.622 | 0.187 | **1.539** | 2.29E-07 | **1.26E-06** |
| ENSG00000139620.15 | **KANSL2** | 54934 | protein_coding | **411.324** | -0.672 | -0.202 | **0.627** | 2.29E-07 | **1.27E-06** |
| ENSG00000177303.12 | **CASKIN2** | 57513 | protein_coding | **47.580** | 1.930 | 0.581 | **3.811** | 2.30E-07 | **1.27E-06** |
| ENSG00000115661.16 | **STK16** | 8576 | protein_coding | **537.965** | 0.576 | 0.173 | **1.491** | 2.32E-07 | **1.28E-06** |
| ENSG00000100906.14 | **NFKBIA** | 4792 | protein_coding | **6340.711** | 0.872 | 0.262 | **1.830** | 2.32E-07 | **1.28E-06** |
| ENSG00000141376.25 | **BCAS3** | 54828 | protein_coding | **199.895** | 1.072 | 0.323 | **2.102** | 2.34E-07 | **1.29E-06** |
| ENSG00000102572.17 | **STK24** | 8428 | protein_coding | **3143.587** | 0.676 | 0.203 | **1.597** | 2.34E-07 | **1.29E-06** |
| ENSG00000183605.19 | **SFXN4** | 119559 | protein_coding | **588.969** | -0.803 | -0.242 | **0.573** | 2.35E-07 | **1.29E-06** |
| ENSG00000275367.1 | **ENSG00000275367** | | lncRNA | **11.160** | 3.138 | 0.945 | **8.804** | 2.37E-07 | **1.31E-06** |
| ENSG00000160298.20 | **C21orf58** | 54058 | protein_coding | **487.286** | -1.045 | -0.315 | **0.485** | 2.39E-07 | **1.32E-06** |
| ENSG00000272735.3 | **ENSG00000272735** | | lncRNA | **13.189** | 2.510 | 0.756 | **5.696** | 2.39E-07 | **1.32E-06** |
| ENSG00000178950.20 | **GAK** | 2580 | protein_coding | **3050.190** | 0.580 | 0.175 | **1.495** | 2.40E-07 | **1.32E-06** |
| ENSG00000167081.20 | **PBX3** | 5090 | protein_coding | **531.111** | -0.687 | -0.207 | **0.621** | 2.40E-07 | **1.32E-06** |
| ENSG00000196391.12 | **ZNF774** | 342132 | protein_coding | **42.896** | 1.548 | 0.466 | **2.925** | 2.41E-07 | **1.33E-06** |
| ENSG00000166557.16 | **TMED3** | 23423 | protein_coding | **855.057** | -0.470 | -0.142 | **0.722** | 2.41E-07 | **1.33E-06** |
| ENSG00000167711.16 | **SERPINF2** | 5345 | protein_coding | **20.178** | 1.911 | 0.575 | **3.762** | 2.42E-07 | **1.33E-06** |
| ENSG00000197302.13 | **KRABD5** | 124411 | protein_coding | **244.605** | -0.575 | -0.173 | **0.671** | 2.42E-07 | **1.33E-06** |
| ENSG00000006555.13 | **TTC22** | 55001 | protein_coding | **66.076** | 1.026 | 0.309 | **2.037** | 2.42E-07 | **1.33E-06** |
| ENSG00000103260.11 | **METRN** | 79006 | protein_coding | **140.299** | 0.949 | 0.286 | **1.931** | 2.43E-07 | **1.34E-06** |
| ENSG00000110536.16 | **PTPMT1** | 114971 | protein_coding | **541.725** | -0.690 | -0.208 | **0.620** | 2.44E-07 | **1.34E-06** |
| ENSG00000185155.13 | **MIXL1** | 83881 | protein_coding | **48.737** | 1.576 | 0.474 | **2.982** | 2.44E-07 | **1.34E-06** |
| ENSG00000214719.14 | **ENSG00000214719** | | lncRNA | **119.543** | -1.017 | -0.306 | **0.494** | 2.44E-07 | **1.34E-06** |
| ENSG00000295762.1 | **ENSG00000295762** | | lncRNA | **88.075** | 0.886 | 0.267 | **1.849** | 2.47E-07 | **1.35E-06** |
| ENSG00000100890.18 | **PRORP** | 9692 | protein_coding | **405.466** | -0.795 | -0.239 | **0.576** | 2.47E-07 | **1.36E-06** |

| | | | | | | | | | |
|---|---|---|---|---|---|---|---|---|---|
| ENSG00000153560.14 | **UBP1** | 7342 | protein_coding | **1031.350** | 0.813 | 0.245 | **1.757** | 2.51E-07 | **1.38E-06** |
| ENSG00000111237.20 | **VPS29** | 51699 | protein_coding | **1267.760** | -0.572 | -0.172 | **0.673** | 2.51E-07 | **1.38E-06** |
| ENSG00000162959.16 | **MEMO1** | 51072 | protein_coding | **724.922** | -0.486 | -0.146 | **0.714** | 2.53E-07 | **1.39E-06** |
| ENSG00000271551.2 | **ENSG0000(** | 115308161 | lncRNA | **26.892** | 1.729 | 0.521 | **3.316** | 2.53E-07 | **1.39E-06** |
| ENSG00000295233.1 | **ENSG00000295233** | | lncRNA | **28.210** | 1.974 | 0.594 | **3.928** | 2.53E-07 | **1.39E-06** |
| ENSG00000111647.15 | **BLTP3B** | 23074 | protein_coding | **139.382** | -0.956 | -0.288 | **0.515** | 2.53E-07 | **1.39E-06** |
| ENSG00000162576.19 | **MXRA8** | 54587 | protein_coding | **9.954** | 2.973 | 0.895 | **7.850** | 2.55E-07 | **1.40E-06** |
| ENSG00000165097.18 | **KDM1B** | 221656 | protein_coding | **176.131** | 0.895 | 0.269 | **1.860** | 2.58E-07 | **1.41E-06** |
| ENSG00000128309.20 | **MPST** | 4357 | protein_coding | **980.112** | 0.759 | 0.228 | **1.692** | 2.58E-07 | **1.41E-06** |
| ENSG00000186469.11 | **GNG2** | 54331 | protein_coding | **564.627** | -0.843 | -0.254 | **0.557** | 2.63E-07 | **1.44E-06** |
| ENSG00000176454.17 | **LPCAT4** | 254531 | protein_coding | **2177.435** | 0.613 | 0.184 | **1.529** | 2.66E-07 | **1.45E-06** |
| ENSG00000111110.14 | **PPM1H** | 57460 | protein_coding | **67.548** | 1.960 | 0.590 | **3.889** | 2.67E-07 | **1.46E-06** |
| ENSG00000227070.3 | **EPS15-AS1** | 10537872( | lncRNA | **19.485** | -1.746 | -0.526 | **0.298** | 2.69E-07 | **1.47E-06** |
| ENSG00000289579.1 | **TMZR1** | | lncRNA | **10.594** | 3.289 | 0.990 | **9.773** | 2.73E-07 | **1.49E-06** |
| ENSG00000110057.11 | **UNC93B1** | 81622 | protein_coding | **1735.530** | -0.628 | -0.189 | **0.647** | 2.73E-07 | **1.49E-06** |
| ENSG00000166780.13 | **BMERB1** | 89927 | protein_coding | **21.230** | 2.282 | 0.687 | **4.864** | 2.74E-07 | **1.50E-06** |
| ENSG00000254997.4 | **KRTAP5-9** | 3846 | protein_coding | **15.095** | 2.735 | 0.823 | **6.659** | 2.75E-07 | **1.50E-06** |
| ENSG00000150961.17 | **SEC24D** | 9871 | protein_coding | **450.380** | -0.821 | -0.247 | **0.566** | 2.75E-07 | **1.50E-06** |
| ENSG00000172724.13 | **CCL19** | 6363 | protein_coding | **28.746** | 2.166 | 0.652 | **4.487** | 2.75E-07 | **1.50E-06** |
| ENSG00000197343.13 | **ZNF655** | 79027 | protein_coding | **794.793** | -0.569 | -0.171 | **0.674** | 2.76E-07 | **1.50E-06** |
| ENSG00000205592.16 | **MUC19** | 283463 | protein_coding | **235.936** | 1.029 | 0.310 | **2.041** | 2.77E-07 | **1.51E-06** |
| ENSG00000183808.14 | **RBM12B** | 389677 | protein_coding | **287.690** | -0.727 | -0.219 | **0.604** | 2.78E-07 | **1.52E-06** |
| ENSG00000287133.2 | **ENSG00000287133** | | lncRNA | **11.619** | 3.615 | 1.088 | **12.256** | 2.79E-07 | **1.52E-06** |
| ENSG00000187522.17 | **HSPA14** | 51182 | protein_coding | **520.608** | -0.621 | -0.187 | **0.650** | 2.80E-07 | **1.53E-06** |
| ENSG00000292277.1 | **ENSG00000292277** | | protein_coding | **71.215** | -1.205 | -0.363 | **0.434** | 2.80E-07 | **1.53E-06** |
| ENSG00000246763.8 | **RGMB-AS1** | 503569 | lncRNA | **16.908** | 2.344 | 0.706 | **5.078** | 2.81E-07 | **1.53E-06** |
| ENSG00000143845.16 | **ETNK2** | 55224 | protein_coding | **7.449** | 4.371 | 1.316 | **20.693** | 2.81E-07 | **1.53E-06** |
| ENSG00000185414.22 | **MRPL30** | 51263 | protein_coding | **609.278** | -0.449 | -0.135 | **0.733** | 2.81E-07 | **1.53E-06** |
| ENSG00000165644.13 | **COMTD1** | 118881 | protein_coding | **411.497** | -0.934 | -0.281 | **0.524** | 2.82E-07 | **1.54E-06** |
| ENSG00000256060.4 | **TRAPPC2B** | 10597 | protein_coding | **128.096** | -0.729 | -0.220 | **0.603** | 2.83E-07 | **1.54E-06** |
| ENSG00000169139.14 | **UBE2V2** | 7336 | protein_coding | **513.175** | -0.568 | -0.171 | **0.674** | 2.84E-07 | **1.54E-06** |

| | | | | | | | | | |
|---|---|---|---|---|---|---|---|---|---|
| ENSG00000198171.15 | **DDRGK1** | 65992 | protein_coding | **878.915** | -0.573 | -0.172 | **0.672** | 2.84E-07 | **1.54E-06** |
| ENSG00000101849.20 | **TBL1X** | 6907 | protein_coding | **669.678** | 0.674 | 0.203 | **1.595** | 2.85E-07 | **1.55E-06** |
| ENSG00000308858.1 | **ENSG00000308858** | | lncRNA | **29.597** | -1.986 | -0.598 | **0.252** | 2.86E-07 | **1.55E-06** |
| ENSG00000250565.9 | **ATP6V1E2** | 90423 | protein_coding | **143.785** | 0.875 | 0.264 | **1.835** | 2.87E-07 | **1.56E-06** |
| ENSG00000188549.15 | **CCDC9B** | 388115 | protein_coding | **30.419** | 2.043 | 0.615 | **4.121** | 2.87E-07 | **1.56E-06** |
| ENSG00000179958.11 | **DCTPP1** | 79077 | protein_coding | **319.481** | -0.836 | -0.252 | **0.560** | 2.88E-07 | **1.56E-06** |
| ENSG00000111711.11 | **GOLT1B** | 51026 | protein_coding | **316.684** | -1.095 | -0.330 | **0.468** | 2.91E-07 | **1.58E-06** |
| ENSG00000203896.12 | **LIME1** | 54923 | protein_coding | **683.251** | -1.186 | -0.357 | **0.439** | 2.92E-07 | **1.58E-06** |
| ENSG00000100994.14 | **PYGB** | 5834 | protein_coding | **965.385** | 0.541 | 0.163 | **1.455** | 2.92E-07 | **1.58E-06** |
| ENSG00000168116.17 | **KIAA1586** | 57691 | protein_coding | **157.640** | -0.831 | -0.250 | **0.562** | 2.92E-07 | **1.59E-06** |
| ENSG00000125910.6 | **S1PR4** | 8698 | protein_coding | **468.307** | -0.733 | -0.221 | **0.602** | 2.93E-07 | **1.59E-06** |
| ENSG00000185010.16 | **F8** | 2157 | protein_coding | **122.757** | 1.303 | 0.392 | **2.468** | 2.96E-07 | **1.60E-06** |
| ENSG00000044115.23 | **CTNNA1** | 1495 | protein_coding | **2452.033** | 0.570 | 0.172 | **1.485** | 2.97E-07 | **1.61E-06** |
| ENSG00000308185.1 | **ENSG0000(** | 12490531Z | lncRNA | **514.346** | -0.608 | -0.183 | **0.656** | 2.97E-07 | **1.61E-06** |
| ENSG00000150779.13 | **TIMM8B** | 26521 | protein_coding | **751.700** | -0.636 | -0.192 | **0.643** | 2.98E-07 | **1.61E-06** |
| ENSG00000065361.19 | **ERBB3** | 2065 | protein_coding | **14.402** | 2.721 | 0.819 | **6.594** | 2.98E-07 | **1.61E-06** |
| ENSG00000198216.15 | **CACNA1E** | 777 | protein_coding | **751.646** | 0.640 | 0.193 | **1.558** | 2.99E-07 | **1.62E-06** |
| ENSG00000198734.14 | **F5** | 2153 | protein_coding | **25.136** | 1.941 | 0.584 | **3.840** | 3.00E-07 | **1.62E-06** |
| ENSG00000120910.17 | **PPP3CC** | 5533 | protein_coding | **822.822** | -0.619 | -0.186 | **0.651** | 3.00E-07 | **1.62E-06** |
| ENSG00000079931.17 | **MOXD1** | 26002 | protein_coding | **397.167** | 0.993 | 0.299 | **1.990** | 3.01E-07 | **1.63E-06** |
| ENSG00000134852.17 | **CLOCK** | 9575 | protein_coding | **160.633** | -1.141 | -0.344 | **0.453** | 3.02E-07 | **1.63E-06** |
| ENSG00000023191.19 | **RNH1** | 6050 | protein_coding | **2091.989** | -0.585 | -0.176 | **0.667** | 3.04E-07 | **1.64E-06** |
| ENSG00000301105.1 | **ENSG0000(** | 12490461Z | lncRNA | **6.302** | 4.666 | 1.405 | **25.395** | 3.04E-07 | **1.64E-06** |
| ENSG00000125977.9 | **EIF2S2** | 8894 | protein_coding | **2384.616** | -0.660 | -0.199 | **0.633** | 3.04E-07 | **1.64E-06** |
| ENSG00000118162.16 | **KPTN** | 11133 | protein_coding | **258.930** | 0.631 | 0.190 | **1.548** | 3.06E-07 | **1.65E-06** |
| ENSG00000183155.7 | **RABIF** | 5877 | protein_coding | **158.397** | -0.675 | -0.203 | **0.626** | 3.07E-07 | **1.66E-06** |
| ENSG00000171940.15 | **ZNF217** | 7764 | protein_coding | **1190.615** | 0.823 | 0.248 | **1.769** | 3.08E-07 | **1.66E-06** |
| ENSG00000228863.9 | **ENSG00000228863** | | lncRNA | **132.013** | 0.695 | 0.209 | **1.619** | 3.08E-07 | **1.66E-06** |
| ENSG00000090932.13 | **DLL3** | 10683 | protein_coding | **37.734** | 1.895 | 0.571 | **3.720** | 3.08E-07 | **1.66E-06** |
| ENSG00000304038.1 | **ENSG00000304038** | | lncRNA | **11.006** | 2.881 | 0.867 | **7.365** | 3.12E-07 | **1.68E-06** |
| ENSG00000283083.3 | **LINC02498** | 10272352E | lncRNA | **19.446** | 3.000 | 0.903 | **7.998** | 3.12E-07 | **1.68E-06** |

| | | | | | | | | | |
|---|---|---|---|---|---|---|---|---|---|
| ENSG00000168411.16 | **RFWD3** | 55159 | protein_coding | **738.109** | -0.612 | -0.184 | **0.654** | 3.17E-07 | **1.71E-06** |
| ENSG00000130787.16 | **HIP1R** | 9026 | protein_coding | **683.204** | 2.237 | 0.674 | **4.715** | 3.19E-07 | **1.72E-06** |
| ENSG00000164008.19 | **C1orf50** | 79078 | protein_coding | **283.411** | -0.611 | -0.184 | **0.655** | 3.20E-07 | **1.73E-06** |
| ENSG00000149599.18 | **DUSP15** | 128853 | protein_coding | **36.785** | 1.610 | 0.485 | **3.052** | 3.21E-07 | **1.73E-06** |
| ENSG00000221821.4 | **PEX39** | 441150 | protein_coding | **126.019** | 0.816 | 0.246 | **1.761** | 3.21E-07 | **1.73E-06** |
| ENSG00000134864.13 | **GGACT** | 87769 | protein_coding | **139.102** | 1.057 | 0.318 | **2.081** | 3.22E-07 | **1.73E-06** |
| ENSG00000164070.14 | **HSPA4L** | 22824 | protein_coding | **230.410** | -0.886 | -0.267 | **0.541** | 3.23E-07 | **1.74E-06** |
| ENSG00000144736.16 | **SHQ1** | 55164 | protein_coding | **349.806** | 0.569 | 0.171 | **1.483** | 3.23E-07 | **1.74E-06** |
| ENSG00000142494.15 | **SLC47A1** | 55244 | protein_coding | **21.980** | 2.894 | 0.871 | **7.432** | 3.23E-07 | **1.74E-06** |
| ENSG00000168028.17 | **RPSA** | 3921 | protein_coding | **43433.175** | -0.480 | -0.145 | **0.717** | 3.24E-07 | **1.74E-06** |
| ENSG00000196502.17 | **SULT1A1** | 6817 | protein_coding | **22.815** | 1.656 | 0.499 | **3.152** | 3.29E-07 | **1.77E-06** |
| ENSG00000242612.9 | **DECR2** | 26063 | protein_coding | **237.473** | 0.613 | 0.185 | **1.530** | 3.29E-07 | **1.77E-06** |
| ENSG00000177599.15 | **ZNF491** | 126069 | protein_coding | **42.639** | 1.319 | 0.397 | **2.494** | 3.33E-07 | **1.79E-06** |
| ENSG00000086504.20 | **MRPL28** | 10573 | protein_coding | **1629.914** | -0.523 | -0.157 | **0.696** | 3.34E-07 | **1.79E-06** |
| ENSG00000111348.11 | **ARHGDIB** | 397 | protein_coding | **9394.110** | -1.024 | -0.308 | **0.492** | 3.34E-07 | **1.79E-06** |
| ENSG00000185585.22 | **OLFML2A** | 169611 | protein_coding | **46.604** | 1.417 | 0.427 | **2.671** | 3.34E-07 | **1.79E-06** |
| ENSG00000059758.10 | **CDK17** | 5128 | protein_coding | **659.435** | 1.157 | 0.348 | **2.230** | 3.35E-07 | **1.80E-06** |
| ENSG00000137078.12 | **SIT1** | 27240 | protein_coding | **652.835** | -0.930 | -0.280 | **0.525** | 3.36E-07 | **1.81E-06** |
| ENSG00000138382.17 | **METTL5** | 29081 | protein_coding | **546.878** | -0.556 | -0.167 | **0.680** | 3.43E-07 | **1.84E-06** |
| ENSG00000243156.10 | **MICAL3** | 57553 | protein_coding | **220.412** | 1.605 | 0.483 | **3.041** | 3.44E-07 | **1.84E-06** |
| ENSG00000244563.1 | **RPS26P19** | | processed_pseudo | **82.425** | -1.411 | -0.425 | **0.376** | 3.46E-07 | **1.86E-06** |
| ENSG00000272894.6 | **MIOS-DT** | | lncRNA | **86.882** | 1.370 | 0.412 | **2.585** | 3.49E-07 | **1.87E-06** |
| ENSG00000169599.15 | **NFU1** | 27247 | protein_coding | **411.021** | -0.556 | -0.167 | **0.680** | 3.51E-07 | **1.88E-06** |
| ENSG00000119900.10 | **OGFRL1** | 79627 | protein_coding | **150.117** | -0.790 | -0.238 | **0.578** | 3.52E-07 | **1.89E-06** |
| ENSG00000120256.13 | **LRP11** | 84918 | protein_coding | **10.817** | 3.237 | 0.974 | **9.426** | 3.52E-07 | **1.89E-06** |
| ENSG00000178209.19 | **PLEC** | 5339 | protein_coding | **3051.221** | 0.618 | 0.186 | **1.534** | 3.54E-07 | **1.89E-06** |
| ENSG00000290988.1 | **ZNF781** | 163115 | lncRNA | **22.482** | 1.971 | 0.593 | **3.920** | 3.54E-07 | **1.89E-06** |
| ENSG00000141699.13 | **RETREG3** | 162427 | protein_coding | **1211.414** | 0.416 | 0.125 | **1.334** | 3.54E-07 | **1.90E-06** |
| ENSG00000161031.15 | **PGLYRP2** | 114770 | protein_coding | **12.289** | 3.076 | 0.926 | **8.431** | 3.55E-07 | **1.90E-06** |
| ENSG00000171827.13 | **ZNF570** | 148268 | protein_coding | **80.741** | -0.866 | -0.261 | **0.549** | 3.55E-07 | **1.90E-06** |
| ENSG00000131149.21 | **GSE1** | 23199 | protein_coding | **1435.495** | 0.541 | 0.163 | **1.455** | 3.55E-07 | **1.90E-06** |

| | | | | | | | | | |
|---|---|---|---|---|---|---|---|---|---|
| ENSG00000136643.14 | **RPS6KC1** | 26750 | protein_coding | **222.411** | 0.724 | 0.218 | **1.652** | 3.57E-07 | **1.91E-06** |
| ENSG00000225231.3 | **LINC02470** | | lncRNA | **15.781** | 2.948 | 0.888 | **7.719** | 3.58E-07 | **1.91E-06** |
| ENSG00000143870.15 | **PDIA6** | 10130 | protein_coding | **2453.915** | -0.674 | -0.203 | **0.627** | 3.58E-07 | **1.91E-06** |
| ENSG00000187033.9 | **SAMD7** | 344658 | protein_coding | **9.449** | 3.198 | 0.963 | **9.180** | 3.59E-07 | **1.92E-06** |
| ENSG00000168081.11 | **PNOC** | 5368 | protein_coding | **362.484** | -1.210 | -0.364 | **0.432** | 3.61E-07 | **1.93E-06** |
| ENSG00000145284.13 | **SCD5** | 79966 | protein_coding | **45.284** | 1.329 | 0.400 | **2.513** | 3.62E-07 | **1.93E-06** |
| ENSG00000136807.15 | **CDK9** | 1025 | protein_coding | **1758.254** | 0.440 | 0.132 | **1.356** | 3.62E-07 | **1.93E-06** |
| ENSG00000149925.23 | **ALDOA** | 226 | protein_coding | **22580.485** | -0.787 | -0.237 | **0.579** | 3.63E-07 | **1.94E-06** |
| ENSG00000078900.18 | **TP73** | 7161 | protein_coding | **590.501** | -0.736 | -0.221 | **0.601** | 3.63E-07 | **1.94E-06** |
| ENSG00000065308.7 | **TRAM2** | 9697 | protein_coding | **493.122** | 0.620 | 0.187 | **1.537** | 3.64E-07 | **1.94E-06** |
| ENSG00000104946.16 | **TBC1D17** | 79735 | protein_coding | **1351.792** | 0.643 | 0.194 | **1.562** | 3.67E-07 | **1.95E-06** |
| ENSG00000105968.21 | **H2AZ2** | 94239 | protein_coding | **2035.531** | -0.492 | -0.148 | **0.711** | 3.67E-07 | **1.96E-06** |
| ENSG00000172239.16 | **PAIP1** | 10605 | protein_coding | **688.012** | -0.608 | -0.183 | **0.656** | 3.68E-07 | **1.96E-06** |
| ENSG00000121680.18 | **PEX16** | 9409 | protein_coding | **629.226** | 0.513 | 0.155 | **1.427** | 3.68E-07 | **1.96E-06** |
| ENSG00000204581.3 | **ACOXL-AS1** | 400997 | lncRNA | **7.512** | 5.073 | 1.527 | **33.668** | 3.70E-07 | **1.97E-06** |
| ENSG00000116752.8 | **BCAS2** | 10286 | protein_coding | **281.947** | -0.740 | -0.223 | **0.599** | 3.74E-07 | **1.99E-06** |
| ENSG00000290576.2 | **ENSG00000290576** | | lncRNA | **575.392** | 0.890 | 0.268 | **1.854** | 3.76E-07 | **2.00E-06** |
| ENSG00000265702.3 | **MARCHF1(** | 10537185 | lncRNA | **18.537** | 2.184 | 0.657 | **4.545** | 3.77E-07 | **2.01E-06** |
| ENSG00000280194.1 | **ENSG00000280194** | | TEC | **21.795** | 1.588 | 0.478 | **3.007** | 3.78E-07 | **2.01E-06** |
| ENSG00000255455.2 | **ENSG00000255455** | | lncRNA | **85.996** | 0.940 | 0.283 | **1.919** | 3.82E-07 | **2.03E-06** |
| ENSG00000288743.2 | **ENSG00000288743** | | lncRNA | **26.693** | 1.722 | 0.518 | **3.300** | 3.82E-07 | **2.03E-06** |
| ENSG00000229921.8 | **KIF25-AS1** | 10050587 | lncRNA | **14.587** | 3.054 | 0.919 | **8.304** | 3.84E-07 | **2.04E-06** |
| ENSG00000128408.10 | **RIBC2** | 26150 | protein_coding | **25.776** | -1.836 | -0.553 | **0.280** | 3.84E-07 | **2.04E-06** |
| ENSG00000149428.23 | **HYOU1** | 10525 | protein_coding | **2073.557** | -0.563 | -0.169 | **0.677** | 3.85E-07 | **2.04E-06** |
| ENSG00000141391.15 | **PRELID3A** | 10650 | protein_coding | **102.419** | -1.024 | -0.308 | **0.492** | 3.88E-07 | **2.06E-06** |
| ENSG00000023909.12 | **GCLM** | 2730 | protein_coding | **107.394** | -1.085 | -0.326 | **0.472** | 3.88E-07 | **2.06E-06** |
| ENSG00000295873.1 | **ENSG00000295873** | | lncRNA | **253.695** | -1.102 | -0.332 | **0.466** | 3.88E-07 | **2.06E-06** |
| ENSG00000116237.18 | **ICMT** | 23463 | protein_coding | **461.119** | -0.672 | -0.202 | **0.628** | 3.89E-07 | **2.07E-06** |
| ENSG00000054277.15 | **OPN3** | 23596 | protein_coding | **259.865** | -0.638 | -0.192 | **0.642** | 3.92E-07 | **2.08E-06** |
| ENSG00000266094.10 | **RASSF5** | 83593 | protein_coding | **1452.898** | -0.528 | -0.159 | **0.694** | 3.97E-07 | **2.11E-06** |
| ENSG00000119335.20 | **SET** | 6418 | protein_coding | **6985.069** | -0.580 | -0.175 | **0.669** | 3.98E-07 | **2.11E-06** |

| | | | | | | | | | |
|---|---|---|---|---|---|---|---|---|---|
| ENSG00000228794.13 | **LINC01128** | 643837 | lncRNA | **308.423** | -0.798 | -0.240 | **0.575** | 4.00E-07 | **2.12E-06** |
| ENSG00000183826.21 | **BTBD9** | 114781 | protein_coding | **154.722** | 0.714 | 0.215 | **1.641** | 4.01E-07 | **2.12E-06** |
| ENSG00000171984.17 | **SHLD1** | 149840 | protein_coding | **193.698** | -0.737 | -0.222 | **0.600** | 4.03E-07 | **2.13E-06** |
| ENSG00000007520.6 | **TSR3** | 115939 | protein_coding | **618.234** | -0.643 | -0.194 | **0.640** | 4.03E-07 | **2.14E-06** |
| ENSG00000168646.16 | **AXIN2** | 8313 | protein_coding | **51.730** | 1.441 | 0.434 | **2.715** | 4.05E-07 | **2.14E-06** |
| ENSG00000145736.17 | **GTF2H2** | 2966 | protein_coding | **194.693** | -0.783 | -0.236 | **0.581** | 4.08E-07 | **2.16E-06** |
| ENSG00000301183.1 | **ENSG00000301183** | | lncRNA | **77.535** | 1.191 | 0.359 | **2.284** | 4.09E-07 | **2.16E-06** |
| ENSG00000234741.11 | **GAS5** | 60674 | lncRNA | **7520.723** | -0.589 | -0.177 | **0.665** | 4.09E-07 | **2.16E-06** |
| ENSG00000172053.20 | **QARS1** | 5859 | protein_coding | **3448.196** | -0.622 | -0.187 | **0.650** | 4.11E-07 | **2.17E-06** |
| ENSG00000186104.13 | **CYP2R1** | 120227 | protein_coding | **493.408** | 0.786 | 0.237 | **1.724** | 4.12E-07 | **2.18E-06** |
| ENSG00000122592.9 | **HOXA7** | 3204 | protein_coding | **10.275** | 3.635 | 1.094 | **12.427** | 4.17E-07 | **2.20E-06** |
| ENSG00000276259.1 | **ENSG00000276259** | | lncRNA | **26.229** | 1.598 | 0.481 | **3.026** | 4.19E-07 | **2.21E-06** |
| ENSG00000167543.18 | **TP53I13** | 90313 | protein_coding | **759.759** | 0.619 | 0.186 | **1.536** | 4.19E-07 | **2.21E-06** |
| ENSG00000173511.12 | **VEGFB** | 7423 | protein_coding | **1057.377** | -0.787 | -0.237 | **0.579** | 4.25E-07 | **2.24E-06** |
| ENSG00000130005.15 | **GAMT** | 2593 | protein_coding | **457.342** | 0.825 | 0.248 | **1.771** | 4.25E-07 | **2.24E-06** |
| ENSG00000267666.3 | **ENSG00000267666** | | lncRNA | **9.318** | 3.073 | 0.925 | **8.414** | 4.28E-07 | **2.26E-06** |
| ENSG00000168060.17 | **NAALADL1** | 10004 | protein_coding | **172.530** | 1.009 | 0.304 | **2.013** | 4.28E-07 | **2.26E-06** |
| ENSG00000304458.1 | **ENSG00000304458** | | lncRNA | **25.080** | 1.455 | 0.438 | **2.742** | 4.32E-07 | **2.28E-06** |
| ENSG00000087266.20 | **SH3BP2** | 6452 | protein_coding | **2200.850** | -0.712 | -0.214 | **0.611** | 4.36E-07 | **2.30E-06** |
| ENSG00000115657.16 | **ABCB6** | 10058 | protein_coding | **747.531** | -0.937 | -0.282 | **0.522** | 4.40E-07 | **2.32E-06** |
| ENSG00000151779.16 | **NBAS** | 51594 | protein_coding | **434.453** | -0.492 | -0.148 | **0.711** | 4.43E-07 | **2.33E-06** |
| ENSG00000118369.15 | **USP35** | 57558 | protein_coding | **271.174** | 0.691 | 0.208 | **1.614** | 4.44E-07 | **2.34E-06** |
| ENSG00000237441.12 | **RGL2** | 5863 | protein_coding | **721.824** | 0.622 | 0.187 | **1.539** | 4.45E-07 | **2.34E-06** |
| ENSG00000235169.13 | **SMIM1** | 388588 | protein_coding | **29.127** | 1.635 | 0.492 | **3.105** | 4.46E-07 | **2.35E-06** |
| ENSG00000106245.13 | **BUD31** | 8896 | protein_coding | **1313.315** | -0.671 | -0.202 | **0.628** | 4.47E-07 | **2.35E-06** |
| ENSG00000175110.16 | **MRPS22** | 56945 | protein_coding | **604.383** | -0.464 | -0.140 | **0.725** | 4.51E-07 | **2.37E-06** |
| ENSG00000005156.14 | **LIG3** | 3980 | protein_coding | **421.646** | -0.728 | -0.219 | **0.604** | 4.51E-07 | **2.37E-06** |
| ENSG00000273604.2 | **EPOP** | 100170841 | protein_coding | **528.308** | 0.719 | 0.216 | **1.646** | 4.51E-07 | **2.37E-06** |
| ENSG00000090612.24 | **ZNF268** | 10795 | protein_coding | **208.699** | -0.848 | -0.255 | **0.556** | 4.53E-07 | **2.38E-06** |
| ENSG00000122034.18 | **GTF3A** | 2971 | protein_coding | **1897.340** | -0.744 | -0.224 | **0.597** | 4.54E-07 | **2.39E-06** |
| ENSG00000176422.15 | **SPRYD4** | 283377 | protein_coding | **328.243** | -0.758 | -0.228 | **0.591** | 4.55E-07 | **2.39E-06** |

| | | | | | | | | | |
|---|---|---|---|---|---|---|---|---|---|
| ENSG00000277369.3 | **ENSG00000277369** | | lncRNA | **21.998** | 1.682 | 0.506 | **3.208** | 4.63E-07 | **2.43E-06** |
| ENSG00000289991.2 | **ENSG00000289991** | | lncRNA | **6.332** | 4.721 | 1.421 | **26.372** | 4.64E-07 | **2.43E-06** |
| ENSG00000287978.2 | **ENSG00000287978** | | lncRNA | **63.986** | 1.377 | 0.414 | **2.597** | 4.68E-07 | **2.46E-06** |
| ENSG00000272829.2 | **ENSG00000272829** | | lncRNA | **19.707** | 2.005 | 0.604 | **4.014** | 4.70E-07 | **2.47E-06** |
| ENSG00000186174.15 | **BCL9L** | 283149 | protein_coding | **256.546** | 1.027 | 0.309 | **2.037** | 4.71E-07 | **2.47E-06** |
| ENSG00000184481.19 | **FOXO4** | 4303 | protein_coding | **180.762** | 0.893 | 0.269 | **1.857** | 4.73E-07 | **2.48E-06** |
| ENSG00000186075.14 | **ZPBP2** | 124626 | protein_coding | **48.318** | 1.214 | 0.366 | **2.320** | 4.74E-07 | **2.48E-06** |
| ENSG00000243649.12 | **CFB** | 629 | protein_coding | **13.955** | 2.874 | 0.865 | **7.331** | 4.74E-07 | **2.49E-06** |
| ENSG00000009950.18 | **MLXIPL** | 51085 | protein_coding | **10.775** | 3.372 | 1.015 | **10.352** | 4.77E-07 | **2.50E-06** |
| ENSG00000108679.15 | **LGALS3BP** | 3959 | protein_coding | **219.922** | 1.283 | 0.386 | **2.433** | 4.77E-07 | **2.50E-06** |
| ENSG00000175866.18 | **BAIAP2** | 10458 | protein_coding | **69.108** | 1.237 | 0.372 | **2.357** | 4.78E-07 | **2.50E-06** |
| ENSG00000122025.17 | **FLT3** | 2322 | protein_coding | **31.221** | 1.921 | 0.578 | **3.788** | 4.84E-07 | **2.53E-06** |
| ENSG00000156931.19 | **VPS8** | 23355 | protein_coding | **599.644** | 0.552 | 0.166 | **1.467** | 4.84E-07 | **2.54E-06** |
| ENSG00000150967.21 | **ABCB9** | 23457 | protein_coding | **88.350** | 1.091 | 0.328 | **2.130** | 4.87E-07 | **2.55E-06** |
| ENSG00000159208.17 | **CIART** | 148523 | protein_coding | **60.072** | -1.093 | -0.329 | **0.469** | 4.89E-07 | **2.56E-06** |
| ENSG00000184178.19 | **SCFD2** | 152579 | protein_coding | **261.074** | -0.756 | -0.227 | **0.592** | 4.89E-07 | **2.56E-06** |
| ENSG00000088305.21 | **DNMT3B** | 1789 | protein_coding | **59.828** | 1.062 | 0.320 | **2.087** | 4.90E-07 | **2.56E-06** |
| ENSG00000223478.3 | **ZDHHC12-DT** | | lncRNA | **127.649** | 0.961 | 0.289 | **1.946** | 4.92E-07 | **2.57E-06** |
| ENSG00000196199.16 | **MPHOSPH** | 54737 | protein_coding | **1517.445** | 0.530 | 0.160 | **1.444** | 4.95E-07 | **2.59E-06** |
| ENSG00000013583.12 | **HEBP1** | 50865 | protein_coding | **48.755** | 1.512 | 0.455 | **2.851** | 4.98E-07 | **2.60E-06** |
| ENSG00000115073.10 | **ACTR1B** | 10120 | protein_coding | **1304.373** | -0.404 | -0.122 | **0.756** | 4.99E-07 | **2.60E-06** |
| ENSG00000185630.21 | **PBX1** | 5087 | protein_coding | **7.938** | 4.481 | 1.349 | **22.338** | 4.99E-07 | **2.61E-06** |
| ENSG00000289021.2 | **ENSG00000289021** | | lncRNA | **24.950** | 1.575 | 0.474 | **2.980** | 5.00E-07 | **2.61E-06** |
| ENSG00000123080.14 | **CDKN2C** | 1031 | protein_coding | **305.381** | -0.559 | -0.168 | **0.679** | 5.02E-07 | **2.62E-06** |
| ENSG00000111676.17 | **ATN1** | 1822 | protein_coding | **1525.485** | -0.735 | -0.221 | **0.601** | 5.11E-07 | **2.66E-06** |
| ENSG00000033011.16 | **ALG1** | 56052 | protein_coding | **259.998** | -0.792 | -0.239 | **0.577** | 5.11E-07 | **2.66E-06** |
| ENSG00000182511.14 | **FES** | 2242 | protein_coding | **144.369** | -1.237 | -0.372 | **0.424** | 5.14E-07 | **2.68E-06** |
| ENSG00000205744.12 | **DENND1C** | 79958 | protein_coding | **2571.083** | -0.846 | -0.255 | **0.556** | 5.16E-07 | **2.69E-06** |
| ENSG00000122085.19 | **MTERF4** | 130916 | protein_coding | **1136.503** | 0.628 | 0.189 | **1.545** | 5.16E-07 | **2.69E-06** |
| ENSG00000087274.21 | **ADD1** | 118 | protein_coding | **3498.780** | 0.631 | 0.190 | **1.549** | 5.19E-07 | **2.70E-06** |
| ENSG00000130413.18 | **STK33** | 65975 | protein_coding | **272.910** | 0.812 | 0.244 | **1.755** | 5.19E-07 | **2.70E-06** |

| | | | | | | | | | |
|---|---|---|---|---|---|---|---|---|---|
| ENSG00000235303.2 | **ENSG00000235303** | | lncRNA | **9.224** | 3.098 | 0.933 | **8.564** | 5.21E-07 | **2.71E-06** |
| ENSG00000213719.10 | **CLIC1** | 1192 | protein_coding | **5427.659** | -0.680 | -0.205 | **0.624** | 5.22E-07 | **2.72E-06** |
| ENSG00000235904.4 | **RBMS3-AS** | 10087397 | lncRNA | **11.922** | 3.768 | 1.134 | **13.626** | 5.23E-07 | **2.72E-06** |
| ENSG00000197261.13 | **C6orf141** | 135398 | protein_coding | **9.068** | 3.205 | 0.965 | **9.219** | 5.27E-07 | **2.74E-06** |
| ENSG00000139946.12 | **PELI2** | 57161 | protein_coding | **14.154** | 2.602 | 0.783 | **6.071** | 5.27E-07 | **2.74E-06** |
| ENSG00000090470.16 | **PDCD7** | 10081 | protein_coding | **593.963** | -0.492 | -0.148 | **0.711** | 5.27E-07 | **2.74E-06** |
| ENSG00000092531.12 | **SNAP23** | 8773 | protein_coding | **1124.232** | 0.810 | 0.244 | **1.753** | 5.29E-07 | **2.75E-06** |
| ENSG00000110422.14 | **HIPK3** | 10114 | protein_coding | **166.684** | -0.763 | -0.230 | **0.589** | 5.29E-07 | **2.75E-06** |
| ENSG00000138621.14 | **PPCDC** | 60490 | protein_coding | **580.963** | 0.609 | 0.183 | **1.525** | 5.30E-07 | **2.75E-06** |
| ENSG00000151883.22 | **PARP8** | 79668 | protein_coding | **551.025** | -0.837 | -0.252 | **0.560** | 5.30E-07 | **2.75E-06** |
| ENSG00000104047.17 | **DTWD1** | 56986 | protein_coding | **196.476** | -0.706 | -0.212 | **0.613** | 5.32E-07 | **2.76E-06** |
| ENSG00000141696.15 | **P3H4** | 10609 | protein_coding | **31.060** | 1.648 | 0.496 | **3.134** | 5.34E-07 | **2.77E-06** |
| ENSG00000163923.12 | **RPL39L** | 116832 | protein_coding | **139.426** | -0.823 | -0.248 | **0.565** | 5.35E-07 | **2.78E-06** |
| ENSG00000101004.18 | **NINL** | 22981 | protein_coding | **32.070** | 1.552 | 0.467 | **2.932** | 5.35E-07 | **2.78E-06** |
| ENSG00000271086.8 | **NAMA** | 10099656 | lncRNA | **37.590** | 1.445 | 0.435 | **2.723** | 5.37E-07 | **2.78E-06** |
| ENSG00000180613.11 | **GSX2** | 170825 | protein_coding | **7.173** | 3.911 | 1.177 | **15.045** | 5.39E-07 | **2.80E-06** |
| ENSG00000275700.8 | **AATF** | 26574 | protein_coding | **1258.865** | -0.550 | -0.166 | **0.683** | 5.40E-07 | **2.80E-06** |
| ENSG00000300095.1 | **ENSG00000300095** | | lncRNA | **12.557** | 2.485 | 0.748 | **5.599** | 5.51E-07 | **2.86E-06** |
| ENSG00000149798.7 | **CDC42EP2** | 10435 | protein_coding | **62.916** | 1.581 | 0.476 | **2.991** | 5.53E-07 | **2.86E-06** |
| ENSG00000033867.20 | **SLC4A7** | 9497 | protein_coding | **106.476** | -0.979 | -0.295 | **0.507** | 5.53E-07 | **2.86E-06** |
| ENSG00000141258.15 | **SGSM2** | 9905 | protein_coding | **1739.394** | 0.767 | 0.231 | **1.702** | 5.54E-07 | **2.87E-06** |
| ENSG00000113273.19 | **ARSB** | 411 | protein_coding | **86.937** | -1.048 | -0.315 | **0.484** | 5.58E-07 | **2.89E-06** |
| ENSG00000173200.15 | **PARP15** | 165631 | protein_coding | **1558.261** | -0.762 | -0.229 | **0.590** | 5.58E-07 | **2.89E-06** |
| ENSG00000103089.11 | **FA2H** | 79152 | protein_coding | **13.227** | 2.941 | 0.885 | **7.678** | 5.59E-07 | **2.89E-06** |
| ENSG00000267228.7 | **ENSG00000267228** | | protein_coding | **69.664** | -1.569 | -0.472 | **0.337** | 5.61E-07 | **2.90E-06** |
| ENSG00000137288.12 | **UQCC2** | 84300 | protein_coding | **798.224** | -0.734 | -0.221 | **0.601** | 5.65E-07 | **2.92E-06** |
| ENSG00000168010.13 | **ATG16L2** | 89849 | protein_coding | **3566.922** | 0.677 | 0.204 | **1.598** | 5.68E-07 | **2.94E-06** |
| ENSG00000152213.6 | **ARL11** | 115761 | protein_coding | **122.716** | -0.819 | -0.247 | **0.567** | 5.70E-07 | **2.95E-06** |
| ENSG00000181652.20 | **ATG9B** | 285973 | protein_coding | **48.322** | 1.719 | 0.517 | **3.292** | 5.71E-07 | **2.95E-06** |
| ENSG00000188886.6 | **ASTL** | 431705 | protein_coding | **44.419** | 1.451 | 0.437 | **2.734** | 5.71E-07 | **2.95E-06** |
| ENSG00000206559.10 | **ZCWPW2** | 152098 | protein_coding | **16.141** | 2.146 | 0.646 | **4.426** | 5.76E-07 | **2.98E-06** |

| | | | | | | | | | |
|---|---|---|---|---|---|---|---|---|---|
| ENSG00000082212.15 | **ME2** | 4200 | protein_coding | **996.708** | -0.743 | -0.224 | **0.597** | 5.77E-07 | **2.98E-06** |
| ENSG00000230415.2 | **LINC01786** | | lncRNA | **21.990** | 1.671 | 0.503 | **3.183** | 5.77E-07 | **2.98E-06** |
| ENSG00000224032.10 | **EPB41L4A-AS1** | | lncRNA | **808.459** | -0.816 | -0.246 | **0.568** | 5.79E-07 | **2.99E-06** |
| ENSG00000105568.21 | **PPP2R1A** | 5518 | protein_coding | **3552.658** | -0.700 | -0.211 | **0.616** | 5.79E-07 | **2.99E-06** |
| ENSG00000158458.23 | **NRG2** | 9542 | protein_coding | **129.901** | 1.306 | 0.393 | **2.472** | 5.84E-07 | **3.01E-06** |
| ENSG00000175606.12 | **TMEM70** | 54968 | protein_coding | **284.324** | -0.591 | -0.178 | **0.664** | 5.86E-07 | **3.02E-06** |
| ENSG00000181577.17 | **LINC03040** | | protein_coding | **746.216** | -1.128 | -0.340 | **0.458** | 5.87E-07 | **3.02E-06** |
| ENSG00000052723.14 | **SIKE1** | 80143 | protein_coding | **504.889** | -1.184 | -0.356 | **0.440** | 5.88E-07 | **3.03E-06** |
| ENSG00000291023.1 | **ENSG00000291023** | | lncRNA | **10.709** | 3.684 | 1.109 | **12.856** | 5.99E-07 | **3.09E-06** |
| ENSG00000133657.19 | **ATP13A3** | 79572 | protein_coding | **475.783** | -0.724 | -0.218 | **0.606** | 6.00E-07 | **3.09E-06** |
| ENSG00000141522.15 | **ARHGDIA** | 396 | protein_coding | **5366.000** | -0.796 | -0.240 | **0.576** | 6.02E-07 | **3.10E-06** |
| ENSG00000295401.1 | **ENSG00000295401** | | lncRNA | **20.904** | 2.252 | 0.678 | **4.765** | 6.03E-07 | **3.11E-06** |
| ENSG00000104881.18 | **PPP1R13L** | 10848 | protein_coding | **482.490** | 1.173 | 0.353 | **2.255** | 6.08E-07 | **3.13E-06** |
| ENSG00000110900.18 | **TSPAN11** | 441631 | protein_coding | **17.923** | -1.929 | -0.581 | **0.263** | 6.08E-07 | **3.13E-06** |
| ENSG00000123136.17 | **DDX39A** | 10212 | protein_coding | **3545.880** | -0.495 | -0.149 | **0.709** | 6.08E-07 | **3.13E-06** |
| ENSG00000214999.4 | **ENSG00000214999** | | lncRNA | **34.708** | 1.533 | 0.462 | **2.894** | 6.13E-07 | **3.15E-06** |
| ENSG00000293514.2 | **CUTALP** | 253039 | lncRNA | **601.684** | 0.769 | 0.232 | **1.704** | 6.13E-07 | **3.15E-06** |
| ENSG00000129474.19 | **AJUBA** | 84962 | protein_coding | **60.501** | 1.123 | 0.338 | **2.178** | 6.13E-07 | **3.15E-06** |
| ENSG00000132958.20 | **TPTE2** | 93492 | protein_coding | **38.199** | 1.300 | 0.391 | **2.462** | 6.19E-07 | **3.18E-06** |
| ENSG00000134109.12 | **EDEM1** | 9695 | protein_coding | **2321.088** | -0.918 | -0.276 | **0.529** | 6.21E-07 | **3.19E-06** |
| ENSG00000118007.15 | **STAG1** | 10274 | protein_coding | **358.166** | -0.632 | -0.190 | **0.645** | 6.22E-07 | **3.19E-06** |
| ENSG00000250222.5 | **TRIM7-AS2** | | lncRNA | **60.714** | 1.068 | 0.321 | **2.096** | 6.23E-07 | **3.20E-06** |
| ENSG00000262160.1 | **ENSG00000262160** | | lncRNA | **8.095** | 3.069 | 0.924 | **8.390** | 6.24E-07 | **3.20E-06** |
| ENSG00000167646.16 | **DNAAF3** | 352909 | protein_coding | **9.130** | 4.399 | 1.324 | **21.098** | 6.28E-07 | **3.22E-06** |
| ENSG00000171388.14 | **APLN** | 8862 | protein_coding | **9.931** | -3.583 | -1.079 | **0.083** | 6.31E-07 | **3.24E-06** |
| ENSG00000274641.2 | **H2BC17** | 8348 | protein_coding | **17.930** | -2.157 | -0.649 | **0.224** | 6.32E-07 | **3.24E-06** |
| ENSG00000228541.4 | **ENSG00000228541** | | lncRNA | **55.274** | 1.439 | 0.433 | **2.712** | 6.32E-07 | **3.24E-06** |
| ENSG00000130023.18 | **ERMARD** | 55780 | protein_coding | **803.850** | 0.587 | 0.177 | **1.502** | 6.32E-07 | **3.24E-06** |
| ENSG00000169100.16 | **SLC25A6** | 293 | protein_coding | **14063.104** | -0.558 | -0.168 | **0.679** | 6.35E-07 | **3.25E-06** |
| ENSG00000105821.17 | **DNAJC2** | 27000 | protein_coding | **560.848** | -0.673 | -0.203 | **0.627** | 6.40E-07 | **3.28E-06** |
| ENSG00000138639.19 | **ARHGAP24** | 83478 | protein_coding | **94.992** | 1.155 | 0.348 | **2.228** | 6.45E-07 | **3.31E-06** |

| | | | | | | | | | |
|---|---|---|---|---|---|---|---|---|---|
| ENSG00000290114.2 | **ENSG00000290114** | | lncRNA | **34.823** | 1.363 | 0.410 | **2.572** | 6.46E-07 | **3.31E-06** |
| ENSG00000197170.12 | **PSMD12** | 5718 | protein_coding | **564.098** | -0.509 | -0.153 | **0.703** | 6.48E-07 | **3.32E-06** |
| ENSG00000229891.5 | **LINC01315** | 10272377! | lncRNA | **6.723** | 3.754 | 1.130 | **13.490** | 6.48E-07 | **3.32E-06** |
| ENSG00000091732.20 | **ZC3HC1** | 51530 | protein_coding | **355.862** | -0.548 | -0.165 | **0.684** | 6.48E-07 | **3.32E-06** |
| ENSG00000225177.7 | **NHSL1-AS1** | 12490021 | lncRNA | **44.143** | 2.150 | 0.647 | **4.439** | 6.49E-07 | **3.32E-06** |
| ENSG00000249790.4 | **LINC02972** | | lncRNA | **108.561** | 1.404 | 0.423 | **2.646** | 6.49E-07 | **3.32E-06** |
| ENSG00000179593.18 | **ALOX15B** | 247 | protein_coding | **21.606** | 1.809 | 0.545 | **3.504** | 6.49E-07 | **3.32E-06** |
| ENSG00000196510.16 | **ANAPC7** | 51434 | protein_coding | **1159.419** | -0.439 | -0.132 | **0.737** | 6.50E-07 | **3.32E-06** |
| ENSG00000162654.11 | **GBP4** | 115361 | protein_coding | **253.317** | 1.049 | 0.316 | **2.070** | 6.51E-07 | **3.33E-06** |
| ENSG00000140830.12 | **TXNL4B** | 54957 | protein_coding | **270.672** | -0.695 | -0.209 | **0.618** | 6.53E-07 | **3.34E-06** |
| ENSG00000165698.18 | **SPACA9** | 11092 | protein_coding | **63.856** | 1.064 | 0.320 | **2.091** | 6.54E-07 | **3.34E-06** |
| ENSG00000258768.2 | **ENSG00000258768** | | lncRNA | **15.676** | 2.144 | 0.645 | **4.419** | 6.54E-07 | **3.34E-06** |
| ENSG00000111581.12 | **NUP107** | 57122 | protein_coding | **985.930** | -0.523 | -0.158 | **0.696** | 6.55E-07 | **3.34E-06** |
| ENSG00000280433.2 | **ENSG0000(** | 10272420( | artifact | **608.934** | 0.495 | 0.149 | **1.410** | 6.59E-07 | **3.36E-06** |
| ENSG00000158764.9 | **ITLN2** | 142683 | protein_coding | **9.305** | 3.503 | 1.055 | **11.338** | 6.59E-07 | **3.36E-06** |
| ENSG00000127084.21 | **FGD3** | 89846 | protein_coding | **620.818** | -0.760 | -0.229 | **0.590** | 6.62E-07 | **3.38E-06** |
| ENSG00000002822.18 | **MAD1L1** | 8379 | protein_coding | **807.506** | -0.874 | -0.263 | **0.546** | 6.63E-07 | **3.38E-06** |
| ENSG00000291263.2 | **SMG1P7** | 10050606( | lncRNA | **171.676** | -0.959 | -0.289 | **0.515** | 6.64E-07 | **3.38E-06** |
| ENSG00000144589.24 | **STK11IP** | 114790 | protein_coding | **451.785** | 0.579 | 0.174 | **1.493** | 6.64E-07 | **3.39E-06** |
| ENSG00000196951.14 | **SCOC-AS1** | 10012985ξ | lncRNA | **135.238** | 0.709 | 0.214 | **1.635** | 6.65E-07 | **3.39E-06** |
| ENSG00000108599.17 | **AKAP10** | 11216 | protein_coding | **317.004** | -0.578 | -0.174 | **0.670** | 6.67E-07 | **3.40E-06** |
| ENSG00000153157.15 | **SYCP2L** | 221711 | protein_coding | **143.847** | 2.534 | 0.763 | **5.792** | 6.70E-07 | **3.41E-06** |
| ENSG00000122122.11 | **SASH3** | 54440 | protein_coding | **2736.431** | -0.654 | -0.197 | **0.636** | 6.73E-07 | **3.42E-06** |
| ENSG00000062598.20 | **ELMO2** | 63916 | protein_coding | **1050.174** | 0.423 | 0.127 | **1.340** | 6.78E-07 | **3.45E-06** |
| ENSG00000276345.1 | **ENSG00000276345** | | protein_coding | **630.425** | -0.815 | -0.245 | **0.568** | 6.78E-07 | **3.45E-06** |
| ENSG00000102970.11 | **CCL17** | 6361 | protein_coding | **38.916** | 1.841 | 0.554 | **3.583** | 6.86E-07 | **3.49E-06** |
| ENSG00000167861.18 | **HID1** | 283987 | protein_coding | **94.944** | 1.174 | 0.353 | **2.256** | 6.88E-07 | **3.50E-06** |
| ENSG00000256646.8 | **ENSG00000256646** | | protein_coding | **241.378** | 0.715 | 0.215 | **1.641** | 6.89E-07 | **3.50E-06** |
| ENSG00000163444.14 | **TMEM183/** | 92703 | protein_coding | **1077.640** | -0.517 | -0.156 | **0.699** | 6.90E-07 | **3.51E-06** |
| ENSG00000152127.11 | **MGAT5** | 4249 | protein_coding | **908.399** | 0.697 | 0.210 | **1.622** | 6.93E-07 | **3.52E-06** |
| ENSG00000131876.20 | **SNRPA1** | 6627 | protein_coding | **1616.244** | -0.632 | -0.190 | **0.645** | 6.94E-07 | **3.53E-06** |

| | | | | | | | | | |
|---|---|---|---|---|---|---|---|---|---|
| ENSG00000299935.1 | **ENSG00000299935** | | lncRNA | **108.381** | -1.099 | -0.331 | **0.467** | 6.94E-07 | **3.53E-06** |
| ENSG00000114270.19 | **COL7A1** | 1294 | protein_coding | **307.115** | 1.040 | 0.313 | **2.056** | 6.97E-07 | **3.54E-06** |
| ENSG00000070915.12 | **SLC12A3** | 6559 | protein_coding | **39.813** | 1.723 | 0.519 | **3.301** | 7.04E-07 | **3.57E-06** |
| ENSG00000188687.19 | **SLC4A5** | 57835 | protein_coding | **61.958** | 1.897 | 0.571 | **3.724** | 7.05E-07 | **3.58E-06** |
| ENSG00000224418.2 | **STK24-AS1** | | lncRNA | **16.002** | 1.877 | 0.565 | **3.674** | 7.06E-07 | **3.58E-06** |
| ENSG00000240990.10 | **HOXA11-A** | 221883 | lncRNA | **10.510** | 3.353 | 1.010 | **10.221** | 7.07E-07 | **3.59E-06** |
| ENSG00000175518.8 | **UBQLNL** | 143630 | protein_coding | **85.207** | 0.968 | 0.291 | **1.956** | 7.10E-07 | **3.60E-06** |
| ENSG00000175048.19 | **ZDHHC14** | 79683 | protein_coding | **605.685** | 0.686 | 0.207 | **1.609** | 7.10E-07 | **3.60E-06** |
| ENSG00000224596.10 | **ZMIZ1-AS1** | 283050 | lncRNA | **20.127** | 2.061 | 0.620 | **4.173** | 7.11E-07 | **3.60E-06** |
| ENSG00000257052.2 | **PRPF19-DT** | | lncRNA | **23.149** | 1.713 | 0.516 | **3.279** | 7.14E-07 | **3.62E-06** |
| ENSG00000105520.13 | **PLPPR2** | 64748 | protein_coding | **235.640** | 0.887 | 0.267 | **1.849** | 7.20E-07 | **3.65E-06** |
| ENSG00000099940.14 | **SNAP29** | 9342 | protein_coding | **578.421** | -0.548 | -0.165 | **0.684** | 7.21E-07 | **3.65E-06** |
| ENSG00000119638.16 | **NEK9** | 91754 | protein_coding | **2007.937** | 0.426 | 0.128 | **1.344** | 7.22E-07 | **3.66E-06** |
| ENSG00000271538.8 | **LINC02427** | | lncRNA | **43.648** | 1.189 | 0.358 | **2.279** | 7.25E-07 | **3.67E-06** |
| ENSG00000170802.19 | **FOXN2** | 3344 | protein_coding | **180.691** | -0.879 | -0.265 | **0.544** | 7.26E-07 | **3.68E-06** |
| ENSG00000175899.17 | **A2M** | 2 | protein_coding | **17.722** | 2.465 | 0.742 | **5.521** | 7.28E-07 | **3.68E-06** |
| ENSG00000165506.16 | **DNAAF2** | 55172 | protein_coding | **253.235** | 0.582 | 0.175 | **1.497** | 7.32E-07 | **3.70E-06** |
| ENSG00000104517.15 | **UBR5** | 51366 | protein_coding | **3666.042** | 0.561 | 0.169 | **1.475** | 7.42E-07 | **3.75E-06** |
| ENSG00000243008.4 | **ENSG0000(** | 10192737! | lncRNA | **7.518** | 3.919 | 1.180 | **15.122** | 7.49E-07 | **3.78E-06** |
| ENSG00000135407.12 | **AVIL** | 10677 | protein_coding | **46.440** | 1.850 | 0.557 | **3.604** | 7.56E-07 | **3.82E-06** |
| ENSG00000149182.17 | **ARFGAP2** | 84364 | protein_coding | **1321.414** | -0.426 | -0.128 | **0.744** | 7.58E-07 | **3.83E-06** |
| ENSG00000077327.18 | **SPAG6** | 9576 | protein_coding | **51.296** | 1.902 | 0.573 | **3.737** | 7.59E-07 | **3.83E-06** |
| ENSG00000272617.3 | **ENSG00000272617** | | protein_coding | **101.213** | -0.869 | -0.262 | **0.547** | 7.62E-07 | **3.85E-06** |
| ENSG00000118596.14 | **SLC16A7** | 9194 | protein_coding | **58.262** | -1.239 | -0.373 | **0.424** | 7.63E-07 | **3.85E-06** |
| ENSG00000089022.16 | **MAPKAPK!** | 8550 | protein_coding | **510.575** | -0.617 | -0.186 | **0.652** | 7.64E-07 | **3.85E-06** |
| ENSG00000096696.18 | **DSP** | 1832 | protein_coding | **22.777** | 2.922 | 0.880 | **7.579** | 7.64E-07 | **3.85E-06** |
| ENSG00000122550.20 | **KLHL7** | 55975 | protein_coding | **170.318** | -0.820 | -0.247 | **0.567** | 7.65E-07 | **3.86E-06** |
| ENSG00000184451.5 | **CCR10** | 2826 | protein_coding | **101.783** | -0.842 | -0.254 | **0.558** | 7.66E-07 | **3.86E-06** |
| ENSG00000123297.22 | **TSFM** | 10102 | protein_coding | **546.013** | -0.525 | -0.158 | **0.695** | 7.78E-07 | **3.92E-06** |
| ENSG00000248596.8 | **ENSG00000248596** | | lncRNA | **6.756** | 4.916 | 1.480 | **30.193** | 7.78E-07 | **3.92E-06** |
| ENSG00000249096.8 | **LINC02362** | | lncRNA | **218.088** | -1.011 | -0.304 | **0.496** | 7.79E-07 | **3.93E-06** |

| | | | | | | | | | |
|---|---|---|---|---|---|---|---|---|---|
| ENSG00000136367.16 | **ZFHX2** | 85446 | protein_coding | **95.290** | 1.548 | 0.466 | **2.923** | 7.83E-07 | **3.94E-06** |
| ENSG00000196878.17 | **LAMB3** | 3914 | protein_coding | **34.207** | 1.814 | 0.546 | **3.516** | 7.84E-07 | **3.95E-06** |
| ENSG00000166963.14 | **MAP1A** | 4130 | protein_coding | **10.165** | 2.962 | 0.892 | **7.793** | 7.88E-07 | **3.96E-06** |
| ENSG00000167302.13 | **TEPSIN** | 146705 | protein_coding | **1076.342** | 0.553 | 0.166 | **1.467** | 7.91E-07 | **3.98E-06** |
| ENSG00000278662.7 | **GOLGA6L1** | 647042 | protein_coding | **120.526** | 1.016 | 0.306 | **2.022** | 7.92E-07 | **3.99E-06** |
| ENSG00000154222.17 | **CC2D1B** | 200014 | protein_coding | **739.447** | -0.442 | -0.133 | **0.736** | 7.95E-07 | **4.00E-06** |
| ENSG00000092140.18 | **G2E3** | 55632 | protein_coding | **141.642** | -1.159 | -0.349 | **0.448** | 7.96E-07 | **4.00E-06** |
| ENSG00000164627.19 | **KIF6** | 221458 | protein_coding | **51.503** | 1.417 | 0.426 | **2.670** | 7.97E-07 | **4.00E-06** |
| ENSG00000137673.11 | **MMP7** | 4316 | protein_coding | **51.871** | -1.803 | -0.543 | **0.287** | 7.99E-07 | **4.02E-06** |
| ENSG00000107742.16 | **SPOCK2** | 9806 | protein_coding | **82.337** | 1.679 | 0.505 | **3.201** | 8.03E-07 | **4.03E-06** |
| ENSG00000198563.16 | **DDX39B** | 7919 | protein_coding | **7199.997** | -0.570 | -0.172 | **0.673** | 8.07E-07 | **4.05E-06** |
| ENSG00000121644.21 | **DESI2** | 51029 | protein_coding | **192.361** | -0.803 | -0.242 | **0.573** | 8.10E-07 | **4.07E-06** |
| ENSG00000074219.16 | **TEAD2** | 8463 | protein_coding | **19.043** | 2.359 | 0.710 | **5.129** | 8.10E-07 | **4.07E-06** |
| ENSG00000297167.1 | **ENSG00000297167** | | lncRNA | **7.685** | 3.607 | 1.086 | **12.185** | 8.20E-07 | **4.12E-06** |
| ENSG00000115738.12 | **ID2** | 3398 | protein_coding | **603.148** | 0.807 | 0.243 | **1.749** | 8.26E-07 | **4.14E-06** |
| ENSG00000290842.2 | **CROCCP3** | 114819 | lncRNA | **75.429** | 1.200 | 0.361 | **2.297** | 8.30E-07 | **4.16E-06** |
| ENSG00000278224.9 | **PRICKLE4** | 29964 | protein_coding | **11.961** | 2.946 | 0.887 | **7.704** | 8.32E-07 | **4.17E-06** |
| ENSG00000174600.16 | **CMKLR1** | 1240 | protein_coding | **162.347** | 1.293 | 0.389 | **2.450** | 8.33E-07 | **4.18E-06** |
| ENSG00000176209.14 | **SMIM19** | 114926 | protein_coding | **583.993** | -0.506 | -0.152 | **0.704** | 8.33E-07 | **4.18E-06** |
| ENSG00000166783.24 | **MARF1** | 9665 | protein_coding | **1048.624** | 0.520 | 0.156 | **1.433** | 8.33E-07 | **4.18E-06** |
| ENSG00000270175.2 | **ENSG00000270175** | | lncRNA | **31.448** | 1.279 | 0.385 | **2.426** | 8.34E-07 | **4.18E-06** |
| ENSG00000165548.13 | **TMEM63C** | 57156 | protein_coding | **207.299** | 3.241 | 0.976 | **9.456** | 8.39E-07 | **4.20E-06** |
| ENSG00000152527.16 | **PLEKHH2** | 130271 | protein_coding | **25.196** | 2.370 | 0.714 | **5.171** | 8.39E-07 | **4.20E-06** |
| ENSG00000109332.22 | **UBE2D3** | 7323 | protein_coding | **2970.380** | -0.674 | -0.203 | **0.627** | 8.41E-07 | **4.21E-06** |
| ENSG00000172113.12 | **NME6** | 10201 | protein_coding | **275.110** | -0.783 | -0.236 | **0.581** | 8.49E-07 | **4.25E-06** |
| ENSG00000186594.17 | **MIR22HG** | 84981 | lncRNA | **149.345** | 0.925 | 0.279 | **1.899** | 8.54E-07 | **4.27E-06** |
| ENSG00000115425.16 | **PECR** | 55825 | protein_coding | **300.144** | 0.601 | 0.181 | **1.517** | 8.57E-07 | **4.29E-06** |
| ENSG00000277476.1 | **ENSG00000277476** | | lncRNA | **74.088** | 1.303 | 0.392 | **2.467** | 8.57E-07 | **4.29E-06** |
| ENSG00000150907.12 | **FOXO1** | 2308 | protein_coding | **321.658** | 0.770 | 0.232 | **1.705** | 8.58E-07 | **4.29E-06** |
| ENSG00000070087.17 | **PFN2** | 5217 | protein_coding | **249.146** | 1.056 | 0.318 | **2.080** | 8.64E-07 | **4.32E-06** |
| ENSG00000120438.14 | **TCP1** | 6950 | protein_coding | **2895.516** | -0.495 | -0.149 | **0.710** | 8.67E-07 | **4.33E-06** |

| | | | | | | | | | |
|---|---|---|---|---|---|---|---|---|---|
| ENSG00000198286.13 | CARD11 | 84433 | protein_coding | 1370.105 | -0.683 | -0.206 | 0.623 | 8.68E-07 | 4.34E-06 |
| ENSG00000186827.13 | TNFRSF4 | 7293 | protein_coding | 201.630 | 1.133 | 0.341 | 2.194 | 8.69E-07 | 4.34E-06 |
| ENSG00000187140.6 | FOXD3 | 27022 | protein_coding | 4.160 | 5.342 | 1.608 | 40.572 | 8.69E-07 | 4.34E-06 |
| ENSG00000105404.13 | RABAC1 | 10567 | protein_coding | 985.960 | 0.654 | 0.197 | 1.573 | 8.69E-07 | 4.34E-06 |
| ENSG00000108669.20 | CYTH1 | 9267 | protein_coding | 5344.821 | 0.549 | 0.165 | 1.463 | 8.70E-07 | 4.34E-06 |
| ENSG00000137054.18 | POLR1E | 64425 | protein_coding | 599.470 | -0.496 | -0.149 | 0.709 | 8.70E-07 | 4.34E-06 |
| ENSG00000072952.23 | IRAG1 | 10335 | protein_coding | 9.234 | 3.537 | 1.065 | 11.609 | 8.73E-07 | 4.35E-06 |
| ENSG00000084731.17 | KIF3C | 3797 | protein_coding | 84.171 | 1.280 | 0.385 | 2.429 | 8.75E-07 | 4.36E-06 |
| ENSG00000055955.19 | ITIH4 | 3700 | protein_coding | 286.803 | 1.097 | 0.330 | 2.140 | 8.79E-07 | 4.38E-06 |
| ENSG00000241852.13 | C8orf58 | 541565 | protein_coding | 147.709 | 0.671 | 0.202 | 1.592 | 8.80E-07 | 4.38E-06 |
| ENSG00000153914.18 | SREK1 | 140890 | protein_coding | 805.303 | -0.730 | -0.220 | 0.603 | 8.81E-07 | 4.39E-06 |
| ENSG00000230698.2 | ENSG00000230698 | | lncRNA | 12.189 | 2.276 | 0.685 | 4.844 | 8.81E-07 | 4.39E-06 |
| ENSG00000197249.16 | SERPINA1 | 5265 | protein_coding | 101.391 | 1.558 | 0.469 | 2.945 | 8.83E-07 | 4.40E-06 |
| ENSG00000232265.9 | LINC02805 | | lncRNA | 68.007 | 1.243 | 0.374 | 2.367 | 8.89E-07 | 4.43E-06 |
| ENSG00000169564.7 | PCBP1 | 5093 | protein_coding | 6871.536 | -0.424 | -0.128 | 0.745 | 8.93E-07 | 4.45E-06 |
| ENSG00000146038.14 | DCDC2 | 51473 | protein_coding | 8.561 | 3.673 | 1.106 | 12.755 | 8.95E-07 | 4.46E-06 |
| ENSG00000108474.18 | PIGL | 9487 | protein_coding | 515.359 | -0.678 | -0.204 | 0.625 | 8.98E-07 | 4.47E-06 |
| ENSG00000087152.18 | ATXN7L3 | 56970 | protein_coding | 1420.956 | 0.489 | 0.147 | 1.404 | 8.99E-07 | 4.47E-06 |
| ENSG00000168528.14 | SERINC2 | 347735 | protein_coding | 395.334 | -1.010 | -0.304 | 0.497 | 9.07E-07 | 4.51E-06 |
| ENSG00000143614.13 | GATAD2B | 57459 | protein_coding | 1408.986 | 0.593 | 0.179 | 1.508 | 9.07E-07 | 4.51E-06 |
| ENSG00000168404.15 | MLKL | 197259 | protein_coding | 293.240 | -0.653 | -0.197 | 0.636 | 9.12E-07 | 4.53E-06 |
| ENSG00000040199.20 | PHLPP2 | 23035 | protein_coding | 205.379 | 0.596 | 0.180 | 1.512 | 9.14E-07 | 4.54E-06 |
| ENSG00000100139.16 | MICALL1 | 85377 | protein_coding | 240.958 | -0.673 | -0.203 | 0.627 | 9.20E-07 | 4.57E-06 |
| ENSG00000106346.14 | USP42 | 84132 | protein_coding | 281.746 | 0.645 | 0.194 | 1.564 | 9.24E-07 | 4.59E-06 |
| ENSG00000123689.8 | G0S2 | 50486 | protein_coding | 141.916 | -1.128 | -0.340 | 0.457 | 9.26E-07 | 4.60E-06 |
| ENSG00000186350.13 | RXRA | 6256 | protein_coding | 949.664 | -0.646 | -0.195 | 0.639 | 9.35E-07 | 4.64E-06 |
| ENSG00000167434.12 | CA4 | 762 | protein_coding | 23.222 | 2.411 | 0.726 | 5.319 | 9.38E-07 | 4.65E-06 |
| ENSG00000166848.9 | TERF2IP | 54386 | protein_coding | 1562.177 | -0.495 | -0.149 | 0.710 | 9.43E-07 | 4.68E-06 |
| ENSG00000163013.13 | FBXO41 | 150726 | protein_coding | 1185.490 | -0.548 | -0.165 | 0.684 | 9.47E-07 | 4.69E-06 |
| ENSG00000173193.17 | PARP14 | 54625 | protein_coding | 3923.896 | -0.530 | -0.160 | 0.693 | 9.49E-07 | 4.71E-06 |
| ENSG00000004139.16 | SARM1 | 23098 | protein_coding | 504.129 | -0.543 | -0.164 | 0.686 | 9.53E-07 | 4.72E-06 |

| | | | | | | | | | |
|---|---|---|---|---|---|---|---|---|---|
| ENSG00000281189.1 | **GHET1** | 102723099 | lncRNA | **11.893** | 2.495 | 0.751 | **5.638** | 9.54E-07 | **4.73E-06** |
| ENSG00000110042.9 | **DTX4** | 23220 | protein_coding | **1181.302** | 0.877 | 0.264 | **1.836** | 9.54E-07 | **4.73E-06** |
| ENSG00000168556.8 | **ING2** | 3622 | protein_coding | **150.529** | -1.245 | -0.375 | **0.422** | 9.57E-07 | **4.74E-06** |
| ENSG00000076555.17 | **ACACB** | 32 | protein_coding | **69.912** | 1.642 | 0.494 | **3.120** | 9.58E-07 | **4.74E-06** |
| ENSG00000187244.14 | **BCAM** | 4059 | protein_coding | **10.292** | 4.136 | 1.245 | **17.577** | 9.60E-07 | **4.75E-06** |
| ENSG00000151135.12 | **TMEM263** | 90488 | protein_coding | **343.083** | -0.607 | -0.183 | **0.656** | 9.60E-07 | **4.75E-06** |
| ENSG00000059769.22 | **DNAJC25** | 548645 | protein_coding | **155.056** | -0.855 | -0.258 | **0.553** | 9.62E-07 | **4.76E-06** |
| ENSG00000175182.17 | **FAM131A** | 131408 | protein_coding | **111.399** | 0.946 | 0.285 | **1.926** | 9.62E-07 | **4.76E-06** |
| ENSG00000179115.13 | **FARSA** | 2193 | protein_coding | **1282.502** | -0.525 | -0.158 | **0.695** | 9.63E-07 | **4.76E-06** |
| ENSG00000100557.12 | **CCDC198** | 55195 | protein_coding | **308.760** | -1.102 | -0.332 | **0.466** | 9.67E-07 | **4.78E-06** |
| ENSG00000145916.21 | **RMND5B** | 64777 | protein_coding | **839.154** | -0.396 | -0.119 | **0.760** | 9.72E-07 | **4.80E-06** |
| ENSG00000103152.13 | **MPG** | 4350 | protein_coding | **1251.605** | -0.468 | -0.141 | **0.723** | 9.72E-07 | **4.80E-06** |
| ENSG00000204815.12 | **ODAD4** | 83538 | protein_coding | **30.037** | 2.037 | 0.613 | **4.104** | 9.73E-07 | **4.81E-06** |
| ENSG00000108370.19 | **RGS9** | 8787 | protein_coding | **227.835** | 1.068 | 0.321 | **2.096** | 9.77E-07 | **4.83E-06** |
| ENSG00000077312.11 | **SNRPA** | 6626 | protein_coding | **2768.163** | -0.674 | -0.203 | **0.627** | 9.79E-07 | **4.83E-06** |
| ENSG00000132286.14 | **TIMM10B** | 26515 | protein_coding | **593.374** | -0.615 | -0.185 | **0.653** | 9.91E-07 | **4.89E-06** |
| ENSG00000161202.22 | **DVL3** | 1857 | protein_coding | **1360.962** | -0.448 | -0.135 | **0.733** | 9.93E-07 | **4.90E-06** |
| ENSG00000175602.4 | **CCDC85B** | 11007 | protein_coding | **1220.050** | -0.675 | -0.203 | **0.626** | 9.96E-07 | **4.91E-06** |
| ENSG00000197965.14 | **MPZL1** | 9019 | protein_coding | **778.398** | 0.431 | 0.130 | **1.348** | 1.00E-06 | **4.93E-06** |
| ENSG00000157350.15 | **ST3GAL2** | 6483 | protein_coding | **1417.477** | 0.668 | 0.201 | **1.589** | 1.00E-06 | **4.94E-06** |
| ENSG00000272677.4 | **HNRNPD-DT** | | lncRNA | **32.334** | 1.244 | 0.374 | **2.368** | 1.00E-06 | **4.94E-06** |
| ENSG00000241015.3 | **TPM3P9** | | transcribed_proce | **12.953** | -4.071 | -1.225 | **0.060** | 1.01E-06 | **4.97E-06** |
| ENSG00000146731.13 | **CCT6A** | 908 | protein_coding | **2643.228** | -0.514 | -0.155 | **0.700** | 1.01E-06 | **4.97E-06** |
| ENSG00000136603.17 | **SKIL** | 6498 | protein_coding | **170.786** | 1.006 | 0.303 | **2.008** | 1.01E-06 | **4.98E-06** |
| ENSG00000204525.20 | **HLA-C** | 3107 | protein_coding | **24036.222** | -0.483 | -0.145 | **0.716** | 1.01E-06 | **4.98E-06** |
| ENSG00000128159.14 | **TUBGCP6** | 85378 | protein_coding | **2447.065** | 0.515 | 0.155 | **1.429** | 1.01E-06 | **4.99E-06** |
| ENSG00000214894.8 | **LINC00243** | 401247 | lncRNA | **6.811** | 4.357 | 1.312 | **20.489** | 1.02E-06 | **5.01E-06** |
| ENSG00000291140.2 | **AMZ2P1** | 201283 | lncRNA | **282.534** | 0.964 | 0.290 | **1.951** | 1.03E-06 | **5.04E-06** |
| ENSG00000106479.12 | **ZNF862** | 643641 | protein_coding | **589.901** | 0.633 | 0.191 | **1.551** | 1.03E-06 | **5.05E-06** |
| ENSG00000141086.18 | **CTRL** | 1506 | protein_coding | **99.101** | -1.154 | -0.347 | **0.450** | 1.03E-06 | **5.06E-06** |
| ENSG00000106305.12 | **AIMP2** | 7965 | protein_coding | **488.880** | -0.578 | -0.174 | **0.670** | 1.03E-06 | **5.08E-06** |

| | | | | | | | | | |
|---|---|---|---|---|---|---|---|---|---|
| ENSG00000256087.9 | ZNF432 | 9668 | protein_coding | 182.764 | -0.672 | -0.202 | 0.627 | 1.03E-06 | 5.09E-06 |
| ENSG00000110169.14 | HPX | 3263 | protein_coding | 23.848 | 2.189 | 0.659 | 4.561 | 1.04E-06 | 5.09E-06 |
| ENSG00000168769.16 | TET2 | 54790 | protein_coding | 387.354 | -0.669 | -0.201 | 0.629 | 1.04E-06 | 5.10E-06 |
| ENSG00000280161.1 | ENSG00000280161 | | TEC | 34.841 | 1.269 | 0.382 | 2.409 | 1.05E-06 | 5.14E-06 |
| ENSG00000033800.16 | PIAS1 | 8554 | protein_coding | 1190.010 | 0.552 | 0.166 | 1.466 | 1.05E-06 | 5.15E-06 |
| ENSG00000088882.10 | CPXM1 | 56265 | protein_coding | 679.904 | -0.963 | -0.290 | 0.513 | 1.05E-06 | 5.16E-06 |
| ENSG00000100029.20 | PES1 | 23481 | protein_coding | 1303.441 | -0.442 | -0.133 | 0.736 | 1.05E-06 | 5.17E-06 |
| ENSG00000119729.14 | RHOQ | 23433 | protein_coding | 1069.246 | -0.603 | -0.182 | 0.658 | 1.07E-06 | 5.27E-06 |
| ENSG00000255252.4 | ENSG00000255252 | | lncRNA | 33.411 | 1.800 | 0.542 | 3.481 | 1.08E-06 | 5.29E-06 |
| ENSG00000198912.13 | C1orf174 | 339448 | protein_coding | 382.286 | -0.554 | -0.167 | 0.681 | 1.08E-06 | 5.29E-06 |
| ENSG00000111674.11 | ENO2 | 2026 | protein_coding | 5048.019 | -0.570 | -0.172 | 0.674 | 1.08E-06 | 5.30E-06 |
| ENSG00000129055.15 | ANAPC13 | 25847 | protein_coding | 421.047 | -0.535 | -0.161 | 0.690 | 1.08E-06 | 5.30E-06 |
| ENSG00000083099.13 | LYRM2 | 57226 | protein_coding | 398.467 | -0.635 | -0.191 | 0.644 | 1.08E-06 | 5.30E-06 |
| ENSG00000007376.10 | RPUSD1 | 113000 | protein_coding | 600.294 | -0.407 | -0.122 | 0.754 | 1.08E-06 | 5.31E-06 |
| ENSG00000138964.20 | PARVG | 64098 | protein_coding | 1632.264 | -0.829 | -0.250 | 0.563 | 1.09E-06 | 5.32E-06 |
| ENSG00000196337.16 | CGB7 | 94027 | protein_coding | 6.495 | 3.613 | 1.088 | 12.236 | 1.09E-06 | 5.33E-06 |
| ENSG00000223914.6 | LINC02471 | | lncRNA | 9.464 | 3.417 | 1.029 | 10.682 | 1.10E-06 | 5.38E-06 |
| ENSG00000099337.7 | KCNK6 | 9424 | protein_coding | 136.563 | -0.746 | -0.224 | 0.596 | 1.10E-06 | 5.39E-06 |
| ENSG00000128791.14 | TWSG1 | 57045 | protein_coding | 379.781 | -0.776 | -0.234 | 0.584 | 1.10E-06 | 5.41E-06 |
| ENSG00000198835.6 | GJC2 | 57165 | protein_coding | 11.062 | 2.213 | 0.666 | 4.636 | 1.11E-06 | 5.41E-06 |
| ENSG00000253824.3 | ENSG0000( | 12490199( | lncRNA | 31.353 | 1.512 | 0.455 | 2.852 | 1.11E-06 | 5.44E-06 |
| ENSG00000168824.17 | NSG1 | 27065 | protein_coding | 277.152 | -0.841 | -0.253 | 0.558 | 1.12E-06 | 5.48E-06 |
| ENSG00000114812.15 | VIPR1 | 7433 | protein_coding | 19.621 | 2.551 | 0.768 | 5.861 | 1.12E-06 | 5.49E-06 |
| ENSG00000058091.19 | CDK14 | 5218 | protein_coding | 175.793 | -0.707 | -0.213 | 0.613 | 1.12E-06 | 5.49E-06 |
| ENSG00000130158.15 | DOCK6 | 57572 | protein_coding | 608.800 | 0.933 | 0.281 | 1.909 | 1.13E-06 | 5.50E-06 |
| ENSG00000134882.18 | UBAC2 | 337867 | protein_coding | 827.806 | 0.641 | 0.193 | 1.560 | 1.13E-06 | 5.51E-06 |
| ENSG00000274376.5 | ADAMTS7P1 | | transcribed_unpro | 7.687 | 4.070 | 1.225 | 16.798 | 1.13E-06 | 5.52E-06 |
| ENSG00000103111.16 | MON1B | 22879 | protein_coding | 756.896 | -0.487 | -0.146 | 0.714 | 1.13E-06 | 5.52E-06 |
| ENSG00000291067.2 | ENSG00000291067 | | lncRNA | 351.605 | 0.699 | 0.210 | 1.623 | 1.14E-06 | 5.56E-06 |
| ENSG00000159871.16 | LYPD5 | 284348 | protein_coding | 16.675 | 2.572 | 0.774 | 5.946 | 1.14E-06 | 5.58E-06 |
| ENSG00000196557.15 | CACNA1H | 8912 | protein_coding | 13.855 | 2.888 | 0.869 | 7.404 | 1.15E-06 | 5.60E-06 |

| | | | | | | | | | |
|---|---|---|---|---|---|---|---|---|---|
| ENSG00000136848.20 | **DAB2IP** | 153090 | protein_coding | **23.107** | 1.607 | 0.484 | **3.046** | 1.15E-06 | **5.60E-06** |
| ENSG00000243716.14 | **NPIPB5** | 100132247 | protein_coding | **18716.196** | 0.453 | 0.136 | **1.369** | 1.15E-06 | **5.61E-06** |
| ENSG00000248663.8 | **LINC00992** | 124900192 | lncRNA | **8.253** | 3.815 | 1.148 | **14.072** | 1.15E-06 | **5.61E-06** |
| ENSG00000197191.7 | **CYSRT1** | 375791 | protein_coding | **38.216** | 1.292 | 0.389 | **2.448** | 1.15E-06 | **5.62E-06** |
| ENSG00000122376.14 | **SHLD2** | 54537 | protein_coding | **367.442** | -0.516 | -0.155 | **0.699** | 1.15E-06 | **5.62E-06** |
| ENSG00000048740.21 | **CELF2** | 10659 | protein_coding | **1542.466** | -0.669 | -0.201 | **0.629** | 1.16E-06 | **5.65E-06** |
| ENSG00000272688.3 | **ENSG00000272688** | | lncRNA | **160.797** | 1.036 | 0.312 | **2.050** | 1.17E-06 | **5.70E-06** |
| ENSG00000272758.8 | **WDR5B-DT** | | lncRNA | **137.170** | 0.891 | 0.268 | **1.855** | 1.17E-06 | **5.71E-06** |
| ENSG00000146676.10 | **PURB** | 5814 | protein_coding | **514.392** | -0.477 | -0.144 | **0.718** | 1.18E-06 | **5.75E-06** |
| ENSG00000166949.19 | **SMAD3** | 4088 | protein_coding | **1420.241** | 0.743 | 0.224 | **1.674** | 1.18E-06 | **5.76E-06** |
| ENSG00000197599.13 | **CCDC154** | 645811 | protein_coding | **36.012** | 1.665 | 0.501 | **3.171** | 1.18E-06 | **5.76E-06** |
| ENSG00000204348.12 | **DXO** | 1797 | protein_coding | **508.342** | -0.750 | -0.226 | **0.594** | 1.19E-06 | **5.77E-06** |
| ENSG00000113758.16 | **DBN1** | 1627 | protein_coding | **777.117** | 0.927 | 0.279 | **1.901** | 1.19E-06 | **5.77E-06** |
| ENSG00000101443.20 | **WFDC2** | 10406 | protein_coding | **303.106** | 1.123 | 0.338 | **2.179** | 1.19E-06 | **5.78E-06** |
| ENSG00000164687.13 | **FABP5** | 2171 | protein_coding | **832.771** | -0.668 | -0.201 | **0.630** | 1.19E-06 | **5.79E-06** |
| ENSG00000144401.16 | **METTL21A** | 151194 | protein_coding | **222.587** | -0.687 | -0.207 | **0.621** | 1.20E-06 | **5.81E-06** |
| ENSG00000203865.13 | **ATP1A1-AS** | 84852 | lncRNA | **110.948** | 0.733 | 0.221 | **1.662** | 1.20E-06 | **5.83E-06** |
| ENSG00000002745.14 | **WNT16** | 51384 | protein_coding | **9.221** | 3.405 | 1.025 | **10.591** | 1.21E-06 | **5.86E-06** |
| ENSG00000072840.15 | **EVC** | 2121 | protein_coding | **384.733** | 0.725 | 0.218 | **1.653** | 1.21E-06 | **5.86E-06** |
| ENSG00000093217.13 | **XYLB** | 9942 | protein_coding | **53.001** | -1.494 | -0.450 | **0.355** | 1.21E-06 | **5.88E-06** |
| ENSG00000227398.7 | **KIF9-AS1** | | lncRNA | **157.986** | 0.684 | 0.206 | **1.607** | 1.21E-06 | **5.89E-06** |
| ENSG00000162384.16 | **CZIB** | 54987 | protein_coding | **605.667** | -0.476 | -0.143 | **0.719** | 1.21E-06 | **5.89E-06** |
| ENSG00000249042.7 | **FAM151B-DT** | | lncRNA | **15.673** | 4.181 | 1.259 | **18.135** | 1.22E-06 | **5.89E-06** |
| ENSG00000156853.15 | **ZNF689** | 115509 | protein_coding | **112.282** | -0.709 | -0.213 | **0.612** | 1.22E-06 | **5.90E-06** |
| ENSG00000174177.15 | **CTU2** | 348180 | protein_coding | **334.505** | -0.826 | -0.249 | **0.564** | 1.22E-06 | **5.92E-06** |
| ENSG00000146223.18 | **RPL7L1** | 285855 | protein_coding | **1624.173** | -0.418 | -0.126 | **0.749** | 1.22E-06 | **5.93E-06** |
| ENSG00000253773.7 | **CFAP418-A** | 10061653( | lncRNA | **13.619** | 2.179 | 0.656 | **4.529** | 1.23E-06 | **5.94E-06** |
| ENSG00000116017.13 | **ARID3A** | 1820 | protein_coding | **1103.238** | -0.501 | -0.151 | **0.706** | 1.23E-06 | **5.96E-06** |
| ENSG00000158109.17 | **TPRG1L** | 127262 | protein_coding | **427.765** | 0.678 | 0.204 | **1.600** | 1.23E-06 | **5.96E-06** |
| ENSG00000120742.13 | **SERP1** | 27230 | protein_coding | **2020.708** | -0.679 | -0.204 | **0.625** | 1.25E-06 | **6.03E-06** |
| ENSG00000176148.18 | **TCP11L1** | 55346 | protein_coding | **158.334** | -0.773 | -0.233 | **0.585** | 1.25E-06 | **6.04E-06** |

| | | | | | | | | | |
|---|---|---|---|---|---|---|---|---|---|
| ENSG00000128596.19 | **CCDC136** | 64753 | protein_coding | **47.832** | 1.364 | 0.410 | **2.573** | 1.26E-06 | **6.07E-06** |
| ENSG00000123843.15 | **C4BPB** | 725 | protein_coding | **62.820** | 1.357 | 0.408 | **2.561** | 1.27E-06 | **6.12E-06** |
| ENSG00000167207.17 | **NOD2** | 64127 | protein_coding | **604.373** | 0.482 | 0.145 | **1.397** | 1.27E-06 | **6.13E-06** |
| ENSG00000117139.20 | **KDM5B** | 10765 | protein_coding | **430.581** | 0.791 | 0.238 | **1.730** | 1.27E-06 | **6.13E-06** |
| ENSG00000150764.16 | **DIXDC1** | 85458 | protein_coding | **17.713** | 2.579 | 0.776 | **5.975** | 1.27E-06 | **6.13E-06** |
| ENSG00000247077.10 | **PGAM5** | 192111 | protein_coding | **685.739** | -0.455 | -0.137 | **0.729** | 1.27E-06 | **6.14E-06** |
| ENSG00000232645.8 | **LINC01431** | | lncRNA | **30.132** | 1.449 | 0.436 | **2.729** | 1.27E-06 | **6.15E-06** |
| ENSG00000240386.4 | **LCE1F** | 353137 | protein_coding | **8.154** | -3.341 | -1.006 | **0.099** | 1.27E-06 | **6.16E-06** |
| ENSG00000143578.18 | **CREB3L4** | 148327 | protein_coding | **320.772** | 0.605 | 0.182 | **1.521** | 1.28E-06 | **6.16E-06** |
| ENSG00000270761.3 | **CD2AP-DT** | | lncRNA | **19.248** | 1.981 | 0.596 | **3.947** | 1.28E-06 | **6.19E-06** |
| ENSG00000285508.3 | **ENSG0000(** | 12870666( | protein_coding | **228.877** | -0.737 | -0.222 | **0.600** | 1.28E-06 | **6.20E-06** |
| ENSG00000135828.13 | **RNASEL** | 6041 | protein_coding | **114.215** | -0.761 | -0.229 | **0.590** | 1.29E-06 | **6.20E-06** |
| ENSG00000204390.12 | **HSPA1L** | 3305 | protein_coding | **50.477** | 1.263 | 0.380 | **2.400** | 1.29E-06 | **6.23E-06** |
| ENSG00000133313.17 | **CNDP2** | 55748 | protein_coding | **1822.903** | -0.412 | -0.124 | **0.752** | 1.29E-06 | **6.24E-06** |
| ENSG00000256116.2 | **ENSG00000256116** | | lncRNA | **16.814** | 1.766 | 0.532 | **3.401** | 1.30E-06 | **6.25E-06** |
| ENSG00000169905.15 | **TOR1AIP2** | 163590 | protein_coding | **680.161** | 0.439 | 0.132 | **1.356** | 1.30E-06 | **6.26E-06** |
| ENSG00000272767.2 | **JMJD1C-AS** | 84989 | lncRNA | **31.294** | 1.601 | 0.482 | **3.034** | 1.31E-06 | **6.29E-06** |
| ENSG00000242866.11 | **STRC** | 161497 | protein_coding | **63.867** | 1.620 | 0.488 | **3.075** | 1.31E-06 | **6.30E-06** |
| ENSG00000108375.15 | **RNF43** | 54894 | protein_coding | **82.640** | 1.284 | 0.387 | **2.435** | 1.31E-06 | **6.32E-06** |
| ENSG00000272602.7 | **ZNF595** | 152687 | protein_coding | **173.538** | -0.752 | -0.226 | **0.594** | 1.32E-06 | **6.36E-06** |
| ENSG00000253304.4 | **TMEM200I** | 399474 | protein_coding | **4.783** | 4.767 | 1.435 | **27.233** | 1.32E-06 | **6.36E-06** |
| ENSG00000101019.24 | **UQCC1** | 55245 | protein_coding | **1042.277** | -0.390 | -0.117 | **0.763** | 1.32E-06 | **6.37E-06** |
| ENSG00000198910.16 | **L1CAM** | 3897 | protein_coding | **69.699** | 1.745 | 0.525 | **3.352** | 1.32E-06 | **6.37E-06** |
| ENSG00000213923.15 | **CSNK1E** | 1454 | protein_coding | **1878.316** | 0.399 | 0.120 | **1.319** | 1.33E-06 | **6.38E-06** |
| ENSG00000288728.3 | **ENSG00000288728** | | lncRNA | **12.593** | -2.031 | -0.611 | **0.245** | 1.33E-06 | **6.40E-06** |
| ENSG00000100395.17 | **L3MBTL2** | 83746 | protein_coding | **746.066** | -0.518 | -0.156 | **0.698** | 1.33E-06 | **6.42E-06** |
| ENSG00000168397.19 | **ATG4B** | 23192 | protein_coding | **2266.195** | 0.498 | 0.150 | **1.412** | 1.34E-06 | **6.43E-06** |
| ENSG00000197961.13 | **ZNF121** | 7675 | protein_coding | **676.162** | -0.465 | -0.140 | **0.724** | 1.37E-06 | **6.58E-06** |
| ENSG00000261416.1 | **CORO1A-AS1** | | lncRNA | **54.290** | -1.440 | -0.433 | **0.369** | 1.38E-06 | **6.62E-06** |
| ENSG00000110876.13 | **SELPLG** | 6404 | protein_coding | **760.683** | 0.645 | 0.194 | **1.563** | 1.38E-06 | **6.63E-06** |
| ENSG00000157036.15 | **EXOG** | 9941 | protein_coding | **201.411** | -0.720 | -0.217 | **0.607** | 1.38E-06 | **6.64E-06** |

| | | | | | | | | | |
|---|---|---|---|---|---|---|---|---|---|
| ENSG00000166763.9 | **STRCP1** | | transcribed_unprc | **46.854** | 1.429 | 0.430 | **2.693** | 1.39E-06 | **6.67E-06** |
| ENSG00000066583.14 | **ISOC1** | 51015 | protein_coding | **303.190** | -0.680 | -0.205 | **0.624** | 1.39E-06 | **6.67E-06** |
| ENSG00000146733.16 | **PSPH** | 5723 | protein_coding | **503.992** | -0.867 | -0.261 | **0.548** | 1.40E-06 | **6.69E-06** |
| ENSG00000126070.22 | **AGO3** | 192669 | protein_coding | **350.624** | -0.673 | -0.203 | **0.627** | 1.40E-06 | **6.70E-06** |
| ENSG00000237550.8 | **ENSG00000237550** | | transcribed_proce | **879.672** | 14.547 | 4.379 | **########** | 1.40E-06 | **6.72E-06** |
| ENSG00000293391.1 | **LDC1P** | 149086 | lncRNA | **13.545** | -2.716 | -0.817 | **0.152** | 1.40E-06 | **6.73E-06** |
| ENSG00000205038.13 | **PKHD1L1** | 93035 | protein_coding | **245.254** | -0.568 | -0.171 | **0.675** | 1.41E-06 | **6.75E-06** |
| ENSG00000113971.23 | **NPHP3** | 27031 | protein_coding | **335.533** | 0.805 | 0.242 | **1.747** | 1.41E-06 | **6.75E-06** |
| ENSG00000278768.5 | **BACE1-AS** | 100379571 | lncRNA | **231.025** | 0.792 | 0.238 | **1.731** | 1.41E-06 | **6.77E-06** |
| ENSG00000198455.5 | **ZXDB** | 158586 | protein_coding | **52.540** | -1.110 | -0.334 | **0.463** | 1.41E-06 | **6.77E-06** |
| ENSG00000135480.19 | **KRT7** | 3855 | protein_coding | **69.408** | -1.366 | -0.411 | **0.388** | 1.42E-06 | **6.78E-06** |
| ENSG00000105647.22 | **PIK3R2** | 5296 | protein_coding | **433.831** | 0.568 | 0.171 | **1.483** | 1.42E-06 | **6.78E-06** |
| ENSG00000183780.14 | **SLC35F3** | 148641 | protein_coding | **35.888** | 1.960 | 0.590 | **3.891** | 1.42E-06 | **6.79E-06** |
| ENSG00000214575.12 | **CPEB1** | 64506 | protein_coding | **102.583** | 1.058 | 0.319 | **2.082** | 1.44E-06 | **6.89E-06** |
| ENSG00000135070.16 | **ISCA1** | 81689 | protein_coding | **260.554** | -0.581 | -0.175 | **0.668** | 1.45E-06 | **6.96E-06** |
| ENSG00000155719.19 | **OTOA** | 146183 | protein_coding | **14.119** | 3.070 | 0.924 | **8.399** | 1.46E-06 | **6.96E-06** |
| ENSG00000172216.7 | **CEBPB** | 1051 | protein_coding | **203.825** | -0.856 | -0.258 | **0.553** | 1.46E-06 | **6.99E-06** |
| ENSG00000270629.8 | **NBPF14** | 25832 | protein_coding | **244.772** | 0.783 | 0.236 | **1.720** | 1.47E-06 | **7.00E-06** |
| ENSG00000272720.1 | **ENSG00000272720** | | lncRNA | **10.001** | 2.319 | 0.698 | **4.990** | 1.47E-06 | **7.04E-06** |
| ENSG00000103335.24 | **PIEZO1** | 9780 | protein_coding | **2897.497** | -0.703 | -0.212 | **0.614** | 1.48E-06 | **7.06E-06** |
| ENSG00000186153.20 | **WWOX** | 51741 | protein_coding | **84.999** | -0.850 | -0.256 | **0.555** | 1.48E-06 | **7.07E-06** |
| ENSG00000152465.20 | **NMT2** | 9397 | protein_coding | **198.298** | 0.631 | 0.190 | **1.548** | 1.48E-06 | **7.07E-06** |
| ENSG00000233610.3 | **LINC00462** | 100129597 | lncRNA | **20.245** | 2.062 | 0.621 | **4.176** | 1.48E-06 | **7.07E-06** |
| ENSG00000289226.3 | **ENSG00000289226** | | lncRNA | **24.312** | 1.329 | 0.400 | **2.512** | 1.48E-06 | **7.07E-06** |
| ENSG00000107372.15 | **ZFAND5** | 7763 | protein_coding | **1401.123** | -0.446 | -0.134 | **0.734** | 1.48E-06 | **7.08E-06** |
| ENSG00000179627.11 | **ZBTB42** | 100128927 | protein_coding | **26.804** | 1.308 | 0.394 | **2.476** | 1.50E-06 | **7.15E-06** |
| ENSG00000119718.13 | **EIF2B2** | 8892 | protein_coding | **412.174** | -0.538 | -0.162 | **0.689** | 1.50E-06 | **7.16E-06** |
| ENSG00000293237.1 | **HTR7P1** | 93164 | TEC | **7.122** | 3.622 | 1.090 | **12.313** | 1.52E-06 | **7.24E-06** |
| ENSG00000177971.9 | **IMP3** | 55272 | protein_coding | **945.052** | -0.615 | -0.185 | **0.653** | 1.54E-06 | **7.32E-06** |
| ENSG00000197084.6 | **LCE1C** | 353133 | protein_coding | **9.467** | -2.514 | -0.757 | **0.175** | 1.54E-06 | **7.33E-06** |
| ENSG00000120833.16 | **SOCS2** | 8835 | protein_coding | **182.174** | 0.883 | 0.266 | **1.844** | 1.54E-06 | **7.34E-06** |

| | | | | | | | | | |
|---|---|---|---|---|---|---|---|---|---|
| ENSG00000198743.8 | **SLC5A3** | 6526 | protein_coding | **217.414** | -0.918 | -0.276 | **0.529** | 1.55E-06 | **7.37E-06** |
| ENSG00000110880.13 | **CORO1C** | 23603 | protein_coding | **2597.666** | -0.505 | -0.152 | **0.705** | 1.56E-06 | **7.41E-06** |
| ENSG00000235890.3 | **TSPEAR-AS1** | | lncRNA | **12.731** | 2.912 | 0.877 | **7.529** | 1.56E-06 | **7.43E-06** |
| ENSG00000103249.21 | **CLCN7** | 1186 | protein_coding | **1869.982** | 0.544 | 0.164 | **1.458** | 1.56E-06 | **7.45E-06** |
| ENSG00000108518.10 | **PFN1** | 5216 | protein_coding | **23907.125** | -0.658 | -0.198 | **0.634** | 1.57E-06 | **7.45E-06** |
| ENSG00000111716.16 | **LDHB** | 3945 | protein_coding | **10077.679** | -0.542 | -0.163 | **0.687** | 1.57E-06 | **7.45E-06** |
| ENSG00000130449.7 | **ZSWIM6** | 57688 | protein_coding | **123.069** | 1.212 | 0.365 | **2.316** | 1.57E-06 | **7.46E-06** |
| ENSG00000183496.7 | **MEX3B** | 84206 | protein_coding | **32.108** | 1.630 | 0.491 | **3.095** | 1.57E-06 | **7.46E-06** |
| ENSG00000165195.17 | **PIGA** | 5277 | protein_coding | **114.706** | -0.750 | -0.226 | **0.594** | 1.57E-06 | **7.48E-06** |
| ENSG00000101413.14 | **RPRD1B** | 58490 | protein_coding | **269.433** | -0.716 | -0.216 | **0.609** | 1.59E-06 | **7.57E-06** |
| ENSG00000138018.20 | **SELENOI** | 85465 | protein_coding | **373.600** | -0.650 | -0.196 | **0.637** | 1.60E-06 | **7.60E-06** |
| ENSG00000288985.3 | **ENSG00000288985** | | lncRNA | **15.771** | 2.543 | 0.766 | **5.829** | 1.60E-06 | **7.61E-06** |
| ENSG00000048471.16 | **SNX29** | 92017 | protein_coding | **1110.223** | 0.698 | 0.210 | **1.622** | 1.61E-06 | **7.66E-06** |
| ENSG00000105409.21 | **ATP1A3** | 478 | protein_coding | **50.465** | 1.573 | 0.474 | **2.975** | 1.61E-06 | **7.66E-06** |
| ENSG00000135482.9 | **ZC3H10** | 84872 | protein_coding | **147.691** | 0.860 | 0.259 | **1.815** | 1.61E-06 | **7.66E-06** |
| ENSG00000103852.15 | **TTC23** | 64927 | protein_coding | **189.462** | 0.670 | 0.202 | **1.591** | 1.62E-06 | **7.70E-06** |
| ENSG00000075914.15 | **EXOSC7** | 23016 | protein_coding | **936.704** | -0.545 | -0.164 | **0.686** | 1.63E-06 | **7.72E-06** |
| ENSG00000187514.19 | **PTMA** | 5757 | protein_coding | **26514.486** | -0.512 | -0.154 | **0.701** | 1.63E-06 | **7.74E-06** |
| ENSG00000150281.8 | **CTF1** | 1489 | protein_coding | **18.655** | 1.969 | 0.593 | **3.916** | 1.63E-06 | **7.75E-06** |
| ENSG00000117682.20 | **DHDDS** | 79947 | protein_coding | **383.899** | -0.522 | -0.157 | **0.697** | 1.63E-06 | **7.75E-06** |
| ENSG00000155016.20 | **CYP2U1** | 113612 | protein_coding | **90.320** | 0.915 | 0.275 | **1.886** | 1.64E-06 | **7.77E-06** |
| ENSG00000197410.15 | **DCHS2** | 54798 | protein_coding | **69.571** | 1.390 | 0.419 | **2.622** | 1.64E-06 | **7.79E-06** |
| ENSG00000149600.14 | **COMMD7** | 149951 | protein_coding | **491.891** | -0.451 | -0.136 | **0.732** | 1.64E-06 | **7.79E-06** |
| ENSG00000164024.14 | **METAP1** | 23173 | protein_coding | **782.697** | -0.450 | -0.135 | **0.732** | 1.65E-06 | **7.81E-06** |
| ENSG00000234350.8 | **ERICH2-DT** | 10192691[illegible] | lncRNA | **17.567** | 1.862 | 0.560 | **3.634** | 1.65E-06 | **7.82E-06** |
| ENSG00000105605.9 | **CACNG7** | 59284 | protein_coding | **3.951** | 5.289 | 1.592 | **39.104** | 1.65E-06 | **7.82E-06** |
| ENSG00000173153.19 | **ESRRA** | 2101 | protein_coding | **957.791** | 0.504 | 0.152 | **1.419** | 1.65E-06 | **7.82E-06** |
| ENSG00000197162.12 | **ZNF785** | 146540 | protein_coding | **172.923** | -0.688 | -0.207 | **0.621** | 1.66E-06 | **7.85E-06** |
| ENSG00000130024.17 | **PHF10** | 55274 | protein_coding | **554.311** | -0.466 | -0.140 | **0.724** | 1.67E-06 | **7.91E-06** |
| ENSG00000104140.9 | **RHOV** | 171177 | protein_coding | **469.497** | 0.860 | 0.259 | **1.815** | 1.68E-06 | **7.92E-06** |
| ENSG00000276293.7 | **PIP4K2B** | 8396 | protein_coding | **925.583** | -0.427 | -0.129 | **0.744** | 1.68E-06 | **7.92E-06** |

| | | | | | | | | | |
|---|---|---|---|---|---|---|---|---|---|
| ENSG00000174796.14 | **THAP6** | 152815 | protein_coding | **415.707** | 0.601 | 0.181 | **1.517** | 1.69E-06 | **7.97E-06** |
| ENSG00000112799.11 | **LY86** | 9450 | protein_coding | **1243.011** | -0.742 | -0.223 | **0.598** | 1.69E-06 | **8.00E-06** |
| ENSG00000117481.14 | **NSUN4** | 387338 | protein_coding | **665.005** | -0.687 | -0.207 | **0.621** | 1.70E-06 | **8.02E-06** |
| ENSG00000162461.10 | **SLC25A34** | 284723 | protein_coding | **46.017** | 1.273 | 0.383 | **2.417** | 1.70E-06 | **8.02E-06** |
| ENSG00000274225.2 | **HEPFAL** | | lncRNA | **16.010** | 1.823 | 0.549 | **3.538** | 1.70E-06 | **8.02E-06** |
| ENSG00000185813.13 | **PCYT2** | 5833 | protein_coding | **864.487** | -0.780 | -0.235 | **0.582** | 1.71E-06 | **8.05E-06** |
| ENSG00000170260.11 | **ZNF212** | 7988 | protein_coding | **372.453** | 0.497 | 0.150 | **1.412** | 1.71E-06 | **8.05E-06** |
| ENSG00000235172.10 | **LINC01366** | | lncRNA | **12.349** | 2.415 | 0.727 | **5.333** | 1.71E-06 | **8.07E-06** |
| ENSG00000102882.14 | **MAPK3** | 5595 | protein_coding | **669.746** | 0.523 | 0.157 | **1.437** | 1.72E-06 | **8.10E-06** |
| ENSG00000129515.22 | **SNX6** | 58533 | protein_coding | **589.939** | -0.578 | -0.174 | **0.670** | 1.72E-06 | **8.13E-06** |
| ENSG00000172795.18 | **DCP2** | 167227 | protein_coding | **550.235** | -0.422 | -0.127 | **0.746** | 1.73E-06 | **8.15E-06** |
| ENSG00000171595.16 | **DNAI2** | 64446 | protein_coding | **22.554** | 1.761 | 0.530 | **3.390** | 1.73E-06 | **8.17E-06** |
| ENSG00000268041.3 | **ERFL** | 390937 | protein_coding | **32.595** | 1.515 | 0.456 | **2.858** | 1.74E-06 | **8.20E-06** |
| ENSG00000080854.17 | **IGSF9B** | 22997 | protein_coding | **70.348** | 1.864 | 0.561 | **3.641** | 1.75E-06 | **8.25E-06** |
| ENSG00000128463.15 | **EMC4** | 51234 | protein_coding | **716.577** | -0.394 | -0.119 | **0.761** | 1.76E-06 | **8.28E-06** |
| ENSG00000122257.22 | **RBBP6** | 5930 | protein_coding | **1216.605** | -0.544 | -0.164 | **0.686** | 1.76E-06 | **8.29E-06** |
| ENSG00000106484.17 | **MEST** | 4232 | protein_coding | **72.234** | 1.486 | 0.447 | **2.802** | 1.76E-06 | **8.30E-06** |
| ENSG00000257556.5 | **LINC02298** | 12490339ٍ8 | lncRNA | **20.076** | 1.908 | 0.574 | **3.752** | 1.76E-06 | **8.30E-06** |
| ENSG00000147100.13 | **SLC16A2** | 6567 | protein_coding | **6.004** | 4.855 | 1.461 | **28.935** | 1.79E-06 | **8.41E-06** |
| ENSG00000095261.16 | **PSMD5** | 5711 | protein_coding | **401.854** | -0.458 | -0.138 | **0.728** | 1.80E-06 | **8.47E-06** |
| ENSG00000174915.14 | **PTDSS2** | 81490 | protein_coding | **647.998** | 0.428 | 0.129 | **1.345** | 1.80E-06 | **8.47E-06** |
| ENSG00000182810.7 | **DDX28** | 55794 | protein_coding | **302.123** | -0.596 | -0.179 | **0.661** | 1.81E-06 | **8.49E-06** |
| ENSG00000293879.1 | **ENSG00000293879** | | lncRNA | **20.639** | 2.865 | 0.863 | **7.286** | 1.81E-06 | **8.50E-06** |
| ENSG00000165434.10 | **PGM2L1** | 283209 | protein_coding | **10.509** | 2.432 | 0.732 | **5.397** | 1.81E-06 | **8.51E-06** |
| ENSG00000142197.15 | **DOP1B** | 9980 | protein_coding | **90.046** | 0.867 | 0.261 | **1.823** | 1.81E-06 | **8.51E-06** |
| ENSG00000255921.2 | **BCAT1-DT** | | lncRNA | **116.100** | 1.254 | 0.378 | **2.386** | 1.82E-06 | **8.55E-06** |
| ENSG00000136504.16 | **KAT7** | 11143 | protein_coding | **557.782** | -0.702 | -0.211 | **0.615** | 1.83E-06 | **8.59E-06** |
| ENSG00000161835.12 | **TAMALIN** | 160622 | protein_coding | **34.295** | 1.714 | 0.516 | **3.281** | 1.83E-06 | **8.61E-06** |
| ENSG00000177374.15 | **HIC1** | 3090 | protein_coding | **160.928** | -1.101 | -0.331 | **0.466** | 1.84E-06 | **8.64E-06** |
| ENSG00000171450.6 | **CDK5R2** | 8941 | protein_coding | **5.953** | 4.374 | 1.317 | **20.741** | 1.85E-06 | **8.66E-06** |
| ENSG00000272217.2 | **ENSG0000(** | 10537502ٍ2 | lncRNA | **32.809** | 1.256 | 0.378 | **2.389** | 1.85E-06 | **8.67E-06** |

| | | | | | | | | | |
|---|---|---|---|---|---|---|---|---|---|
| ENSG00000005812.13 | **FBXL3** | 26224 | protein_coding | **170.729** | -0.819 | -0.247 | **0.567** | 1.85E-06 | **8.67E-06** |
| ENSG00000125503.15 | **PPP1R12C** | 54776 | protein_coding | **2210.674** | 0.594 | 0.179 | **1.510** | 1.85E-06 | **8.67E-06** |
| ENSG00000165556.12 | **CDX2** | 1045 | protein_coding | **7.397** | 4.427 | 1.333 | **21.518** | 1.86E-06 | **8.70E-06** |
| ENSG00000196642.21 | **RABL6** | 55684 | protein_coding | **1616.273** | -0.563 | -0.169 | **0.677** | 1.86E-06 | **8.71E-06** |
| ENSG00000196935.12 | **SRGAP1** | 57522 | protein_coding | **40.521** | 2.398 | 0.722 | **5.271** | 1.87E-06 | **8.76E-06** |
| ENSG00000131055.7 | **COX4I2** | 84701 | protein_coding | **12.773** | 3.130 | 0.942 | **8.752** | 1.88E-06 | **8.79E-06** |
| ENSG00000205726.17 | **ITSN1** | 6453 | protein_coding | **76.449** | 1.401 | 0.422 | **2.641** | 1.89E-06 | **8.83E-06** |
| ENSG00000033050.11 | **ABCF2** | 10061 | protein_coding | **745.287** | -0.742 | -0.223 | **0.598** | 1.89E-06 | **8.84E-06** |
| ENSG00000099622.16 | **CIRBP** | 1153 | protein_coding | **3923.728** | -0.430 | -0.129 | **0.743** | 1.89E-06 | **8.85E-06** |
| ENSG00000274425.1 | **ENSG00000274425** | | lncRNA | **70.460** | 0.883 | 0.266 | **1.844** | 1.89E-06 | **8.86E-06** |
| ENSG00000105323.20 | **HNRNPUL1** | 11100 | protein_coding | **4188.169** | -0.371 | -0.112 | **0.773** | 1.90E-06 | **8.88E-06** |
| ENSG00000290679.2 | **ENSG00000290679** | | lncRNA | **7.215** | 4.634 | 1.395 | **24.823** | 1.92E-06 | **8.99E-06** |
| ENSG00000183850.15 | **ZNF730** | 10012954[illegible] | protein_coding | **27.449** | -1.646 | -0.495 | **0.320** | 1.94E-06 | **9.08E-06** |
| ENSG00000175536.8 | **LIPT2** | 387787 | protein_coding | **42.376** | -1.969 | -0.593 | **0.255** | 1.94E-06 | **9.08E-06** |
| ENSG00000128228.7 | **SDF2L1** | 23753 | protein_coding | **457.540** | -0.727 | -0.219 | **0.604** | 1.95E-06 | **9.12E-06** |
| ENSG00000174788.11 | **PCP2** | 126006 | protein_coding | **70.059** | 1.274 | 0.384 | **2.419** | 1.95E-06 | **9.14E-06** |
| ENSG00000167483.20 | **NIBAN3** | 199786 | protein_coding | **489.292** | 0.875 | 0.263 | **1.834** | 1.96E-06 | **9.18E-06** |
| ENSG00000264235.7 | **MYL12-AS[illegible]** | 10496839[illegible] | lncRNA | **12.173** | 2.523 | 0.760 | **5.749** | 1.97E-06 | **9.19E-06** |
| ENSG00000100625.10 | **SIX4** | 51804 | protein_coding | **12.235** | 3.260 | 0.981 | **9.582** | 1.97E-06 | **9.20E-06** |
| ENSG00000140043.14 | **PTGR2** | 145482 | protein_coding | **231.226** | 0.551 | 0.166 | **1.465** | 1.98E-06 | **9.25E-06** |
| ENSG00000304454.1 | **ENSG00000304454** | | lncRNA | **8.180** | 2.777 | 0.836 | **6.853** | 1.99E-06 | **9.28E-06** |
| ENSG00000153187.23 | **HNRNPU** | 3192 | protein_coding | **7271.311** | -0.474 | -0.143 | **0.720** | 1.99E-06 | **9.29E-06** |
| ENSG00000254911.3 | **SCARNA9** | 619383 | lncRNA | **66.840** | 1.034 | 0.311 | **2.048** | 2.01E-06 | **9.36E-06** |
| ENSG00000124783.16 | **SSR1** | 6745 | protein_coding | **1520.259** | -0.757 | -0.228 | **0.592** | 2.01E-06 | **9.36E-06** |
| ENSG00000171119.5 | **NRTN** | 4902 | protein_coding | **8.943** | 3.377 | 1.016 | **10.386** | 2.04E-06 | **9.51E-06** |
| ENSG00000137076.24 | **TLN1** | 7094 | protein_coding | **4794.715** | 0.364 | 0.109 | **1.287** | 2.05E-06 | **9.55E-06** |
| ENSG00000293920.1 | **ENSG00000293920** | | lncRNA | **25.416** | 2.141 | 0.644 | **4.410** | 2.05E-06 | **9.56E-06** |
| ENSG00000261061.1 | **ENSG00000261061** | | lncRNA | **58.670** | -1.578 | -0.475 | **0.335** | 2.05E-06 | **9.57E-06** |
| ENSG00000135631.19 | **RAB11FIP5** | 26056 | protein_coding | **18.330** | 1.876 | 0.565 | **3.670** | 2.05E-06 | **9.57E-06** |
| ENSG00000115008.8 | **IL1A** | 3552 | protein_coding | **46.820** | -1.384 | -0.417 | **0.383** | 2.07E-06 | **9.64E-06** |
| ENSG00000189376.14 | **C8orf76** | 84933 | protein_coding | **233.644** | -0.624 | -0.188 | **0.649** | 2.07E-06 | **9.65E-06** |

| | | | | | | | | | |
|---|---|---|---|---|---|---|---|---|---|
| ENSG00000187642.12 | **PERM1** | 84808 | protein_coding | **17.771** | 2.786 | 0.839 | **6.897** | 2.08E-06 | **9.68E-06** |
| ENSG00000089116.4 | **LHX5** | 64211 | protein_coding | **6.110** | 3.439 | 1.035 | **10.845** | 2.09E-06 | **9.71E-06** |
| ENSG00000291050.2 | **TEKT4P2** | 10013228ε | lncRNA | **233.812** | 0.850 | 0.256 | **1.803** | 2.09E-06 | **9.73E-06** |
| ENSG00000008517.21 | **IL32** | 9235 | protein_coding | **2550.502** | -0.908 | -0.273 | **0.533** | 2.09E-06 | **9.73E-06** |
| ENSG00000121289.20 | **CEP89** | 84902 | protein_coding | **293.229** | -0.825 | -0.248 | **0.564** | 2.12E-06 | **9.86E-06** |
| ENSG00000279601.1 | **ENSG00000279601** | | TEC | **32.739** | 1.617 | 0.487 | **3.068** | 2.12E-06 | **9.87E-06** |
| ENSG00000230266.3 | **XXYLT1-AS2** | | lncRNA | **182.119** | 0.772 | 0.232 | **1.707** | 2.13E-06 | **9.88E-06** |
| ENSG00000125637.18 | **PSD4** | 23550 | protein_coding | **2746.806** | -0.518 | -0.156 | **0.698** | 2.13E-06 | **9.89E-06** |
| ENSG00000161203.16 | **AP2M1** | 1173 | protein_coding | **2341.306** | -0.597 | -0.180 | **0.661** | 2.13E-06 | **9.91E-06** |
| ENSG00000272138.4 | **LINC01607** | 101927067 | lncRNA | **18.177** | 1.870 | 0.563 | **3.654** | 2.13E-06 | **9.91E-06** |
| ENSG00000121350.18 | **PYROXD1** | 79912 | protein_coding | **231.297** | -0.618 | -0.186 | **0.652** | 2.16E-06 | **1.00E-05** |
| ENSG00000073282.15 | **TP63** | 8626 | protein_coding | **92.209** | 1.301 | 0.392 | **2.464** | 2.18E-06 | **1.01E-05** |
| ENSG00000310191.1 | **ENSG00000310191** | | lncRNA | **16.203** | -1.994 | -0.600 | **0.251** | 2.19E-06 | **1.02E-05** |
| ENSG00000132467.4 | **UTP3** | 57050 | protein_coding | **1003.703** | 0.504 | 0.152 | **1.418** | 2.19E-06 | **1.02E-05** |
| ENSG00000203875.14 | **SNHG5** | | lncRNA | **2493.286** | -0.635 | -0.191 | **0.644** | 2.20E-06 | **1.02E-05** |
| ENSG00000105429.14 | **MEGF8** | 1954 | protein_coding | **588.257** | 0.568 | 0.171 | **1.483** | 2.21E-06 | **1.02E-05** |
| ENSG00000228063.5 | **LYPLAL1-D** | 643723 | lncRNA | **33.671** | 1.474 | 0.444 | **2.778** | 2.23E-06 | **1.03E-05** |
| ENSG00000123892.14 | **RAB38** | 23682 | protein_coding | **171.358** | -0.816 | -0.246 | **0.568** | 2.25E-06 | **1.04E-05** |
| ENSG00000114268.14 | **PFKFB4** | 5210 | protein_coding | **1705.883** | -0.771 | -0.232 | **0.586** | 2.25E-06 | **1.04E-05** |
| ENSG00000152766.8 | **ANKRD22** | 118932 | protein_coding | **12.236** | 2.776 | 0.836 | **6.851** | 2.26E-06 | **1.05E-05** |
| ENSG00000250240.7 | **ENSG00000250240** | | lncRNA | **7.135** | -3.546 | -1.067 | **0.086** | 2.26E-06 | **1.05E-05** |
| ENSG00000104824.20 | **HNRNPL** | 3191 | protein_coding | **5587.455** | -0.635 | -0.191 | **0.644** | 2.28E-06 | **1.05E-05** |
| ENSG00000167779.11 | **IGFBP6** | 3489 | protein_coding | **12.844** | 2.383 | 0.717 | **5.215** | 2.29E-06 | **1.06E-05** |
| ENSG00000163879.13 | **DNALI1** | 7802 | protein_coding | **8.251** | 3.873 | 1.166 | **14.647** | 2.30E-06 | **1.07E-05** |
| ENSG00000262484.3 | **CCER2** | 643669 | protein_coding | **43.458** | 1.554 | 0.468 | **2.937** | 2.33E-06 | **1.08E-05** |
| ENSG00000189067.17 | **LITAF** | 9516 | protein_coding | **2838.881** | 0.688 | 0.207 | **1.611** | 2.33E-06 | **1.08E-05** |
| ENSG00000228414.9 | **REL-DT** | 400957 | lncRNA | **11.213** | 3.063 | 0.922 | **8.355** | 2.33E-06 | **1.08E-05** |
| ENSG00000143061.20 | **IGSF3** | 3321 | protein_coding | **79.513** | 1.173 | 0.353 | **2.255** | 2.34E-06 | **1.08E-05** |
| ENSG00000197020.14 | **ZNF100** | 163227 | protein_coding | **143.200** | -1.005 | -0.303 | **0.498** | 2.35E-06 | **1.09E-05** |
| ENSG00000102531.18 | **FNDC3A** | 22862 | protein_coding | **499.716** | -0.830 | -0.250 | **0.563** | 2.35E-06 | **1.09E-05** |
| ENSG00000125144.16 | **MT1G** | 4495 | protein_coding | **6.452** | 4.235 | 1.275 | **18.826** | 2.39E-06 | **1.10E-05** |

| | | | | | | | | | |
|---|---|---|---|---|---|---|---|---|---|
| ENSG00000007168.17 | **PAFAH1B1** | 5048 | protein_coding | **1903.708** | -0.599 | -0.180 | **0.660** | 2.39E-06 | **1.10E-05** |
| ENSG00000029363.19 | **BCLAF1** | 9774 | protein_coding | **1031.259** | -0.680 | -0.205 | **0.624** | 2.39E-06 | **1.10E-05** |
| ENSG00000132182.15 | **NUP210** | 23225 | protein_coding | **4011.495** | -0.579 | -0.174 | **0.669** | 2.40E-06 | **1.11E-05** |
| ENSG00000160401.16 | **CFAP157** | 286207 | protein_coding | **6.369** | 3.550 | 1.069 | **11.716** | 2.41E-06 | **1.11E-05** |
| ENSG00000205981.11 | **DNAJC19** | 131118 | protein_coding | **461.569** | -0.577 | -0.174 | **0.670** | 2.41E-06 | **1.11E-05** |
| ENSG00000155363.21 | **MOV10** | 4343 | protein_coding | **1787.054** | -0.683 | -0.206 | **0.623** | 2.41E-06 | **1.11E-05** |
| ENSG00000248487.12 | **ABHD14A** | 25864 | protein_coding | **136.886** | -0.864 | -0.260 | **0.550** | 2.41E-06 | **1.11E-05** |
| ENSG00000223764.3 | **LINC02593** | 100130417 | lncRNA | **5.206** | 4.425 | 1.332 | **21.474** | 2.42E-06 | **1.12E-05** |
| ENSG00000140943.19 | **MBTPS1** | 8720 | protein_coding | **1307.148** | 0.462 | 0.139 | **1.377** | 2.42E-06 | **1.12E-05** |
| ENSG00000026103.26 | **FAS** | 355 | protein_coding | **867.544** | -0.962 | -0.290 | **0.513** | 2.43E-06 | **1.12E-05** |
| ENSG00000125812.18 | **GZF1** | 64412 | protein_coding | **231.054** | -0.852 | -0.256 | **0.554** | 2.44E-06 | **1.12E-05** |
| ENSG00000171729.16 | **TMEM51** | 55092 | protein_coding | **124.577** | 1.322 | 0.398 | **2.499** | 2.44E-06 | **1.13E-05** |
| ENSG00000164967.12 | **RPP25L** | 138716 | protein_coding | **178.173** | -0.746 | -0.225 | **0.596** | 2.44E-06 | **1.13E-05** |
| ENSG00000249992.2 | **TMEM158** | 25907 | protein_coding | **5.838** | 3.936 | 1.185 | **15.306** | 2.46E-06 | **1.13E-05** |
| ENSG00000198042.13 | **MAK16** | 84549 | protein_coding | **328.198** | -0.617 | -0.186 | **0.652** | 2.47E-06 | **1.14E-05** |
| ENSG00000170265.14 | **ZNF282** | 8427 | protein_coding | **405.732** | -0.578 | -0.174 | **0.670** | 2.47E-06 | **1.14E-05** |
| ENSG00000172164.16 | **SNTB1** | 6641 | protein_coding | **181.282** | -1.239 | -0.373 | **0.424** | 2.47E-06 | **1.14E-05** |
| ENSG00000072110.19 | **ACTN1** | 87 | protein_coding | **350.559** | 0.829 | 0.250 | **1.777** | 2.48E-06 | **1.14E-05** |
| ENSG00000104808.9 | **DHDH** | 27294 | protein_coding | **9.262** | 2.355 | 0.709 | **5.115** | 2.49E-06 | **1.14E-05** |
| ENSG00000282885.4 | **ENSG00000282885** | | lncRNA | **16.983** | 1.827 | 0.550 | **3.548** | 2.50E-06 | **1.15E-05** |
| ENSG00000145782.15 | **ATG12** | 9140 | protein_coding | **669.258** | -0.452 | -0.136 | **0.731** | 2.50E-06 | **1.15E-05** |
| ENSG00000215014.5 | **SSU72-AS1** | 13400666[illegible] | lncRNA | **27.903** | 1.385 | 0.417 | **2.612** | 2.50E-06 | **1.15E-05** |
| ENSG00000132768.16 | **DPH2** | 1802 | protein_coding | **549.249** | -0.531 | -0.160 | **0.692** | 2.50E-06 | **1.15E-05** |
| ENSG00000139344.10 | **AMDHD1** | 144193 | protein_coding | **24.866** | 1.941 | 0.584 | **3.839** | 2.50E-06 | **1.15E-05** |
| ENSG00000287625.2 | **ENSG00000287625** | | lncRNA | **149.817** | 1.179 | 0.355 | **2.264** | 2.51E-06 | **1.15E-05** |
| ENSG00000145919.13 | **BOD1** | 91272 | protein_coding | **351.375** | -0.546 | -0.164 | **0.685** | 2.51E-06 | **1.15E-05** |
| ENSG00000139211.9 | **AMIGO2** | 347902 | protein_coding | **5.005** | 4.282 | 1.289 | **19.461** | 2.52E-06 | **1.16E-05** |
| ENSG00000288885.3 | **ENSG00000288885** | | lncRNA | **35.082** | 1.848 | 0.556 | **3.599** | 2.53E-06 | **1.16E-05** |
| ENSG00000131370.18 | **SH3BP5** | 9467 | protein_coding | **399.612** | 0.715 | 0.215 | **1.642** | 2.55E-06 | **1.17E-05** |
| ENSG00000105699.19 | **LSR** | 51599 | protein_coding | **610.049** | 0.937 | 0.282 | **1.915** | 2.58E-06 | **1.18E-05** |
| ENSG00000232124.4 | **ENSG0000[illegible]** | 10537283[illegible] | lncRNA | **106.847** | 1.106 | 0.333 | **2.153** | 2.58E-06 | **1.18E-05** |

| | | | | | | | | | |
|---|---|---|---|---|---|---|---|---|---|
| ENSG00000135503.15 | **ACVR1B** | 91 | protein_coding | **70.290** | 1.239 | 0.373 | **2.360** | 2.59E-06 | **1.19E-05** |
| ENSG00000250786.5 | **SNHG18** | | lncRNA | **39.071** | 2.606 | 0.784 | **6.086** | 2.60E-06 | **1.19E-05** |
| ENSG00000228889.10 | **UBAC2-AS:** | 10028937: | lncRNA | **43.072** | 1.184 | 0.356 | **2.272** | 2.60E-06 | **1.19E-05** |
| ENSG00000187193.11 | **MT1X** | 4501 | protein_coding | **148.655** | 1.548 | 0.466 | **2.925** | 2.60E-06 | **1.19E-05** |
| ENSG00000172936.20 | **MYD88** | 4615 | protein_coding | **1202.980** | -0.488 | -0.147 | **0.713** | 2.61E-06 | **1.19E-05** |
| ENSG00000111261.16 | **MANSC1** | 54682 | protein_coding | **10.105** | 3.193 | 0.961 | **9.145** | 2.62E-06 | **1.20E-05** |
| ENSG00000168806.8 | **LCMT2** | 9836 | protein_coding | **159.463** | -0.619 | -0.186 | **0.651** | 2.62E-06 | **1.20E-05** |
| ENSG00000196734.9 | **LCE1B** | 353132 | protein_coding | **9.311** | -2.627 | -0.791 | **0.162** | 2.63E-06 | **1.20E-05** |
| ENSG00000204642.16 | **HLA-F** | 3134 | protein_coding | **7559.037** | -0.428 | -0.129 | **0.743** | 2.63E-06 | **1.20E-05** |
| ENSG00000078124.15 | **ACER3** | 55331 | protein_coding | **136.929** | -0.966 | -0.291 | **0.512** | 2.65E-06 | **1.21E-05** |
| ENSG00000111788.10 | **DDX12B** | | transcribed_unprc | **108.583** | -0.848 | -0.255 | **0.556** | 2.65E-06 | **1.21E-05** |
| ENSG00000145365.14 | **TIFA** | 92610 | protein_coding | **1479.664** | -0.651 | -0.196 | **0.637** | 2.66E-06 | **1.22E-05** |
| ENSG00000100228.16 | **RAB36** | 9609 | protein_coding | **25.923** | 1.943 | 0.585 | **3.845** | 2.66E-06 | **1.22E-05** |
| ENSG00000251533.3 | **LINC00605** | 10013136€ | lncRNA | **8.726** | 3.008 | 0.906 | **8.045** | 2.67E-06 | **1.22E-05** |
| ENSG00000288764.3 | **ENSG00000288764** | | lncRNA | **22.259** | 1.518 | 0.457 | **2.864** | 2.67E-06 | **1.22E-05** |
| ENSG00000116691.13 | **MIIP** | 60672 | protein_coding | **1545.897** | 0.568 | 0.171 | **1.483** | 2.68E-06 | **1.22E-05** |
| ENSG00000165171.13 | **METTL27** | 155368 | protein_coding | **60.699** | 1.415 | 0.426 | **2.666** | 2.70E-06 | **1.23E-05** |
| ENSG00000139618.19 | **BRCA2** | 675 | protein_coding | **388.291** | -0.962 | -0.289 | **0.513** | 2.70E-06 | **1.23E-05** |
| ENSG00000277161.4 | **PIGW** | 284098 | protein_coding | **82.533** | -0.894 | -0.269 | **0.538** | 2.72E-06 | **1.24E-05** |
| ENSG00000213672.10 | **NCKIPSD** | 51517 | protein_coding | **665.629** | -0.521 | -0.157 | **0.697** | 2.74E-06 | **1.25E-05** |
| ENSG00000185022.15 | **MAFF** | 23764 | protein_coding | **159.191** | -1.084 | -0.326 | **0.472** | 2.75E-06 | **1.26E-05** |
| ENSG00000104356.13 | **POP1** | 10940 | protein_coding | **383.995** | -0.646 | -0.194 | **0.639** | 2.77E-06 | **1.26E-05** |
| ENSG00000103932.14 | **RPAP1** | 26015 | protein_coding | **581.639** | -0.503 | -0.151 | **0.706** | 2.79E-06 | **1.27E-05** |
| ENSG00000130182.9 | **ZSCAN10** | 84891 | protein_coding | **6.047** | 4.242 | 1.277 | **18.920** | 2.79E-06 | **1.27E-05** |
| ENSG00000169891.20 | **REPS2** | 9185 | protein_coding | **15.640** | 2.303 | 0.693 | **4.935** | 2.79E-06 | **1.27E-05** |
| ENSG00000159307.22 | **SCUBE1** | 80274 | protein_coding | **7.888** | 3.972 | 1.196 | **15.689** | 2.80E-06 | **1.28E-05** |
| ENSG00000138496.19 | **PARP9** | 83666 | protein_coding | **1271.313** | -0.709 | -0.213 | **0.612** | 2.80E-06 | **1.28E-05** |
| ENSG00000169710.11 | **FASN** | 2194 | protein_coding | **2562.338** | -0.978 | -0.295 | **0.508** | 2.81E-06 | **1.28E-05** |
| ENSG00000116990.13 | **MYCL** | 4610 | protein_coding | **65.561** | 1.284 | 0.387 | **2.435** | 2.81E-06 | **1.28E-05** |
| ENSG00000275185.1 | **ENSG00000275185** | | lncRNA | **29.107** | -1.333 | -0.401 | **0.397** | 2.82E-06 | **1.28E-05** |
| ENSG00000082515.20 | **MRPL22** | 29093 | protein_coding | **884.015** | -0.546 | -0.164 | **0.685** | 2.82E-06 | **1.29E-05** |

| | | | | | | | | | |
|---|---|---|---|---|---|---|---|---|---|
| ENSG00000258429.3 | **PDF** | 64146 | protein_coding | **226.211** | -0.811 | -0.244 | **0.570** | 2.87E-06 | **1.31E-05** |
| ENSG00000106266.13 | **SNX8** | 29886 | protein_coding | **1816.145** | -0.729 | -0.219 | **0.603** | 2.89E-06 | **1.31E-05** |
| ENSG00000231394.1 | **ENSG00000231394** | | lncRNA | **7.000** | 4.249 | 1.279 | **19.009** | 2.89E-06 | **1.32E-05** |
| ENSG00000165240.23 | **ATP7A** | 538 | protein_coding | **214.641** | -0.762 | -0.229 | **0.590** | 2.89E-06 | **1.32E-05** |
| ENSG00000250218.2 | **ALDH1L1-A** | 10087420 | lncRNA | **17.336** | 1.779 | 0.536 | **3.433** | 2.90E-06 | **1.32E-05** |
| ENSG00000175548.10 | **ALG10B** | 144245 | protein_coding | **92.229** | -0.959 | -0.289 | **0.514** | 2.90E-06 | **1.32E-05** |
| ENSG00000104205.18 | **SGK3** | 23678 | protein_coding | **73.881** | 1.032 | 0.311 | **2.045** | 2.90E-06 | **1.32E-05** |
| ENSG00000154589.9 | **LY96** | 23643 | protein_coding | **120.017** | -0.981 | -0.295 | **0.507** | 2.90E-06 | **1.32E-05** |
| ENSG00000151287.19 | **TEX30** | 93081 | protein_coding | **147.948** | -0.680 | -0.205 | **0.624** | 2.91E-06 | **1.32E-05** |
| ENSG00000196388.11 | **INCA1** | 388324 | protein_coding | **33.957** | 1.306 | 0.393 | **2.473** | 2.91E-06 | **1.32E-05** |
| ENSG00000117153.18 | **KLHL12** | 59349 | protein_coding | **642.523** | -0.414 | -0.124 | **0.751** | 2.91E-06 | **1.32E-05** |
| ENSG00000272449.4 | **ENSG00000272449** | | lncRNA | **104.629** | 0.834 | 0.251 | **1.783** | 2.92E-06 | **1.32E-05** |
| ENSG00000124608.7 | **AARS2** | 57505 | protein_coding | **516.971** | -0.445 | -0.134 | **0.734** | 2.93E-06 | **1.33E-05** |
| ENSG00000185963.15 | **BICD2** | 23299 | protein_coding | **496.160** | 0.506 | 0.152 | **1.420** | 2.94E-06 | **1.34E-05** |
| ENSG00000198736.14 | **MSRB1** | 51734 | protein_coding | **629.455** | -0.785 | -0.236 | **0.580** | 2.97E-06 | **1.35E-05** |
| ENSG00000172602.12 | **RND1** | 27289 | protein_coding | **89.685** | 1.168 | 0.352 | **2.247** | 2.99E-06 | **1.35E-05** |
| ENSG00000065183.18 | **WDR3** | 10885 | protein_coding | **414.712** | -0.785 | -0.236 | **0.580** | 3.03E-06 | **1.38E-05** |
| ENSG00000259553.7 | **ENSG00000259553** | | lncRNA | **17.149** | 2.928 | 0.881 | **7.611** | 3.05E-06 | **1.38E-05** |
| ENSG00000119820.13 | **YIPF4** | 84272 | protein_coding | **575.463** | -0.662 | -0.199 | **0.632** | 3.05E-06 | **1.38E-05** |
| ENSG00000128973.15 | **CLN6** | 54982 | protein_coding | **993.555** | -0.616 | -0.185 | **0.652** | 3.06E-06 | **1.38E-05** |
| ENSG00000168389.20 | **MFSD2A** | 84879 | protein_coding | **290.347** | -0.745 | -0.224 | **0.597** | 3.08E-06 | **1.40E-05** |
| ENSG00000286938.2 | **ENSG00000286938** | | lncRNA | **4.734** | 4.592 | 1.382 | **24.111** | 3.09E-06 | **1.40E-05** |
| ENSG00000183077.18 | **AFMID** | 125061 | protein_coding | **616.255** | 0.390 | 0.117 | **1.310** | 3.09E-06 | **1.40E-05** |
| ENSG00000196497.19 | **IPO4** | 79711 | protein_coding | **558.351** | -0.646 | -0.194 | **0.639** | 3.09E-06 | **1.40E-05** |
| ENSG00000262468.9 | **LINC01569** | | lncRNA | **145.682** | 0.885 | 0.266 | **1.846** | 3.10E-06 | **1.40E-05** |
| ENSG00000160404.20 | **TOR2A** | 27433 | protein_coding | **186.223** | -0.698 | -0.210 | **0.616** | 3.10E-06 | **1.40E-05** |
| ENSG00000178764.10 | **ZHX2** | 22882 | protein_coding | **283.263** | -0.713 | -0.215 | **0.610** | 3.15E-06 | **1.42E-05** |
| ENSG00000161921.19 | **CXCL16** | 58191 | protein_coding | **28.888** | 1.796 | 0.541 | **3.474** | 3.15E-06 | **1.42E-05** |
| ENSG00000269397.1 | **ENSG00000269397** | | lncRNA | **19.014** | -1.740 | -0.524 | **0.299** | 3.15E-06 | **1.42E-05** |
| ENSG00000006016.14 | **CRLF1** | 9244 | protein_coding | **7.436** | 5.311 | 1.599 | **39.704** | 3.17E-06 | **1.43E-05** |
| ENSG00000179954.16 | **SSC5D** | 284297 | protein_coding | **17.808** | 1.665 | 0.501 | **3.171** | 3.17E-06 | **1.43E-05** |

| | | | | | | | | | |
|---|---|---|---|---|---|---|---|---|---|
| ENSG00000151065.16 | **DCP1B** | 196513 | protein_coding | **255.574** | -0.569 | -0.171 | **0.674** | 3.17E-06 | **1.43E-05** |
| ENSG00000172058.18 | **SERF1A** | 8293 | protein_coding | **123.592** | -1.127 | -0.339 | **0.458** | 3.18E-06 | **1.44E-05** |
| ENSG00000107077.23 | **KDM4C** | 23081 | protein_coding | **770.823** | 0.673 | 0.203 | **1.594** | 3.19E-06 | **1.44E-05** |
| ENSG00000198858.12 | **R3HDM4** | 91300 | protein_coding | **1956.450** | 0.539 | 0.162 | **1.453** | 3.20E-06 | **1.44E-05** |
| ENSG00000166501.16 | **PRKCB** | 5579 | protein_coding | **1700.014** | 0.856 | 0.258 | **1.810** | 3.20E-06 | **1.44E-05** |
| ENSG00000135441.10 | **BLOC1S1** | 2647 | protein_coding | **901.815** | -0.756 | -0.227 | **0.592** | 3.21E-06 | **1.45E-05** |
| ENSG00000196358.14 | **NTNG2** | 84628 | protein_coding | **56.253** | 2.196 | 0.661 | **4.583** | 3.22E-06 | **1.45E-05** |
| ENSG00000067533.8 | **RRP15** | 51018 | protein_coding | **154.183** | -0.838 | -0.252 | **0.559** | 3.22E-06 | **1.45E-05** |
| ENSG00000147454.15 | **SLC25A37** | 51312 | protein_coding | **869.751** | 0.657 | 0.198 | **1.577** | 3.22E-06 | **1.45E-05** |
| ENSG00000125726.13 | **CD70** | 970 | protein_coding | **4787.180** | -0.506 | -0.152 | **0.704** | 3.23E-06 | **1.45E-05** |
| ENSG00000114857.21 | **NKTR** | 4820 | protein_coding | **3987.454** | 0.449 | 0.135 | **1.365** | 3.24E-06 | **1.46E-05** |
| ENSG00000160191.20 | **PDE9A** | 5152 | protein_coding | **29.807** | 1.444 | 0.435 | **2.722** | 3.26E-06 | **1.47E-05** |
| ENSG00000226251.9 | **LINC02608** | 101929541 | lncRNA | **22.819** | 1.528 | 0.460 | **2.883** | 3.27E-06 | **1.47E-05** |
| ENSG00000183628.16 | **DGCR6** | 8214 | protein_coding | **31.213** | 1.347 | 0.405 | **2.543** | 3.29E-06 | **1.48E-05** |
| ENSG00000142408.7 | **CACNG8** | 59283 | protein_coding | **8.963** | 3.351 | 1.009 | **10.203** | 3.29E-06 | **1.48E-05** |
| ENSG00000182871.18 | **COL18A1** | 80781 | protein_coding | **67.805** | 1.374 | 0.414 | **2.592** | 3.30E-06 | **1.48E-05** |
| ENSG00000205861.14 | **PCOTH** | 542767 | lncRNA | **44.157** | 1.289 | 0.388 | **2.443** | 3.30E-06 | **1.49E-05** |
| ENSG00000285872.4 | **ENSG0000(** | 101929691 | lncRNA | **8.046** | 3.535 | 1.064 | **11.594** | 3.31E-06 | **1.49E-05** |
| ENSG00000198824.10 | **CHAMP1** | 283489 | protein_coding | **407.098** | -0.591 | -0.178 | **0.664** | 3.31E-06 | **1.49E-05** |
| ENSG00000227210.3 | **WDR35-DT** | 10192822 | lncRNA | **6.970** | 3.895 | 1.172 | **14.874** | 3.32E-06 | **1.49E-05** |
| ENSG00000125458.9 | **NT5C** | 30833 | protein_coding | **1199.110** | -0.529 | -0.159 | **0.693** | 3.32E-06 | **1.49E-05** |
| ENSG00000187678.12 | **SPRY4** | 81848 | protein_coding | **108.808** | -0.880 | -0.265 | **0.544** | 3.32E-06 | **1.49E-05** |
| ENSG00000181019.14 | **NQO1** | 1728 | protein_coding | **508.237** | -0.553 | -0.166 | **0.682** | 3.34E-06 | **1.50E-05** |
| ENSG00000261186.2 | **LINC03100** | 12981049 | lncRNA | **9.061** | 2.519 | 0.758 | **5.732** | 3.34E-06 | **1.50E-05** |
| ENSG00000303117.1 | **ENSG00000303117** | | lncRNA | **23.784** | 1.740 | 0.524 | **3.339** | 3.34E-06 | **1.50E-05** |
| ENSG00000127022.18 | **CANX** | 821 | protein_coding | **3681.806** | -0.588 | -0.177 | **0.665** | 3.34E-06 | **1.50E-05** |
| ENSG00000264569.3 | **DCXR-DT** | 11352364( | lncRNA | **8.867** | 2.570 | 0.774 | **5.940** | 3.34E-06 | **1.50E-05** |
| ENSG00000211978.2 | **IGHV5-78** | | IG_V_pseudogene | **54.116** | 1.220 | 0.367 | **2.329** | 3.35E-06 | **1.51E-05** |
| ENSG00000214050.10 | **FBXO16** | 157574 | protein_coding | **127.660** | 1.223 | 0.368 | **2.334** | 3.36E-06 | **1.51E-05** |
| ENSG00000132017.13 | **DCAF15** | 90379 | protein_coding | **1040.963** | -0.668 | -0.201 | **0.629** | 3.36E-06 | **1.51E-05** |
| ENSG00000164086.11 | **DUSP7** | 1849 | protein_coding | **254.765** | -0.595 | -0.179 | **0.662** | 3.37E-06 | **1.51E-05** |

| | | | | | | | | | |
|---|---|---|---|---|---|---|---|---|---|
| ENSG00000169252.6 | **ADRB2** | 154 | protein_coding | **21.229** | 1.716 | 0.517 | **3.286** | 3.38E-06 | **1.51E-05** |
| ENSG00000144228.12 | **SPOPL** | 339745 | protein_coding | **205.337** | -0.650 | -0.196 | **0.637** | 3.39E-06 | **1.52E-05** |
| ENSG00000148408.17 | **CACNA1B** | 774 | protein_coding | **20.274** | 1.993 | 0.600 | **3.979** | 3.39E-06 | **1.52E-05** |
| ENSG00000250644.3 | **ENSG00000250644** | | protein_coding | **23.909** | 2.622 | 0.789 | **6.154** | 3.41E-06 | **1.53E-05** |
| ENSG00000113068.12 | **PFDN1** | 5201 | protein_coding | **915.711** | -0.522 | -0.157 | **0.696** | 3.42E-06 | **1.53E-05** |
| ENSG00000179222.20 | **MAGED1** | 9500 | protein_coding | **1459.600** | -0.434 | -0.131 | **0.740** | 3.42E-06 | **1.53E-05** |
| ENSG00000187862.13 | **TTC24** | 164118 | protein_coding | **28.945** | -1.493 | -0.449 | **0.355** | 3.42E-06 | **1.53E-05** |
| ENSG00000278053.7 | **DDX52** | 11056 | protein_coding | **371.840** | -0.602 | -0.181 | **0.659** | 3.43E-06 | **1.53E-05** |
| ENSG00000165821.13 | **SALL2** | 6297 | protein_coding | **117.534** | 1.055 | 0.318 | **2.078** | 3.43E-06 | **1.54E-05** |
| ENSG00000290832.2 | **ENSG00000290832** | | lncRNA | **746.213** | 0.535 | 0.161 | **1.449** | 3.45E-06 | **1.54E-05** |
| ENSG00000144840.12 | **RABL3** | 285282 | protein_coding | **237.848** | -0.743 | -0.224 | **0.598** | 3.45E-06 | **1.54E-05** |
| ENSG00000178177.19 | **LCORL** | 254251 | protein_coding | **151.792** | -0.991 | -0.298 | **0.503** | 3.46E-06 | **1.55E-05** |
| ENSG00000103942.15 | **HOMER2** | 9455 | protein_coding | **104.944** | 0.900 | 0.271 | **1.866** | 3.46E-06 | **1.55E-05** |
| ENSG00000131269.21 | **ABCB7** | 22 | protein_coding | **406.459** | -0.562 | -0.169 | **0.677** | 3.46E-06 | **1.55E-05** |
| ENSG00000132718.11 | **SYT11** | 23208 | protein_coding | **289.247** | 1.023 | 0.308 | **2.032** | 3.46E-06 | **1.55E-05** |
| ENSG00000160336.19 | **ZNF761** | 388561 | protein_coding | **465.863** | -0.590 | -0.177 | **0.665** | 3.48E-06 | **1.55E-05** |
| ENSG00000172366.22 | **MCRIP2** | 84331 | protein_coding | **996.344** | -0.680 | -0.205 | **0.624** | 3.48E-06 | **1.56E-05** |
| ENSG00000198018.9 | **ENTPD7** | 57089 | protein_coding | **46.670** | -1.030 | -0.310 | **0.490** | 3.49E-06 | **1.56E-05** |
| ENSG00000146707.17 | **POMZP3** | 22932 | protein_coding | **177.268** | -0.659 | -0.199 | **0.633** | 3.52E-06 | **1.57E-05** |
| ENSG00000154781.18 | **CCDC174** | 51244 | protein_coding | **452.614** | -0.696 | -0.210 | **0.617** | 3.53E-06 | **1.58E-05** |
| ENSG00000255693.4 | **LINC02389** | | lncRNA | **33.118** | 1.979 | 0.596 | **3.943** | 3.54E-06 | **1.58E-05** |
| ENSG00000137825.13 | **ITPKA** | 3706 | protein_coding | **62.198** | 1.120 | 0.337 | **2.173** | 3.56E-06 | **1.59E-05** |
| ENSG00000002834.21 | **LASP1** | 3927 | protein_coding | **2726.248** | -0.538 | -0.162 | **0.689** | 3.57E-06 | **1.59E-05** |
| ENSG00000258017.4 | **TUBA1B-AS1** | | lncRNA | **22.062** | -1.736 | -0.523 | **0.300** | 3.57E-06 | **1.59E-05** |
| ENSG00000189007.18 | **ADAT2** | 134637 | protein_coding | **572.609** | -0.448 | -0.135 | **0.733** | 3.58E-06 | **1.59E-05** |
| ENSG00000267041.7 | **ZNF850** | 342892 | protein_coding | **77.999** | -1.408 | -0.424 | **0.377** | 3.58E-06 | **1.59E-05** |
| ENSG00000173230.19 | **GOLGB1** | 2804 | protein_coding | **1839.230** | 0.457 | 0.138 | **1.373** | 3.58E-06 | **1.60E-05** |
| ENSG00000006695.14 | **COX10** | 1352 | protein_coding | **267.988** | -0.641 | -0.193 | **0.641** | 3.58E-06 | **1.60E-05** |
| ENSG00000285625.1 | **ENSG00000285625** | | protein_coding | **41.624** | -2.019 | -0.608 | **0.247** | 3.59E-06 | **1.60E-05** |
| ENSG00000279942.1 | **ENSG00000279942** | | TEC | **234.849** | 0.637 | 0.192 | **1.555** | 3.60E-06 | **1.60E-05** |
| ENSG00000107562.19 | **CXCL12** | 6387 | protein_coding | **14.462** | -2.265 | -0.682 | **0.208** | 3.63E-06 | **1.62E-05** |

| | | | | | | | | | |
|---|---|---|---|---|---|---|---|---|---|
| ENSG00000101464.13 | **PIGU** | 128869 | protein_coding | **357.968** | -0.601 | -0.181 | **0.659** | 3.64E-06 | **1.62E-05** |
| ENSG00000035720.10 | **STAP1** | 26228 | protein_coding | **724.418** | -0.764 | -0.230 | **0.589** | 3.64E-06 | **1.62E-05** |
| ENSG00000101997.15 | **CCDC22** | 28952 | protein_coding | **460.761** | -0.445 | -0.134 | **0.735** | 3.64E-06 | **1.62E-05** |
| ENSG00000038532.19 | **CLEC16A** | 23274 | protein_coding | **1024.766** | 0.498 | 0.150 | **1.412** | 3.64E-06 | **1.62E-05** |
| ENSG00000182934.14 | **SRPRA** | 6734 | protein_coding | **2542.170** | -0.569 | -0.171 | **0.674** | 3.65E-06 | **1.62E-05** |
| ENSG00000126562.19 | **WNK4** | 65266 | protein_coding | **31.837** | 1.672 | 0.503 | **3.186** | 3.65E-06 | **1.62E-05** |
| ENSG00000155254.14 | **MARVELD1** | 83742 | protein_coding | **71.260** | -1.137 | -0.342 | **0.455** | 3.66E-06 | **1.63E-05** |
| ENSG00000253716.9 | **MINCR** | 10050731€ | lncRNA | **137.687** | 0.711 | 0.214 | **1.637** | 3.67E-06 | **1.63E-05** |
| ENSG00000105327.20 | **BBC3** | 27113 | protein_coding | **489.435** | -0.643 | -0.194 | **0.640** | 3.68E-06 | **1.64E-05** |
| ENSG00000178184.17 | **PARD6G** | 84552 | protein_coding | **41.066** | 1.366 | 0.411 | **2.578** | 3.69E-06 | **1.64E-05** |
| ENSG00000140406.4 | **TLNRD1** | 59274 | protein_coding | **394.165** | -0.517 | -0.156 | **0.699** | 3.70E-06 | **1.64E-05** |
| ENSG00000197635.13 | **DPP4** | 1803 | protein_coding | **110.255** | 0.806 | 0.243 | **1.748** | 3.70E-06 | **1.64E-05** |
| ENSG00000005100.14 | **DHX33** | 56919 | protein_coding | **412.664** | -0.525 | -0.158 | **0.695** | 3.71E-06 | **1.64E-05** |
| ENSG00000105639.22 | **JAK3** | 3718 | protein_coding | **1738.303** | -0.715 | -0.215 | **0.609** | 3.71E-06 | **1.65E-05** |
| ENSG00000257027.1 | **ENSG00000257027** | | lncRNA | **69.996** | 1.118 | 0.337 | **2.171** | 3.72E-06 | **1.65E-05** |
| ENSG00000160124.11 | **MIX23** | 131076 | protein_coding | **443.362** | -0.632 | -0.190 | **0.645** | 3.73E-06 | **1.65E-05** |
| ENSG00000169762.19 | **TAPT1** | 202018 | protein_coding | **189.035** | -0.579 | -0.174 | **0.669** | 3.73E-06 | **1.65E-05** |
| ENSG00000133687.18 | **TMTC1** | 83857 | protein_coding | **130.442** | -0.870 | -0.262 | **0.547** | 3.73E-06 | **1.65E-05** |
| ENSG00000112303.15 | **VNN2** | 8875 | protein_coding | **496.697** | 0.660 | 0.199 | **1.580** | 3.74E-06 | **1.66E-05** |
| ENSG00000235052.2 | **ENSG00000235052** | | lncRNA | **12.157** | 2.148 | 0.647 | **4.432** | 3.74E-06 | **1.66E-05** |
| ENSG00000145703.18 | **IQGAP2** | 10788 | protein_coding | **442.020** | -0.691 | -0.208 | **0.619** | 3.75E-06 | **1.66E-05** |
| ENSG00000159199.16 | **ATP5MC1** | 516 | protein_coding | **1615.705** | -0.870 | -0.262 | **0.547** | 3.75E-06 | **1.66E-05** |
| ENSG00000135622.16 | **SEMA4F** | 10505 | protein_coding | **92.907** | 0.850 | 0.256 | **1.802** | 3.76E-06 | **1.66E-05** |
| ENSG00000105656.15 | **ELL** | 8178 | protein_coding | **484.607** | -0.520 | -0.157 | **0.697** | 3.78E-06 | **1.67E-05** |
| ENSG00000173894.12 | **CBX2** | 84733 | protein_coding | **53.078** | 1.482 | 0.446 | **2.794** | 3.79E-06 | **1.68E-05** |
| ENSG00000187240.19 | **DYNC2H1** | 79659 | protein_coding | **7.402** | 3.602 | 1.084 | **12.142** | 3.80E-06 | **1.68E-05** |
| ENSG00000205838.16 | **TTC23L** | 153657 | protein_coding | **6.490** | 3.983 | 1.199 | **15.809** | 3.80E-06 | **1.68E-05** |
| ENSG00000154678.20 | **PDE1C** | 5137 | protein_coding | **41.776** | 1.280 | 0.385 | **2.428** | 3.81E-06 | **1.69E-05** |
| ENSG00000124374.11 | **PAIP2B** | 400961 | protein_coding | **138.875** | 0.783 | 0.236 | **1.720** | 3.81E-06 | **1.69E-05** |
| ENSG00000284946.1 | **ENSG00000284946** | | protein_coding | **187.652** | -1.469 | -0.442 | **0.361** | 3.82E-06 | **1.69E-05** |
| ENSG00000182149.23 | **IST1** | 9798 | protein_coding | **3076.430** | 0.485 | 0.146 | **1.400** | 3.83E-06 | **1.69E-05** |

| | | | | | | | | | |
|---|---|---|---|---|---|---|---|---|---|
| ENSG00000105357.22 | **MYH14** | 79784 | protein_coding | **28.340** | 1.911 | 0.575 | **3.761** | 3.83E-06 | **1.69E-05** |
| ENSG00000304191.1 | **ENSG00000304191** | | lncRNA | **10.672** | 2.693 | 0.811 | **6.465** | 3.84E-06 | **1.70E-05** |
| ENSG00000303202.1 | **ENSG00000303202** | | lncRNA | **9.646** | 2.793 | 0.841 | **6.933** | 3.86E-06 | **1.70E-05** |
| ENSG00000067334.15 | **DNTTIP2** | 30836 | protein_coding | **725.773** | -0.563 | -0.169 | **0.677** | 3.86E-06 | **1.70E-05** |
| ENSG00000272892.2 | **ENSG00000272892** | | lncRNA | **16.529** | 1.771 | 0.533 | **3.414** | 3.87E-06 | **1.71E-05** |
| ENSG00000236859.9 | **NIFK-AS1** | 254128 | lncRNA | **138.500** | 0.640 | 0.193 | **1.559** | 3.87E-06 | **1.71E-05** |
| ENSG00000126705.17 | **AHDC1** | 27245 | protein_coding | **285.388** | 0.725 | 0.218 | **1.653** | 3.88E-06 | **1.71E-05** |
| ENSG00000139132.17 | **FGD4** | 121512 | protein_coding | **10.248** | 2.490 | 0.750 | **5.618** | 3.89E-06 | **1.72E-05** |
| ENSG00000303205.1 | **ENSG00000303205** | | lncRNA | **20.111** | 2.669 | 0.803 | **6.359** | 3.90E-06 | **1.72E-05** |
| ENSG00000230479.2 | **LNCTSI** | | lncRNA | **12.939** | -2.040 | -0.614 | **0.243** | 3.91E-06 | **1.73E-05** |
| ENSG00000100968.15 | **NFATC4** | 4776 | protein_coding | **1447.789** | 0.867 | 0.261 | **1.823** | 3.92E-06 | **1.73E-05** |
| ENSG00000164742.18 | **ADCY1** | 107 | protein_coding | **40.606** | -2.092 | -0.630 | **0.234** | 3.94E-06 | **1.74E-05** |
| ENSG00000143217.11 | **NECTIN4** | 81607 | protein_coding | **11.215** | -2.519 | -0.758 | **0.174** | 3.94E-06 | **1.74E-05** |
| ENSG00000259605.3 | **ENSG00000259605** | | lncRNA | **23.413** | -1.418 | -0.427 | **0.374** | 3.96E-06 | **1.74E-05** |
| ENSG00000245017.5 | **LINC02453** | 643770 | lncRNA | **11.872** | 2.277 | 0.685 | **4.846** | 3.96E-06 | **1.74E-05** |
| ENSG00000135763.12 | **URB2** | 9816 | protein_coding | **176.278** | -0.634 | -0.191 | **0.644** | 3.96E-06 | **1.74E-05** |
| ENSG00000173801.19 | **JUP** | 3728 | protein_coding | **438.529** | 1.193 | 0.359 | **2.287** | 3.96E-06 | **1.75E-05** |
| ENSG00000175324.12 | **LSM1** | 27257 | protein_coding | **454.539** | -0.447 | -0.135 | **0.734** | 3.97E-06 | **1.75E-05** |
| ENSG00000078304.22 | **PPP2R5C** | 5527 | protein_coding | **1984.816** | -0.476 | -0.143 | **0.719** | 3.99E-06 | **1.76E-05** |
| ENSG00000132470.16 | **ITGB4** | 3691 | protein_coding | **40.807** | 1.398 | 0.421 | **2.634** | 3.99E-06 | **1.76E-05** |
| ENSG00000175768.15 | **TOMM5** | 401505 | protein_coding | **1198.906** | -0.627 | -0.189 | **0.647** | 4.00E-06 | **1.76E-05** |
| ENSG00000168152.14 | **THAP9** | 79725 | protein_coding | **90.718** | 0.935 | 0.282 | **1.912** | 4.00E-06 | **1.76E-05** |
| ENSG00000196663.18 | **TECPR2** | 9895 | protein_coding | **319.951** | 0.543 | 0.163 | **1.457** | 4.01E-06 | **1.76E-05** |
| ENSG00000188389.14 | **PDCD1** | 5133 | protein_coding | **269.975** | -0.791 | -0.238 | **0.578** | 4.01E-06 | **1.76E-05** |
| ENSG00000115525.20 | **ST3GAL5** | 8869 | protein_coding | **129.354** | 0.779 | 0.235 | **1.716** | 4.03E-06 | **1.77E-05** |
| ENSG00000053108.20 | **FSTL4** | 23105 | protein_coding | **13.680** | 2.353 | 0.708 | **5.111** | 4.05E-06 | **1.78E-05** |
| ENSG00000213221.7 | **DNLZ** | 728489 | protein_coding | **184.129** | -0.860 | -0.259 | **0.551** | 4.05E-06 | **1.78E-05** |
| ENSG00000163811.14 | **WDR43** | 23160 | protein_coding | **769.571** | -0.802 | -0.241 | **0.574** | 4.06E-06 | **1.78E-05** |
| ENSG00000293205.1 | **TPI1P2** | 286016 | lncRNA | **21.159** | 2.056 | 0.619 | **4.158** | 4.06E-06 | **1.78E-05** |
| ENSG00000089280.22 | **FUS** | 2521 | protein_coding | **6593.524** | -0.592 | -0.178 | **0.663** | 4.08E-06 | **1.79E-05** |
| ENSG00000166439.8 | **RNF169** | 254225 | protein_coding | **244.659** | -0.666 | -0.200 | **0.630** | 4.09E-06 | **1.80E-05** |

| | | | | | | | | | |
|---|---|---|---|---|---|---|---|---|---|
| ENSG00000231607.16 | **DLEU2** | 8847 | lncRNA | **73.334** | -0.974 | -0.293 | **0.509** | 4.09E-06 | **1.80E-05** |
| ENSG00000135272.15 | **MDFIC** | 29969 | protein_coding | **838.578** | -0.796 | -0.239 | **0.576** | 4.12E-06 | **1.81E-05** |
| ENSG00000242258.3 | **LINC00996** | 285972 | lncRNA | **375.931** | -0.685 | -0.206 | **0.622** | 4.13E-06 | **1.81E-05** |
| ENSG00000128789.24 | **PSMG2** | 56984 | protein_coding | **1199.243** | -0.382 | -0.115 | **0.768** | 4.17E-06 | **1.83E-05** |
| ENSG00000116560.13 | **SFPQ** | 6421 | protein_coding | **5527.724** | -0.464 | -0.140 | **0.725** | 4.17E-06 | **1.83E-05** |
| ENSG00000303578.1 | **ENSG00000303578** | | lncRNA | **9.807** | 2.875 | 0.866 | **7.337** | 4.18E-06 | **1.83E-05** |
| ENSG00000196693.17 | **ZNF33B** | 7582 | protein_coding | **269.303** | -0.740 | -0.223 | **0.599** | 4.20E-06 | **1.84E-05** |
| ENSG00000125870.13 | **SNRPB2** | 6629 | protein_coding | **1165.359** | -0.383 | -0.115 | **0.767** | 4.21E-06 | **1.84E-05** |
| ENSG00000100577.21 | **GSTZ1** | 2954 | protein_coding | **587.028** | -0.496 | -0.149 | **0.709** | 4.22E-06 | **1.85E-05** |
| ENSG00000227128.6 | **LBX1-AS1** | | lncRNA | **7.611** | -3.009 | -0.906 | **0.124** | 4.25E-06 | **1.86E-05** |
| ENSG00000128602.13 | **SMO** | 6608 | protein_coding | **261.570** | 0.599 | 0.180 | **1.515** | 4.31E-06 | **1.89E-05** |
| ENSG00000290600.1 | **ENSG00000290600** | | lncRNA | **64.638** | 2.161 | 0.651 | **4.472** | 4.33E-06 | **1.89E-05** |
| ENSG00000110987.11 | **BCL7A** | 605 | protein_coding | **319.715** | -0.677 | -0.204 | **0.626** | 4.34E-06 | **1.90E-05** |
| ENSG00000072858.14 | **SIDT1** | 54847 | protein_coding | **789.164** | -0.510 | -0.153 | **0.702** | 4.34E-06 | **1.90E-05** |
| ENSG00000307088.1 | **ENSG0000(** | 12490546{ | lncRNA | **940.220** | -0.427 | -0.129 | **0.744** | 4.35E-06 | **1.90E-05** |
| ENSG00000138629.18 | **UBL7** | 84993 | protein_coding | **1390.101** | 0.528 | 0.159 | **1.442** | 4.35E-06 | **1.90E-05** |
| ENSG00000232838.5 | **PET117** | 10030375! | protein_coding | **75.105** | -0.936 | -0.282 | **0.523** | 4.35E-06 | **1.90E-05** |
| ENSG00000291117.2 | **ENSG00000291117** | | lncRNA | **111.658** | 0.707 | 0.213 | **1.633** | 4.37E-06 | **1.91E-05** |
| ENSG00000204334.9 | **ERICH2** | 285141 | protein_coding | **13.015** | 2.501 | 0.753 | **5.662** | 4.38E-06 | **1.91E-05** |
| ENSG00000070610.16 | **GBA2** | 57704 | protein_coding | **959.660** | 0.424 | 0.128 | **1.342** | 4.39E-06 | **1.92E-05** |
| ENSG00000117362.15 | **APH1A** | 51107 | protein_coding | **1418.515** | -0.382 | -0.115 | **0.767** | 4.39E-06 | **1.92E-05** |
| ENSG00000004468.15 | **CD38** | 952 | protein_coding | **1282.683** | 1.091 | 0.328 | **2.130** | 4.40E-06 | **1.92E-05** |
| ENSG00000211673.2 | **IGLV3-1** | | IG_V_gene | **2894.879** | -1.224 | -0.369 | **0.428** | 4.40E-06 | **1.92E-05** |
| ENSG00000078369.21 | **GNB1** | 2782 | protein_coding | **4257.273** | -0.319 | -0.096 | **0.802** | 4.41E-06 | **1.93E-05** |
| ENSG00000136450.14 | **SRSF1** | 6426 | protein_coding | **3416.845** | -0.413 | -0.124 | **0.751** | 4.42E-06 | **1.93E-05** |
| ENSG00000153815.19 | **CMIP** | 80790 | protein_coding | **516.733** | 0.558 | 0.168 | **1.472** | 4.46E-06 | **1.95E-05** |
| ENSG00000299307.1 | **ENSG00000299307** | | lncRNA | **14.961** | -2.545 | -0.766 | **0.171** | 4.47E-06 | **1.95E-05** |
| ENSG00000174125.11 | **TLR1** | 7096 | protein_coding | **312.416** | -0.700 | -0.211 | **0.616** | 4.47E-06 | **1.95E-05** |
| ENSG00000134056.14 | **KGD4** | 92259 | protein_coding | **418.179** | -0.658 | -0.198 | **0.634** | 4.47E-06 | **1.95E-05** |
| ENSG00000261693.1 | **ENSG00000261693** | | lncRNA | **7.716** | 2.785 | 0.838 | **6.891** | 4.48E-06 | **1.95E-05** |
| ENSG00000196535.22 | **MYO18A** | 399687 | protein_coding | **2289.114** | -0.527 | -0.159 | **0.694** | 4.49E-06 | **1.96E-05** |

| | | | | | | | | | |
|---|---|---|---|---|---|---|---|---|---|
| ENSG00000275234.1 | **ENSG00000275234** | | lncRNA | **23.530** | 1.580 | 0.476 | **2.991** | 4.49E-06 | **1.96E-05** |
| ENSG00000035664.13 | **DAPK2** | 23604 | protein_coding | **294.247** | 1.139 | 0.343 | **2.202** | 4.50E-06 | **1.96E-05** |
| ENSG00000163659.15 | **TIPARP** | 25976 | protein_coding | **1047.396** | 1.087 | 0.327 | **2.125** | 4.52E-06 | **1.97E-05** |
| ENSG00000245888.8 | **NSMCE1-D** | 400512 | lncRNA | **32.177** | 1.726 | 0.520 | **3.308** | 4.55E-06 | **1.98E-05** |
| ENSG00000265808.6 | **SEC22B** | 9554 | protein_coding | **1021.629** | -0.488 | -0.147 | **0.713** | 4.55E-06 | **1.98E-05** |
| ENSG00000130287.16 | **NCAN** | 1463 | protein_coding | **22.795** | 2.408 | 0.725 | **5.306** | 4.56E-06 | **1.99E-05** |
| ENSG00000217128.15 | **FNIP1** | 96459 | protein_coding | **195.661** | -0.738 | -0.222 | **0.600** | 4.56E-06 | **1.99E-05** |
| ENSG00000116985.13 | **BMP8B** | 656 | protein_coding | **361.278** | 0.542 | 0.163 | **1.455** | 4.56E-06 | **1.99E-05** |
| ENSG00000120533.14 | **ENY2** | 56943 | protein_coding | **1043.378** | -0.537 | -0.162 | **0.689** | 4.57E-06 | **1.99E-05** |
| ENSG00000260565.10 | **ERVK13-1** | | lncRNA | **314.685** | 0.716 | 0.216 | **1.643** | 4.60E-06 | **2.00E-05** |
| ENSG00000142347.23 | **MYO1F** | 4542 | protein_coding | **167.404** | 1.345 | 0.405 | **2.541** | 4.61E-06 | **2.00E-05** |
| ENSG00000115232.15 | **ITGA4** | 3676 | protein_coding | **1393.564** | -0.622 | -0.187 | **0.650** | 4.61E-06 | **2.00E-05** |
| ENSG00000253736.6 | **ENSG0000(** | 10537773( | lncRNA | **13.466** | 2.123 | 0.639 | **4.355** | 4.61E-06 | **2.00E-05** |
| ENSG00000270951.1 | **ENSG00000270951** | | lncRNA | **9.309** | 2.441 | 0.735 | **5.431** | 4.64E-06 | **2.02E-05** |
| ENSG00000174669.14 | **SLC29A2** | 3177 | protein_coding | **577.696** | -0.959 | -0.289 | **0.515** | 4.65E-06 | **2.02E-05** |
| ENSG00000272523.3 | **LINC01023** | | lncRNA | **52.554** | 1.012 | 0.305 | **2.016** | 4.68E-06 | **2.03E-05** |
| ENSG00000136371.13 | **MTHFS** | 10588 | protein_coding | **135.052** | -0.821 | -0.247 | **0.566** | 4.69E-06 | **2.04E-05** |
| ENSG00000133142.21 | **TCEAL4** | 79921 | protein_coding | **780.745** | -0.412 | -0.124 | **0.752** | 4.71E-06 | **2.04E-05** |
| ENSG00000204228.6 | **HSD17B8** | 7923 | protein_coding | **359.227** | 0.854 | 0.257 | **1.808** | 4.71E-06 | **2.04E-05** |
| ENSG00000174989.16 | **FBXW8** | 26259 | protein_coding | **360.841** | 0.494 | 0.149 | **1.409** | 4.71E-06 | **2.05E-05** |
| ENSG00000130529.18 | **TRPM4** | 54795 | protein_coding | **8.074** | 3.483 | 1.048 | **11.181** | 4.73E-06 | **2.05E-05** |
| ENSG00000134072.13 | **CAMK1** | 8536 | protein_coding | **43.378** | -1.582 | -0.476 | **0.334** | 4.74E-06 | **2.06E-05** |
| ENSG00000290718.2 | **FAM95B1** | 10013303( | lncRNA | **42.003** | 2.004 | 0.603 | **4.011** | 4.76E-06 | **2.06E-05** |
| ENSG00000078814.21 | **MYH7B** | 57644 | protein_coding | **51.202** | 1.357 | 0.409 | **2.562** | 4.78E-06 | **2.07E-05** |
| ENSG00000239887.6 | **C1orf226** | 400793 | protein_coding | **4.859** | 4.398 | 1.324 | **21.077** | 4.79E-06 | **2.08E-05** |
| ENSG00000273002.2 | **ARHGEF2-/** | 10798520( | lncRNA | **114.113** | 0.757 | 0.228 | **1.689** | 4.79E-06 | **2.08E-05** |
| ENSG00000144659.15 | **SLC25A38** | 54977 | protein_coding | **795.005** | -0.362 | -0.109 | **0.778** | 4.80E-06 | **2.08E-05** |
| ENSG00000261857.9 | **MIA** | 8190 | protein_coding | **33.471** | 1.630 | 0.491 | **3.096** | 4.81E-06 | **2.08E-05** |
| ENSG00000286408.1 | **ENSG00000286408** | | lncRNA | **48.299** | 1.090 | 0.328 | **2.129** | 4.84E-06 | **2.09E-05** |
| ENSG00000106948.19 | **AKNA** | 80709 | protein_coding | **2497.675** | -0.620 | -0.187 | **0.651** | 4.86E-06 | **2.10E-05** |
| ENSG00000213793.8 | **ZNF888** | 388559 | protein_coding | **96.718** | -0.833 | -0.251 | **0.562** | 4.86E-06 | **2.10E-05** |

| | | | | | | | | | |
|---|---|---|---|---|---|---|---|---|---|
| ENSG00000130347.15 | **RTN4IP1** | 84816 | protein_coding | **172.907** | 0.588 | 0.177 | **1.503** | 4.89E-06 | **2.12E-05** |
| ENSG00000008441.20 | **NFIX** | 4784 | protein_coding | **34.682** | 1.185 | 0.357 | **2.273** | 4.90E-06 | **2.12E-05** |
| ENSG00000172819.19 | **RARG** | 5916 | protein_coding | **281.242** | 0.878 | 0.264 | **1.838** | 4.97E-06 | **2.15E-05** |
| ENSG00000135454.16 | **B4GALNT1** | 2583 | protein_coding | **37.592** | 1.475 | 0.444 | **2.780** | 4.99E-06 | **2.16E-05** |
| ENSG00000176834.16 | **VSIG10** | 54621 | protein_coding | **134.431** | 0.838 | 0.252 | **1.788** | 5.00E-06 | **2.16E-05** |
| ENSG00000173846.16 | **PLK3** | 1263 | protein_coding | **323.475** | -0.647 | -0.195 | **0.639** | 5.02E-06 | **2.17E-05** |
| ENSG00000289213.3 | **ENSG00000289213** | | lncRNA | **27.900** | 1.324 | 0.399 | **2.504** | 5.04E-06 | **2.18E-05** |
| ENSG00000177426.24 | **TGIF1** | 7050 | protein_coding | **140.289** | 0.962 | 0.289 | **1.947** | 5.07E-06 | **2.19E-05** |
| ENSG00000138095.22 | **LRPPRC** | 10128 | protein_coding | **1989.832** | -0.543 | -0.164 | **0.686** | 5.10E-06 | **2.20E-05** |
| ENSG00000092054.16 | **MYH7** | 4625 | protein_coding | **11.887** | 2.584 | 0.778 | **5.995** | 5.12E-06 | **2.21E-05** |
| ENSG00000113790.13 | **EHHADH** | 1962 | protein_coding | **58.772** | 1.151 | 0.347 | **2.221** | 5.13E-06 | **2.21E-05** |
| ENSG00000248367.3 | **ENSG0000(** | 124901151 | lncRNA | **56.474** | 1.112 | 0.335 | **2.162** | 5.13E-06 | **2.21E-05** |
| ENSG00000293487.2 | **CDIPTOSP** | 440356 | lncRNA | **7.054** | 2.892 | 0.870 | **7.421** | 5.14E-06 | **2.22E-05** |
| ENSG00000308538.1 | **ENSG00000308538** | | lncRNA | **13.570** | 2.252 | 0.678 | **4.762** | 5.14E-06 | **2.22E-05** |
| ENSG00000132326.14 | **PER2** | 8864 | protein_coding | **127.111** | -0.787 | -0.237 | **0.580** | 5.17E-06 | **2.23E-05** |
| ENSG00000291015.2 | **ENSG00000291015** | | lncRNA | **108.145** | 0.767 | 0.231 | **1.702** | 5.17E-06 | **2.23E-05** |
| ENSG00000240184.8 | **PCDHGC3** | 5098 | protein_coding | **270.139** | -0.736 | -0.222 | **0.600** | 5.21E-06 | **2.25E-05** |
| ENSG00000115325.16 | **DOK1** | 1796 | protein_coding | **568.251** | -0.496 | -0.149 | **0.709** | 5.22E-06 | **2.25E-05** |
| ENSG00000214013.11 | **GANC** | 2595 | protein_coding | **348.692** | -0.637 | -0.192 | **0.643** | 5.23E-06 | **2.25E-05** |
| ENSG00000121039.12 | **RDH10** | 157506 | protein_coding | **52.982** | 1.347 | 0.405 | **2.544** | 5.26E-06 | **2.27E-05** |
| ENSG00000287737.4 | **ENSG00000287737** | | lncRNA | **22.954** | 1.474 | 0.444 | **2.777** | 5.26E-06 | **2.27E-05** |
| ENSG00000134602.18 | **STK26** | 51765 | protein_coding | **364.939** | -0.783 | -0.236 | **0.581** | 5.26E-06 | **2.27E-05** |
| ENSG00000185133.16 | **INPP5J** | 27124 | protein_coding | **12.335** | 2.636 | 0.794 | **6.218** | 5.26E-06 | **2.27E-05** |
| ENSG00000185808.16 | **PIGP** | 51227 | protein_coding | **221.960** | -0.578 | -0.174 | **0.670** | 5.29E-06 | **2.28E-05** |
| ENSG00000129103.21 | **SUMF2** | 25870 | protein_coding | **997.634** | -0.451 | -0.136 | **0.731** | 5.31E-06 | **2.29E-05** |
| ENSG00000167703.17 | **SLC43A2** | 124935 | protein_coding | **400.257** | -0.634 | -0.191 | **0.644** | 5.31E-06 | **2.29E-05** |
| ENSG00000302621.1 | **ENSG00000302621** | | lncRNA | **61.339** | 0.984 | 0.296 | **1.977** | 5.33E-06 | **2.29E-05** |
| ENSG00000166167.20 | **BTRC** | 8945 | protein_coding | **261.397** | -0.494 | -0.149 | **0.710** | 5.33E-06 | **2.29E-05** |
| ENSG00000100360.17 | **IFT27** | 11020 | protein_coding | **498.209** | 0.618 | 0.186 | **1.535** | 5.36E-06 | **2.30E-05** |
| ENSG00000089163.6 | **SIRT4** | 23409 | protein_coding | **34.023** | 1.298 | 0.391 | **2.459** | 5.36E-06 | **2.30E-05** |
| ENSG00000100105.20 | **PATZ1** | 23598 | protein_coding | **447.766** | -0.593 | -0.179 | **0.663** | 5.36E-06 | **2.30E-05** |

| | | | | | | | | | |
|---|---|---|---|---|---|---|---|---|---|
| ENSG00000167562.16 | **ZNF701** | 55762 | protein_coding | **202.698** | -0.636 | -0.191 | **0.644** | 5.37E-06 | **2.31E-05** |
| ENSG00000116649.12 | **SRM** | 6723 | protein_coding | **1797.311** | -0.929 | -0.280 | **0.525** | 5.39E-06 | **2.31E-05** |
| ENSG00000124216.5 | **SNAI1** | 6615 | protein_coding | **24.715** | 1.884 | 0.567 | **3.690** | 5.39E-06 | **2.32E-05** |
| ENSG00000125388.22 | **GRK4** | 2868 | protein_coding | **55.055** | 1.155 | 0.348 | **2.227** | 5.42E-06 | **2.33E-05** |
| ENSG00000106392.13 | **C1GALT1** | 56913 | protein_coding | **299.932** | -0.760 | -0.229 | **0.590** | 5.45E-06 | **2.34E-05** |
| ENSG00000143742.16 | **SRP9** | 6726 | protein_coding | **1184.916** | -0.645 | -0.194 | **0.640** | 5.47E-06 | **2.35E-05** |
| ENSG00000162511.10 | **LAPTM5** | 7805 | protein_coding | **25001.774** | -0.645 | -0.194 | **0.639** | 5.49E-06 | **2.35E-05** |
| ENSG00000171130.21 | **ATP6V0E2** | 155066 | protein_coding | **386.791** | 0.517 | 0.156 | **1.431** | 5.51E-06 | **2.36E-05** |
| ENSG00000258537.7 | **FRMD6-AS** | 10087418! | lncRNA | **28.940** | 1.366 | 0.411 | **2.577** | 5.53E-06 | **2.37E-05** |
| ENSG00000198467.18 | **TPM2** | 7169 | protein_coding | **497.654** | 0.973 | 0.293 | **1.963** | 5.53E-06 | **2.37E-05** |
| ENSG00000186564.6 | **FOXD2** | 2306 | protein_coding | **7.042** | 2.821 | 0.849 | **7.065** | 5.53E-06 | **2.37E-05** |
| ENSG00000272772.2 | **ENSG00000272772** | | protein_coding | **310.574** | 0.772 | 0.232 | **1.708** | 5.61E-06 | **2.40E-05** |
| ENSG00000055609.22 | **KMT2C** | 58508 | protein_coding | **1081.245** | 0.559 | 0.168 | **1.474** | 5.62E-06 | **2.41E-05** |
| ENSG00000125089.20 | **SH3TC1** | 54436 | protein_coding | **1230.874** | -0.761 | -0.229 | **0.590** | 5.62E-06 | **2.41E-05** |
| ENSG00000168374.13 | **ARF4** | 378 | protein_coding | **1167.950** | -0.480 | -0.145 | **0.717** | 5.63E-06 | **2.41E-05** |
| ENSG00000308602.1 | **OVOS1B** | | lncRNA | **126.011** | -1.107 | -0.333 | **0.464** | 5.65E-06 | **2.42E-05** |
| ENSG00000287562.1 | **ENSG00000287562** | | lncRNA | **66.049** | 1.040 | 0.313 | **2.056** | 5.66E-06 | **2.42E-05** |
| ENSG00000119421.9 | **NDUFA8** | 4702 | protein_coding | **705.736** | -0.554 | -0.167 | **0.681** | 5.69E-06 | **2.43E-05** |
| ENSG00000135749.22 | **PCNX2** | 80003 | protein_coding | **218.585** | 1.079 | 0.325 | **2.113** | 5.69E-06 | **2.44E-05** |
| ENSG00000184224.4 | **NDUFV1-D** | 100505621 | lncRNA | **8.265** | 2.807 | 0.845 | **6.996** | 5.69E-06 | **2.44E-05** |
| ENSG00000155189.14 | **AGPAT5** | 55326 | protein_coding | **1550.745** | 0.590 | 0.178 | **1.505** | 5.71E-06 | **2.44E-05** |
| ENSG00000122481.19 | **RWDD3** | 25950 | protein_coding | **149.500** | -0.725 | -0.218 | **0.605** | 5.71E-06 | **2.44E-05** |
| ENSG00000197632.11 | **SERPINB2** | 5055 | protein_coding | **6.669** | 3.506 | 1.055 | **11.361** | 5.75E-06 | **2.46E-05** |
| ENSG00000134369.19 | **NAV1** | 89796 | protein_coding | **64.925** | 1.185 | 0.357 | **2.274** | 5.75E-06 | **2.46E-05** |
| ENSG00000196263.10 | **ZNF471** | 57573 | protein_coding | **23.911** | 1.984 | 0.597 | **3.955** | 5.78E-06 | **2.47E-05** |
| ENSG00000032389.15 | **EIPR1** | 7260 | protein_coding | **647.437** | -0.461 | -0.139 | **0.726** | 5.79E-06 | **2.47E-05** |
| ENSG00000128563.15 | **PRKRIP1** | 79706 | protein_coding | **2051.531** | 0.532 | 0.160 | **1.446** | 5.82E-06 | **2.49E-05** |
| ENSG00000163510.16 | **CWC22** | 57703 | protein_coding | **400.459** | -0.502 | -0.151 | **0.706** | 5.85E-06 | **2.50E-05** |
| ENSG00000084072.19 | **PPIE** | 10450 | protein_coding | **1088.374** | -0.413 | -0.124 | **0.751** | 5.86E-06 | **2.50E-05** |
| ENSG00000108064.13 | **TFAM** | 7019 | protein_coding | **529.043** | -0.862 | -0.259 | **0.550** | 5.86E-06 | **2.50E-05** |
| ENSG00000255330.9 | **ENSG00000255330** | | protein_coding | **81.718** | 1.251 | 0.377 | **2.380** | 5.88E-06 | **2.51E-05** |

| | | | | | | | | | |
|---|---|---|---|---|---|---|---|---|---|
| ENSG00000258602.3 | **LINC01629** | 10537057ε | lncRNA | **53.618** | -1.305 | -0.393 | **0.405** | 5.89E-06 | **2.51E-05** |
| ENSG00000289151.3 | **ENSG00000289151** | | lncRNA | **15.603** | 1.720 | 0.518 | **3.295** | 5.90E-06 | **2.52E-05** |
| ENSG00000295502.1 | **ENSG00000295502** | | lncRNA | **35.203** | 2.275 | 0.685 | **4.840** | 5.92E-06 | **2.52E-05** |
| ENSG00000065526.14 | **SPEN** | 23013 | protein_coding | **1546.000** | 0.453 | 0.136 | **1.369** | 5.94E-06 | **2.53E-05** |
| ENSG00000154118.14 | **JPH3** | 57338 | protein_coding | **6.293** | 3.839 | 1.156 | **14.307** | 5.94E-06 | **2.53E-05** |
| ENSG00000102032.15 | **RENBP** | 5973 | protein_coding | **137.537** | -0.835 | -0.251 | **0.561** | 5.95E-06 | **2.54E-05** |
| ENSG00000063169.13 | **BICRA** | 29998 | protein_coding | **491.559** | 0.612 | 0.184 | **1.529** | 5.96E-06 | **2.54E-05** |
| ENSG00000183647.13 | **ZNF530** | 348327 | protein_coding | **87.718** | -1.330 | -0.400 | **0.398** | 5.96E-06 | **2.54E-05** |
| ENSG00000141574.10 | **SECTM1** | 6398 | protein_coding | **10.941** | 3.563 | 1.073 | **11.819** | 5.98E-06 | **2.55E-05** |
| ENSG00000160410.17 | **SHKBP1** | 92799 | protein_coding | **1996.463** | 0.620 | 0.187 | **1.537** | 5.98E-06 | **2.55E-05** |
| ENSG00000271743.1 | **ENSG00000271743** | | lncRNA | **19.563** | 1.807 | 0.544 | **3.500** | 5.99E-06 | **2.55E-05** |
| ENSG00000147123.14 | **NDUFB11** | 54539 | protein_coding | **1454.218** | -0.504 | -0.152 | **0.705** | 6.01E-06 | **2.56E-05** |
| ENSG00000197558.14 | **SSPOP** | | transcribed_unita | **8.121** | 3.514 | 1.058 | **11.427** | 6.04E-06 | **2.57E-05** |
| ENSG00000029364.14 | **SLC39A9** | 55334 | protein_coding | **1125.423** | -0.509 | -0.153 | **0.703** | 6.08E-06 | **2.58E-05** |
| ENSG00000286705.3 | **ENSG00000286705** | | lncRNA | **7.675** | 2.725 | 0.820 | **6.611** | 6.08E-06 | **2.59E-05** |
| ENSG00000010292.15 | **NCAPD2** | 9918 | protein_coding | **2344.069** | -0.468 | -0.141 | **0.723** | 6.12E-06 | **2.60E-05** |
| ENSG00000171612.9 | **SLC25A33** | 84275 | protein_coding | **216.376** | -0.554 | -0.167 | **0.681** | 6.15E-06 | **2.61E-05** |
| ENSG00000179021.12 | **C3orf38** | 285237 | protein_coding | **254.761** | -0.659 | -0.198 | **0.633** | 6.17E-06 | **2.62E-05** |
| ENSG00000131019.13 | **ULBP3** | 79465 | protein_coding | **10.034** | 3.011 | 0.906 | **8.059** | 6.19E-06 | **2.63E-05** |
| ENSG00000265458.4 | **NARF-AS2** | 105371941 | lncRNA | **88.042** | 1.055 | 0.318 | **2.078** | 6.24E-06 | **2.65E-05** |
| ENSG00000168066.24 | **SF1** | 7536 | protein_coding | **3829.571** | 0.434 | 0.131 | **1.351** | 6.26E-06 | **2.66E-05** |
| ENSG00000134265.15 | **NAPG** | 8774 | protein_coding | **437.860** | -0.700 | -0.211 | **0.615** | 6.30E-06 | **2.67E-05** |
| ENSG00000126804.16 | **ZBTB1** | 22890 | protein_coding | **796.034** | -0.725 | -0.218 | **0.605** | 6.30E-06 | **2.68E-05** |
| ENSG00000080573.8 | **COL5A3** | 50509 | protein_coding | **14.785** | 1.804 | 0.543 | **3.492** | 6.31E-06 | **2.68E-05** |
| ENSG00000141655.20 | **TNFRSF11/** | 8792 | protein_coding | **135.978** | 1.067 | 0.321 | **2.096** | 6.34E-06 | **2.69E-05** |
| ENSG00000197555.12 | **SIPA1L1** | 26037 | protein_coding | **2102.344** | -0.461 | -0.139 | **0.726** | 6.35E-06 | **2.69E-05** |
| ENSG00000256574.10 | **OR13A1** | 79290 | protein_coding | **30.289** | 1.820 | 0.548 | **3.530** | 6.37E-06 | **2.70E-05** |
| ENSG00000290569.2 | **ENSG00000290569** | | lncRNA | **91.771** | -0.905 | -0.272 | **0.534** | 6.38E-06 | **2.70E-05** |
| ENSG00000226004.3 | **SIMALR** | | lncRNA | **11.369** | 2.764 | 0.832 | **6.792** | 6.39E-06 | **2.71E-05** |
| ENSG00000172296.15 | **SPTLC3** | 55304 | protein_coding | **20.139** | 2.602 | 0.783 | **6.070** | 6.40E-06 | **2.71E-05** |
| ENSG00000104332.14 | **SFRP1** | 6422 | protein_coding | **14.884** | 2.475 | 0.745 | **5.559** | 6.40E-06 | **2.71E-05** |

| | | | | | | | | | |
|---|---|---|---|---|---|---|---|---|---|
| ENSG00000253666.5 | **ENSG00000253666** | | lncRNA | **432.515** | 0.490 | 0.148 | **1.405** | 6.40E-06 | **2.71E-05** |
| ENSG00000065427.17 | **KARS1** | 3735 | protein_coding | **3431.841** | -0.447 | -0.135 | **0.733** | 6.41E-06 | **2.71E-05** |
| ENSG00000100023.21 | **PPIL2** | 23759 | protein_coding | **1641.322** | 0.341 | 0.103 | **1.267** | 6.41E-06 | **2.71E-05** |
| ENSG00000135912.13 | **TTLL4** | 9654 | protein_coding | **477.560** | -0.572 | -0.172 | **0.673** | 6.44E-06 | **2.73E-05** |
| ENSG00000297244.1 | **ENSG00000297244** | | lncRNA | **204.072** | -0.894 | -0.269 | **0.538** | 6.45E-06 | **2.73E-05** |
| ENSG00000115282.23 | **TTC31** | 64427 | protein_coding | **946.499** | 0.391 | 0.118 | **1.312** | 6.46E-06 | **2.73E-05** |
| ENSG00000007541.19 | **PIGQ** | 9091 | protein_coding | **560.396** | 0.463 | 0.139 | **1.379** | 6.48E-06 | **2.74E-05** |
| ENSG00000007944.17 | **MYLIP** | 29116 | protein_coding | **696.097** | 0.688 | 0.207 | **1.612** | 6.48E-06 | **2.74E-05** |
| ENSG00000111412.7 | **SPRING1** | 79794 | protein_coding | **265.920** | -0.957 | -0.288 | **0.515** | 6.51E-06 | **2.75E-05** |
| ENSG00000106853.22 | **PTGR1** | 22949 | protein_coding | **119.867** | 0.930 | 0.280 | **1.906** | 6.54E-06 | **2.76E-05** |
| ENSG00000015592.19 | **STMN4** | 81551 | protein_coding | **8.902** | 2.550 | 0.768 | **5.856** | 6.54E-06 | **2.77E-05** |
| ENSG00000074706.16 | **IPCEF1** | 26034 | protein_coding | **751.927** | 0.876 | 0.264 | **1.835** | 6.55E-06 | **2.77E-05** |
| ENSG00000147647.15 | **DPYS** | 1807 | protein_coding | **12.369** | 2.958 | 0.891 | **7.772** | 6.56E-06 | **2.77E-05** |
| ENSG00000121671.14 | **CRY2** | 1408 | protein_coding | **383.248** | 0.772 | 0.232 | **1.707** | 6.56E-06 | **2.77E-05** |
| ENSG00000174842.19 | **GLMN** | 11146 | protein_coding | **112.423** | -0.683 | -0.206 | **0.623** | 6.58E-06 | **2.78E-05** |
| ENSG00000241163.11 | **LINC00877** | 285286 | lncRNA | **186.405** | 0.746 | 0.225 | **1.678** | 6.59E-06 | **2.78E-05** |
| ENSG00000155366.19 | **RHOC** | 389 | protein_coding | **2924.493** | 0.516 | 0.155 | **1.430** | 6.60E-06 | **2.78E-05** |
| ENSG00000198691.15 | **ABCA4** | 24 | protein_coding | **11.563** | 3.353 | 1.009 | **10.217** | 6.62E-06 | **2.80E-05** |
| ENSG00000133398.6 | **MED10** | 84246 | protein_coding | **644.595** | -0.443 | -0.133 | **0.736** | 6.63E-06 | **2.80E-05** |
| ENSG00000112290.15 | **WASF1** | 8936 | protein_coding | **75.707** | 1.417 | 0.427 | **2.670** | 6.65E-06 | **2.81E-05** |
| ENSG00000034713.9 | **GABARAPL** | 11345 | protein_coding | **1613.420** | 0.539 | 0.162 | **1.453** | 6.65E-06 | **2.81E-05** |
| ENSG00000111321.13 | **LTBR** | 4055 | protein_coding | **175.530** | -1.182 | -0.356 | **0.441** | 6.68E-06 | **2.82E-05** |
| ENSG00000171533.14 | **MAP6** | 4135 | protein_coding | **15.206** | 2.626 | 0.791 | **6.174** | 6.71E-06 | **2.83E-05** |
| ENSG00000173567.17 | **ADGRF3** | 165082 | protein_coding | **31.031** | -2.092 | -0.630 | **0.234** | 6.73E-06 | **2.84E-05** |
| ENSG00000273080.3 | **ENSG00000273080** | | lncRNA | **33.406** | 1.152 | 0.347 | **2.222** | 6.75E-06 | **2.85E-05** |
| ENSG00000101439.11 | **CST3** | 1471 | protein_coding | **1112.673** | 0.712 | 0.214 | **1.638** | 6.77E-06 | **2.85E-05** |
| ENSG00000125482.15 | **TTF1** | 7270 | protein_coding | **315.780** | -0.579 | -0.174 | **0.670** | 6.78E-06 | **2.86E-05** |
| ENSG00000136273.16 | **HUS1** | 3364 | protein_coding | **148.632** | -0.725 | -0.218 | **0.605** | 6.78E-06 | **2.86E-05** |
| ENSG00000153048.13 | **CARHSP1** | 23589 | protein_coding | **1129.453** | -0.560 | -0.169 | **0.678** | 6.78E-06 | **2.86E-05** |
| ENSG00000163281.14 | **GNPDA2** | 132789 | protein_coding | **135.740** | -0.658 | -0.198 | **0.634** | 6.79E-06 | **2.86E-05** |
| ENSG00000170921.17 | **TANC2** | 26115 | protein_coding | **34.101** | 1.230 | 0.370 | **2.346** | 6.80E-06 | **2.86E-05** |

| | | | | | | | | | |
|---|---|---|---|---|---|---|---|---|---|
| ENSG00000160959.10 | **LRRC14** | 9684 | protein_coding | **644.854** | -0.571 | -0.172 | **0.673** | 6.82E-06 | **2.87E-05** |
| ENSG00000228288.11 | **PCAT6** | 10050669€ | lncRNA | **84.634** | 1.035 | 0.312 | **2.049** | 6.82E-06 | **2.87E-05** |
| ENSG00000113141.20 | **IK** | 3550 | protein_coding | **2105.626** | 0.490 | 0.147 | **1.404** | 6.84E-06 | **2.88E-05** |
| ENSG00000278916.2 | **CEP83-DT** | 144486 | lncRNA | **30.318** | 1.294 | 0.390 | **2.453** | 6.88E-06 | **2.89E-05** |
| ENSG00000162836.14 | **ACP6** | 51205 | protein_coding | **317.977** | 0.555 | 0.167 | **1.469** | 6.88E-06 | **2.89E-05** |
| ENSG00000277734.8 | **TRAC** | | TR_C_gene | **1447.405** | -0.799 | -0.240 | **0.575** | 6.90E-06 | **2.90E-05** |
| ENSG00000152778.10 | **IFIT5** | 24138 | protein_coding | **406.613** | -0.725 | -0.218 | **0.605** | 6.90E-06 | **2.90E-05** |
| ENSG00000271614.2 | **ATP2B1-AS1** | | lncRNA | **81.442** | 0.916 | 0.276 | **1.886** | 6.96E-06 | **2.93E-05** |
| ENSG00000136274.10 | **NACAD** | 23148 | protein_coding | **25.300** | 1.986 | 0.598 | **3.961** | 6.96E-06 | **2.93E-05** |
| ENSG00000117640.20 | **MTFR1L** | 56181 | protein_coding | **1130.779** | 0.439 | 0.132 | **1.356** | 6.98E-06 | **2.93E-05** |
| ENSG00000301690.1 | **ENSG00000301690** | | lncRNA | **6.063** | 3.777 | 1.137 | **13.705** | 6.99E-06 | **2.93E-05** |
| ENSG00000173960.16 | **UBXN2A** | 165324 | protein_coding | **241.056** | -0.773 | -0.233 | **0.585** | 7.00E-06 | **2.94E-05** |
| ENSG00000109911.21 | **ELP4** | 26610 | protein_coding | **232.719** | -0.750 | -0.226 | **0.595** | 7.00E-06 | **2.94E-05** |
| ENSG00000186994.14 | **KANK3** | 256949 | protein_coding | **40.796** | 1.827 | 0.550 | **3.547** | 7.05E-06 | **2.96E-05** |
| ENSG00000297976.1 | **ENSG0000(** | 12490521! | lncRNA | **32.015** | 1.261 | 0.380 | **2.396** | 7.06E-06 | **2.96E-05** |
| ENSG00000143621.19 | **ILF2** | 3608 | protein_coding | **2097.039** | -0.528 | -0.159 | **0.694** | 7.07E-06 | **2.96E-05** |
| ENSG00000100567.15 | **PSMA3** | 5684 | protein_coding | **1767.312** | -0.523 | -0.158 | **0.696** | 7.07E-06 | **2.96E-05** |
| ENSG00000305511.1 | **ENSG00000305511** | | lncRNA | **141.439** | -0.681 | -0.205 | **0.624** | 7.08E-06 | **2.97E-05** |
| ENSG00000279453.1 | **ENSG00000279453** | | TEC | **31.166** | 1.164 | 0.350 | **2.240** | 7.08E-06 | **2.97E-05** |
| ENSG00000103160.14 | **HSDL1** | 83693 | protein_coding | **221.275** | 0.771 | 0.232 | **1.707** | 7.09E-06 | **2.97E-05** |
| ENSG00000189079.20 | **ARID2** | 196528 | protein_coding | **225.436** | -0.617 | -0.186 | **0.652** | 7.09E-06 | **2.97E-05** |
| ENSG00000183723.14 | **CMTM4** | 146223 | protein_coding | **13.145** | 2.532 | 0.762 | **5.786** | 7.09E-06 | **2.97E-05** |
| ENSG00000174137.15 | **FAM53A** | 152877 | protein_coding | **33.755** | 1.361 | 0.410 | **2.569** | 7.09E-06 | **2.97E-05** |
| ENSG00000228839.6 | **PIK3IP1-DT** | | lncRNA | **15.491** | 1.764 | 0.531 | **3.395** | 7.14E-06 | **2.99E-05** |
| ENSG00000104903.7 | **LYL1** | 4066 | protein_coding | **288.655** | -0.802 | -0.241 | **0.574** | 7.15E-06 | **2.99E-05** |
| ENSG00000165782.13 | **PIP4P1** | 90809 | protein_coding | **779.868** | 0.427 | 0.129 | **1.344** | 7.16E-06 | **3.00E-05** |
| ENSG00000240857.4 | **RDH14** | 57665 | protein_coding | **219.386** | -0.617 | -0.186 | **0.652** | 7.16E-06 | **3.00E-05** |
| ENSG00000240505.9 | **TNFRSF13E** | 23495 | protein_coding | **3326.484** | -0.638 | -0.192 | **0.643** | 7.17E-06 | **3.00E-05** |
| ENSG00000164754.18 | **RAD21** | 5885 | protein_coding | **1722.617** | -0.563 | -0.169 | **0.677** | 7.19E-06 | **3.01E-05** |
| ENSG00000165637.16 | **VDAC2** | 7417 | protein_coding | **2336.956** | -0.438 | -0.132 | **0.738** | 7.24E-06 | **3.03E-05** |
| ENSG00000289622.3 | **ENSG00000289622** | | lncRNA | **70.902** | 1.182 | 0.356 | **2.269** | 7.25E-06 | **3.03E-05** |

| | | | | | | | | | |
|---|---|---|---|---|---|---|---|---|---|
| ENSG00000115540.17 | **MOB4** | 25843 | protein_coding | **235.617** | -0.508 | -0.153 | **0.703** | 7.26E-06 | **3.03E-05** |
| ENSG00000273320.3 | **NAMPT-AS** | 128266843 | lncRNA | **33.681** | 1.138 | 0.343 | **2.201** | 7.27E-06 | **3.04E-05** |
| ENSG00000198356.14 | **GET3** | 439 | protein_coding | **749.312** | -0.582 | -0.175 | **0.668** | 7.31E-06 | **3.05E-05** |
| ENSG00000058729.13 | **RIOK2** | 55781 | protein_coding | **304.895** | -0.767 | -0.231 | **0.588** | 7.33E-06 | **3.06E-05** |
| ENSG00000198380.15 | **GFPT1** | 2673 | protein_coding | **832.381** | -0.549 | -0.165 | **0.684** | 7.33E-06 | **3.06E-05** |
| ENSG00000205795.6 | **CYS1** | 192668 | protein_coding | **151.080** | 0.770 | 0.232 | **1.706** | 7.35E-06 | **3.07E-05** |
| ENSG00000275052.7 | **PPP4R3B** | 57223 | protein_coding | **700.977** | -0.565 | -0.170 | **0.676** | 7.38E-06 | **3.08E-05** |
| ENSG00000156502.16 | **SUPV3L1** | 6832 | protein_coding | **516.873** | -0.645 | -0.194 | **0.640** | 7.39E-06 | **3.08E-05** |
| ENSG00000142657.23 | **PGD** | 5226 | protein_coding | **1836.314** | -0.615 | -0.185 | **0.653** | 7.43E-06 | **3.10E-05** |
| ENSG00000138867.19 | **GUCD1** | 83606 | protein_coding | **1453.895** | -0.624 | -0.188 | **0.649** | 7.44E-06 | **3.10E-05** |
| ENSG00000188822.9 | **CNR2** | 1269 | protein_coding | **751.017** | -0.812 | -0.245 | **0.569** | 7.48E-06 | **3.12E-05** |
| ENSG00000204947.11 | **ZNF425** | 155054 | protein_coding | **30.777** | 1.252 | 0.377 | **2.382** | 7.53E-06 | **3.14E-05** |
| ENSG00000060138.15 | **YBX3** | 8531 | protein_coding | **2683.380** | -0.423 | -0.127 | **0.746** | 7.56E-06 | **3.15E-05** |
| ENSG00000177000.15 | **MTHFR** | 4524 | protein_coding | **655.146** | 0.578 | 0.174 | **1.493** | 7.57E-06 | **3.16E-05** |
| ENSG00000197724.14 | **PHF2** | 5253 | protein_coding | **365.461** | -0.545 | -0.164 | **0.685** | 7.58E-06 | **3.16E-05** |
| ENSG00000189077.13 | **TMEM120/** | 83862 | protein_coding | **843.707** | 0.868 | 0.261 | **1.825** | 7.58E-06 | **3.16E-05** |
| ENSG00000167972.16 | **ABCA3** | 21 | protein_coding | **236.043** | 1.118 | 0.336 | **2.170** | 7.61E-06 | **3.17E-05** |
| ENSG00000117013.19 | **KCNQ4** | 9132 | protein_coding | **39.716** | 1.131 | 0.340 | **2.190** | 7.70E-06 | **3.20E-05** |
| ENSG00000006744.21 | **ELAC2** | 60528 | protein_coding | **1450.917** | -0.360 | -0.108 | **0.779** | 7.71E-06 | **3.21E-05** |
| ENSG00000267382.2 | **GAD3P** | | transcribed_unprc | **5.937** | 4.505 | 1.356 | **22.702** | 7.73E-06 | **3.22E-05** |
| ENSG00000170852.14 | **KBTBD2** | 25948 | protein_coding | **372.773** | -0.537 | -0.162 | **0.689** | 7.73E-06 | **3.22E-05** |
| ENSG00000164010.17 | **ERMAP** | 114625 | protein_coding | **121.119** | -0.731 | -0.220 | **0.602** | 7.74E-06 | **3.22E-05** |
| ENSG00000144355.16 | **DLX1** | 1745 | protein_coding | **9.807** | 2.917 | 0.878 | **7.552** | 7.75E-06 | **3.22E-05** |
| ENSG00000221988.15 | **PPT2** | 9374 | protein_coding | **103.538** | 0.720 | 0.217 | **1.648** | 7.78E-06 | **3.23E-05** |
| ENSG00000152234.18 | **ATP5F1A** | 498 | protein_coding | **5576.428** | -0.442 | -0.133 | **0.736** | 7.79E-06 | **3.24E-05** |
| ENSG00000235527.8 | **HIPK1-AS1** | 10192884€ | lncRNA | **14.928** | 1.808 | 0.544 | **3.501** | 7.80E-06 | **3.24E-05** |
| ENSG00000246640.3 | **PICART1** | 284080 | lncRNA | **5.922** | 3.948 | 1.189 | **15.435** | 7.82E-06 | **3.25E-05** |
| ENSG00000156535.16 | **CD109** | 135228 | protein_coding | **125.489** | -1.041 | -0.313 | **0.486** | 7.85E-06 | **3.26E-05** |
| ENSG00000236753.8 | **MKLN1-AS** | 100506881 | lncRNA | **73.185** | 1.106 | 0.333 | **2.152** | 7.85E-06 | **3.26E-05** |
| ENSG00000184232.11 | **OAF** | 220323 | protein_coding | **120.791** | 0.859 | 0.259 | **1.814** | 7.88E-06 | **3.27E-05** |
| ENSG00000067221.16 | **STOML1** | 9399 | protein_coding | **201.063** | 0.686 | 0.206 | **1.608** | 7.89E-06 | **3.27E-05** |

| | | | | | | | | | |
|---|---|---|---|---|---|---|---|---|---|
| ENSG00000288858.3 | **ENSG00000288858** | | lncRNA | **24.582** | 1.433 | 0.431 | **2.699** | 7.89E-06 | **3.27E-05** |
| ENSG00000165914.17 | **TTC7B** | 145567 | protein_coding | **17.599** | 2.063 | 0.621 | **4.177** | 7.90E-06 | **3.28E-05** |
| ENSG00000174109.5 | **UQCC4** | 283951 | protein_coding | **136.956** | -0.661 | -0.199 | **0.632** | 7.95E-06 | **3.30E-05** |
| ENSG00000261455.2 | **LINC01003** | | lncRNA | **52.973** | 0.881 | 0.265 | **1.842** | 7.96E-06 | **3.30E-05** |
| ENSG00000185238.15 | **PRMT3** | 10196 | protein_coding | **192.052** | -0.650 | -0.196 | **0.637** | 7.97E-06 | **3.31E-05** |
| ENSG00000176108.10 | **CHMP6** | 79643 | protein_coding | **281.349** | -0.534 | -0.161 | **0.691** | 7.98E-06 | **3.31E-05** |
| ENSG00000174099.14 | **MSRB3** | 253827 | protein_coding | **43.131** | 2.385 | 0.718 | **5.222** | 8.00E-06 | **3.32E-05** |
| ENSG00000156475.21 | **PPP2R2B** | 5521 | protein_coding | **4.836** | 4.153 | 1.250 | **17.793** | 8.02E-06 | **3.32E-05** |
| ENSG00000289904.2 | **ENSG00000289904** | | lncRNA | **10.079** | 3.144 | 0.946 | **8.841** | 8.05E-06 | **3.34E-05** |
| ENSG00000140395.11 | **SKIC8** | 80349 | protein_coding | **564.348** | -0.422 | -0.127 | **0.747** | 8.10E-06 | **3.36E-05** |
| ENSG00000196313.14 | **POM121** | 9883 | protein_coding | **1171.826** | -0.381 | -0.115 | **0.768** | 8.11E-06 | **3.36E-05** |
| ENSG00000253408.7 | **IKBKB-DT** | 10192989 | lncRNA | **6.172** | 3.356 | 1.010 | **10.241** | 8.14E-06 | **3.37E-05** |
| ENSG00000151790.11 | **TDO2** | 6999 | protein_coding | **7.389** | 3.713 | 1.118 | **13.113** | 8.15E-06 | **3.37E-05** |
| ENSG00000244459.5 | **ENSG00000244459** | | lncRNA | **9.636** | 2.684 | 0.808 | **6.428** | 8.17E-06 | **3.38E-05** |
| ENSG00000157106.20 | **SMG1** | 23049 | protein_coding | **1560.722** | 0.714 | 0.215 | **1.640** | 8.18E-06 | **3.39E-05** |
| ENSG00000286619.1 | **PM20D1-AS1** | | lncRNA | **32.638** | 1.771 | 0.533 | **3.414** | 8.20E-06 | **3.39E-05** |
| ENSG00000111737.14 | **RAB35** | 11021 | protein_coding | **1940.732** | -0.460 | -0.139 | **0.727** | 8.20E-06 | **3.39E-05** |
| ENSG00000071051.16 | **NCK2** | 8440 | protein_coding | **817.548** | 0.588 | 0.177 | **1.503** | 8.28E-06 | **3.42E-05** |
| ENSG00000135525.21 | **MAP7** | 9053 | protein_coding | **11.918** | 2.347 | 0.707 | **5.089** | 8.29E-06 | **3.43E-05** |
| ENSG00000064886.16 | **CHI3L2** | 1117 | protein_coding | **5419.451** | -0.939 | -0.283 | **0.522** | 8.31E-06 | **3.43E-05** |
| ENSG00000198723.12 | **SAXO5** | 374877 | protein_coding | **7.138** | 3.412 | 1.027 | **10.648** | 8.34E-06 | **3.45E-05** |
| ENSG00000061794.15 | **MRPS35** | 60488 | protein_coding | **657.963** | -0.439 | -0.132 | **0.738** | 8.35E-06 | **3.45E-05** |
| ENSG00000291132.2 | **MTMR9LP** | 339483 | lncRNA | **141.289** | 1.033 | 0.311 | **2.046** | 8.36E-06 | **3.45E-05** |
| ENSG00000109606.15 | **DHX15** | 1665 | protein_coding | **1483.693** | -0.509 | -0.153 | **0.703** | 8.36E-06 | **3.45E-05** |
| ENSG00000261448.2 | **ENSG0000(** | 10272369 | lncRNA | **241.611** | 0.951 | 0.286 | **1.933** | 8.45E-06 | **3.49E-05** |
| ENSG00000054690.16 | **PLEKHH1** | 57475 | protein_coding | **32.042** | 1.696 | 0.510 | **3.239** | 8.46E-06 | **3.49E-05** |
| ENSG00000249534.3 | **LINC01258** | 10192877 | lncRNA | **12.672** | 2.453 | 0.738 | **5.475** | 8.48E-06 | **3.50E-05** |
| ENSG00000136098.21 | **NEK3** | 4752 | protein_coding | **292.599** | 0.771 | 0.232 | **1.706** | 8.54E-06 | **3.52E-05** |
| ENSG00000116285.15 | **ERRFI1** | 54206 | protein_coding | **26.455** | 1.688 | 0.508 | **3.223** | 8.54E-06 | **3.52E-05** |
| ENSG00000196275.18 | **GTF2IRD2** | 84163 | protein_coding | **140.052** | -1.210 | -0.364 | **0.432** | 8.54E-06 | **3.52E-05** |
| ENSG00000134121.11 | **CHL1** | 10752 | protein_coding | **179.265** | -0.923 | -0.278 | **0.527** | 8.58E-06 | **3.54E-05** |

| | | | | | | | | | |
|---|---|---|---|---|---|---|---|---|---|
| ENSG00000288964.2 | **ENSG0000(** | 10798693[illegible] | lncRNA | **40.278** | 1.549 | 0.466 | **2.926** | 8.59E-06 | **3.54E-05** |
| ENSG00000289952.2 | **ENSG00000289952** | | lncRNA | **28.850** | 1.250 | 0.376 | **2.378** | 8.60E-06 | **3.54E-05** |
| ENSG00000301630.1 | **ENSG0000(** | 10798607[illegible] | lncRNA | **10.051** | 2.418 | 0.728 | **5.346** | 8.63E-06 | **3.56E-05** |
| ENSG00000213578.7 | **CPLX3** | 594855 | protein_coding | **14.889** | 2.111 | 0.635 | **4.319** | 8.64E-06 | **3.56E-05** |
| ENSG00000102934.12 | **PLLP** | 51090 | protein_coding | **31.357** | 1.388 | 0.418 | **2.616** | 8.68E-06 | **3.58E-05** |
| ENSG00000135124.18 | **P2RX4** | 5025 | protein_coding | **593.801** | -0.551 | -0.166 | **0.682** | 8.70E-06 | **3.58E-05** |
| ENSG00000149260.20 | **CAPN5** | 726 | protein_coding | **23.277** | 2.596 | 0.781 | **6.045** | 8.71E-06 | **3.59E-05** |
| ENSG00000108091.13 | **CCDC6** | 8030 | protein_coding | **826.702** | 0.631 | 0.190 | **1.549** | 8.72E-06 | **3.59E-05** |
| ENSG00000273045.9 | **C2orf15** | 150590 | protein_coding | **44.324** | 1.576 | 0.474 | **2.981** | 8.72E-06 | **3.59E-05** |
| ENSG00000172915.21 | **NBEA** | 26960 | protein_coding | **40.048** | 1.233 | 0.371 | **2.350** | 8.72E-06 | **3.59E-05** |
| ENSG00000076108.14 | **BAZ2A** | 11176 | protein_coding | **2544.228** | 0.602 | 0.181 | **1.518** | 8.77E-06 | **3.61E-05** |
| ENSG00000128422.20 | **KRT17** | 3872 | protein_coding | **27.016** | -2.090 | -0.629 | **0.235** | 8.78E-06 | **3.61E-05** |
| ENSG00000188937.9 | **NYX** | 60506 | protein_coding | **4.932** | 4.181 | 1.259 | **18.138** | 8.79E-06 | **3.62E-05** |
| ENSG00000088356.8 | **PDRG1** | 81572 | protein_coding | **372.415** | -0.521 | -0.157 | **0.697** | 8.86E-06 | **3.64E-05** |
| ENSG00000118156.14 | **ZNF541** | 84215 | protein_coding | **8.091** | 2.904 | 0.874 | **7.486** | 8.88E-06 | **3.65E-05** |
| ENSG00000129351.20 | **ILF3** | 3609 | protein_coding | **6168.678** | -0.311 | -0.094 | **0.806** | 8.92E-06 | **3.66E-05** |
| ENSG00000266088.7 | **ENSG0000(** | 10537177[illegible] | lncRNA | **30.617** | -1.454 | -0.438 | **0.365** | 8.92E-06 | **3.66E-05** |
| ENSG00000246145.2 | **RRS1-DT** | 10050567[illegible] | lncRNA | **11.511** | 2.192 | 0.660 | **4.571** | 8.93E-06 | **3.67E-05** |
| ENSG00000267534.4 | **S1PR2** | 9294 | protein_coding | **513.553** | -0.533 | -0.161 | **0.691** | 8.96E-06 | **3.68E-05** |
| ENSG00000147168.15 | **IL2RG** | 3561 | protein_coding | **6326.236** | -0.687 | -0.207 | **0.621** | 8.96E-06 | **3.68E-05** |
| ENSG00000309414.1 | **ENSG00000309414** | | lncRNA | **10.161** | -2.787 | -0.839 | **0.145** | 8.97E-06 | **3.68E-05** |
| ENSG00000188089.14 | **PLA2G4E** | 123745 | protein_coding | **15.331** | 1.779 | 0.536 | **3.432** | 8.97E-06 | **3.68E-05** |
| ENSG00000213468.8 | **FIRRE** | 286467 | lncRNA | **165.123** | -0.744 | -0.224 | **0.597** | 8.99E-06 | **3.69E-05** |
| ENSG00000288908.1 | **ENSG00000288908** | | lncRNA | **65.926** | 0.994 | 0.299 | **1.992** | 9.00E-06 | **3.69E-05** |
| ENSG00000132740.12 | **IGHMBP2** | 3508 | protein_coding | **379.854** | -0.673 | -0.203 | **0.627** | 9.02E-06 | **3.70E-05** |
| ENSG00000142556.21 | **ZNF614** | 80110 | protein_coding | **93.702** | -0.864 | -0.260 | **0.550** | 9.07E-06 | **3.72E-05** |
| ENSG00000255234.7 | **CCDC90B-AS1** | | lncRNA | **7.768** | 3.269 | 0.984 | **9.639** | 9.08E-06 | **3.72E-05** |
| ENSG00000175573.8 | **C11orf68** | 83638 | protein_coding | **505.622** | -0.508 | -0.153 | **0.703** | 9.09E-06 | **3.73E-05** |
| ENSG00000307466.1 | **ENSG00000307466** | | lncRNA | **23.844** | 2.053 | 0.618 | **4.149** | 9.14E-06 | **3.74E-05** |
| ENSG00000144559.13 | **TAMM41** | 132001 | protein_coding | **269.214** | -0.519 | -0.156 | **0.698** | 9.21E-06 | **3.77E-05** |
| ENSG00000165215.6 | **CLDN3** | 1365 | protein_coding | **3.940** | 4.780 | 1.439 | **27.483** | 9.23E-06 | **3.78E-05** |

| | | | | | | | | | |
|---|---|---|---|---|---|---|---|---|---|
| ENSG00000110066.17 | **KMT5B** | 51111 | protein_coding | **492.673** | 0.595 | 0.179 | **1.510** | 9.25E-06 | **3.79E-05** |
| ENSG00000100650.18 | **SRSF5** | 6430 | protein_coding | **7575.328** | 0.388 | 0.117 | **1.309** | 9.26E-06 | **3.79E-05** |
| ENSG00000130119.19 | **GNL3L** | 54552 | protein_coding | **323.740** | -0.643 | -0.193 | **0.641** | 9.29E-06 | **3.80E-05** |
| ENSG00000198482.16 | **ZNF808** | 388558 | protein_coding | **151.123** | -0.770 | -0.232 | **0.586** | 9.31E-06 | **3.81E-05** |
| ENSG00000158169.16 | **FANCC** | 2176 | protein_coding | **163.504** | -0.572 | -0.172 | **0.673** | 9.32E-06 | **3.81E-05** |
| ENSG00000132359.19 | **RAP1GAP2** | 23108 | protein_coding | **64.131** | 1.662 | 0.500 | **3.165** | 9.33E-06 | **3.82E-05** |
| ENSG00000152969.22 | **JAKMIP1** | 152789 | protein_coding | **125.827** | 0.741 | 0.223 | **1.671** | 9.42E-06 | **3.85E-05** |
| ENSG00000310521.1 | **ENSG0000(** | 10537572ξ | lncRNA | **34.448** | 1.123 | 0.338 | **2.178** | 9.42E-06 | **3.85E-05** |
| ENSG00000164867.13 | **NOS3** | 4846 | protein_coding | **32.391** | 1.591 | 0.479 | **3.013** | 9.43E-06 | **3.86E-05** |
| ENSG00000175348.13 | **TMEM9B** | 56674 | protein_coding | **392.968** | 0.447 | 0.135 | **1.363** | 9.45E-06 | **3.86E-05** |
| ENSG00000136378.17 | **ADAMTS7** | 11173 | protein_coding | **167.582** | 0.834 | 0.251 | **1.783** | 9.50E-06 | **3.88E-05** |
| ENSG00000118873.19 | **RAB3GAP2** | 25782 | protein_coding | **769.245** | -0.500 | -0.150 | **0.707** | 9.51E-06 | **3.89E-05** |
| ENSG00000168792.6 | **ABHD15** | 116236 | protein_coding | **199.322** | 0.540 | 0.163 | **1.454** | 9.52E-06 | **3.89E-05** |
| ENSG00000111644.9 | **ACRBP** | 84519 | protein_coding | **65.706** | 1.175 | 0.354 | **2.259** | 9.53E-06 | **3.89E-05** |
| ENSG00000130812.13 | **ANGPTL6** | 83854 | protein_coding | **1032.395** | -0.613 | -0.184 | **0.654** | 9.54E-06 | **3.89E-05** |
| ENSG00000204519.12 | **ZNF551** | 90233 | protein_coding | **68.492** | -0.800 | -0.241 | **0.574** | 9.54E-06 | **3.89E-05** |
| ENSG00000304289.1 | **ENSG00000304289** | | lncRNA | **12.714** | 1.807 | 0.544 | **3.500** | 9.54E-06 | **3.89E-05** |
| ENSG00000290865.1 | **RRN3P3** | 10013199ξ | lncRNA | **137.130** | -0.686 | -0.207 | **0.621** | 9.59E-06 | **3.91E-05** |
| ENSG00000160256.16 | **SLX9** | 85395 | protein_coding | **289.071** | -0.534 | -0.161 | **0.691** | 9.63E-06 | **3.93E-05** |
| ENSG00000124562.12 | **SNRPC** | 6631 | protein_coding | **1993.890** | -0.500 | -0.150 | **0.707** | 9.63E-06 | **3.93E-05** |
| ENSG00000152404.18 | **CWF19L2** | 143884 | protein_coding | **167.083** | -0.673 | -0.203 | **0.627** | 9.66E-06 | **3.94E-05** |
| ENSG00000100124.17 | **ANKRD54** | 129138 | protein_coding | **522.529** | 0.396 | 0.119 | **1.316** | 9.68E-06 | **3.95E-05** |
| ENSG00000177733.8 | **HNRNPA0** | 10949 | protein_coding | **2278.637** | -0.396 | -0.119 | **0.760** | 9.70E-06 | **3.95E-05** |
| ENSG00000254986.10 | **DPP3** | 10072 | protein_coding | **749.031** | -0.696 | -0.210 | **0.617** | 9.70E-06 | **3.95E-05** |
| ENSG00000105339.13 | **DENND3** | 22898 | protein_coding | **2061.871** | 0.559 | 0.168 | **1.473** | 9.77E-06 | **3.98E-05** |
| ENSG00000143028.11 | **SYPL2** | 284612 | protein_coding | **13.812** | 2.233 | 0.672 | **4.700** | 9.79E-06 | **3.99E-05** |
| ENSG00000116984.17 | **MTR** | 4548 | protein_coding | **821.511** | -0.571 | -0.172 | **0.673** | 9.79E-06 | **3.99E-05** |
| ENSG00000234772.4 | **LINC00412** | | lncRNA | **49.159** | 1.392 | 0.419 | **2.624** | 9.84E-06 | **4.01E-05** |
| ENSG00000124444.18 | **ZNF576** | 79177 | protein_coding | **132.698** | -0.756 | -0.227 | **0.592** | 9.84E-06 | **4.01E-05** |
| ENSG00000101084.21 | **RAB5IF** | 55969 | protein_coding | **724.641** | -0.393 | -0.118 | **0.762** | 9.85E-06 | **4.01E-05** |
| ENSG00000134245.19 | **WNT2B** | 7482 | protein_coding | **20.609** | 1.588 | 0.478 | **3.007** | 9.86E-06 | **4.01E-05** |

| | | | | | | | | | |
|---|---|---|---|---|---|---|---|---|---|
| ENSG00000271936.1 | **ENSG00000271936** | | lncRNA | **17.554** | -1.655 | -0.498 | **0.318** | 9.86E-06 | **4.01E-05** |
| ENSG00000119684.18 | **MLH3** | 27030 | protein_coding | **366.961** | -0.574 | -0.173 | **0.672** | 9.89E-06 | **4.02E-05** |
| ENSG00000155099.10 | **PIP4P2** | 55529 | protein_coding | **144.537** | -0.875 | -0.263 | **0.545** | 9.90E-06 | **4.03E-05** |
| ENSG00000122674.14 | **CCZ1** | 51622 | protein_coding | **516.021** | -0.632 | -0.190 | **0.645** | 9.95E-06 | **4.05E-05** |
| ENSG00000127838.17 | **PNKD** | 25953 | protein_coding | **921.195** | 0.394 | 0.119 | **1.314** | 9.98E-06 | **4.05E-05** |
| ENSG00000112812.18 | **PRSS16** | 10279 | protein_coding | **67.416** | 1.291 | 0.389 | **2.447** | 9.99E-06 | **4.06E-05** |
| ENSG00000116127.21 | **ALMS1** | 7840 | protein_coding | **216.714** | -0.604 | -0.182 | **0.658** | 1.00E-05 | **4.07E-05** |
| ENSG00000259940.2 | **ENSG00000259940** | | lncRNA | **14.814** | 1.749 | 0.527 | **3.362** | 1.00E-05 | **4.07E-05** |
| ENSG00000251562.13 | **MALAT1** | 378938 | lncRNA | **19000.521** | 0.828 | 0.249 | **1.775** | 1.00E-05 | **4.08E-05** |
| ENSG00000251615.3 | **ENSG00000251615** | | lncRNA | **46.137** | 0.954 | 0.287 | **1.937** | 1.01E-05 | **4.09E-05** |
| ENSG00000213742.9 | **ZNF337-AS** | 10272482( | lncRNA | **62.746** | 0.941 | 0.283 | **1.920** | 1.01E-05 | **4.09E-05** |
| ENSG00000125355.18 | **TMEM255/** | 55026 | protein_coding | **21.221** | 2.137 | 0.643 | **4.399** | 1.01E-05 | **4.11E-05** |
| ENSG00000169752.19 | **NRG4** | 145957 | protein_coding | **44.665** | 1.769 | 0.533 | **3.408** | 1.02E-05 | **4.12E-05** |
| ENSG00000213654.12 | **GPSM3** | 63940 | protein_coding | **2557.375** | -0.550 | -0.166 | **0.683** | 1.02E-05 | **4.14E-05** |
| ENSG00000114021.14 | **NIT2** | 56954 | protein_coding | **1340.559** | -0.432 | -0.130 | **0.741** | 1.03E-05 | **4.17E-05** |
| ENSG00000110801.16 | **PSMD9** | 5715 | protein_coding | **590.265** | -0.363 | -0.109 | **0.777** | 1.03E-05 | **4.18E-05** |
| ENSG00000136045.14 | **PWP1** | 11137 | protein_coding | **887.241** | -0.448 | -0.135 | **0.733** | 1.03E-05 | **4.18E-05** |
| ENSG00000186767.7 | **SPIN4** | 139886 | protein_coding | **43.052** | -0.953 | -0.287 | **0.517** | 1.03E-05 | **4.19E-05** |
| ENSG00000108405.6 | **P2RX1** | 5023 | protein_coding | **551.902** | -0.837 | -0.252 | **0.560** | 1.04E-05 | **4.20E-05** |
| ENSG00000137496.21 | **IL18BP** | 10068 | protein_coding | **511.827** | -0.707 | -0.213 | **0.613** | 1.04E-05 | **4.21E-05** |
| ENSG00000274929.1 | **ENSG00000274929** | | lncRNA | **22.359** | 1.433 | 0.431 | **2.699** | 1.04E-05 | **4.21E-05** |
| ENSG00000188613.8 | **NANOS1** | 340719 | protein_coding | **40.778** | 1.476 | 0.444 | **2.782** | 1.04E-05 | **4.22E-05** |
| ENSG00000144115.20 | **THNSL2** | 55258 | protein_coding | **24.966** | 2.315 | 0.697 | **4.976** | 1.05E-05 | **4.24E-05** |
| ENSG00000106330.14 | **MOSPD3** | 64598 | protein_coding | **228.234** | 0.615 | 0.185 | **1.532** | 1.05E-05 | **4.25E-05** |
| ENSG00000137166.20 | **FOXP4** | 116113 | protein_coding | **1209.984** | 0.723 | 0.218 | **1.651** | 1.05E-05 | **4.26E-05** |
| ENSG00000272221.2 | **MICA-AS1** | | lncRNA | **103.883** | 0.782 | 0.236 | **1.720** | 1.05E-05 | **4.26E-05** |
| ENSG00000248019.4 | **FAM13A-AS1** | | lncRNA | **90.554** | 0.980 | 0.295 | **1.973** | 1.06E-05 | **4.27E-05** |
| ENSG00000013374.19 | **NUB1** | 51667 | protein_coding | **1598.827** | 0.650 | 0.196 | **1.569** | 1.06E-05 | **4.27E-05** |
| ENSG00000211896.7 | **IGHG1** | | IG_C_gene | **87776.473** | -0.541 | -0.163 | **0.687** | 1.06E-05 | **4.27E-05** |
| ENSG00000084754.14 | **HADHA** | 3030 | protein_coding | **3459.264** | -0.375 | -0.113 | **0.771** | 1.06E-05 | **4.29E-05** |
| ENSG00000162104.12 | **ADCY9** | 115 | protein_coding | **32.821** | 1.288 | 0.388 | **2.442** | 1.06E-05 | **4.30E-05** |

| | | | | | | | | | |
|---|---|---|---|---|---|---|---|---|---|
| ENSG00000197363.13 | **ZNF517** | 340385 | protein_coding | **101.791** | 0.658 | 0.198 | **1.577** | 1.07E-05 | **4.31E-05** |
| ENSG00000121775.20 | **TMEM39B** | 55116 | protein_coding | **574.135** | -0.481 | -0.145 | **0.716** | 1.07E-05 | **4.32E-05** |
| ENSG00000083844.12 | **ZNF264** | 9422 | protein_coding | **265.296** | -0.873 | -0.263 | **0.546** | 1.07E-05 | **4.32E-05** |
| ENSG00000100600.17 | **LGMN** | 5641 | protein_coding | **3186.477** | 0.596 | 0.179 | **1.511** | 1.07E-05 | **4.33E-05** |
| ENSG00000197935.9 | **ZNF311** | 282890 | protein_coding | **13.827** | 2.133 | 0.642 | **4.385** | 1.07E-05 | **4.33E-05** |
| ENSG00000179088.17 | **C12orf42** | 374470 | protein_coding | **35.609** | 1.829 | 0.551 | **3.553** | 1.07E-05 | **4.33E-05** |
| ENSG00000198939.10 | **ZFP2** | 80108 | protein_coding | **10.412** | 2.103 | 0.633 | **4.297** | 1.07E-05 | **4.34E-05** |
| ENSG00000281731.4 | **METTL14-D** | 101929741 | lncRNA | **38.939** | 1.206 | 0.363 | **2.308** | 1.08E-05 | **4.35E-05** |
| ENSG00000170889.16 | **RPS9** | 6203 | protein_coding | **17768.540** | -0.418 | -0.126 | **0.748** | 1.08E-05 | **4.35E-05** |
| ENSG00000182759.4 | **MAFA** | 389692 | protein_coding | **26.087** | 1.759 | 0.530 | **3.386** | 1.08E-05 | **4.37E-05** |
| ENSG00000287780.2 | **ENSG00000287780** | | lncRNA | **35.786** | 1.097 | 0.330 | **2.138** | 1.08E-05 | **4.37E-05** |
| ENSG00000186231.19 | **KLHL32** | 114792 | protein_coding | **22.704** | 1.454 | 0.438 | **2.740** | 1.08E-05 | **4.37E-05** |
| ENSG00000112200.19 | **ZNF451** | 26036 | protein_coding | **521.334** | -0.539 | -0.162 | **0.688** | 1.09E-05 | **4.37E-05** |
| ENSG00000133059.20 | **DSTYK** | 25778 | protein_coding | **47.450** | 1.091 | 0.328 | **2.130** | 1.09E-05 | **4.39E-05** |
| ENSG00000260625.5 | **RUSF1-DT** | 12370652ε | lncRNA | **18.708** | 1.574 | 0.474 | **2.977** | 1.09E-05 | **4.39E-05** |
| ENSG00000246528.5 | **SLCO5A1-A** | 12123392Ε | lncRNA | **58.035** | 0.953 | 0.287 | **1.936** | 1.09E-05 | **4.40E-05** |
| ENSG00000187796.18 | **CARD9** | 64170 | protein_coding | **181.804** | -1.327 | -0.400 | **0.399** | 1.09E-05 | **4.41E-05** |
| ENSG00000291110.2 | **TRIM16L** | 147166 | lncRNA | **112.293** | 0.701 | 0.211 | **1.626** | 1.10E-05 | **4.44E-05** |
| ENSG00000005700.17 | **IBTK** | 25998 | protein_coding | **439.661** | -0.565 | -0.170 | **0.676** | 1.10E-05 | **4.44E-05** |
| ENSG00000274286.2 | **ADRA2B** | 151 | protein_coding | **16.154** | 2.046 | 0.616 | **4.129** | 1.11E-05 | **4.45E-05** |
| ENSG00000155380.15 | **SLC16A1** | 6566 | protein_coding | **442.144** | -0.537 | -0.162 | **0.689** | 1.11E-05 | **4.46E-05** |
| ENSG00000170584.14 | **NUDCD2** | 134492 | protein_coding | **372.123** | -0.543 | -0.163 | **0.686** | 1.12E-05 | **4.50E-05** |
| ENSG00000117586.14 | **TNFSF4** | 7292 | protein_coding | **154.091** | -1.043 | -0.314 | **0.485** | 1.13E-05 | **4.53E-05** |
| ENSG00000120314.20 | **WDR55** | 54853 | protein_coding | **778.620** | -0.460 | -0.139 | **0.727** | 1.14E-05 | **4.57E-05** |
| ENSG00000308294.1 | **ENSG00000308294** | | lncRNA | **26.549** | 1.323 | 0.398 | **2.502** | 1.14E-05 | **4.58E-05** |
| ENSG00000293616.2 | **PCBP1-AS1** | 400960 | lncRNA | **519.263** | 0.593 | 0.179 | **1.509** | 1.15E-05 | **4.60E-05** |
| ENSG00000171617.18 | **ENC1** | 8507 | protein_coding | **390.952** | -0.826 | -0.249 | **0.564** | 1.15E-05 | **4.62E-05** |
| ENSG00000143575.18 | **HAX1** | 10456 | protein_coding | **2389.289** | -0.595 | -0.179 | **0.662** | 1.15E-05 | **4.63E-05** |
| ENSG00000197859.13 | **ADAMTSL2** | 9719 | protein_coding | **62.137** | 1.039 | 0.313 | **2.054** | 1.15E-05 | **4.63E-05** |
| ENSG00000173917.12 | **HOXB2** | 3212 | protein_coding | **133.912** | -0.936 | -0.282 | **0.523** | 1.15E-05 | **4.63E-05** |
| ENSG00000188725.10 | **SMIM15** | 643155 | protein_coding | **407.321** | -0.589 | -0.177 | **0.665** | 1.15E-05 | **4.64E-05** |

| | | | | | | | | | |
|---|---|---|---|---|---|---|---|---|---|
| ENSG00000293314.2 | **SDAD1P1** | 157489 | lncRNA | **42.990** | 1.271 | 0.383 | **2.414** | 1.16E-05 | **4.64E-05** |
| ENSG00000269939.1 | **PCF11-AS1** | | lncRNA | **45.559** | 0.903 | 0.272 | **1.870** | 1.16E-05 | **4.66E-05** |
| ENSG00000065413.22 | **ANKRD44** | 91526 | protein_coding | **436.752** | -0.579 | -0.174 | **0.669** | 1.16E-05 | **4.66E-05** |
| ENSG00000100197.25 | **CYP2D6** | 1565 | protein_coding | **102.635** | 0.844 | 0.254 | **1.795** | 1.17E-05 | **4.69E-05** |
| ENSG00000291211.2 | **ENSG0000(** | 440300 | lncRNA | **10.523** | 2.744 | 0.826 | **6.697** | 1.17E-05 | **4.69E-05** |
| ENSG00000180104.18 | **EXOC3** | 11336 | protein_coding | **1466.694** | 0.397 | 0.119 | **1.317** | 1.18E-05 | **4.73E-05** |
| ENSG00000260997.1 | **ENSG00000260997** | | lncRNA | **793.128** | 0.619 | 0.186 | **1.536** | 1.18E-05 | **4.73E-05** |
| ENSG00000169418.12 | **NPR1** | 4881 | protein_coding | **54.389** | -1.277 | -0.384 | **0.413** | 1.18E-05 | **4.74E-05** |
| ENSG00000092020.13 | **PPP2R3C** | 55012 | protein_coding | **214.693** | -0.692 | -0.208 | **0.619** | 1.18E-05 | **4.74E-05** |
| ENSG00000105668.9 | **UPK1A** | 11045 | protein_coding | **19.654** | 1.975 | 0.594 | **3.930** | 1.18E-05 | **4.74E-05** |
| ENSG00000107738.22 | **VSIR** | 64115 | protein_coding | **370.596** | -0.503 | -0.151 | **0.706** | 1.18E-05 | **4.74E-05** |
| ENSG00000256537.6 | **SMIM10L1** | 100129361 | protein_coding | **397.734** | -0.529 | -0.159 | **0.693** | 1.20E-05 | **4.79E-05** |
| ENSG00000196172.11 | **ZNF681** | 148213 | protein_coding | **73.713** | -1.085 | -0.327 | **0.471** | 1.20E-05 | **4.79E-05** |
| ENSG00000159882.15 | **ZNF230** | 7773 | protein_coding | **130.924** | -0.632 | -0.190 | **0.645** | 1.20E-05 | **4.80E-05** |
| ENSG00000049769.16 | **PPP1R3F** | 89801 | protein_coding | **148.099** | 0.654 | 0.197 | **1.574** | 1.20E-05 | **4.82E-05** |
| ENSG00000258388.7 | **PPT2-EGFL8** | | protein_coding | **20.243** | 2.185 | 0.658 | **4.548** | 1.21E-05 | **4.86E-05** |
| ENSG00000185721.16 | **DRG1** | 4733 | protein_coding | **677.074** | -0.443 | -0.133 | **0.735** | 1.22E-05 | **4.88E-05** |
| ENSG00000102316.19 | **MAGED2** | 10916 | protein_coding | **807.047** | -0.542 | -0.163 | **0.687** | 1.22E-05 | **4.90E-05** |
| ENSG00000169105.8 | **CHST14** | 113189 | protein_coding | **286.781** | -0.647 | -0.195 | **0.639** | 1.23E-05 | **4.90E-05** |
| ENSG00000229847.10 | **EMX2OS** | 196047 | lncRNA | **5.904** | 4.473 | 1.347 | **22.211** | 1.23E-05 | **4.92E-05** |
| ENSG00000165025.17 | **SYK** | 6850 | protein_coding | **926.267** | -0.542 | -0.163 | **0.687** | 1.23E-05 | **4.92E-05** |
| ENSG00000135775.16 | **COG2** | 22796 | protein_coding | **482.051** | 0.483 | 0.145 | **1.397** | 1.24E-05 | **4.95E-05** |
| ENSG00000185088.16 | **RPS27L** | 51065 | protein_coding | **1816.765** | -0.616 | -0.185 | **0.653** | 1.24E-05 | **4.96E-05** |
| ENSG00000104823.11 | **ECH1** | 1891 | protein_coding | **3323.742** | -0.562 | -0.169 | **0.677** | 1.24E-05 | **4.97E-05** |
| ENSG00000248049.9 | **UBA6-DT** | | lncRNA | **298.587** | -0.659 | -0.198 | **0.633** | 1.25E-05 | **4.98E-05** |
| ENSG00000140284.13 | **SLC27A2** | 11001 | protein_coding | **24.181** | 1.589 | 0.478 | **3.009** | 1.25E-05 | **4.98E-05** |
| ENSG00000293263.1 | **ENSG00000293263** | | lncRNA | **10.781** | 2.622 | 0.789 | **6.155** | 1.25E-05 | **4.99E-05** |
| ENSG00000299022.1 | **ENSG00000299022** | | lncRNA | **498.376** | -1.116 | -0.336 | **0.461** | 1.25E-05 | **5.00E-05** |
| ENSG00000170370.13 | **EMX2** | 2018 | protein_coding | **4.167** | 4.551 | 1.370 | **23.438** | 1.25E-05 | **5.00E-05** |
| ENSG00000147894.19 | **C9orf72** | 203228 | protein_coding | **179.191** | 0.692 | 0.208 | **1.616** | 1.26E-05 | **5.02E-05** |
| ENSG00000109787.15 | **KLF3** | 51274 | protein_coding | **207.743** | 0.672 | 0.202 | **1.593** | 1.26E-05 | **5.05E-05** |

| | | | | | | | | | |
|---|---|---|---|---|---|---|---|---|---|
| ENSG00000283235.3 | **ENSG00000283235** | | lncRNA | **7.571** | 3.171 | 0.955 | **9.006** | 1.27E-05 | **5.05E-05** |
| ENSG00000281103.5 | **TRG-AS1** | 10050677€ | lncRNA | **234.170** | -0.651 | -0.196 | **0.637** | 1.27E-05 | **5.06E-05** |
| ENSG00000305051.1 | **ENSG00000305051** | | lncRNA | **20.957** | 1.292 | 0.389 | **2.448** | 1.27E-05 | **5.06E-05** |
| ENSG00000164951.17 | **PDP1** | 54704 | protein_coding | **87.429** | 1.019 | 0.307 | **2.027** | 1.28E-05 | **5.09E-05** |
| ENSG00000105518.16 | **TMEM205** | 374882 | protein_coding | **1038.250** | 0.447 | 0.135 | **1.363** | 1.28E-05 | **5.09E-05** |
| ENSG00000174721.11 | **FGFBP3** | 143282 | protein_coding | **24.919** | 1.465 | 0.441 | **2.761** | 1.29E-05 | **5.14E-05** |
| ENSG00000251192.10 | **ZNF674** | 641339 | protein_coding | **91.463** | 1.157 | 0.348 | **2.230** | 1.29E-05 | **5.15E-05** |
| ENSG00000119408.19 | **NEK6** | 10783 | protein_coding | **2502.718** | -0.639 | -0.192 | **0.642** | 1.29E-05 | **5.15E-05** |
| ENSG00000131067.19 | **GGT7** | 2686 | protein_coding | **380.979** | 0.523 | 0.157 | **1.437** | 1.30E-05 | **5.16E-05** |
| ENSG00000117592.11 | **PRDX6** | 9588 | protein_coding | **1928.015** | -0.384 | -0.116 | **0.766** | 1.31E-05 | **5.20E-05** |
| ENSG00000281344.1 | **HELLPAR** | | lncRNA | **34.380** | -1.199 | -0.361 | **0.436** | 1.31E-05 | **5.20E-05** |
| ENSG00000089094.22 | **KDM2B** | 84678 | protein_coding | **2111.181** | 1.398 | 0.421 | **2.635** | 1.31E-05 | **5.22E-05** |
| ENSG00000225953.6 | **SATB2-AS1** | 150538 | lncRNA | **5.930** | 3.402 | 1.024 | **10.573** | 1.31E-05 | **5.22E-05** |
| ENSG00000173546.9 | **CSPG4** | 1464 | protein_coding | **16.370** | -2.059 | -0.620 | **0.240** | 1.32E-05 | **5.24E-05** |
| ENSG00000206530.12 | **CFAP44** | 55779 | protein_coding | **211.158** | 0.624 | 0.188 | **1.541** | 1.32E-05 | **5.26E-05** |
| ENSG00000010219.15 | **DYRK4** | 8798 | protein_coding | **468.863** | -0.508 | -0.153 | **0.703** | 1.32E-05 | **5.26E-05** |
| ENSG00000084444.16 | **FAM234B** | 57613 | protein_coding | **43.302** | 1.016 | 0.306 | **2.022** | 1.33E-05 | **5.28E-05** |
| ENSG00000074695.8 | **LMAN1** | 3998 | protein_coding | **1094.742** | -0.556 | -0.167 | **0.680** | 1.33E-05 | **5.29E-05** |
| ENSG00000275143.1 | **SCARNA16** | 677781 | scaRNA | **6.667** | 2.898 | 0.873 | **7.456** | 1.34E-05 | **5.31E-05** |
| ENSG00000101040.22 | **ZMYND8** | 23613 | protein_coding | **1288.443** | -0.576 | -0.173 | **0.671** | 1.34E-05 | **5.31E-05** |
| ENSG00000141027.25 | **NCOR1** | 9611 | protein_coding | **1109.775** | -0.497 | -0.150 | **0.708** | 1.34E-05 | **5.32E-05** |
| ENSG00000204138.15 | **PHACTR4** | 65979 | protein_coding | **334.870** | -0.537 | -0.162 | **0.689** | 1.34E-05 | **5.33E-05** |
| ENSG00000278195.4 | **SSTR3** | 6753 | protein_coding | **471.139** | 0.628 | 0.189 | **1.545** | 1.35E-05 | **5.34E-05** |
| ENSG00000259577.3 | **CERNA1** | | lncRNA | **19.887** | 1.615 | 0.486 | **3.063** | 1.35E-05 | **5.35E-05** |
| ENSG00000113716.15 | **HMGXB3** | 22993 | protein_coding | **1419.281** | -0.448 | -0.135 | **0.733** | 1.35E-05 | **5.36E-05** |
| ENSG00000148411.10 | **NACC2** | 138151 | protein_coding | **202.399** | 0.699 | 0.210 | **1.623** | 1.35E-05 | **5.36E-05** |
| ENSG00000006125.20 | **AP2B1** | 163 | protein_coding | **1493.206** | -0.479 | -0.144 | **0.717** | 1.35E-05 | **5.36E-05** |
| ENSG00000198740.11 | **ZNF652** | 22834 | protein_coding | **607.505** | 0.508 | 0.153 | **1.423** | 1.36E-05 | **5.38E-05** |
| ENSG00000140105.20 | **WARS1** | 7453 | protein_coding | **6289.210** | -0.870 | -0.262 | **0.547** | 1.36E-05 | **5.40E-05** |
| ENSG00000204540.11 | **PSORS1C1** | 170679 | protein_coding | **10.209** | 2.264 | 0.682 | **4.803** | 1.36E-05 | **5.40E-05** |
| ENSG00000196782.15 | **MAML3** | 55534 | protein_coding | **34.570** | 2.421 | 0.729 | **5.356** | 1.36E-05 | **5.40E-05** |

| | | | | | | | | | |
|---|---|---|---|---|---|---|---|---|---|
| ENSG00000186567.16 | **CEACAM19** | 56971 | protein_coding | **9.882** | 2.308 | 0.695 | **4.953** | 1.37E-05 | **5.43E-05** |
| ENSG00000253250.5 | **C8orf88** | 100127983 | protein_coding | **70.330** | -0.849 | -0.255 | **0.555** | 1.38E-05 | **5.45E-05** |
| ENSG00000135390.23 | **ATP5MC2** | 517 | protein_coding | **7055.749** | -0.511 | -0.154 | **0.702** | 1.38E-05 | **5.45E-05** |
| ENSG00000152076.21 | **CCDC74B** | 91409 | protein_coding | **123.880** | 1.101 | 0.332 | **2.145** | 1.38E-05 | **5.47E-05** |
| ENSG00000104231.13 | **ZFAND1** | 79752 | protein_coding | **469.792** | -0.666 | -0.201 | **0.630** | 1.38E-05 | **5.47E-05** |
| ENSG00000134202.14 | **GSTM3** | 2947 | protein_coding | **111.327** | 1.286 | 0.387 | **2.438** | 1.39E-05 | **5.52E-05** |
| ENSG00000251676.3 | **SNHG27** | 101927305 | lncRNA | **9.780** | -2.566 | -0.773 | **0.169** | 1.40E-05 | **5.53E-05** |
| ENSG00000148219.20 | **ASTN2** | 23245 | protein_coding | **57.574** | -0.822 | -0.248 | **0.566** | 1.40E-05 | **5.54E-05** |
| ENSG00000244676.6 | **ENSG00000244676** | | lncRNA | **10.278** | 2.898 | 0.872 | **7.456** | 1.41E-05 | **5.57E-05** |
| ENSG00000166987.17 | **MBD6** | 114785 | protein_coding | **1480.273** | 0.620 | 0.187 | **1.536** | 1.41E-05 | **5.57E-05** |
| ENSG00000174672.18 | **BRSK2** | 9024 | protein_coding | **9.570** | 3.292 | 0.991 | **9.795** | 1.41E-05 | **5.57E-05** |
| ENSG00000234965.4 | **SHISA8** | 440829 | protein_coding | **233.431** | -0.927 | -0.279 | **0.526** | 1.41E-05 | **5.58E-05** |
| ENSG00000300585.1 | **ENSG0000(** | 102724883 | lncRNA | **18.645** | 1.706 | 0.513 | **3.262** | 1.42E-05 | **5.60E-05** |
| ENSG00000126088.16 | **UROD** | 7389 | protein_coding | **817.680** | -0.423 | -0.127 | **0.746** | 1.43E-05 | **5.65E-05** |
| ENSG00000149016.18 | **TUT1** | 64852 | protein_coding | **874.471** | 0.441 | 0.133 | **1.358** | 1.43E-05 | **5.65E-05** |
| ENSG00000291114.2 | **SIGLEC17P** | 284367 | lncRNA | **10.587** | 2.393 | 0.720 | **5.252** | 1.43E-05 | **5.66E-05** |
| ENSG00000176170.16 | **SPHK1** | 8877 | protein_coding | **28.805** | 1.301 | 0.392 | **2.464** | 1.43E-05 | **5.66E-05** |
| ENSG00000099956.22 | **SMARCB1** | 6598 | protein_coding | **2032.941** | -0.426 | -0.128 | **0.745** | 1.44E-05 | **5.67E-05** |
| ENSG00000109705.9 | **NKX3-2** | 579 | protein_coding | **5.603** | 3.796 | 1.143 | **13.894** | 1.44E-05 | **5.68E-05** |
| ENSG00000143815.17 | **LBR** | 3930 | protein_coding | **703.609** | -0.593 | -0.179 | **0.663** | 1.44E-05 | **5.70E-05** |
| ENSG00000139364.12 | **TMEM132I** | 114795 | protein_coding | **16.956** | 2.084 | 0.627 | **4.241** | 1.45E-05 | **5.72E-05** |
| ENSG00000178150.12 | **ZNF114** | 163071 | protein_coding | **13.183** | -2.061 | -0.620 | **0.240** | 1.45E-05 | **5.72E-05** |
| ENSG00000152492.17 | **CCDC50** | 152137 | protein_coding | **1720.466** | -0.453 | -0.136 | **0.731** | 1.46E-05 | **5.75E-05** |
| ENSG00000288568.2 | **ZCCHC14-D** | 101928737 | lncRNA | **8.520** | 2.810 | 0.846 | **7.015** | 1.46E-05 | **5.77E-05** |
| ENSG00000164405.13 | **UQCRQ** | 27089 | protein_coding | **1742.085** | -0.531 | -0.160 | **0.692** | 1.47E-05 | **5.79E-05** |
| ENSG00000114354.17 | **TFG** | 10342 | protein_coding | **2373.610** | 0.373 | 0.112 | **1.295** | 1.47E-05 | **5.81E-05** |
| ENSG00000023892.14 | **DEF6** | 50619 | protein_coding | **1747.740** | -0.635 | -0.191 | **0.644** | 1.47E-05 | **5.81E-05** |
| ENSG00000116213.18 | **WRAP73** | 49856 | protein_coding | **915.158** | 0.441 | 0.133 | **1.357** | 1.48E-05 | **5.85E-05** |
| ENSG00000146409.14 | **SLC18B1** | 116843 | protein_coding | **192.581** | 0.754 | 0.227 | **1.686** | 1.49E-05 | **5.87E-05** |
| ENSG00000189431.8 | **RASSF10** | 644943 | protein_coding | **8.113** | 2.707 | 0.815 | **6.528** | 1.50E-05 | **5.91E-05** |
| ENSG00000163162.11 | **RNF149** | 284996 | protein_coding | **1080.534** | 0.504 | 0.152 | **1.418** | 1.50E-05 | **5.91E-05** |

| | | | | | | | | | |
|---|---|---|---|---|---|---|---|---|---|
| ENSG00000196091.17 | **MYBPC1** | 4604 | protein_coding | **4.736** | 3.915 | 1.178 | **15.080** | 1.50E-05 | **5.92E-05** |
| ENSG00000140157.18 | **NIPA2** | 81614 | protein_coding | **1046.349** | -0.465 | -0.140 | **0.725** | 1.50E-05 | **5.92E-05** |
| ENSG00000109062.14 | **NHERF1** | 9368 | protein_coding | **635.723** | 0.436 | 0.131 | **1.353** | 1.50E-05 | **5.92E-05** |
| ENSG00000065485.22 | **PDIA5** | 10954 | protein_coding | **326.013** | -0.632 | -0.190 | **0.645** | 1.51E-05 | **5.93E-05** |
| ENSG00000185650.12 | **ZFP36L1** | 677 | protein_coding | **4922.637** | 0.518 | 0.156 | **1.432** | 1.51E-05 | **5.93E-05** |
| ENSG00000204851.7 | **PNMA8B** | 57469 | protein_coding | **6.284** | 3.848 | 1.158 | **14.397** | 1.51E-05 | **5.94E-05** |
| ENSG00000134910.16 | **STT3A** | 3703 | protein_coding | **1597.248** | -0.505 | -0.152 | **0.705** | 1.51E-05 | **5.95E-05** |
| ENSG00000188677.16 | **PARVB** | 29780 | protein_coding | **2823.863** | -0.560 | -0.169 | **0.678** | 1.51E-05 | **5.96E-05** |
| ENSG00000129993.16 | **CBFA2T3** | 863 | protein_coding | **436.766** | -0.568 | -0.171 | **0.675** | 1.52E-05 | **5.97E-05** |
| ENSG00000107338.12 | **SHB** | 6461 | protein_coding | **81.989** | 1.491 | 0.449 | **2.811** | 1.52E-05 | **5.98E-05** |
| ENSG00000198208.14 | **RPS6KL1** | 83694 | protein_coding | **78.485** | 1.058 | 0.319 | **2.083** | 1.52E-05 | **5.99E-05** |
| ENSG00000168488.21 | **ATXN2L** | 11273 | protein_coding | **4545.905** | -0.668 | -0.201 | **0.629** | 1.53E-05 | **6.00E-05** |
| ENSG00000262874.3 | **C19orf84** | 147646 | protein_coding | **15.637** | 1.747 | 0.526 | **3.357** | 1.53E-05 | **6.02E-05** |
| ENSG00000214140.12 | **PRCD** | 768206 | protein_coding | **19.529** | 1.677 | 0.505 | **3.197** | 1.53E-05 | **6.03E-05** |
| ENSG00000171759.12 | **PAH** | 5053 | protein_coding | **16.007** | 2.339 | 0.704 | **5.059** | 1.54E-05 | **6.06E-05** |
| ENSG00000290760.2 | **UBE2Q2P1** | 10012810ٍ | lncRNA | **124.971** | 0.801 | 0.241 | **1.743** | 1.55E-05 | **6.08E-05** |
| ENSG00000273082.2 | **LARGE1-AS** | 10798556 | lncRNA | **8.088** | 2.721 | 0.819 | **6.595** | 1.55E-05 | **6.09E-05** |
| ENSG00000185973.14 | **TMLHE** | 55217 | protein_coding | **181.881** | -0.667 | -0.201 | **0.630** | 1.55E-05 | **6.09E-05** |
| ENSG00000162896.8 | **PIGR** | 5284 | protein_coding | **217.853** | 1.079 | 0.325 | **2.112** | 1.55E-05 | **6.09E-05** |
| ENSG00000149792.11 | **MRPL49** | 740 | protein_coding | **782.941** | -0.406 | -0.122 | **0.755** | 1.56E-05 | **6.10E-05** |
| ENSG00000309737.1 | **ENSG00000309737** | | lncRNA | **5.562** | 3.251 | 0.979 | **9.519** | 1.56E-05 | **6.11E-05** |
| ENSG00000143387.16 | **CTSK** | 1513 | protein_coding | **149.862** | 0.736 | 0.222 | **1.666** | 1.56E-05 | **6.13E-05** |
| ENSG00000162885.16 | **B3GALNT2** | 148789 | protein_coding | **305.650** | 0.516 | 0.155 | **1.430** | 1.56E-05 | **6.13E-05** |
| ENSG00000296453.1 | **ENSG00000296453** | | lncRNA | **14.355** | 2.077 | 0.625 | **4.220** | 1.57E-05 | **6.14E-05** |
| ENSG00000151376.21 | **ME3** | 10873 | protein_coding | **52.543** | 1.169 | 0.352 | **2.249** | 1.57E-05 | **6.14E-05** |
| ENSG00000182944.20 | **EWSR1** | 2130 | protein_coding | **6619.017** | -0.385 | -0.116 | **0.766** | 1.58E-05 | **6.21E-05** |
| ENSG00000290830.1 | **ENSG00000290830** | | lncRNA | **166.562** | 0.988 | 0.297 | **1.983** | 1.59E-05 | **6.22E-05** |
| ENSG00000153130.20 | **SCOC** | 60592 | protein_coding | **259.470** | -0.451 | -0.136 | **0.732** | 1.60E-05 | **6.25E-05** |
| ENSG00000105991.11 | **HOXA1** | 3198 | protein_coding | **7.680** | 3.078 | 0.927 | **8.444** | 1.60E-05 | **6.26E-05** |
| ENSG00000169213.9 | **RAB3B** | 5865 | protein_coding | **49.338** | 1.387 | 0.418 | **2.616** | 1.60E-05 | **6.27E-05** |
| ENSG00000241404.9 | **EGFL8** | 80864 | protein_coding | **102.935** | 0.874 | 0.263 | **1.833** | 1.61E-05 | **6.30E-05** |

| | | | | | | | | | |
|---|---|---|---|---|---|---|---|---|---|
| ENSG00000184588.19 | **PDE4B** | 5142 | protein_coding | **155.855** | -0.832 | -0.250 | **0.562** | 1.61E-05 | **6.31E-05** |
| ENSG00000139116.21 | **KIF21A** | 55605 | protein_coding | **188.257** | -0.669 | -0.201 | **0.629** | 1.61E-05 | **6.32E-05** |
| ENSG00000233203.8 | **DHCR24-DT** | | lncRNA | **22.260** | -1.849 | -0.557 | **0.278** | 1.61E-05 | **6.32E-05** |
| ENSG00000224078.16 | **SNHG14** | | lncRNA | **507.940** | -0.655 | -0.197 | **0.635** | 1.62E-05 | **6.32E-05** |
| ENSG00000234899.12 | **SOX9-AS1** | 400618 | lncRNA | **17.450** | 1.971 | 0.593 | **3.922** | 1.62E-05 | **6.34E-05** |
| ENSG00000105877.20 | **DNAH11** | 8701 | protein_coding | **20.598** | -2.423 | -0.729 | **0.186** | 1.62E-05 | **6.34E-05** |
| ENSG00000204131.12 | **NHSL2** | 340527 | protein_coding | **19.440** | 1.656 | 0.499 | **3.152** | 1.62E-05 | **6.34E-05** |
| ENSG00000095752.7 | **IL11** | 3589 | protein_coding | **11.209** | 2.158 | 0.650 | **4.463** | 1.62E-05 | **6.34E-05** |
| ENSG00000243725.9 | **TTC4** | 7268 | protein_coding | **412.070** | -0.421 | -0.127 | **0.747** | 1.63E-05 | **6.38E-05** |
| ENSG00000176102.15 | **CSTF3** | 1479 | protein_coding | **859.724** | 0.509 | 0.153 | **1.423** | 1.64E-05 | **6.39E-05** |
| ENSG00000189325.9 | **BNIP5** | 389384 | protein_coding | **7.881** | 2.828 | 0.851 | **7.102** | 1.64E-05 | **6.39E-05** |
| ENSG00000077684.18 | **JADE1** | 79960 | protein_coding | **522.436** | -0.417 | -0.126 | **0.749** | 1.64E-05 | **6.40E-05** |
| ENSG00000289727.1 | **ENSG00000289727** | | lncRNA | **72.213** | 0.904 | 0.272 | **1.872** | 1.64E-05 | **6.40E-05** |
| ENSG00000007255.12 | **TRAPPC6A** | 79090 | protein_coding | **1224.143** | 0.627 | 0.189 | **1.544** | 1.64E-05 | **6.40E-05** |
| ENSG00000157216.19 | **SSBP3** | 23648 | protein_coding | **1216.382** | 0.591 | 0.178 | **1.506** | 1.64E-05 | **6.40E-05** |
| ENSG00000262768.3 | **ENSG00000262768** | | lncRNA | **5.351** | 3.450 | 1.039 | **10.931** | 1.64E-05 | **6.40E-05** |
| ENSG00000205808.7 | **PLPP6** | 403313 | protein_coding | **145.965** | 0.695 | 0.209 | **1.619** | 1.64E-05 | **6.41E-05** |
| ENSG00000174837.16 | **ADGRE1** | 2015 | protein_coding | **150.592** | 1.017 | 0.306 | **2.024** | 1.65E-05 | **6.41E-05** |
| ENSG00000291051.2 | **ENSG00000291051** | | lncRNA | **16.756** | 1.828 | 0.550 | **3.550** | 1.65E-05 | **6.43E-05** |
| ENSG00000181218.6 | **H2AC25** | 92815 | protein_coding | **145.481** | 0.760 | 0.229 | **1.694** | 1.65E-05 | **6.43E-05** |
| ENSG00000186326.4 | **RGS9BP** | 388531 | protein_coding | **7.473** | 2.782 | 0.838 | **6.880** | 1.65E-05 | **6.44E-05** |
| ENSG00000267248.6 | **ENSG0000(** | 10099666( | lncRNA | **42.853** | 1.243 | 0.374 | **2.367** | 1.66E-05 | **6.46E-05** |
| ENSG00000224174.3 | **ENSG00000224174** | | lncRNA | **42.036** | 1.147 | 0.345 | **2.215** | 1.66E-05 | **6.46E-05** |
| ENSG00000115241.13 | **PPM1G** | 5496 | protein_coding | **2849.233** | -0.416 | -0.125 | **0.750** | 1.66E-05 | **6.47E-05** |
| ENSG00000235024.3 | **CDK5R2-AS** | 101928537 | lncRNA | **9.398** | 2.123 | 0.639 | **4.357** | 1.66E-05 | **6.47E-05** |
| ENSG00000228862.6 | **ENSG00000228862** | | lncRNA | **7.214** | 3.546 | 1.067 | **11.681** | 1.67E-05 | **6.49E-05** |
| ENSG00000100154.16 | **TTC28** | 23331 | protein_coding | **40.257** | 1.810 | 0.545 | **3.506** | 1.67E-05 | **6.52E-05** |
| ENSG00000289132.2 | **ENSG0000(** | 12490520z | lncRNA | **40.121** | 1.054 | 0.317 | **2.077** | 1.68E-05 | **6.54E-05** |
| ENSG00000088543.17 | **C3orf18** | 51161 | protein_coding | **79.267** | 0.973 | 0.293 | **1.963** | 1.69E-05 | **6.56E-05** |
| ENSG00000164338.12 | **UTP15** | 84135 | protein_coding | **201.282** | -0.620 | -0.187 | **0.651** | 1.69E-05 | **6.58E-05** |
| ENSG00000100815.14 | **TRIP11** | 9321 | protein_coding | **374.141** | -0.492 | -0.148 | **0.711** | 1.70E-05 | **6.61E-05** |

| | | | | | | | | | |
|---|---|---|---|---|---|---|---|---|---|
| ENSG00000188672.20 | **RHCE** | 6006 | protein_coding | **15.637** | 1.825 | 0.549 | **3.543** | 1.70E-05 | **6.61E-05** |
| ENSG00000144895.14 | **EIF2A** | 83939 | protein_coding | **2330.635** | -0.451 | -0.136 | **0.732** | 1.70E-05 | **6.62E-05** |
| ENSG00000013573.19 | **DDX11** | 1663 | protein_coding | **1505.797** | -0.434 | -0.131 | **0.740** | 1.71E-05 | **6.66E-05** |
| ENSG00000119608.16 | **PROX2** | 283571 | protein_coding | **23.350** | 1.455 | 0.438 | **2.742** | 1.72E-05 | **6.67E-05** |
| ENSG00000179029.17 | **TMEM107** | 84314 | protein_coding | **246.985** | -0.674 | -0.203 | **0.627** | 1.72E-05 | **6.67E-05** |
| ENSG00000169714.19 | **CNBP** | 7555 | protein_coding | **4388.638** | -0.324 | -0.098 | **0.799** | 1.72E-05 | **6.67E-05** |
| ENSG00000302010.1 | **ENSG0000(** | 10798507 | lncRNA | **52.633** | 1.059 | 0.319 | **2.084** | 1.72E-05 | **6.70E-05** |
| ENSG00000154721.17 | **JAM2** | 58494 | protein_coding | **83.197** | -1.016 | -0.306 | **0.495** | 1.73E-05 | **6.70E-05** |
| ENSG00000270755.1 | **ENSG00000270755** | | lncRNA | **6.483** | 3.357 | 1.011 | **10.249** | 1.73E-05 | **6.70E-05** |
| ENSG00000059573.11 | **ALDH18A1** | 5832 | protein_coding | **1205.424** | -0.454 | -0.137 | **0.730** | 1.73E-05 | **6.70E-05** |
| ENSG00000179750.17 | **APOBEC3B** | 9582 | protein_coding | **597.016** | -0.804 | -0.242 | **0.573** | 1.74E-05 | **6.75E-05** |
| ENSG00000135547.10 | **HEY2** | 23493 | protein_coding | **10.987** | 2.733 | 0.823 | **6.647** | 1.75E-05 | **6.78E-05** |
| ENSG00000284707.3 | **ENSG00000284707** | | lncRNA | **1557.365** | -0.752 | -0.226 | **0.594** | 1.75E-05 | **6.78E-05** |
| ENSG00000173451.8 | **THAP2** | 83591 | protein_coding | **69.232** | -0.975 | -0.294 | **0.509** | 1.75E-05 | **6.78E-05** |
| ENSG00000293555.2 | **FAM169BP** | 283777 | lncRNA | **20.447** | 1.981 | 0.596 | **3.949** | 1.76E-05 | **6.82E-05** |
| ENSG00000159348.15 | **CYB5R1** | 51706 | protein_coding | **345.677** | 0.531 | 0.160 | **1.445** | 1.76E-05 | **6.83E-05** |
| ENSG00000179348.15 | **GATA2** | 2624 | protein_coding | **6.769** | 3.217 | 0.968 | **9.299** | 1.77E-05 | **6.87E-05** |
| ENSG00000167536.16 | **DHRS13** | 147015 | protein_coding | **222.592** | -0.677 | -0.204 | **0.626** | 1.78E-05 | **6.90E-05** |
| ENSG00000088451.13 | **TGDS** | 23483 | protein_coding | **200.172** | -0.658 | -0.198 | **0.634** | 1.78E-05 | **6.90E-05** |
| ENSG00000108799.15 | **EZH1** | 2145 | protein_coding | **1162.954** | 0.614 | 0.185 | **1.531** | 1.78E-05 | **6.90E-05** |
| ENSG00000286118.2 | **ENSG00000286118** | | lncRNA | **61.076** | -0.934 | -0.281 | **0.523** | 1.79E-05 | **6.92E-05** |
| ENSG00000166037.13 | **CEP57** | 9702 | protein_coding | **825.963** | -0.658 | -0.198 | **0.634** | 1.79E-05 | **6.93E-05** |
| ENSG00000257337.10 | **TNS2-AS1** | 283335 | lncRNA | **35.693** | 1.095 | 0.330 | **2.137** | 1.80E-05 | **6.95E-05** |
| ENSG00000205560.16 | **CPT1B** | 1375 | protein_coding | **2475.728** | 0.435 | 0.131 | **1.352** | 1.80E-05 | **6.95E-05** |
| ENSG00000018280.20 | **SLC11A1** | 6556 | protein_coding | **16.242** | 1.608 | 0.484 | **3.047** | 1.80E-05 | **6.96E-05** |
| ENSG00000110375.4 | **UPK2** | 7379 | protein_coding | **16.895** | -2.338 | -0.704 | **0.198** | 1.80E-05 | **6.97E-05** |
| ENSG00000079332.18 | **SAR1A** | 56681 | protein_coding | **1021.774** | -0.527 | -0.159 | **0.694** | 1.80E-05 | **6.97E-05** |
| ENSG00000159445.15 | **THEM4** | 117145 | protein_coding | **360.329** | 0.438 | 0.132 | **1.355** | 1.81E-05 | **6.99E-05** |
| ENSG00000153563.18 | **CD8A** | 925 | protein_coding | **6.448** | 3.320 | 0.999 | **9.988** | 1.81E-05 | **6.99E-05** |
| ENSG00000138735.18 | **PDE5A** | 8654 | protein_coding | **5.768** | 4.014 | 1.208 | **16.155** | 1.81E-05 | **6.99E-05** |
| ENSG00000185028.5 | **LRRC14B** | 389257 | protein_coding | **15.038** | -1.831 | -0.551 | **0.281** | 1.81E-05 | **7.00E-05** |

| | | | | | | | | | |
|---|---|---|---|---|---|---|---|---|---|
| ENSG00000171766.19 | **GATM** | 2628 | protein_coding | **537.880** | -0.703 | -0.212 | **0.614** | 1.81E-05 | **7.00E-05** |
| ENSG00000169592.17 | **INO80E** | 283899 | protein_coding | **1553.774** | -0.535 | -0.161 | **0.690** | 1.82E-05 | **7.05E-05** |
| ENSG00000286940.2 | **MICB-DT** | 10272506 | lncRNA | **26.414** | -1.377 | -0.415 | **0.385** | 1.83E-05 | **7.05E-05** |
| ENSG00000288538.1 | **ENSG00000288538** | | lncRNA | **36.020** | 1.197 | 0.360 | **2.292** | 1.84E-05 | **7.09E-05** |
| ENSG00000175699.17 | **CCDC197** | 256369 | protein_coding | **6.943** | 2.868 | 0.863 | **7.302** | 1.85E-05 | **7.12E-05** |
| ENSG00000138081.24 | **FBXO11** | 80204 | protein_coding | **624.451** | -0.496 | -0.149 | **0.709** | 1.85E-05 | **7.13E-05** |
| ENSG00000293281.1 | **ENSG00000293281** | | lncRNA | **5.501** | 3.434 | 1.034 | **10.805** | 1.85E-05 | **7.15E-05** |
| ENSG00000143320.11 | **CRABP2** | 1382 | protein_coding | **43.818** | 1.177 | 0.354 | **2.261** | 1.87E-05 | **7.20E-05** |
| ENSG00000310190.1 | **ENSG00000310190** | | lncRNA | **35.365** | -1.296 | -0.390 | **0.407** | 1.87E-05 | **7.20E-05** |
| ENSG00000128805.17 | **ARHGAP22** | 58504 | protein_coding | **69.009** | 1.264 | 0.381 | **2.402** | 1.87E-05 | **7.21E-05** |
| ENSG00000163344.8 | **PMVK** | 10654 | protein_coding | **689.474** | -0.492 | -0.148 | **0.711** | 1.87E-05 | **7.23E-05** |
| ENSG00000143314.14 | **MRPL24** | 79590 | protein_coding | **1193.098** | -0.598 | -0.180 | **0.661** | 1.88E-05 | **7.26E-05** |
| ENSG00000166226.15 | **CCT2** | 10576 | protein_coding | **2155.595** | -0.370 | -0.111 | **0.774** | 1.89E-05 | **7.27E-05** |
| ENSG00000164924.20 | **YWHAZ** | 7534 | protein_coding | **5081.456** | -0.510 | -0.154 | **0.702** | 1.89E-05 | **7.30E-05** |
| ENSG00000142102.18 | **PGGHG** | 80162 | protein_coding | **5526.378** | 0.543 | 0.163 | **1.457** | 1.90E-05 | **7.31E-05** |
| ENSG00000151012.14 | **SLC7A11** | 23657 | protein_coding | **284.110** | -1.086 | -0.327 | **0.471** | 1.90E-05 | **7.32E-05** |
| ENSG00000267834.1 | **ENSG00000267834** | | lncRNA | **9.684** | 2.348 | 0.707 | **5.091** | 1.90E-05 | **7.32E-05** |
| ENSG00000100027.18 | **YPEL1** | 29799 | protein_coding | **88.686** | 1.012 | 0.305 | **2.017** | 1.91E-05 | **7.36E-05** |
| ENSG00000170379.22 | **TCAF2** | 285966 | protein_coding | **364.451** | -0.801 | -0.241 | **0.574** | 1.91E-05 | **7.37E-05** |
| ENSG00000291124.2 | **ENSG00000291124** | | lncRNA | **353.749** | -0.560 | -0.168 | **0.678** | 1.91E-05 | **7.37E-05** |
| ENSG00000260018.3 | **MBTPS1-DT** | | lncRNA | **28.549** | 1.129 | 0.340 | **2.187** | 1.91E-05 | **7.37E-05** |
| ENSG00000184221.13 | **OLIG1** | 116448 | protein_coding | **21.199** | 1.732 | 0.522 | **3.323** | 1.93E-05 | **7.41E-05** |
| ENSG00000167315.20 | **ACAA2** | 10449 | protein_coding | **808.981** | -0.547 | -0.165 | **0.684** | 1.93E-05 | **7.42E-05** |
| ENSG00000113916.19 | **BCL6** | 604 | protein_coding | **13.943** | 2.159 | 0.650 | **4.465** | 1.93E-05 | **7.42E-05** |
| ENSG00000266402.6 | **SNHG25** | | lncRNA | **206.050** | -0.790 | -0.238 | **0.578** | 1.93E-05 | **7.43E-05** |
| ENSG00000160223.20 | **ICOSLG** | 23308 | protein_coding | **1000.578** | -0.548 | -0.165 | **0.684** | 1.93E-05 | **7.44E-05** |
| ENSG00000189223.16 | **PAX8-AS1** | 654433 | lncRNA | **631.400** | -0.582 | -0.175 | **0.668** | 1.94E-05 | **7.44E-05** |
| ENSG00000162591.19 | **MEGF6** | 1953 | protein_coding | **54.764** | 1.716 | 0.517 | **3.286** | 1.94E-05 | **7.45E-05** |
| ENSG00000197608.14 | **ZNF841** | 284371 | protein_coding | **380.887** | -0.644 | -0.194 | **0.640** | 1.94E-05 | **7.45E-05** |
| ENSG00000110628.18 | **SLC67A1** | 5002 | protein_coding | **294.928** | 0.699 | 0.210 | **1.623** | 1.94E-05 | **7.46E-05** |
| ENSG00000101146.15 | **RAE1** | 8480 | protein_coding | **606.392** | -0.487 | -0.147 | **0.714** | 1.94E-05 | **7.46E-05** |

| | | | | | | | | | |
|---|---|---|---|---|---|---|---|---|---|
| ENSG00000155330.11 | **C16orf87** | 388272 | protein_coding | **229.536** | -0.845 | -0.254 | **0.557** | 1.94E-05 | **7.46E-05** |
| ENSG00000172986.14 | **GXYLT2** | 727936 | protein_coding | **4.422** | 4.722 | 1.421 | **26.392** | 1.94E-05 | **7.46E-05** |
| ENSG00000140650.15 | **PMM2** | 5373 | protein_coding | **914.101** | -0.580 | -0.175 | **0.669** | 1.95E-05 | **7.47E-05** |
| ENSG00000258735.2 | **PPP1R13B-** | 145216 | lncRNA | **92.508** | 0.716 | 0.216 | **1.643** | 1.95E-05 | **7.47E-05** |
| ENSG00000277196.4 | **ENSG00000277196** | | protein_coding | **56.324** | 1.258 | 0.379 | **2.392** | 1.95E-05 | **7.48E-05** |
| ENSG00000119397.21 | **CNTRL** | 11064 | protein_coding | **1216.262** | -0.541 | -0.163 | **0.687** | 1.95E-05 | **7.50E-05** |
| ENSG00000310576.2 | **FAM174C** | 55009 | protein_coding | **213.069** | -0.619 | -0.186 | **0.651** | 1.96E-05 | **7.53E-05** |
| ENSG00000092841.21 | **MYL6** | 4637 | protein_coding | **6162.692** | -0.636 | -0.191 | **0.644** | 1.97E-05 | **7.55E-05** |
| ENSG00000103544.19 | **VPS35L** | 57020 | protein_coding | **519.874** | -0.628 | -0.189 | **0.647** | 1.97E-05 | **7.55E-05** |
| ENSG00000287879.1 | **ENSG00000287879** | | lncRNA | **10.140** | 2.183 | 0.657 | **4.541** | 1.98E-05 | **7.58E-05** |
| ENSG00000135473.18 | **PAN2** | 9924 | protein_coding | **2216.215** | 0.574 | 0.173 | **1.489** | 1.98E-05 | **7.59E-05** |
| ENSG00000041357.18 | **PSMA4** | 5685 | protein_coding | **2095.296** | -0.451 | -0.136 | **0.731** | 1.98E-05 | **7.59E-05** |
| ENSG00000058063.19 | **ATP11B** | 23200 | protein_coding | **717.592** | -0.567 | -0.171 | **0.675** | 1.98E-05 | **7.60E-05** |
| ENSG00000175455.18 | **CCDC14** | 64770 | protein_coding | **1029.960** | -0.540 | -0.163 | **0.688** | 1.98E-05 | **7.60E-05** |
| ENSG00000128274.19 | **A4GALT** | 53947 | protein_coding | **531.066** | 0.676 | 0.204 | **1.598** | 1.99E-05 | **7.62E-05** |
| ENSG00000180098.11 | **TRNAU1AF** | 54952 | protein_coding | **643.685** | -0.528 | -0.159 | **0.694** | 2.00E-05 | **7.64E-05** |
| ENSG00000162735.21 | **PEX19** | 5824 | protein_coding | **556.977** | -0.593 | -0.179 | **0.663** | 2.00E-05 | **7.65E-05** |
| ENSG00000272008.2 | **ENSG0000(** | 12490135! | lncRNA | **26.221** | 1.825 | 0.549 | **3.543** | 2.00E-05 | **7.65E-05** |
| ENSG00000114573.14 | **ATP6V1A** | 523 | protein_coding | **650.088** | 0.800 | 0.241 | **1.741** | 2.01E-05 | **7.67E-05** |
| ENSG00000008056.16 | **SYN1** | 6853 | protein_coding | **37.877** | 1.486 | 0.447 | **2.800** | 2.01E-05 | **7.69E-05** |
| ENSG00000304140.1 | **ENSG00000304140** | | lncRNA | **37.953** | -0.997 | -0.300 | **0.501** | 2.01E-05 | **7.69E-05** |
| ENSG00000074071.17 | **MRPS34** | 65993 | protein_coding | **1318.609** | -0.393 | -0.118 | **0.761** | 2.01E-05 | **7.70E-05** |
| ENSG00000289412.3 | **ZEBTR** | | lncRNA | **17.651** | 1.861 | 0.560 | **3.633** | 2.02E-05 | **7.71E-05** |
| ENSG00000290692.2 | **ENSG00000290692** | | lncRNA | **3.783** | 4.459 | 1.342 | **21.996** | 2.02E-05 | **7.72E-05** |
| ENSG00000124406.19 | **ATP8A1** | 10396 | protein_coding | **912.726** | -0.421 | -0.127 | **0.747** | 2.02E-05 | **7.73E-05** |
| ENSG00000178397.14 | **FAM220A** | 84792 | protein_coding | **154.713** | 0.690 | 0.208 | **1.614** | 2.02E-05 | **7.73E-05** |
| ENSG00000102409.12 | **BEX4** | 56271 | protein_coding | **233.191** | 0.989 | 0.298 | **1.985** | 2.02E-05 | **7.73E-05** |
| ENSG00000118058.26 | **KMT2A** | 4297 | protein_coding | **1578.992** | 0.561 | 0.169 | **1.475** | 2.03E-05 | **7.74E-05** |
| ENSG00000151726.18 | **ACSL1** | 2180 | protein_coding | **1014.714** | 0.520 | 0.157 | **1.434** | 2.03E-05 | **7.75E-05** |
| ENSG00000050820.19 | **BCAR1** | 9564 | protein_coding | **431.177** | 0.575 | 0.173 | **1.490** | 2.03E-05 | **7.75E-05** |
| ENSG00000241258.10 | **CRCP** | 27297 | protein_coding | **403.358** | -0.455 | -0.137 | **0.730** | 2.03E-05 | **7.76E-05** |

| | | | | | | | | | |
|---|---|---|---|---|---|---|---|---|---|
| ENSG00000172339.12 | **ALG14** | 199857 | protein_coding | **110.290** | -0.773 | -0.233 | **0.585** | 2.04E-05 | **7.78E-05** |
| ENSG00000112182.18 | **BACH2** | 60468 | protein_coding | **19.831** | 2.224 | 0.670 | **4.673** | 2.04E-05 | **7.78E-05** |
| ENSG00000246985.9 | **SOCS2-AS1** | 144481 | lncRNA | **96.398** | 0.991 | 0.298 | **1.988** | 2.06E-05 | **7.85E-05** |
| ENSG00000260167.2 | **ENSG00000260167** | | lncRNA | **15.144** | -1.734 | -0.522 | **0.301** | 2.06E-05 | **7.86E-05** |
| ENSG00000067445.24 | **TRO** | 7216 | protein_coding | **244.591** | 0.980 | 0.295 | **1.972** | 2.06E-05 | **7.86E-05** |
| ENSG00000100031.21 | **GGT1** | 2678 | protein_coding | **587.288** | -0.841 | -0.253 | **0.558** | 2.07E-05 | **7.87E-05** |
| ENSG00000214753.6 | **HNRNPUL2** | 221092 | protein_coding | **2969.731** | -0.326 | -0.098 | **0.798** | 2.07E-05 | **7.88E-05** |
| ENSG00000196177.15 | **ACADSB** | 36 | protein_coding | **427.810** | 0.588 | 0.177 | **1.503** | 2.07E-05 | **7.89E-05** |
| ENSG00000067113.19 | **PLPP1** | 8611 | protein_coding | **132.836** | 0.748 | 0.225 | **1.679** | 2.08E-05 | **7.91E-05** |
| ENSG00000163125.18 | **RPRD2** | 23248 | protein_coding | **494.570** | -0.580 | -0.175 | **0.669** | 2.08E-05 | **7.91E-05** |
| ENSG00000161055.6 | **SCGB3A1** | 92304 | protein_coding | **46.424** | 1.992 | 0.600 | **3.977** | 2.08E-05 | **7.91E-05** |
| ENSG00000147799.14 | **ARHGAP39** | 80728 | protein_coding | **17.490** | 1.587 | 0.478 | **3.004** | 2.08E-05 | **7.91E-05** |
| ENSG00000180694.16 | **TMEM64** | 169200 | protein_coding | **106.074** | -0.851 | -0.256 | **0.554** | 2.08E-05 | **7.92E-05** |
| ENSG00000275342.8 | **PRAG1** | 157285 | protein_coding | **342.886** | -0.481 | -0.145 | **0.716** | 2.08E-05 | **7.92E-05** |
| ENSG00000165119.25 | **HNRNPK** | 3190 | protein_coding | **9379.918** | -0.404 | -0.122 | **0.756** | 2.09E-05 | **7.93E-05** |
| ENSG00000129353.17 | **SLC44A2** | 57153 | protein_coding | **2513.504** | -0.529 | -0.159 | **0.693** | 2.09E-05 | **7.94E-05** |
| ENSG00000152642.13 | **GPD1L** | 23171 | protein_coding | **95.916** | 0.731 | 0.220 | **1.660** | 2.09E-05 | **7.94E-05** |
| ENSG00000124659.7 | **TBCC** | 6903 | protein_coding | **441.211** | -0.494 | -0.149 | **0.710** | 2.09E-05 | **7.95E-05** |
| ENSG00000173517.13 | **PEAK1** | 79834 | protein_coding | **393.601** | 0.551 | 0.166 | **1.465** | 2.09E-05 | **7.95E-05** |
| ENSG00000138119.19 | **MYOF** | 26509 | protein_coding | **7.513** | 2.723 | 0.820 | **6.601** | 2.10E-05 | **7.97E-05** |
| ENSG00000150459.15 | **SAP18** | 10284 | protein_coding | **2247.459** | -0.365 | -0.110 | **0.777** | 2.10E-05 | **7.98E-05** |
| ENSG00000088766.14 | **CRLS1** | 54675 | protein_coding | **466.697** | -0.583 | -0.176 | **0.667** | 2.12E-05 | **8.04E-05** |
| ENSG00000164690.8 | **SHH** | 6469 | protein_coding | **4.830** | 4.315 | 1.299 | **19.911** | 2.12E-05 | **8.05E-05** |
| ENSG00000230615.9 | **ENSG00000230615** | | lncRNA | **17.247** | 1.931 | 0.581 | **3.814** | 2.12E-05 | **8.06E-05** |
| ENSG00000124588.24 | **NQO2** | 4835 | protein_coding | **157.430** | 0.653 | 0.197 | **1.573** | 2.13E-05 | **8.09E-05** |
| ENSG00000176945.20 | **MUC20** | 200958 | protein_coding | **109.078** | 1.053 | 0.317 | **2.074** | 2.13E-05 | **8.09E-05** |
| ENSG00000226779.1 | **NAALADL2-AS2** | | lncRNA | **40.061** | 1.398 | 0.421 | **2.636** | 2.13E-05 | **8.10E-05** |
| ENSG00000189180.18 | **ZNF33A** | 7581 | protein_coding | **440.704** | -0.708 | -0.213 | **0.612** | 2.14E-05 | **8.10E-05** |
| ENSG00000167395.12 | **ZNF646** | 9726 | protein_coding | **479.285** | -0.471 | -0.142 | **0.721** | 2.14E-05 | **8.13E-05** |
| ENSG00000013016.18 | **EHD3** | 30845 | protein_coding | **26.724** | 2.115 | 0.637 | **4.333** | 2.14E-05 | **8.13E-05** |
| ENSG00000257327.2 | **ENSG00000257327** | | lncRNA | **5.965** | 3.471 | 1.045 | **11.090** | 2.15E-05 | **8.16E-05** |

| | | | | | | | | | |
|---|---|---|---|---|---|---|---|---|---|
| ENSG00000151532.17 | **VTI1A** | 143187 | protein_coding | **357.431** | -0.581 | -0.175 | **0.669** | 2.16E-05 | **8.20E-05** |
| ENSG00000102349.20 | **KLF8** | 11279 | protein_coding | **253.829** | -0.778 | -0.234 | **0.583** | 2.16E-05 | **8.20E-05** |
| ENSG00000172062.19 | **SMN1** | 6606 | protein_coding | **692.653** | -0.443 | -0.133 | **0.736** | 2.18E-05 | **8.26E-05** |
| ENSG00000224023.11 | **EDRF1-DT** | 399821 | lncRNA | **45.968** | 0.985 | 0.297 | **1.980** | 2.18E-05 | **8.26E-05** |
| ENSG00000139352.5 | **ASCL1** | 429 | protein_coding | **157.165** | 0.862 | 0.259 | **1.818** | 2.18E-05 | **8.27E-05** |
| ENSG00000135722.11 | **FBXL8** | 55336 | protein_coding | **232.096** | 0.698 | 0.210 | **1.622** | 2.19E-05 | **8.28E-05** |
| ENSG00000105723.13 | **GSK3A** | 2931 | protein_coding | **938.442** | -0.621 | -0.187 | **0.650** | 2.19E-05 | **8.28E-05** |
| ENSG00000166340.20 | **TPP1** | 1200 | protein_coding | **1880.270** | 0.544 | 0.164 | **1.458** | 2.19E-05 | **8.29E-05** |
| ENSG00000141505.14 | **ASGR1** | 432 | protein_coding | **11.680** | 1.992 | 0.600 | **3.978** | 2.19E-05 | **8.30E-05** |
| ENSG00000008226.22 | **DLEC1** | 9940 | protein_coding | **9.612** | 2.176 | 0.655 | **4.518** | 2.19E-05 | **8.30E-05** |
| ENSG00000232093.2 | **DCST1-AS1** | | lncRNA | **18.598** | 1.456 | 0.438 | **2.744** | 2.19E-05 | **8.30E-05** |
| ENSG00000172531.18 | **PPP1CA** | 5499 | protein_coding | **3260.844** | -0.414 | -0.125 | **0.750** | 2.20E-05 | **8.31E-05** |
| ENSG00000004897.14 | **CDC27** | 996 | protein_coding | **657.675** | -0.558 | -0.168 | **0.679** | 2.20E-05 | **8.32E-05** |
| ENSG00000196387.12 | **ZNF140** | 7699 | protein_coding | **289.605** | -0.548 | -0.165 | **0.684** | 2.22E-05 | **8.39E-05** |
| ENSG00000169057.27 | **MECP2** | 4204 | protein_coding | **774.751** | -0.464 | -0.140 | **0.725** | 2.23E-05 | **8.42E-05** |
| ENSG00000246090.10 | **ENSG0000(** | 10050705: | lncRNA | **44.235** | 1.150 | 0.346 | **2.220** | 2.24E-05 | **8.45E-05** |
| ENSG00000198168.10 | **SVIP** | 258010 | protein_coding | **421.834** | -0.654 | -0.197 | **0.635** | 2.24E-05 | **8.45E-05** |
| ENSG00000079387.17 | **SENP1** | 29843 | protein_coding | **378.106** | -0.570 | -0.171 | **0.674** | 2.24E-05 | **8.45E-05** |
| ENSG00000273117.2 | **INSIG1-DT** | | lncRNA | **159.693** | 0.906 | 0.273 | **1.874** | 2.24E-05 | **8.46E-05** |
| ENSG00000110921.16 | **MVK** | 4598 | protein_coding | **761.652** | -0.845 | -0.254 | **0.557** | 2.25E-05 | **8.49E-05** |
| ENSG00000131771.16 | **PPP1R1B** | 84152 | protein_coding | **6.503** | 2.787 | 0.839 | **6.904** | 2.25E-05 | **8.49E-05** |
| ENSG00000286656.3 | **ENSG00000286656** | | lncRNA | **33.984** | 1.223 | 0.368 | **2.335** | 2.25E-05 | **8.50E-05** |
| ENSG00000130640.17 | **TUBGCP2** | 10844 | protein_coding | **1709.983** | 0.502 | 0.151 | **1.416** | 2.26E-05 | **8.52E-05** |
| ENSG00000119392.18 | **GLE1** | 2733 | protein_coding | **989.593** | -0.388 | -0.117 | **0.764** | 2.26E-05 | **8.55E-05** |
| ENSG00000133812.20 | **SBF2** | 81846 | protein_coding | **205.118** | 0.785 | 0.236 | **1.723** | 2.27E-05 | **8.58E-05** |
| ENSG00000228408.6 | **ENSG00000228408** | | lncRNA | **7.545** | 4.146 | 1.248 | **17.703** | 2.28E-05 | **8.59E-05** |
| ENSG00000143977.16 | **SNRPG** | 6637 | protein_coding | **1860.006** | -0.615 | -0.185 | **0.653** | 2.28E-05 | **8.59E-05** |
| ENSG00000164989.19 | **CCDC171** | 203238 | protein_coding | **80.996** | 0.824 | 0.248 | **1.770** | 2.28E-05 | **8.59E-05** |
| ENSG00000118160.16 | **SLC8A2** | 6543 | protein_coding | **7.280** | 3.070 | 0.924 | **8.398** | 2.28E-05 | **8.59E-05** |
| ENSG00000155959.14 | **VBP1** | 7411 | protein_coding | **388.800** | -0.690 | -0.208 | **0.620** | 2.28E-05 | **8.59E-05** |
| ENSG00000104915.17 | **STX10** | 8677 | protein_coding | **760.956** | 0.390 | 0.118 | **1.311** | 2.29E-05 | **8.62E-05** |

| | | | | | | | | | |
|---|---|---|---|---|---|---|---|---|---|
| ENSG00000172927.11 | **MYEOV** | 26579 | protein_coding | **20.657** | 2.286 | 0.688 | **4.877** | 2.29E-05 | **8.65E-05** |
| ENSG00000272654.3 | **JTB-DT** | 12370654( | lncRNA | **39.153** | 1.058 | 0.318 | **2.081** | 2.30E-05 | **8.65E-05** |
| ENSG00000130511.18 | **SSBP4** | 170463 | protein_coding | **1155.426** | 0.527 | 0.159 | **1.441** | 2.30E-05 | **8.67E-05** |
| ENSG00000018510.20 | **AGPS** | 8540 | protein_coding | **475.544** | -0.677 | -0.204 | **0.625** | 2.31E-05 | **8.68E-05** |
| ENSG00000188739.17 | **RBM34** | 23029 | protein_coding | **595.782** | -0.493 | -0.148 | **0.711** | 2.31E-05 | **8.70E-05** |
| ENSG00000228221.7 | **LINC00578** | 10050556( | lncRNA | **3.657** | 4.618 | 1.390 | **24.560** | 2.31E-05 | **8.71E-05** |
| ENSG00000132155.16 | **RAF1** | 5894 | protein_coding | **2126.961** | -0.467 | -0.141 | **0.723** | 2.32E-05 | **8.75E-05** |
| ENSG00000310523.1 | **ENSG00000310523** | | lncRNA | **589.618** | 0.628 | 0.189 | **1.546** | 2.33E-05 | **8.76E-05** |
| ENSG00000151746.17 | **BICD1** | 636 | protein_coding | **217.414** | -0.537 | -0.162 | **0.689** | 2.33E-05 | **8.78E-05** |
| ENSG00000151332.22 | **MBIP** | 51562 | protein_coding | **245.075** | -0.580 | -0.175 | **0.669** | 2.35E-05 | **8.85E-05** |
| ENSG00000182389.21 | **CACNB4** | 785 | protein_coding | **154.577** | -1.004 | -0.302 | **0.499** | 2.35E-05 | **8.85E-05** |
| ENSG00000119929.15 | **CUTC** | 51076 | protein_coding | **394.902** | -0.417 | -0.125 | **0.749** | 2.36E-05 | **8.86E-05** |
| ENSG00000140854.15 | **KATNB1** | 10300 | protein_coding | **391.333** | -0.410 | -0.124 | **0.752** | 2.36E-05 | **8.87E-05** |
| ENSG00000214407.4 | **RDUR** | 152225 | lncRNA | **15.863** | 2.965 | 0.892 | **7.806** | 2.36E-05 | **8.87E-05** |
| ENSG00000297784.1 | **ENSG00000297784** | | lncRNA | **7.604** | 2.328 | 0.701 | **5.020** | 2.36E-05 | **8.87E-05** |
| ENSG00000123064.15 | **DDX54** | 79039 | protein_coding | **1262.990** | -0.526 | -0.158 | **0.694** | 2.37E-05 | **8.90E-05** |
| ENSG00000234663.8 | **LINC01934** | | lncRNA | **17.866** | 1.708 | 0.514 | **3.266** | 2.37E-05 | **8.90E-05** |
| ENSG00000224281.7 | **SLC25A5-A** | 10030372{ | lncRNA | **34.067** | 1.126 | 0.339 | **2.182** | 2.39E-05 | **8.96E-05** |
| ENSG00000105426.22 | **PTPRS** | 5802 | protein_coding | **525.326** | 0.511 | 0.154 | **1.425** | 2.39E-05 | **8.98E-05** |
| ENSG00000142684.10 | **ZNF593** | 51042 | protein_coding | **395.666** | -0.561 | -0.169 | **0.678** | 2.39E-05 | **8.98E-05** |
| ENSG00000042980.16 | **ADAM28** | 10863 | protein_coding | **11.772** | 3.102 | 0.934 | **8.588** | 2.40E-05 | **8.99E-05** |
| ENSG00000142252.13 | **GEMIN7** | 79760 | protein_coding | **884.598** | -0.495 | -0.149 | **0.710** | 2.40E-05 | **8.99E-05** |
| ENSG00000293106.2 | **RPL23AP7** | 118433 | lncRNA | **213.688** | -0.584 | -0.176 | **0.667** | 2.40E-05 | **9.00E-05** |
| ENSG00000113196.5 | **HAND1** | 9421 | protein_coding | **6.641** | 3.478 | 1.047 | **11.144** | 2.41E-05 | **9.02E-05** |
| ENSG00000173599.17 | **PC** | 5091 | protein_coding | **57.879** | 1.057 | 0.318 | **2.080** | 2.41E-05 | **9.02E-05** |
| ENSG00000198933.13 | **TBKBP1** | 9755 | protein_coding | **455.138** | 0.709 | 0.213 | **1.634** | 2.41E-05 | **9.05E-05** |
| ENSG00000205078.7 | **SYCE1L** | 10013095{ | protein_coding | **47.534** | 1.202 | 0.362 | **2.300** | 2.41E-05 | **9.05E-05** |
| ENSG00000261606.5 | **ENSG00000261606** | | lncRNA | **6.324** | 3.747 | 1.128 | **13.428** | 2.42E-05 | **9.08E-05** |
| ENSG00000241135.9 | **LINC00881** | | lncRNA | **120.181** | -0.824 | -0.248 | **0.565** | 2.42E-05 | **9.08E-05** |
| ENSG00000137941.19 | **TTLL7** | 79739 | protein_coding | **54.265** | 1.848 | 0.556 | **3.600** | 2.43E-05 | **9.12E-05** |
| ENSG00000140564.15 | **FURIN** | 5045 | protein_coding | **754.593** | 0.564 | 0.170 | **1.479** | 2.43E-05 | **9.12E-05** |

| | | | | | | | | | |
|---|---|---|---|---|---|---|---|---|---|
| ENSG00000286416.2 | **ENSG00000286416** | | lncRNA | **22.283** | 1.412 | 0.425 | **2.660** | 2.44E-05 | **9.12E-05** |
| ENSG00000105711.16 | **SCN1B** | 6324 | protein_coding | **21.171** | 1.368 | 0.412 | **2.581** | 2.44E-05 | **9.13E-05** |
| ENSG00000177853.17 | **ZNF518A** | 9849 | protein_coding | **531.917** | 0.516 | 0.155 | **1.430** | 2.44E-05 | **9.14E-05** |
| ENSG00000125352.6 | **RNF113A** | 7737 | protein_coding | **673.110** | -0.470 | -0.141 | **0.722** | 2.45E-05 | **9.17E-05** |
| ENSG00000077264.19 | **PAK3** | 5063 | protein_coding | **7.859** | 3.309 | 0.996 | **9.909** | 2.45E-05 | **9.17E-05** |
| ENSG00000310458.1 | **ENSG00000310458** | | lncRNA | **8.058** | 2.923 | 0.880 | **7.586** | 2.45E-05 | **9.17E-05** |
| ENSG00000188895.14 | **MSL1** | 339287 | protein_coding | **1759.381** | 0.646 | 0.195 | **1.565** | 2.46E-05 | **9.22E-05** |
| ENSG00000100764.16 | **PSMC1** | 5700 | protein_coding | **1573.579** | -0.392 | -0.118 | **0.762** | 2.46E-05 | **9.22E-05** |
| ENSG00000228172.6 | **ENSG00000228172** | | lncRNA | **12.449** | 1.767 | 0.532 | **3.404** | 2.47E-05 | **9.23E-05** |
| ENSG00000214425.11 | **LRRC37A4P** | | transcribed_unprc | **35.639** | -1.777 | -0.535 | **0.292** | 2.47E-05 | **9.24E-05** |
| ENSG00000167085.16 | **PHB1** | 5245 | protein_coding | **2251.976** | -0.624 | -0.188 | **0.649** | 2.48E-05 | **9.27E-05** |
| ENSG00000073008.18 | **PVR** | 5817 | protein_coding | **49.445** | 1.239 | 0.373 | **2.360** | 2.48E-05 | **9.27E-05** |
| ENSG00000079785.18 | **DDX1** | 1653 | protein_coding | **1008.407** | -0.463 | -0.139 | **0.725** | 2.49E-05 | **9.30E-05** |
| ENSG00000303643.1 | **ENSG00000303643** | | lncRNA | **12.620** | 1.821 | 0.548 | **3.534** | 2.49E-05 | **9.31E-05** |
| ENSG00000282988.2 | **ENSG00000282988** | | protein_coding | **24.025** | -2.618 | -0.788 | **0.163** | 2.50E-05 | **9.35E-05** |
| ENSG00000240449.2 | **REPIN1-AS** | 10537556: | lncRNA | **29.037** | 1.280 | 0.385 | **2.429** | 2.51E-05 | **9.36E-05** |
| ENSG00000011260.16 | **UTP18** | 51096 | protein_coding | **568.423** | -0.370 | -0.111 | **0.774** | 2.51E-05 | **9.38E-05** |
| ENSG00000304119.1 | **ENSG00000304119** | | lncRNA | **52.460** | 1.393 | 0.419 | **2.627** | 2.52E-05 | **9.41E-05** |
| ENSG00000198860.16 | **TSEN15** | 116461 | protein_coding | **446.984** | -0.568 | -0.171 | **0.674** | 2.53E-05 | **9.44E-05** |
| ENSG00000100865.16 | **CINP** | 51550 | protein_coding | **878.471** | -0.584 | -0.176 | **0.667** | 2.53E-05 | **9.45E-05** |
| ENSG00000275759.1 | **ENSG00000275759** | | lncRNA | **10.760** | 2.475 | 0.745 | **5.561** | 2.53E-05 | **9.45E-05** |
| ENSG00000148468.19 | **FAM171A1** | 221061 | protein_coding | **14.501** | 1.766 | 0.532 | **3.401** | 2.53E-05 | **9.45E-05** |
| ENSG00000291300.2 | **PRSS30P** | 124221 | lncRNA | **109.087** | 1.082 | 0.326 | **2.118** | 2.54E-05 | **9.49E-05** |
| ENSG00000269973.1 | **ENSG00000269973** | | lncRNA | **83.147** | 0.792 | 0.239 | **1.732** | 2.55E-05 | **9.49E-05** |
| ENSG00000224383.9 | **PRR29** | 92340 | protein_coding | **14.254** | 2.221 | 0.668 | **4.661** | 2.55E-05 | **9.50E-05** |
| ENSG00000255468.8 | **B4GAT1-D** | 10272406 | lncRNA | **10.210** | 1.993 | 0.600 | **3.980** | 2.55E-05 | **9.50E-05** |
| ENSG00000109113.22 | **RAB34** | 83871 | protein_coding | **269.274** | -0.591 | -0.178 | **0.664** | 2.55E-05 | **9.51E-05** |
| ENSG00000081692.15 | **JMJD4** | 65094 | protein_coding | **316.649** | -0.635 | -0.191 | **0.644** | 2.56E-05 | **9.53E-05** |
| ENSG00000086189.13 | **DIMT1** | 27292 | protein_coding | **238.808** | -0.630 | -0.190 | **0.646** | 2.57E-05 | **9.57E-05** |
| ENSG00000227051.8 | **C14orf132** | 56967 | protein_coding | **15.360** | 1.938 | 0.583 | **3.832** | 2.57E-05 | **9.58E-05** |
| ENSG00000063322.17 | **MED29** | 55588 | protein_coding | **702.864** | 0.387 | 0.116 | **1.307** | 2.57E-05 | **9.58E-05** |

| | | | | | | | | | |
|---|---|---|---|---|---|---|---|---|---|
| ENSG00000122970.18 | **IFT81** | 28981 | protein_coding | **180.306** | 0.680 | 0.205 | **1.602** | 2.59E-05 | **9.65E-05** |
| ENSG00000272970.6 | **ENSG0000(** | 10798616З | lncRNA | **12.179** | 2.440 | 0.735 | **5.427** | 2.60E-05 | **9.68E-05** |
| ENSG00000261611.6 | **ENSG00000261611** | | protein_coding | **44.155** | 1.926 | 0.580 | **3.800** | 2.61E-05 | **9.73E-05** |
| ENSG00000078081.10 | **LAMP3** | 27074 | protein_coding | **1897.440** | 0.642 | 0.193 | **1.561** | 2.61E-05 | **9.73E-05** |
| ENSG00000260830.3 | **ENSG0000(** | 10798464З | lncRNA | **8.437** | -2.103 | -0.633 | **0.233** | 2.62E-05 | **9.76E-05** |
| ENSG00000106462.14 | **EZH2** | 2146 | protein_coding | **1340.524** | -0.502 | -0.151 | **0.706** | 2.63E-05 | **9.78E-05** |
| ENSG00000197054.13 | **ZNF763** | 284390 | protein_coding | **127.170** | 0.888 | 0.267 | **1.851** | 2.64E-05 | **9.83E-05** |
| ENSG00000162441.14 | **LZIC** | 84328 | protein_coding | **284.965** | -0.536 | -0.161 | **0.690** | 2.65E-05 | **9.87E-05** |
| ENSG00000289691.1 | **ENSG00000289691** | | lncRNA | **8.423** | 2.234 | 0.673 | **4.705** | 2.66E-05 | **9.88E-05** |
| ENSG00000131351.17 | **HAUS8** | 93323 | protein_coding | **454.627** | -0.665 | -0.200 | **0.631** | 2.66E-05 | **9.89E-05** |
| ENSG00000134874.20 | **DZIP1** | 22873 | protein_coding | **88.679** | 1.241 | 0.374 | **2.364** | 2.66E-05 | **9.89E-05** |
| ENSG00000145414.11 | **NAF1** | 92345 | protein_coding | **260.147** | -0.605 | -0.182 | **0.657** | 2.66E-05 | **9.89E-05** |
| ENSG00000271797.2 | **ENSG00000271797** | | lncRNA | **11.689** | -1.835 | -0.553 | **0.280** | 2.68E-05 | **9.94E-05** |
| ENSG00000164877.21 | **MICALL2** | 79778 | protein_coding | **79.343** | 1.001 | 0.301 | **2.001** | 2.68E-05 | **9.95E-05** |
| ENSG00000168268.13 | **NT5DC2** | 64943 | protein_coding | **858.767** | -0.839 | -0.253 | **0.559** | 2.68E-05 | **9.95E-05** |
| ENSG00000088808.20 | **PPP1R13B** | 23368 | protein_coding | **272.345** | 0.869 | 0.262 | **1.827** | 2.69E-05 | **9.97E-05** |
| ENSG00000187699.12 | **AKAP19** | 84281 | protein_coding | **70.848** | -1.153 | -0.347 | **0.450** | 2.69E-05 | **1.00E-04** |
| ENSG00000236643.2 | **ENSG00000236643** | | lncRNA | **12.295** | 2.348 | 0.707 | **5.091** | 2.70E-05 | **1.00E-04** |
| ENSG00000184949.19 | **FAM227A** | 646851 | protein_coding | **23.582** | 2.524 | 0.760 | **5.752** | 2.71E-05 | **1.00E-04** |
| ENSG00000159228.14 | **CBR1** | 873 | protein_coding | **366.761** | 0.452 | 0.136 | **1.368** | 2.71E-05 | **1.00E-04** |
| ENSG00000102978.15 | **POLR2C** | 5432 | protein_coding | **785.864** | -0.320 | -0.096 | **0.801** | 2.71E-05 | **1.00E-04** |
| ENSG00000250011.1 | **HMGB1P3** | | processed_pseud( | **84.668** | 0.662 | 0.199 | **1.582** | 2.71E-05 | **1.01E-04** |
| ENSG00000015676.20 | **NUDCD3** | 23386 | protein_coding | **1331.302** | -0.347 | -0.104 | **0.786** | 2.72E-05 | **1.01E-04** |
| ENSG00000232453.8 | **LINC02777** | 10537875З | lncRNA | **11.957** | 2.007 | 0.604 | **4.019** | 2.72E-05 | **1.01E-04** |
| ENSG00000155465.23 | **SLC7A7** | 9056 | protein_coding | **625.916** | 0.678 | 0.204 | **1.600** | 2.75E-05 | **1.02E-04** |
| ENSG00000126216.17 | **TUBGCP3** | 10426 | protein_coding | **501.206** | -0.672 | -0.202 | **0.628** | 2.76E-05 | **1.02E-04** |
| ENSG00000228223.4 | **HCG11** | 493812 | lncRNA | **120.824** | -0.691 | -0.208 | **0.619** | 2.76E-05 | **1.02E-04** |
| ENSG00000246695.10 | **RASSF8-AS1** | | lncRNA | **6.374** | 3.848 | 1.158 | **14.398** | 2.76E-05 | **1.02E-04** |
| ENSG00000219626.11 | **FAM228B** | 375190 | protein_coding | **158.077** | 0.719 | 0.217 | **1.646** | 2.76E-05 | **1.02E-04** |
| ENSG00000128283.9 | **CDC42EP1** | 11135 | protein_coding | **14.579** | 2.676 | 0.806 | **6.391** | 2.77E-05 | **1.03E-04** |
| ENSG00000161249.23 | **DMKN** | 93099 | protein_coding | **434.608** | 1.706 | 0.514 | **3.263** | 2.78E-05 | **1.03E-04** |

| | | | | | | | | | |
|---|---|---|---|---|---|---|---|---|---|
| ENSG00000196411.12 | **EPHB4** | 2050 | protein_coding | **124.281** | 1.158 | 0.349 | **2.232** | 2.78E-05 | **1.03E-04** |
| ENSG00000143226.18 | **FCGR2A** | 2212 | protein_coding | **180.025** | 0.873 | 0.263 | **1.831** | 2.78E-05 | **1.03E-04** |
| ENSG00000130714.21 | **POMT1** | 10585 | protein_coding | **875.439** | 0.535 | 0.161 | **1.449** | 2.79E-05 | **1.03E-04** |
| ENSG00000147121.18 | **KRABD4** | 55634 | protein_coding | **252.739** | -0.612 | -0.184 | **0.654** | 2.79E-05 | **1.03E-04** |
| ENSG00000141428.18 | **RMP24** | 83608 | protein_coding | **281.915** | -0.522 | -0.157 | **0.696** | 2.79E-05 | **1.03E-04** |
| ENSG00000143771.14 | **CNIH4** | 29097 | protein_coding | **477.632** | -0.582 | -0.175 | **0.668** | 2.79E-05 | **1.03E-04** |
| ENSG00000164251.6 | **F2RL1** | 2150 | protein_coding | **4.620** | 3.975 | 1.197 | **15.730** | 2.80E-05 | **1.04E-04** |
| ENSG00000205060.13 | **SLC35B4** | 84912 | protein_coding | **300.020** | -0.623 | -0.187 | **0.649** | 2.82E-05 | **1.04E-04** |
| ENSG00000302789.1 | **ENSG00000302789** | | lncRNA | **6.504** | 3.419 | 1.029 | **10.696** | 2.84E-05 | **1.05E-04** |
| ENSG00000134001.17 | **EIF2S1** | 1965 | protein_coding | **1015.196** | -0.613 | -0.185 | **0.654** | 2.84E-05 | **1.05E-04** |
| ENSG00000188263.11 | **IL17REL** | 400935 | protein_coding | **7.179** | 3.126 | 0.941 | **8.729** | 2.85E-05 | **1.05E-04** |
| ENSG00000179041.4 | **RRS1** | 23212 | protein_coding | **356.154** | -0.552 | -0.166 | **0.682** | 2.85E-05 | **1.05E-04** |
| ENSG00000260077.2 | **ENSG00000260077** | | lncRNA | **89.234** | 0.949 | 0.286 | **1.930** | 2.85E-05 | **1.05E-04** |
| ENSG00000271270.9 | **TMCC1-DT** | 10050703[illegible] | lncRNA | **185.590** | -0.805 | -0.242 | **0.572** | 2.86E-05 | **1.06E-04** |
| ENSG00000101605.16 | **MYOM1** | 8736 | protein_coding | **92.117** | 0.898 | 0.270 | **1.864** | 2.86E-05 | **1.06E-04** |
| ENSG00000139133.8 | **ALG10** | 84920 | protein_coding | **72.116** | -0.777 | -0.234 | **0.584** | 2.86E-05 | **1.06E-04** |
| ENSG00000213203.5 | **GIMAP1** | 170575 | protein_coding | **23.571** | -1.520 | -0.458 | **0.349** | 2.87E-05 | **1.06E-04** |
| ENSG00000096092.8 | **TMEM14A** | 28978 | protein_coding | **208.494** | 0.722 | 0.217 | **1.650** | 2.87E-05 | **1.06E-04** |
| ENSG00000309833.1 | **ENSG00000309833** | | lncRNA | **11.066** | 1.821 | 0.548 | **3.533** | 2.88E-05 | **1.06E-04** |
| ENSG00000118513.23 | **MYB** | 4602 | protein_coding | **304.188** | -0.896 | -0.270 | **0.537** | 2.89E-05 | **1.07E-04** |
| ENSG00000175229.9 | **GAL3ST3** | 89792 | protein_coding | **5.015** | 3.778 | 1.137 | **13.714** | 2.90E-05 | **1.07E-04** |
| ENSG00000188536.15 | **HBA2** | 3040 | protein_coding | **4.694** | 4.081 | 1.229 | **16.927** | 2.91E-05 | **1.07E-04** |
| ENSG00000160209.21 | **PDXK** | 8566 | protein_coding | **2331.279** | -0.497 | -0.150 | **0.709** | 2.91E-05 | **1.07E-04** |
| ENSG00000293025.2 | **ENSG00000293025** | | lncRNA | **25.559** | 1.265 | 0.381 | **2.404** | 2.92E-05 | **1.07E-04** |
| ENSG00000120278.19 | **PLEKHG1** | 57480 | protein_coding | **92.699** | 0.987 | 0.297 | **1.982** | 2.92E-05 | **1.08E-04** |
| ENSG00000179104.10 | **TMTC2** | 160335 | protein_coding | **306.807** | 0.623 | 0.188 | **1.540** | 2.92E-05 | **1.08E-04** |
| ENSG00000112651.14 | **MRPL2** | 51069 | protein_coding | **653.999** | -0.452 | -0.136 | **0.731** | 2.95E-05 | **1.09E-04** |
| ENSG00000127220.8 | **ABHD8** | 79575 | protein_coding | **298.957** | 0.673 | 0.203 | **1.595** | 2.95E-05 | **1.09E-04** |
| ENSG00000181991.18 | **MRPS11** | 64963 | protein_coding | **610.814** | -0.416 | -0.125 | **0.750** | 2.95E-05 | **1.09E-04** |
| ENSG00000130332.17 | **LSM7** | 51690 | protein_coding | **1256.010** | -0.638 | -0.192 | **0.643** | 2.96E-05 | **1.09E-04** |
| ENSG00000240005.8 | **STIM2-AS1** | | lncRNA | **14.696** | 2.225 | 0.670 | **4.674** | 2.97E-05 | **1.09E-04** |

| | | | | | | | | | |
|---|---|---|---|---|---|---|---|---|---|
| ENSG00000179943.10 | **FIZ1** | 84922 | protein_coding | **304.535** | -0.599 | -0.180 | **0.660** | 2.98E-05 | **1.10E-04** |
| ENSG00000166887.18 | **VPS39** | 23339 | protein_coding | **1996.127** | 0.387 | 0.117 | **1.308** | 2.98E-05 | **1.10E-04** |
| ENSG00000140563.18 | **MCTP2** | 55784 | protein_coding | **551.209** | -0.627 | -0.189 | **0.648** | 3.00E-05 | **1.10E-04** |
| ENSG00000284634.2 | **ENSG0000(** | 374443 | lncRNA | **328.143** | 0.688 | 0.207 | **1.611** | 3.01E-05 | **1.10E-04** |
| ENSG00000158373.11 | **H2BC5** | 3017 | protein_coding | **258.986** | 0.698 | 0.210 | **1.622** | 3.01E-05 | **1.11E-04** |
| ENSG00000135624.19 | **CCT7** | 10574 | protein_coding | **3587.072** | -0.578 | -0.174 | **0.670** | 3.01E-05 | **1.11E-04** |
| ENSG00000225138.8 | **SLC9A3-AS1** | | lncRNA | **1724.284** | 0.390 | 0.117 | **1.310** | 3.02E-05 | **1.11E-04** |
| ENSG00000290439.2 | **HSD17B7P:** | 158160 | lncRNA | **302.760** | -0.816 | -0.246 | **0.568** | 3.05E-05 | **1.12E-04** |
| ENSG00000119979.20 | **DENND10** | 404636 | protein_coding | **286.956** | 0.602 | 0.181 | **1.518** | 3.05E-05 | **1.12E-04** |
| ENSG00000275494.3 | **ENSG00000275494** | | lncRNA | **20.461** | 1.493 | 0.449 | **2.814** | 3.08E-05 | **1.13E-04** |
| ENSG00000145725.23 | **PPIP5K2** | 23262 | protein_coding | **255.127** | -1.036 | -0.312 | **0.488** | 3.08E-05 | **1.13E-04** |
| ENSG00000136002.23 | **ARHGEF4** | 50649 | protein_coding | **11.031** | 2.910 | 0.876 | **7.515** | 3.09E-05 | **1.13E-04** |
| ENSG00000138035.17 | **PNPT1** | 87178 | protein_coding | **826.290** | -0.505 | -0.152 | **0.705** | 3.10E-05 | **1.14E-04** |
| ENSG00000189227.7 | **C15orf61** | 145853 | protein_coding | **113.849** | -0.679 | -0.204 | **0.625** | 3.11E-05 | **1.14E-04** |
| ENSG00000011198.12 | **ABHD5** | 51099 | protein_coding | **134.668** | -0.801 | -0.241 | **0.574** | 3.12E-05 | **1.14E-04** |
| ENSG00000179902.15 | **CFAP276** | 127003 | protein_coding | **5.823** | 3.446 | 1.037 | **10.898** | 3.13E-05 | **1.15E-04** |
| ENSG00000146757.15 | **ZNF92** | 168374 | protein_coding | **102.552** | -0.988 | -0.297 | **0.504** | 3.13E-05 | **1.15E-04** |
| ENSG00000265692.1 | **LINC01970** | 10192951 | lncRNA | **15.081** | 1.580 | 0.475 | **2.989** | 3.16E-05 | **1.16E-04** |
| ENSG00000124370.13 | **MCEE** | 84693 | protein_coding | **258.542** | -0.715 | -0.215 | **0.609** | 3.19E-05 | **1.17E-04** |
| ENSG00000168386.20 | **FILIP1L** | 11259 | protein_coding | **52.811** | 1.432 | 0.431 | **2.699** | 3.19E-05 | **1.17E-04** |
| ENSG00000197535.17 | **MYO5A** | 4644 | protein_coding | **740.418** | -0.423 | -0.127 | **0.746** | 3.20E-05 | **1.17E-04** |
| ENSG00000158985.16 | **CDC42SE2** | 56990 | protein_coding | **1982.860** | 0.438 | 0.132 | **1.355** | 3.20E-05 | **1.17E-04** |
| ENSG00000251580.4 | **LINC02482** | | lncRNA | **38.672** | 1.185 | 0.357 | **2.273** | 3.20E-05 | **1.17E-04** |
| ENSG00000141098.15 | **GFOD2** | 81577 | protein_coding | **295.694** | 0.489 | 0.147 | **1.403** | 3.21E-05 | **1.18E-04** |
| ENSG00000162496.12 | **DHRS3** | 9249 | protein_coding | **70.248** | -1.044 | -0.314 | **0.485** | 3.21E-05 | **1.18E-04** |
| ENSG00000243479.4 | **MNX1-AS1** | | lncRNA | **54.320** | 1.003 | 0.302 | **2.005** | 3.21E-05 | **1.18E-04** |
| ENSG00000164647.11 | **STEAP1** | 26872 | protein_coding | **63.418** | -1.193 | -0.359 | **0.437** | 3.23E-05 | **1.18E-04** |
| ENSG00000157796.20 | **WDR19** | 57728 | protein_coding | **354.529** | 0.716 | 0.216 | **1.643** | 3.23E-05 | **1.18E-04** |
| ENSG00000203709.15 | **MIR29B2CHG** | | lncRNA | **397.681** | 0.641 | 0.193 | **1.560** | 3.23E-05 | **1.18E-04** |
| ENSG00000010319.9 | **SEMA3G** | 56920 | protein_coding | **5.452** | 3.194 | 0.961 | **9.151** | 3.24E-05 | **1.18E-04** |
| ENSG00000238269.11 | **PAGE2B** | 389860 | protein_coding | **37.559** | 1.388 | 0.418 | **2.617** | 3.24E-05 | **1.18E-04** |

| | | | | | | | | | |
|---|---|---|---|---|---|---|---|---|---|
| ENSG00000100503.29 | **NIN** | 51199 | protein_coding | **1501.601** | -0.502 | -0.151 | **0.706** | 3.24E-05 | **1.19E-04** |
| ENSG00000306065.1 | **ENSG00000306065** | | lncRNA | **10.385** | 2.887 | 0.869 | **7.399** | 3.25E-05 | **1.19E-04** |
| ENSG00000125347.17 | **IRF1** | 3659 | protein_coding | **1839.341** | 0.823 | 0.248 | **1.769** | 3.25E-05 | **1.19E-04** |
| ENSG00000184408.11 | **KCND2** | 3751 | protein_coding | **34.938** | -1.441 | -0.434 | **0.368** | 3.26E-05 | **1.19E-04** |
| ENSG00000196187.14 | **TMEM63A** | 9725 | protein_coding | **803.061** | -0.368 | -0.111 | **0.775** | 3.26E-05 | **1.19E-04** |
| ENSG00000008311.18 | **AASS** | 10157 | protein_coding | **109.126** | 0.914 | 0.275 | **1.885** | 3.26E-05 | **1.19E-04** |
| ENSG00000284830.1 | **ENSG0000(** | 10192871{ | lncRNA | **7.771** | 2.256 | 0.679 | **4.775** | 3.27E-05 | **1.19E-04** |
| ENSG00000131263.15 | **RLIM** | 51132 | protein_coding | **396.684** | -0.497 | -0.150 | **0.708** | 3.28E-05 | **1.20E-04** |
| ENSG00000156030.16 | **MIDEAS** | 91748 | protein_coding | **1795.879** | 0.560 | 0.168 | **1.474** | 3.28E-05 | **1.20E-04** |
| ENSG00000114923.19 | **SLC4A3** | 6508 | protein_coding | **10.029** | 2.217 | 0.667 | **4.650** | 3.28E-05 | **1.20E-04** |
| ENSG00000168300.16 | **PCMTD1** | 115294 | protein_coding | **601.630** | 0.466 | 0.140 | **1.381** | 3.29E-05 | **1.20E-04** |
| ENSG00000105438.11 | **KDELR1** | 10945 | protein_coding | **2132.668** | -0.452 | -0.136 | **0.731** | 3.29E-05 | **1.20E-04** |
| ENSG00000144455.16 | **SUMF1** | 285362 | protein_coding | **161.290** | 0.593 | 0.179 | **1.508** | 3.29E-05 | **1.20E-04** |
| ENSG00000162972.12 | **MAIP1** | 79568 | protein_coding | **243.807** | -0.561 | -0.169 | **0.678** | 3.30E-05 | **1.20E-04** |
| ENSG00000279879.1 | **ENSG00000279879** | | TEC | **5.892** | 3.650 | 1.099 | **12.554** | 3.30E-05 | **1.20E-04** |
| ENSG00000151465.16 | **CDC123** | 8872 | protein_coding | **1148.437** | -0.405 | -0.122 | **0.755** | 3.31E-05 | **1.21E-04** |
| ENSG00000184368.18 | **MAP7D2** | 256714 | protein_coding | **265.971** | -0.939 | -0.283 | **0.522** | 3.32E-05 | **1.21E-04** |
| ENSG00000274272.1 | **ENSG00000274272** | | lncRNA | **814.486** | 0.499 | 0.150 | **1.413** | 3.32E-05 | **1.21E-04** |
| ENSG00000103495.16 | **MAZ** | 4150 | protein_coding | **2837.759** | -0.401 | -0.121 | **0.757** | 3.34E-05 | **1.22E-04** |
| ENSG00000225889.11 | **ENSG00000225889** | | lncRNA | **996.234** | -0.739 | -0.222 | **0.599** | 3.34E-05 | **1.22E-04** |
| ENSG00000136937.15 | **NCBP1** | 4686 | protein_coding | **584.391** | -0.459 | -0.138 | **0.728** | 3.34E-05 | **1.22E-04** |
| ENSG00000197008.11 | **ZNF138** | 7697 | protein_coding | **182.645** | -0.827 | -0.249 | **0.564** | 3.36E-05 | **1.22E-04** |
| ENSG00000160781.19 | **PAQR6** | 79957 | protein_coding | **264.503** | 0.657 | 0.198 | **1.577** | 3.37E-05 | **1.23E-04** |
| ENSG00000169136.15 | **ATF5** | 22809 | protein_coding | **8265.394** | -0.951 | -0.286 | **0.517** | 3.39E-05 | **1.24E-04** |
| ENSG00000155792.12 | **DEPTOR** | 64798 | protein_coding | **20.871** | 1.770 | 0.533 | **3.410** | 3.40E-05 | **1.24E-04** |
| ENSG00000095794.22 | **CREM** | 1390 | protein_coding | **348.007** | -0.541 | -0.163 | **0.687** | 3.40E-05 | **1.24E-04** |
| ENSG00000119912.20 | **IDE** | 3416 | protein_coding | **336.428** | -0.454 | -0.137 | **0.730** | 3.40E-05 | **1.24E-04** |
| ENSG00000120949.16 | **TNFRSF8** | 943 | protein_coding | **1505.379** | -0.836 | -0.252 | **0.560** | 3.43E-05 | **1.25E-04** |
| ENSG00000303318.1 | **ENSG00000303318** | | lncRNA | **5.770** | 2.971 | 0.894 | **7.840** | 3.44E-05 | **1.25E-04** |
| ENSG00000099250.20 | **NRP1** | 8829 | protein_coding | **12.520** | 2.380 | 0.717 | **5.207** | 3.45E-05 | **1.25E-04** |
| ENSG00000267481.1 | **ENSG00000267481** | | lncRNA | **27.166** | 1.204 | 0.363 | **2.304** | 3.45E-05 | **1.26E-04** |

| | | | | | | | | | |
|---|---|---|---|---|---|---|---|---|---|
| ENSG00000142892.17 | **PIGK** | 10026 | protein_coding | **134.287** | -0.826 | -0.249 | **0.564** | 3.46E-05 | **1.26E-04** |
| ENSG00000113407.16 | **TARS1** | 6897 | protein_coding | **2570.695** | -0.482 | -0.145 | **0.716** | 3.47E-05 | **1.26E-04** |
| ENSG00000129657.18 | **SEC14L1** | 6397 | protein_coding | **1803.266** | 0.358 | 0.108 | **1.282** | 3.48E-05 | **1.26E-04** |
| ENSG00000136478.11 | **TEX2** | 55852 | protein_coding | **192.540** | 0.627 | 0.189 | **1.544** | 3.49E-05 | **1.27E-04** |
| ENSG00000183785.18 | **TUBA8** | 51807 | protein_coding | **224.602** | 1.012 | 0.305 | **2.017** | 3.50E-05 | **1.27E-04** |
| ENSG00000108773.13 | **KAT2A** | 2648 | protein_coding | **2778.310** | 0.448 | 0.135 | **1.365** | 3.52E-05 | **1.28E-04** |
| ENSG00000198056.18 | **PRIM1** | 5557 | protein_coding | **188.268** | -0.722 | -0.217 | **0.606** | 3.52E-05 | **1.28E-04** |
| ENSG00000180228.16 | **PRKRA** | 8575 | protein_coding | **585.848** | -0.431 | -0.130 | **0.742** | 3.52E-05 | **1.28E-04** |
| ENSG00000140104.15 | **CLBA1** | 122616 | protein_coding | **66.198** | 1.017 | 0.306 | **2.023** | 3.54E-05 | **1.29E-04** |
| ENSG00000141447.21 | **OSBPL1A** | 114876 | protein_coding | **8.985** | 2.679 | 0.806 | **6.403** | 3.55E-05 | **1.29E-04** |
| ENSG00000198464.16 | **ZNF480** | 147657 | protein_coding | **417.314** | -0.550 | -0.166 | **0.683** | 3.56E-05 | **1.29E-04** |
| ENSG00000167186.13 | **COQ7** | 10229 | protein_coding | **288.601** | -0.639 | -0.192 | **0.642** | 3.56E-05 | **1.29E-04** |
| ENSG00000054983.19 | **GALC** | 2581 | protein_coding | **100.730** | -0.821 | -0.247 | **0.566** | 3.56E-05 | **1.29E-04** |
| ENSG00000068078.22 | **FGFR3** | 2261 | protein_coding | **19.130** | 2.249 | 0.677 | **4.755** | 3.56E-05 | **1.29E-04** |
| ENSG00000041988.18 | **THAP3** | 90326 | protein_coding | **357.091** | -0.500 | -0.151 | **0.707** | 3.57E-05 | **1.29E-04** |
| ENSG00000204610.16 | **TRIM15** | 89870 | protein_coding | **5.074** | 4.129 | 1.243 | **17.501** | 3.57E-05 | **1.29E-04** |
| ENSG00000106624.13 | **AEBP1** | 165 | protein_coding | **172.300** | 1.384 | 0.417 | **2.609** | 3.58E-05 | **1.30E-04** |
| ENSG00000198498.12 | **TMA16** | 55319 | protein_coding | **373.566** | -0.518 | -0.156 | **0.698** | 3.58E-05 | **1.30E-04** |
| ENSG00000173848.21 | **NET1** | 10276 | protein_coding | **239.658** | -0.855 | -0.257 | **0.553** | 3.59E-05 | **1.30E-04** |
| ENSG00000172172.10 | **MRPL13** | 28998 | protein_coding | **508.722** | -0.535 | -0.161 | **0.690** | 3.60E-05 | **1.30E-04** |
| ENSG00000232653.10 | **GOLGA8N** | 643699 | protein_coding | **40.228** | 1.012 | 0.305 | **2.017** | 3.61E-05 | **1.31E-04** |
| ENSG00000005513.12 | **SOX8** | 30812 | protein_coding | **8.685** | 2.312 | 0.696 | **4.965** | 3.62E-05 | **1.31E-04** |
| ENSG00000225828.2 | **FAM229A** | 100128071 | protein_coding | **222.900** | 0.846 | 0.255 | **1.797** | 3.62E-05 | **1.31E-04** |
| ENSG00000198133.11 | **TMEM229I** | 161145 | protein_coding | **21.109** | 1.475 | 0.444 | **2.779** | 3.63E-05 | **1.31E-04** |
| ENSG00000164465.21 | **DCBLD1** | 285761 | protein_coding | **171.245** | 0.716 | 0.216 | **1.643** | 3.63E-05 | **1.31E-04** |
| ENSG00000287576.2 | **ENSG00000287576** | | lncRNA | **12.064** | 1.822 | 0.549 | **3.536** | 3.64E-05 | **1.32E-04** |
| ENSG00000139173.12 | **TMEM117** | 84216 | protein_coding | **39.372** | -1.074 | -0.323 | **0.475** | 3.66E-05 | **1.32E-04** |
| ENSG00000275023.7 | **MLLT6** | 4302 | protein_coding | **5187.989** | 0.442 | 0.133 | **1.358** | 3.66E-05 | **1.32E-04** |
| ENSG00000179168.16 | **GGN** | 199720 | protein_coding | **11.370** | 3.082 | 0.928 | **8.466** | 3.68E-05 | **1.33E-04** |
| ENSG00000182087.16 | **TMEM259** | 91304 | protein_coding | **4725.993** | 0.391 | 0.118 | **1.311** | 3.68E-05 | **1.33E-04** |
| ENSG00000001617.14 | **SEMA3F** | 6405 | protein_coding | **12.229** | 2.552 | 0.768 | **5.864** | 3.68E-05 | **1.33E-04** |

| | | | | | | | | | |
|---|---|---|---|---|---|---|---|---|---|
| ENSG00000211893.4 | **IGHG2** | | IG_C_gene | **2652.265** | -0.587 | -0.177 | **0.666** | 3.68E-05 | **1.33E-04** |
| ENSG00000129990.17 | **SYT5** | 6861 | protein_coding | **35.800** | 1.646 | 0.495 | **3.129** | 3.69E-05 | **1.33E-04** |
| ENSG00000101639.21 | **CEP192** | 55125 | protein_coding | **513.515** | -0.441 | -0.133 | **0.737** | 3.70E-05 | **1.34E-04** |
| ENSG00000148925.13 | **BTBD10** | 84280 | protein_coding | **258.637** | -0.459 | -0.138 | **0.727** | 3.71E-05 | **1.34E-04** |
| ENSG00000086848.17 | **ALG9** | 79796 | protein_coding | **465.190** | -0.611 | -0.184 | **0.655** | 3.71E-05 | **1.34E-04** |
| ENSG00000147679.14 | **UTP23** | 84294 | protein_coding | **187.991** | -0.677 | -0.204 | **0.626** | 3.71E-05 | **1.34E-04** |
| ENSG00000275799.1 | **ENSG00000275799** | | lncRNA | **41.524** | -1.337 | -0.403 | **0.396** | 3.72E-05 | **1.34E-04** |
| ENSG00000184060.14 | **ADAP2** | 55803 | protein_coding | **47.473** | 1.308 | 0.394 | **2.475** | 3.73E-05 | **1.35E-04** |
| ENSG00000123728.12 | **RAP2C** | 57826 | protein_coding | **368.593** | -0.677 | -0.204 | **0.625** | 3.75E-05 | **1.35E-04** |
| ENSG00000138131.6 | **LOXL4** | 84171 | protein_coding | **9.360** | -2.516 | -0.757 | **0.175** | 3.75E-05 | **1.35E-04** |
| ENSG00000133678.16 | **TMEM254** | 80195 | protein_coding | **202.218** | -0.583 | -0.176 | **0.668** | 3.76E-05 | **1.36E-04** |
| ENSG00000112877.10 | **CEP72** | 55722 | protein_coding | **171.749** | -0.814 | -0.245 | **0.569** | 3.76E-05 | **1.36E-04** |
| ENSG00000205572.12 | **SERF1B** | 728492 | protein_coding | **213.939** | -0.530 | -0.160 | **0.692** | 3.77E-05 | **1.36E-04** |
| ENSG00000112167.12 | **SAYSD1** | 55776 | protein_coding | **116.244** | -0.663 | -0.200 | **0.631** | 3.77E-05 | **1.36E-04** |
| ENSG00000197124.13 | **ZNF682** | 91120 | protein_coding | **86.947** | -1.306 | -0.393 | **0.404** | 3.77E-05 | **1.36E-04** |
| ENSG00000106554.15 | **CHCHD3** | 54927 | protein_coding | **1095.865** | -0.543 | -0.163 | **0.686** | 3.80E-05 | **1.37E-04** |
| ENSG00000242498.10 | **ARPIN** | 348110 | protein_coding | **8.829** | 3.308 | 0.996 | **9.901** | 3.81E-05 | **1.37E-04** |
| ENSG00000267082.3 | **DOCK6-AS1** | | lncRNA | **13.759** | 2.085 | 0.628 | **4.243** | 3.81E-05 | **1.37E-04** |
| ENSG00000175906.7 | **ARL4D** | 379 | protein_coding | **32.189** | 1.051 | 0.317 | **2.073** | 3.81E-05 | **1.37E-04** |
| ENSG00000272356.1 | **REV3L-AS1** | | lncRNA | **41.565** | 0.926 | 0.279 | **1.901** | 3.85E-05 | **1.39E-04** |
| ENSG00000147570.12 | **DNAJC5B** | 85479 | protein_coding | **34.224** | 1.286 | 0.387 | **2.438** | 3.85E-05 | **1.39E-04** |
| ENSG00000183186.10 | **C2CD4C** | 126567 | protein_coding | **4.001** | 4.042 | 1.217 | **16.470** | 3.86E-05 | **1.39E-04** |
| ENSG00000154864.15 | **PIEZO2** | 63895 | protein_coding | **481.885** | 0.618 | 0.186 | **1.535** | 3.87E-05 | **1.39E-04** |
| ENSG00000164659.17 | **ELAPOR2** | 222223 | protein_coding | **89.512** | 1.466 | 0.441 | **2.763** | 3.90E-05 | **1.40E-04** |
| ENSG00000244486.11 | **SCARF2** | 91179 | protein_coding | **40.594** | 0.961 | 0.289 | **1.947** | 3.92E-05 | **1.41E-04** |
| ENSG00000001497.20 | **LAS1L** | 81887 | protein_coding | **1210.412** | -0.402 | -0.121 | **0.757** | 3.93E-05 | **1.41E-04** |
| ENSG00000298255.1 | **ENSG00000298255** | | lncRNA | **17.800** | 1.944 | 0.585 | **3.849** | 3.96E-05 | **1.42E-04** |
| ENSG00000177406.6 | **NINJ2-AS1** | 10004971€ | lncRNA | **327.355** | 0.525 | 0.158 | **1.439** | 3.99E-05 | **1.43E-04** |
| ENSG00000121073.17 | **SLC35B1** | 10237 | protein_coding | **513.304** | -0.514 | -0.155 | **0.700** | 3.99E-05 | **1.43E-04** |
| ENSG00000241837.9 | **ATP5PO** | 539 | protein_coding | **3512.411** | -0.515 | -0.155 | **0.700** | 3.99E-05 | **1.44E-04** |
| ENSG00000203499.13 | **IQANK1** | 642574 | protein_coding | **7.751** | 2.620 | 0.789 | **6.146** | 4.00E-05 | **1.44E-04** |

| | | | | | | | | | |
|---|---|---|---|---|---|---|---|---|---|
| ENSG00000268655.3 | **SAXO3** | 10105994ɛ | protein_coding | **7.330** | 2.642 | 0.795 | **6.242** | 4.01E-05 | **1.44E-04** |
| ENSG00000115524.19 | **SF3B1** | 23451 | protein_coding | **8815.109** | 0.526 | 0.158 | **1.440** | 4.01E-05 | **1.44E-04** |
| ENSG00000108559.15 | **NUP88** | 4927 | protein_coding | **1203.803** | -0.547 | -0.165 | **0.684** | 4.03E-05 | **1.45E-04** |
| ENSG00000266053.4 | **NDUFV2-AS1** | | lncRNA | **30.312** | 1.257 | 0.378 | **2.390** | 4.03E-05 | **1.45E-04** |
| ENSG00000066056.17 | **TIE1** | 7075 | protein_coding | **28.397** | 1.530 | 0.461 | **2.888** | 4.03E-05 | **1.45E-04** |
| ENSG00000174136.15 | **RGMB** | 285704 | protein_coding | **115.634** | 0.690 | 0.208 | **1.614** | 4.03E-05 | **1.45E-04** |
| ENSG00000102265.14 | **TIMP1** | 7076 | protein_coding | **2548.632** | 0.829 | 0.249 | **1.776** | 4.04E-05 | **1.45E-04** |
| ENSG00000165280.20 | **VCP** | 7415 | protein_coding | **4275.678** | -0.495 | -0.149 | **0.710** | 4.06E-05 | **1.46E-04** |
| ENSG00000100219.19 | **XBP1** | 7494 | protein_coding | **3416.406** | -0.678 | -0.204 | **0.625** | 4.06E-05 | **1.46E-04** |
| ENSG00000114026.23 | **OGG1** | 4968 | protein_coding | **269.272** | -0.614 | -0.185 | **0.653** | 4.07E-05 | **1.46E-04** |
| ENSG00000118514.16 | **ALDH8A1** | 64577 | protein_coding | **42.171** | 1.053 | 0.317 | **2.074** | 4.08E-05 | **1.46E-04** |
| ENSG00000276903.2 | **H2AC16** | 8332 | protein_coding | **8.513** | -2.338 | -0.704 | **0.198** | 4.08E-05 | **1.46E-04** |
| ENSG00000164236.13 | **ANKRD33E** | 651746 | protein_coding | **1162.068** | 0.676 | 0.204 | **1.598** | 4.08E-05 | **1.46E-04** |
| ENSG00000100916.16 | **BRMS1L** | 84312 | protein_coding | **73.131** | -0.779 | -0.235 | **0.583** | 4.09E-05 | **1.46E-04** |
| ENSG00000094880.13 | **CDC23** | 8697 | protein_coding | **538.725** | -0.545 | -0.164 | **0.685** | 4.10E-05 | **1.47E-04** |
| ENSG00000196262.17 | **PPIA** | 5478 | protein_coding | **29412.616** | -0.622 | -0.187 | **0.650** | 4.10E-05 | **1.47E-04** |
| ENSG00000011332.24 | **DPF1** | 8193 | protein_coding | **23.453** | 1.267 | 0.381 | **2.407** | 4.10E-05 | **1.47E-04** |
| ENSG00000124383.13 | **MPHOSPH** | 10199 | protein_coding | **549.905** | -0.458 | -0.138 | **0.728** | 4.11E-05 | **1.47E-04** |
| ENSG00000288875.3 | **ENSG00000288875** | | lncRNA | **12.202** | 2.626 | 0.790 | **6.171** | 4.12E-05 | **1.47E-04** |
| ENSG00000104812.17 | **GYS1** | 2997 | protein_coding | **1282.459** | -0.435 | -0.131 | **0.740** | 4.14E-05 | **1.48E-04** |
| ENSG00000258702.6 | **ENSG0000** | 10537064 | lncRNA | **25.377** | 1.269 | 0.382 | **2.409** | 4.14E-05 | **1.48E-04** |
| ENSG00000204116.13 | **CHIC1** | 53344 | protein_coding | **113.853** | -0.696 | -0.210 | **0.617** | 4.14E-05 | **1.48E-04** |
| ENSG00000228395.4 | **ENSG0000** | 10192927 | lncRNA | **16.470** | 1.628 | 0.490 | **3.091** | 4.15E-05 | **1.48E-04** |
| ENSG00000204842.20 | **ATXN2** | 6311 | protein_coding | **799.361** | 0.484 | 0.146 | **1.399** | 4.16E-05 | **1.49E-04** |
| ENSG00000271856.2 | **LINC01215** | 10192962 | lncRNA | **428.609** | 0.825 | 0.248 | **1.772** | 4.16E-05 | **1.49E-04** |
| ENSG00000197646.10 | **PDCD1LG2** | 80380 | protein_coding | **176.553** | 0.654 | 0.197 | **1.574** | 4.16E-05 | **1.49E-04** |
| ENSG00000121653.13 | **MAPK8IP1** | 9479 | protein_coding | **30.186** | 1.237 | 0.372 | **2.357** | 4.17E-05 | **1.49E-04** |
| ENSG00000288009.2 | **ENSG00000288009** | | lncRNA | **10.366** | 1.861 | 0.560 | **3.633** | 4.20E-05 | **1.50E-04** |
| ENSG00000166689.20 | **PLEKHA7** | 144100 | protein_coding | **58.217** | 1.028 | 0.309 | **2.039** | 4.23E-05 | **1.51E-04** |
| ENSG00000122786.22 | **CALD1** | 800 | protein_coding | **20.794** | 1.679 | 0.505 | **3.202** | 4.23E-05 | **1.51E-04** |
| ENSG00000087502.21 | **ERGIC2** | 51290 | protein_coding | **352.400** | -0.567 | -0.171 | **0.675** | 4.23E-05 | **1.51E-04** |

| | | | | | | | | | |
|---|---|---|---|---|---|---|---|---|---|
| ENSG00000100439.13 | **ABHD4** | 63874 | protein_coding | **682.118** | -0.637 | -0.192 | **0.643** | 4.24E-05 | **1.52E-04** |
| ENSG00000273064.1 | **ENSG00000273064** | | lncRNA | **37.580** | 1.194 | 0.359 | **2.288** | 4.25E-05 | **1.52E-04** |
| ENSG00000243629.3 | **LINC00880** | 339894 | lncRNA | **23.871** | 1.209 | 0.364 | **2.311** | 4.26E-05 | **1.52E-04** |
| ENSG00000112232.13 | **KHDRBS2** | 202559 | protein_coding | **4.122** | 3.957 | 1.191 | **15.531** | 4.27E-05 | **1.52E-04** |
| ENSG00000188385.14 | **JAKMIP3** | 282973 | protein_coding | **6.798** | 2.610 | 0.786 | **6.107** | 4.28E-05 | **1.53E-04** |
| ENSG00000205702.14 | **CYP2D7** | 10537720[illegible] | protein_coding | **234.430** | 0.725 | 0.218 | **1.653** | 4.30E-05 | **1.53E-04** |
| ENSG00000124201.16 | **ZNFX1** | 57169 | protein_coding | **1432.108** | 0.514 | 0.155 | **1.428** | 4.30E-05 | **1.53E-04** |
| ENSG00000101247.20 | **NDUFAF5** | 79133 | protein_coding | **214.426** | -0.685 | -0.206 | **0.622** | 4.30E-05 | **1.53E-04** |
| ENSG00000020256.22 | **ZFP64** | 55734 | protein_coding | **223.397** | -0.601 | -0.181 | **0.659** | 4.31E-05 | **1.54E-04** |
| ENSG00000125944.24 | **HNRNPR** | 10236 | protein_coding | **1272.124** | -0.564 | -0.170 | **0.676** | 4.32E-05 | **1.54E-04** |
| ENSG00000135245.12 | **HILPDA** | 29923 | protein_coding | **988.954** | -0.590 | -0.177 | **0.665** | 4.36E-05 | **1.55E-04** |
| ENSG00000147872.12 | **PLIN2** | 123 | protein_coding | **921.842** | -0.613 | -0.184 | **0.654** | 4.37E-05 | **1.56E-04** |
| ENSG00000103021.12 | **CFAP263** | 29070 | protein_coding | **79.364** | 0.863 | 0.260 | **1.819** | 4.37E-05 | **1.56E-04** |
| ENSG00000287743.2 | **ENSG00000287743** | | lncRNA | **6.974** | 3.406 | 1.025 | **10.597** | 4.38E-05 | **1.56E-04** |
| ENSG00000178381.15 | **ZFAND2A** | 90637 | protein_coding | **179.652** | -0.733 | -0.221 | **0.602** | 4.38E-05 | **1.56E-04** |
| ENSG00000111913.22 | **RIPOR2** | 9750 | protein_coding | **992.243** | 0.575 | 0.173 | **1.490** | 4.38E-05 | **1.56E-04** |
| ENSG00000160050.18 | **CCDC28B** | 79140 | protein_coding | **952.085** | -0.491 | -0.148 | **0.711** | 4.40E-05 | **1.57E-04** |
| ENSG00000165591.9 | **FAAH2** | 158584 | protein_coding | **76.724** | 0.677 | 0.204 | **1.599** | 4.41E-05 | **1.57E-04** |
| ENSG00000174799.14 | **CEP135** | 9662 | protein_coding | **134.232** | -0.710 | -0.214 | **0.611** | 4.42E-05 | **1.57E-04** |
| ENSG00000228719.2 | **ENSG00000228719** | | lncRNA | **11.143** | 1.846 | 0.556 | **3.594** | 4.42E-05 | **1.57E-04** |
| ENSG00000224843.9 | **LINC00240** | 10013320[illegible] | lncRNA | **16.000** | 1.890 | 0.569 | **3.706** | 4.42E-05 | **1.57E-04** |
| ENSG00000130821.19 | **SLC6A8** | 6535 | protein_coding | **77.229** | 0.984 | 0.296 | **1.978** | 4.42E-05 | **1.57E-04** |
| ENSG00000188157.17 | **AGRN** | 375790 | protein_coding | **2806.502** | 0.359 | 0.108 | **1.283** | 4.43E-05 | **1.57E-04** |
| ENSG00000165055.18 | **METTL2B** | 55798 | protein_coding | **425.361** | -0.585 | -0.176 | **0.667** | 4.43E-05 | **1.57E-04** |
| ENSG00000198585.13 | **NUDT16** | 131870 | protein_coding | **458.688** | -0.384 | -0.116 | **0.766** | 4.44E-05 | **1.58E-04** |
| ENSG00000105556.14 | **MIER2** | 54531 | protein_coding | **452.633** | -0.552 | -0.166 | **0.682** | 4.44E-05 | **1.58E-04** |
| ENSG00000278845.7 | **MRPL45** | 84311 | protein_coding | **675.817** | -0.443 | -0.133 | **0.736** | 4.44E-05 | **1.58E-04** |
| ENSG00000168610.19 | **STAT3** | 6774 | protein_coding | **3633.876** | 0.556 | 0.167 | **1.470** | 4.44E-05 | **1.58E-04** |
| ENSG00000118965.18 | **WDR35** | 57539 | protein_coding | **215.831** | 0.669 | 0.201 | **1.590** | 4.45E-05 | **1.58E-04** |
| ENSG00000204256.17 | **BRD2** | 6046 | protein_coding | **3920.579** | -0.333 | -0.100 | **0.794** | 4.48E-05 | **1.59E-04** |
| ENSG00000217930.10 | **PAM16** | 51025 | protein_coding | **517.592** | -0.653 | -0.197 | **0.636** | 4.49E-05 | **1.60E-04** |

| | | | | | | | | | |
|---|---|---|---|---|---|---|---|---|---|
| ENSG00000142784.18 | **WDTC1** | 23038 | protein_coding | **1109.682** | 0.455 | 0.137 | **1.371** | 4.52E-05 | **1.60E-04** |
| ENSG00000255072.3 | **PIGY** | 84992 | protein_coding | **102.715** | 12.295 | 3.701 | **5026.764** | 4.54E-05 | **1.61E-04** |
| ENSG00000267106.10 | **ZNF561-AS1** | | lncRNA | **82.890** | -0.961 | -0.289 | **0.514** | 4.55E-05 | **1.61E-04** |
| ENSG00000197774.15 | **EME2** | 197342 | protein_coding | **1084.221** | 0.413 | 0.124 | **1.332** | 4.55E-05 | **1.62E-04** |
| ENSG00000092201.12 | **SUPT16H** | 11198 | protein_coding | **2659.874** | -0.329 | -0.099 | **0.796** | 4.56E-05 | **1.62E-04** |
| ENSG00000188897.10 | **ENSG0000(** | 400499 | protein_coding | **156.923** | 0.844 | 0.254 | **1.795** | 4.59E-05 | **1.63E-04** |
| ENSG00000078018.24 | **MAP2** | 4133 | protein_coding | **7.731** | 4.379 | 1.318 | **20.806** | 4.60E-05 | **1.63E-04** |
| ENSG00000198663.19 | **C6orf89** | 221477 | protein_coding | **852.797** | 0.621 | 0.187 | **1.538** | 4.63E-05 | **1.64E-04** |
| ENSG00000288612.1 | **ENSG00000288612** | | lncRNA | **7.505** | 2.299 | 0.692 | **4.921** | 4.63E-05 | **1.64E-04** |
| ENSG00000065911.14 | **MTHFD2** | 10797 | protein_coding | **2305.575** | -0.847 | -0.255 | **0.556** | 4.64E-05 | **1.65E-04** |
| ENSG00000101558.16 | **VAPA** | 9218 | protein_coding | **1430.263** | -0.557 | -0.168 | **0.680** | 4.64E-05 | **1.65E-04** |
| ENSG00000066044.17 | **ELAVL1** | 1994 | protein_coding | **1698.700** | -0.361 | -0.109 | **0.779** | 4.66E-05 | **1.65E-04** |
| ENSG00000226029.1 | **LINC01772** | 107984921 | lncRNA | **21.084** | 1.335 | 0.402 | **2.522** | 4.66E-05 | **1.65E-04** |
| ENSG00000296549.1 | **ENSG00000296549** | | lncRNA | **6.883** | 2.779 | 0.836 | **6.863** | 4.66E-05 | **1.65E-04** |
| ENSG00000260296.1 | **ENSG00000260296** | | lncRNA | **38.646** | 1.182 | 0.356 | **2.269** | 4.66E-05 | **1.65E-04** |
| ENSG00000257218.8 | **GATC** | 283459 | protein_coding | **629.264** | -0.346 | -0.104 | **0.787** | 4.67E-05 | **1.65E-04** |
| ENSG00000137713.18 | **PPP2R1B** | 5519 | protein_coding | **319.018** | -0.623 | -0.187 | **0.649** | 4.68E-05 | **1.66E-04** |
| ENSG00000107864.17 | **CPEB3** | 22849 | protein_coding | **48.450** | 1.103 | 0.332 | **2.149** | 4.69E-05 | **1.66E-04** |
| ENSG00000116353.18 | **MECR** | 51102 | protein_coding | **331.551** | -0.579 | -0.174 | **0.669** | 4.69E-05 | **1.66E-04** |
| ENSG00000272374.1 | **SCUBE3-AS1** | | lncRNA | **57.251** | 0.784 | 0.236 | **1.722** | 4.69E-05 | **1.66E-04** |
| ENSG00000164708.8 | **PGAM2** | 5224 | protein_coding | **7.675** | 2.536 | 0.763 | **5.798** | 4.70E-05 | **1.66E-04** |
| ENSG00000234719.9 | **NPIPB2** | 729978 | protein_coding | **139.198** | 0.677 | 0.204 | **1.599** | 4.72E-05 | **1.67E-04** |
| ENSG00000293953.1 | **ENSG00000293953** | | lncRNA | **6.875** | 2.583 | 0.777 | **5.990** | 4.75E-05 | **1.68E-04** |
| ENSG00000138363.17 | **ATIC** | 471 | protein_coding | **2314.456** | -0.456 | -0.137 | **0.729** | 4.76E-05 | **1.68E-04** |
| ENSG00000137210.16 | **TMEM14B** | 81853 | protein_coding | **903.397** | -0.373 | -0.112 | **0.772** | 4.76E-05 | **1.68E-04** |
| ENSG00000144635.11 | **DYNC1LI1** | 51143 | protein_coding | **384.359** | -0.463 | -0.139 | **0.726** | 4.77E-05 | **1.69E-04** |
| ENSG00000115694.17 | **STK25** | 10494 | protein_coding | **1728.400** | 0.309 | 0.093 | **1.239** | 4.78E-05 | **1.69E-04** |
| ENSG00000289277.2 | **ENSG00000289277** | | lncRNA | **11.023** | 2.110 | 0.635 | **4.316** | 4.81E-05 | **1.70E-04** |
| ENSG00000289332.3 | **ENSG00000289332** | | lncRNA | **5.484** | 4.194 | 1.263 | **18.306** | 4.81E-05 | **1.70E-04** |
| ENSG00000148798.13 | **INA** | 9118 | protein_coding | **12.226** | 1.753 | 0.528 | **3.371** | 4.83E-05 | **1.71E-04** |
| ENSG00000187987.12 | **ZSCAN23** | 222696 | protein_coding | **109.382** | 0.949 | 0.286 | **1.931** | 4.83E-05 | **1.71E-04** |

| | | | | | | | | | |
|---|---|---|---|---|---|---|---|---|---|
| ENSG00000196998.21 | **WDR45** | 11152 | protein_coding | **491.743** | -0.402 | -0.121 | **0.757** | 4.88E-05 | **1.72E-04** |
| ENSG00000143507.20 | **DUSP10** | 11221 | protein_coding | **807.548** | -0.364 | -0.109 | **0.777** | 4.88E-05 | **1.72E-04** |
| ENSG00000273445.2 | **ENSG00000273445** | | lncRNA | **5.182** | 4.011 | 1.207 | **16.123** | 4.88E-05 | **1.72E-04** |
| ENSG00000136059.18 | **VILL** | 50853 | protein_coding | **262.128** | -0.579 | -0.174 | **0.670** | 4.89E-05 | **1.72E-04** |
| ENSG00000162729.16 | **IGSF8** | 93185 | protein_coding | **489.135** | 0.490 | 0.148 | **1.405** | 4.91E-05 | **1.73E-04** |
| ENSG00000134597.19 | **RBMX2** | 51634 | protein_coding | **577.270** | -0.365 | -0.110 | **0.777** | 4.91E-05 | **1.73E-04** |
| ENSG00000115138.13 | **POMC** | 5443 | protein_coding | **259.600** | 0.611 | 0.184 | **1.528** | 4.92E-05 | **1.74E-04** |
| ENSG00000289370.3 | **ENSG00000289370** | | lncRNA | **33.823** | 1.144 | 0.344 | **2.210** | 4.93E-05 | **1.74E-04** |
| ENSG00000291146.2 | **RP9P** | 441212 | lncRNA | **82.071** | -0.808 | -0.243 | **0.571** | 4.94E-05 | **1.74E-04** |
| ENSG00000101049.20 | **SGK2** | 10110 | protein_coding | **24.882** | 1.234 | 0.372 | **2.353** | 4.96E-05 | **1.75E-04** |
| ENSG00000129467.16 | **ADCY4** | 196883 | protein_coding | **65.272** | 1.065 | 0.321 | **2.093** | 4.97E-05 | **1.75E-04** |
| ENSG00000288926.3 | **ENSG00000288926** | | lncRNA | **37.443** | 1.237 | 0.372 | **2.358** | 4.99E-05 | **1.76E-04** |
| ENSG00000197122.15 | **SRC** | 6714 | protein_coding | **1808.359** | 0.480 | 0.144 | **1.395** | 5.00E-05 | **1.76E-04** |
| ENSG00000145777.15 | **TSLP** | 85480 | protein_coding | **18.186** | 1.905 | 0.574 | **3.746** | 5.00E-05 | **1.76E-04** |
| ENSG00000157349.19 | **DDX19B** | 11269 | protein_coding | **937.865** | -0.467 | -0.141 | **0.723** | 5.01E-05 | **1.76E-04** |
| ENSG00000110955.11 | **ATP5F1B** | 506 | protein_coding | **8583.130** | -0.542 | -0.163 | **0.687** | 5.02E-05 | **1.77E-04** |
| ENSG00000287252.4 | **ENSG00000287252** | | lncRNA | **289.495** | 0.446 | 0.134 | **1.362** | 5.03E-05 | **1.77E-04** |
| ENSG00000272994.3 | **C2orf49-DT** | | lncRNA | **119.396** | 0.583 | 0.175 | **1.498** | 5.04E-05 | **1.77E-04** |
| ENSG00000243417.1 | **RPS3AP18** | | processed_pseud | **32.949** | 1.201 | 0.362 | **2.299** | 5.04E-05 | **1.77E-04** |
| ENSG00000261804.4 | **ENSG00000261804** | | lncRNA | **7.007** | -3.513 | -1.057 | **0.088** | 5.05E-05 | **1.78E-04** |
| ENSG00000159063.16 | **ALG8** | 79053 | protein_coding | **490.824** | -0.859 | -0.259 | **0.551** | 5.06E-05 | **1.78E-04** |
| ENSG00000237489.7 | **C10orf143** | 387723 | protein_coding | **76.814** | 0.724 | 0.218 | **1.651** | 5.06E-05 | **1.78E-04** |
| ENSG00000254166.5 | **CASC19** | 10316461 | lncRNA | **19.593** | 1.435 | 0.432 | **2.704** | 5.08E-05 | **1.78E-04** |
| ENSG00000263002.10 | **ZNF234** | 10780 | protein_coding | **161.889** | -0.577 | -0.174 | **0.670** | 5.09E-05 | **1.79E-04** |
| ENSG00000168090.12 | **COPS6** | 10980 | protein_coding | **1483.042** | -0.465 | -0.140 | **0.725** | 5.09E-05 | **1.79E-04** |
| ENSG00000227857.3 | **ENSG00000227857** | | lncRNA | **7.179** | 2.833 | 0.853 | **7.124** | 5.10E-05 | **1.79E-04** |
| ENSG00000141002.22 | **TCF25** | 22980 | protein_coding | **2097.483** | -0.549 | -0.165 | **0.683** | 5.11E-05 | **1.79E-04** |
| ENSG00000133636.13 | **NTS** | 4922 | protein_coding | **10.305** | 3.271 | 0.985 | **9.650** | 5.12E-05 | **1.80E-04** |
| ENSG00000197880.11 | **MDS2** | 259283 | lncRNA | **21.856** | 1.929 | 0.581 | **3.808** | 5.13E-05 | **1.80E-04** |
| ENSG00000303745.1 | **ENSG00000303745** | | lncRNA | **19.840** | 1.544 | 0.465 | **2.916** | 5.13E-05 | **1.80E-04** |
| ENSG00000204913.8 | **LRRC3C** | 100505591 | protein_coding | **6.762** | 2.911 | 0.876 | **7.520** | 5.15E-05 | **1.81E-04** |

| | | | | | | | | | |
|---|---|---|---|---|---|---|---|---|---|
| ENSG00000308204.1 | **ENSG00000308204** | | lncRNA | **25.650** | 1.205 | 0.363 | **2.305** | 5.16E-05 | **1.81E-04** |
| ENSG00000121022.16 | **COPS5** | 10987 | protein_coding | **849.954** | -0.354 | -0.107 | **0.782** | 5.17E-05 | **1.81E-04** |
| ENSG00000149743.16 | **TRPT1** | 83707 | protein_coding | **1064.441** | 0.535 | 0.161 | **1.449** | 5.18E-05 | **1.82E-04** |
| ENSG00000267673.8 | **FDX2** | 112812 | protein_coding | **165.015** | -0.575 | -0.173 | **0.671** | 5.21E-05 | **1.83E-04** |
| ENSG00000143322.22 | **ABL2** | 27 | protein_coding | **169.656** | 0.714 | 0.215 | **1.641** | 5.21E-05 | **1.83E-04** |
| ENSG00000235513.2 | **L3MBTL2-AS1** | | lncRNA | **12.270** | -2.172 | -0.654 | **0.222** | 5.22E-05 | **1.83E-04** |
| ENSG00000103056.14 | **SMPD3** | 55512 | protein_coding | **14.571** | -1.598 | -0.481 | **0.330** | 5.22E-05 | **1.83E-04** |
| ENSG00000100319.15 | **ZMAT5** | 55954 | protein_coding | **489.541** | 0.453 | 0.136 | **1.369** | 5.23E-05 | **1.83E-04** |
| ENSG00000071127.20 | **WDR1** | 9948 | protein_coding | **3632.041** | -0.358 | -0.108 | **0.780** | 5.23E-05 | **1.83E-04** |
| ENSG00000102967.14 | **DHODH** | 1723 | protein_coding | **258.847** | -0.583 | -0.176 | **0.667** | 5.23E-05 | **1.83E-04** |
| ENSG00000131203.14 | **IDO1** | 3620 | protein_coding | **13.015** | 2.928 | 0.881 | **7.609** | 5.24E-05 | **1.84E-04** |
| ENSG00000290837.2 | **LRRC37A5I** | 652972 | lncRNA | **17.739** | 1.461 | 0.440 | **2.754** | 5.25E-05 | **1.84E-04** |
| ENSG00000164209.19 | **SLC25A46** | 91137 | protein_coding | **474.088** | -0.552 | -0.166 | **0.682** | 5.26E-05 | **1.84E-04** |
| ENSG00000128654.16 | **MTX2** | 10651 | protein_coding | **676.919** | -0.422 | -0.127 | **0.746** | 5.28E-05 | **1.85E-04** |
| ENSG00000267074.1 | **ENSG00000267074** | | lncRNA | **17.803** | -1.720 | -0.518 | **0.304** | 5.29E-05 | **1.85E-04** |
| ENSG00000246863.3 | **GPR176-D1** | 10050553 | lncRNA | **6.232** | 3.352 | 1.009 | **10.208** | 5.30E-05 | **1.85E-04** |
| ENSG00000172046.22 | **USP19** | 10869 | protein_coding | **1098.881** | 0.395 | 0.119 | **1.315** | 5.30E-05 | **1.86E-04** |
| ENSG00000085978.24 | **ATG16L1** | 55054 | protein_coding | **450.392** | 0.522 | 0.157 | **1.436** | 5.31E-05 | **1.86E-04** |
| ENSG00000171208.12 | **NETO2** | 81831 | protein_coding | **303.922** | -0.736 | -0.221 | **0.601** | 5.33E-05 | **1.86E-04** |
| ENSG00000290706.2 | **FAM238C** | 387644 | lncRNA | **51.876** | 1.118 | 0.336 | **2.170** | 5.34E-05 | **1.87E-04** |
| ENSG00000261008.10 | **LINC01572** | 10192795 | lncRNA | **40.322** | -1.140 | -0.343 | **0.454** | 5.34E-05 | **1.87E-04** |
| ENSG00000150093.22 | **ITGB1** | 3688 | protein_coding | **669.401** | 0.422 | 0.127 | **1.340** | 5.35E-05 | **1.87E-04** |
| ENSG00000048140.20 | **TSPAN17** | 26262 | protein_coding | **318.094** | 0.491 | 0.148 | **1.405** | 5.35E-05 | **1.87E-04** |
| ENSG00000168906.15 | **MAT2A** | 4144 | protein_coding | **2033.849** | -0.742 | -0.223 | **0.598** | 5.37E-05 | **1.88E-04** |
| ENSG00000287005.1 | **ENSG00000287005** | | lncRNA | **6.954** | 2.912 | 0.877 | **7.526** | 5.38E-05 | **1.88E-04** |
| ENSG00000178053.22 | **MLF1** | 4291 | protein_coding | **42.488** | 1.273 | 0.383 | **2.416** | 5.38E-05 | **1.88E-04** |
| ENSG00000128590.7 | **DNAJB9** | 4189 | protein_coding | **151.456** | -0.771 | -0.232 | **0.586** | 5.38E-05 | **1.88E-04** |
| ENSG00000303863.1 | **ENSG00000303863** | | lncRNA | **87.744** | 1.061 | 0.319 | **2.086** | 5.39E-05 | **1.88E-04** |
| ENSG00000181264.11 | **TLCD5** | 219902 | protein_coding | **41.612** | 1.430 | 0.430 | **2.694** | 5.40E-05 | **1.88E-04** |
| ENSG00000290848.2 | **ENSG00000290848** | | lncRNA | **18.941** | 1.480 | 0.446 | **2.790** | 5.41E-05 | **1.89E-04** |
| ENSG00000117751.20 | **PPP1R8** | 5511 | protein_coding | **611.271** | -0.536 | -0.161 | **0.690** | 5.42E-05 | **1.89E-04** |

| | | | | | | | | | |
|---|---|---|---|---|---|---|---|---|---|
| ENSG00000300150.1 | **ENSG00000300150** | | lncRNA | **7.330** | 3.281 | 0.988 | **9.718** | 5.44E-05 | **1.90E-04** |
| ENSG00000102878.20 | **HSF4** | 3299 | protein_coding | **790.335** | 0.688 | 0.207 | **1.611** | 5.44E-05 | **1.90E-04** |
| ENSG00000170275.17 | **CRTAP** | 10491 | protein_coding | **791.611** | -0.456 | -0.137 | **0.729** | 5.44E-05 | **1.90E-04** |
| ENSG00000137269.17 | **LRRC1** | 55227 | protein_coding | **101.161** | 1.065 | 0.321 | **2.093** | 5.45E-05 | **1.90E-04** |
| ENSG00000075275.19 | **CELSR1** | 9620 | protein_coding | **64.870** | 0.751 | 0.226 | **1.683** | 5.46E-05 | **1.90E-04** |
| ENSG00000025770.21 | **NCAPH2** | 29781 | protein_coding | **1231.744** | -0.550 | -0.166 | **0.683** | 5.47E-05 | **1.91E-04** |
| ENSG00000139187.11 | **KLRG1** | 10219 | protein_coding | **32.056** | 1.313 | 0.395 | **2.484** | 5.48E-05 | **1.91E-04** |
| ENSG00000163995.23 | **ABLIM2** | 84448 | protein_coding | **16.396** | 1.991 | 0.599 | **3.976** | 5.49E-05 | **1.91E-04** |
| ENSG00000126814.9 | **TRMT5** | 57570 | protein_coding | **267.326** | -0.491 | -0.148 | **0.712** | 5.50E-05 | **1.92E-04** |
| ENSG00000104472.12 | **CHRAC1** | 54108 | protein_coding | **587.728** | -0.616 | -0.185 | **0.652** | 5.51E-05 | **1.92E-04** |
| ENSG00000112981.7 | **NME5** | 8382 | protein_coding | **8.957** | 2.904 | 0.874 | **7.484** | 5.52E-05 | **1.92E-04** |
| ENSG00000196074.14 | **SYCP2** | 10388 | protein_coding | **95.894** | 1.480 | 0.446 | **2.790** | 5.52E-05 | **1.92E-04** |
| ENSG00000044574.11 | **HSPA5** | 3309 | protein_coding | **7313.873** | -0.582 | -0.175 | **0.668** | 5.55E-05 | **1.93E-04** |
| ENSG00000231056.2 | **LINC02522** | | lncRNA | **3.413** | 4.176 | 1.257 | **18.081** | 5.56E-05 | **1.93E-04** |
| ENSG00000102547.21 | **CAB39L** | 81617 | protein_coding | **83.745** | -0.709 | -0.213 | **0.612** | 5.60E-05 | **1.95E-04** |
| ENSG00000100298.17 | **APOBEC3H** | 164668 | protein_coding | **295.368** | 0.667 | 0.201 | **1.588** | 5.61E-05 | **1.95E-04** |
| ENSG00000153832.14 | **FBXO36** | 130888 | protein_coding | **27.271** | 1.110 | 0.334 | **2.158** | 5.62E-05 | **1.96E-04** |
| ENSG00000288638.1 | **LNCDAT** | | lncRNA | **114.803** | 1.363 | 0.410 | **2.572** | 5.62E-05 | **1.96E-04** |
| ENSG00000087237.14 | **CETP** | 1071 | protein_coding | **126.471** | 0.997 | 0.300 | **1.996** | 5.63E-05 | **1.96E-04** |
| ENSG00000149476.18 | **TKFC** | 26007 | protein_coding | **1246.430** | 0.434 | 0.131 | **1.351** | 5.64E-05 | **1.96E-04** |
| ENSG00000174238.17 | **PITPNA** | 5306 | protein_coding | **1448.822** | 0.570 | 0.172 | **1.484** | 5.67E-05 | **1.97E-04** |
| ENSG00000284914.2 | **ENSG00000284914** | | lncRNA | **10.917** | 2.215 | 0.667 | **4.643** | 5.67E-05 | **1.97E-04** |
| ENSG00000147255.21 | **IGSF1** | 3547 | protein_coding | **105.823** | 2.257 | 0.679 | **4.780** | 5.68E-05 | **1.98E-04** |
| ENSG00000070018.11 | **LRP6** | 4040 | protein_coding | **154.545** | 1.087 | 0.327 | **2.125** | 5.70E-05 | **1.98E-04** |
| ENSG00000280187.2 | **ENSG00000280187** | | TEC | **24.818** | 2.321 | 0.699 | **4.996** | 5.70E-05 | **1.98E-04** |
| ENSG00000101557.17 | **USP14** | 9097 | protein_coding | **896.852** | -0.531 | -0.160 | **0.692** | 5.72E-05 | **1.99E-04** |
| ENSG00000243244.9 | **STON1** | 11037 | protein_coding | **59.724** | 1.005 | 0.303 | **2.007** | 5.75E-05 | **2.00E-04** |
| ENSG00000131732.14 | **ZCCHC9** | 84240 | protein_coding | **415.432** | -0.445 | -0.134 | **0.734** | 5.75E-05 | **2.00E-04** |
| ENSG00000232442.4 | **MHENCR** | 100505771 | lncRNA | **1072.068** | 0.661 | 0.199 | **1.581** | 5.78E-05 | **2.01E-04** |
| ENSG00000146425.13 | **DYNLT1** | 6993 | protein_coding | **888.558** | 0.352 | 0.106 | **1.277** | 5.78E-05 | **2.01E-04** |
| ENSG00000109927.11 | **TECTA** | 7007 | protein_coding | **23.601** | 1.896 | 0.571 | **3.721** | 5.79E-05 | **2.01E-04** |

| | | | | | | | | | |
|---|---|---|---|---|---|---|---|---|---|
| ENSG00000150782.15 | **IL18** | 3606 | protein_coding | **28.980** | 1.315 | 0.396 | **2.488** | 5.79E-05 | **2.01E-04** |
| ENSG00000174749.7 | **FAM241A** | 132720 | protein_coding | **184.473** | 0.662 | 0.199 | **1.582** | 5.79E-05 | **2.01E-04** |
| ENSG00000164002.14 | **EXO5** | 64789 | protein_coding | **155.487** | -0.597 | -0.180 | **0.661** | 5.79E-05 | **2.01E-04** |
| ENSG00000183569.20 | **SERHL2** | 253190 | protein_coding | **54.497** | 1.350 | 0.406 | **2.548** | 5.80E-05 | **2.01E-04** |
| ENSG00000125871.16 | **MGME1** | 92667 | protein_coding | **677.291** | -0.461 | -0.139 | **0.727** | 5.81E-05 | **2.02E-04** |
| ENSG00000278869.3 | **ENSG00000278869** | | lncRNA | **25.920** | 1.229 | 0.370 | **2.344** | 5.84E-05 | **2.02E-04** |
| ENSG00000227017.2 | **ENSG00000227017** | | lncRNA | **17.995** | 1.434 | 0.432 | **2.702** | 5.85E-05 | **2.03E-04** |
| ENSG00000172350.12 | **ABCG4** | 64137 | protein_coding | **6.394** | 2.885 | 0.868 | **7.386** | 5.88E-05 | **2.04E-04** |
| ENSG00000305109.1 | **ENSG00000305109** | | lncRNA | **19.012** | 1.357 | 0.408 | **2.561** | 5.89E-05 | **2.04E-04** |
| ENSG00000125107.21 | **CNOT1** | 23019 | protein_coding | **2661.096** | -0.495 | -0.149 | **0.710** | 5.91E-05 | **2.05E-04** |
| ENSG00000149054.18 | **ZNF215** | 7762 | protein_coding | **108.484** | -0.682 | -0.205 | **0.623** | 5.93E-05 | **2.06E-04** |
| ENSG00000100403.14 | **ZC3H7B** | 23264 | protein_coding | **978.730** | -0.486 | -0.146 | **0.714** | 5.94E-05 | **2.06E-04** |
| ENSG00000235034.8 | **C19orf81** | 342918 | protein_coding | **5.765** | 3.162 | 0.952 | **8.952** | 5.94E-05 | **2.06E-04** |
| ENSG00000100721.12 | **TCL1A** | 8115 | protein_coding | **2977.677** | -0.588 | -0.177 | **0.665** | 5.95E-05 | **2.06E-04** |
| ENSG00000124831.22 | **LRRFIP1** | 9208 | protein_coding | **1886.670** | -0.443 | -0.133 | **0.736** | 5.96E-05 | **2.06E-04** |
| ENSG00000166734.22 | **GOLM2** | 113201 | protein_coding | **641.828** | 0.411 | 0.124 | **1.330** | 5.96E-05 | **2.06E-04** |
| ENSG00000165915.16 | **SLC39A13** | 91252 | protein_coding | **492.254** | 0.461 | 0.139 | **1.376** | 5.96E-05 | **2.06E-04** |
| ENSG00000073067.15 | **CYP2W1** | 54905 | protein_coding | **6.420** | 3.185 | 0.959 | **9.092** | 5.98E-05 | **2.07E-04** |
| ENSG00000237298.12 | **TTN-AS1** | 10050686€ | lncRNA | **272.137** | 0.669 | 0.201 | **1.590** | 5.99E-05 | **2.07E-04** |
| ENSG00000135750.17 | **KCNK1** | 3775 | protein_coding | **110.353** | -0.838 | -0.252 | **0.559** | 5.99E-05 | **2.07E-04** |
| ENSG00000117118.14 | **SDHB** | 6390 | protein_coding | **1107.556** | -0.453 | -0.136 | **0.731** | 6.00E-05 | **2.08E-04** |
| ENSG00000164934.17 | **DCAF13** | 25879 | protein_coding | **684.217** | -0.487 | -0.147 | **0.713** | 6.03E-05 | **2.09E-04** |
| ENSG00000258744.4 | **ENSG00000258744** | | lncRNA | **61.916** | 0.728 | 0.219 | **1.657** | 6.04E-05 | **2.09E-04** |
| ENSG00000092094.13 | **OSGEP** | 55644 | protein_coding | **1128.571** | 0.350 | 0.105 | **1.274** | 6.05E-05 | **2.09E-04** |
| ENSG00000154978.15 | **VOPP1** | 81552 | protein_coding | **4439.455** | 0.543 | 0.163 | **1.457** | 6.07E-05 | **2.10E-04** |
| ENSG00000283515.1 | **ENSG00000283515** | | protein_coding | **143.738** | -0.938 | -0.282 | **0.522** | 6.09E-05 | **2.10E-04** |
| ENSG00000135336.19 | **ORC3** | 23595 | protein_coding | **568.986** | -0.434 | -0.131 | **0.740** | 6.11E-05 | **2.11E-04** |
| ENSG00000125351.16 | **UPF3B** | 65109 | protein_coding | **528.607** | -0.558 | -0.168 | **0.679** | 6.11E-05 | **2.11E-04** |
| ENSG00000151474.24 | **FRMD4A** | 55691 | protein_coding | **366.885** | -0.906 | -0.273 | **0.534** | 6.16E-05 | **2.13E-04** |
| ENSG00000137168.10 | **PPIL1** | 51645 | protein_coding | **426.437** | -0.532 | -0.160 | **0.692** | 6.18E-05 | **2.13E-04** |
| ENSG00000243709.3 | **LEFTY1** | 10637 | protein_coding | **4.480** | 3.727 | 1.122 | **13.238** | 6.23E-05 | **2.15E-04** |

| | | | | | | | | | |
|---|---|---|---|---|---|---|---|---|---|
| ENSG00000197165.13 | **SULT1A2** | 6799 | protein_coding | **6.439** | 2.825 | 0.850 | **7.085** | 6.24E-05 | **2.15E-04** |
| ENSG00000072756.20 | **TRNT1** | 51095 | protein_coding | **426.682** | -0.410 | -0.123 | **0.753** | 6.25E-05 | **2.16E-04** |
| ENSG00000239282.10 | **CASTOR1** | 652968 | protein_coding | **193.667** | 1.024 | 0.308 | **2.034** | 6.26E-05 | **2.16E-04** |
| ENSG00000113761.14 | **ZNF346** | 23567 | protein_coding | **309.522** | -0.449 | -0.135 | **0.732** | 6.28E-05 | **2.16E-04** |
| ENSG00000261359.3 | **PYCARD-A!** | 10065274( | lncRNA | **37.466** | 1.139 | 0.343 | **2.202** | 6.29E-05 | **2.17E-04** |
| ENSG00000100024.17 | **UPB1** | 51733 | protein_coding | **237.241** | 0.839 | 0.253 | **1.789** | 6.30E-05 | **2.17E-04** |
| ENSG00000253853.4 | **ENSG00000253853** | | lncRNA | **19.807** | -1.629 | -0.490 | **0.323** | 6.30E-05 | **2.17E-04** |
| ENSG00000109390.15 | **NDUFC1** | 4717 | protein_coding | **457.920** | -0.550 | -0.166 | **0.683** | 6.31E-05 | **2.17E-04** |
| ENSG00000277938.1 | **ENSG00000277938** | | lncRNA | **23.589** | 1.309 | 0.394 | **2.477** | 6.31E-05 | **2.17E-04** |
| ENSG00000300386.1 | **ENSG00000300386** | | lncRNA | **7.503** | 2.730 | 0.822 | **6.635** | 6.34E-05 | **2.18E-04** |
| ENSG00000101574.17 | **METTL4** | 64863 | protein_coding | **140.597** | -0.742 | -0.223 | **0.598** | 6.34E-05 | **2.18E-04** |
| ENSG00000139163.18 | **ETNK1** | 55500 | protein_coding | **572.696** | -0.463 | -0.139 | **0.726** | 6.38E-05 | **2.20E-04** |
| ENSG00000163728.13 | **TTC14** | 151613 | protein_coding | **601.420** | 0.519 | 0.156 | **1.433** | 6.39E-05 | **2.20E-04** |
| ENSG00000088247.20 | **KHSRP** | 8570 | protein_coding | **4240.178** | -0.332 | -0.100 | **0.795** | 6.40E-05 | **2.20E-04** |
| ENSG00000116668.15 | **SWT1** | 54823 | protein_coding | **99.839** | 0.793 | 0.239 | **1.733** | 6.40E-05 | **2.20E-04** |
| ENSG00000237172.6 | **B3GNT9** | 84752 | protein_coding | **196.031** | 0.531 | 0.160 | **1.444** | 6.41E-05 | **2.20E-04** |
| ENSG00000144118.16 | **RALB** | 5899 | protein_coding | **976.273** | -0.407 | -0.123 | **0.754** | 6.41E-05 | **2.21E-04** |
| ENSG00000168214.24 | **RBPJ** | 3516 | protein_coding | **1050.791** | -0.435 | -0.131 | **0.740** | 6.41E-05 | **2.21E-04** |
| ENSG00000179841.9 | **AKAP5** | 9495 | protein_coding | **38.470** | -0.908 | -0.273 | **0.533** | 6.43E-05 | **2.21E-04** |
| ENSG00000185267.12 | **CDNF** | 441549 | protein_coding | **35.973** | 1.355 | 0.408 | **2.558** | 6.44E-05 | **2.22E-04** |
| ENSG00000126091.23 | **ST3GAL3** | 6487 | protein_coding | **214.492** | -0.644 | -0.194 | **0.640** | 6.45E-05 | **2.22E-04** |
| ENSG00000101347.13 | **SAMHD1** | 25939 | protein_coding | **503.781** | 0.541 | 0.163 | **1.455** | 6.46E-05 | **2.22E-04** |
| ENSG00000166261.13 | **ZNF202** | 7753 | protein_coding | **319.065** | -0.579 | -0.174 | **0.669** | 6.47E-05 | **2.22E-04** |
| ENSG00000297423.1 | **ENSG00000297423** | | lncRNA | **63.171** | 1.084 | 0.326 | **2.120** | 6.47E-05 | **2.22E-04** |
| ENSG00000280587.4 | **LINC01348** | 731656 | lncRNA | **19.336** | 1.560 | 0.470 | **2.948** | 6.48E-05 | **2.23E-04** |
| ENSG00000168675.21 | **LDLRAD4** | 753 | protein_coding | **934.090** | -0.544 | -0.164 | **0.686** | 6.50E-05 | **2.23E-04** |
| ENSG00000056050.9 | **HPF1** | 54969 | protein_coding | **275.796** | -0.493 | -0.148 | **0.711** | 6.51E-05 | **2.24E-04** |
| ENSG00000165609.16 | **NUDT5** | 11164 | protein_coding | **1481.762** | -0.429 | -0.129 | **0.743** | 6.51E-05 | **2.24E-04** |
| ENSG00000177788.7 | **RAB4A-AS:** | 10537315( | lncRNA | **42.075** | 1.036 | 0.312 | **2.051** | 6.53E-05 | **2.24E-04** |
| ENSG00000133246.14 | **PRAM1** | 84106 | protein_coding | **16.584** | 1.830 | 0.551 | **3.556** | 6.55E-05 | **2.25E-04** |
| ENSG00000291143.2 | **ENSG00000291143** | | lncRNA | **301.724** | -0.629 | -0.189 | **0.647** | 6.56E-05 | **2.25E-04** |

| | | | | | | | | | |
|---|---|---|---|---|---|---|---|---|---|
| ENSG00000163932.18 | **PRKCD** | 5580 | protein_coding | **3021.987** | 0.343 | 0.103 | **1.269** | 6.59E-05 | **2.26E-04** |
| ENSG00000134780.12 | **DAGLA** | 747 | protein_coding | **5.744** | 3.102 | 0.934 | **8.585** | 6.60E-05 | **2.27E-04** |
| ENSG00000233785.2 | **SAT1-DT** | 10537314ε | lncRNA | **17.792** | 1.655 | 0.498 | **3.149** | 6.62E-05 | **2.27E-04** |
| ENSG00000119915.6 | **ELOVL3** | 83401 | protein_coding | **6.423** | 3.213 | 0.967 | **9.270** | 6.62E-05 | **2.27E-04** |
| ENSG00000107771.19 | **CCSER2** | 54462 | protein_coding | **218.989** | 0.614 | 0.185 | **1.530** | 6.62E-05 | **2.27E-04** |
| ENSG00000133393.15 | **CEP20** | 123811 | protein_coding | **440.388** | -0.403 | -0.121 | **0.756** | 6.64E-05 | **2.28E-04** |
| ENSG00000267737.2 | **ENSG00000267737** | | lncRNA | **4.010** | 3.795 | 1.142 | **13.879** | 6.66E-05 | **2.28E-04** |
| ENSG00000272153.1 | **ENSG00000272153** | | lncRNA | **5.595** | 2.846 | 0.857 | **7.192** | 6.70E-05 | **2.30E-04** |
| ENSG00000166997.10 | **CNPY4** | 245812 | protein_coding | **150.886** | 0.516 | 0.155 | **1.430** | 6.71E-05 | **2.30E-04** |
| ENSG00000160551.14 | **TAOK1** | 57551 | protein_coding | **414.649** | -0.509 | -0.153 | **0.703** | 6.72E-05 | **2.30E-04** |
| ENSG00000235385.4 | **LINC02154** | 10972916¢ | lncRNA | **17.095** | 1.923 | 0.579 | **3.791** | 6.73E-05 | **2.30E-04** |
| ENSG00000180815.17 | **MAP3K15** | 389840 | protein_coding | **27.840** | 1.242 | 0.374 | **2.365** | 6.73E-05 | **2.31E-04** |
| ENSG00000166922.11 | **SCG5** | 6447 | protein_coding | **26.974** | -1.429 | -0.430 | **0.371** | 6.79E-05 | **2.32E-04** |
| ENSG00000132570.17 | **PCBD2** | 84105 | protein_coding | **147.012** | -0.756 | -0.228 | **0.592** | 6.80E-05 | **2.33E-04** |
| ENSG00000249437.10 | **NAIP** | 4671 | protein_coding | **524.464** | 0.539 | 0.162 | **1.453** | 6.82E-05 | **2.33E-04** |
| ENSG00000271040.2 | **TMEM242-DT** | | lncRNA | **6.388** | 3.007 | 0.905 | **8.038** | 6.83E-05 | **2.34E-04** |
| ENSG00000077463.17 | **SIRT6** | 51548 | protein_coding | **834.742** | 0.449 | 0.135 | **1.366** | 6.83E-05 | **2.34E-04** |
| ENSG00000159403.20 | **C1R** | 715 | protein_coding | **141.982** | 0.766 | 0.231 | **1.700** | 6.84E-05 | **2.34E-04** |
| ENSG00000226711.10 | **FAM66C** | | lncRNA | **27.907** | 1.334 | 0.402 | **2.521** | 6.86E-05 | **2.35E-04** |
| ENSG00000188234.17 | **AGAP4** | 119016 | protein_coding | **579.786** | 0.419 | 0.126 | **1.337** | 6.86E-05 | **2.35E-04** |
| ENSG00000107672.17 | **NSMCE4A** | 54780 | protein_coding | **728.190** | -0.391 | -0.118 | **0.763** | 6.88E-05 | **2.35E-04** |
| ENSG00000303651.1 | **ENSG00000303651** | | lncRNA | **8.015** | 2.839 | 0.855 | **7.158** | 6.88E-05 | **2.35E-04** |
| ENSG00000082641.18 | **NFE2L1** | 4779 | protein_coding | **2977.444** | -0.634 | -0.191 | **0.644** | 6.89E-05 | **2.35E-04** |
| ENSG00000258947.9 | **TUBB3** | 10381 | protein_coding | **44.131** | 1.042 | 0.314 | **2.059** | 6.89E-05 | **2.36E-04** |
| ENSG00000145362.23 | **ANK2** | 287 | protein_coding | **130.205** | -0.799 | -0.240 | **0.575** | 6.92E-05 | **2.36E-04** |
| ENSG00000248839.2 | **CLDND1-A!** | 13935520€ | lncRNA | **7.861** | 3.375 | 1.016 | **10.376** | 6.93E-05 | **2.37E-04** |
| ENSG00000278318.7 | **ZNF229** | 7772 | protein_coding | **31.220** | 1.559 | 0.469 | **2.946** | 6.94E-05 | **2.37E-04** |
| ENSG00000120333.6 | **MRPS14** | 63931 | protein_coding | **407.098** | -0.545 | -0.164 | **0.685** | 6.95E-05 | **2.37E-04** |
| ENSG00000112033.16 | **PPARD** | 5467 | protein_coding | **699.355** | 0.479 | 0.144 | **1.394** | 7.00E-05 | **2.39E-04** |
| ENSG00000306172.1 | **ENSG0000(** | 12490102€ | lncRNA | **4.855** | 3.996 | 1.203 | **15.959** | 7.00E-05 | **2.39E-04** |
| ENSG00000197782.17 | **ZNF780A** | 284323 | protein_coding | **351.439** | -0.646 | -0.195 | **0.639** | 7.01E-05 | **2.39E-04** |

| | | | | | | | | | |
|---|---|---|---|---|---|---|---|---|---|
| ENSG00000169813.19 | **HNRNPF** | 3185 | protein_coding | **3716.627** | -0.511 | -0.154 | **0.702** | 7.04E-05 | **2.40E-04** |
| ENSG00000142046.17 | **TMEM91** | 641649 | protein_coding | **295.111** | 0.716 | 0.215 | **1.642** | 7.06E-05 | **2.41E-04** |
| ENSG00000155858.7 | **LSM11** | 134353 | protein_coding | **69.339** | -0.728 | -0.219 | **0.604** | 7.07E-05 | **2.41E-04** |
| ENSG00000289589.3 | **ENSG00000289589** | | lncRNA | **15.098** | 1.469 | 0.442 | **2.768** | 7.09E-05 | **2.42E-04** |
| ENSG00000089195.17 | **TRMT6** | 51605 | protein_coding | **241.792** | -0.457 | -0.138 | **0.728** | 7.09E-05 | **2.42E-04** |
| ENSG00000297504.1 | **ENSG00000297504** | | lncRNA | **4.038** | 3.915 | 1.178 | **15.083** | 7.10E-05 | **2.42E-04** |
| ENSG00000186952.17 | **TMEM232** | 642987 | protein_coding | **14.206** | 1.781 | 0.536 | **3.437** | 7.10E-05 | **2.42E-04** |
| ENSG00000258940.3 | **MIA2-AS1** | 10028884( | lncRNA | **16.965** | 1.557 | 0.469 | **2.942** | 7.10E-05 | **2.42E-04** |
| ENSG00000138138.16 | **ATAD1** | 84896 | protein_coding | **898.912** | -0.465 | -0.140 | **0.724** | 7.11E-05 | **2.42E-04** |
| ENSG00000173578.10 | **XCR1** | 2829 | protein_coding | **8.212** | 3.282 | 0.988 | **9.726** | 7.12E-05 | **2.42E-04** |
| ENSG00000119720.20 | **NRDE2** | 55051 | protein_coding | **197.492** | -0.550 | -0.166 | **0.683** | 7.12E-05 | **2.43E-04** |
| ENSG00000100243.24 | **CYB5R3** | 1727 | protein_coding | **2585.521** | 0.414 | 0.125 | **1.332** | 7.14E-05 | **2.43E-04** |
| ENSG00000235280.2 | **MCF2L-AS1** | | lncRNA | **7.840** | 3.278 | 0.987 | **9.702** | 7.16E-05 | **2.44E-04** |
| ENSG00000112983.19 | **BRD8** | 10902 | protein_coding | **521.005** | -0.395 | -0.119 | **0.761** | 7.19E-05 | **2.45E-04** |
| ENSG00000287166.2 | **ENSG00000287166** | | lncRNA | **6.459** | 2.682 | 0.807 | **6.416** | 7.21E-05 | **2.45E-04** |
| ENSG00000293175.2 | **ENSG00000293175** | | lncRNA | **24.839** | 1.309 | 0.394 | **2.479** | 7.21E-05 | **2.45E-04** |
| ENSG00000198796.9 | **ALPK2** | 115701 | protein_coding | **398.623** | 1.048 | 0.316 | **2.068** | 7.22E-05 | **2.46E-04** |
| ENSG00000231527.8 | **FAM27C** | 10013294ε | lncRNA | **73.052** | -1.219 | -0.367 | **0.430** | 7.23E-05 | **2.46E-04** |
| ENSG00000272121.2 | **ENSG00000272121** | | lncRNA | **17.895** | 1.787 | 0.538 | **3.451** | 7.25E-05 | **2.46E-04** |
| ENSG00000257642.1 | **C12orf75-A** | 10536995Z | lncRNA | **8.985** | -2.243 | -0.675 | **0.211** | 7.26E-05 | **2.47E-04** |
| ENSG00000176953.15 | **NFATC2IP** | 84901 | protein_coding | **1468.647** | -0.439 | -0.132 | **0.738** | 7.27E-05 | **2.47E-04** |
| ENSG00000289440.2 | **ENSG00000289440** | | lncRNA | **38.046** | 0.977 | 0.294 | **1.968** | 7.28E-05 | **2.48E-04** |
| ENSG00000215481.10 | **BCRP3** | 644165 | lncRNA | **21.470** | 1.651 | 0.497 | **3.140** | 7.29E-05 | **2.48E-04** |
| ENSG00000275857.2 | **ENSG00000275857** | | lncRNA | **19.150** | 1.345 | 0.405 | **2.539** | 7.32E-05 | **2.49E-04** |
| ENSG00000197548.15 | **ATG7** | 10533 | protein_coding | **260.584** | -0.523 | -0.157 | **0.696** | 7.33E-05 | **2.49E-04** |
| ENSG00000081320.14 | **STK17B** | 9262 | protein_coding | **1025.303** | 0.505 | 0.152 | **1.419** | 7.33E-05 | **2.49E-04** |
| ENSG00000274461.3 | **LINC02930** | 10537851( | lncRNA | **8.411** | 3.350 | 1.008 | **10.195** | 7.35E-05 | **2.49E-04** |
| ENSG00000189152.13 | **GRAPL** | 400581 | protein_coding | **269.985** | -0.540 | -0.162 | **0.688** | 7.36E-05 | **2.50E-04** |
| ENSG00000230487.10 | **PSMG3-AS** | 114796 | lncRNA | **161.110** | 0.614 | 0.185 | **1.531** | 7.36E-05 | **2.50E-04** |
| ENSG00000224578.5 | **HNRNPA1I** | 642659 | protein_coding | **53.003** | -1.063 | -0.320 | **0.479** | 7.40E-05 | **2.51E-04** |
| ENSG00000197958.15 | **RPL12** | 6136 | protein_coding | **43896.791** | -0.431 | -0.130 | **0.742** | 7.44E-05 | **2.53E-04** |

| | | | | | | | | | |
|---|---|---|---|---|---|---|---|---|---|
| ENSG00000291106.2 | **SNHG33** | 100505687 | lncRNA | **30.176** | 1.038 | 0.313 | **2.054** | 7.46E-05 | **2.53E-04** |
| ENSG00000158806.16 | **NPM2** | 10361 | protein_coding | **154.359** | 0.700 | 0.211 | **1.625** | 7.49E-05 | **2.54E-04** |
| ENSG00000304417.1 | **ENSG00000304417** | | lncRNA | **5.112** | 3.376 | 1.016 | **10.381** | 7.50E-05 | **2.54E-04** |
| ENSG00000143384.15 | **MCL1** | 4170 | protein_coding | **4232.868** | 0.472 | 0.142 | **1.387** | 7.50E-05 | **2.55E-04** |
| ENSG00000162032.18 | **SPSB3** | 90864 | protein_coding | **822.721** | 0.446 | 0.134 | **1.363** | 7.51E-05 | **2.55E-04** |
| ENSG00000134077.18 | **THUMPD3** | 25917 | protein_coding | **446.843** | -0.606 | -0.182 | **0.657** | 7.52E-05 | **2.55E-04** |
| ENSG00000101290.16 | **CDS2** | 8760 | protein_coding | **826.243** | -0.414 | -0.124 | **0.751** | 7.52E-05 | **2.55E-04** |
| ENSG00000226149.7 | **ENSG00000226149** | | lncRNA | **13.626** | 1.968 | 0.592 | **3.912** | 7.52E-05 | **2.55E-04** |
| ENSG00000067066.19 | **SP100** | 6672 | protein_coding | **4182.679** | 0.363 | 0.109 | **1.286** | 7.55E-05 | **2.56E-04** |
| ENSG00000124151.21 | **NCOA3** | 8202 | protein_coding | **3054.332** | 0.517 | 0.156 | **1.431** | 7.55E-05 | **2.56E-04** |
| ENSG00000289207.2 | **ENSG00000289207** | | lncRNA | **12.687** | 2.194 | 0.661 | **4.577** | 7.55E-05 | **2.56E-04** |
| ENSG00000223695.2 | **MYH9-DT** | | lncRNA | **9.572** | 2.137 | 0.643 | **4.399** | 7.57E-05 | **2.57E-04** |
| ENSG00000169991.12 | **IFFO2** | 126917 | protein_coding | **565.024** | 0.482 | 0.145 | **1.396** | 7.63E-05 | **2.58E-04** |
| ENSG00000038210.16 | **PI4K2B** | 55300 | protein_coding | **773.473** | -0.576 | -0.173 | **0.671** | 7.65E-05 | **2.59E-04** |
| ENSG00000156804.9 | **FBXO32** | 114907 | protein_coding | **89.851** | 0.717 | 0.216 | **1.644** | 7.67E-05 | **2.60E-04** |
| ENSG00000173226.19 | **IQCB1** | 9657 | protein_coding | **1017.389** | -0.437 | -0.131 | **0.739** | 7.68E-05 | **2.60E-04** |
| ENSG00000293494.3 | **ZPLD2P** | 101928303 | lncRNA | **5.557** | 3.134 | 0.943 | **8.778** | 7.72E-05 | **2.61E-04** |
| ENSG00000002586.22 | **CD99** | 4267 | protein_coding | **2043.471** | -0.489 | -0.147 | **0.712** | 7.73E-05 | **2.62E-04** |
| ENSG00000236618.4 | **PITPNA-AS1** | | lncRNA | **176.175** | 0.508 | 0.153 | **1.422** | 7.77E-05 | **2.63E-04** |
| ENSG00000101977.23 | **MCF2** | 4168 | protein_coding | **6.628** | -2.986 | -0.899 | **0.126** | 7.78E-05 | **2.63E-04** |
| ENSG00000182117.8 | **NOP10** | 55505 | protein_coding | **1886.632** | -0.412 | -0.124 | **0.752** | 7.79E-05 | **2.63E-04** |
| ENSG00000230155.9 | **CEP250-AS1** | | lncRNA | **60.043** | -1.099 | -0.331 | **0.467** | 7.81E-05 | **2.64E-04** |
| ENSG00000287769.2 | **ENSG0000(** | 12490220[illegible] | lncRNA | **8.072** | 2.397 | 0.722 | **5.269** | 7.82E-05 | **2.64E-04** |
| ENSG00000117407.19 | **ARTN** | 9048 | protein_coding | **42.937** | -1.508 | -0.454 | **0.352** | 7.82E-05 | **2.64E-04** |
| ENSG00000125384.9 | **PTGER2** | 5732 | protein_coding | **13.956** | 1.957 | 0.589 | **3.884** | 7.83E-05 | **2.64E-04** |
| ENSG00000149136.11 | **SSRP1** | 6749 | protein_coding | **2417.587** | -0.525 | -0.158 | **0.695** | 7.84E-05 | **2.65E-04** |
| ENSG00000286431.2 | **ENSG00000286431** | | lncRNA | **18.531** | 1.453 | 0.437 | **2.738** | 7.85E-05 | **2.65E-04** |
| ENSG00000087470.23 | **DNM1L** | 10059 | protein_coding | **911.199** | -0.357 | -0.107 | **0.781** | 7.85E-05 | **2.65E-04** |
| ENSG00000090520.14 | **DNAJB11** | 51726 | protein_coding | **1338.719** | -0.527 | -0.159 | **0.694** | 7.85E-05 | **2.65E-04** |
| ENSG00000196756.16 | **SNHG17** | 388796 | lncRNA | **848.557** | -0.824 | -0.248 | **0.565** | 7.87E-05 | **2.66E-04** |
| ENSG00000263731.1 | **LINC03149** | | lncRNA | **66.895** | -0.779 | -0.234 | **0.583** | 7.89E-05 | **2.66E-04** |

| | | | | | | | | | |
|---|---|---|---|---|---|---|---|---|---|
| ENSG00000140367.14 | **UBE2Q2** | 92912 | protein_coding | **137.037** | -0.727 | -0.219 | **0.604** | 7.89E-05 | **2.66E-04** |
| ENSG00000139579.15 | **NABP2** | 79035 | protein_coding | **587.036** | -0.320 | -0.096 | **0.801** | 7.92E-05 | **2.67E-04** |
| ENSG00000155368.20 | **DBI** | 1622 | protein_coding | **4762.984** | -0.513 | -0.154 | **0.701** | 7.93E-05 | **2.67E-04** |
| ENSG00000287031.2 | **ENSG00000287031** | | lncRNA | **15.254** | 1.527 | 0.460 | **2.883** | 8.00E-05 | **2.70E-04** |
| ENSG00000132530.19 | **XAF1** | 54739 | protein_coding | **2497.323** | -0.772 | -0.232 | **0.586** | 8.00E-05 | **2.70E-04** |
| ENSG00000164134.16 | **NAA15** | 80155 | protein_coding | **446.497** | -0.753 | -0.227 | **0.594** | 8.01E-05 | **2.70E-04** |
| ENSG00000169442.10 | **CD52** | 1043 | protein_coding | **6197.402** | -0.695 | -0.209 | **0.618** | 8.03E-05 | **2.71E-04** |
| ENSG00000246582.4 | **TNFRSF10/** | 389641 | lncRNA | **23.708** | 1.304 | 0.392 | **2.469** | 8.04E-05 | **2.71E-04** |
| ENSG00000235173.8 | **HGH1** | 51236 | protein_coding | **509.458** | -0.414 | -0.125 | **0.751** | 8.05E-05 | **2.71E-04** |
| ENSG00000028528.17 | **SNX1** | 6642 | protein_coding | **2025.165** | 0.398 | 0.120 | **1.318** | 8.07E-05 | **2.72E-04** |
| ENSG00000123472.15 | **ATPAF1** | 64756 | protein_coding | **958.427** | -0.415 | -0.125 | **0.750** | 8.08E-05 | **2.72E-04** |
| ENSG00000156299.17 | **TIAM1** | 7074 | protein_coding | **103.271** | 1.149 | 0.346 | **2.217** | 8.09E-05 | **2.72E-04** |
| ENSG00000083635.9 | **NUFIP1** | 26747 | protein_coding | **146.372** | -0.617 | -0.186 | **0.652** | 8.10E-05 | **2.73E-04** |
| ENSG00000062822.18 | **POLD1** | 5424 | protein_coding | **1375.540** | -0.868 | -0.261 | **0.548** | 8.14E-05 | **2.74E-04** |
| ENSG00000157227.16 | **MMP14** | 4323 | protein_coding | **56.039** | 1.021 | 0.307 | **2.029** | 8.14E-05 | **2.74E-04** |
| ENSG00000108932.15 | **SLC16A6** | 9120 | protein_coding | **91.204** | -0.776 | -0.234 | **0.584** | 8.15E-05 | **2.74E-04** |
| ENSG00000163600.15 | **ICOS** | 29851 | protein_coding | **8.124** | 3.708 | 1.116 | **13.066** | 8.16E-05 | **2.75E-04** |
| ENSG00000146955.13 | **RAB19** | 401409 | protein_coding | **6.907** | 2.308 | 0.695 | **4.951** | 8.17E-05 | **2.75E-04** |
| ENSG00000178201.5 | **VN1R1** | 57191 | protein_coding | **7.164** | 2.475 | 0.745 | **5.559** | 8.18E-05 | **2.75E-04** |
| ENSG00000155511.20 | **GRIA1** | 2890 | protein_coding | **22.856** | -1.991 | -0.599 | **0.252** | 8.19E-05 | **2.75E-04** |
| ENSG00000261236.10 | **BOP1** | 23246 | protein_coding | **1350.771** | -0.604 | -0.182 | **0.658** | 8.22E-05 | **2.76E-04** |
| ENSG00000144136.13 | **SLC20A1** | 6574 | protein_coding | **753.405** | -0.403 | -0.121 | **0.756** | 8.22E-05 | **2.76E-04** |
| ENSG00000236204.8 | **LINC01376** | 400945 | lncRNA | **32.807** | 1.200 | 0.361 | **2.297** | 8.22E-05 | **2.76E-04** |
| ENSG00000214290.9 | **POU2AF3** | 120376 | protein_coding | **80.493** | 0.943 | 0.284 | **1.923** | 8.22E-05 | **2.76E-04** |
| ENSG00000120526.14 | **NUDCD1** | 84955 | protein_coding | **272.121** | -0.664 | -0.200 | **0.631** | 8.23E-05 | **2.76E-04** |
| ENSG00000160683.6 | **CXCR5** | 643 | protein_coding | **82.214** | 1.083 | 0.326 | **2.119** | 8.24E-05 | **2.77E-04** |
| ENSG00000233396.10 | **LINC01719** | | lncRNA | **17.177** | 1.447 | 0.436 | **2.726** | 8.25E-05 | **2.77E-04** |
| ENSG00000052749.16 | **RRP12** | 23223 | protein_coding | **543.665** | -0.399 | -0.120 | **0.759** | 8.26E-05 | **2.77E-04** |
| ENSG00000185090.17 | **MANEAL** | 149175 | protein_coding | **132.788** | -0.687 | -0.207 | **0.621** | 8.27E-05 | **2.78E-04** |
| ENSG00000215041.12 | **NEURL4** | 84461 | protein_coding | **746.890** | 0.422 | 0.127 | **1.340** | 8.28E-05 | **2.78E-04** |
| ENSG00000116120.13 | **FARSB** | 10056 | protein_coding | **1278.381** | -0.557 | -0.168 | **0.680** | 8.28E-05 | **2.78E-04** |

| | | | | | | | | | |
|---|---|---|---|---|---|---|---|---|---|
| ENSG00000010244.21 | **ZNF207** | 7756 | protein_coding | **2336.720** | -0.397 | -0.120 | **0.759** | 8.31E-05 | **2.79E-04** |
| ENSG00000100368.17 | **CSF2RB** | 1439 | protein_coding | **132.524** | -0.826 | -0.249 | **0.564** | 8.32E-05 | **2.79E-04** |
| ENSG00000303266.1 | **ENSG00000303266** | | lncRNA | **8.572** | 2.846 | 0.857 | **7.188** | 8.32E-05 | **2.79E-04** |
| ENSG00000259248.8 | **USP3-AS1** | | lncRNA | **24.300** | 1.155 | 0.348 | **2.227** | 8.35E-05 | **2.80E-04** |
| ENSG00000099985.5 | **OSM** | 5008 | protein_coding | **26.834** | -1.034 | -0.311 | **0.488** | 8.37E-05 | **2.81E-04** |
| ENSG00000285533.5 | **RELA-DT** | 105369347 | lncRNA | **123.459** | 0.748 | 0.225 | **1.680** | 8.38E-05 | **2.81E-04** |
| ENSG00000037241.9 | **RPL26L1** | 51121 | protein_coding | **471.132** | -0.431 | -0.130 | **0.742** | 8.41E-05 | **2.82E-04** |
| ENSG00000062650.22 | **WAPL** | 23063 | protein_coding | **660.627** | -0.463 | -0.139 | **0.725** | 8.44E-05 | **2.83E-04** |
| ENSG00000166762.20 | **CATSPER2** | 117155 | protein_coding | **172.123** | 0.659 | 0.198 | **1.579** | 8.48E-05 | **2.84E-04** |
| ENSG00000170421.15 | **KRT8** | 3856 | protein_coding | **26.445** | 1.854 | 0.558 | **3.614** | 8.49E-05 | **2.84E-04** |
| ENSG00000185262.11 | **UBALD2** | 283991 | protein_coding | **2610.251** | -0.528 | -0.159 | **0.693** | 8.50E-05 | **2.85E-04** |
| ENSG00000103035.13 | **PSMD7** | 5713 | protein_coding | **1025.916** | -0.351 | -0.106 | **0.784** | 8.52E-05 | **2.85E-04** |
| ENSG00000198363.20 | **ASPH** | 444 | protein_coding | **174.322** | 0.554 | 0.167 | **1.468** | 8.53E-05 | **2.86E-04** |
| ENSG00000173473.13 | **SMARCC1** | 6599 | protein_coding | **1779.971** | -0.416 | -0.125 | **0.749** | 8.56E-05 | **2.86E-04** |
| ENSG00000004478.10 | **FKBP4** | 2288 | protein_coding | **1337.634** | -0.538 | -0.162 | **0.689** | 8.56E-05 | **2.87E-04** |
| ENSG00000187824.11 | **TMEM220** | 388335 | protein_coding | **23.780** | -1.490 | -0.449 | **0.356** | 8.57E-05 | **2.87E-04** |
| ENSG00000232876.2 | **ENSG0000(** | 124901405 | lncRNA | **19.629** | 1.265 | 0.381 | **2.404** | 8.57E-05 | **2.87E-04** |
| ENSG00000277287.2 | **ENSG00000277287** | | lncRNA | **16.339** | 1.497 | 0.450 | **2.822** | 8.59E-05 | **2.87E-04** |
| ENSG00000269293.6 | **ZSCAN16-A** | 100129195 | lncRNA | **195.700** | 0.727 | 0.219 | **1.655** | 8.59E-05 | **2.87E-04** |
| ENSG00000090861.19 | **AARS1** | 16 | protein_coding | **4227.842** | -0.606 | -0.182 | **0.657** | 8.60E-05 | **2.88E-04** |
| ENSG00000186501.17 | **TMEM222** | 84065 | protein_coding | **521.667** | -0.465 | -0.140 | **0.724** | 8.63E-05 | **2.89E-04** |
| ENSG00000182141.12 | **ZNF708** | 7562 | protein_coding | **147.075** | -0.736 | -0.222 | **0.600** | 8.63E-05 | **2.89E-04** |
| ENSG00000036672.18 | **USP2** | 9099 | protein_coding | **8.522** | 2.228 | 0.671 | **4.685** | 8.63E-05 | **2.89E-04** |
| ENSG00000138135.7 | **CH25H** | 9023 | protein_coding | **3.374** | 4.226 | 1.272 | **18.716** | 8.67E-05 | **2.90E-04** |
| ENSG00000304941.1 | **ENSG00000304941** | | lncRNA | **5.784** | 3.054 | 0.919 | **8.305** | 8.67E-05 | **2.90E-04** |
| ENSG00000214530.12 | **STARD10** | 10809 | protein_coding | **602.878** | 0.450 | 0.136 | **1.366** | 8.67E-05 | **2.90E-04** |
| ENSG00000175318.14 | **GRAMD2A** | 196996 | protein_coding | **5.597** | 3.403 | 1.024 | **10.579** | 8.68E-05 | **2.90E-04** |
| ENSG00000171357.7 | **LURAP1** | 541468 | protein_coding | **9.703** | 2.066 | 0.622 | **4.187** | 8.71E-05 | **2.91E-04** |
| ENSG00000113048.21 | **MRPS27** | 23107 | protein_coding | **1162.226** | -0.437 | -0.131 | **0.739** | 8.73E-05 | **2.92E-04** |
| ENSG00000185436.14 | **IFNLR1** | 163702 | protein_coding | **281.824** | 0.537 | 0.162 | **1.451** | 8.73E-05 | **2.92E-04** |
| ENSG00000203872.7 | **C6orf163** | 206412 | protein_coding | **64.285** | 1.093 | 0.329 | **2.134** | 8.73E-05 | **2.92E-04** |

| | | | | | | | | | |
|---|---|---|---|---|---|---|---|---|---|
| ENSG00000140538.19 | **NTRK3** | 4916 | protein_coding | **8.291** | 4.651 | 1.400 | **25.123** | 8.75E-05 | **2.92E-04** |
| ENSG00000164976.11 | **MYORG** | 57462 | protein_coding | **17.044** | 1.539 | 0.463 | **2.905** | 8.77E-05 | **2.93E-04** |
| ENSG00000302660.1 | **ENSG00000302660** | | lncRNA | **12.280** | 2.388 | 0.719 | **5.234** | 8.79E-05 | **2.93E-04** |
| ENSG00000134438.11 | **RAX** | 30062 | protein_coding | **8.038** | 2.733 | 0.823 | **6.650** | 8.89E-05 | **2.97E-04** |
| ENSG00000095637.25 | **SORBS1** | 10580 | protein_coding | **30.548** | 1.776 | 0.535 | **3.425** | 8.91E-05 | **2.97E-04** |
| ENSG00000166797.14 | **CIAO2A** | 84191 | protein_coding | **962.513** | -0.311 | -0.094 | **0.806** | 8.96E-05 | **2.99E-04** |
| ENSG00000186019.13 | **ZNF225-AS1** | | lncRNA | **24.792** | 1.166 | 0.351 | **2.244** | 9.00E-05 | **3.00E-04** |
| ENSG00000187902.14 | **SHISA7** | 729956 | protein_coding | **6.349** | 3.223 | 0.970 | **9.335** | 9.03E-05 | **3.01E-04** |
| ENSG00000161905.14 | **ALOX15** | 246 | protein_coding | **4.544** | 3.564 | 1.073 | **11.823** | 9.04E-05 | **3.01E-04** |
| ENSG00000165704.17 | **HPRT1** | 3251 | protein_coding | **626.440** | -0.592 | -0.178 | **0.664** | 9.04E-05 | **3.01E-04** |
| ENSG00000257433.9 | **RPAP3-DT** | | lncRNA | **54.023** | 0.868 | 0.261 | **1.825** | 9.04E-05 | **3.01E-04** |
| ENSG00000196743.11 | **GM2A** | 2760 | protein_coding | **3209.262** | 0.389 | 0.117 | **1.309** | 9.05E-05 | **3.02E-04** |
| ENSG00000103723.18 | **AP3B2** | 8120 | protein_coding | **12.532** | 2.359 | 0.710 | **5.131** | 9.06E-05 | **3.02E-04** |
| ENSG00000081791.11 | **DELE1** | 9812 | protein_coding | **1181.734** | -0.345 | -0.104 | **0.787** | 9.09E-05 | **3.03E-04** |
| ENSG00000137038.10 | **DMAC1** | 90871 | protein_coding | **573.144** | -0.676 | -0.204 | **0.626** | 9.11E-05 | **3.03E-04** |
| ENSG00000243749.4 | **TMEM35B** | 10050614 | protein_coding | **225.630** | -0.592 | -0.178 | **0.663** | 9.15E-05 | **3.05E-04** |
| ENSG00000280128.1 | **ENSG00000280128** | | TEC | **136.650** | 0.600 | 0.180 | **1.515** | 9.16E-05 | **3.05E-04** |
| ENSG00000118689.17 | **FOXO3** | 2309 | protein_coding | **122.399** | 0.905 | 0.272 | **1.873** | 9.23E-05 | **3.07E-04** |
| ENSG00000130749.10 | **ZC3H4** | 23211 | protein_coding | **711.494** | -0.351 | -0.106 | **0.784** | 9.24E-05 | **3.07E-04** |
| ENSG00000290588.2 | **ENSG00000290588** | | lncRNA | **11.717** | 1.969 | 0.593 | **3.914** | 9.24E-05 | **3.07E-04** |
| ENSG00000125457.17 | **MIF4GD** | 57409 | protein_coding | **516.707** | 0.565 | 0.170 | **1.480** | 9.25E-05 | **3.08E-04** |
| ENSG00000284652.1 | **ENSG00000284652** | | lncRNA | **5.771** | 3.272 | 0.985 | **9.661** | 9.26E-05 | **3.08E-04** |
| ENSG00000180008.11 | **SOCS4** | 122809 | protein_coding | **308.726** | -0.594 | -0.179 | **0.662** | 9.29E-05 | **3.09E-04** |
| ENSG00000136522.16 | **MRPL47** | 57129 | protein_coding | **742.723** | -0.339 | -0.102 | **0.791** | 9.39E-05 | **3.12E-04** |
| ENSG00000165526.11 | **RPUSD4** | 84881 | protein_coding | **478.477** | -0.434 | -0.131 | **0.740** | 9.41E-05 | **3.13E-04** |
| ENSG00000291111.2 | **ENSG00000291111** | | lncRNA | **31.106** | 1.637 | 0.493 | **3.111** | 9.43E-05 | **3.13E-04** |
| ENSG00000213523.14 | **SRA1** | 10011 | protein_coding | **1171.002** | -0.333 | -0.100 | **0.794** | 9.44E-05 | **3.14E-04** |
| ENSG00000136758.23 | **YME1L1** | 10730 | protein_coding | **1203.227** | -0.568 | -0.171 | **0.675** | 9.44E-05 | **3.14E-04** |
| ENSG00000101935.14 | **AMMECR1** | 9949 | protein_coding | **178.777** | -0.708 | -0.213 | **0.612** | 9.47E-05 | **3.15E-04** |
| ENSG00000138286.17 | **FAM149B1** | 317662 | protein_coding | **270.664** | 0.485 | 0.146 | **1.399** | 9.48E-05 | **3.15E-04** |
| ENSG00000164244.23 | **PRRC1** | 133619 | protein_coding | **427.015** | -0.480 | -0.144 | **0.717** | 9.48E-05 | **3.15E-04** |

| | | | | | | | | | |
|---|---|---|---|---|---|---|---|---|---|
| ENSG00000233922.5 | **LINC01694** | 10537284( | lncRNA | **260.061** | 0.746 | 0.225 | **1.677** | 9.49E-05 | **3.15E-04** |
| ENSG00000160310.22 | **PRMT2** | 3275 | protein_coding | **864.775** | -0.375 | -0.113 | **0.771** | 9.51E-05 | **3.15E-04** |
| ENSG00000148297.18 | **MED22** | 6837 | protein_coding | **507.501** | -0.386 | -0.116 | **0.765** | 9.51E-05 | **3.16E-04** |
| ENSG00000204536.15 | **CCHCR1** | 54535 | protein_coding | **925.087** | 0.417 | 0.126 | **1.336** | 9.52E-05 | **3.16E-04** |
| ENSG00000308327.1 | **ENSG00000308327** | | lncRNA | **11.829** | 1.744 | 0.525 | **3.349** | 9.55E-05 | **3.17E-04** |
| ENSG00000293240.2 | **ENSG00000293240** | | lncRNA | **13.135** | -2.279 | -0.686 | **0.206** | 9.69E-05 | **3.21E-04** |
| ENSG00000275765.7 | **ENSG0000(** | 10065275ε | lncRNA | **89.866** | 0.719 | 0.217 | **1.646** | 9.70E-05 | **3.22E-04** |
| ENSG00000103253.21 | **HAGHL** | 84264 | protein_coding | **439.366** | -0.553 | -0.167 | **0.681** | 9.71E-05 | **3.22E-04** |
| ENSG00000293573.1 | **ENSG0000(** | 12490158( | lncRNA | **8.341** | 3.454 | 1.040 | **10.957** | 9.72E-05 | **3.22E-04** |
| ENSG00000143157.14 | **POGK** | 57645 | protein_coding | **604.031** | -0.429 | -0.129 | **0.743** | 9.72E-05 | **3.22E-04** |
| ENSG00000228639.9 | **ROCR** | | lncRNA | **36.360** | 1.385 | 0.417 | **2.611** | 9.74E-05 | **3.23E-04** |
| ENSG00000181896.13 | **ZNF101** | 94039 | protein_coding | **241.731** | -0.568 | -0.171 | **0.674** | 9.75E-05 | **3.23E-04** |
| ENSG00000224420.6 | **ADM5** | 199800 | protein_coding | **198.988** | 0.623 | 0.188 | **1.540** | 9.84E-05 | **3.26E-04** |
| ENSG00000260257.2 | **ENSG0000(** | 11974655ξ | lncRNA | **183.281** | 0.716 | 0.216 | **1.643** | 9.87E-05 | **3.27E-04** |
| ENSG00000171421.15 | **MRPL36** | 64979 | protein_coding | **683.038** | -0.415 | -0.125 | **0.750** | 9.91E-05 | **3.28E-04** |
| ENSG00000281327.5 | **LINC01338** | | lncRNA | **12.239** | 2.146 | 0.646 | **4.427** | 9.92E-05 | **3.28E-04** |
| ENSG00000136014.14 | **USP44** | 84101 | protein_coding | **5.261** | 3.270 | 0.984 | **9.645** | 9.93E-05 | **3.29E-04** |
| ENSG00000310005.1 | **ENSG00000310005** | | lncRNA | **6.349** | 2.655 | 0.799 | **6.297** | 9.93E-05 | **3.29E-04** |
| ENSG00000254887.3 | **LNROP** | 12807154( | lncRNA | **49.688** | -1.174 | -0.353 | **0.443** | 9.94E-05 | **3.29E-04** |
| ENSG00000205339.12 | **IPO7** | 10527 | protein_coding | **505.067** | -0.494 | -0.149 | **0.710** | 9.94E-05 | **3.29E-04** |
| ENSG00000144827.11 | **ABHD10** | 55347 | protein_coding | **463.635** | -0.403 | -0.121 | **0.756** | 9.94E-05 | **3.29E-04** |
| ENSG00000132970.16 | **WASF3** | 10810 | protein_coding | **5.933** | 3.082 | 0.928 | **8.470** | 9.95E-05 | **3.29E-04** |
| ENSG00000297098.1 | **ENSG00000297098** | | lncRNA | **4.386** | 3.369 | 1.014 | **10.333** | 9.96E-05 | **3.30E-04** |
| ENSG00000183054.15 | **RGPD6** | 729540 | protein_coding | **239.460** | 0.630 | 0.190 | **1.547** | 9.97E-05 | **3.30E-04** |
| ENSG00000298624.1 | **ENSG00000298624** | | lncRNA | **52.430** | -1.127 | -0.339 | **0.458** | 1.00E-04 | **3.31E-04** |
| ENSG00000136271.13 | **DDX56** | 54606 | protein_coding | **1628.404** | -0.631 | -0.190 | **0.646** | 1.00E-04 | **3.31E-04** |
| ENSG00000278041.2 | **ENSG00000278041** | | lncRNA | **31.681** | 1.187 | 0.357 | **2.277** | 1.00E-04 | **3.31E-04** |
| ENSG00000162302.15 | **RPS6KA4** | 8986 | protein_coding | **569.367** | 0.492 | 0.148 | **1.406** | 1.00E-04 | **3.31E-04** |
| ENSG00000304359.1 | **ENSG00000304359** | | lncRNA | **8.373** | 2.028 | 0.611 | **4.079** | 1.00E-04 | **3.32E-04** |
| ENSG00000160183.19 | **TMPRSS3** | 64699 | protein_coding | **413.152** | 1.881 | 0.566 | **3.684** | 1.01E-04 | **3.32E-04** |
| ENSG00000130638.19 | **ATXN10** | 25814 | protein_coding | **1066.486** | -0.591 | -0.178 | **0.664** | 1.01E-04 | **3.33E-04** |

| | | | | | | | | | |
|---|---|---|---|---|---|---|---|---|---|
| ENSG00000226648.3 | **PLCG1-AS1** | | lncRNA | **34.268** | 1.144 | 0.345 | **2.211** | 1.01E-04 | **3.33E-04** |
| ENSG00000181544.18 | **FANCB** | 2187 | protein_coding | **40.236** | -1.423 | -0.428 | **0.373** | 1.01E-04 | **3.34E-04** |
| ENSG00000198677.15 | **SKIC3** | 9652 | protein_coding | **1216.116** | 0.469 | 0.141 | **1.384** | 1.01E-04 | **3.34E-04** |
| ENSG00000306802.1 | **ENSG00000306802** | | lncRNA | **10.934** | -1.712 | -0.515 | **0.305** | 1.01E-04 | **3.34E-04** |
| ENSG00000270194.2 | **GOLGA4-A** | 152048 | lncRNA | **11.047** | -1.838 | -0.553 | **0.280** | 1.01E-04 | **3.34E-04** |
| ENSG00000132334.19 | **PTPRE** | 5791 | protein_coding | **182.121** | -0.624 | -0.188 | **0.649** | 1.01E-04 | **3.34E-04** |
| ENSG00000308095.1 | **ENSG00000308095** | | lncRNA | **8.352** | 2.423 | 0.730 | **5.364** | 1.01E-04 | **3.34E-04** |
| ENSG00000261971.10 | **MMP25-AS** | 124900372 | lncRNA | **395.950** | 0.460 | 0.138 | **1.375** | 1.01E-04 | **3.34E-04** |
| ENSG00000105793.18 | **GTPBP10** | 85865 | protein_coding | **339.303** | -0.736 | -0.222 | **0.600** | 1.01E-04 | **3.35E-04** |
| ENSG00000259291.2 | **ZNF710-AS** | 109729181 | lncRNA | **179.490** | 0.564 | 0.170 | **1.479** | 1.02E-04 | **3.37E-04** |
| ENSG00000185015.9 | **CA13** | 377677 | protein_coding | **42.237** | 1.016 | 0.306 | **2.022** | 1.02E-04 | **3.37E-04** |
| ENSG00000198759.14 | **EGFL6** | 25975 | protein_coding | **6.946** | 3.396 | 1.022 | **10.529** | 1.02E-04 | **3.38E-04** |
| ENSG00000213186.10 | **TRIM59** | 286827 | protein_coding | **209.318** | -0.504 | -0.152 | **0.705** | 1.03E-04 | **3.39E-04** |
| ENSG00000283757.2 | **ENSG00000283757** | | lncRNA | **9.711** | 3.119 | 0.939 | **8.688** | 1.03E-04 | **3.40E-04** |
| ENSG00000285521.2 | **ENSG00000285521** | | lncRNA | **4.029** | 3.795 | 1.142 | **13.877** | 1.03E-04 | **3.41E-04** |
| ENSG00000159592.13 | **GPBP1L1** | 60313 | protein_coding | **1321.800** | -0.389 | -0.117 | **0.764** | 1.03E-04 | **3.41E-04** |
| ENSG00000075856.14 | **SART3** | 9733 | protein_coding | **1687.842** | 0.487 | 0.147 | **1.402** | 1.03E-04 | **3.41E-04** |
| ENSG00000290838.2 | **NCF1B** | 654816 | lncRNA | **980.624** | -0.542 | -0.163 | **0.687** | 1.04E-04 | **3.41E-04** |
| ENSG00000308740.1 | **ENSG00000308740** | | lncRNA | **10.632** | 2.359 | 0.710 | **5.130** | 1.04E-04 | **3.41E-04** |
| ENSG00000117862.16 | **TXNDC12** | 51060 | protein_coding | **654.501** | -0.481 | -0.145 | **0.717** | 1.04E-04 | **3.42E-04** |
| ENSG00000240445.6 | **FOXO3B** | 2310 | protein_coding | **42.381** | -1.084 | -0.326 | **0.472** | 1.04E-04 | **3.42E-04** |
| ENSG00000186660.17 | **ZFP91** | 80829 | protein_coding | **665.015** | -0.619 | -0.186 | **0.651** | 1.04E-04 | **3.42E-04** |
| ENSG00000180539.8 | **LINC02908** | 401563 | lncRNA | **442.024** | 1.735 | 0.522 | **3.329** | 1.05E-04 | **3.46E-04** |
| ENSG00000070182.23 | **SPTB** | 6710 | protein_coding | **15.616** | 2.434 | 0.733 | **5.405** | 1.05E-04 | **3.46E-04** |
| ENSG00000108309.16 | **RUNDC3A** | 10900 | protein_coding | **9.117** | 2.505 | 0.754 | **5.678** | 1.05E-04 | **3.46E-04** |
| ENSG00000229388.2 | **TAF12-DT** | | lncRNA | **14.408** | 2.030 | 0.611 | **4.083** | 1.06E-04 | **3.49E-04** |
| ENSG00000288833.3 | **ENSG00000288833** | | lncRNA | **41.241** | 1.073 | 0.323 | **2.104** | 1.06E-04 | **3.49E-04** |
| ENSG00000152217.22 | **SETBP1** | 26040 | protein_coding | **118.893** | 0.826 | 0.249 | **1.773** | 1.06E-04 | **3.49E-04** |
| ENSG00000174792.12 | **ODAPH** | 152816 | protein_coding | **9.315** | 2.317 | 0.697 | **4.982** | 1.06E-04 | **3.49E-04** |
| ENSG00000253730.2 | **ENSG00000253730** | | lncRNA | **6.691** | 2.653 | 0.799 | **6.291** | 1.06E-04 | **3.50E-04** |
| ENSG00000205710.6 | **C17orf107** | 100130311 | protein_coding | **13.739** | 1.493 | 0.450 | **2.815** | 1.06E-04 | **3.50E-04** |

| | | | | | | | | | |
|---|---|---|---|---|---|---|---|---|---|
| ENSG00000095203.17 | **EPB41L4B** | 54566 | protein_coding | **28.360** | 1.302 | 0.392 | **2.466** | 1.07E-04 | **3.50E-04** |
| ENSG00000144034.18 | **TPRKB** | 51002 | protein_coding | **315.416** | -0.463 | -0.139 | **0.725** | 1.07E-04 | **3.50E-04** |
| ENSG00000166523.9 | **CLEC4E** | 26253 | protein_coding | **5.277** | 3.750 | 1.129 | **13.455** | 1.07E-04 | **3.51E-04** |
| ENSG00000276791.1 | **ENSG00000276791** | | lncRNA | **48.857** | 0.993 | 0.299 | **1.990** | 1.07E-04 | **3.51E-04** |
| ENSG00000028310.20 | **BRD9** | 65980 | protein_coding | **984.466** | -0.406 | -0.122 | **0.755** | 1.07E-04 | **3.51E-04** |
| ENSG00000100036.14 | **SLC35E4** | 339665 | protein_coding | **40.566** | 1.150 | 0.346 | **2.219** | 1.07E-04 | **3.52E-04** |
| ENSG00000180304.15 | **OAZ2** | 4947 | protein_coding | **1025.267** | 0.479 | 0.144 | **1.394** | 1.07E-04 | **3.52E-04** |
| ENSG00000109536.15 | **FRG1** | 2483 | protein_coding | **612.562** | -0.396 | -0.119 | **0.760** | 1.07E-04 | **3.52E-04** |
| ENSG00000168811.9 | **IL12A** | 3592 | protein_coding | **141.578** | 0.707 | 0.213 | **1.632** | 1.08E-04 | **3.53E-04** |
| ENSG00000257839.2 | **PHLDA1-DT** | | lncRNA | **7.292** | 2.843 | 0.856 | **7.173** | 1.08E-04 | **3.54E-04** |
| ENSG00000290376.2 | **HERC2P3** | 283755 | lncRNA | **86.943** | 0.780 | 0.235 | **1.717** | 1.08E-04 | **3.54E-04** |
| ENSG00000147642.19 | **SYBU** | 55638 | protein_coding | **42.437** | 1.260 | 0.379 | **2.395** | 1.08E-04 | **3.55E-04** |
| ENSG00000198189.13 | **HSD17B11** | 51170 | protein_coding | **258.689** | -0.655 | -0.197 | **0.635** | 1.08E-04 | **3.55E-04** |
| ENSG00000186591.15 | **UBE2H** | 7328 | protein_coding | **1863.351** | 0.483 | 0.145 | **1.398** | 1.09E-04 | **3.57E-04** |
| ENSG00000123094.18 | **RASSF8** | 11228 | protein_coding | **6.621** | 3.567 | 1.074 | **11.851** | 1.09E-04 | **3.57E-04** |
| ENSG00000140848.19 | **CPNE2** | 221184 | protein_coding | **118.535** | 0.784 | 0.236 | **1.722** | 1.09E-04 | **3.57E-04** |
| ENSG00000137876.13 | **RSL24D1** | 51187 | protein_coding | **2543.563** | -0.376 | -0.113 | **0.771** | 1.09E-04 | **3.58E-04** |
| ENSG00000237950.4 | **LINC02918** | 10192960[illegible] | lncRNA | **82.580** | -0.852 | -0.257 | **0.554** | 1.10E-04 | **3.59E-04** |
| ENSG00000289970.2 | **ENSG00000289970** | | lncRNA | **14.484** | 1.626 | 0.490 | **3.087** | 1.10E-04 | **3.60E-04** |
| ENSG00000119801.16 | **YPEL5** | 51646 | protein_coding | **835.108** | 0.502 | 0.151 | **1.416** | 1.10E-04 | **3.61E-04** |
| ENSG00000068831.21 | **RASGRP2** | 10235 | protein_coding | **137.684** | 1.289 | 0.388 | **2.444** | 1.11E-04 | **3.62E-04** |
| ENSG00000149269.12 | **PAK1** | 5058 | protein_coding | **1703.158** | -0.386 | -0.116 | **0.765** | 1.11E-04 | **3.62E-04** |
| ENSG00000007047.18 | **MARK4** | 57787 | protein_coding | **622.496** | 0.434 | 0.131 | **1.351** | 1.11E-04 | **3.64E-04** |
| ENSG00000164253.16 | **WDR41** | 55255 | protein_coding | **750.318** | -0.442 | -0.133 | **0.736** | 1.11E-04 | **3.64E-04** |
| ENSG00000259343.10 | **TMC3-AS1** | 10537092[illegible] | lncRNA | **44.864** | 1.076 | 0.324 | **2.108** | 1.11E-04 | **3.65E-04** |
| ENSG00000008130.17 | **NADK** | 65220 | protein_coding | **1813.714** | 0.425 | 0.128 | **1.343** | 1.12E-04 | **3.65E-04** |
| ENSG00000288911.3 | **ENSG00000288911** | | lncRNA | **7.888** | 2.366 | 0.712 | **5.156** | 1.12E-04 | **3.66E-04** |
| ENSG00000134873.12 | **CLDN10** | 9071 | protein_coding | **19.314** | 2.257 | 0.679 | **4.780** | 1.12E-04 | **3.66E-04** |
| ENSG00000169241.21 | **SLC50A1** | 55974 | protein_coding | **1049.454** | -0.344 | -0.104 | **0.788** | 1.12E-04 | **3.67E-04** |
| ENSG00000258441.1 | **LINC00641** | 283624 | lncRNA | **121.433** | 0.971 | 0.292 | **1.961** | 1.13E-04 | **3.68E-04** |
| ENSG00000196126.14 | **HLA-DRB1** | 3123 | protein_coding | **29983.156** | 0.648 | 0.195 | **1.567** | 1.13E-04 | **3.70E-04** |

| | | | | | | | | | |
|---|---|---|---|---|---|---|---|---|---|
| ENSG00000048649.15 | **RSF1** | 51773 | protein_coding | **526.305** | -0.487 | -0.147 | **0.714** | 1.13E-04 | **3.70E-04** |
| ENSG00000139537.13 | **DRC2** | 85478 | protein_coding | **5.711** | 3.023 | 0.910 | **8.129** | 1.13E-04 | **3.70E-04** |
| ENSG00000167895.18 | **TMC8** | 147138 | protein_coding | **4902.897** | -0.537 | -0.162 | **0.689** | 1.13E-04 | **3.70E-04** |
| ENSG00000107679.17 | **PLEKHA1** | 59338 | protein_coding | **49.209** | 0.964 | 0.290 | **1.951** | 1.14E-04 | **3.71E-04** |
| ENSG00000129757.17 | **CDKN1C** | 1028 | protein_coding | **30.674** | 1.970 | 0.593 | **3.917** | 1.14E-04 | **3.71E-04** |
| ENSG00000272599.2 | **ENSG00000272599** | | lncRNA | **30.700** | -1.106 | -0.333 | **0.465** | 1.14E-04 | **3.71E-04** |
| ENSG00000134627.13 | **PIWIL4** | 143689 | protein_coding | **23.360** | 1.437 | 0.432 | **2.707** | 1.14E-04 | **3.72E-04** |
| ENSG00000285755.3 | **ENSG00000285755** | | lncRNA | **9.654** | 1.881 | 0.566 | **3.683** | 1.14E-04 | **3.73E-04** |
| ENSG00000146830.13 | **GIGYF1** | 64599 | protein_coding | **2398.363** | 0.292 | 0.088 | **1.224** | 1.14E-04 | **3.73E-04** |
| ENSG00000197928.13 | **ZNF677** | 342926 | protein_coding | **48.223** | 0.980 | 0.295 | **1.972** | 1.15E-04 | **3.74E-04** |
| ENSG00000105227.18 | **PRX** | 57716 | protein_coding | **59.689** | 1.265 | 0.381 | **2.403** | 1.15E-04 | **3.74E-04** |
| ENSG00000143554.16 | **SLC27A3** | 11000 | protein_coding | **325.367** | -0.674 | -0.203 | **0.627** | 1.15E-04 | **3.74E-04** |
| ENSG00000112130.19 | **RNF8** | 9025 | protein_coding | **292.583** | -0.482 | -0.145 | **0.716** | 1.15E-04 | **3.75E-04** |
| ENSG00000129654.10 | **FOXJ1** | 2302 | protein_coding | **10.174** | 2.316 | 0.697 | **4.980** | 1.15E-04 | **3.75E-04** |
| ENSG00000165283.18 | **STOML2** | 30968 | protein_coding | **1461.382** | -0.480 | -0.145 | **0.717** | 1.15E-04 | **3.76E-04** |
| ENSG00000267454.6 | **ZNF582-DT** | 386758 | lncRNA | **51.170** | 1.129 | 0.340 | **2.187** | 1.16E-04 | **3.76E-04** |
| ENSG00000120049.22 | **KCNIP2** | 30819 | protein_coding | **185.989** | 0.760 | 0.229 | **1.693** | 1.16E-04 | **3.77E-04** |
| ENSG00000273026.1 | **ENSG00000273026** | | lncRNA | **15.495** | 1.533 | 0.461 | **2.893** | 1.16E-04 | **3.77E-04** |
| ENSG00000143434.19 | **SEMA6C** | 10500 | protein_coding | **107.399** | 1.020 | 0.307 | **2.028** | 1.17E-04 | **3.79E-04** |
| ENSG00000112110.12 | **MRPL18** | 29074 | protein_coding | **855.822** | -0.447 | -0.135 | **0.733** | 1.18E-04 | **3.83E-04** |
| ENSG00000170017.14 | **ALCAM** | 214 | protein_coding | **603.288** | -0.503 | -0.151 | **0.706** | 1.18E-04 | **3.83E-04** |
| ENSG00000113240.16 | **CLK4** | 57396 | protein_coding | **276.640** | -0.577 | -0.174 | **0.670** | 1.18E-04 | **3.84E-04** |
| ENSG00000112667.15 | **DNPH1** | 10591 | protein_coding | **628.713** | -0.595 | -0.179 | **0.662** | 1.18E-04 | **3.84E-04** |
| ENSG00000196436.9 | **NPIPB15** | 440348 | protein_coding | **294.236** | 0.762 | 0.229 | **1.696** | 1.18E-04 | **3.84E-04** |
| ENSG00000143674.13 | **MAP3K21** | 84451 | protein_coding | **21.084** | 1.468 | 0.442 | **2.767** | 1.18E-04 | **3.84E-04** |
| ENSG00000276256.1 | **ENSG00000276256** | | protein_coding | **207.682** | -0.617 | -0.186 | **0.652** | 1.18E-04 | **3.84E-04** |
| ENSG00000115649.18 | **CNPPD1** | 27013 | protein_coding | **1227.429** | -0.329 | -0.099 | **0.796** | 1.18E-04 | **3.84E-04** |
| ENSG00000247679.4 | **ENSG00000247679** | | lncRNA | **69.318** | 0.984 | 0.296 | **1.978** | 1.18E-04 | **3.85E-04** |
| ENSG00000122862.7 | **SRGN** | 5552 | protein_coding | **8743.625** | 0.891 | 0.268 | **1.854** | 1.19E-04 | **3.86E-04** |
| ENSG00000103227.21 | **LMF1** | 64788 | protein_coding | **185.823** | 0.699 | 0.210 | **1.623** | 1.20E-04 | **3.89E-04** |
| ENSG00000157212.22 | **PAXIP1** | 22976 | protein_coding | **388.051** | -0.447 | -0.135 | **0.734** | 1.20E-04 | **3.89E-04** |

| | | | | | | | | | |
|---|---|---|---|---|---|---|---|---|---|
| ENSG00000196419.15 | **XRCC6** | 2547 | protein_coding | **5266.642** | -0.393 | -0.118 | **0.761** | 1.20E-04 | **3.91E-04** |
| ENSG00000081923.17 | **ATP8B1** | 5205 | protein_coding | **29.515** | 1.522 | 0.458 | **2.872** | 1.21E-04 | **3.92E-04** |
| ENSG00000078043.18 | **PIAS2** | 9063 | protein_coding | **470.883** | -0.367 | -0.110 | **0.775** | 1.21E-04 | **3.92E-04** |
| ENSG00000183208.15 | **GDPGP1** | 390637 | protein_coding | **72.119** | 0.746 | 0.225 | **1.678** | 1.21E-04 | **3.92E-04** |
| ENSG00000205213.16 | **LGR4** | 55366 | protein_coding | **20.576** | 1.277 | 0.384 | **2.424** | 1.21E-04 | **3.93E-04** |
| ENSG00000004700.18 | **RECQL** | 5965 | protein_coding | **779.595** | -0.505 | -0.152 | **0.705** | 1.21E-04 | **3.93E-04** |
| ENSG00000236871.8 | **LINC00106** | 751580 | lncRNA | **46.274** | 0.911 | 0.274 | **1.881** | 1.21E-04 | **3.94E-04** |
| ENSG00000118894.17 | **EEF2KMT** | 196483 | protein_coding | **181.019** | -0.573 | -0.173 | **0.672** | 1.22E-04 | **3.95E-04** |
| ENSG00000164118.15 | **CEP44** | 80817 | protein_coding | **275.536** | -0.458 | -0.138 | **0.728** | 1.22E-04 | **3.96E-04** |
| ENSG00000123352.20 | **SPATS2** | 65244 | protein_coding | **1870.656** | 0.431 | 0.130 | **1.348** | 1.22E-04 | **3.96E-04** |
| ENSG00000169567.15 | **HINT1** | 3094 | protein_coding | **3352.978** | -0.444 | -0.134 | **0.735** | 1.22E-04 | **3.96E-04** |
| ENSG00000103269.16 | **RHBDL1** | 9028 | protein_coding | **53.533** | 0.981 | 0.295 | **1.974** | 1.22E-04 | **3.97E-04** |
| ENSG00000198551.11 | **ZNF627** | 199692 | protein_coding | **245.549** | 0.515 | 0.155 | **1.429** | 1.23E-04 | **3.99E-04** |
| ENSG00000130725.10 | **UBE2M** | 9040 | protein_coding | **1201.259** | -0.368 | -0.111 | **0.775** | 1.23E-04 | **3.99E-04** |
| ENSG00000304706.1 | **ENSG00000304706** | | lncRNA | **3.488** | 3.913 | 1.178 | **15.063** | 1.23E-04 | **3.99E-04** |
| ENSG00000197837.4 | **H4C16** | 121504 | protein_coding | **34.475** | -1.072 | -0.323 | **0.476** | 1.23E-04 | **3.99E-04** |
| ENSG00000181754.9 | **AMIGO1** | 57463 | protein_coding | **30.241** | 1.082 | 0.326 | **2.118** | 1.23E-04 | **4.00E-04** |
| ENSG00000148488.18 | **ST8SIA6** | 338596 | protein_coding | **8.854** | 3.164 | 0.952 | **8.964** | 1.24E-04 | **4.00E-04** |
| ENSG00000104228.15 | **TRIM35** | 23087 | protein_coding | **341.872** | 0.426 | 0.128 | **1.344** | 1.24E-04 | **4.01E-04** |
| ENSG00000140553.20 | **UNC45A** | 55898 | protein_coding | **1179.982** | 0.346 | 0.104 | **1.271** | 1.24E-04 | **4.03E-04** |
| ENSG00000197323.14 | **TRIM33** | 51592 | protein_coding | **332.005** | -0.739 | -0.222 | **0.599** | 1.24E-04 | **4.03E-04** |
| ENSG00000290871.2 | **GTF2IP1** | 2970 | lncRNA | **2233.870** | -0.480 | -0.144 | **0.717** | 1.25E-04 | **4.04E-04** |
| ENSG00000100138.16 | **SNU13** | 4809 | protein_coding | **2868.923** | -0.460 | -0.139 | **0.727** | 1.25E-04 | **4.04E-04** |
| ENSG00000143515.22 | **ATP8B2** | 57198 | protein_coding | **530.755** | 0.613 | 0.184 | **1.529** | 1.25E-04 | **4.04E-04** |
| ENSG00000106638.20 | **TBL2** | 26608 | protein_coding | **415.816** | -0.474 | -0.143 | **0.720** | 1.25E-04 | **4.06E-04** |
| ENSG00000141127.17 | **PRPSAP2** | 5636 | protein_coding | **938.131** | -0.385 | -0.116 | **0.766** | 1.25E-04 | **4.06E-04** |
| ENSG00000302981.1 | **ENSG00000302981** | | lncRNA | **4.510** | 3.737 | 1.125 | **13.331** | 1.26E-04 | **4.07E-04** |
| ENSG00000162073.16 | **PAQR4** | 124222 | protein_coding | **466.074** | -0.552 | -0.166 | **0.682** | 1.26E-04 | **4.07E-04** |
| ENSG00000154710.20 | **RABGEF1** | 27342 | protein_coding | **720.408** | -0.453 | -0.136 | **0.730** | 1.26E-04 | **4.09E-04** |
| ENSG00000166348.22 | **USP54** | 159195 | protein_coding | **105.502** | 0.620 | 0.187 | **1.537** | 1.27E-04 | **4.10E-04** |
| ENSG00000111664.12 | **GNB3** | 2784 | protein_coding | **18.499** | 1.543 | 0.465 | **2.914** | 1.27E-04 | **4.10E-04** |

| | | | | | | | | | |
|---|---|---|---|---|---|---|---|---|---|
| ENSG00000152683.16 | **SLC30A6** | 55676 | protein_coding | **362.522** | 0.522 | 0.157 | **1.436** | 1.27E-04 | **4.11E-04** |
| ENSG00000156675.18 | **RAB11FIP1** | 80223 | protein_coding | **940.327** | 0.375 | 0.113 | **1.297** | 1.28E-04 | **4.13E-04** |
| ENSG00000182324.8 | **KCNJ14** | 3770 | protein_coding | **49.989** | 0.881 | 0.265 | **1.841** | 1.28E-04 | **4.14E-04** |
| ENSG00000309490.1 | **ENSG0000(** | 10192808( | lncRNA | **29.666** | 1.771 | 0.533 | **3.412** | 1.28E-04 | **4.14E-04** |
| ENSG00000125630.17 | **POLR1B** | 84172 | protein_coding | **410.368** | -0.652 | -0.196 | **0.636** | 1.28E-04 | **4.14E-04** |
| ENSG00000072778.22 | **ACADVL** | 37 | protein_coding | **7845.262** | 0.480 | 0.144 | **1.395** | 1.28E-04 | **4.14E-04** |
| ENSG00000020633.20 | **RUNX3** | 864 | protein_coding | **5095.525** | 0.449 | 0.135 | **1.365** | 1.28E-04 | **4.14E-04** |
| ENSG00000286177.1 | **ENSG00000286177** | | lncRNA | **28.383** | 0.975 | 0.293 | **1.965** | 1.28E-04 | **4.14E-04** |
| ENSG00000089154.13 | **GCN1** | 10985 | protein_coding | **2921.075** | 0.473 | 0.142 | **1.388** | 1.28E-04 | **4.14E-04** |
| ENSG00000189362.14 | **NEMP2** | 100131211 | protein_coding | **105.090** | -0.727 | -0.219 | **0.604** | 1.28E-04 | **4.14E-04** |
| ENSG00000184357.5 | **H1-5** | 3009 | protein_coding | **9.093** | -2.198 | -0.662 | **0.218** | 1.28E-04 | **4.14E-04** |
| ENSG00000299773.1 | **ENSG00000299773** | | lncRNA | **6.623** | -2.377 | -0.715 | **0.193** | 1.28E-04 | **4.14E-04** |
| ENSG00000293423.2 | **ABCA17P** | 650655 | lncRNA | **8.793** | 2.258 | 0.680 | **4.782** | 1.29E-04 | **4.15E-04** |
| ENSG00000305731.1 | **ENSG00000305731** | | lncRNA | **193.433** | 0.586 | 0.176 | **1.501** | 1.29E-04 | **4.15E-04** |
| ENSG00000135046.17 | **ANXA1** | 301 | protein_coding | **169.803** | 0.795 | 0.239 | **1.735** | 1.29E-04 | **4.15E-04** |
| ENSG00000167680.20 | **SEMA6B** | 10501 | protein_coding | **11.459** | 2.667 | 0.803 | **6.351** | 1.29E-04 | **4.15E-04** |
| ENSG00000116005.14 | **PCYOX1** | 51449 | protein_coding | **141.524** | 0.702 | 0.211 | **1.627** | 1.29E-04 | **4.16E-04** |
| ENSG00000012061.19 | **ERCC1** | 2067 | protein_coding | **1450.333** | 0.472 | 0.142 | **1.387** | 1.29E-04 | **4.17E-04** |
| ENSG00000268401.2 | **ENSG00000268401** | | lncRNA | **15.007** | 1.545 | 0.465 | **2.917** | 1.29E-04 | **4.17E-04** |
| ENSG00000003056.10 | **M6PR** | 4074 | protein_coding | **2932.319** | 0.345 | 0.104 | **1.270** | 1.30E-04 | **4.17E-04** |
| ENSG00000295353.1 | **ENSG00000295353** | | lncRNA | **4.644** | 3.277 | 0.987 | **9.696** | 1.30E-04 | **4.20E-04** |
| ENSG00000156453.15 | **PCDH1** | 5097 | protein_coding | **7.255** | 2.351 | 0.708 | **5.101** | 1.31E-04 | **4.23E-04** |
| ENSG00000154451.17 | **GBP5** | 115362 | protein_coding | **393.309** | 0.791 | 0.238 | **1.731** | 1.31E-04 | **4.23E-04** |
| ENSG00000204347.5 | **BTBD17** | 388419 | protein_coding | **4.138** | 3.422 | 1.030 | **10.719** | 1.32E-04 | **4.24E-04** |
| ENSG00000110435.15 | **PDHX** | 8050 | protein_coding | **279.838** | -0.649 | -0.195 | **0.638** | 1.32E-04 | **4.24E-04** |
| ENSG00000170606.18 | **HSPA4** | 3308 | protein_coding | **2568.530** | -0.450 | -0.135 | **0.732** | 1.32E-04 | **4.26E-04** |
| ENSG00000040608.15 | **RTN4R** | 65078 | protein_coding | **16.073** | -1.648 | -0.496 | **0.319** | 1.32E-04 | **4.26E-04** |
| ENSG00000092108.24 | **SCFD1** | 23256 | protein_coding | **853.492** | -0.435 | -0.131 | **0.740** | 1.33E-04 | **4.27E-04** |
| ENSG00000272752.7 | **STAG3L5P-** | 10175239( | lncRNA | **521.561** | 0.993 | 0.299 | **1.991** | 1.33E-04 | **4.28E-04** |
| ENSG00000103994.19 | **ZNF106** | 64397 | protein_coding | **583.614** | -0.393 | -0.118 | **0.761** | 1.33E-04 | **4.29E-04** |
| ENSG00000180834.10 | **MAP6D1** | 79929 | protein_coding | **15.079** | 2.961 | 0.891 | **7.786** | 1.34E-04 | **4.32E-04** |

| | | | | | | | | | |
|---|---|---|---|---|---|---|---|---|---|
| ENSG00000151694.17 | **ADAM17** | 6868 | protein_coding | **538.339** | 0.472 | 0.142 | **1.387** | 1.35E-04 | **4.34E-04** |
| ENSG00000169750.11 | **RAC3** | 5881 | protein_coding | **79.401** | 0.819 | 0.247 | **1.764** | 1.35E-04 | **4.34E-04** |
| ENSG00000283667.1 | **ENSG00000283667** | | lncRNA | **32.418** | 1.088 | 0.327 | **2.125** | 1.35E-04 | **4.34E-04** |
| ENSG00000173083.18 | **HPSE** | 10855 | protein_coding | **58.287** | 1.408 | 0.424 | **2.655** | 1.35E-04 | **4.34E-04** |
| ENSG00000177692.15 | **DNAJC28** | 54943 | protein_coding | **38.076** | 1.672 | 0.503 | **3.186** | 1.35E-04 | **4.35E-04** |
| ENSG00000187621.17 | **TCL6** | 124903372 | lncRNA | **110.906** | -0.953 | -0.287 | **0.517** | 1.35E-04 | **4.35E-04** |
| ENSG00000091317.10 | **CMTM6** | 54918 | protein_coding | **522.164** | 0.512 | 0.154 | **1.426** | 1.36E-04 | **4.37E-04** |
| ENSG00000105771.16 | **SMG9** | 56006 | protein_coding | **846.050** | -0.388 | -0.117 | **0.764** | 1.36E-04 | **4.38E-04** |
| ENSG00000272902.2 | **TBC1D8-AS** | 10050628€ | lncRNA | **8.599** | 2.170 | 0.653 | **4.501** | 1.37E-04 | **4.39E-04** |
| ENSG00000106628.15 | **POLD2** | 5425 | protein_coding | **1616.801** | -0.425 | -0.128 | **0.745** | 1.37E-04 | **4.41E-04** |
| ENSG00000143630.12 | **HCN3** | 57657 | protein_coding | **253.908** | 0.782 | 0.235 | **1.720** | 1.38E-04 | **4.43E-04** |
| ENSG00000101350.10 | **KIF3B** | 9371 | protein_coding | **656.630** | 0.553 | 0.166 | **1.467** | 1.38E-04 | **4.44E-04** |
| ENSG00000234476.6 | **LINC02765** | | lncRNA | **4.125** | 3.667 | 1.104 | **12.702** | 1.39E-04 | **4.46E-04** |
| ENSG00000065809.14 | **FAM107B** | 83641 | protein_coding | **1180.091** | -0.683 | -0.206 | **0.623** | 1.39E-04 | **4.46E-04** |
| ENSG00000179299.19 | **NSUN7** | 79730 | protein_coding | **93.882** | 1.370 | 0.413 | **2.585** | 1.39E-04 | **4.47E-04** |
| ENSG00000294982.1 | **ENSG00000294982** | | lncRNA | **34.331** | 1.250 | 0.376 | **2.379** | 1.39E-04 | **4.47E-04** |
| ENSG00000126790.14 | **L3HYPDH** | 112849 | protein_coding | **536.297** | 0.457 | 0.138 | **1.373** | 1.39E-04 | **4.47E-04** |
| ENSG00000104154.9 | **SLC30A4** | 7782 | protein_coding | **28.480** | 1.045 | 0.315 | **2.063** | 1.40E-04 | **4.49E-04** |
| ENSG00000105849.8 | **POLR1F** | 221830 | protein_coding | **188.864** | -0.580 | -0.175 | **0.669** | 1.40E-04 | **4.50E-04** |
| ENSG00000121210.18 | **TMEM131l** | 23240 | protein_coding | **4353.024** | 0.346 | 0.104 | **1.271** | 1.41E-04 | **4.51E-04** |
| ENSG00000185101.15 | **ANO9** | 338440 | protein_coding | **2480.738** | 0.451 | 0.136 | **1.367** | 1.41E-04 | **4.51E-04** |
| ENSG00000169718.20 | **DUS1L** | 64118 | protein_coding | **2044.402** | -0.559 | -0.168 | **0.679** | 1.41E-04 | **4.52E-04** |
| ENSG00000114650.23 | **SCAP** | 22937 | protein_coding | **1304.032** | 0.388 | 0.117 | **1.309** | 1.41E-04 | **4.53E-04** |
| ENSG00000285589.1 | **ENSG00000285589** | | protein_coding | **175.429** | 0.589 | 0.177 | **1.504** | 1.42E-04 | **4.54E-04** |
| ENSG00000276846.1 | **ENSG00000276846** | | lncRNA | **6.807** | 2.172 | 0.654 | **4.505** | 1.42E-04 | **4.55E-04** |
| ENSG00000291234.2 | **ENSG00000291234** | | lncRNA | **5.714** | 2.495 | 0.751 | **5.639** | 1.42E-04 | **4.55E-04** |
| ENSG00000139117.16 | **CPNE8** | 144402 | protein_coding | **279.621** | -0.662 | -0.199 | **0.632** | 1.42E-04 | **4.55E-04** |
| ENSG00000177030.21 | **DEAF1** | 10522 | protein_coding | **546.541** | -0.566 | -0.170 | **0.675** | 1.42E-04 | **4.55E-04** |
| ENSG00000099625.16 | **CBARP** | 255057 | protein_coding | **47.819** | 1.247 | 0.375 | **2.373** | 1.42E-04 | **4.56E-04** |
| ENSG00000176542.11 | **USF3** | 205717 | protein_coding | **110.526** | -0.573 | -0.172 | **0.672** | 1.42E-04 | **4.56E-04** |
| ENSG00000185245.10 | **GP1BA** | 2811 | protein_coding | **19.432** | 1.689 | 0.509 | **3.225** | 1.42E-04 | **4.56E-04** |

| | | | | | | | | | |
|---|---|---|---|---|---|---|---|---|---|
| ENSG00000008853.20 | **RHOBTB2** | 23221 | protein_coding | **694.338** | -0.602 | -0.181 | **0.659** | 1.43E-04 | **4.56E-04** |
| ENSG00000234616.11 | **JRK** | 8629 | protein_coding | **341.717** | -0.439 | -0.132 | **0.738** | 1.43E-04 | **4.56E-04** |
| ENSG00000164172.22 | **MOCS2** | 4338 | protein_coding | **281.194** | -0.532 | -0.160 | **0.692** | 1.43E-04 | **4.57E-04** |
| ENSG00000180938.8 | **ZNF572** | 137209 | protein_coding | **41.154** | 0.932 | 0.281 | **1.908** | 1.44E-04 | **4.61E-04** |
| ENSG00000184254.20 | **ALDH1A3** | 220 | protein_coding | **9.012** | 2.565 | 0.772 | **5.916** | 1.45E-04 | **4.62E-04** |
| ENSG00000102445.22 | **RUBCNL** | 80183 | protein_coding | **725.618** | 0.506 | 0.152 | **1.420** | 1.45E-04 | **4.62E-04** |
| ENSG00000138615.8 | **CILP** | 8483 | protein_coding | **12.574** | 1.671 | 0.503 | **3.185** | 1.45E-04 | **4.63E-04** |
| ENSG00000225855.8 | **RUSC1-AS1** | 284618 | lncRNA | **25.032** | 1.071 | 0.323 | **2.102** | 1.45E-04 | **4.63E-04** |
| ENSG00000151151.8 | **IPMK** | 253430 | protein_coding | **102.866** | -0.619 | -0.186 | **0.651** | 1.45E-04 | **4.63E-04** |
| ENSG00000167114.15 | **SLC27A4** | 10999 | protein_coding | **206.312** | 0.473 | 0.142 | **1.388** | 1.45E-04 | **4.64E-04** |
| ENSG00000258405.12 | **ZNF578** | 147660 | protein_coding | **14.766** | 1.830 | 0.551 | **3.556** | 1.45E-04 | **4.64E-04** |
| ENSG00000134253.10 | **TRIM45** | 80263 | protein_coding | **69.614** | 0.995 | 0.300 | **1.993** | 1.46E-04 | **4.65E-04** |
| ENSG00000105397.18 | **TYK2** | 7297 | protein_coding | **5289.992** | 0.525 | 0.158 | **1.439** | 1.46E-04 | **4.65E-04** |
| ENSG00000160803.10 | **UBQLN4** | 56893 | protein_coding | **697.524** | 0.355 | 0.107 | **1.279** | 1.46E-04 | **4.66E-04** |
| ENSG00000174227.18 | **PIGG** | 54872 | protein_coding | **703.131** | -0.313 | -0.094 | **0.805** | 1.47E-04 | **4.68E-04** |
| ENSG00000049323.18 | **LTBP1** | 4052 | protein_coding | **28.876** | 1.413 | 0.425 | **2.663** | 1.47E-04 | **4.68E-04** |
| ENSG00000070669.19 | **ASNS** | 440 | protein_coding | **1107.094** | -0.840 | -0.253 | **0.559** | 1.47E-04 | **4.68E-04** |
| ENSG00000078808.22 | **SDF4** | 51150 | protein_coding | **1408.489** | -0.297 | -0.089 | **0.814** | 1.47E-04 | **4.70E-04** |
| ENSG00000132361.20 | **CLUH** | 23277 | protein_coding | **1713.997** | -0.391 | -0.118 | **0.762** | 1.48E-04 | **4.71E-04** |
| ENSG00000289250.1 | **ENSG00000289250** | | lncRNA | **9.338** | 1.959 | 0.590 | **3.888** | 1.48E-04 | **4.72E-04** |
| ENSG00000132823.13 | **OSER1** | 51526 | protein_coding | **818.737** | -0.383 | -0.115 | **0.767** | 1.48E-04 | **4.72E-04** |
| ENSG00000142676.16 | **RPL11** | 6135 | protein_coding | **44589.331** | 0.416 | 0.125 | **1.334** | 1.49E-04 | **4.75E-04** |
| ENSG00000185614.9 | **INKA1** | 389119 | protein_coding | **33.389** | -1.226 | -0.369 | **0.428** | 1.49E-04 | **4.75E-04** |
| ENSG00000166532.18 | **RIMKLB** | 57494 | protein_coding | **334.029** | 0.559 | 0.168 | **1.474** | 1.50E-04 | **4.78E-04** |
| ENSG00000100580.10 | **TMED8** | 283578 | protein_coding | **579.825** | -0.443 | -0.133 | **0.736** | 1.50E-04 | **4.78E-04** |
| ENSG00000181852.20 | **RNF41** | 10193 | protein_coding | **523.307** | -0.694 | -0.209 | **0.618** | 1.50E-04 | **4.79E-04** |
| ENSG00000159761.16 | **C16orf86** | 388284 | protein_coding | **81.989** | 0.865 | 0.260 | **1.821** | 1.51E-04 | **4.79E-04** |
| ENSG00000243414.8 | **TICAM2** | 353376 | protein_coding | **47.495** | -1.349 | -0.406 | **0.393** | 1.52E-04 | **4.83E-04** |
| ENSG00000213977.10 | **TAX1BP3** | 30851 | protein_coding | **323.253** | 0.527 | 0.159 | **1.441** | 1.52E-04 | **4.83E-04** |
| ENSG00000227765.4 | **MYL12BP2** | | processed_pseud | **107.225** | 0.925 | 0.279 | **1.899** | 1.52E-04 | **4.85E-04** |
| ENSG00000188603.24 | **CLN3** | 1201 | protein_coding | **603.261** | 0.555 | 0.167 | **1.469** | 1.53E-04 | **4.86E-04** |

| | | | | | | | | | |
|---|---|---|---|---|---|---|---|---|---|
| ENSG00000142733.20 | **MAP3K6** | 9064 | protein_coding | **103.253** | 1.024 | 0.308 | **2.033** | 1.53E-04 | **4.88E-04** |
| ENSG00000176476.11 | **SGF29** | 112869 | protein_coding | **346.093** | 0.621 | 0.187 | **1.538** | 1.53E-04 | **4.88E-04** |
| ENSG00000025796.16 | **SEC63** | 11231 | protein_coding | **887.127** | -0.481 | -0.145 | **0.717** | 1.53E-04 | **4.88E-04** |
| ENSG00000185269.14 | **NOTUM** | 147111 | protein_coding | **9.738** | 2.035 | 0.613 | **4.100** | 1.53E-04 | **4.88E-04** |
| ENSG00000285793.2 | **ANAPC1P2** | 285074 | lncRNA | **258.910** | -0.702 | -0.211 | **0.615** | 1.54E-04 | **4.91E-04** |
| ENSG00000138594.16 | **TMOD3** | 29766 | protein_coding | **424.333** | -0.568 | -0.171 | **0.674** | 1.55E-04 | **4.93E-04** |
| ENSG00000205189.14 | **ZBTB10** | 65986 | protein_coding | **125.768** | 0.834 | 0.251 | **1.783** | 1.55E-04 | **4.94E-04** |
| ENSG00000148935.14 | **GAS2** | 2620 | protein_coding | **53.025** | -1.150 | -0.346 | **0.451** | 1.56E-04 | **4.95E-04** |
| ENSG00000287420.2 | **ENSG00000287420** | | lncRNA | **14.175** | 1.668 | 0.502 | **3.178** | 1.56E-04 | **4.97E-04** |
| ENSG00000145912.12 | **NHP2** | 55651 | protein_coding | **1415.075** | -0.599 | -0.180 | **0.660** | 1.56E-04 | **4.97E-04** |
| ENSG00000141452.13 | **RMC1** | 29919 | protein_coding | **1065.486** | 0.416 | 0.125 | **1.334** | 1.56E-04 | **4.97E-04** |
| ENSG00000253738.4 | **OTUD6B-AS1** | | lncRNA | **153.173** | -0.492 | -0.148 | **0.711** | 1.57E-04 | **4.98E-04** |
| ENSG00000173809.19 | **TDRD12** | 91646 | protein_coding | **49.046** | 1.118 | 0.337 | **2.170** | 1.57E-04 | **4.99E-04** |
| ENSG00000146242.11 | **TPBG** | 7162 | protein_coding | **295.569** | 0.711 | 0.214 | **1.637** | 1.57E-04 | **4.99E-04** |
| ENSG00000286650.3 | **ENSG00000286650** | | lncRNA | **11.185** | 1.721 | 0.518 | **3.297** | 1.58E-04 | **5.01E-04** |
| ENSG00000267395.6 | **DM1-AS** | | lncRNA | **28.495** | -1.261 | -0.380 | **0.417** | 1.58E-04 | **5.01E-04** |
| ENSG00000297225.1 | **ENSG0000(** | 10537453ε | lncRNA | **9.860** | 2.068 | 0.623 | **4.194** | 1.58E-04 | **5.02E-04** |
| ENSG00000145982.15 | **FARS2** | 10667 | protein_coding | **163.494** | -0.667 | -0.201 | **0.630** | 1.58E-04 | **5.02E-04** |
| ENSG00000289585.3 | **ENSG00000289585** | | lncRNA | **18.121** | 1.200 | 0.361 | **2.297** | 1.59E-04 | **5.03E-04** |
| ENSG00000083814.15 | **ZNF671** | 79891 | protein_coding | **240.839** | -0.438 | -0.132 | **0.738** | 1.59E-04 | **5.03E-04** |
| ENSG00000103196.14 | **CRISPLD2** | 83716 | protein_coding | **11.032** | 2.704 | 0.814 | **6.514** | 1.59E-04 | **5.03E-04** |
| ENSG00000143994.16 | **ABHD1** | 84696 | protein_coding | **10.258** | 1.893 | 0.570 | **3.715** | 1.59E-04 | **5.03E-04** |
| ENSG00000307950.1 | **ENSG00000307950** | | lncRNA | **5.462** | 3.421 | 1.030 | **10.709** | 1.59E-04 | **5.04E-04** |
| ENSG00000113966.13 | **ARL6** | 84100 | protein_coding | **53.860** | 1.095 | 0.330 | **2.136** | 1.59E-04 | **5.05E-04** |
| ENSG00000079277.24 | **MKNK1** | 8569 | protein_coding | **735.080** | 0.509 | 0.153 | **1.423** | 1.59E-04 | **5.05E-04** |
| ENSG00000287530.2 | **ENSG00000287530** | | lncRNA | **4.906** | 3.291 | 0.991 | **9.787** | 1.60E-04 | **5.06E-04** |
| ENSG00000146540.17 | **CHLSN** | 84310 | protein_coding | **2568.126** | 0.341 | 0.103 | **1.266** | 1.60E-04 | **5.06E-04** |
| ENSG00000137266.16 | **SLC22A23** | 63027 | protein_coding | **37.856** | 1.658 | 0.499 | **3.155** | 1.60E-04 | **5.07E-04** |
| ENSG00000159082.20 | **SYNJ1** | 8867 | protein_coding | **127.173** | 0.793 | 0.239 | **1.733** | 1.60E-04 | **5.07E-04** |
| ENSG00000089820.19 | **ARHGAP4** | 393 | protein_coding | **2875.535** | -0.551 | -0.166 | **0.682** | 1.60E-04 | **5.07E-04** |
| ENSG00000307275.1 | **ENSG00000307275** | | lncRNA | **4.577** | 3.250 | 0.978 | **9.515** | 1.60E-04 | **5.07E-04** |

| | | | | | | | | | |
|---|---|---|---|---|---|---|---|---|---|
| ENSG00000096384.22 | **HSP90AB1** | 3326 | protein_coding | **17943.421** | -0.518 | -0.156 | **0.698** | 1.60E-04 | **5.08E-04** |
| ENSG00000152433.17 | **ZNF547** | 284306 | protein_coding | **129.393** | 0.636 | 0.191 | **1.554** | 1.60E-04 | **5.08E-04** |
| ENSG00000184162.17 | **NR2C2AP** | 126382 | protein_coding | **358.220** | -0.472 | -0.142 | **0.721** | 1.60E-04 | **5.08E-04** |
| ENSG00000101363.15 | **MANBAL** | 63905 | protein_coding | **635.989** | 0.386 | 0.116 | **1.307** | 1.61E-04 | **5.08E-04** |
| ENSG00000164219.11 | **PGGT1B** | 5229 | protein_coding | **216.152** | -0.860 | -0.259 | **0.551** | 1.61E-04 | **5.08E-04** |
| ENSG00000137098.16 | **SPAG8** | 26206 | protein_coding | **34.094** | 1.297 | 0.391 | **2.458** | 1.61E-04 | **5.10E-04** |
| ENSG00000164136.19 | **IL15** | 3600 | protein_coding | **276.770** | 0.576 | 0.173 | **1.491** | 1.61E-04 | **5.11E-04** |
| ENSG00000138435.16 | **CHRNA1** | 1134 | protein_coding | **4.286** | 3.426 | 1.031 | **10.748** | 1.62E-04 | **5.11E-04** |
| ENSG00000279138.1 | **ENSG00000279138** | | TEC | **8.220** | 2.295 | 0.691 | **4.908** | 1.62E-04 | **5.11E-04** |
| ENSG00000134014.20 | **ELP3** | 55140 | protein_coding | **632.929** | -0.543 | -0.164 | **0.686** | 1.62E-04 | **5.11E-04** |
| ENSG00000265491.7 | **RNF115** | 27246 | protein_coding | **887.188** | -0.379 | -0.114 | **0.769** | 1.62E-04 | **5.12E-04** |
| ENSG00000163626.19 | **COX18** | 285521 | protein_coding | **329.351** | -0.462 | -0.139 | **0.726** | 1.62E-04 | **5.12E-04** |
| ENSG00000175215.12 | **CTDSP2** | 10106 | protein_coding | **1453.979** | 0.501 | 0.151 | **1.416** | 1.62E-04 | **5.12E-04** |
| ENSG00000166986.17 | **MARS1** | 4141 | protein_coding | **3073.107** | -0.668 | -0.201 | **0.630** | 1.63E-04 | **5.14E-04** |
| ENSG00000223486.2 | **LINC03053** | 12490517[illegible] | lncRNA | **7.241** | 2.768 | 0.833 | **6.812** | 1.63E-04 | **5.14E-04** |
| ENSG00000196544.8 | **BORCS6** | 54785 | protein_coding | **391.847** | 0.450 | 0.135 | **1.366** | 1.63E-04 | **5.15E-04** |
| ENSG00000110911.19 | **SLC11A2** | 4891 | protein_coding | **407.753** | 0.569 | 0.171 | **1.483** | 1.63E-04 | **5.15E-04** |
| ENSG00000157322.20 | **CLEC18A** | 348174 | protein_coding | **30.344** | 1.497 | 0.451 | **2.822** | 1.63E-04 | **5.16E-04** |
| ENSG00000156136.12 | **DCK** | 1633 | protein_coding | **126.298** | -0.692 | -0.208 | **0.619** | 1.65E-04 | **5.20E-04** |
| ENSG00000254006.6 | **ENSG0000[illegible]** | 10537587[illegible] | lncRNA | **104.420** | 1.211 | 0.365 | **2.316** | 1.65E-04 | **5.20E-04** |
| ENSG00000057757.12 | **PITHD1** | 57095 | protein_coding | **734.252** | -0.347 | -0.104 | **0.786** | 1.65E-04 | **5.20E-04** |
| ENSG00000213780.13 | **GTF2H4** | 2968 | protein_coding | **549.455** | 0.457 | 0.137 | **1.372** | 1.65E-04 | **5.21E-04** |
| ENSG00000136709.14 | **WDR33** | 55339 | protein_coding | **1437.967** | -0.386 | -0.116 | **0.765** | 1.65E-04 | **5.21E-04** |
| ENSG00000185379.23 | **RAD51D** | 5892 | protein_coding | **359.633** | -0.583 | -0.175 | **0.668** | 1.65E-04 | **5.21E-04** |
| ENSG00000272636.7 | **DOC2B** | 8447 | protein_coding | **5.798** | 3.322 | 1.000 | **10.002** | 1.65E-04 | **5.21E-04** |
| ENSG00000288909.3 | **ENSG00000288909** | | lncRNA | **8.529** | 2.134 | 0.642 | **4.388** | 1.66E-04 | **5.22E-04** |
| ENSG00000125952.23 | **MAX** | 4149 | protein_coding | **1515.873** | 0.311 | 0.094 | **1.241** | 1.66E-04 | **5.23E-04** |
| ENSG00000304876.1 | **ENSG00000304876** | | lncRNA | **4.327** | 3.553 | 1.070 | **11.740** | 1.66E-04 | **5.24E-04** |
| ENSG00000226125.3 | **LINC01907** | 151475 | lncRNA | **11.049** | 2.151 | 0.647 | **4.441** | 1.67E-04 | **5.25E-04** |
| ENSG00000104529.21 | **EEF1D** | 1936 | protein_coding | **7347.938** | -0.361 | -0.109 | **0.779** | 1.67E-04 | **5.25E-04** |
| ENSG00000273314.1 | **ENSG00000273314** | | lncRNA | **51.742** | 0.854 | 0.257 | **1.808** | 1.67E-04 | **5.26E-04** |

| | | | | | | | | | |
|---|---|---|---|---|---|---|---|---|---|
| ENSG00000007314.12 | **SCN4A** | 6329 | protein_coding | **41.544** | 1.122 | 0.338 | **2.177** | 1.67E-04 | **5.26E-04** |
| ENSG00000163354.15 | **DCST2** | 127579 | protein_coding | **34.613** | 1.661 | 0.500 | **3.161** | 1.67E-04 | **5.26E-04** |
| ENSG00000130770.20 | **ATP5IF1** | 93974 | protein_coding | **3071.452** | -0.523 | -0.157 | **0.696** | 1.68E-04 | **5.28E-04** |
| ENSG00000115946.10 | **PNO1** | 56902 | protein_coding | **269.878** | -0.607 | -0.183 | **0.657** | 1.68E-04 | **5.28E-04** |
| ENSG00000070388.12 | **FGF22** | 27006 | protein_coding | **4.842** | 3.333 | 1.003 | **10.075** | 1.69E-04 | **5.31E-04** |
| ENSG00000251867.4 | **ENSG00000251867** | | lncRNA | **27.582** | 0.966 | 0.291 | **1.953** | 1.71E-04 | **5.38E-04** |
| ENSG00000240497.4 | **ENSG0000(** | 10272447\ | lncRNA | **106.199** | 0.571 | 0.172 | **1.486** | 1.71E-04 | **5.38E-04** |
| ENSG00000189060.6 | **H1-0** | 3005 | protein_coding | **1057.824** | 1.485 | 0.447 | **2.800** | 1.71E-04 | **5.38E-04** |
| ENSG00000293422.2 | **PRORSD1P** | 344405 | lncRNA | **73.678** | -1.045 | -0.315 | **0.485** | 1.72E-04 | **5.41E-04** |
| ENSG00000267698.3 | **ENSG0000(** | 10192757z | lncRNA | **19.852** | -1.462 | -0.440 | **0.363** | 1.72E-04 | **5.42E-04** |
| ENSG00000183617.7 | **MRPL54** | 116541 | protein_coding | **750.053** | -0.386 | -0.116 | **0.765** | 1.72E-04 | **5.42E-04** |
| ENSG00000140307.13 | **GTF2A2** | 2958 | protein_coding | **591.331** | -0.515 | -0.155 | **0.700** | 1.72E-04 | **5.42E-04** |
| ENSG00000159212.15 | **CLIC6** | 54102 | protein_coding | **666.782** | -0.529 | -0.159 | **0.693** | 1.72E-04 | **5.42E-04** |
| ENSG00000138166.8 | **DUSP5** | 1847 | protein_coding | **1880.623** | 0.448 | 0.135 | **1.364** | 1.73E-04 | **5.43E-04** |
| ENSG00000102786.17 | **INTS6** | 26512 | protein_coding | **610.660** | -0.654 | -0.197 | **0.636** | 1.73E-04 | **5.44E-04** |
| ENSG00000285747.1 | **ENSG00000285747** | | lncRNA | **6.966** | 2.154 | 0.649 | **4.452** | 1.73E-04 | **5.44E-04** |
| ENSG00000177731.18 | **FLII** | 2314 | protein_coding | **1799.388** | -0.360 | -0.108 | **0.779** | 1.73E-04 | **5.45E-04** |
| ENSG00000264575.2 | **LINC00526** | | lncRNA | **83.350** | 0.635 | 0.191 | **1.553** | 1.74E-04 | **5.46E-04** |
| ENSG00000120647.12 | **CCDC77** | 84318 | protein_coding | **280.557** | -0.468 | -0.141 | **0.723** | 1.74E-04 | **5.47E-04** |
| ENSG00000255992.2 | **PUS1-AS1** | | lncRNA | **9.360** | 2.331 | 0.702 | **5.030** | 1.74E-04 | **5.47E-04** |
| ENSG00000135127.14 | **BICDL1** | 92558 | protein_coding | **234.537** | 0.757 | 0.228 | **1.690** | 1.74E-04 | **5.48E-04** |
| ENSG00000149591.19 | **TAGLN** | 6876 | protein_coding | **347.619** | 0.793 | 0.239 | **1.732** | 1.75E-04 | **5.49E-04** |
| ENSG00000175854.16 | **SWI5** | 375757 | protein_coding | **190.256** | -0.498 | -0.150 | **0.708** | 1.75E-04 | **5.50E-04** |
| ENSG00000308283.1 | **ENSG00000308283** | | lncRNA | **14.539** | 1.744 | 0.525 | **3.349** | 1.75E-04 | **5.50E-04** |
| ENSG00000296598.1 | **ENSG00000296598** | | lncRNA | **7.581** | 2.448 | 0.737 | **5.456** | 1.75E-04 | **5.50E-04** |
| ENSG00000170222.14 | **ADPRM** | 56985 | protein_coding | **126.879** | -0.542 | -0.163 | **0.687** | 1.76E-04 | **5.50E-04** |
| ENSG00000182749.8 | **PAQR7** | 164091 | protein_coding | **194.620** | -0.488 | -0.147 | **0.713** | 1.76E-04 | **5.51E-04** |
| ENSG00000006210.8 | **CX3CL1** | 6376 | protein_coding | **9.964** | 1.973 | 0.594 | **3.927** | 1.76E-04 | **5.51E-04** |
| ENSG00000101266.21 | **CSNK2A1** | 1457 | protein_coding | **1724.433** | -0.418 | -0.126 | **0.749** | 1.76E-04 | **5.51E-04** |
| ENSG00000258512.3 | **LINC00239** | 145200 | lncRNA | **21.537** | 1.373 | 0.413 | **2.590** | 1.76E-04 | **5.52E-04** |
| ENSG00000185344.17 | **ATP6V0A2** | 23545 | protein_coding | **674.463** | 0.376 | 0.113 | **1.298** | 1.76E-04 | **5.52E-04** |

| | | | | | | | | | |
|---|---|---|---|---|---|---|---|---|---|
| ENSG00000113721.16 | **PDGFRB** | 5159 | protein_coding | **67.084** | 1.203 | 0.362 | **2.302** | 1.76E-04 | **5.52E-04** |
| ENSG00000115685.17 | **PPP1R7** | 5510 | protein_coding | **1655.728** | -0.371 | -0.112 | **0.773** | 1.76E-04 | **5.53E-04** |
| ENSG00000134470.23 | **IL15RA** | 3601 | protein_coding | **363.645** | -0.482 | -0.145 | **0.716** | 1.77E-04 | **5.53E-04** |
| ENSG00000138433.18 | **CIRSR** | 9541 | protein_coding | **903.334** | 0.388 | 0.117 | **1.309** | 1.77E-04 | **5.55E-04** |
| ENSG00000163521.18 | **GLB1L** | 79411 | protein_coding | **113.132** | 0.749 | 0.226 | **1.681** | 1.77E-04 | **5.56E-04** |
| ENSG00000261794.2 | **GOLGA8H** | 728498 | protein_coding | **8.972** | 2.524 | 0.760 | **5.752** | 1.78E-04 | **5.57E-04** |
| ENSG00000156113.26 | **KCNMA1** | 3778 | protein_coding | **34.199** | -1.034 | -0.311 | **0.488** | 1.78E-04 | **5.57E-04** |
| ENSG00000110274.18 | **CEP164** | 22897 | protein_coding | **631.964** | 0.438 | 0.132 | **1.355** | 1.78E-04 | **5.58E-04** |
| ENSG00000100302.9 | **RASD2** | 23551 | protein_coding | **5.069** | 3.577 | 1.077 | **11.938** | 1.78E-04 | **5.58E-04** |
| ENSG00000024048.13 | **UBR2** | 23304 | protein_coding | **976.564** | 0.442 | 0.133 | **1.358** | 1.78E-04 | **5.58E-04** |
| ENSG00000289298.3 | **ENSG00000289298** | | lncRNA | **41.726** | 1.145 | 0.345 | **2.212** | 1.79E-04 | **5.61E-04** |
| ENSG00000116750.16 | **UCHL5** | 51377 | protein_coding | **586.004** | -0.628 | -0.189 | **0.647** | 1.80E-04 | **5.62E-04** |
| ENSG00000178035.15 | **IMPDH2** | 3615 | protein_coding | **3579.958** | -0.532 | -0.160 | **0.692** | 1.80E-04 | **5.62E-04** |
| ENSG00000198853.14 | **RUSC2** | 9853 | protein_coding | **46.634** | 0.981 | 0.295 | **1.973** | 1.80E-04 | **5.63E-04** |
| ENSG00000022567.12 | **SLC45A4** | 57210 | protein_coding | **103.189** | 0.751 | 0.226 | **1.683** | 1.81E-04 | **5.64E-04** |
| ENSG00000173156.9 | **RHOD** | 29984 | protein_coding | **27.721** | 1.041 | 0.313 | **2.058** | 1.81E-04 | **5.65E-04** |
| ENSG00000100578.19 | **KIAA0586** | 9786 | protein_coding | **287.427** | -0.501 | -0.151 | **0.707** | 1.81E-04 | **5.65E-04** |
| ENSG00000260948.1 | **ENSG00000260948** | | lncRNA | **20.041** | 1.220 | 0.367 | **2.330** | 1.81E-04 | **5.66E-04** |
| ENSG00000282419.3 | **TEX13D** | 10013201 | protein_coding | **15.059** | 1.962 | 0.591 | **3.896** | 1.82E-04 | **5.69E-04** |
| ENSG00000281991.3 | **TMEM265** | 10086267 | protein_coding | **768.688** | -0.727 | -0.219 | **0.604** | 1.82E-04 | **5.70E-04** |
| ENSG00000274220.1 | **ENSG00000274220** | | lncRNA | **20.398** | 1.160 | 0.349 | **2.235** | 1.83E-04 | **5.70E-04** |
| ENSG00000125454.14 | **SLC25A19** | 60386 | protein_coding | **721.425** | -0.562 | -0.169 | **0.677** | 1.83E-04 | **5.72E-04** |
| ENSG00000141526.20 | **SLC16A3** | 9123 | protein_coding | **1808.162** | -0.519 | -0.156 | **0.698** | 1.83E-04 | **5.73E-04** |
| ENSG00000105329.13 | **TGFB1** | 7040 | protein_coding | **1527.491** | -0.450 | -0.136 | **0.732** | 1.83E-04 | **5.73E-04** |
| ENSG00000165322.20 | **ARHGAP12** | 94134 | protein_coding | **338.565** | 0.671 | 0.202 | **1.592** | 1.84E-04 | **5.74E-04** |
| ENSG00000173039.22 | **RELA** | 5970 | protein_coding | **1688.139** | 0.339 | 0.102 | **1.265** | 1.84E-04 | **5.75E-04** |
| ENSG00000272282.2 | **LINC02084** | | lncRNA | **6.607** | 2.867 | 0.863 | **7.296** | 1.84E-04 | **5.75E-04** |
| ENSG00000287998.2 | **ENSG00000287998** | | lncRNA | **18.405** | 1.596 | 0.480 | **3.022** | 1.84E-04 | **5.75E-04** |
| ENSG00000235366.3 | **LINC01055** | | lncRNA | **150.461** | -0.689 | -0.208 | **0.620** | 1.85E-04 | **5.76E-04** |
| ENSG00000067369.16 | **TP53BP1** | 7158 | protein_coding | **960.992** | 0.495 | 0.149 | **1.409** | 1.85E-04 | **5.76E-04** |
| ENSG00000135913.13 | **USP37** | 57695 | protein_coding | **264.574** | -0.564 | -0.170 | **0.677** | 1.85E-04 | **5.76E-04** |

| | | | | | | | | | |
|---|---|---|---|---|---|---|---|---|---|
| ENSG00000293516.2 | **ZNF271P** | 10778 | lncRNA | **349.261** | -0.520 | -0.156 | **0.698** | 1.85E-04 | **5.76E-04** |
| ENSG00000197457.12 | **STMN3** | 50861 | protein_coding | **143.639** | 0.681 | 0.205 | **1.603** | 1.85E-04 | **5.78E-04** |
| ENSG00000130307.14 | **USHBP1** | 83878 | protein_coding | **14.044** | 1.631 | 0.491 | **3.098** | 1.86E-04 | **5.79E-04** |
| ENSG00000235078.2 | **ENSG00000235078** | | lncRNA | **14.042** | 1.530 | 0.461 | **2.888** | 1.86E-04 | **5.80E-04** |
| ENSG00000167447.15 | **SMG8** | 55181 | protein_coding | **175.267** | -0.665 | -0.200 | **0.631** | 1.86E-04 | **5.81E-04** |
| ENSG00000100628.14 | **ASB2** | 51676 | protein_coding | **261.640** | 1.084 | 0.326 | **2.120** | 1.87E-04 | **5.83E-04** |
| ENSG00000168887.13 | **C2orf68** | 388969 | protein_coding | **691.169** | -0.561 | -0.169 | **0.678** | 1.87E-04 | **5.83E-04** |
| ENSG00000260852.2 | **FBXL19-AS** | 283932 | lncRNA | **12.682** | -1.598 | -0.481 | **0.330** | 1.87E-04 | **5.83E-04** |
| ENSG00000219481.11 | **NBPF1** | 55672 | protein_coding | **44.887** | 1.267 | 0.381 | **2.407** | 1.89E-04 | **5.87E-04** |
| ENSG00000289028.3 | **ENSG00000289028** | | lncRNA | **13.495** | 1.512 | 0.455 | **2.852** | 1.90E-04 | **5.91E-04** |
| ENSG00000070614.17 | **NDST1** | 3340 | protein_coding | **438.600** | -0.517 | -0.156 | **0.699** | 1.91E-04 | **5.96E-04** |
| ENSG00000114841.18 | **DNAH1** | 25981 | protein_coding | **500.491** | 0.467 | 0.141 | **1.382** | 1.92E-04 | **5.97E-04** |
| ENSG00000115590.16 | **IL1R2** | 7850 | protein_coding | **37.025** | 1.444 | 0.435 | **2.721** | 1.93E-04 | **5.99E-04** |
| ENSG00000241935.11 | **HOGA1** | 112817 | protein_coding | **19.743** | 1.595 | 0.480 | **3.020** | 1.93E-04 | **6.01E-04** |
| ENSG00000267575.10 | **LINC02987** | 101927151 | lncRNA | **114.398** | -0.589 | -0.177 | **0.665** | 1.93E-04 | **6.01E-04** |
| ENSG00000151498.14 | **ACAD8** | 27034 | protein_coding | **470.906** | 0.478 | 0.144 | **1.393** | 1.94E-04 | **6.02E-04** |
| ENSG00000232732.12 | **ENSG0000(** | 101927687 | lncRNA | **5.521** | -2.999 | -0.903 | **0.125** | 1.94E-04 | **6.03E-04** |
| ENSG00000142233.15 | **NTN5** | 126147 | protein_coding | **10.865** | 2.234 | 0.673 | **4.705** | 1.94E-04 | **6.04E-04** |
| ENSG00000245468.6 | **LINC02447** | | lncRNA | **6.139** | 2.598 | 0.782 | **6.056** | 1.95E-04 | **6.05E-04** |
| ENSG00000111490.15 | **TBC1D30** | 23329 | protein_coding | **212.239** | 1.161 | 0.349 | **2.236** | 1.95E-04 | **6.07E-04** |
| ENSG00000254389.4 | **RHPN1-AS:** | 78998 | lncRNA | **7.380** | 2.373 | 0.714 | **5.180** | 1.96E-04 | **6.09E-04** |
| ENSG00000175054.18 | **ATR** | 545 | protein_coding | **494.212** | -0.429 | -0.129 | **0.743** | 1.97E-04 | **6.11E-04** |
| ENSG00000180712.6 | **LINC02363** | | lncRNA | **506.899** | 0.529 | 0.159 | **1.443** | 1.97E-04 | **6.11E-04** |
| ENSG00000067955.17 | **CBFB** | 865 | protein_coding | **386.652** | -0.476 | -0.143 | **0.719** | 1.97E-04 | **6.12E-04** |
| ENSG00000103342.15 | **GSPT1** | 2935 | protein_coding | **1470.506** | -0.433 | -0.130 | **0.741** | 1.97E-04 | **6.12E-04** |
| ENSG00000088682.16 | **COQ9** | 57017 | protein_coding | **620.768** | -0.396 | -0.119 | **0.760** | 1.97E-04 | **6.12E-04** |
| ENSG00000291041.2 | **ENSG00000291041** | | lncRNA | **9.751** | 2.036 | 0.613 | **4.101** | 1.97E-04 | **6.12E-04** |
| ENSG00000225791.8 | **TRAM2-AS** | 401264 | lncRNA | **97.510** | 0.865 | 0.260 | **1.822** | 1.97E-04 | **6.12E-04** |
| ENSG00000130305.19 | **NSUN5** | 55695 | protein_coding | **842.637** | -0.355 | -0.107 | **0.782** | 1.97E-04 | **6.13E-04** |
| ENSG00000228427.6 | **ENSG00000228427** | | lncRNA | **32.639** | -0.898 | -0.270 | **0.537** | 1.98E-04 | **6.14E-04** |
| ENSG00000001460.20 | **STPG1** | 90529 | protein_coding | **52.847** | 0.827 | 0.249 | **1.774** | 1.99E-04 | **6.18E-04** |

| | | | | | | | | | |
|---|---|---|---|---|---|---|---|---|---|
| ENSG00000171206.19 | **TRIM8** | 81603 | protein_coding | **864.558** | -0.464 | -0.140 | **0.725** | 1.99E-04 | **6.18E-04** |
| ENSG00000060069.20 | **CTDP1** | 9150 | protein_coding | **610.020** | 0.358 | 0.108 | **1.282** | 2.00E-04 | **6.19E-04** |
| ENSG00000198598.9 | **MMP17** | 4326 | protein_coding | **209.721** | 1.056 | 0.318 | **2.079** | 2.00E-04 | **6.22E-04** |
| ENSG00000179840.5 | **PIK3CD-AS** | 644997 | lncRNA | **48.718** | -0.892 | -0.269 | **0.539** | 2.01E-04 | **6.24E-04** |
| ENSG00000067829.22 | **IDH3G** | 3421 | protein_coding | **1287.771** | -0.490 | -0.148 | **0.712** | 2.02E-04 | **6.26E-04** |
| ENSG00000075391.20 | **RASAL2** | 9462 | protein_coding | **13.496** | 1.977 | 0.595 | **3.936** | 2.02E-04 | **6.26E-04** |
| ENSG00000296642.1 | **ENSG00000296642** | | lncRNA | **64.093** | 0.752 | 0.227 | **1.685** | 2.02E-04 | **6.27E-04** |
| ENSG00000030066.16 | **NUP160** | 23279 | protein_coding | **528.062** | -0.384 | -0.116 | **0.766** | 2.03E-04 | **6.29E-04** |
| ENSG00000291175.2 | **ENSG00000291175** | | lncRNA | **210.787** | -0.792 | -0.238 | **0.578** | 2.03E-04 | **6.29E-04** |
| ENSG00000250318.1 | **RPL13AP26** | | unprocessed_pseu | **20.197** | 1.203 | 0.362 | **2.302** | 2.04E-04 | **6.31E-04** |
| ENSG00000110324.14 | **IL10RA** | 3587 | protein_coding | **795.913** | 0.565 | 0.170 | **1.480** | 2.04E-04 | **6.32E-04** |
| ENSG00000122126.20 | **OCRL** | 4952 | protein_coding | **224.169** | 0.604 | 0.182 | **1.520** | 2.04E-04 | **6.32E-04** |
| ENSG00000099365.13 | **STX1B** | 112755 | protein_coding | **89.320** | 0.727 | 0.219 | **1.656** | 2.04E-04 | **6.33E-04** |
| ENSG00000180616.10 | **SSTR2** | 6752 | protein_coding | **209.266** | 0.830 | 0.250 | **1.778** | 2.05E-04 | **6.35E-04** |
| ENSG00000170632.16 | **ARMC10** | 83787 | protein_coding | **403.389** | -0.607 | -0.183 | **0.657** | 2.05E-04 | **6.35E-04** |
| ENSG00000136205.20 | **TNS3** | 64759 | protein_coding | **120.844** | 0.626 | 0.188 | **1.543** | 2.05E-04 | **6.35E-04** |
| ENSG00000053438.12 | **NNAT** | 4826 | protein_coding | **16.445** | 2.196 | 0.661 | **4.584** | 2.05E-04 | **6.35E-04** |
| ENSG00000181007.11 | **ZFP82** | 284406 | protein_coding | **107.252** | -0.584 | -0.176 | **0.667** | 2.05E-04 | **6.35E-04** |
| ENSG00000166855.11 | **CLPX** | 10845 | protein_coding | **546.227** | -0.699 | -0.210 | **0.616** | 2.06E-04 | **6.36E-04** |
| ENSG00000130021.17 | **PUDP** | 8226 | protein_coding | **379.264** | -0.390 | -0.117 | **0.763** | 2.06E-04 | **6.37E-04** |
| ENSG00000102898.15 | **NUTF2** | 10204 | protein_coding | **1673.230** | -0.434 | -0.131 | **0.740** | 2.06E-04 | **6.37E-04** |
| ENSG00000152582.16 | **SPEF2** | 79925 | protein_coding | **55.041** | 0.921 | 0.277 | **1.894** | 2.06E-04 | **6.37E-04** |
| ENSG00000135999.15 | **EPC2** | 26122 | protein_coding | **144.600** | -0.788 | -0.237 | **0.579** | 2.06E-04 | **6.37E-04** |
| ENSG00000303898.1 | **ENSG00000303898** | | lncRNA | **7.084** | 2.209 | 0.665 | **4.623** | 2.06E-04 | **6.38E-04** |
| ENSG00000184349.15 | **EFNA5** | 1946 | protein_coding | **150.206** | -0.831 | -0.250 | **0.562** | 2.07E-04 | **6.40E-04** |
| ENSG00000140876.13 | **NUDT7** | 283927 | protein_coding | **67.400** | 0.733 | 0.221 | **1.662** | 2.08E-04 | **6.43E-04** |
| ENSG00000140829.14 | **DHX38** | 9785 | protein_coding | **1426.949** | -0.395 | -0.119 | **0.761** | 2.09E-04 | **6.45E-04** |
| ENSG00000294141.1 | **ENSG00000294141** | | lncRNA | **8.665** | 2.156 | 0.649 | **4.458** | 2.10E-04 | **6.47E-04** |
| ENSG00000285012.2 | **ENSG00000285012** | | lncRNA | **71.068** | 0.941 | 0.283 | **1.919** | 2.10E-04 | **6.48E-04** |
| ENSG00000125650.5 | **PSPN** | 5623 | protein_coding | **67.985** | 1.047 | 0.315 | **2.067** | 2.10E-04 | **6.50E-04** |
| ENSG00000204444.12 | **APOM** | 55937 | protein_coding | **77.484** | -0.756 | -0.228 | **0.592** | 2.11E-04 | **6.50E-04** |

| | | | | | | | | | |
|---|---|---|---|---|---|---|---|---|---|
| ENSG00000271889.2 | **C2orf74-AS** | 10537475 | lncRNA | **191.937** | -0.613 | -0.185 | **0.654** | 2.11E-04 | **6.50E-04** |
| ENSG00000151881.17 | **TMEM267** | 64417 | protein_coding | **159.952** | -0.830 | -0.250 | **0.562** | 2.11E-04 | **6.50E-04** |
| ENSG00000266028.10 | **SRGAP2** | 23380 | protein_coding | **315.659** | 0.597 | 0.180 | **1.512** | 2.11E-04 | **6.50E-04** |
| ENSG00000179271.5 | **GADD45GI** | 90480 | protein_coding | **1168.892** | -0.502 | -0.151 | **0.706** | 2.11E-04 | **6.51E-04** |
| ENSG00000108984.18 | **MAP2K6** | 5608 | protein_coding | **372.926** | -1.464 | -0.441 | **0.363** | 2.11E-04 | **6.52E-04** |
| ENSG00000231074.12 | **HCG18** | 414777 | lncRNA | **342.885** | -0.586 | -0.177 | **0.666** | 2.12E-04 | **6.54E-04** |
| ENSG00000164715.8 | **LMTK2** | 22853 | protein_coding | **269.115** | 0.500 | 0.150 | **1.414** | 2.12E-04 | **6.54E-04** |
| ENSG00000240758.3 | **HILPDA-AS1** | | lncRNA | **46.495** | 1.005 | 0.302 | **2.007** | 2.13E-04 | **6.55E-04** |
| ENSG00000180914.12 | **OXTR** | 5021 | protein_coding | **162.351** | -0.719 | -0.216 | **0.608** | 2.13E-04 | **6.57E-04** |
| ENSG00000104432.17 | **IL7** | 3574 | protein_coding | **49.271** | -0.915 | -0.275 | **0.530** | 2.14E-04 | **6.58E-04** |
| ENSG00000104825.19 | **NFKBIB** | 4793 | protein_coding | **636.951** | -0.444 | -0.134 | **0.735** | 2.14E-04 | **6.58E-04** |
| ENSG00000107833.13 | **NPM3** | 10360 | protein_coding | **559.285** | -0.563 | -0.170 | **0.677** | 2.14E-04 | **6.59E-04** |
| ENSG00000173868.13 | **PHOSPHO1** | 162466 | protein_coding | **10.442** | 2.201 | 0.663 | **4.598** | 2.14E-04 | **6.59E-04** |
| ENSG00000243696.5 | **ENSG00000243696** | | protein_coding | **158.452** | 0.664 | 0.200 | **1.584** | 2.14E-04 | **6.59E-04** |
| ENSG00000248727.7 | **LINC01948** | | lncRNA | **7.913** | 3.058 | 0.920 | **8.326** | 2.14E-04 | **6.60E-04** |
| ENSG00000163686.17 | **ABHD6** | 57406 | protein_coding | **227.738** | 0.568 | 0.171 | **1.482** | 2.15E-04 | **6.62E-04** |
| ENSG00000080345.20 | **RIF1** | 55183 | protein_coding | **475.379** | -0.445 | -0.134 | **0.735** | 2.15E-04 | **6.62E-04** |
| ENSG00000170633.19 | **RNF34** | 80196 | protein_coding | **595.076** | -0.324 | -0.098 | **0.799** | 2.16E-04 | **6.63E-04** |
| ENSG00000089682.20 | **RBM41** | 55285 | protein_coding | **291.935** | -0.601 | -0.181 | **0.659** | 2.16E-04 | **6.65E-04** |
| ENSG00000185551.17 | **NR2F2** | 7026 | protein_coding | **159.600** | 0.858 | 0.258 | **1.813** | 2.16E-04 | **6.65E-04** |
| ENSG00000234156.4 | **ENSG00000234156** | | lncRNA | **18.668** | 1.784 | 0.537 | **3.444** | 2.17E-04 | **6.66E-04** |
| ENSG00000104921.16 | **FCER2** | 2208 | protein_coding | **21224.160** | 0.616 | 0.185 | **1.533** | 2.17E-04 | **6.67E-04** |
| ENSG00000162819.14 | **BROX** | 148362 | protein_coding | **289.740** | 0.477 | 0.143 | **1.391** | 2.17E-04 | **6.68E-04** |
| ENSG00000110619.20 | **CARS1** | 833 | protein_coding | **1637.283** | -0.621 | -0.187 | **0.650** | 2.21E-04 | **6.78E-04** |
| ENSG00000295910.1 | **ENSG00000295910** | | lncRNA | **6.284** | 2.311 | 0.696 | **4.963** | 2.21E-04 | **6.79E-04** |
| ENSG00000230679.2 | **ENO1-AS1** | | lncRNA | **14.048** | 1.537 | 0.463 | **2.902** | 2.21E-04 | **6.80E-04** |
| ENSG00000144048.14 | **DUSP11** | 8446 | protein_coding | **353.704** | -0.493 | -0.148 | **0.711** | 2.21E-04 | **6.80E-04** |
| ENSG00000170439.9 | **TMT1B** | 196410 | protein_coding | **11.863** | 1.893 | 0.570 | **3.714** | 2.21E-04 | **6.80E-04** |
| ENSG00000259802.4 | **MARCHF6-DT** | | lncRNA | **43.141** | 0.881 | 0.265 | **1.841** | 2.21E-04 | **6.80E-04** |
| ENSG00000151835.18 | **SACS** | 26278 | protein_coding | **456.232** | -0.537 | -0.162 | **0.689** | 2.21E-04 | **6.80E-04** |
| ENSG00000152078.12 | **TLCD4** | 148534 | protein_coding | **8.990** | 1.986 | 0.598 | **3.961** | 2.21E-04 | **6.80E-04** |

| | | | | | | | | | |
|---|---|---|---|---|---|---|---|---|---|
| ENSG00000275409.1 | **ENSG00000275409** | | lncRNA | **7.498** | 2.026 | 0.610 | **4.073** | 2.22E-04 | **6.81E-04** |
| ENSG00000229989.7 | **MIR181A1HG** | | lncRNA | **26.021** | 1.612 | 0.485 | **3.056** | 2.22E-04 | **6.81E-04** |
| ENSG00000282807.4 | **ENSG00000282807** | | lncRNA | **7.348** | 3.275 | 0.986 | **9.677** | 2.24E-04 | **6.89E-04** |
| ENSG00000186432.12 | **KPNA4** | 3840 | protein_coding | **626.109** | -0.486 | -0.146 | **0.714** | 2.24E-04 | **6.89E-04** |
| ENSG00000136167.17 | **LCP1** | 3936 | protein_coding | **29770.540** | 0.534 | 0.161 | **1.448** | 2.24E-04 | **6.89E-04** |
| ENSG00000205129.11 | **CFAP96** | 441054 | protein_coding | **57.028** | -1.190 | -0.358 | **0.438** | 2.25E-04 | **6.89E-04** |
| ENSG00000296743.1 | **ENSG00000296743** | | lncRNA | **14.207** | 1.667 | 0.502 | **3.176** | 2.25E-04 | **6.90E-04** |
| ENSG00000155428.13 | **TRIM74** | 378108 | protein_coding | **19.638** | 1.613 | 0.485 | **3.058** | 2.25E-04 | **6.92E-04** |
| ENSG00000166598.18 | **HSP90B1** | 7184 | protein_coding | **13610.891** | -0.474 | -0.143 | **0.720** | 2.26E-04 | **6.93E-04** |
| ENSG00000260025.3 | **CRIM1-DT** | | lncRNA | **16.999** | 1.762 | 0.531 | **3.392** | 2.26E-04 | **6.93E-04** |
| ENSG00000166902.7 | **MRPL16** | 54948 | protein_coding | **1304.233** | -0.465 | -0.140 | **0.725** | 2.26E-04 | **6.93E-04** |
| ENSG00000102760.15 | **RGCC** | 28984 | protein_coding | **84.893** | -0.838 | -0.252 | **0.560** | 2.26E-04 | **6.93E-04** |
| ENSG00000102796.14 | **DHRS12** | 79758 | protein_coding | **275.015** | 0.505 | 0.152 | **1.419** | 2.26E-04 | **6.93E-04** |
| ENSG00000123130.19 | **ACOT9** | 23597 | protein_coding | **880.965** | 0.346 | 0.104 | **1.271** | 2.27E-04 | **6.94E-04** |
| ENSG00000132688.13 | **NES** | 10763 | protein_coding | **4.814** | 2.998 | 0.902 | **7.988** | 2.27E-04 | **6.94E-04** |
| ENSG00000163121.12 | **NEURL3** | 93082 | protein_coding | **166.728** | -0.698 | -0.210 | **0.616** | 2.27E-04 | **6.96E-04** |
| ENSG00000119227.10 | **PIGZ** | 80235 | protein_coding | **215.398** | 0.587 | 0.177 | **1.502** | 2.28E-04 | **6.97E-04** |
| ENSG00000126821.10 | **SGPP1** | 81537 | protein_coding | **249.981** | 0.628 | 0.189 | **1.545** | 2.28E-04 | **6.97E-04** |
| ENSG00000163644.17 | **PPM1K** | 152926 | protein_coding | **663.671** | -0.527 | -0.159 | **0.694** | 2.28E-04 | **6.97E-04** |
| ENSG00000122484.11 | **RPAP2** | 79871 | protein_coding | **312.806** | -0.513 | -0.154 | **0.701** | 2.29E-04 | **7.00E-04** |
| ENSG00000181192.14 | **DHTKD1** | 55526 | protein_coding | **1220.498** | 0.391 | 0.118 | **1.311** | 2.29E-04 | **7.00E-04** |
| ENSG00000185104.23 | **FAF1** | 11124 | protein_coding | **978.927** | -0.344 | -0.104 | **0.788** | 2.29E-04 | **7.01E-04** |
| ENSG00000007402.14 | **CACNA2D2** | 9254 | protein_coding | **5.704** | 2.565 | 0.772 | **5.919** | 2.29E-04 | **7.01E-04** |
| ENSG00000188846.15 | **RPL14** | 9045 | protein_coding | **23926.480** | -0.445 | -0.134 | **0.734** | 2.30E-04 | **7.02E-04** |
| ENSG00000007923.19 | **DNAJC11** | 55735 | protein_coding | **1307.281** | -0.336 | -0.101 | **0.792** | 2.30E-04 | **7.03E-04** |
| ENSG00000168952.18 | **STXBP6** | 29091 | protein_coding | **63.922** | 1.147 | 0.345 | **2.215** | 2.30E-04 | **7.04E-04** |
| ENSG00000177182.13 | **CLVS1** | 157807 | protein_coding | **35.504** | 1.485 | 0.447 | **2.800** | 2.31E-04 | **7.06E-04** |
| ENSG00000099194.8 | **SCD** | 6319 | protein_coding | **5564.751** | -0.786 | -0.237 | **0.580** | 2.31E-04 | **7.06E-04** |
| ENSG00000196353.15 | **CPNE4** | 131034 | protein_coding | **5.754** | 3.020 | 0.909 | **8.114** | 2.32E-04 | **7.08E-04** |
| ENSG00000204472.15 | **AIF1** | 199 | protein_coding | **267.621** | -0.672 | -0.202 | **0.628** | 2.32E-04 | **7.08E-04** |
| ENSG00000230432.2 | **ENSG00000230432** | | lncRNA | **10.421** | -2.150 | -0.647 | **0.225** | 2.33E-04 | **7.12E-04** |

| | | | | | | | | | |
|---|---|---|---|---|---|---|---|---|---|
| ENSG00000133997.14 | **MED6** | 10001 | protein_coding | **496.221** | -0.330 | -0.099 | **0.796** | 2.33E-04 | **7.12E-04** |
| ENSG00000167657.16 | **DAPK3** | 1613 | protein_coding | **581.902** | 0.415 | 0.125 | **1.334** | 2.35E-04 | **7.19E-04** |
| ENSG00000179240.14 | **GVQW3** | 10050612[illegible] | protein_coding | **141.154** | 0.693 | 0.209 | **1.616** | 2.36E-04 | **7.19E-04** |
| ENSG00000263033.2 | **ENSG00000263033** | | lncRNA | **10.879** | 1.798 | 0.541 | **3.476** | 2.37E-04 | **7.22E-04** |
| ENSG00000293463.2 | **ENSG00000293463** | | lncRNA | **3.279** | -4.406 | -1.326 | **0.047** | 2.37E-04 | **7.23E-04** |
| ENSG00000245248.11 | **USP2-AS1** | | lncRNA | **5.178** | -3.424 | -1.031 | **0.093** | 2.37E-04 | **7.23E-04** |
| ENSG00000307533.1 | **ENSG00000307533** | | lncRNA | **6.746** | 2.371 | 0.714 | **5.173** | 2.38E-04 | **7.26E-04** |
| ENSG00000188404.12 | **SELL** | 6402 | protein_coding | **3270.289** | -0.550 | -0.165 | **0.683** | 2.38E-04 | **7.27E-04** |
| ENSG00000179820.19 | **MYADM** | 91663 | protein_coding | **354.255** | 0.474 | 0.143 | **1.389** | 2.39E-04 | **7.30E-04** |
| ENSG00000140326.15 | **CDAN1** | 146059 | protein_coding | **258.227** | -0.418 | -0.126 | **0.748** | 2.39E-04 | **7.30E-04** |
| ENSG00000275437.1 | **ENSG00000275437** | | lncRNA | **21.208** | -1.087 | -0.327 | **0.471** | 2.40E-04 | **7.31E-04** |
| ENSG00000235288.4 | **ENSG00000235288** | | lncRNA | **187.630** | -0.650 | -0.196 | **0.637** | 2.40E-04 | **7.33E-04** |
| ENSG00000173482.18 | **PTPRM** | 5797 | protein_coding | **39.989** | 1.447 | 0.436 | **2.727** | 2.41E-04 | **7.34E-04** |
| ENSG00000225285.2 | **LINC01770** | 10272431[illegible] | lncRNA | **28.200** | 1.471 | 0.443 | **2.773** | 2.41E-04 | **7.35E-04** |
| ENSG00000103657.15 | **HERC1** | 8925 | protein_coding | **1551.303** | 0.773 | 0.233 | **1.709** | 2.42E-04 | **7.37E-04** |
| ENSG00000153029.18 | **MR1** | 3140 | protein_coding | **770.629** | -0.564 | -0.170 | **0.676** | 2.42E-04 | **7.38E-04** |
| ENSG00000114757.20 | **PEX5L** | 51555 | protein_coding | **85.298** | -1.341 | -0.404 | **0.395** | 2.43E-04 | **7.40E-04** |
| ENSG00000141504.14 | **SAT2** | 112483 | protein_coding | **836.877** | -0.494 | -0.149 | **0.710** | 2.43E-04 | **7.40E-04** |
| ENSG00000213199.10 | **ASIC3** | 9311 | protein_coding | **117.179** | 0.789 | 0.238 | **1.728** | 2.44E-04 | **7.42E-04** |
| ENSG00000158769.21 | **F11R** | 50848 | protein_coding | **1229.513** | 0.449 | 0.135 | **1.365** | 2.44E-04 | **7.43E-04** |
| ENSG00000124120.13 | **TTPAL** | 79183 | protein_coding | **317.188** | 0.476 | 0.143 | **1.391** | 2.44E-04 | **7.43E-04** |
| ENSG00000142606.17 | **MMEL1** | 79258 | protein_coding | **18.211** | 2.217 | 0.667 | **4.648** | 2.45E-04 | **7.45E-04** |
| ENSG00000177173.5 | **NAP1L4P1** | | processed_pseud[illegible] | **15.510** | -1.736 | -0.523 | **0.300** | 2.45E-04 | **7.46E-04** |
| ENSG00000173436.17 | **MICOS10** | 440574 | protein_coding | **897.261** | -0.579 | -0.174 | **0.669** | 2.46E-04 | **7.50E-04** |
| ENSG00000130203.12 | **APOE** | 348 | protein_coding | **44.133** | 1.246 | 0.375 | **2.372** | 2.47E-04 | **7.50E-04** |
| ENSG00000124508.19 | **BTN2A2** | 10385 | protein_coding | **953.511** | 0.400 | 0.120 | **1.319** | 2.47E-04 | **7.52E-04** |
| ENSG00000177485.9 | **ZBTB33** | 10009 | protein_coding | **202.107** | -0.484 | -0.146 | **0.715** | 2.47E-04 | **7.52E-04** |
| ENSG00000008869.14 | **HEATR5B** | 54497 | protein_coding | **531.428** | 0.497 | 0.150 | **1.411** | 2.47E-04 | **7.53E-04** |
| ENSG00000141985.11 | **SH3GL1** | 6455 | protein_coding | **1382.435** | -0.457 | -0.138 | **0.728** | 2.48E-04 | **7.54E-04** |
| ENSG00000101182.17 | **PSMA7** | 5688 | protein_coding | **1782.125** | -0.423 | -0.127 | **0.746** | 2.48E-04 | **7.55E-04** |
| ENSG00000264964.3 | **TWSG1-DT** | 10272439[illegible] | lncRNA | **30.933** | 1.089 | 0.328 | **2.127** | 2.49E-04 | **7.57E-04** |

| | | | | | | | | | |
|---|---|---|---|---|---|---|---|---|---|
| ENSG00000115107.22 | **STEAP3** | 55240 | protein_coding | **206.279** | -0.609 | -0.183 | **0.656** | 2.49E-04 | **7.57E-04** |
| ENSG00000254416.7 | **LINC02732** | 124902752 | lncRNA | **45.310** | 1.189 | 0.358 | **2.279** | 2.49E-04 | **7.58E-04** |
| ENSG00000161653.13 | **NAGS** | 162417 | protein_coding | **33.498** | 0.977 | 0.294 | **1.968** | 2.50E-04 | **7.60E-04** |
| ENSG00000276672.1 | **ENSG00000276672** | | lncRNA | **24.338** | -1.089 | -0.328 | **0.470** | 2.50E-04 | **7.60E-04** |
| ENSG00000235072.3 | **ARNILA** | | lncRNA | **3.717** | 3.560 | 1.072 | **11.792** | 2.51E-04 | **7.64E-04** |
| ENSG00000053918.22 | **KCNQ1** | 3784 | protein_coding | **28.975** | 1.318 | 0.397 | **2.493** | 2.52E-04 | **7.65E-04** |
| ENSG00000173992.11 | **CCS** | 9973 | protein_coding | **1037.356** | 0.413 | 0.124 | **1.332** | 2.52E-04 | **7.66E-04** |
| ENSG00000203710.14 | **CR1** | 1378 | protein_coding | **97.135** | -0.842 | -0.253 | **0.558** | 2.52E-04 | **7.66E-04** |
| ENSG00000142669.17 | **SH3BGRL3** | 83442 | protein_coding | **4305.809** | 0.662 | 0.199 | **1.582** | 2.53E-04 | **7.67E-04** |
| ENSG00000257595.4 | **LINC02356** | 10536998[illegible] | lncRNA | **7.054** | 2.262 | 0.681 | **4.797** | 2.53E-04 | **7.68E-04** |
| ENSG00000213347.13 | **MXD3** | 83463 | protein_coding | **631.502** | -0.417 | -0.125 | **0.749** | 2.54E-04 | **7.69E-04** |
| ENSG00000259705.2 | **FBN1-DT** | | lncRNA | **14.916** | 1.467 | 0.442 | **2.764** | 2.54E-04 | **7.70E-04** |
| ENSG00000168461.15 | **RAB31** | 11031 | protein_coding | **555.068** | 0.498 | 0.150 | **1.412** | 2.54E-04 | **7.70E-04** |
| ENSG00000162490.9 | **DRAXIN** | 374946 | protein_coding | **8.465** | -2.328 | -0.701 | **0.199** | 2.54E-04 | **7.72E-04** |
| ENSG00000300688.1 | **ENSG00000300688** | | lncRNA | **9.896** | 1.913 | 0.576 | **3.767** | 2.55E-04 | **7.73E-04** |
| ENSG00000141499.20 | **WRAP53** | 55135 | protein_coding | **327.534** | -0.623 | -0.187 | **0.650** | 2.55E-04 | **7.73E-04** |
| ENSG00000196562.18 | **SULF2** | 55959 | protein_coding | **148.431** | -1.256 | -0.378 | **0.419** | 2.55E-04 | **7.75E-04** |
| ENSG00000133424.24 | **LARGE1** | 9215 | protein_coding | **215.239** | 0.583 | 0.175 | **1.498** | 2.56E-04 | **7.75E-04** |
| ENSG00000061676.18 | **NCKAP1** | 10787 | protein_coding | **152.538** | 0.881 | 0.265 | **1.842** | 2.56E-04 | **7.76E-04** |
| ENSG00000163029.18 | **SMC6** | 79677 | protein_coding | **1036.448** | 0.650 | 0.196 | **1.569** | 2.56E-04 | **7.77E-04** |
| ENSG00000284906.3 | **ENSG00000284906** | | protein_coding | **60.603** | -1.444 | -0.435 | **0.367** | 2.56E-04 | **7.77E-04** |
| ENSG00000176915.16 | **ANKLE2** | 23141 | protein_coding | **1787.518** | -0.297 | -0.089 | **0.814** | 2.57E-04 | **7.79E-04** |
| ENSG00000260063.2 | **ENSG00000260063** | | lncRNA | **27.839** | 1.106 | 0.333 | **2.152** | 2.57E-04 | **7.80E-04** |
| ENSG00000297800.1 | **ENSG00000297800** | | lncRNA | **5.115** | 3.435 | 1.034 | **10.813** | 2.58E-04 | **7.82E-04** |
| ENSG00000164938.16 | **TP53INP1** | 94241 | protein_coding | **538.530** | 0.573 | 0.173 | **1.488** | 2.58E-04 | **7.83E-04** |
| ENSG00000270972.1 | **ENSG00000270972** | | lncRNA | **27.489** | 1.127 | 0.339 | **2.184** | 2.59E-04 | **7.83E-04** |
| ENSG00000231925.16 | **TAPBP** | 6892 | protein_coding | **5219.350** | -0.470 | -0.141 | **0.722** | 2.59E-04 | **7.83E-04** |
| ENSG00000100528.14 | **CNIH1** | 10175 | protein_coding | **371.661** | -0.724 | -0.218 | **0.605** | 2.59E-04 | **7.84E-04** |
| ENSG00000137770.16 | **CTDSPL2** | 51496 | protein_coding | **299.973** | -0.598 | -0.180 | **0.661** | 2.60E-04 | **7.88E-04** |
| ENSG00000101335.12 | **MYL9** | 10398 | protein_coding | **189.680** | 0.749 | 0.225 | **1.680** | 2.61E-04 | **7.89E-04** |
| ENSG00000087842.13 | **PIR** | 8544 | protein_coding | **347.225** | -0.688 | -0.207 | **0.621** | 2.61E-04 | **7.90E-04** |

| | | | | | | | | | |
|---|---|---|---|---|---|---|---|---|---|
| ENSG00000187581.4 | **COX8C** | 341947 | protein_coding | **8.386** | 3.220 | 0.969 | **9.318** | 2.61E-04 | **7.91E-04** |
| ENSG00000111642.18 | **CHD4** | 1108 | protein_coding | **4084.061** | 0.283 | 0.085 | **1.217** | 2.62E-04 | **7.92E-04** |
| ENSG00000234380.4 | **LINC01426** | 10050638 | lncRNA | **7.033** | 3.098 | 0.933 | **8.563** | 2.63E-04 | **7.95E-04** |
| ENSG00000273456.3 | **ENSG00000273456** | | lncRNA | **59.827** | 0.871 | 0.262 | **1.828** | 2.64E-04 | **7.98E-04** |
| ENSG00000261335.2 | **ENSG0000** | 10527430 | lncRNA | **15.197** | 1.725 | 0.519 | **3.306** | 2.64E-04 | **7.98E-04** |
| ENSG00000245937.11 | **SLC12A2-D** | 644873 | lncRNA | **639.578** | -0.471 | -0.142 | **0.721** | 2.64E-04 | **7.99E-04** |
| ENSG00000115844.12 | **DLX2** | 1746 | protein_coding | **8.672** | 2.041 | 0.615 | **4.117** | 2.65E-04 | **8.01E-04** |
| ENSG00000259630.2 | **FABP5P9** | | transcribed_proce | **10.895** | 1.756 | 0.529 | **3.378** | 2.66E-04 | **8.02E-04** |
| ENSG00000132024.20 | **CC2D1A** | 54862 | protein_coding | **1048.941** | 0.404 | 0.122 | **1.323** | 2.66E-04 | **8.04E-04** |
| ENSG00000163666.12 | **HESX1** | 8820 | protein_coding | **58.142** | 0.966 | 0.291 | **1.953** | 2.66E-04 | **8.05E-04** |
| ENSG00000310465.1 | **ENSG00000310465** | | lncRNA | **4.993** | 2.587 | 0.779 | **6.007** | 2.67E-04 | **8.07E-04** |
| ENSG00000251485.1 | **NRBF2P6** | | processed_pseud | **24.042** | 1.179 | 0.355 | **2.265** | 2.67E-04 | **8.07E-04** |
| ENSG00000231663.4 | **COA6-AS1** | | lncRNA | **77.461** | 0.819 | 0.247 | **1.764** | 2.68E-04 | **8.10E-04** |
| ENSG00000197872.13 | **CYRIA** | 81553 | protein_coding | **17.103** | 2.116 | 0.637 | **4.335** | 2.69E-04 | **8.11E-04** |
| ENSG00000101911.15 | **PRPS2** | 5634 | protein_coding | **628.362** | -0.430 | -0.129 | **0.742** | 2.69E-04 | **8.12E-04** |
| ENSG00000167131.19 | **DNAAF19** | 388389 | protein_coding | **40.852** | 1.017 | 0.306 | **2.024** | 2.70E-04 | **8.14E-04** |
| ENSG00000261126.7 | **RBFADN** | | lncRNA | **13.527** | 1.724 | 0.519 | **3.302** | 2.71E-04 | **8.16E-04** |
| ENSG00000163794.6 | **UCN** | 7349 | protein_coding | **67.858** | 0.862 | 0.259 | **1.818** | 2.71E-04 | **8.16E-04** |
| ENSG00000150768.20 | **DLAT** | 1737 | protein_coding | **288.714** | -0.708 | -0.213 | **0.612** | 2.71E-04 | **8.17E-04** |
| ENSG00000285840.3 | **ENSG00000285840** | | lncRNA | **100.146** | -0.919 | -0.277 | **0.529** | 2.71E-04 | **8.17E-04** |
| ENSG00000178977.5 | **LINC00324** | 284029 | lncRNA | **63.044** | 0.748 | 0.225 | **1.680** | 2.71E-04 | **8.18E-04** |
| ENSG00000187801.17 | **ZFP69B** | 65243 | protein_coding | **45.826** | -0.968 | -0.291 | **0.511** | 2.71E-04 | **8.18E-04** |
| ENSG00000300133.1 | **ENSG00000300133** | | lncRNA | **72.343** | 0.934 | 0.281 | **1.911** | 2.71E-04 | **8.18E-04** |
| ENSG00000008735.16 | **MAPK8IP2** | 23542 | protein_coding | **23.057** | 1.316 | 0.396 | **2.490** | 2.72E-04 | **8.19E-04** |
| ENSG00000158786.6 | **PLA2G2F** | 64600 | protein_coding | **6.704** | 2.273 | 0.684 | **4.832** | 2.73E-04 | **8.22E-04** |
| ENSG00000152422.18 | **XRCC4** | 7518 | protein_coding | **161.282** | -0.632 | -0.190 | **0.645** | 2.73E-04 | **8.22E-04** |
| ENSG00000269534.5 | **ENSG00000269534** | | lncRNA | **3.490** | 4.626 | 1.393 | **24.699** | 2.73E-04 | **8.22E-04** |
| ENSG00000049656.16 | **CLPTM1L** | 81037 | protein_coding | **1016.935** | -0.319 | -0.096 | **0.802** | 2.73E-04 | **8.23E-04** |
| ENSG00000148335.17 | **NTMT1** | 28989 | protein_coding | **364.079** | -0.490 | -0.147 | **0.712** | 2.73E-04 | **8.23E-04** |
| ENSG00000143793.15 | **C1orf35** | 79169 | protein_coding | **516.590** | 0.376 | 0.113 | **1.297** | 2.74E-04 | **8.25E-04** |
| ENSG00000125841.16 | **NRSN2** | 80023 | protein_coding | **419.035** | -0.609 | -0.183 | **0.656** | 2.74E-04 | **8.26E-04** |

| | | | | | | | | | |
|---|---|---|---|---|---|---|---|---|---|
| ENSG00000274294.1 | **ENSG00000274294** | | lncRNA | **5.525** | 2.621 | 0.789 | **6.152** | 2.75E-04 | **8.27E-04** |
| ENSG00000273274.4 | **ZBTB8B** | 728116 | protein_coding | **6.211** | 2.817 | 0.848 | **7.048** | 2.75E-04 | **8.27E-04** |
| ENSG00000271553.1 | **ENSG00000271553** | | lncRNA | **16.708** | 1.297 | 0.390 | **2.456** | 2.76E-04 | **8.31E-04** |
| ENSG00000197111.18 | **PCBP2** | 5094 | protein_coding | **7849.324** | -0.321 | -0.096 | **0.801** | 2.77E-04 | **8.33E-04** |
| ENSG00000188938.18 | **FAM120AC** | 158293 | protein_coding | **840.484** | -0.401 | -0.121 | **0.757** | 2.77E-04 | **8.34E-04** |
| ENSG00000149922.13 | **TBX6** | 6911 | protein_coding | **107.406** | 0.764 | 0.230 | **1.698** | 2.78E-04 | **8.35E-04** |
| ENSG00000147124.15 | **ZNF41** | 7592 | protein_coding | **176.746** | -0.501 | -0.151 | **0.707** | 2.78E-04 | **8.35E-04** |
| ENSG00000132517.17 | **SLC52A1** | 55065 | protein_coding | **7.668** | -2.455 | -0.739 | **0.182** | 2.78E-04 | **8.37E-04** |
| ENSG00000261824.9 | **LINC00662** | 148189 | lncRNA | **134.587** | -0.659 | -0.198 | **0.633** | 2.79E-04 | **8.38E-04** |
| ENSG00000168502.20 | **MTCL1** | 23255 | protein_coding | **8.293** | 2.456 | 0.739 | **5.488** | 2.79E-04 | **8.38E-04** |
| ENSG00000260778.6 | **MIR3677HG** | | lncRNA | **47.125** | 0.915 | 0.275 | **1.885** | 2.79E-04 | **8.39E-04** |
| ENSG00000232828.2 | **ENSG00000232828** | | lncRNA | **12.637** | 1.529 | 0.460 | **2.886** | 2.79E-04 | **8.40E-04** |
| ENSG00000179632.13 | **MAF1** | 84232 | protein_coding | **1237.148** | -0.302 | -0.091 | **0.811** | 2.79E-04 | **8.40E-04** |
| ENSG00000298816.1 | **ENSG00000298816** | | lncRNA | **50.803** | 0.866 | 0.261 | **1.822** | 2.80E-04 | **8.41E-04** |
| ENSG00000135436.8 | **FAM186B** | 84070 | protein_coding | **20.523** | 1.227 | 0.369 | **2.342** | 2.80E-04 | **8.41E-04** |
| ENSG00000134575.15 | **ACP2** | 53 | protein_coding | **285.606** | 0.439 | 0.132 | **1.356** | 2.80E-04 | **8.42E-04** |
| ENSG00000275763.3 | **ZNF516-DT** | 400658 | lncRNA | **12.001** | 1.483 | 0.446 | **2.796** | 2.81E-04 | **8.44E-04** |
| ENSG00000167977.11 | **KCTD5** | 54442 | protein_coding | **939.653** | -0.367 | -0.110 | **0.775** | 2.81E-04 | **8.44E-04** |
| ENSG00000009954.12 | **BAZ1B** | 9031 | protein_coding | **1677.183** | -0.375 | -0.113 | **0.771** | 2.81E-04 | **8.45E-04** |
| ENSG00000125485.20 | **DDX31** | 64794 | protein_coding | **541.903** | -0.412 | -0.124 | **0.752** | 2.81E-04 | **8.45E-04** |
| ENSG00000172331.14 | **BPGM** | 669 | protein_coding | **279.820** | 0.416 | 0.125 | **1.334** | 2.82E-04 | **8.45E-04** |
| ENSG00000070081.20 | **NUCB2** | 4925 | protein_coding | **397.779** | -0.447 | -0.135 | **0.734** | 2.83E-04 | **8.49E-04** |
| ENSG00000254508.6 | **FBXO3-DT** | 10537661 | lncRNA | **5.672** | 2.940 | 0.885 | **7.674** | 2.84E-04 | **8.52E-04** |
| ENSG00000180543.5 | **TSPYL5** | 85453 | protein_coding | **107.864** | 0.647 | 0.195 | **1.566** | 2.86E-04 | **8.59E-04** |
| ENSG00000281398.4 | **SNHG4** | 724102 | lncRNA | **794.615** | -0.405 | -0.122 | **0.755** | 2.87E-04 | **8.60E-04** |
| ENSG00000303375.1 | **ENSG00000303375** | | lncRNA | **7.511** | 3.199 | 0.963 | **9.183** | 2.87E-04 | **8.60E-04** |
| ENSG00000126003.8 | **PLAGL2** | 5326 | protein_coding | **638.909** | -0.506 | -0.152 | **0.704** | 2.88E-04 | **8.64E-04** |
| ENSG00000100591.10 | **AHSA1** | 10598 | protein_coding | **2160.031** | -0.515 | -0.155 | **0.700** | 2.88E-04 | **8.65E-04** |
| ENSG00000166619.17 | **BLCAP** | 10904 | protein_coding | **970.758** | 0.361 | 0.109 | **1.284** | 2.89E-04 | **8.66E-04** |
| ENSG00000071794.18 | **HLTF** | 6596 | protein_coding | **682.640** | 0.460 | 0.138 | **1.375** | 2.89E-04 | **8.67E-04** |
| ENSG00000143363.19 | **PRUNE1** | 58497 | protein_coding | **228.218** | 0.463 | 0.139 | **1.378** | 2.89E-04 | **8.67E-04** |

| | | | | | | | | | |
|---|---|---|---|---|---|---|---|---|---|
| ENSG00000302808.1 | **ENSG0000(** | 10537139[illegible] | lncRNA | **8.549** | 1.963 | 0.591 | **3.898** | 2.90E-04 | **8.70E-04** |
| ENSG00000236104.6 | **ZBTB22** | 9278 | protein_coding | **395.716** | 0.405 | 0.122 | **1.324** | 2.91E-04 | **8.71E-04** |
| ENSG00000278730.1 | **ENSG00000278730** | | lncRNA | **81.387** | 0.826 | 0.249 | **1.773** | 2.92E-04 | **8.74E-04** |
| ENSG00000288848.3 | **ENSG00000288848** | | lncRNA | **11.940** | 2.151 | 0.647 | **4.441** | 2.92E-04 | **8.74E-04** |
| ENSG00000091947.12 | **TMEM101** | 84336 | protein_coding | **393.125** | -0.397 | -0.119 | **0.760** | 2.92E-04 | **8.74E-04** |
| ENSG00000173163.13 | **COMMD1** | 150684 | protein_coding | **567.766** | -0.462 | -0.139 | **0.726** | 2.93E-04 | **8.76E-04** |
| ENSG00000131467.14 | **PSME3** | 10197 | protein_coding | **1487.532** | -0.394 | -0.119 | **0.761** | 2.93E-04 | **8.76E-04** |
| ENSG00000186395.10 | **KRT10** | 3858 | protein_coding | **314.436** | -0.472 | -0.142 | **0.721** | 2.93E-04 | **8.76E-04** |
| ENSG00000224220.2 | **DTNB-AS1** | | lncRNA | **789.195** | -0.686 | -0.207 | **0.621** | 2.93E-04 | **8.77E-04** |
| ENSG00000010404.20 | **IDS** | 3423 | protein_coding | **1782.253** | 0.389 | 0.117 | **1.309** | 2.94E-04 | **8.78E-04** |
| ENSG00000243679.1 | **ENSG00000243679** | | transcribed_proce | **222.464** | -0.759 | -0.229 | **0.591** | 2.94E-04 | **8.78E-04** |
| ENSG00000104611.14 | **SH2D4A** | 63898 | protein_coding | **28.426** | 1.282 | 0.386 | **2.432** | 2.94E-04 | **8.80E-04** |
| ENSG00000273033.3 | **LINC02035** | 10012955( | lncRNA | **25.036** | 1.423 | 0.428 | **2.681** | 2.94E-04 | **8.80E-04** |
| ENSG00000236824.2 | **BCYRN1** | 618 | lncRNA | **32.488** | 1.028 | 0.310 | **2.040** | 2.94E-04 | **8.80E-04** |
| ENSG00000204802.4 | **FAM88C** | | lncRNA | **17.548** | -2.099 | -0.632 | **0.233** | 2.95E-04 | **8.81E-04** |
| ENSG00000248746.8 | **ACTN3** | 89 | protein_coding | **27.572** | 1.304 | 0.393 | **2.469** | 2.96E-04 | **8.84E-04** |
| ENSG00000137500.12 | **CCDC90B** | 60492 | protein_coding | **416.755** | -0.367 | -0.110 | **0.775** | 2.96E-04 | **8.84E-04** |
| ENSG00000166819.14 | **PLIN1** | 5346 | protein_coding | **16.093** | -1.868 | -0.562 | **0.274** | 2.98E-04 | **8.90E-04** |
| ENSG00000074755.17 | **ZZEF1** | 23140 | protein_coding | **1142.636** | 0.449 | 0.135 | **1.365** | 2.98E-04 | **8.91E-04** |
| ENSG00000153037.17 | **SRP19** | 6728 | protein_coding | **573.317** | -0.360 | -0.108 | **0.779** | 2.98E-04 | **8.91E-04** |
| ENSG00000114779.22 | **ABHD14B** | 84836 | protein_coding | **1276.992** | -0.496 | -0.149 | **0.709** | 2.99E-04 | **8.93E-04** |
| ENSG00000168890.16 | **TMEM150/** | 129303 | protein_coding | **100.935** | 0.676 | 0.204 | **1.598** | 3.00E-04 | **8.96E-04** |
| ENSG00000172732.14 | **MUS81** | 80198 | protein_coding | **994.866** | -0.380 | -0.114 | **0.768** | 3.01E-04 | **8.97E-04** |
| ENSG00000131748.18 | **STARD3** | 10948 | protein_coding | **1085.229** | 0.340 | 0.102 | **1.266** | 3.01E-04 | **8.98E-04** |
| ENSG00000011007.15 | **ELOA** | 6924 | protein_coding | **943.997** | 0.430 | 0.129 | **1.347** | 3.01E-04 | **8.98E-04** |
| ENSG00000213920.11 | **MDP1** | 145553 | protein_coding | **160.602** | -0.514 | -0.155 | **0.700** | 3.01E-04 | **8.99E-04** |
| ENSG00000101751.13 | **POLI** | 11201 | protein_coding | **463.953** | 0.484 | 0.146 | **1.399** | 3.02E-04 | **9.01E-04** |
| ENSG00000213096.12 | **ZNF254** | 9534 | protein_coding | **107.893** | -0.820 | -0.247 | **0.567** | 3.03E-04 | **9.04E-04** |
| ENSG00000186792.19 | **HYAL3** | 8372 | protein_coding | **58.344** | 0.707 | 0.213 | **1.632** | 3.04E-04 | **9.07E-04** |
| ENSG00000106244.15 | **PDAP1** | 11333 | protein_coding | **2274.279** | -0.343 | -0.103 | **0.788** | 3.04E-04 | **9.07E-04** |
| ENSG00000287804.4 | **ENSG00000287804** | | lncRNA | **7.760** | 2.325 | 0.700 | **5.012** | 3.05E-04 | **9.08E-04** |

| | | | | | | | | | |
|---|---|---|---|---|---|---|---|---|---|
| ENSG00000188042.9 | **ARL4C** | 10123 | protein_coding | **106.664** | -0.747 | -0.225 | **0.596** | 3.07E-04 | **9.14E-04** |
| ENSG00000106483.14 | **SFRP4** | 6424 | protein_coding | **8.127** | 2.737 | 0.824 | **6.666** | 3.08E-04 | **9.19E-04** |
| ENSG00000105254.14 | **TBCB** | 1155 | protein_coding | **1329.849** | 0.469 | 0.141 | **1.384** | 3.08E-04 | **9.19E-04** |
| ENSG00000223865.14 | **HLA-DPB1** | 3115 | protein_coding | **9897.711** | 0.428 | 0.129 | **1.345** | 3.09E-04 | **9.22E-04** |
| ENSG00000172354.12 | **GNB2** | 2783 | protein_coding | **2609.709** | -0.362 | -0.109 | **0.778** | 3.09E-04 | **9.22E-04** |
| ENSG00000165732.16 | **DDX21** | 9188 | protein_coding | **932.383** | -0.685 | -0.206 | **0.622** | 3.11E-04 | **9.27E-04** |
| ENSG00000167962.15 | **ZNF598** | 90850 | protein_coding | **1103.617** | -0.402 | -0.121 | **0.757** | 3.12E-04 | **9.30E-04** |
| ENSG00000180340.7 | **FZD2** | 2535 | protein_coding | **53.740** | 1.165 | 0.351 | **2.242** | 3.12E-04 | **9.30E-04** |
| ENSG00000095370.22 | **SH2D3C** | 10044 | protein_coding | **949.216** | -0.512 | -0.154 | **0.701** | 3.13E-04 | **9.31E-04** |
| ENSG00000227500.13 | **SCAMP4** | 113178 | protein_coding | **888.896** | 0.427 | 0.129 | **1.345** | 3.13E-04 | **9.33E-04** |
| ENSG00000227256.2 | **MIS18A-AS** | 10087420 | lncRNA | **17.865** | 1.287 | 0.387 | **2.440** | 3.15E-04 | **9.37E-04** |
| ENSG00000273108.1 | **ENSG00000273108** | | lncRNA | **4.366** | 3.019 | 0.909 | **8.108** | 3.16E-04 | **9.40E-04** |
| ENSG00000100353.20 | **EIF3D** | 8664 | protein_coding | **4393.386** | -0.423 | -0.127 | **0.746** | 3.17E-04 | **9.44E-04** |
| ENSG00000137463.7 | **MGARP** | 84709 | protein_coding | **6.278** | 2.230 | 0.671 | **4.692** | 3.18E-04 | **9.46E-04** |
| ENSG00000185339.12 | **TCN2** | 6948 | protein_coding | **106.983** | 0.907 | 0.273 | **1.875** | 3.18E-04 | **9.46E-04** |
| ENSG00000167770.15 | **OTUB1** | 55611 | protein_coding | **2121.641** | -0.338 | -0.102 | **0.791** | 3.18E-04 | **9.47E-04** |
| ENSG00000145247.14 | **OCIAD2** | 132299 | protein_coding | **641.301** | -0.374 | -0.113 | **0.771** | 3.19E-04 | **9.47E-04** |
| ENSG00000297944.1 | **ENSG00000297944** | | lncRNA | **4.672** | 3.605 | 1.085 | **12.169** | 3.19E-04 | **9.50E-04** |
| ENSG00000183258.14 | **DDX41** | 51428 | protein_coding | **2093.491** | -0.587 | -0.177 | **0.666** | 3.19E-04 | **9.50E-04** |
| ENSG00000157613.13 | **CREB3L1** | 90993 | protein_coding | **174.289** | 0.691 | 0.208 | **1.615** | 3.20E-04 | **9.51E-04** |
| ENSG00000130590.15 | **SAMD10** | 140700 | protein_coding | **1534.049** | -0.561 | -0.169 | **0.678** | 3.20E-04 | **9.52E-04** |
| ENSG00000129038.18 | **LOXL1** | 4016 | protein_coding | **5.333** | 3.047 | 0.917 | **8.263** | 3.21E-04 | **9.53E-04** |
| ENSG00000238120.3 | **LINC01589** | 10050673 | lncRNA | **8.079** | 2.532 | 0.762 | **5.783** | 3.22E-04 | **9.55E-04** |
| ENSG00000301524.1 | **ENSG00000301524** | | lncRNA | **8.756** | 1.938 | 0.583 | **3.832** | 3.22E-04 | **9.56E-04** |
| ENSG00000169764.18 | **UGP2** | 7360 | protein_coding | **614.273** | -0.456 | -0.137 | **0.729** | 3.22E-04 | **9.56E-04** |
| ENSG00000165983.17 | **PTER** | 9317 | protein_coding | **182.833** | -0.683 | -0.206 | **0.623** | 3.22E-04 | **9.57E-04** |
| ENSG00000225077.4 | **ICMT-DT** | 148645 | lncRNA | **12.064** | 1.604 | 0.483 | **3.040** | 3.23E-04 | **9.58E-04** |
| ENSG00000213246.8 | **SUPT4H1** | 6827 | protein_coding | **948.717** | 0.552 | 0.166 | **1.466** | 3.23E-04 | **9.58E-04** |
| ENSG00000127920.7 | **GNG11** | 2791 | protein_coding | **52.780** | 0.827 | 0.249 | **1.774** | 3.23E-04 | **9.60E-04** |
| ENSG00000213533.15 | **STIMATE** | 375346 | protein_coding | **234.433** | -0.878 | -0.264 | **0.544** | 3.24E-04 | **9.63E-04** |
| ENSG00000173258.15 | **ZNF483** | 158399 | protein_coding | **9.853** | 2.083 | 0.627 | **4.238** | 3.25E-04 | **9.64E-04** |

| | | | | | | | | | |
|---|---|---|---|---|---|---|---|---|---|
| ENSG00000271795.1 | **ENSG00000271795** | | lncRNA | **17.713** | 1.322 | 0.398 | **2.500** | 3.26E-04 | **9.67E-04** |
| ENSG00000115350.14 | **POLE4** | 56655 | protein_coding | **332.590** | -0.598 | -0.180 | **0.661** | 3.27E-04 | **9.71E-04** |
| ENSG00000131148.11 | **EMC8** | 10328 | protein_coding | **470.392** | -0.368 | -0.111 | **0.775** | 3.28E-04 | **9.72E-04** |
| ENSG00000170209.6 | **ANKK1** | 255239 | protein_coding | **14.250** | 1.635 | 0.492 | **3.106** | 3.28E-04 | **9.72E-04** |
| ENSG00000124226.13 | **RNF114** | 55905 | protein_coding | **1596.638** | 0.312 | 0.094 | **1.241** | 3.29E-04 | **9.74E-04** |
| ENSG00000140548.12 | **ZNF710** | 374655 | protein_coding | **481.315** | -0.619 | -0.186 | **0.651** | 3.31E-04 | **9.81E-04** |
| ENSG00000243364.9 | **EFNA4** | 1945 | protein_coding | **90.372** | 0.661 | 0.199 | **1.581** | 3.31E-04 | **9.82E-04** |
| ENSG00000167723.16 | **TRPV3** | 162514 | protein_coding | **140.837** | -0.754 | -0.227 | **0.593** | 3.33E-04 | **9.86E-04** |
| ENSG00000166171.15 | **DPCD** | 25911 | protein_coding | **219.187** | -0.485 | -0.146 | **0.715** | 3.33E-04 | **9.86E-04** |
| ENSG00000269473.1 | **ZNF132-DT** | | lncRNA | **12.170** | 1.829 | 0.551 | **3.552** | 3.34E-04 | **9.90E-04** |
| ENSG00000110047.19 | **EHD1** | 10938 | protein_coding | **4003.670** | 0.451 | 0.136 | **1.367** | 3.35E-04 | **9.92E-04** |
| ENSG00000187942.12 | **LDLRAD2** | 401944 | protein_coding | **17.052** | 1.375 | 0.414 | **2.594** | 3.35E-04 | **9.93E-04** |
| ENSG00000226824.9 | **ENSG00000226824** | | lncRNA | **44.528** | -1.021 | -0.307 | **0.493** | 3.36E-04 | **9.94E-04** |
| ENSG00000109063.16 | **MYH3** | 4621 | protein_coding | **177.144** | 0.870 | 0.262 | **1.827** | 3.36E-04 | **9.95E-04** |
| ENSG00000134987.15 | **WDR36** | 134430 | protein_coding | **518.031** | -0.485 | -0.146 | **0.715** | 3.37E-04 | **9.96E-04** |
| ENSG00000117143.16 | **UAP1** | 6675 | protein_coding | **260.085** | -0.517 | -0.156 | **0.699** | 3.37E-04 | **9.98E-04** |
| ENSG00000104879.7 | **CKM** | 1158 | protein_coding | **7.327** | 2.411 | 0.726 | **5.320** | 3.38E-04 | **9.99E-04** |
| ENSG00000264745.2 | **TTC39C-AS1** | | lncRNA | **28.274** | 1.223 | 0.368 | **2.335** | 3.38E-04 | **1.00E-03** |
| ENSG00000259826.2 | **CDK13-DT** | | lncRNA | **28.204** | 1.220 | 0.367 | **2.330** | 3.39E-04 | **1.00E-03** |
| ENSG00000169255.17 | **B3GALNT1** | 8706 | protein_coding | **39.192** | -1.250 | -0.376 | **0.421** | 3.39E-04 | **1.00E-03** |
| ENSG00000227308.3 | **LINC02832** | | lncRNA | **7.260** | 2.273 | 0.684 | **4.833** | 3.39E-04 | **1.00E-03** |
| ENSG00000236901.6 | **MIR600HG** | 81571 | lncRNA | **620.709** | 0.374 | 0.112 | **1.296** | 3.41E-04 | **1.01E-03** |
| ENSG00000272195.2 | **EFCAB2-AS1** | | lncRNA | **18.575** | 1.200 | 0.361 | **2.298** | 3.41E-04 | **1.01E-03** |
| ENSG00000309154.1 | **ENSG00000309154** | | lncRNA | **16.016** | 2.773 | 0.835 | **6.834** | 3.41E-04 | **1.01E-03** |
| ENSG00000211669.3 | **IGLV3-10** | | IG_V_gene | **59.185** | -1.621 | -0.488 | **0.325** | 3.42E-04 | **1.01E-03** |
| ENSG00000162817.8 | **C1orf115** | 79762 | protein_coding | **149.906** | 0.874 | 0.263 | **1.833** | 3.42E-04 | **1.01E-03** |
| ENSG00000230989.10 | **HSBP1** | 3281 | protein_coding | **641.855** | -0.454 | -0.137 | **0.730** | 3.42E-04 | **1.01E-03** |
| ENSG00000273619.3 | **RPS21-DT** | 12866722[illegible] | lncRNA | **50.125** | 0.835 | 0.251 | **1.784** | 3.43E-04 | **1.01E-03** |
| ENSG00000125166.15 | **GOT2** | 2806 | protein_coding | **1921.306** | -0.340 | -0.102 | **0.790** | 3.43E-04 | **1.01E-03** |
| ENSG00000301866.3 | **ENSG00000301866** | | lncRNA | **6.870** | 2.400 | 0.723 | **5.279** | 3.45E-04 | **1.02E-03** |
| ENSG00000104221.15 | **BRF2** | 55290 | protein_coding | **149.161** | -0.488 | -0.147 | **0.713** | 3.45E-04 | **1.02E-03** |

| | | | | | | | | | |
|---|---|---|---|---|---|---|---|---|---|
| ENSG00000084090.16 | **STARD7** | 56910 | protein_coding | **1948.680** | 0.458 | 0.138 | **1.373** | 3.46E-04 | **1.02E-03** |
| ENSG00000197580.15 | **BCO2** | 83875 | protein_coding | **60.501** | 0.933 | 0.281 | **1.909** | 3.46E-04 | **1.02E-03** |
| ENSG00000298831.1 | **ENSG00000298831** | | lncRNA | **9.260** | -1.646 | -0.496 | **0.319** | 3.47E-04 | **1.02E-03** |
| ENSG00000162599.20 | **NFIA** | 4774 | protein_coding | **21.516** | 1.780 | 0.536 | **3.433** | 3.47E-04 | **1.02E-03** |
| ENSG00000145331.16 | **TRMT10A** | 93587 | protein_coding | **140.274** | 1.020 | 0.307 | **2.028** | 3.47E-04 | **1.03E-03** |
| ENSG00000141977.9 | **CIB3** | 117286 | protein_coding | **4.664** | 3.361 | 1.012 | **10.278** | 3.48E-04 | **1.03E-03** |
| ENSG00000253200.1 | **ENSG00000253200** | | lncRNA | **20.571** | 1.355 | 0.408 | **2.559** | 3.48E-04 | **1.03E-03** |
| ENSG00000087495.19 | **PHACTR3** | 116154 | protein_coding | **10.904** | 1.842 | 0.555 | **3.586** | 3.49E-04 | **1.03E-03** |
| ENSG00000239713.11 | **APOBEC3G** | 60489 | protein_coding | **2360.495** | -0.347 | -0.104 | **0.786** | 3.49E-04 | **1.03E-03** |
| ENSG00000303936.1 | **ENSG00000303936** | | lncRNA | **28.296** | 1.049 | 0.316 | **2.070** | 3.50E-04 | **1.03E-03** |
| ENSG00000118960.15 | **HS1BP3** | 64342 | protein_coding | **321.427** | 0.408 | 0.123 | **1.326** | 3.50E-04 | **1.03E-03** |
| ENSG00000260719.2 | **ENSG00000260719** | | lncRNA | **79.397** | -0.875 | -0.263 | **0.545** | 3.50E-04 | **1.03E-03** |
| ENSG00000197062.14 | **ZSCAN26** | 7741 | protein_coding | **301.394** | 0.392 | 0.118 | **1.312** | 3.52E-04 | **1.04E-03** |
| ENSG00000173253.18 | **DMRT2** | 10655 | protein_coding | **4.578** | 3.593 | 1.081 | **12.064** | 3.53E-04 | **1.04E-03** |
| ENSG00000108852.18 | **MPP2** | 4355 | protein_coding | **29.786** | 1.455 | 0.438 | **2.741** | 3.56E-04 | **1.05E-03** |
| ENSG00000116171.21 | **SCP2** | 6342 | protein_coding | **687.143** | -0.511 | -0.154 | **0.702** | 3.56E-04 | **1.05E-03** |
| ENSG00000245680.13 | **ZNF585B** | 92285 | protein_coding | **287.456** | 0.493 | 0.148 | **1.407** | 3.57E-04 | **1.05E-03** |
| ENSG00000272142.5 | **LYRM4-AS:** | 100129461 | lncRNA | **27.504** | 0.983 | 0.296 | **1.976** | 3.57E-04 | **1.05E-03** |
| ENSG00000175701.12 | **MTLN** | 205251 | protein_coding | **288.998** | -0.501 | -0.151 | **0.707** | 3.60E-04 | **1.06E-03** |
| ENSG00000114405.13 | **CEP15** | 57415 | protein_coding | **33.804** | -1.038 | -0.312 | **0.487** | 3.61E-04 | **1.06E-03** |
| ENSG00000074800.18 | **ENO1** | 2023 | protein_coding | **34055.377** | -0.525 | -0.158 | **0.695** | 3.62E-04 | **1.07E-03** |
| ENSG00000083544.18 | **TDRD3** | 81550 | protein_coding | **412.906** | -0.459 | -0.138 | **0.728** | 3.64E-04 | **1.07E-03** |
| ENSG00000284969.1 | **ENSG00000284969** | | protein_coding | **224.705** | -0.735 | -0.221 | **0.601** | 3.64E-04 | **1.07E-03** |
| ENSG00000272899.5 | **SPMIP1** | 10013070! | protein_coding | **14.617** | 1.373 | 0.413 | **2.590** | 3.64E-04 | **1.07E-03** |
| ENSG00000197056.14 | **ZMYM1** | 79830 | protein_coding | **62.875** | 0.924 | 0.278 | **1.898** | 3.65E-04 | **1.07E-03** |
| ENSG00000165113.15 | **GKAP1** | 80318 | protein_coding | **94.851** | 0.842 | 0.254 | **1.793** | 3.65E-04 | **1.07E-03** |
| ENSG00000187391.24 | **MAGI2** | 9863 | protein_coding | **13.988** | 1.559 | 0.469 | **2.947** | 3.66E-04 | **1.07E-03** |
| ENSG00000104067.19 | **TJP1** | 7082 | protein_coding | **26.823** | 1.811 | 0.545 | **3.509** | 3.66E-04 | **1.07E-03** |
| ENSG00000107815.11 | **TWNK** | 56652 | protein_coding | **347.697** | -0.452 | -0.136 | **0.731** | 3.66E-04 | **1.08E-03** |
| ENSG00000230124.10 | **ACBD6** | 84320 | protein_coding | **1016.098** | -0.366 | -0.110 | **0.776** | 3.68E-04 | **1.08E-03** |
| ENSG00000148308.20 | **GTF3C5** | 9328 | protein_coding | **1031.462** | -0.299 | -0.090 | **0.813** | 3.68E-04 | **1.08E-03** |

| | | | | | | | | | |
|---|---|---|---|---|---|---|---|---|---|
| ENSG00000117215.16 | **PLA2G2D** | 26279 | protein_coding | **7.446** | 2.411 | 0.726 | **5.317** | 3.68E-04 | **1.08E-03** |
| ENSG00000105982.19 | **RNF32** | 140545 | protein_coding | **60.649** | 0.758 | 0.228 | **1.692** | 3.68E-04 | **1.08E-03** |
| ENSG00000272666.2 | **KLHDC7B-I** | 10537309 | lncRNA | **29.161** | 1.066 | 0.321 | **2.093** | 3.68E-04 | **1.08E-03** |
| ENSG00000154813.12 | **DPH3** | 285381 | protein_coding | **282.352** | -0.534 | -0.161 | **0.690** | 3.68E-04 | **1.08E-03** |
| ENSG00000132518.7 | **GUCY2D** | 3000 | protein_coding | **3.564** | 3.566 | 1.073 | **11.840** | 3.69E-04 | **1.08E-03** |
| ENSG00000230513.5 | **THAP7-AS1** | | lncRNA | **93.693** | 0.820 | 0.247 | **1.765** | 3.69E-04 | **1.08E-03** |
| ENSG00000036530.11 | **CYP46A1** | 10858 | protein_coding | **5.228** | 4.166 | 1.254 | **17.953** | 3.69E-04 | **1.08E-03** |
| ENSG00000205309.16 | **NT5M** | 56953 | protein_coding | **201.254** | 0.672 | 0.202 | **1.594** | 3.71E-04 | **1.09E-03** |
| ENSG00000096717.14 | **SIRT1** | 23411 | protein_coding | **154.592** | -0.677 | -0.204 | **0.625** | 3.71E-04 | **1.09E-03** |
| ENSG00000122026.14 | **RPL21** | 6144 | protein_coding | **16724.797** | -0.343 | -0.103 | **0.788** | 3.71E-04 | **1.09E-03** |
| ENSG00000160801.16 | **PTH1R** | 5745 | protein_coding | **10.358** | 2.465 | 0.742 | **5.522** | 3.71E-04 | **1.09E-03** |
| ENSG00000111300.12 | **NAA25** | 80018 | protein_coding | **436.878** | -0.471 | -0.142 | **0.721** | 3.72E-04 | **1.09E-03** |
| ENSG00000230490.5 | **ENSG00000230490** | | lncRNA | **19.186** | 1.188 | 0.358 | **2.279** | 3.72E-04 | **1.09E-03** |
| ENSG00000064199.9 | **SPA17** | 53340 | protein_coding | **93.557** | 0.621 | 0.187 | **1.538** | 3.73E-04 | **1.09E-03** |
| ENSG00000251201.8 | **TMED7-TIC** | 10030273 | protein_coding | **42.747** | -5.731 | -1.725 | **0.019** | 3.75E-04 | **1.10E-03** |
| ENSG00000277775.2 | **H3C7** | 8968 | protein_coding | **6.575** | -2.553 | -0.769 | **0.170** | 3.75E-04 | **1.10E-03** |
| ENSG00000288741.2 | **ENSG00000288741** | | lncRNA | **36.042** | 0.967 | 0.291 | **1.954** | 3.76E-04 | **1.10E-03** |
| ENSG00000151067.24 | **CACNA1C** | 775 | protein_coding | **9.019** | 2.845 | 0.856 | **7.184** | 3.76E-04 | **1.10E-03** |
| ENSG00000115216.16 | **NRBP1** | 29959 | protein_coding | **1528.765** | -0.386 | -0.116 | **0.765** | 3.76E-04 | **1.10E-03** |
| ENSG00000163558.15 | **PRKCI** | 5584 | protein_coding | **189.797** | 0.673 | 0.203 | **1.594** | 3.77E-04 | **1.10E-03** |
| ENSG00000137502.12 | **RAB30** | 27314 | protein_coding | **655.195** | -0.655 | -0.197 | **0.635** | 3.77E-04 | **1.10E-03** |
| ENSG00000167554.17 | **ZNF610** | 162963 | protein_coding | **57.429** | 0.706 | 0.213 | **1.632** | 3.78E-04 | **1.11E-03** |
| ENSG00000116906.15 | **GNPAT** | 8443 | protein_coding | **1045.071** | -0.376 | -0.113 | **0.771** | 3.78E-04 | **1.11E-03** |
| ENSG00000168092.16 | **PAFAH1B2** | 5049 | protein_coding | **669.500** | 0.351 | 0.106 | **1.275** | 3.78E-04 | **1.11E-03** |
| ENSG00000226465.6 | **ENSG00000226465** | | lncRNA | **24.111** | 1.024 | 0.308 | **2.034** | 3.80E-04 | **1.11E-03** |
| ENSG00000011478.14 | **QPCTL** | 54814 | protein_coding | **412.937** | -0.435 | -0.131 | **0.739** | 3.80E-04 | **1.11E-03** |
| ENSG00000155158.22 | **TTC39B** | 158219 | protein_coding | **44.235** | 0.855 | 0.257 | **1.809** | 3.80E-04 | **1.11E-03** |
| ENSG00000175756.16 | **AURKAIP1** | 54998 | protein_coding | **1528.164** | -0.423 | -0.127 | **0.746** | 3.80E-04 | **1.11E-03** |
| ENSG00000286190.4 | **ENSG0000** | 728392 | protein_coding | **184.848** | 1.018 | 0.306 | **2.025** | 3.80E-04 | **1.11E-03** |
| ENSG00000289594.2 | **ENSG00000289594** | | lncRNA | **7.603** | 1.931 | 0.581 | **3.812** | 3.82E-04 | **1.12E-03** |
| ENSG00000197046.13 | **SIGLEC15** | 284266 | protein_coding | **9.411** | -1.915 | -0.576 | **0.265** | 3.82E-04 | **1.12E-03** |

| | | | | | | | | | |
|---|---|---|---|---|---|---|---|---|---|
| ENSG00000005486.19 | **RHBDD2** | 57414 | protein_coding | **438.885** | 0.390 | 0.117 | **1.311** | 3.82E-04 | **1.12E-03** |
| ENSG00000133454.17 | **MYO18B** | 84700 | protein_coding | **48.295** | 1.574 | 0.474 | **2.978** | 3.83E-04 | **1.12E-03** |
| ENSG00000109118.16 | **PHF12** | 57649 | protein_coding | **1434.652** | 0.372 | 0.112 | **1.294** | 3.85E-04 | **1.12E-03** |
| ENSG00000232618.2 | **HIVEP2-DT** | | lncRNA | **17.722** | 1.296 | 0.390 | **2.456** | 3.85E-04 | **1.12E-03** |
| ENSG00000165832.8 | **TRUB1** | 142940 | protein_coding | **108.612** | -0.746 | -0.224 | **0.596** | 3.85E-04 | **1.13E-03** |
| ENSG00000167775.13 | **CD320** | 51293 | protein_coding | **510.540** | -0.461 | -0.139 | **0.726** | 3.85E-04 | **1.13E-03** |
| ENSG00000141741.14 | **MIEN1** | 84299 | protein_coding | **962.983** | 0.471 | 0.142 | **1.386** | 3.85E-04 | **1.13E-03** |
| ENSG00000214212.11 | **C19orf38** | 255809 | protein_coding | **297.559** | 0.703 | 0.212 | **1.628** | 3.86E-04 | **1.13E-03** |
| ENSG00000126903.18 | **SLC10A3** | 8273 | protein_coding | **252.559** | -0.496 | -0.149 | **0.709** | 3.86E-04 | **1.13E-03** |
| ENSG00000170296.12 | **GABARAP** | 11337 | protein_coding | **4009.048** | 0.369 | 0.111 | **1.292** | 3.87E-04 | **1.13E-03** |
| ENSG00000163823.6 | **CCR1** | 1230 | protein_coding | **70.337** | -1.107 | -0.333 | **0.464** | 3.87E-04 | **1.13E-03** |
| ENSG00000106246.20 | **PTCD1** | 26024 | protein_coding | **357.958** | -0.370 | -0.111 | **0.774** | 3.88E-04 | **1.13E-03** |
| ENSG00000160785.16 | **SLC25A44** | 9673 | protein_coding | **689.279** | 0.511 | 0.154 | **1.425** | 3.89E-04 | **1.13E-03** |
| ENSG00000134569.12 | **LRP4** | 4038 | protein_coding | **21.131** | 1.345 | 0.405 | **2.540** | 3.90E-04 | **1.14E-03** |
| ENSG00000181350.14 | **LRRC75A** | 388341 | protein_coding | **8.324** | 1.965 | 0.591 | **3.903** | 3.90E-04 | **1.14E-03** |
| ENSG00000177455.15 | **CD19** | 930 | protein_coding | **3658.430** | -0.397 | -0.120 | **0.759** | 3.90E-04 | **1.14E-03** |
| ENSG00000163362.13 | **INAVA** | 55765 | protein_coding | **338.010** | 0.730 | 0.220 | **1.659** | 3.91E-04 | **1.14E-03** |
| ENSG00000181061.16 | **HIGD1A** | 25994 | protein_coding | **456.933** | -0.427 | -0.129 | **0.744** | 3.91E-04 | **1.14E-03** |
| ENSG00000100321.17 | **SYNGR1** | 9145 | protein_coding | **207.936** | 0.950 | 0.286 | **1.932** | 3.92E-04 | **1.14E-03** |
| ENSG00000011454.20 | **RABGAP1** | 23637 | protein_coding | **831.229** | 0.449 | 0.135 | **1.365** | 3.93E-04 | **1.15E-03** |
| ENSG00000168291.15 | **PDHB** | 5162 | protein_coding | **822.846** | -0.411 | -0.124 | **0.752** | 3.94E-04 | **1.15E-03** |
| ENSG00000141378.17 | **PTRH2** | 51651 | protein_coding | **558.100** | -0.483 | -0.145 | **0.715** | 3.94E-04 | **1.15E-03** |
| ENSG00000079308.22 | **TNS1** | 7145 | protein_coding | **26.656** | 1.722 | 0.518 | **3.298** | 3.95E-04 | **1.15E-03** |
| ENSG00000107485.20 | **GATA3** | 2625 | protein_coding | **176.055** | 0.731 | 0.220 | **1.660** | 3.95E-04 | **1.15E-03** |
| ENSG00000079805.21 | **DNM2** | 1785 | protein_coding | **3915.335** | 0.316 | 0.095 | **1.245** | 3.95E-04 | **1.15E-03** |
| ENSG00000125734.18 | **GPR108** | 56927 | protein_coding | **1771.129** | 0.450 | 0.135 | **1.366** | 3.97E-04 | **1.16E-03** |
| ENSG00000226944.2 | **RNF207-AS1** | | lncRNA | **7.227** | 2.185 | 0.658 | **4.546** | 3.97E-04 | **1.16E-03** |
| ENSG00000287684.2 | **ENSG00000287684** | | lncRNA | **9.855** | 3.941 | 1.186 | **15.362** | 3.98E-04 | **1.16E-03** |
| ENSG00000256771.7 | **ZNF253** | 56242 | protein_coding | **228.188** | -0.691 | -0.208 | **0.619** | 3.98E-04 | **1.16E-03** |
| ENSG00000105220.18 | **GPI** | 2821 | protein_coding | **10132.308** | -0.536 | -0.161 | **0.690** | 3.99E-04 | **1.16E-03** |
| ENSG00000263011.4 | **ENSG0000(** | 12490362( | lncRNA | **6.530** | 2.834 | 0.853 | **7.130** | 3.99E-04 | **1.16E-03** |

| | | | | | | | | | |
|---|---|---|---|---|---|---|---|---|---|
| ENSG00000140795.15 | **MYLK3** | 91807 | protein_coding | **16.910** | 1.325 | 0.399 | **2.504** | 4.00E-04 | **1.16E-03** |
| ENSG00000187624.10 | **LIAT1** | 400566 | protein_coding | **46.211** | 1.040 | 0.313 | **2.056** | 4.00E-04 | **1.16E-03** |
| ENSG00000151490.17 | **PTPRO** | 5800 | protein_coding | **38.519** | -0.942 | -0.283 | **0.521** | 4.00E-04 | **1.16E-03** |
| ENSG00000135052.18 | **GOLM1** | 51280 | protein_coding | **186.628** | 0.498 | 0.150 | **1.412** | 4.00E-04 | **1.16E-03** |
| ENSG00000306852.1 | **ENSG00000306852** | | lncRNA | **5.175** | 2.985 | 0.898 | **7.915** | 4.02E-04 | **1.17E-03** |
| ENSG00000104131.15 | **EIF3J** | 8669 | protein_coding | **791.000** | -0.520 | -0.157 | **0.697** | 4.03E-04 | **1.17E-03** |
| ENSG00000162385.13 | **MAGOH** | 4116 | protein_coding | **715.065** | -0.349 | -0.105 | **0.785** | 4.04E-04 | **1.17E-03** |
| ENSG00000116157.7 | **GPX7** | 2882 | protein_coding | **153.188** | -0.704 | -0.212 | **0.614** | 4.05E-04 | **1.18E-03** |
| ENSG00000302691.1 | **ENSG00000302691** | | lncRNA | **6.314** | 2.324 | 0.700 | **5.006** | 4.05E-04 | **1.18E-03** |
| ENSG00000213281.7 | **NRAS** | 4893 | protein_coding | **650.036** | -0.415 | -0.125 | **0.750** | 4.08E-04 | **1.18E-03** |
| ENSG00000310402.1 | **ENSG00000310402** | | lncRNA | **8.756** | 1.926 | 0.580 | **3.800** | 4.09E-04 | **1.19E-03** |
| ENSG00000158863.24 | **FHIP2B** | 64760 | protein_coding | **1013.076** | 0.324 | 0.097 | **1.252** | 4.11E-04 | **1.19E-03** |
| ENSG00000303013.1 | **ENSG0000(** | 124903731 | lncRNA | **18.785** | 1.459 | 0.439 | **2.748** | 4.11E-04 | **1.19E-03** |
| ENSG00000109814.14 | **UGDH** | 7358 | protein_coding | **232.297** | -0.570 | -0.172 | **0.674** | 4.11E-04 | **1.19E-03** |
| ENSG00000133619.20 | **KRABD3** | 84626 | protein_coding | **364.353** | 0.398 | 0.120 | **1.318** | 4.12E-04 | **1.20E-03** |
| ENSG00000225945.2 | **ZFAND3-DT** | 10798659: | lncRNA | **14.562** | 1.360 | 0.409 | **2.567** | 4.12E-04 | **1.20E-03** |
| ENSG00000148965.12 | **SAA4** | 6291 | protein_coding | **9.726** | 2.571 | 0.774 | **5.942** | 4.13E-04 | **1.20E-03** |
| ENSG00000182183.16 | **SHISAL2A** | 348378 | protein_coding | **16.893** | -1.634 | -0.492 | **0.322** | 4.15E-04 | **1.20E-03** |
| ENSG00000228252.10 | **COL6A4P2** | | transcribed_unita | **9.133** | 2.456 | 0.739 | **5.487** | 4.15E-04 | **1.20E-03** |
| ENSG00000247240.11 | **UBL7-DT** | 440288 | lncRNA | **216.372** | -0.576 | -0.173 | **0.671** | 4.18E-04 | **1.21E-03** |
| ENSG00000166246.16 | **DNAAF8** | 146562 | protein_coding | **9.871** | 1.815 | 0.546 | **3.519** | 4.18E-04 | **1.21E-03** |
| ENSG00000163214.23 | **DHX57** | 90957 | protein_coding | **377.511** | 0.402 | 0.121 | **1.321** | 4.18E-04 | **1.21E-03** |
| ENSG00000175221.18 | **MED16** | 10025 | protein_coding | **669.960** | -0.585 | -0.176 | **0.667** | 4.19E-04 | **1.21E-03** |
| ENSG00000166664.17 | **CHRFAM7A** | 89832 | protein_coding | **6.562** | 3.230 | 0.972 | **9.380** | 4.19E-04 | **1.21E-03** |
| ENSG00000196972.10 | **SMIM10L2** | 644596 | protein_coding | **3.665** | 3.672 | 1.106 | **12.750** | 4.19E-04 | **1.21E-03** |
| ENSG00000015532.12 | **XYLT2** | 64132 | protein_coding | **436.543** | -0.343 | -0.103 | **0.789** | 4.19E-04 | **1.22E-03** |
| ENSG00000004660.17 | **CAMKK1** | 84254 | protein_coding | **131.197** | 0.806 | 0.243 | **1.749** | 4.20E-04 | **1.22E-03** |
| ENSG00000151806.16 | **GUF1** | 60558 | protein_coding | **447.923** | -0.341 | -0.103 | **0.790** | 4.20E-04 | **1.22E-03** |
| ENSG00000102900.15 | **NUP93** | 9688 | protein_coding | **896.744** | -0.428 | -0.129 | **0.743** | 4.20E-04 | **1.22E-03** |
| ENSG00000224184.8 | **MIR3681H** | 100506457 | lncRNA | **116.220** | -0.609 | -0.183 | **0.655** | 4.21E-04 | **1.22E-03** |
| ENSG00000187997.13 | **C17orf99** | 100141515 | protein_coding | **163.080** | 0.809 | 0.243 | **1.752** | 4.21E-04 | **1.22E-03** |

| | | | | | | | | | |
|---|---|---|---|---|---|---|---|---|---|
| ENSG00000259088.2 | **ENSG00000259088** | | lncRNA | **32.740** | 0.910 | 0.274 | **1.879** | 4.24E-04 | **1.23E-03** |
| ENSG00000187688.16 | **TRPV2** | 51393 | protein_coding | **1277.055** | -0.607 | -0.183 | **0.656** | 4.24E-04 | **1.23E-03** |
| ENSG00000239779.9 | **WBP1** | 23559 | protein_coding | **1414.834** | 0.416 | 0.125 | **1.334** | 4.25E-04 | **1.23E-03** |
| ENSG00000146701.14 | **MDH2** | 4191 | protein_coding | **2889.747** | -0.496 | -0.149 | **0.709** | 4.25E-04 | **1.23E-03** |
| ENSG00000189229.13 | **ENSG0000(** | 10537694( | lncRNA | **20.509** | 1.798 | 0.541 | **3.478** | 4.26E-04 | **1.23E-03** |
| ENSG00000188001.12 | **TPRG1** | 285386 | protein_coding | **59.811** | -1.315 | -0.396 | **0.402** | 4.27E-04 | **1.24E-03** |
| ENSG00000128191.19 | **DGCR8** | 54487 | protein_coding | **655.448** | -0.409 | -0.123 | **0.753** | 4.27E-04 | **1.24E-03** |
| ENSG00000094841.16 | **UPRT** | 139596 | protein_coding | **135.394** | -0.658 | -0.198 | **0.634** | 4.29E-04 | **1.24E-03** |
| ENSG00000185803.14 | **SLC52A2** | 79581 | protein_coding | **523.929** | -0.505 | -0.152 | **0.705** | 4.29E-04 | **1.24E-03** |
| ENSG00000169330.11 | **MINAR1** | 23251 | protein_coding | **11.170** | -1.593 | -0.480 | **0.331** | 4.29E-04 | **1.24E-03** |
| ENSG00000066697.17 | **MSANTD3** | 91283 | protein_coding | **120.836** | -0.631 | -0.190 | **0.646** | 4.31E-04 | **1.25E-03** |
| ENSG00000137843.14 | **PAK6** | 56924 | protein_coding | **31.991** | 1.443 | 0.434 | **2.719** | 4.32E-04 | **1.25E-03** |
| ENSG00000085719.15 | **CPNE3** | 8895 | protein_coding | **843.153** | 0.532 | 0.160 | **1.446** | 4.32E-04 | **1.25E-03** |
| ENSG00000267296.3 | **CEBPA-DT** | 80054 | lncRNA | **5.638** | 2.460 | 0.740 | **5.501** | 4.33E-04 | **1.25E-03** |
| ENSG00000157110.18 | **RBPMS** | 11030 | protein_coding | **97.399** | -0.972 | -0.293 | **0.510** | 4.34E-04 | **1.25E-03** |
| ENSG00000183011.16 | **NAA38** | 84316 | protein_coding | **711.345** | -0.451 | -0.136 | **0.732** | 4.34E-04 | **1.25E-03** |
| ENSG00000066557.8 | **LRRC40** | 55631 | protein_coding | **218.112** | -0.560 | -0.169 | **0.678** | 4.35E-04 | **1.26E-03** |
| ENSG00000178821.15 | **TMEM52** | 339456 | protein_coding | **14.330** | 1.694 | 0.510 | **3.235** | 4.35E-04 | **1.26E-03** |
| ENSG00000178665.17 | **ZNF713** | 349075 | protein_coding | **14.614** | -1.611 | -0.485 | **0.327** | 4.35E-04 | **1.26E-03** |
| ENSG00000081177.21 | **EXD2** | 55218 | protein_coding | **238.997** | 0.493 | 0.149 | **1.408** | 4.36E-04 | **1.26E-03** |
| ENSG00000302999.1 | **ENSG00000302999** | | lncRNA | **15.613** | 1.579 | 0.475 | **2.988** | 4.36E-04 | **1.26E-03** |
| ENSG00000007372.26 | **PAX6** | 5080 | protein_coding | **11.827** | 2.079 | 0.626 | **4.224** | 4.36E-04 | **1.26E-03** |
| ENSG00000251493.5 | **FOXD1** | 2297 | protein_coding | **9.341** | 1.994 | 0.600 | **3.984** | 4.37E-04 | **1.26E-03** |
| ENSG00000290545.2 | **INTS4P1** | 285905 | lncRNA | **19.068** | 1.618 | 0.487 | **3.070** | 4.39E-04 | **1.27E-03** |
| ENSG00000185513.19 | **L3MBTL1** | 26013 | protein_coding | **305.844** | 0.593 | 0.179 | **1.508** | 4.40E-04 | **1.27E-03** |
| ENSG00000259426.7 | **KIF23-AS1** | 145694 | lncRNA | **8.506** | 2.188 | 0.659 | **4.558** | 4.42E-04 | **1.28E-03** |
| ENSG00000062194.18 | **GPBP1** | 65056 | protein_coding | **1473.537** | 0.393 | 0.118 | **1.313** | 4.43E-04 | **1.28E-03** |
| ENSG00000163840.11 | **DTX3L** | 151636 | protein_coding | **853.412** | -0.622 | -0.187 | **0.650** | 4.45E-04 | **1.28E-03** |
| ENSG00000125843.14 | **AP5S1** | 55317 | protein_coding | **97.465** | -0.725 | -0.218 | **0.605** | 4.45E-04 | **1.28E-03** |
| ENSG00000162076.15 | **FLYWCH2** | 114984 | protein_coding | **435.553** | 0.382 | 0.115 | **1.303** | 4.47E-04 | **1.29E-03** |
| ENSG00000240682.12 | **ISY1** | 57461 | protein_coding | **727.725** | -0.321 | -0.097 | **0.801** | 4.47E-04 | **1.29E-03** |

| | | | | | | | | | |
|---|---|---|---|---|---|---|---|---|---|
| ENSG00000105852.13 | **PON3** | 5446 | protein_coding | **4.774** | 3.191 | 0.961 | **9.133** | 4.47E-04 | **1.29E-03** |
| ENSG00000131174.9 | **COX7B** | 1349 | protein_coding | **2090.233** | -0.547 | -0.165 | **0.684** | 4.48E-04 | **1.29E-03** |
| ENSG00000294915.1 | **ENSG00000294915** | | lncRNA | **6.206** | 2.510 | 0.756 | **5.695** | 4.49E-04 | **1.29E-03** |
| ENSG00000141542.13 | **RAB40B** | 10966 | protein_coding | **152.672** | 0.710 | 0.214 | **1.636** | 4.51E-04 | **1.30E-03** |
| ENSG00000162545.8 | **CAMK2N1** | 55450 | protein_coding | **9.748** | 1.786 | 0.537 | **3.447** | 4.51E-04 | **1.30E-03** |
| ENSG00000111269.5 | **CREBL2** | 1389 | protein_coding | **377.215** | 0.512 | 0.154 | **1.426** | 4.52E-04 | **1.30E-03** |
| ENSG00000171843.18 | **MLLT3** | 4300 | protein_coding | **829.236** | -0.529 | -0.159 | **0.693** | 4.53E-04 | **1.30E-03** |
| ENSG00000109775.13 | **UFSP2** | 55325 | protein_coding | **363.829** | -0.458 | -0.138 | **0.728** | 4.53E-04 | **1.31E-03** |
| ENSG00000168778.15 | **TCTN2** | 79867 | protein_coding | **179.822** | 0.571 | 0.172 | **1.486** | 4.56E-04 | **1.31E-03** |
| ENSG00000170469.13 | **SPATA24** | 202051 | protein_coding | **129.147** | 0.713 | 0.215 | **1.640** | 4.59E-04 | **1.32E-03** |
| ENSG00000180549.10 | **FUT7** | 2529 | protein_coding | **282.153** | -1.050 | -0.316 | **0.483** | 4.60E-04 | **1.32E-03** |
| ENSG00000092098.20 | **RNF31** | 55072 | protein_coding | **1913.414** | 0.444 | 0.134 | **1.361** | 4.60E-04 | **1.32E-03** |
| ENSG00000280434.1 | **ENSG00000280434** | | lncRNA | **72.144** | -0.788 | -0.237 | **0.579** | 4.60E-04 | **1.33E-03** |
| ENSG00000142207.9 | **URB1** | 9875 | protein_coding | **366.589** | -0.434 | -0.131 | **0.740** | 4.61E-04 | **1.33E-03** |
| ENSG00000198513.17 | **ATL1** | 51062 | protein_coding | **13.550** | 1.714 | 0.516 | **3.281** | 4.62E-04 | **1.33E-03** |
| ENSG00000113732.11 | **ATP6V0E1** | 8992 | protein_coding | **2648.604** | 0.326 | 0.098 | **1.253** | 4.62E-04 | **1.33E-03** |
| ENSG00000235897.2 | **TM4SF19-A** | 10087421 | lncRNA | **10.511** | 1.694 | 0.510 | **3.235** | 4.62E-04 | **1.33E-03** |
| ENSG00000100284.24 | **TOM1** | 10043 | protein_coding | **436.854** | 0.465 | 0.140 | **1.381** | 4.62E-04 | **1.33E-03** |
| ENSG00000131711.16 | **MAP1B** | 4131 | protein_coding | **165.654** | -0.659 | -0.198 | **0.633** | 4.63E-04 | **1.33E-03** |
| ENSG00000158669.13 | **GPAT4** | 137964 | protein_coding | **1808.254** | 0.309 | 0.093 | **1.239** | 4.63E-04 | **1.33E-03** |
| ENSG00000196352.20 | **CD55** | 1604 | protein_coding | **481.436** | -0.512 | -0.154 | **0.701** | 4.64E-04 | **1.33E-03** |
| ENSG00000150636.19 | **CCDC102B** | 79839 | protein_coding | **51.247** | 0.824 | 0.248 | **1.770** | 4.65E-04 | **1.34E-03** |
| ENSG00000136153.23 | **LMO7** | 4008 | protein_coding | **523.708** | -0.545 | -0.164 | **0.685** | 4.65E-04 | **1.34E-03** |
| ENSG00000171914.19 | **TLN2** | 83660 | protein_coding | **59.327** | 1.290 | 0.388 | **2.445** | 4.66E-04 | **1.34E-03** |
| ENSG00000204611.9 | **ZNF616** | 90317 | protein_coding | **116.461** | -0.505 | -0.152 | **0.705** | 4.66E-04 | **1.34E-03** |
| ENSG00000306879.1 | **ENSG00000306879** | | lncRNA | **57.818** | 1.016 | 0.306 | **2.022** | 4.66E-04 | **1.34E-03** |
| ENSG00000138115.17 | **CYP2C8** | 1558 | protein_coding | **17.805** | 1.244 | 0.375 | **2.369** | 4.66E-04 | **1.34E-03** |
| ENSG00000290083.2 | **ENSG00000290083** | | lncRNA | **6.266** | 2.694 | 0.811 | **6.473** | 4.67E-04 | **1.34E-03** |
| ENSG00000135903.21 | **PAX3** | 5077 | protein_coding | **11.857** | 2.007 | 0.604 | **4.021** | 4.67E-04 | **1.34E-03** |
| ENSG00000273983.2 | **H3C8** | 8355 | protein_coding | **8.125** | -2.198 | -0.662 | **0.218** | 4.68E-04 | **1.34E-03** |
| ENSG00000184292.7 | **TACSTD2** | 4070 | protein_coding | **19.232** | 2.029 | 0.611 | **4.082** | 4.68E-04 | **1.34E-03** |

| | | | | | | | | | |
|---|---|---|---|---|---|---|---|---|---|
| ENSG00000136003.18 | **ISCU** | 23479 | protein_coding | **2984.280** | 0.347 | 0.104 | **1.272** | 4.68E-04 | **1.34E-03** |
| ENSG00000184005.13 | **ST6GALNA** | 256435 | protein_coding | **15.439** | 2.461 | 0.741 | **5.508** | 4.70E-04 | **1.35E-03** |
| ENSG00000175352.13 | **NRIP3** | 56675 | protein_coding | **3.727** | 3.470 | 1.045 | **11.083** | 4.70E-04 | **1.35E-03** |
| ENSG00000138085.20 | **ATRAID** | 51374 | protein_coding | **1365.241** | -0.433 | -0.130 | **0.741** | 4.71E-04 | **1.35E-03** |
| ENSG00000305847.1 | **ENSG00000305847** | | lncRNA | **19.430** | -1.235 | -0.372 | **0.425** | 4.71E-04 | **1.35E-03** |
| ENSG00000134317.19 | **GRHL1** | 29841 | protein_coding | **10.436** | 1.735 | 0.522 | **3.330** | 4.72E-04 | **1.35E-03** |
| ENSG00000242686.4 | **PDE6B-AS1** | | lncRNA | **10.026** | 1.868 | 0.562 | **3.651** | 4.72E-04 | **1.35E-03** |
| ENSG00000151131.13 | **NOPCHAP1** | 121053 | protein_coding | **608.291** | -0.383 | -0.115 | **0.767** | 4.72E-04 | **1.35E-03** |
| ENSG00000026950.20 | **BTN3A1** | 11119 | protein_coding | **1375.553** | 0.376 | 0.113 | **1.298** | 4.75E-04 | **1.36E-03** |
| ENSG00000137103.22 | **TMEM8B** | 51754 | protein_coding | **496.553** | 0.440 | 0.132 | **1.356** | 4.75E-04 | **1.36E-03** |
| ENSG00000110651.15 | **CD81** | 975 | protein_coding | **3082.430** | -0.361 | -0.109 | **0.778** | 4.76E-04 | **1.36E-03** |
| ENSG00000273958.1 | **ENSG00000273958** | | lncRNA | **5.792** | 2.915 | 0.878 | **7.543** | 4.76E-04 | **1.36E-03** |
| ENSG00000171680.25 | **PLEKHG5** | 57449 | protein_coding | **30.886** | 1.129 | 0.340 | **2.187** | 4.77E-04 | **1.37E-03** |
| ENSG00000157570.14 | **TSPAN18** | 90139 | protein_coding | **15.753** | 1.896 | 0.571 | **3.721** | 4.79E-04 | **1.37E-03** |
| ENSG00000224079.2 | **CHLSN-AS1** | | lncRNA | **12.083** | -2.179 | -0.656 | **0.221** | 4.80E-04 | **1.38E-03** |
| ENSG00000162946.25 | **DISC1** | 27185 | protein_coding | **25.015** | 1.789 | 0.539 | **3.456** | 4.80E-04 | **1.38E-03** |
| ENSG00000245848.3 | **CEBPA** | 1050 | protein_coding | **16.612** | 1.519 | 0.457 | **2.866** | 4.80E-04 | **1.38E-03** |
| ENSG00000167011.10 | **NAT16** | 375607 | protein_coding | **6.718** | 2.864 | 0.862 | **7.280** | 4.80E-04 | **1.38E-03** |
| ENSG00000257585.2 | **LINC00609** | 10110177 | lncRNA | **6.090** | 2.795 | 0.841 | **6.939** | 4.82E-04 | **1.38E-03** |
| ENSG00000100592.17 | **DAAM1** | 23002 | protein_coding | **440.581** | 0.624 | 0.188 | **1.541** | 4.82E-04 | **1.38E-03** |
| ENSG00000177301.18 | **KCNA2** | 3737 | protein_coding | **8.446** | -1.829 | -0.551 | **0.281** | 4.83E-04 | **1.38E-03** |
| ENSG00000101132.13 | **PFDN4** | 5203 | protein_coding | **271.021** | -0.586 | -0.176 | **0.666** | 4.83E-04 | **1.38E-03** |
| ENSG00000131844.19 | **MCCC2** | 64087 | protein_coding | **1756.034** | -0.335 | -0.101 | **0.793** | 4.84E-04 | **1.38E-03** |
| ENSG00000065613.17 | **SLK** | 9748 | protein_coding | **233.111** | -0.657 | -0.198 | **0.634** | 4.85E-04 | **1.39E-03** |
| ENSG00000116525.15 | **TRIM62** | 55223 | protein_coding | **111.236** | 0.512 | 0.154 | **1.426** | 4.90E-04 | **1.40E-03** |
| ENSG00000166800.12 | **LDHAL6A** | 160287 | protein_coding | **8.799** | 3.035 | 0.914 | **8.196** | 4.90E-04 | **1.40E-03** |
| ENSG00000173762.10 | **CD7** | 924 | protein_coding | **7.979** | 1.956 | 0.589 | **3.879** | 4.90E-04 | **1.40E-03** |
| ENSG00000005102.16 | **MEOX1** | 4222 | protein_coding | **26.907** | 1.334 | 0.402 | **2.521** | 4.90E-04 | **1.40E-03** |
| ENSG00000166478.12 | **ZNF143** | 7702 | protein_coding | **230.388** | 0.581 | 0.175 | **1.496** | 4.90E-04 | **1.40E-03** |
| ENSG00000106415.15 | **GLCCI1** | 113263 | protein_coding | **802.588** | 0.421 | 0.127 | **1.339** | 4.91E-04 | **1.40E-03** |
| ENSG00000305995.1 | **ENSG00000305995** | | lncRNA | **16.449** | 1.484 | 0.447 | **2.797** | 4.91E-04 | **1.40E-03** |

| | | | | | | | | | |
|---|---|---|---|---|---|---|---|---|---|
| ENSG00000138316.13 | **ADAMTS14** | 140766 | protein_coding | **6.903** | 2.554 | 0.769 | **5.873** | 4.92E-04 | **1.40E-03** |
| ENSG00000137752.27 | **CASP1** | 834 | protein_coding | **244.261** | -0.690 | -0.208 | **0.620** | 4.92E-04 | **1.41E-03** |
| ENSG00000290782.2 | **CFL1P1** | 142913 | lncRNA | **30.424** | 0.991 | 0.298 | **1.987** | 4.93E-04 | **1.41E-03** |
| ENSG00000185896.13 | **LAMP1** | 3916 | protein_coding | **1123.421** | 0.515 | 0.155 | **1.429** | 4.93E-04 | **1.41E-03** |
| ENSG00000289596.2 | **ENSG00000289596** | | lncRNA | **7.193** | 2.209 | 0.665 | **4.625** | 4.94E-04 | **1.41E-03** |
| ENSG00000263072.11 | **ZNF213-AS** | 10050745‹ | lncRNA | **291.980** | 0.408 | 0.123 | **1.327** | 4.94E-04 | **1.41E-03** |
| ENSG00000149527.19 | **PLCH2** | 9651 | protein_coding | **151.634** | 0.646 | 0.195 | **1.565** | 4.95E-04 | **1.41E-03** |
| ENSG00000287574.2 | **ENSG00000287574** | | lncRNA | **25.948** | 1.046 | 0.315 | **2.065** | 4.95E-04 | **1.41E-03** |
| ENSG00000172466.18 | **ZNF24** | 7572 | protein_coding | **731.362** | -0.360 | -0.109 | **0.779** | 4.95E-04 | **1.41E-03** |
| ENSG00000170915.11 | **PAQR8** | 85315 | protein_coding | **234.072** | 0.441 | 0.133 | **1.358** | 4.97E-04 | **1.42E-03** |
| ENSG00000123643.16 | **SLC36A1** | 206358 | protein_coding | **133.174** | 0.649 | 0.195 | **1.568** | 4.97E-04 | **1.42E-03** |
| ENSG00000125970.14 | **RALY** | 22913 | protein_coding | **4060.523** | -0.350 | -0.105 | **0.785** | 4.98E-04 | **1.42E-03** |
| ENSG00000065923.12 | **SLC9A7** | 84679 | protein_coding | **1776.389** | 0.458 | 0.138 | **1.374** | 4.98E-04 | **1.42E-03** |
| ENSG00000197757.9 | **HOXC6** | 3223 | protein_coding | **23.027** | 1.807 | 0.544 | **3.499** | 4.98E-04 | **1.42E-03** |
| ENSG00000161270.21 | **NPHS1** | 4868 | protein_coding | **6.268** | 3.469 | 1.044 | **11.076** | 4.98E-04 | **1.42E-03** |
| ENSG00000188883.7 | **KLRG2** | 346689 | protein_coding | **7.384** | 2.888 | 0.869 | **7.400** | 4.98E-04 | **1.42E-03** |
| ENSG00000254996.5 | **ANKHD1-E** | 404734 | protein_coding | **889.097** | 0.381 | 0.115 | **1.303** | 4.99E-04 | **1.42E-03** |
| ENSG00000290911.2 | **ENSG0000(** | 10192876‹ | lncRNA | **7.629** | 2.024 | 0.609 | **4.068** | 5.00E-04 | **1.43E-03** |
| ENSG00000104133.18 | **SPG11** | 80208 | protein_coding | **1391.612** | 0.513 | 0.154 | **1.427** | 5.01E-04 | **1.43E-03** |
| ENSG00000185710.10 | **SMG1P4** | | transcribed_unprc | **11.844** | 3.115 | 0.938 | **8.667** | 5.02E-04 | **1.43E-03** |
| ENSG00000290689.2 | **ENSG00000290689** | | lncRNA | **648.675** | 0.617 | 0.186 | **1.533** | 5.04E-04 | **1.44E-03** |
| ENSG00000133816.22 | **MICAL2** | 9645 | protein_coding | **90.621** | 0.926 | 0.279 | **1.900** | 5.06E-04 | **1.44E-03** |
| ENSG00000241360.3 | **PDXP** | 57026 | protein_coding | **647.427** | -0.548 | -0.165 | **0.684** | 5.07E-04 | **1.44E-03** |
| ENSG00000238039.1 | **ENSG00000238039** | | lncRNA | **25.632** | 1.308 | 0.394 | **2.476** | 5.07E-04 | **1.44E-03** |
| ENSG00000003147.21 | **ICA1** | 3382 | protein_coding | **105.431** | 1.126 | 0.339 | **2.183** | 5.07E-04 | **1.44E-03** |
| ENSG00000089157.18 | **RPLP0** | 6175 | protein_coding | **71237.130** | -0.303 | -0.091 | **0.811** | 5.08E-04 | **1.44E-03** |
| ENSG00000227495.2 | **KIF1C-AS1** | | lncRNA | **50.100** | 0.825 | 0.248 | **1.771** | 5.08E-04 | **1.44E-03** |
| ENSG00000265354.6 | **TIMM23** | 10028793‹ | protein_coding | **1252.020** | -0.318 | -0.096 | **0.802** | 5.08E-04 | **1.45E-03** |
| ENSG00000177225.18 | **GATD1** | 347862 | protein_coding | **905.451** | -0.403 | -0.121 | **0.756** | 5.08E-04 | **1.45E-03** |
| ENSG00000089057.17 | **SLC23A2** | 9962 | protein_coding | **499.458** | 0.488 | 0.147 | **1.403** | 5.09E-04 | **1.45E-03** |
| ENSG00000304176.1 | **ENSG00000304176** | | lncRNA | **239.702** | 0.432 | 0.130 | **1.349** | 5.09E-04 | **1.45E-03** |

| | | | | | | | | | |
|---|---|---|---|---|---|---|---|---|---|
| ENSG00000232818.2 | **RPS2P32** | | processed_pseudo | **34.816** | 0.861 | 0.259 | **1.816** | 5.10E-04 | **1.45E-03** |
| ENSG00000300109.1 | **ENSG00000300109** | | lncRNA | **49.535** | -0.864 | -0.260 | **0.549** | 5.11E-04 | **1.45E-03** |
| ENSG00000279161.1 | **ENSG00000279161** | | TEC | **6.609** | 2.705 | 0.814 | **6.520** | 5.11E-04 | **1.45E-03** |
| ENSG00000115353.12 | **TACR1** | 6869 | protein_coding | **15.956** | 2.273 | 0.684 | **4.834** | 5.12E-04 | **1.45E-03** |
| ENSG00000100836.13 | **PABPN1** | 8106 | protein_coding | **7634.968** | 0.298 | 0.090 | **1.229** | 5.13E-04 | **1.46E-03** |
| ENSG00000205571.17 | **SMN2** | 6607 | protein_coding | **672.987** | -0.509 | -0.153 | **0.703** | 5.13E-04 | **1.46E-03** |
| ENSG00000125648.17 | **SLC25A23** | 79085 | protein_coding | **3173.513** | 0.484 | 0.146 | **1.398** | 5.13E-04 | **1.46E-03** |
| ENSG00000286294.3 | **ENSG00000286294** | | lncRNA | **6.939** | 2.056 | 0.619 | **4.158** | 5.14E-04 | **1.46E-03** |
| ENSG00000112739.19 | **PRP4K** | 8899 | protein_coding | **709.680** | -0.392 | -0.118 | **0.762** | 5.14E-04 | **1.46E-03** |
| ENSG00000279207.1 | **ENSG00000279207** | | TEC | **46.000** | 0.982 | 0.296 | **1.976** | 5.17E-04 | **1.47E-03** |
| ENSG00000111907.22 | **TPD52L1** | 7164 | protein_coding | **26.633** | 1.701 | 0.512 | **3.251** | 5.18E-04 | **1.47E-03** |
| ENSG00000255920.5 | **CCND2-AS1** | 10375258 | lncRNA | **14.687** | -1.433 | -0.432 | **0.370** | 5.18E-04 | **1.47E-03** |
| ENSG00000037897.19 | **METTL1** | 4234 | protein_coding | **295.590** | -0.612 | -0.184 | **0.654** | 5.19E-04 | **1.47E-03** |
| ENSG00000255987.1 | **TOMM20P2** | | processed_pseudo | **6.090** | -3.915 | -1.178 | **0.066** | 5.19E-04 | **1.47E-03** |
| ENSG00000106992.21 | **AK1** | 203 | protein_coding | **286.500** | 0.564 | 0.170 | **1.478** | 5.20E-04 | **1.48E-03** |
| ENSG00000277399.6 | **GPR179** | 440435 | protein_coding | **9.562** | 1.662 | 0.500 | **3.166** | 5.20E-04 | **1.48E-03** |
| ENSG00000286594.2 | **ENSG00000286594** | | lncRNA | **6.222** | 2.265 | 0.682 | **4.808** | 5.21E-04 | **1.48E-03** |
| ENSG00000169919.19 | **GUSB** | 2990 | protein_coding | **1016.238** | -0.482 | -0.145 | **0.716** | 5.22E-04 | **1.48E-03** |
| ENSG00000009830.15 | **POMT2** | 29954 | protein_coding | **292.261** | 0.566 | 0.170 | **1.480** | 5.23E-04 | **1.48E-03** |
| ENSG00000205707.13 | **ETFRF1** | 144363 | protein_coding | **125.881** | 0.554 | 0.167 | **1.469** | 5.23E-04 | **1.48E-03** |
| ENSG00000179144.7 | **GIMAP7** | 168537 | protein_coding | **21.436** | -1.294 | -0.390 | **0.408** | 5.25E-04 | **1.49E-03** |
| ENSG00000103978.19 | **TMEM87A** | 25963 | protein_coding | **403.906** | -0.487 | -0.147 | **0.714** | 5.25E-04 | **1.49E-03** |
| ENSG00000175189.5 | **INHBC** | 3626 | protein_coding | **10.362** | 1.614 | 0.486 | **3.061** | 5.26E-04 | **1.49E-03** |
| ENSG00000119943.15 | **PYROXD2** | 84795 | protein_coding | **137.917** | 0.747 | 0.225 | **1.678** | 5.26E-04 | **1.49E-03** |
| ENSG00000246228.8 | **CASC8** | 727677 | lncRNA | **4.206** | 3.783 | 1.139 | **13.769** | 5.27E-04 | **1.49E-03** |
| ENSG00000136628.20 | **EPRS1** | 2058 | protein_coding | **1946.384** | -0.436 | -0.131 | **0.739** | 5.28E-04 | **1.50E-03** |
| ENSG00000171877.22 | **FRMD5** | 84978 | protein_coding | **15.849** | 1.853 | 0.558 | **3.613** | 5.28E-04 | **1.50E-03** |
| ENSG00000153558.19 | **FBXL2** | 25827 | protein_coding | **6.059** | 2.243 | 0.675 | **4.735** | 5.29E-04 | **1.50E-03** |
| ENSG00000109220.13 | **CHIC2** | 26511 | protein_coding | **168.424** | -0.560 | -0.169 | **0.678** | 5.29E-04 | **1.50E-03** |
| ENSG00000176723.12 | **ZNF843** | 283933 | protein_coding | **6.871** | 2.265 | 0.682 | **4.807** | 5.30E-04 | **1.50E-03** |
| ENSG00000100033.19 | **PRODH** | 5625 | protein_coding | **36.344** | 1.316 | 0.396 | **2.490** | 5.30E-04 | **1.50E-03** |

| | | | | | | | | | |
|---|---|---|---|---|---|---|---|---|---|
| ENSG00000229140.13 | **CCDC26** | 137196 | lncRNA | **57.219** | -0.855 | -0.257 | **0.553** | 5.30E-04 | **1.50E-03** |
| ENSG00000290875.2 | **ENSG00000290875** | | lncRNA | **8.043** | 2.257 | 0.679 | **4.780** | 5.30E-04 | **1.50E-03** |
| ENSG00000016391.13 | **CHDH** | 55349 | protein_coding | **569.073** | -0.395 | -0.119 | **0.761** | 5.30E-04 | **1.50E-03** |
| ENSG00000260526.1 | **AP1AR-DT** | | lncRNA | **15.726** | 1.258 | 0.379 | **2.392** | 5.31E-04 | **1.50E-03** |
| ENSG00000088205.15 | **DDX18** | 8886 | protein_coding | **959.588** | -0.420 | -0.126 | **0.747** | 5.31E-04 | **1.50E-03** |
| ENSG00000102678.8 | **FGF9** | 2254 | protein_coding | **11.998** | 2.280 | 0.686 | **4.857** | 5.31E-04 | **1.50E-03** |
| ENSG00000226179.6 | **LINC00685** | | lncRNA | **35.498** | 0.874 | 0.263 | **1.833** | 5.32E-04 | **1.50E-03** |
| ENSG00000075790.14 | **BCAP29** | 55973 | protein_coding | **248.326** | -0.523 | -0.158 | **0.696** | 5.33E-04 | **1.51E-03** |
| ENSG00000163104.20 | **SMARCAD1** | 56916 | protein_coding | **285.160** | -0.552 | -0.166 | **0.682** | 5.33E-04 | **1.51E-03** |
| ENSG00000158234.15 | **FAIM** | 55179 | protein_coding | **94.666** | -0.754 | -0.227 | **0.593** | 5.34E-04 | **1.51E-03** |
| ENSG00000297296.1 | **ENSG00000297296** | | lncRNA | **3.848** | 3.249 | 0.978 | **9.507** | 5.35E-04 | **1.51E-03** |
| ENSG00000162604.15 | **TM2D1** | 83941 | protein_coding | **250.366** | -0.566 | -0.170 | **0.676** | 5.36E-04 | **1.51E-03** |
| ENSG00000166821.10 | **PEX11A** | 8800 | protein_coding | **100.543** | -0.630 | -0.190 | **0.646** | 5.37E-04 | **1.52E-03** |
| ENSG00000155085.17 | **AK9** | 221264 | protein_coding | **89.136** | 0.676 | 0.204 | **1.598** | 5.38E-04 | **1.52E-03** |
| ENSG00000299019.1 | **ENSG00000299019** | | lncRNA | **9.454** | -3.212 | -0.967 | **0.108** | 5.38E-04 | **1.52E-03** |
| ENSG00000143393.19 | **PI4KB** | 5298 | protein_coding | **968.082** | -0.410 | -0.123 | **0.753** | 5.38E-04 | **1.52E-03** |
| ENSG00000228701.4 | **TNKS2-DT** | | lncRNA | **10.631** | 1.668 | 0.502 | **3.178** | 5.40E-04 | **1.52E-03** |
| ENSG00000170604.5 | **IRF2BP1** | 26145 | protein_coding | **393.869** | 0.477 | 0.144 | **1.392** | 5.41E-04 | **1.53E-03** |
| ENSG00000211950.2 | **IGHV1-24** | | IG_V_gene | **11.116** | 2.468 | 0.743 | **5.531** | 5.42E-04 | **1.53E-03** |
| ENSG00000263280.5 | **ENSG0000(** | 10192956( | lncRNA | **20.563** | 1.174 | 0.353 | **2.256** | 5.42E-04 | **1.53E-03** |
| ENSG00000015568.15 | **RGPD5** | 84220 | protein_coding | **161.712** | 0.924 | 0.278 | **1.898** | 5.42E-04 | **1.53E-03** |
| ENSG00000139574.11 | **NPFF** | 8620 | protein_coding | **98.117** | 0.722 | 0.217 | **1.649** | 5.42E-04 | **1.53E-03** |
| ENSG00000165282.18 | **PIGO** | 84720 | protein_coding | **315.683** | -0.351 | -0.106 | **0.784** | 5.43E-04 | **1.53E-03** |
| ENSG00000089248.9 | **ERP29** | 10961 | protein_coding | **3384.222** | -0.328 | -0.099 | **0.796** | 5.43E-04 | **1.53E-03** |
| ENSG00000225484.8 | **NUTM2B-A** | 101060691 | lncRNA | **348.833** | -0.363 | -0.109 | **0.777** | 5.45E-04 | **1.54E-03** |
| ENSG00000101306.13 | **MYLK2** | 85366 | protein_coding | **9.970** | -1.892 | -0.570 | **0.269** | 5.46E-04 | **1.54E-03** |
| ENSG00000250312.9 | **ZNF718** | 255403 | protein_coding | **104.220** | -0.661 | -0.199 | **0.632** | 5.47E-04 | **1.54E-03** |
| ENSG00000115286.23 | **NDUFS7** | 374291 | protein_coding | **1063.294** | -0.476 | -0.143 | **0.719** | 5.48E-04 | **1.54E-03** |
| ENSG00000309149.1 | **ENSG00000309149** | | lncRNA | **7.036** | 2.136 | 0.643 | **4.396** | 5.49E-04 | **1.55E-03** |
| ENSG00000211899.10 | **IGHM** | | IG_C_gene | **31534.704** | -0.419 | -0.126 | **0.748** | 5.50E-04 | **1.55E-03** |
| ENSG00000308311.1 | **ENSG00000308311** | | lncRNA | **10.968** | 1.639 | 0.493 | **3.114** | 5.51E-04 | **1.55E-03** |

| | | | | | | | | | |
|---|---|---|---|---|---|---|---|---|---|
| ENSG00000169989.5 | **TIGD4** | 201798 | protein_coding | **10.845** | 1.804 | 0.543 | **3.493** | 5.51E-04 | **1.55E-03** |
| ENSG00000126107.17 | **HECTD3** | 79654 | protein_coding | **1146.551** | 0.318 | 0.096 | **1.247** | 5.51E-04 | **1.55E-03** |
| ENSG00000116991.14 | **SIPA1L2** | 57568 | protein_coding | **10.676** | 2.356 | 0.709 | **5.118** | 5.52E-04 | **1.55E-03** |
| ENSG00000047249.20 | **ATP6V1H** | 51606 | protein_coding | **608.530** | 0.374 | 0.113 | **1.296** | 5.53E-04 | **1.56E-03** |
| ENSG00000228824.9 | **MIR4500H** | 642345 | lncRNA | **8.537** | 2.744 | 0.826 | **6.698** | 5.53E-04 | **1.56E-03** |
| ENSG00000205791.5 | **LOH12CR2** | 503693 | lncRNA | **12.985** | 1.408 | 0.424 | **2.653** | 5.54E-04 | **1.56E-03** |
| ENSG00000241484.12 | **ARHGAP8** | 23779 | protein_coding | **18.099** | 2.179 | 0.656 | **4.528** | 5.54E-04 | **1.56E-03** |
| ENSG00000293460.1 | **LINC02210** | 147081 | lncRNA | **355.173** | 0.469 | 0.141 | **1.384** | 5.54E-04 | **1.56E-03** |
| ENSG00000288752.3 | **ENSG00000288752** | | lncRNA | **41.154** | 0.806 | 0.243 | **1.749** | 5.55E-04 | **1.56E-03** |
| ENSG00000293310.2 | **ENSG00000293310** | | lncRNA | **54.841** | -0.777 | -0.234 | **0.584** | 5.56E-04 | **1.56E-03** |
| ENSG00000290107.1 | **ENSG00000290107** | | lncRNA | **3.888** | -3.362 | -1.012 | **0.097** | 5.56E-04 | **1.56E-03** |
| ENSG00000119508.19 | **NR4A3** | 8013 | protein_coding | **60.203** | 0.829 | 0.250 | **1.777** | 5.57E-04 | **1.57E-03** |
| ENSG00000158636.19 | **EMSY** | 56946 | protein_coding | **233.692** | -0.763 | -0.230 | **0.589** | 5.57E-04 | **1.57E-03** |
| ENSG00000253948.5 | **VPS13B-DT** | | lncRNA | **48.461** | 0.802 | 0.241 | **1.743** | 5.58E-04 | **1.57E-03** |
| ENSG00000291208.2 | **ENSG00000291208** | | lncRNA | **384.837** | 0.462 | 0.139 | **1.378** | 5.59E-04 | **1.57E-03** |
| ENSG00000271254.7 | **ENSG0000** | 12490556 | protein_coding | **50.094** | 1.041 | 0.314 | **2.058** | 5.59E-04 | **1.57E-03** |
| ENSG00000273419.1 | **ENSG00000273419** | | lncRNA | **5.788** | 2.494 | 0.751 | **5.633** | 5.61E-04 | **1.58E-03** |
| ENSG00000167077.15 | **MEI1** | 150365 | protein_coding | **497.157** | -0.624 | -0.188 | **0.649** | 5.61E-04 | **1.58E-03** |
| ENSG00000170873.21 | **MTSS1** | 9788 | protein_coding | **55.925** | 1.154 | 0.347 | **2.226** | 5.62E-04 | **1.58E-03** |
| ENSG00000218018.5 | **RBM38-AS1** | | lncRNA | **24.225** | -0.984 | -0.296 | **0.506** | 5.63E-04 | **1.58E-03** |
| ENSG00000183513.11 | **COA5** | 493753 | protein_coding | **528.311** | -0.552 | -0.166 | **0.682** | 5.63E-04 | **1.58E-03** |
| ENSG00000224738.3 | **ENSG00000224738** | | lncRNA | **6.833** | -2.270 | -0.683 | **0.207** | 5.64E-04 | **1.58E-03** |
| ENSG00000170242.21 | **USP47** | 55031 | protein_coding | **903.549** | -0.433 | -0.130 | **0.741** | 5.64E-04 | **1.58E-03** |
| ENSG00000257221.5 | **ENSG00000257221** | | lncRNA | **440.791** | 0.620 | 0.187 | **1.537** | 5.65E-04 | **1.59E-03** |
| ENSG00000196361.12 | **ELAVL3** | 1995 | protein_coding | **6.454** | 2.408 | 0.725 | **5.308** | 5.65E-04 | **1.59E-03** |
| ENSG00000159921.22 | **GNE** | 10020 | protein_coding | **219.663** | -0.485 | -0.146 | **0.714** | 5.66E-04 | **1.59E-03** |
| ENSG00000184436.14 | **THAP7** | 80764 | protein_coding | **502.446** | -0.422 | -0.127 | **0.747** | 5.67E-04 | **1.59E-03** |
| ENSG00000272476.1 | **ENSG00000272476** | | lncRNA | **13.774** | 1.677 | 0.505 | **3.197** | 5.68E-04 | **1.59E-03** |
| ENSG00000116191.21 | **RALGPS2** | 55103 | protein_coding | **323.890** | -0.832 | -0.250 | **0.562** | 5.68E-04 | **1.59E-03** |
| ENSG00000198876.15 | **DCAF12** | 25853 | protein_coding | **568.592** | -0.347 | -0.104 | **0.786** | 5.70E-04 | **1.60E-03** |
| ENSG00000168924.17 | **LETM1** | 3954 | protein_coding | **710.619** | -0.394 | -0.119 | **0.761** | 5.70E-04 | **1.60E-03** |

| | | | | | | | | | |
|---|---|---|---|---|---|---|---|---|---|
| ENSG00000110719.12 | **TCIRG1** | 10312 | protein_coding | **6027.542** | -0.426 | -0.128 | **0.744** | 5.71E-04 | **1.60E-03** |
| ENSG00000196591.14 | **HDAC2** | 3066 | protein_coding | **1143.413** | -0.338 | -0.102 | **0.791** | 5.72E-04 | **1.60E-03** |
| ENSG00000135720.15 | **DYNC1LI2** | 1783 | protein_coding | **835.212** | 0.528 | 0.159 | **1.442** | 5.72E-04 | **1.60E-03** |
| ENSG00000186897.7 | **C1QL4** | 338761 | protein_coding | **5.749** | 2.639 | 0.794 | **6.227** | 5.75E-04 | **1.61E-03** |
| ENSG00000114867.24 | **EIF4G1** | 1981 | protein_coding | **3350.954** | -0.389 | -0.117 | **0.764** | 5.75E-04 | **1.61E-03** |
| ENSG00000285712.2 | **ENSG00000285712** | | lncRNA | **9.662** | 2.237 | 0.673 | **4.713** | 5.75E-04 | **1.61E-03** |
| ENSG00000164897.16 | **TMUB1** | 83590 | protein_coding | **1297.154** | -0.305 | -0.092 | **0.810** | 5.77E-04 | **1.62E-03** |
| ENSG00000177337.10 | **DLGAP1-AS1** | | lncRNA | **66.402** | 0.826 | 0.249 | **1.773** | 5.79E-04 | **1.62E-03** |
| ENSG00000116544.13 | **DLGAP3** | 58512 | protein_coding | **37.298** | 1.148 | 0.346 | **2.216** | 5.81E-04 | **1.63E-03** |
| ENSG00000126005.19 | **MMP24OS** | 10141053[illegible] | protein_coding | **491.178** | -0.524 | -0.158 | **0.695** | 5.81E-04 | **1.63E-03** |
| ENSG00000101445.12 | **PPP1R16B** | 26051 | protein_coding | **1496.718** | -0.527 | -0.159 | **0.694** | 5.82E-04 | **1.63E-03** |
| ENSG00000171044.12 | **XKR6** | 286046 | protein_coding | **108.302** | -0.696 | -0.210 | **0.617** | 5.83E-04 | **1.63E-03** |
| ENSG00000087263.19 | **OGFOD1** | 55239 | protein_coding | **708.599** | -0.415 | -0.125 | **0.750** | 5.84E-04 | **1.63E-03** |
| ENSG00000068615.22 | **REEP1** | 65055 | protein_coding | **18.089** | 1.401 | 0.422 | **2.641** | 5.84E-04 | **1.63E-03** |
| ENSG00000008324.14 | **SS18L2** | 51188 | protein_coding | **683.984** | -0.430 | -0.130 | **0.742** | 5.87E-04 | **1.64E-03** |
| ENSG00000130935.12 | **NOL11** | 25926 | protein_coding | **494.903** | -0.503 | -0.151 | **0.706** | 5.89E-04 | **1.65E-03** |
| ENSG00000171453.22 | **POLR1C** | 9533 | protein_coding | **474.973** | -0.388 | -0.117 | **0.764** | 5.90E-04 | **1.65E-03** |
| ENSG00000163785.16 | **RYK** | 6259 | protein_coding | **362.455** | 0.669 | 0.201 | **1.590** | 5.91E-04 | **1.65E-03** |
| ENSG00000091262.19 | **ABCC6** | 368 | protein_coding | **3.823** | 3.610 | 1.087 | **12.206** | 5.92E-04 | **1.65E-03** |
| ENSG00000135899.22 | **SP110** | 3431 | protein_coding | **2482.420** | 0.433 | 0.130 | **1.350** | 5.93E-04 | **1.66E-03** |
| ENSG00000204406.14 | **MBD5** | 55777 | protein_coding | **366.577** | -0.569 | -0.171 | **0.674** | 5.95E-04 | **1.66E-03** |
| ENSG00000123388.4 | **HOXC11** | 3227 | protein_coding | **5.738** | 3.101 | 0.933 | **8.580** | 5.95E-04 | **1.66E-03** |
| ENSG00000291137.2 | **GBA1LP** | 2630 | lncRNA | **375.770** | -0.741 | -0.223 | **0.598** | 5.97E-04 | **1.67E-03** |
| ENSG00000111231.11 | **GPN3** | 51184 | protein_coding | **202.553** | -0.775 | -0.233 | **0.584** | 5.97E-04 | **1.67E-03** |
| ENSG00000280157.1 | **ENSG00000280157** | | TEC | **18.882** | 1.323 | 0.398 | **2.502** | 5.97E-04 | **1.67E-03** |
| ENSG00000214546.5 | **ENSG00000214546** | | lncRNA | **7.791** | 2.104 | 0.633 | **4.300** | 5.97E-04 | **1.67E-03** |
| ENSG00000162739.16 | **SLAMF6** | 114836 | protein_coding | **1624.073** | -0.421 | -0.127 | **0.747** | 5.98E-04 | **1.67E-03** |
| ENSG00000249249.3 | **TICAM2-AS1** | | lncRNA | **67.855** | 0.847 | 0.255 | **1.798** | 5.98E-04 | **1.67E-03** |
| ENSG00000118600.15 | **RXYLT1** | 10329 | protein_coding | **289.685** | -0.403 | -0.121 | **0.756** | 5.98E-04 | **1.67E-03** |
| ENSG00000175787.20 | **ZNF169** | 169841 | protein_coding | **159.129** | 0.486 | 0.146 | **1.401** | 6.00E-04 | **1.67E-03** |
| ENSG00000136930.15 | **PSMB7** | 5695 | protein_coding | **2038.667** | -0.349 | -0.105 | **0.785** | 6.00E-04 | **1.68E-03** |

| | | | | | | | | | |
|---|---|---|---|---|---|---|---|---|---|
| ENSG00000135083.19 | **CCNJL** | 79616 | protein_coding | **9.586** | 1.890 | 0.569 | **3.707** | 6.00E-04 | **1.68E-03** |
| ENSG00000261242.2 | **ENSG00000261242** | | lncRNA | **5.690** | 2.168 | 0.653 | **4.494** | 6.01E-04 | **1.68E-03** |
| ENSG00000100129.19 | **EIF3L** | 51386 | protein_coding | **7637.568** | -0.512 | -0.154 | **0.701** | 6.02E-04 | **1.68E-03** |
| ENSG00000113249.16 | **HAVCR1** | 26762 | protein_coding | **7.259** | 1.940 | 0.584 | **3.837** | 6.03E-04 | **1.68E-03** |
| ENSG00000134152.13 | **KATNBL1** | 79768 | protein_coding | **245.930** | -0.448 | -0.135 | **0.733** | 6.06E-04 | **1.69E-03** |
| ENSG00000116754.16 | **SRSF11** | 9295 | protein_coding | **3494.239** | 0.531 | 0.160 | **1.445** | 6.07E-04 | **1.69E-03** |
| ENSG00000101888.14 | **NXT2** | 55916 | protein_coding | **123.959** | 0.568 | 0.171 | **1.483** | 6.08E-04 | **1.70E-03** |
| ENSG00000205426.11 | **KRT81** | 3887 | protein_coding | **4.786** | 3.439 | 1.035 | **10.845** | 6.09E-04 | **1.70E-03** |
| ENSG00000198837.12 | **DENND4B** | 9909 | protein_coding | **2430.391** | -0.370 | -0.111 | **0.774** | 6.11E-04 | **1.70E-03** |
| ENSG00000290810.2 | **YWHAEP7** | 284100 | lncRNA | **7.748** | 1.771 | 0.533 | **3.413** | 6.12E-04 | **1.71E-03** |
| ENSG00000257497.2 | **GLIPR1-AS1** | | lncRNA | **39.350** | -0.905 | -0.272 | **0.534** | 6.12E-04 | **1.71E-03** |
| ENSG00000243317.10 | **STMP1** | 647087 | protein_coding | **634.129** | -0.512 | -0.154 | **0.701** | 6.16E-04 | **1.71E-03** |
| ENSG00000290767.2 | **ENSG00000290767** | | lncRNA | **13.986** | 1.355 | 0.408 | **2.558** | 6.16E-04 | **1.71E-03** |
| ENSG00000116161.20 | **CACYBP** | 27101 | protein_coding | **1312.183** | -0.466 | -0.140 | **0.724** | 6.17E-04 | **1.72E-03** |
| ENSG00000105364.16 | **MRPL4** | 51073 | protein_coding | **1556.126** | -0.282 | -0.085 | **0.823** | 6.17E-04 | **1.72E-03** |
| ENSG00000186111.13 | **PIP5K1C** | 23396 | protein_coding | **1212.336** | 0.470 | 0.141 | **1.385** | 6.18E-04 | **1.72E-03** |
| ENSG00000137959.19 | **IFI44L** | 10964 | protein_coding | **6944.409** | -0.585 | -0.176 | **0.666** | 6.21E-04 | **1.73E-03** |
| ENSG00000204084.16 | **INPP5B** | 3633 | protein_coding | **950.784** | -0.342 | -0.103 | **0.789** | 6.21E-04 | **1.73E-03** |
| ENSG00000171100.18 | **MTM1** | 4534 | protein_coding | **92.421** | 0.702 | 0.211 | **1.626** | 6.21E-04 | **1.73E-03** |
| ENSG00000165092.15 | **ALDH1A1** | 216 | protein_coding | **4.552** | 3.584 | 1.079 | **11.994** | 6.23E-04 | **1.73E-03** |
| ENSG00000272941.1 | **ENSG00000272941** | | lncRNA | **175.520** | -0.586 | -0.176 | **0.666** | 6.23E-04 | **1.73E-03** |
| ENSG00000259033.3 | **ENSG00000259033** | | lncRNA | **17.088** | 1.725 | 0.519 | **3.305** | 6.23E-04 | **1.73E-03** |
| ENSG00000224903.2 | **RNF32-AS1** | | lncRNA | **43.284** | 0.955 | 0.288 | **1.939** | 6.25E-04 | **1.74E-03** |
| ENSG00000166825.17 | **ANPEP** | 290 | protein_coding | **18.444** | 1.827 | 0.550 | **3.548** | 6.26E-04 | **1.74E-03** |
| ENSG00000142798.21 | **HSPG2** | 3339 | protein_coding | **137.306** | 0.763 | 0.230 | **1.697** | 6.26E-04 | **1.74E-03** |
| ENSG00000300103.1 | **ENSG00000300103** | | lncRNA | **19.083** | -1.222 | -0.368 | **0.429** | 6.27E-04 | **1.74E-03** |
| ENSG00000176619.15 | **LMNB2** | 84823 | protein_coding | **2089.383** | -0.490 | -0.147 | **0.712** | 6.27E-04 | **1.74E-03** |
| ENSG00000254319.7 | **LINC03021** | 10192781! | lncRNA | **37.021** | -1.088 | -0.327 | **0.471** | 6.27E-04 | **1.74E-03** |
| ENSG00000134248.16 | **LAMTOR5** | 10542 | protein_coding | **999.074** | -0.458 | -0.138 | **0.728** | 6.27E-04 | **1.74E-03** |
| ENSG00000162542.16 | **TMCO4** | 255104 | protein_coding | **471.393** | -0.506 | -0.152 | **0.704** | 6.27E-04 | **1.74E-03** |
| ENSG00000174945.16 | **AMZ1** | 155185 | protein_coding | **83.285** | -0.797 | -0.240 | **0.575** | 6.29E-04 | **1.75E-03** |

| | | | | | | | | | |
|---|---|---|---|---|---|---|---|---|---|
| ENSG00000268785.1 | **RPL7P50** | | processed_pseudo | **13.341** | 1.324 | 0.399 | **2.503** | 6.29E-04 | **1.75E-03** |
| ENSG00000248098.14 | **BCKDHA** | 593 | protein_coding | **827.466** | -0.499 | -0.150 | **0.708** | 6.30E-04 | **1.75E-03** |
| ENSG00000290045.2 | **ENSG00000290045** | | lncRNA | **15.561** | 1.208 | 0.364 | **2.311** | 6.31E-04 | **1.75E-03** |
| ENSG00000119778.17 | **ATAD2B** | 54454 | protein_coding | **240.762** | -0.591 | -0.178 | **0.664** | 6.31E-04 | **1.75E-03** |
| ENSG00000257621.10 | **PSMA3-AS** | 379025 | lncRNA | **955.785** | 0.322 | 0.097 | **1.250** | 6.32E-04 | **1.75E-03** |
| ENSG00000080819.11 | **CPOX** | 1371 | protein_coding | **897.466** | 0.742 | 0.223 | **1.672** | 6.34E-04 | **1.76E-03** |
| ENSG00000136514.4 | **RTP4** | 64108 | protein_coding | **95.741** | -0.654 | -0.197 | **0.635** | 6.35E-04 | **1.76E-03** |
| ENSG00000112902.14 | **SEMA5A** | 9037 | protein_coding | **22.382** | 1.578 | 0.475 | **2.987** | 6.36E-04 | **1.77E-03** |
| ENSG00000158406.6 | **H4C8** | 8365 | protein_coding | **122.207** | -0.646 | -0.195 | **0.639** | 6.38E-04 | **1.77E-03** |
| ENSG00000132763.17 | **MMACHC** | 25974 | protein_coding | **146.730** | -0.620 | -0.187 | **0.651** | 6.40E-04 | **1.78E-03** |
| ENSG00000291237.3 | **SOD2** | 6648 | protein_coding | **487.955** | -0.403 | -0.121 | **0.756** | 6.41E-04 | **1.78E-03** |
| ENSG00000280426.1 | **ENSG00000280426** | | TEC | **10.066** | 1.949 | 0.587 | **3.860** | 6.42E-04 | **1.78E-03** |
| ENSG00000287042.4 | **ENSG00000287042** | | lncRNA | **31.659** | 0.906 | 0.273 | **1.874** | 6.42E-04 | **1.78E-03** |
| ENSG00000070495.17 | **JMJD6** | 23210 | protein_coding | **609.561** | -0.400 | -0.120 | **0.758** | 6.43E-04 | **1.78E-03** |
| ENSG00000160299.20 | **PCNT** | 5116 | protein_coding | **1339.948** | -0.386 | -0.116 | **0.765** | 6.44E-04 | **1.79E-03** |
| ENSG00000104312.10 | **RIPK2** | 8767 | protein_coding | **408.237** | -0.408 | -0.123 | **0.754** | 6.46E-04 | **1.79E-03** |
| ENSG00000106404.15 | **CLDN15** | 24146 | protein_coding | **383.520** | 0.415 | 0.125 | **1.333** | 6.46E-04 | **1.79E-03** |
| ENSG00000158161.18 | **EYA3** | 2140 | protein_coding | **856.602** | 0.290 | 0.087 | **1.223** | 6.46E-04 | **1.79E-03** |
| ENSG00000013375.19 | **PGM3** | 5238 | protein_coding | **419.723** | -0.466 | -0.140 | **0.724** | 6.48E-04 | **1.79E-03** |
| ENSG00000286219.3 | **NOTCH2NL** | 10099671 | protein_coding | **16.710** | 1.262 | 0.380 | **2.399** | 6.48E-04 | **1.79E-03** |
| ENSG00000291168.2 | **ANKRD19P** | 138649 | lncRNA | **4.794** | 2.692 | 0.810 | **6.463** | 6.49E-04 | **1.80E-03** |
| ENSG00000115020.19 | **PIKFYVE** | 200576 | protein_coding | **512.045** | 0.571 | 0.172 | **1.486** | 6.50E-04 | **1.80E-03** |
| ENSG00000294760.1 | **ENSG00000294760** | | lncRNA | **7.588** | 1.900 | 0.572 | **3.731** | 6.50E-04 | **1.80E-03** |
| ENSG00000257379.1 | **ENSG00000257379** | | lncRNA | **28.133** | 4.384 | 1.320 | **20.875** | 6.53E-04 | **1.81E-03** |
| ENSG00000116521.14 | **SCAMP3** | 10067 | protein_coding | **748.031** | -0.377 | -0.113 | **0.770** | 6.55E-04 | **1.81E-03** |
| ENSG00000163191.8 | **S100A11** | 6282 | protein_coding | **905.920** | -0.742 | -0.223 | **0.598** | 6.55E-04 | **1.81E-03** |
| ENSG00000271737.3 | **ENSG00000271737** | | lncRNA | **31.676** | 0.937 | 0.282 | **1.915** | 6.56E-04 | **1.82E-03** |
| ENSG00000131477.13 | **RAMP2** | 10266 | protein_coding | **10.493** | 1.680 | 0.506 | **3.205** | 6.57E-04 | **1.82E-03** |
| ENSG00000272681.3 | **FAM223B** | 286967 | lncRNA | **25.278** | 1.061 | 0.319 | **2.086** | 6.58E-04 | **1.82E-03** |
| ENSG00000257261.8 | **SLC38A4-A** | 10028879 | lncRNA | **126.200** | 0.674 | 0.203 | **1.596** | 6.59E-04 | **1.82E-03** |
| ENSG00000175166.19 | **PSMD2** | 5708 | protein_coding | **1603.726** | -0.409 | -0.123 | **0.753** | 6.61E-04 | **1.83E-03** |

| | | | | | | | | | |
|---|---|---|---|---|---|---|---|---|---|
| ENSG00000145216.19 | **FIP1L1** | 81608 | protein_coding | **1134.860** | -0.376 | -0.113 | **0.771** | 6.62E-04 | **1.83E-03** |
| ENSG00000006638.15 | **TBXA2R** | 6915 | protein_coding | **9.971** | 2.137 | 0.643 | **4.399** | 6.62E-04 | **1.83E-03** |
| ENSG00000026297.19 | **RNASET2** | 8635 | protein_coding | **6572.902** | -0.590 | -0.178 | **0.664** | 6.63E-04 | **1.83E-03** |
| ENSG00000067208.19 | **EVI5** | 7813 | protein_coding | **52.319** | 0.823 | 0.248 | **1.770** | 6.63E-04 | **1.83E-03** |
| ENSG00000135778.14 | **NTPCR** | 84284 | protein_coding | **574.727** | -0.470 | -0.141 | **0.722** | 6.63E-04 | **1.83E-03** |
| ENSG00000277701.6 | **ENSG00000277701** | | lncRNA | **58.026** | -0.707 | -0.213 | **0.613** | 6.65E-04 | **1.84E-03** |
| ENSG00000170638.12 | **TRABD** | 80305 | protein_coding | **2236.555** | 0.325 | 0.098 | **1.253** | 6.66E-04 | **1.84E-03** |
| ENSG00000168994.15 | **PXDC1** | 221749 | protein_coding | **112.692** | 0.731 | 0.220 | **1.660** | 6.66E-04 | **1.84E-03** |
| ENSG00000211897.9 | **IGHG3** | | IG_C_gene | **15992.328** | -0.482 | -0.145 | **0.716** | 6.70E-04 | **1.85E-03** |
| ENSG00000186468.15 | **RPS23** | 6228 | protein_coding | **29171.249** | -0.395 | -0.119 | **0.761** | 6.70E-04 | **1.85E-03** |
| ENSG00000167524.19 | **RSKR** | 124923 | protein_coding | **455.544** | 0.376 | 0.113 | **1.297** | 6.72E-04 | **1.86E-03** |
| ENSG00000196810.6 | **CTBP1-DT** | | lncRNA | **273.794** | 0.370 | 0.111 | **1.293** | 6.73E-04 | **1.86E-03** |
| ENSG00000073464.15 | **CLCN4** | 1183 | protein_coding | **40.061** | 1.464 | 0.441 | **2.758** | 6.74E-04 | **1.86E-03** |
| ENSG00000124784.11 | **RIOK1** | 83732 | protein_coding | **331.595** | -0.498 | -0.150 | **0.708** | 6.75E-04 | **1.86E-03** |
| ENSG00000272871.1 | **ENSG00000272871** | | lncRNA | **64.435** | 0.693 | 0.209 | **1.616** | 6.75E-04 | **1.86E-03** |
| ENSG00000106009.18 | **BRAT1** | 221927 | protein_coding | **1175.767** | 0.327 | 0.099 | **1.255** | 6.76E-04 | **1.86E-03** |
| ENSG00000117477.15 | **CCDC181** | 57821 | protein_coding | **5.330** | 3.209 | 0.966 | **9.247** | 6.76E-04 | **1.87E-03** |
| ENSG00000295769.1 | **ENSG00000295769** | | lncRNA | **31.161** | 0.976 | 0.294 | **1.968** | 6.78E-04 | **1.87E-03** |
| ENSG00000104960.15 | **PTOV1** | 53635 | protein_coding | **1721.721** | 0.491 | 0.148 | **1.406** | 6.80E-04 | **1.87E-03** |
| ENSG00000139174.14 | **PRICKLE1** | 144165 | protein_coding | **69.209** | 0.979 | 0.295 | **1.971** | 6.80E-04 | **1.88E-03** |
| ENSG00000119760.18 | **SUPT7L** | 9913 | protein_coding | **1067.052** | -0.291 | -0.088 | **0.817** | 6.82E-04 | **1.88E-03** |
| ENSG00000130035.11 | **GALNT8** | 26290 | protein_coding | **14.023** | 1.821 | 0.548 | **3.534** | 6.82E-04 | **1.88E-03** |
| ENSG00000135823.15 | **STX6** | 10228 | protein_coding | **673.492** | 0.352 | 0.106 | **1.277** | 6.82E-04 | **1.88E-03** |
| ENSG00000171847.13 | **FAM90A1** | 55138 | protein_coding | **31.581** | 0.964 | 0.290 | **1.950** | 6.84E-04 | **1.88E-03** |
| ENSG00000105131.10 | **EPHX3** | 79852 | protein_coding | **6.981** | 2.171 | 0.653 | **4.502** | 6.84E-04 | **1.89E-03** |
| ENSG00000168255.20 | **POLR2J3** | 12325839 | protein_coding | **568.456** | 0.431 | 0.130 | **1.348** | 6.87E-04 | **1.89E-03** |
| ENSG00000112699.13 | **GMDS** | 2762 | protein_coding | **648.677** | 0.411 | 0.124 | **1.330** | 6.87E-04 | **1.89E-03** |
| ENSG00000150593.20 | **PDCD4** | 27250 | protein_coding | **3442.001** | 0.508 | 0.153 | **1.422** | 6.88E-04 | **1.89E-03** |
| ENSG00000160883.13 | **HK3** | 3101 | protein_coding | **5.147** | -2.469 | -0.743 | **0.181** | 6.88E-04 | **1.90E-03** |
| ENSG00000145191.17 | **EIF2B5** | 8893 | protein_coding | **1169.384** | -0.289 | -0.087 | **0.819** | 6.89E-04 | **1.90E-03** |
| ENSG00000225265.3 | **TAF1A-AS1** | | lncRNA | **40.382** | 1.058 | 0.318 | **2.082** | 6.89E-04 | **1.90E-03** |

| | | | | | | | | | |
|---|---|---|---|---|---|---|---|---|---|
| ENSG00000182873.5 | **PRKCZ-AS1** | 10050650[illegible] | lncRNA | **5.223** | 2.946 | 0.887 | **7.706** | 6.89E-04 | **1.90E-03** |
| ENSG00000255571.10 | **MIR9-3HG** | 254559 | lncRNA | **19.475** | 1.496 | 0.450 | **2.820** | 6.91E-04 | **1.90E-03** |
| ENSG00000075292.22 | **ZNF638** | 27332 | protein_coding | **1987.473** | -0.437 | -0.132 | **0.738** | 6.91E-04 | **1.90E-03** |
| ENSG00000122859.7 | **NEUROG3** | 50674 | protein_coding | **7.188** | -2.145 | -0.646 | **0.226** | 6.92E-04 | **1.90E-03** |
| ENSG00000249846.8 | **LINC02021** | 10537410[illegible] | lncRNA | **99.040** | 0.692 | 0.208 | **1.615** | 6.93E-04 | **1.91E-03** |
| ENSG00000298035.1 | **ENSG00000298035** | | lncRNA | **27.364** | 1.013 | 0.305 | **2.018** | 6.93E-04 | **1.91E-03** |
| ENSG00000253522.7 | **MIR3142H[illegible]** | 10707511[illegible] | lncRNA | **1524.312** | 0.664 | 0.200 | **1.585** | 6.94E-04 | **1.91E-03** |
| ENSG00000250742.7 | **LINC02381** | 400043 | lncRNA | **25.609** | 1.094 | 0.329 | **2.135** | 6.94E-04 | **1.91E-03** |
| ENSG00000169756.19 | **LIMS1** | 3987 | protein_coding | **276.619** | -0.498 | -0.150 | **0.708** | 6.95E-04 | **1.91E-03** |
| ENSG00000110925.9 | **CSRNP2** | 81566 | protein_coding | **333.230** | 0.545 | 0.164 | **1.459** | 6.97E-04 | **1.92E-03** |
| ENSG00000170324.22 | **FRMPD2** | 143162 | protein_coding | **25.571** | -1.290 | -0.388 | **0.409** | 6.97E-04 | **1.92E-03** |
| ENSG00000172456.20 | **FGGY** | 55277 | protein_coding | **111.153** | 0.593 | 0.178 | **1.508** | 6.98E-04 | **1.92E-03** |
| ENSG00000214338.12 | **MTCL3** | 387104 | protein_coding | **11.289** | 1.989 | 0.599 | **3.971** | 6.98E-04 | **1.92E-03** |
| ENSG00000146112.14 | **PPP1R18** | 170954 | protein_coding | **2707.961** | -0.490 | -0.148 | **0.712** | 6.99E-04 | **1.92E-03** |
| ENSG00000213214.8 | **ARHGEF35** | 445328 | protein_coding | **39.894** | -1.143 | -0.344 | **0.453** | 6.99E-04 | **1.92E-03** |
| ENSG00000170522.12 | **ELOVL6** | 79071 | protein_coding | **354.305** | 0.814 | 0.245 | **1.759** | 7.00E-04 | **1.92E-03** |
| ENSG00000294789.1 | **ENSG00000294789** | | lncRNA | **131.577** | -0.722 | -0.217 | **0.606** | 7.00E-04 | **1.92E-03** |
| ENSG00000181929.14 | **PRKAG1** | 5571 | protein_coding | **813.705** | -0.333 | -0.100 | **0.794** | 7.01E-04 | **1.92E-03** |
| ENSG00000186814.17 | **ZSCAN30** | 10010146[illegible] | protein_coding | **582.195** | 0.396 | 0.119 | **1.316** | 7.02E-04 | **1.93E-03** |
| ENSG00000161057.15 | **PSMC2** | 5701 | protein_coding | **1438.233** | -0.280 | -0.084 | **0.824** | 7.02E-04 | **1.93E-03** |
| ENSG00000100884.12 | **CPNE6** | 9362 | protein_coding | **10.327** | 1.801 | 0.542 | **3.485** | 7.02E-04 | **1.93E-03** |
| ENSG00000106526.14 | **ACTR3C** | 653857 | protein_coding | **22.938** | -1.133 | -0.341 | **0.456** | 7.02E-04 | **1.93E-03** |
| ENSG00000197603.18 | **CPLANE1** | 65250 | protein_coding | **106.465** | -0.801 | -0.241 | **0.574** | 7.04E-04 | **1.93E-03** |
| ENSG00000143603.21 | **KCNN3** | 3782 | protein_coding | **1348.888** | -0.727 | -0.219 | **0.604** | 7.06E-04 | **1.94E-03** |
| ENSG00000118418.18 | **HMGN3** | 9324 | protein_coding | **2700.645** | 0.390 | 0.118 | **1.311** | 7.06E-04 | **1.94E-03** |
| ENSG00000235269.3 | **LINC02331** | 10537050[illegible] | lncRNA | **14.781** | -2.283 | -0.687 | **0.206** | 7.07E-04 | **1.94E-03** |
| ENSG00000004534.17 | **RBM6** | 10180 | protein_coding | **5689.230** | 0.382 | 0.115 | **1.303** | 7.07E-04 | **1.94E-03** |
| ENSG00000121454.8 | **LHX4** | 89884 | protein_coding | **8.719** | 2.052 | 0.618 | **4.148** | 7.10E-04 | **1.95E-03** |
| ENSG00000114395.13 | **CYB561D2** | 11068 | protein_coding | **221.595** | -0.435 | -0.131 | **0.740** | 7.11E-04 | **1.95E-03** |
| ENSG00000120265.23 | **PCMT1** | 5110 | protein_coding | **718.691** | 0.384 | 0.116 | **1.305** | 7.12E-04 | **1.95E-03** |
| ENSG00000170540.18 | **ARL6IP1** | 23204 | protein_coding | **1238.069** | -0.341 | -0.103 | **0.790** | 7.12E-04 | **1.95E-03** |

| | | | | | | | | | |
|---|---|---|---|---|---|---|---|---|---|
| ENSG00000308554.1 | **ENSG00000308554** | | lncRNA | **45.843** | 0.788 | 0.237 | **1.727** | 7.14E-04 | **1.96E-03** |
| ENSG00000286546.2 | **ENSG0000(** | 613266 | lncRNA | **94.453** | -1.153 | -0.347 | **0.450** | 7.15E-04 | **1.96E-03** |
| ENSG00000152147.13 | **GEMIN6** | 79833 | protein_coding | **314.552** | -0.485 | -0.146 | **0.715** | 7.16E-04 | **1.96E-03** |
| ENSG00000114993.18 | **RTKN** | 6242 | protein_coding | **387.143** | 0.414 | 0.124 | **1.332** | 7.18E-04 | **1.97E-03** |
| ENSG00000277496.1 | **SLCO4A1-AS2** | | lncRNA | **155.488** | 0.517 | 0.156 | **1.431** | 7.19E-04 | **1.97E-03** |
| ENSG00000279348.1 | **ENSG00000279348** | | TEC | **18.051** | 1.115 | 0.336 | **2.166** | 7.20E-04 | **1.97E-03** |
| ENSG00000162241.15 | **SLC25A45** | 283130 | protein_coding | **607.288** | -0.535 | -0.161 | **0.690** | 7.24E-04 | **1.98E-03** |
| ENSG00000086666.22 | **ZFAND6** | 54469 | protein_coding | **563.102** | 0.841 | 0.253 | **1.791** | 7.24E-04 | **1.98E-03** |
| ENSG00000151164.20 | **RAD9B** | 144715 | protein_coding | **23.146** | -1.240 | -0.373 | **0.423** | 7.26E-04 | **1.99E-03** |
| ENSG00000179134.18 | **SAMD4B** | 55095 | protein_coding | **1120.834** | 0.545 | 0.164 | **1.459** | 7.27E-04 | **1.99E-03** |
| ENSG00000148339.15 | **SLC25A25** | 114789 | protein_coding | **236.207** | -0.602 | -0.181 | **0.659** | 7.27E-04 | **1.99E-03** |
| ENSG00000064999.18 | **ANKS1A** | 23294 | protein_coding | **201.817** | 0.452 | 0.136 | **1.368** | 7.27E-04 | **1.99E-03** |
| ENSG00000285967.2 | **NIPBL-DT** | 646719 | lncRNA | **231.723** | -0.669 | -0.201 | **0.629** | 7.28E-04 | **1.99E-03** |
| ENSG00000231119.3 | **ENSG0000(** | 10192737 | lncRNA | **8.028** | -2.064 | -0.621 | **0.239** | 7.28E-04 | **1.99E-03** |
| ENSG00000255441.1 | **SIGLEC10-AS1** | | lncRNA | **15.574** | -1.347 | -0.406 | **0.393** | 7.28E-04 | **1.99E-03** |
| ENSG00000121057.15 | **AKAP1** | 8165 | protein_coding | **661.068** | 0.537 | 0.162 | **1.450** | 7.29E-04 | **1.99E-03** |
| ENSG00000151327.16 | **FAM177A1** | 283635 | protein_coding | **272.561** | -0.443 | -0.133 | **0.736** | 7.30E-04 | **2.00E-03** |
| ENSG00000137841.14 | **PLCB2** | 5330 | protein_coding | **1122.425** | -0.542 | -0.163 | **0.687** | 7.32E-04 | **2.00E-03** |
| ENSG00000118307.22 | **DNAI7** | 55259 | protein_coding | **8.085** | 1.826 | 0.550 | **3.546** | 7.33E-04 | **2.00E-03** |
| ENSG00000120063.12 | **GNA13** | 10672 | protein_coding | **471.018** | -0.427 | -0.128 | **0.744** | 7.36E-04 | **2.01E-03** |
| ENSG00000122335.19 | **SERAC1** | 84947 | protein_coding | **38.032** | 1.040 | 0.313 | **2.056** | 7.36E-04 | **2.01E-03** |
| ENSG00000275645.1 | **NEIL1-AS1** | | lncRNA | **7.500** | 1.787 | 0.538 | **3.451** | 7.37E-04 | **2.01E-03** |
| ENSG00000198502.9 | **HLA-DRB5** | 3127 | protein_coding | **6365.220** | 0.746 | 0.225 | **1.678** | 7.38E-04 | **2.02E-03** |
| ENSG00000197892.14 | **KIF13B** | 23303 | protein_coding | **430.134** | 0.541 | 0.163 | **1.455** | 7.38E-04 | **2.02E-03** |
| ENSG00000107796.16 | **ACTA2** | 59 | protein_coding | **473.518** | -0.416 | -0.125 | **0.749** | 7.41E-04 | **2.02E-03** |
| ENSG00000167904.18 | **TMEM68** | 137695 | protein_coding | **291.453** | -0.358 | -0.108 | **0.780** | 7.44E-04 | **2.03E-03** |
| ENSG00000231346.5 | **LINC01160** | | lncRNA | **116.496** | -0.768 | -0.231 | **0.587** | 7.44E-04 | **2.03E-03** |
| ENSG00000165629.22 | **ATP5F1C** | 509 | protein_coding | **2699.647** | -0.499 | -0.150 | **0.708** | 7.45E-04 | **2.03E-03** |
| ENSG00000121552.5 | **CSTA** | 1475 | protein_coding | **14.079** | 2.268 | 0.683 | **4.817** | 7.46E-04 | **2.04E-03** |
| ENSG00000132716.21 | **DCAF8** | 50717 | protein_coding | **2253.012** | -0.375 | -0.113 | **0.771** | 7.46E-04 | **2.04E-03** |
| ENSG00000135148.14 | **TRAFD1** | 10906 | protein_coding | **1798.061** | 0.498 | 0.150 | **1.413** | 7.47E-04 | **2.04E-03** |

| | | | | | | | | | |
|---|---|---|---|---|---|---|---|---|---|
| ENSG00000168282.7 | **MGAT2** | 4247 | protein_coding | **613.366** | -0.457 | -0.137 | **0.729** | 7.47E-04 | **2.04E-03** |
| ENSG00000177042.18 | **TMEM80** | 283232 | protein_coding | **269.645** | 0.348 | 0.105 | **1.272** | 7.50E-04 | **2.05E-03** |
| ENSG00000157045.10 | **NTAN1** | 123803 | protein_coding | **622.371** | -0.415 | -0.125 | **0.750** | 7.52E-04 | **2.05E-03** |
| ENSG00000104899.8 | **AMH** | 268 | protein_coding | **23.604** | 1.108 | 0.334 | **2.155** | 7.52E-04 | **2.05E-03** |
| ENSG00000233237.10 | **LINC00472** | 79940 | lncRNA | **9.390** | 3.350 | 1.009 | **10.198** | 7.52E-04 | **2.05E-03** |
| ENSG00000162639.18 | **HENMT1** | 113802 | protein_coding | **62.113** | 0.899 | 0.271 | **1.865** | 7.52E-04 | **2.05E-03** |
| ENSG00000164167.15 | **LSM6** | 11157 | protein_coding | **693.861** | -0.385 | -0.116 | **0.766** | 7.53E-04 | **2.05E-03** |
| ENSG00000231769.5 | **ENSG00000231769** | | lncRNA | **25.166** | 1.037 | 0.312 | **2.052** | 7.55E-04 | **2.06E-03** |
| ENSG00000282608.3 | **ADORA3** | 140 | protein_coding | **7.198** | 2.327 | 0.701 | **5.019** | 7.56E-04 | **2.06E-03** |
| ENSG00000179743.5 | **SPEN-AS1** | 729614 | lncRNA | **95.631** | 0.578 | 0.174 | **1.493** | 7.56E-04 | **2.06E-03** |
| ENSG00000168785.10 | **TSPAN5** | 10098 | protein_coding | **38.324** | 1.006 | 0.303 | **2.009** | 7.58E-04 | **2.06E-03** |
| ENSG00000228727.12 | **SAPCD1** | 401251 | protein_coding | **362.164** | -0.577 | -0.174 | **0.670** | 7.60E-04 | **2.07E-03** |
| ENSG00000227486.5 | **ENSG00000227486** | | lncRNA | **86.493** | -0.565 | -0.170 | **0.676** | 7.60E-04 | **2.07E-03** |
| ENSG00000130313.9 | **PGLS** | 25796 | protein_coding | **1452.144** | -0.472 | -0.142 | **0.721** | 7.61E-04 | **2.07E-03** |
| ENSG00000142188.19 | **TMEM50B** | 757 | protein_coding | **548.541** | 0.702 | 0.211 | **1.627** | 7.63E-04 | **2.08E-03** |
| ENSG00000131462.11 | **TUBG1** | 7283 | protein_coding | **1649.940** | -0.365 | -0.110 | **0.777** | 7.63E-04 | **2.08E-03** |
| ENSG00000114120.16 | **SLC25A36** | 55186 | protein_coding | **1020.356** | -0.369 | -0.111 | **0.774** | 7.65E-04 | **2.08E-03** |
| ENSG00000124587.16 | **PEX6** | 5190 | protein_coding | **808.442** | 0.410 | 0.124 | **1.329** | 7.67E-04 | **2.09E-03** |
| ENSG00000075151.25 | **EIF4G3** | 8672 | protein_coding | **870.593** | -0.424 | -0.128 | **0.745** | 7.68E-04 | **2.09E-03** |
| ENSG00000021355.15 | **SERPINB1** | 1992 | protein_coding | **285.561** | 0.682 | 0.205 | **1.605** | 7.73E-04 | **2.10E-03** |
| ENSG00000095596.13 | **CYP26A1** | 1592 | protein_coding | **4.267** | 3.577 | 1.077 | **11.935** | 7.75E-04 | **2.11E-03** |
| ENSG00000286366.2 | **ENSG00000286366** | | lncRNA | **30.222** | 0.987 | 0.297 | **1.982** | 7.75E-04 | **2.11E-03** |
| ENSG00000175066.17 | **GK5** | 256356 | protein_coding | **219.788** | -0.531 | -0.160 | **0.692** | 7.76E-04 | **2.11E-03** |
| ENSG00000278903.6 | **ENSG0000(** | 124905527 | lncRNA | **106.459** | -0.800 | -0.241 | **0.574** | 7.77E-04 | **2.11E-03** |
| ENSG00000296703.1 | **ENSG00000296703** | | lncRNA | **7.877** | 2.457 | 0.740 | **5.491** | 7.78E-04 | **2.11E-03** |
| ENSG00000159899.18 | **NPR2** | 4882 | protein_coding | **223.268** | -0.524 | -0.158 | **0.695** | 7.78E-04 | **2.11E-03** |
| ENSG00000114520.13 | **SNX4** | 8723 | protein_coding | **456.187** | -0.599 | -0.180 | **0.660** | 7.81E-04 | **2.12E-03** |
| ENSG00000094755.19 | **GABRP** | 2568 | protein_coding | **3.583** | 4.015 | 1.209 | **16.172** | 7.84E-04 | **2.13E-03** |
| ENSG00000089558.10 | **KCNH4** | 23415 | protein_coding | **16.367** | 1.179 | 0.355 | **2.264** | 7.85E-04 | **2.13E-03** |
| ENSG00000259205.2 | **PRKXP1** | | processed_pseudo | **28.168** | 0.949 | 0.286 | **1.930** | 7.85E-04 | **2.13E-03** |
| ENSG00000132016.14 | **BRME1** | 79173 | protein_coding | **29.045** | 1.085 | 0.327 | **2.122** | 7.88E-04 | **2.14E-03** |

| | | | | | | | | | |
|---|---|---|---|---|---|---|---|---|---|
| ENSG00000153485.8 | **LYSET** | 26175 | protein_coding | **149.415** | 0.466 | 0.140 | **1.381** | 7.93E-04 | **2.15E-03** |
| ENSG00000290878.2 | **HLA-DRB6** | 3128 | lncRNA | **524.585** | 0.812 | 0.244 | **1.755** | 7.93E-04 | **2.15E-03** |
| ENSG00000170325.18 | **PRDM10** | 56980 | protein_coding | **82.507** | -0.626 | -0.188 | **0.648** | 7.96E-04 | **2.16E-03** |
| ENSG00000229117.11 | **RPL41** | 6171 | protein_coding | **35201.489** | -0.402 | -0.121 | **0.757** | 7.99E-04 | **2.17E-03** |
| ENSG00000163608.17 | **RMP64** | 25871 | protein_coding | **872.485** | 0.344 | 0.104 | **1.270** | 8.01E-04 | **2.17E-03** |
| ENSG00000243566.9 | **UPK3B** | 10537535 | protein_coding | **19.334** | 1.611 | 0.485 | **3.055** | 8.01E-04 | **2.17E-03** |
| ENSG00000170364.15 | **SETMAR** | 6419 | protein_coding | **102.311** | -0.625 | -0.188 | **0.648** | 8.01E-04 | **2.18E-03** |
| ENSG00000134480.19 | **CCNH** | 902 | protein_coding | **560.909** | -0.467 | -0.141 | **0.723** | 8.02E-04 | **2.18E-03** |
| ENSG00000196696.14 | **PDXDC2P-I** | 283970 | lncRNA | **280.517** | 0.759 | 0.229 | **1.693** | 8.02E-04 | **2.18E-03** |
| ENSG00000198589.17 | **LRBA** | 987 | protein_coding | **952.282** | 0.378 | 0.114 | **1.299** | 8.03E-04 | **2.18E-03** |
| ENSG00000178996.16 | **SNX18** | 112574 | protein_coding | **10.638** | 1.663 | 0.501 | **3.167** | 8.05E-04 | **2.18E-03** |
| ENSG00000131828.16 | **PDHA1** | 5160 | protein_coding | **2256.078** | -0.445 | -0.134 | **0.735** | 8.06E-04 | **2.19E-03** |
| ENSG00000163660.14 | **CCNL1** | 57018 | protein_coding | **2777.120** | 0.348 | 0.105 | **1.272** | 8.06E-04 | **2.19E-03** |
| ENSG00000132196.18 | **HSD17B7** | 51478 | protein_coding | **685.597** | -0.572 | -0.172 | **0.672** | 8.07E-04 | **2.19E-03** |
| ENSG00000145780.10 | **FEM1C** | 56929 | protein_coding | **167.766** | -0.610 | -0.184 | **0.655** | 8.08E-04 | **2.19E-03** |
| ENSG00000261786.1 | **ENSG00000261786** | | lncRNA | **388.054** | 0.549 | 0.165 | **1.463** | 8.08E-04 | **2.19E-03** |
| ENSG00000204632.15 | **HLA-G** | 3135 | protein_coding | **57.806** | 0.776 | 0.234 | **1.712** | 8.11E-04 | **2.20E-03** |
| ENSG00000117228.13 | **GBP1** | 2633 | protein_coding | **407.620** | -0.663 | -0.199 | **0.632** | 8.12E-04 | **2.20E-03** |
| ENSG00000307320.1 | **ENSG00000307320** | | lncRNA | **7.424** | 1.715 | 0.516 | **3.283** | 8.13E-04 | **2.20E-03** |
| ENSG00000043143.24 | **JADE2** | 23338 | protein_coding | **845.619** | -0.574 | -0.173 | **0.672** | 8.13E-04 | **2.20E-03** |
| ENSG00000297188.1 | **ENSG00000297188** | | lncRNA | **36.510** | -1.027 | -0.309 | **0.491** | 8.14E-04 | **2.20E-03** |
| ENSG00000289415.1 | **ENSG00000289415** | | lncRNA | **4.961** | 2.531 | 0.762 | **5.779** | 8.14E-04 | **2.20E-03** |
| ENSG00000119950.22 | **MXI1** | 4601 | protein_coding | **596.786** | 0.608 | 0.183 | **1.524** | 8.14E-04 | **2.20E-03** |
| ENSG00000183347.17 | **GBP6** | 163351 | protein_coding | **8.997** | 2.553 | 0.769 | **5.870** | 8.15E-04 | **2.21E-03** |
| ENSG00000156304.17 | **SCAF4** | 57466 | protein_coding | **1298.384** | 0.298 | 0.090 | **1.229** | 8.17E-04 | **2.21E-03** |
| ENSG00000180822.14 | **PSMG4** | 389362 | protein_coding | **482.953** | -0.490 | -0.147 | **0.712** | 8.18E-04 | **2.21E-03** |
| ENSG00000095485.20 | **CWF19L1** | 55280 | protein_coding | **692.612** | -0.301 | -0.091 | **0.811** | 8.19E-04 | **2.22E-03** |
| ENSG00000152601.20 | **MBNL1** | 4154 | protein_coding | **1973.441** | -0.432 | -0.130 | **0.741** | 8.21E-04 | **2.22E-03** |
| ENSG00000182158.18 | **CREB3L2** | 64764 | protein_coding | **1012.063** | -0.493 | -0.148 | **0.711** | 8.25E-04 | **2.23E-03** |
| ENSG00000198142.5 | **SOWAHC** | 65124 | protein_coding | **24.872** | 1.161 | 0.350 | **2.237** | 8.26E-04 | **2.23E-03** |
| ENSG00000151240.20 | **DIP2C** | 22982 | protein_coding | **21.491** | 1.320 | 0.397 | **2.496** | 8.27E-04 | **2.24E-03** |

| | | | | | | | | | |
|---|---|---|---|---|---|---|---|---|---|
| ENSG00000160714.13 | **UBE2Q1** | 55585 | protein_coding | **1612.234** | -0.288 | -0.087 | **0.819** | 8.30E-04 | **2.24E-03** |
| ENSG00000183091.22 | **NEB** | 4703 | protein_coding | **18.969** | 1.330 | 0.400 | **2.514** | 8.32E-04 | **2.25E-03** |
| ENSG00000254860.8 | **TMEM9B-AS1** | | lncRNA | **58.649** | 0.860 | 0.259 | **1.815** | 8.35E-04 | **2.26E-03** |
| ENSG00000108848.17 | **LUC7L3** | 51747 | protein_coding | **3488.865** | 0.451 | 0.136 | **1.367** | 8.35E-04 | **2.26E-03** |
| ENSG00000290861.2 | **KRT42P** | 284116 | lncRNA | **3.947** | 3.285 | 0.989 | **9.745** | 8.36E-04 | **2.26E-03** |
| ENSG00000115289.16 | **PCGF1** | 84759 | protein_coding | **316.825** | -0.356 | -0.107 | **0.781** | 8.36E-04 | **2.26E-03** |
| ENSG00000197332.10 | **ZNF833P** | 401898 | lncRNA | **21.735** | -1.091 | -0.328 | **0.470** | 8.37E-04 | **2.26E-03** |
| ENSG00000279145.2 | **ENSG00000279145** | | lncRNA | **8.723** | 1.688 | 0.508 | **3.221** | 8.37E-04 | **2.26E-03** |
| ENSG00000183889.15 | **NPIPA6** | 13167579( | protein_coding | **1429.240** | -0.424 | -0.128 | **0.745** | 8.39E-04 | **2.27E-03** |
| ENSG00000279117.1 | **ENSG00000279117** | | TEC | **33.742** | 1.268 | 0.382 | **2.408** | 8.40E-04 | **2.27E-03** |
| ENSG00000204516.11 | **MICB** | 4277 | protein_coding | **505.301** | -0.314 | -0.095 | **0.804** | 8.41E-04 | **2.27E-03** |
| ENSG00000136810.15 | **TXN** | 7295 | protein_coding | **4632.261** | -0.527 | -0.159 | **0.694** | 8.41E-04 | **2.27E-03** |
| ENSG00000178802.20 | **MPI** | 4351 | protein_coding | **1000.730** | -0.389 | -0.117 | **0.764** | 8.41E-04 | **2.27E-03** |
| ENSG00000173041.14 | **ZNF680** | 340252 | protein_coding | **106.959** | -0.597 | -0.180 | **0.661** | 8.42E-04 | **2.27E-03** |
| ENSG00000120913.26 | **PDLIM2** | 64236 | protein_coding | **115.825** | -0.681 | -0.205 | **0.624** | 8.43E-04 | **2.27E-03** |
| ENSG00000152784.18 | **PRDM8** | 56978 | protein_coding | **12.532** | 1.559 | 0.469 | **2.947** | 8.43E-04 | **2.27E-03** |
| ENSG00000273812.3 | **LINC02970** | | lncRNA | **10.050** | -2.464 | -0.742 | **0.181** | 8.45E-04 | **2.28E-03** |
| ENSG00000164898.14 | **FMC1** | 154791 | protein_coding | **122.657** | -0.602 | -0.181 | **0.659** | 8.45E-04 | **2.28E-03** |
| ENSG00000184967.9 | **NOC4L** | 79050 | protein_coding | **554.443** | -0.476 | -0.143 | **0.719** | 8.46E-04 | **2.28E-03** |
| ENSG00000306014.1 | **ENSG00000306014** | | lncRNA | **14.268** | 1.348 | 0.406 | **2.545** | 8.47E-04 | **2.28E-03** |
| ENSG00000289319.2 | **ENSG0000(** | 12490295( | lncRNA | **14.120** | 1.222 | 0.368 | **2.332** | 8.47E-04 | **2.28E-03** |
| ENSG00000172493.25 | **AFF1** | 4299 | protein_coding | **1856.518** | 0.359 | 0.108 | **1.283** | 8.49E-04 | **2.29E-03** |
| ENSG00000233977.1 | **ENSG00000233977** | | lncRNA | **4.151** | 3.219 | 0.969 | **9.310** | 8.50E-04 | **2.29E-03** |
| ENSG00000187243.19 | **MAGED4B** | 81557 | protein_coding | **9.705** | 3.431 | 1.033 | **10.787** | 8.50E-04 | **2.29E-03** |
| ENSG00000136868.13 | **SLC31A1** | 1317 | protein_coding | **334.412** | -0.454 | -0.137 | **0.730** | 8.51E-04 | **2.29E-03** |
| ENSG00000239569.6 | **KMT2E-AS** | 723809 | lncRNA | **329.075** | 0.434 | 0.131 | **1.351** | 8.52E-04 | **2.30E-03** |
| ENSG00000272274.6 | **ARGLU1-DT** | | lncRNA | **10.658** | 1.453 | 0.438 | **2.739** | 8.53E-04 | **2.30E-03** |
| ENSG00000120832.13 | **MTERF2** | 80298 | protein_coding | **255.686** | 0.432 | 0.130 | **1.349** | 8.54E-04 | **2.30E-03** |
| ENSG00000122884.15 | **P4HA1** | 5033 | protein_coding | **1241.127** | -0.427 | -0.129 | **0.744** | 8.54E-04 | **2.30E-03** |
| ENSG00000178015.5 | **GPR150** | 285601 | protein_coding | **14.616** | 1.320 | 0.397 | **2.496** | 8.54E-04 | **2.30E-03** |
| ENSG00000121236.23 | **TRIM6** | 117854 | protein_coding | **106.902** | 0.587 | 0.177 | **1.502** | 8.57E-04 | **2.31E-03** |

| | | | | | | | | | |
|---|---|---|---|---|---|---|---|---|---|
| ENSG00000146535.17 | **GNA12** | 2768 | protein_coding | **650.802** | 0.524 | 0.158 | **1.438** | 8.58E-04 | **2.31E-03** |
| ENSG00000047634.17 | **SCML1** | 6322 | protein_coding | **164.207** | 0.757 | 0.228 | **1.690** | 8.58E-04 | **2.31E-03** |
| ENSG00000092445.14 | **TYRO3** | 7301 | protein_coding | **13.688** | 1.429 | 0.430 | **2.693** | 8.58E-04 | **2.31E-03** |
| ENSG00000272143.3 | **FGF14-AS2** | 283481 | lncRNA | **24.051** | 0.920 | 0.277 | **1.892** | 8.59E-04 | **2.31E-03** |
| ENSG00000277851.3 | **LINC02391** | 10193002: | lncRNA | **29.030** | 0.970 | 0.292 | **1.958** | 8.62E-04 | **2.32E-03** |
| ENSG00000180035.14 | **ZNF48** | 197407 | protein_coding | **172.634** | 0.557 | 0.168 | **1.472** | 8.63E-04 | **2.32E-03** |
| ENSG00000249700.10 | **SRD5A3-AS** | 10050646. | lncRNA | **8.048** | 2.043 | 0.615 | **4.121** | 8.67E-04 | **2.33E-03** |
| ENSG00000181396.15 | **OGFOD3** | 79701 | protein_coding | **409.553** | -0.470 | -0.142 | **0.722** | 8.67E-04 | **2.33E-03** |
| ENSG00000075420.15 | **FNDC3B** | 64778 | protein_coding | **217.725** | 0.525 | 0.158 | **1.439** | 8.68E-04 | **2.33E-03** |
| ENSG00000197299.15 | **BLM** | 641 | protein_coding | **560.909** | -0.672 | -0.202 | **0.628** | 8.70E-04 | **2.34E-03** |
| ENSG00000090661.15 | **CERS4** | 79603 | protein_coding | **807.022** | -0.468 | -0.141 | **0.723** | 8.70E-04 | **2.34E-03** |
| ENSG00000261373.1 | **VPS9D1-AS** | 100128881 | lncRNA | **73.064** | -0.669 | -0.201 | **0.629** | 8.71E-04 | **2.34E-03** |
| ENSG00000260101.3 | **AFTPH-DT** | | lncRNA | **106.976** | -0.844 | -0.254 | **0.557** | 8.71E-04 | **2.34E-03** |
| ENSG00000150990.11 | **DHX37** | 57647 | protein_coding | **675.890** | -0.341 | -0.103 | **0.790** | 8.73E-04 | **2.34E-03** |
| ENSG00000138398.19 | **PPIG** | 9360 | protein_coding | **862.281** | -0.452 | -0.136 | **0.731** | 8.73E-04 | **2.35E-03** |
| ENSG00000215105.4 | **TTC3P1** | | transcribed_proce | **5.065** | 2.306 | 0.694 | **4.945** | 8.73E-04 | **2.35E-03** |
| ENSG00000163536.15 | **SERPINI1** | 5274 | protein_coding | **99.189** | 0.724 | 0.218 | **1.652** | 8.76E-04 | **2.35E-03** |
| ENSG00000237943.8 | **PRKCQ-AS:** | 439949 | lncRNA | **8.831** | 1.933 | 0.582 | **3.819** | 8.77E-04 | **2.36E-03** |
| ENSG00000226510.3 | **UPK1A-AS1** | | lncRNA | **9.420** | 1.854 | 0.558 | **3.614** | 8.78E-04 | **2.36E-03** |
| ENSG00000261879.10 | **ZNF594-DT** | 10013095( | lncRNA | **75.413** | 0.613 | 0.185 | **1.529** | 8.80E-04 | **2.36E-03** |
| ENSG00000100348.12 | **TXN2** | 25828 | protein_coding | **1862.566** | -0.285 | -0.086 | **0.821** | 8.83E-04 | **2.37E-03** |
| ENSG00000179715.15 | **PCED1B** | 91523 | protein_coding | **194.368** | -0.648 | -0.195 | **0.638** | 8.85E-04 | **2.37E-03** |
| ENSG00000196323.16 | **ZBTB44** | 29068 | protein_coding | **410.263** | -0.425 | -0.128 | **0.745** | 8.88E-04 | **2.38E-03** |
| ENSG00000100450.15 | **GZMH** | 2999 | protein_coding | **12.644** | 1.849 | 0.556 | **3.601** | 8.89E-04 | **2.38E-03** |
| ENSG00000295377.1 | **ENSG00000295377** | | lncRNA | **8.638** | 1.875 | 0.565 | **3.669** | 8.89E-04 | **2.38E-03** |
| ENSG00000135047.18 | **CTSL** | 1514 | protein_coding | **6.245** | 2.035 | 0.613 | **4.098** | 8.89E-04 | **2.38E-03** |
| ENSG00000173674.13 | **EIF1AX** | 1964 | protein_coding | **881.092** | -0.383 | -0.115 | **0.767** | 8.91E-04 | **2.39E-03** |
| ENSG00000136193.19 | **SCRN1** | 9805 | protein_coding | **1024.090** | -0.474 | -0.143 | **0.720** | 8.92E-04 | **2.39E-03** |
| ENSG00000166863.15 | **TAC3** | 6866 | protein_coding | **23.967** | 1.496 | 0.450 | **2.821** | 8.93E-04 | **2.39E-03** |
| ENSG00000290960.2 | **ENSG00000290960** | | lncRNA | **8.410** | 2.071 | 0.623 | **4.202** | 8.95E-04 | **2.40E-03** |
| ENSG00000132254.15 | **ARFIP2** | 23647 | protein_coding | **1049.552** | 0.329 | 0.099 | **1.256** | 8.95E-04 | **2.40E-03** |

| | | | | | | | | | |
|---|---|---|---|---|---|---|---|---|---|
| ENSG00000223442.2 | **TH2LCRR** | 101927761 | lncRNA | **23.987** | 1.350 | 0.406 | **2.549** | 9.00E-04 | **2.41E-03** |
| ENSG00000105278.14 | **ZFR2** | 23217 | protein_coding | **15.080** | 1.706 | 0.513 | **3.262** | 9.05E-04 | **2.43E-03** |
| ENSG00000129691.18 | **ASH2L** | 9070 | protein_coding | **538.851** | -0.360 | -0.109 | **0.779** | 9.08E-04 | **2.43E-03** |
| ENSG00000146205.17 | **ANO7** | 50636 | protein_coding | **24.460** | -0.965 | -0.290 | **0.512** | 9.09E-04 | **2.44E-03** |
| ENSG00000282034.2 | **ENSG00000282034** | | protein_coding | **526.759** | -0.438 | -0.132 | **0.738** | 9.10E-04 | **2.44E-03** |
| ENSG00000226419.10 | **SLC16A1-A** | 100506392 | lncRNA | **219.013** | -0.470 | -0.141 | **0.722** | 9.10E-04 | **2.44E-03** |
| ENSG00000129484.16 | **PARP2** | 10038 | protein_coding | **667.568** | -0.368 | -0.111 | **0.775** | 9.10E-04 | **2.44E-03** |
| ENSG00000029534.23 | **ANK1** | 286 | protein_coding | **155.056** | -0.601 | -0.181 | **0.659** | 9.11E-04 | **2.44E-03** |
| ENSG00000223653.7 | **BCL10-AS1** | | lncRNA | **10.679** | -1.578 | -0.475 | **0.335** | 9.11E-04 | **2.44E-03** |
| ENSG00000140988.18 | **RPS2** | 6187 | protein_coding | **79794.694** | -0.383 | -0.115 | **0.767** | 9.12E-04 | **2.44E-03** |
| ENSG00000145996.13 | **CDKAL1** | 54901 | protein_coding | **270.659** | -0.416 | -0.125 | **0.750** | 9.12E-04 | **2.44E-03** |
| ENSG00000125814.19 | **NAPB** | 63908 | protein_coding | **454.174** | 0.426 | 0.128 | **1.344** | 9.12E-04 | **2.44E-03** |
| ENSG00000228709.3 | **LINC02575** | | lncRNA | **6.965** | 2.404 | 0.724 | **5.293** | 9.14E-04 | **2.44E-03** |
| ENSG00000182197.14 | **EXT1** | 2131 | protein_coding | **155.189** | 0.504 | 0.152 | **1.418** | 9.14E-04 | **2.45E-03** |
| ENSG00000137575.14 | **SDCBP** | 6386 | protein_coding | **250.826** | -0.532 | -0.160 | **0.692** | 9.15E-04 | **2.45E-03** |
| ENSG00000196081.11 | **ZNF724** | 440519 | protein_coding | **25.619** | -1.502 | -0.452 | **0.353** | 9.16E-04 | **2.45E-03** |
| ENSG00000117448.16 | **AKR1A1** | 10327 | protein_coding | **2548.733** | -0.442 | -0.133 | **0.736** | 9.22E-04 | **2.46E-03** |
| ENSG00000048342.20 | **CC2D2A** | 57545 | protein_coding | **13.190** | 2.250 | 0.677 | **4.756** | 9.22E-04 | **2.47E-03** |
| ENSG00000169933.17 | **FRMPD4** | 9758 | protein_coding | **6.163** | 3.408 | 1.026 | **10.618** | 9.23E-04 | **2.47E-03** |
| ENSG00000170892.15 | **TSEN34** | 79042 | protein_coding | **438.277** | 0.474 | 0.143 | **1.389** | 9.24E-04 | **2.47E-03** |
| ENSG00000170043.14 | **TRAPPC1** | 58485 | protein_coding | **2767.219** | 0.462 | 0.139 | **1.377** | 9.27E-04 | **2.48E-03** |
| ENSG00000309919.1 | **ENSG00000309919** | | lncRNA | **4.059** | 2.984 | 0.898 | **7.913** | 9.30E-04 | **2.48E-03** |
| ENSG00000258819.3 | **LINC02289** | 102724190 | lncRNA | **6.269** | 2.006 | 0.604 | **4.015** | 9.31E-04 | **2.49E-03** |
| ENSG00000165731.24 | **RET** | 5979 | protein_coding | **6.011** | 2.465 | 0.742 | **5.522** | 9.33E-04 | **2.49E-03** |
| ENSG00000177854.10 | **TMEM187** | 8269 | protein_coding | **63.851** | 0.705 | 0.212 | **1.630** | 9.34E-04 | **2.50E-03** |
| ENSG00000122299.13 | **ZC3H7A** | 29066 | protein_coding | **1690.988** | -0.316 | -0.095 | **0.803** | 9.35E-04 | **2.50E-03** |
| ENSG00000180992.9 | **MRPL14** | 64928 | protein_coding | **776.296** | -0.487 | -0.147 | **0.713** | 9.37E-04 | **2.50E-03** |
| ENSG00000283913.2 | **ENSG00000283913** | | lncRNA | **13.387** | 1.829 | 0.551 | **3.554** | 9.39E-04 | **2.51E-03** |
| ENSG00000145506.16 | **NKD2** | 85409 | protein_coding | **43.907** | 1.467 | 0.442 | **2.765** | 9.41E-04 | **2.51E-03** |
| ENSG00000172671.22 | **ZFAND4** | 93550 | protein_coding | **190.468** | -0.515 | -0.155 | **0.700** | 9.43E-04 | **2.52E-03** |
| ENSG00000004455.20 | **AK2** | 204 | protein_coding | **3943.683** | -0.331 | -0.100 | **0.795** | 9.48E-04 | **2.53E-03** |

| | | | | | | | | | |
|---|---|---|---|---|---|---|---|---|---|
| ENSG00000288934.3 | **ENSG00000288934** | | lncRNA | **23.880** | -1.025 | -0.308 | **0.492** | 9.49E-04 | **2.53E-03** |
| ENSG00000165929.15 | **TC2N** | 123036 | protein_coding | **186.794** | 0.927 | 0.279 | **1.901** | 9.51E-04 | **2.54E-03** |
| ENSG00000189120.7 | **SP6** | 80320 | protein_coding | **8.002** | 1.604 | 0.483 | **3.040** | 9.52E-04 | **2.54E-03** |
| ENSG00000176720.8 | **BOK** | 666 | protein_coding | **4.519** | 3.165 | 0.953 | **8.972** | 9.52E-04 | **2.54E-03** |
| ENSG00000152749.10 | **GPR180** | 160897 | protein_coding | **115.543** | -0.701 | -0.211 | **0.615** | 9.53E-04 | **2.54E-03** |
| ENSG00000167588.15 | **GPD1** | 2819 | protein_coding | **19.065** | 1.147 | 0.345 | **2.215** | 9.53E-04 | **2.54E-03** |
| ENSG00000184677.20 | **ZBTB40** | 9923 | protein_coding | **1205.017** | 0.408 | 0.123 | **1.327** | 9.54E-04 | **2.54E-03** |
| ENSG00000253320.8 | **MAILR** | | lncRNA | **240.966** | 0.378 | 0.114 | **1.300** | 9.57E-04 | **2.55E-03** |
| ENSG00000125633.13 | **CCDC93** | 54520 | protein_coding | **483.304** | -0.494 | -0.149 | **0.710** | 9.57E-04 | **2.55E-03** |
| ENSG00000178952.13 | **TUFM** | 7284 | protein_coding | **3499.472** | -0.339 | -0.102 | **0.791** | 9.59E-04 | **2.55E-03** |
| ENSG00000137478.17 | **FCHSD2** | 9873 | protein_coding | **1172.902** | -0.464 | -0.140 | **0.725** | 9.62E-04 | **2.56E-03** |
| ENSG00000141298.20 | **SSH2** | 85464 | protein_coding | **839.629** | -0.435 | -0.131 | **0.740** | 9.62E-04 | **2.56E-03** |
| ENSG00000245812.6 | **LINC02202** | 10192774( | lncRNA | **68.246** | -0.729 | -0.220 | **0.603** | 9.63E-04 | **2.57E-03** |
| ENSG00000308739.1 | **ENSG00000308739** | | lncRNA | **9.674** | 1.542 | 0.464 | **2.912** | 9.64E-04 | **2.57E-03** |
| ENSG00000117505.15 | **DR1** | 1810 | protein_coding | **598.235** | -0.720 | -0.217 | **0.607** | 9.65E-04 | **2.57E-03** |
| ENSG00000259488.2 | **DUT-AS1** | | lncRNA | **20.054** | -1.123 | -0.338 | **0.459** | 9.67E-04 | **2.57E-03** |
| ENSG00000157895.13 | **C12orf43** | 64897 | protein_coding | **282.204** | -0.378 | -0.114 | **0.769** | 9.68E-04 | **2.58E-03** |
| ENSG00000175779.6 | **LINC02694** | 400359 | lncRNA | **13.093** | 1.525 | 0.459 | **2.877** | 9.69E-04 | **2.58E-03** |
| ENSG00000277149.7 | **TYW1B** | 441250 | protein_coding | **68.360** | 0.687 | 0.207 | **1.609** | 9.69E-04 | **2.58E-03** |
| ENSG00000274265.6 | **ENSG00000274265** | | lncRNA | **130.436** | -0.663 | -0.200 | **0.631** | 9.69E-04 | **2.58E-03** |
| ENSG00000105559.15 | **PLEKHA4** | 57664 | protein_coding | **226.167** | 0.560 | 0.169 | **1.475** | 9.70E-04 | **2.58E-03** |
| ENSG00000175274.21 | **TP53I11** | 9537 | protein_coding | **13.377** | 1.644 | 0.495 | **3.125** | 9.72E-04 | **2.59E-03** |
| ENSG00000123146.22 | **ADGRE5** | 976 | protein_coding | **1457.055** | -0.471 | -0.142 | **0.722** | 9.73E-04 | **2.59E-03** |
| ENSG00000177103.15 | **DSCAML1** | 57453 | protein_coding | **37.925** | -1.328 | -0.400 | **0.398** | 9.74E-04 | **2.59E-03** |
| ENSG00000181240.14 | **SLC25A41** | 284427 | protein_coding | **19.697** | 1.399 | 0.421 | **2.636** | 9.74E-04 | **2.59E-03** |
| ENSG00000177311.14 | **ZBTB38** | 253461 | protein_coding | **1231.803** | -0.758 | -0.228 | **0.591** | 9.77E-04 | **2.60E-03** |
| ENSG00000198429.12 | **ZNF69** | 7620 | protein_coding | **154.465** | 0.717 | 0.216 | **1.644** | 9.79E-04 | **2.60E-03** |
| ENSG00000287603.1 | **ENSG00000287603** | | lncRNA | **8.863** | -2.685 | -0.808 | **0.156** | 9.84E-04 | **2.62E-03** |
| ENSG00000282508.3 | **LINC01002** | | lncRNA | **273.744** | 0.361 | 0.109 | **1.284** | 9.84E-04 | **2.62E-03** |
| ENSG00000179899.8 | **PHC1P1** | | processed_pseud( | **19.064** | 1.623 | 0.489 | **3.081** | 9.90E-04 | **2.63E-03** |
| ENSG00000059691.14 | **GATB** | 5188 | protein_coding | **420.386** | -0.492 | -0.148 | **0.711** | 9.96E-04 | **2.65E-03** |

| | | | | | | | | | |
|---|---|---|---|---|---|---|---|---|---|
| ENSG00000198556.16 | **ZNF789** | 285989 | protein_coding | **320.452** | -0.536 | -0.161 | **0.690** | 9.97E-04 | **2.65E-03** |
| ENSG00000285601.2 | **ENSG00000285601** | | lncRNA | **13.602** | 1.464 | 0.441 | **2.759** | 9.97E-04 | **2.65E-03** |
| ENSG00000293317.2 | **ZNF702P** | 79986 | lncRNA | **11.603** | 2.097 | 0.631 | **4.277** | 9.98E-04 | **2.65E-03** |
| ENSG00000177096.11 | **PHETA2** | 150368 | protein_coding | **45.930** | -0.751 | -0.226 | **0.594** | 1.00E-03 | **2.66E-03** |
| ENSG00000134250.23 | **NOTCH2** | 4853 | protein_coding | **86.937** | 1.284 | 0.386 | **2.435** | 1.00E-03 | **2.66E-03** |
| ENSG00000073921.20 | **PICALM** | 8301 | protein_coding | **882.161** | 0.486 | 0.146 | **1.401** | 1.01E-03 | **2.67E-03** |
| ENSG00000241973.13 | **PI4KA** | 5297 | protein_coding | **1389.375** | 0.310 | 0.093 | **1.239** | 1.01E-03 | **2.67E-03** |
| ENSG00000110721.14 | **CHKA** | 1119 | protein_coding | **1210.625** | -0.278 | -0.084 | **0.825** | 1.01E-03 | **2.68E-03** |
| ENSG00000291079.2 | **SDHAP3** | 728609 | lncRNA | **1002.843** | -0.467 | -0.141 | **0.724** | 1.01E-03 | **2.68E-03** |
| ENSG00000171604.14 | **CXXC5** | 51523 | protein_coding | **1269.650** | -0.393 | -0.118 | **0.762** | 1.01E-03 | **2.69E-03** |
| ENSG00000267892.1 | **ENSG00000267892** | | lncRNA | **17.979** | 1.422 | 0.428 | **2.680** | 1.02E-03 | **2.69E-03** |
| ENSG00000230138.2 | **LINC02812** | | lncRNA | **5.265** | -2.785 | -0.839 | **0.145** | 1.02E-03 | **2.70E-03** |
| ENSG00000197989.16 | **SNHG12** | 85028 | lncRNA | **1500.746** | -0.480 | -0.145 | **0.717** | 1.02E-03 | **2.71E-03** |
| ENSG00000131437.18 | **KIF3A** | 11127 | protein_coding | **595.713** | 0.551 | 0.166 | **1.465** | 1.02E-03 | **2.72E-03** |
| ENSG00000242759.10 | **LINC00882** | | lncRNA | **18.485** | 1.196 | 0.360 | **2.290** | 1.02E-03 | **2.72E-03** |
| ENSG00000274443.5 | **C8orf89** | 100130301 | protein_coding | **29.430** | 1.032 | 0.311 | **2.045** | 1.02E-03 | **2.72E-03** |
| ENSG00000164985.17 | **PSIP1** | 11168 | protein_coding | **3280.836** | 0.480 | 0.144 | **1.395** | 1.02E-03 | **2.72E-03** |
| ENSG00000227388.4 | **ENSG00000227388** | | lncRNA | **57.499** | -0.815 | -0.245 | **0.569** | 1.03E-03 | **2.72E-03** |
| ENSG00000120899.20 | **PTK2B** | 2185 | protein_coding | **9789.600** | 0.557 | 0.168 | **1.471** | 1.03E-03 | **2.72E-03** |
| ENSG00000185651.18 | **UBE2L3** | 7332 | protein_coding | **2640.665** | -0.366 | -0.110 | **0.776** | 1.03E-03 | **2.72E-03** |
| ENSG00000259584.2 | **ENSG0000(** | 105370787 | lncRNA | **7.801** | 2.312 | 0.696 | **4.965** | 1.03E-03 | **2.72E-03** |
| ENSG00000160062.17 | **ZBTB8A** | 653121 | protein_coding | **23.768** | -1.063 | -0.320 | **0.479** | 1.03E-03 | **2.72E-03** |
| ENSG00000304841.1 | **ENSG00000304841** | | lncRNA | **11.704** | 1.348 | 0.406 | **2.546** | 1.03E-03 | **2.73E-03** |
| ENSG00000106682.17 | **EIF4H** | 7458 | protein_coding | **4999.766** | -0.319 | -0.096 | **0.802** | 1.03E-03 | **2.73E-03** |
| ENSG00000118046.20 | **STK11** | 6794 | protein_coding | **1721.880** | 0.281 | 0.085 | **1.215** | 1.03E-03 | **2.74E-03** |
| ENSG00000168884.17 | **TNIP2** | 79155 | protein_coding | **1513.799** | 0.383 | 0.115 | **1.304** | 1.04E-03 | **2.75E-03** |
| ENSG00000126709.17 | **IFI6** | 2537 | protein_coding | **2676.126** | 0.470 | 0.141 | **1.385** | 1.04E-03 | **2.75E-03** |
| ENSG00000225190.14 | **PLEKHM1** | 9842 | protein_coding | **335.146** | 0.402 | 0.121 | **1.321** | 1.04E-03 | **2.76E-03** |
| ENSG00000115866.13 | **DARS1** | 1615 | protein_coding | **1862.903** | -0.370 | -0.112 | **0.774** | 1.04E-03 | **2.76E-03** |
| ENSG00000127124.16 | **HIVEP3** | 59269 | protein_coding | **425.561** | -0.577 | -0.174 | **0.670** | 1.04E-03 | **2.76E-03** |
| ENSG00000101457.15 | **DNTTIP1** | 116092 | protein_coding | **457.270** | -0.322 | -0.097 | **0.800** | 1.04E-03 | **2.76E-03** |

| | | | | | | | | | |
|---|---|---|---|---|---|---|---|---|---|
| ENSG00000184613.13 | **NELL2** | 4753 | protein_coding | **6.377** | 3.253 | 0.979 | **9.533** | 1.04E-03 | **2.76E-03** |
| ENSG00000115091.14 | **ACTR3** | 10096 | protein_coding | **3342.304** | -0.371 | -0.112 | **0.773** | 1.04E-03 | **2.76E-03** |
| ENSG00000225921.9 | **NOL7** | 51406 | protein_coding | **1103.143** | -0.329 | -0.099 | **0.796** | 1.05E-03 | **2.77E-03** |
| ENSG00000007080.13 | **CCDC124** | 115098 | protein_coding | **1240.629** | 0.414 | 0.125 | **1.332** | 1.05E-03 | **2.77E-03** |
| ENSG00000230587.6 | **LINC02580** | 10798587( | lncRNA | **4.182** | 3.306 | 0.995 | **9.891** | 1.05E-03 | **2.78E-03** |
| ENSG00000115761.19 | **NOL10** | 79954 | protein_coding | **473.596** | -0.359 | -0.108 | **0.780** | 1.05E-03 | **2.78E-03** |
| ENSG00000237522.1 | **NONOP2** | | processed_pseud( | **14.434** | 1.470 | 0.442 | **2.770** | 1.06E-03 | **2.79E-03** |
| ENSG00000136527.21 | **TRA2B** | 6434 | protein_coding | **1916.545** | -0.371 | -0.112 | **0.773** | 1.06E-03 | **2.79E-03** |
| ENSG00000154079.7 | **SDHAF4** | 135154 | protein_coding | **285.032** | -0.409 | -0.123 | **0.753** | 1.06E-03 | **2.79E-03** |
| ENSG00000215883.13 | **CYB5RL** | 606495 | protein_coding | **148.108** | -0.483 | -0.145 | **0.715** | 1.06E-03 | **2.79E-03** |
| ENSG00000100221.13 | **JOSD1** | 9929 | protein_coding | **868.085** | -0.378 | -0.114 | **0.769** | 1.06E-03 | **2.80E-03** |
| ENSG00000230844.4 | **ZNF674-AS1** | | lncRNA | **103.388** | 0.601 | 0.181 | **1.516** | 1.06E-03 | **2.80E-03** |
| ENSG00000143633.15 | **FSAF1** | 128061 | protein_coding | **216.080** | -0.574 | -0.173 | **0.672** | 1.06E-03 | **2.80E-03** |
| ENSG00000170832.15 | **USP32** | 84669 | protein_coding | **555.010** | -0.329 | -0.099 | **0.796** | 1.06E-03 | **2.81E-03** |
| ENSG00000066629.21 | **EML1** | 2009 | protein_coding | **10.072** | 1.942 | 0.584 | **3.841** | 1.06E-03 | **2.81E-03** |
| ENSG00000189171.18 | **S100A13** | 6284 | protein_coding | **311.406** | 0.451 | 0.136 | **1.367** | 1.06E-03 | **2.81E-03** |
| ENSG00000163874.13 | **ZC3H12A** | 80149 | protein_coding | **905.467** | 0.396 | 0.119 | **1.316** | 1.06E-03 | **2.81E-03** |
| ENSG00000158711.16 | **ELK4** | 2005 | protein_coding | **829.779** | -0.355 | -0.107 | **0.782** | 1.06E-03 | **2.81E-03** |
| ENSG00000224609.9 | **FGGY-DT** | | lncRNA | **14.987** | 1.486 | 0.447 | **2.801** | 1.07E-03 | **2.81E-03** |
| ENSG00000163597.18 | **SNHG16** | 10050724( | lncRNA | **2200.902** | -0.398 | -0.120 | **0.759** | 1.07E-03 | **2.83E-03** |
| ENSG00000268836.1 | **ENSG00000268836** | | lncRNA | **19.648** | 1.128 | 0.339 | **2.185** | 1.07E-03 | **2.83E-03** |
| ENSG00000070159.16 | **PTPN3** | 5774 | protein_coding | **21.821** | 1.567 | 0.472 | **2.963** | 1.07E-03 | **2.83E-03** |
| ENSG00000006025.15 | **OSBPL7** | 114881 | protein_coding | **718.776** | 0.467 | 0.141 | **1.382** | 1.07E-03 | **2.83E-03** |
| ENSG00000155229.23 | **MMS19** | 64210 | protein_coding | **1615.263** | -0.407 | -0.123 | **0.754** | 1.07E-03 | **2.83E-03** |
| ENSG00000286753.4 | **ENSG0000(** | 10192920( | lncRNA | **42.547** | 0.844 | 0.254 | **1.795** | 1.08E-03 | **2.84E-03** |
| ENSG00000187808.5 | **SOWAHD** | 347454 | protein_coding | **102.308** | -0.783 | -0.236 | **0.581** | 1.08E-03 | **2.84E-03** |
| ENSG00000148926.12 | **ADM** | 133 | protein_coding | **528.486** | -0.472 | -0.142 | **0.721** | 1.08E-03 | **2.84E-03** |
| ENSG00000226859.1 | **LINC02051** | | lncRNA | **8.607** | -2.004 | -0.603 | **0.249** | 1.08E-03 | **2.85E-03** |
| ENSG00000066739.14 | **ATG2B** | 55102 | protein_coding | **608.177** | 0.648 | 0.195 | **1.567** | 1.08E-03 | **2.85E-03** |
| ENSG00000108010.14 | **GLRX3** | 10539 | protein_coding | **1008.652** | -0.322 | -0.097 | **0.800** | 1.08E-03 | **2.86E-03** |
| ENSG00000175792.14 | **RUVBL1** | 8607 | protein_coding | **785.286** | -0.456 | -0.137 | **0.729** | 1.09E-03 | **2.86E-03** |

| | | | | | | | | | |
|---|---|---|---|---|---|---|---|---|---|
| ENSG00000211679.2 | **IGLC3** | | IG_C_gene | **9723.926** | -0.708 | -0.213 | **0.612** | 1.09E-03 | **2.87E-03** |
| ENSG00000119636.18 | **BBOF1** | 80127 | protein_coding | **15.041** | 1.315 | 0.396 | **2.487** | 1.09E-03 | **2.88E-03** |
| ENSG00000164258.14 | **NDUFS4** | 4724 | protein_coding | **529.007** | -0.415 | -0.125 | **0.750** | 1.09E-03 | **2.88E-03** |
| ENSG00000172243.20 | **CLEC7A** | 64581 | protein_coding | **18.263** | 2.344 | 0.706 | **5.077** | 1.09E-03 | **2.88E-03** |
| ENSG00000005471.21 | **ABCB4** | 5244 | protein_coding | **180.286** | -0.972 | -0.293 | **0.510** | 1.10E-03 | **2.89E-03** |
| ENSG00000009413.18 | **REV3L** | 5980 | protein_coding | **295.456** | -0.482 | -0.145 | **0.716** | 1.10E-03 | **2.89E-03** |
| ENSG00000003393.18 | **ALS2** | 57679 | protein_coding | **307.874** | 0.482 | 0.145 | **1.397** | 1.10E-03 | **2.89E-03** |
| ENSG00000258325.3 | **ITFG2-AS1** | 283440 | lncRNA | **33.760** | 0.834 | 0.251 | **1.783** | 1.10E-03 | **2.89E-03** |
| ENSG00000290720.1 | **ZNF542P** | 147947 | lncRNA | **108.081** | 0.548 | 0.165 | **1.462** | 1.10E-03 | **2.90E-03** |
| ENSG00000260806.2 | **PAPOLA-DT** | | lncRNA | **60.390** | 0.928 | 0.279 | **1.902** | 1.10E-03 | **2.91E-03** |
| ENSG00000161860.10 | **SYCE2** | 256126 | protein_coding | **29.062** | -1.045 | -0.315 | **0.485** | 1.10E-03 | **2.91E-03** |
| ENSG00000188647.15 | **PTAR1** | 375743 | protein_coding | **206.177** | -0.597 | -0.180 | **0.661** | 1.11E-03 | **2.92E-03** |
| ENSG00000223496.3 | **EXOSC6** | 118460 | protein_coding | **471.496** | -0.493 | -0.148 | **0.711** | 1.11E-03 | **2.92E-03** |
| ENSG00000196295.14 | **GARS1-DT** | | lncRNA | **1173.301** | 0.384 | 0.115 | **1.305** | 1.11E-03 | **2.92E-03** |
| ENSG00000228606.5 | **DCAF8-DT** | 10028704[illegible] | lncRNA | **11.266** | -1.463 | -0.440 | **0.363** | 1.11E-03 | **2.92E-03** |
| ENSG00000269713.11 | **NBPF9** | 400818 | protein_coding | **1120.599** | 0.392 | 0.118 | **1.313** | 1.11E-03 | **2.92E-03** |
| ENSG00000255036.6 | **SUGT1P4-S** | 57653 | lncRNA | **19.705** | 2.060 | 0.620 | **4.171** | 1.11E-03 | **2.92E-03** |
| ENSG00000269982.1 | **ENSG00000269982** | | lncRNA | **15.655** | 1.167 | 0.351 | **2.245** | 1.11E-03 | **2.93E-03** |
| ENSG00000178896.11 | **EXOSC4** | 54512 | protein_coding | **260.054** | -0.449 | -0.135 | **0.733** | 1.11E-03 | **2.93E-03** |
| ENSG00000149196.19 | **HIKESHI** | 51501 | protein_coding | **549.879** | -0.325 | -0.098 | **0.798** | 1.11E-03 | **2.93E-03** |
| ENSG00000129355.8 | **CDKN2D** | 1032 | protein_coding | **212.218** | 0.612 | 0.184 | **1.529** | 1.11E-03 | **2.93E-03** |
| ENSG00000108509.23 | **CAMTA2** | 23125 | protein_coding | **1530.836** | 0.387 | 0.117 | **1.308** | 1.12E-03 | **2.93E-03** |
| ENSG00000223638.6 | **RFPL4A** | 342931 | protein_coding | **10.049** | 2.085 | 0.627 | **4.241** | 1.12E-03 | **2.94E-03** |
| ENSG00000256525.10 | **POLG2** | 11232 | protein_coding | **398.423** | 0.420 | 0.126 | **1.338** | 1.12E-03 | **2.94E-03** |
| ENSG00000100991.14 | **TRPC4AP** | 26133 | protein_coding | **1567.988** | -0.305 | -0.092 | **0.809** | 1.12E-03 | **2.94E-03** |
| ENSG00000115009.14 | **CCL20** | 6364 | protein_coding | **12.492** | 1.505 | 0.453 | **2.838** | 1.12E-03 | **2.94E-03** |
| ENSG00000243276.7 | **ENSG0000[illegible]** | 10537405[illegible] | lncRNA | **120.150** | 0.745 | 0.224 | **1.676** | 1.12E-03 | **2.94E-03** |
| ENSG00000174804.5 | **FZD4** | 8322 | protein_coding | **9.174** | 2.300 | 0.692 | **4.923** | 1.12E-03 | **2.94E-03** |
| ENSG00000153234.18 | **NR4A2** | 4929 | protein_coding | **57.082** | 0.625 | 0.188 | **1.542** | 1.12E-03 | **2.94E-03** |
| ENSG00000291091.2 | **ENSG00000291091** | | lncRNA | **475.953** | 0.323 | 0.097 | **1.251** | 1.12E-03 | **2.94E-03** |
| ENSG00000178127.15 | **NDUFV2** | 4729 | protein_coding | **3847.372** | -0.348 | -0.105 | **0.786** | 1.12E-03 | **2.94E-03** |

| | | | | | | | | | |
|---|---|---|---|---|---|---|---|---|---|
| ENSG00000110583.15 | **NAA40** | 79829 | protein_coding | **732.615** | 0.343 | 0.103 | **1.268** | 1.12E-03 | **2.95E-03** |
| ENSG00000211653.2 | **IGLV1-40** | | IG_V_gene | **1144.297** | -0.716 | -0.215 | **0.609** | 1.13E-03 | **2.95E-03** |
| ENSG00000130377.15 | **ACSBG2** | 81616 | protein_coding | **7.800** | 1.984 | 0.597 | **3.956** | 1.13E-03 | **2.95E-03** |
| ENSG00000163322.16 | **ABRAXAS1** | 84142 | protein_coding | **126.037** | 0.547 | 0.165 | **1.461** | 1.13E-03 | **2.95E-03** |
| ENSG00000068878.17 | **PSME4** | 23198 | protein_coding | **1379.377** | 0.416 | 0.125 | **1.335** | 1.13E-03 | **2.95E-03** |
| ENSG00000100596.10 | **SPTLC2** | 9517 | protein_coding | **681.219** | -0.434 | -0.131 | **0.740** | 1.13E-03 | **2.95E-03** |
| ENSG00000287910.2 | **ENSG00000287910** | | lncRNA | **6.753** | 2.246 | 0.676 | **4.743** | 1.13E-03 | **2.96E-03** |
| ENSG00000188229.8 | **TUBB4B** | 10383 | protein_coding | **4490.336** | -0.692 | -0.208 | **0.619** | 1.13E-03 | **2.96E-03** |
| ENSG00000283415.3 | **ENSG00000283415** | | lncRNA | **6.572** | -3.140 | -0.945 | **0.113** | 1.13E-03 | **2.96E-03** |
| ENSG00000141026.7 | **MED9** | 55090 | protein_coding | **226.367** | -0.439 | -0.132 | **0.738** | 1.13E-03 | **2.96E-03** |
| ENSG00000154447.17 | **SH3RF1** | 57630 | protein_coding | **175.056** | -0.548 | -0.165 | **0.684** | 1.13E-03 | **2.96E-03** |
| ENSG00000302941.1 | **ENSG00000302941** | | lncRNA | **6.546** | 2.231 | 0.672 | **4.694** | 1.14E-03 | **2.97E-03** |
| ENSG00000124688.16 | **MAD2L1BP** | 9587 | protein_coding | **323.500** | -0.373 | -0.112 | **0.772** | 1.14E-03 | **2.97E-03** |
| ENSG00000259439.4 | **LINC01833** | 10798587 | lncRNA | **48.031** | 1.299 | 0.391 | **2.460** | 1.14E-03 | **2.98E-03** |
| ENSG00000303180.1 | **ENSG00000303180** | | lncRNA | **88.317** | 0.789 | 0.237 | **1.727** | 1.14E-03 | **2.98E-03** |
| ENSG00000159625.17 | **DRC7** | 84229 | protein_coding | **8.604** | 2.157 | 0.649 | **4.461** | 1.14E-03 | **2.98E-03** |
| ENSG00000114648.14 | **KLHL18** | 23276 | protein_coding | **489.255** | -0.428 | -0.129 | **0.743** | 1.14E-03 | **2.99E-03** |
| ENSG00000236444.6 | **UBE2L5** | 171222 | protein_coding | **15.097** | 1.500 | 0.451 | **2.828** | 1.14E-03 | **2.99E-03** |
| ENSG00000104886.14 | **PLEKHJ1** | 55111 | protein_coding | **962.702** | -0.281 | -0.084 | **0.823** | 1.15E-03 | **3.00E-03** |
| ENSG00000178233.19 | **TMEM151** | 441151 | protein_coding | **7.260** | 1.709 | 0.514 | **3.269** | 1.15E-03 | **3.00E-03** |
| ENSG00000187147.21 | **RNF220** | 55182 | protein_coding | **1036.737** | -0.311 | -0.094 | **0.806** | 1.15E-03 | **3.01E-03** |
| ENSG00000138814.19 | **PPP3CA** | 5530 | protein_coding | **236.995** | 0.613 | 0.184 | **1.529** | 1.15E-03 | **3.01E-03** |
| ENSG00000245317.3 | **MRNIP-DT** | 10099641 | lncRNA | **31.921** | 0.985 | 0.297 | **1.980** | 1.15E-03 | **3.01E-03** |
| ENSG00000120798.19 | **NR2C1** | 7181 | protein_coding | **414.875** | 0.529 | 0.159 | **1.443** | 1.15E-03 | **3.01E-03** |
| ENSG00000105085.12 | **MED26** | 9441 | protein_coding | **620.974** | 0.304 | 0.092 | **1.235** | 1.16E-03 | **3.02E-03** |
| ENSG00000237436.2 | **CAMTA1-DT** | | lncRNA | **8.283** | 1.549 | 0.466 | **2.927** | 1.16E-03 | **3.02E-03** |
| ENSG00000204020.7 | **LIPN** | 643418 | protein_coding | **7.019** | 2.348 | 0.707 | **5.090** | 1.16E-03 | **3.02E-03** |
| ENSG00000147601.17 | **TERF1** | 7013 | protein_coding | **201.275** | -0.525 | -0.158 | **0.695** | 1.16E-03 | **3.02E-03** |
| ENSG00000160007.22 | **ARHGAP35** | 2909 | protein_coding | **611.505** | -0.395 | -0.119 | **0.761** | 1.16E-03 | **3.03E-03** |
| ENSG00000172057.12 | **ORMDL3** | 94103 | protein_coding | **1908.001** | 0.461 | 0.139 | **1.377** | 1.16E-03 | **3.04E-03** |
| ENSG00000249471.10 | **ZNF324B** | 388569 | protein_coding | **87.569** | -0.556 | -0.167 | **0.680** | 1.16E-03 | **3.04E-03** |

| | | | | | | | | | |
|---|---|---|---|---|---|---|---|---|---|
| ENSG00000289073.2 | **ENSG00000289073** | | lncRNA | **9.595** | 1.477 | 0.445 | **2.784** | 1.17E-03 | **3.05E-03** |
| ENSG00000197818.14 | **SLC9A8** | 23315 | protein_coding | **663.653** | 0.487 | 0.147 | **1.402** | 1.17E-03 | **3.05E-03** |
| ENSG00000154305.20 | **MIA3** | 375056 | protein_coding | **1062.322** | 0.428 | 0.129 | **1.346** | 1.17E-03 | **3.05E-03** |
| ENSG00000261115.8 | **TMEM178I** | 100507421 | protein_coding | **23.489** | 1.434 | 0.432 | **2.702** | 1.17E-03 | **3.05E-03** |
| ENSG00000054611.16 | **TBC1D22A** | 25771 | protein_coding | **994.259** | -0.377 | -0.113 | **0.770** | 1.17E-03 | **3.05E-03** |
| ENSG00000140332.20 | **TLE3** | 7090 | protein_coding | **1086.986** | -0.388 | -0.117 | **0.764** | 1.17E-03 | **3.05E-03** |
| ENSG00000295196.1 | **ENSG00000295196** | | lncRNA | **51.492** | -0.882 | -0.266 | **0.543** | 1.17E-03 | **3.05E-03** |
| ENSG00000225793.6 | **TMEM30A-DT** | | lncRNA | **43.622** | 0.761 | 0.229 | **1.695** | 1.17E-03 | **3.06E-03** |
| ENSG00000300184.1 | **ENSG00000300184** | | lncRNA | **7.124** | -2.251 | -0.678 | **0.210** | 1.17E-03 | **3.06E-03** |
| ENSG00000204220.14 | **PFDN6** | 10471 | protein_coding | **2011.122** | 0.443 | 0.133 | **1.360** | 1.17E-03 | **3.06E-03** |
| ENSG00000257526.4 | **ENSG00000257526** | | lncRNA | **13.714** | 1.638 | 0.493 | **3.113** | 1.17E-03 | **3.06E-03** |
| ENSG00000128917.10 | **DLL4** | 54567 | protein_coding | **29.157** | 1.392 | 0.419 | **2.624** | 1.18E-03 | **3.07E-03** |
| ENSG00000258289.12 | **CHURC1** | 91612 | protein_coding | **928.992** | -0.369 | -0.111 | **0.774** | 1.18E-03 | **3.07E-03** |
| ENSG00000172315.7 | **TP53RK** | 112858 | protein_coding | **286.543** | -0.387 | -0.116 | **0.765** | 1.18E-03 | **3.07E-03** |
| ENSG00000266340.1 | **ENSG00000266340** | | lncRNA | **59.124** | 0.943 | 0.284 | **1.922** | 1.18E-03 | **3.08E-03** |
| ENSG00000123636.22 | **BAZ2B** | 29994 | protein_coding | **193.928** | -0.706 | -0.212 | **0.613** | 1.18E-03 | **3.08E-03** |
| ENSG00000236519.3 | **LINC01424** | | lncRNA | **46.616** | 0.745 | 0.224 | **1.676** | 1.18E-03 | **3.09E-03** |
| ENSG00000133703.16 | **KRAS** | 3845 | protein_coding | **492.021** | 0.541 | 0.163 | **1.455** | 1.19E-03 | **3.10E-03** |
| ENSG00000291139.2 | **DPY19L2P2** | 349152 | lncRNA | **133.428** | -0.718 | -0.216 | **0.608** | 1.19E-03 | **3.10E-03** |
| ENSG00000300661.1 | **ENSG00000300661** | | lncRNA | **14.196** | 1.175 | 0.354 | **2.259** | 1.19E-03 | **3.10E-03** |
| ENSG00000125741.7 | **OPA3** | 80207 | protein_coding | **404.550** | -0.352 | -0.106 | **0.784** | 1.19E-03 | **3.10E-03** |
| ENSG00000203880.15 | **PCMTD2** | 55251 | protein_coding | **450.506** | 0.659 | 0.198 | **1.579** | 1.19E-03 | **3.10E-03** |
| ENSG00000276966.3 | **H4C5** | 8367 | protein_coding | **9.143** | -1.902 | -0.573 | **0.268** | 1.19E-03 | **3.10E-03** |
| ENSG00000143164.18 | **DCAF6** | 55827 | protein_coding | **1106.589** | 0.347 | 0.104 | **1.271** | 1.19E-03 | **3.10E-03** |
| ENSG00000168175.18 | **MAPK1IP1** | 93487 | protein_coding | **1753.513** | -0.429 | -0.129 | **0.743** | 1.20E-03 | **3.11E-03** |
| ENSG00000166669.16 | **ATF7IP2** | 80063 | protein_coding | **387.483** | 0.519 | 0.156 | **1.433** | 1.20E-03 | **3.11E-03** |
| ENSG00000100997.22 | **ABHD12** | 26090 | protein_coding | **861.407** | -0.413 | -0.124 | **0.751** | 1.20E-03 | **3.12E-03** |
| ENSG00000229619.6 | **MBNL1-AS** | 401093 | lncRNA | **24.923** | 1.128 | 0.340 | **2.186** | 1.20E-03 | **3.12E-03** |
| ENSG00000220008.6 | **LINGO3** | 645191 | protein_coding | **16.552** | 1.263 | 0.380 | **2.400** | 1.20E-03 | **3.12E-03** |
| ENSG00000005436.16 | **GCFC2** | 6936 | protein_coding | **226.076** | -0.587 | -0.177 | **0.666** | 1.20E-03 | **3.13E-03** |
| ENSG00000181610.15 | **MRPS23** | 51649 | protein_coding | **643.653** | -0.445 | -0.134 | **0.734** | 1.21E-03 | **3.14E-03** |

| | | | | | | | | | |
|---|---|---|---|---|---|---|---|---|---|
| ENSG00000132010.17 | **ZNF20** | 7568 | protein_coding | **48.456** | 1.053 | 0.317 | **2.074** | 1.21E-03 | **3.14E-03** |
| ENSG00000247134.9 | **ENSG00000247134** | | lncRNA | **181.421** | -0.436 | -0.131 | **0.739** | 1.21E-03 | **3.14E-03** |
| ENSG00000159069.16 | **FBXW5** | 54461 | protein_coding | **1490.654** | -0.281 | -0.085 | **0.823** | 1.21E-03 | **3.14E-03** |
| ENSG00000171812.14 | **COL8A2** | 1296 | protein_coding | **4.698** | 2.780 | 0.837 | **6.867** | 1.21E-03 | **3.15E-03** |
| ENSG00000105974.15 | **CAV1** | 857 | protein_coding | **137.305** | 0.751 | 0.226 | **1.683** | 1.21E-03 | **3.15E-03** |
| ENSG00000146950.14 | **SHROOM2** | 357 | protein_coding | **8.938** | 2.026 | 0.610 | **4.074** | 1.21E-03 | **3.15E-03** |
| ENSG00000225756.1 | **DBH-AS1** | 138948 | lncRNA | **9.224** | 2.206 | 0.664 | **4.614** | 1.21E-03 | **3.15E-03** |
| ENSG00000093072.21 | **ADA2** | 51816 | protein_coding | **999.933** | 0.480 | 0.144 | **1.395** | 1.21E-03 | **3.15E-03** |
| ENSG00000106799.17 | **TGFBR1** | 7046 | protein_coding | **160.404** | -0.473 | -0.143 | **0.720** | 1.21E-03 | **3.16E-03** |
| ENSG00000047056.20 | **WDR37** | 22884 | protein_coding | **453.486** | 0.404 | 0.122 | **1.323** | 1.22E-03 | **3.16E-03** |
| ENSG00000262165.2 | **C17orf114** | 11913990! | protein_coding | **7.428** | 2.147 | 0.646 | **4.428** | 1.22E-03 | **3.16E-03** |
| ENSG00000286241.2 | **ENSG00000286241** | | lncRNA | **7.751** | 2.054 | 0.618 | **4.152** | 1.22E-03 | **3.16E-03** |
| ENSG00000205269.8 | **TMEM170I** | 10011340 | protein_coding | **92.272** | 0.570 | 0.172 | **1.485** | 1.22E-03 | **3.17E-03** |
| ENSG00000178028.16 | **DMAP1** | 55929 | protein_coding | **818.955** | 0.319 | 0.096 | **1.247** | 1.22E-03 | **3.17E-03** |
| ENSG00000103494.18 | **RPGRIP1L** | 23322 | protein_coding | **74.087** | -0.900 | -0.271 | **0.536** | 1.22E-03 | **3.17E-03** |
| ENSG00000182173.16 | **TSEN54** | 283989 | protein_coding | **457.024** | 0.440 | 0.132 | **1.356** | 1.22E-03 | **3.17E-03** |
| ENSG00000189164.17 | **ZNF527** | 84503 | protein_coding | **64.281** | -0.784 | -0.236 | **0.581** | 1.23E-03 | **3.18E-03** |
| ENSG00000302714.1 | **ENSG00000302714** | | lncRNA | **6.423** | 2.261 | 0.681 | **4.793** | 1.23E-03 | **3.19E-03** |
| ENSG00000175356.16 | **SCUBE2** | 57758 | protein_coding | **12.928** | 2.310 | 0.695 | **4.958** | 1.23E-03 | **3.19E-03** |
| ENSG00000074966.14 | **TXK** | 7294 | protein_coding | **181.681** | 0.620 | 0.187 | **1.537** | 1.23E-03 | **3.20E-03** |
| ENSG00000099942.15 | **CRKL** | 1399 | protein_coding | **1414.317** | -0.363 | -0.109 | **0.777** | 1.24E-03 | **3.20E-03** |
| ENSG00000229536.4 | **LINC02572** | 10537361! | lncRNA | **6.105** | 1.999 | 0.602 | **3.998** | 1.24E-03 | **3.21E-03** |
| ENSG00000146267.13 | **FAXC** | 84553 | protein_coding | **7.658** | 3.382 | 1.018 | **10.427** | 1.24E-03 | **3.21E-03** |
| ENSG00000072832.16 | **CRMP1** | 1400 | protein_coding | **8.252** | 2.305 | 0.694 | **4.942** | 1.24E-03 | **3.23E-03** |
| ENSG00000236915.3 | **CLCA4-AS1** | 10537882 | lncRNA | **19.141** | 1.287 | 0.388 | **2.441** | 1.25E-03 | **3.23E-03** |
| ENSG00000152284.7 | **TCF7L1** | 83439 | protein_coding | **6.538** | 1.959 | 0.590 | **3.888** | 1.25E-03 | **3.24E-03** |
| ENSG00000159579.16 | **RSPRY1** | 89970 | protein_coding | **517.131** | 0.499 | 0.150 | **1.413** | 1.26E-03 | **3.25E-03** |
| ENSG00000235488.1 | **JARID2-AS1** | | lncRNA | **6.701** | 1.872 | 0.563 | **3.660** | 1.26E-03 | **3.25E-03** |
| ENSG00000236514.1 | **ZNF248-AS1** | | lncRNA | **8.957** | 1.771 | 0.533 | **3.413** | 1.26E-03 | **3.26E-03** |
| ENSG00000264672.7 | **SEPTIN4-AS1** | | lncRNA | **18.384** | 1.157 | 0.348 | **2.231** | 1.26E-03 | **3.26E-03** |
| ENSG00000163518.11 | **FCRL4** | 83417 | protein_coding | **106.148** | -1.097 | -0.330 | **0.467** | 1.26E-03 | **3.26E-03** |

| | | | | | | | | | |
|---|---|---|---|---|---|---|---|---|---|
| ENSG00000101596.19 | **SMCHD1** | 23347 | protein_coding | **936.521** | -0.454 | -0.137 | **0.730** | 1.26E-03 | **3.26E-03** |
| ENSG00000121743.6 | **GJA3** | 2700 | protein_coding | **42.500** | 1.024 | 0.308 | **2.033** | 1.26E-03 | **3.27E-03** |
| ENSG00000291203.2 | **ENSG00000291203** | | lncRNA | **250.394** | -0.465 | -0.140 | **0.724** | 1.26E-03 | **3.27E-03** |
| ENSG00000267783.4 | **ENSG00000267783** | | lncRNA | **8.568** | 1.787 | 0.538 | **3.451** | 1.26E-03 | **3.27E-03** |
| ENSG00000261652.4 | **PIERCE2** | 145788 | protein_coding | **20.698** | 1.190 | 0.358 | **2.282** | 1.27E-03 | **3.28E-03** |
| ENSG00000065491.10 | **TBC1D22B** | 55633 | protein_coding | **502.525** | 0.313 | 0.094 | **1.242** | 1.27E-03 | **3.29E-03** |
| ENSG00000129083.15 | **COPB1** | 1315 | protein_coding | **1607.108** | -0.273 | -0.082 | **0.828** | 1.27E-03 | **3.29E-03** |
| ENSG00000130561.17 | **SAG** | 6295 | protein_coding | **5.907** | 2.728 | 0.821 | **6.625** | 1.27E-03 | **3.29E-03** |
| ENSG00000232098.4 | **ZNF584-DT** | | lncRNA | **144.052** | 0.600 | 0.181 | **1.516** | 1.27E-03 | **3.30E-03** |
| ENSG00000284526.4 | **ENSG0000(** | 12245534: | protein_coding | **9.402** | 2.433 | 0.732 | **5.400** | 1.28E-03 | **3.30E-03** |
| ENSG00000113552.18 | **GNPDA1** | 10007 | protein_coding | **473.768** | -0.363 | -0.109 | **0.777** | 1.28E-03 | **3.30E-03** |
| ENSG00000107949.19 | **BCCIP** | 56647 | protein_coding | **945.651** | -0.314 | -0.095 | **0.804** | 1.28E-03 | **3.31E-03** |
| ENSG00000285920.2 | **ENSG00000285920** | | protein_coding | **69.177** | -5.504 | -1.657 | **0.022** | 1.28E-03 | **3.31E-03** |
| ENSG00000254473.3 | **UBQLN1-AS1** | | lncRNA | **51.274** | 0.760 | 0.229 | **1.694** | 1.28E-03 | **3.31E-03** |
| ENSG00000226950.8 | **DANCR** | 57291 | lncRNA | **1261.970** | -0.405 | -0.122 | **0.755** | 1.29E-03 | **3.33E-03** |
| ENSG00000280417.1 | **ENSG00000280417** | | TEC | **7.589** | 1.601 | 0.482 | **3.033** | 1.29E-03 | **3.33E-03** |
| ENSG00000145029.17 | **NICN1** | 84276 | protein_coding | **278.822** | 0.453 | 0.136 | **1.369** | 1.29E-03 | **3.34E-03** |
| ENSG00000108379.13 | **WNT3** | 7473 | protein_coding | **8.845** | 1.791 | 0.539 | **3.460** | 1.29E-03 | **3.34E-03** |
| ENSG00000246067.11 | **RAB30-DT** | | lncRNA | **232.177** | 0.457 | 0.138 | **1.373** | 1.29E-03 | **3.34E-03** |
| ENSG00000162881.7 | **OXER1** | 165140 | protein_coding | **313.414** | -0.664 | -0.200 | **0.631** | 1.29E-03 | **3.34E-03** |
| ENSG00000139926.18 | **FRMD6** | 122786 | protein_coding | **63.792** | 0.859 | 0.259 | **1.814** | 1.30E-03 | **3.34E-03** |
| ENSG00000181827.18 | **RFX7** | 64864 | protein_coding | **220.557** | -0.557 | -0.168 | **0.680** | 1.30E-03 | **3.35E-03** |
| ENSG00000215114.12 | **UBXN2B** | 137886 | protein_coding | **166.944** | -0.528 | -0.159 | **0.693** | 1.30E-03 | **3.35E-03** |
| ENSG00000300245.1 | **ENSG00000300245** | | lncRNA | **9.554** | 1.494 | 0.450 | **2.817** | 1.30E-03 | **3.35E-03** |
| ENSG00000112039.8 | **FANCE** | 2178 | protein_coding | **130.493** | -0.423 | -0.127 | **0.746** | 1.30E-03 | **3.36E-03** |
| ENSG00000144744.20 | **UBA3** | 9039 | protein_coding | **938.779** | 0.478 | 0.144 | **1.393** | 1.30E-03 | **3.36E-03** |
| ENSG00000137100.18 | **DCTN3** | 11258 | protein_coding | **668.187** | -0.264 | -0.079 | **0.833** | 1.30E-03 | **3.36E-03** |
| ENSG00000127328.24 | **RAB3IP** | 117177 | protein_coding | **321.477** | -0.448 | -0.135 | **0.733** | 1.31E-03 | **3.37E-03** |
| ENSG00000182472.9 | **CAPN12** | 147968 | protein_coding | **478.088** | 0.584 | 0.176 | **1.499** | 1.31E-03 | **3.37E-03** |
| ENSG00000240038.8 | **AMY2B** | 280 | protein_coding | **850.816** | 0.473 | 0.142 | **1.388** | 1.31E-03 | **3.38E-03** |
| ENSG00000148450.15 | **MSRB2** | 22921 | protein_coding | **248.439** | -0.439 | -0.132 | **0.738** | 1.31E-03 | **3.38E-03** |

| | | | | | | | | | |
|---|---|---|---|---|---|---|---|---|---|
| ENSG00000168818.12 | **STX18** | 53407 | protein_coding | **665.762** | -0.398 | -0.120 | **0.759** | 1.31E-03 | **3.39E-03** |
| ENSG00000236810.9 | **ELOA-AS1** | 10050696: | lncRNA | **194.406** | 0.519 | 0.156 | **1.432** | 1.31E-03 | **3.39E-03** |
| ENSG00000166788.12 | **SAAL1** | 113174 | protein_coding | **352.208** | -0.474 | -0.143 | **0.720** | 1.31E-03 | **3.39E-03** |
| ENSG00000224470.11 | **ATXN1L** | 342371 | protein_coding | **712.304** | 0.424 | 0.127 | **1.341** | 1.32E-03 | **3.39E-03** |
| ENSG00000259768.9 | **ZFHX3-AS1** | | lncRNA | **236.730** | 0.447 | 0.135 | **1.363** | 1.32E-03 | **3.40E-03** |
| ENSG00000063587.16 | **ZNF275** | 10838 | protein_coding | **766.932** | -0.406 | -0.122 | **0.755** | 1.32E-03 | **3.40E-03** |
| ENSG00000183971.11 | **NPW** | 283869 | protein_coding | **13.878** | 1.369 | 0.412 | **2.583** | 1.32E-03 | **3.40E-03** |
| ENSG00000179314.19 | **WSCD1** | 23302 | protein_coding | **7.444** | 2.447 | 0.737 | **5.454** | 1.32E-03 | **3.40E-03** |
| ENSG00000125686.14 | **MED1** | 5469 | protein_coding | **553.618** | -0.385 | -0.116 | **0.766** | 1.32E-03 | **3.41E-03** |
| ENSG00000272384.3 | **ZHX2-AS1** | | lncRNA | **14.918** | 1.231 | 0.370 | **2.347** | 1.32E-03 | **3.41E-03** |
| ENSG00000104763.22 | **ASAH1** | 427 | protein_coding | **1496.418** | -0.311 | -0.094 | **0.806** | 1.33E-03 | **3.42E-03** |
| ENSG00000274512.8 | **TBC1D3L** | 10106037( | protein_coding | **1742.473** | 0.419 | 0.126 | **1.337** | 1.33E-03 | **3.42E-03** |
| ENSG00000289222.3 | **ENSG00000289222** | | lncRNA | **14.433** | 1.255 | 0.378 | **2.386** | 1.33E-03 | **3.42E-03** |
| ENSG00000167470.16 | **MIDN** | 90007 | protein_coding | **1008.907** | -0.359 | -0.108 | **0.780** | 1.33E-03 | **3.42E-03** |
| ENSG00000109436.10 | **TBC1D9** | 23158 | protein_coding | **62.277** | 0.966 | 0.291 | **1.954** | 1.33E-03 | **3.43E-03** |
| ENSG00000143799.16 | **PARP1** | 142 | protein_coding | **4752.086** | -0.379 | -0.114 | **0.769** | 1.34E-03 | **3.44E-03** |
| ENSG00000058085.17 | **LAMC2** | 3918 | protein_coding | **11.842** | 2.089 | 0.629 | **4.255** | 1.34E-03 | **3.44E-03** |
| ENSG00000241106.10 | **HLA-DOB** | 3112 | protein_coding | **698.397** | 0.321 | 0.097 | **1.249** | 1.34E-03 | **3.44E-03** |
| ENSG00000205090.10 | **TMEM240** | 339453 | protein_coding | **4.856** | 2.762 | 0.831 | **6.781** | 1.34E-03 | **3.45E-03** |
| ENSG00000197889.12 | **MEIG1** | 644890 | protein_coding | **8.251** | 1.780 | 0.536 | **3.434** | 1.34E-03 | **3.45E-03** |
| ENSG00000144935.18 | **TRPC1** | 7220 | protein_coding | **39.097** | 0.888 | 0.267 | **1.851** | 1.34E-03 | **3.45E-03** |
| ENSG00000275481.1 | **ENSG00000275481** | | lncRNA | **17.059** | 1.126 | 0.339 | **2.183** | 1.35E-03 | **3.46E-03** |
| ENSG00000158604.18 | **TMED4** | 222068 | protein_coding | **841.858** | -0.317 | -0.095 | **0.803** | 1.35E-03 | **3.48E-03** |
| ENSG00000204381.14 | **LAYN** | 143903 | protein_coding | **9.715** | 2.230 | 0.671 | **4.690** | 1.36E-03 | **3.49E-03** |
| ENSG00000263528.10 | **IKBKE** | 9641 | protein_coding | **670.353** | 0.340 | 0.102 | **1.266** | 1.36E-03 | **3.49E-03** |
| ENSG00000136161.15 | **RCBTB2** | 1102 | protein_coding | **818.914** | 0.443 | 0.133 | **1.359** | 1.36E-03 | **3.49E-03** |
| ENSG00000123395.16 | **ATG101** | 60673 | protein_coding | **423.630** | 0.410 | 0.123 | **1.329** | 1.36E-03 | **3.50E-03** |
| ENSG00000168070.14 | **MAJIN** | 283129 | protein_coding | **4.427** | 2.896 | 0.872 | **7.444** | 1.37E-03 | **3.52E-03** |
| ENSG00000118496.7 | **FBXO30** | 84085 | protein_coding | **154.624** | -0.530 | -0.160 | **0.692** | 1.37E-03 | **3.52E-03** |
| ENSG00000288880.3 | **ENSG0000(** | 11226823( | lncRNA | **47.408** | 0.777 | 0.234 | **1.714** | 1.38E-03 | **3.53E-03** |
| ENSG00000173681.19 | **BCLAF3** | 256643 | protein_coding | **236.235** | 0.521 | 0.157 | **1.435** | 1.38E-03 | **3.53E-03** |

| | | | | | | | | | |
|---|---|---|---|---|---|---|---|---|---|
| ENSG00000140153.20 | **WDR20** | 91833 | protein_coding | **261.010** | -0.444 | -0.134 | **0.735** | 1.38E-03 | **3.53E-03** |
| ENSG00000294588.1 | **ENSG00000294588** | | lncRNA | **5.899** | 2.129 | 0.641 | **4.375** | 1.38E-03 | **3.54E-03** |
| ENSG00000121577.16 | **POPDC2** | 64091 | protein_coding | **185.701** | 0.555 | 0.167 | **1.469** | 1.39E-03 | **3.56E-03** |
| ENSG00000182676.7 | **PPP1R27** | 116729 | protein_coding | **10.222** | -2.592 | -0.780 | **0.166** | 1.39E-03 | **3.56E-03** |
| ENSG00000124942.15 | **AHNAK** | 79026 | protein_coding | **2585.674** | 0.551 | 0.166 | **1.465** | 1.39E-03 | **3.57E-03** |
| ENSG00000110934.15 | **BIN2** | 51411 | protein_coding | **1342.541** | 0.465 | 0.140 | **1.380** | 1.40E-03 | **3.58E-03** |
| ENSG00000148842.20 | **CNNM2** | 54805 | protein_coding | **143.381** | 0.537 | 0.162 | **1.451** | 1.40E-03 | **3.58E-03** |
| ENSG00000119655.13 | **NPC2** | 10577 | protein_coding | **1602.288** | -0.395 | -0.119 | **0.761** | 1.40E-03 | **3.59E-03** |
| ENSG00000282936.4 | **ENSG0000(** | 12252678( | protein_coding | **9.833** | -2.402 | -0.723 | **0.189** | 1.40E-03 | **3.59E-03** |
| ENSG00000058866.17 | **DGKG** | 1608 | protein_coding | **68.295** | 0.916 | 0.276 | **1.887** | 1.41E-03 | **3.61E-03** |
| ENSG00000104863.14 | **LIN7B** | 64130 | protein_coding | **31.634** | 0.994 | 0.299 | **1.991** | 1.41E-03 | **3.61E-03** |
| ENSG00000269335.9 | **IKBKG** | 8517 | protein_coding | **458.296** | 0.679 | 0.204 | **1.601** | 1.41E-03 | **3.61E-03** |
| ENSG00000131142.16 | **CCL25** | 6370 | protein_coding | **97.318** | 1.014 | 0.305 | **2.020** | 1.41E-03 | **3.61E-03** |
| ENSG00000140948.14 | **ZCCHC14** | 23174 | protein_coding | **164.232** | 0.660 | 0.199 | **1.580** | 1.41E-03 | **3.62E-03** |
| ENSG00000261000.1 | **ENSG00000261000** | | lncRNA | **14.751** | 1.341 | 0.404 | **2.533** | 1.42E-03 | **3.63E-03** |
| ENSG00000198846.8 | **TOX** | 9760 | protein_coding | **58.600** | -0.868 | -0.261 | **0.548** | 1.42E-03 | **3.63E-03** |
| ENSG00000181789.16 | **COPG1** | 22820 | protein_coding | **2575.365** | -0.321 | -0.097 | **0.801** | 1.42E-03 | **3.63E-03** |
| ENSG00000119977.24 | **TCTN3** | 26123 | protein_coding | **555.318** | -0.306 | -0.092 | **0.809** | 1.42E-03 | **3.63E-03** |
| ENSG00000280153.1 | **ENSG00000280153** | | TEC | **124.534** | 2.328 | 0.701 | **5.022** | 1.42E-03 | **3.63E-03** |
| ENSG00000267496.5 | **FAM215A** | | lncRNA | **28.947** | 0.880 | 0.265 | **1.840** | 1.43E-03 | **3.65E-03** |
| ENSG00000267199.1 | **ENSG00000267199** | | lncRNA | **51.992** | -1.408 | -0.424 | **0.377** | 1.43E-03 | **3.65E-03** |
| ENSG00000261202.3 | **TNRC6B-DT** | | lncRNA | **9.529** | 1.640 | 0.494 | **3.117** | 1.43E-03 | **3.66E-03** |
| ENSG00000263335.1 | **ENSG00000263335** | | lncRNA | **27.148** | 1.209 | 0.364 | **2.312** | 1.43E-03 | **3.66E-03** |
| ENSG00000127837.12 | **AAMP** | 14 | protein_coding | **1420.739** | 0.429 | 0.129 | **1.346** | 1.43E-03 | **3.67E-03** |
| ENSG00000128059.10 | **PPAT** | 5471 | protein_coding | **345.127** | -0.418 | -0.126 | **0.749** | 1.43E-03 | **3.67E-03** |
| ENSG00000134138.24 | **MEIS2** | 4212 | protein_coding | **55.480** | -0.710 | -0.214 | **0.611** | 1.44E-03 | **3.68E-03** |
| ENSG00000144567.13 | **RETREG2** | 79137 | protein_coding | **800.169** | -0.314 | -0.095 | **0.804** | 1.44E-03 | **3.69E-03** |
| ENSG00000023608.7 | **SNAPC1** | 6617 | protein_coding | **172.384** | -0.559 | -0.168 | **0.679** | 1.44E-03 | **3.69E-03** |
| ENSG00000105552.17 | **BCAT2** | 587 | protein_coding | **1043.008** | -0.385 | -0.116 | **0.766** | 1.45E-03 | **3.70E-03** |
| ENSG00000260552.4 | **COSMOC** | 10192780( | lncRNA | **9.510** | 1.824 | 0.549 | **3.540** | 1.45E-03 | **3.70E-03** |
| ENSG00000226754.4 | **MAGOH-DT** | | lncRNA | **24.095** | 1.101 | 0.332 | **2.146** | 1.45E-03 | **3.70E-03** |

| | | | | | | | | | |
|---|---|---|---|---|---|---|---|---|---|
| ENSG00000185009.15 | **AP3M1** | 26985 | protein_coding | **623.435** | -0.447 | -0.135 | **0.733** | 1.45E-03 | **3.70E-03** |
| ENSG00000113073.17 | **SLC4A9** | 83697 | protein_coding | **7.299** | 2.376 | 0.715 | **5.192** | 1.45E-03 | **3.70E-03** |
| ENSG00000164972.15 | **SPMIP6** | 84688 | protein_coding | **6.269** | 2.772 | 0.835 | **6.833** | 1.45E-03 | **3.70E-03** |
| ENSG00000158825.8 | **CDA** | 978 | protein_coding | **14.764** | 2.106 | 0.634 | **4.305** | 1.46E-03 | **3.72E-03** |
| ENSG00000174013.9 | **FBXO45** | 200933 | protein_coding | **204.629** | -0.369 | -0.111 | **0.774** | 1.46E-03 | **3.72E-03** |
| ENSG00000182545.7 | **RNASE10** | 338879 | protein_coding | **8.945** | 1.734 | 0.522 | **3.327** | 1.46E-03 | **3.72E-03** |
| ENSG00000171867.20 | **PRNP** | 5621 | protein_coding | **135.112** | -0.745 | -0.224 | **0.597** | 1.46E-03 | **3.74E-03** |
| ENSG00000184206.13 | **GOLGA6L4** | 643707 | protein_coding | **13.008** | 1.425 | 0.429 | **2.685** | 1.46E-03 | **3.74E-03** |
| ENSG00000160679.14 | **CHTOP** | 26097 | protein_coding | **1312.049** | -0.300 | -0.090 | **0.812** | 1.47E-03 | **3.74E-03** |
| ENSG00000141971.16 | **MVB12A** | 93343 | protein_coding | **635.565** | 0.363 | 0.109 | **1.286** | 1.47E-03 | **3.75E-03** |
| ENSG00000206527.12 | **HACD2** | 201562 | protein_coding | **271.613** | -0.535 | -0.161 | **0.690** | 1.47E-03 | **3.75E-03** |
| ENSG00000129219.16 | **PLD2** | 5338 | protein_coding | **360.010** | 0.550 | 0.166 | **1.464** | 1.47E-03 | **3.75E-03** |
| ENSG00000185482.10 | **STAC3** | 246329 | protein_coding | **142.114** | -0.631 | -0.190 | **0.646** | 1.47E-03 | **3.76E-03** |
| ENSG00000073737.19 | **DHRS9** | 10170 | protein_coding | **17.656** | -1.399 | -0.421 | **0.379** | 1.48E-03 | **3.77E-03** |
| ENSG00000289303.3 | **ENSG00000289303** | | lncRNA | **11.078** | 1.591 | 0.479 | **3.013** | 1.48E-03 | **3.77E-03** |
| ENSG00000165895.20 | **ARHGAP42** | 143872 | protein_coding | **50.586** | 1.100 | 0.331 | **2.144** | 1.48E-03 | **3.77E-03** |
| ENSG00000006194.13 | **ZNF263** | 10127 | protein_coding | **655.189** | -0.299 | -0.090 | **0.813** | 1.48E-03 | **3.77E-03** |
| ENSG00000134375.13 | **TIMM17A** | 10440 | protein_coding | **495.110** | -0.507 | -0.153 | **0.703** | 1.48E-03 | **3.77E-03** |
| ENSG00000088812.20 | **ATRN** | 8455 | protein_coding | **204.180** | 0.601 | 0.181 | **1.517** | 1.48E-03 | **3.77E-03** |
| ENSG00000120029.15 | **ARMH3** | 79591 | protein_coding | **566.652** | -0.322 | -0.097 | **0.800** | 1.48E-03 | **3.78E-03** |
| ENSG00000130881.15 | **LRP3** | 4037 | protein_coding | **7.610** | 1.953 | 0.588 | **3.873** | 1.49E-03 | **3.79E-03** |
| ENSG00000270277.1 | **ENSG00000270277** | | lncRNA | **35.994** | 0.925 | 0.279 | **1.899** | 1.49E-03 | **3.79E-03** |
| ENSG00000310469.1 | **HLA-H** | 3136 | lncRNA | **975.742** | -0.735 | -0.221 | **0.601** | 1.49E-03 | **3.79E-03** |
| ENSG00000170653.21 | **ATF7** | 11016 | protein_coding | **511.026** | 0.359 | 0.108 | **1.283** | 1.49E-03 | **3.81E-03** |
| ENSG00000162413.18 | **KLHL21** | 9903 | protein_coding | **309.523** | -0.434 | -0.131 | **0.740** | 1.49E-03 | **3.81E-03** |
| ENSG00000213380.17 | **COG8** | 84342 | protein_coding | **402.809** | 0.495 | 0.149 | **1.409** | 1.50E-03 | **3.81E-03** |
| ENSG00000262202.5 | **ENSG00000262202** | | lncRNA | **84.688** | 0.883 | 0.266 | **1.844** | 1.50E-03 | **3.81E-03** |
| ENSG00000148943.13 | **LIN7C** | 55327 | protein_coding | **158.524** | -0.485 | -0.146 | **0.714** | 1.50E-03 | **3.82E-03** |
| ENSG00000144161.16 | **ZC3H8** | 84524 | protein_coding | **305.283** | -0.692 | -0.208 | **0.619** | 1.50E-03 | **3.82E-03** |
| ENSG00000086102.21 | **NFX1** | 4799 | protein_coding | **1077.558** | -0.314 | -0.095 | **0.804** | 1.50E-03 | **3.82E-03** |
| ENSG00000142546.16 | **NOSIP** | 51070 | protein_coding | **786.332** | 0.266 | 0.080 | **1.203** | 1.50E-03 | **3.82E-03** |

| | | | | | | | | | |
|---|---|---|---|---|---|---|---|---|---|
| ENSG00000308050.1 | **ENSG00000308050** | | lncRNA | **6.169** | -2.268 | -0.683 | **0.208** | 1.50E-03 | **3.83E-03** |
| ENSG00000095380.13 | **NANS** | 54187 | protein_coding | **934.240** | -0.450 | -0.136 | **0.732** | 1.50E-03 | **3.83E-03** |
| ENSG00000173542.10 | **MOB1B** | 92597 | protein_coding | **231.479** | 0.418 | 0.126 | **1.336** | 1.50E-03 | **3.83E-03** |
| ENSG00000297045.1 | **ENSG0000(** | 10537521ξ | lncRNA | **7.524** | -2.024 | -0.609 | **0.246** | 1.50E-03 | **3.83E-03** |
| ENSG00000156983.19 | **BRPF1** | 7862 | protein_coding | **350.199** | -0.355 | -0.107 | **0.782** | 1.51E-03 | **3.84E-03** |
| ENSG00000177885.17 | **GRB2** | 2885 | protein_coding | **3775.893** | 0.276 | 0.083 | **1.211** | 1.51E-03 | **3.85E-03** |
| ENSG00000231889.10 | **TRAF3IP2-/** | 643749 | lncRNA | **91.102** | -0.568 | -0.171 | **0.674** | 1.51E-03 | **3.85E-03** |
| ENSG00000163629.15 | **PTPN13** | 5783 | protein_coding | **118.056** | 1.102 | 0.332 | **2.146** | 1.52E-03 | **3.86E-03** |
| ENSG00000231672.9 | **DIRC3** | 729582 | lncRNA | **16.217** | 1.232 | 0.371 | **2.348** | 1.52E-03 | **3.87E-03** |
| ENSG00000186212.4 | **SOWAHB** | 345079 | protein_coding | **5.260** | 2.196 | 0.661 | **4.582** | 1.53E-03 | **3.88E-03** |
| ENSG00000223443.3 | **USP17L2** | 377630 | protein_coding | **19.565** | 1.562 | 0.470 | **2.952** | 1.53E-03 | **3.89E-03** |
| ENSG00000271601.6 | **LIX1L** | 128077 | protein_coding | **508.555** | -0.280 | -0.084 | **0.824** | 1.53E-03 | **3.89E-03** |
| ENSG00000260306.1 | **ENSG00000260306** | | lncRNA | **21.717** | 1.114 | 0.335 | **2.164** | 1.53E-03 | **3.89E-03** |
| ENSG00000099817.14 | **POLR2E** | 5434 | protein_coding | **3038.860** | 0.407 | 0.122 | **1.326** | 1.53E-03 | **3.89E-03** |
| ENSG00000105726.19 | **ATP13A1** | 57130 | protein_coding | **2850.032** | 0.353 | 0.106 | **1.277** | 1.53E-03 | **3.89E-03** |
| ENSG00000239697.14 | **TNFSF12** | 8742 | protein_coding | **305.165** | 0.614 | 0.185 | **1.530** | 1.54E-03 | **3.92E-03** |
| ENSG00000121594.14 | **CD80** | 941 | protein_coding | **686.686** | -0.599 | -0.180 | **0.660** | 1.54E-03 | **3.92E-03** |
| ENSG00000289334.1 | **ENSG00000289334** | | lncRNA | **34.844** | 0.907 | 0.273 | **1.876** | 1.55E-03 | **3.93E-03** |
| ENSG00000167460.20 | **TPM4** | 7171 | protein_coding | **7404.871** | -0.516 | -0.155 | **0.700** | 1.55E-03 | **3.94E-03** |
| ENSG00000176124.16 | **DLEU1** | 10301 | lncRNA | **148.072** | 0.662 | 0.199 | **1.582** | 1.55E-03 | **3.95E-03** |
| ENSG00000184508.13 | **HDDC3** | 374659 | protein_coding | **529.363** | 0.298 | 0.090 | **1.229** | 1.56E-03 | **3.95E-03** |
| ENSG00000158517.18 | **NCF1** | 653361 | protein_coding | **9174.087** | 0.373 | 0.112 | **1.295** | 1.56E-03 | **3.96E-03** |
| ENSG00000173786.18 | **CNP** | 1267 | protein_coding | **900.065** | 0.313 | 0.094 | **1.243** | 1.56E-03 | **3.96E-03** |
| ENSG00000260285.1 | **TRMT61A-DT** | | lncRNA | **51.299** | 0.792 | 0.238 | **1.731** | 1.56E-03 | **3.96E-03** |
| ENSG00000247828.11 | **TMEM161I** | 10050589⁄ | lncRNA | **596.807** | -0.370 | -0.111 | **0.774** | 1.56E-03 | **3.97E-03** |
| ENSG00000115295.23 | **CLIP4** | 79745 | protein_coding | **278.455** | -0.385 | -0.116 | **0.766** | 1.57E-03 | **3.97E-03** |
| ENSG00000284691.3 | **ENSG0000(** | 728743 | protein_coding | **878.799** | 0.414 | 0.125 | **1.333** | 1.57E-03 | **3.97E-03** |
| ENSG00000164484.14 | **TMEM200/** | 114801 | protein_coding | **35.123** | -1.130 | -0.340 | **0.457** | 1.57E-03 | **3.98E-03** |
| ENSG00000135974.13 | **C2orf49** | 79074 | protein_coding | **372.857** | -0.412 | -0.124 | **0.752** | 1.57E-03 | **3.98E-03** |
| ENSG00000282199.2 | **ENSG00000282199** | | lncRNA | **16.752** | 1.606 | 0.483 | **3.044** | 1.57E-03 | **3.98E-03** |
| ENSG00000235194.11 | **PPP1R3E** | 90673 | protein_coding | **505.944** | -0.618 | -0.186 | **0.652** | 1.57E-03 | **3.98E-03** |

| | | | | | | | | | |
|---|---|---|---|---|---|---|---|---|---|
| ENSG00000198034.13 | **RPS4X** | 6191 | protein_coding | **50265.708** | -0.314 | -0.095 | **0.804** | 1.57E-03 | **3.98E-03** |
| ENSG00000285886.1 | **ENSG00000285886** | | lncRNA | **37.632** | 0.754 | 0.227 | **1.686** | 1.57E-03 | **3.99E-03** |
| ENSG00000288838.2 | **ENSG0000(** | 12490686! | lncRNA | **314.909** | 0.446 | 0.134 | **1.362** | 1.57E-03 | **3.99E-03** |
| ENSG00000308299.1 | **ENSG00000308299** | | lncRNA | **4.707** | -2.867 | -0.863 | **0.137** | 1.58E-03 | **4.00E-03** |
| ENSG00000270179.1 | **ENSG00000270179** | | lncRNA | **10.963** | 2.401 | 0.723 | **5.281** | 1.58E-03 | **4.01E-03** |
| ENSG00000206567.11 | **ENSG0000(** | 55831 | lncRNA | **12.063** | 1.434 | 0.432 | **2.702** | 1.58E-03 | **4.01E-03** |
| ENSG00000125962.18 | **ARMCX5** | 64860 | protein_coding | **119.748** | -0.547 | -0.165 | **0.684** | 1.58E-03 | **4.01E-03** |
| ENSG00000215845.13 | **TSTD1** | 100131187 | protein_coding | **1046.070** | 0.468 | 0.141 | **1.383** | 1.58E-03 | **4.01E-03** |
| ENSG00000135314.16 | **KHDC1** | 80759 | protein_coding | **65.795** | 0.625 | 0.188 | **1.542** | 1.59E-03 | **4.01E-03** |
| ENSG00000135709.16 | **KIAA0513** | 9764 | protein_coding | **188.982** | 0.904 | 0.272 | **1.871** | 1.59E-03 | **4.01E-03** |
| ENSG00000289909.2 | **ENSG00000289909** | | lncRNA | **6.238** | 2.036 | 0.613 | **4.101** | 1.59E-03 | **4.01E-03** |
| ENSG00000150457.11 | **LATS2** | 26524 | protein_coding | **150.469** | 0.542 | 0.163 | **1.456** | 1.59E-03 | **4.02E-03** |
| ENSG00000107882.14 | **SUFU** | 51684 | protein_coding | **212.384** | 0.367 | 0.110 | **1.289** | 1.59E-03 | **4.02E-03** |
| ENSG00000239857.9 | **GET4** | 51608 | protein_coding | **1627.988** | 0.374 | 0.113 | **1.296** | 1.59E-03 | **4.02E-03** |
| ENSG00000104660.20 | **LEPROTL1** | 23484 | protein_coding | **409.456** | -0.559 | -0.168 | **0.679** | 1.59E-03 | **4.04E-03** |
| ENSG00000259305.6 | **ZHX1-C8or** | 10053310( | protein_coding | **31.553** | -0.801 | -0.241 | **0.574** | 1.60E-03 | **4.04E-03** |
| ENSG00000138593.11 | **SECISBP2L** | 9728 | protein_coding | **544.460** | 0.524 | 0.158 | **1.438** | 1.60E-03 | **4.04E-03** |
| ENSG00000105700.13 | **KXD1** | 79036 | protein_coding | **2414.614** | 0.281 | 0.085 | **1.215** | 1.60E-03 | **4.04E-03** |
| ENSG00000183066.17 | **WBP2NL** | 164684 | protein_coding | **36.861** | 0.915 | 0.276 | **1.886** | 1.60E-03 | **4.05E-03** |
| ENSG00000120509.14 | **PDZD11** | 51248 | protein_coding | **574.387** | -0.366 | -0.110 | **0.776** | 1.60E-03 | **4.05E-03** |
| ENSG00000178104.23 | **PDE4DIP** | 9659 | protein_coding | **195.431** | 0.623 | 0.188 | **1.540** | 1.61E-03 | **4.07E-03** |
| ENSG00000146648.23 | **EGFR** | 1956 | protein_coding | **15.954** | 1.579 | 0.475 | **2.988** | 1.61E-03 | **4.07E-03** |
| ENSG00000080503.27 | **SMARCA2** | 6595 | protein_coding | **2077.021** | 0.747 | 0.225 | **1.678** | 1.61E-03 | **4.07E-03** |
| ENSG00000286115.2 | **ENSG00000286115** | | lncRNA | **8.986** | 1.634 | 0.492 | **3.103** | 1.61E-03 | **4.07E-03** |
| ENSG00000272721.6 | **EIF2B5-DT** | | lncRNA | **18.992** | 1.175 | 0.354 | **2.258** | 1.61E-03 | **4.08E-03** |
| ENSG00000070061.18 | **ELP1** | 8518 | protein_coding | **707.300** | 0.341 | 0.103 | **1.267** | 1.61E-03 | **4.08E-03** |
| ENSG00000164440.16 | **TXLNB** | 167838 | protein_coding | **72.267** | 0.859 | 0.259 | **1.814** | 1.61E-03 | **4.08E-03** |
| ENSG00000289032.2 | **ENSG00000289032** | | lncRNA | **15.865** | 1.211 | 0.365 | **2.315** | 1.61E-03 | **4.08E-03** |
| ENSG00000163159.17 | **VPS72** | 6944 | protein_coding | **654.391** | -0.342 | -0.103 | **0.789** | 1.62E-03 | **4.09E-03** |
| ENSG00000114450.13 | **GNB4** | 59345 | protein_coding | **51.676** | -0.820 | -0.247 | **0.566** | 1.63E-03 | **4.10E-03** |
| ENSG00000072506.16 | **HSD17B10** | 3028 | protein_coding | **1238.562** | -0.387 | -0.116 | **0.765** | 1.63E-03 | **4.11E-03** |

| | | | | | | | | | |
|---|---|---|---|---|---|---|---|---|---|
| ENSG00000129636.15 | **ITFG1** | 81533 | protein_coding | **282.002** | -0.535 | -0.161 | **0.690** | 1.63E-03 | **4.11E-03** |
| ENSG00000198252.14 | **STYX** | 6815 | protein_coding | **183.869** | -0.509 | -0.153 | **0.703** | 1.63E-03 | **4.11E-03** |
| ENSG00000183318.13 | **SPDYE4** | 388333 | protein_coding | **6.451** | 2.150 | 0.647 | **4.437** | 1.63E-03 | **4.12E-03** |
| ENSG00000173208.6 | **ABCD2** | 225 | protein_coding | **62.922** | 0.802 | 0.242 | **1.744** | 1.63E-03 | **4.12E-03** |
| ENSG00000225377.7 | **NRSN2-AS:** | 10050745! | lncRNA | **136.469** | -0.567 | -0.171 | **0.675** | 1.64E-03 | **4.13E-03** |
| ENSG00000110711.13 | **AIP** | 9049 | protein_coding | **1822.001** | -0.352 | -0.106 | **0.783** | 1.64E-03 | **4.14E-03** |
| ENSG00000182612.13 | **TSPAN10** | 83882 | protein_coding | **22.618** | 1.023 | 0.308 | **2.033** | 1.65E-03 | **4.15E-03** |
| ENSG00000175137.13 | **SH3BP5L** | 80851 | protein_coding | **400.539** | 0.382 | 0.115 | **1.303** | 1.66E-03 | **4.17E-03** |
| ENSG00000070214.19 | **SLC44A1** | 23446 | protein_coding | **661.211** | 0.384 | 0.115 | **1.305** | 1.66E-03 | **4.19E-03** |
| ENSG00000169814.20 | **BTD** | 686 | protein_coding | **114.644** | 0.691 | 0.208 | **1.614** | 1.66E-03 | **4.20E-03** |
| ENSG00000164163.13 | **ABCE1** | 6059 | protein_coding | **809.281** | -0.486 | -0.146 | **0.714** | 1.67E-03 | **4.20E-03** |
| ENSG00000198515.18 | **CNGA1** | 1259 | protein_coding | **5.597** | 3.395 | 1.022 | **10.519** | 1.67E-03 | **4.21E-03** |
| ENSG00000188010.18 | **MORN2** | 729967 | protein_coding | **91.511** | 0.623 | 0.188 | **1.540** | 1.67E-03 | **4.21E-03** |
| ENSG00000293477.2 | **CRYZL2P** | 730102 | lncRNA | **5.859** | 2.395 | 0.721 | **5.261** | 1.68E-03 | **4.24E-03** |
| ENSG00000301508.1 | **ENSG00000301508** | | lncRNA | **9.323** | 1.575 | 0.474 | **2.980** | 1.68E-03 | **4.24E-03** |
| ENSG00000184203.10 | **PPP1R2** | 5504 | protein_coding | **633.717** | -0.486 | -0.146 | **0.714** | 1.69E-03 | **4.25E-03** |
| ENSG00000136156.17 | **ITM2B** | 9445 | protein_coding | **852.733** | 0.446 | 0.134 | **1.362** | 1.69E-03 | **4.27E-03** |
| ENSG00000234745.16 | **HLA-B** | 3106 | protein_coding | **51341.668** | -0.351 | -0.106 | **0.784** | 1.69E-03 | **4.27E-03** |
| ENSG00000031823.16 | **RANBP3** | 8498 | protein_coding | **1342.180** | -0.319 | -0.096 | **0.801** | 1.70E-03 | **4.27E-03** |
| ENSG00000169871.15 | **TRIM56** | 81844 | protein_coding | **1578.870** | -0.307 | -0.092 | **0.808** | 1.70E-03 | **4.27E-03** |
| ENSG00000302539.1 | **ENSG00000302539** | | lncRNA | **10.482** | -1.591 | -0.479 | **0.332** | 1.70E-03 | **4.28E-03** |
| ENSG00000257017.12 | **HP** | 3240 | protein_coding | **17.596** | 2.585 | 0.778 | **6.000** | 1.71E-03 | **4.31E-03** |
| ENSG00000298585.1 | **ENSG00000298585** | | lncRNA | **4.785** | 2.335 | 0.703 | **5.046** | 1.71E-03 | **4.31E-03** |
| ENSG00000258183.6 | **LINC02392** | | lncRNA | **6.207** | 2.662 | 0.801 | **6.330** | 1.71E-03 | **4.32E-03** |
| ENSG00000260711.2 | **ENSG00000260711** | | lncRNA | **14.275** | 1.559 | 0.469 | **2.947** | 1.72E-03 | **4.32E-03** |
| ENSG00000108641.21 | **B9D1** | 27077 | protein_coding | **149.223** | 0.502 | 0.151 | **1.416** | 1.72E-03 | **4.32E-03** |
| ENSG00000131051.26 | **RBM39** | 9584 | protein_coding | **4370.927** | -0.411 | -0.124 | **0.752** | 1.72E-03 | **4.32E-03** |
| ENSG00000205832.9 | **C16orf96** | 342346 | protein_coding | **5.380** | 2.348 | 0.707 | **5.090** | 1.72E-03 | **4.33E-03** |
| ENSG00000164631.20 | **ZNF12** | 7559 | protein_coding | **437.966** | -0.406 | -0.122 | **0.755** | 1.72E-03 | **4.33E-03** |
| ENSG00000066185.15 | **ZMYND12** | 84217 | protein_coding | **25.360** | 0.970 | 0.292 | **1.959** | 1.72E-03 | **4.33E-03** |
| ENSG00000138439.13 | **FAM117B** | 150864 | protein_coding | **94.673** | 0.628 | 0.189 | **1.546** | 1.73E-03 | **4.35E-03** |

| | | | | | | | | | |
|---|---|---|---|---|---|---|---|---|---|
| ENSG00000309428.1 | **ENSG00000309428** | | lncRNA | **157.138** | -0.488 | -0.147 | **0.713** | 1.73E-03 | **4.36E-03** |
| ENSG00000286535.2 | **ENSG0000(** | 12490202( | lncRNA | **17.652** | 1.412 | 0.425 | **2.662** | 1.73E-03 | **4.36E-03** |
| ENSG00000186026.9 | **ZNF284** | 342909 | protein_coding | **95.284** | 0.502 | 0.151 | **1.416** | 1.74E-03 | **4.36E-03** |
| ENSG00000154310.19 | **TNIK** | 23043 | protein_coding | **899.087** | -0.511 | -0.154 | **0.702** | 1.74E-03 | **4.36E-03** |
| ENSG00000143772.13 | **ITPKB** | 3707 | protein_coding | **586.613** | -0.426 | -0.128 | **0.744** | 1.74E-03 | **4.37E-03** |
| ENSG00000290711.2 | **ENSG00000290711** | | lncRNA | **8.019** | 2.472 | 0.744 | **5.548** | 1.75E-03 | **4.39E-03** |
| ENSG00000306113.1 | **ENSG00000306113** | | lncRNA | **5.235** | 2.274 | 0.684 | **4.836** | 1.75E-03 | **4.39E-03** |
| ENSG00000143774.20 | **GUK1** | 2987 | protein_coding | **2779.508** | -0.329 | -0.099 | **0.796** | 1.75E-03 | **4.39E-03** |
| ENSG00000006534.19 | **ALDH3B1** | 221 | protein_coding | **20.926** | 1.452 | 0.437 | **2.735** | 1.75E-03 | **4.40E-03** |
| ENSG00000217442.5 | **SYCE3** | 644186 | protein_coding | **12.458** | 1.762 | 0.530 | **3.392** | 1.75E-03 | **4.40E-03** |
| ENSG00000039123.18 | **MTREX** | 23517 | protein_coding | **602.039** | -0.478 | -0.144 | **0.718** | 1.76E-03 | **4.41E-03** |
| ENSG00000303988.1 | **ENSG00000303988** | | lncRNA | **21.285** | -1.100 | -0.331 | **0.467** | 1.77E-03 | **4.43E-03** |
| ENSG00000244462.10 | **RBM12** | 10137 | protein_coding | **633.478** | -0.456 | -0.137 | **0.729** | 1.77E-03 | **4.44E-03** |
| ENSG00000198873.14 | **GRK5** | 2869 | protein_coding | **394.965** | -0.365 | -0.110 | **0.776** | 1.77E-03 | **4.45E-03** |
| ENSG00000137726.19 | **FXYD6** | 53826 | protein_coding | **29.023** | 1.333 | 0.401 | **2.520** | 1.78E-03 | **4.46E-03** |
| ENSG00000229043.4 | **ZFAND2A-DT** | | lncRNA | **175.844** | 0.506 | 0.152 | **1.421** | 1.78E-03 | **4.46E-03** |
| ENSG00000262903.1 | **CTNS-AS1** | | lncRNA | **7.554** | 1.640 | 0.494 | **3.116** | 1.78E-03 | **4.46E-03** |
| ENSG00000232485.5 | **RPL37A-DT** | 10050755 | lncRNA | **25.870** | 1.089 | 0.328 | **2.128** | 1.78E-03 | **4.47E-03** |
| ENSG00000273841.8 | **TAF9** | 6880 | protein_coding | **781.979** | -0.376 | -0.113 | **0.771** | 1.78E-03 | **4.47E-03** |
| ENSG00000287966.4 | **LARRPM** | | lncRNA | **21.593** | 1.281 | 0.386 | **2.431** | 1.78E-03 | **4.47E-03** |
| ENSG00000289257.3 | **ENSG00000289257** | | lncRNA | **5.740** | 1.966 | 0.592 | **3.906** | 1.78E-03 | **4.47E-03** |
| ENSG00000182782.8 | **HCAR2** | 338442 | protein_coding | **28.036** | 1.096 | 0.330 | **2.138** | 1.78E-03 | **4.47E-03** |
| ENSG00000187912.12 | **CLEC17A** | 388512 | protein_coding | **1302.633** | -0.407 | -0.123 | **0.754** | 1.78E-03 | **4.47E-03** |
| ENSG00000289410.3 | **ENSG0000(** | 107985767 | lncRNA | **25.142** | 0.930 | 0.280 | **1.906** | 1.79E-03 | **4.48E-03** |
| ENSG00000156374.18 | **PCGF6** | 84108 | protein_coding | **95.931** | -0.579 | -0.174 | **0.669** | 1.79E-03 | **4.48E-03** |
| ENSG00000107140.19 | **TESK1** | 7016 | protein_coding | **663.585** | 0.420 | 0.126 | **1.338** | 1.79E-03 | **4.49E-03** |
| ENSG00000051523.14 | **CYBA** | 1535 | protein_coding | **5992.781** | -0.395 | -0.119 | **0.761** | 1.79E-03 | **4.49E-03** |
| ENSG00000119673.18 | **ACOT2** | 10965 | protein_coding | **277.109** | -0.463 | -0.140 | **0.725** | 1.80E-03 | **4.50E-03** |
| ENSG00000260303.1 | **ENSG00000260303** | | lncRNA | **30.361** | 0.957 | 0.288 | **1.941** | 1.80E-03 | **4.50E-03** |
| ENSG00000138028.18 | **CGREF1** | 10669 | protein_coding | **23.329** | 1.421 | 0.428 | **2.678** | 1.80E-03 | **4.50E-03** |
| ENSG00000116874.14 | **WARS2** | 10352 | protein_coding | **175.081** | -0.474 | -0.143 | **0.720** | 1.80E-03 | **4.50E-03** |

| | | | | | | | | | |
|---|---|---|---|---|---|---|---|---|---|
| ENSG00000309655.1 | **ENSG00000309655** | | lncRNA | **38.781** | 0.766 | 0.231 | **1.701** | 1.80E-03 | **4.51E-03** |
| ENSG00000105447.15 | **GRWD1** | 83743 | protein_coding | **473.323** | -0.345 | -0.104 | **0.787** | 1.81E-03 | **4.54E-03** |
| ENSG00000110074.14 | **FOXRED1** | 55572 | protein_coding | **953.683** | -0.388 | -0.117 | **0.764** | 1.81E-03 | **4.54E-03** |
| ENSG00000293930.1 | **ENSG00000293930** | | lncRNA | **12.854** | 1.202 | 0.362 | **2.301** | 1.82E-03 | **4.54E-03** |
| ENSG00000100442.13 | **FKBP3** | 2287 | protein_coding | **1286.492** | -0.303 | -0.091 | **0.810** | 1.82E-03 | **4.55E-03** |
| ENSG00000259956.2 | **RBM15B** | 29890 | protein_coding | **794.947** | -0.285 | -0.086 | **0.821** | 1.82E-03 | **4.55E-03** |
| ENSG00000213949.11 | **ITGA1** | 3672 | protein_coding | **85.037** | 0.766 | 0.231 | **1.701** | 1.82E-03 | **4.56E-03** |
| ENSG00000196357.14 | **ZNF565** | 147929 | protein_coding | **57.038** | 0.710 | 0.214 | **1.636** | 1.82E-03 | **4.56E-03** |
| ENSG00000175197.14 | **DDIT3** | 1649 | protein_coding | **712.514** | -0.741 | -0.223 | **0.599** | 1.82E-03 | **4.56E-03** |
| ENSG00000204388.7 | **HSPA1B** | 3304 | protein_coding | **385.396** | 0.700 | 0.211 | **1.625** | 1.82E-03 | **4.56E-03** |
| ENSG00000140398.16 | **NEIL1** | 79661 | protein_coding | **298.338** | 0.864 | 0.260 | **1.820** | 1.83E-03 | **4.58E-03** |
| ENSG00000237491.11 | **LINC01409** | | lncRNA | **55.241** | -0.679 | -0.204 | **0.625** | 1.84E-03 | **4.59E-03** |
| ENSG00000090581.13 | **GNPTG** | 84572 | protein_coding | **789.995** | 0.328 | 0.099 | **1.256** | 1.84E-03 | **4.60E-03** |
| ENSG00000167986.17 | **DDB1** | 1642 | protein_coding | **2504.291** | -0.314 | -0.095 | **0.804** | 1.84E-03 | **4.61E-03** |
| ENSG00000083454.25 | **P2RX5** | 5026 | protein_coding | **750.186** | 0.451 | 0.136 | **1.367** | 1.85E-03 | **4.62E-03** |
| ENSG00000170917.15 | **NUDT6** | 11162 | protein_coding | **160.181** | 0.513 | 0.154 | **1.427** | 1.85E-03 | **4.62E-03** |
| ENSG00000060709.19 | **RIMBP2** | 23504 | protein_coding | **197.114** | -0.665 | -0.200 | **0.631** | 1.85E-03 | **4.62E-03** |
| ENSG00000186642.17 | **PDE2A** | 5138 | protein_coding | **11.065** | 1.375 | 0.414 | **2.594** | 1.85E-03 | **4.62E-03** |
| ENSG00000070047.15 | **PHRF1** | 57661 | protein_coding | **1576.106** | 0.286 | 0.086 | **1.219** | 1.86E-03 | **4.64E-03** |
| ENSG00000165669.16 | **FAM204A** | 63877 | protein_coding | **510.824** | -0.448 | -0.135 | **0.733** | 1.86E-03 | **4.64E-03** |
| ENSG00000163956.15 | **LRPAP1** | 4043 | protein_coding | **663.454** | -0.351 | -0.106 | **0.784** | 1.86E-03 | **4.64E-03** |
| ENSG00000255398.3 | **HCAR3** | 8843 | protein_coding | **118.214** | 0.930 | 0.280 | **1.905** | 1.87E-03 | **4.67E-03** |
| ENSG00000306338.1 | **ENSG00000306338** | | lncRNA | **9.241** | 1.634 | 0.492 | **3.103** | 1.87E-03 | **4.67E-03** |
| ENSG00000272335.1 | **ENSG00000272335** | | lncRNA | **117.611** | 0.450 | 0.136 | **1.366** | 1.87E-03 | **4.67E-03** |
| ENSG00000231993.3 | **EP300-AS1** | 10192727[illegible] | lncRNA | **6.343** | 2.052 | 0.618 | **4.147** | 1.87E-03 | **4.67E-03** |
| ENSG00000176261.18 | **ZBTB8OS** | 339487 | protein_coding | **1024.353** | -0.410 | -0.123 | **0.753** | 1.87E-03 | **4.67E-03** |
| ENSG00000139146.16 | **SINHCAF** | 58516 | protein_coding | **1287.602** | -0.384 | -0.115 | **0.766** | 1.87E-03 | **4.67E-03** |
| ENSG00000146021.17 | **KLHL3** | 26249 | protein_coding | **295.056** | -0.376 | -0.113 | **0.771** | 1.87E-03 | **4.67E-03** |
| ENSG00000177807.12 | **KCNJ10** | 3766 | protein_coding | **4.312** | 2.859 | 0.861 | **7.256** | 1.88E-03 | **4.70E-03** |
| ENSG00000119547.6 | **ONECUT2** | 9480 | protein_coding | **7.328** | 2.074 | 0.624 | **4.211** | 1.88E-03 | **4.70E-03** |
| ENSG00000170271.13 | **FAXDC2** | 10826 | protein_coding | **9.468** | 1.662 | 0.500 | **3.165** | 1.89E-03 | **4.71E-03** |

| | | | | | | | | | |
|---|---|---|---|---|---|---|---|---|---|
| ENSG00000236830.9 | **CBR3-AS1** | 10050642₈ | lncRNA | **6.059** | 2.050 | 0.617 | **4.142** | 1.89E-03 | **4.71E-03** |
| ENSG00000179085.10 | **DPM3** | 54344 | protein_coding | **261.305** | 0.459 | 0.138 | **1.374** | 1.89E-03 | **4.71E-03** |
| ENSG00000290993.2 | **ENSG00000290993** | | lncRNA | **555.771** | 0.281 | 0.084 | **1.215** | 1.89E-03 | **4.72E-03** |
| ENSG00000022277.15 | **RTF2** | 51507 | protein_coding | **1686.262** | 0.326 | 0.098 | **1.253** | 1.90E-03 | **4.72E-03** |
| ENSG00000149131.19 | **SERPING1** | 710 | protein_coding | **8.059** | 2.414 | 0.727 | **5.328** | 1.90E-03 | **4.73E-03** |
| ENSG00000135678.14 | **CPM** | 1368 | protein_coding | **42.515** | 1.069 | 0.322 | **2.098** | 1.90E-03 | **4.73E-03** |
| ENSG00000294432.1 | **ENSG00000294432** | | lncRNA | **4.819** | 2.117 | 0.637 | **4.337** | 1.90E-03 | **4.73E-03** |
| ENSG00000273156.2 | **ENSG00000273156** | | lncRNA | **13.759** | 1.152 | 0.347 | **2.223** | 1.90E-03 | **4.73E-03** |
| ENSG00000148688.16 | **RPP30** | 10556 | protein_coding | **507.776** | -0.289 | -0.087 | **0.818** | 1.90E-03 | **4.73E-03** |
| ENSG00000112305.17 | **SMAP1** | 60682 | protein_coding | **354.162** | -0.330 | -0.099 | **0.796** | 1.90E-03 | **4.74E-03** |
| ENSG00000137142.7 | **IGFBPL1** | 347252 | protein_coding | **7.533** | 1.778 | 0.535 | **3.429** | 1.90E-03 | **4.74E-03** |
| ENSG00000168491.11 | **CCDC110** | 256309 | protein_coding | **21.264** | 1.557 | 0.469 | **2.943** | 1.90E-03 | **4.74E-03** |
| ENSG00000137875.4 | **BCL2L10** | 10017 | protein_coding | **99.921** | -0.689 | -0.207 | **0.620** | 1.91E-03 | **4.75E-03** |
| ENSG00000174516.17 | **PELI3** | 246330 | protein_coding | **65.856** | 0.663 | 0.200 | **1.584** | 1.91E-03 | **4.75E-03** |
| ENSG00000223501.11 | **VPS52** | 6293 | protein_coding | **889.569** | -0.360 | -0.108 | **0.779** | 1.91E-03 | **4.76E-03** |
| ENSG00000198478.11 | **SH3BGRL2** | 83699 | protein_coding | **16.679** | 1.153 | 0.347 | **2.223** | 1.92E-03 | **4.77E-03** |
| ENSG00000175344.21 | **CHRNA7** | 1139 | protein_coding | **4.567** | 2.960 | 0.891 | **7.783** | 1.92E-03 | **4.77E-03** |
| ENSG00000224875.3 | **EML4-AS1** | 10272382₄ | lncRNA | **4.875** | -2.817 | -0.848 | **0.142** | 1.92E-03 | **4.78E-03** |
| ENSG00000297460.1 | **ENSG0000(** | 10537662₇ | lncRNA | **50.269** | 0.693 | 0.209 | **1.617** | 1.92E-03 | **4.78E-03** |
| ENSG00000103326.14 | **CAPN15** | 6650 | protein_coding | **1601.032** | -0.525 | -0.158 | **0.695** | 1.93E-03 | **4.80E-03** |
| ENSG00000156482.13 | **RPL30** | 6156 | protein_coding | **16383.152** | -0.352 | -0.106 | **0.784** | 1.93E-03 | **4.80E-03** |
| ENSG00000203666.15 | **DRC8** | 84288 | protein_coding | **73.291** | -0.770 | -0.232 | **0.586** | 1.93E-03 | **4.81E-03** |
| ENSG00000309346.1 | **FCHO1-AS1** | | lncRNA | **8.912** | 1.468 | 0.442 | **2.767** | 1.94E-03 | **4.81E-03** |
| ENSG00000157578.16 | **LCA5L** | 150082 | protein_coding | **4.839** | 3.066 | 0.923 | **8.372** | 1.94E-03 | **4.83E-03** |
| ENSG00000289891.2 | **ENSG00000289891** | | lncRNA | **11.110** | 1.432 | 0.431 | **2.698** | 1.94E-03 | **4.83E-03** |
| ENSG00000176978.18 | **DPP7** | 29952 | protein_coding | **3939.314** | -0.526 | -0.158 | **0.694** | 1.94E-03 | **4.83E-03** |
| ENSG00000235244.7 | **DANT2** | 642776 | lncRNA | **61.962** | -0.912 | -0.275 | **0.531** | 1.94E-03 | **4.83E-03** |
| ENSG00000116641.20 | **DOCK7** | 85440 | protein_coding | **292.198** | -1.488 | -0.448 | **0.356** | 1.95E-03 | **4.84E-03** |
| ENSG00000147162.17 | **OGT** | 8473 | protein_coding | **8971.229** | -0.455 | -0.137 | **0.729** | 1.95E-03 | **4.84E-03** |
| ENSG00000144674.21 | **GOLGA4** | 2803 | protein_coding | **554.384** | -0.360 | -0.108 | **0.779** | 1.95E-03 | **4.84E-03** |
| ENSG00000266173.9 | **STRADA** | 92335 | protein_coding | **1360.224** | -0.356 | -0.107 | **0.782** | 1.95E-03 | **4.84E-03** |

| | | | | | | | | | |
|---|---|---|---|---|---|---|---|---|---|
| ENSG00000067177.17 | **PHKA1** | 5255 | protein_coding | **34.687** | 0.957 | 0.288 | **1.941** | 1.95E-03 | **4.84E-03** |
| ENSG00000204514.12 | **ZNF814** | 730051 | protein_coding | **209.357** | -0.474 | -0.143 | **0.720** | 1.95E-03 | **4.85E-03** |
| ENSG00000118557.17 | **PMFBP1** | 83449 | protein_coding | **44.335** | -0.900 | -0.271 | **0.536** | 1.95E-03 | **4.85E-03** |
| ENSG00000186166.11 | **CENATAC** | 338657 | protein_coding | **878.870** | 0.407 | 0.122 | **1.326** | 1.96E-03 | **4.86E-03** |
| ENSG00000147789.18 | **ZNF7** | 7553 | protein_coding | **276.069** | -0.333 | -0.100 | **0.794** | 1.96E-03 | **4.86E-03** |
| ENSG00000226976.3 | **COX6A1P2** | | processed_pseudo | **5.115** | 2.222 | 0.669 | **4.664** | 1.96E-03 | **4.86E-03** |
| ENSG00000091972.21 | **CD200** | 4345 | protein_coding | **320.428** | -0.545 | -0.164 | **0.685** | 1.96E-03 | **4.87E-03** |
| ENSG00000122565.21 | **CBX3** | 11335 | protein_coding | **2436.203** | -0.326 | -0.098 | **0.798** | 1.97E-03 | **4.88E-03** |
| ENSG00000301079.1 | **ENSG00000301079** | | lncRNA | **8.936** | 1.460 | 0.440 | **2.751** | 1.97E-03 | **4.88E-03** |
| ENSG00000290035.2 | **ENSG00000290035** | | lncRNA | **11.531** | 1.694 | 0.510 | **3.236** | 1.97E-03 | **4.89E-03** |
| ENSG00000152229.20 | **PSTPIP2** | 9050 | protein_coding | **210.514** | -0.368 | -0.111 | **0.775** | 1.97E-03 | **4.90E-03** |
| ENSG00000143149.15 | **ALDH9A1** | 223 | protein_coding | **607.155** | -0.324 | -0.097 | **0.799** | 1.98E-03 | **4.90E-03** |
| ENSG00000008300.17 | **CELSR3** | 1951 | protein_coding | **209.763** | 0.555 | 0.167 | **1.469** | 1.98E-03 | **4.90E-03** |
| ENSG00000290926.2 | **LINC03006** | 441242 | lncRNA | **374.840** | 0.346 | 0.104 | **1.271** | 1.98E-03 | **4.91E-03** |
| ENSG00000118292.11 | **C1orf54** | 79630 | protein_coding | **154.390** | -0.651 | -0.196 | **0.637** | 1.98E-03 | **4.91E-03** |
| ENSG00000153046.20 | **CDYL** | 9425 | protein_coding | **428.813** | 0.513 | 0.155 | **1.427** | 1.98E-03 | **4.92E-03** |
| ENSG00000255062.3 | **TIRAP-AS1** | 105369557 | lncRNA | **14.563** | -1.157 | -0.348 | **0.448** | 1.98E-03 | **4.92E-03** |
| ENSG00000113615.15 | **SEC24A** | 10802 | protein_coding | **363.404** | -0.553 | -0.167 | **0.681** | 1.99E-03 | **4.93E-03** |
| ENSG00000251661.4 | **IFITM3-AS1** | | lncRNA | **176.611** | 0.500 | 0.151 | **1.414** | 1.99E-03 | **4.94E-03** |
| ENSG00000221887.7 | **HMSD** | 284293 | protein_coding | **502.501** | 0.494 | 0.149 | **1.409** | 2.00E-03 | **4.94E-03** |
| ENSG00000134901.16 | **POGLUT2** | 79070 | protein_coding | **122.156** | 0.583 | 0.175 | **1.498** | 2.00E-03 | **4.95E-03** |
| ENSG00000188171.18 | **ZNF626** | 199777 | protein_coding | **178.174** | 0.447 | 0.135 | **1.363** | 2.00E-03 | **4.95E-03** |
| ENSG00000165527.10 | **ARF6** | 382 | protein_coding | **1785.438** | -0.350 | -0.105 | **0.784** | 2.00E-03 | **4.96E-03** |
| ENSG00000167978.20 | **SRRM2** | 23524 | protein_coding | **21052.653** | 0.389 | 0.117 | **1.309** | 2.02E-03 | **5.01E-03** |
| ENSG00000106686.20 | **SPATA6L** | 55064 | protein_coding | **31.641** | 0.944 | 0.284 | **1.924** | 2.03E-03 | **5.02E-03** |
| ENSG00000198840.2 | **MT-ND3** | 4537 | protein_coding | **28306.239** | 0.557 | 0.168 | **1.471** | 2.03E-03 | **5.03E-03** |
| ENSG00000096093.18 | **EFHC1** | 114327 | protein_coding | **172.050** | 0.770 | 0.232 | **1.705** | 2.04E-03 | **5.04E-03** |
| ENSG00000180855.16 | **ZNF443** | 10224 | protein_coding | **60.352** | -0.648 | -0.195 | **0.638** | 2.05E-03 | **5.06E-03** |
| ENSG00000265148.8 | **TSPOAP1-A** | 100506779 | lncRNA | **320.547** | -0.341 | -0.103 | **0.790** | 2.05E-03 | **5.06E-03** |
| ENSG00000074621.16 | **SLC24A1** | 9187 | protein_coding | **74.150** | 0.697 | 0.210 | **1.622** | 2.05E-03 | **5.06E-03** |
| ENSG00000132694.22 | **ARHGEF11** | 9826 | protein_coding | **375.017** | 0.483 | 0.145 | **1.398** | 2.05E-03 | **5.08E-03** |

| | | | | | | | | | |
|---|---|---|---|---|---|---|---|---|---|
| ENSG00000143198.15 | **MGST3** | 4259 | protein_coding | **940.499** | 0.453 | 0.136 | **1.369** | 2.06E-03 | **5.09E-03** |
| ENSG00000126524.14 | **SBDS** | 51119 | protein_coding | **432.321** | -0.341 | -0.103 | **0.790** | 2.06E-03 | **5.09E-03** |
| ENSG00000270800.4 | **RPS10-NUI** | 10052923( | protein_coding | **9.341** | -3.026 | -0.911 | **0.123** | 2.07E-03 | **5.11E-03** |
| ENSG00000178222.15 | **RNF212** | 285498 | protein_coding | **9.555** | 1.733 | 0.522 | **3.324** | 2.07E-03 | **5.11E-03** |
| ENSG00000133104.16 | **SPART** | 23111 | protein_coding | **158.721** | 0.700 | 0.211 | **1.625** | 2.07E-03 | **5.13E-03** |
| ENSG00000289688.2 | **ENSG00000289688** | | lncRNA | **14.321** | 2.277 | 0.685 | **4.846** | 2.08E-03 | **5.14E-03** |
| ENSG00000137509.14 | **PRCP** | 5547 | protein_coding | **633.843** | 0.407 | 0.123 | **1.326** | 2.08E-03 | **5.14E-03** |
| ENSG00000290758.3 | **POLR2J4** | 84820 | lncRNA | **776.535** | 0.313 | 0.094 | **1.242** | 2.08E-03 | **5.14E-03** |
| ENSG00000225978.4 | **HAR1A** | | lncRNA | **32.107** | 0.896 | 0.270 | **1.862** | 2.08E-03 | **5.14E-03** |
| ENSG00000278158.3 | **ENSG00000278158** | | lncRNA | **33.709** | -0.903 | -0.272 | **0.535** | 2.09E-03 | **5.15E-03** |
| ENSG00000293430.2 | **ENSG00000293430** | | lncRNA | **5.784** | 2.062 | 0.621 | **4.176** | 2.09E-03 | **5.16E-03** |
| ENSG00000287445.2 | **ENSG0000(** | 10537290( | lncRNA | **43.498** | 1.022 | 0.308 | **2.031** | 2.09E-03 | **5.16E-03** |
| ENSG00000260917.1 | **ENSG0000(** | 103344931 | lncRNA | **104.600** | 0.582 | 0.175 | **1.497** | 2.09E-03 | **5.16E-03** |
| ENSG00000144645.18 | **OSBPL10** | 114884 | protein_coding | **5.917** | 2.643 | 0.796 | **6.248** | 2.09E-03 | **5.16E-03** |
| ENSG00000134160.17 | **TRPM1** | 4308 | protein_coding | **5.583** | 2.900 | 0.873 | **7.466** | 2.10E-03 | **5.17E-03** |
| ENSG00000121989.17 | **ACVR2A** | 92 | protein_coding | **33.541** | 1.445 | 0.435 | **2.722** | 2.10E-03 | **5.18E-03** |
| ENSG00000204231.13 | **RXRB** | 6257 | protein_coding | **905.291** | 0.366 | 0.110 | **1.289** | 2.11E-03 | **5.19E-03** |
| ENSG00000170448.14 | **NFXL1** | 152518 | protein_coding | **113.461** | -0.621 | -0.187 | **0.650** | 2.11E-03 | **5.20E-03** |
| ENSG00000139278.11 | **GLIPR1** | 11010 | protein_coding | **331.185** | -0.481 | -0.145 | **0.716** | 2.11E-03 | **5.20E-03** |
| ENSG00000064961.21 | **HMG20B** | 10362 | protein_coding | **1909.707** | 0.401 | 0.121 | **1.321** | 2.11E-03 | **5.20E-03** |
| ENSG00000258875.7 | **ENSG00000258875** | | lncRNA | **25.688** | -0.963 | -0.290 | **0.513** | 2.11E-03 | **5.21E-03** |
| ENSG00000278834.3 | **ENSG00000278834** | | lncRNA | **79.299** | 0.621 | 0.187 | **1.538** | 2.12E-03 | **5.22E-03** |
| ENSG00000072786.15 | **STK10** | 6793 | protein_coding | **2505.345** | -0.280 | -0.084 | **0.824** | 2.12E-03 | **5.23E-03** |
| ENSG00000253755.1 | **IGHGP** | | IG_C_pseudogene | **467.801** | -0.489 | -0.147 | **0.712** | 2.13E-03 | **5.24E-03** |
| ENSG00000111880.18 | **RNGTT** | 8732 | protein_coding | **366.957** | -0.579 | -0.174 | **0.670** | 2.13E-03 | **5.25E-03** |
| ENSG00000148248.16 | **SURF4** | 6836 | protein_coding | **1457.372** | -0.366 | -0.110 | **0.776** | 2.13E-03 | **5.25E-03** |
| ENSG00000144354.17 | **CDCA7** | 83879 | protein_coding | **578.762** | -0.387 | -0.116 | **0.765** | 2.13E-03 | **5.25E-03** |
| ENSG00000171246.8 | **NPTX1** | 4884 | protein_coding | **17.308** | 1.586 | 0.478 | **3.003** | 2.13E-03 | **5.25E-03** |
| ENSG00000270505.1 | **IGHV1OR15-1** | | IG_V_gene | **203.292** | -0.613 | -0.184 | **0.654** | 2.13E-03 | **5.26E-03** |
| ENSG00000224046.4 | **DMTF1-AS** | 10192742( | lncRNA | **28.980** | 0.922 | 0.277 | **1.894** | 2.13E-03 | **5.26E-03** |
| ENSG00000275198.2 | **LINC02960** | | lncRNA | **15.973** | 1.167 | 0.351 | **2.245** | 2.14E-03 | **5.27E-03** |

| | | | | | | | | | |
|---|---|---|---|---|---|---|---|---|---|
| ENSG00000272808.5 | **GCAWKR** | 105369201 | lncRNA | **5.263** | 2.765 | 0.832 | **6.796** | 2.14E-03 | **5.27E-03** |
| ENSG00000287001.4 | **ENSG00000287001** | | lncRNA | **3.834** | 3.387 | 1.020 | **10.463** | 2.14E-03 | **5.28E-03** |
| ENSG00000186073.16 | **CDIN1** | 84529 | protein_coding | **120.305** | -0.490 | -0.147 | **0.712** | 2.15E-03 | **5.28E-03** |
| ENSG00000259366.3 | **ENSG00000259366** | | lncRNA | **16.822** | -1.158 | -0.349 | **0.448** | 2.15E-03 | **5.30E-03** |
| ENSG00000127951.9 | **FGL2** | 10875 | protein_coding | **20.805** | -1.223 | -0.368 | **0.428** | 2.16E-03 | **5.31E-03** |
| ENSG00000157191.22 | **NECAP2** | 55707 | protein_coding | **1318.305** | 0.373 | 0.112 | **1.295** | 2.16E-03 | **5.32E-03** |
| ENSG00000187742.17 | **SECISBP2** | 79048 | protein_coding | **1021.529** | 0.294 | 0.088 | **1.226** | 2.16E-03 | **5.32E-03** |
| ENSG00000106991.17 | **ENG** | 2022 | protein_coding | **240.787** | -0.461 | -0.139 | **0.727** | 2.16E-03 | **5.32E-03** |
| ENSG00000177051.8 | **FBXO46** | 23403 | protein_coding | **1157.388** | 0.286 | 0.086 | **1.219** | 2.16E-03 | **5.32E-03** |
| ENSG00000188693.9 | **CYP51A1-A** | 613126 | lncRNA | **16.289** | 1.062 | 0.320 | **2.087** | 2.16E-03 | **5.32E-03** |
| ENSG00000168159.16 | **RNF187** | 149603 | protein_coding | **1630.543** | -0.337 | -0.101 | **0.792** | 2.17E-03 | **5.33E-03** |
| ENSG00000154814.16 | **OXNAD1** | 92106 | protein_coding | **224.690** | -0.437 | -0.131 | **0.739** | 2.17E-03 | **5.34E-03** |
| ENSG00000196967.13 | **ZNF585A** | 199704 | protein_coding | **240.356** | -0.425 | -0.128 | **0.745** | 2.17E-03 | **5.34E-03** |
| ENSG00000278996.2 | **ENSG00000278996** | | lncRNA | **65.462** | -1.108 | -0.334 | **0.464** | 2.17E-03 | **5.35E-03** |
| ENSG00000088038.22 | **CNOT3** | 4849 | protein_coding | **1451.037** | -0.513 | -0.154 | **0.701** | 2.18E-03 | **5.35E-03** |
| ENSG00000286874.2 | **ENSG00000286874** | | lncRNA | **11.632** | 1.377 | 0.415 | **2.598** | 2.18E-03 | **5.35E-03** |
| ENSG00000133858.18 | **ZFC3H1** | 196441 | protein_coding | **1014.930** | 0.460 | 0.138 | **1.375** | 2.18E-03 | **5.36E-03** |
| ENSG00000135829.19 | **DHX9** | 1660 | protein_coding | **1794.443** | -0.300 | -0.090 | **0.813** | 2.18E-03 | **5.36E-03** |
| ENSG00000010610.12 | **CD4** | 920 | protein_coding | **76.203** | 0.878 | 0.264 | **1.838** | 2.18E-03 | **5.36E-03** |
| ENSG00000101342.12 | **TLDC2** | 140711 | protein_coding | **43.558** | -1.056 | -0.318 | **0.481** | 2.19E-03 | **5.39E-03** |
| ENSG00000300574.1 | **ENSG00000300574** | | lncRNA | **6.846** | 2.019 | 0.608 | **4.054** | 2.20E-03 | **5.41E-03** |
| ENSG00000175463.14 | **TBC1D10C** | 374403 | protein_coding | **1510.847** | -0.618 | -0.186 | **0.652** | 2.20E-03 | **5.41E-03** |
| ENSG00000232006.10 | **ENSG00000232006** | | lncRNA | **11.846** | 1.653 | 0.498 | **3.144** | 2.21E-03 | **5.42E-03** |
| ENSG00000067182.11 | **TNFRSF1A** | 7132 | protein_coding | **279.164** | -0.520 | -0.157 | **0.697** | 2.21E-03 | **5.42E-03** |
| ENSG00000260352.2 | **SMG1-DT** | | lncRNA | **5.888** | 1.974 | 0.594 | **3.928** | 2.21E-03 | **5.43E-03** |
| ENSG00000053371.15 | **AKR7A2** | 8574 | protein_coding | **451.950** | -0.459 | -0.138 | **0.727** | 2.22E-03 | **5.44E-03** |
| ENSG00000075043.23 | **KCNQ2** | 3785 | protein_coding | **104.886** | 1.731 | 0.521 | **3.319** | 2.22E-03 | **5.44E-03** |
| ENSG00000115556.16 | **PLCD4** | 84812 | protein_coding | **10.906** | 1.808 | 0.544 | **3.503** | 2.22E-03 | **5.45E-03** |
| ENSG00000004848.9 | **ARX** | 170302 | protein_coding | **4.896** | 2.584 | 0.778 | **5.997** | 2.22E-03 | **5.45E-03** |
| ENSG00000203879.14 | **GDI1** | 2664 | protein_coding | **3057.146** | 0.336 | 0.101 | **1.262** | 2.22E-03 | **5.45E-03** |
| ENSG00000289045.1 | **ENSG00000289045** | | lncRNA | **12.936** | 1.504 | 0.453 | **2.836** | 2.22E-03 | **5.45E-03** |

| | | | | | | | | | |
|---|---|---|---|---|---|---|---|---|---|
| ENSG00000080815.22 | **PSEN1** | 5663 | protein_coding | **957.419** | 0.385 | 0.116 | **1.305** | 2.23E-03 | **5.46E-03** |
| ENSG00000163466.18 | **ARPC2** | 10109 | protein_coding | **4598.203** | -0.394 | -0.119 | **0.761** | 2.24E-03 | **5.49E-03** |
| ENSG00000171970.16 | **ZNF57** | 126295 | protein_coding | **123.278** | 0.552 | 0.166 | **1.466** | 2.24E-03 | **5.50E-03** |
| ENSG00000172831.16 | **CES2** | 8824 | protein_coding | **457.777** | -0.300 | -0.090 | **0.812** | 2.24E-03 | **5.50E-03** |
| ENSG00000077238.16 | **IL4R** | 3566 | protein_coding | **1010.607** | 0.419 | 0.126 | **1.337** | 2.25E-03 | **5.51E-03** |
| ENSG00000177425.13 | **PAWR** | 5074 | protein_coding | **345.815** | -0.466 | -0.140 | **0.724** | 2.25E-03 | **5.52E-03** |
| ENSG00000085831.18 | **TTC39A** | 22996 | protein_coding | **37.118** | 0.806 | 0.243 | **1.748** | 2.26E-03 | **5.54E-03** |
| ENSG00000120733.17 | **KDM3B** | 51780 | protein_coding | **968.950** | -0.384 | -0.116 | **0.766** | 2.26E-03 | **5.55E-03** |
| ENSG00000178623.15 | **GPR35** | 2859 | protein_coding | **166.086** | 0.642 | 0.193 | **1.560** | 2.27E-03 | **5.56E-03** |
| ENSG00000104613.14 | **INTS10** | 55174 | protein_coding | **541.032** | -0.461 | -0.139 | **0.726** | 2.27E-03 | **5.56E-03** |
| ENSG00000288704.2 | **ENSG00000288704** | | lncRNA | **15.061** | 1.361 | 0.410 | **2.569** | 2.27E-03 | **5.56E-03** |
| ENSG00000174437.20 | **ATP2A2** | 488 | protein_coding | **1274.200** | -0.510 | -0.153 | **0.702** | 2.27E-03 | **5.57E-03** |
| ENSG00000122033.17 | **MTIF3** | 219402 | protein_coding | **625.571** | 0.321 | 0.097 | **1.250** | 2.28E-03 | **5.58E-03** |
| ENSG00000204271.14 | **SPIN3** | 169981 | protein_coding | **315.332** | 0.321 | 0.097 | **1.249** | 2.28E-03 | **5.58E-03** |
| ENSG00000156471.15 | **PTDSS1** | 9791 | protein_coding | **1253.955** | -0.341 | -0.103 | **0.790** | 2.28E-03 | **5.58E-03** |
| ENSG00000112293.17 | **GPLD1** | 2822 | protein_coding | **34.061** | 0.861 | 0.259 | **1.816** | 2.28E-03 | **5.58E-03** |
| ENSG00000177873.16 | **ZNF619** | 285267 | protein_coding | **76.356** | -0.651 | -0.196 | **0.637** | 2.29E-03 | **5.61E-03** |
| ENSG00000132485.16 | **ZRANB2** | 9406 | protein_coding | **961.358** | -0.374 | -0.112 | **0.772** | 2.29E-03 | **5.61E-03** |
| ENSG00000164307.15 | **ERAP1** | 51752 | protein_coding | **655.059** | -0.601 | -0.181 | **0.659** | 2.29E-03 | **5.61E-03** |
| ENSG00000273987.1 | **ENSG00000273987** | | lncRNA | **18.093** | 1.125 | 0.339 | **2.181** | 2.29E-03 | **5.61E-03** |
| ENSG00000092964.19 | **DPYSL2** | 1808 | protein_coding | **474.940** | 0.620 | 0.187 | **1.537** | 2.30E-03 | **5.61E-03** |
| ENSG00000135632.14 | **SMYD5** | 10322 | protein_coding | **519.171** | -0.355 | -0.107 | **0.782** | 2.30E-03 | **5.61E-03** |
| ENSG00000115112.10 | **TFCP2L1** | 29842 | protein_coding | **10.654** | -1.563 | -0.471 | **0.338** | 2.30E-03 | **5.62E-03** |
| ENSG00000166002.9 | **SMCO4** | 56935 | protein_coding | **389.961** | -0.456 | -0.137 | **0.729** | 2.30E-03 | **5.62E-03** |
| ENSG00000205307.13 | **SAP25** | 10031690 | protein_coding | **495.839** | 0.570 | 0.172 | **1.485** | 2.30E-03 | **5.62E-03** |
| ENSG00000296546.1 | **ENSG0000(** | 12490701 | lncRNA | **5.596** | 2.431 | 0.732 | **5.393** | 2.30E-03 | **5.62E-03** |
| ENSG00000081479.16 | **LRP2** | 4036 | protein_coding | **16.731** | 1.814 | 0.546 | **3.516** | 2.31E-03 | **5.66E-03** |
| ENSG00000216863.12 | **LY86-AS1** | 285780 | lncRNA | **548.126** | -0.351 | -0.106 | **0.784** | 2.32E-03 | **5.66E-03** |
| ENSG00000133131.17 | **MORC4** | 79710 | protein_coding | **19.371** | 1.325 | 0.399 | **2.506** | 2.32E-03 | **5.66E-03** |
| ENSG00000139675.13 | **HNRNPA1L** | 144983 | protein_coding | **280.163** | 0.504 | 0.152 | **1.418** | 2.32E-03 | **5.67E-03** |
| ENSG00000260911.2 | **ENSG00000260911** | | lncRNA | **7.179** | 1.884 | 0.567 | **3.692** | 2.32E-03 | **5.68E-03** |

| | | | | | | | | | |
|---|---|---|---|---|---|---|---|---|---|
| ENSG00000152804.13 | **HHEX** | 3087 | protein_coding | **22.514** | 1.265 | 0.381 | **2.404** | 2.33E-03 | **5.68E-03** |
| ENSG00000257067.7 | **LINC02703** | 10272416 | lncRNA | **9.831** | 1.645 | 0.495 | **3.128** | 2.33E-03 | **5.70E-03** |
| ENSG00000107719.12 | **PALD1** | 27143 | protein_coding | **411.344** | 0.478 | 0.144 | **1.393** | 2.33E-03 | **5.70E-03** |
| ENSG00000230373.9 | **GOLGA6L5** | 10013322 | transcribed_unpro | **17.735** | 1.401 | 0.422 | **2.642** | 2.33E-03 | **5.70E-03** |
| ENSG00000147003.9 | **CLTRN** | 57393 | protein_coding | **8.490** | 1.584 | 0.477 | **2.998** | 2.34E-03 | **5.71E-03** |
| ENSG00000173402.15 | **DAG1** | 1605 | protein_coding | **303.110** | 0.476 | 0.143 | **1.391** | 2.34E-03 | **5.71E-03** |
| ENSG00000011638.13 | **LDAF1** | 57146 | protein_coding | **203.325** | 0.390 | 0.118 | **1.311** | 2.34E-03 | **5.71E-03** |
| ENSG00000303385.1 | **ENSG00000303385** | | lncRNA | **18.491** | 1.239 | 0.373 | **2.361** | 2.34E-03 | **5.71E-03** |
| ENSG00000137992.17 | **DBT** | 1629 | protein_coding | **229.546** | -0.393 | -0.118 | **0.762** | 2.34E-03 | **5.71E-03** |
| ENSG00000259673.7 | **IQCH-AS1** | 10050668 | lncRNA | **120.881** | -0.508 | -0.153 | **0.703** | 2.34E-03 | **5.72E-03** |
| ENSG00000164332.10 | **UBLCP1** | 134510 | protein_coding | **239.587** | 0.530 | 0.160 | **1.444** | 2.35E-03 | **5.72E-03** |
| ENSG00000309183.1 | **ENSG00000309183** | | lncRNA | **4.936** | 2.241 | 0.675 | **4.726** | 2.35E-03 | **5.73E-03** |
| ENSG00000127804.14 | **METTL16** | 79066 | protein_coding | **561.110** | -0.415 | -0.125 | **0.750** | 2.35E-03 | **5.74E-03** |
| ENSG00000101407.15 | **TTI1** | 9675 | protein_coding | **495.724** | -0.410 | -0.124 | **0.752** | 2.35E-03 | **5.74E-03** |
| ENSG00000176919.15 | **C8G** | 733 | protein_coding | **15.325** | 1.338 | 0.403 | **2.528** | 2.35E-03 | **5.74E-03** |
| ENSG00000101343.17 | **CRNKL1** | 51340 | protein_coding | **294.280** | -0.435 | -0.131 | **0.740** | 2.36E-03 | **5.74E-03** |
| ENSG00000186951.19 | **PPARA** | 5465 | protein_coding | **300.922** | -0.363 | -0.109 | **0.778** | 2.36E-03 | **5.75E-03** |
| ENSG00000172893.20 | **DHCR7** | 1717 | protein_coding | **1137.090** | -0.652 | -0.196 | **0.637** | 2.36E-03 | **5.76E-03** |
| ENSG00000163684.13 | **RPP14** | 11102 | protein_coding | **103.584** | -0.952 | -0.287 | **0.517** | 2.37E-03 | **5.77E-03** |
| ENSG00000162078.14 | **ZG16B** | 124220 | protein_coding | **49.104** | -0.705 | -0.212 | **0.614** | 2.37E-03 | **5.77E-03** |
| ENSG00000166323.15 | **C11orf65** | 160140 | protein_coding | **19.562** | 0.981 | 0.295 | **1.974** | 2.37E-03 | **5.77E-03** |
| ENSG00000246095.3 | **LINC01096** | | lncRNA | **4.502** | 2.854 | 0.859 | **7.229** | 2.38E-03 | **5.79E-03** |
| ENSG00000144648.18 | **ACKR2** | 1238 | protein_coding | **12.507** | 1.232 | 0.371 | **2.349** | 2.38E-03 | **5.79E-03** |
| ENSG00000164542.15 | **MATCAP2** | 23366 | protein_coding | **5.231** | 2.180 | 0.656 | **4.530** | 2.38E-03 | **5.81E-03** |
| ENSG00000274333.5 | **LINC03104** | 10272421 | lncRNA | **190.201** | -0.991 | -0.298 | **0.503** | 2.39E-03 | **5.82E-03** |
| ENSG00000261147.1 | **ENSG00000261147** | | protein_coding | **9.732** | 2.232 | 0.672 | **4.698** | 2.39E-03 | **5.82E-03** |
| ENSG00000284693.3 | **LINC02606** | 10050602 | lncRNA | **12.649** | 1.481 | 0.446 | **2.791** | 2.39E-03 | **5.82E-03** |
| ENSG00000174567.10 | **GOLT1A** | 127845 | protein_coding | **10.615** | 1.746 | 0.526 | **3.354** | 2.39E-03 | **5.82E-03** |
| ENSG00000105325.17 | **FZR1** | 51343 | protein_coding | **663.667** | -0.326 | -0.098 | **0.798** | 2.39E-03 | **5.82E-03** |
| ENSG00000112378.13 | **PERP** | 64065 | protein_coding | **53.696** | 0.812 | 0.244 | **1.755** | 2.40E-03 | **5.83E-03** |
| ENSG00000305205.1 | **FAM88D** | | lncRNA | **22.142** | -1.371 | -0.413 | **0.387** | 2.40E-03 | **5.84E-03** |

| | | | | | | | | | |
|---|---|---|---|---|---|---|---|---|---|
| ENSG00000270177.2 | **PPP2CA-DT** | | lncRNA | **9.963** | 1.290 | 0.388 | **2.445** | 2.40E-03 | **5.84E-03** |
| ENSG00000187189.11 | **TSPYL4** | 23270 | protein_coding | **231.463** | 0.671 | 0.202 | **1.592** | 2.41E-03 | **5.85E-03** |
| ENSG00000075240.19 | **GRAMD4** | 23151 | protein_coding | **294.814** | 0.519 | 0.156 | **1.433** | 2.42E-03 | **5.88E-03** |
| ENSG00000125826.24 | **RBCK1** | 10616 | protein_coding | **3728.276** | -0.359 | -0.108 | **0.780** | 2.42E-03 | **5.88E-03** |
| ENSG00000163531.19 | **NFASC** | 23114 | protein_coding | **5.694** | 2.022 | 0.609 | **4.063** | 2.42E-03 | **5.88E-03** |
| ENSG00000166562.12 | **SEC11C** | 90701 | protein_coding | **1506.841** | -0.313 | -0.094 | **0.805** | 2.42E-03 | **5.88E-03** |
| ENSG00000272156.1 | **ENSG00000272156** | | lncRNA | **10.628** | 1.302 | 0.392 | **2.466** | 2.42E-03 | **5.88E-03** |
| ENSG00000272779.1 | **BMS1P20** | | transcribed_unprc | **125.868** | -0.520 | -0.157 | **0.697** | 2.42E-03 | **5.88E-03** |
| ENSG00000160392.16 | **C19orf47** | 126526 | protein_coding | **245.743** | -0.390 | -0.117 | **0.763** | 2.43E-03 | **5.90E-03** |
| ENSG00000072134.18 | **EPN2** | 22905 | protein_coding | **176.452** | 0.503 | 0.151 | **1.417** | 2.43E-03 | **5.91E-03** |
| ENSG00000069248.14 | **NUP133** | 55746 | protein_coding | **422.257** | -0.300 | -0.090 | **0.812** | 2.43E-03 | **5.91E-03** |
| ENSG00000197429.13 | **IPP** | 3652 | protein_coding | **142.051** | -0.475 | -0.143 | **0.720** | 2.43E-03 | **5.91E-03** |
| ENSG00000242125.3 | **SNHG3** | 8420 | lncRNA | **1365.330** | -0.510 | -0.154 | **0.702** | 2.44E-03 | **5.93E-03** |
| ENSG00000236404.13 | **VLDLR-AS1** | 401491 | lncRNA | **30.402** | -1.162 | -0.350 | **0.447** | 2.44E-03 | **5.93E-03** |
| ENSG00000197888.5 | **UGT2B17** | 7367 | protein_coding | **261.822** | -0.603 | -0.182 | **0.658** | 2.44E-03 | **5.93E-03** |
| ENSG00000178567.8 | **EPM2AIP1** | 9852 | protein_coding | **579.939** | -0.325 | -0.098 | **0.798** | 2.45E-03 | **5.94E-03** |
| ENSG00000185880.16 | **TRIM69** | 140691 | protein_coding | **2095.627** | -0.386 | -0.116 | **0.765** | 2.46E-03 | **5.98E-03** |
| ENSG00000102100.18 | **SLC35A2** | 7355 | protein_coding | **335.929** | 0.387 | 0.116 | **1.307** | 2.47E-03 | **5.99E-03** |
| ENSG00000140443.15 | **IGF1R** | 3480 | protein_coding | **102.015** | 0.780 | 0.235 | **1.717** | 2.47E-03 | **6.00E-03** |
| ENSG00000143248.15 | **RGS5** | 8490 | protein_coding | **33.002** | -1.600 | -0.482 | **0.330** | 2.47E-03 | **6.00E-03** |
| ENSG00000298681.1 | **ENSG00000298681** | | lncRNA | **6.392** | 1.860 | 0.560 | **3.631** | 2.48E-03 | **6.02E-03** |
| ENSG00000137055.17 | **PLAA** | 9373 | protein_coding | **423.225** | -0.415 | -0.125 | **0.750** | 2.48E-03 | **6.02E-03** |
| ENSG00000066136.23 | **NFYC** | 4802 | protein_coding | **1373.236** | -0.381 | -0.115 | **0.768** | 2.48E-03 | **6.03E-03** |
| ENSG00000293315.2 | **ENSG00000293315** | | lncRNA | **15.207** | 1.466 | 0.441 | **2.763** | 2.48E-03 | **6.03E-03** |
| ENSG00000300379.1 | **ENSG00000300379** | | lncRNA | **25.886** | 0.953 | 0.287 | **1.936** | 2.49E-03 | **6.04E-03** |
| ENSG00000104687.16 | **GSR** | 2936 | protein_coding | **870.810** | -0.281 | -0.085 | **0.823** | 2.49E-03 | **6.05E-03** |
| ENSG00000104957.16 | **YJU2B** | 81576 | protein_coding | **1018.434** | 0.404 | 0.122 | **1.323** | 2.50E-03 | **6.06E-03** |
| ENSG00000188626.7 | **GOLGA8M** | 653720 | protein_coding | **7.037** | 2.522 | 0.759 | **5.745** | 2.50E-03 | **6.06E-03** |
| ENSG00000233456.2 | **LINC01077** | | lncRNA | **15.438** | 1.547 | 0.466 | **2.922** | 2.50E-03 | **6.06E-03** |
| ENSG00000131981.18 | **LGALS3** | 3958 | protein_coding | **1799.292** | -0.427 | -0.129 | **0.744** | 2.51E-03 | **6.07E-03** |
| ENSG00000253293.7 | **HOXA10** | 3206 | protein_coding | **8.106** | 1.713 | 0.516 | **3.279** | 2.51E-03 | **6.07E-03** |

| | | | | | | | | | |
|---|---|---|---|---|---|---|---|---|---|
| ENSG00000160013.12 | **PTGIR** | 5739 | protein_coding | **319.161** | 0.594 | 0.179 | **1.509** | 2.51E-03 | **6.09E-03** |
| ENSG00000151388.12 | **ADAMTS1** | 81792 | protein_coding | **172.741** | 0.687 | 0.207 | **1.610** | 2.51E-03 | **6.09E-03** |
| ENSG00000124786.15 | **SLC35B3** | 51000 | protein_coding | **273.331** | -0.428 | -0.129 | **0.743** | 2.52E-03 | **6.09E-03** |
| ENSG00000132623.18 | **ANKEF1** | 63926 | protein_coding | **30.465** | 0.835 | 0.251 | **1.784** | 2.52E-03 | **6.11E-03** |
| ENSG00000138083.6 | **SIX3** | 6496 | protein_coding | **140.824** | 1.043 | 0.314 | **2.060** | 2.53E-03 | **6.12E-03** |
| ENSG00000132507.20 | **EIF5A** | 1984 | protein_coding | **12177.992** | -0.365 | -0.110 | **0.776** | 2.53E-03 | **6.12E-03** |
| ENSG00000188785.14 | **ZNF548** | 147694 | protein_coding | **394.373** | -0.380 | -0.114 | **0.769** | 2.53E-03 | **6.13E-03** |
| ENSG00000243819.4 | **RN7SL832P** | | lncRNA | **23.048** | 0.895 | 0.269 | **1.860** | 2.54E-03 | **6.14E-03** |
| ENSG00000084207.20 | **GSTP1** | 2950 | protein_coding | **11507.910** | 0.398 | 0.120 | **1.318** | 2.54E-03 | **6.14E-03** |
| ENSG00000180787.8 | **ZFP3** | 124961 | protein_coding | **132.323** | 0.509 | 0.153 | **1.423** | 2.54E-03 | **6.15E-03** |
| ENSG00000180626.12 | **ZNF594** | 84622 | protein_coding | **309.864** | 0.478 | 0.144 | **1.392** | 2.54E-03 | **6.15E-03** |
| ENSG00000141349.12 | **G6PC3** | 92579 | protein_coding | **652.902** | -0.472 | -0.142 | **0.721** | 2.54E-03 | **6.16E-03** |
| ENSG00000166147.17 | **FBN1** | 2200 | protein_coding | **286.501** | 0.601 | 0.181 | **1.517** | 2.55E-03 | **6.17E-03** |
| ENSG00000172380.8 | **GNG12** | 55970 | protein_coding | **11.336** | 1.861 | 0.560 | **3.633** | 2.55E-03 | **6.17E-03** |
| ENSG00000107537.16 | **PHYH** | 5264 | protein_coding | **146.553** | 0.472 | 0.142 | **1.387** | 2.56E-03 | **6.18E-03** |
| ENSG00000158874.14 | **APOA2** | 336 | protein_coding | **9.035** | 3.219 | 0.969 | **9.311** | 2.56E-03 | **6.18E-03** |
| ENSG00000181798.5 | **LINC00471** | 151477 | lncRNA | **28.954** | -0.916 | -0.276 | **0.530** | 2.57E-03 | **6.22E-03** |
| ENSG00000123360.14 | **PDE1B** | 5153 | protein_coding | **353.072** | 0.362 | 0.109 | **1.285** | 2.57E-03 | **6.22E-03** |
| ENSG00000297660.1 | **ENSG00000297660** | | lncRNA | **13.881** | 1.586 | 0.477 | **3.002** | 2.58E-03 | **6.23E-03** |
| ENSG00000131043.15 | **AAR2** | 25980 | protein_coding | **436.393** | -0.317 | -0.095 | **0.803** | 2.58E-03 | **6.24E-03** |
| ENSG00000257660.6 | **ADCY6-DT** | 10050612 | lncRNA | **6.624** | 2.050 | 0.617 | **4.142** | 2.59E-03 | **6.25E-03** |
| ENSG00000115596.5 | **WNT6** | 7475 | protein_coding | **75.314** | 0.949 | 0.286 | **1.930** | 2.59E-03 | **6.25E-03** |
| ENSG00000286592.3 | **LINC02885** | | lncRNA | **9.818** | 2.075 | 0.625 | **4.213** | 2.59E-03 | **6.25E-03** |
| ENSG00000255267.5 | **LINC02687** | 399886 | lncRNA | **6.122** | 2.313 | 0.696 | **4.970** | 2.59E-03 | **6.26E-03** |
| ENSG00000188223.9 | **ENSG00000188223** | | protein_coding | **23.187** | 8.950 | 2.694 | **494.610** | 2.61E-03 | **6.29E-03** |
| ENSG00000077549.21 | **CAPZB** | 832 | protein_coding | **6486.643** | 0.306 | 0.092 | **1.237** | 2.61E-03 | **6.31E-03** |
| ENSG00000103264.20 | **FBXO31** | 79791 | protein_coding | **699.505** | 0.336 | 0.101 | **1.262** | 2.61E-03 | **6.31E-03** |
| ENSG00000196914.11 | **ARHGEF12** | 23365 | protein_coding | **446.294** | 0.455 | 0.137 | **1.371** | 2.62E-03 | **6.31E-03** |
| ENSG00000151651.18 | **ADAM8** | 101 | protein_coding | **2300.412** | 0.539 | 0.162 | **1.453** | 2.62E-03 | **6.32E-03** |
| ENSG00000303041.1 | **ENSG00000303041** | | lncRNA | **13.066** | 1.265 | 0.381 | **2.403** | 2.62E-03 | **6.32E-03** |
| ENSG00000140374.19 | **ETFA** | 2108 | protein_coding | **1341.963** | -0.440 | -0.133 | **0.737** | 2.62E-03 | **6.33E-03** |

| | | | | | | | | | |
|---|---|---|---|---|---|---|---|---|---|
| ENSG00000128203.9 | **ASPHD2** | 57168 | protein_coding | **261.060** | 0.519 | 0.156 | **1.433** | 2.63E-03 | **6.34E-03** |
| ENSG00000197969.15 | **VPS13A** | 23230 | protein_coding | **554.217** | -0.396 | -0.119 | **0.760** | 2.63E-03 | **6.35E-03** |
| ENSG00000146842.19 | **TMEM209** | 84928 | protein_coding | **189.870** | -0.490 | -0.148 | **0.712** | 2.63E-03 | **6.35E-03** |
| ENSG00000205740.3 | **ENSG00000205740** | | lncRNA | **25.491** | 1.151 | 0.347 | **2.221** | 2.64E-03 | **6.38E-03** |
| ENSG00000119878.9 | **CRIPT** | 9419 | protein_coding | **211.263** | -0.430 | -0.129 | **0.742** | 2.65E-03 | **6.38E-03** |
| ENSG00000005483.24 | **KMT2E** | 55904 | protein_coding | **1053.923** | 0.335 | 0.101 | **1.262** | 2.65E-03 | **6.38E-03** |
| ENSG00000130520.13 | **LSM4** | 25804 | protein_coding | **2331.795** | -0.360 | -0.108 | **0.779** | 2.65E-03 | **6.39E-03** |
| ENSG00000145358.8 | **DDIT4L** | 115265 | protein_coding | **11.201** | 1.600 | 0.482 | **3.032** | 2.66E-03 | **6.40E-03** |
| ENSG00000155304.8 | **HSPA13** | 6782 | protein_coding | **170.783** | -0.592 | -0.178 | **0.663** | 2.66E-03 | **6.42E-03** |
| ENSG00000173171.16 | **MTX1** | 4580 | protein_coding | **585.716** | -0.387 | -0.116 | **0.765** | 2.66E-03 | **6.42E-03** |
| ENSG00000269680.1 | **TRIP10-AS1** | | lncRNA | **9.964** | 1.302 | 0.392 | **2.467** | 2.66E-03 | **6.42E-03** |
| ENSG00000110367.16 | **DDX6** | 1656 | protein_coding | **1681.347** | -0.305 | -0.092 | **0.809** | 2.66E-03 | **6.42E-03** |
| ENSG00000075568.19 | **TMEM131** | 23505 | protein_coding | **941.265** | 0.435 | 0.131 | **1.352** | 2.67E-03 | **6.42E-03** |
| ENSG00000129675.18 | **ARHGEF6** | 9459 | protein_coding | **441.153** | -0.435 | -0.131 | **0.740** | 2.67E-03 | **6.42E-03** |
| ENSG00000102886.16 | **GDPD3** | 79153 | protein_coding | **186.326** | 0.602 | 0.181 | **1.518** | 2.67E-03 | **6.43E-03** |
| ENSG00000285413.2 | **ENSG00000285413** | | lncRNA | **11.739** | 1.998 | 0.602 | **3.996** | 2.68E-03 | **6.45E-03** |
| ENSG00000143013.15 | **LMO4** | 8543 | protein_coding | **793.966** | -0.582 | -0.175 | **0.668** | 2.68E-03 | **6.45E-03** |
| ENSG00000125735.14 | **TNFSF14** | 8740 | protein_coding | **716.290** | 0.461 | 0.139 | **1.377** | 2.68E-03 | **6.45E-03** |
| ENSG00000271869.3 | **DCTN6-DT** | 118597835 | lncRNA | **30.554** | 0.838 | 0.252 | **1.787** | 2.68E-03 | **6.45E-03** |
| ENSG00000275835.7 | **TUBGCP5** | 114791 | protein_coding | **229.400** | -0.476 | -0.143 | **0.719** | 2.68E-03 | **6.46E-03** |
| ENSG00000130208.12 | **APOC1** | 341 | protein_coding | **53.231** | -0.991 | -0.298 | **0.503** | 2.68E-03 | **6.46E-03** |
| ENSG00000182359.17 | **KBTBD3** | 143879 | protein_coding | **66.975** | 0.824 | 0.248 | **1.770** | 2.69E-03 | **6.46E-03** |
| ENSG00000174791.13 | **RIN1** | 9610 | protein_coding | **40.504** | 0.824 | 0.248 | **1.770** | 2.69E-03 | **6.47E-03** |
| ENSG00000140465.17 | **CYP1A1** | 1543 | protein_coding | **115.365** | 0.891 | 0.268 | **1.855** | 2.69E-03 | **6.47E-03** |
| ENSG00000166164.18 | **BRD7** | 29117 | protein_coding | **1018.702** | -0.338 | -0.102 | **0.791** | 2.70E-03 | **6.49E-03** |
| ENSG00000231473.6 | **RB1-DT** | | lncRNA | **13.150** | 1.306 | 0.393 | **2.472** | 2.70E-03 | **6.49E-03** |
| ENSG00000224565.3 | **LINC01754** | | lncRNA | **85.219** | -0.776 | -0.234 | **0.584** | 2.71E-03 | **6.51E-03** |
| ENSG00000100422.16 | **CERK** | 64781 | protein_coding | **638.243** | 0.467 | 0.140 | **1.382** | 2.71E-03 | **6.52E-03** |
| ENSG00000145936.11 | **KCNMB1** | 3779 | protein_coding | **21.216** | -1.479 | -0.445 | **0.359** | 2.72E-03 | **6.53E-03** |
| ENSG00000230578.4 | **TMLHEP1** | | transcribed_unprc | **5.184** | -2.807 | -0.845 | **0.143** | 2.72E-03 | **6.53E-03** |
| ENSG00000135643.6 | **KCNMB4** | 27345 | protein_coding | **29.997** | 1.045 | 0.314 | **2.063** | 2.72E-03 | **6.53E-03** |

| | | | | | | | | | |
|---|---|---|---|---|---|---|---|---|---|
| ENSG00000145860.14 | **RNF145** | 153830 | protein_coding | **624.664** | -0.397 | -0.120 | **0.759** | 2.72E-03 | **6.55E-03** |
| ENSG00000231856.5 | **ENSG00000231856** | | lncRNA | **29.374** | 0.934 | 0.281 | **1.910** | 2.74E-03 | **6.57E-03** |
| ENSG00000115977.23 | **AAK1** | 22848 | protein_coding | **471.613** | 0.350 | 0.105 | **1.274** | 2.74E-03 | **6.59E-03** |
| ENSG00000113494.19 | **PRLR** | 5618 | protein_coding | **29.793** | -1.066 | -0.321 | **0.478** | 2.75E-03 | **6.60E-03** |
| ENSG00000168765.21 | **GSTM4** | 2948 | protein_coding | **605.036** | 0.318 | 0.096 | **1.247** | 2.76E-03 | **6.62E-03** |
| ENSG00000122490.21 | **SLC66A2** | 80148 | protein_coding | **805.317** | 0.371 | 0.112 | **1.293** | 2.76E-03 | **6.62E-03** |
| ENSG00000121905.12 | **HPCA** | 3208 | protein_coding | **9.477** | 2.111 | 0.635 | **4.320** | 2.76E-03 | **6.62E-03** |
| ENSG00000182791.5 | **CCDC87** | 55231 | protein_coding | **6.519** | 1.680 | 0.506 | **3.204** | 2.76E-03 | **6.62E-03** |
| ENSG00000171224.11 | **FAM241B** | 219738 | protein_coding | **16.036** | 1.098 | 0.330 | **2.140** | 2.76E-03 | **6.62E-03** |
| ENSG00000196345.15 | **ZKSCAN7** | 55888 | protein_coding | **82.570** | 0.591 | 0.178 | **1.506** | 2.76E-03 | **6.64E-03** |
| ENSG00000124615.23 | **MOCS1** | 4337 | protein_coding | **35.064** | 1.116 | 0.336 | **2.167** | 2.77E-03 | **6.64E-03** |
| ENSG00000110429.16 | **FBXO3** | 26273 | protein_coding | **368.599** | 0.507 | 0.153 | **1.422** | 2.77E-03 | **6.66E-03** |
| ENSG00000290796.1 | **ENSG0000(** | 91355 | lncRNA | **412.147** | 0.479 | 0.144 | **1.394** | 2.77E-03 | **6.66E-03** |
| ENSG00000160563.16 | **MED27** | 9442 | protein_coding | **291.462** | -0.346 | -0.104 | **0.787** | 2.79E-03 | **6.69E-03** |
| ENSG00000229869.2 | **LARP4B-DT** | 10192776 | lncRNA | **27.998** | 0.929 | 0.280 | **1.904** | 2.82E-03 | **6.75E-03** |
| ENSG00000159200.19 | **RCAN1** | 1827 | protein_coding | **156.256** | 0.521 | 0.157 | **1.435** | 2.82E-03 | **6.75E-03** |
| ENSG00000172590.20 | **MRPL52** | 122704 | protein_coding | **1105.178** | -0.343 | -0.103 | **0.788** | 2.82E-03 | **6.76E-03** |
| ENSG00000154832.18 | **CXXC1** | 30827 | protein_coding | **2366.973** | -0.380 | -0.114 | **0.769** | 2.82E-03 | **6.76E-03** |
| ENSG00000187815.12 | **ZFP69** | 339559 | protein_coding | **106.519** | -0.592 | -0.178 | **0.663** | 2.83E-03 | **6.77E-03** |
| ENSG00000247675.9 | **LRP4-AS1** | 100507401 | lncRNA | **7.711** | 1.644 | 0.495 | **3.125** | 2.83E-03 | **6.78E-03** |
| ENSG00000298139.1 | **ENSG00000298139** | | lncRNA | **8.215** | 1.917 | 0.577 | **3.775** | 2.83E-03 | **6.78E-03** |
| ENSG00000284624.2 | **ENSG00000284624** | | lncRNA | **8.468** | 1.716 | 0.516 | **3.284** | 2.83E-03 | **6.78E-03** |
| ENSG00000250490.1 | **LINC02145** | 401172 | lncRNA | **6.032** | 2.256 | 0.679 | **4.776** | 2.83E-03 | **6.79E-03** |
| ENSG00000213593.13 | **TMX2** | 51075 | protein_coding | **868.566** | -0.282 | -0.085 | **0.822** | 2.84E-03 | **6.81E-03** |
| ENSG00000077009.16 | **NMRK2** | 27231 | protein_coding | **5.091** | -2.826 | -0.851 | **0.141** | 2.84E-03 | **6.81E-03** |
| ENSG00000164494.14 | **PDSS2** | 57107 | protein_coding | **130.573** | -0.472 | -0.142 | **0.721** | 2.85E-03 | **6.82E-03** |
| ENSG00000263620.1 | **ENSG00000263620** | | protein_coding | **13.766** | -1.327 | -0.399 | **0.399** | 2.85E-03 | **6.82E-03** |
| ENSG00000237187.10 | **NR2F1-AS1** | 441094 | lncRNA | **11.929** | 1.737 | 0.523 | **3.332** | 2.85E-03 | **6.83E-03** |
| ENSG00000105393.18 | **BABAM1** | 29086 | protein_coding | **1473.706** | -0.273 | -0.082 | **0.828** | 2.86E-03 | **6.84E-03** |
| ENSG00000137507.13 | **LRRC32** | 2615 | protein_coding | **46.887** | -1.115 | -0.336 | **0.462** | 2.86E-03 | **6.85E-03** |
| ENSG00000134490.16 | **SLC35D4** | 85019 | protein_coding | **209.363** | 0.452 | 0.136 | **1.368** | 2.86E-03 | **6.85E-03** |

| | | | | | | | | | |
|---|---|---|---|---|---|---|---|---|---|
| ENSG00000023318.11 | **ERP44** | 23071 | protein_coding | **1925.021** | -0.333 | -0.100 | **0.794** | 2.86E-03 | **6.86E-03** |
| ENSG00000221994.14 | **ZNF630** | 57232 | protein_coding | **57.261** | -0.566 | -0.171 | **0.675** | 2.87E-03 | **6.86E-03** |
| ENSG00000149636.18 | **DSN1** | 79980 | protein_coding | **356.422** | -0.367 | -0.110 | **0.775** | 2.87E-03 | **6.86E-03** |
| ENSG00000286369.3 | **ENSG0000(** | 10536936: | lncRNA | **68.367** | -0.642 | -0.193 | **0.641** | 2.87E-03 | **6.87E-03** |
| ENSG00000101152.14 | **DNAJC5** | 80331 | protein_coding | **1351.604** | 0.297 | 0.089 | **1.228** | 2.87E-03 | **6.87E-03** |
| ENSG00000166189.8 | **HPS6** | 79803 | protein_coding | **330.088** | -0.337 | -0.101 | **0.792** | 2.87E-03 | **6.87E-03** |
| ENSG00000267387.1 | **ENSG00000267387** | | lncRNA | **43.565** | -0.707 | -0.213 | **0.613** | 2.87E-03 | **6.87E-03** |
| ENSG00000189369.9 | **GSPT2** | 23708 | protein_coding | **134.053** | -0.511 | -0.154 | **0.702** | 2.88E-03 | **6.89E-03** |
| ENSG00000096070.21 | **BRPF3** | 27154 | protein_coding | **158.199** | -0.524 | -0.158 | **0.696** | 2.89E-03 | **6.90E-03** |
| ENSG00000205978.8 | **NYNRIN** | 57523 | protein_coding | **9.488** | 1.731 | 0.521 | **3.321** | 2.89E-03 | **6.91E-03** |
| ENSG00000157911.13 | **PEX10** | 5192 | protein_coding | **422.260** | -0.385 | -0.116 | **0.766** | 2.89E-03 | **6.91E-03** |
| ENSG00000286883.1 | **ENSG00000286883** | | lncRNA | **13.062** | 1.294 | 0.389 | **2.451** | 2.89E-03 | **6.91E-03** |
| ENSG00000123595.10 | **RAB9A** | 9367 | protein_coding | **1173.728** | 0.385 | 0.116 | **1.306** | 2.90E-03 | **6.92E-03** |
| ENSG00000145012.17 | **LPP** | 4026 | protein_coding | **38.470** | 0.881 | 0.265 | **1.842** | 2.90E-03 | **6.92E-03** |
| ENSG00000132953.20 | **XPO4** | 64328 | protein_coding | **340.344** | -0.422 | -0.127 | **0.746** | 2.90E-03 | **6.93E-03** |
| ENSG00000187144.12 | **SPATA21** | 374955 | protein_coding | **12.910** | 1.627 | 0.490 | **3.088** | 2.91E-03 | **6.95E-03** |
| ENSG00000228397.4 | **LINC01635** | 10192804: | lncRNA | **14.961** | 1.629 | 0.490 | **3.092** | 2.91E-03 | **6.95E-03** |
| ENSG00000108861.11 | **DUSP3** | 1845 | protein_coding | **278.325** | -0.499 | -0.150 | **0.708** | 2.92E-03 | **6.96E-03** |
| ENSG00000290945.2 | **ENSG00000290945** | | lncRNA | **54.805** | 0.749 | 0.226 | **1.681** | 2.92E-03 | **6.96E-03** |
| ENSG00000032742.20 | **IFT88** | 8100 | protein_coding | **114.461** | 0.577 | 0.174 | **1.492** | 2.92E-03 | **6.97E-03** |
| ENSG00000169249.16 | **ZRSR2** | 8233 | protein_coding | **885.500** | -0.479 | -0.144 | **0.717** | 2.92E-03 | **6.98E-03** |
| ENSG00000293484.2 | **TEX14BP** | 644050 | lncRNA | **6.436** | 2.106 | 0.634 | **4.304** | 2.93E-03 | **6.98E-03** |
| ENSG00000152684.13 | **PELO** | 53918 | protein_coding | **189.927** | -0.490 | -0.147 | **0.712** | 2.93E-03 | **6.99E-03** |
| ENSG00000226942.2 | **IL9RP3** | | transcribed_unpro | **11.159** | -1.535 | -0.462 | **0.345** | 2.94E-03 | **7.01E-03** |
| ENSG00000278970.5 | **HEIH** | 10085993( | lncRNA | **632.665** | -0.352 | -0.106 | **0.783** | 2.94E-03 | **7.01E-03** |
| ENSG00000119686.12 | **FLVCR2** | 55640 | protein_coding | **112.782** | 0.659 | 0.198 | **1.579** | 2.94E-03 | **7.01E-03** |
| ENSG00000164142.18 | **FHIP1A** | 729830 | protein_coding | **35.544** | 1.344 | 0.404 | **2.538** | 2.94E-03 | **7.01E-03** |
| ENSG00000249307.9 | **LINC01088** | | lncRNA | **5.512** | 2.534 | 0.763 | **5.793** | 2.94E-03 | **7.01E-03** |
| ENSG00000242220.11 | **TCP10L** | 140290 | protein_coding | **22.095** | 1.755 | 0.528 | **3.375** | 2.94E-03 | **7.02E-03** |
| ENSG00000286116.1 | **ENSG00000286116** | | lncRNA | **26.869** | -0.921 | -0.277 | **0.528** | 2.95E-03 | **7.03E-03** |
| ENSG00000270112.6 | **ST8SIA5-D** | 10537209 | lncRNA | **10.340** | 2.364 | 0.712 | **5.148** | 2.95E-03 | **7.03E-03** |

| | | | | | | | | | |
|---|---|---|---|---|---|---|---|---|---|
| ENSG00000163006.14 | **CCDC138** | 165055 | protein_coding | **84.098** | -0.624 | -0.188 | **0.649** | 2.95E-03 | **7.03E-03** |
| ENSG00000248092.9 | **NNT-AS1** | | lncRNA | **219.907** | -0.477 | -0.143 | **0.719** | 2.95E-03 | **7.03E-03** |
| ENSG00000188313.15 | **PLSCR1** | 5359 | protein_coding | **762.794** | -0.619 | -0.186 | **0.651** | 2.96E-03 | **7.05E-03** |
| ENSG00000307319.1 | **ENSG00000307319** | | lncRNA | **6.919** | 1.539 | 0.463 | **2.905** | 2.96E-03 | **7.05E-03** |
| ENSG00000268573.1 | **ENSG00000268573** | | lncRNA | **48.246** | -0.685 | -0.206 | **0.622** | 2.97E-03 | **7.08E-03** |
| ENSG00000003402.23 | **CFLAR** | 8837 | protein_coding | **3258.651** | 0.461 | 0.139 | **1.376** | 2.97E-03 | **7.08E-03** |
| ENSG00000293506.2 | **ENSG00000293506** | | lncRNA | **12.595** | 1.304 | 0.392 | **2.468** | 2.97E-03 | **7.08E-03** |
| ENSG00000152348.18 | **ATG10** | 83734 | protein_coding | **88.557** | -0.583 | -0.176 | **0.667** | 2.98E-03 | **7.09E-03** |
| ENSG00000266967.9 | **AARSD1** | 80755 | protein_coding | **645.603** | -0.389 | -0.117 | **0.764** | 2.98E-03 | **7.09E-03** |
| ENSG00000115274.16 | **INO80B** | 83444 | protein_coding | **505.491** | 0.346 | 0.104 | **1.271** | 2.98E-03 | **7.10E-03** |
| ENSG00000011275.20 | **RNF216** | 54476 | protein_coding | **1435.377** | 0.338 | 0.102 | **1.264** | 2.99E-03 | **7.11E-03** |
| ENSG00000166928.12 | **MS4A14** | 84689 | protein_coding | **197.830** | 0.645 | 0.194 | **1.563** | 2.99E-03 | **7.12E-03** |
| ENSG00000256128.8 | **LINC00944** | 100996671 | lncRNA | **37.187** | 1.027 | 0.309 | **2.037** | 3.01E-03 | **7.15E-03** |
| ENSG00000163931.19 | **TKT** | 7086 | protein_coding | **5667.175** | -0.374 | -0.113 | **0.772** | 3.01E-03 | **7.16E-03** |
| ENSG00000139266.7 | **MARCHF9** | 92979 | protein_coding | **1078.467** | 0.354 | 0.107 | **1.278** | 3.01E-03 | **7.16E-03** |
| ENSG00000070367.19 | **EXOC5** | 10640 | protein_coding | **454.509** | -0.357 | -0.107 | **0.781** | 3.01E-03 | **7.17E-03** |
| ENSG00000295728.1 | **ENSG00000295728** | | lncRNA | **13.376** | 1.102 | 0.332 | **2.147** | 3.01E-03 | **7.17E-03** |
| ENSG00000127463.18 | **EMC1** | 23065 | protein_coding | **575.461** | -0.494 | -0.149 | **0.710** | 3.02E-03 | **7.18E-03** |
| ENSG00000288670.1 | **ENSG00000288670** | | lncRNA | **34.851** | 0.798 | 0.240 | **1.739** | 3.03E-03 | **7.21E-03** |
| ENSG00000182195.10 | **LDOC1** | 23641 | protein_coding | **29.404** | 1.448 | 0.436 | **2.729** | 3.04E-03 | **7.24E-03** |
| ENSG00000172139.16 | **SLC9C1** | 285335 | protein_coding | **5.122** | 2.687 | 0.809 | **6.441** | 3.04E-03 | **7.24E-03** |
| ENSG00000181555.24 | **SETD2** | 29072 | protein_coding | **1703.635** | 0.321 | 0.097 | **1.249** | 3.05E-03 | **7.24E-03** |
| ENSG00000115942.11 | **ORC2** | 4999 | protein_coding | **476.517** | -0.362 | -0.109 | **0.778** | 3.05E-03 | **7.24E-03** |
| ENSG00000198780.15 | **FAM169A** | 26049 | protein_coding | **23.125** | -1.070 | -0.322 | **0.476** | 3.05E-03 | **7.26E-03** |
| ENSG00000289551.3 | **ENSG00000289551** | | lncRNA | **7.116** | 1.808 | 0.544 | **3.502** | 3.05E-03 | **7.26E-03** |
| ENSG00000288061.4 | **ENSG00000288061** | | lncRNA | **128.830** | 0.710 | 0.214 | **1.636** | 3.06E-03 | **7.26E-03** |
| ENSG00000117016.12 | **RIMS3** | 9783 | protein_coding | **139.382** | 0.805 | 0.242 | **1.747** | 3.06E-03 | **7.26E-03** |
| ENSG00000262919.10 | **CCNQ** | 92002 | protein_coding | **226.535** | -0.461 | -0.139 | **0.726** | 3.06E-03 | **7.26E-03** |
| ENSG00000121690.13 | **DEPDC7** | 91614 | protein_coding | **62.594** | 0.833 | 0.251 | **1.782** | 3.07E-03 | **7.28E-03** |
| ENSG00000047849.24 | **MAP4** | 4134 | protein_coding | **3705.693** | 0.300 | 0.090 | **1.231** | 3.07E-03 | **7.30E-03** |
| ENSG00000255240.7 | **GLYATL1-A** | 283194 | lncRNA | **70.802** | 0.610 | 0.184 | **1.526** | 3.08E-03 | **7.31E-03** |

| | | | | | | | | | |
|---|---|---|---|---|---|---|---|---|---|
| ENSG00000222009.11 | **BTBD19** | 149478 | protein_coding | **93.286** | 0.754 | 0.227 | **1.686** | 3.08E-03 | **7.32E-03** |
| ENSG00000275963.1 | **ENSG00000275963** | | lncRNA | **6.577** | 1.822 | 0.549 | **3.536** | 3.08E-03 | **7.32E-03** |
| ENSG00000294512.1 | **ENSG00000294512** | | lncRNA | **211.155** | -0.439 | -0.132 | **0.738** | 3.09E-03 | **7.33E-03** |
| ENSG00000105993.17 | **DNAJB6** | 10049 | protein_coding | **1509.199** | 0.385 | 0.116 | **1.306** | 3.09E-03 | **7.33E-03** |
| ENSG00000138785.18 | **INTS12** | 57117 | protein_coding | **466.158** | -0.387 | -0.116 | **0.765** | 3.09E-03 | **7.33E-03** |
| ENSG00000174574.18 | **AKIRIN1** | 79647 | protein_coding | **847.308** | 0.449 | 0.135 | **1.365** | 3.09E-03 | **7.33E-03** |
| ENSG00000257127.6 | **CLLU1** | 574028 | lncRNA | **4.928** | 3.025 | 0.910 | **8.137** | 3.09E-03 | **7.34E-03** |
| ENSG00000196150.16 | **ZNF250** | 58500 | protein_coding | **189.343** | 0.433 | 0.130 | **1.350** | 3.09E-03 | **7.34E-03** |
| ENSG00000261613.2 | **ENSG00000261613** | | lncRNA | **104.065** | 0.593 | 0.179 | **1.509** | 3.10E-03 | **7.36E-03** |
| ENSG00000100387.11 | **RBX1** | 9978 | protein_coding | **1790.132** | -0.510 | -0.154 | **0.702** | 3.10E-03 | **7.36E-03** |
| ENSG00000269190.8 | **FBXO17** | 115290 | protein_coding | **38.987** | 1.061 | 0.319 | **2.087** | 3.11E-03 | **7.38E-03** |
| ENSG00000261474.1 | **ENSG00000261474** | | lncRNA | **33.474** | 0.830 | 0.250 | **1.777** | 3.11E-03 | **7.38E-03** |
| ENSG00000167394.15 | **ZNF668** | 79759 | protein_coding | **203.737** | -0.513 | -0.154 | **0.701** | 3.11E-03 | **7.38E-03** |
| ENSG00000259551.2 | **ENSG00000259551** | | lncRNA | **81.207** | 0.777 | 0.234 | **1.714** | 3.12E-03 | **7.39E-03** |
| ENSG00000186409.20 | **CCDC30** | 728621 | protein_coding | **52.322** | 0.904 | 0.272 | **1.871** | 3.12E-03 | **7.40E-03** |
| ENSG00000253632.2 | **ENSG0000(** | 10537935( | lncRNA | **30.727** | 1.041 | 0.313 | **2.057** | 3.14E-03 | **7.44E-03** |
| ENSG00000163374.23 | **YY1AP1** | 55249 | protein_coding | **1798.884** | -0.296 | -0.089 | **0.814** | 3.14E-03 | **7.44E-03** |
| ENSG00000137309.23 | **HMGA1** | 3159 | protein_coding | **7787.070** | -0.410 | -0.123 | **0.753** | 3.15E-03 | **7.47E-03** |
| ENSG00000143303.14 | **METTL25B** | 51093 | protein_coding | **807.161** | 0.390 | 0.117 | **1.311** | 3.16E-03 | **7.48E-03** |
| ENSG00000258634.3 | **ENSG00000258634** | | lncRNA | **118.255** | 0.619 | 0.186 | **1.536** | 3.16E-03 | **7.49E-03** |
| ENSG00000142082.17 | **SIRT3** | 23410 | protein_coding | **623.132** | 0.274 | 0.082 | **1.209** | 3.16E-03 | **7.49E-03** |
| ENSG00000106125.16 | **MINDY4** | 84182 | protein_coding | **38.700** | -0.946 | -0.285 | **0.519** | 3.17E-03 | **7.50E-03** |
| ENSG00000148843.17 | **PDCD11** | 22984 | protein_coding | **1168.650** | -0.352 | -0.106 | **0.784** | 3.17E-03 | **7.51E-03** |
| ENSG00000144182.19 | **LIPT1** | 51601 | protein_coding | **76.325** | -0.586 | -0.176 | **0.666** | 3.17E-03 | **7.51E-03** |
| ENSG00000071909.20 | **MYO3B** | 140469 | protein_coding | **128.553** | 0.589 | 0.177 | **1.504** | 3.18E-03 | **7.52E-03** |
| ENSG00000270136.6 | **MICOS10-NBL1** | | protein_coding | **29.373** | -0.906 | -0.273 | **0.533** | 3.18E-03 | **7.52E-03** |
| ENSG00000310475.1 | **ENSG00000310475** | | lncRNA | **16.031** | 1.139 | 0.343 | **2.202** | 3.19E-03 | **7.55E-03** |
| ENSG00000166337.12 | **TAF10** | 6881 | protein_coding | **2223.438** | 0.392 | 0.118 | **1.312** | 3.19E-03 | **7.56E-03** |
| ENSG00000129518.12 | **EAPP** | 55837 | protein_coding | **408.271** | 0.472 | 0.142 | **1.387** | 3.20E-03 | **7.56E-03** |
| ENSG00000167693.18 | **NXN** | 64359 | protein_coding | **57.669** | 0.901 | 0.271 | **1.867** | 3.21E-03 | **7.59E-03** |
| ENSG00000237693.7 | **IRGM** | 345611 | protein_coding | **10.574** | 1.851 | 0.557 | **3.608** | 3.21E-03 | **7.59E-03** |

| | | | | | | | | | |
|---|---|---|---|---|---|---|---|---|---|
| ENSG00000146776.16 | **ATXN7L1** | 222255 | protein_coding | **91.268** | -0.608 | -0.183 | **0.656** | 3.21E-03 | **7.60E-03** |
| ENSG00000307568.1 | **ENSG00000307568** | | lncRNA | **10.509** | 1.543 | 0.464 | **2.913** | 3.21E-03 | **7.60E-03** |
| ENSG00000138759.22 | **FRAS1** | 80144 | protein_coding | **10.848** | 2.043 | 0.615 | **4.121** | 3.24E-03 | **7.65E-03** |
| ENSG00000306471.1 | **ENSG00000306471** | | lncRNA | **4.464** | 2.537 | 0.764 | **5.803** | 3.24E-03 | **7.66E-03** |
| ENSG00000189144.17 | **ZNF573** | 126231 | protein_coding | **13.120** | -1.335 | -0.402 | **0.396** | 3.24E-03 | **7.67E-03** |
| ENSG00000286186.2 | **ENSG00000286186** | | lncRNA | **15.935** | -1.131 | -0.341 | **0.456** | 3.25E-03 | **7.67E-03** |
| ENSG00000181790.14 | **ADGRB1** | 575 | protein_coding | **82.732** | 0.964 | 0.290 | **1.951** | 3.25E-03 | **7.67E-03** |
| ENSG00000203761.5 | **MSTO2P** | | transcribed_unpro | **7.911** | -2.174 | -0.654 | **0.222** | 3.25E-03 | **7.68E-03** |
| ENSG00000290999.2 | **ENSG00000290999** | | lncRNA | **22.515** | 0.926 | 0.279 | **1.900** | 3.26E-03 | **7.70E-03** |
| ENSG00000163634.13 | **THOC7** | 80145 | protein_coding | **1187.500** | -0.328 | -0.099 | **0.796** | 3.26E-03 | **7.70E-03** |
| ENSG00000135392.21 | **DNAJC14** | 85406 | protein_coding | **756.404** | 0.338 | 0.102 | **1.264** | 3.27E-03 | **7.72E-03** |
| ENSG00000270164.4 | **LINC01480** | | lncRNA | **1209.860** | -0.573 | -0.173 | **0.672** | 3.27E-03 | **7.73E-03** |
| ENSG00000070718.15 | **AP3M2** | 10947 | protein_coding | **829.677** | 0.425 | 0.128 | **1.343** | 3.27E-03 | **7.73E-03** |
| ENSG00000090376.13 | **IRAK3** | 11213 | protein_coding | **289.313** | 0.339 | 0.102 | **1.265** | 3.27E-03 | **7.73E-03** |
| ENSG00000177464.7 | **GPR4** | 2828 | protein_coding | **8.487** | 1.664 | 0.501 | **3.170** | 3.28E-03 | **7.75E-03** |
| ENSG00000277182.3 | **ENSG0000(** | 10028780 | lncRNA | **11.683** | 1.373 | 0.413 | **2.590** | 3.29E-03 | **7.76E-03** |
| ENSG00000237604.1 | **ENSG00000237604** | | lncRNA | **14.073** | 1.626 | 0.490 | **3.087** | 3.29E-03 | **7.76E-03** |
| ENSG00000230943.5 | **LINC02541** | 10192768 | lncRNA | **190.757** | 0.588 | 0.177 | **1.503** | 3.29E-03 | **7.76E-03** |
| ENSG00000240057.9 | **NEPRO-AS** | 10272343 | lncRNA | **15.384** | -1.180 | -0.355 | **0.441** | 3.30E-03 | **7.78E-03** |
| ENSG00000247157.8 | **LINC01252** | 338817 | lncRNA | **28.379** | -0.852 | -0.257 | **0.554** | 3.30E-03 | **7.78E-03** |
| ENSG00000289018.3 | **ENSG00000289018** | | lncRNA | **14.134** | 1.241 | 0.374 | **2.364** | 3.30E-03 | **7.79E-03** |
| ENSG00000213585.13 | **VDAC1** | 7416 | protein_coding | **6384.221** | -0.401 | -0.121 | **0.757** | 3.30E-03 | **7.79E-03** |
| ENSG00000143183.20 | **TMCO1** | 54499 | protein_coding | **858.534** | -0.291 | -0.088 | **0.817** | 3.31E-03 | **7.81E-03** |
| ENSG00000134255.18 | **CEPT1** | 10390 | protein_coding | **1108.964** | -0.419 | -0.126 | **0.748** | 3.31E-03 | **7.81E-03** |
| ENSG00000196132.16 | **MYT1** | 4661 | protein_coding | **13.634** | 1.873 | 0.564 | **3.664** | 3.31E-03 | **7.82E-03** |
| ENSG00000307331.1 | **ENSG00000307331** | | lncRNA | **4.801** | 2.584 | 0.778 | **5.994** | 3.32E-03 | **7.82E-03** |
| ENSG00000100316.18 | **RPL3** | 6122 | protein_coding | **66016.733** | -0.333 | -0.100 | **0.794** | 3.32E-03 | **7.83E-03** |
| ENSG00000273047.1 | **ENSG00000273047** | | protein_coding | **114.794** | -0.885 | -0.266 | **0.542** | 3.34E-03 | **7.86E-03** |
| ENSG00000235009.2 | **LINC01883** | 10537446 | lncRNA | **37.090** | 0.692 | 0.208 | **1.616** | 3.34E-03 | **7.87E-03** |
| ENSG00000156239.14 | **HEMK2** | 29104 | protein_coding | **166.824** | -0.371 | -0.112 | **0.773** | 3.34E-03 | **7.88E-03** |
| ENSG00000106615.12 | **RHEB** | 6009 | protein_coding | **584.100** | -0.280 | -0.084 | **0.824** | 3.35E-03 | **7.88E-03** |

| | | | | | | | | | |
|---|---|---|---|---|---|---|---|---|---|
| ENSG00000198816.9 | **ZNF358** | 140467 | protein_coding | **435.571** | 0.616 | 0.185 | **1.533** | 3.37E-03 | **7.93E-03** |
| ENSG00000126883.21 | **NUP214** | 8021 | protein_coding | **1201.973** | -0.359 | -0.108 | **0.780** | 3.37E-03 | **7.93E-03** |
| ENSG00000272791.1 | **ENSG00000272791** | | lncRNA | **26.641** | 0.942 | 0.284 | **1.921** | 3.37E-03 | **7.93E-03** |
| ENSG00000183022.5 | **TPM3P8** | | processed_pseudo | **4.132** | 2.606 | 0.784 | **6.088** | 3.38E-03 | **7.95E-03** |
| ENSG00000264345.3 | **LINC01894** | | lncRNA | **9.159** | 2.087 | 0.628 | **4.248** | 3.38E-03 | **7.95E-03** |
| ENSG00000226979.10 | **LTA** | 4049 | protein_coding | **3370.344** | -0.631 | -0.190 | **0.646** | 3.39E-03 | **7.97E-03** |
| ENSG00000121058.7 | **COIL** | 8161 | protein_coding | **258.135** | -0.341 | -0.103 | **0.789** | 3.39E-03 | **7.98E-03** |
| ENSG00000144824.23 | **PHLDB2** | 90102 | protein_coding | **9.066** | 1.813 | 0.546 | **3.514** | 3.39E-03 | **7.98E-03** |
| ENSG00000289827.2 | **ENSG00000289827** | | lncRNA | **16.348** | 1.106 | 0.333 | **2.152** | 3.39E-03 | **7.99E-03** |
| ENSG00000118620.15 | **ZNF430** | 80264 | protein_coding | **97.962** | -0.613 | -0.184 | **0.654** | 3.40E-03 | **8.00E-03** |
| ENSG00000105792.21 | **CFAP69** | 79846 | protein_coding | **21.605** | 1.087 | 0.327 | **2.124** | 3.41E-03 | **8.02E-03** |
| ENSG00000049167.18 | **ERCC8** | 1161 | protein_coding | **149.301** | -0.518 | -0.156 | **0.699** | 3.42E-03 | **8.05E-03** |
| ENSG00000271848.3 | **SYNPO2L-AS1** | | lncRNA | **8.311** | 1.695 | 0.510 | **3.237** | 3.42E-03 | **8.05E-03** |
| ENSG00000136859.12 | **ANGPTL2** | 23452 | protein_coding | **25.704** | -1.100 | -0.331 | **0.466** | 3.42E-03 | **8.05E-03** |
| ENSG00000111605.20 | **CPSF6** | 11052 | protein_coding | **1074.046** | -0.337 | -0.101 | **0.792** | 3.43E-03 | **8.06E-03** |
| ENSG00000196646.14 | **ZNF136** | 7695 | protein_coding | **145.975** | -0.453 | -0.137 | **0.730** | 3.43E-03 | **8.07E-03** |
| ENSG00000245105.5 | **A2M-AS1** | 144571 | lncRNA | **3.111** | 3.288 | 0.990 | **9.769** | 3.45E-03 | **8.12E-03** |
| ENSG00000146574.18 | **CCZ1B** | 221960 | protein_coding | **952.834** | -0.329 | -0.099 | **0.796** | 3.46E-03 | **8.13E-03** |
| ENSG00000142675.20 | **CNKSR1** | 10256 | protein_coding | **69.294** | 0.651 | 0.196 | **1.570** | 3.46E-03 | **8.13E-03** |
| ENSG00000203876.11 | **ADD3-AS1** | 10050593 | lncRNA | **7.039** | 2.028 | 0.610 | **4.077** | 3.46E-03 | **8.13E-03** |
| ENSG00000272277.2 | **ENSG00000272277** | | lncRNA | **10.729** | 1.401 | 0.422 | **2.640** | 3.47E-03 | **8.14E-03** |
| ENSG00000162777.19 | **DENND2D** | 79961 | protein_coding | **1994.250** | -0.442 | -0.133 | **0.736** | 3.47E-03 | **8.15E-03** |
| ENSG00000297741.1 | **ENSG00000297741** | | lncRNA | **5.461** | 2.000 | 0.602 | **4.000** | 3.47E-03 | **8.15E-03** |
| ENSG00000286277.2 | **ENSG00000286277** | | lncRNA | **8.859** | -1.518 | -0.457 | **0.349** | 3.47E-03 | **8.15E-03** |
| ENSG00000204644.12 | **ZFP57** | 346171 | protein_coding | **47.581** | 1.730 | 0.521 | **3.317** | 3.47E-03 | **8.16E-03** |
| ENSG00000149177.16 | **PTPRJ** | 5795 | protein_coding | **93.478** | 0.672 | 0.202 | **1.594** | 3.48E-03 | **8.17E-03** |
| ENSG00000081760.19 | **AACS** | 65985 | protein_coding | **745.050** | -0.278 | -0.084 | **0.825** | 3.48E-03 | **8.18E-03** |
| ENSG00000204599.18 | **TRIM39** | 56658 | protein_coding | **309.405** | -0.320 | -0.096 | **0.801** | 3.49E-03 | **8.18E-03** |
| ENSG00000047648.24 | **ARHGAP6** | 395 | protein_coding | **122.236** | -0.603 | -0.181 | **0.659** | 3.49E-03 | **8.19E-03** |
| ENSG00000113580.17 | **NR3C1** | 2908 | protein_coding | **1764.897** | 0.432 | 0.130 | **1.349** | 3.50E-03 | **8.20E-03** |
| ENSG00000298055.1 | **ENSG00000298055** | | lncRNA | **17.142** | 1.077 | 0.324 | **2.110** | 3.50E-03 | **8.20E-03** |

| | | | | | | | | | |
|---|---|---|---|---|---|---|---|---|---|
| ENSG00000302344.1 | **ENSG00000302344** | | lncRNA | **86.097** | 0.497 | 0.150 | **1.412** | 3.50E-03 | **8.21E-03** |
| ENSG00000184584.16 | **STING1** | 340061 | protein_coding | **275.582** | 0.458 | 0.138 | **1.374** | 3.50E-03 | **8.22E-03** |
| ENSG00000129596.7 | **CDO1** | 1036 | protein_coding | **5.900** | 2.250 | 0.677 | **4.755** | 3.51E-03 | **8.24E-03** |
| ENSG00000293505.1 | **GNRHR2** | 114814 | lncRNA | **21.691** | 1.102 | 0.332 | **2.147** | 3.52E-03 | **8.26E-03** |
| ENSG00000143442.25 | **POGZ** | 23126 | protein_coding | **1975.962** | 0.304 | 0.092 | **1.235** | 3.53E-03 | **8.27E-03** |
| ENSG00000185507.23 | **IRF7** | 3665 | protein_coding | **5856.427** | -0.415 | -0.125 | **0.750** | 3.53E-03 | **8.28E-03** |
| ENSG00000005889.18 | **ZFX** | 7543 | protein_coding | **289.094** | -0.507 | -0.153 | **0.704** | 3.54E-03 | **8.30E-03** |
| ENSG00000101493.13 | **ZNF516** | 9658 | protein_coding | **571.947** | 0.516 | 0.155 | **1.430** | 3.54E-03 | **8.30E-03** |
| ENSG00000163467.14 | **TSACC** | 128229 | protein_coding | **4.773** | 1.944 | 0.585 | **3.847** | 3.54E-03 | **8.30E-03** |
| ENSG00000300534.1 | **ENSG00000300534** | | lncRNA | **8.150** | -1.710 | -0.515 | **0.306** | 3.55E-03 | **8.32E-03** |
| ENSG00000138459.11 | **SLC35A5** | 55032 | protein_coding | **288.447** | 0.469 | 0.141 | **1.384** | 3.55E-03 | **8.32E-03** |
| ENSG00000173621.11 | **LRFN4** | 78999 | protein_coding | **313.985** | -0.431 | -0.130 | **0.742** | 3.55E-03 | **8.33E-03** |
| ENSG00000293801.1 | **ENSG00000293801** | | lncRNA | **6.681** | 1.995 | 0.601 | **3.987** | 3.55E-03 | **8.33E-03** |
| ENSG00000174327.8 | **SLC16A13** | 201232 | protein_coding | **179.669** | 0.398 | 0.120 | **1.318** | 3.55E-03 | **8.33E-03** |
| ENSG00000115239.26 | **ASB3** | 51130 | protein_coding | **300.602** | -0.393 | -0.118 | **0.762** | 3.56E-03 | **8.33E-03** |
| ENSG00000164615.8 | **CAMLG** | 819 | protein_coding | **573.883** | -0.273 | -0.082 | **0.828** | 3.56E-03 | **8.33E-03** |
| ENSG00000152292.19 | **SH2D6** | 284948 | protein_coding | **9.755** | 1.687 | 0.508 | **3.219** | 3.56E-03 | **8.33E-03** |
| ENSG00000152767.19 | **FARP1** | 10160 | protein_coding | **398.117** | 0.484 | 0.146 | **1.399** | 3.56E-03 | **8.34E-03** |
| ENSG00000253819.4 | **LINC01151** | | lncRNA | **7.085** | 2.947 | 0.887 | **7.713** | 3.57E-03 | **8.36E-03** |
| ENSG00000163946.16 | **TASOR** | 23272 | protein_coding | **908.046** | -0.365 | -0.110 | **0.776** | 3.58E-03 | **8.37E-03** |
| ENSG00000153786.15 | **ZDHHC7** | 55625 | protein_coding | **626.751** | 0.279 | 0.084 | **1.213** | 3.58E-03 | **8.38E-03** |
| ENSG00000070756.19 | **PABPC1** | 26986 | protein_coding | **20063.065** | -0.326 | -0.098 | **0.798** | 3.59E-03 | **8.39E-03** |
| ENSG00000260027.6 | **HOXB7** | 3217 | protein_coding | **95.584** | 0.526 | 0.158 | **1.440** | 3.59E-03 | **8.41E-03** |
| ENSG00000004487.20 | **KDM1A** | 23028 | protein_coding | **1797.510** | -0.303 | -0.091 | **0.810** | 3.59E-03 | **8.41E-03** |
| ENSG00000163512.16 | **AZI2** | 64343 | protein_coding | **371.491** | -0.578 | -0.174 | **0.670** | 3.60E-03 | **8.42E-03** |
| ENSG00000258944.1 | **PAPLN-AS2** | | lncRNA | **19.554** | 0.945 | 0.284 | **1.925** | 3.60E-03 | **8.42E-03** |
| ENSG00000166913.15 | **YWHAB** | 7529 | protein_coding | **3248.694** | -0.347 | -0.104 | **0.786** | 3.60E-03 | **8.42E-03** |
| ENSG00000004779.12 | **NDUFAB1** | 4706 | protein_coding | **766.835** | -0.438 | -0.132 | **0.738** | 3.60E-03 | **8.43E-03** |
| ENSG00000167136.9 | **ENDOG** | 2021 | protein_coding | **188.480** | -0.539 | -0.162 | **0.688** | 3.60E-03 | **8.43E-03** |
| ENSG00000246273.10 | **SBF2-AS1** | 283104 | lncRNA | **87.037** | -0.558 | -0.168 | **0.679** | 3.61E-03 | **8.43E-03** |
| ENSG00000276462.3 | **LINC03025** | 440896 | lncRNA | **12.930** | 1.489 | 0.448 | **2.806** | 3.61E-03 | **8.43E-03** |

| | | | | | | | | | |
|---|---|---|---|---|---|---|---|---|---|
| ENSG00000198755.13 | **RPL10A** | 4736 | protein_coding | **15075.813** | -0.378 | -0.114 | **0.769** | 3.62E-03 | **8.46E-03** |
| ENSG00000118503.20 | **TNFAIP3** | 7128 | protein_coding | **6352.416** | 0.454 | 0.137 | **1.370** | 3.62E-03 | **8.46E-03** |
| ENSG00000008382.18 | **MPND** | 84954 | protein_coding | **285.225** | 0.367 | 0.111 | **1.290** | 3.62E-03 | **8.46E-03** |
| ENSG00000156050.11 | **FAM161B** | 145483 | protein_coding | **34.394** | 0.877 | 0.264 | **1.837** | 3.63E-03 | **8.48E-03** |
| ENSG00000289755.2 | **ENSG0000(** | 6817 | lncRNA | **3.987** | 3.680 | 1.108 | **12.818** | 3.64E-03 | **8.50E-03** |
| ENSG00000115541.13 | **HSPE1** | 3336 | protein_coding | **2044.771** | -0.551 | -0.166 | **0.683** | 3.64E-03 | **8.50E-03** |
| ENSG00000124493.16 | **GRM4** | 2914 | protein_coding | **10.899** | 1.607 | 0.484 | **3.046** | 3.64E-03 | **8.51E-03** |
| ENSG00000289537.3 | **ENSG00000289537** | | lncRNA | **11.843** | 1.300 | 0.391 | **2.463** | 3.65E-03 | **8.52E-03** |
| ENSG00000141040.16 | **ZNF287** | 57336 | protein_coding | **138.028** | 0.596 | 0.179 | **1.511** | 3.66E-03 | **8.54E-03** |
| ENSG00000106560.12 | **GIMAP2** | 26157 | protein_coding | **228.048** | -0.483 | -0.145 | **0.715** | 3.66E-03 | **8.54E-03** |
| ENSG00000159346.15 | **ADIPOR1** | 51094 | protein_coding | **1274.905** | -0.315 | -0.095 | **0.804** | 3.66E-03 | **8.55E-03** |
| ENSG00000196504.23 | **PRPF40A** | 55660 | protein_coding | **1291.967** | -0.390 | -0.117 | **0.763** | 3.66E-03 | **8.55E-03** |
| ENSG00000241058.5 | **NSUN6** | 221078 | protein_coding | **376.899** | 0.392 | 0.118 | **1.312** | 3.67E-03 | **8.57E-03** |
| ENSG00000138399.20 | **FASTKD1** | 79675 | protein_coding | **779.909** | -0.405 | -0.122 | **0.755** | 3.68E-03 | **8.58E-03** |
| ENSG00000113583.10 | **C5orf15** | 56951 | protein_coding | **668.948** | -0.285 | -0.086 | **0.821** | 3.68E-03 | **8.58E-03** |
| ENSG00000139547.10 | **RDH16** | 8608 | protein_coding | **30.845** | 0.954 | 0.287 | **1.937** | 3.68E-03 | **8.59E-03** |
| ENSG00000120705.15 | **ETF1** | 2107 | protein_coding | **1435.103** | -0.298 | -0.090 | **0.814** | 3.70E-03 | **8.62E-03** |
| ENSG00000127774.9 | **EMC6** | 83460 | protein_coding | **603.803** | -0.306 | -0.092 | **0.809** | 3.70E-03 | **8.63E-03** |
| ENSG00000183978.9 | **COA3** | 28958 | protein_coding | **449.340** | -0.424 | -0.128 | **0.745** | 3.70E-03 | **8.63E-03** |
| ENSG00000285884.4 | **ENSG0000(** | 10537826ε | lncRNA | **9.346** | 1.597 | 0.481 | **3.025** | 3.70E-03 | **8.64E-03** |
| ENSG00000268119.9 | **ENSG0000(** | 105372321 | lncRNA | **10.796** | -1.765 | -0.531 | **0.294** | 3.71E-03 | **8.65E-03** |
| ENSG00000293431.1 | **MBL1P** | 8512 | lncRNA | **13.331** | 1.326 | 0.399 | **2.507** | 3.71E-03 | **8.66E-03** |
| ENSG00000269892.1 | **ENSG00000269892** | | lncRNA | **8.624** | 1.504 | 0.453 | **2.835** | 3.74E-03 | **8.71E-03** |
| ENSG00000166912.19 | **MTMR10** | 54893 | protein_coding | **300.095** | 0.391 | 0.118 | **1.311** | 3.74E-03 | **8.71E-03** |
| ENSG00000143570.21 | **SLC39A1** | 27173 | protein_coding | **1630.749** | -0.304 | -0.091 | **0.810** | 3.74E-03 | **8.71E-03** |
| ENSG00000147465.14 | **STAR** | 6770 | protein_coding | **11.517** | -1.544 | -0.465 | **0.343** | 3.74E-03 | **8.72E-03** |
| ENSG00000203279.7 | **ENSG00000203279** | | lncRNA | **151.027** | 0.533 | 0.160 | **1.447** | 3.77E-03 | **8.78E-03** |
| ENSG00000175087.12 | **PDIK1L** | 149420 | protein_coding | **65.031** | -0.772 | -0.232 | **0.586** | 3.77E-03 | **8.79E-03** |
| ENSG00000253582.2 | **OXR1-AS1** | | lncRNA | **5.055** | 2.300 | 0.692 | **4.925** | 3.78E-03 | **8.81E-03** |
| ENSG00000163166.17 | **IWS1** | 55677 | protein_coding | **1002.066** | -0.284 | -0.086 | **0.821** | 3.79E-03 | **8.82E-03** |
| ENSG00000120332.19 | **TNN** | 63923 | protein_coding | **14.368** | -1.092 | -0.329 | **0.469** | 3.80E-03 | **8.84E-03** |

| | | | | | | | | | |
|---|---|---|---|---|---|---|---|---|---|
| ENSG00000172888.13 | **ZNF621** | 285268 | protein_coding | **366.337** | -0.442 | -0.133 | **0.736** | 3.81E-03 | **8.86E-03** |
| ENSG00000163218.16 | **PGLYRP4** | 57115 | protein_coding | **17.282** | -1.135 | -0.342 | **0.455** | 3.81E-03 | **8.88E-03** |
| ENSG00000188730.7 | **VWC2** | 375567 | protein_coding | **5.553** | 2.329 | 0.701 | **5.025** | 3.82E-03 | **8.88E-03** |
| ENSG00000291083.2 | **CYP2B7P** | 1556 | lncRNA | **5.909** | 2.376 | 0.715 | **5.191** | 3.82E-03 | **8.90E-03** |
| ENSG00000279140.2 | **ENSG00000279140** | | lncRNA | **27.209** | 0.859 | 0.259 | **1.813** | 3.83E-03 | **8.91E-03** |
| ENSG00000175600.18 | **SUGCT** | 79783 | protein_coding | **11.107** | 1.354 | 0.408 | **2.557** | 3.83E-03 | **8.91E-03** |
| ENSG00000156711.19 | **MAPK13** | 5603 | protein_coding | **164.335** | -0.520 | -0.157 | **0.697** | 3.83E-03 | **8.92E-03** |
| ENSG00000204060.8 | **FOXO6** | 10013207[illegible] | protein_coding | **10.207** | 1.321 | 0.398 | **2.499** | 3.84E-03 | **8.93E-03** |
| ENSG00000261420.1 | **MPC1-DT** | | lncRNA | **6.989** | 1.615 | 0.486 | **3.064** | 3.84E-03 | **8.93E-03** |
| ENSG00000289641.2 | **ENSG00000289641** | | lncRNA | **7.165** | 1.638 | 0.493 | **3.113** | 3.85E-03 | **8.94E-03** |
| ENSG00000125149.13 | **PHAF1** | 80262 | protein_coding | **315.638** | 0.339 | 0.102 | **1.265** | 3.85E-03 | **8.94E-03** |
| ENSG00000130165.13 | **ELOF1** | 84337 | protein_coding | **1280.028** | 0.295 | 0.089 | **1.227** | 3.85E-03 | **8.94E-03** |
| ENSG00000172663.11 | **TMEM134** | 80194 | protein_coding | **556.739** | -0.376 | -0.113 | **0.770** | 3.85E-03 | **8.96E-03** |
| ENSG00000213062.6 | **ENSG00000213062** | | lncRNA | **45.515** | 0.704 | 0.212 | **1.629** | 3.86E-03 | **8.96E-03** |
| ENSG00000107736.24 | **CDH23** | 64072 | protein_coding | **31.824** | 1.119 | 0.337 | **2.173** | 3.86E-03 | **8.98E-03** |
| ENSG00000011590.16 | **ZBTB32** | 27033 | protein_coding | **1488.457** | -0.394 | -0.119 | **0.761** | 3.87E-03 | **8.99E-03** |
| ENSG00000185909.17 | **KLHDC8B** | 200942 | protein_coding | **76.449** | 0.801 | 0.241 | **1.743** | 3.89E-03 | **9.03E-03** |
| ENSG00000129473.13 | **BCL2L2** | 599 | protein_coding | **480.586** | -0.400 | -0.121 | **0.758** | 3.89E-03 | **9.03E-03** |
| ENSG00000197417.10 | **SHPK** | 23729 | protein_coding | **103.081** | -0.666 | -0.201 | **0.630** | 3.89E-03 | **9.03E-03** |
| ENSG00000115526.13 | **CHST10** | 9486 | protein_coding | **240.653** | -0.351 | -0.106 | **0.784** | 3.89E-03 | **9.03E-03** |
| ENSG00000121749.18 | **TBC1D15** | 64786 | protein_coding | **303.866** | -0.429 | -0.129 | **0.743** | 3.89E-03 | **9.04E-03** |
| ENSG00000169860.7 | **P2RY1** | 5028 | protein_coding | **29.211** | -0.860 | -0.259 | **0.551** | 3.91E-03 | **9.07E-03** |
| ENSG00000171435.16 | **KSR2** | 283455 | protein_coding | **7.267** | 1.701 | 0.512 | **3.252** | 3.91E-03 | **9.07E-03** |
| ENSG00000136261.18 | **BZW2** | 28969 | protein_coding | **48.596** | -0.687 | -0.207 | **0.621** | 3.91E-03 | **9.08E-03** |
| ENSG00000163207.8 | **IVL** | 3713 | protein_coding | **48.326** | 0.780 | 0.235 | **1.717** | 3.92E-03 | **9.09E-03** |
| ENSG00000168135.6 | **KCNJ4** | 3761 | protein_coding | **3.329** | 2.517 | 0.758 | **5.725** | 3.92E-03 | **9.11E-03** |
| ENSG00000280620.2 | **SCAANT1** | 10086156[illegible] | lncRNA | **5.840** | 1.894 | 0.570 | **3.717** | 3.93E-03 | **9.11E-03** |
| ENSG00000172292.17 | **CERS6** | 253782 | protein_coding | **787.925** | 0.479 | 0.144 | **1.394** | 3.93E-03 | **9.11E-03** |
| ENSG00000134516.22 | **DOCK2** | 1794 | protein_coding | **2130.467** | -0.423 | -0.127 | **0.746** | 3.93E-03 | **9.11E-03** |
| ENSG00000050555.21 | **LAMC3** | 10319 | protein_coding | **7.757** | 2.245 | 0.676 | **4.739** | 3.93E-03 | **9.12E-03** |
| ENSG00000165406.18 | **MARCHF8** | 220972 | protein_coding | **465.924** | 0.488 | 0.147 | **1.402** | 3.93E-03 | **9.12E-03** |

| | | | | | | | | | |
|---|---|---|---|---|---|---|---|---|---|
| ENSG00000101336.19 | **HCK** | 3055 | protein_coding | **510.509** | -0.530 | -0.160 | **0.692** | 3.93E-03 | **9.12E-03** |
| ENSG00000100209.13 | **HSCB** | 150274 | protein_coding | **256.531** | -0.396 | -0.119 | **0.760** | 3.94E-03 | **9.13E-03** |
| ENSG00000233493.6 | **TMEM238** | 388564 | protein_coding | **24.711** | -0.971 | -0.292 | **0.510** | 3.94E-03 | **9.15E-03** |
| ENSG00000113356.15 | **POLR3G** | 10622 | protein_coding | **140.394** | -0.610 | -0.183 | **0.655** | 3.95E-03 | **9.15E-03** |
| ENSG00000135740.19 | **SLC9A5** | 6553 | protein_coding | **245.444** | 0.617 | 0.186 | **1.533** | 3.95E-03 | **9.15E-03** |
| ENSG00000133028.14 | **SCO1** | 6341 | protein_coding | **490.955** | -0.385 | -0.116 | **0.766** | 3.96E-03 | **9.17E-03** |
| ENSG00000270012.1 | **ENSG00000270012** | | lncRNA | **72.111** | -0.555 | -0.167 | **0.680** | 3.96E-03 | **9.17E-03** |
| ENSG00000130958.15 | **SLC35D2** | 11046 | protein_coding | **187.349** | 0.524 | 0.158 | **1.438** | 3.96E-03 | **9.17E-03** |
| ENSG00000136144.14 | **RCBTB1** | 55213 | protein_coding | **229.268** | 0.464 | 0.140 | **1.379** | 3.96E-03 | **9.18E-03** |
| ENSG00000142937.14 | **RPS8** | 6202 | protein_coding | **49813.697** | -0.266 | -0.080 | **0.831** | 3.96E-03 | **9.18E-03** |
| ENSG00000235652.12 | **EPM2A-DT** | | lncRNA | **54.520** | 0.823 | 0.248 | **1.769** | 3.97E-03 | **9.19E-03** |
| ENSG00000187566.6 | **NHLRC1** | 378884 | protein_coding | **10.315** | 1.366 | 0.411 | **2.578** | 3.97E-03 | **9.20E-03** |
| ENSG00000138326.23 | **RPS24** | 6229 | protein_coding | **34114.715** | -0.338 | -0.102 | **0.791** | 3.97E-03 | **9.20E-03** |
| ENSG00000196712.21 | **NF1** | 4763 | protein_coding | **668.903** | 0.420 | 0.126 | **1.338** | 3.98E-03 | **9.21E-03** |
| ENSG00000198088.12 | **NUP62CL** | 54830 | protein_coding | **16.950** | -1.350 | -0.406 | **0.392** | 3.98E-03 | **9.22E-03** |
| ENSG00000253433.2 | **ENSG0000(** | 10192784! | lncRNA | **3.354** | 2.581 | 0.777 | **5.985** | 3.98E-03 | **9.22E-03** |
| ENSG00000107438.11 | **PDLIM1** | 9124 | protein_coding | **5485.164** | -0.411 | -0.124 | **0.752** | 3.99E-03 | **9.22E-03** |
| ENSG00000248489.5 | **CHD1-DT** | 10028923( | lncRNA | **72.395** | 0.663 | 0.199 | **1.583** | 3.99E-03 | **9.23E-03** |
| ENSG00000291064.2 | **ENSG00000291064** | | lncRNA | **119.160** | -0.507 | -0.153 | **0.704** | 3.99E-03 | **9.23E-03** |
| ENSG00000184304.17 | **PRKD1** | 5587 | protein_coding | **7.044** | 1.868 | 0.562 | **3.650** | 3.99E-03 | **9.23E-03** |
| ENSG00000270673.3 | **YTHDF3-DT** | | lncRNA | **23.707** | 0.811 | 0.244 | **1.754** | 3.99E-03 | **9.23E-03** |
| ENSG00000273382.1 | **TMEM167B-DT** | | lncRNA | **14.981** | 1.052 | 0.317 | **2.074** | 3.99E-03 | **9.24E-03** |
| ENSG00000307857.1 | **ENSG00000307857** | | lncRNA | **9.360** | 1.908 | 0.574 | **3.753** | 4.00E-03 | **9.25E-03** |
| ENSG00000272908.1 | **ENSG00000272908** | | lncRNA | **10.399** | 1.359 | 0.409 | **2.566** | 4.01E-03 | **9.26E-03** |
| ENSG00000163820.18 | **FYCO1** | 79443 | protein_coding | **355.643** | 0.304 | 0.091 | **1.234** | 4.01E-03 | **9.27E-03** |
| ENSG00000082146.15 | **STRADB** | 55437 | protein_coding | **238.065** | 0.430 | 0.130 | **1.348** | 4.01E-03 | **9.28E-03** |
| ENSG00000105538.12 | **RASIP1** | 54922 | protein_coding | **8.570** | 2.569 | 0.773 | **5.932** | 4.02E-03 | **9.29E-03** |
| ENSG00000147138.5 | **GPR174** | 84636 | protein_coding | **291.319** | -0.462 | -0.139 | **0.726** | 4.02E-03 | **9.29E-03** |
| ENSG00000168096.17 | **ANKS3** | 124401 | protein_coding | **504.239** | 0.383 | 0.115 | **1.304** | 4.03E-03 | **9.31E-03** |
| ENSG00000182013.20 | **PNMA8A** | 55228 | protein_coding | **15.717** | 1.550 | 0.467 | **2.929** | 4.03E-03 | **9.31E-03** |
| ENSG00000286019.5 | **NOTCH2NI** | 10099676 | protein_coding | **15.010** | 1.528 | 0.460 | **2.884** | 4.03E-03 | **9.31E-03** |

| | | | | | | | | | |
|---|---|---|---|---|---|---|---|---|---|
| ENSG00000135824.14 | **RGS8** | 85397 | protein_coding | **3.166** | 3.276 | 0.986 | **9.689** | 4.04E-03 | **9.33E-03** |
| ENSG00000226508.2 | **LINC01918** | 10537398 | lncRNA | **4.789** | 2.412 | 0.726 | **5.321** | 4.04E-03 | **9.34E-03** |
| ENSG00000224805.3 | **LINC00853** | 10087425 | lncRNA | **7.079** | 1.794 | 0.540 | **3.468** | 4.05E-03 | **9.35E-03** |
| ENSG00000078237.9 | **TIGAR** | 57103 | protein_coding | **276.735** | -0.500 | -0.150 | **0.707** | 4.05E-03 | **9.36E-03** |
| ENSG00000160799.15 | **CCDC12** | 151903 | protein_coding | **1227.044** | 0.272 | 0.082 | **1.208** | 4.05E-03 | **9.36E-03** |
| ENSG00000148399.15 | **DPH7** | 92715 | protein_coding | **958.105** | 0.408 | 0.123 | **1.327** | 4.06E-03 | **9.37E-03** |
| ENSG00000216937.15 | **CCDC7** | 79741 | protein_coding | **63.775** | -0.637 | -0.192 | **0.643** | 4.06E-03 | **9.38E-03** |
| ENSG00000176624.13 | **MEX3C** | 51320 | protein_coding | **248.782** | -0.591 | -0.178 | **0.664** | 4.08E-03 | **9.42E-03** |
| ENSG00000174564.15 | **IL20RB** | 53833 | protein_coding | **16.760** | -1.073 | -0.323 | **0.475** | 4.08E-03 | **9.42E-03** |
| ENSG00000131116.14 | **ZNF428** | 126299 | protein_coding | **549.198** | 0.323 | 0.097 | **1.251** | 4.08E-03 | **9.42E-03** |
| ENSG00000280383.1 | **ENSG00000280383** | | lncRNA | **19.726** | -1.039 | -0.313 | **0.487** | 4.10E-03 | **9.46E-03** |
| ENSG00000168779.21 | **SHOX2** | 6474 | protein_coding | **60.923** | 0.874 | 0.263 | **1.832** | 4.10E-03 | **9.46E-03** |
| ENSG00000170264.15 | **FAM161A** | 84140 | protein_coding | **84.652** | 0.538 | 0.162 | **1.452** | 4.10E-03 | **9.47E-03** |
| ENSG00000158691.18 | **ZSCAN12** | 9753 | protein_coding | **191.600** | -0.432 | -0.130 | **0.741** | 4.10E-03 | **9.47E-03** |
| ENSG00000214078.15 | **CPNE1** | 8904 | protein_coding | **3005.487** | 0.426 | 0.128 | **1.343** | 4.11E-03 | **9.48E-03** |
| ENSG00000149328.18 | **GLB1L2** | 89944 | protein_coding | **134.096** | 0.578 | 0.174 | **1.493** | 4.11E-03 | **9.49E-03** |
| ENSG00000290846.2 | **ENSG0000** | 107985911 | lncRNA | **138.932** | -0.490 | -0.147 | **0.712** | 4.12E-03 | **9.49E-03** |
| ENSG00000226359.1 | **ACTG1P24** | | processed_pseud | **5.829** | 1.731 | 0.521 | **3.321** | 4.12E-03 | **9.49E-03** |
| ENSG00000257524.6 | **ENSG00000257524** | | protein_coding | **322.628** | 0.487 | 0.147 | **1.402** | 4.12E-03 | **9.49E-03** |
| ENSG00000181004.11 | **BBS12** | 166379 | protein_coding | **30.701** | 0.787 | 0.237 | **1.725** | 4.13E-03 | **9.51E-03** |
| ENSG00000165046.15 | **LETM2** | 137994 | protein_coding | **66.543** | 0.571 | 0.172 | **1.486** | 4.13E-03 | **9.51E-03** |
| ENSG00000225968.9 | **ELFN1** | 392617 | protein_coding | **9.507** | 1.549 | 0.466 | **2.925** | 4.13E-03 | **9.52E-03** |
| ENSG00000140379.10 | **BCL2A1** | 597 | protein_coding | **1485.485** | 0.556 | 0.167 | **1.470** | 4.14E-03 | **9.53E-03** |
| ENSG00000124575.7 | **H1-3** | 3007 | protein_coding | **16.058** | -1.253 | -0.377 | **0.420** | 4.14E-03 | **9.54E-03** |
| ENSG00000243710.10 | **CFAP57** | 149465 | protein_coding | **149.165** | 0.517 | 0.156 | **1.431** | 4.15E-03 | **9.55E-03** |
| ENSG00000149043.18 | **SYT8** | 90019 | protein_coding | **5.165** | -2.846 | -0.857 | **0.139** | 4.15E-03 | **9.56E-03** |
| ENSG00000183741.14 | **CBX6** | 23466 | protein_coding | **1845.491** | 0.317 | 0.095 | **1.246** | 4.15E-03 | **9.57E-03** |
| ENSG00000118402.7 | **ELOVL4** | 6785 | protein_coding | **4.216** | 2.309 | 0.695 | **4.957** | 4.17E-03 | **9.59E-03** |
| ENSG00000147854.20 | **UHRF2** | 115426 | protein_coding | **942.972** | 0.444 | 0.134 | **1.360** | 4.17E-03 | **9.61E-03** |
| ENSG00000149548.17 | **CCDC15** | 80071 | protein_coding | **43.288** | -1.093 | -0.329 | **0.469** | 4.18E-03 | **9.63E-03** |
| ENSG00000300104.1 | **ENSG00000300104** | | lncRNA | **9.130** | 1.997 | 0.601 | **3.991** | 4.19E-03 | **9.65E-03** |

| | | | | | | | | | |
|---|---|---|---|---|---|---|---|---|---|
| ENSG00000118263.18 | **KLF7** | 8609 | protein_coding | **285.091** | 0.461 | 0.139 | **1.376** | 4.19E-03 | **9.65E-03** |
| ENSG00000167393.20 | **PPP2R3B** | 28227 | protein_coding | **576.017** | -0.459 | -0.138 | **0.727** | 4.19E-03 | **9.65E-03** |
| ENSG00000236502.4 | **SIX3-AS1** | 10050610ẟ | lncRNA | **13.389** | 1.582 | 0.476 | **2.993** | 4.21E-03 | **9.69E-03** |
| ENSG00000173812.13 | **EIF1** | 10209 | protein_coding | **12693.099** | -0.317 | -0.096 | **0.803** | 4.21E-03 | **9.69E-03** |
| ENSG00000140323.7 | **DISP2** | 85455 | protein_coding | **6.981** | 2.468 | 0.743 | **5.531** | 4.21E-03 | **9.69E-03** |
| ENSG00000234147.5 | **ENSG00000234147** | | lncRNA | **22.701** | 0.999 | 0.301 | **1.999** | 4.22E-03 | **9.70E-03** |
| ENSG00000256806.6 | **C17orf100** | 388327 | protein_coding | **73.618** | 0.561 | 0.169 | **1.476** | 4.22E-03 | **9.70E-03** |
| ENSG00000225434.6 | **LINC01504** | 10050754( | lncRNA | **8.381** | 1.528 | 0.460 | **2.884** | 4.23E-03 | **9.73E-03** |
| ENSG00000148057.18 | **IDNK** | 414328 | protein_coding | **184.388** | -0.351 | -0.106 | **0.784** | 4.24E-03 | **9.74E-03** |
| ENSG00000233791.8 | **BTG2-DT** | 730227 | lncRNA | **36.271** | 0.881 | 0.265 | **1.842** | 4.24E-03 | **9.74E-03** |
| ENSG00000161217.14 | **PCYT1A** | 5130 | protein_coding | **1220.367** | 0.303 | 0.091 | **1.234** | 4.25E-03 | **9.76E-03** |
| ENSG00000181751.12 | **MACIR** | 90355 | protein_coding | **205.347** | -0.399 | -0.120 | **0.759** | 4.25E-03 | **9.76E-03** |
| ENSG00000272699.1 | **ENSG00000272699** | | lncRNA | **7.610** | 1.597 | 0.481 | **3.026** | 4.25E-03 | **9.76E-03** |
| ENSG00000304314.1 | **ENSG00000304314** | | lncRNA | **4.464** | 2.254 | 0.678 | **4.769** | 4.26E-03 | **9.78E-03** |
| ENSG00000172465.16 | **TCEAL1** | 9338 | protein_coding | **122.521** | -0.444 | -0.134 | **0.735** | 4.26E-03 | **9.78E-03** |
| ENSG00000225470.10 | **JPX** | 554203 | lncRNA | **455.755** | -0.343 | -0.103 | **0.789** | 4.26E-03 | **9.79E-03** |
| ENSG00000075303.15 | **SLC25A40** | 55972 | protein_coding | **172.883** | -0.574 | -0.173 | **0.672** | 4.28E-03 | **9.82E-03** |
| ENSG00000160201.14 | **U2AF1** | 7307 | protein_coding | **2093.469** | -0.459 | -0.138 | **0.728** | 4.28E-03 | **9.83E-03** |
| ENSG00000169925.19 | **BRD3** | 8019 | protein_coding | **453.828** | 0.328 | 0.099 | **1.256** | 4.28E-03 | **9.83E-03** |
| ENSG00000079435.11 | **LIPE** | 3991 | protein_coding | **94.577** | -0.542 | -0.163 | **0.687** | 4.28E-03 | **9.83E-03** |
| ENSG00000156411.12 | **ATP5MJ** | 9556 | protein_coding | **1781.773** | -0.435 | -0.131 | **0.740** | 4.29E-03 | **9.84E-03** |
| ENSG00000258733.8 | **LINC02328** | | lncRNA | **44.730** | 0.633 | 0.191 | **1.551** | 4.30E-03 | **9.86E-03** |
| ENSG00000117308.17 | **GALE** | 2582 | protein_coding | **713.444** | -0.435 | -0.131 | **0.740** | 4.30E-03 | **9.86E-03** |
| ENSG00000197121.17 | **PGAP1** | 80055 | protein_coding | **37.674** | 0.742 | 0.223 | **1.672** | 4.31E-03 | **9.88E-03** |
| ENSG00000276368.2 | **H2AC14** | 8331 | protein_coding | **13.283** | -1.386 | -0.417 | **0.383** | 4.31E-03 | **9.89E-03** |
| ENSG00000003509.18 | **NDUFAF7** | 55471 | protein_coding | **474.878** | 0.362 | 0.109 | **1.285** | 4.32E-03 | **9.90E-03** |
| ENSG00000265781.1 | **ENSG00000265781** | | lncRNA | **11.042** | 1.596 | 0.481 | **3.024** | 4.32E-03 | **9.90E-03** |
| ENSG00000180998.14 | **GPR137C** | 283554 | protein_coding | **8.415** | 1.641 | 0.494 | **3.119** | 4.32E-03 | **9.91E-03** |
| ENSG00000261434.1 | **ENSG00000261434** | | lncRNA | **6.988** | -2.935 | -0.883 | **0.131** | 4.32E-03 | **9.92E-03** |
| ENSG00000100344.13 | **PNPLA3** | 80339 | protein_coding | **5.624** | 2.298 | 0.692 | **4.917** | 4.33E-03 | **9.92E-03** |
| ENSG00000309098.1 | **ENSG00000309098** | | lncRNA | **23.770** | -1.052 | -0.317 | **0.482** | 4.33E-03 | **9.93E-03** |

| | | | | | | | | | |
|---|---|---|---|---|---|---|---|---|---|
| ENSG00000276509.2 | **ENSG00000276509** | | lncRNA | **5.934** | 2.395 | 0.721 | **5.259** | 4.34E-03 | **9.94E-03** |
| ENSG00000284633.3 | **ENSG00000284633** | | lncRNA | **18.707** | 1.213 | 0.365 | **2.318** | 4.35E-03 | **9.97E-03** |
| ENSG00000255052.6 | **FAM66D** | 10013292З | lncRNA | **13.556** | 1.312 | 0.395 | **2.483** | 4.35E-03 | **9.97E-03** |
| ENSG00000242880.3 | **ENSG00000242880** | | lncRNA | **20.134** | 1.065 | 0.321 | **2.092** | 4.35E-03 | **9.97E-03** |
| ENSG00000135404.14 | **CD63** | 967 | protein_coding | **1848.933** | -0.396 | -0.119 | **0.760** | 4.36E-03 | **1.00E-02** |
| ENSG00000143756.13 | **FBXO28** | 23219 | protein_coding | **170.821** | -0.595 | -0.179 | **0.662** | 4.36E-03 | **1.00E-02** |
| ENSG00000112312.12 | **GMNN** | 51053 | protein_coding | **544.340** | -0.347 | -0.104 | **0.786** | 4.37E-03 | **1.00E-02** |
| ENSG00000221838.14 | **AP4M1** | 9179 | protein_coding | **186.036** | -0.409 | -0.123 | **0.753** | 4.38E-03 | **1.00E-02** |
| ENSG00000287382.2 | **EXOC1-AS1** | 10537766З | lncRNA | **18.994** | -1.291 | -0.388 | **0.409** | 4.39E-03 | **1.01E-02** |
| ENSG00000100614.20 | **PPM1A** | 5494 | protein_coding | **1005.840** | -0.318 | -0.096 | **0.802** | 4.40E-03 | **1.01E-02** |
| ENSG00000254802.2 | **ENSG00000254802** | | lncRNA | **79.466** | 0.729 | 0.219 | **1.657** | 4.40E-03 | **1.01E-02** |
| ENSG00000189410.14 | **SH2D5** | 400745 | protein_coding | **36.865** | 0.983 | 0.296 | **1.976** | 4.40E-03 | **1.01E-02** |
| ENSG00000175773.16 | **ZBTB44-DT** | | lncRNA | **12.647** | 1.263 | 0.380 | **2.400** | 4.40E-03 | **1.01E-02** |
| ENSG00000109445.13 | **ZNF330** | 27309 | protein_coding | **740.629** | -0.431 | -0.130 | **0.742** | 4.40E-03 | **1.01E-02** |
| ENSG00000074935.16 | **TUBE1** | 51175 | protein_coding | **152.597** | -0.616 | -0.185 | **0.653** | 4.41E-03 | **1.01E-02** |
| ENSG00000137203.15 | **TFAP2A** | 7020 | protein_coding | **29.927** | 1.378 | 0.415 | **2.600** | 4.41E-03 | **1.01E-02** |
| ENSG00000168813.19 | **ZNF507** | 22847 | protein_coding | **304.066** | -0.409 | -0.123 | **0.753** | 4.41E-03 | **1.01E-02** |
| ENSG00000117425.16 | **PTCH2** | 8643 | protein_coding | **25.094** | 1.007 | 0.303 | **2.010** | 4.42E-03 | **1.01E-02** |
| ENSG00000274653.1 | **ENSG00000274653** | | lncRNA | **10.356** | 1.432 | 0.431 | **2.698** | 4.42E-03 | **1.01E-02** |
| ENSG00000029153.17 | **BMAL2** | 56938 | protein_coding | **925.127** | -0.436 | -0.131 | **0.739** | 4.42E-03 | **1.01E-02** |
| ENSG00000165757.10 | **JCAD** | 57608 | protein_coding | **4.969** | 2.236 | 0.673 | **4.710** | 4.42E-03 | **1.01E-02** |
| ENSG00000156265.19 | **MAP3K7CL** | 56911 | protein_coding | **185.684** | 0.416 | 0.125 | **1.334** | 4.44E-03 | **1.02E-02** |
| ENSG00000281091.4 | **LL0XNC01-** | 100506901 | lncRNA | **34.715** | -0.730 | -0.220 | **0.603** | 4.45E-03 | **1.02E-02** |
| ENSG00000152104.13 | **PTPN14** | 5784 | protein_coding | **41.085** | 1.253 | 0.377 | **2.383** | 4.45E-03 | **1.02E-02** |
| ENSG00000198954.11 | **KIFBP** | 26128 | protein_coding | **293.105** | 0.352 | 0.106 | **1.276** | 4.46E-03 | **1.02E-02** |
| ENSG00000214367.11 | **HAUS3** | 79441 | protein_coding | **265.998** | -0.398 | -0.120 | **0.759** | 4.46E-03 | **1.02E-02** |
| ENSG00000120686.14 | **UFM1** | 51569 | protein_coding | **988.238** | -0.322 | -0.097 | **0.800** | 4.47E-03 | **1.02E-02** |
| ENSG00000233154.6 | **LINC01762** | 10192902З | lncRNA | **5.662** | -1.955 | -0.589 | **0.258** | 4.47E-03 | **1.02E-02** |
| ENSG00000157107.16 | **FCHO2** | 115548 | protein_coding | **10.848** | 1.395 | 0.420 | **2.630** | 4.48E-03 | **1.02E-02** |
| ENSG00000167515.13 | **TRAPPC2L** | 51693 | protein_coding | **801.164** | -0.356 | -0.107 | **0.781** | 4.49E-03 | **1.03E-02** |
| ENSG00000233862.9 | **ENSG00000233862** | | lncRNA | **5.882** | 1.714 | 0.516 | **3.280** | 4.50E-03 | **1.03E-02** |

| | | | | | | | | | |
|---|---|---|---|---|---|---|---|---|---|
| ENSG00000160886.14 | **LY6K** | 54742 | protein_coding | **7.331** | -2.292 | -0.690 | **0.204** | 4.51E-03 | **1.03E-02** |
| ENSG00000290592.2 | **ENSG00000290592** | | lncRNA | **6.636** | 2.559 | 0.770 | **5.892** | 4.51E-03 | **1.03E-02** |
| ENSG00000162849.17 | **KIF26B** | 55083 | protein_coding | **40.079** | -1.011 | -0.304 | **0.496** | 4.51E-03 | **1.03E-02** |
| ENSG00000108406.12 | **DHX40** | 79665 | protein_coding | **442.369** | 0.335 | 0.101 | **1.262** | 4.51E-03 | **1.03E-02** |
| ENSG00000285825.3 | **CCDC15-D1** | 105369547 | lncRNA | **27.986** | 0.879 | 0.265 | **1.840** | 4.53E-03 | **1.03E-02** |
| ENSG00000250790.4 | **ENSG00000250790** | | lncRNA | **11.738** | 1.151 | 0.347 | **2.221** | 4.53E-03 | **1.03E-02** |
| ENSG00000145734.21 | **BDP1** | 55814 | protein_coding | **195.618** | -0.546 | -0.164 | **0.685** | 4.53E-03 | **1.03E-02** |
| ENSG00000100647.10 | **SUSD6** | 9766 | protein_coding | **998.378** | 0.384 | 0.116 | **1.305** | 4.53E-03 | **1.03E-02** |
| ENSG00000174365.23 | **SNHG11** | 128439 | lncRNA | **423.042** | 0.442 | 0.133 | **1.359** | 4.54E-03 | **1.04E-02** |
| ENSG00000179406.11 | **LINC00174** | 285908 | lncRNA | **654.348** | 0.370 | 0.111 | **1.293** | 4.55E-03 | **1.04E-02** |
| ENSG00000277383.3 | **BICRA-AS2** | 12866722[illegible] | lncRNA | **36.559** | 0.755 | 0.227 | **1.688** | 4.56E-03 | **1.04E-02** |
| ENSG00000105926.18 | **PALS2** | 51678 | protein_coding | **282.112** | -0.494 | -0.149 | **0.710** | 4.57E-03 | **1.04E-02** |
| ENSG00000183044.14 | **ABAT** | 18 | protein_coding | **107.703** | 0.506 | 0.152 | **1.420** | 4.57E-03 | **1.04E-02** |
| ENSG00000163738.21 | **MTHFD2L** | 441024 | protein_coding | **111.770** | -0.543 | -0.164 | **0.686** | 4.57E-03 | **1.04E-02** |
| ENSG00000197976.14 | **AKAP17A** | 8227 | protein_coding | **3648.776** | 0.354 | 0.106 | **1.278** | 4.58E-03 | **1.04E-02** |
| ENSG00000286835.1 | **ENSG00000286835** | | lncRNA | **29.692** | 0.752 | 0.226 | **1.684** | 4.59E-03 | **1.05E-02** |
| ENSG00000182362.16 | **YBEY** | 54059 | protein_coding | **400.578** | 0.382 | 0.115 | **1.303** | 4.61E-03 | **1.05E-02** |
| ENSG00000197951.13 | **ZNF71** | 58491 | protein_coding | **98.367** | -0.446 | -0.134 | **0.734** | 4.62E-03 | **1.05E-02** |
| ENSG00000100351.19 | **GRAP2** | 9402 | protein_coding | **152.041** | -0.837 | -0.252 | **0.560** | 4.63E-03 | **1.05E-02** |
| ENSG00000163900.13 | **TMEM41A** | 90407 | protein_coding | **309.449** | -0.293 | -0.088 | **0.816** | 4.64E-03 | **1.06E-02** |
| ENSG00000178917.19 | **ZNF852** | 285346 | protein_coding | **255.586** | -0.344 | -0.104 | **0.788** | 4.64E-03 | **1.06E-02** |
| ENSG00000182240.18 | **BACE2** | 25825 | protein_coding | **29.576** | 0.886 | 0.267 | **1.848** | 4.65E-03 | **1.06E-02** |
| ENSG00000282386.1 | **ENSG00000282386** | | lncRNA | **16.918** | 1.010 | 0.304 | **2.014** | 4.65E-03 | **1.06E-02** |
| ENSG00000187186.17 | **ENSG0000(** | 730098 | protein_coding | **58.271** | 0.777 | 0.234 | **1.713** | 4.65E-03 | **1.06E-02** |
| ENSG00000204839.10 | **MROH6** | 642475 | protein_coding | **168.581** | 0.652 | 0.196 | **1.572** | 4.65E-03 | **1.06E-02** |
| ENSG00000272343.1 | **ENSG00000272343** | | lncRNA | **9.689** | 1.422 | 0.428 | **2.680** | 4.65E-03 | **1.06E-02** |
| ENSG00000291173.2 | **CASP4LP** | 643733 | lncRNA | **7.018** | -2.140 | -0.644 | **0.227** | 4.65E-03 | **1.06E-02** |
| ENSG00000299973.1 | **ENSG00000299973** | | lncRNA | **55.913** | 0.946 | 0.285 | **1.927** | 4.67E-03 | **1.06E-02** |
| ENSG00000151303.13 | **ENSG00000151303** | | lncRNA | **10.222** | 1.788 | 0.538 | **3.454** | 4.68E-03 | **1.06E-02** |
| ENSG00000165806.23 | **CASP7** | 840 | protein_coding | **361.645** | 0.403 | 0.121 | **1.322** | 4.68E-03 | **1.07E-02** |
| ENSG00000132692.21 | **BCAN** | 63827 | protein_coding | **10.218** | 1.533 | 0.462 | **2.894** | 4.69E-03 | **1.07E-02** |

| | | | | | | | | | |
|---|---|---|---|---|---|---|---|---|---|
| ENSG00000122778.12 | **KIAA1549** | 57670 | protein_coding | **52.082** | 0.655 | 0.197 | **1.575** | 4.70E-03 | **1.07E-02** |
| ENSG00000289047.3 | **ENSG0000(** | 10029414! | lncRNA | **263.647** | -0.450 | -0.136 | **0.732** | 4.70E-03 | **1.07E-02** |
| ENSG00000007264.17 | **MATK** | 4145 | protein_coding | **10.183** | 1.503 | 0.453 | **2.835** | 4.70E-03 | **1.07E-02** |
| ENSG00000134590.14 | **RTL8C** | 8933 | protein_coding | **592.103** | 0.408 | 0.123 | **1.326** | 4.71E-03 | **1.07E-02** |
| ENSG00000131042.17 | **LILRB2** | 10288 | protein_coding | **21.695** | 1.133 | 0.341 | **2.193** | 4.71E-03 | **1.07E-02** |
| ENSG00000303580.1 | **ENSG00000303580** | | lncRNA | **7.458** | 3.110 | 0.936 | **8.632** | 4.72E-03 | **1.07E-02** |
| ENSG00000136286.18 | **MYO1G** | 64005 | protein_coding | **6169.520** | -0.453 | -0.136 | **0.731** | 4.73E-03 | **1.07E-02** |
| ENSG00000160588.11 | **MPZL3** | 196264 | protein_coding | **92.325** | -0.633 | -0.190 | **0.645** | 4.73E-03 | **1.08E-02** |
| ENSG00000151491.16 | **EPS8** | 2059 | protein_coding | **147.736** | -0.717 | -0.216 | **0.608** | 4.73E-03 | **1.08E-02** |
| ENSG00000138769.13 | **CDKL2** | 8999 | protein_coding | **8.048** | 1.930 | 0.581 | **3.809** | 4.74E-03 | **1.08E-02** |
| ENSG00000233775.2 | **ENSG00000233775** | | lncRNA | **5.649** | 1.921 | 0.578 | **3.787** | 4.74E-03 | **1.08E-02** |
| ENSG00000083223.20 | **TUT7** | 79670 | protein_coding | **1054.960** | 0.329 | 0.099 | **1.256** | 4.75E-03 | **1.08E-02** |
| ENSG00000156345.20 | **CDK20** | 23552 | protein_coding | **5.747** | 1.792 | 0.539 | **3.463** | 4.76E-03 | **1.08E-02** |
| ENSG00000001084.15 | **GCLC** | 2729 | protein_coding | **200.186** | 0.555 | 0.167 | **1.469** | 4.76E-03 | **1.08E-02** |
| ENSG00000083535.18 | **PIBF1** | 10464 | protein_coding | **195.657** | -0.498 | -0.150 | **0.708** | 4.77E-03 | **1.08E-02** |
| ENSG00000077150.23 | **NFKB2** | 4791 | protein_coding | **8891.009** | 0.410 | 0.123 | **1.328** | 4.77E-03 | **1.08E-02** |
| ENSG00000153896.20 | **ZNF599** | 148103 | protein_coding | **63.557** | 0.543 | 0.163 | **1.457** | 4.77E-03 | **1.08E-02** |
| ENSG00000153531.16 | **ADPRHL1** | 113622 | protein_coding | **22.717** | 1.103 | 0.332 | **2.149** | 4.78E-03 | **1.08E-02** |
| ENSG00000080493.21 | **SLC4A4** | 8671 | protein_coding | **9.153** | 1.779 | 0.536 | **3.432** | 4.78E-03 | **1.08E-02** |
| ENSG00000150527.20 | **MIA2** | 4253 | protein_coding | **479.713** | -0.458 | -0.138 | **0.728** | 4.78E-03 | **1.08E-02** |
| ENSG00000064933.21 | **PMS1** | 5378 | protein_coding | **344.622** | -0.340 | -0.102 | **0.790** | 4.78E-03 | **1.09E-02** |
| ENSG00000204524.8 | **ZNF805** | 390980 | protein_coding | **79.085** | -0.572 | -0.172 | **0.673** | 4.79E-03 | **1.09E-02** |
| ENSG00000184281.17 | **TSSC4** | 10078 | protein_coding | **412.998** | -0.302 | -0.091 | **0.811** | 4.79E-03 | **1.09E-02** |
| ENSG00000256654.5 | **ENSG00000256654** | | lncRNA | **9.242** | 1.601 | 0.482 | **3.033** | 4.80E-03 | **1.09E-02** |
| ENSG00000174628.19 | **IQCK** | 124152 | protein_coding | **26.258** | 0.963 | 0.290 | **1.949** | 4.81E-03 | **1.09E-02** |
| ENSG00000133065.13 | **SLC41A1** | 254428 | protein_coding | **123.994** | 0.578 | 0.174 | **1.493** | 4.81E-03 | **1.09E-02** |
| ENSG00000150054.21 | **MPP7** | 143098 | protein_coding | **10.870** | 1.376 | 0.414 | **2.595** | 4.82E-03 | **1.09E-02** |
| ENSG00000302646.1 | **ENSG00000302646** | | lncRNA | **8.608** | 1.637 | 0.493 | **3.111** | 4.83E-03 | **1.09E-02** |
| ENSG00000082701.19 | **GSK3B** | 2932 | protein_coding | **837.222** | -0.362 | -0.109 | **0.778** | 4.83E-03 | **1.09E-02** |
| ENSG00000151320.13 | **AKAP6** | 9472 | protein_coding | **27.476** | 1.320 | 0.397 | **2.496** | 4.84E-03 | **1.10E-02** |
| ENSG00000273137.1 | **SELENOO-AS1** | | lncRNA | **4.681** | 2.352 | 0.708 | **5.104** | 4.85E-03 | **1.10E-02** |

| | | | | | | | | | |
|---|---|---|---|---|---|---|---|---|---|
| ENSG00000078668.16 | **VDAC3** | 7419 | protein_coding | **2680.980** | -0.326 | -0.098 | **0.798** | 4.87E-03 | **1.10E-02** |
| ENSG00000158571.13 | **PFKFB1** | 5207 | protein_coding | **49.909** | 0.686 | 0.207 | **1.609** | 4.89E-03 | **1.11E-02** |
| ENSG00000111801.18 | **BTN3A3** | 10384 | protein_coding | **767.590** | -0.282 | -0.085 | **0.822** | 4.89E-03 | **1.11E-02** |
| ENSG00000130348.13 | **QRSL1** | 55278 | protein_coding | **392.216** | -0.371 | -0.112 | **0.773** | 4.89E-03 | **1.11E-02** |
| ENSG00000213799.15 | **ZNF845** | 91664 | protein_coding | **99.610** | -0.426 | -0.128 | **0.744** | 4.89E-03 | **1.11E-02** |
| ENSG00000137944.19 | **KYAT3** | 56267 | protein_coding | **356.179** | -0.427 | -0.129 | **0.744** | 4.89E-03 | **1.11E-02** |
| ENSG00000175395.18 | **ZNF25** | 219749 | protein_coding | **102.518** | -0.447 | -0.135 | **0.733** | 4.90E-03 | **1.11E-02** |
| ENSG00000130052.15 | **STARD8** | 9754 | protein_coding | **17.676** | 1.094 | 0.329 | **2.135** | 4.90E-03 | **1.11E-02** |
| ENSG00000307699.1 | **ENSG00000307699** | | lncRNA | **28.777** | 0.840 | 0.253 | **1.790** | 4.90E-03 | **1.11E-02** |
| ENSG00000179178.13 | **TMEM125** | 128218 | protein_coding | **4.638** | 2.381 | 0.717 | **5.210** | 4.91E-03 | **1.11E-02** |
| ENSG00000250666.2 | **LINC01596** | 10537761 | lncRNA | **13.406** | -1.576 | -0.474 | **0.335** | 4.91E-03 | **1.11E-02** |
| ENSG00000223648.4 | **IGHV3-64** | | IG_V_gene | **8.840** | 2.788 | 0.839 | **6.907** | 4.91E-03 | **1.11E-02** |
| ENSG00000184402.16 | **SS18L1** | 26039 | protein_coding | **495.591** | 0.395 | 0.119 | **1.315** | 4.92E-03 | **1.11E-02** |
| ENSG00000102908.25 | **NFAT5** | 10725 | protein_coding | **516.779** | 0.538 | 0.162 | **1.452** | 4.93E-03 | **1.12E-02** |
| ENSG00000269955.2 | **FMC1-LUC** | 10099692 | protein_coding | **231.675** | -0.699 | -0.210 | **0.616** | 4.94E-03 | **1.12E-02** |
| ENSG00000104331.11 | **BPNT2** | 54928 | protein_coding | **371.085** | -0.368 | -0.111 | **0.775** | 4.94E-03 | **1.12E-02** |
| ENSG00000275936.1 | **ENSG00000275936** | | lncRNA | **15.387** | 1.045 | 0.315 | **2.063** | 4.94E-03 | **1.12E-02** |
| ENSG00000167112.12 | **TRUB2** | 26995 | protein_coding | **455.876** | -0.337 | -0.101 | **0.792** | 4.95E-03 | **1.12E-02** |
| ENSG00000223573.8 | **TINCR** | 257000 | protein_coding | **30.218** | 1.053 | 0.317 | **2.075** | 4.96E-03 | **1.12E-02** |
| ENSG00000163093.14 | **BBS5** | 129880 | protein_coding | **73.043** | 0.517 | 0.156 | **1.431** | 4.96E-03 | **1.12E-02** |
| ENSG00000165934.15 | **CPSF2** | 53981 | protein_coding | **712.409** | -0.646 | -0.195 | **0.639** | 4.96E-03 | **1.12E-02** |
| ENSG00000164751.17 | **PEX2** | 5828 | protein_coding | **525.830** | -0.507 | -0.153 | **0.704** | 4.97E-03 | **1.12E-02** |
| ENSG00000105669.16 | **COPE** | 11316 | protein_coding | **3569.500** | -0.423 | -0.127 | **0.746** | 4.98E-03 | **1.13E-02** |
| ENSG00000165156.17 | **ZHX1** | 11244 | protein_coding | **176.484** | -0.517 | -0.156 | **0.699** | 4.98E-03 | **1.13E-02** |
| ENSG00000212916.6 | **MAP10** | 54627 | protein_coding | **52.368** | 0.566 | 0.170 | **1.480** | 4.99E-03 | **1.13E-02** |
| ENSG00000168234.16 | **TTC39C** | 125488 | protein_coding | **854.745** | 0.311 | 0.094 | **1.240** | 5.00E-03 | **1.13E-02** |
| ENSG00000137628.20 | **DDX60** | 55601 | protein_coding | **253.934** | -0.612 | -0.184 | **0.654** | 5.01E-03 | **1.13E-02** |
| ENSG00000272967.1 | **ENSG00000272967** | | lncRNA | **6.204** | 1.655 | 0.498 | **3.148** | 5.01E-03 | **1.13E-02** |
| ENSG00000186522.17 | **SEPTIN10** | 151011 | protein_coding | **202.102** | 0.558 | 0.168 | **1.472** | 5.02E-03 | **1.13E-02** |
| ENSG00000104964.17 | **TLE5** | 166 | protein_coding | **7634.655** | -0.394 | -0.119 | **0.761** | 5.02E-03 | **1.13E-02** |
| ENSG00000118197.16 | **DDX59** | 83479 | protein_coding | **401.490** | -0.321 | -0.097 | **0.801** | 5.02E-03 | **1.13E-02** |

| | | | | | | | | | |
|---|---|---|---|---|---|---|---|---|---|
| ENSG00000011451.23 | **WIZ** | 58525 | protein_coding | **751.551** | -0.266 | -0.080 | **0.832** | 5.07E-03 | **1.14E-02** |
| ENSG00000124164.18 | **VAPB** | 9217 | protein_coding | **532.499** | -0.308 | -0.093 | **0.808** | 5.07E-03 | **1.14E-02** |
| ENSG00000163743.16 | **RCHY1** | 25898 | protein_coding | **111.752** | -0.621 | -0.187 | **0.650** | 5.07E-03 | **1.15E-02** |
| ENSG00000176946.15 | **THAP4** | 51078 | protein_coding | **680.859** | -0.304 | -0.092 | **0.810** | 5.08E-03 | **1.15E-02** |
| ENSG00000010310.11 | **GIPR** | 2696 | protein_coding | **1000.096** | 0.355 | 0.107 | **1.279** | 5.08E-03 | **1.15E-02** |
| ENSG00000305387.1 | **ENSG00000305387** | | lncRNA | **15.104** | 1.199 | 0.361 | **2.296** | 5.08E-03 | **1.15E-02** |
| ENSG00000288591.1 | **ENSG00000288591** | | protein_coding | **68.207** | 1.348 | 0.406 | **2.546** | 5.09E-03 | **1.15E-02** |
| ENSG00000254635.7 | **WAC-AS1** | | lncRNA | **428.420** | 0.389 | 0.117 | **1.310** | 5.10E-03 | **1.15E-02** |
| ENSG00000261801.9 | **LOXL1-AS1** | 10028761€ | lncRNA | **5.435** | 2.797 | 0.842 | **6.952** | 5.10E-03 | **1.15E-02** |
| ENSG00000228784.10 | **LINC00954** | 400946 | lncRNA | **57.337** | -0.595 | -0.179 | **0.662** | 5.10E-03 | **1.15E-02** |
| ENSG00000258609.5 | **LINC-ROR** | 10088577§ | lncRNA | **14.912** | 1.390 | 0.418 | **2.621** | 5.11E-03 | **1.15E-02** |
| ENSG00000273141.1 | **ENSG00000273141** | | lncRNA | **6.371** | 1.752 | 0.527 | **3.368** | 5.12E-03 | **1.15E-02** |
| ENSG00000105865.13 | **DUS4L** | 11062 | protein_coding | **105.425** | -0.438 | -0.132 | **0.738** | 5.13E-03 | **1.16E-02** |
| ENSG00000180257.16 | **ZNF816** | 125893 | protein_coding | **121.413** | -0.444 | -0.134 | **0.735** | 5.13E-03 | **1.16E-02** |
| ENSG00000287558.2 | **ENSG00000287558** | | lncRNA | **8.476** | 2.052 | 0.618 | **4.147** | 5.15E-03 | **1.16E-02** |
| ENSG00000212127.6 | **TAS2R14** | 50840 | protein_coding | **26.089** | 0.855 | 0.257 | **1.808** | 5.16E-03 | **1.16E-02** |
| ENSG00000204370.16 | **SDHD** | 6392 | protein_coding | **876.303** | -0.290 | -0.087 | **0.818** | 5.17E-03 | **1.16E-02** |
| ENSG00000241860.8 | **ENSG00000241860** | | lncRNA | **92.830** | 0.603 | 0.181 | **1.519** | 5.17E-03 | **1.16E-02** |
| ENSG00000140044.15 | **JDP2** | 122953 | protein_coding | **390.036** | -0.496 | -0.149 | **0.709** | 5.17E-03 | **1.16E-02** |
| ENSG00000163399.18 | **ATP1A1** | 476 | protein_coding | **7023.277** | -0.348 | -0.105 | **0.786** | 5.17E-03 | **1.17E-02** |
| ENSG00000289304.3 | **ENSG00000289304** | | lncRNA | **9.064** | 1.432 | 0.431 | **2.698** | 5.20E-03 | **1.17E-02** |
| ENSG00000151229.15 | **SLC2A13** | 114134 | protein_coding | **49.514** | 0.674 | 0.203 | **1.595** | 5.20E-03 | **1.17E-02** |
| ENSG00000167881.17 | **SRP68** | 6730 | protein_coding | **1028.653** | -0.294 | -0.089 | **0.815** | 5.21E-03 | **1.17E-02** |
| ENSG00000254923.1 | **ENSG00000254923** | | processed_pseudo | **6.610** | 2.107 | 0.634 | **4.309** | 5.22E-03 | **1.18E-02** |
| ENSG00000137216.21 | **TMEM63B** | 55362 | protein_coding | **487.727** | 0.380 | 0.114 | **1.301** | 5.24E-03 | **1.18E-02** |
| ENSG00000153404.17 | **PLEKHG4B** | 153478 | protein_coding | **9.416** | 2.368 | 0.713 | **5.162** | 5.25E-03 | **1.18E-02** |
| ENSG00000111850.13 | **SMIM8** | 57150 | protein_coding | **172.466** | 0.382 | 0.115 | **1.303** | 5.25E-03 | **1.18E-02** |
| ENSG00000128951.16 | **DUT** | 1854 | protein_coding | **1186.049** | -0.369 | -0.111 | **0.774** | 5.25E-03 | **1.18E-02** |
| ENSG00000273183.1 | **ENSG00000273183** | | lncRNA | **24.069** | 0.931 | 0.280 | **1.906** | 5.25E-03 | **1.18E-02** |
| ENSG00000104983.10 | **CCDC61** | 729440 | protein_coding | **507.289** | 0.328 | 0.099 | **1.255** | 5.26E-03 | **1.18E-02** |
| ENSG00000122986.16 | **HVCN1** | 84329 | protein_coding | **3446.263** | -0.361 | -0.109 | **0.779** | 5.26E-03 | **1.18E-02** |

| | | | | | | | | | |
|---|---|---|---|---|---|---|---|---|---|
| ENSG00000154099.20 | **DNAAF1** | 123872 | protein_coding | **10.302** | 1.995 | 0.600 | **3.985** | 5.26E-03 | **1.18E-02** |
| ENSG00000138613.16 | **APH1B** | 83464 | protein_coding | **234.881** | -0.450 | -0.135 | **0.732** | 5.26E-03 | **1.18E-02** |
| ENSG00000126945.11 | **HNRNPH2** | 3188 | protein_coding | **695.205** | 0.560 | 0.169 | **1.474** | 5.28E-03 | **1.19E-02** |
| ENSG00000103522.19 | **IL21R** | 50615 | protein_coding | **2240.802** | -0.391 | -0.118 | **0.763** | 5.28E-03 | **1.19E-02** |
| ENSG00000259112.2 | **NDUFC2-K(** | 10053272( | protein_coding | **282.764** | -0.488 | -0.147 | **0.713** | 5.28E-03 | **1.19E-02** |
| ENSG00000279170.4 | **TSTD3** | 10013089( | transcribed_unita | **40.392** | -0.664 | -0.200 | **0.631** | 5.29E-03 | **1.19E-02** |
| ENSG00000280832.4 | **GSEC** | | lncRNA | **51.950** | -0.663 | -0.200 | **0.632** | 5.29E-03 | **1.19E-02** |
| ENSG00000101871.19 | **MID1** | 4281 | protein_coding | **5.057** | -2.947 | -0.887 | **0.130** | 5.29E-03 | **1.19E-02** |
| ENSG00000290015.2 | **ENSG00000290015** | | lncRNA | **17.664** | 0.960 | 0.289 | **1.946** | 5.29E-03 | **1.19E-02** |
| ENSG00000306285.1 | **ENSG00000306285** | | lncRNA | **27.123** | 1.068 | 0.321 | **2.096** | 5.30E-03 | **1.19E-02** |
| ENSG00000268889.3 | **LIM2-AS1** | | lncRNA | **6.070** | -1.774 | -0.534 | **0.292** | 5.31E-03 | **1.19E-02** |
| ENSG00000145934.18 | **TENM2** | 57451 | protein_coding | **8.362** | 1.911 | 0.575 | **3.761** | 5.31E-03 | **1.19E-02** |
| ENSG00000293318.1 | **ENSG00000293318** | | lncRNA | **3.936** | 2.626 | 0.791 | **6.175** | 5.32E-03 | **1.19E-02** |
| ENSG00000273447.1 | **ENSG00000273447** | | lncRNA | **5.920** | 1.791 | 0.539 | **3.461** | 5.32E-03 | **1.20E-02** |
| ENSG00000256043.7 | **CTSO** | 1519 | protein_coding | **120.195** | -0.635 | -0.191 | **0.644** | 5.34E-03 | **1.20E-02** |
| ENSG00000189114.10 | **BLOC1S3** | 388552 | protein_coding | **322.703** | -0.402 | -0.121 | **0.757** | 5.34E-03 | **1.20E-02** |
| ENSG00000100926.17 | **TM9SF1** | 10548 | protein_coding | **350.124** | -0.377 | -0.113 | **0.770** | 5.35E-03 | **1.20E-02** |
| ENSG00000141556.24 | **TBCD** | 6904 | protein_coding | **1410.514** | -0.276 | -0.083 | **0.826** | 5.35E-03 | **1.20E-02** |
| ENSG00000279333.1 | **ENSG00000279333** | | TEC | **5.016** | 1.998 | 0.601 | **3.994** | 5.36E-03 | **1.20E-02** |
| ENSG00000173221.16 | **GLRX** | 2745 | protein_coding | **1857.243** | -0.269 | -0.081 | **0.830** | 5.36E-03 | **1.20E-02** |
| ENSG00000150977.14 | **RILPL2** | 196383 | protein_coding | **2827.856** | -0.285 | -0.086 | **0.820** | 5.37E-03 | **1.20E-02** |
| ENSG00000101052.15 | **IFT52** | 51098 | protein_coding | **241.275** | 0.334 | 0.100 | **1.260** | 5.37E-03 | **1.21E-02** |
| ENSG00000183828.17 | **NUDT14** | 256281 | protein_coding | **214.720** | 0.522 | 0.157 | **1.436** | 5.37E-03 | **1.21E-02** |
| ENSG00000162419.15 | **GMEB1** | 10691 | protein_coding | **218.377** | -0.388 | -0.117 | **0.764** | 5.38E-03 | **1.21E-02** |
| ENSG00000088179.11 | **PTPN4** | 5775 | protein_coding | **147.088** | -0.523 | -0.157 | **0.696** | 5.39E-03 | **1.21E-02** |
| ENSG00000121104.9 | **FAM117A** | 81558 | protein_coding | **1135.311** | 0.306 | 0.092 | **1.236** | 5.40E-03 | **1.21E-02** |
| ENSG00000136026.16 | **CKAP4** | 10970 | protein_coding | **565.345** | -0.439 | -0.132 | **0.738** | 5.40E-03 | **1.21E-02** |
| ENSG00000293141.2 | **ENSG00000293141** | | lncRNA | **331.132** | 0.408 | 0.123 | **1.326** | 5.40E-03 | **1.21E-02** |
| ENSG00000304625.1 | **ENSG00000304625** | | lncRNA | **9.177** | 1.528 | 0.460 | **2.883** | 5.42E-03 | **1.22E-02** |
| ENSG00000235831.9 | **BHLHE40-A** | 10050758Z | lncRNA | **38.676** | 0.657 | 0.198 | **1.577** | 5.42E-03 | **1.22E-02** |
| ENSG00000106263.21 | **EIF3B** | 8662 | protein_coding | **3754.969** | -0.263 | -0.079 | **0.833** | 5.43E-03 | **1.22E-02** |

| | | | | | | | | | |
|---|---|---|---|---|---|---|---|---|---|
| ENSG00000272434.2 | **ENSG00000272434** | | lncRNA | **6.771** | 1.684 | 0.507 | **3.213** | 5.43E-03 | **1.22E-02** |
| ENSG00000166747.15 | **AP1G1** | 164 | protein_coding | **1445.338** | 0.473 | 0.142 | **1.388** | 5.44E-03 | **1.22E-02** |
| ENSG00000290537.2 | **ENSG00000290537** | | lncRNA | **71.670** | 0.555 | 0.167 | **1.469** | 5.45E-03 | **1.22E-02** |
| ENSG00000100731.17 | **PCNX1** | 22990 | protein_coding | **771.345** | -0.335 | -0.101 | **0.793** | 5.45E-03 | **1.22E-02** |
| ENSG00000235106.13 | **BRD3OS** | 266655 | protein_coding | **256.741** | -0.490 | -0.147 | **0.712** | 5.47E-03 | **1.22E-02** |
| ENSG00000183484.14 | **GPR132** | 29933 | protein_coding | **618.551** | 0.473 | 0.142 | **1.388** | 5.47E-03 | **1.23E-02** |
| ENSG00000230303.7 | **ENSG00000230303** | | lncRNA | **5.434** | 2.133 | 0.642 | **4.386** | 5.47E-03 | **1.23E-02** |
| ENSG00000263874.4 | **LASP1NB** | 10050557( | protein_coding | **18.536** | 0.980 | 0.295 | **1.973** | 5.47E-03 | **1.23E-02** |
| ENSG00000271335.7 | **CCNY-AS1** | | lncRNA | **67.790** | 0.514 | 0.155 | **1.428** | 5.48E-03 | **1.23E-02** |
| ENSG00000134278.18 | **SPIRE1** | 56907 | protein_coding | **100.047** | 0.676 | 0.203 | **1.597** | 5.48E-03 | **1.23E-02** |
| ENSG00000173757.13 | **STAT5B** | 6777 | protein_coding | **1384.919** | -0.288 | -0.087 | **0.819** | 5.50E-03 | **1.23E-02** |
| ENSG00000291053.2 | **CATSPER2I** | 440278 | lncRNA | **32.242** | 0.752 | 0.226 | **1.684** | 5.50E-03 | **1.23E-02** |
| ENSG00000130702.16 | **LAMA5** | 3911 | protein_coding | **453.956** | 0.527 | 0.159 | **1.441** | 5.51E-03 | **1.23E-02** |
| ENSG00000265817.5 | **FSBP** | 10086141z | protein_coding | **154.299** | -0.571 | -0.172 | **0.673** | 5.51E-03 | **1.23E-02** |
| ENSG00000184009.15 | **ACTG1** | 71 | protein_coding | **45256.063** | -0.416 | -0.125 | **0.749** | 5.52E-03 | **1.23E-02** |
| ENSG00000114166.10 | **KAT2B** | 8850 | protein_coding | **177.227** | -0.523 | -0.157 | **0.696** | 5.54E-03 | **1.24E-02** |
| ENSG00000288983.3 | **ENSG00000288983** | | lncRNA | **37.831** | 0.621 | 0.187 | **1.538** | 5.56E-03 | **1.24E-02** |
| ENSG00000289286.2 | **ENSG00000289286** | | lncRNA | **4.678** | 1.959 | 0.590 | **3.888** | 5.58E-03 | **1.25E-02** |
| ENSG00000108771.15 | **DHX58** | 79132 | protein_coding | **876.284** | -0.456 | -0.137 | **0.729** | 5.59E-03 | **1.25E-02** |
| ENSG00000197713.17 | **RPE** | 6120 | protein_coding | **487.378** | -0.519 | -0.156 | **0.698** | 5.59E-03 | **1.25E-02** |
| ENSG00000116016.16 | **EPAS1** | 2034 | protein_coding | **28.769** | 1.045 | 0.315 | **2.064** | 5.60E-03 | **1.25E-02** |
| ENSG00000302545.1 | **ENSG00000302545** | | lncRNA | **8.025** | -1.412 | -0.425 | **0.376** | 5.60E-03 | **1.25E-02** |
| ENSG00000079691.21 | **CARMIL1** | 55604 | protein_coding | **160.579** | 0.482 | 0.145 | **1.397** | 5.60E-03 | **1.25E-02** |
| ENSG00000198836.13 | **OPA1** | 4976 | protein_coding | **759.813** | -0.585 | -0.176 | **0.667** | 5.60E-03 | **1.25E-02** |
| ENSG00000213588.8 | **ZBTB9** | 221504 | protein_coding | **176.594** | 0.397 | 0.119 | **1.316** | 5.61E-03 | **1.25E-02** |
| ENSG00000230606.12 | **APPAT** | | lncRNA | **117.882** | -0.477 | -0.144 | **0.718** | 5.61E-03 | **1.25E-02** |
| ENSG00000181649.10 | **PHLDA2** | 7262 | protein_coding | **33.495** | 0.967 | 0.291 | **1.955** | 5.62E-03 | **1.26E-02** |
| ENSG00000152454.5 | **ZNF256** | 10172 | protein_coding | **164.341** | -0.407 | -0.122 | **0.754** | 5.63E-03 | **1.26E-02** |
| ENSG00000068976.16 | **PYGM** | 5837 | protein_coding | **30.180** | 0.957 | 0.288 | **1.942** | 5.63E-03 | **1.26E-02** |
| ENSG00000100167.21 | **SEPTIN3** | 55964 | protein_coding | **186.518** | 0.674 | 0.203 | **1.595** | 5.64E-03 | **1.26E-02** |
| ENSG00000092051.20 | **JPH4** | 84502 | protein_coding | **78.891** | 0.752 | 0.226 | **1.684** | 5.65E-03 | **1.26E-02** |

| | | | | | | | | | |
|---|---|---|---|---|---|---|---|---|---|
| ENSG00000289260.3 | **ENSG00000289260** | | lncRNA | **34.345** | -0.851 | -0.256 | **0.554** | 5.66E-03 | **1.26E-02** |
| ENSG00000119401.13 | **TRIM32** | 22954 | protein_coding | **167.772** | -0.361 | -0.109 | **0.779** | 5.66E-03 | **1.26E-02** |
| ENSG00000130717.15 | **UCK1** | 83549 | protein_coding | **418.408** | -0.290 | -0.087 | **0.818** | 5.66E-03 | **1.26E-02** |
| ENSG00000141582.17 | **CBX4** | 8535 | protein_coding | **657.615** | 0.434 | 0.131 | **1.351** | 5.67E-03 | **1.27E-02** |
| ENSG00000163697.20 | **APBB2** | 323 | protein_coding | **246.039** | -0.470 | -0.142 | **0.722** | 5.68E-03 | **1.27E-02** |
| ENSG00000142444.8 | **TIMM29** | 90580 | protein_coding | **190.713** | -0.382 | -0.115 | **0.767** | 5.70E-03 | **1.27E-02** |
| ENSG00000148053.20 | **NTRK2** | 4915 | protein_coding | **238.893** | 0.604 | 0.182 | **1.520** | 5.70E-03 | **1.27E-02** |
| ENSG00000267498.3 | **UQCRFS1-DT** | | lncRNA | **5.832** | 1.862 | 0.561 | **3.635** | 5.70E-03 | **1.27E-02** |
| ENSG00000289007.3 | **LINC03112** | 10192809 | lncRNA | **116.360** | 0.419 | 0.126 | **1.337** | 5.71E-03 | **1.27E-02** |
| ENSG00000081803.19 | **CADPS2** | 93664 | protein_coding | **40.348** | -1.145 | -0.345 | **0.452** | 5.71E-03 | **1.27E-02** |
| ENSG00000277453.1 | **ENSG00000277453** | | lncRNA | **6.190** | 1.678 | 0.505 | **3.201** | 5.73E-03 | **1.28E-02** |
| ENSG00000275601.1 | **ENSG00000275601** | | lncRNA | **18.910** | 0.982 | 0.296 | **1.976** | 5.73E-03 | **1.28E-02** |
| ENSG00000172005.13 | **MAL** | 4118 | protein_coding | **409.245** | 0.644 | 0.194 | **1.562** | 5.74E-03 | **1.28E-02** |
| ENSG00000229422.2 | **ENSG00000229422** | | lncRNA | **5.270** | 2.024 | 0.609 | **4.068** | 5.75E-03 | **1.28E-02** |
| ENSG00000127720.10 | **METTL25** | 84190 | protein_coding | **60.661** | -0.625 | -0.188 | **0.648** | 5.75E-03 | **1.28E-02** |
| ENSG00000197043.17 | **ANXA6** | 309 | protein_coding | **11422.670** | 0.309 | 0.093 | **1.239** | 5.76E-03 | **1.28E-02** |
| ENSG00000162976.14 | **SLC66A3** | 130814 | protein_coding | **97.812** | 0.445 | 0.134 | **1.361** | 5.77E-03 | **1.29E-02** |
| ENSG00000171823.9 | **FBXL14** | 144699 | protein_coding | **71.806** | 0.771 | 0.232 | **1.706** | 5.79E-03 | **1.29E-02** |
| ENSG00000149823.12 | **VPS51** | 738 | protein_coding | **1697.905** | -0.408 | -0.123 | **0.754** | 5.79E-03 | **1.29E-02** |
| ENSG00000290562.2 | **ENSG00000290562** | | lncRNA | **5.244** | 2.781 | 0.837 | **6.871** | 5.80E-03 | **1.29E-02** |
| ENSG00000109184.17 | **DCUN1D4** | 23142 | protein_coding | **404.854** | -0.432 | -0.130 | **0.741** | 5.80E-03 | **1.29E-02** |
| ENSG00000166529.18 | **ZSCAN21** | 7589 | protein_coding | **207.585** | 0.409 | 0.123 | **1.327** | 5.81E-03 | **1.29E-02** |
| ENSG00000146587.20 | **RBAK** | 57786 | protein_coding | **262.839** | -0.362 | -0.109 | **0.778** | 5.81E-03 | **1.29E-02** |
| ENSG00000178026.15 | **LRRC75B** | 388886 | protein_coding | **128.269** | 0.776 | 0.234 | **1.712** | 5.81E-03 | **1.29E-02** |
| ENSG00000231312.9 | **MAP4K3-DT** | | lncRNA | **94.331** | 0.524 | 0.158 | **1.438** | 5.81E-03 | **1.29E-02** |
| ENSG00000139239.7 | **RPL14P1** | | processed_pseud | **35.853** | -0.672 | -0.202 | **0.627** | 5.81E-03 | **1.29E-02** |
| ENSG00000225614.4 | **ZNF469** | 84627 | protein_coding | **26.024** | -0.793 | -0.239 | **0.577** | 5.82E-03 | **1.30E-02** |
| ENSG00000185024.20 | **BRF1** | 2972 | protein_coding | **767.872** | 0.321 | 0.097 | **1.249** | 5.83E-03 | **1.30E-02** |
| ENSG00000087111.24 | **PIGS** | 94005 | protein_coding | **712.992** | 0.365 | 0.110 | **1.288** | 5.83E-03 | **1.30E-02** |
| ENSG00000131389.20 | **SLC6A6** | 6533 | protein_coding | **439.615** | -0.452 | -0.136 | **0.731** | 5.84E-03 | **1.30E-02** |
| ENSG00000110696.12 | **C11orf58** | 10944 | protein_coding | **1463.559** | -0.414 | -0.125 | **0.751** | 5.84E-03 | **1.30E-02** |

| | | | | | | | | | |
|---|---|---|---|---|---|---|---|---|---|
| ENSG00000149346.17 | **SLX4IP** | 128710 | protein_coding | **284.151** | 0.371 | 0.112 | **1.293** | 5.85E-03 | **1.30E-02** |
| ENSG00000198015.16 | **MRPL42** | 28977 | protein_coding | **761.852** | -0.354 | -0.106 | **0.783** | 5.86E-03 | **1.30E-02** |
| ENSG00000258667.5 | **ENSG00000258667** | | lncRNA | **187.823** | -0.606 | -0.183 | **0.657** | 5.86E-03 | **1.30E-02** |
| ENSG00000233975.3 | **LINC02574** | | lncRNA | **13.251** | 1.408 | 0.424 | **2.654** | 5.87E-03 | **1.30E-02** |
| ENSG00000051620.13 | **HEBP2** | 23593 | protein_coding | **427.680** | 0.488 | 0.147 | **1.402** | 5.87E-03 | **1.31E-02** |
| ENSG00000156603.22 | **MED19** | 219541 | protein_coding | **216.443** | 0.415 | 0.125 | **1.334** | 5.88E-03 | **1.31E-02** |
| ENSG00000137449.17 | **CPEB2** | 132864 | protein_coding | **124.628** | -0.787 | -0.237 | **0.579** | 5.88E-03 | **1.31E-02** |
| ENSG00000130244.15 | **FAM98C** | 147965 | protein_coding | **272.545** | -0.424 | -0.128 | **0.745** | 5.88E-03 | **1.31E-02** |
| ENSG00000246792.3 | **ENSG0000(** | 12490671‹ | lncRNA | **6.268** | -1.679 | -0.505 | **0.312** | 5.88E-03 | **1.31E-02** |
| ENSG00000128829.14 | **EIF2AK4** | 440275 | protein_coding | **873.748** | 0.355 | 0.107 | **1.279** | 5.88E-03 | **1.31E-02** |
| ENSG00000288886.3 | **ENSG00000288886** | | lncRNA | **64.102** | -0.815 | -0.245 | **0.568** | 5.88E-03 | **1.31E-02** |
| ENSG00000240429.1 | **LRRFIP1P1** | | processed_pseud( | **25.987** | 1.115 | 0.336 | **2.166** | 5.89E-03 | **1.31E-02** |
| ENSG00000112210.14 | **RAB23** | 51715 | protein_coding | **60.685** | 0.849 | 0.255 | **1.801** | 5.91E-03 | **1.31E-02** |
| ENSG00000198081.13 | **ZBTB14** | 7541 | protein_coding | **221.358** | -0.511 | -0.154 | **0.702** | 5.91E-03 | **1.31E-02** |
| ENSG00000169981.13 | **ZNF35** | 7584 | protein_coding | **109.237** | -0.431 | -0.130 | **0.742** | 5.91E-03 | **1.31E-02** |
| ENSG00000278214.2 | **LINC02139** | 400548 | lncRNA | **3.511** | 3.180 | 0.957 | **9.066** | 5.94E-03 | **1.32E-02** |
| ENSG00000162736.20 | **NCSTN** | 23385 | protein_coding | **1171.360** | -0.296 | -0.089 | **0.815** | 5.95E-03 | **1.32E-02** |
| ENSG00000066084.15 | **DIP2B** | 57609 | protein_coding | **491.922** | -0.476 | -0.143 | **0.719** | 5.97E-03 | **1.33E-02** |
| ENSG00000080709.18 | **KCNN2** | 3781 | protein_coding | **7.183** | 1.733 | 0.522 | **3.325** | 5.99E-03 | **1.33E-02** |
| ENSG00000273249.2 | **WDR5-DT** | | lncRNA | **21.995** | 0.887 | 0.267 | **1.849** | 5.99E-03 | **1.33E-02** |
| ENSG00000100206.13 | **DMC1** | 11144 | protein_coding | **44.343** | -0.932 | -0.281 | **0.524** | 6.00E-03 | **1.33E-02** |
| ENSG00000160746.15 | **ANO10** | 55129 | protein_coding | **243.252** | 0.429 | 0.129 | **1.346** | 6.00E-03 | **1.33E-02** |
| ENSG00000065135.14 | **GNAI3** | 2773 | protein_coding | **695.298** | -0.399 | -0.120 | **0.758** | 6.01E-03 | **1.33E-02** |
| ENSG00000113300.15 | **CNOT6** | 57472 | protein_coding | **527.620** | 0.332 | 0.100 | **1.259** | 6.02E-03 | **1.34E-02** |
| ENSG00000134744.16 | **TUT4** | 23318 | protein_coding | **974.745** | 0.303 | 0.091 | **1.234** | 6.02E-03 | **1.34E-02** |
| ENSG00000178502.7 | **KLHL11** | 55175 | protein_coding | **66.157** | -0.531 | -0.160 | **0.692** | 6.02E-03 | **1.34E-02** |
| ENSG00000152503.12 | **TRIM36** | 55521 | protein_coding | **6.323** | 1.850 | 0.557 | **3.605** | 6.04E-03 | **1.34E-02** |
| ENSG00000032444.17 | **PNPLA6** | 10908 | protein_coding | **1310.433** | 0.417 | 0.126 | **1.335** | 6.05E-03 | **1.34E-02** |
| ENSG00000163156.14 | **SCNM1** | 79005 | protein_coding | **701.761** | -0.329 | -0.099 | **0.796** | 6.05E-03 | **1.34E-02** |
| ENSG00000177697.21 | **CD151** | 977 | protein_coding | **442.493** | 0.450 | 0.135 | **1.366** | 6.05E-03 | **1.34E-02** |
| ENSG00000132205.13 | **EMILIN2** | 84034 | protein_coding | **292.381** | -0.354 | -0.107 | **0.782** | 6.06E-03 | **1.34E-02** |

| | | | | | | | | | |
|---|---|---|---|---|---|---|---|---|---|
| ENSG00000111641.15 | **NOP2** | 4839 | protein_coding | **1448.446** | -0.378 | -0.114 | **0.770** | 6.06E-03 | **1.34E-02** |
| ENSG00000285968.2 | **ENSG00000285968** | | lncRNA | **14.931** | 1.287 | 0.387 | **2.440** | 6.06E-03 | **1.34E-02** |
| ENSG00000228873.2 | **ENSG00000228873** | | lncRNA | **4.322** | 2.432 | 0.732 | **5.398** | 6.07E-03 | **1.35E-02** |
| ENSG00000125868.18 | **DSTN** | 11034 | protein_coding | **678.724** | 0.313 | 0.094 | **1.242** | 6.08E-03 | **1.35E-02** |
| ENSG00000196866.3 | **H2AC7** | 3013 | protein_coding | **5.306** | -3.743 | -1.127 | **0.075** | 6.08E-03 | **1.35E-02** |
| ENSG00000196735.15 | **HLA-DQA1** | 3117 | protein_coding | **14629.644** | 0.513 | 0.154 | **1.427** | 6.08E-03 | **1.35E-02** |
| ENSG00000139725.9 | **RHOF** | 54509 | protein_coding | **1586.931** | 0.428 | 0.129 | **1.345** | 6.09E-03 | **1.35E-02** |
| ENSG00000139842.17 | **CUL4A** | 8451 | protein_coding | **1336.737** | 0.587 | 0.177 | **1.502** | 6.11E-03 | **1.35E-02** |
| ENSG00000018699.15 | **TTC27** | 55622 | protein_coding | **252.684** | -0.530 | -0.160 | **0.692** | 6.11E-03 | **1.35E-02** |
| ENSG00000262412.2 | **ENSG00000262412** | | lncRNA | **8.274** | 1.473 | 0.443 | **2.776** | 6.11E-03 | **1.35E-02** |
| ENSG00000271851.1 | **ENSG00000271851** | | lncRNA | **17.781** | 0.867 | 0.261 | **1.823** | 6.12E-03 | **1.35E-02** |
| ENSG00000147586.12 | **MRPS28** | 28957 | protein_coding | **524.064** | 0.282 | 0.085 | **1.216** | 6.12E-03 | **1.35E-02** |
| ENSG00000214389.2 | **RPS3AP26** | | processed_pseudo | **135.454** | 0.886 | 0.267 | **1.848** | 6.12E-03 | **1.35E-02** |
| ENSG00000259725.2 | **ENSG00000259725** | | lncRNA | **2.881** | 3.707 | 1.116 | **13.058** | 6.15E-03 | **1.36E-02** |
| ENSG00000134982.21 | **APC** | 324 | protein_coding | **182.827** | -0.505 | -0.152 | **0.705** | 6.16E-03 | **1.36E-02** |
| ENSG00000204290.12 | **BTNL2** | 56244 | protein_coding | **5.739** | 1.857 | 0.559 | **3.622** | 6.16E-03 | **1.36E-02** |
| ENSG00000229017.8 | **LINC01277** | | lncRNA | **11.706** | 1.533 | 0.461 | **2.893** | 6.17E-03 | **1.36E-02** |
| ENSG00000139372.16 | **TDG** | 6996 | protein_coding | **469.758** | -0.284 | -0.086 | **0.821** | 6.17E-03 | **1.36E-02** |
| ENSG00000089234.18 | **BRAP** | 8315 | protein_coding | **406.350** | -0.338 | -0.102 | **0.791** | 6.17E-03 | **1.37E-02** |
| ENSG00000287024.2 | **ENSG0000(** | 10537333 | lncRNA | **13.199** | 1.430 | 0.430 | **2.694** | 6.18E-03 | **1.37E-02** |
| ENSG00000198964.16 | **SGMS1** | 259230 | protein_coding | **527.703** | 0.493 | 0.148 | **1.407** | 6.19E-03 | **1.37E-02** |
| ENSG00000243811.14 | **APOBEC3D** | 140564 | protein_coding | **555.850** | -0.391 | -0.118 | **0.762** | 6.19E-03 | **1.37E-02** |
| ENSG00000179523.8 | **EIF3J-DT** | 645212 | lncRNA | **166.803** | 0.377 | 0.114 | **1.299** | 6.20E-03 | **1.37E-02** |
| ENSG00000150316.14 | **CWC15** | 51503 | protein_coding | **1317.287** | -0.289 | -0.087 | **0.818** | 6.20E-03 | **1.37E-02** |
| ENSG00000270441.1 | **LAMB2P1** | | unprocessed_pseu | **12.062** | 1.103 | 0.332 | **2.149** | 6.21E-03 | **1.37E-02** |
| ENSG00000227885.5 | **ENSG0000(** | 12490133 | lncRNA | **6.976** | 1.861 | 0.560 | **3.632** | 6.22E-03 | **1.37E-02** |
| ENSG00000148832.18 | **PAOX** | 196743 | protein_coding | **296.102** | -0.340 | -0.102 | **0.790** | 6.23E-03 | **1.38E-02** |
| ENSG00000215045.9 | **GRID2IP** | 392862 | protein_coding | **7.564** | 1.761 | 0.530 | **3.389** | 6.25E-03 | **1.38E-02** |
| ENSG00000125869.12 | **LAMP5** | 24141 | protein_coding | **240.435** | -0.530 | -0.159 | **0.693** | 6.26E-03 | **1.38E-02** |
| ENSG00000259687.5 | **LINC01220** | | lncRNA | **5.071** | 2.502 | 0.753 | **5.665** | 6.26E-03 | **1.38E-02** |
| ENSG00000250132.7 | **ENSG00000250132** | | lncRNA | **9.011** | 1.347 | 0.405 | **2.543** | 6.27E-03 | **1.38E-02** |

| | | | | | | | | | |
|---|---|---|---|---|---|---|---|---|---|
| ENSG00000114013.17 | **CD86** | 942 | protein_coding | **983.523** | 0.421 | 0.127 | **1.339** | 6.27E-03 | **1.38E-02** |
| ENSG00000272853.3 | **SUV39H2-DT** | | lncRNA | **23.677** | 0.861 | 0.259 | **1.817** | 6.27E-03 | **1.38E-02** |
| ENSG00000180535.6 | **BHLHA15** | 168620 | protein_coding | **155.988** | -0.651 | -0.196 | **0.637** | 6.27E-03 | **1.38E-02** |
| ENSG00000078269.16 | **SYNJ2** | 8871 | protein_coding | **115.232** | -0.463 | -0.139 | **0.726** | 6.28E-03 | **1.38E-02** |
| ENSG00000179532.14 | **DNHD1** | 144132 | protein_coding | **1437.233** | -0.319 | -0.096 | **0.802** | 6.28E-03 | **1.39E-02** |
| ENSG00000129317.17 | **PUS7L** | 83448 | protein_coding | **200.704** | -0.391 | -0.118 | **0.763** | 6.29E-03 | **1.39E-02** |
| ENSG00000267855.8 | **NDUFA7** | 4701 | protein_coding | **1234.152** | -0.385 | -0.116 | **0.766** | 6.30E-03 | **1.39E-02** |
| ENSG00000269194.2 | **ENSG0000(** | 10537243! | lncRNA | **6.758** | 1.741 | 0.524 | **3.343** | 6.30E-03 | **1.39E-02** |
| ENSG00000286786.1 | **ENSG00000286786** | | lncRNA | **6.753** | 1.738 | 0.523 | **3.336** | 6.31E-03 | **1.39E-02** |
| ENSG00000160190.16 | **SLC37A1** | 54020 | protein_coding | **770.957** | 0.328 | 0.099 | **1.255** | 6.31E-03 | **1.39E-02** |
| ENSG00000196981.5 | **WDR5B** | 54554 | protein_coding | **122.949** | 0.434 | 0.131 | **1.351** | 6.32E-03 | **1.39E-02** |
| ENSG00000273472.3 | **ENSG00000273472** | | lncRNA | **9.542** | 1.764 | 0.531 | **3.396** | 6.32E-03 | **1.39E-02** |
| ENSG00000168101.17 | **NUDT16L1** | 84309 | protein_coding | **328.612** | -0.369 | -0.111 | **0.774** | 6.32E-03 | **1.39E-02** |
| ENSG00000008516.21 | **MMP25** | 64386 | protein_coding | **18.113** | 0.966 | 0.291 | **1.954** | 6.33E-03 | **1.40E-02** |
| ENSG00000169926.13 | **KLF13** | 51621 | protein_coding | **2022.907** | 0.305 | 0.092 | **1.235** | 6.34E-03 | **1.40E-02** |
| ENSG00000272368.2 | **ENSG00000272368** | | lncRNA | **14.497** | 1.008 | 0.304 | **2.012** | 6.34E-03 | **1.40E-02** |
| ENSG00000304092.1 | **ENSG00000304092** | | lncRNA | **4.851** | 2.377 | 0.715 | **5.194** | 6.35E-03 | **1.40E-02** |
| ENSG00000164970.17 | **FAM219A** | 203259 | protein_coding | **105.296** | 0.546 | 0.164 | **1.460** | 6.35E-03 | **1.40E-02** |
| ENSG00000069329.21 | **VPS35** | 55737 | protein_coding | **1086.867** | -0.270 | -0.081 | **0.829** | 6.35E-03 | **1.40E-02** |
| ENSG00000000003.18 | **TSPAN6** | 7105 | protein_coding | **7.248** | 1.833 | 0.552 | **3.564** | 6.35E-03 | **1.40E-02** |
| ENSG00000295937.1 | **ENSG00000295937** | | lncRNA | **21.917** | 1.268 | 0.382 | **2.408** | 6.36E-03 | **1.40E-02** |
| ENSG00000164880.18 | **INTS1** | 26173 | protein_coding | **2032.046** | 0.341 | 0.103 | **1.266** | 6.36E-03 | **1.40E-02** |
| ENSG00000078967.15 | **UBE2D4** | 51619 | protein_coding | **268.581** | 0.295 | 0.089 | **1.227** | 6.37E-03 | **1.40E-02** |
| ENSG00000175279.23 | **CENPS** | 378708 | protein_coding | **109.448** | -0.401 | -0.121 | **0.757** | 6.39E-03 | **1.41E-02** |
| ENSG00000088256.10 | **GNA11** | 2767 | protein_coding | **212.536** | 0.567 | 0.171 | **1.482** | 6.42E-03 | **1.41E-02** |
| ENSG00000121988.20 | **ZRANB3** | 84083 | protein_coding | **91.018** | -0.512 | -0.154 | **0.701** | 6.43E-03 | **1.42E-02** |
| ENSG00000289253.3 | **ENSG00000289253** | | lncRNA | **6.905** | 1.476 | 0.444 | **2.781** | 6.44E-03 | **1.42E-02** |
| ENSG00000257702.6 | **LBX2-AS1** | 151534 | lncRNA | **77.005** | -0.756 | -0.228 | **0.592** | 6.45E-03 | **1.42E-02** |
| ENSG00000128567.19 | **PODXL** | 5420 | protein_coding | **24.980** | 0.949 | 0.286 | **1.930** | 6.45E-03 | **1.42E-02** |
| ENSG00000123200.19 | **ZC3H13** | 23091 | protein_coding | **774.273** | -0.271 | -0.082 | **0.829** | 6.46E-03 | **1.42E-02** |
| ENSG00000213445.13 | **SIPA1** | 6494 | protein_coding | **2699.406** | -0.532 | -0.160 | **0.692** | 6.46E-03 | **1.42E-02** |

| | | | | | | | | | |
|---|---|---|---|---|---|---|---|---|---|
| ENSG00000101844.20 | **ATG4A** | 115201 | protein_coding | **234.895** | 0.495 | 0.149 | **1.409** | 6.46E-03 | **1.42E-02** |
| ENSG00000204149.17 | **AGAP6** | 414189 | protein_coding | **1289.452** | 0.291 | 0.088 | **1.224** | 6.47E-03 | **1.42E-02** |
| ENSG00000305126.1 | **ENSG00000305126** | | lncRNA | **68.893** | 0.773 | 0.233 | **1.709** | 6.48E-03 | **1.43E-02** |
| ENSG00000140319.13 | **SRP14** | 6727 | protein_coding | **3911.710** | -0.353 | -0.106 | **0.783** | 6.49E-03 | **1.43E-02** |
| ENSG00000116741.9 | **RGS2** | 5997 | protein_coding | **220.441** | 0.532 | 0.160 | **1.446** | 6.49E-03 | **1.43E-02** |
| ENSG00000101470.11 | **TNNC2** | 7125 | protein_coding | **8.784** | 1.697 | 0.511 | **3.243** | 6.49E-03 | **1.43E-02** |
| ENSG00000132436.15 | **FIGNL1** | 63979 | protein_coding | **327.759** | -0.470 | -0.142 | **0.722** | 6.49E-03 | **1.43E-02** |
| ENSG00000196408.14 | **NOXO1** | 124056 | protein_coding | **14.097** | 1.461 | 0.440 | **2.753** | 6.49E-03 | **1.43E-02** |
| ENSG00000290027.2 | **ENSG00000290027** | | lncRNA | **43.518** | -0.711 | -0.214 | **0.611** | 6.50E-03 | **1.43E-02** |
| ENSG00000272669.1 | **ENSG00000272669** | | lncRNA | **151.210** | -0.580 | -0.175 | **0.669** | 6.51E-03 | **1.43E-02** |
| ENSG00000167778.11 | **SPRYD3** | 84926 | protein_coding | **428.634** | -0.374 | -0.113 | **0.771** | 6.51E-03 | **1.43E-02** |
| ENSG00000249241.3 | **LINC02265** | 10106049 | lncRNA | **7.020** | 1.603 | 0.483 | **3.038** | 6.51E-03 | **1.43E-02** |
| ENSG00000214026.13 | **MRPL23** | 6150 | protein_coding | **1873.250** | -0.421 | -0.127 | **0.747** | 6.52E-03 | **1.43E-02** |
| ENSG00000135469.16 | **COQ10A** | 93058 | protein_coding | **142.125** | 0.425 | 0.128 | **1.343** | 6.55E-03 | **1.44E-02** |
| ENSG00000296013.1 | **ENSG00000296013** | | lncRNA | **5.795** | -2.130 | -0.641 | **0.228** | 6.56E-03 | **1.44E-02** |
| ENSG00000111731.15 | **C2CD5** | 9847 | protein_coding | **434.059** | -0.381 | -0.115 | **0.768** | 6.58E-03 | **1.44E-02** |
| ENSG00000184840.14 | **TMED9** | 54732 | protein_coding | **2042.076** | -0.383 | -0.115 | **0.767** | 6.58E-03 | **1.44E-02** |
| ENSG00000130589.16 | **HELZ2** | 85441 | protein_coding | **1037.208** | -0.468 | -0.141 | **0.723** | 6.58E-03 | **1.45E-02** |
| ENSG00000108510.12 | **MED13** | 9969 | protein_coding | **413.533** | -0.453 | -0.136 | **0.731** | 6.59E-03 | **1.45E-02** |
| ENSG00000164933.13 | **SLC25A32** | 81034 | protein_coding | **275.267** | -0.353 | -0.106 | **0.783** | 6.61E-03 | **1.45E-02** |
| ENSG00000113520.12 | **IL4** | 3565 | protein_coding | **5.239** | -2.014 | -0.606 | **0.248** | 6.61E-03 | **1.45E-02** |
| ENSG00000180979.12 | **LRRC57** | 255252 | protein_coding | **197.700** | -0.349 | -0.105 | **0.785** | 6.62E-03 | **1.45E-02** |
| ENSG00000012223.15 | **LTF** | 4057 | protein_coding | **6.564** | 2.554 | 0.769 | **5.873** | 6.62E-03 | **1.45E-02** |
| ENSG00000146530.17 | **VWDE** | 221806 | protein_coding | **10.690** | 1.653 | 0.498 | **3.145** | 6.62E-03 | **1.45E-02** |
| ENSG00000077721.19 | **UBE2A** | 7319 | protein_coding | **749.299** | -0.310 | -0.093 | **0.807** | 6.63E-03 | **1.45E-02** |
| ENSG00000254876.6 | **SUGT1P4-S** | 10049948 | lncRNA | **6.383** | 1.739 | 0.523 | **3.337** | 6.63E-03 | **1.45E-02** |
| ENSG00000171045.18 | **TSNARE1** | 203062 | protein_coding | **348.470** | -0.265 | -0.080 | **0.832** | 6.64E-03 | **1.46E-02** |
| ENSG00000038274.20 | **MAT2B** | 27430 | protein_coding | **868.798** | 0.462 | 0.139 | **1.377** | 6.65E-03 | **1.46E-02** |
| ENSG00000213999.19 | **MEF2B** | 10027184 | protein_coding | **1381.823** | -0.589 | -0.177 | **0.665** | 6.67E-03 | **1.46E-02** |
| ENSG00000166579.18 | **NDEL1** | 81565 | protein_coding | **506.235** | -0.330 | -0.099 | **0.796** | 6.68E-03 | **1.46E-02** |
| ENSG00000272525.3 | **BTF3-DT** | | lncRNA | **29.081** | 0.867 | 0.261 | **1.824** | 6.68E-03 | **1.46E-02** |

| | | | | | | | | | |
|---|---|---|---|---|---|---|---|---|---|
| ENSG00000166471.13 | **TMEM41B** | 440026 | protein_coding | **382.179** | 0.394 | 0.119 | **1.314** | 6.68E-03 | **1.46E-02** |
| ENSG00000082781.14 | **ITGB5** | 3693 | protein_coding | **225.094** | 0.566 | 0.170 | **1.480** | 6.69E-03 | **1.47E-02** |
| ENSG00000133027.20 | **PEMT** | 10400 | protein_coding | **205.343** | -0.469 | -0.141 | **0.722** | 6.69E-03 | **1.47E-02** |
| ENSG00000250564.3 | **LINC02999** | | lncRNA | **7.923** | 1.456 | 0.438 | **2.744** | 6.70E-03 | **1.47E-02** |
| ENSG00000050393.14 | **MCUR1** | 63933 | protein_coding | **489.785** | -0.446 | -0.134 | **0.734** | 6.70E-03 | **1.47E-02** |
| ENSG00000152926.19 | **ZNF117** | 51351 | protein_coding | **213.340** | 0.503 | 0.151 | **1.417** | 6.71E-03 | **1.47E-02** |
| ENSG00000149679.14 | **CABLES2** | 81928 | protein_coding | **290.733** | -0.381 | -0.115 | **0.768** | 6.71E-03 | **1.47E-02** |
| ENSG00000253559.2 | **OSGEPL1-A** | 10140925{ | lncRNA | **14.020** | 1.011 | 0.304 | **2.015** | 6.72E-03 | **1.47E-02** |
| ENSG00000233016.10 | **SNHG7** | 84973 | lncRNA | **1184.893** | -0.604 | -0.182 | **0.658** | 6.74E-03 | **1.48E-02** |
| ENSG00000099992.18 | **TBC1D10A** | 83874 | protein_coding | **295.163** | 0.404 | 0.122 | **1.323** | 6.75E-03 | **1.48E-02** |
| ENSG00000121797.12 | **CCRL2** | 9034 | protein_coding | **17.985** | 1.014 | 0.305 | **2.020** | 6.75E-03 | **1.48E-02** |
| ENSG00000175115.14 | **PACS1** | 55690 | protein_coding | **2662.685** | 0.339 | 0.102 | **1.265** | 6.77E-03 | **1.48E-02** |
| ENSG00000247746.7 | **USP51** | 158880 | protein_coding | **33.798** | -0.739 | -0.222 | **0.599** | 6.78E-03 | **1.48E-02** |
| ENSG00000168434.15 | **COG7** | 91949 | protein_coding | **431.164** | -0.323 | -0.097 | **0.800** | 6.80E-03 | **1.49E-02** |
| ENSG00000163590.15 | **PPM1L** | 151742 | protein_coding | **14.315** | 1.203 | 0.362 | **2.302** | 6.85E-03 | **1.50E-02** |
| ENSG00000170166.7 | **HOXD4** | 3233 | protein_coding | **5.887** | 2.033 | 0.612 | **4.093** | 6.87E-03 | **1.50E-02** |
| ENSG00000198944.6 | **SOWAHA** | 134548 | protein_coding | **8.004** | 1.711 | 0.515 | **3.274** | 6.89E-03 | **1.51E-02** |
| ENSG00000173991.7 | **TCAP** | 8557 | protein_coding | **23.374** | 1.000 | 0.301 | **2.000** | 6.89E-03 | **1.51E-02** |
| ENSG00000138640.17 | **FAM13A** | 10144 | protein_coding | **162.645** | 0.559 | 0.168 | **1.473** | 6.90E-03 | **1.51E-02** |
| ENSG00000247774.10 | **PCED1B-AS** | 10023320{ | lncRNA | **238.539** | -0.507 | -0.153 | **0.704** | 6.90E-03 | **1.51E-02** |
| ENSG00000132952.14 | **USPL1** | 10208 | protein_coding | **538.673** | 0.292 | 0.088 | **1.224** | 6.90E-03 | **1.51E-02** |
| ENSG00000205209.8 | **SCGB2B2** | 284402 | protein_coding | **18.545** | 0.879 | 0.265 | **1.839** | 6.91E-03 | **1.51E-02** |
| ENSG00000303442.1 | **ENSG00000303442** | | lncRNA | **8.808** | 1.307 | 0.394 | **2.475** | 6.93E-03 | **1.51E-02** |
| ENSG00000163702.23 | **IL17RC** | 84818 | protein_coding | **30.129** | 0.868 | 0.261 | **1.825** | 6.95E-03 | **1.52E-02** |
| ENSG00000197272.3 | **IL27** | 246778 | protein_coding | **6.500** | 1.873 | 0.564 | **3.664** | 6.96E-03 | **1.52E-02** |
| ENSG00000159588.17 | **CCDC17** | 149483 | protein_coding | **54.165** | -0.616 | -0.185 | **0.653** | 6.96E-03 | **1.52E-02** |
| ENSG00000145779.9 | **TNFAIP8** | 25816 | protein_coding | **1378.040** | -0.342 | -0.103 | **0.789** | 6.97E-03 | **1.52E-02** |
| ENSG00000188501.13 | **LCTL** | 197021 | protein_coding | **4.652** | 2.566 | 0.773 | **5.923** | 6.98E-03 | **1.52E-02** |
| ENSG00000116604.21 | **MEF2D** | 4209 | protein_coding | **1349.306** | 0.296 | 0.089 | **1.227** | 7.00E-03 | **1.53E-02** |
| ENSG00000228037.2 | **ENSG00000228037** | | lncRNA | **149.341** | -0.393 | -0.118 | **0.762** | 7.00E-03 | **1.53E-02** |
| ENSG00000076650.9 | **GPATCH1** | 55094 | protein_coding | **239.861** | -0.328 | -0.099 | **0.797** | 7.00E-03 | **1.53E-02** |

| | | | | | | | | | |
|---|---|---|---|---|---|---|---|---|---|
| ENSG00000269399.3 | **ENSG00000269399** | | lncRNA | **98.587** | 0.772 | 0.232 | **1.708** | 7.01E-03 | **1.53E-02** |
| ENSG00000204392.13 | **LSM2** | 57819 | protein_coding | **919.119** | -0.331 | -0.100 | **0.795** | 7.03E-03 | **1.53E-02** |
| ENSG00000161036.15 | **LRWD1** | 222229 | protein_coding | **441.859** | 0.270 | 0.081 | **1.206** | 7.03E-03 | **1.53E-02** |
| ENSG00000263680.2 | **ENSG00000263680** | | lncRNA | **134.365** | 0.508 | 0.153 | **1.422** | 7.05E-03 | **1.54E-02** |
| ENSG00000124097.8 | **HMGB1P1** | | protein_coding | **24.647** | 1.112 | 0.335 | **2.161** | 7.06E-03 | **1.54E-02** |
| ENSG00000261740.7 | **BOLA2-SM** | 107282092 | protein_coding | **155.729** | -0.522 | -0.157 | **0.697** | 7.06E-03 | **1.54E-02** |
| ENSG00000181016.11 | **LSMEM1** | 286006 | protein_coding | **16.741** | 1.503 | 0.453 | **2.835** | 7.06E-03 | **1.54E-02** |
| ENSG00000147684.12 | **NDUFB9** | 4715 | protein_coding | **1551.944** | -0.275 | -0.083 | **0.826** | 7.07E-03 | **1.54E-02** |
| ENSG00000170417.17 | **TMEM182** | 130827 | protein_coding | **22.593** | 0.933 | 0.281 | **1.909** | 7.09E-03 | **1.55E-02** |
| ENSG00000230322.1 | **CELF2-AS3** | | lncRNA | **7.546** | 1.551 | 0.467 | **2.931** | 7.10E-03 | **1.55E-02** |
| ENSG00000164574.18 | **GALNT10** | 55568 | protein_coding | **1255.476** | 0.325 | 0.098 | **1.253** | 7.10E-03 | **1.55E-02** |
| ENSG00000126522.19 | **ASL** | 435 | protein_coding | **597.088** | 0.441 | 0.133 | **1.358** | 7.12E-03 | **1.55E-02** |
| ENSG00000105141.6 | **CASP14** | 23581 | protein_coding | **10.424** | 1.342 | 0.404 | **2.535** | 7.12E-03 | **1.55E-02** |
| ENSG00000182247.13 | **UBE2E2** | 7325 | protein_coding | **493.365** | 0.319 | 0.096 | **1.247** | 7.13E-03 | **1.55E-02** |
| ENSG00000164919.13 | **COX6C** | 1345 | protein_coding | **2486.361** | -0.302 | -0.091 | **0.811** | 7.13E-03 | **1.55E-02** |
| ENSG00000158615.10 | **PPP1R15B** | 84919 | protein_coding | **630.961** | -0.496 | -0.149 | **0.709** | 7.14E-03 | **1.55E-02** |
| ENSG00000183682.10 | **BMP8A** | 353500 | protein_coding | **20.628** | 1.041 | 0.313 | **2.058** | 7.14E-03 | **1.56E-02** |
| ENSG00000101310.19 | **SEC23B** | 10483 | protein_coding | **599.443** | -0.486 | -0.146 | **0.714** | 7.15E-03 | **1.56E-02** |
| ENSG00000197776.10 | **KLHDC1** | 122773 | protein_coding | **26.318** | 0.941 | 0.283 | **1.920** | 7.16E-03 | **1.56E-02** |
| ENSG00000302734.1 | **ENSG00000302734** | | lncRNA | **28.405** | 1.245 | 0.375 | **2.370** | 7.16E-03 | **1.56E-02** |
| ENSG00000268350.10 | **FAM156A** | 29057 | protein_coding | **1093.804** | -0.438 | -0.132 | **0.738** | 7.18E-03 | **1.56E-02** |
| ENSG00000102580.17 | **DNAJC3** | 5611 | protein_coding | **591.399** | -0.360 | -0.108 | **0.779** | 7.18E-03 | **1.56E-02** |
| ENSG00000187049.13 | **TMEM216** | 51259 | protein_coding | **169.039** | -0.394 | -0.118 | **0.761** | 7.18E-03 | **1.56E-02** |
| ENSG00000100218.13 | **RSPH14** | 27156 | protein_coding | **9.834** | 1.734 | 0.522 | **3.326** | 7.19E-03 | **1.57E-02** |
| ENSG00000257285.8 | **PRMT5-DT** | 10192693 | lncRNA | **14.874** | 1.249 | 0.376 | **2.377** | 7.20E-03 | **1.57E-02** |
| ENSG00000189334.11 | **S100A14** | 57402 | protein_coding | **5.612** | 1.726 | 0.520 | **3.309** | 7.21E-03 | **1.57E-02** |
| ENSG00000235954.10 | **TTC28-AS1** | 284900 | lncRNA | **114.644** | -0.431 | -0.130 | **0.742** | 7.21E-03 | **1.57E-02** |
| ENSG00000149541.12 | **B3GAT3** | 26229 | protein_coding | **718.137** | -0.273 | -0.082 | **0.828** | 7.22E-03 | **1.57E-02** |
| ENSG00000151413.19 | **NUBPL** | 80224 | protein_coding | **104.997** | -0.459 | -0.138 | **0.727** | 7.23E-03 | **1.57E-02** |
| ENSG00000146411.8 | **SLC2A12** | 154091 | protein_coding | **13.675** | 1.336 | 0.402 | **2.524** | 7.23E-03 | **1.57E-02** |
| ENSG00000275464.6 | **ENSG0000(** | 102724159 | artifact | **373.128** | -0.444 | -0.134 | **0.735** | 7.25E-03 | **1.58E-02** |

| | | | | | | | | | |
|---|---|---|---|---|---|---|---|---|---|
| ENSG00000148719.18 | **DNAJB12** | 54788 | protein_coding | **695.516** | -0.265 | -0.080 | **0.832** | 7.26E-03 | **1.58E-02** |
| ENSG00000128039.15 | **SRD5A3** | 79644 | protein_coding | **101.409** | -0.551 | -0.166 | **0.683** | 7.27E-03 | **1.58E-02** |
| ENSG00000159479.19 | **MED8** | 112950 | protein_coding | **724.044** | -0.376 | -0.113 | **0.770** | 7.28E-03 | **1.58E-02** |
| ENSG00000183308.7 | **ENSG00000183308** | | lncRNA | **10.702** | -1.143 | -0.344 | **0.453** | 7.28E-03 | **1.58E-02** |
| ENSG00000278784.2 | **ENSG00000278784** | | lncRNA | **155.294** | 0.445 | 0.134 | **1.361** | 7.30E-03 | **1.59E-02** |
| ENSG00000151552.15 | **QDPR** | 5860 | protein_coding | **317.750** | -0.346 | -0.104 | **0.787** | 7.30E-03 | **1.59E-02** |
| ENSG00000085224.24 | **ATRX** | 546 | protein_coding | **632.310** | -0.619 | -0.186 | **0.651** | 7.30E-03 | **1.59E-02** |
| ENSG00000230082.2 | **PRRT3-AS1** | 100874032 | lncRNA | **11.744** | 1.140 | 0.343 | **2.204** | 7.30E-03 | **1.59E-02** |
| ENSG00000249167.3 | **SEMA6A-A** | 105379132 | lncRNA | **5.543** | 1.814 | 0.546 | **3.517** | 7.30E-03 | **1.59E-02** |
| ENSG00000301217.1 | **ENSG00000301217** | | lncRNA | **6.038** | 1.734 | 0.522 | **3.327** | 7.31E-03 | **1.59E-02** |
| ENSG00000148824.21 | **MTG1** | 92170 | protein_coding | **1056.994** | 0.330 | 0.099 | **1.257** | 7.31E-03 | **1.59E-02** |
| ENSG00000153982.11 | **GDPD1** | 284161 | protein_coding | **12.071** | 1.169 | 0.352 | **2.248** | 7.32E-03 | **1.59E-02** |
| ENSG00000081052.15 | **COL4A4** | 1286 | protein_coding | **228.496** | 0.685 | 0.206 | **1.608** | 7.33E-03 | **1.59E-02** |
| ENSG00000189058.11 | **APOD** | 347 | protein_coding | **7.444** | 1.722 | 0.518 | **3.300** | 7.33E-03 | **1.59E-02** |
| ENSG00000174844.16 | **DNAH12** | 201625 | protein_coding | **9.008** | 1.362 | 0.410 | **2.571** | 7.33E-03 | **1.59E-02** |
| ENSG00000220201.8 | **ZGLP1** | 100125288 | protein_coding | **105.424** | 0.593 | 0.178 | **1.508** | 7.34E-03 | **1.59E-02** |
| ENSG00000159388.7 | **BTG2** | 7832 | protein_coding | **2658.856** | -0.285 | -0.086 | **0.821** | 7.34E-03 | **1.59E-02** |
| ENSG00000255236.4 | **CHKA-DT** | | lncRNA | **12.302** | 1.270 | 0.382 | **2.411** | 7.34E-03 | **1.59E-02** |
| ENSG00000197006.17 | **METTL9** | 51108 | protein_coding | **477.132** | -0.453 | -0.136 | **0.731** | 7.34E-03 | **1.59E-02** |
| ENSG00000185250.18 | **PPIL6** | 285755 | protein_coding | **5.738** | 2.003 | 0.603 | **4.007** | 7.35E-03 | **1.59E-02** |
| ENSG00000116132.13 | **PRRX1** | 5396 | protein_coding | **24.775** | -0.999 | -0.301 | **0.500** | 7.35E-03 | **1.60E-02** |
| ENSG00000167664.11 | **TMIGD2** | 126259 | protein_coding | **7.024** | 1.926 | 0.580 | **3.799** | 7.37E-03 | **1.60E-02** |
| ENSG00000102385.15 | **DRP2** | 1821 | protein_coding | **12.039** | -1.231 | -0.371 | **0.426** | 7.38E-03 | **1.60E-02** |
| ENSG00000172007.9 | **RAB33B** | 83452 | protein_coding | **80.163** | -0.601 | -0.181 | **0.659** | 7.39E-03 | **1.60E-02** |
| ENSG00000140450.11 | **ARRDC4** | 91947 | protein_coding | **11.418** | 1.786 | 0.538 | **3.449** | 7.42E-03 | **1.61E-02** |
| ENSG00000135090.16 | **TAOK3** | 51347 | protein_coding | **825.955** | -0.351 | -0.106 | **0.784** | 7.43E-03 | **1.61E-02** |
| ENSG00000100142.18 | **POLR2F** | 5435 | protein_coding | **556.217** | -0.334 | -0.100 | **0.794** | 7.43E-03 | **1.61E-02** |
| ENSG00000260772.1 | **ENSG00000260772** | | lncRNA | **571.906** | 0.455 | 0.137 | **1.370** | 7.45E-03 | **1.61E-02** |
| ENSG00000247728.3 | **ARHGAP11** | 102725021 | lncRNA | **95.070** | 0.681 | 0.205 | **1.603** | 7.45E-03 | **1.62E-02** |
| ENSG00000038382.24 | **TRIO** | 7204 | protein_coding | **1561.831** | -0.328 | -0.099 | **0.797** | 7.46E-03 | **1.62E-02** |
| ENSG00000115207.17 | **GTF3C2** | 2976 | protein_coding | **1376.197** | -0.311 | -0.094 | **0.806** | 7.47E-03 | **1.62E-02** |

| | | | | | | | | | |
|---|---|---|---|---|---|---|---|---|---|
| ENSG00000182489.10 | **XKRX** | 402415 | protein_coding | **9.114** | 1.417 | 0.427 | **2.670** | 7.48E-03 | **1.62E-02** |
| ENSG00000226091.8 | **LINC00937** | 389634 | lncRNA | **54.452** | 0.800 | 0.241 | **1.741** | 7.48E-03 | **1.62E-02** |
| ENSG00000261832.6 | **ENSG0000(** | 1201 | protein_coding | **16.112** | 1.374 | 0.414 | **2.591** | 7.49E-03 | **1.62E-02** |
| ENSG00000214654.10 | **B3GALT9** | 10028884‹ | protein_coding | **120.357** | 0.430 | 0.129 | **1.347** | 7.49E-03 | **1.62E-02** |
| ENSG00000265185.6 | **SNORD3B-1** | | snoRNA | **32.494** | 0.854 | 0.257 | **1.807** | 7.49E-03 | **1.62E-02** |
| ENSG00000228775.8 | **WEE2-AS1** | 285962 | lncRNA | **32.339** | 0.821 | 0.247 | **1.766** | 7.49E-03 | **1.62E-02** |
| ENSG00000267519.7 | **MIR23AHG** | | lncRNA | **237.193** | -0.418 | -0.126 | **0.748** | 7.50E-03 | **1.62E-02** |
| ENSG00000271147.8 | **ENSG00000271147** | | lncRNA | **25.870** | -1.001 | -0.301 | **0.500** | 7.51E-03 | **1.63E-02** |
| ENSG00000166073.13 | **GPR176** | 11245 | protein_coding | **10.873** | 1.480 | 0.446 | **2.790** | 7.51E-03 | **1.63E-02** |
| ENSG00000052126.19 | **PLEKHA5** | 54477 | protein_coding | **11.057** | 1.756 | 0.529 | **3.379** | 7.52E-03 | **1.63E-02** |
| ENSG00000229419.1 | **RALGAPA1P1** | | processed_pseud( | **10.357** | 1.341 | 0.404 | **2.533** | 7.52E-03 | **1.63E-02** |
| ENSG00000238132.2 | **GOLM2P1** | | processed_pseud( | **14.714** | 1.085 | 0.327 | **2.122** | 7.53E-03 | **1.63E-02** |
| ENSG00000028203.21 | **VEZT** | 55591 | protein_coding | **810.736** | -0.371 | -0.112 | **0.773** | 7.53E-03 | **1.63E-02** |
| ENSG00000173598.16 | **NUDT4** | 11163 | protein_coding | **1442.553** | -0.277 | -0.083 | **0.826** | 7.54E-03 | **1.63E-02** |
| ENSG00000184924.8 | **PTRHD1** | 391356 | protein_coding | **616.368** | -0.392 | -0.118 | **0.762** | 7.54E-03 | **1.63E-02** |
| ENSG00000100030.17 | **MAPK1** | 5594 | protein_coding | **1146.133** | 0.283 | 0.085 | **1.216** | 7.56E-03 | **1.63E-02** |
| ENSG00000294159.1 | **ENSG00000294159** | | lncRNA | **19.597** | -0.995 | -0.300 | **0.502** | 7.57E-03 | **1.64E-02** |
| ENSG00000293068.2 | **ENSG00000293068** | | lncRNA | **24.333** | 1.302 | 0.392 | **2.465** | 7.58E-03 | **1.64E-02** |
| ENSG00000154479.14 | **CFAP210** | 129881 | protein_coding | **6.221** | 2.217 | 0.667 | **4.648** | 7.60E-03 | **1.64E-02** |
| ENSG00000138794.11 | **CASP6** | 839 | protein_coding | **148.498** | -0.452 | -0.136 | **0.731** | 7.61E-03 | **1.64E-02** |
| ENSG00000260329.3 | **TMEM263-** | 11986387( | lncRNA | **18.231** | 0.969 | 0.292 | **1.957** | 7.62E-03 | **1.65E-02** |
| ENSG00000196526.13 | **AFAP1** | 60312 | protein_coding | **17.942** | 1.387 | 0.417 | **2.615** | 7.62E-03 | **1.65E-02** |
| ENSG00000185917.16 | **SETD4** | 54093 | protein_coding | **339.932** | -0.304 | -0.092 | **0.810** | 7.63E-03 | **1.65E-02** |
| ENSG00000244558.7 | **KCNK15-AS** | 79015 | lncRNA | **12.641** | 1.055 | 0.318 | **2.077** | 7.63E-03 | **1.65E-02** |
| ENSG00000008405.14 | **CRY1** | 1407 | protein_coding | **99.629** | 0.759 | 0.228 | **1.692** | 7.64E-03 | **1.65E-02** |
| ENSG00000248673.3 | **LINC01331** | 104310351 | lncRNA | **4.050** | 2.984 | 0.898 | **7.913** | 7.64E-03 | **1.65E-02** |
| ENSG00000148343.21 | **MIGA2** | 84895 | protein_coding | **641.270** | -0.274 | -0.083 | **0.827** | 7.65E-03 | **1.65E-02** |
| ENSG00000278249.1 | **SCARNA2** | 677766 | scaRNA | **12.776** | 1.105 | 0.333 | **2.151** | 7.65E-03 | **1.65E-02** |
| ENSG00000085365.20 | **SCAMP1** | 9522 | protein_coding | **103.032** | -0.447 | -0.135 | **0.734** | 7.66E-03 | **1.65E-02** |
| ENSG00000305244.1 | **ENSG00000305244** | | lncRNA | **5.680** | 1.773 | 0.534 | **3.419** | 7.67E-03 | **1.66E-02** |
| ENSG00000261553.5 | **ENSG00000261553** | | lncRNA | **8.640** | 2.219 | 0.668 | **4.655** | 7.67E-03 | **1.66E-02** |

| | | | | | | | | | |
|---|---|---|---|---|---|---|---|---|---|
| ENSG00000180481.13 | **GLIPR1L2** | 144321 | protein_coding | **5.807** | 1.774 | 0.534 | **3.420** | 7.67E-03 | **1.66E-02** |
| ENSG00000104325.9 | **DECR1** | 1666 | protein_coding | **1240.931** | -0.321 | -0.097 | **0.800** | 7.68E-03 | **1.66E-02** |
| ENSG00000100376.14 | **FAM118A** | 55007 | protein_coding | **1245.789** | 0.291 | 0.087 | **1.223** | 7.68E-03 | **1.66E-02** |
| ENSG00000087258.18 | **GNAO1** | 2775 | protein_coding | **7.340** | 1.642 | 0.494 | **3.122** | 7.69E-03 | **1.66E-02** |
| ENSG00000187922.16 | **LCN10** | 414332 | protein_coding | **68.807** | 0.715 | 0.215 | **1.642** | 7.69E-03 | **1.66E-02** |
| ENSG00000178562.20 | **CD28** | 940 | protein_coding | **138.802** | 0.531 | 0.160 | **1.445** | 7.69E-03 | **1.66E-02** |
| ENSG00000198040.13 | **ZNF84** | 7637 | protein_coding | **686.917** | -0.356 | -0.107 | **0.781** | 7.70E-03 | **1.66E-02** |
| ENSG00000285976.5 | **ENSG0000(** | 12812582: | protein_coding | **21.926** | -1.440 | -0.433 | **0.369** | 7.70E-03 | **1.66E-02** |
| ENSG00000284959.4 | **TSHR-AS1** | | lncRNA | **10.024** | 1.508 | 0.454 | **2.844** | 7.71E-03 | **1.66E-02** |
| ENSG00000113269.16 | **RNF130** | 55819 | protein_coding | **32.347** | 0.900 | 0.271 | **1.866** | 7.71E-03 | **1.66E-02** |
| ENSG00000204397.13 | **CARD16** | 114769 | protein_coding | **191.518** | -0.355 | -0.107 | **0.782** | 7.73E-03 | **1.67E-02** |
| ENSG00000291219.2 | **ENSG00000291219** | | lncRNA | **26.523** | -0.815 | -0.245 | **0.569** | 7.73E-03 | **1.67E-02** |
| ENSG00000173653.9 | **RCE1** | 9986 | protein_coding | **315.448** | -0.275 | -0.083 | **0.826** | 7.74E-03 | **1.67E-02** |
| ENSG00000103415.14 | **HMOX2** | 3163 | protein_coding | **590.610** | -0.279 | -0.084 | **0.824** | 7.75E-03 | **1.67E-02** |
| ENSG00000080546.16 | **SESN1** | 27244 | protein_coding | **112.196** | -0.555 | -0.167 | **0.681** | 7.75E-03 | **1.67E-02** |
| ENSG00000103313.16 | **MEFV** | 4210 | protein_coding | **6.912** | 1.645 | 0.495 | **3.128** | 7.76E-03 | **1.67E-02** |
| ENSG00000262691.1 | **ENSG00000262691** | | lncRNA | **18.298** | 1.006 | 0.303 | **2.008** | 7.80E-03 | **1.68E-02** |
| ENSG00000138592.17 | **USP8** | 9101 | protein_coding | **473.291** | 0.345 | 0.104 | **1.270** | 7.81E-03 | **1.68E-02** |
| ENSG00000158813.19 | **EDA** | 1896 | protein_coding | **6.079** | 2.647 | 0.797 | **6.262** | 7.81E-03 | **1.68E-02** |
| ENSG00000138032.23 | **PPM1B** | 5495 | protein_coding | **483.696** | 0.391 | 0.118 | **1.311** | 7.81E-03 | **1.68E-02** |
| ENSG00000164975.18 | **SNAPC3** | 6619 | protein_coding | **490.465** | -0.302 | -0.091 | **0.811** | 7.82E-03 | **1.68E-02** |
| ENSG00000273473.1 | **ENSG00000273473** | | lncRNA | **5.018** | 1.955 | 0.588 | **3.876** | 7.83E-03 | **1.69E-02** |
| ENSG00000137494.16 | **ANKRD42** | 338699 | protein_coding | **98.202** | 0.610 | 0.184 | **1.526** | 7.84E-03 | **1.69E-02** |
| ENSG00000114023.17 | **FAM162A** | 26355 | protein_coding | **2452.493** | -0.415 | -0.125 | **0.750** | 7.85E-03 | **1.69E-02** |
| ENSG00000178691.14 | **SUZ12** | 23512 | protein_coding | **402.173** | -0.363 | -0.109 | **0.777** | 7.87E-03 | **1.70E-02** |
| ENSG00000023171.22 | **GRAMD1B** | 57476 | protein_coding | **228.425** | 0.504 | 0.152 | **1.418** | 7.88E-03 | **1.70E-02** |
| ENSG00000188483.8 | **IER5L** | 389792 | protein_coding | **34.778** | 0.872 | 0.263 | **1.831** | 7.88E-03 | **1.70E-02** |
| ENSG00000125845.9 | **BMP2** | 650 | protein_coding | **8.850** | 1.373 | 0.413 | **2.591** | 7.92E-03 | **1.71E-02** |
| ENSG00000117298.17 | **ECE1** | 1889 | protein_coding | **268.036** | 0.450 | 0.135 | **1.366** | 7.93E-03 | **1.71E-02** |
| ENSG00000251323.4 | **LINC02728** | 10192886! | lncRNA | **9.752** | 1.390 | 0.419 | **2.621** | 7.94E-03 | **1.71E-02** |
| ENSG00000233264.2 | **ENSG00000233264** | | processed_pseud( | **25.239** | 0.756 | 0.228 | **1.689** | 7.94E-03 | **1.71E-02** |

| | | | | | | | | | |
|---|---|---|---|---|---|---|---|---|---|
| ENSG00000104894.14 | **CD37** | 951 | protein_coding | **7502.266** | -0.417 | -0.125 | **0.749** | 7.94E-03 | **1.71E-02** |
| ENSG00000173660.14 | **UQCRH** | 7388 | protein_coding | **4859.250** | 0.322 | 0.097 | **1.250** | 7.95E-03 | **1.71E-02** |
| ENSG00000107625.16 | **DDX50** | 79009 | protein_coding | **757.041** | -0.312 | -0.094 | **0.805** | 7.96E-03 | **1.71E-02** |
| ENSG00000196208.16 | **GREB1** | 9687 | protein_coding | **11.005** | 1.333 | 0.401 | **2.520** | 7.97E-03 | **1.71E-02** |
| ENSG00000196189.16 | **SEMA4A** | 64218 | protein_coding | **3484.164** | 0.379 | 0.114 | **1.301** | 7.98E-03 | **1.71E-02** |
| ENSG00000158163.17 | **DZIP1L** | 199221 | protein_coding | **6.069** | 2.324 | 0.700 | **5.007** | 7.99E-03 | **1.72E-02** |
| ENSG00000196670.15 | **ZFP62** | 643836 | protein_coding | **426.026** | -0.360 | -0.108 | **0.779** | 8.00E-03 | **1.72E-02** |
| ENSG00000163131.14 | **CTSS** | 1520 | protein_coding | **1623.834** | -0.345 | -0.104 | **0.787** | 8.02E-03 | **1.72E-02** |
| ENSG00000163866.11 | **SMIM12** | 113444 | protein_coding | **400.134** | -0.321 | -0.097 | **0.801** | 8.03E-03 | **1.73E-02** |
| ENSG00000141682.14 | **PMAIP1** | 5366 | protein_coding | **1432.716** | 0.559 | 0.168 | **1.473** | 8.04E-03 | **1.73E-02** |
| ENSG00000175003.17 | **SLC22A1** | 6580 | protein_coding | **40.883** | 1.201 | 0.362 | **2.299** | 8.04E-03 | **1.73E-02** |
| ENSG00000106772.21 | **PRUNE2** | 158471 | protein_coding | **8.253** | 1.914 | 0.576 | **3.769** | 8.05E-03 | **1.73E-02** |
| ENSG00000100216.8 | **TOMM22** | 56993 | protein_coding | **1255.403** | -0.333 | -0.100 | **0.794** | 8.05E-03 | **1.73E-02** |
| ENSG00000182973.22 | **CNOT10** | 25904 | protein_coding | **608.380** | -0.306 | -0.092 | **0.809** | 8.06E-03 | **1.73E-02** |
| ENSG00000236423.9 | **LINC01134** | 10013361[illegible] | lncRNA | **10.127** | -1.230 | -0.370 | **0.426** | 8.07E-03 | **1.73E-02** |
| ENSG00000214110.3 | **LDHAP4** | | transcribed_proce | **138.204** | -0.725 | -0.218 | **0.605** | 8.08E-03 | **1.73E-02** |
| ENSG00000133422.16 | **MORC2** | 22880 | protein_coding | **1019.046** | 0.370 | 0.112 | **1.293** | 8.08E-03 | **1.73E-02** |
| ENSG00000031698.15 | **SARS1** | 6301 | protein_coding | **2456.616** | -0.469 | -0.141 | **0.723** | 8.11E-03 | **1.74E-02** |
| ENSG00000303299.1 | **ENSG00000303299** | | lncRNA | **3.226** | 3.107 | 0.935 | **8.615** | 8.12E-03 | **1.74E-02** |
| ENSG00000168495.15 | **POLR3D** | 661 | protein_coding | **407.851** | -0.545 | -0.164 | **0.685** | 8.12E-03 | **1.74E-02** |
| ENSG00000010278.17 | **CD9** | 928 | protein_coding | **145.915** | 0.516 | 0.155 | **1.430** | 8.13E-03 | **1.74E-02** |
| ENSG00000258651.2 | **SEC23A-AS** | 10537045[illegible] | lncRNA | **8.565** | 1.497 | 0.451 | **2.823** | 8.14E-03 | **1.75E-02** |
| ENSG00000069998.14 | **HDHD5** | 27440 | protein_coding | **681.012** | -0.276 | -0.083 | **0.826** | 8.15E-03 | **1.75E-02** |
| ENSG00000154309.11 | **DISP1** | 84976 | protein_coding | **101.694** | 0.504 | 0.152 | **1.418** | 8.15E-03 | **1.75E-02** |
| ENSG00000288739.2 | **ENSG00000288739** | | lncRNA | **13.475** | 1.059 | 0.319 | **2.083** | 8.17E-03 | **1.75E-02** |
| ENSG00000275479.2 | **ENSG00000275479** | | lncRNA | **5.990** | 1.909 | 0.575 | **3.757** | 8.19E-03 | **1.76E-02** |
| ENSG00000163041.15 | **H3-3A** | 3020 | protein_coding | **3191.714** | -0.377 | -0.113 | **0.770** | 8.19E-03 | **1.76E-02** |
| ENSG00000154743.21 | **TSEN2** | 80746 | protein_coding | **186.903** | -0.420 | -0.127 | **0.747** | 8.19E-03 | **1.76E-02** |
| ENSG00000159556.11 | **ISL2** | 64843 | protein_coding | **208.002** | 0.382 | 0.115 | **1.303** | 8.20E-03 | **1.76E-02** |
| ENSG00000151150.24 | **ANK3** | 288 | protein_coding | **14.658** | 1.797 | 0.541 | **3.474** | 8.23E-03 | **1.76E-02** |
| ENSG00000272455.1 | **MRPL20-DT** | | lncRNA | **93.435** | 0.477 | 0.144 | **1.392** | 8.23E-03 | **1.76E-02** |

| | | | | | | | | | |
|---|---|---|---|---|---|---|---|---|---|
| ENSG00000239382.13 | **ALKBH6** | 84964 | protein_coding | **808.330** | -0.383 | -0.115 | **0.767** | 8.23E-03 | **1.76E-02** |
| ENSG00000176268.5 | **CYCSP34** | | processed_pseud | **6.197** | 1.522 | 0.458 | **2.871** | 8.24E-03 | **1.76E-02** |
| ENSG00000204427.14 | **ABHD16A** | 7920 | protein_coding | **397.272** | -0.300 | -0.090 | **0.812** | 8.24E-03 | **1.77E-02** |
| ENSG00000278126.1 | **ENSG00000278126** | | lncRNA | **10.863** | -1.068 | -0.321 | **0.477** | 8.25E-03 | **1.77E-02** |
| ENSG00000164091.14 | **WDR82** | 80335 | protein_coding | **1024.611** | -0.304 | -0.091 | **0.810** | 8.26E-03 | **1.77E-02** |
| ENSG00000121807.8 | **CCR2** | 729230 | protein_coding | **5.396** | 2.057 | 0.619 | **4.161** | 8.27E-03 | **1.77E-02** |
| ENSG00000290983.2 | **ENSG00000290983** | | lncRNA | **12.373** | 1.608 | 0.484 | **3.047** | 8.28E-03 | **1.77E-02** |
| ENSG00000113532.15 | **ST8SIA4** | 7903 | protein_coding | **388.811** | -0.403 | -0.121 | **0.756** | 8.29E-03 | **1.77E-02** |
| ENSG00000137343.20 | **ATAT1** | 79969 | protein_coding | **308.430** | 0.448 | 0.135 | **1.364** | 8.29E-03 | **1.78E-02** |
| ENSG00000146085.10 | **MMUT** | 4594 | protein_coding | **279.547** | -0.381 | -0.115 | **0.768** | 8.30E-03 | **1.78E-02** |
| ENSG00000143353.14 | **LYPLAL1** | 127018 | protein_coding | **458.616** | -0.428 | -0.129 | **0.743** | 8.30E-03 | **1.78E-02** |
| ENSG00000293471.2 | **ENSG00000293471** | | lncRNA | **5.516** | 1.710 | 0.515 | **3.272** | 8.30E-03 | **1.78E-02** |
| ENSG00000006756.18 | **ARSD** | 414 | protein_coding | **90.701** | -0.610 | -0.184 | **0.655** | 8.31E-03 | **1.78E-02** |
| ENSG00000170191.6 | **NANP** | 140838 | protein_coding | **58.829** | -0.499 | -0.150 | **0.707** | 8.31E-03 | **1.78E-02** |
| ENSG00000147231.15 | **RADX** | 55086 | protein_coding | **142.224** | 0.726 | 0.218 | **1.654** | 8.31E-03 | **1.78E-02** |
| ENSG00000263080.2 | **ENSG0000** | 10537108 | lncRNA | **8.031** | 1.663 | 0.501 | **3.167** | 8.32E-03 | **1.78E-02** |
| ENSG00000118640.13 | **VAMP8** | 8673 | protein_coding | **1454.196** | -0.319 | -0.096 | **0.802** | 8.33E-03 | **1.78E-02** |
| ENSG00000117036.13 | **ETV3** | 2117 | protein_coding | **186.555** | 0.450 | 0.135 | **1.366** | 8.33E-03 | **1.78E-02** |
| ENSG00000290717.2 | **ZNF658B** | 401509 | lncRNA | **17.709** | 1.316 | 0.396 | **2.490** | 8.34E-03 | **1.78E-02** |
| ENSG00000227741.2 | **PEA15-AS1** | 729867 | lncRNA | **7.374** | -1.832 | -0.552 | **0.281** | 8.34E-03 | **1.78E-02** |
| ENSG00000175591.14 | **P2RY2** | 5029 | protein_coding | **5.581** | 1.981 | 0.596 | **3.947** | 8.37E-03 | **1.79E-02** |
| ENSG00000179935.11 | **LINC00652** | | lncRNA | **4.709** | 2.222 | 0.669 | **4.666** | 8.38E-03 | **1.79E-02** |
| ENSG00000069943.12 | **PIGB** | 9488 | protein_coding | **207.689** | -0.410 | -0.123 | **0.753** | 8.38E-03 | **1.79E-02** |
| ENSG00000236021.5 | **COP1-DT** | | lncRNA | **14.726** | 0.918 | 0.276 | **1.889** | 8.40E-03 | **1.79E-02** |
| ENSG00000187079.21 | **TEAD1** | 7003 | protein_coding | **44.036** | 0.946 | 0.285 | **1.927** | 8.40E-03 | **1.80E-02** |
| ENSG00000162645.15 | **GBP2** | 2634 | protein_coding | **707.020** | -0.279 | -0.084 | **0.824** | 8.41E-03 | **1.80E-02** |
| ENSG00000174405.17 | **LIG4** | 3981 | protein_coding | **219.421** | 0.538 | 0.162 | **1.452** | 8.41E-03 | **1.80E-02** |
| ENSG00000215193.16 | **PEX26** | 55670 | protein_coding | **876.949** | -0.417 | -0.126 | **0.749** | 8.41E-03 | **1.80E-02** |
| ENSG00000104953.22 | **TLE6** | 79816 | protein_coding | **159.743** | 0.518 | 0.156 | **1.432** | 8.42E-03 | **1.80E-02** |
| ENSG00000064102.17 | **INTS13** | 55726 | protein_coding | **946.598** | -0.336 | -0.101 | **0.792** | 8.44E-03 | **1.80E-02** |
| ENSG00000107960.14 | **STN1** | 79991 | protein_coding | **447.518** | -0.418 | -0.126 | **0.749** | 8.44E-03 | **1.80E-02** |

| | | | | | | | | | |
|---|---|---|---|---|---|---|---|---|---|
| ENSG00000013441.19 | **CLK1** | 1195 | protein_coding | **1523.985** | -0.350 | -0.105 | **0.784** | 8.44E-03 | **1.80E-02** |
| ENSG00000289701.1 | **ENSG00000289701** | | protein_coding | **43.140** | -0.820 | -0.247 | **0.566** | 8.45E-03 | **1.80E-02** |
| ENSG00000289943.2 | **ENSG00000289943** | | lncRNA | **8.578** | 1.319 | 0.397 | **2.494** | 8.45E-03 | **1.80E-02** |
| ENSG00000281332.1 | **LINC00997** | | lncRNA | **20.392** | 1.059 | 0.319 | **2.084** | 8.45E-03 | **1.80E-02** |
| ENSG00000153064.13 | **BANK1** | 55024 | protein_coding | **43.366** | -0.601 | -0.181 | **0.659** | 8.46E-03 | **1.80E-02** |
| ENSG00000144369.16 | **FAM171B** | 165215 | protein_coding | **12.590** | 1.129 | 0.340 | **2.187** | 8.47E-03 | **1.81E-02** |
| ENSG00000230882.1 | **ENSG00000230882** | | processed_pseud | **16.578** | -0.978 | -0.294 | **0.508** | 8.49E-03 | **1.81E-02** |
| ENSG00000162458.15 | **FBLIM1** | 54751 | protein_coding | **5.219** | 1.879 | 0.566 | **3.677** | 8.50E-03 | **1.81E-02** |
| ENSG00000232119.10 | **MCTS1** | 28985 | protein_coding | **759.694** | -0.536 | -0.161 | **0.690** | 8.51E-03 | **1.82E-02** |
| ENSG00000135940.9 | **COX5B** | 1329 | protein_coding | **2804.883** | -0.287 | -0.086 | **0.819** | 8.52E-03 | **1.82E-02** |
| ENSG00000213465.10 | **ARL2** | 402 | protein_coding | **2016.061** | 0.386 | 0.116 | **1.307** | 8.52E-03 | **1.82E-02** |
| ENSG00000138780.17 | **GSTCD** | 79807 | protein_coding | **121.096** | -0.477 | -0.144 | **0.719** | 8.52E-03 | **1.82E-02** |
| ENSG00000099910.19 | **KLHL22** | 84861 | protein_coding | **363.289** | 0.265 | 0.080 | **1.202** | 8.54E-03 | **1.82E-02** |
| ENSG00000079974.21 | **RABL2B** | 11158 | protein_coding | **414.224** | 0.295 | 0.089 | **1.227** | 8.55E-03 | **1.82E-02** |
| ENSG00000058056.12 | **USP13** | 8975 | protein_coding | **187.384** | -0.697 | -0.210 | **0.617** | 8.56E-03 | **1.83E-02** |
| ENSG00000145708.13 | **CRHBP** | 1393 | protein_coding | **8.378** | 1.822 | 0.549 | **3.537** | 8.60E-03 | **1.83E-02** |
| ENSG00000082074.21 | **FYB1** | 2533 | protein_coding | **70.535** | 0.857 | 0.258 | **1.812** | 8.60E-03 | **1.83E-02** |
| ENSG00000122729.21 | **ACO1** | 48 | protein_coding | **235.315** | 0.363 | 0.109 | **1.286** | 8.61E-03 | **1.83E-02** |
| ENSG00000196586.19 | **MYO6** | 4646 | protein_coding | **676.142** | 0.408 | 0.123 | **1.327** | 8.61E-03 | **1.83E-02** |
| ENSG00000156650.16 | **KAT6B** | 23522 | protein_coding | **431.835** | -0.418 | -0.126 | **0.749** | 8.63E-03 | **1.84E-02** |
| ENSG00000162804.17 | **SNED1** | 25992 | protein_coding | **46.841** | 0.906 | 0.273 | **1.874** | 8.65E-03 | **1.84E-02** |
| ENSG00000289149.2 | **ENSG00000289149** | | lncRNA | **6.275** | -1.510 | -0.455 | **0.351** | 8.65E-03 | **1.84E-02** |
| ENSG00000059728.13 | **MXD1** | 4084 | protein_coding | **238.694** | -0.626 | -0.188 | **0.648** | 8.65E-03 | **1.84E-02** |
| ENSG00000297424.1 | **ENSG00000297424** | | lncRNA | **17.232** | 0.966 | 0.291 | **1.953** | 8.66E-03 | **1.84E-02** |
| ENSG00000289011.2 | **ENSG0000** | 10537174 | lncRNA | **13.418** | 1.263 | 0.380 | **2.401** | 8.67E-03 | **1.84E-02** |
| ENSG00000226806.2 | **LCT-AS1** | 10050760 | lncRNA | **12.138** | -1.439 | -0.433 | **0.369** | 8.67E-03 | **1.84E-02** |
| ENSG00000137434.14 | **C6orf52** | 347744 | protein_coding | **28.868** | 0.817 | 0.246 | **1.762** | 8.67E-03 | **1.85E-02** |
| ENSG00000167685.17 | **ZNF444** | 55311 | protein_coding | **972.175** | 0.303 | 0.091 | **1.234** | 8.68E-03 | **1.85E-02** |
| ENSG00000272502.1 | **SNTB1-AS1** | | lncRNA | **14.709** | -1.102 | -0.332 | **0.466** | 8.68E-03 | **1.85E-02** |
| ENSG00000169499.17 | **PLEKHA2** | 59339 | protein_coding | **790.338** | -0.324 | -0.097 | **0.799** | 8.68E-03 | **1.85E-02** |
| ENSG00000290397.2 | **ENSG00000290397** | | lncRNA | **6.875** | -1.944 | -0.585 | **0.260** | 8.68E-03 | **1.85E-02** |

| | | | | | | | | | |
|---|---|---|---|---|---|---|---|---|---|
| ENSG00000148516.25 | **ZEB1** | 6935 | protein_coding | **409.860** | -0.466 | -0.140 | **0.724** | 8.69E-03 | **1.85E-02** |
| ENSG00000145868.19 | **FBXO38** | 81545 | protein_coding | **551.368** | 0.426 | 0.128 | **1.344** | 8.69E-03 | **1.85E-02** |
| ENSG00000287710.1 | **ENSG00000287710** | | lncRNA | **27.216** | 0.840 | 0.253 | **1.791** | 8.71E-03 | **1.85E-02** |
| ENSG00000171824.16 | **EXOSC10** | 5394 | protein_coding | **1339.421** | -0.353 | -0.106 | **0.783** | 8.72E-03 | **1.85E-02** |
| ENSG00000234851.4 | **RPL23AP42** | | processed_pseud | **717.292** | -3.335 | -1.004 | **0.099** | 8.72E-03 | **1.85E-02** |
| ENSG00000167377.21 | **ZNF23** | 7571 | protein_coding | **442.584** | 0.380 | 0.114 | **1.301** | 8.72E-03 | **1.85E-02** |
| ENSG00000120658.16 | **ENOX1** | 55068 | protein_coding | **9.218** | 1.552 | 0.467 | **2.932** | 8.72E-03 | **1.85E-02** |
| ENSG00000162825.19 | **NBPF20** | 10028814 | protein_coding | **113.902** | 0.556 | 0.167 | **1.470** | 8.77E-03 | **1.86E-02** |
| ENSG00000169299.16 | **PGM2** | 55276 | protein_coding | **174.108** | -0.448 | -0.135 | **0.733** | 8.77E-03 | **1.86E-02** |
| ENSG00000176788.11 | **BASP1** | 10409 | protein_coding | **1570.856** | 0.373 | 0.112 | **1.295** | 8.77E-03 | **1.86E-02** |
| ENSG00000146215.16 | **CRIP3** | 401262 | protein_coding | **311.584** | 0.372 | 0.112 | **1.294** | 8.78E-03 | **1.87E-02** |
| ENSG00000231050.4 | **GNB1-DT** | 10537894 | lncRNA | **17.685** | 1.094 | 0.329 | **2.135** | 8.79E-03 | **1.87E-02** |
| ENSG00000092009.11 | **CMA1** | 1215 | protein_coding | **32.497** | 0.859 | 0.259 | **1.813** | 8.79E-03 | **1.87E-02** |
| ENSG00000101680.18 | **LAMA1** | 284217 | protein_coding | **6.222** | 2.295 | 0.691 | **4.907** | 8.79E-03 | **1.87E-02** |
| ENSG00000205593.14 | **DENND6B** | 414918 | protein_coding | **1019.974** | 0.392 | 0.118 | **1.312** | 8.80E-03 | **1.87E-02** |
| ENSG00000131115.18 | **ZNF227** | 7770 | protein_coding | **220.930** | -0.355 | -0.107 | **0.782** | 8.80E-03 | **1.87E-02** |
| ENSG00000130764.12 | **LRRC47** | 57470 | protein_coding | **671.011** | 0.332 | 0.100 | **1.259** | 8.81E-03 | **1.87E-02** |
| ENSG00000173085.16 | **COQ2** | 27235 | protein_coding | **259.527** | -0.353 | -0.106 | **0.783** | 8.81E-03 | **1.87E-02** |
| ENSG00000180011.8 | **PTGR3** | 284273 | protein_coding | **621.297** | -0.354 | -0.106 | **0.783** | 8.83E-03 | **1.87E-02** |
| ENSG00000147459.19 | **DOCK5** | 80005 | protein_coding | **119.587** | 0.777 | 0.234 | **1.714** | 8.84E-03 | **1.88E-02** |
| ENSG00000135535.19 | **CD164** | 8763 | protein_coding | **1636.403** | -0.430 | -0.129 | **0.742** | 8.84E-03 | **1.88E-02** |
| ENSG00000166401.17 | **SERPINB8** | 5271 | protein_coding | **201.684** | -0.623 | -0.188 | **0.649** | 8.85E-03 | **1.88E-02** |
| ENSG00000290951.2 | **ENSG00000290951** | | lncRNA | **304.921** | 0.320 | 0.096 | **1.249** | 8.85E-03 | **1.88E-02** |
| ENSG00000105229.9 | **PIAS4** | 51588 | protein_coding | **378.933** | -0.331 | -0.100 | **0.795** | 8.86E-03 | **1.88E-02** |
| ENSG00000158482.10 | **SNX29P1** | | transcribed_unpr | **17.654** | -1.138 | -0.343 | **0.454** | 8.88E-03 | **1.88E-02** |
| ENSG00000196757.9 | **ZNF700** | 90592 | protein_coding | **717.292** | -0.278 | -0.084 | **0.825** | 8.88E-03 | **1.88E-02** |
| ENSG00000237836.6 | **PHKA2-AS1** | | lncRNA | **35.628** | 0.918 | 0.276 | **1.889** | 8.89E-03 | **1.88E-02** |
| ENSG00000177082.15 | **WDR73** | 84942 | protein_coding | **430.365** | 0.339 | 0.102 | **1.265** | 8.89E-03 | **1.88E-02** |
| ENSG00000234494.9 | **SP2-AS1** | | lncRNA | **53.195** | 0.621 | 0.187 | **1.538** | 8.89E-03 | **1.88E-02** |
| ENSG00000125834.15 | **STK35** | 140901 | protein_coding | **457.523** | -0.311 | -0.094 | **0.806** | 8.89E-03 | **1.89E-02** |
| ENSG00000268895.6 | **A1BG-AS1** | 503538 | lncRNA | **223.083** | -0.373 | -0.112 | **0.772** | 8.93E-03 | **1.89E-02** |

| | | | | | | | | | |
|---|---|---|---|---|---|---|---|---|---|
| ENSG00000148204.14 | **CRB2** | 286204 | protein_coding | **14.152** | -1.269 | -0.382 | **0.415** | 8.93E-03 | **1.89E-02** |
| ENSG00000135040.18 | **NAA35** | 60560 | protein_coding | **545.255** | -0.329 | -0.099 | **0.796** | 8.94E-03 | **1.89E-02** |
| ENSG00000272398.7 | **CD24** | 100133941 | protein_coding | **147.906** | 0.525 | 0.158 | **1.439** | 8.95E-03 | **1.90E-02** |
| ENSG00000171223.6 | **JUNB** | 3726 | protein_coding | **2784.754** | 0.397 | 0.120 | **1.317** | 8.96E-03 | **1.90E-02** |
| ENSG00000289324.3 | **ENSG00000289324** | | lncRNA | **7.500** | 1.581 | 0.476 | **2.992** | 8.98E-03 | **1.90E-02** |
| ENSG00000024422.14 | **EHD2** | 30846 | protein_coding | **4.772** | 2.872 | 0.864 | **7.320** | 9.01E-03 | **1.91E-02** |
| ENSG00000204673.13 | **AKT1S1** | 84335 | protein_coding | **1184.367** | -0.347 | -0.104 | **0.786** | 9.04E-03 | **1.92E-02** |
| ENSG00000308970.1 | **ENSG00000308970** | | lncRNA | **7.689** | 2.140 | 0.644 | **4.408** | 9.05E-03 | **1.92E-02** |
| ENSG00000183423.14 | **LRIT3** | 345193 | protein_coding | **11.404** | 1.081 | 0.325 | **2.115** | 9.06E-03 | **1.92E-02** |
| ENSG00000162769.15 | **FLVCR1** | 28982 | protein_coding | **357.626** | -0.669 | -0.201 | **0.629** | 9.07E-03 | **1.92E-02** |
| ENSG00000105784.18 | **RUNDC3B** | 154661 | protein_coding | **5.270** | 2.012 | 0.606 | **4.034** | 9.07E-03 | **1.92E-02** |
| ENSG00000168542.20 | **COL3A1** | 1281 | protein_coding | **9.228** | 1.478 | 0.445 | **2.785** | 9.08E-03 | **1.92E-02** |
| ENSG00000080007.10 | **DDX43** | 55510 | protein_coding | **41.591** | 0.761 | 0.229 | **1.695** | 9.08E-03 | **1.92E-02** |
| ENSG00000296810.1 | **ENSG00000296810** | | lncRNA | **20.428** | 0.767 | 0.231 | **1.702** | 9.09E-03 | **1.92E-02** |
| ENSG00000290548.2 | **ENSG00000290548** | | lncRNA | **61.760** | -0.562 | -0.169 | **0.677** | 9.09E-03 | **1.92E-02** |
| ENSG00000270757.1 | **HSPE1-MO** | 100529241 | protein_coding | **61.447** | -1.088 | -0.327 | **0.470** | 9.10E-03 | **1.93E-02** |
| ENSG00000291144.2 | **ENSG00000291144** | | lncRNA | **8.960** | 1.553 | 0.467 | **2.934** | 9.10E-03 | **1.93E-02** |
| ENSG00000258082.2 | **LINC03108** | | lncRNA | **6.745** | 1.554 | 0.468 | **2.936** | 9.11E-03 | **1.93E-02** |
| ENSG00000015153.17 | **YAF2** | 10138 | protein_coding | **193.517** | -0.506 | -0.152 | **0.704** | 9.12E-03 | **1.93E-02** |
| ENSG00000115839.22 | **RAB3GAP1** | 22930 | protein_coding | **1037.369** | -0.286 | -0.086 | **0.820** | 9.13E-03 | **1.93E-02** |
| ENSG00000302960.1 | **ENSG00000302960** | | lncRNA | **11.516** | 1.172 | 0.353 | **2.253** | 9.14E-03 | **1.93E-02** |
| ENSG00000213057.5 | **C1orf220** | 400798 | lncRNA | **17.342** | -0.920 | -0.277 | **0.528** | 9.15E-03 | **1.93E-02** |
| ENSG00000227036.11 | **LINC00511** | 400619 | lncRNA | **77.470** | 0.529 | 0.159 | **1.443** | 9.15E-03 | **1.94E-02** |
| ENSG00000118690.15 | **ARMC2** | 84071 | protein_coding | **8.585** | 1.396 | 0.420 | **2.632** | 9.17E-03 | **1.94E-02** |
| ENSG00000141564.16 | **RPTOR** | 57521 | protein_coding | **384.822** | -0.363 | -0.109 | **0.778** | 9.18E-03 | **1.94E-02** |
| ENSG00000167083.9 | **GNGT2** | 2793 | protein_coding | **679.055** | -0.376 | -0.113 | **0.770** | 9.20E-03 | **1.94E-02** |
| ENSG00000239264.11 | **TXNDC5** | 81567 | protein_coding | **5402.551** | -0.368 | -0.111 | **0.775** | 9.22E-03 | **1.95E-02** |
| ENSG00000155111.17 | **CDK19** | 23097 | protein_coding | **544.069** | 0.280 | 0.084 | **1.214** | 9.23E-03 | **1.95E-02** |
| ENSG00000116212.18 | **LRRC42** | 115353 | protein_coding | **346.799** | -0.442 | -0.133 | **0.736** | 9.24E-03 | **1.95E-02** |
| ENSG00000290121.2 | **ENSG00000290121** | | lncRNA | **14.183** | 1.224 | 0.368 | **2.336** | 9.26E-03 | **1.96E-02** |
| ENSG00000173588.18 | **CEP83** | 51134 | protein_coding | **299.936** | -0.386 | -0.116 | **0.765** | 9.26E-03 | **1.96E-02** |

| | | | | | | | | | |
|---|---|---|---|---|---|---|---|---|---|
| ENSG00000293738.1 | **ENSG00000293738** | | lncRNA | **15.545** | 1.074 | 0.323 | **2.106** | 9.30E-03 | **1.96E-02** |
| ENSG00000291129.2 | **PPIEL** | 728448 | lncRNA | **430.535** | 0.377 | 0.113 | **1.299** | 9.31E-03 | **1.96E-02** |
| ENSG00000136770.13 | **DNAJC1** | 64215 | protein_coding | **480.561** | -0.331 | -0.100 | **0.795** | 9.31E-03 | **1.97E-02** |
| ENSG00000155265.14 | **GOLGA7B** | 401647 | protein_coding | **7.215** | 1.767 | 0.532 | **3.403** | 9.32E-03 | **1.97E-02** |
| ENSG00000279168.3 | **ENSG00000279168** | | lncRNA | **96.775** | 0.720 | 0.217 | **1.648** | 9.35E-03 | **1.97E-02** |
| ENSG00000100564.11 | **PIGH** | 5283 | protein_coding | **223.641** | 0.489 | 0.147 | **1.404** | 9.36E-03 | **1.97E-02** |
| ENSG00000291185.2 | **ENSG00000291185** | | lncRNA | **5.804** | 2.323 | 0.699 | **5.004** | 9.36E-03 | **1.97E-02** |
| ENSG00000277218.1 | **ENSG00000277218** | | lncRNA | **12.329** | 1.099 | 0.331 | **2.142** | 9.37E-03 | **1.98E-02** |
| ENSG00000229106.1 | **BTBD6P1** | | processed_pseud | **16.551** | -0.947 | -0.285 | **0.519** | 9.38E-03 | **1.98E-02** |
| ENSG00000198690.12 | **FAN1** | 22909 | protein_coding | **810.249** | 0.278 | 0.084 | **1.212** | 9.38E-03 | **1.98E-02** |
| ENSG00000168298.7 | **H1-4** | 3008 | protein_coding | **17.755** | -1.258 | -0.379 | **0.418** | 9.40E-03 | **1.98E-02** |
| ENSG00000289311.3 | **ENSG00000289311** | | lncRNA | **7.293** | 1.448 | 0.436 | **2.728** | 9.40E-03 | **1.98E-02** |
| ENSG00000271380.1 | **PYGO2-AS1** | | lncRNA | **29.219** | 0.646 | 0.194 | **1.565** | 9.40E-03 | **1.98E-02** |
| ENSG00000259943.3 | **ZMPSTE24-DT** | | lncRNA | **144.898** | 0.428 | 0.129 | **1.345** | 9.42E-03 | **1.99E-02** |
| ENSG00000121933.20 | **TMIGD3** | 57413 | protein_coding | **10.272** | 1.203 | 0.362 | **2.302** | 9.48E-03 | **2.00E-02** |
| ENSG00000259985.1 | **ENSG00000259985** | | lncRNA | **32.648** | 0.674 | 0.203 | **1.595** | 9.48E-03 | **2.00E-02** |
| ENSG00000262728.7 | **ARHGAP11** | 10192778 | lncRNA | **66.036** | 0.598 | 0.180 | **1.513** | 9.48E-03 | **2.00E-02** |
| ENSG00000260317.1 | **LINC02986** | | lncRNA | **9.172** | 1.342 | 0.404 | **2.536** | 9.48E-03 | **2.00E-02** |
| ENSG00000259775.1 | **EIF5-DT** | | lncRNA | **6.838** | 1.390 | 0.418 | **2.621** | 9.49E-03 | **2.00E-02** |
| ENSG00000246740.3 | **PLA2G4E-A** | 10192838 | lncRNA | **5.427** | 1.802 | 0.543 | **3.488** | 9.49E-03 | **2.00E-02** |
| ENSG00000212864.6 | **RNF208** | 727800 | protein_coding | **225.522** | 0.488 | 0.147 | **1.403** | 9.50E-03 | **2.00E-02** |
| ENSG00000221923.11 | **ZNF880** | 400713 | protein_coding | **241.852** | -0.512 | -0.154 | **0.701** | 9.50E-03 | **2.00E-02** |
| ENSG00000198695.2 | **MT-ND6** | 4541 | protein_coding | **1169.507** | -0.454 | -0.137 | **0.730** | 9.51E-03 | **2.00E-02** |
| ENSG00000157625.18 | **TAB3** | 257397 | protein_coding | **230.890** | 0.467 | 0.141 | **1.382** | 9.52E-03 | **2.00E-02** |
| ENSG00000293246.2 | **ZNF321P** | 399669 | lncRNA | **35.351** | -0.760 | -0.229 | **0.591** | 9.53E-03 | **2.01E-02** |
| ENSG00000137869.18 | **CYP19A1** | 1588 | protein_coding | **12.595** | 1.694 | 0.510 | **3.235** | 9.53E-03 | **2.01E-02** |
| ENSG00000275632.1 | **ENSG00000275632** | | lncRNA | **33.961** | -0.776 | -0.234 | **0.584** | 9.54E-03 | **2.01E-02** |
| ENSG00000225931.3 | **ENSG00000225931** | | TEC | **15.625** | -1.079 | -0.325 | **0.473** | 9.55E-03 | **2.01E-02** |
| ENSG00000297420.1 | **ENSG00000297420** | | lncRNA | **7.970** | 1.786 | 0.538 | **3.449** | 9.56E-03 | **2.01E-02** |
| ENSG00000282381.3 | **ENSG00000282381** | | lncRNA | **11.738** | -1.287 | -0.387 | **0.410** | 9.59E-03 | **2.02E-02** |
| ENSG00000143545.12 | **RAB13** | 5872 | protein_coding | **1188.867** | 0.309 | 0.093 | **1.239** | 9.61E-03 | **2.02E-02** |

| | | | | | | | | | |
|---|---|---|---|---|---|---|---|---|---|
| ENSG00000055917.19 | **PUM2** | 23369 | protein_coding | **1083.654** | -0.292 | -0.088 | **0.817** | 9.62E-03 | **2.02E-02** |
| ENSG00000128513.18 | **POT1** | 25913 | protein_coding | **269.820** | -0.519 | -0.156 | **0.698** | 9.62E-03 | **2.02E-02** |
| ENSG00000125878.8 | **TCF15** | 6939 | protein_coding | **13.279** | 1.267 | 0.382 | **2.407** | 9.63E-03 | **2.02E-02** |
| ENSG00000272333.9 | **KMT2B** | 9757 | protein_coding | **1498.312** | -0.378 | -0.114 | **0.770** | 9.63E-03 | **2.03E-02** |
| ENSG00000108582.14 | **CPD** | 1362 | protein_coding | **204.479** | 0.365 | 0.110 | **1.288** | 9.69E-03 | **2.04E-02** |
| ENSG00000291127.2 | **ANKRD36B** | 645784 | lncRNA | **116.882** | -0.572 | -0.172 | **0.673** | 9.71E-03 | **2.04E-02** |
| ENSG00000171103.13 | **TRMT61B** | 55006 | protein_coding | **149.270** | -0.446 | -0.134 | **0.734** | 9.71E-03 | **2.04E-02** |
| ENSG00000198046.15 | **ZNF667** | 63934 | protein_coding | **18.049** | 1.336 | 0.402 | **2.525** | 9.72E-03 | **2.04E-02** |
| ENSG00000198919.15 | **DZIP3** | 9666 | protein_coding | **435.320** | 0.276 | 0.083 | **1.211** | 9.73E-03 | **2.04E-02** |
| ENSG00000273213.4 | **H3-7** | 440686 | protein_coding | **7.594** | 1.294 | 0.389 | **2.452** | 9.74E-03 | **2.05E-02** |
| ENSG00000278677.2 | **H2AC17** | 8336 | protein_coding | **12.660** | -1.076 | -0.324 | **0.474** | 9.75E-03 | **2.05E-02** |
| ENSG00000111859.19 | **NEDD9** | 4739 | protein_coding | **594.361** | 0.468 | 0.141 | **1.383** | 9.76E-03 | **2.05E-02** |
| ENSG00000052802.15 | **MSMO1** | 6307 | protein_coding | **927.091** | -0.533 | -0.160 | **0.691** | 9.76E-03 | **2.05E-02** |
| ENSG00000305228.1 | **ENSG00000305228** | | lncRNA | **5.256** | 2.153 | 0.648 | **4.447** | 9.76E-03 | **2.05E-02** |
| ENSG00000155760.3 | **FZD7** | 8324 | protein_coding | **33.913** | 0.733 | 0.221 | **1.662** | 9.77E-03 | **2.05E-02** |
| ENSG00000118564.17 | **FBXL5** | 26234 | protein_coding | **352.124** | -0.297 | -0.089 | **0.814** | 9.78E-03 | **2.05E-02** |
| ENSG00000161558.13 | **TMEM143** | 55260 | protein_coding | **251.096** | -0.400 | -0.120 | **0.758** | 9.79E-03 | **2.05E-02** |
| ENSG00000178537.12 | **SLC25A20** | 788 | protein_coding | **300.211** | 0.375 | 0.113 | **1.297** | 9.79E-03 | **2.06E-02** |
| ENSG00000159593.17 | **NAE1** | 8883 | protein_coding | **545.822** | -0.274 | -0.083 | **0.827** | 9.82E-03 | **2.06E-02** |
| ENSG00000160213.12 | **CSTB** | 1476 | protein_coding | **2702.740** | 0.390 | 0.117 | **1.310** | 9.82E-03 | **2.06E-02** |
| ENSG00000100258.20 | **LMF2** | 91289 | protein_coding | **1266.848** | -0.294 | -0.088 | **0.816** | 9.82E-03 | **2.06E-02** |
| ENSG00000288586.2 | **ENSG00000288586** | | lncRNA | **14.434** | 0.971 | 0.292 | **1.961** | 9.83E-03 | **2.06E-02** |
| ENSG00000251022.6 | **THAP9-AS1** | | lncRNA | **1159.754** | 0.331 | 0.100 | **1.258** | 9.83E-03 | **2.06E-02** |
| ENSG00000284825.1 | **ENSG00000284825** | | lncRNA | **9.159** | 1.478 | 0.445 | **2.785** | 9.84E-03 | **2.06E-02** |
| ENSG00000295613.1 | **ENSG00000295613** | | lncRNA | **9.543** | 1.316 | 0.396 | **2.489** | 9.86E-03 | **2.07E-02** |
| ENSG00000117543.24 | **DPH5** | 51611 | protein_coding | **602.839** | -0.324 | -0.097 | **0.799** | 9.87E-03 | **2.07E-02** |
| ENSG00000232434.4 | **AJM1** | 389813 | protein_coding | **46.807** | 0.571 | 0.172 | **1.486** | 9.87E-03 | **2.07E-02** |
| ENSG00000066032.20 | **CTNNA2** | 1496 | protein_coding | **17.751** | -0.978 | -0.294 | **0.508** | 9.88E-03 | **2.07E-02** |
| ENSG00000143952.22 | **VPS54** | 51542 | protein_coding | **338.530** | 0.294 | 0.088 | **1.226** | 9.88E-03 | **2.07E-02** |
| ENSG00000184905.11 | **TCEAL2** | 140597 | protein_coding | **14.732** | 1.303 | 0.392 | **2.467** | 9.88E-03 | **2.07E-02** |
| ENSG00000049759.22 | **NEDD4L** | 23327 | protein_coding | **312.263** | 0.379 | 0.114 | **1.300** | 9.91E-03 | **2.08E-02** |

| | | | | | | | | | |
|---|---|---|---|---|---|---|---|---|---|
| ENSG00000259363.7 | **ENSG00000259363** | | lncRNA | **105.768** | 0.475 | 0.143 | **1.390** | 9.91E-03 | **2.08E-02** |
| ENSG00000273102.3 | **ENSG00000273102** | | lncRNA | **7.822** | 1.415 | 0.426 | **2.667** | 9.91E-03 | **2.08E-02** |
| ENSG00000300702.1 | **ENSG00000300702** | | lncRNA | **23.229** | -0.906 | -0.273 | **0.534** | 9.92E-03 | **2.08E-02** |
| ENSG00000276445.1 | **ENSG00000276445** | | lncRNA | **14.353** | 1.006 | 0.303 | **2.008** | 9.94E-03 | **2.08E-02** |
| ENSG00000272663.2 | **PPP1R21-DT** | | lncRNA | **7.868** | 1.453 | 0.437 | **2.737** | 9.94E-03 | **2.08E-02** |
| ENSG00000157077.18 | **ZFYVE9** | 9372 | protein_coding | **101.077** | 0.496 | 0.149 | **1.410** | 9.95E-03 | **2.08E-02** |
| ENSG00000149218.7 | **ENDOD1** | 23052 | protein_coding | **552.097** | 0.408 | 0.123 | **1.327** | 9.96E-03 | **2.09E-02** |
| ENSG00000176244.8 | **ACBD7** | 414149 | protein_coding | **11.097** | 1.556 | 0.469 | **2.941** | 9.98E-03 | **2.09E-02** |
| ENSG00000226314.10 | **ZKSCAN8P1** | | protein_coding | **44.734** | 0.562 | 0.169 | **1.477** | 9.98E-03 | **2.09E-02** |
| ENSG00000149289.12 | **ZC3H12C** | 85463 | protein_coding | **351.240** | -0.419 | -0.126 | **0.748** | 9.99E-03 | **2.09E-02** |
| ENSG00000273541.1 | **ENSG00000273541** | | misc_RNA | **40.559** | 0.681 | 0.205 | **1.603** | 1.00E-02 | **2.09E-02** |
| ENSG00000154642.13 | **C21orf91** | 54149 | protein_coding | **272.816** | -0.452 | -0.136 | **0.731** | 1.00E-02 | **2.09E-02** |
| ENSG00000103489.14 | **XYLT1** | 64131 | protein_coding | **116.611** | -0.611 | -0.184 | **0.655** | 1.00E-02 | **2.10E-02** |
| ENSG00000185442.15 | **FAM174B** | 400451 | protein_coding | **661.526** | 0.427 | 0.129 | **1.344** | 1.00E-02 | **2.10E-02** |
| ENSG00000170919.17 | **TPT1-AS1** | | lncRNA | **703.276** | 0.310 | 0.093 | **1.239** | 1.00E-02 | **2.10E-02** |
| ENSG00000183475.15 | **ASB7** | 140460 | protein_coding | **348.826** | 0.269 | 0.081 | **1.205** | 1.01E-02 | **2.10E-02** |
| ENSG00000162623.18 | **TYW3** | 127253 | protein_coding | **785.884** | -0.497 | -0.150 | **0.709** | 1.01E-02 | **2.10E-02** |
| ENSG00000197119.14 | **SLC25A29** | 123096 | protein_coding | **794.347** | 0.387 | 0.117 | **1.308** | 1.01E-02 | **2.10E-02** |
| ENSG00000165672.9 | **PRDX3** | 10935 | protein_coding | **1221.330** | -0.273 | -0.082 | **0.828** | 1.01E-02 | **2.11E-02** |
| ENSG00000153391.18 | **INO80C** | 125476 | protein_coding | **469.481** | 0.359 | 0.108 | **1.283** | 1.01E-02 | **2.11E-02** |
| ENSG00000176463.15 | **SLCO3A1** | 28232 | protein_coding | **14.431** | 1.445 | 0.435 | **2.723** | 1.01E-02 | **2.11E-02** |
| ENSG00000177189.16 | **RPS6KA3** | 6197 | protein_coding | **444.078** | -0.362 | -0.109 | **0.778** | 1.01E-02 | **2.11E-02** |
| ENSG00000307060.1 | **ENSG00000307060** | | lncRNA | **5.456** | 1.886 | 0.568 | **3.696** | 1.01E-02 | **2.11E-02** |
| ENSG00000164096.15 | **ARLN** | 401152 | protein_coding | **555.394** | -0.428 | -0.129 | **0.743** | 1.01E-02 | **2.11E-02** |
| ENSG00000168564.8 | **CDKN2AIP** | 55602 | protein_coding | **172.105** | -0.571 | -0.172 | **0.673** | 1.01E-02 | **2.11E-02** |
| ENSG00000171307.21 | **ZDHHC16** | 84287 | protein_coding | **597.074** | -0.289 | -0.087 | **0.819** | 1.01E-02 | **2.12E-02** |
| ENSG00000293258.1 | **ENSG00000293258** | | lncRNA | **13.588** | -1.134 | -0.341 | **0.456** | 1.01E-02 | **2.12E-02** |
| ENSG00000262624.2 | **ENSG00000262624** | | lncRNA | **8.365** | -1.307 | -0.394 | **0.404** | 1.02E-02 | **2.12E-02** |
| ENSG00000274020.6 | **LINC01138** | 388685 | lncRNA | **250.091** | -0.398 | -0.120 | **0.759** | 1.02E-02 | **2.12E-02** |
| ENSG00000167257.13 | **RNF214** | 257160 | protein_coding | **383.138** | 0.384 | 0.115 | **1.305** | 1.02E-02 | **2.12E-02** |
| ENSG00000130726.14 | **TRIM28** | 10155 | protein_coding | **4381.278** | -0.348 | -0.105 | **0.786** | 1.02E-02 | **2.12E-02** |

| | | | | | | | | | |
|---|---|---|---|---|---|---|---|---|---|
| ENSG00000021826.20 | **CPS1** | 1373 | protein_coding | **57.082** | 0.769 | 0.232 | **1.704** | 1.02E-02 | **2.12E-02** |
| ENSG00000180628.17 | **PCGF5** | 84333 | protein_coding | **929.496** | -0.279 | -0.084 | **0.824** | 1.02E-02 | **2.13E-02** |
| ENSG00000085832.20 | **EPS15** | 2060 | protein_coding | **1807.384** | 0.299 | 0.090 | **1.230** | 1.02E-02 | **2.13E-02** |
| ENSG00000143919.17 | **CAMKMT** | 79823 | protein_coding | **124.446** | 0.390 | 0.117 | **1.310** | 1.02E-02 | **2.13E-02** |
| ENSG00000299597.1 | **ENSG00000299597** | | lncRNA | **5.976** | 1.528 | 0.460 | **2.885** | 1.02E-02 | **2.13E-02** |
| ENSG00000288966.2 | **ENSG00000288966** | | lncRNA | **21.131** | 0.912 | 0.274 | **1.881** | 1.02E-02 | **2.14E-02** |
| ENSG00000289758.1 | **ENSG00000289758** | | protein_coding | **12.686** | 1.729 | 0.521 | **3.316** | 1.03E-02 | **2.14E-02** |
| ENSG00000109083.16 | **IFT20** | 90410 | protein_coding | **396.995** | 0.326 | 0.098 | **1.253** | 1.03E-02 | **2.14E-02** |
| ENSG00000118705.20 | **RPN2** | 6185 | protein_coding | **2347.284** | -0.364 | -0.110 | **0.777** | 1.03E-02 | **2.14E-02** |
| ENSG00000010818.13 | **HIVEP2** | 3097 | protein_coding | **483.861** | 0.332 | 0.100 | **1.258** | 1.03E-02 | **2.15E-02** |
| ENSG00000127995.19 | **CASD1** | 64921 | protein_coding | **85.377** | -0.465 | -0.140 | **0.725** | 1.03E-02 | **2.15E-02** |
| ENSG00000133114.20 | **GPALPP1** | 55425 | protein_coding | **336.079** | -0.348 | -0.105 | **0.786** | 1.03E-02 | **2.15E-02** |
| ENSG00000275180.3 | **ENSG00000275180** | | lncRNA | **53.028** | -0.732 | -0.220 | **0.602** | 1.04E-02 | **2.16E-02** |
| ENSG00000161671.18 | **EMC10** | 284361 | protein_coding | **1201.783** | -0.275 | -0.083 | **0.826** | 1.04E-02 | **2.16E-02** |
| ENSG00000171763.22 | **AFG2B** | 79029 | protein_coding | **158.578** | -0.480 | -0.145 | **0.717** | 1.04E-02 | **2.17E-02** |
| ENSG00000291027.2 | **UBE2Q2P1** | 388165 | lncRNA | **36.095** | 1.393 | 0.419 | **2.626** | 1.04E-02 | **2.17E-02** |
| ENSG00000105204.15 | **DYRK1B** | 9149 | protein_coding | **267.314** | 0.340 | 0.102 | **1.266** | 1.04E-02 | **2.17E-02** |
| ENSG00000149926.16 | **TLCD3B** | 83723 | protein_coding | **15.949** | 1.016 | 0.306 | **2.023** | 1.04E-02 | **2.17E-02** |
| ENSG00000278558.6 | **TMEM191I** | 728229 | protein_coding | **56.646** | -0.671 | -0.202 | **0.628** | 1.04E-02 | **2.18E-02** |
| ENSG00000146802.15 | **TMEM168** | 64418 | protein_coding | **309.256** | -0.456 | -0.137 | **0.729** | 1.04E-02 | **2.18E-02** |
| ENSG00000232593.10 | **KANTR** | 102723508 | protein_coding | **425.592** | 0.286 | 0.086 | **1.219** | 1.05E-02 | **2.18E-02** |
| ENSG00000119723.19 | **COQ6** | 51004 | protein_coding | **333.966** | -0.293 | -0.088 | **0.816** | 1.05E-02 | **2.18E-02** |
| ENSG00000182919.17 | **C11orf54** | 28970 | protein_coding | **409.739** | 0.428 | 0.129 | **1.345** | 1.05E-02 | **2.18E-02** |
| ENSG00000271754.1 | **ENSG00000271754** | | lncRNA | **16.285** | 1.190 | 0.358 | **2.281** | 1.05E-02 | **2.18E-02** |
| ENSG00000161381.16 | **PLXDC1** | 57125 | protein_coding | **14.897** | 1.037 | 0.312 | **2.051** | 1.05E-02 | **2.18E-02** |
| ENSG00000298040.1 | **ENSG00000298040** | | lncRNA | **4.422** | 2.063 | 0.621 | **4.179** | 1.05E-02 | **2.19E-02** |
| ENSG00000309954.1 | **ENSG00000309954** | | lncRNA | **6.713** | 1.963 | 0.591 | **3.898** | 1.06E-02 | **2.20E-02** |
| ENSG00000185875.15 | **THNSL1** | 79896 | protein_coding | **79.561** | -0.480 | -0.144 | **0.717** | 1.06E-02 | **2.20E-02** |
| ENSG00000284523.3 | **ENSG0000(** | 105375421 | lncRNA | **13.766** | 0.914 | 0.275 | **1.885** | 1.06E-02 | **2.20E-02** |
| ENSG00000109180.17 | **OCIAD1** | 54940 | protein_coding | **1548.581** | -0.299 | -0.090 | **0.813** | 1.06E-02 | **2.21E-02** |
| ENSG00000212123.5 | **PRR22** | 163154 | protein_coding | **52.090** | -0.668 | -0.201 | **0.629** | 1.06E-02 | **2.21E-02** |

| | | | | | | | | | |
|---|---|---|---|---|---|---|---|---|---|
| ENSG00000130985.19 | **UBA1** | 7317 | protein_coding | **4423.954** | -0.279 | -0.084 | **0.824** | 1.07E-02 | **2.22E-02** |
| ENSG00000281207.1 | **SLFNL1-AS:** | 10050717ε | lncRNA | **13.061** | -1.019 | -0.307 | **0.493** | 1.07E-02 | **2.22E-02** |
| ENSG00000165802.25 | **NSMF** | 26012 | protein_coding | **1484.237** | 0.282 | 0.085 | **1.216** | 1.07E-02 | **2.23E-02** |
| ENSG00000125505.19 | **MBOAT7** | 79143 | protein_coding | **633.809** | 0.314 | 0.094 | **1.243** | 1.07E-02 | **2.23E-02** |
| ENSG00000175414.9 | **ARL10** | 285598 | protein_coding | **245.873** | 0.664 | 0.200 | **1.584** | 1.07E-02 | **2.23E-02** |
| ENSG00000258057.7 | **BCDIN3D-A** | 10028684 | lncRNA | **13.584** | 1.173 | 0.353 | **2.255** | 1.07E-02 | **2.23E-02** |
| ENSG00000230648.2 | **ENSG00000230648** | | lncRNA | **24.108** | 0.713 | 0.215 | **1.639** | 1.08E-02 | **2.23E-02** |
| ENSG00000141429.16 | **GALNT1** | 2589 | protein_coding | **368.562** | -0.531 | -0.160 | **0.692** | 1.08E-02 | **2.24E-02** |
| ENSG00000168661.17 | **ZNF30** | 90075 | protein_coding | **74.645** | 0.535 | 0.161 | **1.449** | 1.08E-02 | **2.24E-02** |
| ENSG00000064652.13 | **SNX24** | 28966 | protein_coding | **86.269** | 0.505 | 0.152 | **1.419** | 1.08E-02 | **2.24E-02** |
| ENSG00000080802.23 | **CNOT4** | 4850 | protein_coding | **407.102** | -0.285 | -0.086 | **0.821** | 1.08E-02 | **2.24E-02** |
| ENSG00000135637.17 | **CCDC142** | 84865 | protein_coding | **117.042** | 0.551 | 0.166 | **1.466** | 1.08E-02 | **2.24E-02** |
| ENSG00000304520.1 | **ENSG00000304520** | | lncRNA | **6.500** | 1.798 | 0.541 | **3.476** | 1.08E-02 | **2.25E-02** |
| ENSG00000172009.17 | **THOP1** | 7064 | protein_coding | **2240.742** | -0.357 | -0.107 | **0.781** | 1.08E-02 | **2.25E-02** |
| ENSG00000140386.15 | **SCAPER** | 49855 | protein_coding | **162.593** | 0.454 | 0.137 | **1.370** | 1.08E-02 | **2.25E-02** |
| ENSG00000079215.17 | **SLC1A3** | 6507 | protein_coding | **20.375** | 1.330 | 0.400 | **2.514** | 1.08E-02 | **2.25E-02** |
| ENSG00000164168.10 | **TMEM184(** | 55751 | protein_coding | **262.007** | 0.528 | 0.159 | **1.442** | 1.08E-02 | **2.25E-02** |
| ENSG00000295087.1 | **ENSG00000295087** | | lncRNA | **39.903** | -0.722 | -0.217 | **0.606** | 1.08E-02 | **2.25E-02** |
| ENSG00000142687.21 | **KIAA0319L** | 79932 | protein_coding | **1651.227** | 0.271 | 0.081 | **1.206** | 1.09E-02 | **2.26E-02** |
| ENSG00000136874.13 | **STX17** | 55014 | protein_coding | **558.898** | 0.297 | 0.089 | **1.229** | 1.09E-02 | **2.26E-02** |
| ENSG00000167034.11 | **NKX3-1** | 4824 | protein_coding | **7.467** | 1.414 | 0.426 | **2.665** | 1.09E-02 | **2.26E-02** |
| ENSG00000290935.2 | **ENSG00000290935** | | lncRNA | **4.127** | 2.739 | 0.825 | **6.677** | 1.09E-02 | **2.26E-02** |
| ENSG00000304801.1 | **ENSG00000304801** | | lncRNA | **7.067** | 1.532 | 0.461 | **2.892** | 1.09E-02 | **2.27E-02** |
| ENSG00000262801.7 | **ENSG00000262801** | | lncRNA | **15.506** | 1.199 | 0.361 | **2.296** | 1.09E-02 | **2.27E-02** |
| ENSG00000286681.1 | **ENSG00000286681** | | lncRNA | **26.966** | 0.678 | 0.204 | **1.600** | 1.10E-02 | **2.27E-02** |
| ENSG00000132541.13 | **RIDA** | 10247 | protein_coding | **287.157** | -0.319 | -0.096 | **0.802** | 1.10E-02 | **2.27E-02** |
| ENSG00000104814.14 | **MAP4K1** | 11184 | protein_coding | **4281.104** | -0.556 | -0.167 | **0.680** | 1.10E-02 | **2.28E-02** |
| ENSG00000267383.10 | **ENSG00000267383** | | lncRNA | **14.307** | -1.240 | -0.373 | **0.423** | 1.10E-02 | **2.28E-02** |
| ENSG00000291225.2 | **ENSG00000291225** | | lncRNA | **43.541** | 1.282 | 0.386 | **2.431** | 1.10E-02 | **2.28E-02** |
| ENSG00000004399.15 | **PLXND1** | 23129 | protein_coding | **39.761** | 1.106 | 0.333 | **2.152** | 1.10E-02 | **2.28E-02** |
| ENSG00000119314.19 | **PTBP3** | 9991 | protein_coding | **885.595** | -0.313 | -0.094 | **0.805** | 1.10E-02 | **2.28E-02** |

| | | | | | | | | | |
|---|---|---|---|---|---|---|---|---|---|
| ENSG00000084110.13 | **HAL** | 3034 | protein_coding | **8.490** | 1.755 | 0.528 | **3.375** | 1.10E-02 | **2.28E-02** |
| ENSG00000156500.18 | **PABIR3** | 159091 | protein_coding | **144.179** | -0.446 | -0.134 | **0.734** | 1.10E-02 | **2.28E-02** |
| ENSG00000259171.1 | **ENSG00000259171** | | protein_coding | **18.942** | 1.151 | 0.347 | **2.221** | 1.10E-02 | **2.28E-02** |
| ENSG00000213853.12 | **EMP2** | 2013 | protein_coding | **21.284** | 1.172 | 0.353 | **2.254** | 1.10E-02 | **2.29E-02** |
| ENSG00000070761.10 | **CFAP20** | 29105 | protein_coding | **404.594** | -0.279 | -0.084 | **0.824** | 1.10E-02 | **2.29E-02** |
| ENSG00000302317.1 | **ENSG0000(** | 10798625( | lncRNA | **20.884** | 0.873 | 0.263 | **1.832** | 1.10E-02 | **2.29E-02** |
| ENSG00000123610.6 | **TNFAIP6** | 7130 | protein_coding | **20.487** | 1.159 | 0.349 | **2.233** | 1.11E-02 | **2.29E-02** |
| ENSG00000260924.4 | **LINC01311** | | lncRNA | **62.475** | -0.583 | -0.175 | **0.668** | 1.11E-02 | **2.30E-02** |
| ENSG00000178234.16 | **GALNT11** | 63917 | protein_coding | **219.434** | -0.367 | -0.110 | **0.775** | 1.11E-02 | **2.30E-02** |
| ENSG00000163138.21 | **PACRGL** | 133015 | protein_coding | **86.914** | -0.418 | -0.126 | **0.748** | 1.11E-02 | **2.30E-02** |
| ENSG00000293160.2 | **ENSG0000(** | 10537338: | lncRNA | **20.552** | 0.869 | 0.262 | **1.826** | 1.11E-02 | **2.30E-02** |
| ENSG00000196793.16 | **ZNF239** | 8187 | protein_coding | **33.661** | 0.955 | 0.287 | **1.938** | 1.11E-02 | **2.31E-02** |
| ENSG00000111319.15 | **SCNN1A** | 6337 | protein_coding | **7.829** | 1.923 | 0.579 | **3.791** | 1.12E-02 | **2.31E-02** |
| ENSG00000160145.17 | **KALRN** | 8997 | protein_coding | **59.233** | 0.828 | 0.249 | **1.776** | 1.12E-02 | **2.31E-02** |
| ENSG00000197629.7 | **MPEG1** | 219972 | protein_coding | **217.019** | -0.610 | -0.184 | **0.655** | 1.12E-02 | **2.31E-02** |
| ENSG00000103037.14 | **SETD6** | 79918 | protein_coding | **353.458** | 0.330 | 0.099 | **1.257** | 1.12E-02 | **2.31E-02** |
| ENSG00000110700.9 | **RPS13** | 6207 | protein_coding | **15767.343** | -0.289 | -0.087 | **0.819** | 1.12E-02 | **2.31E-02** |
| ENSG00000008282.12 | **SYPL1** | 6856 | protein_coding | **282.658** | -0.551 | -0.166 | **0.682** | 1.12E-02 | **2.31E-02** |
| ENSG00000067900.10 | **ROCK1** | 6093 | protein_coding | **493.205** | -0.414 | -0.125 | **0.751** | 1.12E-02 | **2.31E-02** |
| ENSG00000126756.14 | **UXT** | 8409 | protein_coding | **1636.242** | -0.265 | -0.080 | **0.832** | 1.12E-02 | **2.32E-02** |
| ENSG00000165389.8 | **SPTSSA** | 171546 | protein_coding | **269.573** | -0.291 | -0.088 | **0.817** | 1.12E-02 | **2.32E-02** |
| ENSG00000289471.2 | **ENSG00000289471** | | lncRNA | **16.148** | -1.128 | -0.339 | **0.458** | 1.12E-02 | **2.32E-02** |
| ENSG00000164543.9 | **STK17A** | 9263 | protein_coding | **864.036** | 0.425 | 0.128 | **1.342** | 1.12E-02 | **2.32E-02** |
| ENSG00000143590.16 | **EFNA3** | 1944 | protein_coding | **125.650** | 0.554 | 0.167 | **1.468** | 1.13E-02 | **2.33E-02** |
| ENSG00000224424.9 | **PRKAR2A-/** | 100506637 | lncRNA | **64.706** | 0.554 | 0.167 | **1.468** | 1.13E-02 | **2.33E-02** |
| ENSG00000071889.19 | **FAM3A** | 60343 | protein_coding | **680.016** | -0.263 | -0.079 | **0.833** | 1.13E-02 | **2.33E-02** |
| ENSG00000293902.1 | **ENSG00000293902** | | lncRNA | **10.497** | 1.073 | 0.323 | **2.104** | 1.13E-02 | **2.33E-02** |
| ENSG00000140839.15 | **CLEC18B** | 497190 | protein_coding | **12.858** | 1.554 | 0.468 | **2.936** | 1.13E-02 | **2.34E-02** |
| ENSG00000183150.10 | **GPR19** | 2842 | protein_coding | **58.500** | -0.816 | -0.246 | **0.568** | 1.13E-02 | **2.34E-02** |
| ENSG00000272913.3 | **LINC03052** | 12713886! | lncRNA | **116.139** | -0.403 | -0.121 | **0.756** | 1.14E-02 | **2.36E-02** |
| ENSG00000286912.1 | **ENSG00000286912** | | lncRNA | **6.830** | -1.466 | -0.441 | **0.362** | 1.14E-02 | **2.36E-02** |

| | | | | | | | | | |
|---|---|---|---|---|---|---|---|---|---|
| ENSG00000141759.16 | **TXNL4A** | 10907 | protein_coding | **1246.323** | -0.295 | -0.089 | **0.815** | 1.14E-02 | **2.36E-02** |
| ENSG00000289555.2 | **ENSG00000289555** | | lncRNA | **5.950** | 1.551 | 0.467 | **2.931** | 1.15E-02 | **2.36E-02** |
| ENSG00000138336.11 | **TET1** | 80312 | protein_coding | **10.010** | -1.313 | -0.395 | **0.402** | 1.15E-02 | **2.37E-02** |
| ENSG00000198646.16 | **NCOA6** | 23054 | protein_coding | **596.944** | -0.366 | -0.110 | **0.776** | 1.15E-02 | **2.37E-02** |
| ENSG00000175265.21 | **GOLGA8A** | 23015 | protein_coding | **12123.824** | 0.288 | 0.087 | **1.221** | 1.15E-02 | **2.37E-02** |
| ENSG00000135297.19 | **MTO1** | 25821 | protein_coding | **419.573** | -0.315 | -0.095 | **0.804** | 1.15E-02 | **2.38E-02** |
| ENSG00000214145.9 | **LINC00887** | | lncRNA | **6.760** | 1.689 | 0.508 | **3.224** | 1.15E-02 | **2.38E-02** |
| ENSG00000183421.13 | **RIPK4** | 54101 | protein_coding | **26.621** | 0.973 | 0.293 | **1.963** | 1.15E-02 | **2.38E-02** |
| ENSG00000145794.18 | **MEGF10** | 84466 | protein_coding | **2.909** | 3.543 | 1.067 | **11.655** | 1.16E-02 | **2.38E-02** |
| ENSG00000162444.14 | **RBP7** | 116362 | protein_coding | **4.978** | 1.970 | 0.593 | **3.918** | 1.16E-02 | **2.38E-02** |
| ENSG00000221916.4 | **C19orf73** | 55150 | protein_coding | **18.526** | 0.833 | 0.251 | **1.781** | 1.16E-02 | **2.38E-02** |
| ENSG00000214253.11 | **FIS1** | 51024 | protein_coding | **816.817** | 0.276 | 0.083 | **1.211** | 1.16E-02 | **2.39E-02** |
| ENSG00000169994.22 | **MYO7B** | 4648 | protein_coding | **205.757** | -0.526 | -0.158 | **0.694** | 1.16E-02 | **2.39E-02** |
| ENSG00000107130.12 | **NCS1** | 23413 | protein_coding | **9.141** | -1.446 | -0.435 | **0.367** | 1.16E-02 | **2.39E-02** |
| ENSG00000136802.14 | **LRRC8A** | 56262 | protein_coding | **165.287** | -0.332 | -0.100 | **0.795** | 1.16E-02 | **2.39E-02** |
| ENSG00000134324.14 | **LPIN1** | 23175 | protein_coding | **945.517** | 0.447 | 0.135 | **1.364** | 1.16E-02 | **2.40E-02** |
| ENSG00000286162.5 | **INCR1** | 12490211 | lncRNA | **167.234** | 0.614 | 0.185 | **1.530** | 1.16E-02 | **2.40E-02** |
| ENSG00000099256.21 | **PRTFDC1** | 56952 | protein_coding | **27.790** | -0.860 | -0.259 | **0.551** | 1.16E-02 | **2.40E-02** |
| ENSG00000108381.13 | **ASPA** | 443 | protein_coding | **10.066** | -1.434 | -0.432 | **0.370** | 1.17E-02 | **2.40E-02** |
| ENSG00000228340.9 | **MIR646HG** | 284757 | lncRNA | **27.552** | -1.106 | -0.333 | **0.464** | 1.17E-02 | **2.41E-02** |
| ENSG00000225964.6 | **NRIR** | | lncRNA | **13.255** | -1.119 | -0.337 | **0.461** | 1.17E-02 | **2.41E-02** |
| ENSG00000228826.4 | **ENSG00000228826** | | lncRNA | **3.411** | 2.979 | 0.897 | **7.883** | 1.17E-02 | **2.41E-02** |
| ENSG00000124191.20 | **TOX2** | 84969 | protein_coding | **678.359** | 0.404 | 0.122 | **1.323** | 1.17E-02 | **2.41E-02** |
| ENSG00000291093.2 | **SVIL-AS1** | 10272431 | lncRNA | **149.700** | -0.340 | -0.102 | **0.790** | 1.17E-02 | **2.41E-02** |
| ENSG00000135318.14 | **NT5E** | 4907 | protein_coding | **101.381** | 0.496 | 0.149 | **1.410** | 1.17E-02 | **2.41E-02** |
| ENSG00000290394.3 | **PKD1P6-NI** | 10536915 | lncRNA | **504.642** | -0.396 | -0.119 | **0.760** | 1.17E-02 | **2.41E-02** |
| ENSG00000175283.8 | **DOLK** | 22845 | protein_coding | **114.879** | -0.491 | -0.148 | **0.711** | 1.17E-02 | **2.41E-02** |
| ENSG00000138463.11 | **SLC49A4** | 84925 | protein_coding | **26.728** | 0.663 | 0.200 | **1.584** | 1.17E-02 | **2.42E-02** |
| ENSG00000268751.3 | **SCGB1B2P** | | lncRNA | **24.187** | 0.809 | 0.243 | **1.751** | 1.17E-02 | **2.42E-02** |
| ENSG00000232721.4 | **ENSG00000232721** | | lncRNA | **7.897** | 1.724 | 0.519 | **3.304** | 1.18E-02 | **2.42E-02** |
| ENSG00000306637.1 | **ENSG00000306637** | | lncRNA | **19.930** | 0.864 | 0.260 | **1.820** | 1.18E-02 | **2.43E-02** |

| | | | | | | | | | |
|---|---|---|---|---|---|---|---|---|---|
| ENSG00000155100.13 | **OTUD6B** | 51633 | protein_coding | **94.559** | -0.518 | -0.156 | **0.698** | 1.18E-02 | **2.43E-02** |
| ENSG00000279696.1 | **ENSG00000279696** | | TEC | **20.739** | 0.781 | 0.235 | **1.718** | 1.18E-02 | **2.43E-02** |
| ENSG00000290602.2 | **ENSG00000290602** | | lncRNA | **16.301** | -1.032 | -0.311 | **0.489** | 1.18E-02 | **2.43E-02** |
| ENSG00000288815.1 | **ENSG00000288815** | | lncRNA | **12.401** | 1.121 | 0.337 | **2.174** | 1.18E-02 | **2.43E-02** |
| ENSG00000196605.9 | **ZNF846** | 162993 | protein_coding | **102.312** | -0.466 | -0.140 | **0.724** | 1.18E-02 | **2.43E-02** |
| ENSG00000275055.3 | **ENSG00000275055** | | lncRNA | **16.091** | -1.069 | -0.322 | **0.477** | 1.18E-02 | **2.43E-02** |
| ENSG00000123570.6 | **RAB9B** | 51209 | protein_coding | **18.984** | 1.034 | 0.311 | **2.048** | 1.18E-02 | **2.44E-02** |
| ENSG00000272463.3 | **ENSG00000272463** | | lncRNA | **19.109** | 0.841 | 0.253 | **1.791** | 1.19E-02 | **2.44E-02** |
| ENSG00000123144.12 | **TRIR** | 79002 | protein_coding | **4274.613** | -0.284 | -0.085 | **0.821** | 1.19E-02 | **2.44E-02** |
| ENSG00000112514.20 | **CUTA** | 51596 | protein_coding | **2624.058** | 0.297 | 0.089 | **1.229** | 1.19E-02 | **2.44E-02** |
| ENSG00000277639.4 | **CHD9NB** | 10537126 | protein_coding | **12.757** | -1.039 | -0.313 | **0.487** | 1.19E-02 | **2.44E-02** |
| ENSG00000184730.14 | **APOBR** | 55911 | protein_coding | **71.866** | -0.609 | -0.183 | **0.656** | 1.19E-02 | **2.45E-02** |
| ENSG00000170298.18 | **LGALS9B** | 284194 | protein_coding | **17.691** | -1.555 | -0.468 | **0.340** | 1.19E-02 | **2.45E-02** |
| ENSG00000173545.7 | **ZNF622** | 90441 | protein_coding | **466.973** | -0.296 | -0.089 | **0.814** | 1.19E-02 | **2.45E-02** |
| ENSG00000155329.14 | **ZCCHC10** | 54819 | protein_coding | **166.474** | -0.413 | -0.124 | **0.751** | 1.19E-02 | **2.45E-02** |
| ENSG00000284642.1 | **ENSG00000284642** | | lncRNA | **7.359** | 1.410 | 0.424 | **2.657** | 1.19E-02 | **2.45E-02** |
| ENSG00000113391.21 | **ARB2A** | 83989 | protein_coding | **137.428** | -0.451 | -0.136 | **0.732** | 1.19E-02 | **2.45E-02** |
| ENSG00000107854.8 | **TNKS2** | 80351 | protein_coding | **282.571** | 0.460 | 0.138 | **1.375** | 1.20E-02 | **2.46E-02** |
| ENSG00000132471.13 | **WBP2** | 23558 | protein_coding | **2590.439** | 0.268 | 0.081 | **1.204** | 1.20E-02 | **2.46E-02** |
| ENSG00000258056.4 | **CD63-AS1** | 10536977 | lncRNA | **83.513** | -0.423 | -0.127 | **0.746** | 1.20E-02 | **2.46E-02** |
| ENSG00000148600.17 | **CDHR1** | 92211 | protein_coding | **137.805** | 0.681 | 0.205 | **1.603** | 1.20E-02 | **2.46E-02** |
| ENSG00000182700.5 | **IGIP** | 492311 | protein_coding | **78.362** | 0.549 | 0.165 | **1.463** | 1.20E-02 | **2.46E-02** |
| ENSG00000251191.9 | **LINC00589** | 619351 | lncRNA | **8.482** | 1.466 | 0.441 | **2.762** | 1.20E-02 | **2.46E-02** |
| ENSG00000170322.17 | **NFRKB** | 4798 | protein_coding | **638.766** | 0.302 | 0.091 | **1.233** | 1.20E-02 | **2.46E-02** |
| ENSG00000289294.3 | **ENSG00000289294** | | lncRNA | **41.985** | 0.612 | 0.184 | **1.528** | 1.20E-02 | **2.46E-02** |
| ENSG00000189190.13 | **ZNF600** | 162966 | protein_coding | **366.345** | -0.341 | -0.103 | **0.790** | 1.20E-02 | **2.47E-02** |
| ENSG00000291065.2 | **ENSG0000(** | 441666 | lncRNA | **15.640** | -1.197 | -0.360 | **0.436** | 1.21E-02 | **2.47E-02** |
| ENSG00000293550.1 | **ENSG00000293550** | | lncRNA | **9.605** | 1.464 | 0.441 | **2.758** | 1.21E-02 | **2.49E-02** |
| ENSG00000107201.14 | **RIGI** | 23586 | protein_coding | **271.666** | -0.491 | -0.148 | **0.711** | 1.21E-02 | **2.49E-02** |
| ENSG00000240288.8 | **GHRLOS** | 10012679 | lncRNA | **43.385** | -0.586 | -0.177 | **0.666** | 1.22E-02 | **2.50E-02** |
| ENSG00000133983.18 | **COX16** | 51241 | protein_coding | **658.535** | -0.362 | -0.109 | **0.778** | 1.22E-02 | **2.50E-02** |

| | | | | | | | | | |
|---|---|---|---|---|---|---|---|---|---|
| ENSG00000160284.17 | **SPATC1L** | 84221 | protein_coding | **374.702** | 0.486 | 0.146 | **1.400** | 1.22E-02 | **2.50E-02** |
| ENSG00000117419.18 | **ERI3** | 79033 | protein_coding | **620.759** | -0.303 | -0.091 | **0.810** | 1.22E-02 | **2.50E-02** |
| ENSG00000115970.21 | **THADA** | 63892 | protein_coding | **420.192** | -0.355 | -0.107 | **0.782** | 1.22E-02 | **2.50E-02** |
| ENSG00000008086.14 | **CDKL5** | 6792 | protein_coding | **58.971** | 0.572 | 0.172 | **1.486** | 1.22E-02 | **2.50E-02** |
| ENSG00000236617.3 | **CHFR-DT** | | lncRNA | **7.382** | 1.368 | 0.412 | **2.581** | 1.22E-02 | **2.50E-02** |
| ENSG00000125459.20 | **MSTO1** | 55154 | protein_coding | **791.807** | -0.315 | -0.095 | **0.804** | 1.22E-02 | **2.51E-02** |
| ENSG00000152942.21 | **RAD17** | 5884 | protein_coding | **694.756** | 0.275 | 0.083 | **1.210** | 1.22E-02 | **2.51E-02** |
| ENSG00000279489.1 | **ENSG00000279489** | | TEC | **8.041** | 1.498 | 0.451 | **2.824** | 1.23E-02 | **2.51E-02** |
| ENSG00000134440.14 | **NARS1** | 4677 | protein_coding | **1137.794** | -0.422 | -0.127 | **0.746** | 1.23E-02 | **2.51E-02** |
| ENSG00000260898.7 | **ADPGK-AS** | 10028755 | lncRNA | **9.423** | -1.572 | -0.473 | **0.336** | 1.23E-02 | **2.52E-02** |
| ENSG00000185837.6 | **HDHD5-AS** | 10013071 | lncRNA | **6.131** | 1.471 | 0.443 | **2.771** | 1.23E-02 | **2.52E-02** |
| ENSG00000243646.13 | **IL10RB** | 3588 | protein_coding | **996.520** | 0.287 | 0.086 | **1.220** | 1.23E-02 | **2.52E-02** |
| ENSG00000157224.18 | **CLDN12** | 9069 | protein_coding | **109.901** | 0.362 | 0.109 | **1.285** | 1.23E-02 | **2.52E-02** |
| ENSG00000185905.6 | **C16orf54** | 283897 | protein_coding | **366.140** | -0.383 | -0.115 | **0.767** | 1.23E-02 | **2.53E-02** |
| ENSG00000204316.15 | **MRPL38** | 64978 | protein_coding | **1228.874** | -0.322 | -0.097 | **0.800** | 1.24E-02 | **2.53E-02** |
| ENSG00000248810.5 | **LINC02432** | 10050763 | lncRNA | **17.801** | 4.054 | 1.221 | **16.616** | 1.24E-02 | **2.53E-02** |
| ENSG00000247950.10 | **SEC24B-AS** | 10053318 | lncRNA | **35.078** | 0.662 | 0.199 | **1.583** | 1.24E-02 | **2.53E-02** |
| ENSG00000197808.15 | **ZNF461** | 92283 | protein_coding | **111.196** | -0.378 | -0.114 | **0.769** | 1.24E-02 | **2.53E-02** |
| ENSG00000100147.17 | **CCDC134** | 79879 | protein_coding | **336.434** | -0.309 | -0.093 | **0.807** | 1.24E-02 | **2.54E-02** |
| ENSG00000129158.13 | **SERGEF** | 26297 | protein_coding | **365.794** | -0.405 | -0.122 | **0.755** | 1.24E-02 | **2.54E-02** |
| ENSG00000161955.18 | **TNFSF13** | 8741 | protein_coding | **426.721** | -0.463 | -0.139 | **0.725** | 1.24E-02 | **2.54E-02** |
| ENSG00000100938.21 | **GMPR2** | 51292 | protein_coding | **1082.381** | 0.289 | 0.087 | **1.221** | 1.24E-02 | **2.54E-02** |
| ENSG00000125912.13 | **NCLN** | 56926 | protein_coding | **1366.398** | -0.345 | -0.104 | **0.788** | 1.24E-02 | **2.54E-02** |
| ENSG00000291214.1 | **ENSG00000291214** | | lncRNA | **10.581** | 1.097 | 0.330 | **2.139** | 1.25E-02 | **2.55E-02** |
| ENSG00000271554.1 | **ENSG00000271554** | | lncRNA | **12.605** | -1.113 | -0.335 | **0.462** | 1.25E-02 | **2.55E-02** |
| ENSG00000176868.3 | **ENSG00000176868** | | lncRNA | **6.619** | 1.473 | 0.443 | **2.776** | 1.25E-02 | **2.55E-02** |
| ENSG00000289236.2 | **ENSG00000289236** | | lncRNA | **6.100** | 1.629 | 0.490 | **3.093** | 1.25E-02 | **2.55E-02** |
| ENSG00000150773.14 | **PIH1D2** | 120379 | protein_coding | **16.039** | 0.923 | 0.278 | **1.897** | 1.25E-02 | **2.55E-02** |
| ENSG00000296016.1 | **ENSG00000296016** | | lncRNA | **7.068** | 1.400 | 0.421 | **2.639** | 1.25E-02 | **2.55E-02** |
| ENSG00000119946.14 | **CNNM1** | 26507 | protein_coding | **60.246** | -0.560 | -0.169 | **0.678** | 1.25E-02 | **2.56E-02** |
| ENSG00000085760.17 | **MTIF2** | 4528 | protein_coding | **628.086** | -0.303 | -0.091 | **0.810** | 1.25E-02 | **2.56E-02** |

| | | | | | | | | | |
|---|---|---|---|---|---|---|---|---|---|
| ENSG00000132510.13 | **KDM6B** | 23135 | protein_coding | **1012.332** | -0.285 | -0.086 | **0.821** | 1.25E-02 | **2.56E-02** |
| ENSG00000251442.9 | **LINC01094** | 10192889З | lncRNA | **12.194** | 1.502 | 0.452 | **2.832** | 1.26E-02 | **2.57E-02** |
| ENSG00000170027.8 | **YWHAG** | 7532 | protein_coding | **1670.547** | -0.287 | -0.086 | **0.820** | 1.26E-02 | **2.57E-02** |
| ENSG00000244405.10 | **ETV5** | 2119 | protein_coding | **212.766** | -0.546 | -0.165 | **0.685** | 1.26E-02 | **2.57E-02** |
| ENSG00000299938.1 | **ENSG00000299938** | | lncRNA | **7.068** | 1.352 | 0.407 | **2.553** | 1.26E-02 | **2.57E-02** |
| ENSG00000223350.2 | **IGLV9-49** | | IG_V_gene | **62.221** | -0.932 | -0.281 | **0.524** | 1.26E-02 | **2.57E-02** |
| ENSG00000266313.3 | **LINC03146** | | lncRNA | **5.729** | 1.629 | 0.490 | **3.092** | 1.26E-02 | **2.57E-02** |
| ENSG00000287046.2 | **ENSG00000287046** | | lncRNA | **20.555** | 0.795 | 0.239 | **1.735** | 1.26E-02 | **2.57E-02** |
| ENSG00000148337.24 | **CIZ1** | 25792 | protein_coding | **1181.750** | -0.334 | -0.101 | **0.793** | 1.26E-02 | **2.58E-02** |
| ENSG00000182150.22 | **ERCC6L2** | 375748 | protein_coding | **190.356** | -0.343 | -0.103 | **0.789** | 1.26E-02 | **2.58E-02** |
| ENSG00000060339.16 | **CCAR1** | 55749 | protein_coding | **1120.610** | -0.294 | -0.089 | **0.815** | 1.27E-02 | **2.58E-02** |
| ENSG00000129465.18 | **RIPK3** | 11035 | protein_coding | **266.768** | 0.312 | 0.094 | **1.241** | 1.27E-02 | **2.59E-02** |
| ENSG00000072864.18 | **NDE1** | 54820 | protein_coding | **1135.593** | -0.439 | -0.132 | **0.737** | 1.27E-02 | **2.59E-02** |
| ENSG00000188760.13 | **TMEM198** | 130612 | protein_coding | **188.128** | 0.478 | 0.144 | **1.393** | 1.27E-02 | **2.59E-02** |
| ENSG00000135519.10 | **KCNH3** | 23416 | protein_coding | **33.800** | 1.063 | 0.320 | **2.089** | 1.27E-02 | **2.59E-02** |
| ENSG00000253797.3 | **UTP14C** | 9724 | protein_coding | **31.260** | -1.260 | -0.379 | **0.418** | 1.27E-02 | **2.59E-02** |
| ENSG00000162992.6 | **NEUROD1** | 4760 | protein_coding | **6.083** | 2.268 | 0.683 | **4.817** | 1.27E-02 | **2.59E-02** |
| ENSG00000181381.17 | **DDX60L** | 91351 | protein_coding | **93.788** | -0.769 | -0.231 | **0.587** | 1.27E-02 | **2.59E-02** |
| ENSG00000197549.10 | **PRAMENP** | | transcribed_unprc | **18.512** | 1.044 | 0.314 | **2.062** | 1.27E-02 | **2.59E-02** |
| ENSG00000151640.15 | **DPYSL4** | 10570 | protein_coding | **79.355** | -0.740 | -0.223 | **0.599** | 1.27E-02 | **2.60E-02** |
| ENSG00000290117.2 | **ENSG00000290117** | | lncRNA | **23.781** | 0.715 | 0.215 | **1.642** | 1.28E-02 | **2.60E-02** |
| ENSG00000257093.8 | **DENND11** | 57189 | protein_coding | **858.494** | -0.365 | -0.110 | **0.776** | 1.28E-02 | **2.60E-02** |
| ENSG00000108256.11 | **NUFIP2** | 57532 | protein_coding | **827.337** | -0.270 | -0.081 | **0.830** | 1.28E-02 | **2.61E-02** |
| ENSG00000233654.3 | **NEMP2-DT** | 10537674ᴇ | lncRNA | **20.054** | -0.934 | -0.281 | **0.523** | 1.28E-02 | **2.61E-02** |
| ENSG00000136932.18 | **TRMO** | 51531 | protein_coding | **160.047** | -0.299 | -0.090 | **0.813** | 1.28E-02 | **2.61E-02** |
| ENSG00000254910.2 | **ENSG00000254910** | | lncRNA | **158.500** | 0.468 | 0.141 | **1.383** | 1.29E-02 | **2.62E-02** |
| ENSG00000188816.5 | **HMX2** | 3167 | protein_coding | **67.425** | -1.004 | -0.302 | **0.499** | 1.29E-02 | **2.62E-02** |
| ENSG00000224292.2 | **PORCN-DT** | | lncRNA | **9.961** | 1.104 | 0.332 | **2.150** | 1.29E-02 | **2.62E-02** |
| ENSG00000162702.10 | **ZNF281** | 23528 | protein_coding | **214.559** | 0.493 | 0.148 | **1.407** | 1.29E-02 | **2.62E-02** |
| ENSG00000289929.2 | **ENSG00000289929** | | lncRNA | **146.271** | 0.654 | 0.197 | **1.573** | 1.29E-02 | **2.62E-02** |
| ENSG00000110455.16 | **ACCS** | 84680 | protein_coding | **533.646** | 0.476 | 0.143 | **1.390** | 1.29E-02 | **2.63E-02** |

| | | | | | | | | | |
|---|---|---|---|---|---|---|---|---|---|
| ENSG00000108465.17 | **CDK5RAP3** | 80279 | protein_coding | **6600.196** | 0.327 | 0.098 | **1.254** | 1.30E-02 | **2.64E-02** |
| ENSG00000272086.3 | **GOLPH3-DT** | | lncRNA | **34.449** | 0.745 | 0.224 | **1.676** | 1.30E-02 | **2.64E-02** |
| ENSG00000139289.16 | **PHLDA1** | 22822 | protein_coding | **83.003** | 0.560 | 0.169 | **1.474** | 1.30E-02 | **2.64E-02** |
| ENSG00000271855.2 | **ENSG00000271855** | | lncRNA | **29.856** | 0.676 | 0.203 | **1.597** | 1.30E-02 | **2.64E-02** |
| ENSG00000281026.1 | **N4BP2L2-I** | 116828 | lncRNA | **119.573** | 0.398 | 0.120 | **1.318** | 1.30E-02 | **2.64E-02** |
| ENSG00000055950.19 | **MRPL43** | 84545 | protein_coding | **870.449** | 0.297 | 0.089 | **1.229** | 1.30E-02 | **2.64E-02** |
| ENSG00000091157.16 | **WDR7** | 23335 | protein_coding | **150.036** | 0.386 | 0.116 | **1.307** | 1.30E-02 | **2.65E-02** |
| ENSG00000185869.17 | **ZNF829** | 374899 | protein_coding | **59.700** | -0.598 | -0.180 | **0.661** | 1.30E-02 | **2.65E-02** |
| ENSG00000107593.17 | **PKD2L1** | 9033 | protein_coding | **8.301** | 2.014 | 0.606 | **4.038** | 1.30E-02 | **2.65E-02** |
| ENSG00000185920.19 | **PTCH1** | 5727 | protein_coding | **200.819** | 0.462 | 0.139 | **1.378** | 1.30E-02 | **2.65E-02** |
| ENSG00000245526.14 | **MIR9-2HG** | 645323 | lncRNA | **9.975** | 1.338 | 0.403 | **2.528** | 1.31E-02 | **2.65E-02** |
| ENSG00000146857.6 | **STRA8** | 346673 | protein_coding | **5.671** | 2.094 | 0.630 | **4.270** | 1.31E-02 | **2.66E-02** |
| ENSG00000115998.10 | **C2orf42** | 54980 | protein_coding | **192.785** | 0.342 | 0.103 | **1.267** | 1.31E-02 | **2.66E-02** |
| ENSG00000089009.18 | **RPL6** | 6128 | protein_coding | **41459.924** | -0.266 | -0.080 | **0.832** | 1.31E-02 | **2.66E-02** |
| ENSG00000285554.2 | **LINC02988** | | lncRNA | **29.558** | 0.670 | 0.202 | **1.592** | 1.31E-02 | **2.66E-02** |
| ENSG00000197291.10 | **RAMP2-AS** | 10019093 | lncRNA | **7.377** | 1.330 | 0.400 | **2.514** | 1.31E-02 | **2.67E-02** |
| ENSG00000279865.1 | **ENSG00000279865** | | TEC | **12.730** | 1.068 | 0.322 | **2.097** | 1.32E-02 | **2.67E-02** |
| ENSG00000226025.12 | **ENSG00000226025** | | transcribed_unpro | **11.404** | -1.555 | -0.468 | **0.340** | 1.32E-02 | **2.67E-02** |
| ENSG00000248275.5 | **TRIM52-AS** | 10050760 | lncRNA | **293.920** | 0.371 | 0.112 | **1.293** | 1.32E-02 | **2.68E-02** |
| ENSG00000171124.15 | **FUT3** | 2525 | protein_coding | **14.355** | 1.163 | 0.350 | **2.239** | 1.32E-02 | **2.68E-02** |
| ENSG00000165752.19 | **STK32C** | 282974 | protein_coding | **227.159** | 0.383 | 0.115 | **1.304** | 1.32E-02 | **2.69E-02** |
| ENSG00000143878.10 | **RHOB** | 388 | protein_coding | **134.425** | 0.483 | 0.145 | **1.398** | 1.32E-02 | **2.69E-02** |
| ENSG00000204396.12 | **VWA7** | 80737 | protein_coding | **13.308** | 1.101 | 0.331 | **2.145** | 1.32E-02 | **2.69E-02** |
| ENSG00000295667.1 | **ENSG00000295667** | | lncRNA | **25.191** | 0.736 | 0.222 | **1.666** | 1.32E-02 | **2.69E-02** |
| ENSG00000230533.6 | **LINC03004** | | lncRNA | **10.078** | 1.448 | 0.436 | **2.728** | 1.33E-02 | **2.69E-02** |
| ENSG00000120341.21 | **SEC16B** | 89866 | protein_coding | **7.184** | 2.366 | 0.712 | **5.154** | 1.33E-02 | **2.70E-02** |
| ENSG00000171840.15 | **NINJ2** | 4815 | protein_coding | **90.870** | -0.477 | -0.144 | **0.718** | 1.33E-02 | **2.70E-02** |
| ENSG00000289494.3 | **ENSG00000289494** | | lncRNA | **25.352** | 0.776 | 0.234 | **1.712** | 1.34E-02 | **2.71E-02** |
| ENSG00000196937.14 | **FAM3C** | 10447 | protein_coding | **572.905** | 0.421 | 0.127 | **1.339** | 1.34E-02 | **2.71E-02** |
| ENSG00000171159.6 | **BBLN** | 79095 | protein_coding | **1386.596** | 0.367 | 0.111 | **1.290** | 1.34E-02 | **2.71E-02** |
| ENSG00000146243.15 | **IRAK1BP1** | 134728 | protein_coding | **79.038** | 0.626 | 0.188 | **1.543** | 1.34E-02 | **2.71E-02** |

| | | | | | | | | | |
|---|---|---|---|---|---|---|---|---|---|
| ENSG00000240498.10 | **CDKN2B-A!** | 10004891Z | lncRNA | **17.735** | -0.918 | -0.276 | **0.529** | 1.34E-02 | **2.72E-02** |
| ENSG00000152056.20 | **AP1S3** | 130340 | protein_coding | **254.878** | 0.382 | 0.115 | **1.303** | 1.34E-02 | **2.72E-02** |
| ENSG00000279254.1 | **ENSG00000279254** | | TEC | **8.173** | 1.496 | 0.450 | **2.820** | 1.34E-02 | **2.72E-02** |
| ENSG00000108774.17 | **RAB5C** | 5878 | protein_coding | **897.376** | 0.360 | 0.108 | **1.283** | 1.34E-02 | **2.72E-02** |
| ENSG00000250850.4 | **ENSG00000250850** | | lncRNA | **31.944** | -1.023 | -0.308 | **0.492** | 1.34E-02 | **2.72E-02** |
| ENSG00000104549.14 | **SQLE** | 6713 | protein_coding | **1003.170** | -0.537 | -0.162 | **0.689** | 1.35E-02 | **2.73E-02** |
| ENSG00000108528.16 | **SLC25A11** | 8402 | protein_coding | **969.307** | -0.300 | -0.090 | **0.812** | 1.35E-02 | **2.74E-02** |
| ENSG00000144026.16 | **ZNF514** | 84874 | protein_coding | **569.251** | 0.404 | 0.122 | **1.323** | 1.35E-02 | **2.74E-02** |
| ENSG00000289469.3 | **ENSG0000(** | 105374101 | lncRNA | **16.654** | 0.904 | 0.272 | **1.872** | 1.35E-02 | **2.74E-02** |
| ENSG00000138030.16 | **KHK** | 3795 | protein_coding | **89.771** | 0.463 | 0.139 | **1.378** | 1.36E-02 | **2.75E-02** |
| ENSG00000129968.19 | **ABHD17A** | 81926 | protein_coding | **1894.058** | 0.324 | 0.098 | **1.252** | 1.36E-02 | **2.75E-02** |
| ENSG00000215417.15 | **MIR17HG** | 407975 | lncRNA | **74.322** | -0.564 | -0.170 | **0.677** | 1.36E-02 | **2.75E-02** |
| ENSG00000286732.2 | **ENSG00000286732** | | lncRNA | **3.746** | 2.369 | 0.713 | **5.166** | 1.36E-02 | **2.76E-02** |
| ENSG00000233435.2 | **AGGF1P2** | | processed_pseud( | **14.180** | 1.057 | 0.318 | **2.080** | 1.36E-02 | **2.76E-02** |
| ENSG00000185736.17 | **ADARB2** | 105 | protein_coding | **37.915** | 0.891 | 0.268 | **1.854** | 1.36E-02 | **2.76E-02** |
| ENSG00000122512.19 | **PMS2** | 5395 | protein_coding | **595.341** | -0.360 | -0.108 | **0.779** | 1.36E-02 | **2.76E-02** |
| ENSG00000010626.17 | **LRRC23** | 10233 | protein_coding | **198.423** | 0.393 | 0.118 | **1.313** | 1.36E-02 | **2.76E-02** |
| ENSG00000105755.10 | **ETHE1** | 23474 | protein_coding | **543.291** | -0.313 | -0.094 | **0.805** | 1.37E-02 | **2.76E-02** |
| ENSG00000102225.19 | **CDK16** | 5127 | protein_coding | **1475.441** | -0.358 | -0.108 | **0.780** | 1.37E-02 | **2.76E-02** |
| ENSG00000273308.2 | **ENSG00000273308** | | lncRNA | **13.250** | 1.047 | 0.315 | **2.066** | 1.37E-02 | **2.77E-02** |
| ENSG00000153250.23 | **RBMS1** | 5937 | protein_coding | **56.053** | 0.725 | 0.218 | **1.652** | 1.37E-02 | **2.77E-02** |
| ENSG00000262879.8 | **ENSG00000262879** | | lncRNA | **304.139** | -0.316 | -0.095 | **0.803** | 1.37E-02 | **2.77E-02** |
| ENSG00000172878.16 | **METAP1D** | 254042 | protein_coding | **204.130** | -0.364 | -0.110 | **0.777** | 1.37E-02 | **2.77E-02** |
| ENSG00000111245.20 | **MYL2** | 4633 | protein_coding | **32.508** | 0.741 | 0.223 | **1.672** | 1.37E-02 | **2.77E-02** |
| ENSG00000064309.18 | **CDON** | 50937 | protein_coding | **22.085** | 1.239 | 0.373 | **2.360** | 1.37E-02 | **2.77E-02** |
| ENSG00000224152.3 | **BAZ2B-AS1** | | lncRNA | **21.148** | -0.972 | -0.292 | **0.510** | 1.37E-02 | **2.77E-02** |
| ENSG00000244480.1 | **ENSG00000244480** | | transcribed_proce | **55.300** | 0.702 | 0.211 | **1.627** | 1.37E-02 | **2.78E-02** |
| ENSG00000153363.16 | **LINC00467** | | lncRNA | **282.300** | -0.431 | -0.130 | **0.742** | 1.38E-02 | **2.78E-02** |
| ENSG00000287808.2 | **DHX9-AS1** | | lncRNA | **22.210** | 0.772 | 0.232 | **1.708** | 1.38E-02 | **2.78E-02** |
| ENSG00000106617.18 | **PRKAG2** | 51422 | protein_coding | **150.981** | 0.405 | 0.122 | **1.324** | 1.38E-02 | **2.79E-02** |
| ENSG00000128011.7 | **LRFN1** | 57622 | protein_coding | **91.534** | 0.466 | 0.140 | **1.381** | 1.38E-02 | **2.79E-02** |

| | | | | | | | | | |
|---|---|---|---|---|---|---|---|---|---|
| ENSG00000255966.2 | **GAPDH-DT** | | lncRNA | **73.537** | -0.661 | -0.199 | **0.633** | 1.38E-02 | **2.79E-02** |
| ENSG00000259891.1 | **ENSG00000259891** | | lncRNA | **37.276** | 0.698 | 0.210 | **1.622** | 1.38E-02 | **2.79E-02** |
| ENSG00000263766.7 | **KPNB1-DT** | | lncRNA | **36.812** | 0.598 | 0.180 | **1.514** | 1.38E-02 | **2.79E-02** |
| ENSG00000136840.21 | **ST6GALNA** | 27090 | protein_coding | **745.213** | -0.343 | -0.103 | **0.788** | 1.38E-02 | **2.80E-02** |
| ENSG00000188542.11 | **DUSP28** | 285193 | protein_coding | **132.534** | -0.411 | -0.124 | **0.752** | 1.39E-02 | **2.80E-02** |
| ENSG00000071054.19 | **MAP4K4** | 9448 | protein_coding | **97.341** | 0.551 | 0.166 | **1.465** | 1.39E-02 | **2.80E-02** |
| ENSG00000198185.14 | **ZNF334** | 55713 | protein_coding | **204.193** | 0.513 | 0.154 | **1.427** | 1.39E-02 | **2.80E-02** |
| ENSG00000144214.12 | **LYG1** | 129530 | protein_coding | **86.877** | 0.545 | 0.164 | **1.459** | 1.39E-02 | **2.81E-02** |
| ENSG00000162783.11 | **IER5** | 51278 | protein_coding | **3292.988** | 0.279 | 0.084 | **1.213** | 1.39E-02 | **2.81E-02** |
| ENSG00000212719.16 | **LINC02693** | 339263 | lncRNA | **7.194** | 1.431 | 0.431 | **2.696** | 1.39E-02 | **2.81E-02** |
| ENSG00000005339.16 | **CREBBP** | 1387 | protein_coding | **715.278** | -0.379 | -0.114 | **0.769** | 1.39E-02 | **2.81E-02** |
| ENSG00000154727.13 | **GABPA** | 2551 | protein_coding | **234.159** | -0.425 | -0.128 | **0.745** | 1.39E-02 | **2.81E-02** |
| ENSG00000291178.2 | **ENSG00000291178** | | lncRNA | **767.821** | -0.311 | -0.094 | **0.806** | 1.39E-02 | **2.81E-02** |
| ENSG00000289405.3 | **ENSG00000289405** | | lncRNA | **84.944** | -0.614 | -0.185 | **0.653** | 1.40E-02 | **2.82E-02** |
| ENSG00000305300.1 | **ENSG00000305300** | | lncRNA | **5.875** | 1.661 | 0.500 | **3.162** | 1.40E-02 | **2.82E-02** |
| ENSG00000224376.4 | **ENSG00000224376** | | lncRNA | **140.032** | -0.558 | -0.168 | **0.679** | 1.40E-02 | **2.82E-02** |
| ENSG00000274615.1 | **NPEPPSP1** | 440434 | transcribed_unpro | **44.345** | -2.625 | -0.790 | **0.162** | 1.40E-02 | **2.82E-02** |
| ENSG00000166908.20 | **PIP4K2C** | 79837 | protein_coding | **370.404** | 0.309 | 0.093 | **1.239** | 1.40E-02 | **2.82E-02** |
| ENSG00000204267.21 | **TAP2** | 6891 | protein_coding | **2890.659** | 0.291 | 0.088 | **1.224** | 1.40E-02 | **2.83E-02** |
| ENSG00000309043.1 | **ENSG00000309043** | | lncRNA | **5.627** | 2.033 | 0.612 | **4.094** | 1.40E-02 | **2.83E-02** |
| ENSG00000196422.13 | **PPP1R26** | 9858 | protein_coding | **409.904** | 0.549 | 0.165 | **1.463** | 1.40E-02 | **2.83E-02** |
| ENSG00000100320.26 | **RBFOX2** | 23543 | protein_coding | **19.748** | 1.029 | 0.310 | **2.041** | 1.40E-02 | **2.83E-02** |
| ENSG00000132294.17 | **EFR3A** | 23167 | protein_coding | **550.859** | 0.414 | 0.125 | **1.333** | 1.40E-02 | **2.83E-02** |
| ENSG00000130294.20 | **KIF1A** | 547 | protein_coding | **36.699** | 1.084 | 0.326 | **2.120** | 1.40E-02 | **2.83E-02** |
| ENSG00000169436.20 | **COL22A1** | 169044 | protein_coding | **11.842** | 1.949 | 0.587 | **3.862** | 1.40E-02 | **2.83E-02** |
| ENSG00000277511.3 | **VILMIR** | | lncRNA | **47.926** | -0.641 | -0.193 | **0.641** | 1.40E-02 | **2.83E-02** |
| ENSG00000114978.20 | **MOB1A** | 55233 | protein_coding | **2765.097** | 0.296 | 0.089 | **1.227** | 1.41E-02 | **2.83E-02** |
| ENSG00000087302.11 | **RTRAF** | 51637 | protein_coding | **2932.877** | -0.328 | -0.099 | **0.797** | 1.41E-02 | **2.83E-02** |
| ENSG00000235590.10 | **GNAS-AS1** | 149775 | lncRNA | **10.062** | -0.996 | -0.300 | **0.501** | 1.41E-02 | **2.83E-02** |
| ENSG00000158716.11 | **DUSP23** | 54935 | protein_coding | **170.254** | 0.390 | 0.117 | **1.310** | 1.41E-02 | **2.83E-02** |
| ENSG00000288044.4 | **ENSG00000288044** | | lncRNA | **119.200** | -0.614 | -0.185 | **0.653** | 1.41E-02 | **2.84E-02** |

| | | | | | | | | | |
|---|---|---|---|---|---|---|---|---|---|
| ENSG00000299860.1 | **ENSG00000299860** | | lncRNA | **15.865** | 1.077 | 0.324 | **2.109** | 1.41E-02 | **2.84E-02** |
| ENSG00000127364.4 | **TAS2R4** | 50832 | protein_coding | **20.221** | 0.806 | 0.243 | **1.749** | 1.41E-02 | **2.84E-02** |
| ENSG00000086200.19 | **IPO11** | 51194 | protein_coding | **173.072** | -0.372 | -0.112 | **0.772** | 1.41E-02 | **2.84E-02** |
| ENSG00000084652.18 | **TXLNA** | 200081 | protein_coding | **1509.330** | -0.306 | -0.092 | **0.809** | 1.41E-02 | **2.84E-02** |
| ENSG00000169087.13 | **HSPBAP1** | 79663 | protein_coding | **252.391** | 0.366 | 0.110 | **1.288** | 1.41E-02 | **2.84E-02** |
| ENSG00000175793.12 | **SFN** | 2810 | protein_coding | **44.071** | -0.747 | -0.225 | **0.596** | 1.41E-02 | **2.84E-02** |
| ENSG00000184076.15 | **UQCR10** | 29796 | protein_coding | **1587.435** | -0.342 | -0.103 | **0.789** | 1.42E-02 | **2.85E-02** |
| ENSG00000141665.15 | **FBXO15** | 201456 | protein_coding | **9.744** | 1.316 | 0.396 | **2.490** | 1.42E-02 | **2.85E-02** |
| ENSG00000198727.2 | **MT-CYB** | 4519 | protein_coding | **103692.766** | 0.442 | 0.133 | **1.359** | 1.42E-02 | **2.85E-02** |
| ENSG00000197021.11 | **EOLA2** | 541578 | protein_coding | **600.776** | 0.376 | 0.113 | **1.298** | 1.42E-02 | **2.85E-02** |
| ENSG00000198843.15 | **SELENOT** | 51714 | protein_coding | **930.049** | -0.460 | -0.139 | **0.727** | 1.42E-02 | **2.85E-02** |
| ENSG00000293476.1 | **ENSG00000293476** | | lncRNA | **170.989** | -0.368 | -0.111 | **0.775** | 1.42E-02 | **2.85E-02** |
| ENSG00000244219.9 | **TMEM225I** | 10028918 | protein_coding | **25.800** | 0.744 | 0.224 | **1.675** | 1.42E-02 | **2.86E-02** |
| ENSG00000139890.12 | **REM2** | 161253 | protein_coding | **56.096** | 0.517 | 0.156 | **1.431** | 1.42E-02 | **2.86E-02** |
| ENSG00000116675.16 | **DNAJC6** | 9829 | protein_coding | **26.652** | 0.847 | 0.255 | **1.799** | 1.43E-02 | **2.87E-02** |
| ENSG00000158156.10 | **XKR8** | 55113 | protein_coding | **107.553** | 0.392 | 0.118 | **1.313** | 1.43E-02 | **2.87E-02** |
| ENSG00000153140.11 | **CETN3** | 1070 | protein_coding | **350.644** | -0.344 | -0.103 | **0.788** | 1.43E-02 | **2.87E-02** |
| ENSG00000269918.1 | **ENSG00000269918** | | lncRNA | **240.196** | 0.305 | 0.092 | **1.235** | 1.43E-02 | **2.87E-02** |
| ENSG00000013561.20 | **RNF14** | 9604 | protein_coding | **247.646** | -0.322 | -0.097 | **0.800** | 1.43E-02 | **2.87E-02** |
| ENSG00000137124.10 | **ALDH1B1** | 219 | protein_coding | **348.977** | -0.446 | -0.134 | **0.734** | 1.43E-02 | **2.87E-02** |
| ENSG00000233290.5 | **ENSG00000233290** | | lncRNA | **31.699** | 0.798 | 0.240 | **1.739** | 1.43E-02 | **2.88E-02** |
| ENSG00000171115.6 | **GIMAP8** | 155038 | protein_coding | **179.636** | 0.392 | 0.118 | **1.313** | 1.44E-02 | **2.88E-02** |
| ENSG00000203546.7 | **ENSG00000203546** | | protein_coding | **7.811** | -1.476 | -0.444 | **0.359** | 1.44E-02 | **2.88E-02** |
| ENSG00000119705.12 | **SLIRP** | 81892 | protein_coding | **854.574** | -0.385 | -0.116 | **0.766** | 1.44E-02 | **2.89E-02** |
| ENSG00000178074.7 | **C2orf69** | 205327 | protein_coding | **111.846** | -0.444 | -0.134 | **0.735** | 1.44E-02 | **2.89E-02** |
| ENSG00000147509.15 | **RGS20** | 8601 | protein_coding | **320.232** | -0.396 | -0.119 | **0.760** | 1.44E-02 | **2.90E-02** |
| ENSG00000232613.8 | **LINC02576** | 10272423 | lncRNA | **39.660** | -0.988 | -0.298 | **0.504** | 1.44E-02 | **2.90E-02** |
| ENSG00000275297.2 | **ENSG00000275297** | | lncRNA | **21.448** | -1.192 | -0.359 | **0.438** | 1.44E-02 | **2.90E-02** |
| ENSG00000102003.14 | **SYP** | 6855 | protein_coding | **45.944** | 0.541 | 0.163 | **1.455** | 1.44E-02 | **2.90E-02** |
| ENSG00000224531.8 | **SMIM13** | 221710 | protein_coding | **107.430** | -0.453 | -0.136 | **0.731** | 1.45E-02 | **2.90E-02** |
| ENSG00000148120.21 | **AOPEP** | 84909 | protein_coding | **114.658** | 0.353 | 0.106 | **1.278** | 1.45E-02 | **2.90E-02** |

| | | | | | | | | | |
|---|---|---|---|---|---|---|---|---|---|
| ENSG00000119862.15 | **LGALSL** | 29094 | protein_coding | **73.570** | 0.602 | 0.181 | **1.518** | 1.45E-02 | **2.90E-02** |
| ENSG00000275393.2 | **ENSG00000275393** | | lncRNA | **17.638** | 1.099 | 0.331 | **2.141** | 1.45E-02 | **2.90E-02** |
| ENSG00000198178.13 | **CLEC4C** | 170482 | protein_coding | **52.372** | -0.943 | -0.284 | **0.520** | 1.45E-02 | **2.91E-02** |
| ENSG00000234362.7 | **LINC01914** | | lncRNA | **10.535** | 1.476 | 0.444 | **2.782** | 1.45E-02 | **2.91E-02** |
| ENSG00000143553.13 | **SNAPIN** | 23557 | protein_coding | **531.679** | 0.292 | 0.088 | **1.225** | 1.45E-02 | **2.91E-02** |
| ENSG00000273828.3 | **ENSG0000(** | 12490491 | lncRNA | **6.017** | 1.834 | 0.552 | **3.565** | 1.45E-02 | **2.91E-02** |
| ENSG00000272037.1 | **ENSG00000272037** | | lncRNA | **8.595** | 1.180 | 0.355 | **2.265** | 1.45E-02 | **2.91E-02** |
| ENSG00000187486.9 | **KCNJ11** | 3767 | protein_coding | **20.127** | 0.952 | 0.287 | **1.935** | 1.45E-02 | **2.91E-02** |
| ENSG00000225914.6 | **TSBP1-AS1** | 414764 | lncRNA | **18.711** | 0.942 | 0.283 | **1.921** | 1.46E-02 | **2.92E-02** |
| ENSG00000235655.3 | **H3P6** | | processed_pseud | **667.371** | -0.495 | -0.149 | **0.710** | 1.46E-02 | **2.92E-02** |
| ENSG00000302434.1 | **ENSG00000302434** | | lncRNA | **7.691** | 1.270 | 0.382 | **2.411** | 1.46E-02 | **2.92E-02** |
| ENSG00000251034.2 | **ENSG00000251034** | | lncRNA | **5.648** | 2.081 | 0.626 | **4.231** | 1.46E-02 | **2.92E-02** |
| ENSG00000204520.16 | **MICA** | 10050743 | protein_coding | **335.391** | 0.272 | 0.082 | **1.207** | 1.46E-02 | **2.92E-02** |
| ENSG00000104129.12 | **DNAJC17** | 55192 | protein_coding | **365.506** | -0.381 | -0.115 | **0.768** | 1.46E-02 | **2.92E-02** |
| ENSG00000175390.17 | **EIF3F** | 8665 | protein_coding | **4412.033** | -0.268 | -0.081 | **0.831** | 1.46E-02 | **2.93E-02** |
| ENSG00000167074.17 | **TEF** | 7008 | protein_coding | **119.528** | 0.357 | 0.107 | **1.280** | 1.46E-02 | **2.93E-02** |
| ENSG00000060749.18 | **QSER1** | 79832 | protein_coding | **204.912** | -0.497 | -0.150 | **0.709** | 1.47E-02 | **2.94E-02** |
| ENSG00000137767.16 | **SQOR** | 58472 | protein_coding | **988.215** | 0.357 | 0.107 | **1.281** | 1.47E-02 | **2.94E-02** |
| ENSG00000188185.14 | **LINC00265** | | lncRNA | **359.708** | 0.307 | 0.092 | **1.237** | 1.47E-02 | **2.94E-02** |
| ENSG00000266910.2 | **ENSG00000266910** | | processed_pseud | **21.313** | 0.789 | 0.238 | **1.728** | 1.47E-02 | **2.95E-02** |
| ENSG00000295516.1 | **ENSG00000295516** | | lncRNA | **20.354** | 0.728 | 0.219 | **1.656** | 1.48E-02 | **2.96E-02** |
| ENSG00000267980.1 | **SH3GL1-AS1** | | lncRNA | **10.938** | 1.163 | 0.350 | **2.240** | 1.48E-02 | **2.96E-02** |
| ENSG00000263934.5 | **SNORD3A** | | snoRNA | **382.001** | 0.571 | 0.172 | **1.486** | 1.48E-02 | **2.96E-02** |
| ENSG00000147202.20 | **DIAPH2** | 1730 | protein_coding | **224.855** | -0.352 | -0.106 | **0.783** | 1.48E-02 | **2.96E-02** |
| ENSG00000134871.22 | **COL4A2** | 1284 | protein_coding | **34.984** | 1.021 | 0.307 | **2.029** | 1.48E-02 | **2.97E-02** |
| ENSG00000196912.14 | **ANKRD36E** | 57730 | protein_coding | **306.155** | -0.429 | -0.129 | **0.743** | 1.48E-02 | **2.97E-02** |
| ENSG00000196653.14 | **ZNF502** | 91392 | protein_coding | **27.140** | 0.696 | 0.210 | **1.620** | 1.48E-02 | **2.97E-02** |
| ENSG00000153487.15 | **ING1** | 3621 | protein_coding | **385.867** | -0.467 | -0.141 | **0.723** | 1.49E-02 | **2.97E-02** |
| ENSG00000129007.18 | **CALML4** | 91860 | protein_coding | **199.992** | -0.462 | -0.139 | **0.726** | 1.49E-02 | **2.98E-02** |
| ENSG00000284626.1 | **ENSG00000284626** | | protein_coding | **17.071** | 1.220 | 0.367 | **2.329** | 1.49E-02 | **2.98E-02** |
| ENSG00000233461.6 | **CYDAER** | 12252678 | lncRNA | **85.203** | -0.536 | -0.161 | **0.690** | 1.49E-02 | **2.98E-02** |

| | | | | | | | | | |
|---|---|---|---|---|---|---|---|---|---|
| ENSG00000163382.15 | **NAXE** | 128240 | protein_coding | **863.895** | -0.331 | -0.100 | **0.795** | 1.49E-02 | **2.98E-02** |
| ENSG00000147316.17 | **MCPH1** | 79648 | protein_coding | **492.389** | 0.410 | 0.123 | **1.329** | 1.49E-02 | **2.98E-02** |
| ENSG00000197933.15 | **ZNF823** | 55552 | protein_coding | **77.664** | -0.461 | -0.139 | **0.726** | 1.49E-02 | **2.98E-02** |
| ENSG00000283765.1 | **ENSG00000283765** | | protein_coding | **6.175** | -4.827 | -1.453 | **0.035** | 1.49E-02 | **2.98E-02** |
| ENSG00000166927.15 | **MS4A7** | 58475 | protein_coding | **111.864** | 0.616 | 0.185 | **1.533** | 1.49E-02 | **2.99E-02** |
| ENSG00000170627.13 | **GTSF1** | 121355 | protein_coding | **99.550** | 0.532 | 0.160 | **1.446** | 1.50E-02 | **2.99E-02** |
| ENSG00000145416.15 | **MARCHF1** | 55016 | protein_coding | **12.291** | -1.368 | -0.412 | **0.387** | 1.50E-02 | **2.99E-02** |
| ENSG00000065802.14 | **ASB1** | 51665 | protein_coding | **423.939** | -0.305 | -0.092 | **0.810** | 1.50E-02 | **2.99E-02** |
| ENSG00000103197.22 | **TSC2** | 7249 | protein_coding | **1298.265** | 0.312 | 0.094 | **1.241** | 1.50E-02 | **2.99E-02** |
| ENSG00000099822.3 | **HCN2** | 610 | protein_coding | **38.833** | 0.713 | 0.215 | **1.639** | 1.50E-02 | **2.99E-02** |
| ENSG00000242474.4 | **LINC03015** | 102723672 | lncRNA | **52.210** | 0.553 | 0.166 | **1.467** | 1.50E-02 | **2.99E-02** |
| ENSG00000140365.18 | **COMMD4** | 54939 | protein_coding | **959.383** | -0.308 | -0.093 | **0.808** | 1.50E-02 | **2.99E-02** |
| ENSG00000184719.14 | **RNLS** | 55328 | protein_coding | **46.501** | -0.598 | -0.180 | **0.661** | 1.50E-02 | **3.00E-02** |
| ENSG00000247287.4 | **PRKCH-AS1** | | lncRNA | **20.360** | -0.766 | -0.231 | **0.588** | 1.50E-02 | **3.00E-02** |
| ENSG00000179195.18 | **ZNF664** | 144348 | protein_coding | **748.891** | 0.366 | 0.110 | **1.289** | 1.50E-02 | **3.00E-02** |
| ENSG00000136720.8 | **HS6ST1** | 9394 | protein_coding | **222.925** | 0.405 | 0.122 | **1.324** | 1.50E-02 | **3.00E-02** |
| ENSG00000100802.17 | **C14orf93** | 60686 | protein_coding | **297.044** | 0.299 | 0.090 | **1.230** | 1.51E-02 | **3.01E-02** |
| ENSG00000132481.9 | **TRIM47** | 91107 | protein_coding | **72.856** | 0.578 | 0.174 | **1.493** | 1.51E-02 | **3.01E-02** |
| ENSG00000143190.25 | **POU2F1** | 5451 | protein_coding | **542.966** | -0.339 | -0.102 | **0.791** | 1.51E-02 | **3.02E-02** |
| ENSG00000128908.21 | **INO80** | 54617 | protein_coding | **474.717** | -0.363 | -0.109 | **0.778** | 1.51E-02 | **3.02E-02** |
| ENSG00000103811.20 | **CTSH** | 1512 | protein_coding | **8854.481** | -0.338 | -0.102 | **0.791** | 1.52E-02 | **3.02E-02** |
| ENSG00000162438.13 | **CTRC** | 11330 | protein_coding | **5.667** | 1.879 | 0.566 | **3.679** | 1.52E-02 | **3.02E-02** |
| ENSG00000115155.19 | **OTOF** | 9381 | protein_coding | **13.119** | 1.436 | 0.432 | **2.705** | 1.52E-02 | **3.02E-02** |
| ENSG00000108523.18 | **RNF167** | 26001 | protein_coding | **2478.006** | 0.286 | 0.086 | **1.219** | 1.52E-02 | **3.03E-02** |
| ENSG00000137831.17 | **UACA** | 55075 | protein_coding | **96.113** | 0.562 | 0.169 | **1.477** | 1.52E-02 | **3.03E-02** |
| ENSG00000182512.7 | **GLRX5** | 51218 | protein_coding | **571.806** | -0.312 | -0.094 | **0.805** | 1.52E-02 | **3.03E-02** |
| ENSG00000224349.3 | **FAM135A-** | 10537784 | lncRNA | **8.619** | 1.137 | 0.342 | **2.200** | 1.52E-02 | **3.03E-02** |
| ENSG00000291066.2 | **ENSG00000291066** | | lncRNA | **400.717** | 0.337 | 0.102 | **1.264** | 1.52E-02 | **3.03E-02** |
| ENSG00000104055.18 | **TGM5** | 9333 | protein_coding | **116.154** | 0.552 | 0.166 | **1.466** | 1.52E-02 | **3.03E-02** |
| ENSG00000050030.17 | **NEXMIF** | 340533 | protein_coding | **39.912** | -0.716 | -0.215 | **0.609** | 1.52E-02 | **3.04E-02** |
| ENSG00000289843.2 | **BMS1P14** | 10192995 | lncRNA | **63.835** | -0.554 | -0.167 | **0.681** | 1.52E-02 | **3.04E-02** |

| | | | | | | | | | |
|---|---|---|---|---|---|---|---|---|---|
| ENSG00000310068.1 | **ENSG00000310068** | | lncRNA | **7.424** | 1.279 | 0.385 | **2.426** | 1.53E-02 | **3.04E-02** |
| ENSG00000289038.3 | **ENSG00000289038** | | lncRNA | **7.822** | 1.246 | 0.375 | **2.372** | 1.53E-02 | **3.04E-02** |
| ENSG00000048707.16 | **VPS13D** | 55187 | protein_coding | **464.659** | 0.329 | 0.099 | **1.257** | 1.53E-02 | **3.04E-02** |
| ENSG00000197905.11 | **TEAD4** | 7004 | protein_coding | **127.662** | -0.353 | -0.106 | **0.783** | 1.53E-02 | **3.05E-02** |
| ENSG00000106804.12 | **C5** | 727 | protein_coding | **82.169** | 0.503 | 0.151 | **1.417** | 1.54E-02 | **3.06E-02** |
| ENSG00000141543.14 | **EIF4A3** | 9775 | protein_coding | **1641.679** | 0.311 | 0.094 | **1.241** | 1.54E-02 | **3.06E-02** |
| ENSG00000137815.17 | **RTF1** | 23168 | protein_coding | **870.459** | 0.355 | 0.107 | **1.279** | 1.54E-02 | **3.06E-02** |
| ENSG00000172262.14 | **ZNF131** | 7690 | protein_coding | **292.250** | -0.383 | -0.115 | **0.767** | 1.54E-02 | **3.07E-02** |
| ENSG00000135426.18 | **TESPA1** | 9840 | protein_coding | **874.231** | -0.324 | -0.097 | **0.799** | 1.54E-02 | **3.07E-02** |
| ENSG00000185515.17 | **BRCC3** | 79184 | protein_coding | **376.142** | -0.298 | -0.090 | **0.813** | 1.54E-02 | **3.07E-02** |
| ENSG00000241728.5 | **ENSG00000241728** | | lncRNA | **19.531** | 0.817 | 0.246 | **1.762** | 1.55E-02 | **3.08E-02** |
| ENSG00000102870.8 | **ZNF629** | 23361 | protein_coding | **188.759** | -0.300 | -0.090 | **0.812** | 1.55E-02 | **3.09E-02** |
| ENSG00000236397.3 | **DDX11L2** | | transcribed_unprc | **25.761** | 4.620 | 1.391 | **24.586** | 1.55E-02 | **3.09E-02** |
| ENSG00000226659.3 | **TSPAN14-AS1** | | lncRNA | **17.820** | -0.945 | -0.284 | **0.520** | 1.55E-02 | **3.09E-02** |
| ENSG00000140905.13 | **GCSH** | 2653 | protein_coding | **199.540** | -0.584 | -0.176 | **0.667** | 1.55E-02 | **3.09E-02** |
| ENSG00000290847.2 | **PRAMENP** | 649179 | lncRNA | **25.618** | 0.869 | 0.262 | **1.826** | 1.56E-02 | **3.09E-02** |
| ENSG00000276533.1 | **ENSG00000276533** | | lncRNA | **8.733** | -1.156 | -0.348 | **0.449** | 1.56E-02 | **3.10E-02** |
| ENSG00000185000.14 | **DGAT1** | 8694 | protein_coding | **572.707** | 0.335 | 0.101 | **1.262** | 1.56E-02 | **3.10E-02** |
| ENSG00000258102.5 | **MAP1LC3B** | 643246 | protein_coding | **32.578** | 0.666 | 0.201 | **1.587** | 1.56E-02 | **3.10E-02** |
| ENSG00000150275.21 | **PCDH15** | 65217 | protein_coding | **4.985** | 1.830 | 0.551 | **3.555** | 1.56E-02 | **3.11E-02** |
| ENSG00000309294.1 | **ENSG00000309294** | | lncRNA | **12.985** | 0.950 | 0.286 | **1.931** | 1.56E-02 | **3.11E-02** |
| ENSG00000177369.12 | **FLJ40194** | 124871 | lncRNA | **28.737** | 0.733 | 0.221 | **1.662** | 1.58E-02 | **3.13E-02** |
| ENSG00000291063.2 | **ENSG0000(** | 10798497( | lncRNA | **48.199** | -0.568 | -0.171 | **0.675** | 1.58E-02 | **3.14E-02** |
| ENSG00000061273.20 | **HDAC7** | 51564 | protein_coding | **569.213** | -0.271 | -0.081 | **0.829** | 1.58E-02 | **3.14E-02** |
| ENSG00000276728.3 | **ENSG00000276728** | | lncRNA | **52.707** | 0.516 | 0.155 | **1.430** | 1.58E-02 | **3.14E-02** |
| ENSG00000198205.7 | **ZXDA** | 7789 | protein_coding | **19.396** | -0.930 | -0.280 | **0.525** | 1.58E-02 | **3.14E-02** |
| ENSG00000076604.17 | **TRAF4** | 9618 | protein_coding | **2163.152** | -0.279 | -0.084 | **0.824** | 1.58E-02 | **3.15E-02** |
| ENSG00000109501.17 | **WFS1** | 7466 | protein_coding | **131.676** | 0.556 | 0.167 | **1.470** | 1.59E-02 | **3.15E-02** |
| ENSG00000158062.23 | **UBXN11** | 91544 | protein_coding | **569.472** | -0.340 | -0.102 | **0.790** | 1.59E-02 | **3.16E-02** |
| ENSG00000130803.17 | **ZNF317** | 57693 | protein_coding | **740.742** | -0.368 | -0.111 | **0.775** | 1.59E-02 | **3.16E-02** |
| ENSG00000131558.18 | **EXOC4** | 60412 | protein_coding | **587.543** | -0.277 | -0.084 | **0.825** | 1.60E-02 | **3.17E-02** |

| | | | | | | | | | |
|---|---|---|---|---|---|---|---|---|---|
| ENSG00000115762.19 | **PLEKHB2** | 55041 | protein_coding | **706.843** | 0.343 | 0.103 | **1.268** | 1.60E-02 | **3.17E-02** |
| ENSG00000290126.2 | **ENSG00000290126** | | lncRNA | **41.440** | 0.591 | 0.178 | **1.506** | 1.60E-02 | **3.17E-02** |
| ENSG00000183625.18 | **CCR3** | 1232 | protein_coding | **13.502** | 1.230 | 0.370 | **2.346** | 1.60E-02 | **3.17E-02** |
| ENSG00000102984.17 | **ZNF821** | 55565 | protein_coding | **399.546** | 0.435 | 0.131 | **1.352** | 1.60E-02 | **3.17E-02** |
| ENSG00000181894.17 | **ZNF329** | 79673 | protein_coding | **92.998** | 0.476 | 0.143 | **1.391** | 1.60E-02 | **3.18E-02** |
| ENSG00000141570.12 | **CBX8** | 57332 | protein_coding | **181.265** | -0.352 | -0.106 | **0.784** | 1.60E-02 | **3.18E-02** |
| ENSG00000139515.7 | **PDX1** | 3651 | protein_coding | **65.201** | 0.824 | 0.248 | **1.771** | 1.60E-02 | **3.18E-02** |
| ENSG00000104413.20 | **ESRP1** | 54845 | protein_coding | **4.119** | 1.982 | 0.597 | **3.951** | 1.60E-02 | **3.18E-02** |
| ENSG00000108433.18 | **GOSR2** | 9570 | protein_coding | **792.277** | -0.277 | -0.083 | **0.826** | 1.61E-02 | **3.19E-02** |
| ENSG00000116830.14 | **TTF2** | 8458 | protein_coding | **901.522** | -0.406 | -0.122 | **0.755** | 1.61E-02 | **3.19E-02** |
| ENSG00000189420.11 | **ZFP92** | 139735 | protein_coding | **65.729** | 0.503 | 0.152 | **1.417** | 1.61E-02 | **3.19E-02** |
| ENSG00000014123.12 | **UFL1** | 23376 | protein_coding | **373.617** | -0.482 | -0.145 | **0.716** | 1.61E-02 | **3.19E-02** |
| ENSG00000206560.13 | **ANKRD28** | 23243 | protein_coding | **772.814** | -0.526 | -0.158 | **0.695** | 1.62E-02 | **3.20E-02** |
| ENSG00000167065.16 | **DUSP18** | 150290 | protein_coding | **40.551** | 0.733 | 0.221 | **1.662** | 1.62E-02 | **3.20E-02** |
| ENSG00000258701.1 | **VESTAR** | 196872 | lncRNA | **20.142** | 0.794 | 0.239 | **1.734** | 1.62E-02 | **3.21E-02** |
| ENSG00000180902.20 | **D2HGDH** | 728294 | protein_coding | **1154.438** | 0.281 | 0.085 | **1.215** | 1.62E-02 | **3.21E-02** |
| ENSG00000116704.10 | **SLC35D1** | 23169 | protein_coding | **147.624** | -0.474 | -0.143 | **0.720** | 1.62E-02 | **3.21E-02** |
| ENSG00000165917.12 | **RAPSN** | 5913 | protein_coding | **12.224** | -1.046 | -0.315 | **0.484** | 1.62E-02 | **3.21E-02** |
| ENSG00000226085.3 | **UQCRFS1P1** | | processed_pseud | **27.285** | -0.724 | -0.218 | **0.606** | 1.62E-02 | **3.22E-02** |
| ENSG00000167994.15 | **RAB3IL1** | 5866 | protein_coding | **5.356** | 1.895 | 0.570 | **3.719** | 1.63E-02 | **3.22E-02** |
| ENSG00000273210.1 | **ENSG00000273210** | | lncRNA | **20.374** | 0.828 | 0.249 | **1.776** | 1.63E-02 | **3.22E-02** |
| ENSG00000140279.14 | **DUOX2** | 50506 | protein_coding | **13.222** | 1.425 | 0.429 | **2.685** | 1.63E-02 | **3.22E-02** |
| ENSG00000293507.2 | **FBXL9P** | 26231 | lncRNA | **57.785** | 0.580 | 0.175 | **1.495** | 1.63E-02 | **3.22E-02** |
| ENSG00000256948.2 | **IQSEC3-AS2** | | lncRNA | **11.386** | 1.487 | 0.448 | **2.803** | 1.63E-02 | **3.22E-02** |
| ENSG00000105717.16 | **PBX4** | 80714 | protein_coding | **168.343** | -0.574 | -0.173 | **0.672** | 1.64E-02 | **3.24E-02** |
| ENSG00000148429.17 | **USP6NL** | 9712 | protein_coding | **600.866** | 0.369 | 0.111 | **1.292** | 1.64E-02 | **3.24E-02** |
| ENSG00000273443.2 | **ENSG00000273443** | | lncRNA | **87.381** | -0.504 | -0.152 | **0.705** | 1.64E-02 | **3.24E-02** |
| ENSG00000272662.1 | **TIMMDC1-DT** | | lncRNA | **5.588** | 1.545 | 0.465 | **2.918** | 1.64E-02 | **3.24E-02** |
| ENSG00000285744.2 | **MITA1** | | lncRNA | **68.679** | 0.492 | 0.148 | **1.406** | 1.64E-02 | **3.25E-02** |
| ENSG00000184307.19 | **ZDHHC23** | 254887 | protein_coding | **92.844** | 0.608 | 0.183 | **1.524** | 1.64E-02 | **3.25E-02** |
| ENSG00000250626.4 | **ENSG0000** | 10193002 | lncRNA | **9.494** | -1.776 | -0.535 | **0.292** | 1.64E-02 | **3.25E-02** |

| | | | | | | | | | |
|---|---|---|---|---|---|---|---|---|---|
| ENSG00000274565.2 | **ENSG0000(** | 10798502( | lncRNA | **7.874** | 1.541 | 0.464 | **2.909** | 1.64E-02 | **3.25E-02** |
| ENSG00000144747.19 | **TMF1** | 7110 | protein_coding | **524.830** | -0.481 | -0.145 | **0.716** | 1.64E-02 | **3.25E-02** |
| ENSG00000288109.2 | **ENSG0000(** | 10192804$ | lncRNA | **5.721** | 1.415 | 0.426 | **2.667** | 1.64E-02 | **3.25E-02** |
| ENSG00000128595.19 | **CALU** | 813 | protein_coding | **676.272** | -0.332 | -0.100 | **0.794** | 1.65E-02 | **3.26E-02** |
| ENSG00000230202.1 | **RPL29P4** | | processed_pseud( | **213.056** | -0.850 | -0.256 | **0.555** | 1.65E-02 | **3.26E-02** |
| ENSG00000272279.2 | **ENSG0000(** | 10272394⁄ | lncRNA | **4.294** | 2.103 | 0.633 | **4.295** | 1.65E-02 | **3.26E-02** |
| ENSG00000253734.3 | **LINC01289** | 286184 | lncRNA | **4.622** | 2.021 | 0.608 | **4.059** | 1.65E-02 | **3.26E-02** |
| ENSG00000196507.13 | **TCEAL3** | 85012 | protein_coding | **316.290** | -0.290 | -0.087 | **0.818** | 1.65E-02 | **3.27E-02** |
| ENSG00000143643.15 | **TTC13** | 79573 | protein_coding | **462.574** | -0.308 | -0.093 | **0.808** | 1.65E-02 | **3.27E-02** |
| ENSG00000140983.17 | **RHOT2** | 89941 | protein_coding | **2274.183** | -0.270 | -0.081 | **0.829** | 1.65E-02 | **3.27E-02** |
| ENSG00000162594.18 | **IL23R** | 149233 | protein_coding | **25.287** | 0.859 | 0.259 | **1.813** | 1.66E-02 | **3.27E-02** |
| ENSG00000294791.1 | **ENSG00000294791** | | lncRNA | **6.085** | 1.760 | 0.530 | **3.387** | 1.66E-02 | **3.28E-02** |
| ENSG00000054793.16 | **ATP9A** | 10079 | protein_coding | **6.172** | 1.463 | 0.440 | **2.756** | 1.66E-02 | **3.28E-02** |
| ENSG00000104142.13 | **VPS18** | 57617 | protein_coding | **326.692** | 0.272 | 0.082 | **1.208** | 1.66E-02 | **3.28E-02** |
| ENSG00000182963.13 | **GJC1** | 10052 | protein_coding | **65.089** | -0.531 | -0.160 | **0.692** | 1.66E-02 | **3.28E-02** |
| ENSG00000128165.10 | **ADM2** | 79924 | protein_coding | **249.383** | -0.573 | -0.173 | **0.672** | 1.66E-02 | **3.28E-02** |
| ENSG00000068912.17 | **ERLEC1** | 27248 | protein_coding | **362.368** | -0.375 | -0.113 | **0.771** | 1.66E-02 | **3.28E-02** |
| ENSG00000220161.5 | **LINC02076** | 400579 | lncRNA | **5.463** | -2.125 | -0.640 | **0.229** | 1.66E-02 | **3.28E-02** |
| ENSG00000301807.1 | **ENSG00000301807** | | lncRNA | **18.288** | 0.913 | 0.275 | **1.882** | 1.66E-02 | **3.28E-02** |
| ENSG00000101166.18 | **PRELID3B** | 51012 | protein_coding | **816.947** | -0.341 | -0.103 | **0.789** | 1.66E-02 | **3.28E-02** |
| ENSG00000168827.17 | **GFM1** | 85476 | protein_coding | **438.867** | -0.323 | -0.097 | **0.799** | 1.66E-02 | **3.29E-02** |
| ENSG00000128915.14 | **ICE2** | 79664 | protein_coding | **430.851** | -0.318 | -0.096 | **0.802** | 1.67E-02 | **3.29E-02** |
| ENSG00000296560.1 | **ENSG00000296560** | | lncRNA | **6.369** | 1.356 | 0.408 | **2.560** | 1.67E-02 | **3.29E-02** |
| ENSG00000136100.16 | **VPS36** | 51028 | protein_coding | **436.337** | -0.324 | -0.098 | **0.799** | 1.67E-02 | **3.30E-02** |
| ENSG00000248360.7 | **LINC00504** | 201853 | lncRNA | **9.888** | 1.214 | 0.366 | **2.320** | 1.67E-02 | **3.30E-02** |
| ENSG00000290746.2 | **CA5BP1** | 340591 | lncRNA | **253.163** | -0.411 | -0.124 | **0.752** | 1.67E-02 | **3.30E-02** |
| ENSG00000304654.1 | **ENSG00000304654** | | lncRNA | **12.517** | 1.022 | 0.308 | **2.030** | 1.67E-02 | **3.30E-02** |
| ENSG00000290058.2 | **ENSG00000290058** | | lncRNA | **5.254** | 1.723 | 0.519 | **3.302** | 1.67E-02 | **3.30E-02** |
| ENSG00000119699.10 | **TGFB3** | 7043 | protein_coding | **71.466** | 0.583 | 0.175 | **1.498** | 1.68E-02 | **3.30E-02** |
| ENSG00000288093.1 | **ENSG00000288093** | | lncRNA | **9.525** | 1.207 | 0.363 | **2.308** | 1.68E-02 | **3.31E-02** |
| ENSG00000168334.11 | **XIRP1** | 165904 | protein_coding | **22.109** | 0.962 | 0.290 | **1.948** | 1.68E-02 | **3.31E-02** |

| | | | | | | | | | |
|---|---|---|---|---|---|---|---|---|---|
| ENSG00000233452.10 | **STXBP5-AS1** | | lncRNA | **7.541** | -1.365 | -0.411 | **0.388** | 1.68E-02 | **3.31E-02** |
| ENSG00000086061.18 | **DNAJA1** | 3301 | protein_coding | **2399.366** | -0.321 | -0.097 | **0.800** | 1.68E-02 | **3.31E-02** |
| ENSG00000223799.4 | **IL10RB-DT** | | lncRNA | **11.163** | 1.109 | 0.334 | **2.157** | 1.68E-02 | **3.32E-02** |
| ENSG00000272690.7 | **LINC02018** | 10798610( | lncRNA | **14.064** | 1.038 | 0.313 | **2.054** | 1.68E-02 | **3.32E-02** |
| ENSG00000154174.10 | **TOMM70** | 9868 | protein_coding | **865.471** | -0.269 | -0.081 | **0.830** | 1.68E-02 | **3.32E-02** |
| ENSG00000251002.10 | **TRD-AS1** | 105370401 | lncRNA | **376.536** | 0.436 | 0.131 | **1.353** | 1.68E-02 | **3.32E-02** |
| ENSG00000213231.15 | **TCL1B** | 9623 | protein_coding | **17.353** | -1.326 | -0.399 | **0.399** | 1.69E-02 | **3.33E-02** |
| ENSG00000107789.17 | **MINPP1** | 9562 | protein_coding | **143.493** | 0.466 | 0.140 | **1.381** | 1.69E-02 | **3.33E-02** |
| ENSG00000274290.3 | **H2BC6** | 8344 | protein_coding | **18.487** | 1.198 | 0.361 | **2.294** | 1.70E-02 | **3.34E-02** |
| ENSG00000181722.20 | **ZBTB20** | 26137 | protein_coding | **99.909** | 1.306 | 0.393 | **2.472** | 1.70E-02 | **3.34E-02** |
| ENSG00000140807.8 | **NKD1** | 85407 | protein_coding | **24.587** | 0.947 | 0.285 | **1.928** | 1.70E-02 | **3.35E-02** |
| ENSG00000065615.16 | **CYB5R4** | 51167 | protein_coding | **197.240** | -0.476 | -0.143 | **0.719** | 1.70E-02 | **3.35E-02** |
| ENSG00000010327.12 | **STAB1** | 23166 | protein_coding | **5.387** | 2.411 | 0.726 | **5.318** | 1.70E-02 | **3.35E-02** |
| ENSG00000105997.25 | **HOXA3** | 3200 | protein_coding | **10.440** | 1.489 | 0.448 | **2.807** | 1.70E-02 | **3.35E-02** |
| ENSG00000163249.15 | **CCNYL1** | 151195 | protein_coding | **168.288** | 0.451 | 0.136 | **1.367** | 1.70E-02 | **3.35E-02** |
| ENSG00000153975.12 | **ZUP1** | 221302 | protein_coding | **322.013** | -0.361 | -0.109 | **0.779** | 1.70E-02 | **3.35E-02** |
| ENSG00000286129.1 | **ENSG00000286129** | | lncRNA | **271.816** | 0.387 | 0.117 | **1.308** | 1.71E-02 | **3.35E-02** |
| ENSG00000134996.14 | **OSTF1** | 26578 | protein_coding | **624.915** | -0.269 | -0.081 | **0.830** | 1.71E-02 | **3.35E-02** |
| ENSG00000170954.14 | **ZNF415** | 55786 | protein_coding | **52.932** | -0.606 | -0.182 | **0.657** | 1.71E-02 | **3.36E-02** |
| ENSG00000254783.1 | **ENSG00000254783** | | processed_pseud( | **7.729** | 1.134 | 0.341 | **2.195** | 1.71E-02 | **3.37E-02** |
| ENSG00000177410.14 | **ZFAS1** | 441951 | lncRNA | **4011.707** | -0.317 | -0.096 | **0.802** | 1.71E-02 | **3.37E-02** |
| ENSG00000133056.16 | **PIK3C2B** | 5287 | protein_coding | **611.884** | -0.268 | -0.081 | **0.830** | 1.71E-02 | **3.37E-02** |
| ENSG00000160446.21 | **ZDHHC12** | 84885 | protein_coding | **916.017** | -0.323 | -0.097 | **0.800** | 1.71E-02 | **3.37E-02** |
| ENSG00000111215.14 | **PRR4** | 11272 | protein_coding | **63.117** | -0.676 | -0.203 | **0.626** | 1.72E-02 | **3.38E-02** |
| ENSG00000184678.11 | **H2BC21** | 8349 | protein_coding | **116.238** | 0.660 | 0.199 | **1.580** | 1.72E-02 | **3.38E-02** |
| ENSG00000180901.11 | **KCTD2** | 23510 | protein_coding | **431.000** | 0.268 | 0.081 | **1.204** | 1.72E-02 | **3.38E-02** |
| ENSG00000172264.20 | **MACROD2** | 140733 | protein_coding | **523.950** | -0.415 | -0.125 | **0.750** | 1.72E-02 | **3.38E-02** |
| ENSG00000268713.1 | **ENSG00000268713** | | lncRNA | **78.008** | -0.477 | -0.143 | **0.719** | 1.72E-02 | **3.38E-02** |
| ENSG00000237232.10 | **ZNF295-AS** | 150142 | lncRNA | **64.174** | 0.633 | 0.191 | **1.551** | 1.73E-02 | **3.40E-02** |
| ENSG00000197622.15 | **CDC42SE1** | 56882 | protein_coding | **2001.938** | 0.340 | 0.102 | **1.266** | 1.73E-02 | **3.40E-02** |
| ENSG00000270379.6 | **HEATR9** | 256957 | protein_coding | **4.858** | 1.731 | 0.521 | **3.320** | 1.73E-02 | **3.40E-02** |

| | | | | | | | | | |
|---|---|---|---|---|---|---|---|---|---|
| ENSG00000277767.2 | **ENSG00000277767** | | lncRNA | **36.462** | 0.549 | 0.165 | **1.463** | 1.73E-02 | **3.40E-02** |
| ENSG00000141068.18 | **KSR1** | 8844 | protein_coding | **2161.142** | 0.350 | 0.105 | **1.274** | 1.73E-02 | **3.40E-02** |
| ENSG00000064607.19 | **SUGP2** | 10147 | protein_coding | **3512.258** | -0.322 | -0.097 | **0.800** | 1.74E-02 | **3.41E-02** |
| ENSG00000166272.20 | **WBP1L** | 54838 | protein_coding | **436.899** | 0.285 | 0.086 | **1.218** | 1.74E-02 | **3.41E-02** |
| ENSG00000258603.6 | **ENSG00000258603** | | lncRNA | **7.121** | 1.450 | 0.437 | **2.733** | 1.74E-02 | **3.42E-02** |
| ENSG00000074181.11 | **NOTCH3** | 4854 | protein_coding | **13.930** | 1.794 | 0.540 | **3.468** | 1.74E-02 | **3.42E-02** |
| ENSG00000243701.9 | **DUBR** | 344595 | lncRNA | **24.831** | 0.770 | 0.232 | **1.705** | 1.74E-02 | **3.42E-02** |
| ENSG00000198838.16 | **RYR3** | 6263 | protein_coding | **6.533** | 1.736 | 0.522 | **3.330** | 1.74E-02 | **3.42E-02** |
| ENSG00000242086.9 | **MUC20-OT** | 727956 | lncRNA | **574.478** | 0.268 | 0.081 | **1.204** | 1.75E-02 | **3.43E-02** |
| ENSG00000277879.1 | **ENSG00000277879** | | lncRNA | **19.944** | -0.913 | -0.275 | **0.531** | 1.75E-02 | **3.43E-02** |
| ENSG00000234572.2 | **LINC01800** | 10192743 | lncRNA | **86.953** | -0.696 | -0.210 | **0.617** | 1.75E-02 | **3.43E-02** |
| ENSG00000279602.2 | **ENSG00000279602** | | lncRNA | **21.447** | -0.732 | -0.220 | **0.602** | 1.75E-02 | **3.44E-02** |
| ENSG00000180233.13 | **ZNRF2** | 223082 | protein_coding | **135.757** | -0.442 | -0.133 | **0.736** | 1.75E-02 | **3.44E-02** |
| ENSG00000022976.18 | **ZNF839** | 55778 | protein_coding | **554.973** | -0.331 | -0.099 | **0.795** | 1.76E-02 | **3.44E-02** |
| ENSG00000205057.7 | **CLLU1-AS1** | 574016 | lncRNA | **108.844** | 0.849 | 0.256 | **1.802** | 1.76E-02 | **3.45E-02** |
| ENSG00000291122.2 | **CASTOR3P** | 352954 | lncRNA | **90.327** | 0.536 | 0.161 | **1.450** | 1.76E-02 | **3.45E-02** |
| ENSG00000164039.17 | **BDH2** | 56898 | protein_coding | **353.304** | -0.322 | -0.097 | **0.800** | 1.76E-02 | **3.45E-02** |
| ENSG00000140479.19 | **PCSK6** | 5046 | protein_coding | **15.441** | 1.161 | 0.349 | **2.236** | 1.76E-02 | **3.45E-02** |
| ENSG00000145819.19 | **ARHGAP26** | 23092 | protein_coding | **610.616** | 0.286 | 0.086 | **1.219** | 1.76E-02 | **3.45E-02** |
| ENSG00000152484.16 | **USP12** | 219333 | protein_coding | **885.351** | 0.292 | 0.088 | **1.225** | 1.76E-02 | **3.45E-02** |
| ENSG00000080845.21 | **DLGAP4** | 22839 | protein_coding | **1396.818** | 0.322 | 0.097 | **1.250** | 1.77E-02 | **3.46E-02** |
| ENSG00000286638.3 | **ENSG00000286638** | | lncRNA | **4.933** | 1.900 | 0.572 | **3.733** | 1.77E-02 | **3.46E-02** |
| ENSG00000184260.6 | **H2AC20** | 8338 | protein_coding | **23.728** | -0.960 | -0.289 | **0.514** | 1.77E-02 | **3.46E-02** |
| ENSG00000186654.23 | **PRR5** | 55615 | protein_coding | **238.711** | 0.418 | 0.126 | **1.336** | 1.77E-02 | **3.47E-02** |
| ENSG00000259212.1 | **ENSG00000259212** | | lncRNA | **6.506** | 1.300 | 0.391 | **2.463** | 1.77E-02 | **3.47E-02** |
| ENSG00000294916.1 | **ENSG00000294916** | | lncRNA | **6.351** | 2.487 | 0.749 | **5.607** | 1.78E-02 | **3.48E-02** |
| ENSG00000129472.16 | **RAB2B** | 84932 | protein_coding | **548.317** | 0.269 | 0.081 | **1.205** | 1.78E-02 | **3.48E-02** |
| ENSG00000133597.13 | **ADCK2** | 90956 | protein_coding | **303.188** | 0.294 | 0.089 | **1.226** | 1.78E-02 | **3.49E-02** |
| ENSG00000229186.4 | **ADAM1A** | | unitary_pseudoge | **41.370** | 0.666 | 0.200 | **1.587** | 1.78E-02 | **3.49E-02** |
| ENSG00000186615.13 | **KTN1-AS1** | 10012907 | lncRNA | **58.965** | 0.504 | 0.152 | **1.418** | 1.78E-02 | **3.49E-02** |
| ENSG00000183648.13 | **NDUFB1** | 4707 | protein_coding | **1336.304** | -0.329 | -0.099 | **0.796** | 1.78E-02 | **3.49E-02** |

| | | | | | | | | | |
|---|---|---|---|---|---|---|---|---|---|
| ENSG00000121417.16 | **ZNF211** | 10520 | protein_coding | **419.597** | 0.324 | 0.098 | **1.252** | 1.79E-02 | **3.50E-02** |
| ENSG00000310788.1 | **ENSG00000310788** | | lncRNA | **17.032** | 1.558 | 0.469 | **2.944** | 1.79E-02 | **3.50E-02** |
| ENSG00000086570.13 | **FAT2** | 2196 | protein_coding | **7.537** | 1.333 | 0.401 | **2.519** | 1.79E-02 | **3.50E-02** |
| ENSG00000197061.5 | **H4C3** | 8364 | protein_coding | **6.070** | -1.442 | -0.434 | **0.368** | 1.79E-02 | **3.51E-02** |
| ENSG00000160285.17 | **LSS** | 4047 | protein_coding | **2505.145** | 0.372 | 0.112 | **1.294** | 1.79E-02 | **3.51E-02** |
| ENSG00000289088.1 | **ENSG00000289088** | | lncRNA | **3.705** | 2.400 | 0.722 | **5.278** | 1.79E-02 | **3.51E-02** |
| ENSG00000151923.20 | **TIAL1** | 7073 | protein_coding | **2241.542** | -0.289 | -0.087 | **0.818** | 1.80E-02 | **3.51E-02** |
| ENSG00000130731.18 | **METTL26** | 84326 | protein_coding | **1608.443** | -0.380 | -0.115 | **0.768** | 1.80E-02 | **3.51E-02** |
| ENSG00000134851.15 | **TMEM165** | 55858 | protein_coding | **767.938** | -0.279 | -0.084 | **0.824** | 1.80E-02 | **3.51E-02** |
| ENSG00000167130.20 | **DOLPP1** | 57171 | protein_coding | **171.286** | -0.372 | -0.112 | **0.773** | 1.80E-02 | **3.52E-02** |
| ENSG00000125246.18 | **CLYBL** | 171425 | protein_coding | **161.287** | 0.387 | 0.116 | **1.307** | 1.80E-02 | **3.52E-02** |
| ENSG00000160796.20 | **NBEAL2** | 23218 | protein_coding | **6406.152** | 0.301 | 0.091 | **1.232** | 1.80E-02 | **3.52E-02** |
| ENSG00000163602.12 | **RYBP** | 23429 | protein_coding | **330.440** | 0.474 | 0.143 | **1.389** | 1.81E-02 | **3.53E-02** |
| ENSG00000290970.2 | **ENSG0000(** | 10537925Z | lncRNA | **13.852** | -2.010 | -0.605 | **0.248** | 1.81E-02 | **3.53E-02** |
| ENSG00000120837.10 | **NFYB** | 4801 | protein_coding | **403.111** | -0.393 | -0.118 | **0.761** | 1.81E-02 | **3.53E-02** |
| ENSG00000171988.21 | **JMJD1C** | 221037 | protein_coding | **888.371** | 0.398 | 0.120 | **1.318** | 1.82E-02 | **3.55E-02** |
| ENSG00000142449.15 | **FBN3** | 84467 | protein_coding | **9.867** | 2.166 | 0.652 | **4.488** | 1.82E-02 | **3.55E-02** |
| ENSG00000273706.7 | **LHX1** | 3975 | protein_coding | **5.742** | 1.658 | 0.499 | **3.156** | 1.82E-02 | **3.55E-02** |
| ENSG00000079134.15 | **THOC1** | 9984 | protein_coding | **911.621** | -0.300 | -0.090 | **0.812** | 1.82E-02 | **3.55E-02** |
| ENSG00000291081.2 | **ENSG00000291081** | | lncRNA | **128.813** | 0.416 | 0.125 | **1.334** | 1.82E-02 | **3.56E-02** |
| ENSG00000261226.1 | **ENSG00000261226** | | lncRNA | **12.732** | 1.154 | 0.347 | **2.225** | 1.82E-02 | **3.56E-02** |
| ENSG00000293414.2 | **TMEM277I** | 284365 | lncRNA | **73.847** | -0.569 | -0.171 | **0.674** | 1.83E-02 | **3.56E-02** |
| ENSG00000103066.15 | **PLA2G15** | 23659 | protein_coding | **246.197** | 0.386 | 0.116 | **1.307** | 1.83E-02 | **3.57E-02** |
| ENSG00000128311.16 | **TST** | 7263 | protein_coding | **201.450** | 0.287 | 0.086 | **1.220** | 1.83E-02 | **3.57E-02** |
| ENSG00000293167.1 | **WDR5CP** | | lncRNA | **6.411** | 1.761 | 0.530 | **3.390** | 1.84E-02 | **3.59E-02** |
| ENSG00000214226.10 | **C17orf67** | 339210 | protein_coding | **116.430** | 0.459 | 0.138 | **1.374** | 1.84E-02 | **3.59E-02** |
| ENSG00000284753.4 | **EEF1AKMT** | 11059956 | protein_coding | **95.888** | -0.438 | -0.132 | **0.738** | 1.84E-02 | **3.60E-02** |
| ENSG00000303878.1 | **ENSG00000303878** | | lncRNA | **8.597** | 1.081 | 0.325 | **2.116** | 1.85E-02 | **3.60E-02** |
| ENSG00000144476.8 | **ACKR3** | 57007 | protein_coding | **209.497** | 0.536 | 0.161 | **1.450** | 1.85E-02 | **3.60E-02** |
| ENSG00000257698.4 | **GIHCG** | | lncRNA | **173.691** | -0.364 | -0.109 | **0.777** | 1.85E-02 | **3.61E-02** |
| ENSG00000119917.16 | **IFIT3** | 3437 | protein_coding | **2397.807** | 0.461 | 0.139 | **1.377** | 1.85E-02 | **3.61E-02** |

| | | | | | | | | | |
|---|---|---|---|---|---|---|---|---|---|
| ENSG00000221817.13 | **PPP3CB-AS** | 101929145 | lncRNA | **104.939** | -0.460 | -0.138 | **0.727** | 1.85E-02 | **3.61E-02** |
| ENSG00000255153.1 | **TOLLIP-DT** | 255512 | lncRNA | **28.240** | 0.689 | 0.207 | **1.612** | 1.86E-02 | **3.62E-02** |
| ENSG00000128564.10 | **VGF** | 7425 | protein_coding | **10.983** | 1.070 | 0.322 | **2.100** | 1.86E-02 | **3.62E-02** |
| ENSG00000173276.16 | **ZBTB21** | 49854 | protein_coding | **311.918** | 0.396 | 0.119 | **1.315** | 1.86E-02 | **3.63E-02** |
| ENSG00000270720.1 | **ENSG00000270720** | | lncRNA | **5.681** | 1.450 | 0.437 | **2.733** | 1.86E-02 | **3.63E-02** |
| ENSG00000290002.1 | **ENSG00000290002** | | lncRNA | **9.178** | 1.264 | 0.380 | **2.402** | 1.86E-02 | **3.63E-02** |
| ENSG00000243199.1 | **RPL6P10** | | processed_pseud | **225.055** | -0.319 | -0.096 | **0.802** | 1.86E-02 | **3.63E-02** |
| ENSG00000187800.16 | **PEAR1** | 375033 | protein_coding | **5.399** | 1.986 | 0.598 | **3.961** | 1.87E-02 | **3.63E-02** |
| ENSG00000286136.3 | **ENSG00000286136** | | lncRNA | **6.915** | -1.362 | -0.410 | **0.389** | 1.87E-02 | **3.64E-02** |
| ENSG00000290450.2 | **ENSG00000290450** | | lncRNA | **237.461** | -0.508 | -0.153 | **0.703** | 1.87E-02 | **3.64E-02** |
| ENSG00000127922.12 | **SEM1** | 7979 | protein_coding | **1874.114** | -0.358 | -0.108 | **0.780** | 1.87E-02 | **3.64E-02** |
| ENSG00000065548.21 | **ZC3H15** | 55854 | protein_coding | **839.165** | -0.317 | -0.096 | **0.803** | 1.87E-02 | **3.64E-02** |
| ENSG00000198839.12 | **ZNF277** | 11179 | protein_coding | **473.809** | 0.324 | 0.098 | **1.252** | 1.87E-02 | **3.65E-02** |
| ENSG00000144741.21 | **SLC25A26** | 115286 | protein_coding | **267.960** | -0.452 | -0.136 | **0.731** | 1.87E-02 | **3.65E-02** |
| ENSG00000166387.16 | **PPFIBP2** | 8495 | protein_coding | **800.016** | -0.287 | -0.086 | **0.819** | 1.87E-02 | **3.65E-02** |
| ENSG00000290460.2 | **ENSG00000290460** | | lncRNA | **48.337** | 0.686 | 0.206 | **1.609** | 1.88E-02 | **3.65E-02** |
| ENSG00000306369.1 | **ENSG00000306369** | | lncRNA | **16.288** | 0.852 | 0.257 | **1.805** | 1.88E-02 | **3.66E-02** |
| ENSG00000275221.3 | **H2AC15** | 8330 | protein_coding | **16.276** | -0.821 | -0.247 | **0.566** | 1.88E-02 | **3.66E-02** |
| ENSG00000293232.1 | **ENSG0000** | 10013436 | lncRNA | **15.533** | 0.837 | 0.252 | **1.787** | 1.88E-02 | **3.66E-02** |
| ENSG00000120925.18 | **RNF170** | 81790 | protein_coding | **160.562** | 0.429 | 0.129 | **1.347** | 1.88E-02 | **3.66E-02** |
| ENSG00000138777.22 | **PPA2** | 27068 | protein_coding | **990.911** | -0.326 | -0.098 | **0.798** | 1.89E-02 | **3.67E-02** |
| ENSG00000263327.7 | **TAPT1-AS1** | | lncRNA | **44.789** | 0.600 | 0.181 | **1.516** | 1.89E-02 | **3.67E-02** |
| ENSG00000286522.3 | **H3C2** | 8358 | protein_coding | **8.302** | -1.505 | -0.453 | **0.352** | 1.89E-02 | **3.67E-02** |
| ENSG00000164576.14 | **SAP30L** | 79685 | protein_coding | **444.918** | -0.421 | -0.127 | **0.747** | 1.89E-02 | **3.67E-02** |
| ENSG00000163960.15 | **UBXN7** | 26043 | protein_coding | **1080.926** | 0.301 | 0.091 | **1.232** | 1.89E-02 | **3.68E-02** |
| ENSG00000278002.2 | **ENSG00000278002** | | lncRNA | **5.246** | 1.483 | 0.446 | **2.795** | 1.90E-02 | **3.69E-02** |
| ENSG00000103404.17 | **USP31** | 57478 | protein_coding | **137.404** | -0.513 | -0.154 | **0.701** | 1.90E-02 | **3.69E-02** |
| ENSG00000136875.15 | **PRPF4** | 9128 | protein_coding | **470.397** | -0.269 | -0.081 | **0.830** | 1.90E-02 | **3.69E-02** |
| ENSG00000273399.2 | **ENSG00000273399** | | lncRNA | **12.943** | 0.989 | 0.298 | **1.985** | 1.90E-02 | **3.69E-02** |
| ENSG00000165804.18 | **ZNF219** | 51222 | protein_coding | **110.247** | 0.401 | 0.121 | **1.320** | 1.90E-02 | **3.69E-02** |
| ENSG00000299450.1 | **ENSG00000299450** | | lncRNA | **43.351** | -0.685 | -0.206 | **0.622** | 1.90E-02 | **3.70E-02** |

| | | | | | | | | | |
|---|---|---|---|---|---|---|---|---|---|
| ENSG00000118707.13 | **TGIF2** | 60436 | protein_coding | **735.363** | -0.307 | -0.092 | **0.808** | 1.90E-02 | **3.70E-02** |
| ENSG00000196576.18 | **PLXNB2** | 23654 | protein_coding | **3110.663** | -0.361 | -0.109 | **0.778** | 1.90E-02 | **3.70E-02** |
| ENSG00000117242.7 | **PINK1-AS** | 10086154{ | lncRNA | **65.744** | 0.428 | 0.129 | **1.346** | 1.90E-02 | **3.70E-02** |
| ENSG00000163812.18 | **ZDHHC3** | 51304 | protein_coding | **635.724** | -0.278 | -0.084 | **0.825** | 1.91E-02 | **3.70E-02** |
| ENSG00000266923.1 | **ENSG00000266923** | | lncRNA | **183.724** | -0.447 | -0.134 | **0.734** | 1.91E-02 | **3.70E-02** |
| ENSG00000119280.19 | **C1orf198** | 84886 | protein_coding | **63.547** | 0.537 | 0.162 | **1.451** | 1.91E-02 | **3.70E-02** |
| ENSG00000213983.14 | **AP1G2** | 8906 | protein_coding | **4308.153** | 0.396 | 0.119 | **1.316** | 1.91E-02 | **3.71E-02** |
| ENSG00000099991.20 | **CABIN1** | 23523 | protein_coding | **1276.057** | -0.323 | -0.097 | **0.799** | 1.91E-02 | **3.71E-02** |
| ENSG00000233942.2 | **ENSG0000(** | 10798682: | lncRNA | **4.584** | 2.508 | 0.755 | **5.687** | 1.91E-02 | **3.71E-02** |
| ENSG00000149403.15 | **GRIK4** | 2900 | protein_coding | **36.870** | 0.598 | 0.180 | **1.514** | 1.91E-02 | **3.72E-02** |
| ENSG00000213648.13 | **SULT1A4** | 445329 | protein_coding | **194.845** | 0.621 | 0.187 | **1.538** | 1.92E-02 | **3.72E-02** |
| ENSG00000147604.16 | **RPL7** | 6129 | protein_coding | **26132.056** | -0.311 | -0.094 | **0.806** | 1.92E-02 | **3.73E-02** |
| ENSG00000247271.10 | **ZBED5-AS1** | 729013 | lncRNA | **186.050** | 0.342 | 0.103 | **1.268** | 1.92E-02 | **3.73E-02** |
| ENSG00000171295.14 | **ZNF440** | 126070 | protein_coding | **197.708** | 0.302 | 0.091 | **1.233** | 1.93E-02 | **3.73E-02** |
| ENSG00000122705.20 | **CLTA** | 1211 | protein_coding | **1368.224** | -0.287 | -0.086 | **0.819** | 1.93E-02 | **3.74E-02** |
| ENSG00000170819.5 | **BFSP2** | 8419 | protein_coding | **163.135** | 0.562 | 0.169 | **1.477** | 1.93E-02 | **3.74E-02** |
| ENSG00000290574.3 | **ENSG00000290574** | | lncRNA | **40.559** | -0.625 | -0.188 | **0.648** | 1.93E-02 | **3.74E-02** |
| ENSG00000169570.12 | **DTWD2** | 285605 | protein_coding | **87.870** | 0.412 | 0.124 | **1.330** | 1.93E-02 | **3.74E-02** |
| ENSG00000113712.20 | **CSNK1A1** | 1452 | protein_coding | **1128.427** | -0.379 | -0.114 | **0.769** | 1.93E-02 | **3.75E-02** |
| ENSG00000293707.1 | **ENSG00000293707** | | lncRNA | **26.365** | 0.663 | 0.200 | **1.583** | 1.94E-02 | **3.76E-02** |
| ENSG00000118961.17 | **LDAH** | 60526 | protein_coding | **407.769** | -0.275 | -0.083 | **0.826** | 1.94E-02 | **3.76E-02** |
| ENSG00000104728.18 | **ARHGEF10** | 9639 | protein_coding | **79.448** | 0.981 | 0.295 | **1.973** | 1.94E-02 | **3.76E-02** |
| ENSG00000214756.9 | **CSKMT** | 751071 | protein_coding | **86.974** | -0.497 | -0.150 | **0.708** | 1.94E-02 | **3.77E-02** |
| ENSG00000237292.2 | **LINC01732** | 10192897: | lncRNA | **33.615** | 0.622 | 0.187 | **1.539** | 1.94E-02 | **3.77E-02** |
| ENSG00000196338.17 | **NLGN3** | 54413 | protein_coding | **79.112** | 0.432 | 0.130 | **1.349** | 1.95E-02 | **3.77E-02** |
| ENSG00000003249.17 | **DBNDD1** | 79007 | protein_coding | **407.029** | -0.472 | -0.142 | **0.721** | 1.95E-02 | **3.77E-02** |
| ENSG00000103174.15 | **NAGPA** | 51172 | protein_coding | **176.338** | -0.421 | -0.127 | **0.747** | 1.95E-02 | **3.78E-02** |
| ENSG00000176406.26 | **RIMS2** | 9699 | protein_coding | **15.477** | 1.146 | 0.345 | **2.213** | 1.95E-02 | **3.78E-02** |
| ENSG00000131759.20 | **RARA** | 5914 | protein_coding | **1613.267** | 0.271 | 0.082 | **1.207** | 1.95E-02 | **3.78E-02** |
| ENSG00000168874.16 | **ATOH8** | 84913 | protein_coding | **5.030** | 2.047 | 0.616 | **4.134** | 1.96E-02 | **3.79E-02** |
| ENSG00000291115.2 | **ENSG00000291115** | | lncRNA | **4.300** | 2.485 | 0.748 | **5.597** | 1.96E-02 | **3.79E-02** |

| | | | | | | | | | |
|---|---|---|---|---|---|---|---|---|---|
| ENSG00000283782.2 | **ENSG00000283782** | | protein_coding | **198.888** | -1.598 | -0.481 | **0.330** | 1.96E-02 | **3.79E-02** |
| ENSG00000253537.4 | **PCDHGA7** | 56108 | protein_coding | **15.509** | 0.842 | 0.253 | **1.792** | 1.97E-02 | **3.82E-02** |
| ENSG00000172586.9 | **CHCHD1** | 118487 | protein_coding | **590.027** | -0.287 | -0.086 | **0.819** | 1.98E-02 | **3.82E-02** |
| ENSG00000232586.1 | **KIAA1614-** | 10334492 | lncRNA | **22.476** | 0.679 | 0.204 | **1.601** | 1.98E-02 | **3.82E-02** |
| ENSG00000122482.24 | **ZNF644** | 84146 | protein_coding | **203.195** | -0.357 | -0.108 | **0.781** | 1.98E-02 | **3.82E-02** |
| ENSG00000230325.1 | **ENSG00000230325** | | lncRNA | **6.218** | 1.349 | 0.406 | **2.548** | 1.98E-02 | **3.83E-02** |
| ENSG00000166278.18 | **C2** | 717 | protein_coding | **12.343** | 1.044 | 0.314 | **2.061** | 1.98E-02 | **3.83E-02** |
| ENSG00000290923.2 | **PGM5P2** | 595135 | lncRNA | **28.102** | -0.675 | -0.203 | **0.626** | 1.98E-02 | **3.83E-02** |
| ENSG00000130695.18 | **CEP85** | 64793 | protein_coding | **364.221** | -0.325 | -0.098 | **0.798** | 1.98E-02 | **3.83E-02** |
| ENSG00000157654.21 | **PALM2AKA** | 445815 | protein_coding | **2128.468** | 0.455 | 0.137 | **1.371** | 1.99E-02 | **3.84E-02** |
| ENSG00000270049.5 | **ATP6V0D1-** | 10192783 | lncRNA | **83.627** | -0.353 | -0.106 | **0.783** | 1.99E-02 | **3.84E-02** |
| ENSG00000287825.4 | **ENSG00000287825** | | lncRNA | **58.805** | 0.654 | 0.197 | **1.573** | 1.99E-02 | **3.84E-02** |
| ENSG00000180089.8 | **TMEM86B** | 255043 | protein_coding | **155.083** | -0.383 | -0.115 | **0.767** | 1.99E-02 | **3.85E-02** |
| ENSG00000073969.21 | **NSF** | 4905 | protein_coding | **340.201** | 0.369 | 0.111 | **1.292** | 1.99E-02 | **3.85E-02** |
| ENSG00000106211.12 | **HSPB1** | 3315 | protein_coding | **2801.526** | 0.312 | 0.094 | **1.241** | 1.99E-02 | **3.85E-02** |
| ENSG00000087095.15 | **NLK** | 51701 | protein_coding | **438.253** | 0.386 | 0.116 | **1.307** | 1.99E-02 | **3.85E-02** |
| ENSG00000186056.12 | **MATN1-AS** | 10012919 | lncRNA | **148.667** | 0.474 | 0.143 | **1.389** | 2.00E-02 | **3.86E-02** |
| ENSG00000136997.22 | **MYC** | 4609 | protein_coding | **1571.814** | -0.378 | -0.114 | **0.769** | 2.00E-02 | **3.86E-02** |
| ENSG00000106400.14 | **ZNHIT1** | 10467 | protein_coding | **1886.886** | 0.340 | 0.102 | **1.266** | 2.00E-02 | **3.86E-02** |
| ENSG00000128266.11 | **GNAZ** | 2781 | protein_coding | **13.785** | 1.097 | 0.330 | **2.140** | 2.00E-02 | **3.86E-02** |
| ENSG00000162194.14 | **LBHD1** | 79081 | protein_coding | **303.727** | 0.265 | 0.080 | **1.201** | 2.00E-02 | **3.86E-02** |
| ENSG00000259736.2 | **CRTC3-AS1** | | lncRNA | **16.479** | 0.852 | 0.256 | **1.805** | 2.00E-02 | **3.87E-02** |
| ENSG00000091490.13 | **SEL1L3** | 23231 | protein_coding | **1144.402** | -0.413 | -0.124 | **0.751** | 2.00E-02 | **3.87E-02** |
| ENSG00000157800.20 | **SLC37A3** | 84255 | protein_coding | **211.727** | 0.363 | 0.109 | **1.286** | 2.01E-02 | **3.87E-02** |
| ENSG00000273711.2 | **ENSG00000273711** | | lncRNA | **38.696** | 1.292 | 0.389 | **2.448** | 2.01E-02 | **3.87E-02** |
| ENSG00000180530.12 | **NRIP1** | 8204 | protein_coding | **87.147** | 0.560 | 0.169 | **1.474** | 2.01E-02 | **3.88E-02** |
| ENSG00000235781.2 | **LINC02569** | 11108299 | lncRNA | **5.503** | 1.531 | 0.461 | **2.890** | 2.01E-02 | **3.88E-02** |
| ENSG00000166682.15 | **TMPRSS5** | 80975 | protein_coding | **13.538** | -1.234 | -0.372 | **0.425** | 2.01E-02 | **3.88E-02** |
| ENSG00000225733.8 | **FGD5-AS1** | 100505641 | lncRNA | **796.278** | 0.307 | 0.092 | **1.237** | 2.01E-02 | **3.88E-02** |
| ENSG00000258725.2 | **PRC1-AS1** | 10050711 | lncRNA | **26.761** | 0.695 | 0.209 | **1.619** | 2.01E-02 | **3.88E-02** |
| ENSG00000277782.2 | **ENSG00000277782** | | lncRNA | **42.017** | 0.565 | 0.170 | **1.479** | 2.01E-02 | **3.88E-02** |

| | | | | | | | | | |
|---|---|---|---|---|---|---|---|---|---|
| ENSG00000134755.21 | **DSC2** | 1824 | protein_coding | **31.213** | -0.968 | -0.291 | **0.511** | 2.01E-02 | **3.89E-02** |
| ENSG00000232472.1 | **EEF1B2P3** | | processed_pseudo | **6.716** | -1.249 | -0.376 | **0.421** | 2.02E-02 | **3.89E-02** |
| ENSG00000163704.15 | **PRRT3** | 285368 | protein_coding | **262.954** | 0.418 | 0.126 | **1.336** | 2.02E-02 | **3.89E-02** |
| ENSG00000164300.20 | **SERINC5** | 256987 | protein_coding | **61.252** | 0.512 | 0.154 | **1.426** | 2.02E-02 | **3.90E-02** |
| ENSG00000133639.7 | **BTG1** | 694 | protein_coding | **6729.641** | 0.338 | 0.102 | **1.264** | 2.02E-02 | **3.90E-02** |
| ENSG00000131697.20 | **NPHP4** | 261734 | protein_coding | **195.514** | 0.305 | 0.092 | **1.235** | 2.03E-02 | **3.90E-02** |
| ENSG00000198920.12 | **KIAA0753** | 9851 | protein_coding | **259.820** | -0.290 | -0.087 | **0.818** | 2.03E-02 | **3.91E-02** |
| ENSG00000258072.2 | **PDCD2P2** | | transcribed_proce | **47.361** | 0.547 | 0.165 | **1.461** | 2.03E-02 | **3.91E-02** |
| ENSG00000173065.16 | **FAM222B** | 55731 | protein_coding | **277.929** | 0.302 | 0.091 | **1.233** | 2.03E-02 | **3.91E-02** |
| ENSG00000151093.10 | **OXSM** | 54995 | protein_coding | **272.638** | 0.295 | 0.089 | **1.227** | 2.03E-02 | **3.91E-02** |
| ENSG00000224505.5 | **HEXIM2-AS1** | | lncRNA | **213.357** | 0.296 | 0.089 | **1.228** | 2.03E-02 | **3.92E-02** |
| ENSG00000123091.7 | **RNF11** | 26994 | protein_coding | **233.039** | 0.595 | 0.179 | **1.510** | 2.04E-02 | **3.93E-02** |
| ENSG00000114030.15 | **KPNA1** | 3836 | protein_coding | **752.044** | 0.336 | 0.101 | **1.262** | 2.04E-02 | **3.93E-02** |
| ENSG00000076554.18 | **TPD52** | 7163 | protein_coding | **838.877** | -0.342 | -0.103 | **0.789** | 2.04E-02 | **3.93E-02** |
| ENSG00000168826.18 | **ZBTB49** | 166793 | protein_coding | **86.601** | -0.452 | -0.136 | **0.731** | 2.04E-02 | **3.93E-02** |
| ENSG00000223874.4 | **TESHL** | | lncRNA | **5.696** | 1.860 | 0.560 | **3.631** | 2.05E-02 | **3.94E-02** |
| ENSG00000255750.7 | **ENSG00000255750** | | lncRNA | **8.447** | 1.232 | 0.371 | **2.349** | 2.05E-02 | **3.95E-02** |
| ENSG00000183128.7 | **CALHM3** | 119395 | protein_coding | **11.755** | 1.076 | 0.324 | **2.108** | 2.05E-02 | **3.95E-02** |
| ENSG00000288817.2 | **ENSG00000288817** | | lncRNA | **28.345** | 0.668 | 0.201 | **1.589** | 2.05E-02 | **3.95E-02** |
| ENSG00000149654.13 | **CDH22** | 64405 | protein_coding | **5.042** | 1.901 | 0.572 | **3.736** | 2.05E-02 | **3.96E-02** |
| ENSG00000256576.4 | **LINC02361** | 10099624 | lncRNA | **14.623** | 1.008 | 0.303 | **2.011** | 2.06E-02 | **3.96E-02** |
| ENSG00000281852.2 | **LINC00891** | | lncRNA | **50.600** | 0.829 | 0.250 | **1.777** | 2.06E-02 | **3.96E-02** |
| ENSG00000120008.17 | **WDR11** | 55717 | protein_coding | **1085.613** | 0.322 | 0.097 | **1.250** | 2.06E-02 | **3.96E-02** |
| ENSG00000071242.15 | **RPS6KA2** | 6196 | protein_coding | **247.717** | 0.460 | 0.139 | **1.376** | 2.06E-02 | **3.97E-02** |
| ENSG00000285219.5 | **HULC** | 728655 | lncRNA | **22.356** | 0.919 | 0.277 | **1.891** | 2.06E-02 | **3.97E-02** |
| ENSG00000100485.14 | **SOS2** | 6655 | protein_coding | **248.999** | -0.322 | -0.097 | **0.800** | 2.06E-02 | **3.97E-02** |
| ENSG00000130270.17 | **ATP8B3** | 148229 | protein_coding | **76.199** | 0.468 | 0.141 | **1.383** | 2.07E-02 | **3.98E-02** |
| ENSG00000262712.1 | **ENSG00000262712** | | lncRNA | **11.322** | -0.966 | -0.291 | **0.512** | 2.07E-02 | **3.98E-02** |
| ENSG00000240303.10 | **ACAD11** | 84129 | protein_coding | **1119.571** | 0.274 | 0.083 | **1.209** | 2.07E-02 | **3.98E-02** |
| ENSG00000129946.13 | **SHC2** | 25759 | protein_coding | **7.794** | 1.412 | 0.425 | **2.660** | 2.07E-02 | **3.98E-02** |
| ENSG00000051382.12 | **PIK3CB** | 5291 | protein_coding | **182.190** | 0.290 | 0.087 | **1.223** | 2.07E-02 | **3.99E-02** |

| | | | | | | | | | |
|---|---|---|---|---|---|---|---|---|---|
| ENSG00000110697.16 | **PITPNM1** | 9600 | protein_coding | **961.567** | 0.335 | 0.101 | **1.261** | 2.08E-02 | **3.99E-02** |
| ENSG00000065057.11 | **NTHL1** | 4913 | protein_coding | **357.990** | 0.301 | 0.090 | **1.232** | 2.08E-02 | **4.00E-02** |
| ENSG00000174607.14 | **UGT8** | 7368 | protein_coding | **12.200** | 1.236 | 0.372 | **2.355** | 2.09E-02 | **4.01E-02** |
| ENSG00000175691.11 | **ZNF77** | 58492 | protein_coding | **86.758** | 0.479 | 0.144 | **1.394** | 2.09E-02 | **4.01E-02** |
| ENSG00000228203.9 | **GRASLND** | 386597 | lncRNA | **9.124** | 1.240 | 0.373 | **2.363** | 2.09E-02 | **4.01E-02** |
| ENSG00000132792.23 | **CTNNBL1** | 56259 | protein_coding | **1053.199** | -0.281 | -0.085 | **0.823** | 2.09E-02 | **4.02E-02** |
| ENSG00000109452.15 | **INPP4B** | 8821 | protein_coding | **39.285** | 0.753 | 0.227 | **1.685** | 2.09E-02 | **4.02E-02** |
| ENSG00000119616.12 | **FCF1** | 51077 | protein_coding | **307.490** | -0.393 | -0.118 | **0.762** | 2.09E-02 | **4.02E-02** |
| ENSG00000100092.25 | **SH3BP1** | 23616 | protein_coding | **992.679** | -0.287 | -0.086 | **0.820** | 2.09E-02 | **4.02E-02** |
| ENSG00000238062.7 | **SPATA3-AS** | 348761 | lncRNA | **65.263** | 0.630 | 0.190 | **1.548** | 2.09E-02 | **4.02E-02** |
| ENSG00000113302.7 | **IL12B** | 3593 | protein_coding | **16.991** | -0.941 | -0.283 | **0.521** | 2.10E-02 | **4.03E-02** |
| ENSG00000230709.1 | **ENSG0000** | 284191 | lncRNA | **26.246** | -0.682 | -0.205 | **0.623** | 2.10E-02 | **4.03E-02** |
| ENSG00000297670.1 | **ENSG0000** | 10537525 | lncRNA | **22.433** | 0.676 | 0.204 | **1.598** | 2.10E-02 | **4.03E-02** |
| ENSG00000272186.2 | **VPS11-DT** | 12490276 | lncRNA | **11.003** | -0.909 | -0.274 | **0.533** | 2.10E-02 | **4.03E-02** |
| ENSG00000136040.10 | **PLXNC1** | 10154 | protein_coding | **416.923** | -0.431 | -0.130 | **0.742** | 2.10E-02 | **4.04E-02** |
| ENSG00000244607.9 | **CCDC13** | 152206 | protein_coding | **24.201** | 0.873 | 0.263 | **1.831** | 2.11E-02 | **4.04E-02** |
| ENSG00000188620.12 | **HMX3** | 340784 | protein_coding | **141.624** | -0.435 | -0.131 | **0.740** | 2.11E-02 | **4.04E-02** |
| ENSG00000169031.23 | **COL4A3** | 1285 | protein_coding | **599.586** | 0.448 | 0.135 | **1.365** | 2.11E-02 | **4.05E-02** |
| ENSG00000132680.13 | **KHDC4** | 22889 | protein_coding | **1303.959** | 0.264 | 0.079 | **1.201** | 2.11E-02 | **4.05E-02** |
| ENSG00000234608.10 | **MAPKAPK** | 51275 | lncRNA | **383.689** | -0.274 | -0.083 | **0.827** | 2.11E-02 | **4.05E-02** |
| ENSG00000304265.1 | **ENSG00000304265** | | lncRNA | **14.580** | 0.826 | 0.249 | **1.772** | 2.12E-02 | **4.06E-02** |
| ENSG00000211592.8 | **IGKC** | | IG_C_gene | **63674.456** | -0.345 | -0.104 | **0.787** | 2.12E-02 | **4.07E-02** |
| ENSG00000160606.12 | **TLCD1** | 116238 | protein_coding | **95.284** | 0.408 | 0.123 | **1.327** | 2.12E-02 | **4.07E-02** |
| ENSG00000261136.1 | **ENSG00000261136** | | lncRNA | **28.727** | 0.701 | 0.211 | **1.625** | 2.12E-02 | **4.07E-02** |
| ENSG00000167604.17 | **NFKBID** | 84807 | protein_coding | **973.309** | -0.449 | -0.135 | **0.732** | 2.13E-02 | **4.08E-02** |
| ENSG00000131779.13 | **PEX11B** | 8799 | protein_coding | **549.209** | -0.271 | -0.082 | **0.829** | 2.13E-02 | **4.08E-02** |
| ENSG00000224272.3 | **ENSG0000** | 12490534 | lncRNA | **3.861** | -1.951 | -0.587 | **0.259** | 2.13E-02 | **4.08E-02** |
| ENSG00000275163.1 | **ENSG0000** | 10242 | protein_coding | **17.460** | -3.520 | -1.060 | **0.087** | 2.13E-02 | **4.08E-02** |
| ENSG00000078142.16 | **PIK3C3** | 5289 | protein_coding | **556.995** | 0.323 | 0.097 | **1.251** | 2.13E-02 | **4.08E-02** |
| ENSG00000255435.8 | **TTC36-AS1** | 10192908 | lncRNA | **6.155** | 1.518 | 0.457 | **2.864** | 2.13E-02 | **4.08E-02** |
| ENSG00000170100.16 | **ZNF778** | 197320 | protein_coding | **196.439** | 0.554 | 0.167 | **1.468** | 2.13E-02 | **4.08E-02** |

| | | | | | | | | | |
|---|---|---|---|---|---|---|---|---|---|
| ENSG00000163689.23 | **CFAP20DC** | 200844 | protein_coding | **22.861** | -0.823 | -0.248 | **0.565** | 2.13E-02 | **4.08E-02** |
| ENSG00000175505.12 | **CLCF1** | 23529 | protein_coding | **90.445** | -0.648 | -0.195 | **0.638** | 2.13E-02 | **4.08E-02** |
| ENSG00000253865.2 | **ENSG00000253865** | | lncRNA | **14.428** | 0.960 | 0.289 | **1.945** | 2.13E-02 | **4.09E-02** |
| ENSG00000146576.15 | **INTS15** | 79034 | protein_coding | **292.017** | -0.310 | -0.093 | **0.807** | 2.13E-02 | **4.09E-02** |
| ENSG00000290110.2 | **ENSG00000290110** | | lncRNA | **9.756** | 1.142 | 0.344 | **2.207** | 2.13E-02 | **4.09E-02** |
| ENSG00000254093.11 | **PINX1** | 54984 | protein_coding | **323.459** | -0.280 | -0.084 | **0.824** | 2.13E-02 | **4.09E-02** |
| ENSG00000151208.19 | **DLG5** | 9231 | protein_coding | **148.612** | 0.472 | 0.142 | **1.387** | 2.13E-02 | **4.09E-02** |
| ENSG00000245281.10 | **ASAH1-AS1** | | lncRNA | **21.979** | 0.909 | 0.274 | **1.878** | 2.14E-02 | **4.09E-02** |
| ENSG00000291104.2 | **BTN2A3P** | 54718 | lncRNA | **104.851** | 0.346 | 0.104 | **1.271** | 2.15E-02 | **4.11E-02** |
| ENSG00000132704.18 | **FCRL2** | 79368 | protein_coding | **7.493** | 2.083 | 0.627 | **4.238** | 2.15E-02 | **4.11E-02** |
| ENSG00000198715.15 | **GLMP** | 112770 | protein_coding | **378.822** | -0.337 | -0.102 | **0.792** | 2.15E-02 | **4.11E-02** |
| ENSG00000279668.5 | **ENSG00000279668** | | lncRNA | **7.584** | 1.415 | 0.426 | **2.667** | 2.15E-02 | **4.11E-02** |
| ENSG00000165533.21 | **TTC8** | 123016 | protein_coding | **43.698** | 0.648 | 0.195 | **1.567** | 2.15E-02 | **4.11E-02** |
| ENSG00000276101.1 | **ENSG00000276101** | | lncRNA | **5.786** | 1.539 | 0.463 | **2.907** | 2.15E-02 | **4.12E-02** |
| ENSG00000132394.13 | **EEFSEC** | 60678 | protein_coding | **428.970** | -0.289 | -0.087 | **0.819** | 2.15E-02 | **4.12E-02** |
| ENSG00000002330.16 | **BAD** | 572 | protein_coding | **413.288** | -0.277 | -0.083 | **0.825** | 2.16E-02 | **4.12E-02** |
| ENSG00000268049.1 | **ZNF497-AS2** | | lncRNA | **5.497** | 1.401 | 0.422 | **2.641** | 2.16E-02 | **4.13E-02** |
| ENSG00000230426.6 | **LINC01036** | 104169671 | lncRNA | **11.622** | -1.045 | -0.315 | **0.485** | 2.16E-02 | **4.13E-02** |
| ENSG00000215910.9 | **C1orf167** | 284498 | protein_coding | **3.961** | 2.242 | 0.675 | **4.731** | 2.16E-02 | **4.13E-02** |
| ENSG00000173064.15 | **HECTD4** | 283450 | protein_coding | **1364.683** | 0.331 | 0.100 | **1.258** | 2.16E-02 | **4.13E-02** |
| ENSG00000134775.18 | **FHOD3** | 80206 | protein_coding | **282.573** | -0.352 | -0.106 | **0.783** | 2.17E-02 | **4.14E-02** |
| ENSG00000161940.13 | **BCL6B** | 255877 | protein_coding | **7.824** | 1.340 | 0.403 | **2.531** | 2.17E-02 | **4.14E-02** |
| ENSG00000126467.12 | **TSKS** | 60385 | protein_coding | **5.877** | 1.581 | 0.476 | **2.992** | 2.17E-02 | **4.14E-02** |
| ENSG00000132911.7 | **NMUR2** | 56923 | protein_coding | **3.677** | 2.170 | 0.653 | **4.500** | 2.17E-02 | **4.14E-02** |
| ENSG00000122912.17 | **SLC25A16** | 8034 | protein_coding | **272.542** | -0.421 | -0.127 | **0.747** | 2.17E-02 | **4.14E-02** |
| ENSG00000174917.11 | **MICOS13** | 125988 | protein_coding | **914.434** | -0.280 | -0.084 | **0.824** | 2.17E-02 | **4.15E-02** |
| ENSG00000183696.16 | **UPP1** | 7378 | protein_coding | **208.502** | -0.504 | -0.152 | **0.705** | 2.17E-02 | **4.15E-02** |
| ENSG00000293107.1 | **ENSG00000293107** | | lncRNA | **106.744** | 0.411 | 0.124 | **1.330** | 2.18E-02 | **4.16E-02** |
| ENSG00000204745.4 | **ENSG00000204745** | | transcribed_unprc | **81.412** | -0.552 | -0.166 | **0.682** | 2.18E-02 | **4.16E-02** |
| ENSG00000175040.7 | **CHST2** | 9435 | protein_coding | **598.914** | 0.338 | 0.102 | **1.264** | 2.18E-02 | **4.16E-02** |
| ENSG00000310796.1 | **ENSG00000310796** | | protein_coding | **1304.595** | -0.292 | -0.088 | **0.817** | 2.18E-02 | **4.17E-02** |

| | | | | | | | | | |
|---|---|---|---|---|---|---|---|---|---|
| ENSG00000108960.11 | **MMD** | 23531 | protein_coding | **511.175** | 0.282 | 0.085 | **1.216** | 2.18E-02 | **4.17E-02** |
| ENSG00000136404.18 | **TM6SF1** | 53346 | protein_coding | **119.036** | -0.604 | -0.182 | **0.658** | 2.19E-02 | **4.17E-02** |
| ENSG00000213185.9 | **FAM24B** | 196792 | protein_coding | **155.020** | -0.494 | -0.149 | **0.710** | 2.19E-02 | **4.17E-02** |
| ENSG00000203995.12 | **ZYG11A** | 440590 | protein_coding | **119.763** | -0.501 | -0.151 | **0.707** | 2.19E-02 | **4.18E-02** |
| ENSG00000113657.15 | **DPYSL3** | 1809 | protein_coding | **13.011** | 1.213 | 0.365 | **2.317** | 2.19E-02 | **4.19E-02** |
| ENSG00000136895.20 | **GARNL3** | 84253 | protein_coding | **93.503** | 0.590 | 0.178 | **1.505** | 2.20E-02 | **4.19E-02** |
| ENSG00000148835.14 | **TAF5** | 6877 | protein_coding | **152.452** | -0.410 | -0.123 | **0.753** | 2.20E-02 | **4.19E-02** |
| ENSG00000108187.18 | **PBLD** | 64081 | protein_coding | **28.488** | -0.769 | -0.231 | **0.587** | 2.20E-02 | **4.19E-02** |
| ENSG00000293473.2 | **CENPBD1P** | 92806 | lncRNA | **110.535** | 0.357 | 0.107 | **1.281** | 2.20E-02 | **4.20E-02** |
| ENSG00000289406.2 | **ENSG00000289406** | | lncRNA | **14.404** | -0.945 | -0.284 | **0.519** | 2.20E-02 | **4.20E-02** |
| ENSG00000215915.10 | **ATAD3C** | 219293 | protein_coding | **60.152** | 0.857 | 0.258 | **1.811** | 2.20E-02 | **4.20E-02** |
| ENSG00000164054.19 | **SHISA5** | 51246 | protein_coding | **1603.560** | -0.354 | -0.106 | **0.783** | 2.20E-02 | **4.20E-02** |
| ENSG00000255931.3 | **ENSG00000255931** | | lncRNA | **5.041** | 1.985 | 0.598 | **3.958** | 2.21E-02 | **4.21E-02** |
| ENSG00000107957.18 | **SH3PXD2A** | 9644 | protein_coding | **800.163** | -0.269 | -0.081 | **0.830** | 2.21E-02 | **4.21E-02** |
| ENSG00000269947.1 | **ENSG00000269947** | | lncRNA | **20.113** | -0.741 | -0.223 | **0.598** | 2.21E-02 | **4.21E-02** |
| ENSG00000138670.21 | **RASGEF1B** | 153020 | protein_coding | **7.138** | 1.467 | 0.442 | **2.764** | 2.21E-02 | **4.22E-02** |
| ENSG00000099246.21 | **RAB18** | 22931 | protein_coding | **371.425** | -0.328 | -0.099 | **0.796** | 2.22E-02 | **4.22E-02** |
| ENSG00000105221.20 | **AKT2** | 208 | protein_coding | **2311.231** | -0.274 | -0.083 | **0.827** | 2.22E-02 | **4.22E-02** |
| ENSG00000049883.17 | **PTCD2** | 79810 | protein_coding | **252.607** | -0.471 | -0.142 | **0.721** | 2.22E-02 | **4.23E-02** |
| ENSG00000137877.10 | **SPTBN5** | 51332 | protein_coding | **32.091** | 0.869 | 0.262 | **1.827** | 2.22E-02 | **4.23E-02** |
| ENSG00000243147.10 | **MRPL33** | 9553 | protein_coding | **325.717** | -0.286 | -0.086 | **0.820** | 2.22E-02 | **4.23E-02** |
| ENSG00000197614.13 | **MFAP5** | 8076 | protein_coding | **9.069** | 1.407 | 0.423 | **2.651** | 2.22E-02 | **4.24E-02** |
| ENSG00000258982.2 | **YY1-DT** | | lncRNA | **25.101** | 0.757 | 0.228 | **1.690** | 2.23E-02 | **4.24E-02** |
| ENSG00000160229.15 | **ZNF66** | 7617 | protein_coding | **37.101** | -0.670 | -0.202 | **0.628** | 2.23E-02 | **4.24E-02** |
| ENSG00000177294.8 | **FBXO39** | 162517 | protein_coding | **6.956** | 1.535 | 0.462 | **2.898** | 2.23E-02 | **4.24E-02** |
| ENSG00000290427.2 | **ENSG00000290427** | | lncRNA | **24.960** | 0.975 | 0.294 | **1.966** | 2.23E-02 | **4.25E-02** |
| ENSG00000186812.15 | **ZNF397** | 84307 | protein_coding | **400.761** | -0.265 | -0.080 | **0.832** | 2.23E-02 | **4.25E-02** |
| ENSG00000175198.19 | **PCCA** | 5095 | protein_coding | **181.690** | 0.288 | 0.087 | **1.221** | 2.23E-02 | **4.25E-02** |
| ENSG00000197885.13 | **NKIRAS1** | 28512 | protein_coding | **185.128** | 0.385 | 0.116 | **1.306** | 2.24E-02 | **4.26E-02** |
| ENSG00000198692.12 | **EIF1AY** | 9086 | protein_coding | **217.931** | -0.756 | -0.227 | **0.592** | 2.24E-02 | **4.26E-02** |
| ENSG00000006459.13 | **KDM7A** | 80853 | protein_coding | **108.184** | 0.503 | 0.151 | **1.417** | 2.24E-02 | **4.26E-02** |

| | | | | | | | | | |
|---|---|---|---|---|---|---|---|---|---|
| ENSG00000092208.21 | **GEMIN2** | 8487 | protein_coding | **224.040** | -0.414 | -0.125 | **0.751** | 2.24E-02 | **4.27E-02** |
| ENSG00000231475.3 | **IGHV4-31** | | IG_V_gene | **23.062** | 0.986 | 0.297 | **1.981** | 2.25E-02 | **4.27E-02** |
| ENSG00000156273.18 | **BACH1** | 571 | protein_coding | **252.179** | 0.410 | 0.123 | **1.329** | 2.25E-02 | **4.27E-02** |
| ENSG00000151948.14 | **GLT1D1** | 144423 | protein_coding | **166.708** | -0.466 | -0.140 | **0.724** | 2.25E-02 | **4.27E-02** |
| ENSG00000167987.13 | **VPS37C** | 55048 | protein_coding | **334.336** | 0.315 | 0.095 | **1.244** | 2.25E-02 | **4.27E-02** |
| ENSG00000143390.20 | **RFX5** | 5993 | protein_coding | **2491.594** | 0.267 | 0.080 | **1.203** | 2.25E-02 | **4.28E-02** |
| ENSG00000175105.10 | **ZNF654** | 55279 | protein_coding | **159.126** | -0.345 | -0.104 | **0.787** | 2.25E-02 | **4.28E-02** |
| ENSG00000139354.13 | **GAS2L3** | 283431 | protein_coding | **46.811** | -0.797 | -0.240 | **0.575** | 2.25E-02 | **4.28E-02** |
| ENSG00000188807.15 | **TMEM201** | 199953 | protein_coding | **316.582** | -0.365 | -0.110 | **0.777** | 2.26E-02 | **4.29E-02** |
| ENSG00000280441.4 | **ENSG0000(** | 10537950 | lncRNA | **120.642** | -0.536 | -0.161 | **0.690** | 2.26E-02 | **4.29E-02** |
| ENSG00000260498.8 | **ENSG00000260498** | | lncRNA | **6.771** | -1.490 | -0.448 | **0.356** | 2.26E-02 | **4.30E-02** |
| ENSG00000114737.18 | **CISH** | 1154 | protein_coding | **68.288** | -0.517 | -0.156 | **0.699** | 2.26E-02 | **4.30E-02** |
| ENSG00000141293.19 | **SKAP1** | 8631 | protein_coding | **1260.772** | -0.337 | -0.101 | **0.792** | 2.26E-02 | **4.30E-02** |
| ENSG00000115365.15 | **LANCL1** | 10314 | protein_coding | **279.684** | -0.391 | -0.118 | **0.763** | 2.26E-02 | **4.30E-02** |
| ENSG00000120756.15 | **PLS1** | 5357 | protein_coding | **14.032** | 1.090 | 0.328 | **2.129** | 2.27E-02 | **4.31E-02** |
| ENSG00000005238.22 | **ATOSB** | 80256 | protein_coding | **265.741** | 0.338 | 0.102 | **1.264** | 2.27E-02 | **4.31E-02** |
| ENSG00000109016.20 | **DHRS7B** | 25979 | protein_coding | **279.177** | -0.339 | -0.102 | **0.790** | 2.27E-02 | **4.31E-02** |
| ENSG00000230724.10 | **LINC01001** | | lncRNA | **528.710** | 0.307 | 0.093 | **1.237** | 2.27E-02 | **4.32E-02** |
| ENSG00000172530.24 | **BANP** | 54971 | protein_coding | **295.255** | 0.287 | 0.086 | **1.220** | 2.28E-02 | **4.33E-02** |
| ENSG00000031081.13 | **ARHGAP31** | 57514 | protein_coding | **641.972** | 0.403 | 0.121 | **1.322** | 2.28E-02 | **4.33E-02** |
| ENSG00000290933.1 | **ENSG00000290933** | | lncRNA | **91.598** | -0.652 | -0.196 | **0.636** | 2.28E-02 | **4.33E-02** |
| ENSG00000289931.2 | **ENSG00000289931** | | lncRNA | **8.171** | 1.124 | 0.338 | **2.179** | 2.28E-02 | **4.33E-02** |
| ENSG00000130997.17 | **POLN** | 353497 | protein_coding | **18.669** | 1.096 | 0.330 | **2.137** | 2.28E-02 | **4.33E-02** |
| ENSG00000085491.19 | **SLC25A24** | 29957 | protein_coding | **21.801** | 0.750 | 0.226 | **1.682** | 2.28E-02 | **4.33E-02** |
| ENSG00000036549.15 | **ZZZ3** | 26009 | protein_coding | **351.427** | -0.272 | -0.082 | **0.828** | 2.29E-02 | **4.34E-02** |
| ENSG00000105887.13 | **MTPN** | 136319 | protein_coding | **1416.871** | 0.320 | 0.096 | **1.249** | 2.29E-02 | **4.34E-02** |
| ENSG00000131507.13 | **NDFIP1** | 80762 | protein_coding | **228.284** | 0.468 | 0.141 | **1.383** | 2.30E-02 | **4.36E-02** |
| ENSG00000260136.7 | **ENSG0000(** | 12490366 | lncRNA | **4.977** | -1.543 | -0.465 | **0.343** | 2.30E-02 | **4.36E-02** |
| ENSG00000267809.1 | **NDUFV2P1** | | processed_pseud( | **26.345** | -0.646 | -0.194 | **0.639** | 2.30E-02 | **4.37E-02** |
| ENSG00000281100.1 | **ENSG00000281100** | | TEC | **7.132** | 1.220 | 0.367 | **2.330** | 2.30E-02 | **4.37E-02** |
| ENSG00000116337.22 | **AMPD2** | 271 | protein_coding | **897.877** | -0.278 | -0.084 | **0.825** | 2.31E-02 | **4.37E-02** |

| | | | | | | | | | |
|---|---|---|---|---|---|---|---|---|---|
| ENSG00000133195.14 | **SLC39A11** | 201266 | protein_coding | **283.750** | 0.348 | 0.105 | **1.273** | 2.31E-02 | **4.38E-02** |
| ENSG00000181982.22 | **CCDC149** | 91050 | protein_coding | **7.216** | 1.538 | 0.463 | **2.904** | 2.31E-02 | **4.38E-02** |
| ENSG00000267871.7 | **ZNF460-AS** | 10537247( | lncRNA | **25.690** | -0.719 | -0.216 | **0.608** | 2.31E-02 | **4.38E-02** |
| ENSG00000281903.5 | **ASMER1** | 10798729! | lncRNA | **12.232** | 1.261 | 0.380 | **2.397** | 2.32E-02 | **4.39E-02** |
| ENSG00000244187.10 | **TMEM141** | 85014 | protein_coding | **431.817** | 0.291 | 0.088 | **1.224** | 2.32E-02 | **4.39E-02** |
| ENSG00000198925.14 | **ATG9A** | 79065 | protein_coding | **615.880** | -0.295 | -0.089 | **0.815** | 2.32E-02 | **4.39E-02** |
| ENSG00000157181.18 | **ODR4** | 54953 | protein_coding | **319.549** | -0.412 | -0.124 | **0.752** | 2.32E-02 | **4.39E-02** |
| ENSG00000143702.19 | **CEP170** | 9859 | protein_coding | **657.431** | 0.429 | 0.129 | **1.347** | 2.32E-02 | **4.39E-02** |
| ENSG00000266256.4 | **LINC00908** | 284276 | lncRNA | **60.193** | -0.473 | -0.142 | **0.720** | 2.32E-02 | **4.39E-02** |
| ENSG00000165272.18 | **AQP3** | 360 | protein_coding | **49.147** | 0.746 | 0.224 | **1.677** | 2.32E-02 | **4.39E-02** |
| ENSG00000177335.14 | **LY6S-AS1** | 286122 | lncRNA | **34.003** | -0.648 | -0.195 | **0.638** | 2.32E-02 | **4.39E-02** |
| ENSG00000136143.18 | **SUCLA2** | 8803 | protein_coding | **238.661** | -0.476 | -0.143 | **0.719** | 2.32E-02 | **4.39E-02** |
| ENSG00000230457.3 | **PA2G4P4** | | processed_pseud( | **15.895** | 0.805 | 0.242 | **1.747** | 2.32E-02 | **4.39E-02** |
| ENSG00000140474.16 | **ULK3** | 25989 | protein_coding | **851.079** | 0.285 | 0.086 | **1.218** | 2.32E-02 | **4.40E-02** |
| ENSG00000214274.12 | **ANG** | 283 | protein_coding | **56.538** | 0.623 | 0.187 | **1.540** | 2.33E-02 | **4.40E-02** |
| ENSG00000167785.11 | **ZNF558** | 148156 | protein_coding | **775.928** | 0.268 | 0.081 | **1.204** | 2.33E-02 | **4.40E-02** |
| ENSG00000169288.20 | **MRPL1** | 65008 | protein_coding | **262.425** | -0.292 | -0.088 | **0.817** | 2.33E-02 | **4.40E-02** |
| ENSG00000267940.1 | **ENSG00000267940** | | lncRNA | **14.171** | 0.883 | 0.266 | **1.844** | 2.33E-02 | **4.41E-02** |
| ENSG00000172809.15 | **RPL38** | 6169 | protein_coding | **15682.000** | 0.290 | 0.087 | **1.223** | 2.33E-02 | **4.41E-02** |
| ENSG00000239911.5 | **PRKAG2-AS1** | | lncRNA | **23.527** | 1.266 | 0.381 | **2.405** | 2.33E-02 | **4.41E-02** |
| ENSG00000006704.13 | **GTF2IRD1** | 9569 | protein_coding | **216.958** | 0.649 | 0.195 | **1.568** | 2.34E-02 | **4.43E-02** |
| ENSG00000170006.15 | **TMEM154** | 201799 | protein_coding | **262.869** | -0.387 | -0.116 | **0.765** | 2.34E-02 | **4.43E-02** |
| ENSG00000049192.16 | **ADAMTS6** | 11174 | protein_coding | **111.821** | 0.638 | 0.192 | **1.556** | 2.35E-02 | **4.44E-02** |
| ENSG00000184545.13 | **DUSP8** | 1850 | protein_coding | **28.338** | 0.692 | 0.208 | **1.616** | 2.35E-02 | **4.44E-02** |
| ENSG00000198931.13 | **APRT** | 353 | protein_coding | **2827.184** | -0.311 | -0.094 | **0.806** | 2.35E-02 | **4.44E-02** |
| ENSG00000124839.15 | **RAB17** | 64284 | protein_coding | **14.366** | -1.186 | -0.357 | **0.439** | 2.35E-02 | **4.44E-02** |
| ENSG00000224848.5 | **ENSG00000224848** | | lncRNA | **7.092** | 1.977 | 0.595 | **3.936** | 2.35E-02 | **4.44E-02** |
| ENSG00000296923.1 | **ENSG00000296923** | | lncRNA | **5.536** | -1.693 | -0.510 | **0.309** | 2.35E-02 | **4.44E-02** |
| ENSG00000228315.13 | **GUSBP11** | 91316 | lncRNA | **890.967** | 0.373 | 0.112 | **1.295** | 2.35E-02 | **4.44E-02** |
| ENSG00000136052.12 | **SLC41A2** | 84102 | protein_coding | **237.165** | -0.447 | -0.134 | **0.734** | 2.35E-02 | **4.45E-02** |
| ENSG00000289421.2 | **ENSG00000289421** | | lncRNA | **8.472** | -1.055 | -0.318 | **0.481** | 2.36E-02 | **4.45E-02** |

| | | | | | | | | | |
|---|---|---|---|---|---|---|---|---|---|
| ENSG00000273513.3 | **TBC1D3K** | 101060351 | protein_coding | **33.154** | 1.331 | 0.401 | **2.516** | 2.36E-02 | **4.45E-02** |
| ENSG00000076053.14 | **RBM7** | 10179 | protein_coding | **115.128** | -0.614 | -0.185 | **0.653** | 2.37E-02 | **4.47E-02** |
| ENSG00000291153.2 | **STAG3L4** | 64940 | lncRNA | **223.996** | -0.307 | -0.092 | **0.808** | 2.37E-02 | **4.47E-02** |
| ENSG00000166527.10 | **CLEC4D** | 338339 | protein_coding | **8.208** | 1.762 | 0.530 | **3.392** | 2.37E-02 | **4.47E-02** |
| ENSG00000132702.15 | **HAPLN2** | 60484 | protein_coding | **9.170** | 1.117 | 0.336 | **2.169** | 2.37E-02 | **4.48E-02** |
| ENSG00000273345.6 | **ENSG00000273345** | | protein_coding | **1187.005** | -0.277 | -0.083 | **0.826** | 2.37E-02 | **4.48E-02** |
| ENSG00000299227.1 | **ENSG00000299227** | | lncRNA | **20.431** | 0.784 | 0.236 | **1.721** | 2.37E-02 | **4.48E-02** |
| ENSG00000088832.20 | **FKBP1A** | 2280 | protein_coding | **3757.263** | -0.300 | -0.090 | **0.812** | 2.39E-02 | **4.50E-02** |
| ENSG00000147905.20 | **ZCCHC7** | 84186 | protein_coding | **1073.553** | -0.390 | -0.117 | **0.763** | 2.39E-02 | **4.51E-02** |
| ENSG00000142700.14 | **DMRTA2** | 63950 | protein_coding | **3.237** | 2.345 | 0.706 | **5.079** | 2.39E-02 | **4.51E-02** |
| ENSG00000102230.15 | **PCYT1B** | 9468 | protein_coding | **6.781** | 1.401 | 0.422 | **2.641** | 2.39E-02 | **4.51E-02** |
| ENSG00000273297.5 | **LINC03060** | 10537551 | lncRNA | **6.665** | 1.368 | 0.412 | **2.582** | 2.39E-02 | **4.52E-02** |
| ENSG00000107897.22 | **ACBD5** | 91452 | protein_coding | **199.628** | -0.437 | -0.132 | **0.739** | 2.40E-02 | **4.53E-02** |
| ENSG00000168487.21 | **BMP1** | 649 | protein_coding | **45.016** | 0.569 | 0.171 | **1.483** | 2.40E-02 | **4.53E-02** |
| ENSG00000245648.3 | **KLRK1-AS1** | 10192810 | lncRNA | **16.322** | -0.958 | -0.288 | **0.515** | 2.40E-02 | **4.53E-02** |
| ENSG00000261150.4 | **EPPK1** | 83481 | protein_coding | **7.200** | 1.481 | 0.446 | **2.791** | 2.41E-02 | **4.53E-02** |
| ENSG00000175782.13 | **SLC35E3** | 55508 | protein_coding | **293.121** | -0.404 | -0.122 | **0.756** | 2.41E-02 | **4.54E-02** |
| ENSG00000287787.1 | **ENSG00000287787** | | lncRNA | **6.274** | 1.572 | 0.473 | **2.973** | 2.41E-02 | **4.54E-02** |
| ENSG00000258674.5 | **ENSG00000258674** | | protein_coding | **28.211** | 0.799 | 0.240 | **1.739** | 2.41E-02 | **4.54E-02** |
| ENSG00000102043.17 | **MTMR8** | 55613 | protein_coding | **29.165** | -0.856 | -0.258 | **0.552** | 2.41E-02 | **4.54E-02** |
| ENSG00000273271.1 | **ENSG00000273271** | | lncRNA | **12.632** | 0.888 | 0.267 | **1.850** | 2.41E-02 | **4.54E-02** |
| ENSG00000213753.12 | **CENPBD2P** | | transcribed_proce | **33.820** | -0.788 | -0.237 | **0.579** | 2.41E-02 | **4.54E-02** |
| ENSG00000177570.17 | **SAMD12** | 401474 | protein_coding | **139.046** | 0.626 | 0.188 | **1.543** | 2.41E-02 | **4.54E-02** |
| ENSG00000160321.17 | **ZNF208** | 7757 | protein_coding | **30.461** | 1.065 | 0.321 | **2.092** | 2.41E-02 | **4.55E-02** |
| ENSG00000242616.6 | **GNG10** | 2790 | protein_coding | **183.708** | -0.323 | -0.097 | **0.799** | 2.41E-02 | **4.55E-02** |
| ENSG00000164144.18 | **ARFIP1** | 27236 | protein_coding | **178.515** | 0.395 | 0.119 | **1.315** | 2.41E-02 | **4.55E-02** |
| ENSG00000225051.6 | **HMGB3P22** | | transcribed_proce | **4.653** | 2.704 | 0.814 | **6.516** | 2.41E-02 | **4.55E-02** |
| ENSG00000170231.18 | **FABP6** | 2172 | protein_coding | **28.823** | 0.823 | 0.248 | **1.769** | 2.42E-02 | **4.55E-02** |
| ENSG00000131015.7 | **ULBP2** | 80328 | protein_coding | **4.922** | 2.290 | 0.689 | **4.889** | 2.42E-02 | **4.55E-02** |
| ENSG00000282851.2 | **BISPR** | 10522169 | lncRNA | **464.252** | -0.267 | -0.080 | **0.831** | 2.42E-02 | **4.56E-02** |
| ENSG00000128438.11 | **TBC1D27P** | | transcribed_unpro | **66.679** | 0.651 | 0.196 | **1.570** | 2.43E-02 | **4.57E-02** |

| | | | | | | | | | |
|---|---|---|---|---|---|---|---|---|---|
| ENSG00000182544.10 | **SLC61A1** | 84975 | protein_coding | **224.529** | -0.269 | -0.081 | **0.830** | 2.43E-02 | **4.57E-02** |
| ENSG00000104883.10 | **PEX11G** | 92960 | protein_coding | **25.576** | 0.654 | 0.197 | **1.574** | 2.43E-02 | **4.57E-02** |
| ENSG00000159339.14 | **PADI4** | 23569 | protein_coding | **10.328** | -1.179 | -0.355 | **0.442** | 2.43E-02 | **4.58E-02** |
| ENSG00000115828.19 | **QPCT** | 25797 | protein_coding | **19.370** | 0.789 | 0.237 | **1.727** | 2.44E-02 | **4.58E-02** |
| ENSG00000271425.10 | **NBPF10** | 10013240( | protein_coding | **37.036** | 0.814 | 0.245 | **1.758** | 2.44E-02 | **4.58E-02** |
| ENSG00000162757.6 | **C1orf74** | 148304 | protein_coding | **135.366** | 0.324 | 0.098 | **1.252** | 2.44E-02 | **4.59E-02** |
| ENSG00000188487.14 | **INSC** | 387755 | protein_coding | **6.119** | 1.375 | 0.414 | **2.593** | 2.44E-02 | **4.59E-02** |
| ENSG00000080298.18 | **RFX3** | 5991 | protein_coding | **246.022** | -0.441 | -0.133 | **0.737** | 2.44E-02 | **4.60E-02** |
| ENSG00000237310.4 | **LINC03011** | 10028909{ | lncRNA | **56.310** | 0.531 | 0.160 | **1.445** | 2.45E-02 | **4.60E-02** |
| ENSG00000171659.18 | **GPR34** | 2857 | protein_coding | **4.821** | -1.792 | -0.540 | **0.289** | 2.45E-02 | **4.61E-02** |
| ENSG00000286011.2 | **ENSG00000286011** | | lncRNA | **12.689** | 0.943 | 0.284 | **1.923** | 2.45E-02 | **4.61E-02** |
| ENSG00000298862.1 | **ENSG00000298862** | | lncRNA | **6.572** | 1.270 | 0.382 | **2.412** | 2.46E-02 | **4.62E-02** |
| ENSG00000221944.9 | **TIGD1** | 200765 | protein_coding | **107.387** | -0.408 | -0.123 | **0.754** | 2.46E-02 | **4.63E-02** |
| ENSG00000140743.10 | **CDR2** | 1039 | protein_coding | **399.912** | -0.300 | -0.090 | **0.812** | 2.46E-02 | **4.63E-02** |
| ENSG00000225331.3 | **LINC01678** | | lncRNA | **22.070** | 0.723 | 0.218 | **1.650** | 2.47E-02 | **4.64E-02** |
| ENSG00000250548.8 | **LINC01303** | 10192778( | lncRNA | **4.410** | 2.505 | 0.754 | **5.678** | 2.47E-02 | **4.64E-02** |
| ENSG00000134716.13 | **CYP2J2** | 1573 | protein_coding | **34.517** | 0.720 | 0.217 | **1.647** | 2.47E-02 | **4.64E-02** |
| ENSG00000171574.20 | **ZNF584** | 201514 | protein_coding | **111.362** | -0.363 | -0.109 | **0.778** | 2.47E-02 | **4.64E-02** |
| ENSG00000206190.14 | **ATP10A** | 57194 | protein_coding | **9.880** | -1.340 | -0.403 | **0.395** | 2.48E-02 | **4.66E-02** |
| ENSG00000174332.8 | **GLIS1** | 148979 | protein_coding | **5.534** | 1.988 | 0.598 | **3.966** | 2.48E-02 | **4.66E-02** |
| ENSG00000139921.15 | **TMX1** | 81542 | protein_coding | **324.082** | -0.363 | -0.109 | **0.777** | 2.48E-02 | **4.66E-02** |
| ENSG00000264207.1 | **ENSG00000264207** | | lncRNA | **65.041** | 0.478 | 0.144 | **1.392** | 2.49E-02 | **4.67E-02** |
| ENSG00000103356.18 | **EARS2** | 124454 | protein_coding | **501.462** | -0.365 | -0.110 | **0.776** | 2.49E-02 | **4.67E-02** |
| ENSG00000096401.10 | **CDC5L** | 988 | protein_coding | **552.371** | -0.308 | -0.093 | **0.808** | 2.49E-02 | **4.68E-02** |
| ENSG00000159216.22 | **RUNX1** | 861 | protein_coding | **238.733** | 0.358 | 0.108 | **1.282** | 2.49E-02 | **4.68E-02** |
| ENSG00000125462.21 | **MIR9-1HG** | 10485 | lncRNA | **50.462** | 0.624 | 0.188 | **1.541** | 2.50E-02 | **4.69E-02** |
| ENSG00000151967.20 | **SCHIP1** | 29970 | protein_coding | **76.727** | 0.885 | 0.266 | **1.847** | 2.50E-02 | **4.70E-02** |
| ENSG00000088726.17 | **TMEM40** | 55287 | protein_coding | **15.198** | 1.046 | 0.315 | **2.064** | 2.50E-02 | **4.70E-02** |
| ENSG00000266916.10 | **ZNF793-AS** | 10192772( | lncRNA | **22.757** | -0.729 | -0.220 | **0.603** | 2.51E-02 | **4.70E-02** |
| ENSG00000197530.15 | **MIB2** | 142678 | protein_coding | **1064.016** | 0.385 | 0.116 | **1.306** | 2.51E-02 | **4.70E-02** |
| ENSG00000171497.7 | **PPID** | 5481 | protein_coding | **636.206** | -0.311 | -0.094 | **0.806** | 2.51E-02 | **4.71E-02** |

| | | | | | | | | | |
|---|---|---|---|---|---|---|---|---|---|
| ENSG00000183207.17 | **RUVBL2** | 10856 | protein_coding | **1742.123** | -0.315 | -0.095 | **0.804** | 2.51E-02 | **4.71E-02** |
| ENSG00000114853.16 | **ZBTB47** | 92999 | protein_coding | **76.019** | 0.510 | 0.153 | **1.424** | 2.52E-02 | **4.72E-02** |
| ENSG00000258454.2 | **ENSG00000258454** | | lncRNA | **5.564** | 1.439 | 0.433 | **2.712** | 2.52E-02 | **4.72E-02** |
| ENSG00000218891.7 | **ZNF579** | 163033 | protein_coding | **362.542** | -0.322 | -0.097 | **0.800** | 2.52E-02 | **4.72E-02** |
| ENSG00000268658.5 | **LINC00664** | 400680 | lncRNA | **11.251** | -1.471 | -0.443 | **0.361** | 2.52E-02 | **4.73E-02** |
| ENSG00000214113.13 | **LYRM4** | 57128 | protein_coding | **556.443** | -0.284 | -0.085 | **0.821** | 2.52E-02 | **4.73E-02** |
| ENSG00000216895.9 | **RBM33-DT** | 100506302 | lncRNA | **136.328** | 0.356 | 0.107 | **1.280** | 2.52E-02 | **4.73E-02** |
| ENSG00000082458.14 | **DLG3** | 1741 | protein_coding | **324.973** | 0.395 | 0.119 | **1.315** | 2.53E-02 | **4.74E-02** |
| ENSG00000138381.11 | **ASNSD1** | 54529 | protein_coding | **436.716** | -0.280 | -0.084 | **0.824** | 2.53E-02 | **4.74E-02** |
| ENSG00000151239.16 | **TWF1** | 5756 | protein_coding | **174.197** | -0.386 | -0.116 | **0.765** | 2.53E-02 | **4.75E-02** |
| ENSG00000180871.10 | **CXCR2** | 3579 | protein_coding | **6.678** | -1.854 | -0.558 | **0.277** | 2.53E-02 | **4.75E-02** |
| ENSG00000256235.5 | **SMIM3** | 85027 | protein_coding | **217.798** | -0.428 | -0.129 | **0.743** | 2.53E-02 | **4.75E-02** |
| ENSG00000160051.14 | **IQCC** | 55721 | protein_coding | **81.553** | -0.379 | -0.114 | **0.769** | 2.54E-02 | **4.75E-02** |
| ENSG00000288988.2 | **ENSG00000288988** | | lncRNA | **30.145** | -0.746 | -0.225 | **0.596** | 2.54E-02 | **4.75E-02** |
| ENSG00000108883.15 | **EFTUD2** | 9343 | protein_coding | **1578.724** | -0.302 | -0.091 | **0.811** | 2.54E-02 | **4.75E-02** |
| ENSG00000140459.20 | **CYP11A1** | 1583 | protein_coding | **28.883** | -0.850 | -0.256 | **0.555** | 2.54E-02 | **4.76E-02** |
| ENSG00000295213.1 | **ENSG00000295213** | | lncRNA | **7.091** | 1.285 | 0.387 | **2.437** | 2.55E-02 | **4.77E-02** |
| ENSG00000110427.19 | **KIAA1549L** | 25758 | protein_coding | **126.988** | 0.571 | 0.172 | **1.486** | 2.55E-02 | **4.77E-02** |
| ENSG00000188779.12 | **SKOR1** | 390598 | protein_coding | **13.224** | 0.876 | 0.264 | **1.836** | 2.55E-02 | **4.77E-02** |
| ENSG00000157869.18 | **RAB28** | 9364 | protein_coding | **164.771** | -0.486 | -0.146 | **0.714** | 2.55E-02 | **4.77E-02** |
| ENSG00000293352.2 | **RPLP0P2** | 113157 | lncRNA | **5.756** | 1.936 | 0.583 | **3.825** | 2.55E-02 | **4.77E-02** |
| ENSG00000163520.16 | **FBLN2** | 2199 | protein_coding | **4.969** | 1.549 | 0.466 | **2.926** | 2.55E-02 | **4.77E-02** |
| ENSG00000154127.11 | **UBASH3B** | 84959 | protein_coding | **28.364** | -0.827 | -0.249 | **0.564** | 2.55E-02 | **4.78E-02** |
| ENSG00000250673.4 | **REELD1** | 345051 | protein_coding | **38.298** | 0.742 | 0.223 | **1.673** | 2.56E-02 | **4.78E-02** |
| ENSG00000184163.4 | **C1QTNF12** | 388581 | protein_coding | **51.589** | 0.683 | 0.206 | **1.606** | 2.56E-02 | **4.78E-02** |
| ENSG00000295161.1 | **ENSG00000295161** | | lncRNA | **6.020** | 1.430 | 0.431 | **2.695** | 2.56E-02 | **4.78E-02** |
| ENSG00000105829.14 | **BET1** | 10282 | protein_coding | **291.071** | -0.290 | -0.087 | **0.818** | 2.56E-02 | **4.78E-02** |
| ENSG00000297268.1 | **ENSG00000297268** | | lncRNA | **12.085** | 0.927 | 0.279 | **1.901** | 2.56E-02 | **4.78E-02** |
| ENSG00000095574.14 | **IKZF5** | 64376 | protein_coding | **242.522** | -0.397 | -0.120 | **0.759** | 2.56E-02 | **4.78E-02** |
| ENSG00000186300.14 | **ZNF555** | 148254 | protein_coding | **74.471** | 0.438 | 0.132 | **1.355** | 2.56E-02 | **4.79E-02** |
| ENSG00000266998.3 | **ENSG00000266998** | | lncRNA | **3.830** | 1.998 | 0.601 | **3.994** | 2.57E-02 | **4.80E-02** |

| | | | | | | | | | |
|---|---|---|---|---|---|---|---|---|---|
| ENSG00000293063.1 | **ENSG00000293063** | | lncRNA | **3.863** | 2.203 | 0.663 | **4.603** | 2.57E-02 | **4.81E-02** |
| ENSG00000253317.1 | **ENSG00000253317** | | lncRNA | **21.581** | 0.710 | 0.214 | **1.635** | 2.57E-02 | **4.81E-02** |
| ENSG00000204217.17 | **BMPR2** | 659 | protein_coding | **337.530** | 0.386 | 0.116 | **1.306** | 2.58E-02 | **4.82E-02** |
| ENSG00000103150.9 | **MLYCD** | 23417 | protein_coding | **159.565** | -0.349 | -0.105 | **0.785** | 2.58E-02 | **4.83E-02** |
| ENSG00000110172.14 | **CHORDC1** | 26973 | protein_coding | **451.898** | -0.323 | -0.097 | **0.799** | 2.59E-02 | **4.83E-02** |
| ENSG00000225556.3 | **C2CD4D** | 10019104( | protein_coding | **9.836** | 1.045 | 0.314 | **2.063** | 2.59E-02 | **4.83E-02** |
| ENSG00000064393.18 | **HIPK2** | 28996 | protein_coding | **889.318** | -0.342 | -0.103 | **0.789** | 2.59E-02 | **4.83E-02** |
| ENSG00000273428.5 | **ENSG00000273428** | | lncRNA | **22.311** | 0.710 | 0.214 | **1.636** | 2.59E-02 | **4.84E-02** |
| ENSG00000249855.1 | **EEF1A1P19** | | processed_pseudo | **14.270** | -1.050 | -0.316 | **0.483** | 2.60E-02 | **4.85E-02** |
| ENSG00000253704.5 | **VIRMA-DT** | 10192697 | lncRNA | **41.848** | 0.451 | 0.136 | **1.367** | 2.60E-02 | **4.85E-02** |
| ENSG00000204616.13 | **TRIM31** | 11074 | protein_coding | **12.291** | 1.311 | 0.395 | **2.480** | 2.60E-02 | **4.85E-02** |
| ENSG00000293757.1 | **ENSG00000293757** | | lncRNA | **5.458** | 1.706 | 0.513 | **3.262** | 2.60E-02 | **4.86E-02** |
| ENSG00000104412.10 | **EMC2** | 9694 | protein_coding | **526.857** | -0.304 | -0.091 | **0.810** | 2.60E-02 | **4.86E-02** |
| ENSG00000227492.3 | **CHUK-DT** | | lncRNA | **4.283** | 1.680 | 0.506 | **3.205** | 2.60E-02 | **4.86E-02** |
| ENSG00000138587.8 | **MNS1** | 55329 | protein_coding | **23.737** | -0.747 | -0.225 | **0.596** | 2.60E-02 | **4.86E-02** |
| ENSG00000141527.20 | **CARD14** | 79092 | protein_coding | **6.268** | 1.812 | 0.545 | **3.510** | 2.61E-02 | **4.86E-02** |
| ENSG00000120162.12 | **MOB3B** | 79817 | protein_coding | **75.231** | 0.710 | 0.214 | **1.636** | 2.61E-02 | **4.86E-02** |
| ENSG00000054654.21 | **SYNE2** | 23224 | protein_coding | **2059.540** | 0.307 | 0.092 | **1.237** | 2.61E-02 | **4.86E-02** |
| ENSG00000197180.5 | **ATP6AP1-DT** | | lncRNA | **69.870** | 0.575 | 0.173 | **1.490** | 2.61E-02 | **4.87E-02** |
| ENSG00000162408.13 | **NOL9** | 79707 | protein_coding | **319.506** | -0.267 | -0.080 | **0.831** | 2.61E-02 | **4.87E-02** |
| ENSG00000279821.1 | **ENSG00000279821** | | TEC | **6.944** | 1.425 | 0.429 | **2.685** | 2.61E-02 | **4.87E-02** |
| ENSG00000188153.16 | **COL4A5** | 1287 | protein_coding | **12.942** | 1.309 | 0.394 | **2.479** | 2.62E-02 | **4.88E-02** |
| ENSG00000251409.3 | **ENSG00000251409** | | lncRNA | **9.909** | 1.019 | 0.307 | **2.027** | 2.62E-02 | **4.89E-02** |
| ENSG00000174738.16 | **NR1D2** | 9975 | protein_coding | **103.728** | 0.468 | 0.141 | **1.383** | 2.62E-02 | **4.89E-02** |
| ENSG00000184937.18 | **WT1** | 7490 | protein_coding | **7.336** | 1.360 | 0.409 | **2.567** | 2.63E-02 | **4.89E-02** |
| ENSG00000240065.11 | **PSMB9** | 5698 | protein_coding | **4028.947** | 0.277 | 0.083 | **1.212** | 2.63E-02 | **4.89E-02** |
| ENSG00000091592.18 | **NLRP1** | 22861 | protein_coding | **2708.022** | 0.304 | 0.092 | **1.235** | 2.63E-02 | **4.90E-02** |
| ENSG00000064547.16 | **LPAR2** | 9170 | protein_coding | **312.696** | -0.376 | -0.113 | **0.771** | 2.63E-02 | **4.90E-02** |
| ENSG00000077157.25 | **PPP1R12B** | 4660 | protein_coding | **203.356** | 0.378 | 0.114 | **1.299** | 2.63E-02 | **4.90E-02** |
| ENSG00000246174.11 | **KCTD21-AS** | 10028938 | lncRNA | **38.052** | 0.545 | 0.164 | **1.459** | 2.63E-02 | **4.91E-02** |
| ENSG00000232112.5 | **TMA7** | 51372 | protein_coding | **4201.189** | -0.306 | -0.092 | **0.809** | 2.64E-02 | **4.91E-02** |

| | | | | | | | | | |
|---|---|---|---|---|---|---|---|---|---|
| ENSG00000116729.16 | **WLS** | 79971 | protein_coding | **5.150** | 1.628 | 0.490 | **3.091** | 2.64E-02 | **4.91E-02** |
| ENSG00000181744.10 | **DIPK2A** | 205428 | protein_coding | **114.006** | -0.353 | -0.106 | **0.783** | 2.64E-02 | **4.92E-02** |
| ENSG00000278383.1 | **ENSG00000278383** | | lncRNA | **9.913** | 1.077 | 0.324 | **2.110** | 2.64E-02 | **4.92E-02** |
| ENSG00000204475.11 | **NCR3** | 259197 | protein_coding | **190.269** | 0.478 | 0.144 | **1.392** | 2.64E-02 | **4.92E-02** |
| ENSG00000105523.6 | **FAM83E** | 54854 | protein_coding | **8.170** | 1.090 | 0.328 | **2.129** | 2.65E-02 | **4.93E-02** |
| ENSG00000205085.13 | **GARIN1A** | 346653 | protein_coding | **37.745** | 0.641 | 0.193 | **1.559** | 2.65E-02 | **4.94E-02** |
| ENSG00000243896.5 | **OR2A7** | 401427 | protein_coding | **7.362** | -1.587 | -0.478 | **0.333** | 2.65E-02 | **4.94E-02** |
| ENSG00000186566.15 | **GPATCH8** | 23131 | protein_coding | **1067.139** | 0.304 | 0.092 | **1.235** | 2.66E-02 | **4.94E-02** |
| ENSG00000100478.19 | **AP4S1** | 11154 | protein_coding | **112.529** | -0.562 | -0.169 | **0.677** | 2.66E-02 | **4.94E-02** |
| ENSG00000267317.2 | **APC2-AS1** | | lncRNA | **98.790** | 0.339 | 0.102 | **1.265** | 2.66E-02 | **4.95E-02** |
| ENSG00000164916.13 | **FOXK1** | 221937 | protein_coding | **626.558** | 0.308 | 0.093 | **1.238** | 2.66E-02 | **4.95E-02** |
| ENSG00000258469.3 | **CHMP4BP1** | | protein_coding | **16.908** | -0.858 | -0.258 | **0.552** | 2.66E-02 | **4.95E-02** |
| ENSG00000289740.2 | **TALAM1** | 10913657 | lncRNA | **23.579** | 0.842 | 0.254 | **1.793** | 2.66E-02 | **4.95E-02** |
| ENSG00000259917.1 | **HNRNPLP2** | | transcribed_unpro | **4.330** | 1.936 | 0.583 | **3.828** | 2.68E-02 | **4.97E-02** |
| ENSG00000236069.3 | **ENSG00000236069** | | lncRNA | **9.925** | 1.192 | 0.359 | **2.285** | 2.69E-02 | **4.99E-02** |
| ENSG00000239445.7 | **ST3GAL6-AS1** | | lncRNA | **47.445** | 0.699 | 0.210 | **1.623** | 2.69E-02 | **5.00E-02** |
| ENSG00000227252.3 | **COPS8-DT** | 105373951 | lncRNA | **34.445** | 0.562 | 0.169 | **1.477** | 2.69E-02 | **5.00E-02** |
| ENSG00000166192.17 | **SENP8** | 123228 | protein_coding | **42.391** | 0.756 | 0.228 | **1.689** | 2.69E-02 | **5.00E-02** |
