## Supplemental Table 2 for "RNA-sequencing of butyrate-treated human lymphoblastoid cell lines shows dysregulated genes of multiple disease-related pathways"

| Gene_symbol | NCBI_Gene_ID | Ensembl_gene_ID | log2 FD (butyrate/ control) | Linear FD (butyrate/ control) | padj | Direction |
|---|---|---|---|---|---|---|
| **SEPTIN4** | 5414 | ENSG00000108387 | 4.962 | **31.162** | **3.38E-61** | up |
| **DTYMK** | 1841 | ENSG00000168393 | -1.617 | **0.326** | **1.24E-58** | down |
| **MTFR2** | 113115 | ENSG00000146410 | -2.133 | **0.228** | **2.32E-41** | down |
| **FAM136A** | 84908 | ENSG00000035141 | -1.296 | **0.407** | **1.01E-38** | down |
| **SFXN1** | 94081 | ENSG00000164466 | -1.219 | **0.430** | **4.47E-38** | down |
| **TRIAP1** | 51499 | ENSG00000170855 | -1.350 | **0.392** | **1.39E-32** | down |
| **PAICS** | 10606 | ENSG00000128050 | -1.490 | **0.356** | **1.02E-31** | down |
| **MMAB** | 326625 | ENSG00000139428 | -1.652 | **0.318** | **1.35E-31** | down |
| **MRPL15** | 29088 | ENSG00000137547 | -1.116 | **0.461** | **1.48E-31** | down |
| **AK4** | 205 | ENSG00000162433 | -1.593 | **0.332** | **7.15E-30** | down |
| **POLQ** | 10721 | ENSG00000051341 | -1.583 | **0.334** | **1.45E-28** | down |
| **IVD** | 3712 | ENSG00000128928 | 0.902 | **1.868** | **3.73E-28** | up |
| **SFXN3** | 81855 | ENSG00000107819 | 1.441 | **2.715** | **8.05E-27** | up |
| **GPX1** | 2876 | ENSG00000233276 | -1.085 | **0.471** | **1.08E-26** | down |
| **NNT** | 23530 | ENSG00000112992 | -1.264 | **0.417** | **7.88E-26** | down |
| **MRPL11** | 65003 | ENSG00000174547 | -1.119 | **0.461** | **2.47E-25** | down |
| **ABCD1** | 215 | ENSG00000101986 | 1.508 | **2.845** | **6.95E-25** | up |
| **OXCT1** | 5019 | ENSG00000083720 | -1.383 | **0.383** | **1.26E-24** | down |
| **CYC1** | 1537 | ENSG00000179091 | -1.192 | **0.438** | **1.33E-24** | down |
| **PIF1** | 80119 | ENSG00000140451 | -1.430 | **0.371** | **1.95E-24** | down |
| **AIFM2** | 84883 | ENSG00000042286 | -1.261 | **0.417** | **2.06E-24** | down |
| **FAM210B** | 116151 | ENSG00000124098 | 1.863 | **3.639** | **2.62E-24** | up |
| **PDK3** | 5165 | ENSG00000067992 | -1.497 | **0.354** | **7.60E-24** | down |
| **ALDH1L2** | 160428 | ENSG00000136010 | -2.199 | **0.218** | **1.01E-23** | down |
| **PDK2** | 5164 | ENSG00000005882 | 1.614 | **3.060** | **1.39E-23** | up |
| **GLYCTK** | 132158 | ENSG00000168237 | -2.107 | **0.232** | **3.40E-23** | down |
| **KMO** | 8564 | ENSG00000117009 | -2.827 | **0.141** | **2.35E-22** | down |
| **TRMT2B** | 79979 | ENSG00000188917 | -1.316 | **0.402** | **2.47E-22** | down |
| **COX6B2** | 125965 | ENSG00000160471 | 4.833 | **28.511** | **2.92E-22** | up |
| **LDHD** | 197257 | ENSG00000166816 | 4.523 | **22.983** | **4.58E-22** | up |
| **DHRS1** | 115817 | ENSG00000157379 | 1.022 | **2.031** | **5.58E-22** | up |
| **MTFP1** | 51537 | ENSG00000242114 | -1.545 | **0.343** | **1.32E-21** | down |
| **MTHFD1L** | 25902 | ENSG00000120254 | -1.241 | **0.423** | **2.30E-21** | down |
| **METTL8** | 79828 | ENSG00000123600 | -1.260 | **0.418** | **3.71E-21** | down |
| **MRPL35** | 51318 | ENSG00000132313 | -1.080 | **0.473** | **3.89E-21** | down |
| **ATAD3A** | 55210 | ENSG00000197785 | -0.961 | **0.514** | **4.14E-21** | down |
| **MRPS16** | 51021 | ENSG00000182180 | -1.067 | **0.477** | **4.65E-21** | down |
| **MRPS7** | 51081 | ENSG00000125445 | -0.913 | **0.531** | **4.49E-20** | down |
| **SHMT2** | 6472 | ENSG00000182199 | -1.406 | **0.377** | **4.56E-20** | down |
| **RNASEH1** | 246243 | ENSG00000171865 | -1.005 | **0.498** | **4.81E-20** | down |
| **PLD6** | 201164 | ENSG00000179598 | -1.356 | **0.391** | **4.82E-20** | down |
| **LONP1** | 9361 | ENSG00000196365 | -1.028 | **0.491** | **9.09E-20** | down |
| **RECQL4** | 9401 | ENSG00000160957 | -2.217 | **0.215** | **1.52E-19** | down |
| **NIT1** | 4817 | ENSG00000158793 | 0.863 | **1.818** | **5.29E-19** | up |

| | | | | | | |
|---|---|---|---|---|---|---|
| **ECSIT** | 51295 | ENSG00000130159 | -1.226 | **0.428** | **6.29E-19** | down |
| **PPIF** | 10105 | ENSG00000108179 | -1.067 | **0.477** | **7.19E-19** | down |
| **PYCR1** | 5831 | ENSG00000183010 | -1.180 | **0.441** | **1.04E-18** | down |
| **MGST1** | 4257 | ENSG00000008394 | -1.489 | **0.356** | **1.09E-18** | down |
| **CPT1C** | 126129 | ENSG00000169169 | 4.701 | **26.019** | **1.23E-18** | up |
| **PINK1** | 65018 | ENSG00000158828 | 1.777 | **3.427** | **3.27E-18** | up |
| **CLPP** | 8192 | ENSG00000125656 | -0.930 | **0.525** | **5.51E-18** | down |
| **ALDH2** | 217 | ENSG00000111275 | 1.666 | **3.174** | **6.92E-18** | up |
| **MRM3** | 55178 | ENSG00000171861 | -0.911 | **0.532** | **9.70E-18** | down |
| **SLC25A5** | 292 | ENSG00000005022 | -1.149 | **0.451** | **1.23E-17** | down |
| **YARS2** | 51067 | ENSG00000139131 | -1.246 | **0.421** | **1.62E-17** | down |
| **MRPS30** | 10884 | ENSG00000112996 | -0.980 | **0.507** | **1.70E-17** | down |
| **MRPL37** | 51253 | ENSG00000116221 | -1.030 | **0.490** | **1.77E-17** | down |
| **ECHDC3** | 79746 | ENSG00000134463 | 1.423 | **2.682** | **1.85E-17** | up |
| **RCC1L** | 81554 | ENSG00000274523 | -1.056 | **0.481** | **5.86E-17** | down |
| **CHCHD4** | 131474 | ENSG00000163528 | -1.234 | **0.425** | **7.12E-17** | down |
| **BDH1** | 622 | ENSG00000161267 | -0.871 | **0.547** | **1.35E-16** | down |
| **MPC2** | 25874 | ENSG00000143158 | -0.799 | **0.575** | **2.01E-16** | down |
| **ACOT7** | 11332 | ENSG00000097021 | -1.274 | **0.413** | **3.26E-16** | down |
| **AFG3L2** | 10939 | ENSG00000141385 | -0.666 | **0.630** | **4.47E-16** | down |
| **PDSS1** | 23590 | ENSG00000148459 | -1.348 | **0.393** | **5.85E-16** | down |
| **MRPL50** | 54534 | ENSG00000136897 | -1.103 | **0.465** | **8.17E-16** | down |
| **FAHD2A** | 51011 | ENSG00000115042 | -0.806 | **0.572** | **8.25E-16** | down |
| **MRPL17** | 63875 | ENSG00000158042 | -0.996 | **0.501** | **1.20E-15** | down |
| **NDUFB2** | 4708 | ENSG00000090266 | -0.938 | **0.522** | **2.90E-15** | down |
| **NDUFS1** | 4719 | ENSG00000023228 | -0.734 | **0.601** | **2.92E-15** | down |
| **ADHFE1** | 137872 | ENSG00000147576 | 1.978 | **3.938** | **3.31E-15** | up |
| **HPDL** | 84842 | ENSG00000186603 | -1.580 | **0.335** | **3.85E-15** | down |
| **PCK2** | 5106 | ENSG00000100889 | -1.297 | **0.407** | **3.95E-15** | down |
| **ECI1** | 1632 | ENSG00000167969 | -0.879 | **0.544** | **5.15E-15** | down |
| **C1QBP** | 708 | ENSG00000108561 | -0.888 | **0.540** | **6.13E-15** | down |
| **RPIA** | 22934 | ENSG00000153574 | -0.842 | **0.558** | **6.19E-15** | down |
| **SUCLG2** | 8801 | ENSG00000172340 | -0.864 | **0.549** | **9.48E-15** | down |
| **TIMM10** | 26519 | ENSG00000134809 | -0.887 | **0.541** | **1.40E-14** | down |
| **MRPL12** | 6182 | ENSG00000262814 | -1.022 | **0.492** | **1.43E-14** | down |
| **ARMCX2** | 9823 | ENSG00000184867 | 1.196 | **2.291** | **2.46E-14** | up |
| **DNA2** | 1763 | ENSG00000138346 | -1.460 | **0.363** | **2.89E-14** | down |
| **MRPL39** | 54148 | ENSG00000154719 | -0.715 | **0.609** | **3.02E-14** | down |
| **SLC25A35** | 399512 | ENSG00000125434 | 1.154 | **2.226** | **3.04E-14** | up |
| **MRM1** | 79922 | ENSG00000278619 | -1.061 | **0.479** | **3.32E-14** | down |
| **TIMM21** | 29090 | ENSG00000075336 | -1.080 | **0.473** | **4.25E-14** | down |
| **TMEM126A** | 84233 | ENSG00000171202 | -1.105 | **0.465** | **4.47E-14** | down |
| **SLC25A27** | 9481 | ENSG00000153291 | 5.076 | **33.728** | **4.80E-14** | up |
| **BPHL** | 670 | ENSG00000137274 | 1.308 | **2.476** | **4.80E-14** | up |
| **TIMMDC1** | 51300 | ENSG00000113845 | 0.636 | **1.554** | **5.93E-14** | up |
| **GRHPR** | 9380 | ENSG00000137106 | 0.835 | **1.784** | **9.44E-14** | up |
| **GLUD1** | 2746 | ENSG00000148672 | -0.643 | **0.641** | **1.13E-13** | down |
| **BAX** | 581 | ENSG00000087088 | -0.880 | **0.544** | **1.35E-13** | down |

| | | | | | | |
|---|---|---|---|---|---|---|
| **ACAD9** | 28976 | ENSG00000177646 | -0.819 | **0.567** | **2.28E-13** | down |
| **HSPA9** | 3313 | ENSG00000113013 | -0.957 | **0.515** | **2.52E-13** | down |
| **RAB24** | 53917 | ENSG00000169228 | 0.911 | **1.880** | **2.54E-13** | up |
| **GARS1** | 2617 | ENSG00000106105 | -1.257 | **0.418** | **3.92E-13** | down |
| **UNG** | 7374 | ENSG00000076248 | -1.252 | **0.420** | **5.09E-13** | down |
| **BOLA3** | 388962 | ENSG00000163170 | -1.086 | **0.471** | **6.04E-13** | down |
| **FKBP10** | 60681 | ENSG00000141756 | 2.029 | **4.080** | **6.63E-13** | up |
| **UQCRC1** | 7384 | ENSG00000010256 | -0.872 | **0.546** | **7.19E-13** | down |
| **GRPEL2** | 134266 | ENSG00000164284 | -1.192 | **0.438** | **8.56E-13** | down |
| **PCCB** | 5096 | ENSG00000114054 | -0.795 | **0.576** | **1.02E-12** | down |
| **MSRA** | 4482 | ENSG00000175806 | -1.134 | **0.456** | **1.14E-12** | down |
| **HADH** | 3033 | ENSG00000138796 | -0.737 | **0.600** | **1.30E-12** | down |
| **PDK1** | 5163 | ENSG00000152256 | -0.800 | **0.574** | **1.42E-12** | down |
| **C8orf82** | 414919 | ENSG00000213563 | -0.988 | **0.504** | **1.52E-12** | down |
| **TRMT1** | 55621 | ENSG00000104907 | -1.139 | **0.454** | **1.77E-12** | down |
| **TXNRD2** | 10587 | ENSG00000184470 | -1.079 | **0.473** | **1.80E-12** | down |
| **TACO1** | 51204 | ENSG00000136463 | -0.824 | **0.565** | **1.89E-12** | down |
| **IDH2** | 3418 | ENSG00000182054 | -0.758 | **0.591** | **2.17E-12** | down |
| **MRPS6** | 64968 | ENSG00000243927 | -0.595 | **0.662** | **2.32E-12** | down |
| **GFER** | 2671 | ENSG00000127554 | -0.739 | **0.599** | **2.63E-12** | down |
| **TK2** | 7084 | ENSG00000166548 | 1.282 | **2.431** | **3.18E-12** | up |
| **NAT8L** | 339983 | ENSG00000185818 | 3.251 | **9.524** | **1.07E-11** | up |
| **COA6** | 388753 | ENSG00000168275 | -0.856 | **0.552** | **1.24E-11** | down |
| **SDSL** | 113675 | ENSG00000139410 | 1.535 | **2.898** | **1.35E-11** | up |
| **ACSS1** | 84532 | ENSG00000154930 | -0.668 | **0.629** | **1.47E-11** | down |
| **GRPEL1** | 80273 | ENSG00000109519 | -0.677 | **0.626** | **1.63E-11** | down |
| **NDUFS3** | 4722 | ENSG00000213619 | -0.934 | **0.523** | **2.09E-11** | down |
| **SLC8B1** | 80024 | ENSG00000089060 | 0.950 | **1.931** | **2.23E-11** | up |
| **GSTK1** | 373156 | ENSG00000197448 | -0.602 | **0.659** | **2.40E-11** | down |
| **NDUFA6** | 4700 | ENSG00000184983 | -0.893 | **0.538** | **2.43E-11** | down |
| **HDHD3** | 81932 | ENSG00000119431 | 0.600 | **1.516** | **3.80E-11** | up |
| **CLPB** | 81570 | ENSG00000162129 | -0.859 | **0.551** | **4.20E-11** | down |
| **RMDN3** | 55177 | ENSG00000137824 | 0.710 | **1.636** | **5.81E-11** | up |
| **TIMM22** | 29928 | ENSG00000177370 | -0.730 | **0.603** | **6.03E-11** | down |
| **TIMM8A** | 1678 | ENSG00000126953 | -1.524 | **0.348** | **6.52E-11** | down |
| **ETFBKMT** | 254013 | ENSG00000139160 | 1.928 | **3.805** | **6.76E-11** | up |
| **NDUFB8** | 4714 | ENSG00000166136 | -0.818 | **0.567** | **8.25E-11** | down |
| **NDUFA9** | 4704 | ENSG00000139180 | -0.850 | **0.555** | **8.40E-11** | down |
| **ISOC2** | 79763 | ENSG00000063241 | -0.860 | **0.551** | **9.62E-11** | down |
| **COMT** | 1312 | ENSG00000093010 | -0.853 | **0.554** | **9.64E-11** | down |
| **GCDH** | 2639 | ENSG00000105607 | -0.656 | **0.635** | **1.13E-10** | down |
| **COQ8A** | 56997 | ENSG00000163050 | 0.766 | **1.701** | **1.26E-10** | up |
| **RBFA** | 79863 | ENSG00000101546 | -0.678 | **0.625** | **1.54E-10** | down |
| **TMEM126B** | 55863 | ENSG00000171204 | -0.687 | **0.621** | **1.62E-10** | down |
| **MRPS9** | 64965 | ENSG00000135972 | -0.632 | **0.645** | **1.68E-10** | down |
| **DHRS2** | 10202 | ENSG00000100867 | 3.476 | **11.130** | **2.59E-10** | up |
| **SFXN2** | 118980 | ENSG00000156398 | -0.770 | **0.587** | **2.71E-10** | down |
| **MMADHC** | 27249 | ENSG00000168288 | -0.914 | **0.531** | **2.90E-10** | down |

| | | | | | | |
|---|---|---|---|---|---|---|
| **SARDH** | 1757 | ENSG00000123453 | 1.849 | **3.602** | **4.48E-10** | up |
| **MRPL3** | 11222 | ENSG00000114686 | -0.883 | **0.542** | **4.57E-10** | down |
| **TIMM50** | 92609 | ENSG00000105197 | -0.585 | **0.667** | **5.91E-10** | down |
| **POLDIP2** | 26073 | ENSG00000004142 | -0.537 | **0.689** | **5.97E-10** | down |
| **MRPS5** | 64969 | ENSG00000144029 | -0.438 | **0.738** | **6.67E-10** | down |
| **EPHX2** | 2053 | ENSG00000120915 | 0.940 | **1.919** | **6.91E-10** | up |
| **MRM2** | 29960 | ENSG00000122687 | -0.582 | **0.668** | **7.69E-10** | down |
| **HSPD1** | 3329 | ENSG00000144381 | -0.715 | **0.609** | **8.58E-10** | down |
| **SLC25A10** | 1468 | ENSG00000183048 | -1.279 | **0.412** | **1.03E-09** | down |
| **LYRM9** | 201229 | ENSG00000232859 | 0.838 | **1.788** | **1.16E-09** | up |
| **YRDC** | 79693 | ENSG00000196449 | -0.638 | **0.642** | **1.71E-09** | down |
| **DUS2** | 54920 | ENSG00000167264 | -0.654 | **0.636** | **1.87E-09** | down |
| **SPG7** | 6687 | ENSG00000197912 | 0.757 | **1.691** | **2.26E-09** | up |
| **FDXR** | 2232 | ENSG00000161513 | -0.688 | **0.621** | **2.31E-09** | down |
| **UQCC3** | 790955 | ENSG00000204922 | -0.944 | **0.520** | **2.47E-09** | down |
| **MTPAP** | 55149 | ENSG00000107951 | -0.775 | **0.584** | **2.59E-09** | down |
| **BNIP3** | 664 | ENSG00000176171 | -0.794 | **0.577** | **3.05E-09** | down |
| **COA7** | 65260 | ENSG00000162377 | -1.273 | **0.414** | **3.11E-09** | down |
| **AGPAT4** | 56895 | ENSG00000026652 | 0.872 | **1.830** | **3.59E-09** | up |
| **MRPL57** | 78988 | ENSG00000173141 | -0.847 | **0.556** | **4.00E-09** | down |
| **METTL15** | 196074 | ENSG00000169519 | -0.805 | **0.572** | **4.49E-09** | down |
| **AGMAT** | 79814 | ENSG00000116771 | -1.447 | **0.367** | **4.54E-09** | down |
| **ACAD10** | 80724 | ENSG00000111271 | 0.611 | **1.528** | **4.70E-09** | up |
| **USP30** | 84749 | ENSG00000135093 | 0.928 | **1.903** | **4.99E-09** | up |
| **MTERF3** | 51001 | ENSG00000156469 | -0.652 | **0.636** | **5.00E-09** | down |
| **NUDT8** | 254552 | ENSG00000167799 | -0.963 | **0.513** | **5.15E-09** | down |
| **HSD17B4** | 3295 | ENSG00000133835 | 0.632 | **1.550** | **7.05E-09** | up |
| **MRPL51** | 51258 | ENSG00000111639 | -0.682 | **0.623** | **7.12E-09** | down |
| **MRPS18B** | 28973 | ENSG00000204568 | -0.475 | **0.720** | **9.12E-09** | down |
| **NDUFAF8** | 284184 | ENSG00000224877 | -0.902 | **0.535** | **1.11E-08** | down |
| **BNIP3L** | 665 | ENSG00000104765 | -0.864 | **0.549** | **1.34E-08** | down |
| **COX8A** | 1351 | ENSG00000176340 | -0.806 | **0.572** | **1.47E-08** | down |
| **APOOL** | 139322 | ENSG00000155008 | -1.064 | **0.478** | **1.64E-08** | down |
| **MRPS12** | 6183 | ENSG00000128626 | -0.671 | **0.628** | **1.81E-08** | down |
| **NIPSNAP2** | 2631 | ENSG00000146729 | -0.718 | **0.608** | **1.84E-08** | down |
| **NLRX1** | 79671 | ENSG00000160703 | -0.882 | **0.543** | **2.00E-08** | down |
| **MRPL19** | 9801 | ENSG00000115364 | -0.772 | **0.586** | **2.31E-08** | down |
| **PRKACA** | 5566 | ENSG00000072062 | 0.724 | **1.651** | **2.52E-08** | up |
| **GLRX2** | 51022 | ENSG00000023572 | -0.839 | **0.559** | **2.57E-08** | down |
| **PRELID2** | 153768 | ENSG00000186314 | -1.625 | **0.324** | **2.72E-08** | down |
| **NDUFS6** | 4726 | ENSG00000145494 | -0.679 | **0.624** | **2.79E-08** | down |
| **TOMM40** | 10452 | ENSG00000130204 | -0.689 | **0.620** | **2.89E-08** | down |
| **FAM210A** | 125228 | ENSG00000177150 | -0.889 | **0.540** | **2.90E-08** | down |
| **PUSL1** | 126789 | ENSG00000169972 | -0.971 | **0.510** | **2.96E-08** | down |
| **AFG1L** | 246269 | ENSG00000135537 | -1.141 | **0.453** | **3.06E-08** | down |
| **FH** | 2271 | ENSG00000091483 | -0.514 | **0.700** | **3.12E-08** | down |
| **THG1L** | 54974 | ENSG00000113272 | -0.643 | **0.640** | **3.14E-08** | down |
| **ALDH5A1** | 7915 | ENSG00000112294 | -1.119 | **0.460** | **3.41E-08** | down |

| | | | | | | |
|---|---|---|---|---|---|---|
| **ACAT1** | 38 | ENSG00000075239 | -0.555 | **0.681** | **4.24E-08** | down |
| **MTCH2** | 23788 | ENSG00000109919 | -0.827 | **0.564** | **4.97E-08** | down |
| **OAT** | 4942 | ENSG00000065154 | 1.331 | **2.516** | **5.25E-08** | up |
| **PRDX4** | 10549 | ENSG00000123131 | -0.905 | **0.534** | **5.26E-08** | down |
| **MRPS17** | 51373 | ENSG00000239789 | -0.766 | **0.588** | **5.56E-08** | down |
| **GPAT2** | 150763 | ENSG00000186281 | -1.676 | **0.313** | **5.84E-08** | down |
| **FASTKD2** | 22868 | ENSG00000118246 | -0.723 | **0.606** | **5.87E-08** | down |
| **SLC25A3** | 5250 | ENSG00000075415 | -0.715 | **0.609** | **5.93E-08** | down |
| **MRPL48** | 51642 | ENSG00000175581 | -0.668 | **0.629** | **5.98E-08** | down |
| **COA4** | 51287 | ENSG00000181924 | -0.864 | **0.550** | **5.98E-08** | down |
| **MRPS15** | 64960 | ENSG00000116898 | -0.895 | **0.538** | **5.98E-08** | down |
| **MRPL27** | 51264 | ENSG00000108826 | -0.867 | **0.548** | **6.13E-08** | down |
| **GHITM** | 27069 | ENSG00000165678 | -0.627 | **0.648** | **7.33E-08** | down |
| **ACSF2** | 80221 | ENSG00000167107 | 0.760 | **1.693** | **7.36E-08** | up |
| **MRPL46** | 26589 | ENSG00000259494 | -0.633 | **0.645** | **9.27E-08** | down |
| **MRPS18C** | 51023 | ENSG00000163319 | -0.735 | **0.601** | **9.90E-08** | down |
| **TCAIM** | 285343 | ENSG00000179152 | -0.664 | **0.631** | **1.05E-07** | down |
| **MRPS2** | 51116 | ENSG00000122140 | -0.608 | **0.656** | **1.08E-07** | down |
| **RSAD1** | 55316 | ENSG00000136444 | 0.572 | **1.487** | **1.12E-07** | up |
| **PGS1** | 9489 | ENSG00000087157 | 0.667 | **1.588** | **1.21E-07** | up |
| **LARS2** | 23395 | ENSG00000011376 | -0.703 | **0.614** | **1.28E-07** | down |
| **TIMM13** | 26517 | ENSG00000099800 | -0.548 | **0.684** | **1.37E-07** | down |
| **NDUFAF2** | 91942 | ENSG00000164182 | -0.713 | **0.610** | **1.43E-07** | down |
| **MRPS24** | 64951 | ENSG00000062582 | -0.675 | **0.626** | **1.55E-07** | down |
| **DARS2** | 55157 | ENSG00000117593 | -0.652 | **0.637** | **1.65E-07** | down |
| **HIBADH** | 11112 | ENSG00000106049 | 0.562 | **1.476** | **1.95E-07** | up |
| **MRPS25** | 64432 | ENSG00000131368 | -0.570 | **0.673** | **1.96E-07** | down |
| **TRMU** | 55687 | ENSG00000100416 | -0.645 | **0.640** | **2.17E-07** | down |
| **CHCHD2** | 51142 | ENSG00000106153 | -0.597 | **0.661** | **2.39E-07** | down |
| **IDH3A** | 3419 | ENSG00000166411 | -0.574 | **0.672** | **2.52E-07** | down |
| **ACO2** | 50 | ENSG00000100412 | -0.567 | **0.675** | **3.12E-07** | down |
| **TIMM44** | 10469 | ENSG00000104980 | -0.938 | **0.522** | **3.24E-07** | down |
| **CRAT** | 1384 | ENSG00000095321 | 0.769 | **1.704** | **3.36E-07** | up |
| **UQCRFS1** | 7386 | ENSG00000169021 | -0.804 | **0.573** | **3.38E-07** | down |
| **TOP3A** | 7156 | ENSG00000177302 | -0.695 | **0.618** | **3.38E-07** | down |
| **ATP5MF** | 9551 | ENSG00000241468 | -0.796 | **0.576** | **3.40E-07** | down |
| **MRPS26** | 64949 | ENSG00000125901 | -0.605 | **0.657** | **4.00E-07** | down |
| **TMEM11** | 8834 | ENSG00000178307 | -0.664 | **0.631** | **4.05E-07** | down |
| **NDUFA10** | 4705 | ENSG00000130414 | -0.512 | **0.701** | **4.17E-07** | down |
| **NIPSNAP3A** | 25934 | ENSG00000136783 | -0.802 | **0.573** | **4.29E-07** | down |
| **CHCHD7** | 79145 | ENSG00000170791 | -0.660 | **0.633** | **4.33E-07** | down |
| **NSUN3** | 63899 | ENSG00000178694 | -0.647 | **0.639** | **4.41E-07** | down |
| **CYCS** | 54205 | ENSG00000172115 | -0.596 | **0.661** | **4.64E-07** | down |
| **COA1** | 55744 | ENSG00000106603 | 0.397 | **1.317** | **5.56E-07** | up |
| **NDUFAF4** | 29078 | ENSG00000123545 | -0.631 | **0.646** | **5.78E-07** | down |
| **LYPLA1** | 10434 | ENSG00000120992 | -0.753 | **0.593** | **6.02E-07** | down |
| **LETMD1** | 25875 | ENSG00000050426 | -0.439 | **0.738** | **6.04E-07** | down |
| **MRRF** | 92399 | ENSG00000148187 | -0.548 | **0.684** | **6.12E-07** | down |

| | | | | | | |
|---|---|---|---|---|---|---|
| **ACLY** | 47 | ENSG00000131473 | -0.560 | **0.678** | **6.21E-07** | down |
| **SAMM50** | 25813 | ENSG00000100347 | -0.595 | **0.662** | **6.28E-07** | down |
| **TIMM9** | 26520 | ENSG00000100575 | -0.462 | **0.726** | **6.32E-07** | down |
| **NIF3L1** | 60491 | ENSG00000196290 | -0.503 | **0.706** | **7.08E-07** | down |
| **MRPL58** | 3396 | ENSG00000167862 | -0.528 | **0.693** | **7.52E-07** | down |
| **ERAL1** | 26284 | ENSG00000132591 | -0.660 | **0.633** | **7.65E-07** | down |
| **SLC25A43** | 203427 | ENSG00000077713 | -0.637 | **0.643** | **8.15E-07** | down |
| **PARL** | 55486 | ENSG00000175193 | -0.672 | **0.628** | **8.61E-07** | down |
| **FDPS** | 2224 | ENSG00000160752 | -1.035 | **0.488** | **8.62E-07** | down |
| **GPT2** | 84706 | ENSG00000166123 | -1.143 | **0.453** | **8.66E-07** | down |
| **HEMK1** | 51409 | ENSG00000114735 | 0.532 | **1.446** | **9.20E-07** | up |
| **NUDT19** | 390916 | ENSG00000213965 | -0.655 | **0.635** | **9.46E-07** | down |
| **CMPK2** | 129607 | ENSG00000134326 | -1.240 | **0.423** | **9.96E-07** | down |
| **ACACA** | 31 | ENSG00000278540 | -0.647 | **0.639** | **1.02E-06** | down |
| **SSBP1** | 6742 | ENSG00000106028 | -0.515 | **0.700** | **1.07E-06** | down |
| **GPX4** | 2879 | ENSG00000167468 | 0.590 | **1.506** | **1.15E-06** | up |
| **MRPL9** | 65005 | ENSG00000143436 | -0.839 | **0.559** | **1.18E-06** | down |
| **TRAP1** | 10131 | ENSG00000126602 | -0.561 | **0.678** | **1.23E-06** | down |
| **CBR3** | 874 | ENSG00000159231 | 2.358 | **5.127** | **1.24E-06** | up |
| **CMC2** | 56942 | ENSG00000103121 | -0.513 | **0.701** | **1.26E-06** | down |
| **SFXN4** | 119559 | ENSG00000183605 | -0.803 | **0.573** | **1.29E-06** | down |
| **PTPMT1** | 114971 | ENSG00000110536 | -0.690 | **0.620** | **1.34E-06** | down |
| **PRORP** | 9692 | ENSG00000100890 | -0.795 | **0.576** | **1.36E-06** | down |
| **MPST** | 4357 | ENSG00000128309 | 0.759 | **1.692** | **1.41E-06** | up |
| **MRPL30** | 51263 | ENSG00000185414 | -0.449 | **0.733** | **1.53E-06** | down |
| **COMTD1** | 118881 | ENSG00000165644 | -0.934 | **0.524** | **1.54E-06** | down |
| **TIMM8B** | 26521 | ENSG00000150779 | -0.636 | **0.643** | **1.61E-06** | down |
| **MRPL28** | 10573 | ENSG00000086504 | -0.523 | **0.696** | **1.79E-06** | down |
| **METTL5** | 29081 | ENSG00000138382 | -0.556 | **0.680** | **1.84E-06** | down |
| **NFU1** | 27247 | ENSG00000169599 | -0.556 | **0.680** | **1.88E-06** | down |
| **PRELID3A** | 10650 | ENSG00000141391 | -1.024 | **0.492** | **2.06E-06** | down |
| **ABCB6** | 10058 | ENSG00000115657 | -0.937 | **0.522** | **2.32E-06** | down |
| **MRPS22** | 56945 | ENSG00000175110 | -0.464 | **0.725** | **2.37E-06** | down |
| **LIG3** | 3980 | ENSG00000005156 | -0.728 | **0.604** | **2.37E-06** | down |
| **SPRYD4** | 283377 | ENSG00000176422 | -0.758 | **0.591** | **2.39E-06** | down |
| **HEBP1** | 50865 | ENSG00000013583 | 1.512 | **2.851** | **2.60E-06** | up |
| **MTERF4** | 130916 | ENSG00000122085 | 0.628 | **1.545** | **2.69E-06** | up |
| **UQCC2** | 84300 | ENSG00000137288 | -0.734 | **0.601** | **2.92E-06** | down |
| **ME2** | 4200 | ENSG00000082212 | -0.743 | **0.597** | **2.98E-06** | down |
| **TMEM70** | 54968 | ENSG00000175606 | -0.591 | **0.664** | **3.02E-06** | down |
| **SLC25A6** | 293 | ENSG00000169100 | -0.558 | **0.679** | **3.25E-06** | down |
| **AKAP10** | 11216 | ENSG00000108599 | -0.578 | **0.670** | **3.40E-06** | down |
| **SNAP29** | 9342 | ENSG00000099940 | -0.548 | **0.684** | **3.65E-06** | down |
| **TSFM** | 10102 | ENSG00000123297 | -0.525 | **0.695** | **3.92E-06** | down |
| **NME6** | 10201 | ENSG00000172113 | -0.783 | **0.581** | **4.25E-06** | down |
| **ACACB** | 32 | ENSG00000076555 | 1.642 | **3.120** | **4.74E-06** | up |
| **TIMM10B** | 26515 | ENSG00000132286 | -0.615 | **0.653** | **4.89E-06** | down |
| **LYRM2** | 57226 | ENSG00000083099 | -0.635 | **0.644** | **5.30E-06** | down |

| | | | | | | |
|---|---|---|---|---|---|---|
| **PGAM5** | 192111 | ENSG00000247077 | -0.455 | **0.729** | **6.14E-06** | down |
| **UQCC1** | 55245 | ENSG00000101019 | -0.390 | **0.763** | **6.37E-06** | down |
| **EXOG** | 9941 | ENSG00000157036 | -0.720 | **0.607** | **6.64E-06** | down |
| **ISCA1** | 81689 | ENSG00000135070 | -0.581 | **0.668** | **6.96E-06** | down |
| **LDHB** | 3945 | ENSG00000111716 | -0.542 | **0.687** | **7.45E-06** | down |
| **NSUN4** | 387338 | ENSG00000117481 | -0.687 | **0.621** | **8.02E-06** | down |
| **SLC25A34** | 284723 | ENSG00000162461 | 1.273 | **2.417** | **8.02E-06** | up |
| **DDX28** | 55794 | ENSG00000182810 | -0.596 | **0.661** | **8.49E-06** | down |
| **COX4I2** | 84701 | ENSG00000131055 | 3.130 | **8.752** | **8.79E-06** | up |
| **LIPT2** | 387787 | ENSG00000175536 | -1.969 | **0.255** | **9.08E-06** | down |
| **CEP89** | 84902 | ENSG00000121289 | -0.825 | **0.564** | **9.86E-06** | down |
| **DNAJC19** | 131118 | ENSG00000205981 | -0.577 | **0.670** | **1.11E-05** | down |
| **FASN** | 2194 | ENSG00000169710 | -0.978 | **0.508** | **1.28E-05** | down |
| **MRPL22** | 29093 | ENSG00000082515 | -0.546 | **0.685** | **1.29E-05** | down |
| **PDF** | 64146 | ENSG00000258429 | -0.811 | **0.570** | **1.31E-05** | down |
| **AARS2** | 57505 | ENSG00000124608 | -0.445 | **0.734** | **1.33E-05** | down |
| **BLOC1S1** | 2647 | ENSG00000135441 | -0.756 | **0.592** | **1.45E-05** | down |
| **SLC25A37** | 51312 | ENSG00000147454 | 0.657 | **1.577** | **1.45E-05** | up |
| **ABCB7** | 22 | ENSG00000131269 | -0.562 | **0.677** | **1.55E-05** | down |
| **MCRIP2** | 84331 | ENSG00000172366 | -0.680 | **0.624** | **1.56E-05** | down |
| **COX10** | 1352 | ENSG00000006695 | -0.641 | **0.641** | **1.60E-05** | down |
| **BBC3** | 27113 | ENSG00000105327 | -0.643 | **0.640** | **1.64E-05** | down |
| **MIX23** | 131076 | ENSG00000160124 | -0.632 | **0.645** | **1.65E-05** | down |
| **ATP5MC1** | 516 | ENSG00000159199 | -0.870 | **0.547** | **1.66E-05** | down |
| **TOMM5** | 401505 | ENSG00000175768 | -0.627 | **0.647** | **1.76E-05** | down |
| **DNLZ** | 728489 | ENSG00000213221 | -0.860 | **0.551** | **1.78E-05** | down |
| **GSTZ1** | 2954 | ENSG00000100577 | -0.496 | **0.709** | **1.85E-05** | down |
| **PET117** | 100303755 | ENSG00000232838 | -0.936 | **0.523** | **1.90E-05** | down |
| **KGD4** | 92259 | ENSG00000134056 | -0.658 | **0.634** | **1.95E-05** | down |
| **MTHFS** | 10588 | ENSG00000136371 | -0.821 | **0.566** | **2.04E-05** | down |
| **HSD17B8** | 7923 | ENSG00000204228 | 0.854 | **1.808** | **2.04E-05** | up |
| **SLC25A38** | 54977 | ENSG00000144659 | -0.362 | **0.778** | **2.08E-05** | down |
| **RTN4IP1** | 84816 | ENSG00000130347 | 0.588 | **1.503** | **2.12E-05** | up |
| **LRPPRC** | 10128 | ENSG00000138095 | -0.543 | **0.686** | **2.20E-05** | down |
| **EHHADH** | 1962 | ENSG00000113790 | 1.151 | **2.221** | **2.21E-05** | up |
| **SIRT4** | 23409 | ENSG00000089163 | 1.298 | **2.459** | **2.30E-05** | up |
| **NDUFA8** | 4702 | ENSG00000119421 | -0.554 | **0.681** | **2.43E-05** | down |
| **AGPAT5** | 55326 | ENSG00000155189 | 0.590 | **1.505** | **2.44E-05** | up |
| **TFAM** | 7019 | ENSG00000108064 | -0.862 | **0.550** | **2.50E-05** | down |
| **NDUFB11** | 54539 | ENSG00000147123 | -0.504 | **0.705** | **2.56E-05** | down |
| **SLC25A33** | 84275 | ENSG00000171612 | -0.554 | **0.681** | **2.61E-05** | down |
| **KARS1** | 3735 | ENSG00000065427 | -0.447 | **0.733** | **2.71E-05** | down |
| **ACP6** | 51205 | ENSG00000162836 | 0.555 | **1.469** | **2.89E-05** | up |
| **MTFR1L** | 56181 | ENSG00000117640 | 0.439 | **1.356** | **2.93E-05** | up |
| **HSDL1** | 83693 | ENSG00000103160 | 0.771 | **1.707** | **2.97E-05** | up |
| **RDH14** | 57665 | ENSG00000240857 | -0.617 | **0.652** | **3.00E-05** | down |
| **VDAC2** | 7417 | ENSG00000165637 | -0.438 | **0.738** | **3.03E-05** | down |
| **SUPV3L1** | 6832 | ENSG00000156502 | -0.645 | **0.640** | **3.08E-05** | down |

| | | | | | | |
|---|---|---|---|---|---|---|
| **ELAC2** | 60528 | ENSG00000006744 | -0.360 | **0.779** | **3.21E-05** | down |
| **ATP5F1A** | 498 | ENSG00000152234 | -0.442 | **0.736** | **3.24E-05** | down |
| **UQCC4** | 283951 | ENSG00000174109 | -0.661 | **0.632** | **3.30E-05** | down |
| **MSRB3** | 253827 | ENSG00000174099 | 2.385 | **5.222** | **3.32E-05** | up |
| **MRPS35** | 60488 | ENSG00000061794 | -0.439 | **0.738** | **3.45E-05** | down |
| **TAMM41** | 132001 | ENSG00000144559 | -0.519 | **0.698** | **3.77E-05** | down |
| **RAB5IF** | 55969 | ENSG00000101084 | -0.393 | **0.762** | **4.01E-05** | down |
| **PNKD** | 25953 | ENSG00000127838 | 0.394 | **1.314** | **4.05E-05** | up |
| **NIT2** | 56954 | ENSG00000114021 | -0.432 | **0.741** | **4.17E-05** | down |
| **HADHA** | 3030 | ENSG00000084754 | -0.375 | **0.771** | **4.29E-05** | down |
| **ECH1** | 1891 | ENSG00000104823 | -0.562 | **0.677** | **4.97E-05** | down |
| **PDP1** | 54704 | ENSG00000164951 | 1.019 | **2.027** | **5.09E-05** | up |
| **TMEM205** | 374882 | ENSG00000105518 | 0.447 | **1.363** | **5.09E-05** | up |
| **PRDX6** | 9588 | ENSG00000117592 | -0.384 | **0.766** | **5.20E-05** | down |
| **ATP5MC2** | 517 | ENSG00000135390 | -0.511 | **0.702** | **5.45E-05** | down |
| **UQCRQ** | 27089 | ENSG00000164405 | -0.531 | **0.692** | **5.79E-05** | down |
| **TMLHE** | 55217 | ENSG00000185973 | -0.667 | **0.630** | **6.09E-05** | down |
| **MRPL49** | 740 | ENSG00000149792 | -0.406 | **0.755** | **6.10E-05** | down |
| **ME3** | 10873 | ENSG00000151376 | 1.169 | **2.249** | **6.14E-05** | up |
| **ALDH18A1** | 5832 | ENSG00000059573 | -0.454 | **0.730** | **6.70E-05** | down |
| **CPT1B** | 1375 | ENSG00000205560 | 0.435 | **1.352** | **6.95E-05** | up |
| **THEM4** | 117145 | ENSG00000159445 | 0.438 | **1.355** | **6.99E-05** | up |
| **GATM** | 2628 | ENSG00000171766 | -0.703 | **0.614** | **7.00E-05** | down |
| **MRPL24** | 79590 | ENSG00000143314 | -0.598 | **0.661** | **7.26E-05** | down |
| **ACAA2** | 10449 | ENSG00000167315 | -0.547 | **0.684** | **7.42E-05** | down |
| **MRPS34** | 65993 | ENSG00000074071 | -0.393 | **0.761** | **7.70E-05** | down |
| **ACSL1** | 2180 | ENSG00000151726 | 0.520 | **1.434** | **7.75E-05** | up |
| **ACADSB** | 36 | ENSG00000196177 | 0.588 | **1.503** | **7.89E-05** | up |
| **CRLS1** | 54675 | ENSG00000088766 | -0.583 | **0.667** | **8.04E-05** | down |
| **PC** | 5091 | ENSG00000173599 | 1.057 | **2.080** | **9.02E-05** | up |
| **PHB1** | 5245 | ENSG00000167085 | -0.624 | **0.649** | **9.27E-05** | down |
| **NT5DC2** | 64943 | ENSG00000168268 | -0.839 | **0.559** | **9.95E-05** | down |
| **MRPL2** | 51069 | ENSG00000112651 | -0.452 | **0.731** | **1.09E-04** | down |
| **MRPS11** | 64963 | ENSG00000181991 | -0.416 | **0.750** | **1.09E-04** | down |
| **PNPT1** | 87178 | ENSG00000138035 | -0.505 | **0.705** | **1.14E-04** | down |
| **C15orf61** | 145853 | ENSG00000189227 | -0.679 | **0.625** | **1.14E-04** | down |
| **MCEE** | 84693 | ENSG00000124370 | -0.715 | **0.609** | **1.17E-04** | down |
| **AASS** | 10157 | ENSG00000008311 | 0.914 | **1.885** | **1.19E-04** | up |
| **MAIP1** | 79568 | ENSG00000162972 | -0.561 | **0.678** | **1.20E-04** | down |
| **IDE** | 3416 | ENSG00000119912 | -0.454 | **0.730** | **1.24E-04** | down |
| **OSBPL1A** | 114876 | ENSG00000141447 | 2.679 | **6.403** | **1.29E-04** | up |
| **COQ7** | 10229 | ENSG00000167186 | -0.639 | **0.642** | **1.29E-04** | down |
| **MRPL13** | 28998 | ENSG00000172172 | -0.535 | **0.690** | **1.30E-04** | down |
| **CHCHD3** | 54927 | ENSG00000106554 | -0.543 | **0.686** | **1.37E-04** | down |
| **ATP5PO** | 539 | ENSG00000241837 | -0.515 | **0.700** | **1.44E-04** | down |
| **OGG1** | 4968 | ENSG00000114026 | -0.614 | **0.653** | **1.46E-04** | down |
| **NDUFAF5** | 79133 | ENSG00000101247 | -0.685 | **0.622** | **1.53E-04** | down |
| **MRPL45** | 84311 | ENSG00000278845 | -0.443 | **0.736** | **1.58E-04** | down |

| | | | | | | |
|---|---|---|---|---|---|---|
| **PAM16** | 51025 | ENSG00000217930 | -0.653 | **0.636** | **1.60E-04** | down |
| **MTHFD2** | 10797 | ENSG00000065911 | -0.847 | **0.556** | **1.65E-04** | down |
| **GATC** | 283459 | ENSG00000257218 | -0.346 | **0.787** | **1.65E-04** | down |
| **MECR** | 51102 | ENSG00000116353 | -0.579 | **0.669** | **1.66E-04** | down |
| **ATP5F1B** | 506 | ENSG00000110955 | -0.542 | **0.687** | **1.77E-04** | down |
| **FDX2** | 112812 | ENSG00000267673 | -0.575 | **0.671** | **1.83E-04** | down |
| **DHODH** | 1723 | ENSG00000102967 | -0.583 | **0.667** | **1.83E-04** | down |
| **SLC25A46** | 91137 | ENSG00000164209 | -0.552 | **0.682** | **1.84E-04** | down |
| **MTX2** | 10651 | ENSG00000128654 | -0.422 | **0.746** | **1.85E-04** | down |
| **TRMT5** | 57570 | ENSG00000126814 | -0.491 | **0.712** | **1.92E-04** | down |
| **SERHL2** | 253190 | ENSG00000183569 | 1.350 | **2.548** | **2.01E-04** | up |
| **MGME1** | 92667 | ENSG00000125871 | -0.461 | **0.727** | **2.02E-04** | down |
| **SDHB** | 6390 | ENSG00000117118 | -0.453 | **0.731** | **2.08E-04** | down |
| **TRNT1** | 51095 | ENSG00000072756 | -0.410 | **0.753** | **2.16E-04** | down |
| **NDUFC1** | 4717 | ENSG00000109390 | -0.550 | **0.683** | **2.17E-04** | down |
| **METTL4** | 64863 | ENSG00000101574 | -0.742 | **0.598** | **2.18E-04** | down |
| **NUDT5** | 11164 | ENSG00000165609 | -0.429 | **0.743** | **2.24E-04** | down |
| **PCBD2** | 84105 | ENSG00000132570 | -0.756 | **0.592** | **2.33E-04** | down |
| **MRPS14** | 63931 | ENSG00000120333 | -0.545 | **0.685** | **2.37E-04** | down |
| **ATAD1** | 84896 | ENSG00000138138 | -0.465 | **0.724** | **2.42E-04** | down |
| **CYB5R3** | 1727 | ENSG00000100243 | 0.414 | **1.332** | **2.43E-04** | up |
| **MCL1** | 4170 | ENSG00000143384 | 0.472 | **1.387** | **2.55E-04** | up |
| **DNM1L** | 10059 | ENSG00000087470 | -0.357 | **0.781** | **2.65E-04** | down |
| **DBI** | 1622 | ENSG00000155368 | -0.513 | **0.701** | **2.67E-04** | down |
| **ATPAF1** | 64756 | ENSG00000123472 | -0.415 | **0.750** | **2.72E-04** | down |
| **MRPS27** | 23107 | ENSG00000113048 | -0.437 | **0.739** | **2.92E-04** | down |
| **DELE1** | 9812 | ENSG00000081791 | -0.345 | **0.787** | **3.03E-04** | down |
| **DMAC1** | 90871 | ENSG00000137038 | -0.676 | **0.626** | **3.03E-04** | down |
| **MRPL47** | 57129 | ENSG00000136522 | -0.339 | **0.791** | **3.12E-04** | down |
| **RPUSD4** | 84881 | ENSG00000165526 | -0.434 | **0.740** | **3.13E-04** | down |
| **YME1L1** | 10730 | ENSG00000136758 | -0.568 | **0.675** | **3.14E-04** | down |
| **MRPL36** | 64979 | ENSG00000171421 | -0.415 | **0.750** | **3.28E-04** | down |
| **ABHD10** | 55347 | ENSG00000144827 | -0.403 | **0.756** | **3.29E-04** | down |
| **GTPBP10** | 85865 | ENSG00000105793 | -0.736 | **0.600** | **3.35E-04** | down |
| **STOML2** | 30968 | ENSG00000165283 | -0.480 | **0.717** | **3.76E-04** | down |
| **MRPL18** | 29074 | ENSG00000112110 | -0.447 | **0.733** | **3.83E-04** | down |
| **HINT1** | 3094 | ENSG00000169567 | -0.444 | **0.735** | **3.96E-04** | down |
| **ACADVL** | 37 | ENSG00000072778 | 0.480 | **1.395** | **4.14E-04** | up |
| **PDHX** | 8050 | ENSG00000110435 | -0.649 | **0.638** | **4.24E-04** | down |
| **DNAJC28** | 54943 | ENSG00000177692 | 1.672 | **3.186** | **4.35E-04** | up |
| **FARS2** | 10667 | ENSG00000145982 | -0.667 | **0.630** | **5.02E-04** | down |
| **COX18** | 285521 | ENSG00000163626 | -0.462 | **0.726** | **5.12E-04** | down |
| **ATP5IF1** | 93974 | ENSG00000130770 | -0.523 | **0.696** | **5.28E-04** | down |
| **MRPL54** | 116541 | ENSG00000183617 | -0.386 | **0.765** | **5.42E-04** | down |
| **SLC25A19** | 60386 | ENSG00000125454 | -0.562 | **0.677** | **5.72E-04** | down |
| **HOGA1** | 112817 | ENSG00000241935 | 1.595 | **3.020** | **6.01E-04** | up |
| **ACAD8** | 27034 | ENSG00000151498 | 0.478 | **1.393** | **6.02E-04** | up |
| **COQ9** | 57017 | ENSG00000088682 | -0.396 | **0.760** | **6.12E-04** | down |

| | | | | | | |
|---|---|---|---|---|---|---|
| **IDH3G** | 3421 | ENSG00000067829 | -0.490 | **0.712** | **6.26E-04** | down |
| **ARMC10** | 83787 | ENSG00000170632 | -0.607 | **0.657** | **6.35E-04** | down |
| **CLPX** | 10845 | ENSG00000166855 | -0.699 | **0.616** | **6.36E-04** | down |
| **GADD45GIP1** | 90480 | ENSG00000179271 | -0.502 | **0.706** | **6.51E-04** | down |
| **MRPL16** | 54948 | ENSG00000166902 | -0.465 | **0.725** | **6.93E-04** | down |
| **ACOT9** | 23597 | ENSG00000123130 | 0.346 | **1.271** | **6.94E-04** | up |
| **PPM1K** | 152926 | ENSG00000163644 | -0.527 | **0.694** | **6.97E-04** | down |
| **DHTKD1** | 55526 | ENSG00000181192 | 0.391 | **1.311** | **7.00E-04** | up |
| **DNAJC11** | 55735 | ENSG00000007923 | -0.336 | **0.792** | **7.03E-04** | down |
| **MICOS10** | 440574 | ENSG00000173436 | -0.579 | **0.669** | **7.50E-04** | down |
| **NAGS** | 162417 | ENSG00000161653 | 0.977 | **1.968** | **7.60E-04** | up |
| **COX8C** | 341947 | ENSG00000187581 | 3.220 | **9.318** | **7.91E-04** | up |
| **DLAT** | 1737 | ENSG00000150768 | -0.708 | **0.612** | **8.17E-04** | down |
| **CCDC90B** | 60492 | ENSG00000137500 | -0.367 | **0.775** | **8.84E-04** | down |
| **MGARP** | 84709 | ENSG00000137463 | 2.230 | **4.692** | **9.46E-04** | up |
| **OCIAD2** | 132299 | ENSG00000145247 | -0.374 | **0.771** | **9.47E-04** | down |
| **GOT2** | 2806 | ENSG00000125166 | -0.340 | **0.790** | **1.01E-03** | down |
| **STARD7** | 56910 | ENSG00000084090 | 0.458 | **1.373** | **1.02E-03** | up |
| **BCO2** | 83875 | ENSG00000197580 | 0.933 | **1.909** | **1.02E-03** | up |
| **SCP2** | 6342 | ENSG00000116171 | -0.511 | **0.702** | **1.05E-03** | down |
| **TWNK** | 56652 | ENSG00000107815 | -0.452 | **0.731** | **1.08E-03** | down |
| **NT5M** | 56953 | ENSG00000205309 | 0.672 | **1.594** | **1.09E-03** | up |
| **AURKAIP1** | 54998 | ENSG00000175756 | -0.423 | **0.746** | **1.11E-03** | down |
| **PTCD1** | 26024 | ENSG00000106246 | -0.370 | **0.774** | **1.13E-03** | down |
| **SLC25A44** | 9673 | ENSG00000160785 | 0.511 | **1.425** | **1.13E-03** | up |
| **HIGD1A** | 25994 | ENSG00000181061 | -0.427 | **0.744** | **1.14E-03** | down |
| **PDHB** | 5162 | ENSG00000168291 | -0.411 | **0.752** | **1.15E-03** | down |
| **PTRH2** | 51651 | ENSG00000141378 | -0.483 | **0.715** | **1.15E-03** | down |
| **GUF1** | 60558 | ENSG00000151806 | -0.341 | **0.790** | **1.22E-03** | down |
| **MDH2** | 4191 | ENSG00000146701 | -0.496 | **0.709** | **1.23E-03** | down |
| **EXD2** | 55218 | ENSG00000081177 | 0.493 | **1.408** | **1.26E-03** | up |
| **COX7B** | 1349 | ENSG00000131174 | -0.547 | **0.684** | **1.29E-03** | down |
| **ISCU** | 23479 | ENSG00000136003 | 0.347 | **1.272** | **1.34E-03** | up |
| **MCCC2** | 64087 | ENSG00000131844 | -0.335 | **0.793** | **1.38E-03** | down |
| **TIMM23** | 100287932 | ENSG00000265354 | -0.318 | **0.802** | **1.45E-03** | down |
| **SLC25A23** | 79085 | ENSG00000125648 | 0.484 | **1.398** | **1.46E-03** | up |
| **ETFRF1** | 144363 | ENSG00000205707 | 0.554 | **1.469** | **1.48E-03** | up |
| **PRODH** | 5625 | ENSG00000100033 | 1.316 | **2.490** | **1.50E-03** | up |
| **CHDH** | 55349 | ENSG00000016391 | -0.395 | **0.761** | **1.50E-03** | down |
| **NDUFS7** | 374291 | ENSG00000115286 | -0.476 | **0.719** | **1.54E-03** | down |
| **COA5** | 493753 | ENSG00000183513 | -0.552 | **0.682** | **1.58E-03** | down |
| **LETM1** | 3954 | ENSG00000168924 | -0.394 | **0.761** | **1.60E-03** | down |
| **MRPL4** | 51073 | ENSG00000105364 | -0.282 | **0.823** | **1.72E-03** | down |
| **BCKDHA** | 593 | ENSG00000248098 | -0.499 | **0.708** | **1.75E-03** | down |
| **CPOX** | 1371 | ENSG00000080819 | 0.742 | **1.672** | **1.76E-03** | up |
| **SLC25A45** | 283130 | ENSG00000162241 | -0.535 | **0.690** | **1.98E-03** | down |
| **SLC25A25** | 114789 | ENSG00000148339 | -0.602 | **0.659** | **1.99E-03** | down |
| **AKAP1** | 8165 | ENSG00000121057 | 0.537 | **1.450** | **1.99E-03** | up |

| | | | | | | |
|---|---|---|---|---|---|---|
| **SERAC1** | 84947 | ENSG00000122335 | 1.040 | **2.056** | **2.01E-03** | up |
| **ATP5F1C** | 509 | ENSG00000165629 | -0.499 | **0.708** | **2.03E-03** | down |
| **SLC25A36** | 55186 | ENSG00000114120 | -0.369 | **0.774** | **2.08E-03** | down |
| **PDHA1** | 5160 | ENSG00000131828 | -0.445 | **0.735** | **2.19E-03** | down |
| **FMC1** | 154791 | ENSG00000164898 | -0.602 | **0.659** | **2.28E-03** | down |
| **MTERF2** | 80298 | ENSG00000120832 | 0.432 | **1.349** | **2.30E-03** | up |
| **TXN2** | 25828 | ENSG00000100348 | -0.285 | **0.821** | **2.37E-03** | down |
| **MRPL14** | 64928 | ENSG00000180992 | -0.487 | **0.713** | **2.50E-03** | down |
| **AK2** | 204 | ENSG00000004455 | -0.331 | **0.795** | **2.53E-03** | down |
| **BOK** | 666 | ENSG00000176720 | 3.165 | **8.972** | **2.54E-03** | up |
| **TUFM** | 7284 | ENSG00000178952 | -0.339 | **0.791** | **2.55E-03** | down |
| **SLC25A41** | 284427 | ENSG00000181240 | 1.399 | **2.636** | **2.59E-03** | up |
| **GATB** | 5188 | ENSG00000059691 | -0.492 | **0.711** | **2.65E-03** | down |
| **SDHAF4** | 135154 | ENSG00000154079 | -0.409 | **0.753** | **2.79E-03** | down |
| **NDUFS4** | 4724 | ENSG00000164258 | -0.415 | **0.750** | **2.88E-03** | down |
| **POLG2** | 11232 | ENSG00000256525 | 0.420 | **1.338** | **2.94E-03** | up |
| **NDUFV2** | 4729 | ENSG00000178127 | -0.348 | **0.786** | **2.94E-03** | down |
| **SPTLC2** | 9517 | ENSG00000100596 | -0.434 | **0.740** | **2.95E-03** | down |
| **OPA3** | 80207 | ENSG00000125741 | -0.352 | **0.784** | **3.10E-03** | down |
| **MRPS23** | 51649 | ENSG00000181610 | -0.445 | **0.734** | **3.14E-03** | down |
| **MSRB2** | 22921 | ENSG00000148450 | -0.439 | **0.738** | **3.38E-03** | down |
| **BCAT2** | 587 | ENSG00000105552 | -0.385 | **0.766** | **3.70E-03** | down |
| **TIMM17A** | 10440 | ENSG00000134375 | -0.507 | **0.703** | **3.77E-03** | down |
| **TSTD1** | 100131187 | ENSG00000215845 | 0.468 | **1.383** | **4.01E-03** | up |
| **HSD17B10** | 3028 | ENSG00000072506 | -0.387 | **0.765** | **4.11E-03** | down |
| **ABCD2** | 225 | ENSG00000173208 | 0.802 | **1.744** | **4.12E-03** | up |
| **GUK1** | 2987 | ENSG00000143774 | -0.329 | **0.796** | **4.39E-03** | down |
| **ACOT2** | 10965 | ENSG00000119673 | -0.463 | **0.725** | **4.50E-03** | down |
| **WARS2** | 10352 | ENSG00000116874 | -0.474 | **0.720** | **4.50E-03** | down |
| **FOXRED1** | 55572 | ENSG00000110074 | -0.388 | **0.764** | **4.54E-03** | down |
| **NUDT6** | 11162 | ENSG00000170917 | 0.513 | **1.427** | **4.62E-03** | up |
| **PDE2A** | 5138 | ENSG00000186642 | 1.375 | **2.594** | **4.62E-03** | up |
| **BCL2L10** | 10017 | ENSG00000137875 | -0.689 | **0.620** | **4.75E-03** | down |
| **ALDH9A1** | 223 | ENSG00000143149 | -0.324 | **0.799** | **4.90E-03** | down |
| **MT-ND3** | 4537 | ENSG00000198840 | 0.557 | **1.471** | **5.03E-03** | up |
| **MGST3** | 4259 | ENSG00000143198 | 0.453 | **1.369** | **5.09E-03** | up |
| **OXNAD1** | 92106 | ENSG00000154814 | -0.437 | **0.739** | **5.34E-03** | down |
| **AKR7A2** | 8574 | ENSG00000053371 | -0.459 | **0.727** | **5.44E-03** | down |
| **MTIF3** | 219402 | ENSG00000122033 | 0.321 | **1.250** | **5.58E-03** | up |
| **DBT** | 1629 | ENSG00000137992 | -0.393 | **0.762** | **5.71E-03** | down |
| **GSR** | 2936 | ENSG00000104687 | -0.281 | **0.823** | **6.05E-03** | down |
| **PHYH** | 5264 | ENSG00000107537 | 0.472 | **1.387** | **6.18E-03** | up |
| **ETFA** | 2108 | ENSG00000140374 | -0.440 | **0.737** | **6.33E-03** | down |
| **MTX1** | 4580 | ENSG00000173171 | -0.387 | **0.765** | **6.42E-03** | down |
| **MOCS1** | 4337 | ENSG00000124615 | 1.116 | **2.167** | **6.64E-03** | up |
| **MRPL52** | 122704 | ENSG00000172590 | -0.343 | **0.788** | **6.76E-03** | down |
| **PDSS2** | 57107 | ENSG00000164494 | -0.472 | **0.721** | **6.82E-03** | down |
| **SIRT3** | 23410 | ENSG00000142082 | 0.274 | **1.209** | **7.49E-03** | up |

| | | | | | | |
|---|---|---|---|---|---|---|
| **LIPT1** | 51601 | ENSG00000144182 | -0.586 | **0.666** | **7.51E-03** | down |
| **VDAC1** | 7416 | ENSG00000213585 | -0.401 | **0.757** | **7.79E-03** | down |
| **NDUFAB1** | 4706 | ENSG00000004779 | -0.438 | **0.738** | **8.43E-03** | down |
| **ENDOG** | 2021 | ENSG00000167136 | -0.539 | **0.688** | **8.43E-03** | down |
| **HSPE1** | 3336 | ENSG00000115541 | -0.551 | **0.683** | **8.50E-03** | down |
| **FASTKD1** | 79675 | ENSG00000138399 | -0.405 | **0.755** | **8.58E-03** | down |
| **COA3** | 28958 | ENSG00000183978 | -0.424 | **0.745** | **8.63E-03** | down |
| **STAR** | 6770 | ENSG00000147465 | -1.544 | **0.343** | **8.72E-03** | down |
| **SUGCT** | 79783 | ENSG00000175600 | 1.354 | **2.557** | **8.91E-03** | up |
| **BCL2L2** | 599 | ENSG00000129473 | -0.400 | **0.758** | **9.03E-03** | down |
| **HSCB** | 150274 | ENSG00000100209 | -0.396 | **0.760** | **9.13E-03** | down |
| **SCO1** | 6341 | ENSG00000133028 | -0.385 | **0.766** | **9.17E-03** | down |
| **LETM2** | 137994 | ENSG00000165046 | 0.571 | **1.486** | **9.51E-03** | up |
| **BCL2A1** | 597 | ENSG00000140379 | 0.556 | **1.470** | **9.53E-03** | up |
| **SLC25A40** | 55972 | ENSG00000075303 | -0.574 | **0.672** | **9.82E-03** | down |
| **ATP5MJ** | 9556 | ENSG00000156411 | -0.435 | **0.740** | **9.84E-03** | down |
| **NDUFAF7** | 55471 | ENSG00000003509 | 0.362 | **1.285** | **9.90E-03** | up |
| **ABAT** | 18 | ENSG00000183044 | 0.506 | **1.420** | **1.04E-02** | up |
| **MTHFD2L** | 441024 | ENSG00000163738 | -0.543 | **0.686** | **1.04E-02** | down |
| **YBEY** | 54059 | ENSG00000182362 | 0.382 | **1.303** | **1.05E-02** | up |
| **VDAC3** | 7419 | ENSG00000078668 | -0.326 | **0.798** | **1.10E-02** | down |
| **QRSL1** | 55278 | ENSG00000130348 | -0.371 | **0.773** | **1.11E-02** | down |
| **KYAT3** | 56267 | ENSG00000137944 | -0.427 | **0.744** | **1.11E-02** | down |
| **TRUB2** | 26995 | ENSG00000167112 | -0.337 | **0.792** | **1.12E-02** | down |
| **SDHD** | 6392 | ENSG00000204370 | -0.290 | **0.818** | **1.16E-02** | down |
| **SMIM8** | 57150 | ENSG00000111850 | 0.382 | **1.303** | **1.18E-02** | up |
| **DUT** | 1854 | ENSG00000128951 | -0.369 | **0.774** | **1.18E-02** | down |
| **TSTD3** | 100130890 | ENSG00000279170 | -0.664 | **0.631** | **1.19E-02** | down |
| **SPIRE1** | 56907 | ENSG00000134278 | 0.676 | **1.597** | **1.23E-02** | up |
| **OPA1** | 4976 | ENSG00000198836 | -0.585 | **0.667** | **1.25E-02** | down |
| **TIMM29** | 90580 | ENSG00000142444 | -0.382 | **0.767** | **1.27E-02** | down |
| **MRPL42** | 28977 | ENSG00000198015 | -0.354 | **0.783** | **1.30E-02** | down |
| **MRPS28** | 28957 | ENSG00000147586 | 0.282 | **1.216** | **1.35E-02** | up |
| **NDUFA7** | 4701 | ENSG00000267855 | -0.385 | **0.766** | **1.39E-02** | down |
| **MRPL23** | 6150 | ENSG00000214026 | -0.421 | **0.747** | **1.43E-02** | down |
| **COQ10A** | 93058 | ENSG00000135469 | 0.425 | **1.343** | **1.44E-02** | up |
| **SLC25A32** | 81034 | ENSG00000164933 | -0.353 | **0.783** | **1.45E-02** | down |
| **MCUR1** | 63933 | ENSG00000050393 | -0.446 | **0.734** | **1.47E-02** | down |
| **NDUFB9** | 4715 | ENSG00000147684 | -0.275 | **0.826** | **1.54E-02** | down |
| **COX6C** | 1345 | ENSG00000164919 | -0.302 | **0.811** | **1.55E-02** | down |
| **NUBPL** | 80224 | ENSG00000151413 | -0.459 | **0.727** | **1.57E-02** | down |
| **QDPR** | 5860 | ENSG00000151552 | -0.346 | **0.787** | **1.59E-02** | down |
| **MTG1** | 92170 | ENSG00000148824 | 0.330 | **1.257** | **1.59E-02** | up |
| **CRY1** | 1407 | ENSG00000008405 | 0.759 | **1.692** | **1.65E-02** | up |
| **MIGA2** | 84895 | ENSG00000148343 | -0.274 | **0.827** | **1.65E-02** | down |
| **DECR1** | 1666 | ENSG00000104325 | -0.321 | **0.800** | **1.66E-02** | down |
| **FAM162A** | 26355 | ENSG00000114023 | -0.415 | **0.750** | **1.69E-02** | down |
| **UQCRH** | 7388 | ENSG00000173660 | 0.322 | **1.250** | **1.71E-02** | up |

| | | | | | | |
|---|---|---|---|---|---|---|
| **PMAIP1** | 5366 | ENSG00000141682 | 0.559 | **1.473** | **1.73E-02** | up |
| **TOMM22** | 56993 | ENSG00000100216 | -0.333 | **0.794** | **1.73E-02** | down |
| **HDHD5** | 27440 | ENSG00000069998 | -0.276 | **0.826** | **1.75E-02** | down |
| **MMUT** | 4594 | ENSG00000146085 | -0.381 | **0.768** | **1.78E-02** | down |
| **LYPLAL1** | 127018 | ENSG00000143353 | -0.428 | **0.743** | **1.78E-02** | down |
| **COX5B** | 1329 | ENSG00000135940 | -0.287 | **0.819** | **1.82E-02** | down |
| **ARL2** | 402 | ENSG00000213465 | 0.386 | **1.307** | **1.82E-02** | up |
| **COQ2** | 27235 | ENSG00000173085 | -0.353 | **0.783** | **1.87E-02** | down |
| **PTGR3** | 284273 | ENSG00000180011 | -0.354 | **0.783** | **1.87E-02** | down |
| **MT-ND6** | 4541 | ENSG00000198695 | -0.454 | **0.730** | **2.00E-02** | down |
| **TRMT61B** | 55006 | ENSG00000171103 | -0.446 | **0.734** | **2.04E-02** | down |
| **TMEM143** | 55260 | ENSG00000161558 | -0.400 | **0.758** | **2.05E-02** | down |
| **SLC25A20** | 788 | ENSG00000178537 | 0.375 | **1.297** | **2.06E-02** | up |
| **SLC25A29** | 123096 | ENSG00000197119 | 0.387 | **1.308** | **2.10E-02** | up |
| **PRDX3** | 10935 | ENSG00000165672 | -0.273 | **0.828** | **2.11E-02** | down |
| **CPS1** | 1373 | ENSG00000021826 | 0.769 | **1.704** | **2.12E-02** | up |
| **COQ6** | 51004 | ENSG00000119723 | -0.293 | **0.816** | **2.18E-02** | down |
| **THNSL1** | 79896 | ENSG00000185875 | -0.480 | **0.717** | **2.20E-02** | down |
| **OCIAD1** | 54940 | ENSG00000109180 | -0.299 | **0.813** | **2.21E-02** | down |
| **STX17** | 55014 | ENSG00000136874 | 0.297 | **1.229** | **2.26E-02** | up |
| **RIDA** | 10247 | ENSG00000132541 | -0.319 | **0.802** | **2.27E-02** | down |
| **MTO1** | 25821 | ENSG00000135297 | -0.315 | **0.804** | **2.38E-02** | down |
| **FIS1** | 51024 | ENSG00000214253 | 0.276 | **1.211** | **2.39E-02** | up |
| **COX16** | 51241 | ENSG00000133983 | -0.362 | **0.778** | **2.50E-02** | down |
| **MRPL38** | 64978 | ENSG00000204316 | -0.322 | **0.800** | **2.53E-02** | down |
| **MTIF2** | 4528 | ENSG00000085760 | -0.303 | **0.810** | **2.56E-02** | down |
| **ACCS** | 84680 | ENSG00000110455 | 0.476 | **1.390** | **2.63E-02** | up |
| **MRPL43** | 84545 | ENSG00000055950 | 0.297 | **1.229** | **2.64E-02** | up |
| **SLC25A11** | 8402 | ENSG00000108528 | -0.300 | **0.812** | **2.74E-02** | down |
| **ETHE1** | 23474 | ENSG00000105755 | -0.313 | **0.805** | **2.76E-02** | down |
| **METAP1D** | 254042 | ENSG00000172878 | -0.364 | **0.777** | **2.77E-02** | down |
| **UQCR10** | 29796 | ENSG00000184076 | -0.342 | **0.789** | **2.85E-02** | down |
| **MT-CYB** | 4519 | ENSG00000198727 | 0.442 | **1.359** | **2.85E-02** | up |
| **ALDH1B1** | 219 | ENSG00000137124 | -0.446 | **0.734** | **2.87E-02** | down |
| **SLIRP** | 81892 | ENSG00000119705 | -0.385 | **0.766** | **2.89E-02** | down |
| **C2orf69** | 205327 | ENSG00000178074 | -0.444 | **0.735** | **2.89E-02** | down |
| **SQOR** | 58472 | ENSG00000137767 | 0.357 | **1.281** | **2.94E-02** | up |
| **NAXE** | 128240 | ENSG00000163382 | -0.331 | **0.795** | **2.98E-02** | down |
| **ENSG00000283765** | | ENSG00000283765 | -4.827 | **0.035** | **2.98E-02** | down |
| **GLRX5** | 51218 | ENSG00000182512 | -0.312 | **0.805** | **3.03E-02** | down |
| **VPS13D** | 55187 | ENSG00000048707 | 0.329 | **1.257** | **3.04E-02** | up |
| **GCSH** | 2653 | ENSG00000140905 | -0.584 | **0.667** | **3.09E-02** | down |
| **D2HGDH** | 728294 | ENSG00000180902 | 0.281 | **1.215** | **3.21E-02** | up |
| **RHOT2** | 89941 | ENSG00000140983 | -0.270 | **0.829** | **3.27E-02** | down |
| **PRELID3B** | 51012 | ENSG00000101166 | -0.341 | **0.789** | **3.28E-02** | down |
| **GFM1** | 85476 | ENSG00000168827 | -0.323 | **0.799** | **3.29E-02** | down |
| **TOMM70** | 9868 | ENSG00000154174 | -0.269 | **0.830** | **3.32E-02** | down |
| **ADCK2** | 90956 | ENSG00000133597 | 0.294 | **1.226** | **3.49E-02** | up |

| | | | | | | |
|---|---|---|---|---|---|---|
| **NDUFB1** | 4707 | ENSG00000183648 | -0.329 | **0.796** | **3.49E-02** | down |
| **CLYBL** | 171425 | ENSG00000125246 | 0.387 | **1.307** | **3.52E-02** | up |
| **TST** | 7263 | ENSG00000128311 | 0.287 | **1.220** | **3.57E-02** | up |
| **SLC25A26** | 115286 | ENSG00000144741 | -0.452 | **0.731** | **3.65E-02** | down |
| **PPA2** | 27068 | ENSG00000138777 | -0.326 | **0.798** | **3.67E-02** | down |
| **CSKMT** | 751071 | ENSG00000214756 | -0.497 | **0.708** | **3.77E-02** | down |
| **CHCHD1** | 118487 | ENSG00000172586 | -0.287 | **0.819** | **3.82E-02** | down |
| **OXSM** | 54995 | ENSG00000151093 | 0.295 | **1.227** | **3.91E-02** | up |
| **ACAD11** | 84129 | ENSG00000240303 | 0.274 | **1.209** | **3.98E-02** | up |
| **NTHL1** | 4913 | ENSG00000065057 | 0.301 | **1.232** | **4.00E-02** | up |
| **PEX11B** | 8799 | ENSG00000131779 | -0.271 | **0.829** | **4.08E-02** | down |
| **BAD** | 572 | ENSG00000002330 | -0.277 | **0.825** | **4.12E-02** | down |
| **SLC25A16** | 8034 | ENSG00000122912 | -0.421 | **0.747** | **4.14E-02** | down |
| **MICOS13** | 125988 | ENSG00000174917 | -0.280 | **0.824** | **4.15E-02** | down |
| **PTCD2** | 79810 | ENSG00000049883 | -0.471 | **0.721** | **4.23E-02** | down |
| **MRPL33** | 9553 | ENSG00000243147 | -0.286 | **0.820** | **4.23E-02** | down |
| **PCCA** | 5095 | ENSG00000175198 | 0.288 | **1.221** | **4.25E-02** | up |
| **DHRS7B** | 25979 | ENSG00000109016 | -0.339 | **0.790** | **4.31E-02** | down |
| **SLC25A24** | 29957 | ENSG00000085491 | 0.750 | **1.682** | **4.33E-02** | up |
| **SUCLA2** | 8803 | ENSG00000136143 | -0.476 | **0.719** | **4.39E-02** | down |
| **MRPL1** | 65008 | ENSG00000169288 | -0.292 | **0.817** | **4.40E-02** | down |
| **EARS2** | 124454 | ENSG00000103356 | -0.365 | **0.776** | **4.67E-02** | down |
| **LYRM4** | 57128 | ENSG00000214113 | -0.284 | **0.821** | **4.73E-02** | down |
| **CYP11A1** | 1583 | ENSG00000140459 | -0.850 | **0.555** | **4.76E-02** | down |
| **MLYCD** | 23417 | ENSG00000103150 | -0.349 | **0.785** | **4.83E-02** | down |
