## Supplemental Table 3 for "RNA-sequencing of butyrate-treated human lymphoblastoid cell lines shows dysregulated genes of multiple disease-related pathways"

**Table S3: Forward and Reverse primer sequences used for RT-qPCR analysis**

(note: RT-qPCR findings are not displayed for some genes in this table)

| Gene | Forward primer | Reverse primer |
|---|---|---|
| GAPDH | GGAGCGAGATCCCTCCAAAAT | GGCTGTTGTCATACTTCTCATGG |
| SLC6A6 | TGGTGGAGGTGCGTTTCTC | AGAGGTGTACTGGCCTATGATG |
| BATF2 | CCTCATTGCTCTTTGGACTAGG | TGTGTGAAACCTGGGTGAAG |
| CD99 | AACCCACCCAAACCGATGCC | TGAAAAGCTACCGGAGGAACTA |
| LY6E | GGGAATCTCGTGACATTTGGC | ACACCAACATTGACGCCTTCT |
| ISG15 | CTCTGAGCATCCTGGTGAGGAA | AAGGTCAGCCAGAACAGGTCGT |
| HDAC1 | GGTCCAAATGCAGGCGATTCCT | TCGGAGAACTCTTCCTCACAGG |
| HDAC2 | CTCATGCACCTGGTGTCCAGAT | GCTATCCGCTTGTCTGATGCTC |
| HDAC3 | GAGTTCTGCTCGCGTTACACAG | CGTTGACATAGCAGAAGCCAGAG |
| HDAC5 | CGCTGAGAATGGCTTTACTGGC | GTGTAGAGGCTGAACTGGTTGG |
| HDAC6 | GCCTCAATCACTGAGACCATCC | GGTGCCTTCTTGGTGACCAACT |
| RPL19 | TCACAGCCTGTACCTGAAGGTG | CGTGCTTCCTTGGTCTTAGACC |
| RPL32 | ACAAAGCACATGCTGCCCAGTG | TTCCACGATGGCTTTGCGGTTC |
| ASMTL | CTGCGGATATTGCCAGCAAAGTG | GAGTATTCGCCATCCGATGCCA |
| IER3 | GCAGCCGCAGGGTTCTCTACC | CTCTTCAGCCATCAGGATCTGG |
| BCAS1 | CCAAGAACCTCTCTGATGGCGT | TTCAGGCTCTGGCGGTGTGATA |
| HMGB1 | GCGAAGAAACTGGGAGAGATGTG | GCATCAGGCTTTCCTTTAGCTCG |
| HMGN2 | AAACCTGCTCCTCCAAAGCCAG | CTTGCCAGCATCAGCTTTTCCC |
| HDGF | CCAAAGACCTCTTCCCTTACGAG | TGGTTCAGGCTCTTCCACACAG |
